# Beyond the ink: cellular and molecular effects of iron-based pigments on macrophages

**DOI:** 10.1101/2025.06.30.662256

**Authors:** Marianne Vitipon, Esther Akingbagbohun, Fabienne Devime, Hélène Diemer, Aurélie Hirschler, Daphna Fenel, Stéphane Ravanel, Christine Carapito, Thierry Rabilloud

## Abstract

As ochre, iron oxide is among the most ancient pigments used by mankind for different purposes, including tattooing as demonstrated on tattoed mummies. Iron oxides are still used in tattooing nowadays and especially in dermopigmentation, an area of medical tattoing aiming at restoring the color of skin. This ancient use of iron oxide does not mean that it has no effect on cells, and especially on macrophages, the cells that maintain pigments particles on site in tattoos. We thus investigated in vitro the delayed/sustained effects of iron oxide pigments on macrophages, i.e. the effects occurring a few days after the exposure to pigments, on pigments-loaded macrophages but in a pigment-free medium, mimicking the status of tattooed skin after all the pigment particles have been captured. By combining proteomic and targeted approaches, we determined that red iron oxide (but not black iron oxide) induces perturbations in mitochondria, altering the mitochondrial transmembrane potential. Red iron oxide also induces oxidative stress and the secretion of pro-inflammatory cytokines such as interleukin 6 and tumor necrosis factor. Thus, red iron oxide induces adverse effects on macrophages that may persist over time, owing to its low intracellular dissolution.

## 1. Introduction

Today, tattooing is increasingly popular across Europe. It is estimated that over 60 million people - approximately 12% of the European population - have at least one tattoo. This number is steadily rising, particularly among younger people [1]. Tattooing is defined as the practice of creating permanent markings in the skin through the intradermal injection of inks made of colorants and auxiliary substances [2]. Technically, tattoo inks are dispersions, composed of insoluble pigments (dispersed phase) finely distributed within a continuous liquid medium. This medium typically consists of water, simple alcohols, polyols (such as glycerin and propylene glycol), and binders [3]. Binders, like polyvinylpyrrolidone or natural gums, enhance pigment stability and prevent sedimentation by maintaining uniform dispersion of the pigment in the liquid phase [4]. The selection of these components is critical to ensure the ink’s stability, ease of application, and safety for intradermal use [2].

Although tattoo inks are administered intradermally, they are not classified as medical or pharmaceutical substances and thus are not subject to the rigorous certification standards applied to injectable products [1]. Consequently, their ingredients often lack comprehensive testing, and the quality of tattoo inks largely depends on individual manufacturers’ standards. The composition of tattoo inks is frequently undefined and highly variable, potentially including allergenic, toxic or potentially carcinogenic compounds [3]. For instance, inks may contain heavy metals such as chromium, cobalt, lead, nickel, and mercury, as well as aromatic amines, phthalates, and polycyclic aromatic hydrocarbons, which are associated with various health risks [1].

One subfield of tattooing, referred to as dermopigmentation, micropigmentation, or dermatography is applied for example in permanent makeup, and currently one of the fastest-growing fields in the beauty industry [2]. Dermopigmentation also encompasses medical tattooing, which aims to camouflage post-surgical scars, correcting pigment loss in vitiligo, or restoring appearance after disease or trauma - such as nipple-areola complex reconstruction following breast cancer surgery [5,6].

Medical tattooing holds its therapeutic purpose and falls under the purview of medical device regulations, highlighting the necessity of adherence to stringent standards that ensure safety and efficacy. This classification mandates that manufacturers and practitioners comply with both medical device legislation and relevant chemical safety standards [2].

Iron oxide pigments, historically used in art and tattooing, remain widely applied today, particularly in the permanent makeup and medical dermopigmentation industries. Indeed, those pigments can take on various shades - red iron oxide Fe_2_O_3_ (CI77491/very stable), black iron oxide Fe_3_O_4_ (CI77499/stable), yellow iron oxide FeOOH (CI77492/unstable) – and be used to match skin tone shades [2,6,7]. The benefits of iron oxide pigments are their low price, excellent colour robustness toward sun exposure and other ambient conditions. Although iron oxide pigments are generally not associated with immediate allergic reactions, adverse reactions can occur under specific conditions [8], a precautionary recommendation is to ensure adequate iron levels in iron-deficient patients prior to dermopigmentation, as transdermal absorption of iron from the tattoo site may occur [6].

Besides their use in dermopigmentation, iron oxide nanoparticles are of particular interest in biomedical applications, as they can be utilised for diagnostic imaging – such as tumor visualisation – as well as for drug delivery, treatment of anaemia [9], or experimental cancer therapies [10]. Therefore, iron oxide toxicity has been extensively investigated in recent years on different tissues [11] and different cell types [12–14]. Depending on the specific cell type, concentration, duration of exposure and the nature of the iron oxide particles, significant adverse effects can also be induced, including the generation of reactive oxygen species (ROS), inflammatory responses, and mitochondrial dysfunction, as exemplified on neural tissue [15]. Thus, iron oxide particles may not be devoid of adverse effects, even in the frame of medical tattooing.

Tattooing, including techniques like dermopigmentation, disrupts tissue integrity and damages both the epidermis and dermis [1]. It was long assumed that dermal fibroblasts served as the primary long-term reservoir for pigment particles within the skin. However, more recent studies have demonstrated that, although many cell types are found to contain pigment particles in tattooed skin [16], pigment retention at the injection site is mainly mediated by dermal macrophages, known as melanophage [17,18]. Indeed the “capture-release-recapture” model has been refined by integrating the concept of continuous macrophage renewal in the dermis. This renewal is driven by the recruitment of circulating monocytes, which replenish the pool of dermal macrophages to compensate for the loss of cells through natural turnover. Therefore, following an intradermal injection of ink, macrophages engulf the pigment and ensure the tattoo’s persistence over time, through this cycle of capture (by macrophages present on site), release (upon macrophage death) and recapture by new macrophages recruited on site. However, the recapture step is not perfect, which explains how pigment particles can be found in short lived cells such as keratinocytes, for which some pigments can be quite toxic [19], even in old tattoos [16] . Furthermore, it is currently not clear whether this weakness of the pigment recapture contributes significantly to the presence of pigments in the draining lymph nodes of tattooed individuals [20], or if the pigment present in lymph nodes derives from the uninternalized pigment during the tattooing process, pigment that will be carried directly by the lymph to the lymph nodes and then further in liver Kupffer cells [21]. It should be kept in mind that particles-loaded macrophages can migrate in the body [22,23], so that it cannot be surely derived that the pigment found in Kupffer cells has been carried as free particles in the blood.

This updated model raises new questions regarding the impact of pigment phagocytosis on macrophage cellular physiology as iron oxide pigments have been found intracellularly in at least mouse dermal macrophages ninety days post-tattooing [24]. As macrophages are key players in inflammation and tissue homeostasis [25,26], it is worth investigating the impact of iron oxide particles on macrophages functions, especially in view of the known presence of pigments in non dermal macrophages [20,21], sometimes with associated clinical signs [27] .

The vast majority of the work investigating the effects of iron oxide nanoparticles on macrophages (e.g. in [13,14] has been carried out immediately after exposure of the cells to the particles. In the case of tattooing, delayed effects are very relevant to investigate, in the frame of the “capture-release-recapture” model [17], and of the persistence of iron oxide over time in tattoos [24]. To this purpose, we implemented a system in which we exposed J774A.1 murine macrophages to iron oxide pigments for 24 hours, followed by a five-day recovery period in fresh medium to mimic post-exposure persistence within dermal tissue. Two pigments were selected for analysis: PR101 (red iron oxide) and PBk11 (black iron oxide), both representative of pigments frequently used in tattoo formulations for dermopigmentation. In order to obtain a wide appraisal of macrophage responses after the recovery period, we used a proteomic analysis, which we complemented with functional assays to evaluate proinflammatory effects, including altered responses to bacterial challenge, changes in mitochondrial membrane potential, and intracellular glutathione (GSH) levels. Together, these approaches provide insight into the cellular adaptations and physiological consequences associated with pigment retention in macrophages, contributing to a better understanding of the long-term impact of tattoo inks on dermis homeostasis.

## 2. Material and methods

### 2.1 Pigment particles preparation and characterization

Pigment Red 101 (red iron oxide, Fe2O3, ref. PS-MI0050) was purchased from Kama Pigments (Montreal, Canada). Pigment Black 11 (black iron oxide, Fe3O4, ref. 48422) was purchased from Kremer Pigment (Aichstetten, Germany). Corundum (aluminium oxide spinel, Al2O3, ref. ERMFD066) was purchased from Merck (Saint-Quentin-Fallavier, France). The pigments were received as dry powders and dispersed at 100 mg/mL in an aqueous solution of arabic gum (100 mg/ml), previously sterilised overnight at 80°C in humid atmosphere. Pigment’s dispersions were then re-sterilised under the same conditions to minimise microbial contamination. To reduce agglomerates and aggregates, the dispersions were sonicated in a Vibra Cell VC 750 sonicator (VWR, Fontenay-sous-Bois, France) equipped with a Cup Horn probe. Sonication was carried out in pulse mode for 30 minutes (1 s ON/ 1 s OFF) at 60% amplitude, corresponding to 90W per pulse, in volume not exceeding 3 mL to ensure efficient energy transfer. Dispersions were stored at 4 °C and gently stirred before each experiment to avoid sediment accumulation on tube walls. Prior to use, pigment dispersions were sonicated for 15 minutes in an ultrasonic bath, then diluted in sterile water to intermediate concentrations as required. To limit potential sedimentation and microbial contamination over time, fresh pigment preparations were renewed at least once per month.

For the in solution characterization of the pigments dispersions, Dynamic Light Scattering (DLS) and Electrophoretic Light Scattering (ELS) were performed using a Litesizer 200 instrument (Anton Paar, Les Ulis, France) equipped with an Omega reusable cuvette (225288, Anton Paar), suitable for both measurements. Before introducing the sample, the zeta potential of the cuvette filled with PBS 0.001X alone was measured to verify cleanliness and exclude background signal or instrument drift. Pigment dispersions were diluted to a final concentration of 10 µg/mL in 0.001X PBS to ensure adequate particle presence for detection while maintaining optimal optical transmittance and ionic strength for electrophoretic mobility analysis. DLS measurements were conducted at 25 °C, and two sequential measurements were recorded with a 1-minute interval to monitor potential sedimentation effects. The hydrodynamic size distribution was expressed as number-weighted intensity. ELS measurements were conducted at 25 °C, three measurements were taken, each separated by a 30-second delay. Optical parameters were automatically adjusted by the instrument to ensure accurate signal acquisition for each sample.

Transmission Electronic Microscopy (TEM) was performed as previously described [28]. Briefly, 10 µL of a 100µg/mL dispersion were added to a glow discharge grid coated with a carbon supporting film for 5 minutes. The excess solution was soaked off using a filter paper and the grid was air-dried. The images were taken under low dose conditions (<10 e-/Å2) with defocus values comprised between 1.2 and 2.5 mm on a Tecnai 12 LaB6 electron microscope at 120 kV accelerating voltage using a 4k x 4k CEMOS TVIPS F416 camera.

### 2.2. Cell culture and treatment

The mouse macrophage cell line J774A.1 was purchased from the European Cell Culture Collection (Salisbury, UK). For cell maintenance, cells were cultured in DMEM supplemented with 10% of fetal bovine serum (FBS) in non-adherent flasks (Cellstar flasks for suspension culture, Greiner Bio One, Les Ulis, France). Cells were split every 2 days at 200,000 cells/ml and harvested at 1 million cells/ml.

#### 2.2.1. Determination of Sublethal Dose (LD_20_)

To determine the sublethal dose 20 (LD_20_) for each pigment, cells were exposed to a range of pigment concentrations, and viability was assessed using the VVBlue assay [29] after a 24-hour exposure period. Briefly, J774A.1 macrophages were seeded at 500,000 cells/mL in 24-well adherent plates in DMEM supplemented with 1% horse serum, 1% HEPES, and 10,000 units/mL penicillin-streptomycin as described before [30] to prevent cell proliferation and pigment dilution through time and let rest for 24 hours before pigment exposure. To account for potential interference from pigment absorbance, blank controls containing pigments without cells were included for each condition. This allowed us to correct for background signal in VVBlue assay. Absorbance readings were obtained using a BMG Labtech FLUOstar Omega® plate reader. The LD_20_ value was defined as the concentration inducing approximately 20% reduction in viability compared to the untreated control. Afterward, the dose used was determined by comparing the cytotoxicity of PR101, PBk11 and ERM.

#### 2.2.2. Recovery exposure settings

Cells were seeded at 500,000 cells/ml in DMEM supplemented with horse serum, as described above. Pigments were added at the selected concentration for 24 further hours as for an acute exposure. Afterwards, the medium was changed in order to remove remaining pigments. The medium was then renewed every 24 hours until the end of the 5 day-recovery period. At the end of the exposure, cells were treated and harvested depending on the functional test performed. For the viability test, the VVBlue assay was used as described above.

### 2.3. Cellular quantification of metals by inductively coupled plasma-mass spectrometry (ICP-MS)

Cells were seeded as previously described into 12-well adherent plates. After the recovery period, cells were rinsed with PBS and harvested into 2 mL Eppendorf tubes. Samples were centrifuged for 5 minutes at 1200 rpm, the supernatant was carefully removed, and 200 µL of ICP lysis buffer (5mM Hepes NaOH pH 7.5, 0.75 mM spermine tetrahydrohloride, 0.1% (w/v) dodecyltrimethylammonio-propane-sulfonate (SB 3-12)) was added to the pellet. The cell pellet was thoroughly vortexed to ensure complete lysis. To separate the soluble and insoluble fractions, lysates were centrifuged at 15,000 g for 30 minutes. The supernatant was transferred into a fresh Eppendorf tube. Samples were stored at -20°C until analysis. The resulting supernatant (soluble fraction) was expected to contain soluble metal ions or degradation products, while the pellet (solid fraction) retained any intracellular, undissolved pigment particles. Both fractions, along with raw pigment suspensions at the working dilution, were analysed by Inductively Coupled Plasma Mass Spectrometry (ICP-MS) to quantify the elemental metal content. Protein concentration in each cell sample was determined using the Bradford assay to normalize metal content to total protein levels. For the ICP-MS analysis, samples were dehydrated to dryness and digested at 90°C for 4 hours using aqua regia made of three parts of 30% (w/v) hydrochloric acid and one part of 65% (w/v) nitric acid for PR101 and PBk11. Mineralized samples were analyzed using an iCAP RQ quadrupole mass spectrometer (Thermo Fisher Scientific). Concentrations were determined using standard curves made from serial dilutions of a multi-element solution (ICP multi-element standard IV, Merck) and corrected using an internal standard solution containing 45Sc and 103Rh, added online. Iron was determined using 56Fe and 57Fe data collected in the helium collision mode. Data integration was done using the Qtegra software (Thermo Fisher Scientific).

### 2.4. Proteomics

Proteomics was carried out essentially as described previously [31]. However, the experimental details are given here for the sake of consistency

#### 2.4.1. Sample preparation

After exposure to the pigment particles and the recovery period, the cells were harvested by flushing the 6 well plates. They were collected by centrifugation (300g, 5 minutes) and rinsed twice in PBS. The cell pellets were lysed in 100 µl of extraction buffer (4M urea, 2.5% cetyltrimethylammonium chloride (CTAC), 100mM sodium phosphate buffer pH 3, 150µM methylene blue). The extraction was let to proceed at room temperature for 30 minutes, after which the lysate was centrifuged (15,000g, 15 minutes) to pellet the nucleic acids. The supernatants were then stored at -20°C until use.

#### 2.4.2. Shotgun proteomics

For the shotgun proteomic analysis, the samples were included in polyacrylamide plugs according to Muller et al. [32] with some modifications to downscale the process [33]. To this purpose, the photopolymerization system using methylene blue, toluene sulfinate and diphenyliodonium chloride was used [34].

As mentioned above, the methylene blue was included in the cell lysis buffer. The other initiator solutions consisted in a 1 M solution of sodium toluene sulfinate in water and in a saturated water solution of diphenyliodonium chloride. The ready-to-use polyacrylamide solution consisted of 1.2 ml of a commercial 40% acrylamide/bis solution (37.5/1) to which 100 µl of diphenyliodonium chloride solution, 100 µl of sodium toluene sulfinate solution and 100 µl of water were added.

To the protein samples (16 µl), 4 µl of acrylamide solution were added and mixed by pipetting in a 500µl conical polypropylene microtube. 100 µl of water-saturated butanol were then layered on top of the samples, and polymerization was carried out under a 1500 lumen 2700K LED lamp for 2 hours, during which the initially blue gel solution discolored. At the end of the polymerization period, the butanol was removed, and the gel plugs were fixed for 1 hr with 200 µl of 30% ethanol 2 % phosphoric acid, followed by 3x 15 minutes washes in 20% ethanol. The fixed gel plugs were then stored at -20°C until use.

Gel plug processing, digestion, peptide extraction and nanoLC-MS/MS was performed as previously described, without the robotic protein handling system and using a nanoACQUITY Ultra-Performance-LC (Waters Corporation, Milford, USA) coupled to a Q-Exactive HF-Plus mass spectrometer (Thermo Fisher Scientific, Bremen, Germany).

For protein identification, the MS/MS data were interpreted using a local Mascot server with MASCOT 2.6.2 algorithm (Matrix Science, London, UK) against an in-house database containing all *Mus musculus* and *Rattus norvegicus* entries from UniProtKB/SwissProt (version 2019_10, 25,156 sequences) and the corresponding 25,156 reverse entries. Spectra were searched with a mass tolerance of 10 ppm for MS and 0.05 Da for MS/MS data, allowing a maximum of one trypsin missed cleavage. Trypsin was specified as enzyme. Acetylation of protein N-termini, carbamidomethylation of cysteine residues and oxidation of methionine residues were specified as variable modifications. Identification results were imported into Proline software version 2.1 (https://www.profiproteomics.fr/proline/) for validation. Peptide Spectrum Matches (PSM) with pretty rank equal to one were retained. False Discovery Rate was then optimized to be below 1% at PSM level using Mascot Adjusted E-value and below 1% at Protein Level using Mascot Mudpit score.

Mass spectrometry data are available via ProteomeXchange Consortium via the PRIDE [35] partner repository with the dataset identifier PXD064985 and 10.6019/PXD064985.

#### 2.4.3. Label Free Quantification

Peptides abundances were extracted thanks to Proline software version 2.2 [36] using a m/z tolerance of 10 ppm. Alignment of the LC-MS runs was performed using Loess smoothing. Cross Assignment was performed within groups only. Protein abundances were computed by sum of peptides abundances (normalized using the median).

#### 2.4.4. Data analysis

For the global analysis of the protein abundances data, missing data were imputed with a low, non-null value. The complete abundance dataset was then analyzed by the PAST software [37].

Proteins were considered as significantly different if their p value in the Mann-Whitney U-test against control values was inferior to 0.05. The selected proteins were then submitted to pathway analysis using the DAVID tool [38], with a cutoff value set at a FDR of 0.2.

### 2.5 Assessment of Cellular Functional Parameters

To evaluate mitochondrial membrane potential, intracellular glutathione content, and reactive oxygen species (ROS) production, cells were seeded in 12-well adherent plates and exposed to iron-based pigments as previously described. After the recovery period, cells were incubated with their respective probes.

For the mitochondrial membrane potential assay, Rhodamine 123 (Rh123) was added at a final concentration of 10 µM and incubated for 30 minutes at 37 °C. Butanedione monoxime (BDM, 30 mM) was used as a positive control and carbonyl cyanide 4-(trifluoromethoxy)phenylhydrazone (FCCP, 5 µM) as a negative control, both added concurrently with Rh123.

For ROS detection, Dihydrorhodamine 123 (DHR123) was used at a final concentration of 100 µM, with cells incubated for 30 minutes at 37 °C. Menadione was used as a positive control at 50 and 75 µM, added 2 hours prior to probe incubation.

To assess intracellular glutathione (GSH) levels, cells were incubated with Monochlorobimane (MCB, 25 µM) for 4 minutes at 37 °C. CDNB (25 µM) was added 30 minutes prior to MCB as a negative control to deplete GSH.

Following incubation with each probe, cold 1X PBS was added to each well, and plates were incubated for 5 minutes at 4 °C to halt further metabolic activity. Approximately two-thirds of the supernatant was removed after a 3 minutes centrifugation at 1200 rpm, and the remaining medium was transferred into 2 ml microtubes. To collect non-adherent cells, wells were rinsed once with cold PBS; this rinse was added to the corresponding tubes. The tubes were centrifuged at 300 × g for 5 minutes at 4 °C. Following centrifugation, 200 µL of freshly prepared CTAC lysis buffer (2.5% CTAC, 10mM Hepes NaOH, pH 7.5, 2mM MgCl2) was added directly into each well. The plates were placed on ice and gently agitated on a rotating table to ensure complete cell lysis. The supernatants from the initial cell collection were discarded, and the lysates from each well were transferred into the corresponding tubes. To separate the soluble protein fraction, samples were centrifuged at 15,000 × g for 30 minutes at 4 °C. The resulting supernatants were transferred into fresh tubes. For each replicate (n = 4 per condition), 50 µL of the lysate was transferred into a black 96-well plate for immediate fluorescence reading using a BMG Labtech FLUOstar Omega® plate reader (excitation/emission filters: MCB [355-20/460 nm], Rho123 and DHR123 [485-12/520 nm]).). The remaining lysates were stored at −20 °C for subsequent protein quantification using the Bradford assay.

For each assay, corresponding no-probe controls were prepared under identical conditions (same incubation times, treatments, and processing) to evaluate autofluorescence originating from cells, residual pigments, and the lysis buffer. Fluorescence values, with or without probes, were normalized to total protein content. Afterward, without probes controls fluorescence values were subtracted from the corresponding samples to correct for background signals. For each experiment, conditions were performed in four replicates.

To assess lysosomal activity, cells were seeded as previously described in 12-well adherent plates. After the recovery period, the medium was replaced and Neutral Red (prepared in 50% ethanol) was added to the fresh medium at a final concentration of 20 µg/ml. Cells were incubated for 2 hours at 37 °C to allow dye accumulation in lysosomes. Following incubation, cells were carefully washed twice with PBS and neutral red was eluted with 1 ml of acid-alcohol solution (1% acetic acid, 50% ethanol v/v). Plates were incubated on a rotating table at room temperature for 30 minutes, and the released red dye was quantified by measuring absorbance at 540 nm using a BMG Labtech FLUOstar Omega® plate reader. To normalize lysosomal activity to cell numbers, the fixed cell layers were rinsed with PBS, and stained with Crystal Violet (4 µg/ml in PBS) for 30 minutes. After two PBS washes, the dye was eluted using 1 ml of the acid-alcohol solution, and absorbance was measured at 590 nm. For each well, the Neutral Red / Crystal Violet absorbance ratio was calculated, and fold changes were expressed relative to unexposed control cells.

### 2.6. Measurement of Cytokines Secretion

To evaluate inflammatory effects and potential disruption of the response to bacterial challenge following the recovery period, cells were either exposed or not to lipopolysaccharide (LPS, 50 ng/mL) for an additional 24 hours. After incubation, supernatants were transfer into a 2mL tube and centrifuged 5 min at 300g. Afterwards, supernatants were transferred into fresh tubes and Tumor Necrosis Factor alpha (TNFα) and Interleukin-6 (IL-6) levels were quantified using murine ELISA kits (Covalab), according to the manufacturer’s instructions; Absorbances were read using a BMG Labtech FLUOstar Omega® plate reader. The remaining adherent cells were lysed as described above and added to the corresponding tubes to determine total protein content. Cytokine concentrations were normalised to the corresponding protein content of each sample. For each experiment, experiments were performed in four replicates.

## 3. Results

### 3.1. Physicochemical characterisation of iron-based pigments

The two iron-based pigments were characterized using Dynamic Light Scattering (DLS) to assess their size distribution, polydispersity index (PI), and sedimentation behaviour. As summarized in **Table 1**, PR101 exhibited a narrower size range (188.76 – 811.09 nm) than PBk11 (83.97 – 1318.64 nm), with corresponding PIs of 16.76% and 28.76%, indicating greater heterogeneity in PBk11. Despite its broader distribution, PR101 had a slightly larger mean hydrodynamic diameter based on number-weighted intensity (339.7 nm) than PBk11 (222.26 nm). Manufacturer data report primary particle sizes of ∼90 nm (PR101) and ∼300 nm (PBk11) in powder form, however larger sizes were observed in dispersions suggesting aggregation. Electrophoretic Light Scattering (ELS) analysis revealed negative zeta potentials (-31.96 mV for PR101, -34.51 mV for PBk11), indicating moderate electrostatic repulsion but insufficient to prevent aggregation. Both pigments sedimented within one minute, with PBk11 settling more rapidly despite its lower density (4.6 g/cm³ for PR101, 5 g/cm³ for PBk11). This may be due to its higher polydispersity, larger particles sizes and more negative zeta potential, reinforcing its strong interparticle interactions. Nonetheless, 90% of particles remained below 500 nm (D90: 484.6 nm for PR101, 350.90 nm for PBk11), confirming that a significant portion of the dispersion remains in the sub-micrometric range.

**Table 1:** Physicochemical characterization of iron oxide pigments.

|  | **PR101** | **PBk11** | **Corundum** |
| --- | --- | --- | --- |
| Size distribution (nm) | [188.76 – 811.09] | [83.97 – 1318.64] | [1400 – 7500] |
| Mean Hydrodynamic diameter (nm) | 339.7 | 222.26 | \ |
| Polydispersity index (%) | 16.76 | 28.76 | \ |
| Peak T0 (nm) | 345.48 | 304.17 | \ |
| Peak T1 (nm) | 333.61 | 117.57 | \ |
| D90 Number (nm) | 484.6 | 350.90 | \ |
| Zeta potential (mV) | -31.96 ± 0.31 | -34.51 ± 0.27 | -34.14 ± 0.60 |

In contrast, representative electron microscopy images in **Figure 1** illustrate the morphological differences between hematite (PR101, Fe_2_O_3_; A) and magnetite (PBk11, Fe_3_O_4_; B). The primary distinction lies in the size of the primary particles. PR101 consists of small spherical particles (<100 nm) that assemble into large aggregates spanning several micrometers, whereas PBk11 is composed of larger spherical particles that also tend to aggregate. This discrepancy suggests that PR101 aggregates more extensively in dispersion, leading to a larger measured size in the DLS analysis despite its smaller primary particle size.

**Figure 1:**
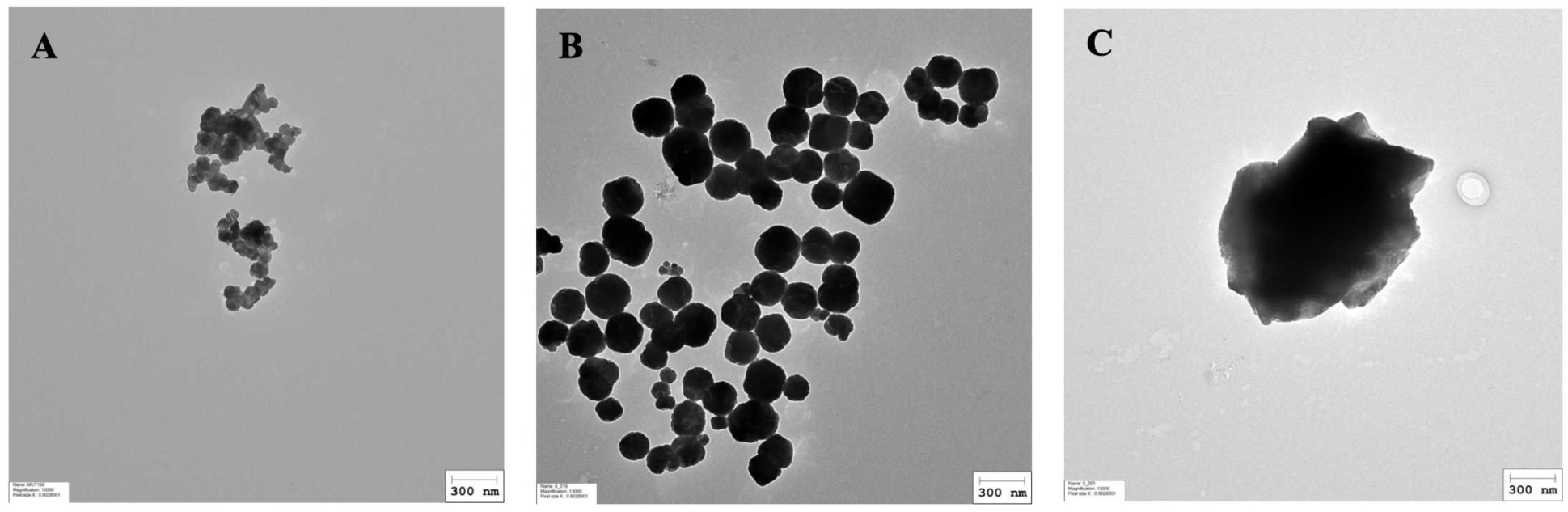
representative electron microscopy images of (A) Pr101, (B) PBk11 and (C) Corundum at 13000x magnification. Particles were prepared as described in section 2.1, diluted at 100µg/mL in sterile water. A 10 µL aliquot was applied to a glow discharge grid coated with a carbon supporting film for 5 minutes. The excess solution was soaked off by a filter paper and the grid was air-dried. The images were taken under low dose conditions (<10 e-/Å2) with defocus values between 1.2 and 2.5 µm on a Tecnai 12 LaB6 electron microscope at 120 kV accelerating voltage using 4k x 4k CEMOS TVIPS F416 camera.

Corundum was used as a negative control to assess pigment-induced cellular effects, based on its well-documented inertness in similar experimental setups [28,39]. This certified reference material (ERM-FD066) has been characterized using standard methods, including laser diffraction and electron microscopy, revealing a size distribution ranging from 1.4 to 7.5 µm [40] . Despite its larger particle size compared to the tested pigments, ELS analysis indicates a comparable surface charge, with a zeta potential of –34.14 mV (Table 1). TEM images further confirm the presence of large, micron-sized particles (Figure 1C).

### 3.2. Cell exposure and dose selection

For pigment exposure, cells were treated as described in **Figure 2A**. Macrophages were seeded at 500,000 cells/mL in DMEM supplemented with 1% horse serum and 1% streptomycin-penicillin in 24-well adherent plates. The following day, pigment dispersions were added for a 24-hour acute exposure.

In a first series of experiments, we determined which concentrations of each pigment could be used on macrophages, by performing viability assays after the acute exposure. In **Figure 2B**, the VVBlue assay [29], chosen to avoid optical interference from pigment opacity, did not reveal any cytotoxicity of the pigments for up to 1400 µg/ml. We then tested the two highest concentrations (1200 and 1400 µg/ml) in the recovery format (.**Figure 2C**). After 5 days post exposure, PR101 showed a 20% lethality at 1200 µg/ml and a 40% lethality at 1400µg/ml. The other pigments did not show any lethality in this format at any of the two tested doses.

Based on these results, 1200 µg/mL was selected for subsequent experiments, which were carried out in the recovery format.

**Figure 2.**
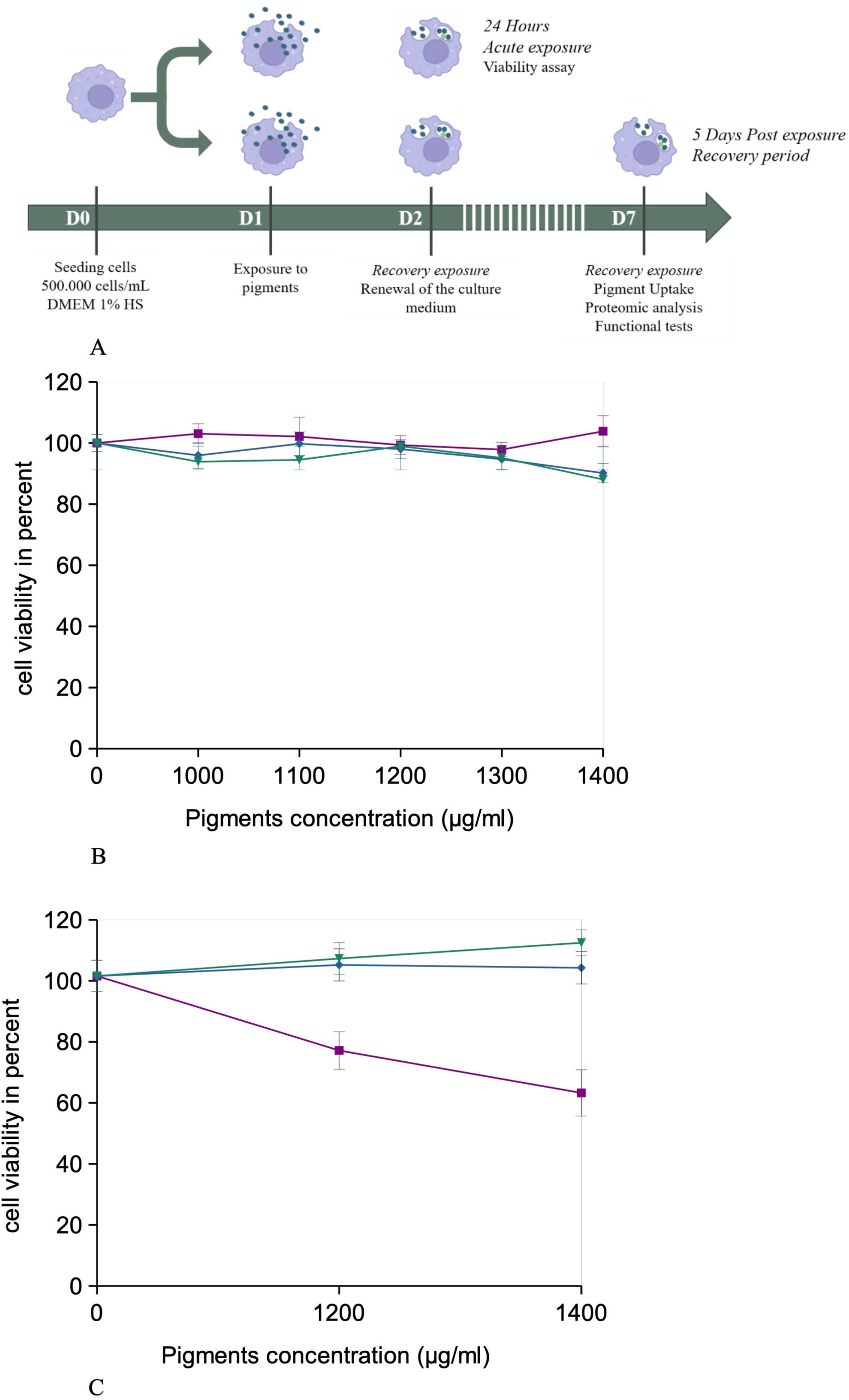
Exposure scheme and viability determination Panel A, details of the recovery exposure scheme Panel B: viability curve immediately after exposure to pigments (24 hours) purple: PR101 blue: PBk11 green: ERM (corundum) Results are expressed as mean ± standard deviation Panel C: viability curve after exposure to pigments for 24 hours followed by 5 days of recovery in a pigment-free medium purple: PR101 blue: PBk11 green: ERM (corundum) Results are expressed as mean ± standard deviation

### 3.3. Pigments uptake and dissolution by cells

In order to obtain a first appraisal of pigment uptake by macrophages, we performed a simple examination of pigments-loaded cells by optical microscopy. The results, displayed on **Figure 3**, showed that the cells were heavily loaded with pigments particles, even after the recovery period. This heavy loading rendered the cells opaque, which explains why we could not apply any optical method, and especially flow cytometry, on the pigments-loaded cells.

**Figure 3:**
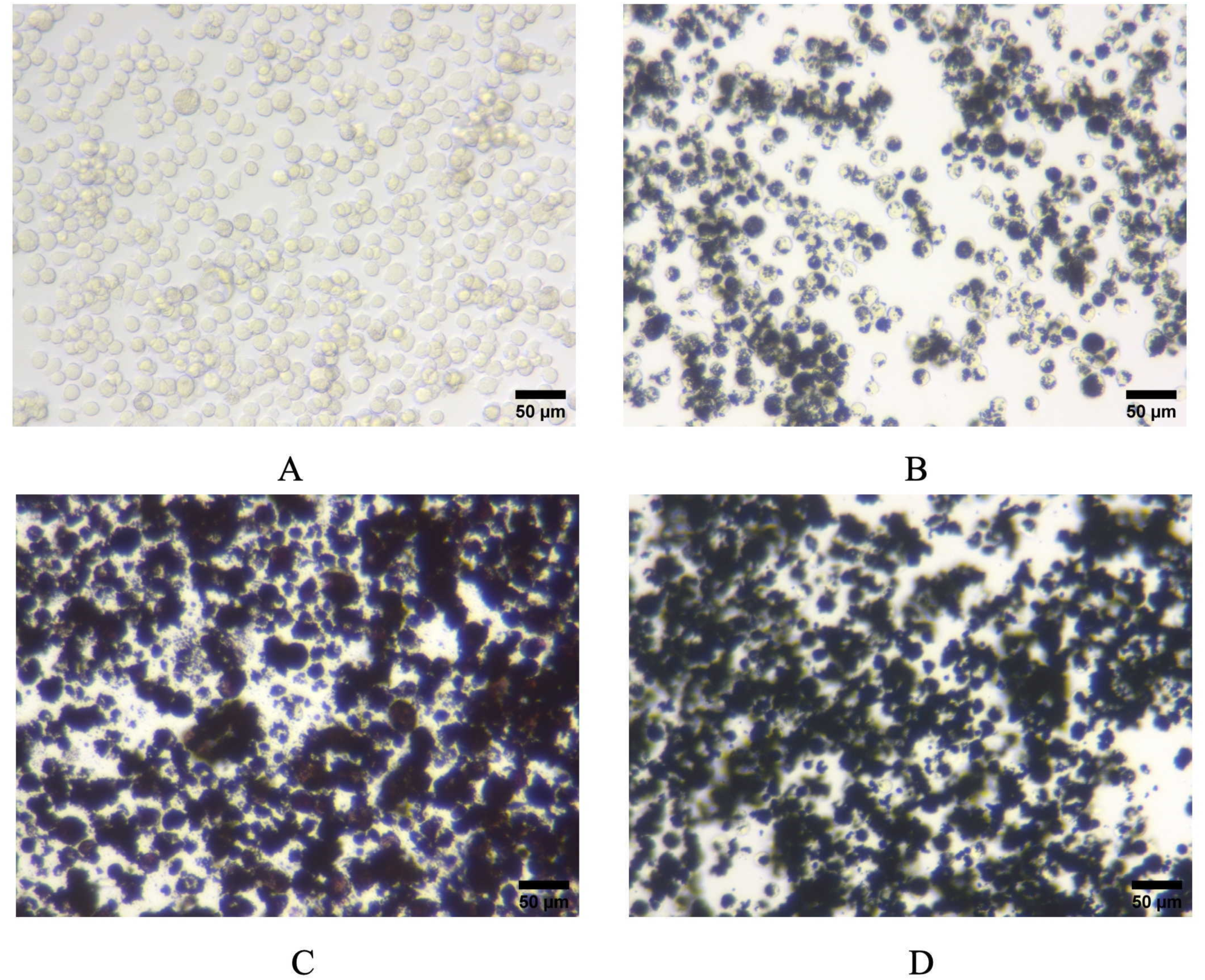
Optical microscopy images of A: control, untreated cells, B: Corundum (ERM066)-treated cells C: PR101-treated cells D: PBk11-treated cells All images were acquired at 10× magnification

To more quantitatively assess pigment uptake after acute exposure and retention following the recovery period, mineralization experiments were conducted on the lysates of the pigments-loaded cells. The supernatant (soluble fraction) gave the proportion of soluble iron, while the pellet (internalized particular fraction) gave the amount of pigment remaining as particular material after the recovery period.

The results, shown in **Table 2**, showed that the pigments were very efficiently internalized by the macrophages, with 78% of the PR101 input and 93% of the PBk11 input being still present in the cells after 5 days of recovery post exposure. As expected for an insoluble pigment the intracellular dissolution was very low, with only 0.008% of the total input iron being detected as soluble for PR101-exposed cells, and 0.03% for the PBk11-exposed cells.

**Table 2:** quantification of internalized and dissolved iron in pigments-exposed macrophages.

|  | control cells | PR101-treated cells | PBk11-treated cells |
| --- | --- | --- | --- |
| Iron input | ND | 734±20 | 783±19 |
| Total cell-associated iron | 0.048±0.015 | 571±44 | 729±22 |
| Non-sedimentable, cell-associated iron | 0.019±0.018 | 0.048±0.015 | 0.22±0.03 |
| Percent iron internalized | ND | 78±6 | 93±3 |
| Percent iron dissolved | ND | 8.5E-05±2.67E-05 | 3.01E-04±3.65E-05 |

The results are expressed in µg iron/culture well
The results are expressed as mean±standard deviation

### 3.4. Global analysis of the proteomic experiments

The shotgun proteomic analysis was able to detect and quantify 3127 proteins (**Table S1**). First, we analyzed the complete dataset via the PAST software, through a principal coordinates analysis. The results, displayed in **Figure 4**, showed a good separation of the PR101-treated cells from the other groups in the first axis of the principal coordinates analysis (describing 56% of the total variance), while the PBk11-treated cells separated from the other groups in the third axis describing 13% of the total variance).

**Figure 4:**
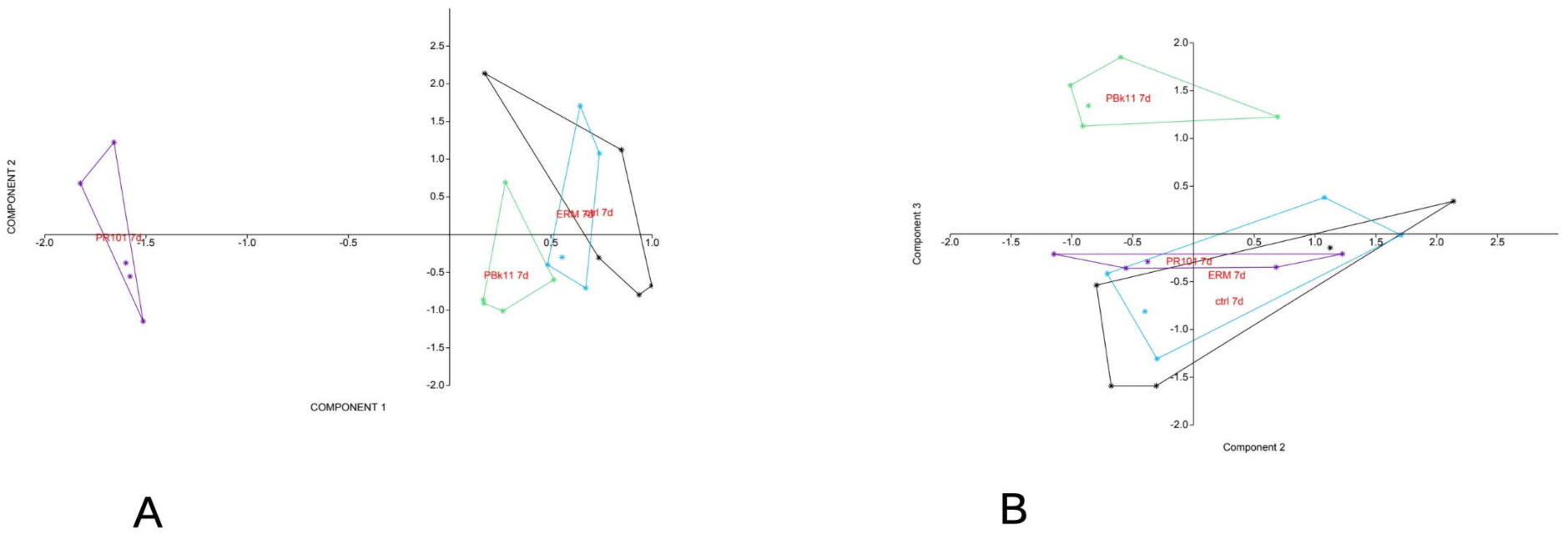
Global analysis of the proteomic data The complete proteomic data table (3127 proteins) was analyzed by Principal Component Analysis, using the PAST software. The results are represented as the X-Y diagram of the first two axes of the Principal Component Analysis, representing 71% of the total variance (A), and the second and third axes, the third axis representing 13% of the total variance (B) . Eigenvalue scale. This representation allows to figure out how, at the global proteome scale, the samples are related to each other. Samples grouped in such a diagram indicate similar proteomes, and the larger the distance between samples are, the more dissimilar their respective proteomes.

The extent of the proteomes modulations is confirmed by the ANalysis Of SIMilarity test [41]. This test showed a p-value (probability by random) of 0.0075 for the PR101 vs. control comparison, 0.014 for the PBk11 vs. control comparison, and 0.007 for the PR101 vs. PBk11 comparison. In the pigment vs corundum (ERM) comparisons, PR101 showed a low p-value (0.0079) while PBk11 showed a higher p-value (0.093). All these data values are consistent with the fact that PR101 induced a much deeper modification of the proteome than PBk11 in our exposure scheme.

In order to select the proteins whose abundances are modulated by the two pigments, we conducted a double Mann-Whitney test. We compared first the pigments-treated cells to the untreated cells, and then the pigments-treated cells to the corundum-treated cells. Only proteins which appeared significantly altered in their abundances in both tests (U≤ 2) were selected (**Tables S2 and S3**).

To gain further insight into the significance of the observed changes, this list of modulated proteins was used to perform pathways analyses using the DAVID software, and the results are shown in **Tables S4 and S5**. The pathway analysis pointed out several cellular processes which were modulated upon treatment with the iron oxide particles. Some were generic, e.g. mitochondria, lysosomes, while some were more specific to macrophages functions, such as immune responses.

Among the generic processes, it was interesting to note proteins that are involved in iron homeostasis (**Table S6**). The two ferritin chains were strongly increased in response to iron oxide particles, and much more in response to PBk11 than to PR101 (22 fold vs 9 fold for the ferritin heavy chain, for example). Heme oxygenase was also strongly induced, but in this case more markedly in response to PR101. In contrast, proteins that deliver iron or help to deliver iron, such as transferrin receptor, ferric-chelate reductase or polyRC binding protein (which is also an iron chaperone [42]), are decreased in response to iron oxide particles.

### 3.5. Mitochondria

Mitochondrial proteins represented an abundant class among the proteins modulated in response to iron oxide particles, with 110 proteins (**Table S7**). Among them, 7 were modulated by both iron oxide particles (5 increases, 2 decreases). These increases encompassed stress proteins (HSP60, P63038 ; HSP70, P38647 ; HSP10, Q64433) and proteins implicated in proteostasis (AFG3L2, Q8JZQ2, a component of the mAAA protease [43] ; PHB1, P67778, a chaperone for inner membrane proteins [44]). In addition, 8 proteins were modulated in response to PBk11 only (5 increases, 3 decreases). Both the PBk11-increased proteins and PBk11-decreased proteins were quite diverse in their functions. Among the 95 proteins that were modulated in response to PR101, 65 were increased and 30 decreased.

As several subunits of the oxidative phosphorylation complexes appeared in the list of modulated proteins such as 5 subunits of ATP synthase, 5 subunits of complex I, 3 subunits of complex III, as well as proteins involved in proteostasis of the inner membrane proteins (mAAA protease subunits, prohibitins) we analyzed the mitochondrial transmembrane potential. The results, displayed on **Figure 5**, showed that PR101 induced a significant increase in the mitochondrial transmembrane potential, similar to the one induced by the positive control butanedione monoxime. The fact that only PR101 induced a detectable change was consistent with the range of mitochondrial proteome alterations induced by this pigment.

**Figure 5:**
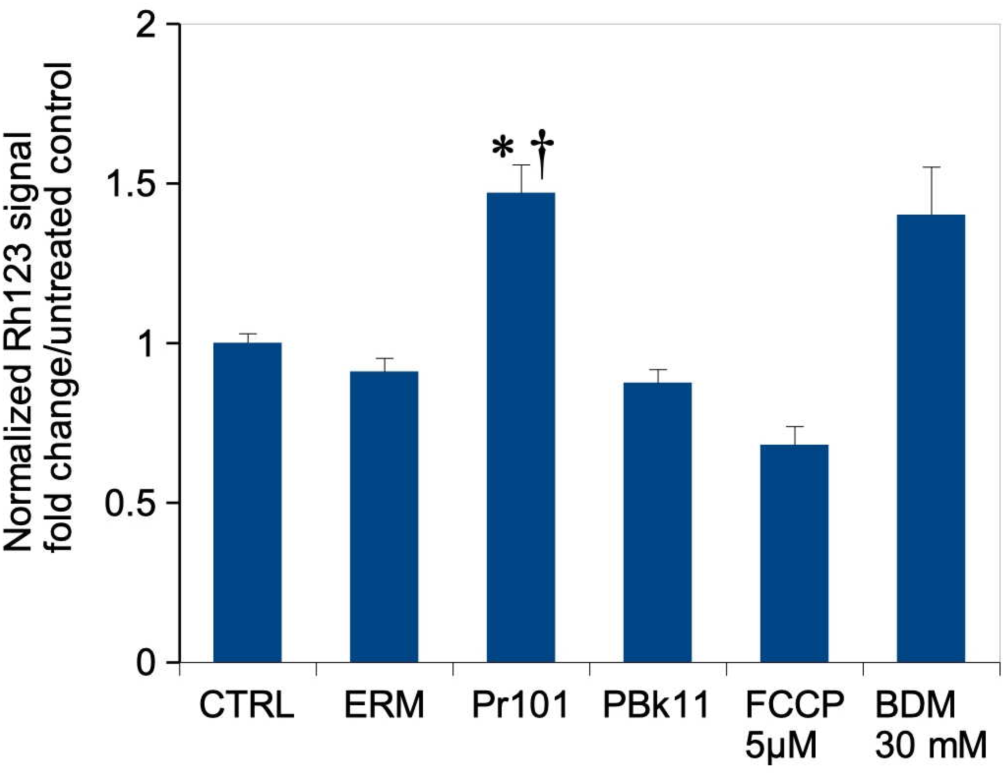
Mitochondrial transmembrane potential The cells were exposed to pigments or control particles for 24 hours, then left to recover for 5 days post exposure without added particles and medium changes every 2 days. Mitochondrial transmembrane potential was then measured via the rhodamine 123 method. All cells were positive for rhodamine 123 internalization in mitochondria, and the mean fluorescence is the displayed parameter. Results are displayed as mean± standard deviation (N=4) CTRL: Unexposed cells ERM: cells exposed to corundum (ERM 066) PR101: cells exposed to Fe_2_O_3_ pigment PBk11: cells exposed to Fe_3_O_4_ pigment FCCP: Cells exposed for 30 minutes to FCCP (induces a decrease in the transmembrane potential) BDM: Cells exposed for 30 minutes to Butanedione monoxime (induces an increase in the transmembrane potential) *: statistically different from the unexposed control (p<0.05, Mann-Whitney U test) †: statistically different from the corundum-exposed cells (p<0.05, Mann-Whitney U test)

### 3.6. Oxidative stress

As an increase in mitochondrial transmembrane potential has been correlated to oxidative stress [45,46], we probed the level of oxidative stress in pigment-exposed cells. The results, displayed in **Figure 6A**, showed a significant increase in the oxidative stress level in PR101-treated cells. The increase in ROS induced by PR101 was not widely different to the effect induced by the minimal menadione concentration used as a positive control, which gives a good appraisal of the magnitude of the effect.

In order to complement this result, we also probed the level of reduced glutathione in cells. This decision was also prompted by the fact that 8 proteins related to glutathione metabolism were found modulated in response to exposure to PR101 (**Table S8**), including the glutathione peroxidase GPx3, glutathione reductase and the regulatory protein GCLM, implicated in the control of glutathione biosynthesis. The results, displayed in **Figure 6B**, showed a moderate but significant increase in the levels of reduced glutathione in PR101 cells, indicating a response against oxidative stress.

### 3.7. Lysosomes

Proteins annotated as related to lysosomes also formed a major cluster in pathway analysis, with 45 proteins present in this cluster **(Table S9)**. The changes detected were mostly decreases (36/45). This indicated a major perturbation in the lysosomal function in PR101-treated cells. As a proxy of lysosomal function, we used the neutral red uptake assay, which measures proton pumping-activity, i.e. the activity of the proton pump in the number of intact lysosomes in the cells. The results of this test, displayed in **Figure 6C**, showed an increase of the lysosomal activity in all particle-treated cells, including the corundum control, but no significant difference between the corundum-treated cells and either of the iron oxide-treated cells. This result was in line with the detailed proteomic results obtained on the proton pump subunits **(Table S9),** as most of them did not change significantly in pigment-treated cells compared to the controls.

**Figure 6:**
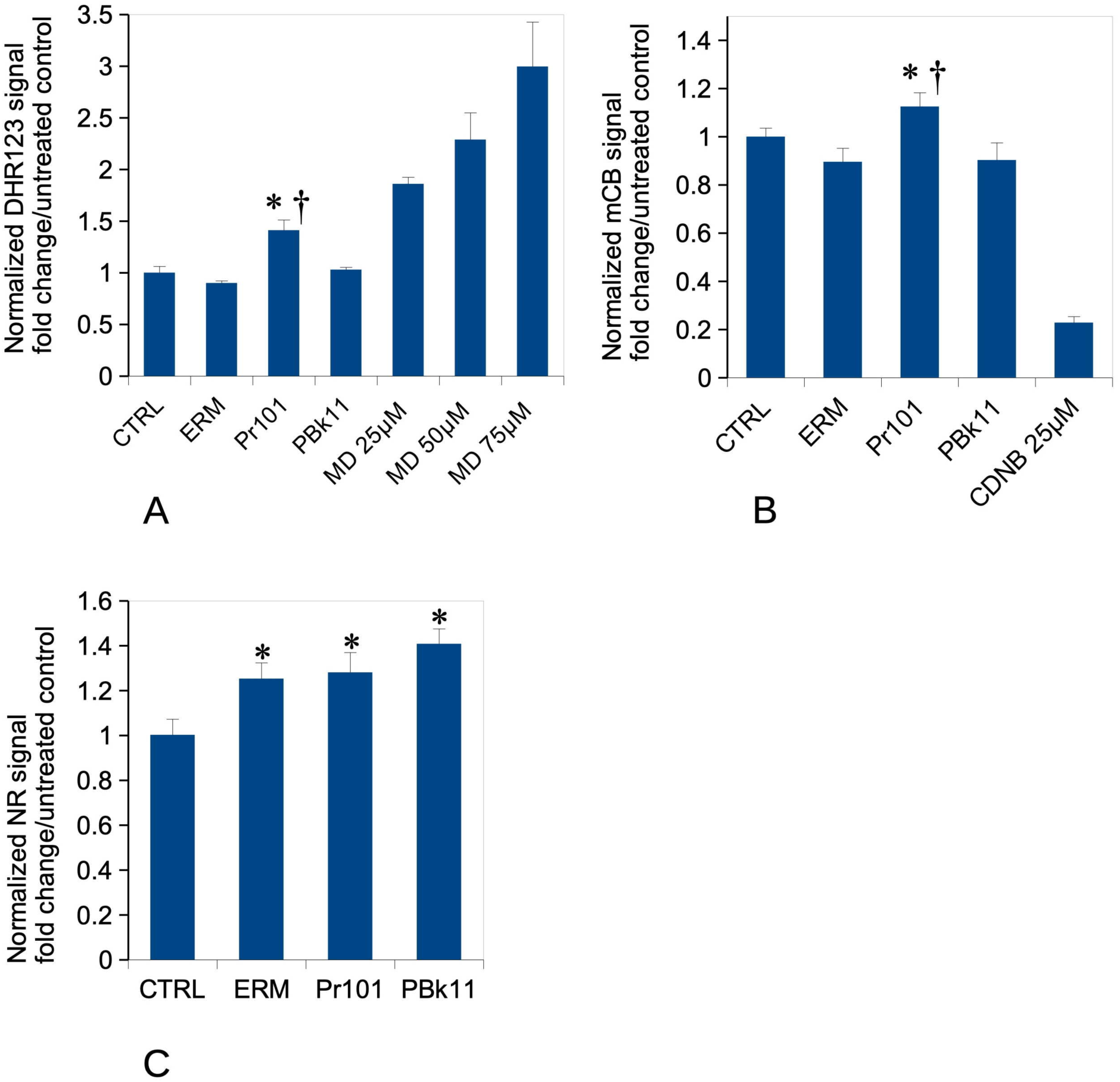
Oxidative stress level, reduced glutathione content, lysosomal activity CTRL: Unexposed cells ERM: cells exposed to corundum (ERM 066) PR101: cells exposed to Fe_2_O_3_ pigment PBk11: cells exposed to Fe_3_O_4_ pigment MD: menadione (added 2 hours before measurement) CDNB: chlorodinitrobenzene (added 30 minutes before measurement) Panel A: oxidative stress level The cells were exposed to pigments or control particles for 24 hours, then left to recover for 5 days post exposure without added particles and medium changes every 2 days. The level of oxidative stress was measured via the DHR123 method. Results are displayed as mean± standard deviation (N=4). *: statistically different from the unexposed control (p<0.05, Mann-Whitney U test) †: statistically different from the corundum-exposed cells (p<0.05, Mann-Whitney U test) Panel B: Reduced glutathione content The cells were exposed to pigments or control particles for 24 hours, then left to recover for 5 days post exposure without added particles and medium changes every 2 days. The level of reduced glutathione was measured via the monochlorobimane conjugation method. Results are displayed as mean± standard deviation (N=4). *: statistically different from the unexposed control (p<0.05, Mann-Whitney U test) †: statistically different from the corundum-exposed cells (p<0.05, Mann-Whitney U test) Panel C : Lysosomal proton pumping (Lysosensor method). The cells were exposed to pigments or control particles for 24 hours, then left to recover for 5 days post exposure without added particles and medium changes every 2 days. All cells were positive for lysosensor internalization in lysosomes, and the mean fluorescence is the displayed parameter. Results are displayed as mean± standard deviation (N=4). *: statistically different from the unexposed control (p<0.05, Mann-Whitney U test) †: statistically different from the corundum-exposed cells (p<0.05, Mann-Whitney U test)

### 3.7. Immune functions

Proteins annotated as related to the innate immune functions also formed an important cluster in pathway analysis, with 55 proteins present in this cluster **(Table S10)**. Of these, 26 were increases and 29 decreases. The cluster could be subdivided into subclusters:

- Proteins implicated in antiviral defences, such as the 2’-5’ oligoadenylate synthases 1A, 2 (P11928 and E9Q9A9, both increased) and 3 (Q8VI93, decreased), DHX15 helicase (B2RRE7, decreased), Endod1 (Q8C522, increased), ISG20 (Q9JL16, decreased), OASL1 (Q8VI94, increased), RIG-I (Q6Q899, decreased), Riok3 (Q9DBU3, increased) Serinc3 (Q9QZI9, increased), TLR7 (P58681, decreased), Tmem43 (Q9DBS1, increased), Trim14 (Q8BVW3, increased), Trim 25 (Q61510, decreased), Trim56 (Q80VI1, decreased), Usp14 (Q9JMA1, decreased).
- Components of the complement pathway such as C1q (P14106, decreased), C3 (P01027, increased) and C4B (P01029, increased)
- Proteins implied the antigen presentation pathway such as ERAP1 (Q9EQH2, decreased), H2D1 (P01900, increased) and H2K1 (P01902, decreased), Ifi30 (Q9ESY9, decreased), Psmb8 (P28063, decreased) .
- Proteins implied directly in pathogen degradation, such as gasdermin (Q9D8T2, decreased) and MPEG1 (A1L3I4, increased)
- Regulators of the inflammatory response, such as Acod1 (P54987, increased), ASC (Q9EPB4, decreased), Bcl10 (Q9Z0H7, decreased), Clec4E (Q9R0Q8, increased), Fcer1G (P20491, increased), GBP1 (Q01514, decreased), Inpp5d (Q9ES52, decreased), Inppl1 (Q6P549, decreased), Lilrb4a (Q64281, increased), Nagk (Q9QZ08, decreased), Slc15A3 (Q8BPX9, increased) TLR2 (Q9QUN7, increased), TLR7 (P58681, decreased), Unc93b1 (Q8VCW4, increased). Due to the role of the cGAS/STING pathway beyond antiviral immunity [47], the modulators of this pathway, such as Endod1, Tmem43, Trim14, Trim25, Trim56 and Usp14 can also be classified in this category.

This long list and the variety of observed modulations between increases and decreases make predictions in the modulation of the macrophage’s immune functions difficult. As a marker of these functions, we tested the release of the two pro-inflammatory cytokines IL6 and TNF alpha under two circumstances. After exposure to the pigments only, to assess the intrinsic pro-inflammatory potential of the iron oxide pigments, or after a stimulation with lipopolysaccharide, mimicking a bacterial challenge, to determine whether pre-exposure to the pigments may alter the potential of macrophages to respond against bacteria. The results, displayed in **Figure 7**, showed that PR101 induced an increase in the secretion of IL6 and TNF even 6 days after the exposure to the pigments (**Figure 7A&B**), demonstrating a pro-inflammatory effect of this pigment, which was not found with PBk11. The results were more contrasted between IL6 and TNF for the response after stimulation with LPS. Both pigments decreased the secretion of IL6 (**Figure 7C**). However, the situation was more complex for TNF, as ERM did induce an increase of TNF secretion after LPS stimulation compared to the particles-free control (**Figure 7D**). Thus, only PBk11 induced a decrease of TNF secretion compared to both controls, while PR101 induced a decrease compared to the ERM control, but not the particles-free control. Nevertheless, the combination of the results on IL6 and TNF suggest that both pigments decrease the response of macrophages to a bacterial challenge.

**Figure 7:**
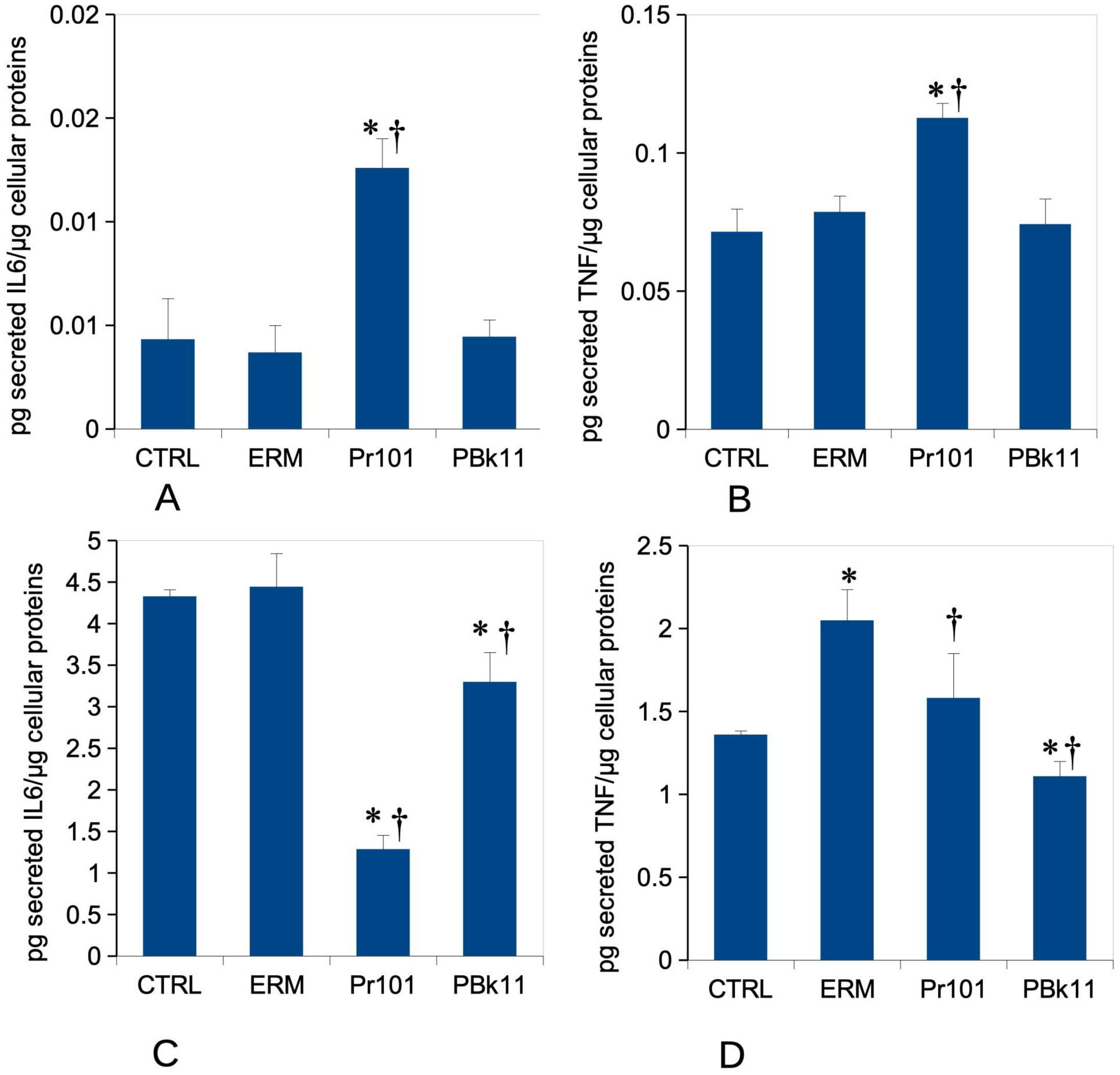
Pro-inflammatory cytokines release The cells were exposed to pigments or control particles for 24 hours, then left to recover for 5 days post exposure without added particles and medium changes every 2 days. The cells were treated (or not) with 50ng/ml lipopolysaccharide in complete cell culture medium for the last 24 hours. The cell medium was then collected for secreted TNF and IL-6 measurements. Results are displayed as mean± standard deviation (N=4). *: statistically different from the unexposed control (p<0.05, Mann-Whitney U test) †: statistically different from the corundum-exposed cells (p<0.05, Mann-Whitney U test) Panel A: IL-6 release without LPS stimulation Panel B: TNF-alpha release without LPS stimulation Panel C: IL-6 release with LPS stimulation (50 ng/ml) Panel D: TNF-alpha release with LPS stimulation (50 ng/ml)

## 4. Discussion

Together with charcoal, iron oxide, in the form of ochre, is one of the most anciently-used pigments by humans, both for paintings and tattoos [48] . More recently in time, black iron oxide (magnetite) has also been found in the tattoos of mummies [49]. However, this ancient use shall not be construed as a marker for innocuity as toxic heavy metals oxides or sulfides such as PbO or HgS have also been found in ancient tattoos [50]. Nevertheless, iron oxide pigments are still widely used today, e.g. in dermopigmentation and in its medical applications, a field where they appear well tolerated [51], which is not the case for other, organic pigments [52–54]. Although iron oxide pigments were not linked to allergic reactions, this does not mean that they are devoid of any side effects, and the purpose of our study was to investigate the effects that iron oxide pigments could induce upon macrophages, i.e. the cells that internalize the pigments in tattoos and maintain them at the tattooing site [17] . In this investigation, we did not intend to study the immediate response of the macrophages, i.e. immediately after treatment with the pigments, which has been well documented in the literature [12–14], but rather the sustained/delayed effects, i.e. the macrophages responses a few days after exposure to the pigments, during the recovery phase post-exposure. This is why we implemented a protocol in which the macrophages were first exposed to the pigments, and then let to recover for 5 days in a pigment-free medium. This in turn posed the question of the dose of pigments that is used, i.e. to figure out what a realistic dose should be. Previous research has determined that pigments are present in tattooed skin at concentrations such as 2mg/cm^2^ [55] or 2 mg/g of skin [24]. For the former publication, which was conducted on human skin, in which the dermis thickness is around 2mm [56], the surface dose translates into a volumetric dermis dose of 10 mg/ml. For the latter publication, the mouse dermis is ca. 250 µm thick [57], for a total skin thickness of ca. 500 µm [58] . This means that the observed concentration translates to a dermal concentration of 4mg/g of dermis, i.e. close to 4 mg/ml. As we decided not to go into toxic doses in order to be able to work on truly live macrophages, we were limited to concentrations close to 1.2 mg/ml, i.e. at the lower end of the real concentrations found in tattoos.

Nevertheless, even at these relatively moderate doses, we found that iron oxide pigments induced adverse effects in macrophages, even a few days after exposure. As the lifespan of dermal macrophages in vivo appears to be relatively short, i.e. a few weeks [17], effects that are detected in vitro close to one week after exposure are rather representative of what happens at the macrophage scale in tattoos. Indeed, we were limited in the time scale that we could investigate post exposure, as pigments-loaded cells showed a strong tendency to detach from the culture plates, which seriously complicated further analysis.

When integrating the proteome results with the pigments dissolution results, a puzzling phenomenon occurred: at equal input doses, the intracellular dissolution of PBk11 liberates more soluble iron than PR101. However, the proteomic and functional effects of PR101 are much more pronounced than those of PBk11. However, it should be noted that the protective protein ferritin is induced to a much stronger extent in response to PBk11 (15-22 fold) than to PR101 (3-9 fold). The fact that the two ferritin chains are not induced to the same extent just reflects the well known heterogeneity of ferritin [59,60] . It is worth noting that the heavy chain, which carries the ferroxidase activity that inactivates iron as a Fenton metal, is more induced in response to both pigments. Thus, the lesser functional effects induced by PBk11 may be explained by a better sequestration of liberated iron from the pigment by a higher amount of ferritin. Conversely, the fact that proteins involved in ferroptosis, such as SLC3A2 (P10852) and ER aminopeptidase 1 (Q9EQH2) are much more induced in response to PR101 also favors the hypothesis that more free cytosolic iron is released by PR101. This hypothesis is further supported by the fact that the lysosomal iron antiporter SLC11A1 (P41251) is much more induced in response to PR101 (close to 4 fold) than in response to PBk11 (less than 1.5 fold). As this antiporter function is to deplete the phagosomes and lysosomes from iron [61] to decrease the viability of internalized pathogen, the landscape suggested by our experiments is that PR101 is, for completely unknown reasons, more perceived as a pathogen than PBk11, which induces an adverse response from the macrophages. This hypothesis is further substantiated by the fact that cytochrome b245 (Q61093), a component of the NADPH oxidase [62] that is activated to destroy pathogens, is induced in response to PR101 (1.8 fold), while its level is constant in response to PBk11.

This adverse response seems to be sensed by the macrophages, as there is a sharp decline in ferric chelate reductase (Q8K385, 0.4 fold change) in response to PR101. As ferric reductases are reducing Fe^3+^ into Fe2^+^ [63], which is required for the transmembrane transport made by SLC11A1, this could be viewed as an attempt to decrease the outward iron ion flow from the lysosomes to the cytosol.

In the same trend, the fact that other proteins that deliver iron or help to deliver iron (e.g.transferrin receptor or polyRC binding protein) are decreased in response to iron oxide particles may be viewed as a way for the cells to decrease the iron fluxes under these conditions of intracellular iron excess.

Nevertheless, there is clearly an excess of soluble iron in pigments-treated cells, as highlighted by the ICP-MS results. From the proteomic results, it is clear that some of this iron accumulates in the mitochondria [64] and perturbs their functioning. As it has been described that iron overload perturbs the organization of mitochondria [65], we checked the expression of mitofusin, mitochondrial fission protein and subunits of the MICOS complex. The antagonist proteins mitofusin (Q811U4) and mitochondrial fission protein (P84817) were decreased in PR101-treated cells, so that a trend in the mitochondrial network could not be inferred from the proteomic results. As the pigments-loaded cells are refractory to optical methods, we could not investigate directly the mitochondrial network. Regarding the MICOS complex, the subunits of the Mic60 subcomplex [66] (i.e. Mic19 (Q9CRB9) Mic25 (Q91VN4) and Mic60 (Q8CAQ8)) were not significantly altered in their levels in response to iron oxide pigments). The only detected subunit of the Mic10 subcomplex, i.e. Mic27 (Q78IK4) was slightly but significantly decreased in PR101 treated cells (0.75 fold).

However, the fact that the iron oxide-treated cells were still completely viable after the recovery period dismisses the hypothesis of ferroptosis. Indeed, the high levels of ferritins observed in the pigments-treated cells are not compatible with the ferritinophagy mechanism observed in macrophage ferroptosis [67] .

Nevertheless, the mitochondria of PR101-treated cells are clearly dysfunctional, as seen from their increased mitochondrial potential. As previously described [45,46], this increase in mitochondrial induces oxidative stress, which we detected. In response to this oxidative stress, an increase of the reduced glutathione levels was detected in PR101-treated cells, consistent with the strong induction (6 fold) of the regulatory subunit of the rate-limiting step of GSH biosynthesis (GCLM, P47791). This mitochondrial production of ROS, driven by an altered functioning of the respiratory chain (which translates to an altered mitochondrial transmembrane potential), has been recently shown to induce a pro-inflammatory phenotype [68]. We thus investigated whether iron oxide pigments, and especially PR101, could induce such a phenotype in macrophages. We detected an increase in the production of IL6 and TNF even after the recovery period, which indicated a low but sustained inflammatory response induced by PR101 alone.

In addition, this pro-inflammatory phenotype may explain in turn the resistance of the iron oxide-treated macrophages to ferroptosis [69].

Beyond the inflammatory response induced by the pigments alone, we also investigated if pigments may alter the inflammatory response to bacteria, which we mimicked by a bacterial challenge. In this case, the iron oxide pigments rather induced a decrease in the efficiency of the inflammatory responses, a phenomenon that we also observed for silver nanoparticles [70], but not for amorphous silica [71].

Taken together, our observations of increased oxidative stress and pro-inflammatory response are highly compatible with the macroscopic observations made on tattooed skin [72]. Inflammation induces a massive infiltration of immune cells, which further increases oxidative stress, which in turn damages dermal proteins, inducing a repair reaction that can be detected macroscopically [72]. In extreme cases, these phenomena can culminate in dermal destruction [73].

In conclusion, iron oxide pigments, although considered as safe, are not without effects on macrophages, and induce sustained oxidative stress and sustained inflammation at least for a fews days after pigment uptake. Taking into account the capture-release-recapture phenomenon that characterizes handling of pigments particles by macrophages [17], this means that such an adverse reaction may reproduce over time at each recapture cycle. Furthermore, it shall be noted that the only particle for which we observed such a lasting inflammatory response in vitro was crystalline silica [30], i.e. a particle that is known to induce extracellular matrix degradation [74] and persistent inflammation [75] . This suggest in turn that some tattoo pigments may be able to produce such phenomena.

However, it is epidemiologically quite clear that tattoos do not induce the same strong adverse reactions than those observed in silicosis. The simple answer to this discrepancy may well lie in the low cell density in the dermis compared to the lung epithelium, and the intrinsic resistance of fibroblasts and macrophages to carcinogenesis, which reduce the probability of strong adverse reactions for tattoos. Nevertheless, the phenomena observed here on macrophages mays explain the rare severe effects that are observed in a time-delayed frame after tattooing (e.g. in [4]). Finally, it would be useful to confirm these in vitro results on in vivo models. Regarding the inflammatory cytokines, the observed increases, although indicative of a low grade inflammation are likely to be way too low to be detectable at the blood level. Regarding the other parameters, such as oxidative stress or mitochondrial potential, their increases may be detectable in vivo, at least in skin, using near infrared probes (e.g. in [76,77]). However, as this requires injection of the probes at the tattoos sites, it is likely that such studies may be ethically limited to animal models.

## Data availability

The proteomic data are available via ProteomeXchange with the identifier PXD064985

The cell biology data are available through the BioStudies database under the identifier S-BSST2099

## Authors contributions

MV, EA, FD, HD, AH, DF: investigation, formal analysis

SR, CC, TR: formal analysis, funding acquisition, project administration

MV, HD, SR, TR: Writing – original draft

TR: conceptualization

All co-authors: Writing – review and editing

