## supplemental Table S1 for "Beyond the ink: cellular and molecular effects of iron-based pigments on macrophages"

Supplementary Table 1: list of proteins detected and quantified by the proteomic screen

| accession | gene_name | description | protein_set_score | coverage | MW | #peptides | abund.CTRL #1 | abund.CTRL #2 | abund.CTRL#3 | abund.CTRL#4 | abund.CTRL#5 | abund.ERM#1 | abund.ERM#2 | abund.ERM#3 | abund.ERM#4 | abund.ERM#5 | abund.PR101#1 | abund.PR101#2 | abund.PR101#3 | abund.PR101#4 | abund.PR101#5 | abund.PBk11#1 | abund.PBk11#2 | abund.PBk11#3 | abund.PBk11#4 | abund.PBk11#5 | U ctrl vs PR101 | U ctrl vs PBk11 | U ERM vs PR101 | U ERM vs PBk11 | Max U PR101 | Max U PBk11 | ratio PR101 Ctrl | ratio PR101 ERM | ratio PBk11 Ctrl | ratio PBK11 ERM |
|---|---|---|---|---|---|---|---|---|---|---|---|---|---|---|---|---|---|---|---|---|---|---|---|---|---|---|---|---|---|---|---|---|---|---|---|---|
| P14046 | A1i3 | Alpha-1-inhibitor 3 | 69.99 | 1.35 | 163773 | 2 | 2 | 2 | 228175.59 | 308055.28 | 267774.09 | 2 | 306388.81 | 476651.59 | 361503.34 | 516319.66 | 804531.81 | 796653.44 | 1528199.62 | 1643296 | 1381671.75 | 2 | 2 | 339412.78 | 583525.38 | 354524.78 | 0 | 8 | 0 | 10 | 0 | 10 | 7.654582133 | 3.705509561 | 1.588871522 | 0.769157416 |
| Q6GQT1 | A2m | Alpha-2-macroglobulin-P | 147.5 | 2.31 | 164353 | 4 | 2402396 | 2334206.5 | 3747582.5 | 2809115 | 4002285.5 | 2903337.75 | 3693804.75 | 4428173 | 5426778.5 | 4046132.25 | 2999354.25 | 7650216.5 | 8354859.5 | 4685579 | 9638795 | 2996478.75 | 2780760.75 | 2867172.5 | 3719682.25 | 2962398 | 2 | 11 | 5 | 4 | 5 | 11 | 2.178981919 | 1.625936012 | 1.002020632 | 0.747698462 |
| P58742 | Aaas | Aladin | 224.85 | 9.16 | 59431 | 4 | 1789176 | 3369475.5 | 3878800.75 | 3795508.5 | 3143381 | 2762686.5 | 2950862 | 3228890.5 | 3289398.75 | 3673388 | 3955719.75 | 4783486 | 5092353 | 3330571.75 | 4749750.5 | 2024188.5 | 562031.56 | 4020494 | 3368866.5 | 3682747.5 | 3 | 11 | 1 | 11 | 3 | 11 | 1.371520549 | 1.377652939 | 0.854909608 | 0.858732109 |
| Q9D2R0 | Aacs | Acetoacetyl-CoA synthetase | 428.19 | 12.95 | 75200 | 9 | 14534950 | 14511162 | 11106037 | 11636410 | 11673466 | 12611744 | 12379490 | 9551637 | 11236144 | 11306804 | 17277032 | 13263274 | 14768595 | 14713797 | 11760832 | 10656766 | 15658030 | 12233195 | 11640896 | 11429758 | 4 | 11 | 2 | 10 | 4 | 11 | 1.13112574 | 1.257466938 | 0.970953023 | 1.079403713 |
| P0C1X8 | Aak1 | AP2-associated protein kinase 1 | 92.63 | 2.91 | 103761 | 2 | 751131.5 | 861356.56 | 1313063.88 | 769912.19 | 925448.12 | 891970.88 | 724858.19 | 1145019.75 | 1303018.5 | 656801.88 | 464786.66 | 528626.75 | 398620.97 | 749964.44 | 2 | 2 | 601887.31 | 962370 | 1107653.88 | 1183237.38 | 0 | 12 | 2 | 10 | 2 | 12 | 0.463545011 | 0.453653301 | 0.83428344 | 0.816480445 |
| Q8BGQ7 | Aars | Alanine--tRNA ligase, cytoplasmic | 2017.73 | 39.77 | 106909 | 30 | 6238024.5 | 8373463 | 6719462.5 | 5620681.5 | 6842751 | 6692189 | 7435547 | 7056463 | 5870239 | 5828923.5 | 14376877 | 14293355 | 10615027 | 10255833 | 10941392 | 8147467.5 | 7186804 | 7815803 | 6506542.5 | 4954248 | 0 | 11 | 0 | 9 | 0 | 11 | 1.789720052 | 1.839303564 | 1.024160302 | 1.052534273 |
| Q3THG9 | Aarsd1 | Alanyl-tRNA editing protein Aarsd1 | 309.71 | 20.15 | 44971 | 6 | 5133612 | 2901265 | 7176543 | 8964312 | 7768755.5 | 5143108 | 4852180 | 7353632 | 7580859.5 | 7955905 | 2229817.5 | 3786541 | 4762729.5 | 4033877.5 | 5324451.5 | 5672113 | 5443880 | 6319979.5 | 8307101 | 7626796.5 | 5 | 12 | 2 | 10 | 5 | 12 | 0.630387856 | 0.612345989 | 1.044620609 | 1.014723291 |
| Q9JKX4 | Aatf | Protein AATF | 177.84 | 14.26 | 59482 | 5 | 2 | 2 | 2 | 2 | 2 | 315934.78 | 2 | 2 | 2 | 2 | 458563.22 | 449100.12 | 320707.81 | 2 | 607283.25 | 481670.5 | 628322.69 | 512804.09 | 589159.5 | 2 | 2.5 | 2.5 | 3 | 3 | 3 | 3 | 183565.64 | 5.810091308 | 221195.878 | 7.001137295 |
| P41233 | Abca1 | ATP-binding cassette sub-family A member 1 | 63.75 | 0.8 | 253912 | 2 | 129360.34 | 686504.06 | 2 | 78050.42 | 453713.12 | 763379.75 | 134580.25 | 87983.62 | 394579.59 | 2 | 812750.25 | 600635.69 | 77764.72 | 2 | 80416.35 | 454717.72 | 204404.94 | 477462.16 | 89663.64 | 2 | 12.5 | 11.5 | 11.5 | 11.5 | 12.5 | 11.5 | 1.166172525 | 1.138384869 | 0.909931149 | 0.888249234 |
| P06795 | Abcb1b | Multidrug resistance protein 1B | 229.5 | 4.86 | 140994 | 3 | 117436.65 | 393292.94 | 415265.25 | 156722.98 | 398207.56 | 2 | 211622.48 | 415003.28 | 508253.75 | 87771.59 | 227969.95 | 554226.69 | 308930.97 | 424152.31 | 2 | 2 | 149662.03 | 266828.78 | 188467.36 | 354663.25 | 11 | 7 | 9.5 | 10.5 | 11 | 10.5 | 1.023199373 | 1.239339204 | 0.647989043 | 0.784869739 |
| Q9DC29 | Abcb6 | ATP-binding cassette sub-family B member 6, mitochondrial | 206.24 | 5.82 | 93770 | 3 | 2 | 232343.08 | 2 | 2 | 2 | 325862.94 | 238962.69 | 259263.44 | 2 | 406558.72 | 2243979.5 | 167298.72 | 2 | 339293.38 | 157919.52 | 26125.08 | 723890.88 | 213296.91 | 2 | 2 | 5 | 8 | 11.5 | 8 | 11.5 | 8 | 12.51766559 | 2.363380016 | 4.145953916 | 0.782770921 |
| Q61102 | Abcb7 | ATP-binding cassette sub-family B member 7, mitochondrial | 387.98 | 11.44 | 82581 | 6 | 4040167.75 | 2327493.5 | 5301460.5 | 4780923 | 5316562 | 3924067.5 | 3975112.75 | 4186394 | 4937932 | 6076769 | 3032426.5 | 4910823.5 | 7157522.5 | 8117256 | 7821633 | 3531760.75 | 3909345 | 6665628 | 6415767.5 | 6165610 | 6 | 8 | 7 | 10 | 7 | 10 | 1.426022065 | 1.343692279 | 1.226103432 | 1.155315725 |
| O35379 | Abcc1 | Multidrug resistance-associated protein 1 | 265.61 | 4.12 | 171185 | 5 | 704874.62 | 584766.38 | 936539.81 | 775012.75 | 1275302.38 | 890507.75 | 1537146.12 | 1418271.88 | 1222654.5 | 733313.62 | 1191033.88 | 744066.19 | 1441774 | 1454719.12 | 820750.12 | 1127077.12 | 1868287.25 | 773213.62 | 1033860.75 | 1324340.75 | 6 | 5 | 12 | 12 | 12 | 12 | 1.321723062 | 0.974223837 | 1.432663465 | 1.055996478 |
| B2RX12 | Abcc3 | Canalicular multispecific organic anion transporter 2 | 332.75 | 6.11 | 169124 | 6 | 2402194.25 | 2003062 | 1745215.62 | 1959752.5 | 2209255.75 | 1794472.12 | 2877274.5 | 1424677.38 | 1889759.88 | 2209339.25 | 2279143.75 | 3273273.5 | 2451957.5 | 2309809 | 2508075.25 | 2627571.25 | 1571461.75 | 2273432.25 | 2405902.75 | 1287580.12 | 2 | 11 | 4 | 12 | 4 | 12 | 1.242529551 | 1.257636203 | 0.985122119 | 0.997099216 |
| D3ZHR2 | Abcd1 | ATP-binding cassette sub-family D member 1 | 252.82 | 9.09 | 81938 | 5 | 743976.5 | 495797.25 | 426802.41 | 542116.38 | 608409.81 | 744396.69 | 592294.06 | 495935.69 | 448847.81 | 763921.38 | 520472.81 | 479638.88 | 1733094.12 | 605360.31 | 522717.88 | 755264 | 1190878 | 772681 | 584396.75 | 777756 | 12 | 2 | 12 | 4 | 12 | 4 | 1.370658045 | 1.267908827 | 1.448643053 | 1.340047812 |
| P55096 | Abcd3 | ATP-binding cassette sub-family D member 3 | 169.16 | 7.28 | 75475 | 3 | 315099.34 | 312355.88 | 424497.69 | 2 | 570307 | 2 | 276493.69 | 574790.44 | 673587.81 | 608229.44 | 1356213.5 | 1148035.38 | 1504144.5 | 1173876.12 | 1367783.25 | 364442.75 | 2 | 978311.38 | 2 | 646735.25 | 0 | 11 | 0 | 12 | 0 | 12 | 4.03760497 | 3.070668216 | 1.226370026 | 0.932675556 |
| O89016 | Abcd4 | ATP-binding cassette sub-family D member 4 | 47.68 | 1.32 | 68583 | 1 | 2 | 2 | 2 | 2 | 2 | 2 | 2 | 2 | 2 | 2 | 2 | 2 | 2 | 2 | 394057.84 | 2 | 182827.59 | 161423.7 | 401253.31 | 2 | 10 | 5 | 10 | 5 | 10 | 5 | 39406.584 | 39406.584 | 74550.86 | 74550.86 |
| P61222 | Abce1 | ATP-binding cassette sub-family E member 1 | 1121.99 | 33.89 | 67314 | 18 | 23447010 | 21605618 | 25485200 | 25220050 | 25898530 | 22405440 | 23385574 | 24610686 | 27128602 | 20725624 | 34438630 | 33029642 | 32882174 | 28954910 | 32443174 | 26514610 | 23552572 | 27166056 | 25107636 | 26451888 | 0 | 6 | 0 | 5 | 0 | 6 | 1.329552078 | 1.367783717 | 1.05865991 | 1.08910197 |
| Q6P542 | Abcf1 | ATP-binding cassette sub-family F member 1 | 222.76 | 6.81 | 94945 | 5 | 5476770.5 | 5479462 | 5195951.5 | 5754986 | 3111324 | 5265897.5 | 6271813 | 6302755.5 | 5246074 | 6349478.5 | 9530251 | 8395944 | 7432936 | 7366759.5 | 8117239.5 | 5481887.5 | 5415308 | 6962911 | 5784079 | 4388544.5 | 0 | 8 | 0 | 11 | 0 | 11 | 1.632517529 | 1.387522229 | 1.120480313 | 0.952327503 |
| Q99LE6 | Abcf2 | ATP-binding cassette sub-family F member 2 | 192.56 | 7.64 | 71782 | 4 | 2303904.5 | 3130276.25 | 3072528.25 | 2674455.25 | 1796052 | 1817489.62 | 1458207.5 | 2525999.25 | 1439237.38 | 2512478 | 3386342.5 | 4271862 | 4888278 | 3023831.5 | 3361912.5 | 2069335.75 | 1333197.25 | 2697136.5 | 2199707.25 | 2128842.25 | 2 | 6 | 0 | 11 | 2 | 11 | 1.458881946 | 1.941087589 | 0.803579042 | 1.06918679 |
| Q64343 | Abcg1 | ATP-binding cassette sub-family G member 1 | 334.09 | 11.56 | 74033 | 6 | 4299607 | 3507407 | 852908.5 | 2239634.25 | 2566109.75 | 4237484.5 | 4902044 | 2974458 | 3308147.5 | 2914781.25 | 2305221 | 3891151.5 | 2079851.5 | 749966.5 | 1711617.75 | 3744562.5 | 1136810.38 | 2509132.25 | 1360180.75 | 2710481.5 | 8 | 11 | 3 | 3 | 8 | 11 | 0.797421223 | 0.585584222 | 0.85114 | 0.625032467 |
| Q7TMS5 | Abcg2 | ATP-binding cassette sub-family G member 2 | 51.04 | 1.52 | 72978 | 1 | 2 | 2 | 2 | 2 | 2 | 2 | 2 | 111994.7 | 2 | 2 | 2 | 377397.78 | 505850.22 | 587229.06 | 586873.25 | 2 | 2 | 2 | 2 | 2 | 2.5 | 12.5 | 3 | 10 | 3 | 12.5 | 205735.231 | 18.36877423 | 1 | 8.928356E-05 |
| Q6AYT7 | Abhd12 | Monoacylglycerol lipase ABHD12 | 637.13 | 35.68 | 45296 | 12 | 4912159 | 5216900.5 | 7965349 | 5556858 | 8213546.5 | 5620483.5 | 5551305 | 9515242 | 7656511 | 7186580.5 | 9928425 | 10001447 | 13420353 | 11155857 | 13529037 | 3605408.25 | 3115539 | 7362916 | 5457305 | 8016842 | 0 | 9 | 0 | 7 | 0 | 9 | 1.821291686 | 1.633406128 | 0.864841424 | 0.775623857 |
| Q6DGG1 | Abhd14b | Protein ABHD14B | 44.28 | 5.24 | 22618 | 1 | 2217031.75 | 2030096.88 | 1218257 | 1802718.75 | 1493621 | 1821519.88 | 1845627.38 | 2 | 1195866.62 | 1863028 | 1008347.31 | 767205.19 | 815692.56 | 737948.44 | 2 | 2167975.25 | 1846568.25 | 1587670.25 | 1439437.62 | 1298375.62 | 0 | 11 | 4.5 | 10 | 4.5 | 11 | 0.379970309 | 0.494970827 | 0.951870394 | 1.239960241 |
| Q6MG55 | Abhd16a | Protein ABHD16A | 49.67 | 2.15 | 63038 | 1 | 572714.56 | 499482.91 | 443458.06 | 558502.12 | 493130.69 | 469616.66 | 461588.75 | 1745414.75 | 2 | 2 | 731815.75 | 459615.69 | 462966.94 | 382573 | 341916.81 | 2 | 311667.41 | 2 | 519829.12 | 457564.47 | 7 | 4 | 12 | 10 | 12 | 10 | 0.926615119 | 0.888764372 | 0.5021115 | 0.481601048 |
| B5DFK7 | Abhd17c | Alpha/beta hydrolase domain-containing protein 17C | 41.87 | 3.44 | 35098 | 1 | 751338.75 | 938515.5 | 540047 | 884215.81 | 793029.94 | 902714.5 | 994029.94 | 897280.12 | 1081535.12 | 2 | 692551.5 | 642242.06 | 948397.25 | 811414.19 | 915764.56 | 370905.81 | 660935.12 | 955076.38 | 1220424.5 | 648983 | 11 | 12 | 9 | 11 | 11 | 12 | 1.026418909 | 1.034784088 | 0.986992506 | 0.995036366 |
| Q8CBW3 | Abi1 | Abl interactor 1 | 184.31 | 9.15 | 52288 | 4 | 4543707.5 | 5116722.5 | 4458697.5 | 2476582.5 | 7211364 | 2204648.75 | 5048914 | 3960553.5 | 5673222 | 2357758 | 623636.25 | 3057447 | 1031224.56 | 357051.44 | 1646344.62 | 4977649.5 | 1997275.25 | 2138443.75 | 1779962.5 | 2423521.5 | 1 | 3 | 2 | 5 | 2 | 5 | 0.282088587 | 0.348956627 | 0.559365359 | 0.691960816 |
| Q5SSL4 | Abr | Active breakpoint cluster region-related protein | 228.57 | 7.45 | 97667 | 5 | 3511817 | 3506653.5 | 2263343.5 | 2084539.75 | 3661150.75 | 2751773.25 | 3672347.25 | 2426181.5 | 3550312 | 2213943.5 | 1298322.5 | 1708274.38 | 951102.44 | 1425104.38 | 1746725.12 | 4205417 | 1823019.62 | 2151851.25 | 2548934 | 2576422 | 0 | 10 | 0 | 9 | 0 | 10 | 0.474431987 | 0.487837474 | 0.88541939 | 0.910437683 |
| Q4KML4 | Abracl | Costars family protein ABRACL | 253.06 | 46.91 | 9030 | 4 | 10122667 | 7522881.5 | 8765203 | 5566618 | 9997868 | 7122345 | 8183080 | 6210514.5 | 7408654 | 6162303 | 3244150 | 3238398.75 | 2356760.75 | 1314134.38 | 2500888 | 7758814 | 5931499.5 | 4284298.5 | 3279311 | 4009400 | 0 | 3 | 0 | 4 | 0 | 4 | 0.301471359 | 0.360656916 | 0.601862539 | 0.72002159 |
| Q3TCJ1 | Abraxas2 | BRISC complex subunit Abraxas 2 | 133.38 | 10.12 | 46943 | 3 | 1530408.25 | 1801312.88 | 2627071 | 2602985.5 | 2853623 | 1197829.12 | 2793171.25 | 2274369.5 | 2311887.5 | 2813128.75 | 1635825 | 1991474.88 | 2476134.25 | 2548495.5 | 2580041.25 | 1853363.25 | 1449047 | 2847105.25 | 3188933.25 | 2218276.75 | 9 | 12 | 11 | 12 | 11 | 12 | 0.983931379 | 0.986092198 | 1.012380194 | 1.014603489 |
| Q921H8 | Acaa1a | 3-ketoacyl-CoA thiolase A, peroxisomal | 1249.45 | 57.31 | 43953 | 17 | 52328034 | 46276826 | 31480616 | 32183094 | 32588936 | 47164458 | 46298130 | 21213058 | 22612490 | 29779798 | 38773466 | 35738034 | 31826420 | 30991394 | 31499924 | 46707338 | 37898378 | 33161128 | 28792982 | 30769972 | 8 | 10 | 10 | 10 | 10 | 10 | 0.866424094 | 1.010542442 | 0.910048587 | 1.061423301 |
| Q8BWT1 | Acaa2 | 3-ketoacyl-CoA thiolase, mitochondrial | 270.64 | 23.93 | 41830 | 5 | 2378756.75 | 1859129 | 3059032.5 | 3265991.25 | 2440298 | 1809124.75 | 1551628.5 | 1755223.75 | 3125772.5 | 2268107 | 1847809.25 | 1249169 | 1955735.25 | 1965275.12 | 1639066.75 | 2197727 | 2 | 2161894.5 | 2323760 | 2073637.75 | 2 | 4 | 10 | 12 | 10 | 12 | 0.665763072 | 0.823708237 | 0.673450858 | 0.833219868 |
| P11497 | Acaca | Acetyl-CoA carboxylase 1 | 960.71 | 8.61 | 265194 | 17 | 11324967 | 10542662 | 8519150 | 10532117 | 13300456 | 12657131 | 11578942 | 9833489 | 11738617 | 9746258 | 8657329 | 8002260.5 | 7694931 | 7734428.5 | 7352655 | 12502676 | 12914949 | 8288740 | 9242459 | 9698856 | 1 | 10 | 0 | 9 | 1 | 10 | 0.727445138 | 0.709963166 | 0.971012711 | 0.947677321 |
| Q9D7B6 | Acad8 | Isobutyryl-CoA dehydrogenase, mitochondrial | 41.86 | 3.39 | 45020 | 1 | 340783.34 | 2 | 591089 | 2 | 2 | 495986.28 | 471410.5 | 2 | 447214.91 | 2 | 2 | 2 | 870876.81 | 743963.69 | 604290.31 | 351943.25 | 2 | 2 | 619185.19 | 2 | 7 | 11.5 | 8 | 10 | 8 | 11.5 | 2.381356787 | 1.568719212 | 1.042125778 | 0.686500543 |
| Q8JZN5 | Acad9 | Complex I assembly factor ACAD9, mitochondrial, | 833.17 | 24.8 | 68722 | 14 | 19367828 | 20783634 | 21649060 | 21483136 | 23708906 | 19850636 | 18651862 | 21238116 | 19475986 | 19100152 | 13395500 | 14618983 | 18045232 | 17511716 | 12765865 | 19926202 | 19981610 | 22734662 | 21098514 | 22717526 | 0 | 12 | 0 | 3 | 0 | 12 | 0.713482257 | 0.776442411 | 0.995008532 | 1.082811544 |
| P51174 | Acadl | Long-chain specific acyl-CoA dehydrogenase, mitochondrial | 1149.78 | 41.86 | 47908 | 18 | 21448768 | 23198336 | 31826866 | 39619526 | 40684934 | 20548086 | 23637334 | 38322274 | 41865094 | 36028206 | 12230342 | 17170336 | 22138502 | 24663898 | 26224310 | 21575894 | 21966078 | 30366708 | 33423068 | 33989606 | 5 | 10 | 5 | 8 | 5 | 10 | 0.653325767 | 0.638570781 | 0.901408147 | 0.881050363 |
| P15650 | Acadl | Long-chain specific acyl-CoA dehydrogenase, mitochondrial | 816.56 | 32.09 | 47873 | 13 | 5464190 | 5253466 | 7947618 | 9395617 | 8140753.5 | 9116898 | 6204901 | 9523621 | 10441448 | 12066101 | 5483321.5 | 4358276 | 11660529 | 10238920 | 9299806 | 6303859 | 6916296 | 10734567 | 10049325 | 10658635 | 9 | 6 | 9 | 12 | 9 | 12 | 1.133673706 | 0.866700724 | 1.2337197 | 0.943186519 |
| P45952 | Acadm | Medium-chain specific acyl-CoA dehydrogenase, mitochondrial | 471.07 | 28.27 | 46481 | 10 | 16555526 | 21097706 | 20319710 | 26387190 | 24250584 | 23797926 | 19384488 | 20730774 | 23363098 | 26513874 | 8270995.5 | 12266707 | 15937865 | 18617272 | 20106558 | 21444492 | 22173790 | 26162712 | 23046294 | 24281606 | 2 | 8 | 1 | 11 | 2 | 11 | 0.692375488 | 0.660860284 | 1.078244379 | 1.029165387 |
| P15651 | Acads | Short-chain specific acyl-CoA dehydrogenase, mitochondrial | 222.03 | 12.86 | 44765 | 4 | 3724981.5 | 3861887 | 2920398.5 | 3786592.5 | 2828520 | 3485947 | 3700279 | 2853695.25 | 2362879.75 | 3231411.75 | 1218151.62 | 2735917 | 1749908.75 | 2826879.75 | 2463983 | 2674835 | 4282079 | 4086214.75 | 4116522.5 | 3641011 | 0 | 8 | 3 | 5 | 3 | 8 | 0.64213272 | 0.703255117 | 1.098016911 | 1.202533351 |
| Q9DBL1 | Acadsb | Short/branched chain specific acyl-CoA dehydrogenase, mitochondrial | 323 | 18.98 | 47874 | 6 | 8399483 | 7976039 | 9273280 | 10739570 | 8866604 | 4318321 | 8085269 | 7542354 | 8587302 | 9382110 | 5618508.5 | 6279296 | 9182612 | 8266776 | 9863688 | 7793067 | 7940559.5 | 10873802 | 10671320 | 10703206 | 8 | 12 | 11 | 6 | 11 | 12 | 0.866443516 | 1.03416886 | 1.060258092 | 1.265501885 |
| P50544 | Acadvl | Very long-chain specific acyl-CoA dehydrogenase, mitochondrial | 477.63 | 20.43 | 70875 | 7 | 2834727.5 | 2884674.25 | 5736382 | 2137595 | 5652236.5 | 2921937 | 4210110 | 3652970 | 4268540 | 4001351.5 | 1612268.75 | 2165679.25 | 2622099.25 | 1751690.5 | 2637655 | 2552521.5 | 837234.56 | 2573010.5 | 1780154.25 | 2339971.5 | 3 | 3 | 0 | 0 | 3 | 3 | 0.560615632 | 0.566226427 | 0.523905948 | 0.529149343 |
| Q8QZT1 | Acat1 | Acetyl-CoA acetyltransferase, mitochondrial | 850.68 | 43.16 | 44816 | 13 | 26204268 | 27220776 | 37970174 | 36559342 | 30341418 | 32688850 | 28288316 | 32972544 | 34100694 | 35013850 | 16894826 | 17765628 | 24847262 | 28959072 | 24038362 | 25668278 | 25894530 | 36957378 | 31649110 | 32498500 | 2 | 10 | 1 | 7 | 2 | 10 | 0.710726523 | 0.689943671 | 0.964445199 | 0.936243182 |
| Q8CAY6 | Acat2 | Acetyl-CoA acetyltransferase, cytosolic | 365.41 | 24.94 | 41298 | 6 | 14564278 | 17322250 | 13904134 | 16146889 | 13305429 | 18757684 | 13281943 | 13431818 | 12677805 | 12988482 | 11740502 | 11273730 | 12236470 | 13253706 | 11692324 | 17763148 | 17505086 | 19557784 | 16864940 | 20933670 | 0 | 1 | 2 | 3 | 2 | 3 | 0.800031206 | 0.846199764 | 1.231006906 | 1.302046402 |
| Q8BMP6 | Acbd3 | Golgi resident protein GCP60 | 316.6 | 12.76 | 60181 | 5 | 3348883.5 | 3237224 | 4394974 | 3736755 | 3997881.5 | 2790568.5 | 3993099 | 4414948.5 | 4488731 | 5584069 | 3492706 | 4827799 | 5059617 | 3305508 | 4772623.5 | 3661755 | 4569853 | 4725572 | 3234970.5 | 5917319.5 | 7 | 8 | 11 | 10 | 11 | 10 | 1.146536483 | 1.008783501 | 1.181331649 | 1.039398129 |
| Q9JIX8 | Acin1 | Apoptotic chromatin condensation inducer in the nucleus | 625.8 | 8.82 | 150719 | 13 | 5637156.5 | 4327341 | 5628949 | 5763103 | 6790344.5 | 5614522.5 | 7231470.5 | 7198500.5 | 6137494 | 5002033 | 3362996.25 | 2986162.75 | 2702165.5 | 3615508 | 3506767.5 | 3966059.75 | 4420869 | 3939507.25 | 4204008 | 4560664.5 | 0 | 2 | 0 | 0 | 0 | 2 | 0.574614023 | 0.518650249 | 0.749322767 | 0.676343466 |
| Q91V92 | Acly | ATP-citrate synthase | 2875.58 | 45.46 | 119728 | 43 | 4895433 | 5268456 | 4241133 | 3473709.5 | 4340311.5 | 4205352.5 | 4933314 | 3662162.5 | 727313.44 | 3532095 | 3798262.25 | 5940838 | 4698195 | 3481527 | 5223822 | 5303044.5 | 3081587.5 | 3467811.25 | 3418454 | 4066835 | 11 | 6 | 7 | 11 | 11 | 11 | 1.041568003 | 1.356525332 | 0.870322464 | 1.133497252 |
| P28271 | Aco1 | Cytoplasmic aconitate hydratase | 338.65 | 8.89 | 98126 | 6 | 7391412 | 6075050 | 6731787 | 7264341 | 6245328.5 | 6967235.5 | 6823889.5 | 5705470.5 | 6380406 | 6895429.5 | 6186463.5 | 6265698.5 | 5551886 | 4792575.5 | 5669717 | 6435557.5 | 11833010 | 6398967 | 9051768 | 5834373 | 3 | 11 | 2 | 10 | 3 | 11 | 0.844500099 | 0.868606314 | 1.17342385 | 1.206919179 |
| Q99KI0 | Aco2 | Aconitate hydratase, mitochondrial | 1836.53 | 38.72 | 85464 | 28 | 98937026 | 108239586 | 112348922 | 117157938 | 110199562 | 94385986 | 102614714 | 117282754 | 108750690 | 104880218 | 78249626 | 88412626 | 101216410 | 100555194 | 111738634 | 121943090 | 108396722 | 133252026 | 131355874 | 143176754 | 5 | 3 | 6 | 2 | 6 | 3 | 0.878016797 | 0.90956512 | 1.166839025 | 1.208765118 |
| P54987 | Acod1 | Cis-aconitate decarboxylase | 942.6 | 46.72 | 53758 | 17 | 3044199 | 4004318.5 | 915954.06 | 3716687.5 | 2212149.5 | 6331263 | 4484360 | 3538024.25 | 3336212.75 | 3481112 | 20002776 | 19841840 | 22009238 | 14388771 | 22568140 | 4476999 | 5382819 | 6600090.5 | 5672641 | 7470627 | 0 | 0 | 0 | 4 | 0 | 4 | 7.11211189 | 4.667275787 | 2.130750668 | 1.398290853 |
| Q9CQR4 | Acot13 | Acyl-coenzyme A thioesterase 13 | 158.32 | 30.71 | 15183 | 3 | 1911034.88 | 1444346.75 | 289815.31 | 626404.56 | 740460.94 | 1674202.12 | 1779092.25 | 577663.88 | 535586.62 | 1034561.12 | 1686226.62 | 526680.38 | 2218970 | 2427570.75 | 1516084.88 | 1959046.75 | 1934234.88 | 1909353.5 | 1498687.12 | 388586.62 | 6 | 6 | 8 | 7 | 8 | 7 | 1.671075077 | 1.495335501 | 1.534280341 | 1.372926862 |
| Q9QYR9 | Acot2 | Acyl-coenzyme A thioesterase 2, mitochondrial | 369.48 | 22.3 | 49657 | 8 | 7293702 | 8286939 | 6672335 | 9951024 | 6500513 | 9967002 | 6756689 | 6678616 | 7720353.5 | 8637121 | 12033325 | 9562777 | 10658222 | 13313819 | 11936642 | 8458113 | 6291363 | 9791124 | 9576298 | 6315352.5 | 1 | 12 | 1 | 11 | 1 | 12 | 1.485738498 | 1.446305358 | 1.044639174 | 1.016913297 |
| Q91V12 | Acot7 | Cytosolic acyl coenzyme A thioester hydrolase | 233.67 | 14.17 | 42537 | 4 | 4301141 | 4548482.5 | 2 | 4211812.5 | 3307630.75 | 4399415 | 4454586 | 3768898 | 3426568 | 2 | 3596047.75 | 3981588 | 3724722.5 | 3431810 | 610878.81 | 5403781 | 4566798.5 | 3488144.75 | 3769611.75 | 3674031.25 | 9 | 9 | 10 | 8 | 10 | 9 | 0.937441665 | 0.95610933 | 1.276942969 | 1.302371265 |
| P58137 | Acot8 | Acyl-coenzyme A thioesterase 8 | 48.21 | 5 | 35827 | 1 | 2 | 370295.44 | 2 | 2 | 2 | 418332.53 | 2 | 2 | 2 | 2 | 2 | 524036.28 | 426576.28 | 2 | 360866.5 | 2 | 2 | 2 | 2 | 281541.53 | 7 | 12 | 7 | 12 | 7 | 12 | 3.54164428 | 3.134965336 | 0.76032113 | 0.673015187 |
| Q9R0X4 | Acot9 | Acyl-coenzyme A thioesterase 9, mitochondrial | 252.71 | 15.95 | 50560 | 6 | 4559140 | 3689021.5 | 2973731.75 | 4642006.5 | 3520807.75 | 5459893 | 4664878.5 | 3926136.5 | 3624524.5 | 4710738 | 5353930 | 4667128.5 | 4899597.5 | 4775519.5 | 4819728 | 3709617.75 | 4607826.5 | 3832383.75 | 4768726 | 4824469.5 | 0 | 5 | 6 | 12 | 6 | 12 | 1.264703298 | 1.095136102 | 1.121658581 | 0.971270343 |
| Q9EPL9 | Acox3 | Peroxisomal acyl-coenzyme A oxidase 3 | 477.35 | 14.57 | 78404 | 8 | 7589337 | 6118906.5 | 3891163.5 | 5622688 | 3634531 | 8223612 | 6255918 | 3190058.25 | 3613107.5 | 4195013.5 | 4689528 | 2775675.5 | 4056003 | 4372478 | 4132876 | 6584745 | 9984524 | 5463659.5 | 7275136 | 2331101 | 8 | 10 | 10 | 9 | 10 | 10 | 0.745684156 | 0.78604243 | 1.178076706 | 1.241837137 |
| Q9D358 | Acp1 | Low molecular weight phosphotyrosine protein phosphatase | 340.25 | 53.8 | 18192 | 5 | 17341212 | 18472974 | 15462322 | 19239262 | 12596592 | 16087477 | 14781451 | 13625287 | 13923661 | 14276070 | 9311773 | 11091374 | 10357466 | 10997143 | 9846320 | 17781652 | 21960460 | 15401542 | 17242272 | 16758012 | 0 | 12 | 0 | 1 | 0 | 12 | 0.620895313 | 0.709881343 | 1.072571346 | 1.226291086 |
| P20611 | Acp2 | Lysosomal acid phosphatase | 78.11 | 2.84 | 48320 | 1 | 1386162 | 1782998.5 | 1162682.38 | 1577526 | 1463096.75 | 1179255.88 | 1302892.12 | 904782.88 | 1425735.88 | 1263860.75 | 551955.5 | 1241376.12 | 1535638.88 | 378118.44 | 1018481.06 | 1611354 | 430187.44 | 1112411 | 1046106.19 | 1200716.75 | 4 | 5 | 8 | 9 | 8 | 9 | 0.640975521 | 0.777676065 | 0.732560266 | 0.888793044 |
| Q8BP40 | Acp6 | Lysophosphatidic acid phosphatase type 6 | 125.46 | 6.94 | 47625 | 2 | 2502912.25 | 2453002 | 2520164.5 | 3186463.75 | 2788453 | 2630213.75 | 2728787.5 | 2865017.25 | 3316034 | 318494.75 | 831219.5 | 2339461.5 | 1192181.88 | 331496.28 | 2893912 | 2975778.5 | 1411069.25 | 3328713 | 3542392.5 | 3300818 | 4 | 6 | 8 | 6 | 8 | 6 | 0.564141974 | 0.639898885 | 1.082356414 | 1.227702765 |
| Q8VCW8 | Acsf2 | Acyl-CoA synthetase family member 2, mitochondrial | 262.22 | 10.41 | 67951 | 5 | 1444531.62 | 1620326 | 2791607 | 2291989 | 5187222 | 1755911.25 | 1880751.12 | 2217638.75 | 3334078.75 | 1788696 | 1290026.25 | 995202.25 | 1109158.5 | 2140125 | 1490058.62 | 996190.75 | 2826621 | 2076567.75 | 1559203.62 | 1556829.62 | 3 | 8 | 3 | 7 | 3 | 8 | 0.526750261 | 0.639930953 | 0.676037195 | 0.821294564 |
| P41216 | Acsl1 | Long-chain-fatty-acid--CoA ligase 1 | 1390.77 | 37.48 | 77951 | 25 | 28522254 | 28125574 | 24715036 | 36527822 | 41999550 | 32389196 | 31803704 | 29200890 | 38275862 | 30569168 | 21980552 | 19971534 | 33236584 | 29047424 | 33988150 | 27057322 | 29331676 | 34245782 | 35493622 | 40999410 | 9 | 11 | 8 | 11 | 9 | 11 | 0.86449459 | 0.851980087 | 1.045265904 | 1.030134539 |
| Q9CZW4 | Acsl3 | Long-chain-fatty-acid--CoA ligase 3 | 1163.46 | 37.08 | 80492 | 21 | 22665738 | 26771378 | 19117168 | 23176984 | 22844330 | 25316566 | 28297000 | 19276960 | 19943004 | 20877052 | 29094542 | 24873816 | 31847766 | 30600920 | 30825496 | 24952532 | 26800570 | 26447298 | 25128348 | 24583282 | 1 | 4 | 2 | 8 | 2 | 8 | 1.285112559 | 1.294888632 | 1.116398537 | 1.124891173 |
| Q9QUJ7 | Acsl4 | Long-chain-fatty-acid--CoA ligase 4 | 1694.28 | 49.79 | 79077 | 31 | 72391778 | 76940946 | 52452486 | 57201658 | 55650418 | 74602466 | 71282586 | 59552758 | 57176118 | 61749586 | 89543818 | 91066066 | 91863090 | 78868394 | 93616482 | 75037770 | 69010842 | 66721334 | 63455134 | 69597050 | 0 | 9 | 0 | 8 | 0 | 9 | 1.414193008 | 1.371787611 | 1.092757106 | 1.05999015 |
| Q8JZR0 | Acsl5 | Long-chain-fatty-acid--CoA ligase 5 | 779.55 | 25.04 | 76206 | 14 | 9897010 | 9563186 | 9765022 | 10934424 | 9152895 | 11918206 | 9156390 | 10322725 | 8545862 | 9701906 | 9097059 | 10575746 | 13848971 | 14339693 | 12183843 | 11119385 | 12480043 | 11224152 | 10461413 | 10584599 | 6 | 2 | 5 | 4 | 6 | 4 | 1.217647999 | 1.209491477 | 1.13296933 | 1.125380035 |
| Q9QXG4 | Acss2 | Acetyl-coenzyme A synthetase, cytoplasmic | 320.05 | 8.99 | 78862 | 6 | 12206158 | 12202398 | 9685869 | 13284103 | 13257877 | 12834568 | 12886325 | 11590474 | 12016646 | 11995372 | 7307326.5 | 7458311 | 8089299 | 8152918 | 9004635 | 15379777 | 12087815 | 12756392 | 12197419 | 12498191 | 0 | 12 | 0 | 8 | 0 | 12 | 0.659875689 | 0.65248338 | 1.070637252 | 1.058643354 |
| P60710 | Actb | Actin, cytoplasmic 1 | 4410.089 | 93.33 | 41737 | 59 | 1903567.75 | 3456343 | 1590567.5 | 2177505.25 | 2814027.25 | 1746062.88 | 2369415 | 1691173.88 | 3222573.5 | 2456823.75 | 4577360 | 5946585 | 4084039.5 | 2183052.5 | 3250668.75 | 912575.69 | 878834.81 | 1134181.62 | 966919.25 | 969713.25 | 3 | 0 | 3 | 0 | 3 | 0 | 1.678252195 | 1.74487378 | 0.407152926 | 0.423315678 |
| Q8BFZ3 | Actbl2 | Beta-actin-like protein 2 | 1246.72 | 35.9 | 42004 | 20 | 2 | 2 | 808316.38 | 906189.62 | 876803.19 | 2 | 2 | 606012.25 | 2 | 506933.25 | 2 | 2 | 2 | 2 | 2 | 2 | 2 | 639027.62 | 619481.44 | 904653.31 | 5 | 10 | 7.5 | 7 | 7.5 | 10 | 3.859047E-06 | 8.985118E-06 | 0.83477612 | 1.943630401 |
| P68033 | Actc1 | Actin, alpha cardiac muscle 1 | 2006.44 | 50.66 | 42019 | 32 | 2 | 2 | 1376759.12 | 849330.44 | 1193910.12 | 2 | 847272 | 13039763 | 2 | 460640.56 | 2 | 723262.31 | 2 | 808958.31 | 858784.94 | 1230617.75 | 2 | 922923.25 | 1002348.5 | 1019162.56 | 9 | 11 | 12 | 8 | 12 | 11 | 0.69912485 | 0.166647823 | 1.220774727 | 0.290991588 |
| P63259 | Actg1 | Actin, cytoplasmic 2 | 4416.749 | 93.33 | 41793 | 59 | 3193824 | 4651257 | 2134905.75 | 2053409.38 | 1554199.38 | 3416460 | 4374765.5 | 2078096.12 | 2568780.75 | 2515026 | 4711988.5 | 4185462.25 | 2221636.5 | 1383974.88 | 2108570.75 | 1735729.88 | 1731756.5 | 1226768.5 | 946602 | 960163.19 | 11 | 2 | 11 | 0 | 11 | 2 | 1.075365606 | 0.977162271 | 0.48581223 | 0.441447429 |
| Q9Z2N8 | Actl6a | Actin-like protein 6A | 580.95 | 29.6 | 47448 | 9 | 10441830 | 9525989 | 10358539 | 11031376 | 10012406 | 10191119 | 9333650 | 8102491 | 7945215 | 10107036 | 9107309 | 9559680 | 9182002 | 6160120.5 | 7102590 | 9089944 | 11636330 | 9325515 | 8137057.5 | 7941510 | 1 | 5 | 7 | 11 | 7 | 11 | 0.800303474 | 0.9000031 | 0.897999431 | 1.009869753 |
| Q7TPR4 | Actn1 | Alpha-actinin-1 | 1445.12 | 29.48 | 103068 | 24 | 16411922 | 15038022 | 9307904 | 10521349 | 10581996 | 11564938 | 13728450 | 9574411 | 9593676 | 7617270.5 | 4884120.5 | 8370109 | 4183680.5 | 3218683 | 4789386 | 14157026 | 9845708 | 5638205 | 10378459 | 9918144 | 0 | 6 | 1 | 11 | 1 | 11 | 0.411339933 | 0.488605836 | 0.807251519 | 0.958885271 |
| P57780 | Actn4 | Alpha-actinin-4 | 2514.16 | 47.15 | 104977 | 37 | 59461186 | 64161754 | 64462126 | 57605290 | 62320282 | 45613078 | 57856514 | 67049682 | 60539498 | 54634886 | 27095216 | 40868066 | 27574836 | 29738726 | 34009346 | 57831258 | 46718858 | 50458038 | 46790306 | 54159386 | 0 | 1 | 0 | 6 | 0 | 6 | 0.517145093 | 0.557541918 | 0.831003266 | 0.895917144 |
| Q9QZB7 | Actr10 | Actin-related protein 10 | 58.45 | 3.12 | 46208 | 1 | 1046604.06 | 933360.69 | 863840.81 | 2 | 719723.69 | 1173531 | 860884.44 | 842592.94 | 723040 | 897445.56 | 357181.41 | 547728.81 | 630646.06 | 918743 | 606898.19 | 1023885.56 | 952655.38 | 1002122.81 | 809299 | 2 | 7 | 10.5 | 4 | 12 | 7 | 12 | 0.859034832 | 0.680645157 | 1.062980646 | 0.842238989 |
| P61164 | Actr1a | Alpha-centractin | 602.51 | 37.23 | 42614 | 10 | 1704927 | 909394.38 | 1580809.62 | 2 | 1346785.88 | 1468555 | 967662.12 | 1149046.75 | 250996.73 | 718745.5 | 190277.38 | 442689.19 | 1773343.38 | 435820.44 | 372124.66 | 978514.06 | 594017.19 | 936963.56 | 915572.5 | 793271.88 | 9 | 8 | 8 | 10 | 9 | 10 | 0.579989552 | 0.705653292 | 0.761169422 | 0.926088593 |
| P61161 | Actr2 | Actin-related protein 2 | 1119.5 | 48.48 | 44761 | 18 | 127818530 | 130561226 | 141169970 | 129621602 | 153819490 | 86996514 | 127854210 | 141904146 | 135838514 | 120815562 | 58896066 | 75830002 | 66273258 | 57209086 | 80672202 | 142445106 | 75853762 | 96370498 | 102546354 | 113596122 | 0 | 4 | 0 | 8 | 0 | 8 | 0.496171552 | 0.552454633 | 0.777187376 | 0.865347409 |
| Q4V7C7 | Actr3 | Actin-related protein 3 | 1128.21 | 49.28 | 47357 | 19 | 132035970 | 134613778 | 138888562 | 138051842 | 129791114 | 122956690 | 117319962 | 125465938 | 115840226 | 128392450 | 91478762 | 96705882 | 84592490 | 75955322 | 86810746 | 121355322 | 116082322 | 121633922 | 119218602 | 129139090 | 0 | 0 | 0 | 12 | 0 | 12 | 0.646800296 | 0.714034202 | 0.902058445 | 0.995826047 |
| Q99JW2 | Acy1 | Aminoacylase-1 | 137.59 | 12.75 | 45781 | 3 | 652034.56 | 883274.12 | 813575.31 | 736385.38 | 648893 | 1104219.25 | 474431.75 | 2 | 485066.56 | 1246737 | 1968802.12 | 1610324.62 | 1079691.62 | 1741335.38 | 1370829.62 | 755020.06 | 871366.75 | 1194566.62 | 1128399.75 | 832918.88 | 0 | 4 | 2 | 8 | 2 | 8 | 2.081051275 | 2.347405326 | 1.280681338 | 1.444595926 |
| P03958 | Ada | Adenosine deaminase | 108.16 | 7.67 | 39992 | 2 | 1037655.44 | 874747.94 | 560212.06 | 387095.91 | 598517.31 | 2 | 720254.69 | 871154.75 | 604136.69 | 478913.19 | 2 | 562009.38 | 2 | 2 | 281743.66 | 994455.44 | 2 | 337045.28 | 2 | 2 | 2 | 4 | 4.5 | 7.5 | 4.5 | 7.5 | 0.243985902 | 0.315487472 | 0.385025645 | 0.497859778 |
| Q9Z0F8 | Adam17 | Disintegrin and metalloproteinase domain-containing protein 17 | 101.54 | 5.44 | 93056 | 3 | 2 | 2 | 203178.44 | 272327.16 | 386541.12 | 151502.56 | 176088.41 | 194177.84 | 273287.34 | 2 | 841951.88 | 645758.12 | 791699.94 | 1167681 | 867609.25 | 299561.25 | 230305.8 | 442023.25 | 15863.36 | 2 | 0 | 10 | 0 | 9.5 | 0 | 10 | 5.005158154 | 5.426898888 | 1.145820817 | 1.242369077 |
| Q05910 | Adam8 | Disintegrin and metalloproteinase domain-containing protein 8 | 630.72 | 18.64 | 90046 | 11 | 581307 | 2812762.75 | 485550.88 | 1150360.25 | 1061625 | 2705873.75 | 2880323.25 | 1014802.94 | 1933687.62 | 406192.75 | 15753664 | 12194551 | 10485853 | 9631999 | 10020924 | 1748590.38 | 1596504.25 | 1306191.38 | 2465615.25 | 1533906.75 | 0 | 5 | 0 | 11 | 0 | 11 | 9.535579311 | 6.496786556 | 1.420119453 | 0.967556629 |
| Q61072 | Adam9 | Disintegrin and metalloproteinase domain-containing protein 9 | 339.66 | 13.25 | 92079 | 7 | 655124.69 | 1759609 | 1136527.5 | 2181835.75 | 1627632.25 | 2465833 | 561307.12 | 1187416.75 | 1374555.12 | 2241418.25 | 1249987.75 | 1361930 | 2570206 | 2065107 | 1979160.25 | 868507.25 | 834843.81 | 1002926.69 | 1455549.25 | 995908 | 8 | 6 | 10 | 7 | 10 | 7 | 1.253461547 | 1.178258779 | 0.700709789 | 0.658669955 |
| Q99MU3 | Adar | Double-stranded RNA-specific adenosine deaminase | 117.02 | 2.04 | 130447 | 2 | 941808.38 | 883819.12 | 766467.25 | 1015624.25 | 506605.5 | 527988.06 | 1262808.25 | 231511.25 | 456389.19 | 1097073.5 | 263054.5 | 596626.88 | 428875.88 | 402751.69 | 2 | 1430000.5 | 419880.16 | 424770.38 | 107269.08 | 380371.38 | 1 | 5 | 6 | 8 | 6 | 8 | 0.411078647 | 0.472992064 | 0.671383966 | 0.772502512 |
| Q61549 | Adgre1 | Adhesion G protein-coupled receptor E1 | 74.89 | 2.26 | 102130 | 1 | 716401.38 | 548108.06 | 897251.62 | 2 | 2 | 910979.31 | 842935.69 | 1072272.12 | 732759.25 | 759813.12 | 2 | 442358.12 | 458233.28 | 2 | 2 | 799720.19 | 786927.94 | 934746.19 | 644675.25 | 627535.12 | 7 | 6 | 0 | 8 | 7 | 8 | 0.416602811 | 0.208531501 | 1.754864467 | 0.878401471 |
| P28474 | Adh5 | Alcohol dehydrogenase class-3 | 781.32 | 54.01 | 39548 | 14 | 24505870 | 34464042 | 23906614 | 35585754 | 29481052 | 25214346 | 26172746 | 30503812 | 25500980 | 29825952 | 25652056 | 26328074 | 19332026 | 17458424 | 22598756 | 32606522 | 21428140 | 26583684 | 34027278 | 32259356 | 4 | 11 | 5 | 7 | 5 | 11 | 0.752783748 | 0.811624343 | 0.992981421 | 1.070596828 |
| P55264 | Adk | Adenosine kinase | 122.23 | 7.48 | 40149 | 2 | 1623180.5 | 1619690.5 | 1748497.38 | 1959624 | 1896789.38 | 1784372.5 | 1804258.12 | 2020351.12 | 2123921.75 | 1988162.25 | 1926698.88 | 1599606.25 | 1394255.75 | 1614727 | 1645159 | 1539440.62 | 1824849.38 | 2082906.62 | 2287610.75 | 2636815.25 | 6 | 7 | 2 | 9 | 6 | 9 | 0.924576024 | 0.841517494 | 1.172228577 | 1.066922383 |
| Q8VDL4 | Adpgk | ADP-dependent glucokinase | 371.03 | 18.95 | 53902 | 6 | 6996739.5 | 7401538 | 10011605 | 6133316.5 | 10773526 | 5801740.5 | 7639626.5 | 8854509 | 9227532 | 6874834.5 | 8057686.5 | 9484150 | 9167536 | 9053632 | 10760194 | 6189829 | 5499076 | 6864655.5 | 7189917 | 7404064 | 9 | 7 | 4 | 6 | 9 | 7 | 1.126013703 | 1.21159708 | 0.802279017 | 0.863256736 |
| P54923 | Adprh | [Protein ADP-ribosylarginine] hydrolase | 112.23 | 7.18 | 40068 | 2 | 1935405.25 | 1293396.88 | 1554724.75 | 1123469.12 | 1790047.25 | 480383.56 | 1519575.25 | 1243428.62 | 1919990.75 | 928472 | 715930.75 | 1140556.5 | 660584.75 | 2 | 226650.09 | 1029000.25 | 385756.53 | 502431.31 | 1440348.75 | 947286.25 | 1 | 2 | 4 | 8 | 4 | 8 | 0.356464684 | 0.450392575 | 0.559282695 | 0.706652817 |
| Q8CG72 | Adprhl2 | Poly(ADP-ribose) glycohydrolase ARH3 | 135.88 | 9.73 | 39414 | 3 | 2094636.75 | 1533325.75 | 1710240.75 | 2144823.5 | 2119500 | 2098537.5 | 1982290.38 | 1159928.12 | 1835109.75 | 1813306.88 | 807874.94 | 1355638 | 2 | 1268929 | 1200876.12 | 2130216.5 | 1763735.25 | 2055935.12 | 1077914 | 2043969 | 0 | 10 | 3 | 11 | 3 | 11 | 0.482510508 | 0.521231869 | 0.944727373 | 1.020541534 |
| Q9JKV1 | Adrm1 | Proteasomal ubiquitin receptor ADRM1 | 107.36 | 7.62 | 42060 | 2 | 2617054.75 | 4937010 | 2169880.25 | 3343936.5 | 3498536.5 | 3641247 | 3617752.25 | 3293737.5 | 3527497.5 | 3600390.25 | 3327410.5 | 3277980.5 | 3434852 | 4550858 | 2994821 | 3019085.75 | 3909211.75 | 3487719.25 | 4487613.5 | 3875802.25 | 12 | 8 | 7 | 9 | 12 | 9 | 1.061540401 | 0.994643713 | 1.133584369 | 1.062147579 |
| P54822 | Adsl | Adenylosuccinate lyase | 553.41 | 29.75 | 54866 | 9 | 18813698 | 20400156 | 22335890 | 22167288 | 21696836 | 18079324 | 20461048 | 25272118 | 23443260 | 24138280 | 20415022 | 22059736 | 22965746 | 21566714 | 24761354 | 21060044 | 21573638 | 22745738 | 25576268 | 27256222 | 8 | 6 | 11 | 9 | 11 | 9 | 1.060283378 | 1.003362317 | 1.121407574 | 1.061205075 |
| P46664 | Adss | Adenylosuccinate synthetase isozyme 2 | 1223.46 | 37.72 | 50021 | 19 | 27595940 | 32264466 | 31360294 | 31027856 | 28939278 | 30509138 | 28644990 | 29652362 | 29312732 | 25360958 | 28031046 | 30244470 | 19468396 | 23898232 | 17288056 | 29295574 | 30876420 | 23417376 | 29709734 | 22462714 | 3 | 6 | 5 | 11 | 5 | 11 | 0.786638692 | 0.828896367 | 0.897967875 | 0.946206075 |
| P28650 | Adssl1 | Adenylosuccinate synthetase isozyme 1 | 392.31 | 18.38 | 50254 | 6 | 4745910 | 6220811.5 | 3927412.5 | 3878442.25 | 6505309 | 1367694 | 3341025 | 6143876 | 6149218.5 | 3751463.25 | 1857254.25 | 3027506.75 | 1957452.25 | 1668408 | 1560749.75 | 4892076.5 | 2450503.25 | 3799608.5 | 3503111 | 3347477 | 0 | 3 | 5 | 11 | 5 | 11 | 0.39842617 | 0.485290642 | 0.711799111 | 0.866984837 |
| Q32PX9 | Afg1l | AFG1-like ATPase | 43.88 | 2.29 | 54503 | 1 | 312712.41 | 557020 | 407141.59 | 2 | 573783 | 2 | 540008.19 | 461040.62 | 523108.53 | 2 | 2 | 372648.09 | 2 | 2 | 257629.09 | 2 | 2 | 81039.35 | 2 | 254759.92 | 4.5 | 3.5 | 7 | 7 | 7 | 7 | 0.340572293 | 0.413527862 | 0.181451726 | 0.220321341 |
| Q920A7 | Afg3l1 | AFG3-like protein 1 | 254.34 | 8.62 | 87047 | 5 | 3626907 | 4021630.5 | 3526595.25 | 3658533.5 | 3200927.75 | 3270029 | 4235210 | 3223704.25 | 1750377.25 | 2905540.5 | 6135459 | 3967930.75 | 6268462.5 | 6547007.5 | 5385317 | 4012403 | 4594071.5 | 2750356 | 4289658 | 2185402.5 | 1 | 11 | 1 | 9 | 1 | 11 | 1.569437978 | 1.839742117 | 0.988760324 | 1.159054411 |
| Q8JZQ2 | Afg3l2 | AFG3-like protein 2 | 410.61 | 12.97 | 89519 | 9 | 6845131.5 | 7766787 | 3979045 | 4973322.5 | 5312291 | 4622065.5 | 6322183 | 5467263 | 5756979 | 7902616 | 7953209.5 | 8205233 | 11657622 | 13026175 | 12836860 | 5787243.5 | 8783274 | 8861670 | 9259989 | 8774111 | 0 | 2 | 0 | 2 | 0 | 2 | 1.858914909 | 1.785072309 | 1.435983479 | 1.378941194 |
| Q64191 | Aga | N(4)-(beta-N-acetylglucosaminyl)-L-asparaginase | 67.4 | 4.34 | 37022 | 1 | 1305845 | 1195316.38 | 861038.69 | 1375179.5 | 2 | 750335.5 | 2 | 2 | 862448.06 | 894581 | 2 | 2 | 2 | 2 | 2 | 950466.81 | 1155577 | 2 | 918098.94 | 634960.5 | 2.5 | 7.5 | 5 | 7 | 5 | 7.5 | 2.110871E-06 | 3.988245E-06 | 0.77238981 | 1.459340804 |
| Q4KLH5 | Agfg1 | Arf-GAP domain and FG repeat-containing protein 1 | 50.02 | 1.78 | 58170 | 1 | 170444.45 | 2 | 427541.94 | 302932.03 | 766653.25 | 2 | 2 | 485402.41 | 517779.25 | 255855.11 | 2 | 239584.86 | 2 | 2 | 284163.09 | 541627 | 2 | 143505.69 | 179190.28 | 127306.91 | 5.5 | 8.5 | 8 | 12 | 8 | 12 | 0.314081446 | 0.415994432 | 0.594655515 | 0.78760903 |
| Q9ESW4 | Agk | Acylglycerol kinase, mitochondrial | 96.19 | 8.79 | 46976 | 2 | 2 | 487690.38 | 482012.44 | 163575.03 | 414769.56 | 2 | 218734.31 | 322614.16 | 573454.56 | 2 | 363847.75 | 860451.12 | 320461.81 | 2 | 645108 | 2 | 2 | 193462.02 | 2 | 93326.72 | 10.5 | 4.5 | 7 | 7 | 10.5 | 7 | 1.41459999 | 1.964349543 | 0.185262007 | 0.257259537 |
| Q8CJG0 | Ago2 | Protein argonaute-2 | 117.82 | 3.26 | 97304 | 2 | 175550.36 | 206889.12 | 138146.7 | 79420.74 | 344054.34 | 73115.12 | 199410.36 | 2 | 259556.38 | 142277.42 | 103657.57 | 271281.34 | 2 | 2 | 2 | 128630.74 | 121053.7 | 314919.19 | 206585.8 | 64167.21 | 5 | 9 | 8.5 | 11 | 8.5 | 11 | 0.397161631 | 0.556000057 | 0.884854273 | 1.23873755 |
| Q8C0I1 | Agps | Alkyldihydroxyacetonephosphate synthase, peroxisomal | 936.43 | 28.84 | 71684 | 17 | 13436782 | 12225078 | 7403059.5 | 6264905.5 | 4724326.5 | 12883685 | 13901180 | 3524863.5 | 6686323 | 6920957 | 12064020 | 9331645 | 7681022 | 12333332 | 12341856 | 13895757 | 13732777 | 3851112.75 | 5647809.5 | 5007409 | 8 | 12 | 10 | 11 | 10 | 12 | 1.220131887 | 1.223942086 | 0.956433476 | 0.959420204 |
| P50247 | Ahcy | Adenosylhomocysteinase | 1144.66 | 41.67 | 47688 | 18 | 193107282 | 183661506 | 140885602 | 175843538 | 160921730 | 201264898 | 179944066 | 145350658 | 143223618 | 156261282 | 110514234 | 115190322 | 94739314 | 101615874 | 91543746 | 188086002 | 158783122 | 152496530 | 161247826 | 152698050 | 0 | 9 | 0 | 11 | 0 | 11 | 0.60111385 | 0.621762479 | 0.951887661 | 0.984585586 |
| B5DFN2 | Ahcyl1 | S-adenosylhomocysteine hydrolase-like protein 1 | 359.2 | 18.01 | 53753 | 8 | 2557811 | 2881662.75 | 3449470.5 | 3998142.75 | 3023553 | 3471000 | 2682895.25 | 3045911.25 | 3017800.5 | 3185049.25 | 2109430.25 | 1988876.12 | 2462204 | 2648915.75 | 2050080.38 | 2812777.75 | 3285509.75 | 3398386.25 | 4335576.5 | 3533494.5 | 1 | 9 | 0 | 6 | 1 | 9 | 0.707671502 | 0.731010698 | 1.091454822 | 1.127451296 |
| Q68FL4 | Ahcyl2 | Putative adenosylhomocysteinase 3 | 386.48 | 11.91 | 66899 | 8 | 4496992 | 3464234 | 4303983.5 | 4629187.5 | 2457063.5 | 5118244 | 4135819.5 | 3306414.5 | 5939327.5 | 6137527 | 4462170 | 4221307.5 | 3513960.75 | 5195994 | 4354206 | 3449659 | 4806764 | 5153246.5 | 5495937.5 | 5538842.5 | 10 | 4 | 10 | 12 | 10 | 12 | 1.123824129 | 0.88271075 | 1.263183701 | 0.992171109 |
| Q8BK64 | Ahsa1 | Activator of 90 kDa heat shock protein ATPase homolog 1 | 404.23 | 23.96 | 38117 | 8 | 16231544 | 17276428 | 14032522 | 11083937 | 20005194 | 12335072 | 18610996 | 17010956 | 18770516 | 10794688 | 11023236 | 17326434 | 11792467 | 7089706 | 11581252 | 15798684 | 7700252.5 | 12828840 | 10461060 | 9717393 | 6 | 3 | 6 | 4 | 6 | 4 | 0.747976288 | 0.758661051 | 0.718637912 | 0.72890358 |
| O70200 | Aif1 | Allograft inflammatory factor 1 | 195.51 | 21.77 | 16911 | 2 | 3191494 | 2254997.25 | 2368332.75 | 1876165 | 1692851.75 | 1773797.12 | 2055104 | 1374937.88 | 1612740 | 1472655.75 | 500938.94 | 367437.47 | 275214.69 | 678180.56 | 2 | 2220180.5 | 1811020 | 1192203.5 | 739106.12 | 986513.06 | 0 | 3 | 0 | 9 | 0 | 9 | 0.160031548 | 0.219775856 | 0.610428706 | 0.838319023 |
| Q9Z0X1 | Aifm1 | Apoptosis-inducing factor 1, mitochondrial | 893.2 | 34.64 | 66765 | 15 | 17471806 | 11055251 | 8922933 | 9479438 | 14023986 | 12407372 | 9613569 | 9593093 | 10154236 | 11390891 | 14782355 | 13595276 | 15525849 | 14726548 | 12846626 | 10819698 | 10319605 | 11904951 | 11460840 | 15574892 | 7 | 11 | 0 | 6 | 7 | 11 | 1.17264398 | 1.344578294 | 0.985670565 | 1.130190636 |
| Q91VJ1 | Aim2 | Interferon-inducible protein AIM2 | 49.53 | 3.11 | 40155 | 1 | 690955.25 | 753650.19 | 742768.38 | 309942.06 | 664235.31 | 2 | 1011668.56 | 675068.25 | 1082591 | 306833.94 | 2 | 534126.69 | 2 | 2 | 2 | 664078.38 | 2 | 309403.5 | 168989.92 | 322002 | 1 | 2 | 4 | 7.5 | 4 | 7.5 | 0.168947032 | 0.173636625 | 0.463214325 | 0.476072121 |
| P31230 | Aimp1 | Aminoacyl tRNA synthase complex-interacting multifunctional protein 1 | 305.23 | 20.32 | 33997 | 6 | 3539439.75 | 5315381.5 | 4061426.5 | 4329355.5 | 3064261.5 | 3875512 | 3291654 | 4137779 | 3777760.5 | 3166413 | 4044107 | 7662512 | 4735076 | 3420773.75 | 3167535.75 | 4942599 | 5362496 | 4263392.5 | 5815965 | 4598845 | 12 | 4 | 8 | 0 | 12 | 4 | 1.133931948 | 1.26197901 | 1.230106542 | 1.369013933 |
| Q32PX2 | Aimp2 | Aminoacyl tRNA synthase complex-interacting multifunctional protein 2 | 181.31 | 13.12 | 35443 | 3 | 2927538.75 | 3083237.5 | 2791327.5 | 2081863.75 | 3028937.5 | 2499184.5 | 3500172 | 2949180.5 | 3394096.5 | 2273724.5 | 4055728 | 3835251.5 | 3285407 | 2155903.75 | 3569344 | 4217157.5 | 2115979.75 | 2147243.75 | 2370713.5 | 2802096 | 4 | 10 | 7 | 8 | 7 | 10 | 1.214817053 | 1.156350594 | 0.981332835 | 0.934103454 |
| O08915 | Aip | AH receptor-interacting protein | 98.18 | 7.58 | 37605 | 2 | 2781631.25 | 2271788.5 | 2608537.5 | 2711404.25 | 4072153.5 | 2971000.25 | 3216023 | 2233197.75 | 2397911.25 | 3269125.5 | 1518733.12 | 1060656.12 | 2095438 | 1822907.25 | 1215229.75 | 3442565 | 2799465 | 3990173.5 | 3298561.5 | 2734841.5 | 0 | 6 | 0 | 6 | 0 | 6 | 0.533934875 | 0.547513532 | 1.125996996 | 1.154632561 |
| Q9WTP6 | Ak2 | Adenylate kinase 2, mitochondrial | 337.36 | 26.78 | 26469 | 6 | 17050968 | 15519310 | 17435626 | 20147874 | 18146264 | 19808836 | 15247666 | 16091462 | 14531410 | 17320546 | 9463196 | 8058540.5 | 7168981.5 | 10841718 | 8515407 | 17559652 | 25295778 | 17166932 | 19596966 | 18396718 | 0 | 7 | 0 | 5 | 0 | 7 | 0.498842832 | 0.530697415 | 1.110033968 | 1.180917355 |
| Q9WTP7 | Ak3 | GTP:AMP phosphotransferase AK3, mitochondrial | 254.87 | 25.11 | 25426 | 4 | 2869026.75 | 3608399.25 | 1702553.88 | 2976899.5 | 1295143.38 | 2699537.75 | 2488632.25 | 1570261.5 | 805948.12 | 2395328.5 | 2107020.25 | 1833234.75 | 1165583.12 | 3294573.25 | 1148918.88 | 2818384 | 2593359 | 3229412.5 | 1962518.5 | 2288731.75 | 8 | 12 | 11 | 7 | 11 | 12 | 0.766889881 | 0.958796195 | 1.035366382 | 1.294456182 |
| Q9DBR0 | Akap8 | A-kinase anchor protein 8 | 85.84 | 3.49 | 76294 | 2 | 1432953.5 | 1785479 | 1329790.25 | 1508696.12 | 901917.25 | 1407714.75 | 1312576.38 | 1204737.75 | 8375441.5 | 555786.94 | 993138.19 | 1098905 | 451184.5 | 1066381.5 | 346840.25 | 621223.88 | 1418938.62 | 1156573.25 | 425347.38 | 1412299 | 3 | 5 | 3 | 10 | 3 | 10 | 0.568550455 | 0.307745041 | 0.723451744 | 0.391590025 |
| Q9JII6 | Akr1a1 | Alcohol dehydrogenase [NADP(+)] | 622.89 | 37.54 | 36587 | 10 | 19711822 | 17667154 | 15135505 | 16445080 | 19842120 | 16524590 | 19193334 | 16395446 | 16004760 | 17782664 | 15848949 | 17674086 | 13576456 | 13345440 | 14835311 | 25035800 | 20129850 | 18165188 | 17939240 | 18929216 | 4 | 6 | 3 | 3 | 4 | 6 | 0.847734425 | 0.876362586 | 1.128349068 | 1.166453642 |
| P45376 | Akr1b1 | Aldose reductase | 692.63 | 43.67 | 35732 | 12 | 81929666 | 77499010 | 82726018 | 81446298 | 69028578 | 69255298 | 66816330 | 78461362 | 71392002 | 77970058 | 110561778 | 98076330 | 67672642 | 60123542 | 80722802 | 87637738 | 65876674 | 80550418 | 70847986 | 76476770 | 12 | 9 | 9 | 10 | 12 | 10 | 1.062469885 | 1.146366498 | 0.971372548 | 1.048075773 |
| P45377 | Akr1b8 | Aldose reductase-related protein 2 | 618.5 | 42.72 | 36121 | 11 | 2524560.5 | 2885487.25 | 3187959.75 | 1935480.88 | 3167691 | 1990540.75 | 2506921.75 | 3207276.5 | 3355851 | 3101955.25 | 2486528.75 | 4144530.5 | 3616262.5 | 2467474.75 | 3603857 | 3122105.75 | 1815385 | 2664712.5 | 1139272.25 | 1401153.88 | 8 | 5 | 8 | 5 | 8 | 5 | 1.191040059 | 1.152240167 | 0.740274184 | 0.716158657 |
| Q8CG76 | Akr7a2 | Aflatoxin B1 aldehyde reductase member 2 | 128.4 | 7.36 | 40612 | 2 | 2218618.25 | 946388.44 | 768876.81 | 2132923.5 | 745168.06 | 1685560 | 2074808.75 | 619052.12 | 766918.25 | 1911446.5 | 687433 | 1316415.75 | 1246593.75 | 1065148.5 | 1204985.75 | 2075534.5 | 1631109.38 | 1564197 | 1450691.38 | 2244714.5 | 12 | 8 | 9 | 9 | 12 | 9 | 0.810422337 | 0.782196718 | 1.316247737 | 1.270405088 |
| P06214 | Alad | Delta-aminolevulinic acid dehydratase | 237.04 | 19.09 | 36032 | 4 | 5574829 | 5852783.5 | 5416799.5 | 4834762.5 | 5282699 | 5294828.5 | 4463786 | 5814810.5 | 6425961.5 | 6040768 | 4085427.75 | 4150240 | 2907144 | 3895298.5 | 3933843.5 | 5965630 | 6557158 | 5666135.5 | 6405606.5 | 5814811 | 0 | 2 | 0 | 8 | 0 | 8 | 0.703658585 | 0.676599473 | 1.127864538 | 1.084492634 |
| Q61490 | Alcam | CD166 antigen | 459.63 | 18.18 | 65092 | 8 | 3060232.25 | 2512798.25 | 1713197 | 2911986.25 | 1999733 | 3103421 | 1990410 | 2472610.25 | 2411045.75 | 2769197 | 10319752 | 10759348 | 7847498 | 9404139 | 10426511 | 3304052 | 3045259.5 | 2667592 | 2930240.5 | 2746924.25 | 0 | 6 | 0 | 6 | 0 | 6 | 3.997168458 | 3.825092706 | 1.204634563 | 1.152775753 |
| Q571I9 | Aldh16a1 | Aldehyde dehydrogenase family 16 member A1 | 307.95 | 6.23 | 84756 | 4 | 1547311.75 | 1339140.75 | 1759207.75 | 1768484 | 1545967.25 | 2043093.38 | 695490.5 | 1943945.5 | 1615397.5 | 1888868 | 3246910.5 | 2585368 | 2347460.5 | 2874486.75 | 2713110.5 | 1514888.75 | 3639365.25 | 2033837.25 | 2299912.5 | 2786625 | 0 | 4 | 0 | 5 | 0 | 5 | 1.729540629 | 1.681651544 | 1.542017188 | 1.499320421 |
| Q9Z110 | Aldh18a1 | Delta-1-pyrroline-5-carboxylate synthase | 1443.39 | 30.94 | 87266 | 24 | 16789276 | 16605424 | 14339203 | 16235999 | 23111060 | 18583576 | 18302608 | 14332950 | 21195324 | 19744596 | 39710534 | 35184546 | 44583054 | 48924290 | 47650102 | 17545054 | 23329384 | 23320368 | 19758636 | 17604710 | 0 | 3 | 0 | 9 | 0 | 9 | 2.481053505 | 2.344344008 | 1.166249771 | 1.101987787 |
| Q9CZS1 | Aldh1b1 | Aldehyde dehydrogenase X, mitochondrial | 179.42 | 10.02 | 57553 | 4 | 1378970 | 726236.25 | 1068729 | 792176.94 | 1079053.88 | 1253182 | 1331695.75 | 1023508.69 | 1247635.5 | 1448747.88 | 939785.62 | 1024898.94 | 1274773 | 1180636.75 | 1045417.56 | 1252886.75 | 1320760.25 | 1544693.38 | 1562737.12 | 1549857 | 11 | 2 | 6 | 5 | 11 | 5 | 1.083316544 | 0.866885235 | 1.433240135 | 1.146899047 |
| Q8R0Y6 | Aldh1l1 | Cytosolic 10-formyltetrahydrofolate dehydrogenase | 262.85 | 8.31 | 98709 | 6 | 3192059.5 | 4120721.25 | 2645054.75 | 3482712 | 3595872.5 | 3460014.5 | 4284520 | 1741004 | 3626590 | 1973139 | 2143437 | 2894498.75 | 1786371 | 2078351.38 | 1540900.75 | 3240518.25 | 2819572.25 | 3533943.5 | 2529236 | 3345470.5 | 1 | 8 | 7 | 11 | 7 | 11 | 0.613013701 | 0.692301869 | 0.907980697 | 1.025420365 |
| P47738 | Aldh2 | Aldehyde dehydrogenase, mitochondrial | 1519.55 | 61.27 | 56538 | 26 | 96247698 | 93425146 | 123559274 | 119905442 | 125399498 | 88216530 | 87492722 | 112421722 | 121685650 | 115895674 | 65628406 | 83335266 | 90636722 | 78417186 | 96696962 | 111042482 | 93563898 | 113638850 | 120345778 | 123997250 | 2 | 12 | 4 | 9 | 4 | 12 | 0.74250139 | 0.788862166 | 1.007253234 | 1.070144754 |
| P47740 | Aldh3a2 | Fatty aldehyde dehydrogenase | 349.23 | 16.12 | 53971 | 7 | 738503 | 1064408.62 | 930854.94 | 638659.06 | 652880.5 | 490524.25 | 952903.69 | 880155.25 | 625864.19 | 304300.81 | 416699.88 | 1338493.38 | 1239254 | 302490.62 | 1254055.5 | 678026.62 | 201776.02 | 328408.97 | 191321.44 | 480612.31 | 10 | 2 | 9 | 5 | 10 | 5 | 1.130595598 | 1.398692558 | 0.467081336 | 0.577839848 |
| Q5XI42 | Aldh3b1 | Aldehyde dehydrogenase family 3 member B1 | 203.38 | 9.19 | 52147 | 4 | 4523064.5 | 1901359.75 | 1953191.75 | 2808889.75 | 2508310 | 2174950.25 | 2176220.25 | 2610861 | 5415790 | 2190015.5 | 5210317.5 | 4590463.5 | 6536507 | 5113454.5 | 5818261.5 | 2318117 | 4490951.5 | 2911517.5 | 6466847 | 7630213 | 0 | 5 | 3 | 4 | 3 | 5 | 1.99119174 | 1.871863613 | 1.739172431 | 1.634947316 |
| Q8CHT0 | Aldh4a1 | Delta-1-pyrroline-5-carboxylate dehydrogenase, mitochondrial | 45.12 | 3.38 | 61841 | 1 | 308800.81 | 163457.16 | 189896.17 | 149361.06 | 163490.69 | 171352.52 | 216010.88 | 161437.25 | 203621.31 | 58791.71 | 2 | 117141.09 | 113651.38 | 2 | 49082.12 | 269982.09 | 61557.95 | 89877.54 | 2 | 29709.31 | 0 | 4 | 2 | 7 | 2 | 7 | 0.28705323 | 0.345012172 | 0.462693502 | 0.556115986 |
| Q9JLJ2 | Aldh9a1 | 4-trimethylaminobutyraldehyde dehydrogenase | 1114.28 | 49.39 | 53515 | 18 | 24936352 | 27214196 | 35795850 | 36142026 | 30274492 | 27163048 | 26450906 | 35995202 | 36507230 | 36353890 | 24949818 | 25984088 | 33086230 | 27835102 | 31143412 | 21789918 | 20929050 | 33446734 | 33719290 | 32359820 | 10 | 9 | 6 | 6 | 10 | 9 | 0.926379559 | 0.880152687 | 0.921496015 | 0.875512835 |
| P05064 | Aldoa | Fructose-bisphosphate aldolase A | 2284.05 | 79.4 | 39356 | 32 | 873672706 | 874774786 | 840246722 | 1013832770 | 885816706 | 1009478018 | 963755266 | 975091330 | 904854978 | 1050313922 | 614135874 | 651428674 | 697288066 | 810012418 | 688416642 | 1032909826 | 1217700098 | 1095962370 | 1275317506 | 1156400898 | 0 | 0 | 0 | 1 | 0 | 1 | 0.771171263 | 0.705880749 | 1.287399339 | 1.178402843 |
| P05063 | Aldoc | Fructose-bisphosphate aldolase C | 1339.98 | 54.82 | 39395 | 20 | 39491382 | 42831134 | 47020754 | 49537006 | 43610570 | 42141506 | 34972970 | 36549770 | 39433734 | 39510246 | 26245102 | 23378532 | 27989454 | 30687114 | 26776524 | 36423338 | 41220870 | 45593106 | 40814906 | 45788706 | 0 | 8 | 0 | 6 | 0 | 8 | 0.607111387 | 0.701302996 | 0.943144088 | 1.089470218 |
| Q9DBE8 | Alg2 | Alpha-1,3/1,6-mannosyltransferase ALG2 | 51.08 | 3.13 | 47405 | 1 | 2 | 2 | 2 | 2 | 2 | 2 | 2 | 2 | 2 | 2 | 2 | 2 | 526264.31 | 2 | 542794.62 | 2 | 2 | 2 | 2 | 2 | 7.5 | 12.5 | 7.5 | 12.5 | 7.5 | 12.5 | 106906.493 | 106906.493 | 1 | 1 |
| Q9DB25 | Alg5 | Dolichyl-phosphate beta-glucosyltransferase | 331.67 | 24.07 | 36791 | 6 | 2779985.5 | 2720061.5 | 3054323 | 3024555.25 | 4804590.5 | 3536512.75 | 2812052 | 3054136.5 | 2508318.75 | 4274237 | 4608453 | 5748151.5 | 6561112.5 | 5310270 | 5853950.5 | 5010623 | 4460241.5 | 5822724 | 4409222.5 | 3988750 | 1 | 3 | 0 | 1 | 1 | 3 | 1.714036104 | 1.735031918 | 1.446060868 | 1.463774162 |
| Q3TSG4 | Alkbh5 | RNA demethylase ALKBH5 | 55.78 | 7.59 | 44411 | 1 | 2 | 2 | 2 | 223495.69 | 186755.2 | 2 | 2 | 419566.78 | 2 | 2 | 2 | 202612.03 | 2 | 2 | 2 | 2 | 280549.72 | 126748.39 | 2 | 190468.34 | 10 | 10 | 12 | 9 | 12 | 10 | 0.493885746 | 0.482917562 | 1.457063768 | 1.424705389 |
| P48999 | Alox5 | Arachidonate 5-lipoxygenase | 348.54 | 10.09 | 77967 | 6 | 5211182 | 5592098.5 | 4024445 | 2274519 | 4344241.5 | 6652062.5 | 6064063.5 | 2567617.75 | 4628740 | 2405004 | 1702517.25 | 2301356 | 1716003.5 | 1800951.38 | 1540111.5 | 5600932 | 5980421.5 | 2033284.75 | 1795194.12 | 2009383 | 1 | 10 | 0 | 6 | 1 | 10 | 0.422490642 | 0.406001775 | 0.812217692 | 0.780518648 |
| P20291 | Alox5ap | Arachidonate 5-lipoxygenase-activating protein | 145.49 | 17.39 | 18070 | 3 | 10269977 | 10889844 | 10442481 | 16335452 | 9600235 | 15143988 | 13944342 | 16089465 | 15919705 | 16195206 | 20617514 | 14206963 | 21946034 | 20310198 | 20350530 | 12444424 | 15443666 | 19811412 | 19613022 | 19128586 | 1 | 2 | 4 | 8 | 4 | 8 | 1.693337579 | 1.260548945 | 1.502331095 | 1.118360509 |
| O08583 | Alyref | THO complex subunit 4 | 345.7 | 29.41 | 26940 | 6 | 4978518.5 | 5950879 | 4480411.5 | 5102544 | 4412668 | 4228137.5 | 5288272 | 4650475 | 3649808 | 5430088.5 | 6266498.5 | 6644567.5 | 6099895 | 7345934 | 6444590 | 5912227.5 | 5891946.5 | 7305514.5 | 5144697.5 | 7032554 | 0 | 3 | 0 | 2 | 0 | 3 | 1.316006313 | 1.411011916 | 1.255242272 | 1.345861175 |

| | | | | | | | | | | | | | | | | | | | | | | | | | | | | | | | | | | | | |
|---|---|---|---|---|---|---|---|---|---|---|---|---|---|---|---|---|---|---|---|---|---|---|---|---|---|---|---|---|---|---|---|---|---|---|---|---|
| Q5BJY6 | Amdhd2 | N-acetylglucosamine-6-phosphate deacetylase | 130.88 | 6.36 | 43539 | 2 | 2956993.25 | 2615948.5 | 3191588.5 | 4365033.5 | 2232129.5 | 2836521 | 3014796.75 | 2663709.25 | 2509087.75 | 3548584.75 | 2314193.5 | 1655677.5 | 2863378 | 3550105.25 | 2625898.75 | 2873196.25 | 2927805.25 | 4785816.5 | 4106513.75 | 3770505.25 | 9 | 8 | 9 | 4 | 9 | 8 | 0.846863219 | 0.89271401 | 1.201940222 | 1.267015559 |
| Q02356 | Ampd2 | AMP deaminase 2 | 132.28 | 3.28 | 94787 | 3 | 1997602.75 | 2406736.75 | 2945163 | 2302399 | 2999662.5 | 2277567.75 | 2586761 | 3273297.25 | 2334285 | 2631462.5 | 2913732.5 | 2166498.5 | 2 | 2578053 | 2250856.5 | 2487561 | 1950797.38 | 2910751.25 | 2927617 | 2986975 | 8 | 12 | 6 | 11 | 8 | 12 | 0.783234587 | 0.756228348 | 1.048384344 | 1.012235638 |
| O08739 | Ampd3 | AMP deaminase 3 | 108.6 | 2.61 | 88652 | 2 | 2 | 275841.75 | 445564.69 | 509218.06 | 767836.69 | 2 | 2 | 512972.81 | 966341.88 | 618622.75 | 748828.62 | 1006023.31 | 1099069.12 | 947782.75 | 1151622.88 | 2 | 2 | 751337 | 472971.81 | 355352.84 | 1 | 10 | 2 | 10 | 2 | 10 | 2.478567884 | 2.361041441 | 0.790440203 | 0.75295984 |
| Q810B6 | Ankfy1 | Rabankyrin-5 | 520.28 | 9.5 | 128653 | 9 | 2260575 | 3142019 | 2444842.75 | 3205509 | 4374895.5 | 2782428.5 | 3659254 | 3882044.75 | 4224823 | 4254227 | 2980541.75 | 3321058.25 | 4235945 | 2906966.5 | 3220835.25 | 3228046.25 | 2685744.75 | 3456721.25 | 2865038.25 | 2247449.75 | 9 | 12 | 8 | 3 | 9 | 12 | 1.080212486 | 0.886323681 | 0.938757407 | 0.770258566 |
| Q99NH0 | Ankrd17 | Ankyrin repeat domain-containing protein 17 | 99.3 | 0.96 | 274213 | 2 | 671293.44 | 873606.69 | 394106.47 | 2 | 463332.28 | 875978.38 | 442771.47 | 366773.06 | 2 | 2 | 2 | 732111.69 | 467974.25 | 2 | 2 | 889723 | 789327.62 | 803588.81 | 2 | 2 | 8.5 | 11 | 11 | 10 | 11 | 11 | 0.499551063 | 0.71199809 | 1.033426792 | 1.472918299 |
| B2RXR6 | Ankrd44 | Serine/threonine-protein phosphatase 6 regulatory ankyrin repeat subunit B | 88.2 | 1.81 | 107385 | 2 | 1104343.88 | 989595.06 | 2501037 | 2367270 | 2583239.5 | 1140419.5 | 1370698.38 | 2338095.25 | 2087410.25 | 1708380 | 2 | 320308.06 | 606909.69 | 591800.19 | 767775.94 | 841410.06 | 437206.31 | 807747.75 | 1102335.12 | 1567634 | 0 | 3 | 0 | 2 | 0 | 3 | 0.239568317 | 0.264522266 | 0.498280917 | 0.550182924 |
| Q8K298 | Anln | Anillin | 266.91 | 6.78 | 122794 | 5 | 2118728.75 | 1980347.5 | 2003021.25 | 1720598.62 | 1435279.5 | 2017354.75 | 1842732.25 | 1817433.5 | 1193087.75 | 1589734.5 | 940191.38 | 1243722.75 | 924029.62 | 315938.81 | 634309.62 | 2038471.75 | 1392028.75 | 467074.84 | 1302800.88 | 1285070.25 | 0 | 4 | 1 | 8 | 1 | 8 | 0.438345525 | 0.479672314 | 0.700525335 | 0.766570181 |
| Q8BH79 | Ano10 | Anoctamin-10 | 55.61 | 1.82 | 76188 | 1 | 2 | 2 | 2 | 417631.09 | 458182 | 2 | 235202.05 | 404734.38 | 383980.5 | 579119.5 | 777151.5 | 852273.19 | 1374865.12 | 1062282.75 | 961519.62 | 290066.38 | 2 | 479395.94 | 503243.41 | 583772.56 | 0 | 5.5 | 0 | 9.5 | 0 | 9.5 | 5.741016881 | 3.136601148 | 2.119707496 | 1.158100926 |
| Q6P9J9 | Ano6 | Anoctamin-6 | 88.88 | 2.96 | 106255 | 2 | 700269.62 | 376152.28 | 519150.06 | 587039 | 2 | 579902 | 445603.59 | 2 | 573507.31 | 562618.81 | 811242.94 | 643856 | 2 | 653596.31 | 2 | 311671.59 | 429004.94 | 633053.94 | 699620.81 | 539340.44 | 11 | 11 | 9 | 11 | 11 | 11 | 0.966135219 | 0.975511827 | 1.197047652 | 1.208665329 |
| O35381 | Anp32a | Acidic leucine-rich nuclear phosphoprotein 32 family member A | 487.19 | 27.53 | 28538 | 8 | 21466994 | 20309486 | 19489360 | 16973700 | 19900896 | 16661476 | 19727032 | 16454370 | 15562468 | 15358286 | 7219836.5 | 15248762 | 7751460.5 | 6406946 | 10426322 | 20829754 | 11469968 | 11301684 | 10228472 | 16302438 | 0 | 4 | 0 | 7 | 0 | 7 | 0.47944893 | 0.561739336 | 0.714611824 | 0.837264506 |
| Q9EST5 | Anp32b | Acidic leucine-rich nuclear phosphoprotein 32 family member B | 385.18 | 27.94 | 31079 | 8 | 39030546 | 36995526 | 36545950 | 25332370 | 39812518 | 27436470 | 42139894 | 33950414 | 34211506 | 21965270 | 11975588 | 22429962 | 10864926 | 7448038 | 14780276 | 34219746 | 17061458 | 17385138 | 21568990 | 21096050 | 0 | 1 | 1 | 4 | 1 | 4 | 0.379810734 | 0.422650519 | 0.62645351 | 0.697112739 |
| P97822 | Anp32e | Acidic leucine-rich nuclear phosphoprotein 32 family member E | 528.18 | 24.23 | 29622 | 6 | 24676742 | 29109652 | 31411002 | 24726266 | 30181986 | 20396028 | 30543850 | 25709610 | 29526012 | 20192414 | 6384944.5 | 18182350 | 12450206 | 10589692 | 14407733 | 30720846 | 13666128 | 15249906 | 10109668 | 17604306 | 0 | 4 | 0 | 5 | 0 | 5 | 0.442629733 | 0.490748985 | 0.623464188 | 0.691242351 |
| P97449 | Anpep | Aminopeptidase N | 492.1 | 10.87 | 109651 | 8 | 1955430.5 | 1906450.75 | 1396597.75 | 1581618.12 | 1337129.5 | 2543075.25 | 2094589.75 | 799782.19 | 2055773.25 | 1967363 | 1323874.25 | 1134846.5 | 1120102.5 | 474381.69 | 1152060.5 | 699963.75 | 1569998.75 | 1905722.62 | 666043.94 | 1949890.12 | 0 | 9 | 4 | 3 | 4 | 9 | 0.636556339 | 0.550205542 | 0.830552887 | 0.717885871 |
| P10107 | Anxa1 | Annexin A1 | 2157.66 | 74.86 | 38734 | 32 | 174471970 | 172956978 | 166713554 | 192902674 | 166016098 | 194338402 | 207584946 | 203405506 | 208188498 | 219677458 | 177384786 | 163166882 | 140475474 | 145612738 | 136844898 | 188397826 | 182167314 | 199673538 | 201427458 | 207455266 | 4 | 2 | 0 | 4 | 4 | 4 | 0.874491632 | 0.738955297 | 1.121480738 | 0.94766388 |
| P97384 | Anxa11 | Annexin A11 | 525.76 | 19.28 | 54079 | 10 | 4232738 | 4239313 | 3797040.5 | 5032036.5 | 4587623 | 2252849 | 6383096 | 6657444 | 5787171 | 6780937 | 4286827.5 | 9345141 | 8221026 | 5617265 | 7392010.5 | 5067412.5 | 2564407.5 | 6211084 | 4293189 | 5322002 | 2 | 7 | 8 | 6 | 8 | 7 | 1.592702571 | 1.251270526 | 1.07169637 | 0.841953862 |
| P07356 | Anxa2 | Annexin A2 | 1880.44 | 71.39 | 38676 | 31 | 114000714 | 120912042 | 147964914 | 183597634 | 191806386 | 118852850 | 137023698 | 224163218 | 227146722 | 212384306 | 129450778 | 164432930 | 199758626 | 171883058 | 203425954 | 120446050 | 111246514 | 173715698 | 199843362 | 170199330 | 7 | 12 | 9 | 7 | 9 | 12 | 1.145947947 | 0.94495318 | 1.02264233 | 0.843274883 |
| O35639 | Anxa3 | Annexin A3 | 1494.45 | 53.56 | 36384 | 23 | 35656430 | 36672018 | 36236422 | 34930710 | 30519452 | 33965946 | 31479576 | 35128338 | 32482574 | 34079086 | 53432270 | 53534506 | 67959578 | 76970082 | 70776042 | 32335598 | 42426562 | 39826022 | 33507966 | 34065218 | 0 | 12 | 0 | 9 | 0 | 12 | 1.854279336 | 1.930603848 | 1.046813967 | 1.08990217 |
| P97429 | Anxa4 | Annexin A4 | 1600.22 | 62.7 | 35916 | 22 | 148119954 | 162758514 | 139373314 | 163878770 | 156132770 | 145660418 | 168292162 | 198155026 | 188185778 | 209214370 | 163118322 | 192222514 | 171091890 | 154688434 | 182650690 | 147971586 | 144600194 | 192766306 | 200701490 | 193230370 | 4 | 8 | 9 | 11 | 9 | 11 | 1.121398131 | 0.949713563 | 1.141518648 | 0.966753656 |
| P48036 | Anxa5 | Annexin A5 | 2631.61 | 84.33 | 35752 | 36 | 207385858 | 179338434 | 118678498 | 107301794 | 223155442 | 157476946 | 180735250 | 127264866 | 223382290 | 140681202 | 213187186 | 315520354 | 230076450 | 118147610 | 235022450 | 151128978 | 131070906 | 132000450 | 118285362 | 114162594 | 5 | 8 | 6 | 4 | 6 | 8 | 1.330311315 | 1.340445678 | 0.773632271 | 0.779525831 |
| P14824 | Anxa6 | Annexin A6 | 1219.39 | 35.96 | 75885 | 21 | 14402894 | 14213114 | 20420662 | 22164726 | 33331874 | 12396656 | 16186389 | 27906926 | 43123746 | 27148036 | 19856814 | 26801352 | 32152504 | 23898442 | 32084770 | 13333851 | 11712870 | 22596918 | 23257744 | 20003008 | 7 | 10 | 11 | 7 | 11 | 10 | 1.289483071 | 1.063363979 | 0.869621614 | 0.71712791 |
| Q07076 | Anxa7 | Annexin A7 | 940.16 | 39.96 | 49925 | 17 | 32629286 | 28557588 | 22625132 | 29040586 | 23805810 | 33397952 | 33327636 | 26154360 | 24897258 | 29094482 | 53988222 | 50497058 | 43796738 | 42667494 | 49499678 | 30753040 | 30077082 | 26561904 | 29410552 | 30364024 | 0 | 7 | 0 | 11 | 0 | 11 | 1.759490719 | 1.637137785 | 1.076893918 | 1.00200797 |
| O35643 | Ap1b1 | AP-1 complex subunit beta-1 | 1481.33 | 29.27 | 103935 | 20 | 14256028 | 14068773 | 17435386 | 15508918 | 17886432 | 13430468 | 11384600 | 15975513 | 16481691 | 14301395 | 6781939.5 | 5704941.5 | 7107218 | 4930555 | 7176565.5 | 11480509 | 10142868 | 12698150 | 11347795 | 13937547 | 0 | 0 | 0 | 4 | 0 | 4 | 0.40049276 | 0.442917358 | 0.753034737 | 0.832804459 |
| P22892 | Ap1g1 | AP-1 complex subunit gamma-1 | 556.66 | 16.67 | 91350 | 10 | 11817118 | 11262903 | 15879660 | 15824366 | 18750032 | 10234695 | 11808577 | 13903822 | 17361028 | 12827280 | 6541864 | 7704800 | 9013904 | 6820035 | 10284966 | 10553962 | 10304549 | 12437823 | 15157330 | 12707895 | 0 | 6 | 1 | 10 | 1 | 10 | 0.548936895 | 0.610347375 | 0.831744408 | 0.924793033 |
| P35585 | Ap1m1 | AP-1 complex subunit mu-1 | 152.28 | 8.27 | 48543 | 3 | 4602712.5 | 3920404 | 3805348 | 4266384.5 | 3566014.25 | 4240057 | 3626939.5 | 3715794.25 | 3690692 | 4747461 | 3099140.5 | 2370494.5 | 2248026.75 | 2022577 | 2302721.5 | 3630415.5 | 3943334.75 | 4156437.25 | 3960636 | 3223179.5 | 0 | 10 | 0 | 10 | 0 | 10 | 0.597343482 | 0.60151811 | 0.938154422 | 0.944710861 |
| Q7TN05 | Ap1s3 | AP-1 complex subunit sigma-3 | 37.91 | 5.84 | 18440 | 1 | 1121027.25 | 1431482.62 | 1720830.25 | 1359817.38 | 1674457.5 | 1512531.62 | 1295067 | 1591734.25 | 1752318.62 | 1578887.75 | 775720.44 | 2 | 2 | 718021 | 2 | 980667 | 854427.5 | 2 | 1452535.88 | 2 | 0 | 3 | 0 | 1 | 0 | 3 | 0.204409707 | 0.1932681 | 0.449891569 | 0.425278791 |
| P17426 | Ap2a1 | AP-2 complex subunit alpha-1 | 1241.83 | 25.28 | 107664 | 23 | 21073554 | 21818626 | 18125148 | 18011586 | 19703046 | 17730580 | 20185044 | 18147686 | 19749938 | 15587065 | 15080250 | 15472211 | 16946672 | 12630575 | 18329464 | 19142806 | 15281752 | 18228386 | 18511746 | 19642468 | 2 | 8 | 4 | 12 | 4 | 12 | 0.794668434 | 0.858412509 | 0.919734177 | 0.993510361 |
| P17427 | Ap2a2 | AP-2 complex subunit alpha-2 | 1015.09 | 24.2 | 104017 | 19 | 4166443.5 | 3762091.5 | 4352758 | 3685424.75 | 4247260 | 3987717.25 | 4134502.5 | 3535753.5 | 4274629 | 2508313 | 1812691 | 2872644.5 | 3925130.25 | 3300816.5 | 3818726 | 3792417 | 2517348 | 3233773 | 3047829 | 3437302.25 | 2 | 6 | 6 | 6 | 6 | 6 | 0.778174808 | 0.852994986 | 0.792949782 | 0.869190549 |
| P62944 | Ap2b1 | AP-2 complex subunit beta | 1609.47 | 29.35 | 104553 | 25 | 20009406 | 19448886 | 20353208 | 16827374 | 22626248 | 15179642 | 19273844 | 13969652 | 20114294 | 20033690 | 14246619 | 16646146 | 15561221 | 13576316 | 17306720 | 20830546 | 15738376 | 20554358 | 16946394 | 18657618 | 1 | 10 | 7 | 9 | 7 | 10 | 0.779095622 | 0.873162948 | 0.934137692 | 1.046924662 |
| P84091 | Ap2m1 | AP-2 complex subunit mu | 486.25 | 26.44 | 49655 | 9 | 5658845 | 4917709 | 5391327.5 | 4390251 | 6677608 | 5231250.5 | 6152359 | 7475665.5 | 7640794 | 4256129 | 8849765 | 6579379 | 7594615.5 | 7362118.5 | 5741760 | 5501944.5 | 5836882 | 7245026 | 6533956 | 7527840.5 | 2 | 4 | 8 | 11 | 8 | 11 | 1.336291788 | 1.174645774 | 1.207499717 | 1.061433178 |
| P62743 | Ap2s1 | AP-2 complex subunit sigma | 71.79 | 13.38 | 17018 | 2 | 2 | 2 | 2 | 2 | 2 | 2 | 2 | 378888.94 | 2 | 2 | 3498136.75 | 1413719.62 | 1262728.5 | 1105073 | 1488366.12 | 2 | 2 | 2 | 2 | 2 | 0 | 12.5 | 0 | 10 | 0 | 12.5 | 876802.399 | 23.14092056 | 1 | 2.63924E-05 |
| Q9Z1T1 | Ap3b1 | AP-3 complex subunit beta-1 | 379.75 | 6.61 | 122740 | 7 | 7169919 | 7866085.5 | 4666507.5 | 4955009 | 5420815 | 6516627 | 6091590 | 5793720 | 6646594.5 | 5866840 | 6482710 | 5622101.5 | 5874252 | 4377503.5 | 6178682.5 | 5264524 | 6635059 | 5067492.5 | 6482422.5 | 6066341 | 12 | 12 | 8 | 9 | 12 | 12 | 0.948697744 | 0.923011697 | 0.981298932 | 0.954730206 |
| O54774 | Ap3d1 | AP-3 complex subunit delta-1 | 315.36 | 6.09 | 135081 | 6 | 2808330 | 2244349 | 2790976.75 | 3073251.5 | 3448824.75 | 2648832.5 | 3038780 | 3250604.5 | 3050053.75 | 2469140 | 2031895.12 | 1865146.25 | 1577748.88 | 1735517.62 | 1667938.25 | 3413284.75 | 1314963.62 | 2922310.25 | 2167795 | 2715637.75 | 0 | 8 | 0 | 9 | 0 | 9 | 0.618015575 | 0.614096554 | 0.872492357 | 0.866959623 |
| P53676 | Ap3m1 | AP-3 complex subunit mu-1 | 146.77 | 13.16 | 46981 | 4 | 3187036.5 | 3048453.75 | 4198279 | 4388599 | 3151330.25 | 3486032 | 3041545 | 5535788 | 4677826 | 5948398 | 2624196.25 | 2960328.5 | 3253078 | 2227640.25 | 1953815.25 | 5280945 | 4531195 | 4684633.5 | 4896457 | 5033286 | 3 | 0 | 1 | 11 | 3 | 11 | 0.724339415 | 0.573789955 | 1.359014479 | 1.076551739 |
| Q3TAP4 | Ap5b1 | AP-5 complex subunit beta-1 | 48.42 | 1.37 | 94679 | 1 | 433972 | 352896.16 | 470055.12 | 557376.81 | 577058.69 | 414740.91 | 366307.19 | 405312.38 | 511404.78 | 492662.5 | 453936.81 | 350582.47 | 490254.75 | 362995.84 | 387247.44 | 446836.53 | 578817.75 | 560189.25 | 537278.38 | 563667.5 | 7 | 7 | 7 | 2 | 7 | 7 | 0.855169591 | 0.933651501 | 1.123540906 | 1.226604894 |
| Q8R5A3 | Apbb1ip | Amyloid beta A4 precursor protein-binding family B member 1-interacting protein | 47.53 | 1.34 | 74319 | 1 | 2 | 629051.94 | 391181.19 | 2 | 2 | 409234.06 | 2 | 2 | 316943.16 | 518750.97 | 2036772.75 | 1898100.88 | 1560816.75 | 1382374.12 | 1703166.25 | 281777.69 | 278846.5 | 2 | 2 | 2 | 0 | 10.5 | 0 | 7 | 0 | 10.5 | 8.410999439 | 6.892930249 | 0.549508614 | 0.450329901 |
| Q8R146 | Apeh | Acylamino-acid-releasing enzyme | 207.34 | 5.74 | 81581 | 4 | 1404984.75 | 3191251.5 | 3056773.25 | 1163976.75 | 3172110.75 | 846654.19 | 2754686.5 | 1796811 | 2998012 | 1793847.88 | 2 | 1837346.25 | 1203354.38 | 539896.81 | 1688638.25 | 2778650 | 1351993.25 | 625281 | 811677.5 | 1268737.5 | 5 | 4 | 5 | 6 | 5 | 6 | 0.439502465 | 0.517098303 | 0.570213024 | 0.670886309 |
| P28352 | Apex1 | DNA-(apurinic or apyrimidinic site) lyase | 338.68 | 25.87 | 35490 | 6 | 6459332 | 7471710.5 | 7719575.5 | 7658523.5 | 8321810 | 6548145.5 | 5652162.5 | 8042180 | 7936422 | 7549343 | 6656400.5 | 6829427 | 6322680 | 5508986.5 | 6189533 | 7875797.5 | 7859562 | 7336946.5 | 8399904 | 8671268 | 2 | 6 | 6 | 7 | 6 | 7 | 0.837263629 | 0.881851878 | 1.066767552 | 1.123577971 |
| O35841 | Api5 | Apoptosis inhibitor 5 | 614.49 | 20.63 | 56785 | 10 | 17806082 | 14854430 | 16057500 | 14692676 | 17485268 | 14489432 | 16112909 | 15038181 | 16515048 | 12646082 | 9813522 | 10870504 | 8826902 | 9062649 | 8833780 | 17293072 | 17253454 | 13930846 | 13168368 | 12390238 | 0 | 6 | 0 | 12 | 0 | 12 | 0.586028763 | 0.633774198 | 0.915199988 | 0.989763943 |
| Q9WVQ5 | Apip | Methylthioribulose-1-phosphate dehydratase | 87.1 | 9.54 | 26949 | 2 | 1622713 | 1408415.62 | 1333911.12 | 636368.81 | 1354877.88 | 664875.19 | 1389966 | 275098.94 | 610960.69 | 714469 | 2 | 708134.38 | 292838 | 554486.31 | 281359.31 | 527126.44 | 314075.72 | 238658.59 | 695445.25 | 706457.69 | 1 | 2 | 6 | 8 | 6 | 8 | 0.288976908 | 0.502496338 | 0.390442394 | 0.678932703 |
| Q9D7N9 | Apmap | Adipocyte plasma membrane-associated protein | 407.21 | 25.3 | 46434 | 7 | 1327941 | 695056.88 | 471551 | 509244.84 | 525800.38 | 704318.12 | 155242.98 | 601353.25 | 396343.69 | 651756.75 | 3951952.5 | 1831665 | 2141260.75 | 1777084.25 | 1997864.38 | 473584.06 | 596468.75 | 1084666.12 | 726964.25 | 766984.25 | 0 | 9 | 0 | 6 | 0 | 9 | 3.314779702 | 4.663115948 | 1.033735701 | 1.454223165 |
| P38483 | Apobec1 | C->U-editing enzyme APOBEC-1 | 114.85 | 14.85 | 27274 | 3 | 1447477.5 | 1624671.75 | 762873.38 | 413801.78 | 847572.5 | 171176.53 | 1844299.5 | 2286023.25 | 2777117.75 | 434185.72 | 1826526.75 | 2270645 | 2315144.75 | 334949.44 | 1011618.25 | 1350939.5 | 2 | 332376.12 | 1727495 | 1550218.12 | 7 | 12 | 12 | 7 | 12 | 12 | 1.522425417 | 1.032754945 | 0.973438849 | 0.660343537 |
| Q8VBT6 | Apobr | Apolipoprotein B receptor | 1095.55 | 19.96 | 102705 | 17 | 16370340 | 16354740 | 17635972 | 22875438 | 20154668 | 14174222 | 14715416 | 21003998 | 17673752 | 16796570 | 6984240 | 8092972.5 | 8685173 | 5829414 | 7365107.5 | 14895387 | 12503558 | 17470098 | 18724398 | 19805150 | 0 | 8 | 0 | 12 | 0 | 12 | 0.395721691 | 0.438065115 | 0.893003072 | 0.988557116 |
| Q78IK4 | Apool | MICOS complex subunit Mic27 | 433.13 | 27.92 | 29261 | 6 | 10763256 | 10324312 | 10618502 | 10179094 | 10640308 | 12087951 | 11637010 | 10536372 | 10380883 | 9318317 | 6968096 | 7585226.5 | 9436677 | 6036326 | 9584420 | 9387035 | 11838509 | 9529942 | 13245232 | 9422321 | 0 | 10 | 2 | 12 | 2 | 12 | 0.754124504 | 0.734068833 | 1.017088223 | 0.990039127 |
| P08592 | App | Amyloid-beta A4 protein | 50.75 | 1.17 | 86704 | 1 | 2 | 389052.5 | 2 | 2 | 583217.81 | 2 | 387140.78 | 2 | 420845.28 | 2 | 520850.06 | 331053.84 | 2 | 2 | 2 | 225186.67 | 211573.52 | 373450.09 | 582583.56 | 495659.06 | 11.5 | 8 | 12.5 | 6 | 12.5 | 8 | 0.876201437 | 1.054354297 | 1.942300641 | 2.33721715 |
| P08030 | Aprt | Adenine phosphoribosyltransferase | 762.63 | 71.11 | 19724 | 11 | 42578694 | 41728886 | 44145282 | 48188598 | 42677686 | 39663918 | 38598678 | 49184478 | 44501390 | 53468682 | 43292590 | 42781118 | 47127538 | 33801958 | 45318010 | 50912974 | 44751794 | 56751270 | 55132522 | 54655306 | 11 | 1 | 10 | 3 | 11 | 3 | 0.968092471 | 0.941903568 | 1.195536688 | 1.163193975 |
| Q8CFQ3 | Aqr | RNA helicase aquarius | 66.29 | 1.08 | 170294 | 2 | 33442296 | 37600186 | 48683214 | 59098126 | 53297946 | 56480222 | 60680702 | 62961922 | 85331058 | 86137426 | 46123562 | 64270870 | 121247786 | 144586818 | 101383730 | 40006734 | 77225498 | 71917706 | 114182186 | 79055666 | 3 | 3 | 7 | 11 | 7 | 11 | 2.057595762 | 1.35843158 | 1.647358597 | 1.087591637 |
| P04627 | Araf | Serine/threonine-protein kinase A-Raf | 127.76 | 4.64 | 67581 | 2 | 227320.25 | 148736.16 | 233387.7 | 172654.81 | 165946.3 | 176058.05 | 143201.81 | 132237.5 | 137493.55 | 116409.99 | 213895.83 | 222334.19 | 329856.81 | 296414.69 | 220991.23 | 132652.19 | 145806.44 | 160021.08 | 188088.27 | 2 | 6 | 4 | 0 | 10 | 6 | 10 | 1.353830728 | 1.819522416 | 0.660907272 | 0.888246641 |
| Q4LDD4 | Arap1 | Arf-GAP with Rho-GAP domain, ANK repeat and PH domain-containing protein 1 | 289.2 | 6.4 | 162276 | 6 | 2247274.5 | 1867383.12 | 3263438.5 | 2656656 | 2484999.25 | 2022382 | 2424993.25 | 2860902 | 3206686 | 4170015.5 | 3477942 | 3725045.75 | 4654903.5 | 4547419 | 3352327 | 2469213.5 | 2679524 | 4336658.5 | 3840138.25 | 2328190 | 0 | 7 | 3 | 11 | 3 | 11 | 1.578117382 | 1.34543179 | 1.250322294 | 1.065968465 |
| Q5XJY5 | Arcn1 | Coatomer subunit delta | 730.1 | 29.35 | 57229 | 14 | 21812922 | 20882020 | 19575370 | 22054354 | 19473418 | 21637182 | 21134904 | 17149868 | 20953602 | 20769078 | 14824155 | 16118553 | 16458550 | 14717467 | 12848068 | 20689190 | 17854228 | 21141150 | 22718240 | 20076190 | 0 | 12 | 0 | 12 | 0 | 12 | 0.722236771 | 0.737538127 | 0.987291808 | 1.008208638 |
| P61205 | Arf3 | ADP-ribosylation factor 3 | 875.98 | 62.98 | 20601 | 14 | 10790344 | 12932249 | 17021412 | 10317325 | 20344346 | 6253486 | 12452902 | 16226819 | 14877004 | 10834996 | 9424043 | 15819853 | 13247378 | 7522325.5 | 14912382 | 13672286 | 8651838 | 9531147 | 10883544 | 13403475 | 9 | 8 | 12 | 10 | 12 | 10 | 0.853237234 | 1.004629789 | 0.78624408 | 0.925749829 |
| P61750 | Arf4 | ADP-ribosylation factor 4 | 590.44 | 42.78 | 20397 | 10 | 8581086 | 8018323 | 7183045.5 | 8083450 | 8121187 | 6495719 | 9799070 | 8562870 | 8039282 | 9276112 | 7597797 | 7638341.5 | 6488763 | 5423394 | 7960250 | 8192907 | 6766664 | 7557488 | 8911570 | 8868593 | 3 | 10 | 3 | 10 | 3 | 10 | 0.877996978 | 0.832487643 | 1.007755765 | 0.955520626 |
| P84083 | Arf5 | ADP-ribosylation factor 5 | 669.76 | 52.78 | 20530 | 11 | 1128082.12 | 1408733.12 | 1240618.62 | 2 | 1614578.5 | 884706 | 1228961.88 | 1438650.25 | 1384001.75 | 580285.88 | 2 | 804672.12 | 282275.03 | 2 | 2 | 1130841.25 | 2 | 596027.44 | 910795 | 873307.75 | 3.5 | 5.5 | 1 | 6 | 3.5 | 6 | 0.20158573 | 0.197032958 | 0.651143192 | 0.636437257 |
| P62331 | Arf6 | ADP-ribosylation factor 6 | 420.41 | 39.43 | 20082 | 7 | 5936702 | 4723340 | 4013333.75 | 5201588.5 | 3779954 | 9401017 | 3813029.25 | 4564431.5 | 4064737.5 | 4963951 | 9785669 | 5817173.5 | 7834184 | 9495791 | 7299880.5 | 3542312.5 | 8498274 | 6600495 | 7093065 | 4831042 | 1 | 7 | 3 | 10 | 3 | 10 | 1.700817461 | 1.50081876 | 1.292128266 | 1.140187225 |
| Q3MID3 | Arfgap2 | ADP-ribosylation factor GTPase-activating protein 2 | 47.09 | 3.46 | 56556 | 1 | 290360.69 | 224143.67 | 173772.69 | 191378.28 | 2 | 2 | 176087.56 | 2 | 130228.29 | 145343.69 | 92963.64 | 198217.08 | 201225.47 | 2 | 2 | 179329.83 | 260586.61 | 227957.19 | 2 | 206677.64 | 8 | 11.5 | 11 | 4 | 11 | 11.5 | 0.559775009 | 1.09021461 | 0.994197672 | 1.936293707 |
| Q4KLN7 | Arfgap3 | ADP-ribosylation factor GTPase-activating protein 3 | 45.32 | 1.52 | 57671 | 1 | 127962.68 | 221857.22 | 344579.12 | 223941.42 | 251123.23 | 2 | 131413.84 | 347979.56 | 305261.22 | 283402.78 | 2 | 142076.48 | 61492.52 | 2 | 85382.2 | 89654.02 | 2 | 158954.36 | 2 | 218740.41 | 1 | 2 | 5 | 6 | 5 | 6 | 0.24708352 | 0.270542256 | 0.39963002 | 0.437571908 |
| D4A631 | Arfgef1 | Brefeldin A-inhibited guanine nucleotide-exchange protein 1 | 129.83 | 1.19 | 207890 | 3 | 900670.5 | 797805 | 476714.38 | 1466965.75 | 1630310.88 | 625216.75 | 936572 | 663550.19 | 871506.31 | 636990.75 | 610203.38 | 294433.03 | 560400.38 | 1286071.5 | 476917.47 | 500447.97 | 651307.88 | 872016.19 | 359640.53 | 565594.75 | 6 | 5 | 5 | 6 | 6 | 6 | 0.612242061 | 0.864533354 | 0.559322153 | 0.789806333 |
| A2A5R2 | Arfgef2 | Brefeldin A-inhibited guanine nucleotide-exchange protein 2 | 143.31 | 2.12 | 202240 | 3 | 962771.88 | 560784.75 | 377357.75 | 846448.19 | 1218363 | 606296 | 1055350 | 356619 | 1176584.62 | 349206.47 | 339366.59 | 662644.19 | 245186.91 | 775767.69 | 1179755 | 1123124 | 695545.88 | 912552.5 | 1142599.75 | 396027.41 | 8 | 11 | 11 | 9 | 11 | 11 | 0.807600103 | 0.903687836 | 1.076688103 | 1.20479175 |
| Q9JHU5 | Arfip1 | Arfaptin-1 | 71.56 | 3.28 | 40779 | 1 | 879414.31 | 812415.62 | 713326.69 | 2 | 701912 | 2 | 852656.69 | 2 | 743389.19 | 2 | 2 | 2 | 2 | 867785.81 | 2 | 660821.75 | 560534.62 | 2 | 156836.5 | 823769 | 6 | 7.5 | 11 | 10.5 | 11 | 10.5 | 0.279296455 | 0.543712783 | 0.708694503 | 1.379631764 |
| Q63055 | Arfrp1 | ADP-ribosylation factor-related protein 1 | 75.32 | 8.96 | 22659 | 1 | 2 | 2 | 2 | 2 | 2 | 2 | 2 | 2 | 2 | 2 | 2 | 2 | 2 | 2 | 2 | 2 | 2 | 2 | 2 | 2 | 12.5 | 12.5 | 12.5 | 12.5 | 12.5 | 12.5 | 1 | 1 | 1 | 1 |
| Q5FWK3 | Arhgap1 | Rho GTPase-activating protein 1 | 97.57 | 4.78 | 50411 | 2 | 655062.69 | 462137.91 | 797268.44 | 761878.56 | 707886.19 | 563888.44 | 435868.88 | 615892.88 | 708208.44 | 461515.66 | 336661.72 | 496085.88 | 460977.66 | 393907.25 | 531017.75 | 413021.84 | 242566.67 | 634972.12 | 646574.12 | 180382.17 | 2 | 2 | 5 | 8 | 5 | 8 | 0.655584217 | 0.796535769 | 0.625700543 | 0.760227062 |
| Q6YSD8 | Arhgap10 | Rho GTPase-activating protein 10 | 51.96 | 0.89 | 89366 | 1 | 765254.31 | 740616.5 | 870887.12 | 713201.06 | 807463.69 | 793611.56 | 710421.81 | 915630.12 | 1235333.88 | 970539.94 | 471807.03 | 459191.34 | 668398.56 | 442574.38 | 2 | 716455.75 | 762442.5 | 910481.75 | 845766.19 | 813679.88 | 0 | 9 | 0 | 8 | 0 | 9 | 0.523929139 | 0.441456456 | 1.038847054 | 0.875320163 |
| Q3UIA2 | Arhgap17 | Rho GTPase-activating protein 17 | 459.8 | 13.71 | 92202 | 8 | 3759071.5 | 4572134.5 | 5820784 | 5900824 | 5833046.5 | 4571993 | 3755844 | 4929673.5 | 5082329.5 | 5510133 | 3673514.75 | 4110693.25 | 2436612.75 | 3373478.25 | 3229381.25 | 4536390 | 3661387 | 3899119.5 | 5247094.5 | 3997795.5 | 1 | 5 | 1 | 7 | 1 | 7 | 0.649917751 | 0.705396197 | 0.824457294 | 0.894834829 |
| Q8BYW1 | Arhgap25 | Rho GTPase-activating protein 25 | 449.72 | 16.05 | 73383 | 8 | 9068028 | 11308087 | 6841170.5 | 11143228 | 11040658 | 5287468.5 | 8856660 | 3947353 | 6129612 | 8532422 | 5507163 | 6332788 | 5864185 | 2625252 | 4876363 | 10754549 | 13866842 | 11182910 | 12826310 | 13105140 | 0 | 4 | 8 | 0 | 8 | 4 | 0.510225775 | 0.769558645 | 1.249681923 | 1.884858772 |
| Q640N3 | Arhgap30 | Rho GTPase-activating protein 30 | 179.92 | 3.18 | 120114 | 3 | 515828.59 | 421009.59 | 2 | 190532.2 | 421268.69 | 278149.56 | 562255.88 | 203631.3 | 513211.44 | 336861.16 | 1026521.44 | 1018189.25 | 1052182.88 | 886350.62 | 899897.5 | 300406.12 | 365594.06 | 164644.23 | 262098.05 | 2 | 0 | 7.5 | 0 | 6 | 0 | 7.5 | 3.153178477 | 2.578067478 | 0.705615059 | 0.576917307 |
| Q3TBD2 | Arhgap45 | Rho GTPase-activating protein 45 | 516.05 | 9.77 | 122902 | 9 | 8338293 | 9331254 | 12721108 | 10986869 | 11600024 | 7408723 | 8299012 | 10014531 | 10033499 | 7926435 | 3740066.25 | 4884473.5 | 5015516 | 4531720.5 | 5416835.5 | 8506425 | 7069089 | 7603472 | 8723605 | 8062815 | 0 | 9 | 0 | 9 | 0 | 9 | 0.445256767 | 0.540005122 | 0.754383838 | 0.914912848 |
| Q5XI73 | Arhgdia | Rho GDP-dissociation inhibitor 1 | 838.57 | 80.39 | 23407 | 13 | 152337906 | 158255778 | 173653458 | 177965010 | 177413554 | 138094322 | 137578802 | 161678738 | 153094642 | 171669378 | 103417106 | 105216970 | 93833882 | 96809474 | 92593586 | 132299490 | 156260386 | 161944370 | 147219794 | 142174338 | 0 | 11 | 0 | 10 | 0 | 11 | 0.585821771 | 0.645401873 | 0.881224065 | 0.970847604 |
| Q61599 | Arhgdib | Rho GDP-dissociation inhibitor 2 | 684.78 | 81 | 22851 | 12 | 40057570 | 43259786 | 55311622 | 61691430 | 54552982 | 41224954 | 18630292 | 54044530 | 48351378 | 63926422 | 56668978 | 45838054 | 50871482 | 54350734 | 57851118 | 40187418 | 46118118 | 63574486 | 61191710 | 56611970 | 11 | 9 | 8 | 10 | 10 | 10 | 1.042008999 | 1.174211744 | 1.050261473 | 1.183511234 |
| Q61210 | Arhgef1 | Rho guanine nucleotide exchange factor 1 | 640.85 | 13.91 | 102805 | 10 | 3345640.5 | 4915514 | 4754303 | 4210843.5 | 4258991 | 3702200.5 | 3019504 | 3578559.75 | 5567374.5 | 2735761 | 5710911 | 7730284.5 | 5326105 | 6046377 | 6065336 | 2426557.75 | 4138239.75 | 4672820 | 3345446.25 | 2955711.5 | 0 | 4 | 1 | 11 | 1 | 11 | 1.437216376 | 1.659858623 | 0.816315424 | 0.942772584 |
| Q60875 | Arhgef2 | Rho guanine nucleotide exchange factor 2 | 888.3 | 18.78 | 111974 | 16 | 8164554.5 | 8601680 | 14205391 | 13209679 | 11703391 | 10065359 | 9392565 | 11762017 | 11664070 | 11778548 | 7907556 | 13050376 | 12506349 | 11941354 | 8978035 | 8846432 | 7823622.5 | 7519660 | 11032365 | 10973491 | 11 | 6 | 10 | 4 | 11 | 6 | 0.97314067 | 0.994897989 | 0.826622926 | 0.845104425 |
| Q8K4I3 | Arhgef6 | Rho guanine nucleotide exchange factor 6 | 257.54 | 8.43 | 87051 | 5 | 2943310.5 | 3567730.5 | 4848394.5 | 3670983.75 | 5111451.5 | 2117899.5 | 3047522.5 | 4456591 | 4791731.5 | 4316519.5 | 1052793 | 2591722.25 | 2837004 | 2 | 2460382.75 | 3985271.75 | 1039048.75 | 3321717 | 2399335.25 | 3607487 | 0 | 6 | 3 | 7 | 3 | 7 | 0.443964052 | 0.477404056 | 0.712588216 | 0.766292443 |
| O55043 | Arhgef7 | Rho guanine nucleotide exchange factor 7 | 156.7 | 7.59 | 73140 | 3 | 2288450.25 | 1883114 | 1764299.25 | 1128315.12 | 1674032.75 | 1586991.5 | 2364424.5 | 1161651.12 | 2009553.25 | 1182021.62 | 1058506.62 | 1492338.38 | 1123380.25 | 424721.34 | 507937.38 | 2041622.75 | 1016375.75 | 1334076.5 | 1203312.25 | 1141759.5 | 1 | 7 | 2 | 8 | 2 | 8 | 0.527211322 | 0.554736011 | 0.770998373 | 0.811250715 |
| E9Q4N7 | Arid1b | AT-rich interactive domain-containing protein 1B | 98.24 | 1.43 | 237795 | 2 | 68764.05 | 2 | 83861.38 | 2 | 157884.11 | 2 | 176447.83 | 2 | 170243.98 | 2 | 2 | 143964.7 | 2 | 2 | 2 | 2 | 2 | 2 | 2 | 2 | 8 | 5 | 9 | 7.5 | 9 | 7.5 | 0.463659974 | 0.415268559 | 3.22047E-05 | 2.88435E-05 |
| E9Q7E2 | Arid2 | AT-rich interactive domain-containing protein 2 | 91.58 | 1.09 | 195988 | 2 | 611723.75 | 757591.56 | 2 | 765499.31 | 493201.84 | 829432.44 | 2 | 2 | 2 | 590827.19 | 2 | 251938.22 | 2 | 592906.56 | 537879.38 | 781196.19 | 719149.69 | 610521.75 | 694043.06 | 2 | 6 | 11.5 | 12 | 7.5 | 12 | 11.5 | 0.526148572 | 0.973570106 | 1.067310878 | 1.974921191 |
| Q9Z1K5 | Arih1 | E3 ubiquitin-protein ligase ARIH1 | 57.81 | 2.52 | 64017 | 1 | 2 | 2 | 560502.06 | 2 | 864848.25 | 416804.66 | 2 | 2 | 1080925.5 | 555089.5 | 2 | 533249.75 | 2 | 2 | 2 | 2 | 2 | 2 | 2 | 2 | 9 | 7.5 | 7 | 5 | 9 | 7.5 | 0.374122419 | 0.259767929 | 7.01579E-06 | 4.871339E-06 |
| P61211 | Arl1 | ADP-ribosylation factor-like protein 1 | 194.61 | 18.78 | 20412 | 3 | 6794352.5 | 6102207.5 | 7031237.5 | 5798666.5 | 6511398 | 4414784.5 | 6193669 | 7026158 | 6993739 | 6451733 | 6944067 | 6961261 | 5691127 | 5466870.5 | 6217114 | 6091723.5 | 4823174.5 | 4488253 | 2653780.5 | 4847360 | 10 | 1 | 10 | 4 | 10 | 4 | 0.970301303 | 1.006446443 | 0.710477993 | 0.736944336 |
| O08697 | Arl2 | ADP-ribosylation factor-like protein 2 | 256.46 | 33.15 | 20836 | 5 | 1036368.12 | 870003.44 | 1055617.75 | 992804.69 | 785456.81 | 565084.88 | 764627.31 | 916559.69 | 582378.62 | 903951.94 | 543298.25 | 620365.69 | 943211.81 | 790171.12 | 1063265.25 | 1107684.88 | 762324.25 | 775670.25 | 1031311.25 | 832068.62 | 8 | 9 | 10 | 7 | 10 | 9 | 0.835464679 | 1.061005608 | 0.95122799 | 1.208020228 |
| P37996 | Arl3 | ADP-ribosylation factor-like protein 3 | 296.39 | 25.82 | 20457 | 5 | 5534767 | 5016219 | 4269733.5 | 4695912 | 6033863.5 | 5345713.5 | 5801201 | 2727162.25 | 6148352 | 3722000 | 2419817.5 | 3056317.25 | 3958308.5 | 3260802 | 3845903.5 | 5409259 | 4870008 | 5604683 | 3883390.5 | 3318504 | 0 | 9 | 6 | 11 | 6 | 11 | 0.64739054 | 0.696632836 | 0.903538053 | 0.97226363 |
| P61208 | Arl4c | ADP-ribosylation factor-like protein 4C | 74.88 | 7.81 | 21487 | 2 | 2 | 1669726.12 | 2 | 888285.94 | 910538.19 | 1145946.38 | 1781947.88 | 813575.12 | 607804.5 | 723885.19 | 2 | 2 | 2 | 2 | 2 | 1370145.88 | 2 | 1124687.75 | 2 | 565863.5 | 5 | [illegible] | [illegible] | [illegible] | [illegible] | [illegible] | 2.883046E-06 | 1.971158E-06 | 0.882414087 | 0.603312667 |
| Q8R5J9 | Arl6ip5 | PRA1 family protein 3 | 251.71 | 28.19 | 21558 | 5 | 1115544.5 | 974755.25 | 2 | 2019444.5 | 2340851.25 | 1518357.88 | 795948.69 | 1839366.75 | 2224351.25 | 3059323 | 3336133 | 2768283.5 | 3202625 | 1664469.25 | 3993938.5 | 2447230.5 | 2691041.5 | 2512065.75 | 2465559 | 2736055 | 2 | 5 | 4 | 5 | 4 | 5 | 2.320009774 | 1.585768579 | 1.992366095 | 1.361818207 |
| Q8VEH3 | Arl8a | ADP-ribosylation factor-like protein 8A | 438.18 | 66.13 | 21390 | 8 | 2962462.75 | 3060492.25 | 2941784.5 | 2678851.5 | 3414514.25 | 3322810.75 | 2497812 | 2800120 | 1414070.62 | 3261186.5 | 1505577.75 | 2298608 | 2472184.5 | 1871146.75 | 2522503.25 | 2763991.25 | 2817719.75 | 3156015.5 | 4203097 | 3787873.5 | 0 | 9 | 6 | 7 | 6 | 9 | 0.708589831 | 0.802498522 | 1.110943025 | 1.258175178 |
| Q66HA6 | Arl8b | ADP-ribosylation factor-like protein 8B | 746.43 | 69.89 | 21539 | 13 | 16140820 | 17599086 | 12992651 | 18322862 | 17161294 | 18104306 | 17872818 | 19239064 | 18055174 | 17643576 | 11088532 | 11897538 | 17079098 | 13560702 | 16783774 | 16316262 | 18449682 | 21577340 | 20074476 | 18802784 | 5 | 3 | 0 | 7 | 5 | 7 | 0.85639089 | 0.774456273 | 1.158165299 | 1.04735862 |
| Q9DBR3 | Armc8 | Armadillo repeat-containing protein 8 | 46.31 | 1.63 | 75364 | 1 | 528077.44 | 389603.75 | 615928.38 | 907055.25 | 737503.31 | 601487.5 | 605624.88 | 1071116.62 | 683141.75 | 837381.5 | 837866.94 | 913727.75 | 1533063.62 | 669610.56 | 1489477.12 | 590990.75 | 834918.19 | 849306.62 | 1134648.5 | 887343.5 | 3 | 6 | 5 | 9 | 5 | 9 | 1.712856516 | 1.433035279 | 1.352102024 | 1.131215535 |
| O88656 | Arpc1b | Actin-related protein 2/3 complex subunit 1B | 663.82 | 29.84 | 41057 | 12 | 84054194 | 100062050 | 120587282 | 130692810 | 108079218 | 86242450 | 83568514 | 115159970 | 102649298 | 123270234 | 94049354 | 83933418 | 97105130 | 89443178 | 102492242 | 91028954 | 68916754 | 114891538 | 113285978 | 119969130 | 5 | 10 | 9 | 12 | 9 | 12 | 0.859321192 | 0.914135912 | 0.934894588 | 0.994523069 |
| Q9CVB6 | Arpc2 | Actin-related protein 2/3 complex subunit 2 | 1257.46 | 67.33 | 34357 | 18 | 54058158 | 54716686 | 57175410 | 46980326 | 53451254 | 38431922 | 49896970 | 48045818 | 60570462 | 50567038 | 30559312 | 51280198 | 35105486 | 20360430 | 41254186 | 45367218 | 31678088 | 39573426 | 33185108 | 35796450 | 1 | 0 | 5 | 2 | 5 | 2 | 0.670317039 | 0.721417388 | 0.696747912 | 0.749863168 |
| Q9JM76 | Arpc3 | Actin-related protein 2/3 complex subunit 3 | 531.34 | 53.37 | 20525 | 8 | 59298366 | 76589202 | 69504898 | 92179162 | 62857198 | 54976134 | 71688482 | 86938546 | 78132810 | 64210686 | 40141946 | 41659018 | 49016626 | 36794566 | 36587090 | 87001578 | 79442010 | 90808738 | 57191478 | 80051026 | 0 | 9 | 0 | 6 | 0 | 9 | 0.566545268 | 0.573679346 | 1.094515204 | 1.108297609 |
| P59999 | Arpc4 | Actin-related protein 2/3 complex subunit 4 | 385.2 | 42.26 | 19667 | 7 | 82401818 | 70296882 | 81580490 | 109245266 | 73231418 | 92669594 | 67081818 | 77432890 | 72965082 | 95274882 | 99847890 | 64067702 | 72986490 | 52478698 | 71689370 | 75392914 | 123250978 | 106038794 | 106504402 | 92299194 | 6 | 8 | 6 | 4 | 8 | 6 | 0.86638287 | 0.89059827 | 1.215306883 | 1.249274709 |
| Q4KLF8 | Arpc5 | Actin-related protein 2/3 complex subunit 5 | 370.03 | 39.07 | 16320 | 6 | 88466962 | 74341330 | 89732802 | 98268490 | 96868810 | 76719690 | 77427354 | 78706650 | 80887434 | 82314210 | 39164490 | 41572970 | 41487010 | 42810470 | 44390294 | 78574538 | 89374226 | 83094882 | 91923106 | 86940682 | 0 | 8 | 0 | 3 | 0 | 8 | 0.467802862 | 0.528777708 | 0.960304182 | 1.085473147 |
| A1L108 | Arpc5l | Actin-related protein 2/3 complex subunit 5-like protein | 160.88 | 21.57 | 17010 | 3 | 3699598 | 4082532.5 | 4234978.5 | 3033092.5 | 6161086 | 2795035.75 | 5528082 | 3737905 | 3934016 | 2661151.5 | 1114495.12 | 2224783.25 | 1138557.88 | 1863533.12 | 1875632 | 4405968 | 2909724.5 | 3047416.25 | 3344755 | 2055927.75 | 0 | 6 | 0 | 10 | 0 | 10 | 0.387388147 | 0.440443695 | 0.74317938 | 0.844963055 |
| P29066 | Arrb1 | Beta-arrestin-1 | 120.39 | 6.7 | 47020 | 3 | 728971 | 702386.69 | 533680.19 | 327714.69 | 610151.88 | 328667.5 | 659856.88 | 543208.44 | 761962.44 | 349985.66 | 2 | 237990.5 | 297565.94 | 66171.43 | 175562.75 | 412344.41 | 167970.64 | 305472.34 | 297133.78 | 305457.84 | 0 | 1 | 0 | 2 | 0 | 2 | 0.267763763 | 0.294019075 | 0.512720634 | 0.562994951 |
| P29067 | Arrb2 | Beta-arrestin-2 | 500.44 | 20.49 | 46340 | 11 | 9895255 | 7191552 | 6662854.5 | 4177038.75 | 6848940.5 | 7203335 | 9796674 | 6731424 | 6953158.5 | 7675919 | 14506648 | 11737890 | 13662688 | 11407377 | 9561296 | 10253096 | 8429972 | 5677624.5 | 6115687.5 | 4089589.25 | 1 | 11 | 1 | 9 | 1 | 11 | 1.750532778 | 1.586941837 | 0.993970737 | 0.901082097 |
| P50428 | Arsa | Arylsulfatase A | 136.2 | 7.31 | 53748 | 2 | 1074395 | 464228 | 495454.47 | 1127842.12 | 1309707.62 | 741076.44 | 617301.31 | 563911.44 | 975495.31 | 730591.12 | 271358.75 | 2 | 2 | 2 | 663678.12 | 1377756 | 1107358 | 1153772.62 | 592369.38 | 2 | 0 | 9 | 0 | 7 | 0 | 9 | 0.060686354 | 0.074790148 | 1.094665026 | 1.349070392 |
| P50429 | Arsb | Arylsulfatase B | 499.22 | 18.16 | 59647 | 9 | 26900540 | 25801130 | 28698422 | 28841544 | 41645642 | 16307439 | 25303292 | 17455828 | 23083110 | 14600540 | 982859.5 | 2694667.25 | 2482796.75 | 1703945.5 | 3113993.75 | 16842870 | 12119315 | 10708555 | 11000842 | 12631576 | 0 | 0 | 0 | 2 | 0 | 2 | 0.072279014 | 0.113470171 | 0.416777223 | 0.654294793 |
| Q91WU5 | As3mt | Arsenite methyltransferase | 94.26 | 3.19 | 41794 | 1 | 733910.88 | 536011.69 | 2 | 2 | 1310471.75 | 2 | 978502.62 | 817006.69 | 984139.19 | 2 | 2 | 2 | 2 | 2 | 2 | 2 | 2 | 2 | 2 | 2 | 2 | 5 | 5 | 2 | 5 | 5 | 3.875371E-06 | 3.597572E-06 | 3.875371E-06 | 3.597572E-06 |
| Q9WV54 | Asah1 | Acid ceramidase | 914.02 | 39.59 | 44670 | 15 | 10032551 | 9483134 | 12095022 | 11631176 | 10700058 | 9734821 | 9004514 | 8800062 | 10457309 | 10576928 | 7946214.5 | 7091462 | 9203370 | 7335724 | 7416714.5 | 6994265 | 9057073 | 9850942 | 10922684 | 8035628 | 0 | 4 | 2 | 10 | 2 | 10 | 0.722878789 | 0.802770594 | 0.831645862 | 0.923558489 |
| Q1AAU6 | Asap1 | Arf-GAP with SH3 domain, ANK repeat and PH domain-containing protein 1 | 354.01 | 6.64 | 127088 | 7 | 4235975 | 5285425 | 3879784.25 | 3843571 | 4061034.25 | 4370210 | 4550113 | 3624084.5 | 4885006 | 4267462 | 1040991.5 | 1979642.75 | 1939111.75 | 1411923.12 | 1728913.25 | 3362983.5 | 1381564.88 | 3573960.5 | 4817299.5 | 4555063 | 0 | 8 | 0 | 8 | 0 | 8 | 0.380205689 | 0.373352484 | 0.830331651 | 0.81536493 |
| E9PZJ8 | Ascc3 | Activating signal cointegrator 1 complex subunit 3 | 38.22 | 0.5 | 250557 | 1 | 490778.41 | 599253 | 390407.34 | 660616.94 | 526985.62 | 583743.81 | 527209.38 | 678654.75 | 508565.44 | 531420.56 | 2 | 2 | 334125.56 | 2 | 466349.5 | 578242.06 | 422132.5 | 618223 | 267297.72 | 488781.44 | 1 | 9 | 0 | 7 | 1 | 9 | 0.300025737 | 0.282896089 | 0.89004496 | 0.839228798 |
| Q91X20 | Ash2l | Set1/Ash2 histone methyltransferase complex subunit ASH2 | 44.43 | 2.41 | 68250 | 1 | 152892.33 | 135582.83 | 181732.14 | 2 | 211452.44 | 187268.86 | 2 | 2 | 179589.27 | 2 | 2 | 209529.39 | 2 | 2 | 243709.59 | 2 | 2 | 2 | 2 | 2 | 6 | 7 | 11 | 11 | 11 | 11 | 0.307392036 | 0.57115802 | 0.357534501 | 0.664326572 |
| Q91Y10 | Asl | Argininosuccinate lyase | 458.78 | 20.26 | 51739 | 8 | 4723808 | 6482170 | 4969352 | 4408539 | 5879507.5 | 3880154 | 2046003.88 | 4669120.5 | 4156282 | 3091497 | 5170706 | 4287915.5 | 3825745.75 | 3732280.25 | 3204946.25 | 5672208 | 2908145 | 4461624.5 | 5622899 | 5447080.5 | 3 | 10 | 10 | 5 | 10 | 10 | 0.764135059 | 1.133003185 | 0.911144389 | 1.351335508 |
| O54984 | Asna1 | ATPase Asna1 | 287.09 | 16.67 | 38823 | 5 | 9940088 | 10366003 | 13467372 | 12293315 | 13029428 | 4553495 | 8612154 | 15124021 | 14339317 | 13279631 | 7705175.5 | 10918571 | 12034798 | 8017750.5 | 13136153 | 9746814 | 5457550 | 9887338 | 10989422 | 11502764 | 8 | 4 | 8 | 9 | 8 | 9 | 0.876747451 | 0.926734551 | 0.805193619 | 0.851101131 |
| Q61024 | Asns | Asparagine synthetase [glutamine-hydrolyzing] | 917.48 | 31.91 | 64283 | 16 | 21166886 | 22116378 | 17689506 | 21294528 | 23271476 | 20062274 | 24133718 | 22337614 | 28311134 | 22627548 | 24710390 | 38047358 | 38547094 | 28788770 | 33352748 | 30047160 | 26982828 | 26724758 | 28907746 | 27183986 | 0 | 0 | 1 | 3 | 1 | 3 | 1.548685415 | 1.391360999 | 1.325072035 | 1.190463559 |
| Q8BSY0 | Asph | Aspartyl/asparaginyl beta-hydroxylase | 1202.3 | 36.03 | 83042 | 23 | 12274598 | 12373160 | 26326034 | 19457568 | 20270126 | 11325267 | 10839667 | 19933972 | 18438630 | 21103562 | 17865252 | 17298840 | 24976056 | 21928000 | 27116356 | 7707458.5 | 6910713 | 16176401 | 14925726 | 13158362 | 8 | 6 | 6 | 6 | 6 | 8 | 1.203778558 | 1.337371822 | 0.649147694 | 0.721188984 |
| Q8C0M9 | Asrgl1 | Isoaspartyl peptidase/L-asparaginase | 200.55 | 10.12 | 33950 | 3 | 3246340.75 | 3407937 | 5489696.5 | 6247459.5 | 5462548.5 | 3874943.5 | 3763051 | 5260210 | 5187130.5 | 6155852 | 1714815.25 | 1842438 | 2685574 | 3444235 | 1311502 | 4174999.75 | 5568604 | 5833915 | 7343994 | 6052836.5 | 2 | 6 | 0 | 6 | 0 | 6 | 0.461078747 | 0.453713931 | 1.214654599 | 1.195252908 |
| P16460 | Ass1 | Argininosuccinate synthase | 929.12 | 37.86 | 46584 | 18 | 49066770 | 49939930 | 23573706 | 32403652 | 38902294 | 56802106 | 55379790 | 35862566 | 46083522 | 39714214 | 10088794 | 93338914 | 56267630 | 48249802 | 55432214 | 53631026 | 67167874 | 37881874 | 40750426 | 40833510 | 2 | 7 | 4 | 12 | 4 | 12 | 1.826731744 | 1.514604109 | 1.23920383 | 1.027465154 |
| Q505J9 | Atad1 | ATPase family AAA domain-containing protein 1 | 50.26 | 3.05 | 40717 | 1 | 608261.25 | 833073.69 | 2 | 2 | 2237790 | 127215.09 | 772673.06 | 2 | 1379066.75 | 2 | 2 | 280925.53 | 174573.78 | 2 | 2 | 1043235.06 | 181624.34 | 2 | 2 | 2 | 7 | 9 | 9 | 10 | 9 | 10 | 0.123807922 | 0.199874298 | 0.332922662 | 0.537467086 |
| Q3KRE0 | Atad3 | ATPase family AAA domain-containing protein 3 | 251.39 | 7.45 | 66759 | 4 | 621306 | 393168.62 | 296415.69 | 397205.06 | 219772.64 | 560855.69 | 268728.28 | 325972.03 | 284027 | 536972.81 | 2887723.75 | 2205978.5 | 3028997.75 | 1823179.25 | 2942015.75 | 377643.38 | 696432 | 580788.56 | 383538.69 | 409622.25 | 0 | 8 | 0 | 6 | 0 | 8 | 6.685050498 | 6.520380014 | 1.269809379 | 1.238530614 |
| Q60765 | Atf3 | Cyclic AMP-dependent transcription factor ATF-3 | 40.53 | 5.52 | 20722 | 1 | 88870.71 | 145690.42 | 2 | 94705.5 | 2 | 158964.55 | 183596.73 | 171215.12 | 89099.57 | 104865.56 | 151170.16 | 2 | 79398.7 | 2 | 2 | 156241.44 | 184022.81 | 107896.02 | 151095.25 | 2 | 10 | 5 | 2 | 11 | 10 | 11 | 0.700259419 | 0.325789642 | 1.819954364 | 0.846718038 |
| Q6AZ50 | Atg3 | Ubiquitin-like-conjugating enzyme ATG3 | 41.79 | 3.18 | 35822 | 1 | 442407.81 | 447700 | 426826.69 | 359552.88 | 344191.47 | 568263.88 | 444223.62 | 421564.59 | 468614.59 | 336222.75 | 645811.62 | 523910.41 | 408196.34 | 348740.41 | 261868.34 | 2 | 459773.09 | 2 | 461212 | 629515.75 | 12 | 10 | 11 | 11 | 12 | 12 | 1.083065288 | 0.977505673 | 0.767318785 | 0.692533012 |
| Q8BGE6 | Atg4b | Cysteine protease ATG4B | 96.76 | 6.11 | 44375 | 2 | 2140978 | 1726990.88 | 1140127.25 | 300654.75 | 984646.5 | 1807843 | 1563572.25 | 848387 | 1114521 | 833993.12 | 801360.5 | 1847952 | 340913.56 | 367458.25 | 316536.62 | 599424.62 | 645674.69 | 908815.5 | 558013.75 | 508966.06 | 8 | 5 | 5 | 2 | 8 | 5 | 0.583821537 | 0.59566026 | 0.511789488 | 0.522167546 |
| Q9D906 | Atg7 | Ubiquitin-like modifier-activating enzyme ATG7 | 694.5 | 18.62 | 77520 | 13 | 10885618 | 8432584 | 12354986 | 13137642 | 16990018 | 10910292 | 10997926 | 11833645 | 17793606 | 13551501 | 11895630 | 12544265 | 14007175 | 11398319 | 13759504 | 13168181 | 9820691 | 13443320 | 14132068 | 14241464 | 10 | 8 | 9 | 11 | 10 | 11 | 1.029191266 | 0.977229283 | 1.048621922 | 0.99567892 |
| Q5FWU3 | Atg9a | Autophagy-related protein 9A | 55.15 | 1.67 | 94488 | 1 | 257398.22 | 267443.38 | 182729.61 | 247786.98 | 388394.81 | 293179.34 | 271979.97 | 208552.42 | 220052.34 | 347156.25 | 256599.89 | 107884.19 | 257187.58 | 342632.78 | 115474.73 | 145678.62 | 564987.38 | 385727.91 | 409291.94 | 248916.58 | 8 | 9 | 8 | 8 | 8 | 9 | 0.803554798 | 0.805252299 | 1.30574773 | 1.308506109 |
| Q9CWJ9 | Atic | Bifunctional purine biosynthesis protein PURH | 1627.75 | 56.93 | 64217 | 27 | 92847410 | 91252874 | 95929274 | 97690610 | 81843730 | 100448026 | 90456354 | 87684738 | 79198474 | 92849434 | 87779410 | 97543322 | 79108082 | 72752042 | 77750570 | 98725042 | 97804050 | 99671570 | 90245378 | 93412826 | 5 | 6 | 6 | 7 | 6 | 7 | 0.902885165 | 0.920770825 | 1.044161363 | 1.064845626 |
| Q6PA06 | Atl2 | Atlastin-2 | 185.18 | 7.38 | 66224 | 4 | 1120680.25 | 1022491.31 | 867998 | 525422.56 | 1905865.25 | 587030.5 | 1429234.12 | 989938.5 | 2063375 | 623368.5 | 2 | 711133.06 | 2 | 2 | 611004 | 1257159.75 | 2 | 2 | 2 | 519381.25 | 2 | 4 | 3 | 3 | 3 | 4 | 0.242931266 | 0.232242308 | 0.326423687 | 0.312061068 |
| Q91YH5 | Atl3 | Atlastin-3 | 673.34 | 28.1 | 60575 | 12 | 4966677.5 | 4648851 | 6676334 | 5626472 | 7383152 | 3783483.5 | 5456209.5 | 5770195.5 | 8741570 | 5659593 | 3735074.25 | 4406012 | 5914866 | 5399062 | 3453327.25 | 6139218 | 4599498 | 8162057 | 6097698 | 6523233.5 | 5 | 11 | 6 | 8 | 6 | 11 | 0.781868361 | 0.778902499 | 1.075844947 | 1.071763942 |
| O08997 | Atox1 | Copper transport protein ATOX1 | 306.47 | 38.24 | 7338 | 4 | 12109069 | 14876994 | 28524254 | 25702970 | 25410484 | 11967905 | 12094744 | 14922848 | 20397342 | 19704474 | 10027890 | 9524634 | 11068398 | 11784971 | 11218188 | 10836040 | 8510422 | 18714182 | 16028714 | 18859590 | 0 | 6 | 0 | 9 | 0 | 9 | 0.50292801 | 0.678036451 | 0.684171525 | 0.92238496 |
| P98197 | Atp11a | Probable phospholipid-transporting ATPase IH | 86.74 | 2.53 | 135502 | 2 | 555895.75 | 967971.38 | 1084637.38 | 1087393 | 1401803.62 | 1091235.88 | 1108126 | 1432006.75 | 1608937.25 | 1368263 | 220010.58 | 682504.88 | 613198 | 367966.72 | 804873.88 | 977776.06 | 1221115.75 | 1043514.62 | 595918.12 | 1504533 | 3 | 11 | 0 | 6 | 3 | 11 | 0.527405195 | 0.406828484 | 1.048091564 | 0.808474217 |
| Q9EPE9 | Atp13a1 | Manganese-transporting ATPase 13A1 | 383.72 | 8.17 | 132388 | 7 | 2567569.75 | 2897184 | 2933726.25 | 3657757.25 | 2806586 | 2898591 | 2172956 | 2380934.5 | 2008960.62 | 2438730.5 | 891656.38 | 2156547.5 | 1976587.62 | 1429738 | 3069301 | 3243403.75 | 4154896.25 | 2524941.25 | 3046438.5 | 3304947 | 4 | 8 | 6 | 1 | 6 | 8 | 0.64078206 | 0.800310282 | 1.094988918 | 1.367595855 |
| P06685 | Atp1a1 | Sodium/potassium-transporting ATPase subunit alpha-1 | 2677.09 | 35.29 | 113054 | 36 | 111862914 | 111305074 | 116626002 | 134508642 | 114016554 | 123949986 | 113490122 | 122476922 | 114040162 | 130560122 | 104652986 | 105464578 | 121282482 | 117163522 | 117825202 | 112306762 | 116826650 | 123740698 | 129453002 | 131084714 | 12 | 7 | 6 | 11 | 12 | 11 | 0.962723609 | 0.936927292 | 1.042651405 | 1.014713412 |
| P06687 | Atp1a3 | Sodium/potassium-transporting ATPase subunit alpha-3 | 1243.26 | 17.67 | 111692 | 16 | 2626267.5 | 2830829.5 | 1191698.5 | 2254248.5 | 1256029.62 | 3494138 | 2375794 | 1511041.88 | 1488216.38 | 1762125.88 | 1802983.25 | 1869962.88 | 1416129.75 | 1456727.25 | 1465381.88 | 2328929 | 3741172.5 | 1981865.5 | 2603430.5 | 2189998.75 | 10 | 10 | 6 | 7 | 10 | 10 | 0.788574363 | 0.75354593 | 1.264425944 | 1.208260208 |
| Q9WV27 | Atp1a4 | Sodium/potassium-transporting ATPase subunit alpha-4 | 514.11 | 7.46 | 114887 | 6 | 1098288.12 | 961781.5 | 774267.44 | 1376343.88 | 2 | 1601339.38 | 1131727.88 | 899116.38 | 2 | 1189128.38 | 1377594 | 1483358.62 | 1676622.88 | 2496708.25 | 1491057 | 1135421.38 | 1205485.88 | 1166380.5 | 1166908.62 | 1304423.62 | 0 | 5 | 3 | 8 | 3 | 8 | 2.024693113 | 1.768260834 | 1.419869433 | 1.240039536 |
| P97370 | Atp1b3 | Sodium/potassium-transporting ATPase subunit beta-3 | 353.62 | 41.37 | 31776 | 7 | 18224662 | 14946475 | 16402326 | 18606252 | 14984770 | 18422382 | 15571598 | 14491956 | 14943833 | 18553568 | 20196828 | 18120950 | 21763448 | 23313734 | 23312618 | 13911182 | 18365040 | 16030410 | 13427631 | 16457997 | 2 | 9 | 2 | 9 | 2 | 9 | 1.283090709 | 1.301576417 | 0.940212159 | 0.953757967 |
| O55143 | Atp2a2 | Sarcoplasmic/endoplasmic reticulum calcium ATPase 2 | 2364.48 | 35.92 | 114858 | 36 | 40242550 | 39812954 | 27019330 | 30278532 | 30126208 | 37049030 | 41430338 | 33271638 | 32640502 | 34645902 | 51009886 | 48171486 | 46003126 | 44210522 | 49306630 | 42423274 | 40996514 | 37055514 | 43775230 | 37127102 | 0 | 4 | 0 | 3 | 0 | 4 | 1.425258283 | 1.333250129 | 1.20240116 | 1.124779642 |
| Q64518 | Atp2a3 | Sarcoplasmic/endoplasmic reticulum calcium ATPase 3 | 929.76 | 21.48 | 113638 | 15 | 2471695 | 3325290 | 6237073 | 6447011.5 | 5239079 | 2757748.5 | 3332570 | 7296791 | 5232757 | 4921322 | 2086031.88 | 1312184.12 | 2259472 | 2117376.5 | 2715179 | 2068664.12 | 1188284.25 | 3657723.5 | 4233920.5 | 3033257.25 | 1 | 5 | 0 | 5 | 1 | 5 | 0.44225033 | 0.445612315 | 0.597881991 | 0.602427087 |
| G5E829 | Atp2b1 | Plasma membrane calcium-transporting ATPase 1 | 1228.97 | 19.43 | 134747 | 20 | 6381927 | 5999291 | 7077566 | 8204443 | 8799870 | 7628784 | 7393511 | 3440140.5 | 10139064 | 10439477 | 6799776 | 9600910 | 12645447 | 12523747 | 11307893 | 6208636.5 | 6845114.5 | 9504923 | 10689728 | 9715169 | 3 | 7 | 6 | 12 | 6 | 12 | 1.450172293 | 1.354417275 | 1.178275422 | 1.10047378 |
| Q64566 | Atp2c1 | Calcium-transporting ATPase type 2C member 1 | 60.89 | 1.2 | 100500 | 1 | 2 | 2 | 2 | 1071339 | 2 | 710277.25 | 2 | 2 | 2 | 1393763 | 724516.81 | 828358.25 | 833204.12 | 1112169.38 | 1115314.38 | 2 | 2 | 2 | 2 | 1073329.75 | 3 | 12 | 5 | 10 | 5 | 12 | 4.306319932 | 2.192709851 | 1.001858175 | 0.510130303 |
| P15999 | Atp5f1a | ATP synthase subunit alpha, mitochondrial | 2944.79 | 63.47 | 59754 | 42 | 7766863.5 | 7417389.5 | 7649394 | 7779600.5 | 6668269.5 | 8189972 | 5242943 | 7285566.5 | 6824187 | 7559088 | 4754507.5 | 5047347.5 | 5240527.5 | 6344206.5 | 5071641.5 | 8549246 | 9591793 | 7875812.5 | 9648108 | 9336178 | 0 | 0 | 1 | 1 | 1 | 1 | 0.709687605 | 0.753758021 | 1.20706294 | 1.282019534 |
| Q03265 | Atp5f1a | ATP synthase subunit alpha, mitochondrial | 3289.939 | 70.71 | 59753 | 47 | 37606202 | 42817578 | 42911478 | 16263805 | 36850714 | 18364842 | 47189854 | 36824338 | 39411474 | 29847400 | 14143793 | 53124714 | 27040114 | 17708120 | 40014466 | 40779622 | 12741386 | 17030938 | 20059026 | 24300356 | 10 | 6 | 10 | 6 | 10 | 6 | 0.861611783 | 0.885767071 | 0.651240993 | 0.669498535 |
| P56480 | Atp5f1b | ATP synthase subunit beta, mitochondrial | 3291.26 | 82.61 | 56300 | 46 | 328098754 | 304828194 | 362188098 | 333184738 | 364119714 | 327507426 | 292508834 | 402535170 | 355526402 | 369485410 | 383475714 | 395858978 | 514923746 | 489951106 | 518279810 | 318385090 | 328095682 | 378541026 | 337576258 | 373592930 | 0 | 10 | 2 | 12 | 2 | 12 | 1.360472009 | 1.317542793 | 1.025863261 | 0.993492507 |
| Q91VR2 | Atp5f1c | ATP synthase subunit gamma, mitochondrial | 681.95 | 41.61 | 32886 | 11 | 21448328 | 21415392 | 27669524 | 27411542 | 24160046 | 26044114 | 20811692 | 33378534 | 24969740 | 34403098 | 40006862 | 30622354 | 40935542 | 44161346 | 46866194 | 22689466 | 24426624 | 34245358 | 34104162 | 29499408 | 0 | 5 | 2 | 12 | 2 | 12 | 1.65916692 | 1.45115961 | 1.187217702 | 1.038377969 |
| P35434 | Atp5f1d | ATP synthase subunit delta, mitochondrial | 130.32 | 13.69 | 17595 | 2 | 22896798 | 23222358 | 26522842 | 27974298 | 22762096 | 23202274 | 22157522 | 25487514 | 24270808 | 26327454 | 29590250 | 24443698 | 30642502 | 31920642 | 31453250 | 21308748 | 26389202 | 27720842 | 26426546 | 24269120 | 2 | 12 | 2 | 8 | 2 | 12 | 1.199969781 | 1.219067435 | 1.022176217 | 1.038444267 |
| P29418 | Atp5f1e | ATP synthase subunit epsilon, mitochondrial | 49.29 | 15.69 | 5767 | 1 | 6547974.5 | 6319299 | 6488471 | 6383379.5 | 5801830 | 7712574.5 | 7063969 | 6951554 | 5727816 | 7624739 | 7462482 | 6368167.5 | 8342842.5 | 9833746 | 7747806.5 | 7195991 | 8685536 | 8201833 | 9197844 | 8701264 | 3 | 0 | 6 | 2 | 6 | 2 | 1.260426191 | 1.133247008 | 1.331046233 | 1.196741366 |
| O35143 | ATP5IF1 | ATPase inhibitor, mitochondrial | 52.12 | 8.49 | 12159 | 1 | 5911805.5 | 5588341.5 | 5616483 | 7182935 | 4234496.5 | 5678476 | 4823111.5 | 5102285.5 | 4403061.5 | 6154048.5 | 3594067.5 | 2 | 2936256 | 3131063.5 | 3892749.25 | 6122254 | 7265000.5 | 5627148 | 5263701.5 | 6616508 | 0 | 8 | 0 | 5 | 0 | 8 | 0.475016087 | 0.518105082 | 1.082727462 | 1.180942322 |
| P56383 | Atp5mc2 | ATP synthase F(0) complex subunit C2, mitochondrial | 82.35 | 21.23 | 15476 | 1 | 383770.97 | 148325.48 | 158194.17 | 194948.06 | 195426.11 | 418742.56 | 259768.89 | 125975.42 | 204331.09 | 176509.06 | 308909 | 217009.56 | 268430.66 | 227928.36 | 210895.39 | 234898.67 | 456898.81 | 133383.91 | 309563.62 | 292268.03 | 5 | 8 | 8 | 8 | 8 | 8 | 1.141124409 | 1.04036519 | 1.320495544 | 1.203898178 |
| Q78IK2 | Atp5md | ATP synthase F(0) complex subunit k, mitochondrial | 228.84 | 44.83 | 6382 | 3 | 7492585 | 6021504 | 8170398 | 8883460 | 7988478 | 7433931 | 6588542.5 | 8779794 | 6479979 | 9199603 | 5081172.5 | 5363472 | 6653119 | 6803642.5 | 6982287 | 9853499 | 11115798 | 9575196 | 10097592 | 9661366 | 0 | 0 | 6 | 0 | 6 | 0 | 0.800999911 | 0.802552201 | 1.304671037 | 1.307199411 |
| Q06185 | Atp5me | ATP synthase subunit e, mitochondrial | 181.84 | 46.48 | 8236 | 4 | 5482912 | 5246346 | 5144053.5 | 5716730 | 3385503.25 | 4677589 | 4356689 | 4690409.5 | 3650966 | 4787940.5 | 5304771 | 5246037 | 5484720.5 | 5816973 | 5497747 | 5538550 | 10016669 | 5085693 | 4537153 | 5408258.5 | 7 | 12 | 0 | 3 | 7 | 12 | 1.095081159 | 1.234016852 | 1.216643072 | 1.37100163 |
| P56135 | Atp5mf | ATP synthase subunit f, mitochondrial | 164.8 | 36.36 | 10344 | 4 | 2061785.62 | 2013454.38 | 2658419 | 2689834.25 | 3139336 | 1995790.38 | 1716912.62 | 3085545 | 2602798 | 3598187.75 | 3438753.75 | 2901202 | 4373426 | 4139347 | 3791171 | 2069861.25 | 1865661 | 2614732.75 | 2590346.5 | 2792263 | 1 | 10 | 3 | 11 | 3 | 11 | 1.484052627 | 1.434230671 | 0.949854866 | 0.917966761 |
| Q9CPQ8 | Atp5mg | ATP synthase subunit g, mitochondrial | 192.01 | 33.01 | 11425 | 3 | 5508447 | 3693086 | 4351526 | 5060654.5 | 3661665.25 | 5403110 | 3750691.25 | 4843292 | 3857783 | 5536140 | 4481579 | 3948701.25 | 5367709.5 | 6374492.5 | 6479766 | 4523345 | 6529227 | 5694536.5 | 6313536 | 5195752 | 6 | 3 | 8 | 5 | 8 | 5 | 1.196489117 | 1.139422416 | 1.268503527 | 1.208002089 |
| Q9CQQ7 | Atp5pb | ATP synthase F(0) complex subunit B1, mitochondrial | 474.73 | 34.38 | 28949 | 8 | 11709389 | 14118780 | 11899932 | 12754266 | 9931300 | 16283180 | 10441256 | 14817111 | 8848232 | 16445706 | 16346674 | 16710861 | 19404772 | 18597346 | 19045068 | 16827610 | 18605342 | 17558612 | 19664778 | 11344490 | 0 | 4 | 1 | 3 | 1 | 4 | 1.491462536 | 1.348156911 | 1.390427633 | 1.256829841 |
| Q9DCX2 | Atp5pd | ATP synthase subunit d, mitochondrial | 631.5 | 68.94 | 18749 | 11 | 25285858 | 21403144 | 20853424 | 20608170 | 22173828 | 19900648 | 20690038 | 24607854 | 24038264 | 22821170 | 21028826 | 23638348 | 27157614 | 24036338 | 27719710 | 24812284 | 18876036 | 21133460 | 20713560 | 22358062 | 5 | 11 | 7 | 11 | 7 | 11 | 1.120158452 | 1.102829469 | 0.977964789 | 0.962835559 |

| | | | | | | | | | | | | | | | | | | | | | | | | | | | | | | | | | | | | |
|---|---|---|---|---|---|---|---|---|---|---|---|---|---|---|---|---|---|---|---|---|---|---|---|---|---|---|---|---|---|---|---|---|---|---|---|---|
| P97450 | Atp5pf | ATP synthase-coupling factor 6, mitochondrial | 99.88 | 17.59 | 12496 | 2 | 2499949.25 | 2417097.75 | 2048014.25 | 1516469 | 2104595.5 | 1241372.5 | 2490039.75 | 2186265.75 | 2027119.25 | 1166291.25 | 294205.72 | 1922693.25 | 482718.25 | 2 | 1993024.25 | 2377239.25 | 248799.89 | 1020759.38 | 1093523.62 | 1476895 | 2 | 3 | 4 | 6 | 4 | 6 | 0.443282423 | 0.515047513 | 0.587298629 | 0.682379184 |
| Q9DB20 | Atp5po | ATP synthase subunit O, mitochondrial | 677.6 | 62.91 | 23364 | 11 | 16187478 | 16990650 | 17998474 | 17914718 | 17137546 | 20092152 | 14922055 | 16871868 | 14500164 | 17585718 | 25401152 | 20806226 | 25484574 | 24161616 | 26546278 | 16202130 | 22864852 | 20146762 | 18568622 | 20435636 | 0 | 4 | 0 | 4 | 0 | 4 | 1.419476466 | 1.457627646 | 1.139038544 | 1.169652411 |
| Q9R1Q9 | Atp6ap1 | V-type proton ATPase subunit S1 | 110.31 | 3.89 | 51008 | 1 | 2 | 2 | 2 | 2 | 2 | 2 | 2 | 2 | 2 | 2 | 2 | 334177.91 | 2 | 2 | 2 | 2 | 2 | 2 | 2 | 2 | 10 | 12.5 | 10 | 12.5 | 10 | 12.5 | 33418.591 | 33418.591 | 1 | 1 |
| Q9CYN9 | Atp6ap2 | Renin receptor | 111.91 | 8.86 | 39092 | 2 | 2 | 2 | 105774.09 | 2 | 2 | 2 | 2 | 2 | 2 | 2 | 338383.62 | 2 | 244025.97 | 2 | 233273.62 | 2 | 2 | 2 | 2 | 2 | 6 | 10 | 5 | 12.5 | 6 | 12.5 | 7.711014313 | 81568.721 | 9.453396E-05 | 1 |
| P25286 | Atp6v0a1 | V-type proton ATPase 116 kDa subunit a isoform 1 | 464.14 | 11.58 | 96328 | 9 | 2377868.75 | 2087131.12 | 2219328.75 | 3551273.25 | 4236371 | 4471018.5 | 3116367.75 | 3177261.5 | 4134850 | 5274171.5 | 3580279.5 | 2690205.5 | 6589345.5 | 6093909.5 | 5121981 | 2167966 | 1798925.12 | 4463713.5 | 6017014 | 4802765 | 3 | 9 | 9 | 12 | 9 | 12 | 1.663610153 | 1.193423006 | 1.330183783 | 0.954233133 |
| P15920 | Atp6v0a2 | V-type proton ATPase 116 kDa subunit a isoform 2 | 54.11 | 1.4 | 98145 | 1 | 2 | 2 | 2 | 2 | 222612.81 | 331589.44 | 211840.06 | 2 | 468985.56 | 429312.09 | 588741 | 697374.12 | 686893.5 | 842727.62 | 707264.38 | 2 | 2 | 450995.16 | 546834.69 | 404734.31 | 0 | 6 | 0 | 12 | 0 | 12 | 15.82511815 | 2.443593944 | 6.300256297 | 0.972837485 |
| P63081 | Atp6v0c | V-type proton ATPase 16 kDa proteolipid subunit | 611.04 | 76.13 | 15808 | 9 | 28849242 | 22735390 | 19705988 | 25861348 | 25707718 | 29197808 | 28271918 | 22172038 | 28414342 | 10499781 | 15228900 | 17450410 | 22643522 | 21652062 | 19005686 | 23545172 | 30670450 | 20456242 | 25209740 | 20585646 | 2 | 11 | 6 | 11 | 6 | 11 | 0.781221108 | 0.809580886 | 0.980527087 | 1.016122042 |
| P51863 | Atp6v0d1 | V-type proton ATPase subunit d 1 | 634.9 | 32.76 | 40301 | 12 | 36721762 | 35938706 | 28560606 | 29036440 | 35449746 | 31480444 | 35763878 | 33304542 | 38601294 | 31449688 | 39251934 | 40080814 | 36919914 | 34656498 | 43638650 | 34107174 | 36012474 | 34306930 | 34114938 | 34026902 | 3 | 12 | 3 | 9 | 3 | 12 | 1.174045181 | 1.140375062 | 1.041405295 | 1.011539119 |
| P50516 | Atp6v1a | V-type proton ATPase catalytic subunit A | 2661.5 | 70.34 | 68326 | 40 | 130367554 | 139879442 | 120649466 | 130742546 | 128159442 | 125779570 | 131879042 | 157385858 | 151241970 | 144571250 | 118775554 | 127295946 | 130197474 | 129379354 | 134654802 | 130936378 | 129538938 | 141135218 | 150386482 | 140851042 | 9 | 4 | 5 | 9 | 9 | 9 | 0.985387284 | 0.900747279 | 1.066250709 | 0.974664926 |
| P62814 | Atp6v1b2 | V-type proton ATPase subunit B, brain isoform | 1677.21 | 66.34 | 56551 | 24 | 98893530 | 92615258 | 79881018 | 90824026 | 76970986 | 101262626 | 95317466 | 91443338 | 85550690 | 91588106 | 76449586 | 75966738 | 80805858 | 79022210 | 80255802 | 91462866 | 108635034 | 92580802 | 102073394 | 99765602 | 5 | 4 | 0 | 6 | 5 | 6 | 0.893701644 | 0.843792062 | 1.125989965 | 1.063108031 |
| Q5FVI6 | Atp6v1c1 | V-type proton ATPase subunit C 1 | 1026.21 | 42.41 | 43901 | 18 | 45145594 | 41812854 | 32763114 | 35760402 | 39985270 | 37281942 | 45677598 | 40949690 | 40814930 | 39301102 | 20291176 | 23482254 | 22020416 | 18243908 | 21647282 | 39976774 | 34725002 | 34369574 | 41254678 | 41174810 | 0 | 10 | 0 | 10 | 0 | 10 | 0.540679038 | 0.517999756 | 0.979708128 | 0.938613366 |
| P57746 | Atp6v1d | V-type proton ATPase subunit D | 271.31 | 20.24 | 28369 | 4 | 2670233 | 2645036.5 | 2076778.25 | 2493996 | 2518211 | 2544734 | 1865017.38 | 3119141.5 | 3331391.25 | 3546643.75 | 4567239 | 2591064.5 | 3728945.75 | 3987166.5 | 3717347 | 2326573 | 2262997 | 2902511.5 | 1443653.75 | 1110404.5 | 2 | 7 | 3 | 4 | 3 | 7 | 1.498821423 | 1.290473785 | 0.809894665 | 0.697313114 |
| P50518 | Atp6v1e1 | V-type proton ATPase subunit E 1 | 572.29 | 41.59 | 26157 | 9 | 22508574 | 29355798 | 23267604 | 25364428 | 12792496 | 26282462 | 25380134 | 28162386 | 26300026 | 32069418 | 26373986 | 25458918 | 29988322 | 28647310 | 30899640 | 20990688 | 25037268 | 33016970 | 28422032 | 27341688 | 3 | 8 | 9 | 12 | 9 | 12 | 1.247855492 | 1.022965832 | 1.189954585 | 0.97549988 |
| P50408 | Atp6v1f | V-type proton ATPase subunit F | 210 | 31.09 | 13370 | 3 | 1006709.12 | 757537.75 | 859774.25 | 1442004.5 | 497129.31 | 678828.88 | 802496.31 | 864014.69 | 762578.75 | 994695.56 | 791504 | 490033.25 | 937209 | 1208967.25 | 426312.16 | 903588.5 | 2320353.75 | 1502832.5 | 1568711.25 | 1338131.75 | 9 | 3 | 11 | 1 | 11 | 3 | 0.844596714 | 0.939407286 | 1.672881563 | 1.86067161 |
| Q9CR51 | Atp6v1g1 | V-type proton ATPase subunit G 1 | 224.01 | 22.88 | 13724 | 3 | 4199630 | 4156320 | 3486056 | 4946154 | 3951064.5 | 3813235.75 | 3855577.75 | 4604022 | 4577084 | 5097580 | 2873990.5 | 3361185.25 | 3526574.5 | 4007931 | 3896498.25 | 4908546 | 3560899.25 | 4716149.5 | 5715410 | 5434745 | 4 | 6 | 4 | 7 | 4 | 7 | 0.851824498 | 0.804929031 | 1.173416574 | 1.108816508 |
| Q8BVE3 | Atp6v1h | V-type proton ATPase subunit H | 637.89 | 25.88 | 55855 | 12 | 7183891 | 10154310 | 8809174 | 7921626 | 12365190 | 7347120 | 7116445.5 | 11549696 | 11255460 | 8644788 | 13085035 | 14729671 | 13944304 | 11467183 | 14185712 | 6137229 | 4406123 | 7390519 | 11056899 | 8463675 | 1 | 7 | 1 | 7 | 1 | 7 | 1.451773005 | 1.468236816 | 0.806613493 | 0.815760882 |
| Q64430 | Atp7a | Copper-transporting ATPase 1 | 286.03 | 6.24 | 161959 | 6 | 1441512.75 | 676571.62 | 1256390.25 | 604031.25 | 644795.25 | 332241.38 | 561094.44 | 948117.88 | 1177116.75 | 1046671.38 | 1551131.25 | 1868661.38 | 2581115.25 | 1878435.5 | 2764956.25 | 1100947.62 | 1291759.25 | 1416808.5 | 1479432.25 | 957584.25 | 0 | 6 | 0 | 3 | 0 | 6 | 2.302315889 | 2.618368125 | 1.351097778 | 1.536570795 |
| P70704 | Atp8a1 | Phospholipid-transporting ATPase IA | 48.44 | 0.77 | 131413 | 1 | 202308.83 | 2 | 2 | 618402.94 | 605559.75 | 468359.81 | 318775 | 491680.84 | 482281.38 | 454228.19 | 143724.52 | 2 | 351835.72 | 504949.19 | 204382.48 | 249744.67 | 236458.97 | 616415.12 | 2 | 340899.97 | 12 | 11 | 6 | 6 | 12 | 11 | 0.844783419 | 0.543890305 | 1.012091079 | 0.651606688 |
| Q99J86 | Atrn | Attractin | 51.08 | 0.63 | 158673 | 1 | 2 | 2 | 2 | 2 | 2 | 2 | 2 | 2 | 2 | 2 | 404401.5 | 378675.28 | 392320.97 | 238304.66 | 532639.81 | 2 | 2 | 2 | 2 | 2 | 0 | 12.5 | 0 | 12.5 | 0 | 12.5 | 194634.222 | 194634.222 | 1 | 1 |
| P28658 | Atxn10 | Ataxin-10 | 676 | 26.32 | 53707 | 11 | 10426426 | 10817685 | 7097845.5 | 7425582.5 | 10273911 | 7688878.5 | 11413341 | 9564340 | 14758628 | 11413026 | 6840681.5 | 11575979 | 8763728 | 5658281 | 9473600 | 8346724.5 | 7641306.5 | 9284154 | 9014321 | 9307452 | 9 | 10 | 6 | 4 | 9 | 10 | 0.919003843 | 0.77158366 | 0.946841553 | 0.794955839 |
| Q7TQH0 | Atxn2l | Ataxin-2-like protein | 342.42 | 6.86 | 110649 | 6 | 2505619.75 | 3644259.5 | 1631564.38 | 2464883.75 | 2719130 | 4290617 | 4640534.5 | 3565687 | 2469923.75 | 4622601 | 5171647.5 | 6613557.5 | 4561293 | 6317220.5 | 3653368.5 | 3631709.75 | 3976522.5 | 4173733.5 | 3687258.5 | 5260763 | 0 | 1 | 5 | 12 | 5 | 12 | 2.029784699 | 1.343437592 | 1.598862781 | 1.058226701 |
| A1L134 | Aup1 | Ancient ubiquitous protein 1 | 86.96 | 7.07 | 46199 | 2 | 2 | 151616.39 | 644697 | 681394.5 | 435610.88 | 1462539.25 | 2 | 158950.23 | 62556.34 | 867775.75 | 1417933.88 | 1196494.38 | 1526791.25 | 1454361 | 1269774 | 2 | 2 | 2 | 2 | 213682.11 | 0 | 4 | 4 | 5 | 4 | 5 | 3.588187939 | 2.690371933 | 0.11168546 | 0.083740158 |
| O55099 | Aurkb | Aurora kinase B | 54.54 | 2.92 | 39234 | 1 | 313606.66 | 412414.94 | 498848.53 | 195868.14 | 453747.72 | 273312.66 | 227879.3 | 307634.72 | 504960.97 | 300610 | 206467.17 | 2 | 228473.78 | 2 | 2 | 2 | 2 | 317109.22 | 2 | 255875.11 | 2 | 3 | 1 | 5 | 2 | 5 | 0.232035317 | 0.269417482 | 0.305678641 | 0.354925151 |
| P01887 | B2m | Beta-2-microglobulin | 150.35 | 35.29 | 13779 | 3 | 6074018.5 | 7360020 | 9407858 | 10438001 | 8587429 | 6761889.5 | 6828585 | 10227463 | 8763791 | 10549246 | 4513722 | 6041876.5 | 7670179.5 | 7068350 | 9455346 | 6457259 | 6342426.5 | 8706771 | 7409681 | 10361858 | 7 | 11 | 7 | 8 | 7 | 11 | 0.829990279 | 0.805673287 | 0.938153897 | 0.910667935 |
| Q09200 | B4galnt1 | Beta-1,4 N-acetylgalactosaminyltransferase 1 | 223.95 | 14.45 | 59212 | 6 | 3611626.75 | 3651131.25 | 5156428.5 | 3497665.5 | 5558101 | 3979804.5 | 3688765 | 4478429 | 4271180 | 3231837.5 | 3284298.5 | 4895242 | 5580359.5 | 1977552.5 | 4913635.5 | 3775486.75 | 1751067.25 | 3005262 | 3024767 | 3977210.5 | 11 | 6 | 9 | 4 | 11 | 6 | 0.961636005 | 1.050945098 | 0.723344703 | 0.790523199 |
| P15535 | B4galt1 | Beta-1,4-galactosyltransferase 1 | 110.41 | 6.77 | 44411 | 2 | 234257.55 | 257301.67 | 2 | 329867.66 | 415033.53 | 304749.88 | 319913.84 | 252402.5 | 294196.59 | 2 | 2 | 2 | 2 | 2 | 2 | 246571.19 | 2 | 2 | 301713.75 | 324450.59 | 2.5 | 9 | 2.5 | 10 | 2.5 | 10 | 8.087589E-06 | 8.537779E-06 | 0.705835877 | 0.745125716 |
| Q6P7Q1 | Babam2 | BRISC and BRCA1-A complex member 2 | 55.06 | 3.66 | 43559 | 1 | 127591.12 | 114612.33 | 2 | 2 | 813637.56 | 2 | 2 | 2 | 604392.56 | 2 | 2 | 2 | 178471.72 | 2 | 2 | 2 | 2 | 184530.81 | 2 | 2 | 8 | 8 | 12 | 12 | 12 | 12 | 0.169039696 | 0.295300388 | 0.174778313 | 0.305325346 |
| Q6MG49 | Bag6 | Large proline-rich protein BAG6 | 185.4 | 3.66 | 120012 | 3 | 1679902.88 | 2133678.75 | 1889555.75 | 877168.12 | 1769627.62 | 509804.97 | 2349261.5 | 1662198.12 | 2997992.5 | 1219981.25 | 230423.88 | 492289.22 | 817690.31 | 2 | 1082244.5 | 1156720.25 | 996902.25 | 687448.88 | 916249.25 | 1181538.25 | 1 | 4 | 2 | 5 | 2 | 5 | 0.314092325 | 0.300100513 | 0.591484843 | 0.565136078 |
| O08734 | Bak1 | Bcl-2 homologous antagonist/killer | 337.7 | 34.45 | 23295 | 6 | 11962536 | 10311900 | 6745248.5 | 8090344.5 | 8270709.5 | 11448770 | 10557729 | 9090908 | 7138449.5 | 8821856 | 10772991 | 11099060 | 12529498 | 12732316 | 10397627 | 11498484 | 13304154 | 11033546 | 11558100 | 11444126 | 3 | 4 | 4 | 2 | 4 | 4 | 1.267751339 | 1.222573069 | 1.296550297 | 1.250345732 |
| O54962 | Banf1 | Barrier-to-autointegration factor | 259.47 | 52.81 | 10103 | 4 | 10598436 | 9011016 | 14254400 | 14565587 | 15305168 | 10615507 | 11217534 | 13989371 | 13986866 | 13936608 | 6724573.5 | 5618558 | 10235035 | 7481617 | 9347490 | 12392250 | 11550010 | 15967121 | 11576564 | 14718053 | 2 | 10 | 0 | 9 | 2 | 10 | 0.618302604 | 0.618193204 | 1.0387449 | 1.038561108 |
| Q91XV3 | Basp1 | Brain acid soluble protein 1 | 1206.97 | 83.19 | 22087 | 15 | 221095810 | 166081298 | 114296738 | 193591650 | 127235106 | 211086002 | 164883106 | 155059042 | 127267466 | 223221042 | 85773626 | 65896242 | 86146162 | 118788858 | 88783970 | 192481586 | 293953730 | 165916290 | 160184034 | 180322370 | 1 | 10 | 0 | 9 | 1 | 10 | 0.541637519 | 0.505252912 | 1.207414913 | 1.126306577 |
| Q07813 | Bax | Apoptosis regulator BAX | 480.43 | 40.62 | 21395 | 6 | 12366928 | 14811585 | 17297056 | 17336868 | 18945826 | 13407121 | 13542912 | 14294791 | 19099628 | 10614139 | 12536887 | 12793572 | 16100512 | 16056856 | 13937917 | 13492765 | 16980228 | 16834172 | 19489586 | 11519991 | 7 | 10 | 12 | 9 | 12 | 10 | 0.884438834 | 1.006583459 | 0.969767539 | 1.103696408 |
| Q9Z277 | Baz1b | Tyrosine-protein kinase BAZ1B | 249.37 | 3.65 | 170651 | 5 | 4189283.25 | 4287778 | 4111482 | 4384761.5 | 4152229.25 | 3728296 | 4585770 | 2615767.75 | 3014232 | 3337199.25 | 887648.38 | 2420304.5 | 1873248.38 | 1942934.25 | 2522276.25 | 2936937 | 1533712.5 | 2919676 | 2949842.5 | 3029727.5 | 0 | 0 | 0 | 5 | 0 | 5 | 0.456623334 | 0.558200558 | 0.632878464 | 0.773664168 |
| Q61335 | Bcap31 | B-cell receptor-associated protein 31 | 138.65 | 15.92 | 27957 | 3 | 1483167.38 | 2593766.5 | 1247654.38 | 2047341.5 | 1426627 | 2987009.5 | 1639476.88 | 1946836.25 | 1957506 | 2432846.25 | 2887474 | 2929106.5 | 3525285.5 | 3567821.5 | 2561308.25 | 1902235.5 | 2282732.5 | 2260634.5 | 2205191 | 2810955.5 | 1 | 5 | 3 | 11 | 3 | 11 | 1.758356077 | 1.411114058 | 1.302685124 | 1.045429487 |
| Q9D287 | Bcas2 | Pre-mRNA-splicing factor SPF27 | 63.21 | 5.78 | 26131 | 1 | 2 | 2 | 923708.31 | 2 | 2 | 1089048.75 | 2 | 2 | 2 | 2 | 2 | 2 | 2 | 2 | 723536.56 | 1426194.12 | 1069288.75 | 1249903.62 | 1219625.5 | 1157161 | 12 | 0 | 12 | 1 | 12 | 1 | 0.783297374 | 0.664377279 | 6.627763225 | 5.62153716 |
| P24288 | Bcat1 | Branched-chain-amino-acid aminotransferase, cytosolic | 219.77 | 11.66 | 42791 | 5 | 3779573 | 2341889.25 | 2175478 | 2311744.25 | 2629626.75 | 5064638 | 5400000 | 3225089 | 2006005.25 | 2819730 | 10205009 | 9245840 | 6986470.5 | 6745172 | 6915704.5 | 6600257 | 6544434.5 | 3549133.25 | 4581167 | 3258155.75 | 0 | 2 | 0 | 6 | 0 | 6 | 3.028950993 | 2.165660001 | 1.853193133 | 1.325008642 |
| O35855 | Bcat2 | Branched-chain-amino-acid aminotransferase, mitochondrial | 340.87 | 15.01 | 44127 | 5 | 20837462 | 20085942 | 18106018 | 23489116 | 17680850 | 22667158 | 19699926 | 20346566 | 18642548 | 19331194 | 13677339 | 13989961 | 17698942 | 17427866 | 17293580 | 16663638 | 19558416 | 19213962 | 22115774 | 21004682 | 1 | 12 | 0 | 11 | 1 | 12 | 0.799283205 | 0.7954093 | 0.983603533 | 0.978836278 |
| Q9CWI3 | Bccip | BRCA2 and CDKN1A-interacting protein | 103.66 | 7.59 | 35942 | 2 | 1719908.5 | 1336848.12 | 1222553.75 | 632402.44 | 1663543.88 | 1304378.38 | 1977416.5 | 777419.88 | 1140548.5 | 538649.38 | 637138.31 | 1471250.75 | 659052.81 | 156501.62 | 1055535 | 1324538 | 1682151.62 | 426873.78 | 411162.31 | 543469.94 | 6 | 6 | 8 | 9 | 8 | 9 | 0.605220249 | 0.693480713 | 0.667380128 | 0.76470549 |
| P35738 | Bckdhb | 2-oxoisovalerate dehydrogenase subunit beta, mitochondrial | 115.63 | 7.18 | 42823 | 2 | 987854 | 1040473 | 663908.88 | 919147.12 | 112160.17 | 1266032.5 | 1092499.25 | 102933.41 | 765661.31 | 106375.62 | 885400.94 | 733101.81 | 713412.75 | 596632.44 | 762999.19 | 827609.56 | 610373.31 | 463591.88 | 2 | 392173.25 | 9 | 5 | 11 | 9 | 11 | 9 | 0.991407098 | 1.107408074 | 0.616012732 | 0.688090164 |
| Q9Z0H7 | Bcl10 | B-cell lymphoma/leukemia 10 | 96.39 | 8.58 | 25948 | 2 | 876198.69 | 834370.25 | 991918.56 | 971190.5 | 1184364.25 | 806719.56 | 1213505.62 | 881183.56 | 1167156 | 920283.12 | 2 | 478889.97 | 2 | 2 | 2 | 875730.75 | 407561.66 | 720654.56 | 652777.75 | 713016.88 | 0 | 1 | 0 | 1 | 0 | 1 | 0.098578387 | 0.095993701 | 0.693641888 | 0.675454873 |
| P10417 | Bcl2 | Apoptosis regulator Bcl-2 | 51.79 | 4.24 | 26407 | 1 | 505046.41 | 505971.88 | 2 | 288652.06 | 1181353.88 | 172188.83 | 574629.44 | 618025.94 | 1272816.5 | 442915.22 | 2 | 366337.12 | 287051.72 | 118937.62 | 258126.03 | 257405.19 | 2 | 2 | 379760.62 | 302732.53 | 5.5 | 6 | 3 | 3 | 5.5 | 6 | 0.415333977 | 0.334500598 | 0.378836116 | 0.305106046 |
| P53563 | Bcl2l1 | Bcl-2-like protein 1 | 116.07 | 8.15 | 26158 | 2 | 5983776 | 5442084 | 4469930 | 5116802 | 4992255.5 | 5435748 | 5390682 | 4588434 | 4811265 | 4647390 | 3178506.5 | 3726258.75 | 3068271.5 | 2736069 | 2172570.5 | 6483247 | 5799423 | 4580708 | 4605383.5 | 4065793 | 0 | 11 | 0 | 11 | 0 | 11 | 0.572265469 | 0.598293963 | 0.981915179 | 1.026575874 |
| P59017 | Bcl2l13 | Bcl-2-like protein 13 | 90.48 | 9.45 | 46719 | 2 | 822995.94 | 851050.56 | 915134.25 | 775541.25 | 949020.94 | 363038 | 779211.5 | 782276.56 | 1813582.88 | 432619.81 | 391183.88 | 197470.09 | 305857.62 | 296963.59 | 326555.19 | 664107.94 | 484148.91 | 428570.44 | 762784 | 497649.94 | 0 | 0 | 1 | 9 | 1 | 9 | 0.351905617 | 0.363972452 | 0.657726079 | 0.680279491 |
| Q8K019 | Bclaf1 | Bcl-2-associated transcription factor 1 | 252 | 5.98 | 106002 | 6 | 6429701.5 | 7750148.5 | 5239115 | 5818503 | 4533822.5 | 7975591 | 7676946.5 | 5962809 | 5212348 | 4260029.5 | 5421957 | 5709003.5 | 3980684.5 | 4925428.5 | 4645798 | 7514888.5 | 5628347 | 5413183 | 5806791.5 | 7187796.5 | 6 | 11 | 6 | 12 | 6 | 12 | 0.829083022 | 0.793974866 | 1.059779605 | 1.014902426 |
| Q9CZP5 | Bcs1l | Mitochondrial chaperone BCS1 | 118.52 | 6.94 | 47406 | 2 | 112066.77 | 353453 | 471989.84 | 266622.44 | 547057.06 | 192240.95 | 300653.25 | 533586.5 | 536282.38 | 427995.12 | 751819.56 | 300422.59 | 1090861.88 | 1103507.75 | 955457.25 | 271012.72 | 363200.5 | 550149.31 | 706591.38 | 487021.34 | 3 | 6 | 4 | 9 | 4 | 9 | 2.399551828 | 2.110788256 | 1.357920305 | 1.194507324 |
| O08539 | Bin1 | Myc box-dependent-interacting protein 1 | 72.24 | 4.59 | 64470 | 2 | 4157506.5 | 3862174 | 3264043.25 | 3437358 | 3223071.5 | 4200793.5 | 3632246.5 | 2921400.5 | 3057574 | 3228750 | 3082021.5 | 2679215.5 | 2398512.25 | 2617835.5 | 2607420.75 | 4310908.5 | 5144775 | 3176149.75 | 3810881 | 3538203.75 | 0 | 9 | 2 | 6 | 2 | 9 | 0.745925724 | 0.785469777 | 1.11350576 | 1.172536479 |
| D3Z6Q9 | Bin2 | Bridging integrator 2 | 268.63 | 10.84 | 52554 | 5 | 4130938.75 | 4873408 | 2885032 | 4729066.5 | 2822600.25 | 4484485.5 | 4546635 | 2864642 | 4688729 | 4481656 | 3334740.5 | 3601795.25 | 2610985.75 | 2663621 | 3626438 | 4491136 | 5766866 | 3898062.5 | 3430298.25 | 4117135.75 | 6 | 11 | 3 | 11 | 6 | 11 | 0.814646543 | 0.751802412 | 1.116375068 | 1.030254749 |
| O88738 | Birc6 | Baculoviral IAP repeat-containing protein 6 | 252.13 | 1.29 | 532170 | 6 | 2305818.25 | 2096187.25 | 1601537.75 | 1320638.12 | 3418059 | 1669573.12 | 2596904 | 1136682.75 | 2551665.75 | 712327.12 | 626229.62 | 1317883.75 | 962871.44 | 2 | 1015251.19 | 2233925.5 | 809477.62 | 1109425 | 787512.12 | 505331.62 | 0 | 3 | 4 | 6 | 4 | 6 | 0.365122904 | 0.452540542 | 0.506940049 | 0.628311514 |
| Q566Q8 | Bles03 | UPF0696 protein C11orf68 homolog | 95.59 | 7.57 | 27493 | 2 | 4445206 | 4077209.5 | 3795122.25 | 2788341.5 | 4359151 | 1993332 | 3673701 | 3701075 | 4043024 | 3071245.25 | 636979.25 | 2494373.5 | 1510295.88 | 533081.88 | 1955884 | 4284398 | 923915.56 | 1870041.25 | 1390528.62 | 1006317.06 | 0 | 3 | 1 | 5 | 1 | 5 | 0.366329485 | 0.432620514 | 0.486780671 | 0.574868561 |
| Q8R016 | Blmh | Bleomycin hydrolase | 460.86 | 26.37 | 52511 | 8 | 3155000.25 | 4288166 | 5613409.5 | 5085691 | 5512334 | 4964528 | 4051054.5 | 4001199.25 | 5409179.5 | 3240309.75 | 2126503.25 | 2848438 | 1940982.75 | 3054732.5 | 2134410.5 | 5263597.5 | 4646043 | 4200817.5 | 2778729 | 4690108.5 | 0 | 8 | 0 | 12 | 0 | 12 | 0.511742605 | 0.558705603 | 0.912266317 | 0.995985673 |
| Q9CY64 | Blvra | Biliverdin reductase A | 742.43 | 49.15 | 33525 | 13 | 34148002 | 35786338 | 31139602 | 32285032 | 31313518 | 34014330 | 31515080 | 35357358 | 31699838 | 33271602 | 27281938 | 28413794 | 28861326 | 25251022 | 23350476 | 31471798 | 28921738 | 30461610 | 31793530 | 30824600 | 0 | 4 | 0 | 2 | 0 | 4 | 0.808626592 | 0.802845742 | 0.931990973 | 0.925328194 |
| Q923D2 | Blvrb | Flavin reductase (NADPH) | 485.1 | 52.43 | 22197 | 6 | 2 | 208048.06 | 129867.8 | 1111843.12 | 587203.44 | 1038699.06 | 404462.66 | 954894.12 | 628881.62 | 198019.28 | 14671318 | 6608738 | 6307858 | 8127343 | 6285399 | 1212510.25 | 1520904.88 | 1950822.5 | 1578359 | 1225602.38 | 0 | 0 | 0 | 0 | 0 | 0 | 20.6192389 | 13.02363392 | 3.676156017 | 2.32195332 |
| Q9D8S9 | Bola1 | BolA-like protein 1 | 52.91 | 11.68 | 14379 | 1 | 1291989 | 1512927.5 | 1583565.75 | 1672233.75 | 1496165.12 | 1506708.88 | 1604919.75 | 1232906 | 1290167.88 | 1557782.12 | 1581180.62 | 1359919.88 | 1187520 | 1436074.38 | 1659663.5 | 1763800.38 | 2460316 | 1676388.62 | 1676782.25 | 2172626.25 | 9 | 0 | 12 | 0 | 12 | 0 | 0.955997357 | 1.004431535 | 1.290203372 | 1.355569598 |
| P97452 | Bop1 | Ribosome biogenesis protein BOP1 | 205.06 | 7.92 | 82546 | 4 | 1502194.88 | 1759157.5 | 2533172 | 1840094.25 | 1440005.75 | 2337324 | 1772804 | 1808883 | 1241538.25 | 1881000.62 | 2562245 | 2813368 | 1937007 | 1770629.5 | 1744445 | 2940848.25 | 2644264.5 | 2276935.5 | 2118041 | 1362665.25 | 6 | 7 | 9 | 6 | 9 | 7 | 1.193183767 | 1.197548502 | 1.249942039 | 1.254514399 |
| Q8R164 | Bphl | Valacyclovir hydrolase | 52.71 | 6.87 | 32851 | 1 | 143080.97 | 2 | 164740.08 | 2 | 142943.8 | 2 | 132503.5 | 2 | 168767.92 | 2 | 2 | 2 | 193442 | 2 | 2 | 179214.7 | 2 | 2 | 2 | 2 | 9 | 9 | 11 | 11 | 11 | 11 | 0.429155653 | 0.642099232 | 0.397593356 | 0.594875978 |
| Q9Z1N4 | Bpnt1 | 3'(2'),5'-bisphosphate nucleotidase 1 | 294.37 | 20.45 | 33174 | 5 | 1145364.75 | 687222.44 | 723300.25 | 839941.12 | 1185456.5 | 1190009.75 | 1022534.81 | 613900.06 | 776222.62 | 477652.47 | 1159556.38 | 623821.88 | 488773.47 | 594000.31 | 620078.62 | 872875.75 | 1350228.88 | 1344169.25 | 426380.78 | 806780.94 | 4 | 10 | 10 | 9 | 10 | 10 | 0.760972219 | 0.854401348 | 1.047836041 | 1.176485163 |
| Q9ESU6 | Brd4 | Bromodomain-containing protein 4 | 169.99 | 3.64 | 155895 | 4 | 2080296.5 | 2404465.5 | 2669863.5 | 2591601 | 2567059 | 2529392 | 2655828.5 | 2779690.5 | 2550171 | 1937684.75 | 769168.62 | 1204432.5 | 1276054.62 | 562412.44 | 919077.69 | 2484575.25 | 2474942.5 | 2230089.5 | 2290493.25 | 2190506.5 | 0 | 7 | 0 | 5 | 0 | 7 | 0.384230989 | 0.379927286 | 0.947806091 | 0.937189882 |
| Q8BXV2 | Bri3bp | BRI3-binding protein | 171.94 | 13.04 | 28263 | 2 | 2 | 2 | 2 | 2 | 2 | 2 | 2 | 2 | 2 | 2 | 2 | 2 | 803731.25 | 2 | 600597.81 | 2 | 2 | 2 | 2 | 2 | 7.5 | 12.5 | 7.5 | 12.5 | 7.5 | 12.5 | 140433.506 | 140433.506 | 1 | 1 |
| Q4QQT6 | Brix1 | Ribosome biogenesis protein BRX1 homolog | 301.04 | 17.33 | 41115 | 5 | 3441228.75 | 2608474.5 | 3397461 | 2166050 | 2405865 | 2835981.25 | 3460996 | 1680778.88 | 1930076.38 | 1929778.75 | 3013490 | 4558545 | 4538242.5 | 2675890.25 | 2998786.5 | 3055638.75 | 3889624.25 | 2975344.75 | 2535866 | 2037355.62 | 6 | 12 | 4 | 6 | 6 | 12 | 1.268624988 | 1.502410736 | 1.033864572 | 1.224388016 |
| Q91VR8 | Brk1 | Protein BRICK1 | 88.76 | 25.33 | 8761 | 2 | 2839962.75 | 2790175.5 | 3936429.75 | 3163345.5 | 4452210 | 2477315 | 3252593 | 3816226.75 | 4241510 | 3348926 | 1727652.25 | 2463406.75 | 2104739.75 | 2744413.25 | 2148803.5 | 2599547.5 | 2296945.25 | 3564559.5 | 3317075.75 | 3073138.25 | 0 | 8 | 1 | 7 | 1 | 8 | 0.651200971 | 0.652932005 | 0.864344052 | 0.866641667 |
| Q4V8K5 | Brox | BRO1 domain-containing protein BROX | 214.19 | 10.46 | 46192 | 3 | 1912412.12 | 1827141.5 | 2566792.5 | 2714611 | 3089391.25 | 1446266.12 | 1231890.5 | 2922309.5 | 2923115.5 | 2254674.5 | 538933.88 | 2215445.5 | 3001685.5 | 2353882.5 | 2635881 | 2400665.75 | 705919.62 | 2364097.5 | 1917153.5 | 2695017.5 | 11 | 9 | 12 | 11 | 12 | 11 | 0.887326116 | 0.996991374 | 0.832581653 | 0.935481005 |
| Q80Y55 | Bsdc1 | BSD domain-containing protein 1 | 131.84 | 7.26 | 46953 | 2 | 208326.16 | 713448.81 | 594767.19 | 451121.25 | 432251.94 | 391390.5 | 690462.38 | 320638.03 | 665693.5 | 513951.84 | 222754.75 | 2 | 265128.97 | 151229.44 | 141569.52 | 330918.25 | 156453.83 | 725670.75 | 371800.41 | 509307 | 2 | 10 | 0 | 9 | 2 | 10 | 0.325296757 | 0.302340622 | 0.872593377 | 0.811014616 |
| P18572 | Bsg | Basigin | 94.16 | 6.43 | 42445 | 2 | 3886586 | 2797314.5 | 3296496.5 | 3538491.5 | 2501834.5 | 5263315 | 3396960 | 3360117 | 2386263.75 | 5216186.5 | 5974050.5 | 5489107.5 | 5946719.5 | 7031816 | 6179538.5 | 5059563 | 6006546.5 | 3882911.25 | 4168914 | 3876563.75 | 0 | 2 | 0 | 8 | 0 | 8 | 1.911351442 | 1.560489128 | 1.435297177 | 1.171823032 |
| Q63072 | Bst1 | ADP-ribosyl cyclase/cyclic ADP-ribose hydrolase 2 | 44.43 | 4.08 | 35131 | 1 | 549269.5 | 408333.66 | 249632.67 | 367619.03 | 2 | 598117.81 | 404270.12 | 281517.03 | 324373.75 | 323263.91 | 313221.91 | 2 | 286790.75 | 2 | 237093.61 | 466364.69 | 551691.94 | 495281.34 | 274136.5 | 328449.44 | 6 | 8 | 2 | 10 | 6 | 10 | 0.531546892 | 0.433389489 | 1.343565859 | 1.095458049 |
| Q8R2Q8 | Bst2 | Bone marrow stromal antigen 2 | 249.12 | 17.44 | 19152 | 4 | 5135926.5 | 6803412.5 | 4709711 | 4146478.5 | 8321500 | 2889279 | 7638503 | 9641277 | 10372434 | 5367512 | 4062948 | 13298285 | 5544086 | 3379985.75 | 6216299 | 5513553 | 2412361.25 | 3028555 | 3321248.75 | 3619952 | 11 | 3 | 11 | 5 | 11 | 5 | 1.116240407 | 0.905110118 | 0.614611824 | 0.498361623 |
| Q64152 | Btf3 | Transcription factor BTF3 | 267.65 | 33.82 | 22031 | 4 | 2 | 371407.34 | 1088299.12 | 1240705.38 | 664365.12 | 600144.12 | 326985.41 | 617994.25 | 456508.38 | 542940.56 | 1159616.38 | 1160822.5 | 2085161 | 2547377.5 | 1916867.5 | 1623086 | 1833734.12 | 1485502.75 | 1652312.88 | 1747117.62 | 2 | 0 | 0 | 0 | 2 | 0 | 2.636085456 | 3.485789504 | 2.479138591 | 3.278253085 |
| P35991 | Btk | Tyrosine-protein kinase BTK | 72.9 | 6.22 | 76437 | 2 | 403223.88 | 241318.52 | 569779.25 | 402548.62 | 888264 | 253235.88 | 415581.12 | 582004.38 | 581420.25 | 541716.12 | 389629.72 | 557886.5 | 447882.12 | 2 | 471562.75 | 466408.78 | 474961.31 | 325838.81 | 488697.97 | 607230.62 | 10 | 11 | 8 | 12 | 10 | 12 | 0.7452547 | 0.786434843 | 0.943317697 | 0.995442101 |
| O08901 | Bub1 | Mitotic checkpoint serine/threonine-protein kinase BUB1 | 39.2 | 0.66 | 119563 | 1 | 53064110 | 56774522 | 67513602 | 72164474 | 58571318 | 58706218 | 50041318 | 64327986 | 53182234 | 67461266 | 31996902 | 39303550 | 40061906 | 48747586 | 40773638 | 61797510 | 77785898 | 65190994 | 67593570 | 75526626 | 0 | 5 | 0 | 3 | 0 | 5 | 0.652033072 | 0.683931128 | 1.12920519 | 1.184446944 |
| Q9WVA3 | Bub3 | Mitotic checkpoint protein BUB3 | 389.95 | 21.78 | 36955 | 7 | 31337964 | 31045182 | 31081166 | 32318698 | 28737694 | 33658702 | 28977592 | 25943822 | 25812254 | 26831082 | 17059766 | 21671838 | 16436074 | 23576734 | 22049990 | 30316034 | 36363978 | 27500792 | 29730698 | 29196570 | 0 | 8 | 0 | 5 | 0 | 8 | 0.652303539 | 0.713722831 | 0.990857976 | 1.08415472 |
| Q9CY21 | Bud23 | Probable 18S rRNA (guanine-N(7))-methyltransferase | 60.54 | 5.69 | 31587 | 1 | 2 | 2 | 2 | 389252.09 | 236317.53 | 2 | 2 | 630910.38 | 2 | 2 | 705836.88 | 579008.94 | 2 | 2 | 2 | 2 | 416004.75 | 2 | 2 | 2 | 10.5 | 11 | 10 | 12 | 10.5 | 12 | 2.053871313 | 2.036478665 | 0.665007933 | 0.659376495 |
| O70454 | Bud31 | Protein BUD31 homolog | 142.78 | 27.08 | 17000 | 3 | 147179.78 | 376540.09 | 631144 | 902939.5 | 888460.12 | 271665.62 | 463129.66 | 880346 | 363468.66 | 1259943.25 | 408733.38 | 571168 | 661206.12 | 728383.19 | 549368.69 | 419910.81 | 904828.06 | 778745.38 | 466988.5 | 488496.97 | 12 | 11 | 11 | 10 | 12 | 11 | 0.99069869 | 0.90128499 | 1.038253955 | 0.944548241 |
| O54825 | Bysl | Bystin | 231.2 | 9.4 | 49784 | 4 | 1523993.5 | 1220600 | 852859.19 | 1177252.38 | 158790.75 | 1404297 | 1084647.75 | 121461.29 | 879509.31 | 1221797.62 | 2902069.5 | 3208845.75 | 2921822.5 | 2149537.5 | 2000373.88 | 1623538.38 | 1731947.75 | 1040340.19 | 1017715.81 | 912993 | 0 | 9 | 0 | 9 | 0 | 9 | 2.672070599 | 2.797846391 | 1.282363533 | 1.342725071 |
| Q6P7P5 | Bzw1 | Basic leucine zipper and W2 domain-containing protein 1 | 873.18 | 45.82 | 48043 | 17 | 11533360 | 12613858 | 15282309 | 11847883 | 21570058 | 7651582.5 | 9371216 | 12835498 | 21636114 | 9415657 | 8933514 | 14767918 | 9340816 | 7163664 | 10894633 | 10885099 | 9684606 | 11419884 | 10404326 | 10193981 | 3 | 0 | 9 | 10 | 9 | 10 | 0.70147318 | 0.83895072 | 0.721890512 | 0.863369524 |
| Q2L4X1 | Bzw2 | Basic leucine zipper and W2 domain-containing protein 2 molossinus (Japanese house mouse) | 185.98 | 11.46 | 48063 | 4 | 1683235.62 | 1429959.12 | 1493424.12 | 1421652.88 | 1153408 | 1486018.62 | 1398478.38 | 726976.62 | 1221753 | 1266509.5 | 1381665.5 | 1510658.12 | 834710.12 | 1152822.12 | 997628.75 | 978487.88 | 1422963.75 | 1081451.38 | 1333912.5 | 1277554.5 | 5 | 4 | 11 | 12 | 11 | 12 | 0.818399709 | 0.96356375 | 0.848599524 | 0.999120272 |
| P14106 | C1qb | Complement C1q subcomponent subunit B | 148.94 | 8.3 | 26717 | 2 | 2376605.75 | 3038769.25 | 1179590.38 | 1368993.75 | 1472854.75 | 2005331.12 | 2429269.25 | 1394878.75 | 947475 | 1065909 | 849015.94 | 657943.25 | 2 | 2 | 2 | 2066759.25 | 2418033.75 | 764220.81 | 447096.72 | 2 | 0 | 7 | 0 | 8 | 0 | 8 | 0.15969004 | 0.192144778 | 0.603605476 | 0.726279733 |
| Q80X80 | C2cd2l | Phospholipid transfer protein C2CD2L | 57.47 | 1.42 | 76329 | 1 | 2 | 349934.97 | 210200.73 | 2 | 249467.64 | 587846.25 | 323152.31 | 267317.59 | 251236.81 | 230962.64 | 234753.31 | 320268.78 | 405684.94 | 2 | 2 | 217320.31 | 2 | 2 | 2 | 2 | 11 | 7 | 8 | 0 | 11 | 7 | 1.186638241 | 0.578561882 | 0.268436684 | 0.130880017 |
| P01027 | C3 | Complement C3 | 317.4 | 4.87 | 186484 | 7 | 1971061.12 | 790038.44 | 2593966.75 | 3325820.5 | 3827975.25 | 2024950.88 | 2489394.75 | 3701321.75 | 5424224 | 3773607.5 | 6384931.5 | 5884645 | 8649081 | 9174553 | 8417449 | 902166.38 | 2294937.5 | 3349028 | 4154984.75 | 3636309.5 | 0 | 9 | 0 | 9 | 0 | 9 | 3.078670091 | 2.211540585 | 1.146181488 | 0.823351253 |
| O09047 | C3ar1 | C3a anaphylatoxin chemotactic receptor | 43.87 | 1.47 | 53576 | 1 | 2405669.5 | 2126790.25 | 909989.69 | 1485782.25 | 1066047.38 | 3251595.75 | 2276333.75 | 1287708.12 | 1360665 | 1787395.25 | 1945238.62 | 2039755.88 | 2413092.5 | 2280999.75 | 2395062.25 | 2157187.25 | 2391238.5 | 1781003.5 | 1755423.75 | 1855759.75 | 6 | 8 | 7 | 11 | 7 | 11 | 1.385259246 | 1.111449699 | 1.243465816 | 0.997683077 |
| P01029 | C4b | Complement C4-B | 82.86 | 1.27 | 192915 | 1 | 2 | 2 | 2 | 2 | 2 | 2 | 2 | 90546.8 | 2 | 2 | 105530.12 | 116954.55 | 478942 | 435041.59 | 458600.28 | 2 | 2 | 277277.56 | 288529.41 | 125331.89 | 0 | 5 | 0 | 6 | 0 | 6 | 159506.854 | 17.61440078 | 69114.286 | 7.632316122 |
| P30993 | C5ar1 | C5a anaphylatoxin chemotactic receptor 1 | 133.86 | 8.26 | 39023 | 3 | 15491035 | 15617265 | 11642071 | 15047504 | 13113567 | 15143459 | 14910852 | 11746356 | 11132380 | 13743704 | 19034822 | 19478004 | 21360574 | 21672352 | 21959542 | 17688686 | 20563804 | 13774112 | 15859223 | 14815232 | 0 | 6 | 0 | 4 | 0 | 6 | 1.459641647 | 1.552344595 | 1.166765964 | 1.240868155 |
| Q9D6N1 | Ca13 | Carbonic anhydrase 13 | 103.86 | 4.2 | 29522 | 2 | 1025524.44 | 443455.53 | 693409.25 | 1746565.38 | 342916.34 | 1757195.25 | 416268.12 | 570204.69 | 323317.94 | 1594671.12 | 739035.69 | 549799.38 | 1077125.5 | 1026206.88 | 825674.88 | 668820.25 | 1756540 | 2013881.25 | 1691226 | 1673171.88 | 9 | 5 | 11 | 5 | 11 | 5 | 0.991996791 | 0.90479463 | 1.835342486 | 1.67400544 |
| P00920 | Ca2 | Carbonic anhydrase 2 | 57.58 | 6.15 | 29033 | 1 | 613407.06 | 361823.91 | 341655.97 | 371467.28 | 487354 | 440425.09 | 330471.5 | 387761.66 | 366198.97 | 373392.84 | 288087.12 | 207276.84 | 226640.8 | 176559.75 | 108265.89 | 338823.22 | 354136.59 | 330817.91 | 248126.77 | 247569.97 | 0 | 1 | 0 | 3 | 0 | 3 | 0.462759846 | 0.530399246 | 0.698381541 | 0.800460641 |
| Q9CXW3 | Cacybp | Calcyclin-binding protein | 572.84 | 45.41 | 26510 | 10 | 30273778 | 31879846 | 32453310 | 24680994 | 30928280 | 25114858 | 31127102 | 28016542 | 29031882 | 25933426 | 16201527 | 25225594 | 17202650 | 15691935 | 14723398 | 30705798 | 21281410 | 23241872 | 22392810 | 24945556 | 1 | 3 | 1 | 4 | 1 | 4 | 0.592779602 | 0.639582439 | 0.815940221 | 0.880362676 |
| B2RQC6 | Cad | CAD protein | 1505.8 | 13.12 | 243238 | 24 | 11373606 | 13200222 | 14757128 | 10609823 | 15202567 | 11242790 | 13707552 | 14560994 | 14823904 | 13316910 | 19145830 | 19989256 | 16531820 | 12658203 | 15761605 | 13599590 | 11210586 | 12101287 | 10892259 | 12122547 | 3 | 9 | 4 | 4 | 4 | 9 | 1.290795134 | 1.242927446 | 0.919913893 | 0.885799919 |
| Q8R5M8 | Cadm1 | Cell adhesion molecule 1 | 188.29 | 10.09 | 49788 | 3 | 420523.19 | 1628873.12 | 1945474.38 | 1681808.5 | 1408268.12 | 1706651.38 | 1794733 | 2151235.25 | 2375443 | 1860244.75 | 1727667.5 | 1974619 | 1990333.38 | 1556761.62 | 1985246.25 | 1635902.5 | 946921.56 | 1446299.12 | 2 | 1524597.25 | 4 | 8 | 10 | 0 | 10 | 8 | 1.303415163 | 0.933893678 | 0.783876321 | 0.561645408 |
| Q8VEC4 | Calhm2 | Calcium homeostasis modulator protein 2 | 91.32 | 9.91 | 35823 | 2 | 194360.58 | 143750.45 | 489369.69 | 191503.53 | 536262.25 | 2 | 451856.19 | 443153.5 | 394499.91 | 163226.17 | 66732 | 2 | 2 | 2 | 2 | 154880.67 | 2 | 110965.61 | 2 | 101088.79 | 0 | 1 | 3 | 4 | 3 | 4 | 0.042912812 | 0.045940845 | 0.235936278 | 0.252584518 |
| P0DP26 | Calm1 | Calmodulin-1 | 484.43 | 63.09 | 16838 | 7 | 3540860.5 | 6075226 | 4435032 | 1087698.25 | 11500129 | 940173.94 | 5972454 | 6970917.5 | 9816110 | 3475069.5 | 942832.38 | 2788222.75 | 1166909 | 132905.23 | 1647890.38 | 6016525 | 1149479.5 | 1702524.38 | 1820192.75 | 2444137.25 | 3 | 7 | 4 | 7 | 4 | 7 | 0.250714116 | 0.245771015 | 0.492994693 | 0.483274768 |
| P14211 | Calr | Calreticulin | 1693.59 | 59.38 | 47995 | 26 | 409721410 | 468065890 | 433742082 | 408013922 | 425892674 | 391748258 | 456689730 | 454773282 | 409010978 | 381036354 | 297192034 | 371624642 | 366896386 | 324613954 | 419252578 | 440619330 | 305931778 | 403019042 | 344045282 | 421763458 | 2 | 6 | 3 | 8 | 3 | 8 | 0.829472244 | 0.85014799 | 0.892769073 | 0.915022582 |
| O35887 | Calu | Calumenin | 361.6 | 24.44 | 37064 | 5 | 6023502.5 | 6352820.5 | 7804990 | 5627518 | 8401052 | 3435656.5 | 6273500.5 | 7649974 | 7081051 | 5047854 | 2800721 | 7600023.5 | 6434267 | 6277330 | 6641233 | 6969859.5 | 3469429.75 | 4623831 | 4149939.75 | 6258223.5 | 11 | 5 | 12 | 8 | 12 | 8 | 0.869736225 | 1.009004957 | 0.744559211 | 0.863783654 |
| Q63450 | Camk1 | Calcium/calmodulin-dependent protein kinase type 1 | 176.08 | 9.89 | 41638 | 3 | 1126003 | 1048995.62 | 733757.75 | 595430.81 | 2162993.5 | 721501.19 | 1714746.75 | 838690.75 | 2201000 | 605865.31 | 2 | 367121.75 | 250972.03 | 140350.34 | 215962.78 | 1612253.12 | 739433.94 | 591837.94 | 723871 | 468101.88 | 0 | 7 | 0 | 7 | 0 | 7 | 0.171938915 | 0.160217084 | 0.729727551 | 0.679978815 |
| Q6PHZ2 | Camk2d | Calcium/calmodulin-dependent protein kinase type II subunit delta | 630.65 | 23.85 | 56369 | 10 | 14761767 | 17290820 | 14772407 | 15316142 | 19107852 | 14276134 | 14820661 | 19050226 | 16972416 | 17902582 | 10608928 | 10671437 | 11223797 | 8371907 | 12397556 | 17281780 | 15815405 | 19507128 | 21090912 | 19628640 | 0 | 4 | 0 | 5 | 0 | 5 | 0.655683551 | 0.641680673 | 1.148615722 | 1.124085708 |
| Q6ZQ38 | Cand1 | Cullin-associated NEDD8-dissociated protein 1 | 1054.47 | 20.65 | 136332 | 18 | 28769238 | 26692090 | 30039876 | 32621932 | 32496030 | 26715000 | 27008500 | 31462830 | 33765678 | 30072296 | 20132430 | 22119792 | 26975474 | 21094742 | 23364316 | 25948690 | 21893548 | 30268794 | 28521350 | 30111600 | 1 | 7 | 1 | 8 | 1 | 8 | 0.754796066 | 0.762873914 | 0.907879028 | 0.917595173 |
| P35564 | Canx | Calnexin | 1774.08 | 46.36 | 67278 | 28 | 46254934 | 41993938 | 45009826 | 47516962 | 45770370 | 43795870 | 47622826 | 51902226 | 50187818 | 58924594 | 92233210 | 77760794 | 81179882 | 63588282 | 83527874 | 54067514 | 54062926 | 52782790 | 50798246 | 50104354 | 0 | 0 | 0 | 8 | 0 | 8 | 1.758097646 | 1.577802882 | 1.155684918 | 1.037168213 |
| P40124 | Cap1 | Adenylyl cyclase-associated protein 1 | 2521.46 | 68.57 | 51565 | 36 | 454731010 | 432073826 | 380845410 | 349042722 | 404554530 | 460949922 | 432430946 | 384106434 | 346042466 | 333121794 | 222119330 | 226758658 | 169648098 | 177702498 | 163722226 | 506864450 | 398874722 | 300150242 | 309237986 | 311801634 | 0 | 7 | 0 | 8 | 0 | 8 | 0.474929869 | 0.490608971 | 0.903862113 | 0.933701774 |
| P24452 | Capg | Macrophage-capping protein | 758.64 | 34.94 | 39240 | 10 | 2 | 882755.81 | 2 | 2 | 2 | 2 | 2 | 2 | 2 | 2 | 6362775 | 4258558.5 | 2413275 | 2430783 | 2980359.25 | 2 | 2 | 2 | 2 | 2 | 0 | 10 | 0 | 12.5 | 0 | 12.5 | 20.89545419 | 1844575.075 | 1.132806E-05 | 1 |
| O35350 | Capn1 | Calpain-1 catalytic subunit | 325.71 | 8.27 | 82106 | 6 | 3334657.25 | 1801237 | 3479715 | 3102148.5 | 4762940 | 2300542.5 | 2546136.5 | 2600073.25 | 3792394.5 | 2979434.75 | 172390.72 | 1760429.38 | 754564.12 | 641610.5 | 1441714.12 | 3422802.5 | 1220637.38 | 3353927.75 | 2379836.5 | 3110247.5 | 0 | 9 | 0 | 12 | 0 | 12 | 0.289472504 | 0.335526356 | 0.818378678 | 0.948579268 |
| O08529 | Capn2 | Calpain-2 catalytic subunit | 516.64 | 19.29 | 79872 | 10 | 6698480.5 | 5953105 | 4311564.5 | 4285090 | 7881370.5 | 5359264 | 6396876.5 | 4802521.5 | 7413379 | 3991678.25 | 4408469 | 3782247 | 3816807 | 2688660 | 2557550.25 | 5292069.5 | 4648910.5 | 3651381.5 | 3251504.5 | 3050047.5 | 2 | 4 | 1 | 3 | 2 | 4 | 0.592309096 | 0.617004237 | 0.682944714 | 0.711418725 |
| O88456 | Capns1 | Calpain small subunit 1 | 454.05 | 32.34 | 28463 | 7 | 12699062 | 11054213 | 10990640 | 8734476 | 11349547 | 8508298 | 10983392 | 9046370 | 11015143 | 8013432 | 2847223 | 6101771.5 | 2315000.75 | 1051738.62 | 4044087.5 | 11144080 | 8247007.5 | 5794765 | 5712035 | 6976809 | 0 | 3 | 0 | 6 | 0 | 6 | 0.298384765 | 0.343934806 | 0.690791919 | 0.796245026 |
| Q5M9G3 | Caprin1 | Caprin-1 | 779.34 | 15.98 | 78121 | 12 | 15963971 | 16758913 | 14540034 | 12160922 | 12115580 | 17465544 | 13887069 | 10984277 | 11899044 | 10893736 | 15070785 | 16985354 | 11169448 | 9520574 | 12014181 | 14861062 | 15668413 | 12277182 | 12093169 | 11806099 | 8 | 8 | 12 | 9 | 12 | 9 | 0.905239964 | 0.994329343 | 0.932435921 | 1.024201796 |
| P47753 | Capza1 | F-actin-capping protein subunit alpha-1 | 493.17 | 36.36 | 32940 | 9 | 17945636 | 17215050 | 15094537 | 14184220 | 17536002 | 17928808 | 17030992 | 14020186 | 15595670 | 14102569 | 11103678 | 9524960 | 8189003 | 7174848 | 9206734 | 16221279 | 17780278 | 13523088 | 12902043 | 13518558 | 0 | 6 | 0 | 7 | 0 | 7 | 0.551375147 | 0.574481987 | 0.90204141 | 0.939843851 |
| P47754 | Capza2 | F-actin-capping protein subunit alpha-2 | 594.84 | 47.55 | 32967 | 11 | 27279572 | 27435490 | 22769862 | 26858764 | 29924502 | 26134616 | 30298978 | 23531586 | 29306442 | 26596066 | 12204783 | 11001484 | 13006926 | 12325875 | 13858605 | 29088270 | 29478204 | 26228878 | 23719384 | 21982158 | 0 | 10 | 0 | 10 | 0 | 10 | 0.464724169 | 0.459253218 | 0.971912215 | 0.96047041 |
| Q5XI32 | Capzb | F-actin-capping protein subunit beta | 964.97 | 44.49 | 30629 | 15 | 78737442 | 87836946 | 92646642 | 98804978 | 90624874 | 82884402 | 86002642 | 91528042 | 82415842 | 83482482 | 39567682 | 46893322 | 44563590 | 46664402 | 48374170 | 91263938 | 91504266 | 86511234 | 93787242 | 92017610 | 0 | 11 | 0 | 3 | 0 | 11 | 0.503873223 | 0.530274584 | 1.014339452 | 1.067487626 |
| A2AIV8 | Card9 | Caspase recruitment domain-containing protein 9 | 74.35 | 1.87 | 62462 | 1 | 459462.38 | 584665.38 | 385252.19 | 284798.56 | 390091.22 | 651487.06 | 488100.56 | 328995.69 | 421552.53 | 238421.42 | 305025.53 | 493807.72 | 301679.38 | 2 | 2 | 616900.69 | 410147.31 | 2 | 2 | 332755.56 | 6 | 9 | 6 | 8 | 6 | 9 | 0.522992188 | 0.517024677 | 0.646213525 | 0.638840019 |
| Q4AE70 | Carm1 | Histone-arginine methyltransferase CARM1 | 37.2 | 2 | 70341 | 1 | 70138.98 | 2 | 2 | 189779.16 | 2 | 2 | 2 | 2 | 2 | 2 | 228177.22 | 214448.17 | 321026.53 | 345460.75 | 214950.98 | 59143.31 | 392782 | 159819.75 | 2 | 181364.05 | 0 | 7.5 | 0 | 2.5 | 0 | 7.5 | 5.094038784 | 132406.365 | 3.051317627 | 79311.111 |
| Q9ER72 | Cars | Cysteine--tRNA ligase, cytoplasmic | 654.48 | 17.57 | 94860 | 13 | 14108915 | 17247996 | 11907128 | 12266642 | 14091791 | 16327042 | 14926280 | 13828549 | 18672276 | 15172915 | 16170358 | 16833418 | 16025664 | 16072440 | 15210744 | 14777747 | 16589348 | 16163338 | 16368988 | 16159511 | 5 | 5 | 9 | 10 | 9 | 10 | 1.153544562 | 1.017554967 | 1.149900739 | 1.014340709 |
| P29452 | Casp1 | Caspase-1 | 510.12 | 29.85 | 45640 | 9 | 11996941 | 11714341 | 14524062 | 15720257 | 15465249 | 14250604 | 13376620 | 17806184 | 17472866 | 17139972 | 15690750 | 17383998 | 15615875 | 12510529 | 15704604 | 11783179 | 8552904 | 14664807 | 14812557 | 14777534 | 6 | 10 | 9 | 6 | 9 | 10 | 1.107819279 | 0.960766555 | 0.930426248 | 0.806920802 |
| P70677 | Casp3 | Caspase-3 | 263.4 | 22.02 | 31475 | 5 | 4880557 | 5311360 | 3660925.25 | 6660492 | 8627440 | 5433385 | 3741856 | 5843686 | 5199035.5 | 5242537 | 3773760.25 | 5904757.5 | 5605546 | 3912849.75 | 5515838 | 3950043.75 | 5830192 | 4321372 | 2385021.75 | 4869058 | 11 | 6 | 10 | 7 | 11 | 7 | 0.848047183 | 0.970631055 | 0.732845576 | 0.83877724 |
| P97864 | Casp7 | Caspase-7 | 48.8 | 3.3 | 34061 | 1 | 819535.62 | 1168690.75 | 1092761.75 | 663838.25 | 2 | 1024731.69 | 998504.12 | 702627.88 | 942937.44 | 310417.59 | 511214 | 839764.5 | 752257.81 | 854774.62 | 677666.69 | 774951.06 | 654607.88 | 629693.38 | 2 | 922090.94 | 11 | 7.5 | 8 | 6 | 11 | 7.5 | 0.970852937 | 0.913666193 | 0.796123337 | 0.749228798 |
| O89110 | Casp8 | Caspase-8 | 238.4 | 9.79 | 55357 | 4 | 6704277 | 7112024.5 | 7250546.5 | 7527506.5 | 7648271.5 | 6182639.5 | 7790562 | 7713102 | 7300013.5 | 7314465.5 | 3254475.25 | 5075877 | 5196326 | 1477953.38 | 6546232.5 | 8036769 | 4251604 | 6537527.5 | 5975522.5 | 7179114.5 | 0 | 7 | 1 | 7 | 1 | 7 | 0.594627556 | 0.59367492 | 0.882401223 | 0.880987552 |
| P51125 | Cast | Calpastatin | 269.72 | 8.88 | 84922 | 4 | 3127957 | 4331275.5 | 5137537 | 5167401.5 | 6031069.5 | 3742020.5 | 5385085 | 5374164.5 | 6012679.5 | 5388009.5 | 2620520 | 2997422 | 2243961.75 | 2436643.75 | 1861749.25 | 4736277 | 4154599.25 | 5415365 | 4734693.5 | 4932092.5 | 0 | 11 | 0 | 8 | 0 | 11 | 0.511039035 | 0.469474017 | 1.007471526 | 0.925529503 |
| P24270 | Cat | Catalase | 586.62 | 27.51 | 59795 | 10 | 1015076.38 | 1016389.5 | 1230331 | 1248781.88 | 1346897.88 | 1443785.5 | 2379220.25 | 2021463.38 | 2010364.75 | 1978505.12 | 3345282 | 5556524 | 4422586 | 3836579.5 | 5190314.5 | 2830377.25 | 1359751.88 | 2928296 | 3977639.75 | 2898963.5 | 0 | 0 | 0 | 5 | 0 | 5 | 3.815855764 | 2.273010826 | 2.389258932 | 1.423222405 |
| Q08024 | Cbfb | Core-binding factor subunit beta | 192.36 | 26.2 | 22030 | 3 | 2462940.75 | 2468117.25 | 1892552.62 | 1680925.75 | 2369409.5 | 1200248 | 2292090.75 | 1877280.12 | 2210026.75 | 1167125.25 | 512553.47 | 1919288 | 2 | 2 | 1061881.88 | 1927835.38 | 1221797.75 | 943255.75 | 858565.06 | 1235551.12 | 2 | 2 | 3 | 7 | 3 | 7 | 0.321293429 | 0.399430533 | 0.568975157 | 0.707347334 |
| P22682 | Cbl | E3 ubiquitin-protein ligase CBL | 57.5 | 1.64 | 100564 | 1 | 2 | 2 | 2 | 2 | 2 | 392315.75 | 2 | 2 | 424867.56 | 719811.56 | 894043.44 | 888159.38 | 853576.31 | 1230390.25 | 790115.19 | 2 | 2 | 552526.38 | 724146.62 | 565990.31 | 0 | 5 | 0 | 10 | 0 | 10 | 465628.457 | 3.029465188 | 184266.731 | 1.198873562 |
| P48758 | Cbr1 | Carbonyl reductase [NADPH] 1 | 221.31 | 15.88 | 30641 | 5 | 2 | 991837.88 | 624977.88 | 868902.94 | 553920.88 | 620139.81 | 594271.88 | 591586.12 | 660429.75 | 582010.12 | 2 | 532189 | 530408.56 | 783789.69 | 438679.25 | 2 | 635388.12 | 735752.38 | 561637.56 | 817985.12 | 6.5 | 11.5 | 5 | 11 | 6.5 | 11.5 | 0.751755903 | 0.749586752 | 0.904963664 | 0.90235244 |
| Q8K354 | Cbr3 | Carbonyl reductase [NADPH] 3 | 41.84 | 3.97 | 30953 | 1 | 723297.19 | 797801.12 | 350954.75 | 431311.88 | 412297.69 | 943404.31 | 2 | 657069.69 | 554684.5 | 782372.25 | 942119.75 | 910504.94 | 823620.25 | 893761.69 | 704873.81 | 1026945.75 | 1060072.38 | 954834.44 | 812542 | 920030.38 | 2 | 0 | 6 | 2 | 6 | 2 | 1.574157406 | 1.455262223 | 1.758106805 | 1.625318033 |
| Q7TS56 | Cbr4 | Carbonyl reductase family member 4 | 102.59 | 8.05 | 25287 | 1 | 568247.06 | 780874.12 | 543262.12 | 695135.94 | 566350.06 | 2 | 717087.38 | 530505.44 | 453963.5 | 734855.19 | 479897.94 | 324699.06 | 516620.44 | 2 | 474756.72 | 756324.69 | 1213676.25 | 787501.94 | 793367.44 | 766045.75 | 0 | 2 | 7.5 | 0 | 7.5 | 2 | 0.569451676 | 0.737139304 | 1.368768221 | 1.771832266 |
| Q8VEH6 | Cbwd1 | COBW domain-containing protein 1 | 159.54 | 9.16 | 43772 | 2 | 174366.45 | 96391.29 | 162194.31 | 102740.75 | 2 | 229284.47 | 177719.64 | 203851.08 | 209489.56 | 200448.36 | 441313.59 | 598301.56 | 585462.88 | 357843.06 | 619200.38 | 167533.69 | 2 | 2 | 95171.94 | 245965.44 | 0 | 11 | 0 | 5 | 0 | 11 | 4.857470093 | 2.549117392 | 0.949561336 | 0.498313581 |
| P83917 | Cbx1 | Chromobox protein homolog 1 | 411.36 | 28.11 | 21418 | 6 | 7805867.5 | 8413875 | 9155328 | 9662774 | 9106639 | 8163976.5 | 8869478 | 8710416 | 6845292 | 8591863 | 4180597.75 | 3956777 | 4060838.5 | 4783398.5 | 4579744 | 9722690 | 11320977 | 7190009.5 | 8184884.5 | 7555478 | 0 | 11 | 0 | 11 | 0 | 11 | 0.488426957 | 0.523575008 | 0.99613894 | 1.067822825 |
| P23198 | Cbx3 | Chromobox protein homolog 3 | 736.21 | 42.08 | 20855 | 10 | 15397651 | 14285073 | 16507842 | 17742556 | 15173822 | 11632485 | 14064014 | 14376407 | 12790562 | 12511768 | 10257938 | 9489630 | 7177663 | 7514349 | 8341837.5 | 10456609 | 16212969 | 11878497 | 12867423 | 11522276 | 0 | 3 | 0 | 9 | 0 | 9 | 0.540804831 | 0.654397905 | 0.795603658 | 0.962715821 |
| Q61686 | Cbx5 | Chromobox protein homolog 5 | 262.84 | 19.37 | 22186 | 4 | 2025084.25 | 2683737.5 | 5088881 | 3586381 | 3396031.5 | 2932006 | 3068397.25 | 3600424.75 | 2997733 | 3293764.75 | 2204987.5 | 2645630.5 | 1109065.38 | 1504620.75 | 1821255.75 | 2163299.5 | 3095867 | 3661661.5 | 3137389 | 2359425.25 | 2 | 10 | 0 | 11 | 2 | 11 | 0.553366872 | 0.584279483 | 0.859209966 | 0.907207823 |
| Q8CH18 | Ccar1 | Cell division cycle and apoptosis regulator protein 1 | 90.62 | 1.57 | 132060 | 2 | 1953924 | 2343466 | 2349308 | 1519248.38 | 1277113.38 | 2524620 | 2177548.75 | 2496702.25 | 2491288.75 | 2504975.5 | 2438428.75 | 2392261.25 | 2565113.5 | 1895756.5 | 2285799.5 | 1886686.25 | 2223858.5 | 2875043 | 2881425.75 | 1428086.38 | 5 | 9 | 8 | 11 | 8 | 11 | 1.226017816 | 0.949342444 | 1.196127121 | 0.92619718 |
| Q8VDP4 | Ccar2 | Cell cycle and apoptosis regulator protein 2 | 250.74 | 8.24 | 103002 | 5 | 2323680 | 2261175.25 | 2173934 | 2850118 | 2670660 | 2255687.75 | 2361709.25 | 1950865.88 | 2554406.75 | 1564996.75 | 462239.41 | 1101663 | 1647431.5 | 179453.59 | 1230882.38 | 2058792.12 | 1810425.75 | 1317677.25 | 1962819.5 | 2876560.5 | 0 | 5 | 1 | 10 | 1 | 10 | 0.376370745 | 0.432430216 | 0.816500689 | 0.938116401 |
| Q8VE99 | Ccdc115 | Coiled-coil domain-containing protein 115 | 55.57 | 8.33 | 19743 | 1 | 2 | 2 | 532941.88 | 353791.28 | 108989.73 | 2 | 2 | 2 | 2 | 2 | 2 | 415867.56 | 2 | 2 | 2 | 2 | 2 | 2 | 342904.03 | 2 | 8 | 7 | 10 | 10 | 10 | 10 | 0.417660268 | 41587.556 | 0.344383619 | 34291.203 |
| P86182 | Ccdc22 | Coiled-coil domain-containing protein 22 | 60.99 | 2.07 | 70855 | 1 | 136174.86 | 190422.66 | 2 | 2 | 122638.77 | 106700.31 | 137919.83 | 117610.15 | 2 | 2 | 303310 | 287356.28 | 256112.25 | 2 | 252160.84 | 2 | 2 | 2 | 2 | 2 | 4 | 5 | 4 | 5 | 4 | 5 | 2.446221754 | 3.033786144 | 2.22598E-05 | 2.760644E-05 |
| Q5U2X6 | Ccdc47 | Coiled-coil domain-containing protein 47 | 579.45 | 23.19 | 55733 | 9 | 7805034 | 6170764 | 6364384.5 | 7851719 | 5837322 | 7950260.5 | 7345815 | 4617109 | 6891481 | 7502878 | 7226651 | 6571964 | 7316931 | 7417080.5 | 7228120 | 6645426 | 10432861 | 7511843.5 | 7483317.5 | 6342074.5 | 10 | 9 | 10 | 11 | 10 | 11 | 1.050883412 | 1.042358119 | 1.128898004 | 1.119739818 |
| Q3URS9 | Ccdc51 | Mitochondrial potassium channel | 42.33 | 2.71 | 45132 | 1 | 148176.33 | 212491.25 | 2 | 261195.73 | 146049.83 | 250172.66 | 176588.73 | 321704.94 | 239003.53 | 243864.48 | 401227.12 | 349304.19 | 310800.06 | 271989.75 | 357688 | 227905.56 | 284536.88 | 238637.48 | 340868.94 | 293878.47 | 0 | 2 | 2 | 10 | 2 | 10 | 2.20207811 | 1.373314351 | 1.804662075 | 1.125467946 |
| Q9JJ89 | Ccdc86 | Coiled-coil domain-containing protein 86 | 124.16 | 10.09 | 46487 | 3 | 900194.5 | 1153214.5 | 930372.25 | 526486 | 443031.09 | 1079218.75 | 1004569.81 | 610488.81 | 994999.62 | 566758.12 | 1522732.5 | 1854703.5 | 1310629.88 | 799534.62 | 1149973.25 | 971875.69 | 317525.62 | 1078837.38 | 2 | 1097382.25 | 4 | 12 | 3 | 11 | 4 | 12 | 1.678996417 | 1.559567433 | 0.876640881 | 0.814284387 |
| Q4QRL3 | Ccdc88b | Coiled-coil domain-containing protein 88B | 101.54 | 1.28 | 166608 | 2 | 2027151.75 | 1981819.12 | 2343968.5 | 2878678 | 2310498 | 2092833.38 | 1970831 | 803781.81 | 2840269.5 | 2377849.5 | 1203866 | 1397363.5 | 2414919 | 1617602.5 | 1027529.69 | 1870676.62 | 1293550.75 | 2240933 | 2187750 | 1899222.25 | 4 | 4 | 8 | 9 | 8 | 9 | 0.663767468 | 0.759628295 | 0.822391071 | 0.941160207 |
| P51670 | Ccl9 | C-C motif chemokine 9 | 76.53 | 17.21 | 13871 | 2 | 2 | 2 | 2 | 2 | 2 | 2 | 2 | 2 | 2 | 2 | 768454.06 | 473038.34 | 2 | 361407.91 | 524812.62 | 2 | 2 | 2 | 2 | 2 | 2.5 | 12.5 | 2.5 | 12.5 | 2.5 | 12.5 | 212771.493 | 212771.493 | 1 | 1 |
| O88874 | Ccnk | Cyclin-K | 48.41 | 4.33 | 61363 | 1 | 2 | 2 | 2 | 2 | 2 | 2 | 2 | 2 | 2 | 2 | 2 | 2 | 2 | 2 | 474553.09 | 2 | 2 | 2 | 2 | 2 | 10 | 12.5 | 10 | 12.5 | 10 | 12.5 | 47456.109 | 47456.109 | 1 | 1 |
| Q9WU84 | Ccs | Copper chaperone for superoxide dismutase | 122.34 | 8.03 | 28912 | 2 | 3169196 | 3745303.75 | 3702982.5 | 4573268 | 4706114 | 3486547.25 | 3555463.5 | 2670772.25 | 3659285.5 | 5210381 | 1896295.75 | 2059991.38 | 2280457.25 | 2863105.25 | 2 | 3474351 | 2367599.5 | 3356016.5 | 3420140.5 | 3407005.25 | 0 | 4 | 1 | 4 | 1 | 4 | 0.457351044 | 0.489701405 | 0.80540896 | 0.862378921 |
| P80314 | Cct2 | T-complex protein 1 subunit beta | 2544.33 | 69.16 | 57477 | 37 | 56762166 | 64877634 | 69440346 | 58687410 | 64747546 | 63375050 | 56664182 | 69936434 | 69255034 | 64379166 | 95542666 | 86454266 | 81077826 | 83109146 | 72384178 | 64904890 | 73764274 | 71461770 | 69159962 | 66055374 | 0 | 3 | 0 | 6 | 0 | 6 | 1.330836196 | 1.293434243 | 1.098027623 | 1.067168545 |
| P80318 | Cct3 | T-complex protein 1 subunit gamma | 1878.19 | 57.61 | 60630 | 33 | 164318450 | 167559762 | 158639618 | 152439490 | 143605010 | 182509266 | 169167938 | 159081362 | 137153986 | 153562786 | 207755090 | 207883170 | 172882914 | 164882162 | 160471362 | 174939874 | 195712674 | 137971954 | 179116674 | 164744450 | 3 | 6 | 5 | 8 | 5 | 8 | 1.161859224 | 1.140240572 | 1.083811916 | 1.063645487 |
| P80315 | Cct4 | T-complex protein 1 subunit delta | 2252.59 | 71.43 | 58066 | 34 | 192388402 | 200680338 | 210853794 | 211036002 | 186079410 | 194990290 | 184808194 | 199630946 | 200083634 | 194008866 | 223421746 | 211773954 | 207268098 | 191843970 | 196663474 | 204393826 | 211377634 | 205277346 | 217870114 | 212157698 | 9 | 4 | 6 | 0 | 9 | 4 | 1.029902259 | 1.059011831 | 1.049986788 | 1.079664038 |
| P80316 | Cct5 | T-complex protein 1 subunit epsilon | 1588.6 | 55.27 | 59624 | 26 | 104454626 | 108642978 | 116604354 | 125296234 | 111454250 | 102842466 | 103676882 | 112351866 | 117870346 | 111255482 | 102561242 | 106093130 | 109526354 | 96813834 | 104288610 | 121286410 | 108184306 | 116819450 | 117899002 | 117132410 | 3 | 8 | 6 | 5 | 6 | 8 | 0.916728628 | 0.947602141 | 1.026249575 | 1.060811525 |
| P80317 | Cct6a | T-complex protein 1 subunit zeta | 1108.26 | 34.09 | 58004 | 17 | 96271530 | 84837266 | 94037386 | 95747938 | 91566642 | 104934242 | 97543874 | 93814290 | 94388266 | 95498754 | 122256258 | 104015354 | 94506410 | 82422602 | 80759218 | 98525506 | 109654722 | 97038626 | 94744338 | 100186738 | 12 | 2 | 11 | 7 | 12 | 7 | 1.046488441 | 0.99543464 | 1.081497007 | 1.028735284 |
| P80313 | Cct7 | T-complex protein 1 subunit eta | 2304.58 | 64.71 | 59652 | 33 | 100202282 | 113481114 | 115497986 | 124026946 | 103497242 | 112411538 | 100522530 | 111683706 | 99555314 | 115008602 | 132479162 | 117021098 | 124916234 | 129594778 | 130966026 | 121111114 | 131870674 | 130732914 | 133505162 | 128493874 | 1 | 1 | 0 | 0 | 1 | 1 | 1.140598069 | 1.177668511 | 1.159883739 | 1.197580982 |
| P42932 | Cct8 | T-complex protein 1 subunit theta | 2337.17 | 63.5 | 59555 | 38 | 164662226 | 173322962 | 203532722 | 188340882 | 175855346 | 167020530 | 162797378 | 177385906 | 165263954 | 175177330 | 162433890 | 199159202 | 171964546 | 149500306 | 177289986 | 180882258 | 165951378 | 175518402 | 186732818 | 191598050 | 8 | 12 | 10 | 4 | 12 | 12 | 0.949911119 | 1.014986027 | 0.994445011 | 1.062570772 |
| P10810 | Cd14 | Monocyte differentiation antigen CD14 | 981.85 | 39.89 | 39204 | 14 | 44822970 | 23070258 | 37402338 | 46775102 | 44336574 | 34790202 | 30279684 | 26963786 | 29876832 | 43874150 | 33514534 | 17662988 | 24579410 | 20153086 | 28354556 | 29748810 | 34525142 | 38671374 | 35704698 | 36633558 | 3 | 6 | 4 | 9 | 4 | 9 | 0.632688351 | 0.749554141 | 0.892449689 | 1.057296787 |
| Q62192 | Cd180 | CD180 antigen | 766.37 | 23.9 | 74302 | 12 | 4437875 | 4176012.5 | 4370339 | 4475864 | 5431020 | 3668693.5 | 4558684.5 | 5223076.5 | 5609415.5 | 4645561.5 | 5108080.5 | 5687245.5 | 6183710 | 4660805 | 6922632.5 | 3729678.75 | 3061317.5 | 3350638.75 | 4287459.5 | 4361064 | 2 | 2 | 4 | 3 | 4 | 3 | 1.247753948 | 1.204891525 | 0.820849583 | 0.792652034 |
| Q9JLQ0 | Cd2ap | CD2-associated protein | 135.81 | 5.18 | 70450 | 3 | 3164551.75 | 3150614.25 | 3373515.75 | 2522259 | 1638692 | 2793771 | 3527361 | 3103920 | 3304581.5 | 2406082 | 1343720.75 | 1774949.25 | 1466846.75 | 960120.69 | 1743996 | 3146648.25 | 2397158.5 | 1897132.75 | 2306739 | 1984848 | 2 | 6 | 0 | 3 | 2 | 6 | 0.526341281 | 0.481618027 | 0.84713629 | 0.775155063 |
| Q9CWK3 | Cd2bp2 | CD2 antigen cytoplasmic tail-binding protein 2 | 116.3 | 7.02 | 37694 | 2 | 651518.12 | 1047302 | 663270.38 | 664443.81 | 409822.41 | 486007.09 | 398941.78 | 452900.88 | 2 | 633913.44 | 1324484.75 | 1229371.88 | 491005.75 | 576757.06 | 495748.44 | 2 | 703877.31 | 984076 | 339777.59 | 604273.75 | 12 | 9 | 3 | 9.5 | 12 | 9.5 | 1.19817825 | 2.088163388 | 0.765929403 | 1.334847914 |
| P15379 | Cd44 | CD44 antigen | 393.52 | 9.13 | 85617 | 6 | 18788770 | 18795434 | 14483571 | 15547624 | 14793470 | 19376338 | 18380050 | 15440320 | 16861716 | 14923534 | 16258136 | 15578939 | 17590092 | 15496937 | 7480847.5 | 16815444 | 16032987 | 13187524 | 9565056 | 12528308 | 11 | 6 | 9 | 4 | 11 | 6 | 0.8786063 | 0.852003804 | 0.826723141 | 0.801691566 |
| Q61735 | Cd47 | Leukocyte surface antigen CD47 | 274.14 | 12.21 | 33098 | 4 | 6876034 | 4850732 | 5177152 | 6004214 | 7596992.5 | 5734179 | 7360339 | 4370006.5 | 6900699 | 6840532 | 2047302.75 | 3407040.5 | 5516295.5 | 2092879.75 | 4898735 | 6061727 | 3002737 | 3622171 | 3139883.75 | 3936099 | 3 | 3 | 2 | 2 | 3 | 3 | 0.588827412 | 0.57560707 | 0.647845832 | 0.63330041 |
| P13379 | Cd5 | T-cell surface glycoprotein CD5 | 422.55 | 19.84 | 53849 | 7 | 3854641.75 | 3472596.25 | 10185818 | 10390520 | 6003488 | 5746199 | 5106958 | 11695387 | 9467299 | 12318346 | 1808639.38 | 1450387.38 | 5091827 | 6637368 | 3355973 | 3743937.25 | 5490896.5 | 9173602 | 8982014 | 7775877 | 5 | 12 | 2 | 7 | 5 | 12 | 0.541013954 | 0.413770843 | 1.037138655 | 0.793210106 |
| P31996 | Cd68 | Macrosialin | 223.3 | 16.87 | 34818 | 5 | 130423794 | 131125242 | 101951330 | 117243442 | 109782354 | 143972674 | 121569034 | 110559586 | 119152986 | 141321970 | 102972922 | 82800450 | 100146946 | 112507394 | 111341354 | 117535170 | 125624178 | 143413730 | 144403026 | 140143698 | 5 | 4 | 2 | 9 | 5 | 9 | 0.863245524 | 0.800798123 | 1.136477679 | 1.054264594 |
| P21855 | Cd72 | B-cell differentiation antigen CD72 | 302.28 | 20.06 | 40347 | 6 | 1974123 | 2726857.25 | 2814198 | 2878806.75 | 2861679.75 | 1990908.38 | 2714641.75 | 3081007.5 | 1920546.75 | 2116273 | 5685576.5 | 4950914 | 5794479 | 5462664.5 | 5506759 | 2771005.75 | 2558361.5 | 4043862.25 | 3980384 | 3514743.75 | 0 | 7 | 0 | 3 | 0 | 7 | 2.067070458 | 2.317475973 | 1.272539519 | 1.426695326 |
| P35762 | Cd81 | CD81 antigen | 95.11 | 9.75 | 25815 | 1 | 2 | 2 | 2 | 2 | 233892.83 | 2 | 150124.03 | 309494.28 | 195581.16 | 2 | 473126.31 | 2 | 829407.94 | 902941.25 | 2 | 2 | 2 | 2 | 2 | 243316.42 | 6 | 12 | 8 | 8 | 8 | 12 | 9.429122163 | 3.366098626 | 1.040288827 | 0.371372301 |
| P40237 | Cd82 | CD82 antigen | 67.24 | 4.14 | 29629 | 1 | 504902.91 | 274562.44 | 2 | 2 | 2 | 492079.59 | 378797.22 | 209738.42 | 364374.88 | 2 | 1528681.12 | 1270900.5 | 1804171.62 | 1707588.62 | 1965921.75 | 709602.69 | 2 | 294829.88 | 2 | 2 | 0 | 11.5 | 0 | 8.5 | 0 | 11.5 | 10.61907357 | 5.728241388 | 1.288615124 | 0.695116993 |
| Q18PI6 | Cd84 | SLAM family member 5 | 54.69 | 3.34 | 37378 | 1 | 1302865.62 | 1638518 | 433313.34 | 827805.12 | 663563.06 | 1608090 | 1448275.88 | 574646.81 | 818320.62 | 612224.75 | 650088.06 | 448481.97 | 437090.22 | 384764.75 | 325667.34 | 1526399.75 | 2248567.25 | 809274.31 | 888675.88 | 981513 | 3 | 8 | 2 | 8 | 3 | 8 | 0.461582876 | 0.443755127 | 1.326416726 | 1.275186437 |

| | | | | | | | | | | | | | | | | | | | | | | | | | | | | | | | | | |
|---|---|---|---|---|---|---|---|---|---|---|---|---|---|---|---|---|---|---|---|---|---|---|---|---|---|---|---|---|---|---|---|---|---|
| P42082 | Cd86 | T-lymphocyte activation antigen CD86 | 52.25 | 4.53 | 34666 | 1 | 1068954.12 | 1366131.38 | 1571653.88 | 1339316.62 | 1673176.12 | 1313359.5 | 1445239.38 | 2 | 1775405.5 | 1832918.5 | 1934949.12 | 2856797.5 | 2557349.5 | 1173975.62 | 3256004 | 985968.75 | 2 | 1417560.25 | 1405367.88 | 1412496.25 | 4 | 9 | 4 | 7.5 | 4 | 9 | 1.678114577 | 1.850041576 | 0.743869848 | 0.820081158 |
| P40240 | Cd9 | CD9 antigen | 140.35 | 16.37 | 25258 | 2 | 2 | 2 | 542973.62 | 354011.75 | 1492847.75 | 2 | 590755.38 | 1081013.38 | 1219159.5 | 1147816 | 1230283 | 1786222.75 | 2162968 | 1529887.12 | 1717307.25 | 2 | 2 | 1564702.5 | 1345326.38 | 1363522.12 | 1 | 10 | 0 | 9 | 1 | 10 | 3.526042863 | 2.086456434 | 1.788220195 | 1.058139018 |
| Q8CII2 | Cdc123 | Cell division cycle protein 123 homolog | 169.24 | 13.99 | 38816 | 3 | 1030981.12 | 1358507.38 | 719521.31 | 729913 | 1184680 | 1144989.62 | 1244999.25 | 293455.31 | 991000.06 | 521377.75 | 962193.81 | 1037081.38 | 456655 | 137821.08 | 949246.62 | 1046318.38 | 1298152.5 | 687407.94 | 610241.88 | 846882.12 | 7 | 9 | 8 | 11 | 8 | 11 | 0.705270306 | 0.844410916 | 0.893582353 | 1.069874468 |
| Q61081 | Cdc37 | Hsp90 co-chaperone Cdc37 | 423.54 | 18.47 | 44593 | 6 | 7332705.5 | 8025065 | 9942470 | 9292617 | 12655509 | 7784860.5 | 8732776 | 9836713 | 12069622 | 9425300 | 5231879.5 | 6011082.5 | 4728049.5 | 4351062 | 5163146 | 9370558 | 10353200 | 8327007.5 | 8593367 | 9637675 | 0 | 11 | 0 | 11 | 0 | 11 | 0.539388372 | 0.532614577 | 0.979543018 | 0.967241633 |
| P60766 | Cdc42 | Cell division control protein 42 homolog | 547.11 | 48.69 | 21259 | 7 | 21899460 | 23785862 | 22278164 | 29773068 | 22477274 | 27670322 | 20865454 | 23521956 | 23168952 | 23629832 | 19090246 | 18736476 | 18718286 | 11450051 | 16932422 | 26588534 | 37559450 | 24744980 | 25928838 | 24017224 | 0 | 4 | 0 | 4 | 0 | 4 | 0.706470149 | 0.714537863 | 1.154933907 | 1.168122966 |
| Q6A068 | Cdc5l | Cell division cycle 5-like protein | 768.38 | 20.57 | 92190 | 13 | 7775831.5 | 7235925.5 | 6064023 | 5808410 | 6076182.5 | 9822750 | 8911626 | 6008874 | 5669278 | 6785833 | 10406056 | 7958027.5 | 6454354 | 6992788 | 6638850 | 4664027 | 6333716 | 6596286.5 | 4832125 | 5642043.5 | 6 | 6 | 10 | 4 | 10 | 6 | 1.166554641 | 1.033649722 | 0.851574053 | 0.754554697 |
| Q4V8C8 | Cdc73 | Parafibromin | 328.82 | 15.44 | 60603 | 7 | 1210095.75 | 2090063.75 | 1224152.12 | 1027044.12 | 598324.88 | 2064874.25 | 2059908 | 888936.25 | 1427338.75 | 1606827.5 | 7694914.5 | 7987646 | 7272199 | 2378364.75 | 2353211.25 | 285681.22 | 1185893.12 | 233703.89 | 996589.5 | 233024.5 | 0 | 3 | 0 | 2 | 0 | 3 | 4.502076971 | 3.440200296 | 0.477243033 | 0.36467871 |
| Q8VDP6 | Cdipt | CDP-diacylglycerol--inositol 3-phosphatidyltransferase | 332.35 | 34.74 | 23599 | 6 | 1292918.5 | 1446556.5 | 2010187.75 | 1894758.5 | 1877103.25 | 1411208.75 | 2116336.25 | 1869134.75 | 1761355.88 | 1772729 | 2581199.5 | 2403590.75 | 4043855 | 4379123 | 3077969.5 | 1526571.25 | 1784353 | 2228243.25 | 2356180.5 | 2231270 | 0 | 6 | 0 | 6 | 0 | 6 | 1.934599584 | 1.845949192 | 1.188357553 | 1.133902686 |
| P11440 | Cdk1 | Cyclin-dependent kinase 1 | 423.31 | 34.34 | 34107 | 9 | 13342826 | 12049056 | 13427033 | 12347816 | 11275599 | 11762978 | 10537082 | 11491982 | 10426408 | 10732894 | 6934705.5 | 8777903 | 6981411 | 6292311 | 6694706.5 | 13258054 | 10062064 | 8526360 | 7900430 | 7271599 | 0 | 3 | 0 | 5 | 0 | 5 | 0.571423856 | 0.649320552 | 0.752990912 | 0.85563889 |
| Q04899 | Cdk18 | Cyclin-dependent kinase 18 | 211.87 | 7.54 | 51848 | 3 | 178587.28 | 309266.94 | 257543.59 | 186507.52 | 231715.66 | 194244.91 | 259092.8 | 326017.66 | 324296.69 | 254981 | 540974.31 | 690629.25 | 570322.06 | 331825.75 | 594809.94 | 223020.81 | 161912.7 | 385973.56 | 280306.88 | 299368.03 | 0 | 10 | 0 | 12 | 0 | 12 | 2.344888356 | 2.008313643 | 1.160671724 | 0.994074132 |
| P49615 | Cdk5 | Cyclin-dependent-like kinase 5 | 163.89 | 10.62 | 33288 | 4 | 3576788 | 3394274 | 2517204.25 | 3917062.75 | 3448207.5 | 3673698.5 | 2959639.25 | 2925539.75 | 2985168.5 | 3059804.5 | 2525073.75 | 2075965 | 2711580.5 | 2722890 | 1591421.25 | 4084771.5 | 4821827.5 | 3766467.75 | 2683098.75 | 3331877 | 3 | 9 | 0 | 6 | 3 | 9 | 0.689880756 | 0.745132139 | 1.108849914 | 1.197655829 |
| Q99LM2 | Cdk5rap3 | CDK5 regulatory subunit-associated protein 3 | 157.67 | 6.36 | 56991 | 3 | 201112.89 | 207598.59 | 285956.34 | 2 | 502622.69 | 2 | 408613.91 | 2 | 331613.84 | 2 | 36134.99 | 336070.5 | 139788.42 | 43291.73 | 1185403.75 | 243648.77 | 102442.14 | 193962.17 | 92331.17 | 188436.89 | 12 | 7 | 7 | 10 | 12 | 10 | 1.453854739 | 2.351540159 | 0.685564416 | 1.108867489 |
| Q64261 | Cdk6 | Cyclin-dependent kinase 6 | 130.25 | 14.72 | 37028 | 3 | 920936 | 923715.75 | 1641359.62 | 1265812 | 744366.75 | 677744.31 | 764081.19 | 1026684.94 | 922965.75 | 1169281.88 | 408242.97 | 787506.31 | 1179779.75 | 397354.97 | 660013.12 | 838173.19 | 519976.22 | 1223241.75 | 689027.94 | 996457.12 | 4 | 7 | 7 | 11 | 7 | 11 | 0.624595774 | 0.752703184 | 0.776333447 | 0.935562938 |
| Q641Z4 | Cdk9 | Cyclin-dependent kinase 9 | 91.4 | 6.45 | 42762 | 2 | 2 | 115545.57 | 2 | 2 | 262900.06 | 2 | 2 | 2 | 78078.34 | 169588.67 | 465585.06 | 367857.44 | 434891.31 | 353494.59 | 376377.28 | 2 | 2 | 2 | 2 | 2 | 0 | 7.5 | 0 | 7.5 | 0 | 7.5 | 5.279950175 | 8.067918866 | 2.642346E-05 | 4.037582E-05 |
| Q5RK03 | Cdkn2aipnl | CDKN2AIP N-terminal-like protein | 53.03 | 8.62 | 13197 | 1 | 1694597.62 | 974687.06 | 963982.81 | 2 | 743701.56 | 2 | 1422638.38 | 1066881 | 2 | 2 | 2 | 2 | 348170.94 | 585019.06 | 2 | 1441775.5 | 1745355.62 | 2 | 2 | 2 | 3.5 | 10.5 | 10.5 | 10.5 | 10.5 | 10.5 | 0.213205888 | 0.37484896 | 0.72816043 | 1.28021877 |
| Q99L43 | Cds2 | Phosphatidate cytidylyltransferase 2 | 143.05 | 8.11 | 51314 | 2 | 2 | 2 | 2 | 2 | 494044.62 | 2 | 566433.38 | 773626.75 | 767878.81 | 881069.88 | 1371127 | 1584039.75 | 1696750.5 | 1849520.25 | 1681232.75 | 2 | 2 | 164354.97 | 1401031.5 | 680030.88 | 0 | 7 | 0 | 9 | 0 | 9 | 16.56234563 | 2.737584687 | 4.544903233 | 0.751225568 |
| Q4VAA2 | Cdv3 | Protein CDV3 | 215.54 | 18.86 | 29729 | 3 | 3035774 | 3331059 | 2842529 | 3883012 | 2955565 | 3486896.5 | 2878235.25 | 3542792.5 | 3258618.25 | 4275869.5 | 3504389 | 3555693.25 | 2752858.75 | 2973024 | 2754446.75 | 3902453 | 5055901 | 4400552.5 | 3951605.5 | 4460441.5 | 10 | 0 | 8 | 2 | 10 | 2 | 0.968374303 | 0.890955434 | 1.356619906 | 1.248161865 |
| P28659 | Celf1 | CUGBP Elav-like family member 1 | 160.88 | 6.58 | 52107 | 3 | 865261.88 | 734913.44 | 745036.25 | 872370.88 | 600001.44 | 906230.38 | 581172.19 | 663452.06 | 734091.81 | 627396.31 | 1389668 | 939876.62 | 979234.5 | 899674.75 | 872242.56 | 880577.31 | 1094080.62 | 772794.38 | 978676.38 | 937467.69 | 1 | 2 | 2 | 2 | 2 | 2 | 1.330867003 | 1.446526376 | 1.221609404 | 1.327773715 |
| Q792H5 | Celf2 | CUGBP Elav-like family member 2 | 183.61 | 7.87 | 54271 | 4 | 2213445 | 2090790.62 | 3250341.75 | 2874037.5 | 2050998.38 | 1770122.38 | 2346200.75 | 2366365.75 | 2699346.25 | 2328005 | 2570252.5 | 1999701.75 | 1630422.12 | 2053102.5 | 1430791.12 | 1809009.75 | 2041641 | 2423808.75 | 1942319.25 | 2271561 | 4 | 6 | 6 | 8 | 6 | 8 | 0.77600722 | 0.841375867 | 0.840437884 | 0.911233986 |
| Q6A065 | Cep170 | Centrosomal protein of 170 kDa | 41.03 | 1.01 | 175050 | 1 | 2 | 452829.25 | 484117.47 | 2 | 2 | 595771.69 | 532372.12 | 569972.38 | 651872.69 | 691226.44 | 368581.75 | 418808.38 | 2 | 2 | 443493.03 | 2 | 2 | 525146.12 | 729453.81 | 838789.06 | 12 | 7 | 0 | 10 | 12 | 10 | 1.313713204 | 0.404735289 | 2.234256804 | 0.688340933 |
| Q8BGS7 | Cept1 | Choline/ethanolaminephosphotransferase 1 | 164.46 | 9.62 | 46434 | 3 | 313879.72 | 282369.75 | 161611.38 | 156998.12 | 133812.2 | 257531.56 | 650846.62 | 349210.75 | 432505.62 | 511876.25 | 1977661.25 | 2127569.5 | 2015721.25 | 2064738.25 | 1956105.62 | 154077.41 | 2 | 242225.23 | 2 | 353603.53 | 0 | 9 | 0 | 2 | 0 | 9 | 9.671092484 | 4.605781271 | 0.715105165 | 0.340563176 |
| Q924Z4 | Cers2 | Ceramide synthase 2 | 108.97 | 9.47 | 45024 | 2 | 272612.88 | 133682.48 | 366419 | 602650.38 | 854853.75 | 2 | 2 | 861838.5 | 153301.5 | 577451 | 1102118.5 | 1426117.38 | 2419867.5 | 1652745.5 | 2182233 | 183355.59 | 873208.75 | 780294.81 | 150521.55 | 78248.88 | 0 | 11 | 0 | 9 | 0 | 11 | 3.938215883 | 5.514950053 | 0.926200545 | 1.297021264 |
| Q9D6J1 | Cers4 | Ceramide synthase 4 | 39.79 | 2.04 | 46017 | 1 | 2 | 273247.97 | 291680.91 | 560015.19 | 2 | 459046.41 | 320864.56 | 440955.06 | 2 | 600065 | 903536.06 | 672312.81 | 1027753.06 | 1234819.38 | 1142286.25 | 2 | 2 | 652073.31 | 795254.31 | 508848.66 | 0 | 9 | 0 | 10 | 0 | 10 | 4.427499982 | 2.735250269 | 1.738907184 | 1.074273599 |
| Q9D6K9 | Cers5 | Ceramide synthase 5 | 237.36 | 14.49 | 48167 | 5 | 652969.5 | 1257981.62 | 2169277.5 | 1404197.62 | 1498431.38 | 885592.81 | 820964.81 | 1867120.75 | 2285121.75 | 1621061.5 | 3270555.5 | 4278985 | 4629232.5 | 3096323.25 | 4330059.5 | 747337.31 | 282292.97 | 1584979.62 | 1185369.75 | 1199595.62 | 0 | 7 | 0 | 6 | 0 | 7 | 2.807612129 | 2.621058617 | 0.715978406 | 0.668404781 |
| Q8VCC2 | Ces1 | Liver carboxylesterase 1 | 66.54 | 2.12 | 62680 | 1 | 2 | 2 | 2 | 2 | 253822.75 | 2 | 452391.16 | 2 | 550390.44 | 2 | 2 | 2 | 2 | 2 | 2 | 2 | 2 | 612343.44 | 876920.62 | 696330.12 | 10 | 6 | 7.5 | 7 | 10 | 7 | 3.939633E-05 | 9.972201E-06 | 8.610454722 | 2.179522543 |
| Q499T7 | Cfap20 | Cilia- and flagella-associated protein 20 | 47.94 | 5.49 | 19132 | 1 | 420250.09 | 377187.91 | 495680.75 | 280988.38 | 622377.38 | 104615.73 | 592011.12 | 377426.44 | 554728.19 | 164516.72 | 2 | 303008.06 | 68334.57 | 2 | 2 | 283015 | 2 | 73445.95 | 2 | 127790.95 | 1 | 1 | 2 | 3 | 2 | 3 | 0.169064989 | 0.207075784 | 0.220468616 | 0.270036461 |
| O88271 | Cfdp1 | Craniofacial development protein 1 | 171.2 | 18.64 | 32921 | 4 | 4046592 | 2674527.5 | 3386712.75 | 3875226.25 | 2422189 | 3938528.75 | 2789958.75 | 4076197 | 1819751.75 | 3673797 | 4016341.5 | 2656047.5 | 2562255 | 3404409 | 2343425.75 | 3008232.75 | 3166851.75 | 4212016.5 | 3362943.75 | 3903388 | 9 | 10 | 9 | 11 | 9 | 11 | 0.913273558 | 0.919270115 | 1.076084512 | 1.083150086 |
| P18760 | Cfl1 | Cofilin-1 | 862.19 | 79.52 | 18560 | 11 | 168372978 | 245837506 | 251746466 | 187882946 | 204472754 | 208324818 | 204240754 | 249966242 | 258456434 | 235133490 | 272290082 | 257495090 | 244209570 | 230999314 | 230175778 | 156317778 | 171828018 | 282216898 | 244786642 | 238269522 | 6 | 12 | 9 | 11 | 9 | 12 | 1.167112416 | 1.068373505 | 1.033171868 | 0.945764466 |
| Q9D0N7 | Chaf1b | Chromatin assembly factor 1 subunit B | 50.13 | 1.92 | 63132 | 1 | 493958.28 | 2 | 466325.69 | 2 | 2 | 321718.41 | 2 | 2 | 2 | 2 | 2 | 384036.75 | 2 | 2 | 239782.55 | 2 | 222478.25 | 249272.97 | 2 | 286771.06 | 10.5 | 12 | 10 | 9 | 10.5 | 12 | 0.649621801 | 1.938993134 | 0.789892953 | 2.357674895 |
| Q9D1L0 | Chchd2 | Coiled-coil-helix-coiled-coil-helix domain-containing protein 2 | 49.21 | 6.54 | 15661 | 1 | 739110.69 | 543792.88 | 954257.19 | 1102282 | 547022.06 | 761881.5 | 477274.19 | 930515.69 | 2 | 1178693.38 | 570024.69 | 443248.09 | 625324.44 | 430303.22 | 558231.94 | 615601 | 481837.25 | 1249957.25 | 1079885.75 | 936080.31 | 6 | 11 | 8 | 8 | 8 | 11 | 0.67596968 | 0.784601141 | 1.122707078 | 1.303131309 |
| Q9CRB9 | Chchd3 | MICOS complex subunit Mic19 | 262.61 | 18.94 | 26335 | 4 | 3551494.5 | 2763113.5 | 2255704.75 | 5392101 | 1543460.62 | 2685772 | 5421232 | 36820898 | 5536525.5 | 4851217 | 2554168.75 | 5647742 | 5602247 | 2536588.75 | 6159106 | 1609952.5 | 3570575 | 5908420 | 4470539 | 7574723 | 6 | 6 | 12 | 10 | 12 | 10 | 1.45105345 | 0.406753871 | 1.491964203 | 0.41822182 |
| Q5BJN5 | Chchd4 | Mitochondrial intermembrane space import and assembly protein 40 | 46.03 | 6.47 | 15467 | 1 | 1301916.5 | 988377.69 | 946600.06 | 873903.06 | 605465.06 | 706845.75 | 2 | 2 | 817566.69 | 924935.69 | 1101618.5 | 1134252.12 | 939054.44 | 1022939.81 | 1017066.88 | 1323056.12 | 1143704.25 | 933937.62 | 1016134.31 | 1090149.75 | 7 | 6 | 0 | 0 | 7 | 6 | 1.10573402 | 2.129106585 | 1.167658119 | 2.24834232 |
| Q91VN4 | Chchd6 | MICOS complex subunit Mic25 | 156.04 | 11.36 | 29852 | 2 | 1561006.88 | 1338348.5 | 429004.28 | 1607536.62 | 1215810.5 | 1767071.25 | 1606590.12 | 1003803.5 | 1309656.38 | 1773198.5 | 2464266.5 | 1570491.5 | 1226194 | 2284742 | 2034428 | 940249.06 | 2153831.75 | 2238334 | 520368.88 | 1636098.25 | 4 | 8 | 7 | 12 | 7 | 12 | 1.557311222 | 1.284143619 | 1.217366531 | 1.003828548 |
| Q6PDQ2 | Chd4 | Chromodomain-helicase-DNA-binding protein 4 | 1024.02 | 11.75 | 217751 | 21 | 16145910 | 16240685 | 15932525 | 22243228 | 14847435 | 19354356 | 15160923 | 14054105 | 13732092 | 17998110 | 12740057 | 9912070 | 11242066 | 13268734 | 10669761 | 16715333 | 18694924 | 14284036 | 17150234 | 16006203 | 0 | 11 | 0 | 10 | 0 | 11 | 0.677120184 | 0.720211534 | 0.970037941 | 1.031770326 |
| Q8CGZ0 | Cherp | Calcium homeostasis endoplasmic reticulum protein | 116.46 | 3.74 | 106169 | 3 | 2 | 2 | 2 | 389241.53 | 509587.81 | 473755.84 | 144901.11 | 350157.12 | 548096.31 | 492518.62 | 389473.69 | 1251214.38 | 1387052.12 | 466049.12 | 1300420.88 | 2 | 2 | 2 | 2 | 411758.41 | 2 | 10 | 6 | 2 | 6 | 10 | 5.333802507 | 2.385856972 | 0.45811106 | 0.204917123 |
| Q9QXG2 | Chm | Rab proteins geranylgeranyltransferase component A 1 | 137.13 | 8.27 | 73977 | 3 | 1364928.25 | 919514.88 | 1833151.25 | 1945976.5 | 1688393.5 | 1480183.75 | 1343914.12 | 1817841 | 2003348.62 | 1239372.75 | 1577801.75 | 1297307.25 | 1027826.5 | 1116054.38 | 1555701.88 | 1590614.75 | 2275660.5 | 2161490.25 | 2281684.25 | 2160465.5 | 7 | 3 | 7 | 2 | 7 | 3 | 0.848132349 | 0.833858601 | 1.350614468 | 1.327884136 |
| Q99LU0 | Chmp1b1 | Charged multivesicular body protein 1b-1 | 48.56 | 4.52 | 22124 | 1 | 2104975.25 | 2 | 2 | 2814125 | 1767743.12 | 2522111.25 | 2326093.25 | 2943831.75 | 2456794.75 | 2751509.75 | 1811624.12 | 1798972.38 | 2848761.5 | 2931375.75 | 2450273.75 | 2774202.75 | 2437395.75 | 2577341.25 | 3588723 | 3341847 | 5 | 3 | 9 | 7 | 9 | 7 | 1.770790755 | 0.910822857 | 2.201263007 | 1.132240303 |
| Q9D8B3 | Chmp4b | Charged multivesicular body protein 4b | 271.41 | 21.43 | 24936 | 4 | 6127320 | 4648010.5 | 3018293 | 3339297.5 | 3749067.5 | 6045805.5 | 3893751.25 | 4444100 | 4055374.75 | 5445003.5 | 6363682 | 6555359 | 6855648.5 | 7694914 | 7289633.5 | 4237323 | 6028029 | 4288894.5 | 3589109.5 | 4791954 | 0 | 9 | 0 | 11 | 0 | 11 | 1.664555892 | 1.455333531 | 1.098329788 | 0.960277859 |
| Q8R1T1 | Chmp7 | Charged multivesicular body protein 7 | 71.31 | 3.77 | 50633 | 1 | 274575.06 | 308542.78 | 323342.66 | 415495.16 | 273087.16 | 567060.31 | 266778.88 | 2 | 330700.19 | 397051.59 | 423202.16 | 283012.41 | 348646.22 | 418540.31 | 287218.25 | 431082.19 | 400408.19 | 481238.78 | 92495.89 | 455552.12 | 7 | 6 | 10 | 8 | 10 | 8 | 1.10380695 | 1.127450868 | 1.166600136 | 1.191589105 |
| D4A4T9 | Chordc1 | Cysteine and histidine-rich domain-containing protein 1 | 195.37 | 11.18 | 37362 | 3 | 6146168.5 | 7141336.5 | 6632346 | 7033641.5 | 5735376.5 | 6435409 | 6380036.5 | 6643235 | 5951059 | 6329205.5 | 3997395 | 5199590 | 4257780.5 | 3451181.5 | 5023869.5 | 8846882 | 6807567.5 | 6789221.5 | 6791580 | 6978797 | 0 | 8 | 0 | 0 | 0 | 8 | 0.670864951 | 0.690943461 | 1.107840348 | 1.14099722 |
| P61022 | Chp1 | Calcineurin B homologous protein 1 | 49.58 | 6.15 | 22432 | 1 | 676339.56 | 872839.06 | 635670.81 | 2 | 753645.69 | 2 | 1115757.25 | 619011 | 863111.12 | 2 | 2 | 2 | 322295.12 | 2 | 592445.81 | 752479.5 | 2 | 137968.28 | 2 | 2 | 3.5 | 5.5 | 7 | 8 | 7 | 8 | 0.311297542 | 0.352112393 | 0.30303034 | 0.342761261 |
| Q9JKP8 | Chrac1 | Chromatin accessibility complex protein 1 | 40.19 | 6.2 | 14127 | 1 | 816233.5 | 837616.06 | 938523.19 | 142932.34 | 693903.56 | 775202.5 | 806112.19 | 875206.38 | 825933.94 | 777584.19 | 880949.25 | 1046067.31 | 166497.09 | 810917.94 | 2 | 860602.88 | 293446.91 | 1280515.38 | 1064222.12 | 923691.75 | 12 | 6 | 12 | 6 | 12 | 6 | 0.846969049 | 0.715370824 | 1.28965003 | 1.089270035 |
| Q9CY57 | Chtop | Chromatin target of PRMT1 protein | 118.72 | 11.65 | 26585 | 2 | 405960.84 | 336235.22 | 401906.53 | 2 | 410551.94 | 1284919.5 | 575524.31 | 2 | 448337 | 889128.62 | 2047352.25 | 1713699.38 | 1087901 | 2159011.75 | 1529772.12 | 416941.78 | 433482.81 | 136987.75 | 885272.38 | 1113177.38 | 0 | 4 | 1 | 10 | 1 | 10 | 5.491718804 | 2.669785167 | 1.920592776 | 0.933691306 |
| Q60680 | Chuk | Inhibitor of nuclear factor kappa-B kinase subunit alpha | 41.82 | 1.21 | 84729 | 1 | 165177.92 | 133902.67 | 182144.25 | 268412.53 | 306009.84 | 215312.94 | 159658.89 | 189189.58 | 272443.31 | 300913.44 | 207300.31 | 317596.16 | 340437.81 | 320701.03 | 299597.78 | 103515.16 | 2 | 195651.03 | 2 | 253419.73 | 3 | 6 | 4 | 5 | 4 | 6 | 1.407319676 | 1.306030218 | 0.523460788 | 0.485785581 |
| Q6DG52 | Churc1 | Protein Churchill | 44.47 | 11.61 | 12858 | 1 | 783743.94 | 2 | 411683.78 | 2 | 656213.75 | 1167551.25 | 2 | 2 | 726206.12 | 653296.88 | 2 | 2 | 558646.88 | 2 | 2 | 2 | 2 | 810444.69 | 2 | 2 | 7 | 9 | 6 | 8 | 7 | 9 | 0.301707259 | 0.219333374 | 0.437693232 | 0.318191659 |
| Q9DCL2 | Ciao2a | Cytosolic iron-sulfur assembly component 2A | 40.7 | 5.62 | 18418 | 1 | 2 | 2 | 519821.47 | 473416.84 | 2 | 978961.06 | 2 | 875561.75 | 137739.8 | 799883.31 | 912658.94 | 2 | 659927.12 | 2 | 330056.41 | 1140805 | 1168665 | 63756.96 | 517876.5 | 2 | 9 | 6.5 | 9 | 11.5 | 9 | 11.5 | 1.915587586 | 0.681427533 | 2.910769718 | 1.035441367 |
| Q8WTY4 | Ciapin1 | Anamorsin | 195.18 | 12.94 | 33429 | 3 | 3957603.75 | 4255422 | 4180778.25 | 4184645.5 | 4409513 | 4698365 | 4326254 | 3408517.5 | 2922341 | 3600711.5 | 2738113.5 | 3124015 | 2138176.5 | 2643184.75 | 2345807.5 | 4416690.5 | 4350587.5 | 3662743.5 | 4235965.5 | 4193105 | 0 | 10 | 1 | 8 | 1 | 10 | 0.618892722 | 0.68522725 | 0.99385979 | 1.10038426 |
| B0K020 | Cisd1 | CDGSH iron-sulfur domain-containing protein 1 | 135.59 | 43.52 | 12097 | 3 | 2 | 862488.88 | 915598.25 | 619513.75 | 573302.38 | 887016 | 636036.25 | 1115960.62 | 717528.62 | 1029605.25 | 2283674.25 | 1725444.12 | 2860612.75 | 2954805.75 | 3399822.25 | 448586.75 | 556483.12 | 1080794.5 | 1025185.44 | 1637559.12 | 0 | 8 | 0 | 12 | 0 | 12 | 4.451289409 | 3.015028886 | 1.59837104 | 1.082637953 |
| Q9CQB5 | Cisd2 | CDGSH iron-sulfur domain-containing protein 2 | 310.42 | 38.52 | 15242 | 6 | 1968381.75 | 2978438.75 | 2783347.5 | 2691126 | 1811388.88 | 3479999.75 | 2753030 | 3421556.5 | 3047259.75 | 3532589.5 | 9108032 | 8995554 | 9684727 | 9927192 | 6099127 | 2177535.5 | 5158616 | 4744773 | 4363574 | 5079340 | 0 | 3 | 0 | 5 | 0 | 5 | 3.581767992 | 2.698870065 | 1.759535395 | 1.325813793 |
| Q8BMK4 | Ckap4 | Cytoskeleton-associated protein 4 | 1200.05 | 35.65 | 63692 | 19 | 11421080 | 10211765 | 16440126 | 15092336 | 18496232 | 9826568 | 10053564 | 17384280 | 19510934 | 16038593 | 9517444 | 15031788 | 20330238 | 12512068 | 21020864 | 10732027 | 8010153 | 14270850 | 15436416 | 16654714 | 11 | 10 | 11 | 9 | 11 | 10 | 1.094204829 | 1.076887243 | 0.908495141 | 0.894116716 |
| A2AGT5 | Ckap5 | Cytoskeleton-associated protein 5 | 1095.6 | 12.89 | 225635 | 22 | 13139344 | 14791759 | 16453442 | 15129965 | 17123694 | 11282931 | 14292536 | 12517392 | 16204204 | 12680436 | 7306344.5 | 10917230 | 6760339.5 | 7577599.5 | 6989689 | 15036311 | 9376876 | 13761917 | 12746102 | 11546591 | 0 | 3 | 0 | 11 | 0 | 11 | 0.516076845 | 0.590514771 | 0.81509996 | 0.932668403 |
| Q04447 | Ckb | Creatine kinase B-type | 1605.93 | 66.14 | 42713 | 20 | 161425218 | 144468994 | 179188242 | 162811410 | 166258082 | 166240818 | 152185426 | 166771394 | 182024082 | 168691330 | 95051754 | 121946738 | 122833034 | 111181650 | 101794042 | 124975410 | 128055778 | 167563298 | 150542530 | 148797714 | 0 | 6 | 0 | 3 | 0 | 6 | 0.678997601 | 0.661321435 | 0.884275636 | 0.861255522 |
| Q8BRT1 | Clasp2 | CLIP-associating protein 2 | 198.86 | 1.79 | 140739 | 3 | 2358136 | 2066481.75 | 1260206.25 | 2030130 | 1873879.5 | 2072245.88 | 2307426.75 | 1792357.5 | 2201624.5 | 1439332.25 | 1510308.62 | 1479561.88 | 2006697 | 1269003 | 1207238.5 | 2531130.5 | 2119692.75 | 1595672.75 | 2087838.5 | 1372310.5 | 5 | 10 | 4 | 12 | 5 | 12 | 0.779324096 | 0.761522367 | 1.012286322 | 0.989163149 |
| Q99LI2 | Clcc1 | Chloride channel CLIC-like protein 1 | 133.9 | 7.24 | 60621 | 2 | 2 | 2 | 228927.38 | 2 | 101215.05 | 2 | 2 | 171322.17 | 429019.94 | 2 | 2 | 148892.03 | 314823.19 | 2 | 2 | 197206.86 | 2 | 2 | 2 | 2 | 11.5 | 10 | 11.5 | 10 | 11.5 | 10 | 1.404584053 | 0.772420554 | 0.597352106 | 0.328500843 |
| P49300 | Clec10a | C-type lectin domain family 10 member A | 486.59 | 25.66 | 34596 | 7 | 2918947.75 | 3485245.75 | 7380997.5 | 2716060.5 | 4543006 | 2731380.25 | 2851810 | 1920872.75 | 944616.81 | 2824410.5 | 3121790.5 | 3432560.25 | 1213550.38 | 3797380 | 3553010 | 2132109 | 2817258.25 | 2381752 | 1508391.75 | 1701248.88 | 10 | 1 | 4 | 9 | 10 | 9 | 0.718404588 | 1.341095539 | 0.500885331 | 0.935037296 |
| Q504P2 | Clec12a | C-type lectin domain family 12 member A | 56.1 | 3.37 | 30757 | 1 | 263360.62 | 225029.8 | 331348.09 | 429261.59 | 531918.88 | 118763.29 | 196170.36 | 310308.44 | 257926.64 | 2 | 2 | 2 | 2 | 76237.41 | 89403.1 | 192764.12 | 184781.22 | 196735.22 | 166749.64 | 262226.22 | 0 | 1 | 3.5 | 12 | 3.5 | 12 | 0.093011817 | 0.187558877 | 0.563336362 | 1.135971094 |
| Q9R0Q8 | Clec4e | C-type lectin domain family 4 member E | 144.93 | 12.15 | 24431 | 2 | 2 | 2 | 293403.88 | 2 | 217834.89 | 2 | 454512.56 | 382112.5 | 434754.38 | 1242368.88 | 1635191.75 | 1565770.75 | 1157819 | 3619881.5 | 3175214 | 305161.75 | 376993.97 | 1157214 | 595455.56 | 1301800 | 0 | 0 | 1 | 10 | 1 | 10 | 21.81709751 | 4.437145929 | 7.308877272 | 1.486474313 |
| Q6MG61 | Clic1 | Chloride intracellular channel protein 1 | 1394.49 | 70.12 | 26981 | 18 | 136910546 | 166408018 | 131178226 | 127645218 | 150754418 | 124493082 | 143770514 | 129884338 | 134248274 | 115611746 | 134687250 | 145821010 | 131227058 | 101523170 | 136267250 | 136855810 | 106561666 | 110956898 | 106481378 | 110436706 | 9 | 2 | 9 | 4 | 9 | 4 | 0.911108142 | 1.002342211 | 0.801368105 | 0.881613342 |
| Q9QYB1 | Clic4 | Chloride intracellular channel protein 4 | 1104.48 | 70.75 | 28729 | 15 | 48247698 | 54755630 | 66288290 | 58809614 | 56968522 | 42686406 | 58248026 | 65038010 | 58653902 | 59253542 | 33220386 | 45854646 | 43411422 | 32159748 | 44227998 | 53929034 | 30784032 | 38691998 | 45939102 | 50606186 | 0 | 2 | 3 | 3 | 3 | 3 | 0.697633464 | 0.700557559 | 0.771566779 | 0.774800762 |
| Q99KN9 | Clint1 | Clathrin interactor 1 | 253.82 | 6.66 | 68513 | 4 | 7828398 | 9154773 | 10557484 | 4486338 | 12086314 | 2794233.75 | 9493212 | 9684301 | 10927460 | 5199250.5 | 1080036.38 | 4311833 | 2031879.88 | 890859.81 | 2423061 | 8177334.5 | 2024662 | 2976371 | 2477946.5 | 5088433.5 | 0 | 3 | 1 | 4 | 1 | 4 | 0.243411134 | 0.281840023 | 0.470260538 | 0.54450361 |
| Q922J3 | Clip1 | CAP-Gly domain-containing linker protein 1 | 44.56 | 1.29 | 155814 | 1 | 82141.94 | 2 | 328469.97 | 299513.22 | 215758.61 | 254254.7 | 2 | 307594.69 | 335862.44 | 321225.81 | 2 | 94309.16 | 2 | 2 | 135258.34 | 2 | 2 | 237799.48 | 340856.75 | 295710.53 | 5.5 | 12 | 3.5 | 9 | 5.5 | 12 | 0.247950141 | 0.188338694 | 0.944361407 | 0.717320802 |
| O88696 | Clpp | ATP-dependent Clp protease proteolytic subunit, mitochondrial | 165.95 | 9.93 | 29800 | 2 | 1193247.25 | 336820.59 | 2 | 425006.72 | 561422.88 | 831027.31 | 609021.25 | 569236.88 | 885965.62 | 531460 | 1632208.38 | 1003611.69 | 1557240.88 | 536764.5 | 953479.75 | 2 | 1346466.5 | 799011.5 | 704701.62 | 2 | 4 | 11 | 4 | 11 | 4 | 11 | 2.258417033 | 1.658530614 | 1.132598551 | 0.831754872 |
| Q8VBZ3 | Clptm1 | Cleft lip and palate transmembrane protein 1 homolog | 431.53 | 14.16 | 75291 | 6 | 2013114.25 | 2520916 | 1965712.75 | 2868268 | 2104517 | 3797316 | 3473973 | 3565440.5 | 1785985 | 4379906 | 4435151.5 | 4877026 | 4646717 | 5908635 | 5385962 | 4286360 | 5302041 | 4720796.5 | 1679070.75 | 4269669 | 0 | 5 | 0 | 7 | 0 | 7 | 2.201214196 | 1.48527055 | 1.765777974 | 1.191459708 |
| Q5U2U0 | Clpx | ATP-dependent Clp protease ATP-binding subunit clpX-like, mitochondrial | 320.12 | 11.06 | 69208 | 6 | 3452242.25 | 2104510 | 3165225.5 | 3736248 | 3093498 | 2909517 | 3210202.75 | 3741153.75 | 3389004 | 1890587 | 2518881.5 | 3376405 | 4773024.5 | 1687549.25 | 4554259 | 2904257 | 2080615.88 | 1649725.62 | 3617662.5 | 5092083.5 | 11 | 10 | 11 | 11 | 11 | 11 | 1.087346941 | 1.116882461 | 0.986665192 | 1.013465901 |
| O08585 | Clta | Clathrin light chain A | 175.38 | 14.04 | 25604 | 4 | 13698329 | 12749071 | 18641834 | 18134450 | 16723630 | 12249745 | 14481654 | 16072367 | 14789170 | 18733798 | 2353338.5 | 1684241.12 | 6685222 | 2313229.5 | 5212441.5 | 15590410 | 15659413 | 14860430 | 19435998 | 16973350 | 0 | 11 | 0 | 7 | 0 | 11 | 0.228256232 | 0.23908363 | 1.032174777 | 1.081136277 |
| P08082 | Cltb | Clathrin light chain B | 99.11 | 6.99 | 25117 | 2 | 961839.88 | 3197542.75 | 3731518 | 8657162 | 3148922.25 | 3480017.5 | 2861465.5 | 7823030 | 7152813 | 8006590.5 | 2603491.75 | 2507395.5 | 3040055.25 | 7224801 | 2902008.25 | 3458653.25 | 9388809 | 3593038.25 | 4321463.5 | 4083089.75 | 8 | 6 | 5 | 12 | 8 | 12 | 0.927946681 | 0.623305272 | 1.261363295 | 0.847262464 |
| Q68FD5 | Cltc | Clathrin heavy chain 1 | 5023.168 | 52.78 | 191557 | 77 | 181695314 | 182202466 | 252729362 | 226583410 | 256627522 | 169952898 | 193245794 | 232686882 | 236377266 | 198491874 | 166678386 | 193706066 | 195643938 | 176924002 | 193306418 | 191662850 | 160606930 | 196361986 | 205646866 | 215153362 | 6 | 8 | 7 | 9 | 7 | 9 | 0.842177437 | 0.898621949 | 0.881431564 | 0.940506971 |
| Q5SW19 | Cluh | Clustered mitochondria protein homolog | 393.92 | 6.46 | 148068 | 8 | 8998213 | 9061088 | 7484216.5 | 7758781.5 | 8036094.5 | 10018414 | 9692210 | 7314415.5 | 6943600.5 | 5759317 | 7997148.5 | 7035787.5 | 7042026.5 | 3783795.5 | 4214167 | 9103781 | 6502864.5 | 6872105 | 7496536 | 5423002 | 2 | 6 | 7 | 8 | 7 | 8 | 0.727481705 | 0.756971344 | 0.856305374 | 0.891017087 |
| Q5I0K3 | Clybl | Citramalyl-CoA lyase, mitochondrial | 185.04 | 13.02 | 37321 | 3 | 1462440.62 | 1477385 | 1253630.88 | 1396889.25 | 1386050.75 | 1274115.38 | 1416498.12 | 1287435.62 | 1699396.75 | 1601771.5 | 893941.31 | 934923.44 | 1158842.25 | 998002.12 | 917607.75 | 1639086.75 | 1520903.62 | 1656044.88 | 1333862.38 | 508131.22 | 0 | 9 | 0 | 12 | 0 | 12 | 0.702843778 | 0.673604952 | 0.954365029 | 0.914662732 |
| Q99KK2 | Cmas | N-acylneuraminate cytidylyltransferase | 344.42 | 17.82 | 48058 | 7 | 7628619.5 | 8435576 | 7128415.5 | 8247266 | 8856793 | 9084448 | 8503447 | 8364436 | 9768020 | 8560030 | 7578269 | 8372716 | 7327367.5 | 7402809.5 | 8226560 | 8730645 | 7588657 | 8802628 | 8205892.5 | 8549007 | 8 | 10 | 1 | 8 | 8 | 10 | 0.965531941 | 0.878667282 | 1.039213153 | 0.94571972 |
| Q9D486 | Cmip | C-Maf-inducing protein | 70.6 | 1.81 | 86259 | 1 | 486473.72 | 9235168 | 376053.09 | 2 | 2 | 728365.44 | 2 | 427944.31 | 2 | 2 | 2 | 2 | 2 | 574185.69 | 1051120.12 | 490631.34 | 580744.31 | 2 | 440784.12 | 2 | 11 | 12 | 11.5 | 10 | 11.5 | 12 | 0.160958634 | 1.40559515 | 0.149753305 | 1.30774295 |
| Q4KM73 | Cmpk1 | UMP-CMP kinase | 467.5 | 46.94 | 22169 | 8 | 7379787 | 8601723 | 10183753 | 8449711 | 9837036 | 7548971 | 8750860 | 6628959.5 | 11664963 | 10351793 | 9379655 | 10785285 | 11574680 | 5203410.5 | 8208064 | 5746842.5 | 5060606.5 | 10521196 | 7852519 | 10073450 | 11 | 10 | 12 | 9 | 12 | 10 | 1.015726724 | 1.004573267 | 0.883078493 | 0.873381615 |
| Q3U5Q7 | Cmpk2 | UMP-CMP kinase 2, mitochondrial | 456.25 | 17.23 | 50036 | 7 | 6470152 | 7021939.5 | 718882.69 | 2061689.12 | 3882863.25 | 6360633 | 6109795 | 4545668 | 2812589 | 3502593.5 | 1785632.75 | 2357508.5 | 1726355.25 | 893184.69 | 1580120 | 6270331 | 5488697 | 1892160 | 1879267.75 | 2321081.5 | 6 | 10 | 0 | 7 | 6 | 10 | 0.413921272 | 0.357580112 | 0.885689451 | 0.765133263 |
| Q9CZ69 | Cmtm6 | CKLF-like MARVEL transmembrane domain-containing protein 6 | 55.17 | 6.01 | 19824 | 1 | 2 | 243956.92 | 2 | 202176.08 | 2 | 500569.84 | 396595.81 | 195349.69 | 294503.06 | 502665.66 | 793798.56 | 452956.16 | 816642.69 | 675500.19 | 2 | 2 | 411731.91 | 489203.66 | 412250.56 | 526344.44 | 3.5 | 3.5 | 7 | 11 | 7 | 11 | 6.139117181 | 1.449395514 | 4.123227447 | 0.973460384 |
| Q5U2Z5 | Cmtr1 | Cap-specific mRNA (nucleoside-2'-O-)-methyltransferase 1 | 40.98 | 0.84 | 95619 | 1 | 1085626.12 | 774060.25 | 514266.25 | 836252.06 | 749231.62 | 1079299.75 | 820770.62 | 686285.75 | 762767.56 | 978239.62 | 736943.31 | 516510.94 | 940061.31 | 672923 | 709298.69 | 823658.75 | 808322.25 | 759968.81 | 745402.25 | 690109 | 8 | 10 | 5 | 8 | 8 | 10 | 0.903092506 | 0.82630854 | 0.966668174 | 0.884478791 |
| P62634 | Cnbp | Cellular nucleic acid-binding protein | 229.3 | 28.25 | 19463 | 4 | 4641721 | 6903768 | 3643641.5 | 2412688.75 | 2240878.25 | 6431674 | 5730008.5 | 2284595.25 | 1628454.5 | 1715741.25 | 8311745.5 | 8615465 | 2864602 | 4049449.75 | 2298376.5 | 5760600 | 4905701 | 2991561 | 2903392.25 | 2842146 | 9 | 11 | 6 | 9 | 9 | 11 | 1.317343005 | 1.469305398 | 0.977861012 | 1.090662385 |
| Q9D1A2 | Cndp2 | Cytosolic non-specific dipeptidase | 1264.78 | 49.68 | 52767 | 21 | 31194912 | 34162302 | 32547142 | 32847594 | 32894854 | 27273016 | 29610304 | 37769886 | 36802870 | 39299498 | 35138390 | 35372074 | 34349694 | 34234850 | 37224278 | 22905788 | 22963992 | 37912970 | 37652862 | 34919514 | 0 | 10 | 11 | 9 | 11 | 10 | 1.077438005 | 1.032582901 | 0.955442588 | 0.915666308 |
| Q9CX13 | Cnih4 | Protein cornichon homolog 4 | 50.23 | 14.39 | 16089 | 1 | 3335949 | 3049061.5 | 4471321 | 5393580 | 2566935.5 | 2365126.5 | 2004584 | 4962764 | 663865.88 | 6816614.5 | 3873597.75 | 850483.25 | 2460939 | 1741198 | 6203534.5 | 2340140.5 | 1998137.38 | 4597129 | 5219979 | 5191262 | 8 | 12 | 12 | 11 | 12 | 12 | 0.804053543 | 0.899886582 | 1.028155667 | 1.150698852 |
| Q6ZQ08 | Cnot1 | CCR4-NOT transcription complex subunit 1 | 264.29 | 2.69 | 266808 | 6 | 2726535.25 | 2069105.5 | 1878851.62 | 2599557.25 | 3685175.25 | 1515409.62 | 1064867 | 2315973.5 | 2993399.75 | 2073539.75 | 2966526.25 | 3460695.75 | 3213673.5 | 1500384 | 3800321.25 | 3026963.25 | 1164183.25 | 1814359.5 | 1230986.75 | 1263971.62 | 8 | 4 | 5 | 10 | 8 | 10 | 1.152970251 | 1.499680456 | 0.65593926 | 0.853187051 |
| Q8C5L3 | Cnot2 | CCR4-NOT transcription complex subunit 2 | 58.02 | 4.26 | 59711 | 1 | 336528.47 | 2 | 316430.91 | 293983.16 | 426009.12 | 431330.75 | 350088.72 | 407516.16 | 355824.69 | 206762.48 | 2 | 211791.84 | 360050.72 | 288337.97 | 253184.67 | 303711.53 | 384181.91 | 308389.53 | 2 | 261133.61 | 7.5 | 9.5 | 6 | 6 | 7.5 | 9.5 | 0.810928462 | 0.635656698 | 0.91584925 | 0.717900207 |
| P16330 | Cnp | 2',3'-cyclic-nucleotide 3'-phosphodiesterase | 125.86 | 5.71 | 47123 | 2 | 726288.56 | 605596.44 | 816636.44 | 912145.94 | 43638.67 | 610027.69 | 725087.31 | 644649.69 | 275365.59 | 495171.56 | 858000.38 | 1002114.5 | 933668.31 | 780884.25 | 759026.5 | 2 | 739843.12 | 450261.94 | 553729.38 | 560177.75 | 5 | 6 | 0 | 10 | 5 | 10 | 1.396026639 | 1.575715755 | 0.742199433 | 0.837731392 |
| Q9QXT0 | Cnpy2 | Protein canopy homolog 2 | 212.07 | 22.53 | 20767 | 3 | 2245234.25 | 2050938.62 | 2180582.5 | 2927464.5 | 2027458.75 | 2472562.5 | 3820137 | 3446818 | 1719250.25 | 4363951 | 4181075 | 3119747 | 3281766.5 | 3225455 | 3602892.5 | 1598208.12 | 2144923.5 | 3683619 | 3607459.25 | 3316933.5 | 0 | 8 | 12 | 9 | 12 | 9 | 1.523042816 | 1.100375749 | 1.255383732 | 0.906996048 |
| Q9DBL7 | Coasy | Bifunctional coenzyme A synthase | 238.66 | 10.48 | 62023 | 5 | 2457853.25 | 2450339.5 | 3027724 | 2557143 | 2764909 | 2422387.5 | 2227375.75 | 2401554.75 | 2767433.75 | 3025586.5 | 3833864.5 | 3114185.25 | 3334404.75 | 3445516.5 | 3371358 | 2565154.25 | 2425627.5 | 2955260.5 | 3116308.75 | 3028344.75 | 0 | 8 | 0 | 5 | 0 | 8 | 1.289739727 | 1.331273645 | 1.062809546 | 1.097035556 |
| Q9Z160 | Cog1 | Conserved oligomeric Golgi complex subunit 1 | 39.92 | 0.92 | 109050 | 1 | 449694.12 | 553511.62 | 329528.31 | 512253.38 | 691699.25 | 322817.25 | 317070.31 | 509539 | 2 | 402908.34 | 2 | 442698.28 | 337484.62 | 2 | 347219.91 | 515644.28 | 92889.74 | 441171.44 | 464732.62 | 417584.62 | 3 | 7 | 11 | 7 | 11 | 7 | 0.444440702 | 0.72626426 | 0.761632375 | 1.244589818 |
| Q8C0L8 | Cog5 | Conserved oligomeric Golgi complex subunit 5 | 112.54 | 2.53 | 91391 | 2 | 1556385.25 | 1289291.25 | 1292609.5 | 1493729.25 | 1275342.12 | 707657.88 | 1210592.12 | 1109672.75 | 1404176.38 | 1223607 | 764312.69 | 1254336.25 | 103298.16 | 614583.5 | 979610.88 | 1093862.75 | 1087591.12 | 788987.25 | 1114035.5 | 1364003.25 | 0 | 3 | 6 | 9 | 6 | 9 | 0.53799757 | 0.657060568 | 0.788793684 | 0.96335979 |
| Q5SU73 | Coil | Coilin | 47.48 | 1.75 | 61917 | 1 | 2 | 301952.28 | 420504.94 | 414717.97 | 545937.88 | 347331.91 | 380120 | 375140.03 | 496678 | 450367.94 | 195745.77 | 285910.34 | 404226.03 | 304308.66 | 181220.53 | 464712.47 | 729493.75 | 371295.44 | 482375.22 | 472462.06 | 7 | 6 | 3 | 7 | 7 | 7 | 0.814805449 | 0.669099329 | 1.497425212 | 1.22965084 |
| Q8K4Q8 | Colec12 | Collectin-12 | 625.75 | 15.36 | 81304 | 10 | 9871772 | 10740815 | 4994966.5 | 3406919 | 6948511 | 10699543 | 8294682.5 | 3373472.5 | 8922013 | 8207232 | 6194816.5 | 7899670 | 8201221 | 4467630 | 5621445.5 | 7901341 | 6594677.5 | 8339430 | 5121473 | 8332831 | 11 | 12 | 5 | 9 | 11 | 12 | 0.900503236 | 0.819931381 | 1.009086259 | 0.918799019 |
| Q8K297 | Colgalt1 | Procollagen galactosyltransferase 1 | 492.62 | 16.21 | 71061 | 10 | 6316465.5 | 6631513.5 | 10341376 | 6919533.5 | 9945237 | 7080995 | 9312300 | 10230210 | 12401221 | 9881968 | 13057111 | 12143810 | 13827149 | 14545194 | 15242906 | 9368100 | 7388491 | 8904424 | 7925501 | 6700500.5 | 0 | 11 | 1 | 5 | 1 | 11 | 1.71380074 | 1.407091021 | 1.003309523 | 0.823752603 |
| Q8BXC6 | Commd2 | COMM domain-containing protein 2 | 40.27 | 4.52 | 22848 | 1 | 1334608 | 1061149.62 | 998805 | 1310639.25 | 877240.81 | 2 | 1214553 | 996653.94 | 1545699.5 | 1037784.62 | 850758.31 | 797708.62 | 1177895.5 | 688663.19 | 2 | 1320903.12 | 1195098.75 | 1014771.88 | 1109951.12 | 798821.31 | 3 | 12 | 6.5 | 12 | 6.5 | 12 | 0.629657629 | 0.733107954 | 0.974402514 | 1.134493097 |
| Q63829 | Commd3 | COMM domain-containing protein 3 | 106.37 | 10.77 | 22037 | 1 | 2 | 635731.69 | 516816.72 | 354304.16 | 704497.88 | 229779.27 | 519543.03 | 704956.56 | 759075.56 | 243746.22 | 228229.81 | 350776.22 | 294882.97 | 189226.98 | 188851.47 | 389973.72 | 456251.62 | 400829.59 | 356067.69 | 613503.31 | 5 | 11 | 4 | 11 | 5 | 11 | 0.56615464 | 0.509530391 | 1.002384731 | 0.902130704 |
| O88587 | Comt | Catechol O-methyltransferase | 222.27 | 35.47 | 29486 | 5 | 658546.75 | 797114.5 | 827455.62 | 660984.31 | 2476013.75 | 253133.11 | 326839 | 475776.31 | 2200330.5 | 724299.62 | 2187135.75 | 1295848.25 | 1793280.12 | 1251820 | 957107 | 255973.73 | 262639.09 | 684912.62 | 638089.31 | 517999.59 | 5 | 2 | 5 | 11 | 5 | 11 | 1.381002288 | 1.880522429 | 0.435343968 | 0.592811542 |
| Q8BIG7 | Comtd1 | Catechol O-methyltransferase domain-containing protein 1 | 372.41 | 41.22 | 28961 | 6 | 5550451 | 4390174 | 2861018.5 | 3596811 | 3763666.25 | 4555779 | 4514774 | 3654933 | 3389992.5 | 4001443 | 8755382 | 5453733 | 6245979.5 | 7598881 | 4628597 | 3880878 | 4534800 | 4036096.25 | 5051660.5 | 4237763 | 2 | 8 | 0 | 8 | 2 | 8 | 1.620988829 | 1.624630911 | 1.078318993 | 1.08074179 |
| Q8CIE6 | Copa | Coatomer subunit alpha | 1820.34 | 28.27 | 138432 | 32 | 62518622 | 56942294 | 55290430 | 56411790 | 68287466 | 58696046 | 60620222 | 58862522 | 67046610 | 59933682 | 40265106 | 40837738 | 44116738 | 36684994 | 40788714 | 57514306 | 61179482 | 55639322 | 53523154 | 53331334 | 0 | 7 | 0 | 4 | 0 | 7 | 0.676883896 | 0.664221719 | 0.93901163 | 0.921445943 |
| P23514 | Copb1 | Coatomer subunit beta | 1047.6 | 21.83 | 107011 | 18 | 22540418 | 16952538 | 20559930 | 21190054 | 24489120 | 17493250 | 22045548 | 22234936 | 22572358 | 21131844 | 9424965 | 15110004 | 15151232 | 9784707 | 13539165 | 20379126 | 17989354 | 16425265 | 19841698 | 17398334 | 0 | 4 | 0 | 3 | 0 | 4 | 0.595941032 | 0.597376811 | 0.87044343 | 0.872540557 |
| O35142 | Copb2 | Coatomer subunit beta' | 1129.96 | 26.52 | 102551 | 19 | 44185318 | 38809958 | 41160438 | 44669362 | 41426026 | 40856970 | 38514082 | 41103786 | 40136366 | 41967262 | 34425702 | 34324630 | 31019382 | 31574358 | 31988566 | 43237810 | 40324462 | 44933562 | 47517874 | 45399338 | 0 | 6 | 0 | 3 | 0 | 6 | 0.776845574 | 0.80626851 | 1.053088635 | 1.092974245 |
| O89079 | Cope | Coatomer subunit epsilon | 381.34 | 34.09 | 34567 | 6 | 4250025 | 4124222.25 | 5507106 | 3586611 | 6616840.5 | 2572399.75 | 4401832.5 | 4360027.5 | 2441046.5 | 2811627.5 | 1267094.25 | 3945221.25 | 2307422.25 | 315464.31 | 2758708.25 | 4894398 | 1278795 | 2668183.5 | 2388468.75 | 3465152.25 | 1 | 3 | 5 | 10 | 5 | 10 | 0.439858675 | 0.638690096 | 0.610135629 | 0.885938156 |
| Q4AEF8 | Copg1 | Coatomer subunit gamma-1 | 1270.15 | 34.9 | 97614 | 22 | 23925522 | 25877720 | 20171306 | 25429790 | 22423046 | 23757074 | 23022774 | 21369818 | 24092130 | 25643276 | 22808506 | 23413056 | 24826118 | 21259120 | 19702938 | 25417088 | 25606708 | 21524350 | 24643314 | 23751844 | 8 | 12 | 7 | 10 | 8 | 12 | 0.950625688 | 0.950160492 | 1.026444786 | 1.025942487 |
| Q9QXK3 | Copg2 | Coatomer subunit gamma-2 | 869.72 | 21.24 | 97681 | 15 | 9186461 | 9970847 | 13850229 | 14476522 | 13414816 | 9906283 | 9627230 | 12448507 | 14305155 | 15116345 | 8000947 | 7802925 | 11701465 | 10308775 | 9804182 | 10300908 | 10367340 | 13182974 | 14495600 | 12797268 | 5 | 12 | 5 | 11 | 5 | 12 | 0.781924034 | 0.775497789 | 1.004026593 | 0.995774998 |
| P61202 | Cops2 | COP9 signalosome complex subunit 2 | 73.28 | 2.93 | 51597 | 1 | 729308 | 494259.12 | 371065.94 | 478869.12 | 853231 | 523918.38 | 484425.03 | 551506.5 | 736759.19 | 2 | 498186.44 | 636485.56 | 559282.25 | 462847.53 | 481562.47 | 538290.94 | 2 | 534516.12 | 2 | 259458.61 | 12 | 6 | 12 | 8 | 12 | 8 | 0.901470714 | 1.148807584 | 0.455207082 | 0.580102426 |
| O88543 | Cops3 | COP9 signalosome complex subunit 3 | 408.53 | 16.55 | 47832 | 6 | 11310011 | 12215404 | 13699483 | 13691289 | 14171999 | 9249122 | 11147893 | 11190612 | 14173865 | 13895108 | 8595662 | 10323890 | 11807488 | 10714190 | 9847526 | 12570096 | 12491075 | 14016354 | 13883269 | 9005503 | 1 | 12 | 6 | 12 | 6 | 12 | 0.787988774 | 0.859733139 | 0.952036012 | 1.038716538 |
| O88544 | Cops4 | COP9 signalosome complex subunit 4 | 556.8 | 27.34 | 46285 | 9 | 7957743.5 | 6619432.5 | 7345180.5 | 9445969 | 8747539 | 4468927 | 6982915 | 7415492 | 6799228.5 | 8158274 | 6565915 | 5518977.5 | 4665305.5 | 5489297 | 7489194 | 5381955 | 8903796 | 8809119 | 9287975 | 8446095 | 2 | 10 | 8 | 4 | 8 | 10 | 0.741070631 | 0.878901188 | 1.017775399 | 1.207069841 |
| O35864 | Cops5 | COP9 signalosome complex subunit 5 | 294.84 | 14.97 | 37549 | 5 | 7283181 | 7133197 | 10666911 | 9320678 | 8299558 | 5031911 | 5869031 | 9183234 | 8595103 | 7817493 | 6558482.5 | 5930446.5 | 7620684.5 | 5256909 | 7190444 | 7997417 | 5369675 | 10436615 | 8066718 | 7307253 | 3 | 10 | 9 | 11 | 9 | 11 | 0.762395294 | 0.892050576 | 0.917434287 | 1.073455976 |
| O88545 | Cops6 | COP9 signalosome complex subunit 6 | 138.07 | 6.17 | 35880 | 3 | 2510431.25 | 2138555.25 | 2893671.75 | 2818218.5 | 3516541 | 1727429.75 | 2445324.75 | 3052965 | 2973350.75 | 3082430.75 | 1819719.25 | 1753233.38 | 1824832.38 | 1483122.38 | 2168775.25 | 2521826.5 | 3226715 | 1432728 | 2724296.25 | 1978221.75 | 1 | 8 | 4 | 10 | 4 | 10 | 0.652115747 | 0.681374992 | 0.85633997 | 0.894762384 |
| Q9CZ04 | Cops7a | COP9 signalosome complex subunit 7a | 275.5 | 16 | 30224 | 3 | 1519955.5 | 1819158.5 | 2171140.75 | 1668375.5 | 1391834.12 | 2267312.5 | 2553711 | 6359844 | 2081494.12 | 1737585.38 | 1980753.88 | 1555785 | 1885048.38 | 1677850 | 1610670.75 | 1534998 | 1767672 | 2003285 | 2415860.75 | 2005650.12 | 10 | 7 | 2 | 6 | 10 | 7 | 1.016293591 | 0.580675919 | 1.134998694 | 0.648500016 |
| Q8BV13 | Cops7b | COP9 signalosome complex subunit 7b | 53.25 | 7.2 | 29689 | 1 | 352103.44 | 508108.44 | 470809.19 | 412315.56 | 377836.09 | 321597.34 | 2 | 203662.64 | 495294.56 | 496978.94 | 2 | 401717.38 | 337943.91 | 282076.16 | 254256.33 | 315462.25 | 412128.91 | 281231.75 | 369828 | 415365.78 | 2 | 6 | 10.5 | 12 | 10.5 | 12 | 0.601552041 | 0.840834232 | 0.845766435 | 1.182190936 |
| Q8VBV7 | Cops8 | COP9 signalosome complex subunit 8 | 161.29 | 22.49 | 23256 | 3 | 3264131.5 | 2439226 | 3143335.75 | 3287645.5 | 1900691.25 | 3708964.75 | 2756684 | 2966241 | 2476320.25 | 2958189.25 | 2007797.5 | 1629273.75 | 2978140.5 | 3772340 | 2620489.25 | 2583826 | 4380320.5 | 3771246 | 3697286 | 3400887.5 | 10 | 3 | 10 | 6 | 10 | 6 | 0.926826733 | 0.874996075 | 1.270646803 | 1.199588798 |
| P61924 | Copz1 | Coatomer subunit zeta-1 | 370.69 | 35.03 | 20198 | 6 | 6112942.5 | 6947260.5 | 7776923 | 7052920.5 | 8207471 | 4109091 | 6819614 | 6502580 | 8067166.5 | 5374916 | 2040044.75 | 3843425.25 | 3240284 | 2022807.75 | 3490172 | 7463687 | 4741617 | 6226048 | 6188686 | 5135334 | 0 | 5 | 0 | 10 | 0 | 10 | 0.405477572 | 0.474089318 | 0.824305217 | 0.963787705 |
| Q566J8 | Coq8b | Atypical kinase COQ8B, mitochondrial | 43.04 | 4.13 | 59232 | 1 | 2 | 2 | 2 | 145243.41 | 2 | 2 | 2 | 2 | 2 | 45529.27 | 111474.27 | 2 | 170878.92 | 214607.64 | 132035.81 | 2 | 183745.5 | 166923.89 | 316249.75 | 76752.51 | 5 | 4 | 3 | 3 | 5 | 4 | 4.330413316 | 13.81283156 | 5.119906581 | 16.33109868 |
| O89053 | Coro1a | Coronin-1A | 1373.28 | 55.31 | 50989 | 23 | 217529282 | 272010914 | 307646658 | 241697474 | 215837650 | 121061522 | 225452690 | 297649058 | 257171346 | 260363714 | 291261282 | 311353250 | 313316226 | 247820722 | 305182146 | 218303906 | 191150738 | 305967266 | 209416002 | 292252354 | 4 | 10 | 4 | 12 | 4 | 12 | 1.170724393 | 1.264470808 | 0.970007928 | 1.047681859 |
| Q9WUM3 | Coro1b | Coronin-1B | 115.99 | 5.37 | 53912 | 3 | 5935097.5 | 6006882.5 | 7908851.5 | 7793590 | 8144940 | 6441612.5 | 5810840 | 6887570 | 6246150 | 8846313 | 4190097.25 | 4199364 | 6947797 | 8664012 | 6941767 | 5512540.5 | 6006953.5 | 9550574 | 10186073 | 8540121 | 9 | 8 | 12 | 10 | 12 | 10 | 0.864587574 | 0.903908577 | 1.111957865 | 1.162529142 |
| Q9WUM4 | Coro1c | Coronin-1C | 769.5 | 29.11 | 53121 | 12 | 34473306 | 34155134 | 33503794 | 40889766 | 29990694 | 34992010 | 33178604 | 32932090 | 27737644 | 36226350 | 19133772 | 18659226 | 17965376 | 19039818 | 18496726 | 37734350 | 34852418 | 37349514 | 37194498 | 41645930 | 0 | 4 | 0 | 2 | 0 | 4 | 0.539237416 | 0.565195276 | 1.091114794 | 1.143638979 |
| Q9D2V7 | Coro7 | Coronin-7 | 303.11 | 5.97 | 100812 | 5 | 9053420 | 10865616 | 6785693.5 | 13387078 | 9046619 | 9451717 | 7881362.5 | 9787965 | 8731591 | 9752732 | 6007479.5 | 6176702.5 | 6958483.5 | 7741362.5 | 6268989 | 8896002 | 9479314 | 7546859 | 10548104 | 11020179 | 2 | 12 | 0 | 10 | 2 | 12 | 0.674686175 | 0.726954278 | 0.966462734 | 1.041334838 |
| Q9CQI6 | Cotl1 | Coactosin-like protein | 552.38 | 50 | 15944 | 10 | 88434530 | 95162810 | 59937286 | 70603034 | 55232158 | 76530770 | 61803874 | 55533270 | 46163806 | 60965430 | 80091930 | 55912134 | 49696922 | 48041758 | 49602418 | 84189530 | 109038978 | 65281310 | 69498570 | 62962222 | 4 | 11 | 10 | 3 | 10 | 11 | 0.767104263 | 0.941354966 | 1.058480122 | 1.29891798 |
| Q5XFV8 | Cox14 | Cytochrome c oxidase assembly protein COX14 | 49.91 | 14.04 | 6361 | 1 | 1472379.38 | 1226660.88 | 2129575.25 | 1884116.75 | 2 | 1686638.62 | 1504982.75 | 1840923.25 | 431509.06 | 2 | 774752.06 | 1318875.38 | 331680.72 | 2 | 1473593.88 | 1588856 | 1143948.5 | 1273861.5 | 2077569.5 | 1716124.75 | 7.5 | 12 | 7.5 | 9 | 7.5 | 12 | 0.580822045 | 0.713554961 | 1.162024288 | 1.427577006 |
| Q8BJ03 | Cox15 | Cytochrome c oxidase assembly protein COX15 homolog | 111.36 | 7.99 | 45853 | 2 | 143721.69 | 133392.56 | 194922.3 | 175618.66 | 169020.78 | 414531.59 | 178891.92 | 648861.19 | 767773.25 | 164309.14 | 145569.52 | 150946.58 | 347091.5 | 1273939.25 | 772121.75 | 152335.41 | 111005.77 | 567692.5 | 577213.56 | 662972.75 | 6 | 8 | 12 | 10 | 12 | 10 | 3.293434156 | 1.236989197 | 2.536158789 | 0.952562242 |
| P56394 | Cox17 | Cytochrome c oxidase copper chaperone | 78.43 | 25.4 | 6784 | 1 | 2177381.75 | 3223873 | 2721935.5 | 3438080.5 | 3145011 | 2621590.5 | 2602594 | 3333132.75 | 3385933.75 | 4164232 | 1817334.25 | 2173081 | 3117996.75 | 3268730 | 3275786.75 | 2892251.25 | 3375194 | 4996023.5 | 4462687 | 4936685 | 10 | 4 | 6 | 5 | 10 | 5 | 0.928373941 | 0.847614041 | 1.405035012 | 1.28281003 |
| P19783 | Cox4i1 | Cytochrome c oxidase subunit 4 isoform 1, mitochondrial | 206.74 | 30.77 | 19530 | 4 | 14667338 | 14778218 | 17991180 | 15664650 | 12697059 | 13551548 | 11134288 | 13447786 | 11237602 | 15498217 | 24113178 | 21657602 | 22750404 | 25271566 | 25269576 | 13697534 | 16728997 | 16813934 | 15888168 | 16324582 | 0 | 8 | 0 | 1 | 0 | 8 | 1.570775311 | 1.835414706 | 1.048216952 | 1.224817322 |
| P11240 | Cox5a | Cytochrome c oxidase subunit 5A, mitochondrial | 388.94 | 38.36 | 16130 | 7 | 44838462 | 35949570 | 40842742 | 44274914 | 39375850 | 47295454 | 38330634 | 34163878 | 39460162 | 50218982 | 33860022 | 39141578 | 47631746 | 56691922 | 54409438 | 48988986 | 67034246 | 56872070 | 60359082 | 57591114 | 9 | 0 | 9 | 1 | 9 | 1 | 1.128862869 | 1.106295367 | 1.416812738 | 1.388488728 |
| P19536 | Cox5b | Cytochrome c oxidase subunit 5B, mitochondrial | 224.71 | 23.44 | 13813 | 4 | 11387051 | 10905558 | 16894578 | 17367186 | 14034855 | 12196214 | 11019126 | 15618282 | 15287958 | 17380704 | 8707589 | 8411784 | 11776748 | 13246962 | 12305454 | 11749818 | 16110477 | 16100510 | 18388082 | 17201318 | 6 | 8 | 5 | 7 | 6 | 8 | 0.771343427 | 0.761493675 | 1.126945389 | 1.11255474 |
| P10818 | Cox6a1 | Cytochrome c oxidase subunit 6A1, mitochondrial | 64.06 | 18.92 | 12301 | 1 | 2 | 2 | 2 | 2 | 2 | 2 | 2 | 2 | 2 | 2 | 300907.22 | 202004.3 | 178053.77 | 2 | 159018.69 | 2 | 2 | 2 | 2 | 2 | 2.5 | 12.5 | 2.5 | 12.5 | 2.5 | 12.5 | 83998.598 | 83998.598 | 1 | 1 |
| P56391 | Cox6b1 | Cytochrome c oxidase subunit 6B1 | 78.3 | 16.28 | 10071 | 1 | 2 | 2 | 2 | 2 | 2 | 2 | 2 | 2 | 2 | 2 | 2 | 2 | 2 | 2 | 2 | 2 | 1008471.25 | 2 | 2 | 2 | 12.5 | 10 | 12.5 | 10 | 12.5 | 10 | 1 | 1 | 100847.925 | 100847.925 |
| Q9CPQ1 | Cox6c | Cytochrome c oxidase subunit 6C | 167.04 | 44.74 | 8469 | 4 | 15238614 | 15157244 | 12939878 | 10488756 | 9233261 | 16095516 | 14748134 | 8125216 | 7785535 | 11591142 | 21348290 | 21548734 | 19244470 | 21701218 | 21927320 | 14318079 | 20776030 | 11594254 | 12461696 | 10714111 | 0 | 11 | 0 | 9 | 0 | 11 | 1.677351745 | 1.81282111 | 1.107939415 | 1.197420855 |
| P48771 | Cox7a2 | Cytochrome c oxidase subunit 7A2, mitochondrial | 64.67 | 12.05 | 9291 | 1 | 2705127.25 | 2008185.75 | 2201341.5 | 2324223.25 | 1986465 | 2947569.5 | 2005671.88 | 1796653.75 | 1396709.12 | 2658039 | 1844403.88 | 1608053.62 | 1637196 | 2278042.25 | 1976359.38 | 3079496.25 | 4113336.75 | 2451436.5 | 2751867 | 2343606.5 | 3 | 2 | 9 | 5 | 9 | 5 | 0.832407111 | 0.864818478 | 1.313077322 | 1.364204505 |
| P17665 | Cox7c | Cytochrome c oxidase subunit 7C, mitochondrial | 39.45 | 14.29 | 7333 | 1 | 2330031 | 1625040.88 | 1735969.25 | 2176814.25 | 1465640.5 | 2730263.25 | 1785429.62 | 1775757.12 | 1497039.75 | 2255774 | 2191252.5 | 1902821.38 | 1744575.5 | 1912935.12 | 2346850.75 | 2685680.75 | 2545020.25 | 1943225.88 | 1581014.38 | 2064135.12 | 7 | 8 | 11 | 10 | 11 | 10 | 1.081956362 | 1.005393278 | 1.159166567 | 1.077139814 |
| O89001 | Cpd | Carboxypeptidase D | 302.47 | 4.28 | 152406 | 6 | 5757836 | 7323969 | 4113734.5 | 5498823 | 8541857 | 5424900.5 | 6386209 | 5074758 | 6348866.5 | 7666359.5 | 8072861 | 9626536 | 9687440 | 9924379 | 11156255 | 5105822.5 | 8358155 | 11883878 | 10465033 | 9911125 | 1 | 5 | 0 | 4 | 1 | 5 | 1.551643309 | 1.568471064 | 1.463813939 | 1.479689173 |
| Q8C166 | Cpne1 | Copine-1 | 501.89 | 20.9 | 58887 | 9 | 11092353 | 16456788 | 12464096 | 14388068 | 10736798 | 13301323 | 12858830 | 11352805 | 9876946 | 10796578 | 19649290 | 13638845 | 17321712 | 12870548 | 15272442 | 10518783 | 14920686 | 12252099 | 12358968 | 12976594 | 5 | 11 | 1 | 9 | 5 | 11 | 1.209013363 | 1.353455894 | 0.967592348 | 1.083191969 |
| Q8BT60 | Cpne3 | Copine-3 | 416.28 | 22.33 | 59585 | 8 | 4859156 | 5256580 | 5785397 | 5752997.5 | 5690934.5 | 5957534 | 5702587 | 6837383 | 5616764 | 6330988 | 4147779.75 | 5400756 | 5004093.5 | 5865918 | 5714047.5 | 5505257 | 5883325 | 4755922.5 | 6282172 | 5011158 | 11 | 12 | 4 | 5 | 11 | 12 | 0.955660363 | 0.85834702 | 1.00339255 | 0.901218715 |
| P36552 | Cpox | Oxygen-dependent coproporphyrinogen-III oxidase, mitochondrial | 183.19 | 9.71 | 49715 | 3 | 1711021.88 | 444992.31 | 408494.16 | 1454743.12 | 1444875.38 | 1656985.88 | 2131817 | 953199.81 | 1377525.25 | 1155773.75 | 802726 | 1192683 | 900899.44 | 665129.69 | 1240385.62 | 1200633.5 | 945907.88 | 776969.62 | 622368.12 | 838599.06 | 10 | 10 | 4 | 2 | 10 | 10 | 0.87879068 | 0.660017131 | 0.802411492 | 0.602652421 |
| Q66H71 | Cpped1 | Serine/threonine-protein phosphatase CPPED1 | 48.8 | 3.53 | 35261 | 1 | 1094485.62 | 1188337.88 | 1173629.38 | 796496.06 | 1381942.75 | 669278.62 | 866744.81 | 1165254.25 | 934586.81 | 853257.75 | 2 | 494528.78 | 2 | 2 | 461718.19 | 1120445.5 | 2 | 2 | 642677.31 | 704329.62 | 0 | 2 | 0 | 5 | 0 | 5 | 0.169702103 | 0.213015578 | 0.437888883 | 0.549652315 |
| Q9WVJ3 | Cpq | Carboxypeptidase Q | 184.04 | 10.43 | 51813 | 3 | 2336854.75 | 1589347.88 | 2066472.75 | 3994095.25 | 3762459.75 | 1654565.25 | 1515363.88 | 1996728.12 | 1901944.25 | 2466873 | 614710.12 | 236829.23 | 614426.19 | 1012373.62 | 417944.97 | 2744713.5 | 3158618 | 3846206.5 | 3411793.5 | 3640248.5 | 0 | 9 | 0 | 0 | 0 | 9 | 0.210650636 | 0.303737809 | 1.222001489 | 1.762007753 |
| Q9EPU4 | Cpsf1 | Cleavage and polyadenylation specificity factor subunit 1 | 339.55 | 5.48 | 160818 | 6 | 562085.5 | 974362.81 | 1233830.75 | 305424.5 | 708388.25 | 833700.31 | 374168.69 | 328803.69 | 1193761.75 | 296670.78 | 309266.38 | 405082.81 | 204548.98 | 420882.5 | 359751.97 | 1250294.12 | 567932.62 | 1105362.62 | 878523.88 | 287864.47 | 4 | 11 | 9 | 9 | 9 | 11 | 0.449125636 | 0.561438244 | 1.080834693 | 1.351118449 |
| Q9QXK7 | Cpsf3 | Cleavage and polyadenylation specificity factor subunit 3 | 153.24 | 5.26 | 77505 | 4 | 6167545 | 4688813.5 | 4825373 | 5948823 | 5190037 | 7435664.5 | 6083842.5 | 8272964 | 5669671.5 | 5342985.5 | 6144600 | 5440998 | 5154150 | 5750976.5 | 5589841.5 | 5515206 | 6375154 | 5401847 | 5698452 | 5904217 | 10 | 8 | 7 | 9 | 10 | 9 | 1.046977879 | 0.855980992 | 1.077339253 | 0.880803635 |
| Q6NVF9 | Cpsf6 | Cleavage and polyadenylation specificity factor subunit 6 | 447.12 | 17.6 | 59153 | 7 | 7367215.5 | 8446803 | 7471223 | 8605520 | 6678760 | 9573927 | 8241998 | 7200472 | 6767321 | 8148053 | 8961622 | 7368817 | 7148781 | 8637070 | 8131952 | 6151530 | 6487039.5 | 6422705 | 7456730.5 | 4155834.5 | 9 | 2 | 12 | 2 | 12 | 2 | 1.043524535 | 1.007925293 | 0.795287012 | 0.768156251 |
| Q5XI29 | Cpsf7 | Cleavage and polyadenylation specificity factor subunit 7 | 281.12 | 13.64 | 51073 | 6 | 3798085 | 1858408.25 | 3715232 | 3341292.25 | 3275650.5 | 3737975.5 | 3104751.5 | 3244838.5 | 1447562.5 | 3015394.25 | 1726687.25 | 1397467.62 | 1767951.25 | 1735990.25 | 2324665.75 | 974677 | 462638.22 | 3194001.75 | 927527.38 | 3079400.25 | 1 | 2 | 4 | 5 | 4 | 5 | 0.559944213 | 0.615288027 | 0.540272936 | 0.593672478 |
| Q64735 | Cr1l | Complement component receptor 1-like protein | 90.52 | 3.73 | 53763 | 1 | 2 | 2 | 2 | 2 | 2 | 2 | 2 | 2 | 2 | 2 | 2 | 497877.94 | 410748.06 | 2 | 604874 | 2 | 2 | 2 | 2 | 2 | 5 | 12.5 | 5 | 12.5 | 5 | 12.5 | 151350.4 | 151350.4 | 1 | 1 |
| O88668 | Creg1 | Protein CREG1 | 232.76 | 17.73 | 24452 | 4 | 2844224.75 | 2328332 | 1808015.5 | 2865231.5 | 2694705.75 | 3335767 | 3588065.75 | 3052793.5 | 4687482 | 4756861 | 3787437.75 | 3041952 | 4271372 | 5075047.5 | 4898390 | 5104928.5 | 6124831 | 5883303 | 5480842.5 | 6140475.5 | 0 | 0 | 9 | 0 | 9 | 0 | 1.680489876 | 1.085126029 | 2.291324806 | 1.4795544 |
| Q9CYA0 | Creld2 | Cysteine-rich with EGF-like domain protein 2 | 263.88 | 14.29 | 38220 | 4 | 346591.62 | 2 | 759090.31 | 448449.78 | 685971.44 | 2 | 752094.31 | 431974.19 | 2 | 761897.94 | 327163.09 | 379874.25 | 301902.09 | 2 | 378383.28 | 431942.16 | 564288.75 | 504445.97 | 487629.25 | 324763.94 | 6.5 | 12 | 9 | 12 | 9 | 12 | 0.619312317 | 0.71292178 | 1.032572096 | 1.188646046 |
| Q63768 | Crk | Adapter molecule crk | 86.79 | 3.95 | 33845 | 1 | 1350827.62 | 1154114.38 | 935866.31 | 899352.81 | 827302.56 | 826106.44 | 973307.75 | 787776.75 | 803028.5 | 396852.34 | 79334.99 | 629721.62 | 487503.28 | 212061.28 | 437925.44 | 1332492.25 | 493588.44 | 618052.19 | 589546.25 | 954405.56 | 0 | 7 | 3 | 12 | 3 | 12 | 0.357340994 | 0.487592187 | 0.771768306 | 1.053078717 |
| P47941 | Crkl | Crk-like protein | 55.85 | 3.96 | 33830 | 1 | 1752709.25 | 1730249.5 | 1659781.5 | 2007394.12 | 1447285.88 | 1669555.25 | 1751374.75 | 1257192.75 | 1395756.75 | 1605954.5 | 2 | 2 | 2 | 2 | 2 | 1751191.25 | 1001856.62 | 1280819.75 | 1675077 | 1781967.75 | 0 | 9 | 0 | 11 | 0 | 11 | 1.16314E-06 | 1.302111E-06 | 0.871297686 | 0.975400298 |
| Q9Z2L7 | Crlf3 | Cytokine receptor-like factor 3 | 56.31 | 4.98 | 49559 | 1 | 2 | 657632.38 | 754862.5 | 630406.31 | 899155.56 | 494105.5 | 2 | 634022.44 | 867738.69 | 694401.06 | 2 | 552917.12 | 371540.56 | 2 | 2 | 879765.25 | 2 | 268162.28 | 460830.94 | 568061.12 | 3.5 | 7.5 | 4.5 | 9.5 | 4.5 | 9.5 | 0.314223392 | 0.343632344 | 0.73989739 | 0.809146235 |

| | | | | | | | | | | | | | | | | | | | | | | | | | | | | | | | | | | | | |
|---|---|---|---|---|---|---|---|---|---|---|---|---|---|---|---|---|---|---|---|---|---|---|---|---|---|---|---|---|---|---|---|---|---|---|---|---|
| Q99KP3 | Cryl1 | Lambda-crystallin homolog | 91.07 | 5.64 | 35209 | 1 | 735152.19 | 686044.88 | 812075.88 | 864720.81 | 714443.25 | 798788.38 | 706216.12 | 527464.81 | 671193 | 717495.12 | 342654.72 | 200679.23 | 350701.22 | 2 | 2 | 624801.38 | 943206.56 | 2 | 822363.12 | 703246.31 | 0 | 10 | 0 | 12 | 0 | 12 | 0.234505952 | 0.261326521 | 0.811454553 | 0.904261038 |
| O54983 | Crym | Ketimine reductase mu-crystallin | 109.69 | 7.35 | 33523 | 2 | 2 | 124006.2 | 2 | 2 | 475602.16 | 303690.78 | 2 | 584975.06 | 2 | 460331.03 | 2 | 2 | 2 | 2 | 380907.62 | 2 | 149488.73 | 542165.62 | 361637.09 | 545453.19 | 10 | 5.5 | 7 | 11 | 10 | 11 | 0.635267674 | 0.282368699 | 2.666291431 | 1.185133876 |
| P47199 | Cryz | Quinone oxidoreductase | 95.66 | 11.18 | 35269 | 2 | 1975823 | 1871215.25 | 1676205 | 901993.62 | 1396860 | 2 | 1985010.5 | 696708.62 | 2727690 | 1127052.75 | 2 | 797851.25 | 255372.94 | 2 | 960799.62 | 2244011.75 | 2 | 587873.81 | 558853.62 | 943467 | 1 | 6 | 6 | 8.5 | 6 | 8.5 | 0.257479272 | 0.308121922 | 0.554097993 | 0.663081487 |
| Q921W4 | Cryzl1 | Quinone oxidoreductase-like protein 1 | 48.1 | 5.75 | 38725 | 1 | 258344.94 | 577172.19 | 297905.03 | 122366.63 | 716214.62 | 154307.73 | 503962.88 | 495091.25 | 1070452.12 | 303899.34 | 2 | 708037.88 | 250041.84 | 2 | 378757.81 | 209746.94 | 2 | 245108.86 | 174038.59 | 2 | 8 | 3 | 7 | 3 | 8 | 3 | 0.677910354 | 0.528873872 | 0.318913439 | 0.248801312 |
| Q9CZU6 | Cs | Citrate synthase, mitochondrial | 1068.53 | 47.63 | 51737 | 19 | 123543586 | 111880586 | 110289450 | 126750690 | 130171442 | 115904770 | 113669610 | 131632634 | 121170794 | 123145458 | 89736122 | 119881162 | 141822898 | 132816386 | 141446530 | 120488354 | 108391042 | 128379050 | 135400242 | 146363714 | 8 | 9 | 8 | 9 | 8 | 9 | 1.038277424 | 1.03332627 | 1.060379172 | 1.055322624 |
| P18395 | Csde1 | Cold shock domain-containing protein E1 | 767.53 | 22.68 | 88895 | 15 | 13324688 | 13540787 | 13771010 | 11039766 | 10099697 | 13382956 | 11447522 | 13586364 | 10761577 | 13809623 | 19450140 | 18738514 | 17474372 | 13413791 | 19336656 | 13286310 | 14525364 | 15547939 | 13829385 | 13714606 | 2 | 4 | 2 | 4 | 2 | 4 | 1.43119573 | 1.403654887 | 1.147754204 | 1.125667694 |
| Q9ERK4 | Cse1l | Exportin-2 | 1290.62 | 27.19 | 110455 | 22 | 35044874 | 26672250 | 27711090 | 28570796 | 31474068 | 28932230 | 30874414 | 27424730 | 31622756 | 25073286 | 23992470 | 29046872 | 17572098 | 18104896 | 18364610 | 39849250 | 37131714 | 28026026 | 28429596 | 24601616 | 3 | 11 | 3 | 11 | 3 | 11 | 0.716389516 | 0.743992694 | 1.057302118 | 1.098040988 |
| P09581 | Csf1r | Macrophage colony-stimulating factor 1 receptor | 600.42 | 12.59 | 109179 | 10 | 7902504 | 7764489 | 16146858 | 13933335 | 13668426 | 8177764 | 7491016 | 7565643 | 8167214 | 8599820 | 5957355.5 | 6134714.5 | 6690032.5 | 5705836 | 7150953.5 | 7398196 | 6294002.5 | 6751680 | 7439528 | 8636705 | 0 | 2 | 0 | 5 | 0 | 5 | 0.532501323 | 0.79094349 | 0.614655143 | 0.912969533 |
| P32577 | Csk | Tyrosine-protein kinase CSK | 539.03 | 24.22 | 50746 | 10 | 14913664 | 13335016 | 19599320 | 17386564 | 16316670 | 11849904 | 12859444 | 15972984 | 14994298 | 13205348 | 9236026 | 9805139 | 9181255 | 8359036 | 8915433 | 15947652 | 14225793 | 15611836 | 14215982 | 13185240 | 0 | 6 | 0 | 9 | 0 | 9 | 0.557893324 | 0.660504973 | 0.897429743 | 1.062491309 |
| P48729 | CSNK1A1 | Casein kinase I isoform alpha Homo sapiens OX=9606 | 50.87 | 2.97 | 38915 | 1 | 2 | 357405.56 | 254658.78 | 2 | 2 | 2 | 2 | 374418.41 | 281235.5 | 2 | 592499.81 | 709105.75 | 859391.88 | 2 | 549601.06 | 2 | 2 | 2 | 2 | 2 | 3.5 | 7.5 | 3.5 | 7.5 | 3.5 | 7.5 | 4.428576787 | 4.134156227 | 1.633799E-05 | 1.525181E-05 |
| Q06486 | Csnk1d | Casein kinase I isoform delta | 81.19 | 3.37 | 47316 | 2 | 570624.5 | 675733.12 | 587066.5 | 603603.56 | 408058.34 | 558225.94 | 681734.62 | 235784.59 | 314713.81 | 410417.69 | 547672.5 | 565516.88 | 317224.41 | 294336.09 | 196975.3 | 519153 | 260707.02 | 487777.81 | 231718.22 | 576836.75 | 2 | 4 | 10 | 11 | 10 | 11 | 0.675454157 | 0.87316351 | 0.729746934 | 0.943348097 |
| P19139 | Csnk2a1 | Casein kinase II subunit alpha | 303.53 | 20.46 | 45073 | 6 | 1631860.12 | 2446861.75 | 3346998 | 3661860.75 | 2934493.5 | 1580936 | 2036629.62 | 1421681.75 | 4263301.5 | 3025523.5 | 5113488 | 7876103 | 5341959 | 2815059.5 | 4370109.5 | 1701003.62 | 1406488 | 3207131.5 | 3393416 | 2398000.25 | 3 | 9 | 2 | 12 | 3 | 12 | 1.819753539 | 2.069806068 | 0.863355825 | 0.981989642 |
| O54833 | Csnk2a2 | Casein kinase II subunit alpha' | 77.33 | 4.57 | 41215 | 1 | 89111.52 | 159020.06 | 387026.72 | 260370.53 | 2 | 2 | 54602.31 | 201558.73 | 2 | 304227.62 | 2 | 344778.31 | 322383.62 | 202140.47 | 141257.19 | 197167.67 | 2 | 318075.66 | 2 | 323689.44 | 11.5 | 12 | 7 | 10 | 11.5 | 12 | 1.128449804 | 1.803309826 | 0.936803895 | 1.497051674 |
| P67871 | Csnk2b | Casein kinase II subunit beta | 42.68 | 5.12 | 24942 | 1 | 1111974.25 | 1691166.88 | 2 | 605168.88 | 460873.38 | 1171398 | 1074849.12 | 2 | 429146.81 | 646010.69 | 1941109.75 | 2076990.12 | 479543.69 | 637139.38 | 473936.66 | 1161514.12 | 1297650.12 | 2 | 427371.03 | 441641.5 | 8 | 10.5 | 9 | 12.5 | 9 | 12.5 | 1.449586679 | 1.688657922 | 0.860175576 | 1.00203894 |
| P97315 | Csrp1 | Cysteine and glycine-rich protein 1 | 159.12 | 25.39 | 20583 | 3 | 3196636.5 | 3660373.5 | 1608936.12 | 2799338 | 3140934 | 4541139 | 4415708 | 4275115.5 | 3221828.75 | 3339409 | 2991064.5 | 2644333 | 1680736 | 1976701.25 | 1454052 | 3309548 | 2588836.25 | 4334892.5 | 3231255.75 | 3875671 | 5 | 6 | 0 | 7 | 5 | 7 | 0.745989451 | 0.542958522 | 1.203661041 | 0.876068715 |
| P14841 | Cst3 | Cystatin-C | 48.61 | 7.86 | 15437 | 1 | 721487.94 | 957791.38 | 1305470.88 | 1350721 | 2 | 678253.25 | 995890.81 | 1684756.75 | 2 | 1712902 | 440886.22 | 2 | 633888.12 | 376103.72 | 644362 | 1123602.25 | 2 | 1130972.5 | 1033212.06 | 1326251.5 | 4.5 | 11.5 | 4.5 | 12.5 | 4.5 | 12.5 | 0.483278748 | 0.413115674 | 1.064252989 | 0.909743273 |
| Q62426 | Cstb | Cystatin-B | 161.85 | 41.84 | 11046 | 3 | 8472974 | 9935852 | 7092119.5 | 10921310 | 9204859 | 11780998 | 8575034 | 9470133 | 7088252 | 9584553 | 10517191 | 7793224 | 9727302 | 13101336 | 9661600 | 13334215 | 23149752 | 15108231 | 13749050 | 16594676 | 9 | 0 | 7 | 0 | 9 | 0 | 1.11338737 | 1.092511361 | 1.795772643 | 1.762101913 |
| Q8BIQ5 | Cstf2 | Cleavage stimulation factor subunit 2 | 193.57 | 10.34 | 61341 | 3 | 2 | 2 | 168919.98 | 175772.8 | 97571.51 | 103544.89 | 198147.36 | 139254.38 | 2 | 124211.99 | 1816623.25 | 1418831.25 | 1615494.25 | 1672684.5 | 1652313.5 | 2 | 171670.06 | 692029.12 | 2 | 2 | 0 | 12 | 0 | 10.5 | 0 | 12 | 18.48639601 | 14.4665896 | 1.952898726 | 1.52824728 |
| Q99LI7 | Cstf3 | Cleavage stimulation factor subunit 3 | 281.31 | 6.97 | 82877 | 4 | 1062641.5 | 746236.88 | 865839.5 | 843042.12 | 530829.88 | 973580.75 | 1103093 | 337176.41 | 961206 | 749848.25 | 1197545.38 | 969946.12 | 1115815.12 | 2 | 304738.56 | 1326596 | 2046384.5 | 1436053.25 | 1102324.5 | 1099629 | 11 | 0 | 12 | 2 | 12 | 2 | 0.886246147 | 0.869849777 | 1.731710906 | 1.69967266 |
| Q8R311 | Ctage5 | Endoplasmic reticulum export factor CTAGE5 | 279.31 | 9.37 | 87718 | 6 | 1121413.75 | 483141.25 | 2456018.5 | 1455946.75 | 3339517 | 992139.44 | 2155917 | 1009294.31 | 351298.28 | 3235322.5 | 1832651.38 | 2794864.75 | 1098086 | 1558942 | 3479078.25 | 335623.38 | 2230957.75 | 2844688 | 3857903.5 | 3321851.5 | 9 | 9 | 7 | 7 | 9 | 9 | 1.215399402 | 1.389935686 | 1.421744712 | 1.625913019 |
| O88712 | Ctbp1 | C-terminal-binding protein 1 | 432.26 | 19.27 | 47745 | 8 | 4837460 | 4373362 | 4441901 | 4639444 | 4206856.5 | 5290492 | 4579939 | 4116397.5 | 3925816.5 | 4041799 | 2912457 | 2926564.25 | 2902575 | 2998265.75 | 1697815 | 4692173 | 4455718.5 | 3836497 | 4373472.5 | 3559868.75 | 0 | 9 | 0 | 10 | 0 | 10 | 0.59725601 | 0.612070932 | 0.929717228 | 0.952778843 |
| P56546 | Ctbp2 | C-terminal-binding protein 2 | 414.83 | 16.63 | 48957 | 8 | 4490715.5 | 4985099 | 4433667.5 | 5545908.5 | 4337867.5 | 6426905 | 5681337 | 4096769.5 | 4995204.5 | 5272288.5 | 5109421.5 | 5305754 | 5572876.5 | 4869056.5 | 4208458 | 2365502.5 | 5248474 | 5909227 | 5392969 | 2430594.75 | 9 | 12 | 10 | 9 | 10 | 12 | 1.053473488 | 0.946852856 | 0.897177144 | 0.806375054 |
| Q5SUQ9 | Ctc1 | CST complex subunit CTC1 | 68.16 | 0.91 | 134028 | 1 | 377259.66 | 305969.53 | 354404.06 | 386156.78 | 445666.09 | 405283.44 | 550210.38 | 417154.94 | 376980.75 | 452127.88 | 175849.25 | 214894.31 | 265026.94 | 129092.23 | 282463 | 225921.61 | 2 | 334925.38 | 398748.81 | 402998.91 | 0 | 9 | 0 | 2 | 0 | 9 | 0.570928474 | 0.484760826 | 0.728873331 | 0.618867781 |
| Q61164 | Ctcf | Transcriptional repressor CTCF | 89.16 | 2.45 | 83745 | 1 | 305099.44 | 267622.34 | 321708.66 | 233637.69 | 295764.78 | 310896.09 | 365388.44 | 220003.52 | 231989.31 | 167699.58 | 2 | 147670.44 | 2 | 2 | 2 | 277983.22 | 331633 | 177337.94 | 2 | 2 | 0 | 7 | 0 | 8 | 0 | 8 | 0.10371894 | 0.113951441 | 0.552704011 | 0.607231607 |
| P58466 | Ctdsp1 | Carboxy-terminal domain RNA polymerase II polypeptide A small phosphatase 1 | 49.79 | 6.13 | 29266 | 1 | 141748.62 | 183201.53 | 158143.12 | 175914.92 | 2 | 83141.16 | 2 | 2 | 2 | 147139.75 | 2 | 2 | 222036.27 | 131487.78 | 136481.08 | 146678.09 | 239400.83 | 179542.11 | 100476.8 | 188952.39 | 8 | 8 | 9 | 2 | 9 | 8 | 0.743553191 | 2.127820161 | 1.297476478 | 3.712977954 |
| Q8VCN5 | Cth | Cystathionine gamma-lyase | 942.24 | 40.95 | 43567 | 15 | 38931342 | 39684586 | 22701240 | 39828798 | 34159694 | 48102042 | 41071706 | 38456990 | 35483394 | 43035890 | 105159770 | 82884922 | 94313754 | 95149810 | 99341730 | 46959098 | 56565026 | 57118294 | 58389214 | 59422318 | 0 | 0 | 0 | 1 | 0 | 1 | 2.720106048 | 2.313121199 | 1.588391099 | 1.350734515 |
| P30999 | Ctnnd1 | Catenin delta-1 | 160.26 | 4.37 | 104925 | 4 | 4337469 | 3043664 | 2904020 | 3821457.75 | 4186951.5 | 1571804.75 | 3800807 | 2764926.5 | 4020418.75 | 3370510.5 | 2963223.75 | 4171463.5 | 2644265.25 | 3377058 | 3418633 | 3702439 | 1471284.62 | 3750646 | 4532169 | 1871139.25 | 8 | 9 | 11 | 12 | 11 | 12 | 0.906036958 | 1.06737149 | 0.837872781 | 0.987069579 |
| P70698 | Ctps1 | CTP synthase 1 | 527.23 | 17.26 | 66682 | 8 | 5857988 | 6179817.5 | 5763454.5 | 7146328 | 8053995 | 5781857 | 6378918.5 | 6035521 | 7448615 | 8946788 | 7920633.5 | 8852870 | 7194070.5 | 11115574 | 7615722 | 6591878 | 5382390 | 6175920.5 | 6901159.5 | 8149298 | 3 | 12 | 5 | 12 | 5 | 12 | 1.293843086 | 1.234367511 | 1.006031923 | 0.959786494 |
| P70303 | Ctps2 | CTP synthase 2 | 139.61 | 6.48 | 65514 | 3 | 670679.94 | 1527360 | 2 | 1164482.12 | 2755551.25 | 455322.06 | 1572176.25 | 1551280.88 | 2498801 | 1100836 | 2 | 1191168.88 | 756978.12 | 2 | 2 | 963527.44 | 2 | 784398.06 | 2 | 2 | 6.5 | 5.5 | 3 | 2 | 6.5 | 5.5 | 0.318425796 | 0.271390366 | 0.285699572 | 0.243498211 |
| P16675 | Ctsa | Lysosomal protective protein | 1131.48 | 43.46 | 53844 | 18 | 133395418 | 130606898 | 129632746 | 148174402 | 166887522 | 105849642 | 124751426 | 111381714 | 116145970 | 101272954 | 26000130 | 26040210 | 26977068 | 24063008 | 26794096 | 146341250 | 131279202 | 121700338 | 124675034 | 132276658 | 0 | 7 | 0 | 2 | 0 | 7 | 0.183258169 | 0.232166814 | 0.926026913 | 1.173168539 |
| P10605 | Ctsb | Cathepsin B | 1390.44 | 59.88 | 37280 | 19 | 227687010 | 237342786 | 144279778 | 188717474 | 282802850 | 167335506 | 245120722 | 215620834 | 222223698 | 180320594 | 47628342 | 80215610 | 64645382 | 39798438 | 80146114 | 250517506 | 154833202 | 152005266 | 150147826 | 187809458 | 0 | 8 | 0 | 7 | 0 | 8 | 0.289068508 | 0.303150992 | 0.828357228 | 0.868712117 |
| P97821 | Ctsc | Dipeptidyl peptidase 1 | 350.48 | 11.04 | 52376 | 5 | 14882918 | 13536377 | 12625771 | 16922566 | 16400908 | 10056369 | 11104187 | 10505408 | 12143344 | 14608572 | 3926110.5 | 3988632.25 | 6805714 | 4995329.5 | 5691353.5 | 12099124 | 14572128 | 7845456.5 | 11764732 | 10175244 | 0 | 2 | 0 | 11 | 0 | 11 | 0.341638275 | 0.434920606 | 0.75914741 | 0.966428164 |
| P24268 | Ctsd | Cathepsin D | 576.48 | 19.66 | 44681 | 10 | 28492142 | 21037556 | 15338439 | 10309086 | 49814782 | 10294403 | 23875046 | 19274632 | 27578042 | 10057534 | 1218670.5 | 4259972 | 1393277.75 | 545463 | 2177100.25 | 2 | 10114845 | 5529561 | 7100870 | 8432397 | 0 | 0 | 0 | 1 | 0 | 1 | 0.076760778 | 0.105341674 | 0.249437354 | 0.34231217 |
| P18242 | Ctsd | Cathepsin D | 1781.51 | 54.63 | 44954 | 28 | 214010386 | 141707890 | 175723282 | 232967138 | 266084386 | 211509906 | 212960738 | 234712674 | 231453986 | 235514466 | 101865506 | 93374226 | 94719018 | 93631594 | 112494722 | 277949826 | 294248354 | 217899106 | 254964818 | 258123506 | 0 | 4 | 0 | 3 | 0 | 4 | 0.481405528 | 0.440513507 | 1.264623347 | 1.15720247 |
| P49935 | Ctsh | Pro-cathepsin H | 106.63 | 4.5 | 37170 | 1 | 404815.09 | 349718.38 | 479154.53 | 561990.44 | 806879.12 | 2 | 659772.94 | 768401.81 | 684899.5 | 2 | 2 | 2 | 2 | 2 | 2 | 328116.28 | 2 | 296866.03 | 375323.16 | 291178.56 | 0 | 1 | 5 | 9 | 5 | 9 | 3.842374E-06 | 4.732432E-06 | 0.496237259 | 0.611187035 |
| P06797 | Ctsl | Cathepsin L1 | 213.78 | 14.97 | 37547 | 3 | 1303869.75 | 1742213.62 | 1560192.88 | 534496.25 | 3286122.75 | 1092375 | 1349676.88 | 1650955.88 | 2302136.75 | 1223378.5 | 317393.03 | 1573316.12 | 2 | 536795.88 | 1155621 | 2531660.75 | 1301984.5 | 444054.16 | 2469060.75 | 1583271 | 5 | 12 | 4 | 10 | 5 | 12 | 0.425201444 | 0.470317938 | 0.988505364 | 1.093391875 |
| O70370 | Ctss | Cathepsin S | 706.33 | 38.53 | 38475 | 9 | 67497538 | 67125162 | 35839430 | 47299234 | 64095410 | 64323454 | 79080074 | 42865774 | 52315650 | 42232778 | 15772356 | 18940196 | 15141739 | 13974331 | 17016190 | 71862594 | 54928322 | 39866838 | 42796870 | 45202370 | 0 | 10 | 0 | 10 | 0 | 10 | 0.286829409 | 0.287890697 | 0.903497867 | 0.906840868 |
| Q9WUU7 | Ctsz | Cathepsin Z | 644.7 | 45.1 | 33996 | 11 | 301307426 | 366608706 | 186394642 | 260748866 | 247046274 | 239288658 | 269751266 | 173890210 | 163964898 | 179187298 | 53963922 | 55904946 | 51712094 | 47698950 | 51216690 | 257881410 | 248935074 | 148243970 | 174790066 | 177257506 | 0 | 4 | 0 | 12 | 0 | 12 | 0.191245482 | 0.253874952 | 0.739375709 | 0.98150801 |
| Q9WTX6 | Cul1 | Cullin-1 | 120.02 | 3.61 | 89692 | 3 | 2595892 | 2429602.5 | 2785762.5 | 3227002.75 | 3033803.25 | 2462422.75 | 2821746.5 | 3171341 | 2665338 | 3417696.5 | 2428842.5 | 4426985 | 4401578 | 4521054 | 5082683.5 | 2193414 | 3232775.25 | 2544411.5 | 3378134 | 2677201.5 | 5 | 12 | 5 | 11 | 5 | 12 | 1.482450938 | 1.434885221 | 0.996722105 | 0.964741416 |
| B5DF89 | Cul3 | Cullin-3 | 116.63 | 3.78 | 88932 | 2 | 641723.06 | 2 | 878661.38 | 2 | 250400.08 | 958666.94 | 908376.56 | 2 | 233033.03 | 598433.56 | 389236.69 | 676814.88 | 690103.75 | 2 | 690315.69 | 780612.12 | 372626.75 | 687108.12 | 2 | 583859.38 | 9 | 10 | 11.5 | 10.5 | 11.5 | 10.5 | 1.381572662 | 0.906600722 | 1.368999371 | 0.898350013 |
| Q3TCH7 | Cul4a | Cullin-4A | 149.43 | 3.82 | 87753 | 3 | 562657.19 | 502596.91 | 662265.19 | 812738.94 | 706983.75 | 470702.91 | 667217.94 | 576083.75 | 674552.75 | 587442.38 | 493629.84 | 508573.59 | 625106.56 | 2 | 530861.12 | 607149.69 | 664649.75 | 669888.75 | 667120.69 | 2 | 4 | 11 | 6 | 12 | 6 | 12 | 0.664617273 | 0.725192643 | 0.803392816 | 0.876616639 |
| A2A432 | Cul4b | Cullin-4B | 205.37 | 5.36 | 110699 | 4 | 300901.47 | 376484.41 | 503217.94 | 312496.94 | 562908.88 | 406709.09 | 432782.97 | 657685.25 | 657973.06 | 697616.75 | 605205.25 | 1080655.25 | 1022405.81 | 386208.97 | 550338 | 570079.75 | 744456.69 | 496191 | 832732.62 | 703741.06 | 3 | 2 | 11 | 6 | 11 | 6 | 1.772760793 | 1.277641366 | 1.628008476 | 1.173317337 |
| Q6MGD0 | Cuta | Protein CutA | 64.27 | 7.91 | 18659 | 1 | 3473019 | 2974854.75 | 2806350.5 | 3537871.5 | 2915292 | 2675715.5 | 3425591.75 | 2715197.75 | 2662199 | 2883027.5 | 1393326.5 | 1102284.75 | 1084233.12 | 1089290.12 | 988306.62 | 3349193 | 3400780.75 | 980415.38 | 2987158.25 | 2824246.25 | 0 | 10 | 0 | 10 | 0 | 10 | 0.360177084 | 0.393924723 | 0.862128945 | 0.942908147 |
| P53564 | Cux1 | Homeobox protein cut-like 1 | 50.56 | 1.32 | 165596 | 1 | 2 | 179823.3 | 196458.42 | 2 | 137282.84 | 2 | 136509.3 | 2 | 191037.09 | 2 | 2 | 2 | 2 | 2 | 2 | 2 | 2 | 2 | 2 | 2 | 5 | 5 | 7.5 | 7.5 | 7.5 | 7.5 | 1.94716E-05 | 3.052947E-05 | 1.94716E-05 | 3.052947E-05 |
| Q5BJP2 | Cwc15 | Spliceosome-associated protein CWC15 homolog | 225.58 | 21.4 | 26638 | 4 | 1968452.88 | 2528611 | 1931807.62 | 2217687.5 | 1427274.62 | 2022129.5 | 1546493.75 | 854838.38 | 1355755.25 | 1705034.75 | 1631119 | 1103685.62 | 430799.22 | 1332095.38 | 1391442.75 | 1292737.25 | 1823698 | 2068344.5 | 1913386.12 | 2094489.75 | 1 | 8 | 7 | 6 | 7 | 8 | 0.584597899 | 0.786871188 | 0.912528037 | 1.228266509 |
| Q8C5N3 | Cwc22 | Pre-mRNA-splicing factor CWC22 homolog | 159.78 | 6.5 | 104773 | 4 | 641660.38 | 571528.12 | 592714.81 | 685339.62 | 942176.56 | 443828.16 | 1003344.62 | 727568.94 | 663087.81 | 596588.12 | 366723.41 | 856099.69 | 863509.94 | 191536.08 | 965737.25 | 1489816 | 133560.67 | 498038.19 | 268183.34 | 417396.69 | 12 | 5 | 12 | 6 | 12 | 6 | 0.944716013 | 0.944441446 | 0.817550811 | 0.817313203 |
| Q8CI33 | Cwf19l1 | CWF19-like protein 1 | 41.29 | 2.23 | 60193 | 1 | 259209.44 | 2 | 238841.05 | 265324.16 | 2 | 262070.48 | 309322.03 | 270036.28 | 214901 | 371850.75 | 2 | 102316.33 | 80858.79 | 2 | 2 | 518500.34 | 326442.97 | 276351.97 | 191231.66 | 442620.62 | 7 | 3 | 0 | 8 | 7 | 8 | 0.239961021 | 0.128261879 | 2.299183452 | 1.228939557 |
| Q9CQX2 | Cyb5b | Cytochrome b5 type B | 316.49 | 58.22 | 16318 | 5 | 1376888.12 | 2354756 | 3105410 | 1008773.75 | 1437789.25 | 1263738 | 1654581.62 | 1468857 | 1813716.25 | 2239677 | 6360090 | 6773005 | 8791580 | 8028418 | 9347346 | 2433243.5 | 2596683 | 3948626 | 2649847.25 | 3138392 | 0 | 3 | 0 | 0 | 0 | 3 | 4.233311057 | 4.656135736 | 1.590629122 | 1.749501749 |
| Q9DB73 | Cyb5r1 | NADH-cytochrome b5 reductase 1 | 682.64 | 50.49 | 34135 | 12 | 14111291 | 15806638 | 7516598.5 | 11459522 | 10617681 | 12623020 | 14057831 | 12205023 | 12645846 | 15056724 | 21034610 | 26214476 | 33145326 | 27885792 | 32187298 | 10676115 | 12902689 | 12910420 | 14340060 | 19072378 | 0 | 8 | 0 | 10 | 0 | 10 | 2.36033301 | 2.109487676 | 1.174586278 | 1.049756651 |
| Q9DCN2 | Cyb5r3 | NADH-cytochrome b5 reductase 3 | 972.82 | 46.18 | 34128 | 14 | 43607954 | 42809978 | 37728798 | 43531538 | 36114694 | 45600294 | 43297914 | 41942534 | 40805942 | 51465694 | 76787674 | 67449250 | 71572978 | 52350722 | 76243866 | 41668962 | 44562422 | 49733694 | 52286794 | 50384954 | 0 | 3 | 0 | 8 | 0 | 8 | 1.689972444 | 1.543636857 | 1.170976778 | 1.069581294 |
| Q61462 | Cyba | Cytochrome b-245 light chain | 198.52 | 12.5 | 20748 | 4 | 7173113.5 | 6877934 | 7528410 | 7729957.5 | 7336593.5 | 6437414.5 | 5594079 | 6391683 | 6008047 | 7804476.5 | 5840125 | 5844569.5 | 10443563 | 9925128 | 9528534 | 5610812 | 1139665.75 | 6574363 | 9012803 | 7192310 | 10 | 7 | 8 | 11 | 10 | 11 | 1.13469164 | 1.289933816 | 0.805816376 | 0.916063673 |
| Q61093 | Cybb | Cytochrome b-245 heavy chain | 839.23 | 29.3 | 65305 | 14 | 7644749.5 | 6439553 | 10833598 | 13610290 | 11816560 | 8851723 | 9517807 | 10575317 | 12291670 | 11611803 | 12012609 | 13311517 | 22758782 | 20157186 | 22831006 | 7401418.5 | 5422021.5 | 12068021 | 12187429 | 12963714 | 2 | 12 | 1 | 12 | 2 | 12 | 1.808949277 | 1.723254401 | 0.993998451 | 0.946910025 |
| Q3TYS2 | Cybc1 | Cytochrome b-245 chaperone 1 | 49.19 | 5.35 | 20921 | 1 | 2 | 2 | 2 | 2 | 1358363.88 | 2 | 2 | 1076995.62 | 1036351.38 | 2 | 2 | 907431.94 | 1018768.56 | 780899.38 | 2 | 2 | 2 | 925975.12 | 2 | 2 | 9 | 12 | 12 | 9 | 12 | 12 | 1.992903357 | 1.280952061 | 0.68168602 | 0.438158282 |
| Q9D0M3 | Cyc1 | Cytochrome c1, heme protein, mitochondrial | 354.02 | 43.69 | 35328 | 7 | 6370051 | 7750000 | 9304479 | 7627395.5 | 9136189 | 6764911 | 6593894.5 | 7329207.5 | 7615947 | 6479375 | 9955296 | 10793203 | 10228164 | 10125768 | 9202940 | 8406211 | 9031330 | 8105640.5 | 8375333 | 9660940 | 1 | 8 | 0 | 0 | 1 | 8 | 1.251747479 | 1.446249217 | 1.084386641 | 1.252883155 |
| P62897 | Cycs | Cytochrome c, somatic | 493.77 | 58.1 | 11605 | 8 | 49704282 | 48117066 | 35428586 | 43605522 | 38304610 | 52847270 | 47305654 | 41012570 | 36324342 | 43760318 | 26555042 | 30518744 | 33549436 | 35179258 | 37381330 | 48252870 | 48410638 | 46447130 | 47034958 | 55866758 | 1 | 6 | 1 | 6 | 1 | 6 | 0.758429819 | 0.73755343 | 1.143392259 | 1.11191947 |
| Q7TMB8 | Cyfip1 | Cytoplasmic FMR1-interacting protein 1 | 1773.73 | 23.86 | 145241 | 32 | 11850161 | 11096112 | 14482419 | 15196704 | 17271088 | 10476219 | 10362880 | 15120561 | 17509638 | 13982860 | 12232101 | 13330688 | 13366751 | 10548557 | 12547584 | 10176378 | 11008725 | 12578221 | 14115264 | 12124053 | 8 | 6 | 10 | 9 | 10 | 9 | 0.887393435 | 0.919550728 | 0.858450062 | 0.889558508 |
| Q5SQX6 | Cyfip2 | Cytoplasmic FMR1-interacting protein 2 | 831.83 | 13.73 | 145659 | 16 | 356855.81 | 397036.72 | 475176.94 | 544225.69 | 691382.12 | 421745.28 | 432188.34 | 622916.75 | 642624.44 | 589192.25 | 212102.48 | 321919.66 | 437719.59 | 494379.47 | 410457.22 | 439697.91 | 305729.66 | 479406 | 503529.75 | 2 | 7 | 8 | 4 | 6 | 7 | 8 | 0.761389102 | 0.692805125 | 0.70125421 | 0.63808703 |
| Q8BKE6 | Cyp20a1 | Cytochrome P450 20A1 | 131.96 | 7.58 | 52149 | 3 | 746238.81 | 643629.56 | 580776.12 | 2 | 681437.25 | 392378.03 | 701323.62 | 579586.31 | 519170.19 | 2 | 1175618.75 | 1897439.75 | 751774.25 | 907403.94 | 979668.5 | 745240.69 | 2 | 2 | 2 | 415797.16 | 0 | 6.5 | 0 | 8.5 | 0 | 8.5 | 2.153742397 | 2.605249263 | 0.437785517 | 0.529562122 |
| Q9DBG1 | Cyp27a1 | Sterol 26-hydroxylase, mitochondrial | 101.64 | 4.69 | 60720 | 2 | 2 | 95629.26 | 106594.07 | 2 | 800773.38 | 2 | 385119.41 | 2 | 472941.25 | 506706.59 | 244058.38 | 543804.44 | 2 | 2 | 812397.5 | 862322.62 | 2 | 611944.81 | 360711.47 | 489336.78 | 10 | 7 | 11 | 8 | 11 | 8 | 1.595476757 | 1.172551312 | 2.317363943 | 1.703082242 |
| Q64654 | Cyp51a1 | Lanosterol 14-alpha demethylase | 88.66 | 4.57 | 56707 | 2 | 852211.69 | 594935.19 | 634266.5 | 2 | 422087.12 | 2 | 565994.44 | 535076.38 | 443097.12 | 274380.28 | 2 | 2 | 312863.56 | 2 | 201642.02 | 591322.12 | 2 | 2 | 2 | 2 | 3.5 | 4 | 4.5 | 7 | 4.5 | 7 | 0.205516703 | 0.282924043 | 0.23620113 | 0.325165681 |
| Q80YW0 | Cyth4 | Cytohesin-4 | 178.59 | 6.36 | 45285 | 2 | 425624.12 | 632663.44 | 748862.19 | 429639.31 | 941681.25 | 469854.16 | 438036.41 | 2 | 765557 | 283545.88 | 754579.69 | 700701.31 | 611196.75 | 469227.91 | 509182.56 | 286548.72 | 386836.62 | 391421.53 | 738521.19 | 370739.06 | 12 | 3 | 6 | 12 | 12 | 12 | 0.957972837 | 1.555899489 | 0.683997932 | 1.110920886 |
| P98078 | Dab2 | Disabled homolog 2 | 975.9 | 28.07 | 82312 | 17 | 9788383 | 11944442 | 18181304 | 14667054 | 23677686 | 12190677 | 14803209 | 22300230 | 25822870 | 19222242 | 12358682 | 13694463 | 14784455 | 15439975 | 17312650 | 12081766 | 9378001 | 20187148 | 15565628 | 22314666 | 12 | 12 | 7 | 9 | 12 | 12 | 0.940343579 | 0.780059648 | 1.016206981 | 0.842991942 |
| P61804 | Dad1 | Dolichyl-diphosphooligosaccharide--protein glycosyltransferase subunit DAD1 | 275.11 | 35.4 | 12497 | 5 | 10470926 | 12811956 | 14223651 | 14010104 | 13973814 | 12306418 | 12763802 | 17467618 | 15727106 | 20385484 | 22323038 | 25797446 | 40525470 | 41786242 | 37469026 | 10679342 | 13241611 | 19977326 | 24555152 | 20216342 | 0 | 7 | 0 | 10 | 0 | 10 | 2.563751195 | 2.134778237 | 1.353934377 | 1.127390852 |
| Q91WC9 | Daglb | Sn1-specific diacylglycerol lipase beta | 98.28 | 3.29 | 73905 | 2 | 270742.56 | 221522.75 | 263942.19 | 154196.7 | 2 | 163851.92 | 176826.39 | 237398.48 | 465706.75 | 230430.31 | 136923.7 | 161810.12 | 178308.97 | 152682.42 | 120174.52 | 201619.17 | 2 | 204885.48 | 128441.18 | 2 | 7 | 6 | 2 | 4 | 7 | 6 | 0.82369796 | 0.588519525 | 0.587594669 | 0.419827355 |
| Q9ER88 | Dap3 | 28S ribosomal protein S29, mitochondrial | 217.34 | 13.3 | 44699 | 4 | 1859439 | 2246164.5 | 1444593.5 | 1848837.12 | 1917494.25 | 1962263.75 | 2055437.75 | 707267.81 | 1711053.5 | 1290800 | 1428478.62 | 2152507 | 2656152 | 1639805.12 | 2633939.25 | 1663648.5 | 2036076 | 2000555.25 | 1890748.62 | 1711469.5 | 10 | 12 | 6 | 9 | 10 | 12 | 1.128197283 | 1.36031099 | 0.998494021 | 1.203922763 |
| Q922B2 | Dars | Aspartate--tRNA ligase, cytoplasmic | 1532.61 | 54.69 | 57147 | 26 | 83683754 | 85106370 | 86362410 | 85487370 | 94836450 | 92638922 | 97067698 | 106057682 | 111598546 | 119959690 | 104408914 | 131346442 | 109350314 | 91293954 | 109818770 | 89728058 | 74409490 | 106290402 | 91385162 | 101458330 | 1 | 7 | 12 | 5 | 12 | 7 | 1.254300926 | 1.035833583 | 1.063826859 | 0.878535258 |
| Q9JII5 | Dazap1 | DAZ-associated protein 1 | 199.49 | 11.33 | 43214 | 3 | 932959.56 | 1309160.88 | 1330261.62 | 1107980.75 | 1627089.75 | 571237.75 | 1402222 | 1029748.44 | 1755761.25 | 974190.81 | 1076537.5 | 860094.06 | 992289.12 | 1089513.62 | 435775.25 | 1322608.75 | 1462385.75 | 1513918.75 | 1295893.88 | 1386141.38 | 3 | 8 | 9 | 8 | 9 | 8 | 0.706182014 | 0.776920469 | 1.106777807 | 1.217644058 |
| P31786 | Dbi | Acyl-CoA-binding protein | 217.95 | 50.57 | 10000 | 3 | 10634191 | 9164357 | 12138297 | 14119228 | 12166713 | 10097446 | 8429456 | 12619304 | 10330353 | 9496982 | 4078808 | 5613093 | 6361463.5 | 4408121 | 6142115.5 | 12271616 | 11459901 | 10699329 | 9096512 | 12448279 | 0 | 12 | 0 | 8 | 0 | 12 | 0.456927654 | 0.521910004 | 0.961404303 | 1.098131225 |
| Q62418 | Dbnl | Drebrin-like protein | 495.07 | 21.1 | 48700 | 7 | 4061173 | 6864657 | 4340706.5 | 6315556.5 | 4282886 | 4715205 | 4502619.5 | 6016644 | 5039675.5 | 4677979.5 | 6147922 | 4721783 | 3479767 | 5067844.5 | 2996988.5 | 4685999 | 5021959.5 | 6540473.5 | 5306071.5 | 5460368 | 9 | 9 | 12 | 7 | 12 | 9 | 0.86658895 | 0.898292484 | 1.044457508 | 1.082668235 |
| P53395 | Dbt | Lipoamide acyltransferase component of branched-chain alpha-keto acid dehydrogenase complex, mitochondrial | 387.79 | 23.44 | 53247 | 8 | 2857530.5 | 4603374 | 6200970.5 | 2497960.75 | 6132585.5 | 4566314.5 | 5067505 | 6868625 | 8490765 | 6623059 | 1718749.62 | 4777044 | 6005269 | 2015249.88 | 7781777 | 5418568.5 | 2627102.5 | 5756863 | 6485373 | 5636080.5 | 11 | 10 | 7 | 8 | 11 | 10 | 1.000254268 | 0.705272651 | 1.162905869 | 0.819957216 |
| Q8BHC4 | Dcakd | Dephospho-CoA kinase domain-containing protein | 295.18 | 23.81 | 26476 | 5 | 3589117 | 4485932.5 | 6453290 | 5145522 | 5348524 | 3950878 | 3036588.5 | 4491708 | 4810211 | 3600944.5 | 2705653.75 | 4380515 | 5666666 | 4052161.75 | 4845612 | 3916203.5 | 3421887.75 | 4849230.5 | 4030911 | 4406623 | 8 | 5 | 9 | 11 | 9 | 11 | 0.865249578 | 1.08849921 | 0.824256175 | 1.036928787 |
| Q9DAR7 | Dcps | m7GpppX diphosphatase | 428.35 | 32.25 | 38988 | 8 | 3966325.75 | 4859376 | 8467326 | 6631604 | 6164466.5 | 4082162.75 | 4391243.5 | 6509106 | 5938030 | 5887626 | 5239376 | 4634130.5 | 4117914.5 | 4994943 | 3842944 | 2601841.5 | 2930651.5 | 6859996 | 5013741 | 5748307 | 6 | 8 | 7 | 9 | 7 | 9 | 0.758723569 | 0.851580301 | 0.769532434 | 0.863712014 |
| O08788 | Dctn1 | Dynactin subunit 1 | 337.82 | 3.75 | 141676 | 4 | 1401434.38 | 1727162.38 | 1181334.62 | 1057623.12 | 1739546.62 | 739752.5 | 1372941.12 | 1484869.38 | 1776677 | 1458482.62 | 605177.44 | 1488474.62 | 2 | 684504.44 | 992307.94 | 1975221 | 646135.56 | 1195636.38 | 1003265.56 | 716392.25 | 3 | 7 | 5 | 7 | 5 | 7 | 0.530521007 | 0.551824895 | 0.779030811 | 0.810313993 |
| Q6AYH5 | Dctn2 | Dynactin subunit 2 | 338.51 | 18.66 | 44148 | 6 | 5959974.5 | 8162913.5 | 9791227 | 8089291.5 | 9545225 | 3924589.5 | 4072486.25 | 9122600 | 10249373 | 8690826 | 2442434.5 | 5585399 | 3070481 | 1729904.88 | 5282295.5 | 4232222 | 3010063.75 | 7366309.5 | 7960053.5 | 7981573 | 0 | 3 | 4 | 8 | 4 | 8 | 0.435887157 | 0.502234548 | 0.735288279 | 0.847208205 |
| Q8CBY8 | Dctn4 | Dynactin subunit 4 | 156.48 | 5.78 | 53057 | 2 | 2073636.5 | 1296095.25 | 2169572 | 2416345.25 | 2020182.25 | 1681754.12 | 1625209.5 | 1824548 | 2325559.75 | 2105989.75 | 1308593.62 | 490002.59 | 627527.62 | 1297798.62 | 1236672.62 | 1325548.75 | 1945475.5 | 2173865.75 | 2504719.25 | 2622946 | 2 | 9 | 0 | 8 | 2 | 9 | 0.497261325 | 0.518724602 | 1.05981697 | 1.105561819 |
| Q9QZ73 | Dcun1d1 | DCN1-like protein 1 | 135.39 | 11.58 | 30097 | 2 | 717088.81 | 214265.86 | 1184051.88 | 870282.56 | 856010.62 | 700984.94 | 653975.56 | 547598.25 | 529388.94 | 932681.25 | 2 | 584134.31 | 664097.69 | 553329.31 | 576940.75 | 732859 | 735583.56 | 1034775.25 | 833052.19 | 890129.31 | 4 | 11 | 9 | 4 | 9 | 11 | 0.619128049 | 0.706914225 | 1.100137857 | 1.256126421 |
| Q3U1J4 | Ddb1 | DNA damage-binding protein 1 | 1623.21 | 31.23 | 126853 | 30 | 7544031.5 | 7297385 | 6805009 | 7817289 | 5373368.5 | 8107798 | 6450220.5 | 5223936.5 | 6262101.5 | 6449267 | 7682286 | 5459999.5 | 5732548.5 | 7065779.5 | 5618304 | 6729133 | 9753752 | 9181531 | 7946559 | 6521367.5 | 9 | 8 | 11 | 3 | 11 | 8 | 0.905900115 | 0.971243139 | 1.152000657 | 1.235095034 |
| A2ADY9 | Ddi2 | Protein DDI1 homolog 2 | 86.89 | 4.76 | 44591 | 1 | 460215.62 | 1660463.88 | 1770114.88 | 1806054.12 | 1609955.75 | 1423775.88 | 1750502.75 | 1380170 | 1722849.88 | 1234616.12 | 728184.12 | 805751.31 | 1061548.38 | 2 | 209183.59 | 1930486.5 | 1573329.62 | 2 | 1139201.62 | 1654532.62 | 3 | 9 | 0 | 11 | 3 | 11 | 0.383843511 | 0.373362789 | 0.861875061 | 0.838341844 |
| O54734 | Ddost | Dolichyl-diphosphooligosaccharide--protein glycosyltransferase 48 kDa subunit | 939.88 | 39.68 | 49028 | 14 | 20915626 | 20031972 | 24709282 | 23269566 | 21556386 | 25729610 | 20606310 | 23082530 | 23075630 | 26319756 | 65929622 | 46711034 | 56523034 | 49633682 | 52499346 | 17429464 | 22019734 | 30279706 | 22128570 | 25773916 | 0 | 9 | 0 | 11 | 0 | 11 | 2.45555543 | 2.283376475 | 1.064702885 | 0.99004791 |
| Q80WW9 | Ddrgk1 | DDRGK domain-containing protein 1 | 242.32 | 18.1 | 35977 | 4 | 1353635.5 | 784227.38 | 520440.5 | 862021.25 | 925777.12 | 2122104.75 | 1441683.75 | 970293 | 730194.5 | 1567458 | 870779.62 | 348400.19 | 701798.75 | 97311.08 | 464659.41 | 1402133.5 | 1730987.75 | 1478475.75 | 1412903.5 | 779939 | 4 | 4 | 1 | 12 | 4 | 12 | 0.558455292 | 0.363443461 | 1.5304282 | 0.996004748 |
| O35215 | Ddt | D-dopachrome decarboxylase | 92.48 | 16.1 | 13077 | 2 | 3125647 | 3987796.75 | 3750068.25 | 5805327 | 4659468 | 4137218.5 | 3829063.25 | 4899268.5 | 4047608.75 | 5310407.5 | 3233126.75 | 3297980.75 | 3235715.25 | 3841709 | 3919854 | 3654239 | 4186800.25 | 5546284.5 | 6873409.5 | 6761061 | 7 | 7 | 2 | 7 | 7 | 7 | 0.821836714 | 0.788729647 | 1.26694511 | 1.215907188 |
| Q641Y8 | Ddx1 | ATP-dependent RNA helicase DDX1 | 817.53 | 20.41 | 82497 | 13 | 13143343 | 12113606 | 10890779 | 11664676 | 12860899 | 13740650 | 10723055 | 11063710 | 9778413 | 12343066 | 11453202 | 10795550 | 11294814 | 6572909.5 | 10780946 | 12098652 | 7134268 | 13634259 | 10882483 | 11017388 | 2 | 8 | 10 | 11 | 10 | 11 | 0.838876721 | 0.882886348 | 0.90265483 | 0.95001042 |
| Q80Y44 | Ddx10 | Probable ATP-dependent RNA helicase DDX10 | 105.88 | 3.31 | 100739 | 2 | 708754.69 | 773512.06 | 647299.31 | 818686.25 | 189144.8 | 1016384.88 | 530037.88 | 184747.17 | 182454.56 | 773671.25 | 1213203.12 | 1159950.62 | 1546014.75 | 983782 | 1147964.88 | 611027.62 | 498868.31 | 701127.62 | 510031.47 | 748771 | 0 | 8 | 1 | 12 | 1 | 12 | 1.928641851 | 2.251674529 | 0.978462691 | 1.142347667 |
| Q501J6 | Ddx17 | Probable ATP-dependent RNA helicase DDX17 | 1195.27 | 31.69 | 72399 | 20 | 16069296 | 21299302 | 22525162 | 22054908 | 25131066 | 16138516 | 19960414 | 21083836 | 19944432 | 18820302 | 9851357 | 9546976 | 11767068 | 7322714 | 11627910 | 21681464 | 17088558 | 17356752 | 18575230 | 20090958 | 0 | 6 | 0 | 12 | 0 | 12 | 0.468025303 | 0.522327575 | 0.88525586 | 0.987966982 |
| Q8K363 | Ddx18 | ATP-dependent RNA helicase DDX18 | 1237.5 | 37.12 | 74181 | 20 | 12644099 | 13999926 | 15630689 | 12616488 | 14841593 | 17911370 | 17860454 | 13264922 | 13356821 | 12151103 | 17694206 | 21976210 | 20626754 | 18435572 | 14893623 | 12731645 | 11474333 | 13492730 | 15163852 | 11176954 | 1 | 8 | 4 | 7 | 4 | 8 | 1.342644662 | 1.255976651 | 0.918355761 | 0.859075692 |
| Q61655 | Ddx19a | ATP-dependent RNA helicase DDX19A | 430.32 | 17.36 | 53933 | 8 | 6650934.5 | 4279058 | 5378205.5 | 6522997.5 | 3363693.5 | 4094296.5 | 4073897.5 | 6928175.5 | 5466416.5 | 3924953.5 | 9063342 | 8575493 | 6998061.5 | 9036992 | 7657545 | 1560178.75 | 4610726.5 | 4420766.5 | 6975163.5 | 3725118 | 0 | 10 | 0 | 11 | 0 | 11 | 1.57784343 | 1.68784193 | 0.812828535 | 0.869494436 |
| Q9JJY4 | Ddx20 | Probable ATP-dependent RNA helicase DDX20 | 64.19 | 1.7 | 91710 | 1 | 2 | 161953.23 | 2 | 2 | 2 | 155947.12 | 2 | 2 | 353766.47 | 226106.3 | 2 | 2 | 263638.5 | 301231.34 | 2 | 291636.31 | 2 | 2 | 2 | 2 | 9 | 12 | 11 | 8 | 11 | 12 | 3.487722586 | 0.767678038 | 1.800704465 | 0.396350695 |
| Q9JIK5 | Ddx21 | Nucleolar RNA helicase 2 | 2452.57 | 48.53 | 93551 | 40 | 63928462 | 66031342 | 63182518 | 49324662 | 53209618 | 72421594 | 58987374 | 53742974 | 50561802 | 53818146 | 96736234 | 84203210 | 69593818 | 70375978 | 69843554 | 62880818 | 74487002 | 49517906 | 54935234 | 54731902 | 0 | 12 | 3 | 10 | 3 | 12 | 1.321554669 | 1.349601918 | 1.002963576 | 1.024249391 |
| Q9ESV0 | Ddx24 | ATP-dependent RNA helicase DDX24 | 165.27 | 5.02 | 96429 | 4 | 1746706.88 | 2478110 | 1033239.25 | 2333603.5 | 1472814.75 | 3299732.5 | 1641726 | 1362157.88 | 2258057 | 2363034.5 | 2440200.75 | 3392901.5 | 2622918.25 | 2111026 | 791364.94 | 3293444.25 | 3245459.5 | 3117024.25 | 3278425.5 | 2739119.5 | 8 | 0 | 10 | 5 | 10 | 5 | 1.253068955 | 1.039699328 | 1.729109967 | 1.434681199 |
| Q921N6 | Ddx27 | Probable ATP-dependent RNA helicase DDX27 | 100.23 | 2.63 | 85939 | 2 | 2 | 321288.94 | 2 | 2 | 2 | 2 | 2 | 413319.34 | 2 | 2 | 1874121.62 | 1293865.62 | 1140336.38 | 1041999.62 | 1162560 | 2 | 2 | 282114.09 | 2 | 2 | 0 | 12 | 0 | 12 | 0 | 12 | 20.27060463 | 15.75720406 | 0.878072757 | 0.682563341 |
| Q6NZQ2 | Ddx31 | Probable ATP-dependent RNA helicase DDX31 | 51.2 | 1.89 | 76913 | 1 | 329581.12 | 279394.38 | 281212.44 | 327733 | 2 | 336157.09 | 420212.44 | 133420.3 | 2 | 2 | 443862.94 | 357087.88 | 214260.23 | 360728.84 | 199943.08 | 2 | 205340.25 | 414259.41 | 103497.82 | 262231.5 | 8 | 8.5 | 6 | 12 | 8 | 12 | 1.293910245 | 1.771065293 | 0.809025717 | 1.107369985 |
| Q5U216 | Ddx39a | ATP-dependent RNA helicase DDX39A | 699.91 | 28.1 | 49109 | 12 | 7287421.5 | 7262019 | 8078872.5 | 8864285 | 7919325.5 | 8447836 | 7088198.5 | 7383833.5 | 6794994 | 8361025 | 11297392 | 10639778 | 11073890 | 9328365 | 5993323.5 | 6169045.5 | 7868360 | 7802997 | 8209812 | 7873956.5 | 5 | 10 | 5 | 12 | 5 | 12 | 1.226348379 | 1.26937945 | 0.962251208 | 0.996015431 |
| Q63413 | Ddx39b | Spliceosome RNA helicase Ddx39b | 1241.82 | 50.93 | 49035 | 20 | 25447358 | 24410644 | 32732670 | 33207510 | 24134258 | 27626986 | 24174278 | 25638444 | 28859106 | 26629166 | 21657576 | 19966526 | 15534338 | 17997040 | 19397272 | 24395200 | 22276408 | 20457640 | 20125906 | 27366250 | 0 | 4 | 0 | 4 | 0 | 4 | 0.675702875 | 0.711308123 | 0.819119598 | 0.862281997 |
| Q62167 | Ddx3x | ATP-dependent RNA helicase DDX3X | 1508.77 | 34.14 | 73101 | 23 | 9833914 | 11959182 | 7842665 | 6302093 | 7432325 | 13665614 | 10400670 | 9578825 | 6878013 | 9299997 | 18000072 | 13465462 | 12661444 | 13179058 | 11714485 | 9971447 | 12800722 | 7113386 | 9639688 | 9491902 | 1 | 9 | 4 | 12 | 4 | 12 | 1.591428087 | 1.385311124 | 1.13020389 | 0.983823293 |
| Q91VN6 | Ddx41 | Probable ATP-dependent RNA helicase DDX41 | 99.48 | 4.98 | 69820 | 2 | 326385.94 | 482943.69 | 228461.88 | 966272.75 | 807081.75 | 175491.55 | 300736.75 | 227187.39 | 509100.34 | 2 | 163497.62 | 231019.2 | 456769.41 | 97371.83 | 196606.52 | 408442.69 | 366351.88 | 2 | 225039.06 | 2 | 3 | 4 | 11 | 11 | 11 | 11 | 0.407401315 | 0.944534062 | 0.35566905 | 0.824596093 |
| Q810A7 | Ddx42 | ATP-dependent RNA helicase DDX42 | 235.6 | 5.81 | 101965 | 5 | 2823988 | 3403991 | 3960422.75 | 4340417 | 3063020.5 | 4146535.75 | 4331446.5 | 2715572 | 3031044.5 | 3183671.75 | 1764379.5 | 1568656.75 | 2493289.25 | 3319938 | 1081724.5 | 3921821 | 3960823 | 3202708.5 | 3557812 | 3295353.25 | 2 | 11 | 3 | 10 | 3 | 11 | 0.581405267 | 0.587536137 | 1.01970678 | 1.030459502 |
| Q569Z5 | Ddx46 | Probable ATP-dependent RNA helicase DDX46 | 592.71 | 13.95 | 117448 | 12 | 12671807 | 10429213 | 11270568 | 13951172 | 11358139 | 12965018 | 12152149 | 12844627 | 11623509 | 10144438 | 11802026 | 10522618 | 12288838 | 10353354 | 10584631 | 9968004 | 12804407 | 13074935 | 13982283 | 12190840 | 8 | 9 | 8 | 9 | 8 | 9 | 0.930808147 | 0.93004701 | 1.03920132 | 1.038351548 |
| Q9CWX9 | Ddx47 | Probable ATP-dependent RNA helicase DDX47 | 473.91 | 19.12 | 50639 | 9 | 4210993.5 | 4199856.5 | 3807033.75 | 1444020 | 3295991 | 4044662 | 3522336 | 2000602.25 | 2435649 | 2248510 | 4814973.5 | 4235272.5 | 4894923 | 4578365 | 4310187 | 3254863 | 2041247.75 | 2079435.62 | 3086381.5 | 2711935 | 0 | 5 | 0 | 11 | 0 | 11 | 1.34649503 | 1.602168588 | 0.776857214 | 0.924367486 |
| Q4FZF3 | Ddx49 | Probable ATP-dependent RNA helicase DDX49 | 44.1 | 2.29 | 54094 | 1 | 307963.88 | 325931.25 | 392271.53 | 167037.45 | 374389.34 | 2 | 2 | 106969.55 | 289833.28 | 153065.86 | 2 | 372615.41 | 274959.22 | 2 | 327342.84 | 257282.69 | 2 | 95382.55 | 2 | 190315.11 | 7 | 2 | 9 | 12 | 9 | 12 | 0.621922393 | 1.772994891 | 0.346380849 | 0.987472846 |
| Q61656 | Ddx5 | Probable ATP-dependent RNA helicase DDX5 | 2002.21 | 54.07 | 69290 | 36 | 65959414 | 69693442 | 78875386 | 72860770 | 66095930 | 70279594 | 69612522 | 53951686 | 58947358 | 56937642 | 60172938 | 60963674 | 54981490 | 44070242 | 51081278 | 72167562 | 64203102 | 63146594 | 65230386 | 65229754 | 0 | 3 | 7 | 8 | 7 | 8 | 0.76741493 | 0.875829501 | 0.93349775 | 1.065375244 |
| Q99MJ9 | Ddx50 | ATP-dependent RNA helicase DDX50 | 64.88 | 2.59 | 82175 | 2 | 1154453.12 | 873487 | 1007801.19 | 262209.03 | 833737 | 335561.94 | 248984.56 | 258115.83 | 1520943.38 | 208939.73 | 278739.16 | 2 | 190598.08 | 2 | 187060.75 | 861428.75 | 642471.44 | 580669.81 | 1097552.62 | 2 | 1 | 8 | 3 | 9 | 3 | 9 | 0.158870199 | 0.255156616 | 0.770175562 | 1.236955651 |
| Q6P9R1 | Ddx51 | ATP-dependent RNA helicase DDX51 | 90.06 | 2.97 | 70368 | 2 | 962990 | 961472.38 | 2 | 2 | 900453.75 | 321291.53 | 1006298.38 | 468710.16 | 407196.88 | 987315.38 | 1080042.88 | 901311.88 | 2 | 771314.44 | 1204833.38 | 799271.81 | 589280.5 | 173139.22 | 802882.12 | 488453.91 | 9 | 10 | 9 | 12 | 9 | 12 | 1.400926185 | 1.240281211 | 1.009949814 | 0.894138315 |
| Q8K301 | Ddx52 | Probable ATP-dependent RNA helicase DDX52 | 101.82 | 5.85 | 67474 | 2 | 395793 | 303490.03 | 243429.06 | 249093.3 | 303660.53 | 292921.59 | 440068.25 | 237330.59 | 247383.03 | 229879.44 | 175156.91 | 359440.53 | 413560.12 | 328853.62 | 421647.88 | 348467.28 | 303592.84 | 230061.81 | 282308.47 | 256078.62 | 7 | 11 | 9 | 10 | 9 | 11 | 1.135872799 | 1.173445099 | 0.949877226 | 0.981297182 |
| Q8K4L0 | Ddx54 | ATP-dependent RNA helicase DDX54 | 250.51 | 6.06 | 97748 | 5 | 1613567 | 1393491.25 | 987139.25 | 3414042.25 | 880233.12 | 1630369.12 | 4042067.5 | 4119862.25 | 955314.62 | 1066131.75 | 3144290 | 1655283.12 | 1769605.25 | 1490215.38 | 1637532.38 | 3149898.75 | 1693384.62 | 1039228.38 | 1109484.62 | 1110224.5 | 6 | 11 | 11 | 11 | 11 | 11 | 1.169929163 | 0.820817271 | 0.977528792 | 0.685829998 |
| Q9D0R4 | Ddx56 | Probable ATP-dependent RNA helicase DDX56 | 205.22 | 9.52 | 61212 | 4 | 1250722.88 | 1271492.12 | 1179409.88 | 2037641.12 | 1315141.38 | 635658.44 | 1485957.38 | 1181388 | 1514121.5 | 494334.31 | 2984934.25 | 1895738.25 | 2850929.5 | 2076176.5 | 2104065 | 1557556.5 | 894174.12 | 1319310.88 | 1196419.38 | 655754.88 | 1 | 9 | 0 | 10 | 1 | 10 | 1.688567566 | 2.242668556 | 0.797120928 | 1.058695001 |
| Q6Q899 | Ddx58 | Antiviral innate immune response receptor RIG-I | 240.08 | 6.8 | 105975 | 5 | 3704273.25 | 3759491 | 3360024.5 | 2085483 | 4515250 | 2445025.5 | 3720996.75 | 5571987.5 | 4484877 | 3936143.5 | 648171.25 | 2362976.75 | 2345742.25 | 539298.12 | 1750258.75 | 2240114.5 | 697283.5 | 1960737.25 | 1394739.5 | 2075040.75 | 2 | 1 | 0 | 0 | 2 | 1 | 0.438832539 | 0.379306297 | 0.480237887 | 0.415095141 |
| P54823 | Ddx6 | Probable ATP-dependent RNA helicase DDX6 | 795.46 | 38.72 | 54192 | 13 | 15458514 | 11830848 | 12595928 | 13948778 | 11489484 | 14909339 | 13251974 | 9563173 | 10549970 | 11008778 | 16142702 | 14185983 | 13297627 | 12164953 | 11894342 | 12513013 | 12820584 | 11782918 | 11512790 | 12479474 | 9 | 9 | 6 | 10 | 9 | 10 | 1.036159317 | 1.141732703 | 0.935478509 | 1.030793614 |
| Q9CQ62 | Decr1 | 2,4-dienoyl-CoA reductase, mitochondrial | 180.86 | 13.43 | 36214 | 4 | 3310948.5 | 3218076.75 | 3981287.75 | 6243086 | 6509166.5 | 2886532.5 | 3101690 | 5641976 | 5950616 | 5512158.5 | 2346062.75 | 2965518.25 | 4195280 | 4584277 | 3976318.25 | 3784221 | 4562648 | 5453762.5 | 5433841 | 5517949 | 8 | 11 | 7 | 11 | 8 | 11 | 0.776675137 | 0.782378962 | 1.064045215 | 1.071859457 |
| Q9WV68 | Decr2 | Peroxisomal 2,4-dienoyl-CoA reductase | 123.81 | 11.64 | 31300 | 2 | 647184.31 | 535967.62 | 555637.94 | 2 | 167560.22 | 766358.75 | 907763 | 570702.75 | 270793.12 | 2 | 780804.06 | 679314.5 | 863221.69 | 693365.81 | 708773 | 886346.38 | 664286.69 | 503452.53 | 649267.38 | 2 | 0 | 7.5 | 8 | 12.5 | 8 | 12.5 | 1.95424501 | 1.480938943 | 1.418077486 | 1.07462788 |
| O09005 | Degs1 | Sphingolipid delta(4)-desaturase DES1 | 145.14 | 10.53 | 38241 | 2 | 2 | 2 | 164094.52 | 334792.81 | 2 | 2 | 2 | 2 | 2 | 2 | 633214.75 | 422370.75 | 1008304.19 | 1006414.75 | 842824.5 | 2 | 2 | 457219.59 | 264788.66 | 358493.06 | 0 | 8 | 0 | 5 | 0 | 8 | 7.843618475 | 391312.894 | 2.165804281 | 108050.531 |
| Q6AXS3 | Dek | Protein DEK | 445.44 | 20.37 | 42892 | 7 | 20493654 | 19553952 | 22402142 | 19361510 | 21256832 | 15014022 | 19694926 | 18151994 | 18670892 | 15232538 | 8541964 | 10528663 | 7752561 | 6916722 | 7525699.5 | 20402430 | 11690834 | 12359950 | 10419368 | 14940667 | 0 | 2 | 0 | 5 | 0 | 5 | 0.400372312 | 0.475605465 | 0.677350759 | 0.804630373 |
| A6H8H2 | Dennd4c | DENN domain-containing protein 4C | 179.86 | 2.31 | 211458 | 4 | 1806835.75 | 1408024.75 | 1728051.38 | 1760529.25 | 1958200.5 | 1782875.88 | 1582826.62 | 1751823.5 | 2685984.75 | 2315327.25 | 2 | 390661 | 819668 | 2 | 804231.69 | 1718979.25 | 1521341.25 | 1183484.5 | 672851.38 | 1322616.5 | 0 | 2 | 0 | 1 | 0 | 2 | 0.232584627 | 0.199090517 | 0.74111504 | 0.634388344 |
| Q9CQJ6 | Denr | Density-regulated protein | 140.13 | 13.13 | 22166 | 2 | 3091532.5 | 3653996.25 | 4545085.5 | 3561991.75 | 4084616.5 | 2365028.5 | 3543943.5 | 3723218 | 3941202.5 | 3293574.5 | 1662105.25 | 2201602.5 | 1880427.38 | 1644686.75 | 1970223.25 | 4741275.5 | 3523439.75 | 2841897.75 | 2781883.25 | 3306993.75 | 0 | 7 | 0 | 11 | 0 | 11 | 0.494214246 | 0.554874218 | 0.908025979 | 1.019477301 |
| Q570Y9 | Deptor | DEP domain-containing mTOR-interacting protein | 54.82 | 2.69 | 46120 | 1 | 492410.44 | 363792.09 | 476437.94 | 601637.69 | 548711.69 | 2 | 2 | 2 | 512737.88 | 503531.38 | 2 | 2 | 235795.75 | 2 | 2 | 845064.06 | 2 | 477692.56 | 2 | 260996.52 | 0 | 7 | 9 | 11 | 9 | 11 | 0.094967666 | 0.232027443 | 0.637842777 | 1.558393875 |
| Q91YP3 | Dera | Deoxyribose-phosphate aldolase | 550.95 | 38.68 | 34975 | 8 | 7695460 | 8233301 | 11354594 | 10450725 | 10117326 | 7929975 | 6764418 | 9045075 | 7238691 | 7363217 | 4697197 | 6102087 | 7447379 | 5639032 | 5996294 | 6954871.5 | 5682194 | 8208457 | 7404533 | 7672693 | 0 | 1 | 3 | 11 | 3 | 11 | 0.624474629 | 0.77936663 | 0.750714587 | 0.936918605 |
| Q99J56 | Derl1 | Derlin-1 | 165.68 | 13.15 | 28835 | 3 | 180752.86 | 2 | 163375.83 | 2 | 39241.09 | 147228.64 | 253868.34 | 330686.69 | 418369.12 | 133970.81 | 1741929.88 | 1817824.25 | 2741966.25 | 1940256.5 | 2248538.5 | 183937.23 | 2 | 375452.97 | 160496.42 | 496785 | 0 | 6 | 0 | 12 | 0 | 12 | 27.36367464 | 8.169396918 | 3.173596327 | 0.947473919 |
| Q9R1C6 | Dgke | Diacylglycerol kinase epsilon | 44.85 | 2.3 | 63635 | 1 | 2 | 229016.44 | 227757.67 | 2 | 2 | 320900.84 | 177000.44 | 2 | 207563.86 | 261207.69 | 237542.75 | 2 | 2 | 2 | 221488.5 | 2 | 2 | 2 | 2 | 2 | 12.5 | 7.5 | 7.5 | 2.5 | 12.5 | 7.5 | 1.004941415 | 0.474862111 | 2.189237E-05 | 1.034474E-05 |
| O08560 | Dgkz | Diacylglycerol kinase zeta | 41.9 | 1.72 | 104009 | 1 | 2 | 2 | 2 | 2 | 2 | 2 | 2 | 2 | 2 | 2 | 264357.81 | 203947.2 | 2 | 2 | 118804.73 | 2 | 2 | 2 | 2 | 2 | 5 | 12.5 | 5 | 12.5 | 5 | 12.5 | 58711.374 | 58711.374 | 1 | 1 |
| Q9QX60 | Dguok | Deoxyguanosine kinase, mitochondrial | 40.25 | 3.25 | 32281 | 1 | 1351655.12 | 1203217.38 | 1570463 | 2041801.5 | 1413646.62 | 2039743.62 | 992293.5 | 1459703.38 | 1479442.75 | 1848014.25 | 1565653.38 | 899801.06 | 1249079.75 | 1300212.62 | 1426885.5 | 1616257.62 | 3064178.25 | 1783709.75 | 1937461.75 | 1661283.25 | 8 | 4 | 6 | 7 | 8 | 7 | 0.849731721 | 0.823822689 | 1.327420901 | 1.286946726 |
| O88455 | Dhcr7 | 7-dehydrocholesterol reductase | 250.55 | 13.38 | 53919 | 5 | 8420224 | 8326428.5 | 6577549 | 9156462 | 7734344 | 7811316.5 | 7582275.5 | 9103084 | 8379915 | 6764625 | 12156979 | 10105335 | 16013407 | 12918322 | 16240539 | 12110634 | 11161273 | 11089765 | 9033594 | 12011790 | 0 | 1 | 0 | 1 | 0 | 1 | 1.676851161 | 1.701122942 | 1.377770625 | 1.397713329 |
| Q9DBB8 | Dhdh | Trans-1,2-dihydrobenzene-1,2-diol dehydrogenase | 115.3 | 10.21 | 36301 | 2 | 3280341.5 | 3367484.75 | 2658273 | 2130175.5 | 2155294.75 | 2473015.5 | 3352970 | 2550494.5 | 1653441.62 | 2335235 | 904253.62 | 2037383.5 | 1237699.25 | 468387.84 | 1401610.88 | 2927108 | 1230199.12 | 2163536 | 1689074.38 | 2085685.62 | 0 | 5 | 1 | 7 | 1 | 7 | 0.445079951 | 0.4892243 | 0.742784203 | 0.816455742 |
| P00375 | Dhfr | Dihydrofolate reductase | 188.71 | 16.04 | 21606 | 3 | 4078497.25 | 3703753.5 | 3185696.75 | 2970118 | 3949301 | 2994072.75 | 3258373 | 2972876.75 | 3482712 | 2778173 | 2069364 | 3449049.75 | 2873216 | 2 | 3145160.5 | 2475836 | 1860836 | 1825632.62 | 2116349.5 | 2612738.5 | 3 | 0 | 8 | 0 | 8 | 0 | 0.644968741 | 0.744972082 | 0.60888743 | 0.703296312 |
| O35435 | Dhodh | Dihydroorotate dehydrogenase (quinone), mitochondrial | 66.55 | 2.28 | 42700 | 1 | 512401.12 | 338291.34 | 297141.38 | 393161.56 | 2 | 491724.28 | 268650.53 | 270610.06 | 301282.22 | 2 | 544420.31 | 222727.06 | 573059.81 | 395086.97 | 264894.28 | 373911.25 | 437308.25 | 443597.81 | 378877.28 | 495525 | 9 | 7 | 9 | 4 | 9 | 7 | 1.297983001 | 1.501339666 | 1.381715238 | 1.59819034 |
| Q99L04 | Dhrs1 | Dehydrogenase/reductase SDR family member 1 | 378.99 | 22.04 | 34005 | 6 | 8729525 | 9051382 | 7550474 | 8024687 | 9113276 | 9452120 | 9545061 | 9067570 | 9805063 | 9368334 | 9596500 | 9423150 | 9067792 | 7126885 | 7815306.5 | 14359098 | 8790312 | 17094356 | 10969040 | 19425144 | 10 | 2 | 7 | 5 | 10 | 5 | 1.013192799 | 0.910908563 | 1.663269157 | 1.495358158 |
| Q5SS80 | Dhrs13 | Dehydrogenase/reductase SDR family member 13 | 53.92 | 2.13 | 40745 | 1 | 421870.28 | 620751.69 | 506313.41 | 741946.62 | 571140.38 | 430340.94 | 404618.88 | 459120.53 | 366782.34 | 529079.12 | 645038.19 | 678320.31 | 801268 | 651572 | 602418.62 | 510321.84 | 749747.62 | 781703.25 | 563030 | 544463.56 | 5 | 9 | 0 | 1 | 5 | 9 | 1.180499895 | 1.542788536 | 1.100363957 | 1.438059338 |
| Q99LB2 | Dhrs4 | Dehydrogenase/reductase SDR family member 4 | 317.79 | 21.86 | 29885 | 5 | 5420507 | 5516992 | 4335593.5 | 5652825.5 | 3679009.5 | 6274157.5 | 4908779 | 3334603.5 | 4218530 | 4869705 | 6880136 | 5470969 | 6126124 | 6592027 | 6812585 | 6238955 | 6500336 | 3780289.75 | 5564403.5 | 4349295.5 | 2 | 8 | 2 | 9 | 2 | 9 | 1.295750252 | 1.350594971 | 1.074308378 | 1.119780213 |
| Q9CXR1 | Dhrs7 | Dehydrogenase/reductase SDR family member 7 | 195.4 | 9.47 | 38167 | 3 | 191769.73 | 344904.69 | 2 | 291593.91 | 839271.38 | 2 | 478300.94 | 482029.91 | 1246240.25 | 2 | 1226951.62 | 1783041.88 | 1788327.62 | 1838871.62 | 1456250 | 2 | 160828.67 | 2 | 42885.94 | 2 | 0 | 3.5 | 1 | 7 | 1 | 7 | 4.853517421 | 3.667875496 | 0.122168225 | 0.092324349 |
| Q99J47 | Dhrs7b | Dehydrogenase/reductase SDR family member 7B | 311.79 | 19.81 | 34987 | 5 | 2487142.25 | 2193360.5 | 2090305.75 | 2643748 | 1884103.62 | 2774615 | 2580111.5 | 1696947.75 | 2079898.75 | 2242783.75 | 3696339.75 | 3548176.5 | 3308227.25 | 3226782 | 3673769 | 2843676.75 | 2885475 | 2441491.5 | 3737674.25 | 2831290.25 | 0 | 2 | 0 | 2 | 0 | 2 | 1.544722499 | 1.534442332 | 1.304544751 | 1.295862973 |
| O35286 | Dhx15 | Pre-mRNA-splicing factor ATP-dependent RNA helicase DHX15 | 1768.96 | 36.86 | 91007 | 28 | 47725702 | 47857806 | 49693378 | 48751798 | 47388102 | 49460418 | 49558526 | 44943334 | 49148714 | 45145742 | 41762806 | 43193062 | 44577294 | 38775022 | 43388310 | 51012074 | 47104654 | 44544570 | 46021050 | 46747242 | 0 | 5 | 0 | 11 | 0 | 11 | 0.876892189 | 0.888522605 | 0.975199753 | 0.988134044 |
| Q6PGC1 | Dhx29 | ATP-dependent RNA helicase DHX29 | 99 | 2.05 | 153975 | 2 | 245253.23 | 104278.86 | 170684.53 | 147342.22 | 590792.31 | 154385.33 | 305893.81 | 434780 | 570886.81 | 157178.97 | 418520.88 | 2 | 2 | 226133.97 | 2 | 276456.16 | 58827.45 | 175282.98 | 135247.95 | 200286.05 | 7 | 11 | 5 | 6 | 7 | 11 | 0.512306005 | 0.397172665 | 0.6723883 | 0.521278787 |
| Q5BJS0 | Dhx30 | ATP-dependent RNA helicase DHX30 | 270.98 | 5.11 | 133997 | 5 | 1709324.5 | 1738087.12 | 2156932.75 | 1943652.62 | 2504464.5 | 1256569.38 | 1763641.75 | 1894142 | 2042912.5 | 1420118.75 | 1600345.5 | 1843060.75 | 1368009.5 | 1561879.38 | 1963736 | 1811653.5 | 679873.62 | 1778676.12 | 1626048.25 | 1577148.75 | 5 | 4 | 12 | 10 | 12 | 10 | 0.829352208 | 0.995183073 | 0.743439828 | 0.892092317 |
| D4A2Z8 | Dhx36 | ATP-dependent DNA/RNA helicase DHX36 | 124.35 | 2.4 | 113843 | 2 | 1371294.25 | 1584176 | 1777847.25 | 446673.16 | 704586 | 476817.06 | 1379184 | 1297904.38 | 580946.06 | 467536.44 | 1392862.38 | 1457107.88 | 427435.12 | 581604.62 | 509048.41 | 2192334.5 | 2095580.75 | 498690.38 | 1167894.38 | 894317.06 | 8 | 10 | 10 | 7 | 10 | 10 | 0.74228932 | 1.039422936 | 1.163858926 | 1.629744128 |
| O70133 | Dhx9 | ATP-dependent RNA helicase A | 2345.91 | 30.29 | 149475 | 39 | 67166050 | 70344378 | 68870618 | 72953946 | 66605398 | 75960938 | 68427018 | 63433690 | 63722650 | 61127874 | 68259330 | 57221318 | 61530354 | 58492314 | 55829758 | 64175634 | 69982858 | 65510050 | 63828854 | 62036930 | 2 | 3 | 4 | 11 | 4 | 11 | 0.871054906 | 0.905795859 | 0.941012774 | 0.97854391 |
| O08808 | Diaph1 | Protein diaphanous homolog 1 | 163.05 | 2.79 | 139343 | 3 | 603431.12 | 345780.53 | 617017.75 | 514112.97 | 547111.19 | 1052081.25 | 633953.81 | 321882.81 | 380761.25 | 648231.44 | 334722.5 | 576252.06 | 444008.69 | 156573.58 | 354733.69 | 934463 | 1154651.5 | 546992.44 | 275116.72 | 223130.7 | 5 | 12 | 6 | 11 | 6 | 12 | 0.710303904 | 0.614535885 | 1.192924742 | 1.03208649 |
| Q8C9B9 | Dido1 | Death-inducer obliterator 1 | 52.52 | 0.58 | 247176 | 1 | 459719.41 | 415242.25 | 612249.12 | 566930.69 | 456856.34 | 804627.81 | 514245.09 | 504927.03 | 573178.88 | 702817.56 | 311777.41 | 2 | 2 | 2 | 350266.28 | 497476.62 | 2 | 438412.22 | 636770.19 | 628040.44 | 0 | 11 | 0 | 6 | 0 | 11 | 0.263660003 | 0.213578445 | 0.876425085 | 0.709950335 |
| Q5M9G7 | Diexf | Digestive organ expansion factor homolog | 105.47 | 2.1 | 87825 | 2 | 840952.25 | 2 | 2 | 417896.5 | 693681.88 | 537862.38 | 503274.59 | 935106.62 | 2 | 659135.19 | 887532.25 | 814091.69 | 764729.06 | 828612.69 | 879633.38 | 482166.44 | 758431.06 | 756458.12 | 2 | 491658.06 | 3 | 10 | 5 | 10.5 | 5 | 10.5 | 2.138040988 | 1.584059162 | 1.274607703 | 0.944347663 |
| Q9D0D4 | Dimt1 | Probable dimethyladenosine transferase | 73.1 | 10.22 | 35274 | 2 | 2 | 2 | 2 | 2 | 247011.91 | 2 | 2 | 2 | 236158.92 | 2 | 430642.16 | 587505.25 | 345660.59 | 288312.94 | 479209.59 | 2 | 2 | 2 | 2 | 2 | 0 | 10 | 0 | 10 | 0 | 10 | 8.628173049 | 9.02467852 | 4.048257E-05 | 4.234293E-05 |
| Q3UH60 | Dip2b | Disco-interacting protein 2 homolog B | 357.83 | 7.12 | 171126 | 8 | 1978190.88 | 2328841.5 | 4505429.5 | 3013527 | 3253059.5 | 3125891 | 2276144 | 2464276.5 | 3319735.5 | 4351012 | 2998841.25 | 2924192 | 3857310.5 | 5794324 | 3082424.5 | 2345700 | 2478313.75 | 3434190 | 9682942 | 3849658 | 9 | 8 | 10 | 9 | 10 | 9 | 1.237285788 | 1.200812345 | 1.445104704 | 1.402505053 |
| Q3USZ8 | Dipk2a | Deleted in autism protein 1 homolog | 116.92 | 5.35 | 49463 | 2 | 2 | 2 | 397896.81 | 226714.94 | 370995.53 | 2 | 2 | 265037.03 | 510568.91 | 597538.75 | 667352 | 502886.34 | 1327066.25 | 973958.88 | 910389.31 | 2 | 2 | 2 | 142025.61 | 271269.19 | 0 | 8 | 2 | 8 | 2 | 8 | 4.400967394 | 3.190952889 | 0.415122657 | 0.300987652 |
| Q9CSH3 | Dis3 | Exosome complex exonuclease RRP44 | 165.9 | 4.18 | 108838 | 3 | 1716495.38 | 1469116.38 | 1882777.88 | 1637750.5 | 1631217.5 | 1181304.62 | 2075360.75 | 1398445.75 | 1895130.38 | 2229666 | 886289.69 | 1403831.12 | 1003215.44 | 338416.78 | 1241249.25 | 1982116.12 | 855803.88 | 1219648.12 | 1198312.75 | 1229353.25 | 0 | 5 | 3 | 6 | 3 | 6 | 0.584478019 | 0.555017496 | 0.777852456 | 0.738644925 |

| | | | | | | | | | | | | | | | | | | | | | | | | | | | | | | | | | | | | |
|---|---|---|---|---|---|---|---|---|---|---|---|---|---|---|---|---|---|---|---|---|---|---|---|---|---|---|---|---|---|---|---|---|---|---|---|---|
| Q9ESX5 | Dkc1 | H/ACA ribonucleoprotein complex subunit DKC1 | 491.3 | 23.18 | 57402 | 8 | 6855922 | 7131872 | 5296310 | 6087363.5 | 6350568.5 | 6325512 | 6275570 | 5156337.5 | 6520654 | 6617723.5 | 11448150 | 10178244 | 9067182 | 7312616.5 | 8735823 | 5460428 | 3884198.75 | 6075846.5 | 4183797.25 | 5407961 | 0 | 3 | 0 | 3 | 0 | 3 | 1.473487247 | 1.512892368 | 0.788481278 | 0.809567447 |
| Q8BMF4 | Dlat | Dihydrolipoyllysine-residue acetyltransferase component of pyruvate dehydrogenase complex, mitochondrial | 775.5 | 26.32 | 67942 | 13 | 28863490 | 31774864 | 31888394 | 34470678 | 37913546 | 31598538 | 30933024 | 31700982 | 32499396 | 37391942 | 47253438 | 47557834 | 54315390 | 50696094 | 54020226 | 29725828 | 29937930 | 40128374 | 34624230 | 38293722 | 0 | 9 | 0 | 11 | 0 | 11 | 1.539272851 | 1.546654752 | 1.047292863 | 1.052315372 |
| O08749 | Dld | Dihydrolipoyl dehydrogenase, mitochondrial | 1330.55 | 42.63 | 54272 | 19 | 70020714 | 76878370 | 57145674 | 71355106 | 67879138 | 81577594 | 74153938 | 68227010 | 67463050 | 68787338 | 78200698 | 77967290 | 90110986 | 87822738 | 89216170 | 87098290 | 83602866 | 73668490 | 81040898 | 79327906 | 0 | 1 | 2 | 4 | 2 | 4 | 1.233159848 | 1.175200965 | 1.179036433 | 1.123621366 |
| B1AZP2 | Dlgap4 | Disks large-associated protein 4 | 81.14 | 1.51 | 108037 | 1 | 692647.69 | 1049372.25 | 752433.12 | 1323032.12 | 689134.94 | 2 | 763384.19 | 829183.06 | 937643.5 | 771631.12 | 2 | 2 | 560638.31 | 2 | 2 | 655219.94 | 2 | 527352.62 | 762498.69 | 816624.12 | 0 | 6 | 3 | 6.5 | 3 | 6.5 | 0.124405052 | 0.169797947 | 0.612809 | 0.836410648 |
| Q01205 | Dlst | Dihydrolipoyllysine-residue succinyltransferase component of 2-oxoglutarate dehydrogenase complex, mitochondrial | 868.78 | 32.16 | 48925 | 14 | 62412966 | 52688806 | 41609106 | 47722082 | 44143446 | 60143174 | 61715730 | 50255858 | 50464742 | 56512398 | 79403370 | 76885458 | 78721682 | 75284986 | 85600338 | 57553314 | 69854426 | 56178822 | 51460630 | 50691002 | 0 | 6 | 0 | 11 | 0 | 11 | 1.592652498 | 1.418514228 | 1.149498452 | 1.023813991 |
| P63036 | Dnaja1 | DnaJ homolog subfamily A member 1 | 252.24 | 22.17 | 44868 | 5 | 1492424 | 2243936.5 | 1384542.12 | 485738.62 | 1530106.5 | 504990.88 | 2040006.5 | 229336.05 | 488154.31 | 550806.12 | 761842.19 | 3567429.5 | 3468442 | 773328.06 | 2890106.25 | 2035686.75 | 256273.92 | 904722.75 | 784140 | 892256.62 | 8 | 7 | 2 | 8 | 8 | 8 | 1.605934302 | 3.005576916 | 0.682815229 | 1.277918833 |
| Q9QYJ0 | Dnaja2 | DnaJ homolog subfamily A member 2 | 603.23 | 43.93 | 45746 | 12 | 3131704.25 | 4848873.5 | 4247618 | 4079323.75 | 3602772.5 | 6211460.5 | 5959330.5 | 3644132.75 | 5210506.5 | 6984792.5 | 16248890 | 15479597 | 14773827 | 16830914 | 15631852 | 4844743 | 4387009 | 3594779.5 | 4470630 | 6007625 | 0 | 7 | 0 | 6 | 0 | 7 | 3.966043291 | 2.819152161 | 1.170489438 | 0.832010038 |
| Q99M87 | Dnaja3 | DnaJ homolog subfamily A member 3, mitochondrial | 110.59 | 4.79 | 52443 | 2 | 2087120.75 | 1925290.25 | 1596821.62 | 1989978 | 1930132.12 | 1742798.5 | 2 | 1913947.38 | 1567977.62 | 2087166.25 | 1713207 | 849665.12 | 2403400.5 | 2266914.25 | 1527414.5 | 1018552.12 | 2 | 2562263.75 | 2416327.75 | 1770414.25 | 11 | 11 | 11 | 10.5 | 11 | 11 | 0.919329025 | 1.198130616 | 0.815120212 | 1.062318773 |
| Q6TUG0 | Dnajb11 | DnaJ homolog subfamily B member 11 | 624.15 | 24.86 | 40495 | 9 | 8929003 | 10125287 | 11404280 | 10382389 | 9216261 | 11871480 | 8138703.5 | 11243629 | 11873576 | 14385044 | 10728126 | 13950039 | 16038953 | 16186789 | 17007066 | 9613229 | 6080915.5 | 11457098 | 8605564 | 8990704 | 1 | 8 | 5 | 5 | 5 | 8 | 1.476529719 | 1.285130359 | 0.8939272 | 0.778049346 |
| Q9QYI4 | Dnajb12 | DnaJ homolog subfamily B member 12 | 64.35 | 2.66 | 41988 | 1 | 82303.97 | 2 | 2 | 2 | 2 | 2 | 2 | 2 | 2 | 2 | 126380.53 | 2 | 2 | 95576.25 | 2 | 2 | 2 | 63773.71 | 2 | 2 | 9 | 12 | 7.5 | 10 | 9 | 12 | 2.696603908 | 22196.278 | 0.7748777 | 6378.171 |
| O54946 | Dnajb6 | DnaJ homolog subfamily B member 6 | 111.63 | 6.3 | 39807 | 2 | 854526.88 | 1619862.12 | 1009167.94 | 904656 | 2 | 1111342.38 | 2 | 532287.25 | 377637.53 | 2 | 1625823 | 1024861.25 | 1247290.5 | 298530.94 | 2 | 1235191.75 | 2 | 1179556.75 | 2 | 1515633.5 | 10.5 | 12 | 8 | 8 | 10.5 | 12 | 0.956313159 | 2.076172546 | 0.895668524 | 1.944511987 |
| Q498R3 | Dnajc10 | DnaJ homolog subfamily C member 10 | 228.02 | 8.32 | 90747 | 5 | 1369493 | 1847762.12 | 786174.94 | 1526362.25 | 545242 | 1860527.38 | 1640256 | 1677857.25 | 952675.31 | 2155436.5 | 2913996.5 | 1802018.25 | 3108334.25 | 2899967 | 2655206 | 1248649.5 | 1623875.75 | 1320857.38 | 991799.81 | 1267014.5 | 1 | 12 | 2 | 5 | 2 | 12 | 2.202378014 | 1.6145676 | 1.062084033 | 0.778615868 |
| Q5U458 | Dnajc11 | DnaJ homolog subfamily C member 11 | 84.52 | 4.65 | 63233 | 2 | 1037263.44 | 1017516.69 | 986345.06 | 1208397.75 | 619565.38 | 1524811.62 | 871355 | 1082608 | 1067692.62 | 1268481.62 | 2473401.5 | 1916764.5 | 2535218 | 2474392.25 | 2228923.25 | 552010.5 | 1039655.12 | 1202565.5 | 1566417 | 1163570.88 | 0 | 8 | 0 | 12 | 0 | 12 | 2.388270398 | 1.999793941 | 1.134548942 | 0.950003024 |
| P54103 | Dnajc2 | DnaJ homolog subfamily C member 2 | 109.32 | 5.64 | 71722 | 2 | 512834.91 | 709481.94 | 511782.47 | 645590.06 | 501571.41 | 1016858.25 | 504980.56 | 470623.25 | 591340.69 | 795982.31 | 999214.5 | 456252.91 | 692980.12 | 1046743.75 | 783978.62 | 498310.78 | 765687.25 | 469273.66 | 2 | 457051.22 | 6 | 5 | 10 | 4 | 10 | 5 | 1.381051626 | 1.177344071 | 0.760196688 | 0.648066333 |
| Q91YW3 | Dnajc3 | DnaJ homolog subfamily C member 3 | 162.41 | 7.14 | 57464 | 3 | 1159718.25 | 1440991.88 | 456577.12 | 956604.06 | 1433667.25 | 1612001.25 | 1509874.12 | 427333.91 | 1260117.25 | 483283.34 | 1206011.5 | 2174254.75 | 563073.44 | 942405.75 | 782138.44 | 2144355.5 | 1669368 | 1071603.25 | 909518.75 | 979985.5 | 11 | 10 | 12 | 9 | 12 | 10 | 1.040444782 | 1.070905285 | 1.243645373 | 1.280054863 |
| Q9QYI3 | Dnajc7 | DnaJ homolog subfamily C member 7 | 542.2 | 19.23 | 56476 | 9 | 7682782 | 9339744 | 8049637.5 | 8068625.5 | 10628348 | 8038142 | 9670347 | 10485377 | 10026463 | 6609929.5 | 6091954.5 | 9289447 | 7513935.5 | 5665913.5 | 7294732 | 9298471 | 7468887 | 5950690 | 6743877.5 | 6481601 | 3 | 3 | 4 | 4 | 4 | 4 | 0.819206979 | 0.799816546 | 0.82120711 | 0.801769334 |
| Q642C0 | Dnajc8 | DnaJ homolog subfamily C member 8 | 104.9 | 9.09 | 29813 | 3 | 4354400.5 | 5994667 | 5928528 | 4735088 | 4878608.5 | 1629393.38 | 5512871.5 | 5041466.5 | 5530726 | 3367557 | 1519193 | 3160000.75 | 845380.12 | 1230680.5 | 2384441.5 | 3804513.5 | 1538477.75 | 2723140.5 | 2289484 | 4020354.75 | 0 | 0 | 2 | 6 | 2 | 6 | 0.353002696 | 0.433530483 | 0.555243458 | 0.681906873 |
| P56542 | Dnase2 | Deoxyribonuclease-2-alpha | 178.03 | 12.18 | 38811 | 4 | 3046905 | 2833234.5 | 5074117.5 | 4271193 | 6219831.5 | 1999076.38 | 2989927.75 | 4601271.5 | 3845534 | 3550079 | 450472.59 | 1073032.38 | 1237456.62 | 866165.62 | 1565803.75 | 3342988.25 | 1418778.38 | 2989654.5 | 1852334.5 | 3520259.5 | 0 | 5 | 0 | 5 | 0 | 5 | 0.24214795 | 0.3057203 | 0.611976818 | 0.772642245 |
| O35303 | Dnm1l | Dynamin-1-like protein | 433.98 | 13.38 | 83908 | 7 | 5164699 | 5705837 | 4926384.5 | 5085202 | 6232730 | 4946726 | 5063236.5 | 9808042 | 7948044 | 2823191.5 | 5493975.5 | 6593368 | 6364929 | 6575300 | 6218024.5 | 4908376 | 3370630.75 | 5099507.5 | 5296486.5 | 6003005 | 3 | 9 | 10 | 11 | 10 | 11 | 1.152342503 | 1.02145712 | 0.910128711 | 0.806754458 |
| P39052 | Dnm2 | Dynamin-2 | 859.67 | 19.89 | 98230 | 16 | 10058868 | 8047391 | 7895961 | 9169608 | 8998679 | 8696856 | 9821837 | 7197201.5 | 7433360.5 | 7264446.5 | 10703142 | 8910760 | 12132471 | 9803323 | 8816499 | 7396770.5 | 8614830 | 9388194 | 9210801 | 8395705 | 7 | 12 | 3 | 9 | 7 | 12 | 1.140267532 | 1.246265329 | 0.973642899 | 1.064151486 |
| P13864 | Dnmt1 | DNA (cytosine-5)-methyltransferase 1 | 1238.12 | 16.11 | 183189 | 22 | 4077360.5 | 3838882.25 | 6004068.5 | 4103041 | 5683431 | 3407801.5 | 4018142.5 | 3528618.5 | 4995618.5 | 2698949.5 | 1810264.5 | 2415443.25 | 3821919 | 7103722 | 8036811.5 | 4016822.25 | 2603217 | 2892519.75 | 3093786.5 | 2681544.5 | 10 | 1 | 12 | 5 | 12 | 5 | 0.978123434 | 1.243390959 | 0.644874078 | 0.819764224 |
| O88508 | Dnmt3a | DNA (cytosine-5)-methyltransferase 3A | 346.73 | 8.37 | 101672 | 6 | 2578082 | 2728381.5 | 3201360.5 | 2745604 | 2859341.75 | 2098206.75 | 2583196.25 | 3262621.75 | 2573124.5 | 2300572.25 | 913542.56 | 944162.5 | 328877.19 | 965993.62 | 561245.38 | 1596132 | 2574168 | 2037701.25 | 1854950.25 | 2259786.75 | 0 | 0 | 0 | 4 | 0 | 4 | 0.263153252 | 0.289741141 | 0.731446657 | 0.805348926 |
| Q9Z2W0 | Dnpep | Aspartyl aminopeptidase | 545.26 | 30.66 | 52207 | 10 | 14633964 | 15319277 | 12877703 | 16423051 | 16297750 | 13351956 | 16245348 | 16680640 | 16991562 | 16896084 | 13226043 | 13924374 | 14662530 | 11260735 | 13669386 | 16586396 | 11338352 | 14971477 | 15618060 | 19578696 | 5 | 10 | 3 | 9 | 5 | 10 | 0.883408689 | 0.832565044 | 1.033635702 | 0.974145902 |
| Q80VJ3 | Dnph1 | 2'-deoxynucleoside 5'-phosphate N-hydrolase 1 | 51.09 | 4.62 | 18977 | 1 | 723878.5 | 628221.81 | 722847.56 | 677860.31 | 537921.38 | 753576.56 | 830447.19 | 746383.25 | 651398.81 | 637537.94 | 282006.5 | 253685.58 | 2 | 288783.31 | 249066.61 | 861323.12 | 1073947.62 | 815715.06 | 751323.69 | 735453.56 | 0 | 0 | 0 | 6 | 0 | 6 | 0.326232825 | 0.296612887 | 1.287788307 | 1.170865036 |
| Q8R2M2 | Dnttip2 | Deoxynucleotidyltransferase terminal-interacting protein 2 | 98.2 | 2.51 | 84277 | 2 | 694752.19 | 714141.38 | 2 | 808786.12 | 2 | 1707187 | 2 | 606196.56 | 383914.88 | 2 | 1169451 | 790328.88 | 358334.16 | 1052718.38 | 1427020.12 | 682234.38 | 968758.5 | 600870.44 | 1252934.38 | 2 | 4 | 10 | 7 | 9 | 7 | 10 | 2.163452147 | 1.778759574 | 1.580387553 | 1.299372161 |
| Q8BZN6 | Dock10 | Dedicator of cytokinesis protein 10 | 475.01 | 5.77 | 245758 | 9 | 1157872.75 | 1906305.12 | 2603382.75 | 2618486.5 | 3318488 | 2380658.5 | 1660096.62 | 2611452 | 3155391.75 | 2591461.25 | 1544579 | 1519382.62 | 2009623.88 | 694019.88 | 1097736.38 | 725135.81 | 1486082.62 | 1737759 | 1270641.12 | 2070959.88 | 4 | 5 | 1 | 2 | 4 | 5 | 0.591608512 | 0.553698562 | 0.62825252 | 0.587994441 |
| Q8C3J5 | Dock2 | Dedicator of cytokinesis protein 2 | 1342.2 | 17.89 | 211704 | 29 | 13044485 | 10426091 | 17956062 | 19724108 | 15553214 | 13276222 | 12106737 | 16885980 | 19843904 | 16451477 | 12693188 | 15427944 | 19141400 | 13850710 | 14106893 | 9646796 | 11080492 | 13022794 | 14382991 | 13031838 | 11 | 5 | 11 | 4 | 11 | 5 | 0.980655171 | 0.957433794 | 0.797415296 | 0.77853294 |
| Q8C147 | Dock8 | Dedicator of cytokinesis protein 8 | 789.21 | 10.81 | 238978 | 16 | 7162572 | 8235190 | 8870338 | 8755393 | 7592066 | 7829460 | 7272329 | 7824176.5 | 8026211 | 8169520 | 4926037 | 5455893.5 | 5184860.5 | 5550080.5 | 4467253 | 8000438 | 4979872 | 6214086 | 7611319.5 | 7042958.5 | 0 | 4 | 0 | 4 | 0 | 4 | 0.629909452 | 0.653962552 | 0.833391804 | 0.865214881 |
| P97465 | Dok1 | Docking protein 1 | 50.24 | 1.87 | 52452 | 1 | 1015767.06 | 1602414.62 | 1086912.75 | 1245170.62 | 827958.5 | 1072158.25 | 811990.88 | 761280.38 | 823165 | 982563 | 750921 | 854288.12 | 772920.75 | 982282.12 | 749303.12 | 902518.44 | 1146289.38 | 1395996.88 | 1002867.94 | 932195 | 2 | 10 | 7 | 5 | 7 | 10 | 0.711241972 | 0.923291324 | 0.931059104 | 1.208644634 |
| O70469 | Dok2 | Docking protein 2 | 100.91 | 5.1 | 45522 | 2 | 509625.75 | 1221942.25 | 1386566.5 | 1205484.25 | 1171502.5 | 1287114 | 852439.62 | 1242184 | 1270431 | 1209209 | 1081236.88 | 900056.38 | 1011168.94 | 1150861.38 | 1178246 | 531306.69 | 1307341 | 1267613.75 | 1038053.81 | 1294963.5 | 6 | 11 | 5 | 11 | 6 | 11 | 0.968417135 | 0.907904238 | 0.989837804 | 0.927986406 |
| Q61103 | Dpf2 | Zinc finger protein ubi-d4 | 204.23 | 8.7 | 44230 | 3 | 2021057.62 | 2242443.25 | 1816908.12 | 2554518.75 | 2242532 | 2774882 | 1906940 | 2185902.5 | 1436040.62 | 1939905.25 | 655157 | 1038061.62 | 430762.44 | 1187797.25 | 1281222 | 1797644 | 2333036 | 2668887.5 | 2065799.62 | 2109442.25 | 0 | 12 | 0 | 10 | 0 | 12 | 0.42224935 | 0.448374474 | 1.008949666 | 1.07137471 |
| O70152 | Dpm1 | Dolichol-phosphate mannosyltransferase subunit 1 | 255.06 | 23.08 | 29175 | 4 | 1999628.25 | 211640.92 | 1527133.12 | 1569813.75 | 2789389 | 2682183 | 2383246.25 | 1961016.75 | 2116420.5 | 1911999 | 4913640 | 2999986 | 3349860.25 | 2796365.25 | 2384457.75 | 1853080.62 | 3279293.5 | 1983776.75 | 2296371 | 2368952.5 | 1 | 6 | 1 | 12 | 1 | 12 | 2.030762079 | 1.487517804 | 1.454933195 | 1.065727518 |
| Q99KK7 | Dpp3 | Dipeptidyl peptidase 3 | 694.81 | 31.17 | 82898 | 13 | 11403524 | 12171415 | 9164446 | 10831242 | 13670032 | 9960280 | 12681797 | 9754570 | 17428356 | 11084307 | 8568424 | 9042664 | 8663324 | 5768870 | 7595527 | 10351509 | 11849008 | 12034576 | 9078905 | 9729000 | 0 | 8 | 0 | 8 | 0 | 8 | 0.692493932 | 0.650784076 | 0.926666445 | 0.870852059 |
| Q9EPB1 | Dpp7 | Dipeptidyl peptidase 2 | 46.43 | 2 | 55114 | 1 | 1325564 | 1413193.75 | 1048353.75 | 814386.75 | 1181039.25 | 1232362 | 1008559.12 | 758895.88 | 1109484.62 | 789596.88 | 2 | 290522.53 | 275817.16 | 298784.75 | 254544.31 | 933929.88 | 1027867.69 | 952214.38 | 677195.62 | 843438.38 | 0 | 4 | 0 | 9 | 0 | 9 | 0.193629657 | 0.228555613 | 0.766903103 | 0.905233278 |
| Q8BVG4 | Dpp9 | Dipeptidyl peptidase 9 | 93.73 | 2.9 | 98001 | 2 | 944856.62 | 722211.81 | 656356.88 | 555557 | 1018953.38 | 634722.5 | 982295.81 | 803284.56 | 795669.5 | 224261.39 | 101126.98 | 554225.81 | 495323.19 | 106688.02 | 2 | 992497.38 | 857045.81 | 443120.25 | 715922.25 | 499796.56 | 0 | 9 | 2 | 12 | 2 | 12 | 0.322572279 | 0.365488536 | 0.900061604 | 1.019809261 |
| O08553 | Dpysl2 | Dihydropyrimidinase-related protein 2 | 1857.08 | 63.29 | 62278 | 26 | 183940514 | 197865314 | 199506082 | 213494050 | 191185666 | 204487522 | 185658178 | 186302738 | 170798274 | 179177954 | 84239362 | 93429706 | 90996938 | 100954730 | 98888978 | 181593186 | 212221810 | 182453154 | 187842802 | 189860258 | 0 | 6 | 0 | 8 | 0 | 8 | 0.475166017 | 0.505718091 | 0.967524657 | 1.029734251 |
| A0JPP1 | Drap1 | Dr1-associated corepressor | 151.34 | 11.22 | 22326 | 3 | 3970337.25 | 3065957.5 | 3521112.5 | 2940893.5 | 4329134.5 | 2115163.25 | 3540594 | 3537646.5 | 4123338.75 | 2532914.25 | 525952.25 | 1961701.88 | 1539841.5 | 752996.88 | 1427972.75 | 4055291.25 | 1803777 | 2340476 | 2886541 | 3029603 | 0 | 5 | 0 | 9 | 0 | 9 | 0.348253418 | 0.391709761 | 0.791795794 | 0.890598987 |
| P32233 | Drg1 | Developmentally-regulated GTP-binding protein 1 | 354.77 | 16.35 | 40512 | 6 | 1374078.5 | 1492018.5 | 1484171.75 | 792938.81 | 1299491.75 | 2174285.5 | 1797074 | 1422477.25 | 916385.44 | 1246717.62 | 3040175.5 | 3152560.5 | 2579766.5 | 2471443.75 | 2619185 | 1313201.88 | 1700365.5 | 1332502.5 | 1173468.5 | 1679211.5 | 0 | 10 | 0 | 11 | 0 | 11 | 2.151758228 | 1.834490098 | 1.11734997 | 0.952601193 |
| Q9QXB9 | Drg2 | Developmentally-regulated GTP-binding protein 2 | 359.15 | 19.23 | 40718 | 6 | 2474294.5 | 2597344 | 3429530.5 | 1902329.75 | 3375858.5 | 1279845.5 | 2310520 | 3109141.75 | 3648389 | 2615181.5 | 2144793.75 | 4508522 | 3092262.75 | 1075935.88 | 3319381.75 | 2715263 | 2 | 1547345 | 570795.69 | 1984779.25 | 12 | 4 | 12 | 5 | 12 | 5 | 1.026237717 | 1.09085947 | 0.494811537 | 0.525969609 |
| O35075 | Dscr3 | Down syndrome critical region protein 3 homolog | 90.62 | 6.73 | 32970 | 2 | 1157519.25 | 1078408.12 | 979522.5 | 1286018.38 | 1303547 | 1338826.75 | 1172271.25 | 537812.31 | 1213272.62 | 1717476 | 883196.75 | 705298.5 | 1343641.62 | 2 | 1252657.12 | 1300071.5 | 1197247.5 | 1438175.25 | 2 | 1345579.75 | 8 | 8 | 9 | 12 | 9 | 12 | 0.720893195 | 0.699838576 | 0.909743691 | 0.883173449 |
| Q9R0P5 | Dstn | Destrin | 548.12 | 60 | 18522 | 9 | 17215930 | 17366074 | 13515709 | 7917258.5 | 9084638 | 16077582 | 15841338 | 11719104 | 9936955 | 10945926 | 17174962 | 16231814 | 9213121 | 6775569 | 7636668 | 15252116 | 9482591 | 11253196 | 10808373 | 10762019 | 8 | 11 | 10 | 7 | 10 | 11 | 0.876074902 | 0.883932642 | 0.884157301 | 0.892087533 |
| Q9DD18 | Dtd1 | D-aminoacyl-tRNA deacylase 1 | 116.36 | 14.83 | 23384 | 2 | 2826744.25 | 2246407.75 | 1247130.25 | 2354354.5 | 1545010.38 | 2827348.5 | 2631227 | 2424835.5 | 1971774.5 | 2803348.25 | 1505392 | 1630732 | 1475067.12 | 1696732.12 | 361838 | 2304150 | 3711935.5 | 3209991.5 | 3342225.5 | 2519277 | 6 | 3 | 0 | 7 | 6 | 7 | 0.652641051 | 0.526898405 | 1.476330768 | 1.191889977 |
| Q3UIR3 | Dtx3l | E3 ubiquitin-protein ligase DTX3L | 251.72 | 8.96 | 83044 | 5 | 2934914.75 | 2278330.5 | 2402400 | 3082405.25 | 3344188.25 | 3569071.75 | 2609245 | 3647822.25 | 3079284.25 | 3088358.5 | 343808.38 | 1032422.75 | 546028.75 | 129778.22 | 701802.62 | 1487467.5 | 3209973 | 2707638.5 | 3046488 | 2762401.5 | 0 | 11 | 0 | 6 | 0 | 11 | 0.19611123 | 0.172181962 | 0.941015798 | 0.826194124 |
| P97930 | Dtymk | Thymidylate kinase | 80.05 | 10.85 | 23915 | 2 | 1323998.88 | 1963514.25 | 3033057.75 | 3334930 | 2313943.25 | 2733958.5 | 1363292.62 | 2567364.75 | 2340055 | 2829004.75 | 1597571 | 2298482.75 | 2817819 | 2649276.75 | 2603109 | 1289882.38 | 1195865.12 | 2953930 | 2702246.5 | 2488745.75 | 12 | 9 | 12 | 10 | 12 | 10 | 0.999733853 | 1.011203863 | 0.888150664 | 0.898340473 |
| Q3KRC5 | Dus3l | tRNA-dihydrouridine(47) synthase [NAD(P)(+)]-like | 47.12 | 2.03 | 71540 | 1 | 2 | 60380.93 | 252810.2 | 90001.62 | 146803.03 | 2 | 2 | 80025.64 | 79208.4 | 88672.42 | 272072.34 | 266907.5 | 314993.41 | 2 | 101711.73 | 2 | 2 | 2 | 2 | 85393.95 | 6.5 | 4 | 4 | 8 | 6.5 | 8 | 1.737619705 | 3.854968362 | 0.155276899 | 0.344487078 |
| Q9D7X3 | Dusp3 | Dual specificity protein phosphatase 3 | 475.25 | 50.81 | 20472 | 7 | 9857928 | 11076486 | 11509561 | 10437444 | 10983042 | 10484899 | 11010231 | 10318837 | 9993306 | 9967412 | 8977778 | 9555490 | 8397068 | 9744912 | 8440193 | 10662021 | 9921944 | 11822272 | 11059273 | 12052829 | 0 | 9 | 0 | 6 | 0 | 9 | 0.837573423 | 0.871380309 | 1.030704438 | 1.072306649 |
| P70583 | Dut | Deoxyuridine 5'-triphosphate nucleotidohydrolase | 66.86 | 5.85 | 22003 | 1 | 2 | 315644.81 | 2 | 2 | 2 | 2 | 2 | 2 | 2 | 2 | 926151 | 393771.91 | 51107.13 | 151629.69 | 412534.69 | 2 | 2 | 2 | 2 | 2 | 2 | 10 | 0 | 12.5 | 2 | 12.5 | 6.130768866 | 193519.442 | 3.168038E-05 | 1 |
| B4F766 | Dym | Dymeclin | 39.31 | 1.48 | 76491 | 1 | 2 | 2 | 165540.11 | 109675.88 | 178387.09 | 2 | 2 | 161300.61 | 124512.67 | 2 | 2 | 142720.83 | 108680.55 | 2 | 89483.12 | 2 | 2 | 2 | 2 | 64816.15 | 9 | 6 | 12 | 9 | 12 | 9 | 0.751506127 | 1.192671467 | 0.142908153 | 0.22680118 |
| Q9JHU4 | Dync1h1 | Cytoplasmic dynein 1 heavy chain 1 | 6142.521 | 27.07 | 532045 | 108 | 145004402 | 139568482 | 146092610 | 143011314 | 134834354 | 119510082 | 135156370 | 142668082 | 141465986 | 140054114 | 97889538 | 117178562 | 104135938 | 77970714 | 96465922 | 132771786 | 114973050 | 121877570 | 114409282 | 117620170 | 0 | 0 | 0 | 2 | 0 | 2 | 0.696729565 | 0.72716698 | 0.84917767 | 0.886274952 |
| O88487 | Dync1i2 | Cytoplasmic dynein 1 intermediate chain 2 | 393.35 | 17.16 | 68394 | 6 | 3277624.75 | 2388330.75 | 2841078.5 | 2868051.25 | 2532928.25 | 2670307 | 2174826.75 | 2531084.75 | 1893661.88 | 2325673 | 2553038.5 | 2519792.5 | 1055186.62 | 281301.78 | 2055111 | 2626277.5 | 3334059 | 1837800.88 | 1874137.25 | 2359453 | 3 | 7 | 8 | 12 | 8 | 12 | 0.608600962 | 0.729972096 | 0.86509318 | 1.037615648 |
| Q8R1Q8 | Dync1li1 | Cytoplasmic dynein 1 light intermediate chain 1 | 415.95 | 13.96 | 56614 | 6 | 2027032.25 | 2084705 | 2724696 | 1873185.75 | 4417844 | 1660826.88 | 3372727.5 | 2619562.5 | 3011225.75 | 1308792.62 | 4413063.5 | 4326550.5 | 2155299.25 | 1648945.75 | 2457707.5 | 3422057.25 | 1455943.5 | 2514982 | 2939937.5 | 1293907.5 | 11 | 11 | 10 | 11 | 11 | 11 | 1.142762048 | 1.252935525 | 0.885687337 | 0.971076289 |
| P63168 | Dynll1 | Dynein light chain 1, cytoplasmic | 95.03 | 37.08 | 10366 | 2 | 846282.88 | 778651.75 | 657885.31 | 355434.91 | 484499.16 | 305563.5 | 1051397.38 | 344684.25 | 984296.94 | 2 | 2 | 842535.69 | 2 | 2 | 2 | 2 | 2 | 2 | 2 | 2 | 4 | 0 | 5 | 2.5 | 5 | 2.5 | 0.269807896 | 0.313686238 | 3.202302E-06 | 3.723086E-06 |
| Q78P75 | Dynll2 | Dynein light chain 2, cytoplasmic | 127.38 | 25.84 | 10350 | 3 | 4234094.5 | 4005410.25 | 4965213 | 2752904.75 | 6888876.5 | 1583480 | 4336261 | 4836842.5 | 280943.94 | 3282770.5 | 2 | 3128723.5 | 1584506.12 | 2 | 2381511.75 | 4787477 | 2 | 2151612.25 | 1229479.88 | 3436873 | 1 | 4 | 6 | 10 | 6 | 10 | 0.310539719 | 0.495432805 | 0.507974729 | 0.810419181 |
| P62627 | Dynlrb1 | Dynein light chain roadblock-type 1 | 48.35 | 16.67 | 10990 | 1 | 654522.94 | 421278.47 | 403428.88 | 2 | 2 | 645714.06 | 334145.81 | 2 | 2 | 558725.31 | 2 | 2 | 2 | 2 | 2 | 2 | 1237818.5 | 595972.44 | 676848.56 | 407504.34 | 5 | 7 | 5 | 7 | 5 | 7 | 6.760254E-06 | 6.499461E-06 | 1.972740802 | 1.896637438 |
| P51807 | Dynlt1 | Dynein light chain Tctex-type 1 | 73.87 | 15.93 | 12483 | 1 | 291664.62 | 306325.31 | 624937.88 | 437678 | 484456.94 | 310499.5 | 259420.14 | 453556.09 | 643586.88 | 512112.06 | 453477.72 | 481512.5 | 477676.19 | 266331.59 | 735712.38 | 454573.03 | 523586.88 | 678633 | 608513.19 | 348281.62 | 11 | 7 | 11 | 7 | 11 | 7 | 1.125706173 | 1.108084824 | 1.21842017 | 1.199347513 |
| Q9D4C5 | Eaf1 | ELL-associated factor 1 | 48.2 | 3.73 | 28967 | 1 | 519954.62 | 919985.69 | 563450.75 | 825078.69 | 602661.31 | 550112.62 | 801002.19 | 602979.75 | 503239.88 | 386831.34 | 492067 | 382478.44 | 403645.78 | 629886.06 | 340531.12 | 891394.75 | 674444.31 | 580829.94 | 934630.44 | 663313.19 | 3 | 8 | 6 | 4 | 6 | 8 | 0.655354855 | 0.790603844 | 1.091363916 | 1.316594362 |
| Q9D903 | Ebna1bp2 | Probable rRNA-processing protein EBP2 | 260.77 | 16.67 | 34703 | 5 | 1573460.5 | 2579066.25 | 4371956 | 4376086 | 2023395.62 | 3041326.75 | 3045855.25 | 3321539.5 | 2831368 | 4802193 | 4297485.5 | 3461097.75 | 2249714 | 1076331.12 | 3217925 | 2290881.5 | 2806181 | 1072985.5 | 3218117.75 | 1745765.88 | 11 | 9 | 11 | 3 | 11 | 9 | 0.958361533 | 0.839239308 | 0.746043836 | 0.653312233 |
| P70245 | Ebp | 3-beta-hydroxysteroid-Delta(8),Delta(7)-isomerase | 39.66 | 4.35 | 26215 | 1 | 216528.83 | 2 | 758600.19 | 2 | 2 | 2 | 2 | 2 | 546564.62 | 2 | 4054871.5 | 3204041 | 2 | 2129757.5 | 5523081 | 149720.5 | 2 | 2 | 2634515.5 | 2 | 3.5 | 12.5 | 3 | 10 | 3.5 | 12.5 | 15.29198798 | 27.2822905 | 2.855237421 | 5.094001964 |
| O35459 | Ech1 | Delta(3,5)-Delta(2,4)-dienoyl-CoA isomerase, mitochondrial | 376.38 | 23.24 | 36118 | 6 | 2524929.25 | 3487035 | 6201564 | 5399724.5 | 5265445.5 | 1970672.88 | 2988957.5 | 4992284.5 | 3767702.75 | 3721606 | 2554676 | 3290287 | 5818557 | 4510396 | 5422029.5 | 1642513 | 2912197.5 | 4980943 | 6103778.5 | 5681810 | 12 | 11 | 8 | 10 | 12 | 11 | 0.943932442 | 1.238212751 | 0.931925487 | 1.222462509 |
| Q6AYG5 | Echdc1 | Ethylmalonyl-CoA decarboxylase | 85.61 | 5.69 | 32631 | 2 | 4555536.5 | 2 | 3088443.25 | 3485659 | 3114415.25 | 3844371.75 | 3571268.25 | 3375894 | 2598997 | 3549826.5 | 2569986.25 | 1784393.38 | 2403620.75 | 2641616.5 | 1964116.38 | 3569492.25 | 4294252.5 | 3603477.5 | 4011271 | 3706170.25 | 5 | 5 | 1 | 4 | 5 | 5 | 0.797787741 | 0.67080835 | 1.346853979 | 1.1324828 |
| Q8BH95 | Echs1 | Enoyl-CoA hydratase, mitochondrial | 468.06 | 35.86 | 31474 | 9 | 9979307 | 6286705 | 5104973 | 6703169 | 6592625.5 | 11275065 | 7426836 | 7789570 | 6736066 | 7310867 | 14751167 | 9621268 | 11057813 | 8953052 | 15725794 | 9080945 | 15836884 | 11572606 | 12389282 | 9650616 | 2 | 2 | 3 | 2 | 3 | 2 | 1.73391053 | 1.482769129 | 1.688369495 | 1.443824305 |
| P42125 | Eci1 | Enoyl-CoA delta isomerase 1, mitochondrial | 465.66 | 47.75 | 32250 | 7 | 3433240.5 | 3336398.25 | 4370804.5 | 2670969.25 | 3691248.5 | 4118672.5 | 4342862.5 | 4328974.5 | 5047957 | 5524563.5 | 7395487 | 6793461 | 8554648 | 9825218 | 9418746 | 2214910 | 2998927.75 | 6200718.5 | 6207767 | 5054152.5 | 0 | 9 | 0 | 11 | 0 | 11 | 2.398924369 | 1.797179561 | 1.295601609 | 0.970613647 |
| Q9WUR2 | Eci2 | Enoyl-CoA delta isomerase 2, mitochondrial | 165.26 | 7.67 | 43268 | 3 | 1868037.75 | 1703416.25 | 1996822.5 | 1986162.5 | 1904222.75 | 1797815 | 1034323.12 | 1922240.25 | 1749721 | 1018204.75 | 1325028.38 | 650874.38 | 1250633.38 | 1833523 | 2119126.75 | 1191405.25 | 1999126.5 | 1038065.75 | 1692437.25 | 1513149.25 | 6 | 5 | 12 | 12 | 12 | 12 | 0.759006515 | 0.954386552 | 0.785965731 | 0.988285488 |
| Q6PDI5 | Ecpas | Proteasome adapter and scaffold protein ECM29 | 473.61 | 6.58 | 203703 | 9 | 3587757.25 | 3416102.75 | 3556305.75 | 4387594 | 5294079.5 | 3854649.25 | 5067150 | 5454739 | 4151909 | 4649420 | 3224975.25 | 4483644.5 | 4342245 | 3334652 | 3733217.5 | 5104602 | 2876043.5 | 4404779.5 | 3572955.25 | 4513394.5 | 10 | 11 | 4 | 8 | 10 | 11 | 0.944515665 | 0.824870297 | 1.011359417 | 0.883246699 |
| Q3UJB9 | Edc4 | Enhancer of mRNA-decapping protein 4 | 522.49 | 7.75 | 152484 | 9 | 7991692 | 7875347 | 9350608 | 8552192 | 8658630 | 8079244 | 8093085.5 | 8713856 | 8554901 | 7415124.5 | 5799534 | 6969027.5 | 7638182.5 | 7639382 | 7359331 | 7600140 | 9313051 | 9223503 | 8368825.5 | 9317376 | 0 | 11 | 2 | 6 | 2 | 11 | 0.834474065 | 0.866586894 | 1.03286535 | 1.072612815 |
| O35474 | Edil3 | EGF-like repeat and discoidin I-like domain-containing protein 3 | 136.03 | 7.29 | 53712 | 3 | 278383.06 | 318849.03 | 60915.11 | 138268.36 | 208195.23 | 326443.91 | 413373.47 | 373618.25 | 1091285.5 | 2 | 966419.81 | 2406893 | 3626450.25 | 1772868 | 3352026 | 2 | 412693.19 | 383384.06 | 157010.42 | 2 | 0 | 12 | 1 | 8 | 1 | 12 | 12.06900939 | 5.499401215 | 0.948717334 | 0.432295401 |
| Q8BL66 | Eea1 | Early endosome antigen 1 | 689.88 | 11.69 | 160915 | 14 | 18843836 | 15539928 | 12045410 | 14729336 | 11680782 | 14896219 | 15195059 | 13786291 | 9988943 | 14282538 | 17800626 | 15360107 | 12953168 | 10698372 | 13710776 | 15312637 | 19872750 | 16972246 | 17930158 | 13849147 | 11 | 7 | 12 | 3 | 12 | 7 | 0.968200638 | 1.034835394 | 1.152357961 | 1.23166703 |
| Q921E6 | Eed | Polycomb protein EED | 50.21 | 2.95 | 50198 | 1 | 729371.5 | 2 | 2 | 2 | 433639.25 | 2 | 2 | 488127.69 | 2 | 599628.06 | 2 | 411295.66 | 2 | 369265.78 | 2 | 438406.12 | 683330.94 | 2 | 708060.94 | 2 | 10.5 | 10 | 10.5 | 9 | 10.5 | 10 | 0.671157522 | 0.717590447 | 1.573323858 | 1.68217167 |
| P10126 | Eef1a1 | Elongation factor 1-alpha 1 | 2160.719 | 61.9 | 50114 | 34 | 1030543618 | 1170794114 | 1197046914 | 1041232386 | 1137700610 | 979320002 | 1061729026 | 1169864706 | 1206226946 | 1098271234 | 1806016386 | 2043400578 | 1524526850 | 1392806274 | 1625211906 | 1178606850 | 1028110082 | 1149407106 | 1067687234 | 1210632066 | 0 | 11 | 0 | 10 | 0 | 11 | 1.504659145 | 1.521547642 | 1.010242504 | 1.021581602 |
| O70251 | Eef1b | Elongation factor 1-beta | 586.81 | 39.56 | 24694 | 9 | 131771842 | 121854370 | 128707994 | 149759090 | 110387106 | 139726210 | 120997874 | 124049722 | 122448898 | 137379346 | 107920082 | 105154066 | 96249626 | 107846266 | 102002162 | 147016594 | 188359506 | 156442082 | 169184274 | 151086946 | 0 | 1 | 0 | 0 | 0 | 1 | 0.808074768 | 0.805415065 | 1.263990932 | 1.259830623 |
| P57776 | Eef1d | Elongation factor 1-delta | 848.2 | 54.09 | 31293 | 13 | 107040474 | 101984394 | 112340730 | 117554170 | 100273450 | 113836402 | 102780050 | 93984738 | 93946170 | 107623186 | 74352722 | 82442754 | 77204906 | 74214418 | 71849442 | 113664906 | 120213322 | 124013970 | 123229770 | 125278306 | 0 | 1 | 0 | 1 | 0 | 1 | 0.704875784 | 0.742065792 | 1.124643734 | 1.183981154 |
| Q9D8N0 | Eef1g | Elongation factor 1-gamma | 1365.66 | 56.06 | 50061 | 21 | 170312690 | 175228626 | 184085762 | 184949970 | 168093346 | 179416610 | 168894690 | 178720290 | 162700370 | 170140674 | 211077890 | 225902082 | 177581234 | 178396434 | 182660818 | 204462722 | 203189874 | 206756370 | 203438162 | 195869618 | 6 | 0 | 4 | 0 | 6 | 0 | 1.105303253 | 1.134608103 | 1.148465784 | 1.178915 |
| P58252 | Eef2 | Elongation factor 2 | 3438.76 | 62.35 | 95314 | 54 | 329060322 | 351599810 | 367255362 | 346052962 | 348820674 | 353570722 | 306345506 | 353023106 | 377328962 | 367936002 | 493306274 | 474916226 | 397677058 | 414912578 | 381094370 | 348080866 | 392179650 | 332216066 | 393574690 | 393441154 | 0 | 7 | 0 | 8 | 0 | 8 | 1.240486568 | 1.229610523 | 1.066963521 | 1.05760885 |
| Q9JHW4 | Eefsec | Selenocysteine-specific elongation factor | 204 | 10.12 | 63539 | 5 | 2168941.75 | 3551377.5 | 2826348.25 | 1687028.88 | 2183126.5 | 2192777.5 | 2213017.75 | 2703186 | 2226457.5 | 3294565.5 | 3698659.5 | 3782455.5 | 3853694.5 | 1937033.62 | 4793248 | 2508414 | 861136.88 | 2453056.75 | 2157062.25 | 2584573.5 | 4 | 10 | 5 | 9 | 5 | 10 | 1.454888364 | 1.430331357 | 0.850800844 | 0.836440208 |
| Q4FZY0 | Efhd2 | EF-hand domain-containing protein D2 | 1184.57 | 59.83 | 26759 | 18 | 67667330 | 70539330 | 81479018 | 61647334 | 77141626 | 50672538 | 67809018 | 73450810 | 80635314 | 72807418 | 43513966 | 47312902 | 40859390 | 45750734 | 45903554 | 58002854 | 44637410 | 67405514 | 62757134 | 66464482 | 0 | 3 | 0 | 4 | 0 | 4 | 0.623030257 | 0.646660826 | 0.834835613 | 0.866499628 |
| Q8C0D5 | Efl1 | Elongation factor-like GTPase 1 | 204.8 | 3.55 | 125777 | 4 | 2273220 | 2338838.5 | 2245106 | 2594092.5 | 2189590.25 | 2409480.5 | 2201101.75 | 2588308.25 | 2680155.5 | 2408180.75 | 1922699.75 | 1855840.25 | 1744362 | 11608290 | 2277296.25 | 2436049.75 | 3479652 | 3438606.5 | 3432538.5 | 9249854 | 8 | 1 | 6 | 2 | 8 | 2 | 1.667274541 | 1.579566215 | 1.893049559 | 1.793464156 |
| O08810 | Eftud2 | 116 kDa U5 small nuclear ribonucleoprotein component | 1077.28 | 23.58 | 109361 | 20 | 15342416 | 12522402 | 13872018 | 12258745 | 16827270 | 14835546 | 14414613 | 13162887 | 16330368 | 11214880 | 11009032 | 10212536 | 10092388 | 9350003 | 11614514 | 16261616 | 13698714 | 12518574 | 12985200 | 11538068 | 0 | 9 | 1 | 9 | 1 | 9 | 0.738158268 | 0.747280558 | 0.946053019 | 0.95774451 |
| Q99MS7 | Ehbp1l1 | EH domain-binding protein 1-like protein 1 | 228.91 | 3.85 | 184834 | 4 | 1763612.75 | 1644569.25 | 1518586 | 1707403.38 | 1389441.88 | 2001114 | 1592359.88 | 1355649.5 | 1207934.62 | 1449296.5 | 1791866.75 | 1280854.5 | 1038001.5 | 1102639.5 | 1094526.25 | 1584950.75 | 2049778.75 | 2060741 | 1945780 | 1630714.62 | 5 | 6 | 5 | 4 | 5 | 6 | 0.786165571 | 0.829291943 | 1.15558475 | 1.218976202 |
| Q641Z6 | Ehd1 | EH domain-containing protein 1 | 1300.64 | 40.64 | 60603 | 22 | 19867912 | 20930430 | 17494354 | 17740606 | 19019650 | 20654194 | 18432034 | 18865630 | 18848330 | 20025234 | 24294008 | 29272462 | 24268062 | 21376980 | 23749172 | 19046954 | 20437218 | 17560422 | 20522000 | 20101454 | 0 | 9 | 0 | 10 | 0 | 10 | 1.293601949 | 1.269921488 | 1.027511991 | 1.008702529 |
| Q9EQP2 | Ehd4 | EH domain-containing protein 4 | 897.71 | 33.46 | 61481 | 16 | 25445490 | 22944646 | 25472126 | 28310800 | 25653818 | 20906108 | 23713010 | 23802804 | 27698488 | 26318510 | 14672710 | 19894054 | 18987064 | 15068005 | 20059944 | 21992832 | 18147290 | 19688014 | 22885084 | 21309906 | 0 | 0 | 0 | 3 | 0 | 3 | 0.6937647 | 0.724294015 | 0.813781311 | 0.849591992 |
| P48024 | Eif1 | Eukaryotic translation initiation factor 1 | 50.39 | 12.39 | 12747 | 1 | 2 | 463190.38 | 336170.81 | 2 | 309167.28 | 2 | 576658.38 | 650339.38 | 621818.88 | 468491.06 | 700219.94 | 669877.44 | 2 | 2 | 2 | 2 | 2 | 467406.91 | 474982.41 | 454262 | 12 | 9 | 11.5 | 5 | 12 | 9 | 1.235961433 | 0.591247419 | 1.259913767 | 0.602705508 |
| Q8BMJ3 | Eif1ax | Eukaryotic translation initiation factor 1A, X-chromosomal | 338.28 | 34.03 | 16460 | 5 | 4508761 | 4159454 | 5248629 | 4131101.25 | 231268.56 | 3106672 | 5022513.5 | 4005260.75 | 3463151.25 | 4016951.5 | 7397335 | 6928750 | 5232928 | 4325605 | 7354487 | 4937021 | 5615868 | 4513107.5 | 4431549.5 | 3603637 | 3 | 8 | 1 | 6 | 3 | 8 | 1.708996094 | 1.59264967 | 1.263795218 | 1.177757541 |
| Q8BJW6 | Eif2a | Eukaryotic translation initiation factor 2A | 353.99 | 15.32 | 64403 | 7 | 10271237 | 9056115 | 12031900 | 12480533 | 10220921 | 9349546 | 9968361 | 12045378 | 11337536 | 10735666 | 8657526 | 10205570 | 8654586 | 9865237 | 10829189 | 10679350 | 8301524 | 11893107 | 12414261 | 12864535 | 5 | 10 | 6 | 9 | 6 | 10 | 0.89181425 | 0.90223199 | 1.038698551 | 1.050832122 |
| Q03963 | Eif2ak2 | Interferon-induced, double-stranded RNA-activated protein kinase | 222.37 | 11.26 | 58280 | 3 | 549470.38 | 1249897.25 | 473650.03 | 1201546 | 455202.72 | 1234601.25 | 317014.47 | 436333.12 | 1253443.88 | 349184.53 | 1041095.94 | 1322828.12 | 1174182.38 | 686308.25 | 1326274.5 | 727941.88 | 690893.69 | 1290163 | 1178447.38 | 1170039.12 | 6 | 8 | 6 | 8 | 6 | 8 | 1.412473072 | 1.545904406 | 1.28696838 | 1.408543729 |
| Q64270 | Eif2b1 | Translation initiation factor eIF-2B subunit alpha | 44.47 | 2.95 | 33678 | 1 | 1111660.12 | 1186169.62 | 2042558.25 | 2250273 | 1607496.12 | 1244731.12 | 959241 | 1598072.38 | 1440366.62 | 1616951.88 | 1531445.12 | 1503546.38 | 1952112.75 | 2071079.38 | 1950225.38 | 1451163.38 | 1473485.38 | 1695639.88 | 1806256.12 | 1885284.12 | 11 | 12 | 4 | 4 | 11 | 12 | 1.098833419 | 1.313301105 | 1.013865527 | 1.211749383 |
| Q99LD9 | Eif2b2 | Translation initiation factor eIF-2B subunit beta | 43.31 | 4.56 | 38898 | 1 | 358795 | 413217.25 | 278451.28 | 369331.28 | 258486.33 | 333236.28 | 355701.22 | 2 | 2 | 360525.62 | 685621.12 | 554450.31 | 536127.88 | 491320.31 | 507121.31 | 2 | 2 | 364145.09 | 475630.75 | 2 | 0 | 8 | 0 | 12 | 0 | 12 | 1.653263487 | 2.643856941 | 0.500382099 | 0.800198333 |
| P70541 | Eif2b3 | Translation initiation factor eIF-2B subunit gamma | 337.64 | 14.82 | 50435 | 6 | 3654245.5 | 4349547.5 | 3522318.5 | 3173389 | 4824320.5 | 2977826.75 | 3826848.5 | 4757094 | 4850989.5 | 4287169.5 | 2006943 | 5226044 | 3176642.75 | 1979511.88 | 2471065.75 | 5702681 | 3316017 | 4521619.5 | 3278222 | 3308500.75 | 6 | 12 | 6 | 11 | 6 | 12 | 0.761132126 | 0.717886903 | 1.030896578 | 0.972324155 |
| Q61749 | Eif2b4 | Translation initiation factor eIF-2B subunit delta | 328.67 | 14.5 | 57624 | 6 | 3247852.5 | 5040503.5 | 5460648 | 4645250 | 4113787.5 | 3081380.25 | 2184176 | 5403548 | 5400072 | 6830617 | 7597456.5 | 5931220.5 | 8772362 | 9960244 | 9340261 | 2214191.25 | 2649621.75 | 5818426 | 7585296 | 6346404.5 | 0 | 10 | 1 | 10 | 1 | 10 | 1.84829693 | 1.816677712 | 1.093562028 | 1.074854224 |
| Q8CHW4 | Eif2b5 | Translation initiation factor eIF-2B subunit epsilon | 96.98 | 2.09 | 80086 | 2 | 1698342.62 | 1842075.38 | 3116361.5 | 4047185 | 3836379 | 3072830 | 3877036.5 | 3190823.5 | 3657961.75 | 3699941 | 3027878.5 | 3447893 | 3621411.5 | 2594289.25 | 3315098 | 4442901 | 2894207.5 | 4554215 | 4896507 | 4868225 | 12 | 3 | 6 | 5 | 12 | 5 | 1.100838522 | 0.914734715 | 1.489377159 | 1.237588406 |
| P68101 | Eif2s1 | Eukaryotic translation initiation factor 2 subunit 1 | 1203.76 | 69.84 | 36108 | 21 | 76437002 | 83664978 | 76139722 | 82376266 | 76909202 | 78707594 | 74851866 | 74594354 | 70109626 | 78879202 | 89454978 | 82565130 | 74207770 | 71941394 | 79066874 | 88145082 | 86855866 | 84762954 | 89289274 | 92938602 | 12 | 0 | 8 | 0 | 12 | 0 | 1.004320755 | 1.053278261 | 1.117475136 | 1.171948565 |
| Q99L45 | Eif2s2 | Eukaryotic translation initiation factor 2 subunit 2 | 677.22 | 48.64 | 38092 | 13 | 30187998 | 35690918 | 28081482 | 22938526 | 23885110 | 31115042 | 30063770 | 30930912 | 28557302 | 32691598 | 18887052 | 17399320 | 15913156 | 18329932 | 20291966 | 32045076 | 30169030 | 32022004 | 33022038 | 35767238 | 0 | 5 | 0 | 5 | 0 | 5 | 0.645111689 | 0.592215968 | 1.157982062 | 1.063033703 |
| Q9Z0N1 | Eif2s3x | Eukaryotic translation initiation factor 2 subunit 3, X-linked | 993.83 | 45.13 | 51065 | 15 | 41026146 | 37746286 | 30992512 | 35576362 | 33689946 | 39430634 | 40416566 | 47899294 | 42414654 | 42234882 | 62572882 | 57265502 | 53799214 | 51660934 | 58259138 | 41309214 | 43896446 | 45763762 | 44792794 | 47832466 | 0 | 0 | 0 | 7 | 0 | 7 | 1.583844535 | 1.335042232 | 1.248914251 | 1.052725336 |
| P23116 | Eif3a | Eukaryotic translation initiation factor 3 subunit A | 1126.26 | 17.04 | 161936 | 22 | 21868080 | 17756718 | 29420770 | 25521554 | 24696070 | 22871688 | 21121368 | 29724916 | 30217030 | 38599058 | 31167766 | 35451650 | 45348810 | 38272146 | 41807478 | 19714406 | 20188680 | 34475358 | 34781530 | 29335166 | 0 | 9 | 3 | 10 | 3 | 10 | 1.610286013 | 1.347382163 | 1.161256358 | 0.971663475 |
| Q8JZQ9 | Eif3b | Eukaryotic translation initiation factor 3 subunit B | 802.36 | 22.42 | 91370 | 15 | 15988394 | 12848348 | 12776970 | 10729108 | 10622604 | 12686530 | 14357862 | 9423728 | 9876042 | 9845633 | 15557657 | 11798723 | 14674587 | 10365191 | 10061481 | 14518064 | 14029498 | 12629252 | 7409394 | 9502003 | 10 | 10 | 6 | 12 | 10 | 12 | 0.991935495 | 1.111547729 | 0.922541409 | 1.033785779 |
| Q8R1B4 | Eif3c | Eukaryotic translation initiation factor 3 subunit C | 1215.83 | 24.59 | 105531 | 22 | 26428630 | 28545168 | 30785486 | 30230136 | 29459800 | 27935156 | 29476892 | 32486432 | 34040622 | 30967914 | 44159542 | 42834982 | 38036718 | 37864214 | 39112274 | 27397046 | 28968726 | 29393904 | 27916540 | 30374588 | 0 | 10 | 0 | 4 | 0 | 10 | 1.388853993 | 1.304057978 | 0.990385538 | 0.929917881 |
| O70194 | Eif3d | Eukaryotic translation initiation factor 3 subunit D | 654.4 | 30.66 | 63989 | 13 | 13827652 | 16328090 | 20177050 | 15381741 | 18003842 | 13909680 | 16230598 | 18825624 | 17180572 | 15205552 | 19397812 | 23323566 | 17398998 | 18116814 | 18487724 | 15438818 | 14459324 | 18957286 | 18378760 | 17440188 | 5 | 11 | 3 | 9 | 5 | 11 | 1.155360624 | 1.188967488 | 1.011419249 | 1.04083918 |
| P60229 | Eif3e | Eukaryotic translation initiation factor 3 subunit E | 690.78 | 32.81 | 52221 | 13 | 32631162 | 31372758 | 35847458 | 42737162 | 41754094 | 30146848 | 32536070 | 41172746 | 38754554 | 41144142 | 36452978 | 40762606 | 45260774 | 40210070 | 42872134 | 37818722 | 43446714 | 41427250 | 49232814 | 39317622 | 6 | 6 | 7 | 5 | 7 | 6 | 1.115089643 | 1.118659508 | 1.145926568 | 1.149595155 |
| Q9DCH4 | Eif3f | Eukaryotic translation initiation factor 3 subunit F | 633.19 | 28.25 | 37984 | 8 | 15227971 | 13969049 | 13148398 | 13985797 | 18631434 | 14228249 | 14074022 | 14056471 | 13176866 | 15623680 | 17381032 | 18585592 | 19262400 | 17073548 | 18236878 | 15292737 | 18627134 | 13408169 | 15343273 | 14304050 | 4 | 9 | 0 | 7 | 4 | 9 | 1.207794164 | 1.272349015 | 1.026849558 | 1.081733181 |
| Q9Z1D1 | Eif3g | Eukaryotic translation initiation factor 3 subunit G | 412.07 | 21.25 | 35638 | 7 | 13172290 | 12824299 | 13440771 | 16176288 | 12311142 | 12888302 | 12036649 | 13953353 | 11801702 | 15783256 | 11248912 | 10792458 | 9923438 | 8955490 | 10937724 | 12803637 | 11926550 | 14637110 | 17033636 | 15425285 | 0 | 11 | 0 | 9 | 0 | 11 | 0.763462382 | 0.78025093 | 1.057437469 | 1.080690532 |
| Q91WK2 | Eif3h | Eukaryotic translation initiation factor 3 subunit H | 561.51 | 30.68 | 39832 | 10 | 16404936 | 15878410 | 17663426 | 17950262 | 16166766 | 12865312 | 15049301 | 14907003 | 16781334 | 17775796 | 7557405 | 18624424 | 8687761 | 4062247 | 11807630 | 17246390 | 5571945 | 14422410 | 9750929 | 18897710 | 5 | 8 | 5 | 10 | 5 | 10 | 0.603582838 | 0.655728732 | 0.783802112 | 0.851517857 |
| B0BNA7 | Eif3i | Eukaryotic translation initiation factor 3 subunit I | 460.44 | 31.69 | 36461 | 9 | 2621145 | 2257842 | 2700000.5 | 2388992 | 2740422.5 | 2645388 | 2068384.62 | 2512597.75 | 3021868.5 | 1766340.38 | 7253555.5 | 5549140 | 6511433 | 6064594 | 6546310 | 3024335.5 | 3507737 | 3520692.5 | 3247407.5 | 2735716.5 | 0 | 1 | 0 | 1 | 0 | 1 | 2.512120131 | 2.657191054 | 1.261833628 | 1.334702503 |
| A0JPM9 | Eif3j | Eukaryotic translation initiation factor 3 subunit J | 281.79 | 17.76 | 29187 | 5 | 6021665.5 | 7876358 | 5645608 | 5808510.5 | 5412088 | 5565797 | 7611918.5 | 6912893.5 | 6753360 | 5093955 | 6242201.5 | 7644659 | 5762563 | 4796532 | 5125825 | 8990424 | 9273580 | 6056627 | 6886050 | 6986509 | 10 | 3 | 10 | 6 | 10 | 6 | 0.961239092 | 0.925914299 | 1.241480447 | 1.195857001 |
| Q9DBZ5 | Eif3k | Eukaryotic translation initiation factor 3 subunit K | 174.56 | 20.64 | 25087 | 3 | 1582692 | 1414594.5 | 1585392.25 | 984350.06 | 1154802.38 | 925539.19 | 1459901.88 | 1771032.88 | 1526499 | 977223 | 2723163.75 | 3300840.5 | 1918106.75 | 985365.06 | 3260164 | 1581193.62 | 787797.25 | 1232181 | 1674245.75 | 928024.81 | 4 | 10 | 3 | 11 | 4 | 11 | 1.813142835 | 1.829922145 | 0.922879831 | 0.931420408 |
| Q8QZY1 | Eif3l | Eukaryotic translation initiation factor 3 subunit L | 966.92 | 28.37 | 66613 | 15 | 5480722.5 | 5829148.5 | 8327198 | 8944528 | 8713883 | 4721745 | 6392368 | 8097364 | 10770226 | 7273625 | 11395667 | 13219535 | 14181032 | 12278080 | 15104551 | 6440975.5 | 6879478.5 | 7448142.5 | 7800854.5 | 8006947 | 0 | 10 | 0 | 12 | 0 | 12 | 1.774447332 | 1.776359746 | 0.980719326 | 0.981776298 |
| Q99JX4 | Eif3m | Eukaryotic translation initiation factor 3 subunit M | 555.22 | 35.29 | 42517 | 8 | 6211157 | 7257501 | 6181710 | 6009001 | 6440972 | 6154293 | 6465750.5 | 6832967 | 5808833.5 | 5091135 | 9042360 | 8652593 | 6217959.5 | 5688922 | 8186182.5 | 5610883 | 6202477 | 5840398.5 | 5115403 | 6062122 | 7 | 3 | 6 | 9 | 7 | 9 | 1.177184286 | 1.244952497 | 0.898161284 | 0.949866684 |
| P60843 | Eif4a1 | Eukaryotic initiation factor 4A-I | 2043.49 | 77.09 | 46154 | 28 | 46953838 | 47131014 | 57355494 | 28802696 | 93267698 | 18077458 | 60568050 | 72864314 | 88334090 | 39397298 | 21281174 | 75120010 | 32212552 | 21102308 | 44699558 | 54673766 | 19814160 | 33490392 | 25062486 | 33150438 | 6 | 5 | 9 | 6 | 9 | 6 | 0.710815239 | 0.696228189 | 0.607622362 | 0.595152993 |
| Q91VC3 | Eif4a3 | Eukaryotic initiation factor 4A-III | 1330.12 | 50.12 | 46840 | 22 | 47629754 | 45397810 | 49000850 | 41542466 | 50426294 | 43652706 | 49422302 | 52445638 | 52945362 | 45960322 | 38884346 | 45033154 | 40377630 | 31224234 | 44650270 | 48865194 | 37435774 | 35312734 | 41539406 | 43012858 | 2 | 4 | 2 | 2 | 2 | 4 | 0.855436117 | 0.818936462 | 0.881061777 | 0.843468729 |
| Q8BGD9 | Eif4b | Eukaryotic translation initiation factor 4B | 741.04 | 26.19 | 68840 | 10 | 17748954 | 22763940 | 19662652 | 17461068 | 16760939 | 19318154 | 20006486 | 18544922 | 15732971 | 19276450 | 19987752 | 25720972 | 15633768 | 19967308 | 14971535 | 20989858 | 20544872 | 21025750 | 23335838 | 21346908 | 12 | 4 | 12 | 0 | 12 | 4 | 1.019955835 | 1.036632098 | 1.136080572 | 1.154655472 |
| P63073 | Eif4e | Eukaryotic translation initiation factor 4E | 126.91 | 8.29 | 25053 | 3 | 1180814.75 | 1638962.75 | 2004141.75 | 1020816.12 | 2911861 | 1019402.94 | 1236599.25 | 2191592.75 | 1887025.5 | 1106056 | 1462283.25 | 2141848.75 | 1636690.25 | 1445629 | 1137809.25 | 1580823.25 | 1100882.75 | 2254527.75 | 2101227.5 | 2820021.5 | 11 | 10 | 10 | 7 | 11 | 10 | 0.893527596 | 1.05155231 | 1.125720809 | 1.324810026 |
| Q6NZJ6 | Eif4g1 | Eukaryotic translation initiation factor 4 gamma 1 | 1825.47 | 21.25 | 176077 | 33 | 33705538 | 33268010 | 35229510 | 30982480 | 31683804 | 35002890 | 31832786 | 30486514 | 31611836 | 27608602 | 39694238 | 42621478 | 33852582 | 31758800 | 32402220 | 39420674 | 30922212 | 34192534 | 34152078 | 36692018 | 7 | 7 | 4 | 5 | 7 | 7 | 1.093771078 | 1.151950241 | 1.063748505 | 1.120330726 |
| Q62448 | Eif4g2 | Eukaryotic translation initiation factor 4 gamma 2 | 941.63 | 22.08 | 102106 | 20 | 10674454 | 11612718 | 12654184 | 8810106 | 13159489 | 8165306 | 11064697 | 10153783 | 12431849 | 6713764 | 21811288 | 28220834 | 20006296 | 17815760 | 21311028 | 11309470 | 7730185 | 8962351 | 6634666 | 9799681 | 0 | 4 | 0 | 9 | 0 | 9 | 1.918175748 | 2.249465443 | 0.780804963 | 0.915658424 |
| Q80XI3 | Eif4g3 | Eukaryotic translation initiation factor 4 gamma 3 | 482.53 | 5.64 | 174890 | 9 | 1293757.25 | 1187060.5 | 1393129.12 | 1041817.75 | 839202.25 | 1406657.62 | 1058526 | 1283696.5 | 900467.25 | 904453.88 | 1077885.38 | 1158982.62 | 1256659.5 | 1243003.75 | 853417 | 1292911.88 | 1374019.5 | 1312425.12 | 1239225.75 | 1361708.62 | 11 | 7 | 12 | 6 | 12 | 7 | 0.971325878 | 1.006508515 | 1.14341073 | 1.184826495 |
| Q5XI72 | Eif4h | Eukaryotic translation initiation factor 4H | 498.97 | 47.58 | 27324 | 9 | 15012649 | 17119632 | 14884545 | 15396530 | 18953138 | 13795413 | 15898705 | 15485674 | 18240350 | 17776976 | 12265596 | 15499071 | 14113599 | 11309629 | 11994262 | 14069565 | 14806264 | 15644080 | 15710211 | 16631547 | 3 | 9 | 3 | 9 | 3 | 9 | 0.801093347 | 0.802764416 | 0.944635356 | 0.946605851 |
| P59325 | Eif5 | Eukaryotic translation initiation factor 5 | 263.76 | 10.72 | 48968 | 5 | 7082615 | 7857169.5 | 5683370 | 5490265 | 6287326.5 | 5610213 | 7571847 | 4959265.5 | 4069476.75 | 5503212 | 5615752 | 7354366.5 | 5347855 | 4903392 | 6115410 | 8047330.5 | 5568204 | 4828064 | 5043112 | 5716819 | 7 | 8 | 10 | 10 | 10 | 10 | 0.905435187 | 1.058553815 | 0.90132275 | 1.053745922 |
| P63242 | Eif5a | Eukaryotic translation initiation factor 5A-1 | 822.47 | 83.12 | 16832 | 12 | 93043074 | 103897738 | 107832522 | 86984962 | 133373298 | 87807586 | 101914802 | 101696210 | 138076034 | 81486618 | 65055850 | 91225234 | 84518298 | 77611162 | 81487674 | 100465074 | 78328450 | 101274898 | 76279154 | 107457962 | 1 | 7 | 4 | 8 | 4 | 8 | 0.761520012 | 0.782608399 | 0.883217737 | 0.907676237 |
| Q05D44 | Eif5b | Eukaryotic translation initiation factor 5B | 1089.44 | 22.53 | 137616 | 22 | 14216167 | 12673791 | 15250753 | 13277721 | 12549709 | 12063978 | 15050880 | 14282578 | 15033705 | 13362029 | 14839363 | 16073922 | 15886626 | 12622184 | 16144581 | 11746661 | 9628013 | 15506236 | 14325501 | 15157369 | 5 | 12 | 6 | 12 | 6 | 12 | 1.11179554 | 1.08272308 | 0.976395397 | 0.95086353 |
| O55135 | Eif6 | Eukaryotic translation initiation factor 6 | 345.48 | 26.94 | 26511 | 4 | 18018272 | 18007302 | 14240986 | 16138637 | 14202554 | 18133672 | 17437594 | 15082950 | 19283738 | 15859921 | 18379562 | 18082514 | 17013190 | 17601362 | 17987540 | 18055366 | 22712560 | 19711514 | 20161832 | 20063878 | 6 | 0 | 10 | 2 | 10 | 2 | 1.104908236 | 1.038069626 | 1.249323406 | 1.173748767 |
| Q5PPK9 | Eipr1 | EARP and GARP complex-interacting protein 1 | 47.73 | 2.33 | 43157 | 1 | 1313429 | 1378099.88 | 1441632.38 | 1912876.12 | 1297479.12 | 1302592.25 | 1347871.62 | 1246203.88 | 1672854.38 | 1849405.75 | 888687 | 2 | 737996.25 | 2 | 2 | 1206113.5 | 862123 | 1289087.75 | 1354703.5 | 1374582.12 | 0 | 4 | 0 | 7 | 0 | 7 | 0.221513664 | 0.219262038 | 0.828841315 | 0.820416369 |
| Q80Y81 | Elac2 | Zinc phosphodiesterase ELAC protein 2 | 64.54 | 2.29 | 92719 | 1 | 411921.41 | 539142.81 | 423149.16 | 353542.72 | 469474.38 | 341959.69 | 492800.47 | 493854.12 | 509729.25 | 365801.47 | 2 | 438509.03 | 234501.08 | 2 | 383863.69 | 437258.81 | 2 | 450497 | 398537.56 | 338735.84 | 4 | 7 | 4 | 6 | 4 | 7 | 0.481004523 | 0.479495587 | 0.739581589 | 0.737261482 |
| B5DF91 | Elavl1 | ELAV-like protein 1 | 776.42 | 43.25 | 36169 | 15 | 34766586 | 31866640 | 32819010 | 33502792 | 29180594 | 36762278 | 34003106 | 27135548 | 26841338 | 30704490 | 29494540 | 25853546 | 23527778 | 21310742 | 23892150 | 33660830 | 39922698 | 30203866 | 31638644 | 32283148 | 1 | 12 | 2 | 9 | 2 | 12 | 0.765277577 | 0.798207412 | 1.034375937 | 1.078885054 |
| Q8BPU7 | Elmo1 | Engulfment and cell motility protein 1 | 455.51 | 12.65 | 83936 | 9 | 10474585 | 11417404 | 12314479 | 12679575 | 11089144 | 9561596 | 10135867 | 11454115 | 11787528 | 11515365 | 5902627 | 8792404 | 5802504 | 2810125.75 | 8685426 | 10600521 | 8862784 | 8724269 | 8454968 | 6767100 | 0 | 1 | 0 | 2 | 0 | 2 | 0.551841027 | 0.587519926 | 0.748762432 | 0.797173147 |
| Q8BHL5 | Elmo2 | Engulfment and cell motility protein 2 | 100.94 | 4.78 | 83887 | 3 | 2 | 2 | 2 | 2 | 2 | 2 | 2 | 127470.35 | 2 | 2 | 131182.66 | 2 | 129922.59 | 2 | 2 | 2 | 2 | 159017.23 | 2 | 80496.93 | 7.5 | 7.5 | 9 | 10 | 9 | 10 | 26111.125 | 2.048279178 | 23952.016 | 1.878908536 |
| Q8CB77 | Eloa | Elongin-A | 102.91 | 5.43 | 87161 | 2 | 310114.16 | 339946.28 | 478889.5 | 317594.78 | 499651.5 | 355039.66 | 450818.81 | 450546.59 | 348876.03 | 263342.84 | 102164.05 | 330001.5 | 79621.67 | 216488.08 | 2 | 352338.78 | 146348.56 | 209238.62 | 300729.91 | 341288.5 | 2 | 6 | 1 | 4 | 2 | 6 | 0.374205485 | 0.389739898 | 0.693632202 | 0.722426995 |
| P62869 | Elob | Elongin-B | 421.8 | 49.15 | 13170 | 7 | 10117124 | 11082114 | 11185229 | 11492752 | 11318186 | 9141868 | 10096282 | 11598962 | 8021507.5 | 8515193 | 7229582 | 7636213 | 8266598 | 5479339 | 8369458 | 10867246 | 11081651 | 10892940 | 12324654 | 12227384 | 0 | 12 | 2 | 3 | 2 | 12 | 0.670004867 | 0.780625161 | 1.039830671 | 1.211510579 |
| P83940 | Eloc | Elongin-C | 145.55 | 17.86 | 12473 | 1 | 1388173 | 1240285.38 | 1037827 | 848497.56 | 1216629.5 | 1177662.88 | 1549580.5 | 1239994.62 | 1081936.88 | 1011246.12 | 503042.78 | 784307.25 | 891560.31 | 523465.25 | 1030187.38 | 1091573.12 | 1079793.75 | 910376.19 | 1016080.25 | 935421.5 | 2 | 7 | 1 | 4 | 2 | 7 | 0.651246618 | 0.615891696 | 0.878185764 | 0.830510753 |
| Q9JLJ5 | Elovl1 | Elongation of very long chain fatty acids protein 1 | 161.51 | 12.54 | 32678 | 3 | 232668.17 | 162598.12 | 146736.78 | 63342.02 | 248787.86 | 128353.64 | 2 | 405928.25 | 331378.53 | 60543.55 | 1397396.25 | 1239972.75 | 633588 | 1194960.5 | 978077.25 | 191717.67 | 260532.53 | 335626.31 | 432603.5 | 621376.75 | 0 | 2 | 0 | 5 | 0 | 5 | 6.373708859 | 5.877736623 | 2.156405229 | 1.988603852 |
| Q7TT37 | Elp1 | Elongator complex protein 1 | 94.95 | 1.5 | 149584 | 2 | 1031274.44 | 1091911.75 | 1102650.5 | 2 | 1134935 | 763406.69 | 1065633.62 | 975952.38 | 953457.75 | 526756.31 | 1702377.88 | 1914105.25 | 1894216.88 | 1295163.5 | 1324139 | 1253002.25 | 771725.19 | 2 | 648942.75 | 732505.38 | 0 | 8.5 | 0 | 9 | 0 | 9 | 1.864348643 | 1.897225265 | 0.781094781 | 0.794868899 |
| Q9CZX0 | Elp3 | Elongator complex protein 3 | 230.62 | 10.42 | 62385 | 4 | 238420.45 | 872307 | 637537.25 | 171146.84 | 1226126.25 | 326355.09 | 862631.88 | 813968.12 | 1139762.25 | 259868.66 | 2 | 900241.25 | 2 | 112565.6 | 263228.34 | 637706.56 | 176281.05 | 188139.91 | 131090.69 | 305348.5 | 6 | 7 | 5 | 3 | 6 | 7 | 0.405666463 | 0.375020408 | 0.457335695 | 0.42278629 |
| Q8C7X2 | Emc1 | ER membrane protein complex subunit 1 | 1552.84 | 33.3 | 111605 | 26 | 18481804 | 17367878 | 19549058 | 16301832 | 15415819 | 19980334 | 20930014 | 16942430 | 16584822 | 19260798 | 34832042 | 31491022 | 30645086 | 31147820 | 33525678 | 21983420 | 21412606 | 22343084 | 21251526 | 19101662 | 0 | 1 | 0 | 3 | 0 | 3 | 1.855467681 | 1.725127125 | 1.217822465 | 1.132274407 |
| B0BNG0 | Emc2 | ER membrane protein complex subunit 2 | 215.4 | 17.51 | 34870 | 4 | 3389873.75 | 2336424 | 2317062.25 | 1303853.75 | 2490317.5 | 2473223.75 | 3715608.5 | 3923817.5 | 3470784.5 | 3128738.5 | 3839772.5 | 5214697 | 6114067.5 | 5526436 | 6913969.5 | 3083924.25 | 2043757.5 | 3501314.5 | 2826593 | 3243633.25 | 0 | 7 | 1 | 7 | 1 | 7 | 2.332322671 | 1.652025916 | 1.241747303 | 0.879551852 |

| | | | | | | | | | | | | | | | | | | | | | | | | | | | | | | | | | | | | |
|---|---|---|---|---|---|---|---|---|---|---|---|---|---|---|---|---|---|---|---|---|---|---|---|---|---|---|---|---|---|---|---|---|---|---|---|---|
| Q5U2V8 | Emc3 | ER membrane protein complex subunit 3 | 308.13 | 27.2 | 29980 | 6 | 4587161 | 2964865.5 | 2797672.75 | 3408000 | 3217148.25 | 3201659.75 | 4357719.5 | 2878517.25 | 3319192.5 | 3109040 | 7005619 | 4582609.5 | 6556658 | 6691622.5 | 6354937.5 | 4056032.5 | 6490242.5 | 3744880.75 | 3916222 | 1642932.25 | 1 | 8 | 0 | 8 | 1 | 8 | 1.837509674 | 1.8493542 | 1.169395484 | 1.176933367 |
| O70378 | Emc8 | ER membrane protein complex subunit 8 | 291.31 | 26.09 | 23348 | 4 | 4785793 | 4529128.5 | 4848187.5 | 5255306 | 4241928 | 6027472 | 4414625.5 | 4325981.5 | 4057900.5 | 5482924 | 7291946.5 | 7445748 | 7710460 | 5920541 | 7935277.5 | 4050450.75 | 6183388 | 6369153 | 6755309.5 | 6744128 | 0 | 5 | 1 | 5 | 1 | 5 | 1.534380672 | 1.493443462 | 1.272273578 | 1.238329374 |
| O08579 | Emd | Emerin | 173.46 | 16.99 | 29436 | 3 | 2954163 | 1767117.62 | 5941719 | 6490122 | 3816134 | 3573880.5 | 3987903.25 | 3976799.25 | 3752722.5 | 4636045.5 | 2272246.25 | 3424664 | 3100316 | 3047099.5 | 2824053.5 | 4253365 | 4479351 | 5143451 | 4510100 | 4837664 | 8 | 10 | 0 | 3 | 8 | 10 | 0.699518358 | 0.736092783 | 1.10752291 | 1.165429916 |
| O35130 | Emg1 | Ribosomal RNA small subunit methyltransferase NEP1 | 233.85 | 24.59 | 26974 | 4 | 426419.81 | 1011391.56 | 737607.38 | 536937.12 | 555231.12 | 736113.19 | 550164 | 923984.62 | 604435.62 | 725618.69 | 2476015.75 | 2024096.12 | 2218142.75 | 2108873.25 | 2135799 | 270581.66 | 404482.84 | 941546.81 | 632047.25 | 370633.56 | 0 | 7 | 0 | 7 | 0 | 7 | 3.355052797 | 3.096595473 | 0.801598283 | 0.73984696 |
| Q6P6T4 | Eml2 | Echinoderm microtubule-associated protein-like 2 | 89.11 | 4.78 | 70711 | 2 | 1969360.75 | 1691764.62 | 2002754.88 | 2089358.88 | 2504038 | 1593188.5 | 1930653 | 2120536.25 | 1854927.5 | 1967721.62 | 1378754.5 | 330893.44 | 1791958.88 | 1088378.88 | 1757108.25 | 2057380.5 | 995826 | 2375247 | 1986140.5 | 2268018.25 | 2 | 12 | 2 | 7 | 2 | 12 | 0.61878936 | 0.670442161 | 0.943974909 | 1.022772237 |
| Q8C522 | Endod1 | Endonuclease domain-containing 1 protein | 56.85 | 2.2 | 55262 | 1 | 2 | 2 | 2 | 2 | 2 | 2 | 2 | 2 | 2 | 2 | 315772.56 | 381703.91 | 737180.94 | 646391.44 | 684715.44 | 2 | 2 | 2 | 2 | 427658.72 | 0 | 10 | 0 | 10 | 0 | 10 | 276576.429 | 276576.429 | 42766.672 | 42766.672 |
| P17182 | Eno1 | Alpha-enolase | 2661.16 | 82.26 | 47141 | 36 | 346858626 | 385324514 | 274449090 | 258186658 | 256981954 | 321404098 | 360957698 | 260688434 | 245639106 | 275950978 | 395460034 | 419464482 | 279993058 | 280040226 | 314703650 | 406430562 | 320031234 | 299586210 | 288790914 | 292881250 | 6 | 8 | 6 | 8 | 6 | 8 | 1.110303926 | 1.153635766 | 1.056458983 | 1.097689415 |
| P17183 | Eno2 | Gamma-enolase | 694.65 | 29.26 | 47297 | 9 | 836248 | 1038044.12 | 1051601 | 2293538 | 1909041.62 | 1650298.25 | 2044021.38 | 1918787.5 | 2093685.38 | 2107583.5 | 1372220.25 | 1432470 | 2597595 | 3143730 | 2885113 | 3261036 | 2103995.5 | 686086.38 | 1532272.5 | 2084500 | 4 | 9 | 10 | 12 | 10 | 12 | 1.603587285 | 1.164733065 | 1.356235863 | 0.985074382 |
| Q8BGB7 | Enoph1 | Enolase-phosphatase E1 | 90.27 | 7.39 | 28600 | 2 | 1434683.25 | 1709498 | 1446047.25 | 1567400.62 | 1252940.5 | 1184888.12 | 1256180 | 1550899.75 | 1339288.38 | 1164832.12 | 735044.44 | 1004465.5 | 791417.62 | 852323.69 | 1324226.38 | 1318631.75 | 1078959 | 1390313.75 | 1245705.25 | 1394067.12 | 1 | 3 | 3 | 12 | 3 | 12 | 0.635238298 | 0.724663422 | 0.8673661 | 0.989468816 |
| Q8BTJ4 | Enpp4 | Bis(5'-adenosyl)-triphosphatase enpp4 | 128.44 | 4.39 | 51611 | 2 | 901444.88 | 962755.5 | 907018.75 | 1066310.88 | 1216423.62 | 1368656.38 | 1372424.5 | 2 | 1872384.25 | 1357201 | 42848.02 | 2 | 1453047.5 | 987235.81 | 580072.44 | 707661.12 | 848582.38 | 1412496.75 | 1513049.5 | 1257581.12 | 8 | 10 | 7.5 | 11 | 8 | 11 | 0.606100885 | 0.513042377 | 1.13562001 | 0.961261076 |
| Q8BYW9 | Eogt | EGF domain-specific O-linked N-acetylglucosamine transferase | 128.92 | 7.21 | 61446 | 3 | 1039750 | 1454381.75 | 858395.88 | 905750.12 | 730591.06 | 1035952.69 | 1152567.25 | 743724.06 | 829187.31 | 581477.31 | 620405.75 | 1387387.88 | 1628354.62 | 1723437 | 1179177.25 | 1103472 | 982782.62 | 893486.31 | 633919.12 | 746233.75 | 7 | 10 | 4 | 12 | 7 | 12 | 1.310670364 | 1.505618255 | 0.873924324 | 1.003911015 |
| P48193 | Epb41 | Protein 4.1 | 340.92 | 11.42 | 95911 | 7 | 1613756.38 | 895739.12 | 2162445.25 | 3595866.5 | 1620107.75 | 2126523.5 | 1644746 | 2232761 | 1390715.5 | 3603132 | 3679732.75 | 2325441 | 2738787 | 3438329.5 | 3238924.75 | 555991.38 | 2353635.5 | 1327287.75 | 3482920 | 2199728.25 | 4 | 12 | 4 | 11 | 4 | 12 | 1.559602302 | 1.402199133 | 1.003200663 | 0.901952438 |
| O70318 | Epb41l2 | Band 4.1-like protein 2 | 504.18 | 11.23 | 109940 | 10 | 5549793 | 5539100 | 4900328.5 | 6415429.5 | 4459572 | 6495210.5 | 6190053 | 7265797 | 8781715 | 7491951.5 | 6980459 | 7368151 | 10194744 | 9359352 | 10478642 | 4761176.5 | 3394900.25 | 6647906.5 | 4111196.25 | 6561500.5 | 0 | 11 | 5 | 4 | 5 | 11 | 1.65206148 | 1.225167218 | 0.948349781 | 0.703295293 |
| Q03145 | Epha2 | Ephrin type-A receptor 2 | 51.16 | 1.33 | 108852 | 1 | 101988.66 | 2 | 73858.43 | 2 | 2 | 2 | 186186.83 | 43279.79 | 2 | 2 | 2 | 132657.59 | 2 | 2 | 26644.54 | 2 | 2 | 2 | 2 | 2 | 12.5 | 7.5 | 11.5 | 7.5 | 12.5 | 7.5 | 0.905916012 | 0.694235896 | 5.686565E-05 | 4.357818E-05 |
| Q8CGC7 | Eprs | Bifunctional glutamate/proline--tRNA ligase | 2976.59 | 37.43 | 170079 | 47 | 69261714 | 79808250 | 73123498 | 78397066 | 72556914 | 75767354 | 73700258 | 73577666 | 83873530 | 83770346 | 154616690 | 150332530 | 132186002 | 144640738 | 126726594 | 79761218 | 94751962 | 91702866 | 86288674 | 89401194 | 0 | 1 | 0 | 2 | 0 | 2 | 1.898720115 | 1.813468705 | 1.184266229 | 1.131093376 |
| P42567 | Eps15 | Epidermal growth factor receptor substrate 15 | 124.2 | 1.34 | 98471 | 2 | 503380.38 | 638999.12 | 496923.94 | 543143 | 467423.22 | 900699.44 | 674197.06 | 864895.81 | 555644.5 | 442000.72 | 559712 | 477932.94 | 413505.47 | 177853.97 | 294519.47 | 712215.38 | 1204418.25 | 950695.38 | 587881.81 | 507547.12 | 5 | 3 | 3 | 9 | 5 | 9 | 0.72589376 | 0.559580744 | 1.495453908 | 1.15282326 |
| Q60902 | Eps15l1 | Epidermal growth factor receptor substrate 15-like 1 | 228.95 | 5.18 | 99309 | 4 | 942557.06 | 923638.31 | 677712.81 | 852186.44 | 1231511.25 | 1071731.12 | 869466.25 | 770452.56 | 1328456.88 | 845377.62 | 893462.12 | 953161.5 | 401200.41 | 494681.59 | 948246.56 | 960518.62 | 1152637.25 | 1061164.5 | 897015.31 | 916371.69 | 10 | 9 | 9 | 9 | 10 | 9 | 0.797551106 | 0.755452654 | 1.07781594 | 1.020923808 |
| Q08509 | Eps8 | Epidermal growth factor receptor kinase substrate 8 | 42.14 | 3.29 | 91737 | 1 | 2 | 144826.5 | 2 | 188466.95 | 210776.78 | 2 | 2 | 2 | 190396.11 | 259387.08 | 2 | 2 | 2 | 2 | 73390.84 | 243249.39 | 2 | 161200.89 | 2 | 2 | 6 | 11 | 9 | 11.5 | 9 | 11.5 | 0.13490593 | 0.16318498 | 0.743384372 | 0.899212985 |
| Q9EQH2 | Erap1 | Endoplasmic reticulum aminopeptidase 1 | 1195.16 | 27.85 | 106599 | 23 | 15928867 | 17380890 | 20719174 | 21131884 | 21456402 | 16762797 | 19238832 | 20854920 | 23770178 | 20499010 | 17647346 | 20107688 | 28858864 | 26812716 | 24815430 | 15239297 | 12707638 | 16510649 | 12291273 | 16813998 | 6 | 2 | 7 | 1 | 7 | 2 | 1.223819601 | 1.169257674 | 0.761384537 | 0.727439494 |
| Q80TH2 | Erbin | Erbin | 45.36 | 0.86 | 157248 | 1 | 503436.59 | 418326.12 | 650377.25 | 715211.25 | 545994.56 | 417698.62 | 378373.84 | 461853.22 | 506947.28 | 685470.06 | 418373.72 | 232612.19 | 263356.53 | 231893.73 | 299813.72 | 442198.59 | 589745.81 | 447277.34 | 516479.06 | 486257.12 | 1 | 8 | 2 | 10 | 2 | 10 | 0.510368309 | 0.59014182 | 0.875981303 | 1.012902234 |
| Q9DC16 | Ergic1 | Endoplasmic reticulum-Golgi intermediate compartment protein 1 | 152.45 | 9.66 | 32562 | 3 | 3864363.75 | 4039279.5 | 4870981 | 4051686 | 3267568 | 4070570 | 4009455.5 | 4029218 | 4189291.5 | 4597938 | 6824986 | 7070171 | 8384725 | 7331553 | 8133665 | 4438117 | 4922859 | 5580743 | 5117602 | 5355347.5 | 0 | 1 | 0 | 1 | 0 | 1 | 1.878437777 | 1.806290468 | 1.264796581 | 1.216218091 |
| Q9CQE7 | Ergic3 | Endoplasmic reticulum-Golgi intermediate compartment protein 3 | 101.81 | 4.44 | 43208 | 2 | 239605.31 | 38680.39 | 73586.16 | 458031.59 | 78549.29 | 40155.32 | 2 | 93574.88 | 88107.98 | 544705.62 | 874711.56 | 763483 | 1259647.88 | 1116063.88 | 1332706.88 | 33148.82 | 28714.62 | 344427.19 | 242481.55 | 55899.18 | 0 | 9 | 0 | 12 | 0 | 12 | 6.017892634 | 6.974942919 | 0.79314445 | 0.919281483 |
| P84089 | Erh | Enhancer of rudimentary homolog | 40.22 | 10.58 | 12259 | 1 | 1039916.62 | 863728.75 | 1239884.62 | 925381.56 | 1056677.88 | 435675.44 | 946246.12 | 1104444.62 | 1078584.5 | 997470.44 | 2 | 2 | 943242.62 | 2 | 998449.5 | 2 | 2 | 649528.38 | 686350.31 | 897683.38 | 4 | 1 | 4 | 3 | 4 | 3 | 0.378824357 | 0.425585028 | 0.435767652 | 0.489557192 |
| Q8BFZ9 | Erlin2 | Erlin-2 | 342.08 | 20.59 | 37873 | 6 | 1728519.5 | 1841829 | 1739276 | 2221873.5 | 1800745.12 | 2264106.5 | 2847378 | 1488473.75 | 1977487.5 | 2851249 | 3116842.25 | 2314407.5 | 3461254.5 | 2139068 | 2436342 | 1977631.5 | 2420823.5 | 2092903 | 2153617.5 | 1837317 | 1 | 5 | 7 | 10 | 7 | 10 | 1.443159386 | 1.178429781 | 1.123233971 | 0.917190697 |
| Q3UVK0 | Ermp1 | Endoplasmic reticulum metallopeptidase 1 | 206.72 | 3.34 | 100148 | 3 | 161155.33 | 2 | 505708.59 | 2 | 2 | 2 | 720145.38 | 147366.53 | 484164.84 | 812815.81 | 1083981.5 | 995245.62 | 1707930.25 | 1775645.5 | 1830113.75 | 865565.75 | 158379.88 | 743712.69 | 597387.44 | 540757.5 | 0 | 2 | 0 | 8 | 0 | 8 | 11.08599503 | 3.415539478 | 4.357376413 | 1.342485823 |
| Q8R180 | Ero1a | ERO1-like protein alpha | 1431.74 | 51.08 | 54084 | 24 | 9343198 | 11848098 | 5633499 | 7986201.5 | 9563434 | 10706812 | 11630947 | 8544218 | 11171371 | 7705960 | 12178094 | 14558365 | 14588018 | 8101916 | 16403024 | 11504066 | 8890058 | 12459617 | 7930224.5 | 9310266 | 3 | 11 | 4 | 11 | 4 | 11 | 1.483498859 | 1.322956843 | 1.128898578 | 1.006730871 |
| Q8R2E9 | Ero1b | ERO1-like protein beta | 511.92 | 20.34 | 53518 | 9 | 334002.38 | 178092.41 | 2 | 553115.06 | 685351.75 | 335402.03 | 186964.03 | 499835.78 | 713461.81 | 423474.69 | 526690.31 | 336254.88 | 681641.44 | 691303.25 | 120321.91 | 375880.12 | 753472 | 773353.62 | 473072.22 | 598010 | 9 | 5 | 11 | 6 | 11 | 6 | 1.345973257 | 1.091274119 | 1.698760308 | 1.377303114 |
| P57759 | Erp29 | Endoplasmic reticulum resident protein 29 | 525.23 | 43.89 | 28823 | 9 | 6102849 | 8488076 | 8270290 | 5082312.5 | 7252702.5 | 4310599.5 | 5652218.5 | 4488450 | 6211205 | 5150833.5 | 12503332 | 12514592 | 8984732 | 7726597.5 | 8493772 | 5623980 | 4721745 | 3698637.5 | 4587794 | 3901948 | 2 | 1 | 0 | 7 | 2 | 7 | 1.426943326 | 1.945625428 | 0.640241995 | 0.872964666 |
| Q9D1Q6 | Erp44 | Endoplasmic reticulum resident protein 44 | 209.65 | 14.04 | 46853 | 5 | 743651.56 | 1132591 | 1018326.06 | 1098364.62 | 5861482.5 | 1506237.5 | 1262323.38 | 2496077 | 1610554.25 | 2 | 2988105 | 3615954.5 | 697417.31 | 3435912.75 | 7232588.5 | 2100106.5 | 1426366 | 1762723.75 | 2886158 | 1772545.88 | 8 | 5 | 4 | 6 | 8 | 6 | 1.823545762 | 2.613741186 | 1.009486548 | 1.446926435 |
| Q9R0P3 | Esd | S-formylglutathione hydrolase | 1127.09 | 71.63 | 31320 | 17 | 65296098 | 76205842 | 46468122 | 51129030 | 57718778 | 73324242 | 75570362 | 63368618 | 69002194 | 71205010 | 184859842 | 195729378 | 162199010 | 148743954 | 158067074 | 78615178 | 86860162 | 105089498 | 92808050 | 95206258 | 0 | 0 | 0 | 0 | 0 | 0 | 2.862358853 | 2.41041289 | 1.544984963 | 1.301042902 |
| Q3V1V3 | Esf1 | ESF1 homolog | 159.04 | 3.31 | 98048 | 4 | 3474920 | 1369858.12 | 887191 | 938334.19 | 105750.74 | 1045037.25 | 1294056.25 | 962155.38 | 2409629.25 | 2699111.25 | 1252299.12 | 739553.75 | 749757.56 | 531608.75 | 538057.94 | 1197939.5 | 1397631.12 | 618322.75 | 900848.94 | 712479.19 | 7 | 11 | 2 | 5 | 7 | 11 | 0.562462621 | 0.453184534 | 0.712394185 | 0.573986634 |
| Q3U7R1 | Esyt1 | Extended synaptotagmin-1 | 708.63 | 17.67 | 121554 | 12 | 9560872 | 8811495 | 5688600.5 | 6542616.5 | 5472162 | 8356517 | 9430540 | 6008448 | 7311805.5 | 8862323 | 24161692 | 17831466 | 15742915 | 13596386 | 15048117 | 7582002.5 | 8500848 | 8491353 | 9051297 | 7323864 | 0 | 9 | 0 | 11 | 0 | 11 | 2.394422447 | 2.161155068 | 1.135094046 | 1.024511884 |
| Q3TZZ7 | Esyt2 | Extended synaptotagmin-2 | 128.63 | 4.5 | 94139 | 2 | 765410.88 | 936944.62 | 683521.81 | 743574.25 | 587209.62 | 784736.44 | 861169.12 | 595385.19 | 678289.12 | 694121.25 | 727619 | 906901.44 | 887414.75 | 873353.25 | 624123.88 | 588383.44 | 499751.5 | 756551.25 | 913992.56 | 678618.12 | 10 | 9 | 6 | 10 | 10 | 10 | 1.081457826 | 1.11227027 | 0.924834604 | 0.951184604 |
| Q5U2Q7 | Etf1 | Eukaryotic peptide chain release factor subunit 1 | 569.69 | 26.32 | 49031 | 8 | 14656244 | 13094521 | 13011825 | 11538765 | 11515465 | 14184750 | 11284379 | 11254099 | 11758474 | 11377132 | 14581926 | 16804494 | 10663666 | 16646267 | 11082548 | 14796204 | 20441950 | 12839705 | 9387451 | 10893315 | 11 | 12 | 10 | 11 | 11 | 12 | 1.093424915 | 1.165724361 | 1.071169403 | 1.14199727 |
| Q99LC5 | Etfa | Electron transfer flavoprotein subunit alpha, mitochondrial | 1169.67 | 66.07 | 35009 | 18 | 29268418 | 33642562 | 38704126 | 34139986 | 39466202 | 31783694 | 31156590 | 35219858 | 39502510 | 36962462 | 27348888 | 27622364 | 36280666 | 38791338 | 41312154 | 34745222 | 31334146 | 36413122 | 38176874 | 43791254 | 12 | 10 | 12 | 10 | 12 | 10 | 0.977937134 | 0.981275866 | 1.052729459 | 1.056323537 |
| Q9DCW4 | Etfb | Electron transfer flavoprotein subunit beta | 850.34 | 50.2 | 27623 | 14 | 2698308.5 | 2494337 | 3455976 | 3658862.25 | 3804868.75 | 1481980.38 | 2777067 | 2172282.5 | 1482692.5 | 1708794 | 2586553.25 | 2407901.75 | 4131574 | 3398074.25 | 3317246 | 3460985 | 2476880.75 | 4171511 | 2571631 | 4014003.75 | 10 | 11 | 2 | 2 | 10 | 11 | 0.983180405 | 1.646227947 | 1.036162255 | 1.734940255 |
| Q921G7 | Etfdh | Electron transfer flavoprotein-ubiquinone oxidoreductase, mitochondrial | 181.61 | 6.66 | 68091 | 4 | 2748641.25 | 4834111 | 5682282 | 1785930.88 | 3444230 | 3794879 | 5265131 | 3312131.5 | 3828473.75 | 3675330 | 3478872.5 | 2442373.5 | 4120526 | 6959800 | 4036128.5 | 4810143.5 | 4781353.5 | 6436035 | 3697743.5 | 3713538 | 10 | 8 | 11 | 8 | 11 | 8 | 1.137468426 | 1.058450314 | 1.267292036 | 1.179255286 |
| Q9DCM0 | Ethe1 | Persulfide dioxygenase ETHE1, mitochondrial | 96.94 | 9.06 | 27739 | 2 | 3020438 | 1731596.5 | 1231122.25 | 947761.19 | 1067297.62 | 2805768.5 | 1396086.25 | 408427.53 | 1250503.38 | 1415078.5 | 2201612 | 2306142.5 | 1694395.25 | 1678644.75 | 871545.69 | 1572759.62 | 448840.09 | 1110562.25 | 949757.88 | 458056.41 | 11 | 6 | 8 | 8 | 11 | 8 | 1.09428661 | 1.202927927 | 0.567623643 | 0.623977599 |
| P20934 | Evi2a | Protein EVI2A | 165.69 | 9.42 | 24087 | 2 | 119787.56 | 429897.19 | 103185.62 | 308999.88 | 2 | 455494 | 131082.31 | 646475.25 | 174416.25 | 567103.06 | 1167299.75 | 1300729.25 | 1863121.12 | 2205133.25 | 1681069.88 | 2 | 2 | 2 | 363274.22 | 2 | 0 | 6 | 0 | 2 | 0 | 6 | 8.543081735 | 4.161589424 | 0.3776824 | 0.18398034 |
| Q61545 | Ewsr1 | RNA-binding protein EWS | 315.63 | 10.38 | 68462 | 5 | 7554345 | 8367296.5 | 10341932 | 10419818 | 7753610 | 7825319.5 | 8322902 | 9293542 | 8896522 | 9594298 | 6154129 | 6936670.5 | 8051136 | 9076508 | 7791989 | 8924164 | 9928575 | 11577642 | 11047064 | 12792944 | 7 | 4 | 4 | 2 | 7 | 4 | 0.855377978 | 0.865199118 | 1.221288277 | 1.235310666 |
| Q8VEG4 | Exd2 | Exonuclease 3'-5' domain-containing protein 2 | 49.04 | 2 | 74332 | 1 | 222700.02 | 182885.28 | 225938 | 234069.62 | 149773.81 | 264727.19 | 165069.14 | 185012.58 | 225833.27 | 197412.7 | 2 | 137431.47 | 2 | 2 | 101095.98 | 199459.58 | 177371.64 | 344801.78 | 2 | 124948.03 | 0 | 8 | 0 | 9 | 0 | 9 | 0.234923445 | 0.229788862 | 0.833770701 | 0.815547469 |
| Q9D4H1 | Exoc2 | Exocyst complex component 2 | 136.02 | 3.25 | 103959 | 3 | 1244137.75 | 1130721.5 | 941944.31 | 1113141.62 | 1246965 | 1172813 | 1149914.5 | 1172264.38 | 1318192.75 | 180331.39 | 1185845.5 | 1182448.25 | 1239825.88 | 898651.44 | 973462.88 | 1180276.12 | 1219564.12 | 682746.62 | 1152382.75 | 837524.81 | 10 | 9 | 11 | 12 | 11 | 12 | 0.965355057 | 1.097469985 | 0.893530857 | 1.01581619 |
| O35382 | Exoc4 | Exocyst complex component 4 | 222.2 | 6.36 | 110545 | 4 | 3135023.25 | 1035354.75 | 2545278.75 | 2034473.5 | 3854070.5 | 2391853.75 | 3733664 | 1469199.38 | 3103638.75 | 3275448.25 | 631066.06 | 1607231.25 | 1634081.25 | 1728335 | 2180893.75 | 1403569.75 | 899838.81 | 1915578 | 2452006 | 2106854 | 5 | 6 | 4 | 4 | 5 | 6 | 0.617382051 | 0.556871074 | 0.696422307 | 0.62816442 |
| O54924 | Exoc8 | Exocyst complex component 8 | 76.12 | 3.35 | 81043 | 2 | 1100021.62 | 819506.5 | 817617.5 | 991021.44 | 900820.56 | 1090983.75 | 828897.94 | 818560.44 | 898674.38 | 933237.56 | 671809.69 | 2 | 586981.44 | 757944.44 | 532290.06 | 858672.44 | 1469939.88 | 2 | 992163.44 | 872366.88 | 0 | 12 | 0 | 12 | 0 | 12 | 0.550666331 | 0.557730887 | 0.905844859 | 0.917466038 |
| Q6PGF7 | Exoc8 | Exocyst complex component 8 | 89.28 | 5.31 | 81035 | 2 | 492014.72 | 2 | 129348.39 | 282761.59 | 2 | 508684.91 | 313458.69 | 2 | 643078.25 | 293087.06 | 233043.36 | 2 | 2 | 2 | 2 | 2 | 563109.06 | 489792.38 | 283969.19 | 303489.22 | 7 | 7 | 3 | 10.5 | 7 | 10.5 | 0.25776348 | 0.132542748 | 1.814301271 | 0.93291911 |
| Q9DAA6 | Exosc1 | Exosome complex component CSL4 | 39.81 | 5.64 | 21424 | 1 | 374710.69 | 435638.91 | 350818.19 | 467606.12 | 351473.19 | 443275.66 | 2 | 2 | 396776.75 | 414255.28 | 688142.31 | 631663.06 | 658752.69 | 566467.94 | 874838.88 | 499370.97 | 469931.16 | 411223.62 | 319517.53 | 516584.34 | 0 | 7 | 0 | 5 | 0 | 7 | 1.72698896 | 2.72648729 | 1.119369204 | 1.767206379 |
| P56960 | Exosc10 | Exosome component 10 | 47.74 | 1.35 | 100942 | 1 | 319489.91 | 346451.91 | 2 | 2 | 580267.19 | 2 | 508599 | 2 | 651874 | 308936.5 | 300113.31 | 343355.19 | 332249.84 | 2 | 125266.17 | 292564.5 | 217778 | 235307.61 | 2 | 2 | 11 | 8 | 11 | 8 | 11 | 8 | 0.883465749 | 0.749269358 | 0.598336002 | 0.507450156 |
| Q8VBV3 | Exosc2 | Exosome complex component RRP4 | 194.6 | 20.48 | 32632 | 3 | 1355925.75 | 1153806 | 1082270 | 928889.38 | 1024476.75 | 1187933.88 | 1007959 | 772161.75 | 939180.06 | 1190207 | 1908278.62 | 1079418.25 | 1533969.12 | 1132451.5 | 730309.25 | 1001795.25 | 504951.44 | 1185491 | 935281.38 | 1207898.38 | 10 | 10 | 9 | 11 | 10 | 11 | 1.151308061 | 1.25247666 | 0.871974151 | 0.948596913 |
| Q7TQK4 | Exosc3 | Exosome complex component RRP40 | 47.43 | 2.55 | 29546 | 1 | 1331918.5 | 1223750.38 | 1445939.75 | 1178195.5 | 1869278.62 | 909982.81 | 1309312 | 1751148 | 2141620 | 1266648.5 | 1166133 | 1095303.5 | 1136751.75 | 1260258 | 1143269.62 | 1677827.88 | 541615.06 | 576465.12 | 2 | 863008.5 | 2 | 4 | 5 | 3 | 5 | 4 | 0.823045505 | 0.786277661 | 0.519063074 | 0.495875012 |
| Q921I9 | Exosc4 | Exosome complex component RRP41 | 63.89 | 5.71 | 26250 | 1 | 2 | 2 | 177167.25 | 241198.77 | 361354.81 | 2 | 2 | 142994.33 | 2 | 2 | 120190.34 | 2 | 430753.03 | 240156.91 | 606094.56 | 2 | 410667.19 | 250248.12 | 2 | 162903.41 | 9 | 12 | 4 | 6 | 9 | 12 | 1.791910154 | 9.770448076 | 1.056555708 | 5.760904175 |
| Q9CRA8 | Exosc5 | Exosome complex component RRP46 | 142.82 | 13.62 | 25194 | 3 | 4972220 | 4417322 | 4709684 | 4646499 | 4535027 | 4459453 | 4270309.5 | 3294096.75 | 4256702 | 3550968 | 4256778 | 3495309.5 | 3885376 | 3939702.75 | 4618467 | 3155823 | 4759253 | 4999486.5 | 4940345 | 3877278.5 | 2 | 12 | 12 | 8 | 12 | 12 | 0.867481998 | 1.018359855 | 0.933482991 | 1.095840151 |
| Q8BTW3 | Exosc6 | Exosome complex component MTR3 | 111.5 | 11.36 | 28370 | 2 | 784200.94 | 2 | 2 | 1548933.25 | 1261206.75 | 702942.88 | 820891.69 | 2 | 1371478.38 | 1225940.5 | 1481039 | 2101587.5 | 1965669 | 1744897.25 | 2113186.5 | 2 | 2 | 1436280.75 | 1609791.75 | 1646646.12 | 1 | 9 | 0 | 9 | 1 | 9 | 2.61699403 | 2.282406263 | 1.305584939 | 1.13866337 |
| Q9D0M0 | Exosc7 | Exosome complex exonuclease RRP42 | 56.24 | 3.44 | 31825 | 1 | 2 | 1093149.62 | 1230564 | 1263445.75 | 876985.62 | 873281.94 | 687960.56 | 898532 | 1136048.5 | 951728.44 | 652744.69 | 895159.94 | 1196272.75 | 653126.75 | 1223286.75 | 717737.88 | 482341.19 | 2 | 847099.5 | 912309.69 | 10 | 5.5 | 12 | 5 | 12 | 5.5 | 1.03504452 | 1.016061267 | 0.662946419 | 0.650787638 |
| Q9D753 | Exosc8 | Exosome complex component RRP43 | 226.9 | 24.28 | 29949 | 4 | 2457807 | 4149332.25 | 3106492 | 4693998.5 | 2920708.5 | 3535860.5 | 5184467 | 4839099.5 | 3074162.5 | 2825014.5 | 1414173.12 | 2709830.25 | 2067312.25 | 1788193.25 | 3333128 | 4699421 | 5095370.5 | 4093217 | 4960326 | 4603003.5 | 4 | 3 | 2 | 8 | 4 | 8 | 0.652840261 | 0.581369397 | 1.353351814 | 1.205191184 |
| Q4QR75 | Exosc9 | Exosome complex component RRP45 | 169.08 | 6.64 | 48882 | 3 | 3010553 | 2531281.75 | 3416912.5 | 3671124.5 | 2985614.5 | 3744560.5 | 2579763.75 | 2390352.75 | 3174093.75 | 3009663.5 | 349630.75 | 2775358.5 | 3013266.5 | 2400031 | 2160635.25 | 4050316 | 2365154.25 | 3271808 | 2527833.5 | 2168443 | 4 | 8 | 6 | 10 | 6 | 10 | 0.685148181 | 0.71812392 | 0.921108348 | 0.965440697 |
| P97480 | Eya3 | Eyes absent homolog 3 | 82.62 | 3.92 | 55973 | 1 | 2 | 123240.95 | 143349.48 | 2 | 214379.3 | 2 | 246482.67 | 135494.61 | 184037.22 | 2 | 2 | 2 | 2 | 2 | 2 | 151322.11 | 2 | 2 | 2 | 2 | 5 | 8 | 5 | 7 | 5 | 8 | 2.079116E-05 | 1.766727E-05 | 0.314632797 | 0.267358947 |
| P26040 | Ezr | Ezrin | 952.83 | 22.01 | 69407 | 16 | 516538.62 | 167470.72 | 528685.94 | 207979 | 1529681.5 | 2 | 1868561.25 | 904027.56 | 2066017.25 | 867004.81 | 2 | 1717982.75 | 2 | 2 | 814227.88 | 1411009.12 | 829467.88 | 2 | 1353396 | 261661.41 | 9 | 11 | 5.5 | 8.5 | 9 | 11 | 0.858275008 | 0.443811504 | 1.30680389 | 0.675744481 |
| P04117 | Fabp4 | Fatty acid-binding protein, adipocyte | 392.87 | 53.79 | 14650 | 6 | 3712428.25 | 2877863 | 2 | 2046855 | 1112137.38 | 5807552 | 1698319.62 | 2265547.5 | 3326754 | 4107701.75 | 4041946.75 | 1927676.75 | 4303404.5 | 3237206.75 | 3981523 | 4768674.5 | 7796943.5 | 5605300 | 6866908.5 | 6254466.5 | 4 | 0 | 12 | 2 | 12 | 2 | 1.794157892 | 1.016615422 | 3.209701119 | 1.818698162 |
| Q05816 | Fabp5 | Fatty acid-binding protein 5 | 928.7 | 64.44 | 15137 | 13 | 191853570 | 210988242 | 56844282 | 58440734 | 95613362 | 130072682 | 208743970 | 106004130 | 100556994 | 78155066 | 113326226 | 147489522 | 58975490 | 32803920 | 63840962 | 208814722 | 101468146 | 83156146 | 88210810 | 41588922 | 10 | 11 | 7 | 9 | 10 | 11 | 0.678521835 | 0.667865575 | 0.852541115 | 0.839151863 |
| Q61160 | Fadd | FAS-associated death domain protein | 115.93 | 17.56 | 22960 | 2 | 436401.88 | 479072.94 | 503734.28 | 495696.88 | 426399.5 | 2 | 374201.16 | 596738.69 | 623616.62 | 582899.25 | 2 | 346383.44 | 231784.34 | 284386.38 | 288653.81 | 468299.06 | 2 | 189455.33 | 394570.22 | 311929.84 | 0 | 2 | 4.5 | 6.5 | 4.5 | 6.5 | 0.491695757 | 0.528694523 | 0.582690495 | 0.626536367 |
| Q920L1 | Fads1 | Fatty acid desaturase 1 | 44.94 | 4.03 | 52323 | 1 | 2 | 2 | 2 | 2 | 145702.39 | 2 | 2 | 2 | 148912.25 | 2 | 2 | 2 | 2 | 2 | 2 | 2 | 2 | 2 | 2 | 2 | 7.5 | 7.5 | 10 | 10 | 10 | 10 | 5.094547E-05 | 6.715004E-05 | 5.094547E-05 | 6.715004E-05 |
| P54731 | Faf1 | FAS-associated factor 1 | 81.01 | 3.7 | 73863 | 2 | 340468.84 | 420659.66 | 203571.56 | 2 | 358664.56 | 182830.88 | 338651.94 | 352013 | 336611.12 | 119243.5 | 352869.03 | 398453.22 | 403587.75 | 368619.19 | 358714 | 243774.25 | 2 | 118284.84 | 119141 | 131690.03 | 6 | 5.5 | 0 | 3 | 6 | 5.5 | 1.422314241 | 1.415911962 | 0.463131011 | 0.461046314 |
| Q3TDN2 | Faf2 | FAS-associated factor 2 | 246.44 | 10.56 | 52471 | 4 | 3293456.75 | 3358714.5 | 2055496.38 | 2579102.5 | 2287342.75 | 4323803.5 | 3314669.75 | 3225560.75 | 2152926.25 | 3576681.25 | 3542725.75 | 3760342.5 | 4763339 | 2474971.5 | 3828731.25 | 2669925.5 | 2418138 | 1842752.25 | 2844320.75 | 2158089 | 3 | 9 | 8 | 4 | 8 | 9 | 1.353319378 | 1.107057182 | 0.879116419 | 0.719144469 |
| P35505 | Fah | Fumarylacetoacetase | 136.15 | 8.83 | 46176 | 3 | 2168883.25 | 3525974.25 | 3840234.5 | 3910919 | 3219120 | 3889039 | 3035370 | 3492837.25 | 3775399.25 | 4430304.5 | 6703918.5 | 6432458 | 4880465 | 5707814.5 | 5494775 | 3935399 | 3314026 | 5310956.5 | 3996080.5 | 2235877 | 0 | 7 | 0 | 11 | 0 | 11 | 1.753327412 | 1.569001206 | 1.127644241 | 1.009095713 |
| Q9WUD8 | Faim | Fas apoptotic inhibitory molecule 1 | 38.93 | 4.47 | 20201 | 1 | 1049024.88 | 874190.88 | 986749.31 | 900874.5 | 984936.31 | 831822.94 | 1004101.38 | 901134.12 | 2 | 1375180.38 | 372012.66 | 780776.62 | 642710.69 | 2 | 786242.56 | 954338.19 | 674268.94 | 870523.94 | 756216.88 | 898727.56 | 0 | 3 | 4.5 | 9 | 4.5 | 9 | 0.538337194 | 0.627819392 | 0.866194671 | 1.01017321 |
| Q8VE88 | Fam114a2 | Protein FAM114A2 | 99.94 | 7.65 | 54045 | 2 | 1009979.75 | 1037351.75 | 815890.31 | 317230.69 | 1080864.5 | 548480.75 | 1123763.12 | 1016401.25 | 1097146.62 | 893140.25 | 259426.77 | 638725.88 | 219288.16 | 2 | 314136.81 | 1018743.62 | 279006.47 | 356725.5 | 347987.53 | 549989.75 | 1 | 6 | 1 | 4 | 1 | 6 | 0.335947694 | 0.305962904 | 0.598982162 | 0.545520404 |
| Q6A0A9 | FAM120A | Constitutive coactivator of PPAR-gamma-like protein 1 | 201.14 | 5.04 | 121646 | 4 | 1714103 | 1443574.5 | 1938383.75 | 2204931.5 | 1090596.75 | 1356811.38 | 1483470.5 | 1849403.75 | 577661.38 | 1653859.75 | 1091461.5 | 1626401.62 | 523039.19 | 1136253 | 371845.5 | 1830859.5 | 2544234.25 | 1690182.12 | 211196.7 | 2183387.5 | 4 | 11 | 5 | 7 | 5 | 11 | 0.565923871 | 0.686152137 | 1.008135595 | 1.222309976 |
| Q3UW53 | Fam129a | Protein Niban | 1130 | 23.97 | 102649 | 20 | 27753622 | 29459116 | 45440442 | 45648230 | 42554526 | 29725276 | 32800302 | 38268090 | 37539042 | 37875650 | 29376072 | 32437684 | 35972790 | 40694230 | 34342654 | 28710122 | 27512178 | 33660474 | 38671218 | 37471302 | 9 | 7 | 10 | 9 | 10 | 9 | 0.905517709 | 0.980790185 | 0.869898508 | 0.942210086 |
| Q8R1F1 | Fam129b | Niban-like protein 1 | 817.84 | 25.1 | 84819 | 14 | 22757864 | 24378526 | 13081068 | 19918774 | 20886366 | 23740672 | 22287242 | 23910618 | 26164402 | 24780830 | 17352278 | 19705594 | 19564168 | 14600426 | 17031170 | 24088632 | 21411562 | 27586916 | 27939216 | 30478582 | 5 | 3 | 0 | 7 | 5 | 7 | 0.873602914 | 0.730070219 | 1.301737538 | 1.087862453 |
| Q4QQV3 | Fam162a | Protein FAM162A | 67.82 | 15.48 | 17827 | 2 | 139769.56 | 405552.72 | 412854.28 | 505323.53 | 141133.23 | 335212.5 | 292857.47 | 466097.34 | 461709.72 | 749852.25 | 1208003.25 | 1038436.88 | 1530561.25 | 1643258.62 | 1716172.25 | 312297.22 | 298463.75 | 578279 | 392708.16 | 2 | 0 | 11 | 0 | 8 | 0 | 11 | 4.447391289 | 3.095086796 | 0.985739303 | 0.686008606 |
| Q501J2 | Fam173a | Protein N-lysine methyltransferase FAM173A | 46.41 | 4.37 | 24740 | 1 | 311884.78 | 2 | 148855.58 | 2 | 120626.86 | 270232.56 | 2 | 2 | 157802.17 | 146983.77 | 886694.75 | 509860.28 | 549598.94 | 590867.19 | 2 | 2 | 2 | 2 | 2 | 160784.08 | 4 | 8 | 4 | 8 | 4 | 8 | 4.363860942 | 4.412041546 | 0.276573856 | 0.279627458 |
| Q91WE2 | Fam192a | PSME3-interacting protein | 52.9 | 6.69 | 28702 | 2 | 3724813.75 | 3188321 | 4955081 | 4370472 | 5392265.5 | 2394564 | 3562254.25 | 4849926 | 4759753 | 4310643 | 295968.12 | 3639295 | 2534063.25 | 1495064.25 | 3784767.25 | 887721.81 | 1610604.75 | 3175485.25 | 4125459 | 3690795.5 | 3 | 3 | 5 | 5 | 5 | 5 | 0.543164128 | 0.591088946 | 0.623646409 | 0.678672392 |
| Q5MJS3 | Fam20c | Extracellular serine/threonine protein kinase FAM20C | 39.15 | 1.21 | 65766 | 1 | 927201.44 | 1300365.12 | 946487.69 | 1008050.25 | 1258117.25 | 902702.94 | 1306456.12 | 1788612.75 | 1975976.75 | 1802999.38 | 1798020.25 | 2634195.75 | 2284483 | 1846741.75 | 2886629 | 862346.69 | 856929.88 | 1566673.25 | 2097341.25 | 1625976.88 | 0 | 10 | 3 | 9 | 3 | 10 | 2.10470644 | 1.472346775 | 1.288415854 | 0.901310934 |
| Q9D8B6 | Fam210b | Protein FAM210B, mitochondrial | 52.75 | 6.84 | 20344 | 1 | 2 | 2 | 2 | 2 | 2 | 2 | 2 | 2 | 2 | 2 | 541837.56 | 218933.42 | 304704.22 | 247822.86 | 242151.64 | 2 | 2 | 2 | 2 | 2 | 0 | 12.5 | 0 | 12.5 | 0 | 12.5 | 155544.97 | 155544.97 | 1 | 1 |
| Q91VU0 | Fam3c | Protein FAM3C | 166 | 17.18 | 24753 | 3 | 2885714 | 2997622 | 3390319.5 | 1745503.88 | 2139384.5 | 2825196.25 | 2791998 | 3259756.5 | 2397497.25 | 3235518.75 | 2463396 | 2637331 | 3356495.5 | 3790351.5 | 3853885 | 2032673.38 | 3125627.25 | 3589505.25 | 2994778.5 | 2956610 | 7 | 9 | 8 | 11 | 8 | 11 | 1.223650515 | 1.109682694 | 1.117083662 | 1.013041217 |
| Q9D8N2 | Fam45a | Protein FAM45A | 66.99 | 3.08 | 40421 | 1 | 985810.88 | 755390 | 670868 | 1088930 | 1080205.62 | 815665.38 | 877998.25 | 2 | 2 | 2 | 2 | 617114.56 | 2 | 796950.25 | 2 | 2 | 804021 | 903304.25 | 2 | 1114480.88 | 2 | 9 | 10.5 | 9 | 10.5 | 9 | 0.308667908 | 0.834915372 | 0.615953758 | 1.666092419 |
| Q8BHZ0 | Fam49a | Protein FAM49A | 175.38 | 11.46 | 37343 | 3 | 2 | 486893.59 | 701138.19 | 2 | 783933.62 | 2 | 2 | 505449.56 | 697751.88 | 355635.72 | 2 | 165464.81 | 2 | 2 | 2 | 2 | 359451.66 | 2 | 2 | 2 | 6 | 6 | 6 | 7 | 6 | 7 | 0.083912463 | 0.106151168 | 0.182284603 | 0.230594155 |
| Q921M7 | Fam49b | CYFIP-related Rac1 interactor B | 689.92 | 38.89 | 36776 | 10 | 12861141 | 14445928 | 17839438 | 17254798 | 18573588 | 13732099 | 15099142 | 15022374 | 16201185 | 13642780 | 8373122 | 10964909 | 11621341 | 9739654 | 7291179 | 15004552 | 14686504 | 14977756 | 10381945 | 14345370 | 0 | 7 | 0 | 8 | 0 | 8 | 0.592655368 | 0.651177488 | 0.857007949 | 0.941633728 |
| Q80VD1 | Fam98b | Protein FAM98B | 92.21 | 6.29 | 45349 | 2 | 2 | 1540346 | 807066.69 | 2 | 2795332.5 | 2 | 1548263.25 | 1106479.12 | 2847251.5 | 2 | 463732.31 | 1595423 | 408948.44 | 2 | 1375082.5 | 852870.06 | 2 | 653999.06 | 696365 | 954038.06 | 12 | 11 | 12 | 9 | 12 | 11 | 0.747302291 | 0.698507768 | 0.613927311 | 0.573841404 |
| Q922J9 | Far1 | Fatty acyl-CoA reductase 1 | 1092.32 | 49.51 | 59435 | 19 | 16641804 | 10686342 | 10341209 | 13125131 | 12728094 | 16481948 | 14269302 | 13616062 | 11013636 | 11615346 | 25758894 | 29708074 | 31270736 | 29192340 | 32006064 | 15893355 | 16773403 | 14739104 | 14015383 | 13724871 | 0 | 4 | 0 | 6 | 0 | 6 | 2.328874362 | 2.208123751 | 1.182982744 | 1.121645863 |
| Q8C0C7 | Farsa | Phenylalanine--tRNA ligase alpha subunit | 361.26 | 10.83 | 57599 | 6 | 6220320.5 | 3151881.25 | 5886567.5 | 4777731 | 5214309 | 9068458 | 5078745.5 | 3233915 | 709975.88 | 2626928 | 4550385 | 7046714 | 5915142 | 4840465 | 3112960.25 | 3720447.25 | 6101020.5 | 7134325 | 1231256.38 | 2265810.75 | 12 | 10 | 9 | 12 | 12 | 12 | 1.008508915 | 1.229155263 | 0.809988293 | 0.98720136 |
| Q9WUA2 | Farsb | Phenylalanine--tRNA ligase beta subunit | 656.07 | 23.94 | 65697 | 14 | 21526610 | 23939552 | 25638974 | 24736648 | 22805866 | 25088838 | 19701588 | 20885812 | 22240146 | 25957974 | 30056540 | 29085762 | 28121212 | 30596406 | 28850276 | 21601754 | 28802046 | 28916514 | 28906888 | 27575658 | 0 | 4 | 0 | 3 | 0 | 4 | 1.236520032 | 1.288351465 | 1.144589547 | 1.192567514 |
| P19096 | Fasn | Fatty acid synthase | 4937.68 | 38.86 | 272428 | 77 | 107550322 | 118319858 | 97874042 | 107639506 | 90715490 | 109689138 | 103837714 | 87221586 | 77386050 | 87895890 | 131663682 | 111603674 | 94461834 | 87466802 | 82958570 | 145836946 | 133732866 | 103817090 | 94996650 | 108801506 | 10 | 8 | 9 | 5 | 10 | 8 | 0.973291176 | 1.090389352 | 1.124661822 | 1.259971636 |
| P62862 | Fau | 40S ribosomal protein S30 | 105.29 | 18.64 | 6648 | 2 | 11825293 | 9815767 | 9039790 | 8051095.5 | 6598297 | 11625420 | 11015620 | 6550601 | 7970864.5 | 10400472 | 9222162 | 6287528.5 | 6444130 | 5507620 | 5054627 | 8586045 | 10317706 | 6947158 | 5651602 | 6522821 | 3 | 7 | 2 | 5 | 3 | 7 | 0.717315102 | 0.683642388 | 0.838851281 | 0.79947333 |
| P22509 | Fbl | rRNA 2'-O-methyltransferase fibrillarin | 960.05 | 63.3 | 34222 | 16 | 21404730 | 27398548 | 25358232 | 28009130 | 22077558 | 24209648 | 22411264 | 24153646 | 22849104 | 21489198 | 27786556 | 32750108 | 30876884 | 35032110 | 30618022 | 26485854 | 27545930 | 28453998 | 27057136 | 31108998 | 1 | 5 | 0 | 0 | 1 | 5 | 1.264112337 | 1.364432089 | 1.13202379 | 1.221861015 |
| Q9CZV8 | Fbxl20 | F-box/LRR-repeat protein 20 | 39.45 | 1.83 | 48396 | 1 | 416583.5 | 473573.5 | 411313.62 | 628199.38 | 519762 | 337046.28 | 339828.78 | 324579.56 | 499710.34 | 346274.69 | 2 | 458946.47 | 535308.88 | 2 | 446758.69 | 512054.69 | 529999 | 386230.06 | 588527.38 | 418636.06 | 8 | 12 | 12 | 2 | 12 | 12 | 0.58830702 | 0.780008181 | 0.994290591 | 1.318282408 |
| Q78JE5 | Fbxo22 | F-box only protein 22 | 100.97 | 5.72 | 44203 | 2 | 1744519.75 | 1228619.25 | 1186829.62 | 1305072.12 | 926742.94 | 1558242.5 | 1126986.25 | 1423077.75 | 953394.38 | 1035274.38 | 1242054.62 | 1361490.88 | 1166886 | 1127633 | 992140.31 | 1074102.62 | 821160.81 | 1179965.62 | 1309753.75 | 1071576.88 | 10 | 7 | 12 | 10 | 12 | 10 | 0.921527559 | 0.966086389 | 0.853683409 | 0.894961755 |
| P20491 | Fcer1g | High affinity immunoglobulin epsilon receptor subunit gamma | 172.4 | 25.58 | 9652 | 3 | 7293618 | 10586340 | 7397920 | 8877770 | 3926941.75 | 10143556 | 8692348 | 10305551 | 6647899 | 10310799 | 38111330 | 33805394 | 41004698 | 33926402 | 44153050 | 6364498 | 8049496.5 | 12716928 | 10700599 | 12268295 | 0 | 6 | 0 | 9 | 0 | 9 | 5.015438164 | 4.143172236 | 1.315556973 | 1.086760309 |
| P26151 | Fcgr1 | High affinity immunoglobulin gamma Fc receptor I | 328.69 | 19.31 | 44888 | 7 | 10358322 | 6828544 | 4298951.5 | 6643145 | 6816944 | 11241287 | 8880291 | 5772027.5 | 7733214 | 8040725 | 7087239.5 | 5918700 | 6640044 | 4677350 | 6716819.5 | 7896921 | 9852431 | 8755662 | 5049418 | 7269359.5 | 9 | 8 | 4 | 10 | 9 | 10 | 0.888234306 | 0.744947977 | 1.110968219 | 0.931751366 |
| P08101 | Fcgr2 | Low affinity immunoglobulin gamma Fc region receptor II | 334.24 | 29.7 | 36695 | 6 | 2 | 1566742.75 | 1805839.25 | 651330.38 | 1088704.75 | 2 | 950864.69 | 529951.75 | 508079.88 | 140311.62 | 1274329.38 | 2832895.25 | 2282229.75 | 1992983.12 | 2232350 | 2009672.25 | 837369.56 | 1357900 | 725059.12 | 1491630.75 | 2 | 10 | 0 | 2 | 2 | 10 | 2.076193675 | 4.985317465 | 1.256035609 | 3.015969238 |
| P08508 | Fcgr3 | Low affinity immunoglobulin gamma Fc region receptor III | 91.83 | 8.05 | 30036 | 2 | 844036.62 | 867694 | 1226969.25 | 1141360.12 | 254584.95 | 2 | 771743.69 | 906477.38 | 907945.31 | 909634 | 2 | 416349.38 | 2 | 464971.03 | 331288.97 | 462230.62 | 848998.44 | 2 | 2 | 2 | 3 | 3 | 4 | 4.5 | 4 | 4.5 | 0.279749183 | 0.346876982 | 0.302501146 | 0.37508844 |
| A0A0B4 | Fcgr4 | Low affinity immunoglobulin gamma Fc region receptor IV | 94.91 | 9.24 | 28398 | 2 | 1649004.75 | 1917487.75 | 1929528.62 | 2847039.5 | 1669290.38 | 1579356.88 | 1295349.88 | 2 | 1008217.06 | 1693868.88 | 1942101.5 | 1858925.25 | 1960374.75 | 2 | 1438510.75 | 1322013.5 | 1936462.88 | 1863998 | 1636967.62 | 1817837.75 | 10 | 8 | 6.5 | 3 | 10 | 8 | 0.719103261 | 1.291048826 | 0.856669902 | 1.538030394 |
| D3ZYR1 | Fcho2 | F-BAR domain only protein 2 | 129.67 | 4.94 | 88712 | 3 | 1307975 | 399185.5 | 497183.09 | 2009178.88 | 1528999.25 | 1080505 | 1367427.88 | 1057453 | 1572927.25 | 1222218.75 | 530939 | 682274.5 | 942747.44 | 534359.81 | 646829.75 | 2 | 843269.38 | 759333.62 | 911355.12 | 2 | 10 | 6 | 0 | 0 | 10 | 6 | 0.5811298 | 0.529661712 | 0.437780167 | 0.39900792 |
| P53798 | Fdft1 | Squalene synthase | 72.3 | 3.37 | 48154 | 1 | 2 | 422874.5 | 282221.22 | 182876.92 | 2 | 238775.31 | 619984 | 265362.03 | 2 | 2 | 2 | 363016.44 | 2 | 2 | 2 | 535110.56 | 2 | 190282.44 | 87057.04 | 225020.62 | 8 | 11 | 8 | 11 | 8 | 11 | 0.408826665 | 0.322939469 | 1.16836896 | 0.922915464 |
| Q920E5 | Fdps | Farnesyl pyrophosphate synthase | 1259.5 | 46.74 | 40582 | 18 | 46967846 | 49762690 | 40426382 | 37639654 | 53651678 | 38857490 | 49193742 | 48109350 | 52983382 | 38715762 | 38766594 | 47789630 | 33512242 | 31230466 | 38119214 | 60291926 | 49872854 | 45849542 | 50171722 | 44168870 | 5 | 8 | 3 | 8 | 5 | 8 | 0.829151224 | 0.831292784 | 1.095893333 | 1.098723844 |
| P24483 | Fdx1 | Adrenodoxin, mitochondrial | 97.76 | 9.04 | 20135 | 2 | 805551.12 | 1089451.38 | 496347.47 | 291768 | 1028971.19 | 357153.78 | 227476.5 | 1235398.5 | 356417.5 | 1115426.12 | 1335928 | 1057570.75 | 1445007.75 | 1547390.5 | 1595789.12 | 2 | 360429 | 1114900.25 | 368067.06 | 1126759.88 | 1 | 12 | 2 | 12 | 2 | 12 | 1.88079699 | 2.12088601 | 0.800131156 | 0.902270146 |
| P39749 | Fen1 | Flap endonuclease 1 | 588.82 | 22.75 | 42315 | 9 | 18350324 | 18601704 | 25782404 | 22303604 | 22462160 | 15569011 | 15663790 | 20306910 | 16889446 | 17730516 | 12328346 | 12408609 | 8817527 | 8912347 | 12664128 | 18775342 | 12994049 | 17653564 | 11721029 | 17579514 | 0 | 2 | 0 | 10 | 0 | 10 | 0.512845177 | 0.639869551 | 0.732310274 | 0.913692996 |
| O88842 | Fgd3 | FYVE, RhoGEF and PH domain-containing protein 3 | 103.42 | 4.09 | 80624 | 2 | 257465.56 | 209773.81 | 289933.03 | 294180.38 | 687988.94 | 180611 | 190514.11 | 304908.06 | 329735.28 | 165982.56 | 52244.48 | 181715.31 | 316432 | 2 | 157659.53 | 229582.14 | 195987.78 | 99969.27 | 377311.16 | 111950.43 | 4 | 5 | 6 | 11 | 6 | 11 | 0.407081203 | 0.604269434 | 0.583439567 | 0.866054965 |
| P97807 | Fh | Fumarate hydratase, mitochondrial | 1716.07 | 50.69 | 54357 | 25 | 81698874 | 79065034 | 99973650 | 94571130 | 89467466 | 81434226 | 85056538 | 92308890 | 92726682 | 94678298 | 72491626 | 85886386 | 98722074 | 100804082 | 102974626 | 74993810 | 81696674 | 100019482 | 89219098 | 104039786 | 9 | 12 | 8 | 12 | 9 | 12 | 1.036203919 | 1.032886615 | 1.011674853 | 1.008436076 |
| Q9R059 | Fhl3 | Four and a half LIM domains protein 3 | 115 | 7.27 | 31795 | 2 | 6108417 | 8043971 | 10975038 | 10602851 | 9599254 | 7585582 | 6140138 | 11079656 | 9687932 | 10608502 | 4699449 | 5271089 | 7163241.5 | 6543798.5 | 7528317 | 6213408 | 6406752 | 10050405 | 9918200 | 11404989 | 3 | 12 | 3 | 12 | 3 | 12 | 0.688423072 | 0.69189895 | 0.970531859 | 0.975432117 |
| Q5U317 | Fip1l1 | Pre-mRNA 3'-end-processing factor FIP1 | 120.86 | 6.72 | 60188 | 2 | 287048.75 | 401842.03 | 411933.41 | 232768.22 | 286111.28 | 369010.44 | 397540.34 | 608279 | 308700.59 | 366115.81 | 334682.16 | 490848.59 | 599018.94 | 450339.72 | 549436.75 | 136411.34 | 125895.72 | 397171.47 | 362531.94 | 2 | 2 | 6 | 8 | 4 | 8 | 6 | 1.4967714 | 1.182802273 | 0.630987307 | 0.498628729 |
| P84817 | Fis1 | Mitochondrial fission 1 protein | 421.25 | 44.74 | 16995 | 6 | 2273554.75 | 3469042.5 | 4548336.5 | 3376777.5 | 4943141.5 | 2082440.38 | 3233517.5 | 4000489 | 2830745.75 | 1699995.12 | 385355.56 | 986572.75 | 778236.81 | 999233.12 | 894572.5 | 3490586.5 | 2396232.25 | 2365250.5 | 3037691 | 3024747.5 | 0 | 7 | 0 | 11 | 0 | 11 | 0.217290997 | 0.292042746 | 0.769148407 | 1.033748369 |
| Q6P9Q6 | Fkbp15 | FK506-binding protein 15 | 44.07 | 1.56 | 132961 | 1 | 400036.34 | 678246.5 | 399313.22 | 477087.44 | 320379.12 | 396883.75 | 460948.53 | 609284.5 | 408377 | 478115.53 | 310378.62 | 282808.72 | 2 | 321161.12 | 2 | 516595.5 | 656064.25 | 454333.09 | 451608 | 512084.19 | 1 | 7 | 0 | 8 | 1 | 8 | 0.401902107 | 0.388489481 | 1.138731307 | 1.100728578 |
| P26883 | Fkbp1a | Peptidyl-prolyl cis-trans isomerase FKBP1A | 143.76 | 25 | 11923 | 2 | 2 | 2 | 2 | 2 | 2 | 1336983.12 | 1785448.75 | 2 | 2 | 2 | 5570327 | 5588228.5 | 3596534.25 | 4939032.5 | 5138845.5 | 2 | 1320234.75 | 2 | 2 | 2 | 0 | 10 | 0 | 9 | 0 | 10 | 2483296.775 | 7.953070256 | 132024.275 | 0.422824346 |
| P45878 | Fkbp2 | Peptidyl-prolyl cis-trans isomerase FKBP2 | 368.69 | 50.71 | 15344 | 7 | 6430538 | 5080311 | 4120020 | 4964810.5 | 2856015.25 | 6341819 | 4867467 | 4722144 | 4243504 | 5609145.5 | 5313909.5 | 4218743.5 | 4205917 | 5662995 | 3615702.5 | 5062218 | 8114076.5 | 6165580.5 | 5183519 | 4245302.5 | 12 | 7 | 7 | 9 | 12 | 9 | 0.981475656 | 0.892693008 | 1.226806711 | 1.115831826 |
| Q62446 | Fkbp3 | Peptidyl-prolyl cis-trans isomerase FKBP3 | 37.54 | 4.91 | 25148 | 1 | 2 | 2 | 2 | 2 | 2 | 2 | 2 | 2 | 341681.88 | 2 | 430973.72 | 519518.72 | 346974.66 | 2 | 2 | 2 | 2 | 2 | 2 | 2 | 5 | 12.5 | 6 | 10 | 6 | 12.5 | 129747.11 | 3.797218402 | 1 | 2.92663E-05 |
| P30416 | Fkbp4 | Peptidyl-prolyl cis-trans isomerase FKBP4 | 1137.45 | 52.62 | 51572 | 21 | 58496002 | 58823514 | 74111858 | 67042838 | 67602466 | 54306094 | 57344730 | 73713882 | 53234890 | 65177846 | 47690214 | 58452262 | 46141450 | 50789750 | 52916850 | 71528754 | 51579778 | 67963714 | 62790490 | 62753102 | 0 | 12 | 3 | 11 | 3 | 12 | 0.785062359 | 0.842691032 | 0.970985843 | 1.042262506 |
| Q64378 | Fkbp5 | Peptidyl-prolyl cis-trans isomerase FKBP5 | 252.82 | 12.94 | 50966 | 5 | 7184186 | 6872824 | 3961716 | 7377362 | 5974424 | 7911398 | 5037068.5 | 6519845 | 5821131 | 6613109.5 | 5814555 | 7430776 | 6549724 | 7178513.5 | 7174450.5 | 6105715 | 7460308 | 5199176 | 5941596.5 | 5733834 | 11 | 10 | 9 | 10 | 11 | 10 | 1.088538784 | 1.070385184 | 0.970358071 | 0.954175374 |
| O35465 | Fkbp8 | Peptidyl-prolyl cis-trans isomerase FKBP8 | 459.64 | 17.16 | 43529 | 6 | 5714374 | 7134131 | 8565982 | 7907022 | 5871518 | 5692394.5 | 4995112.5 | 7691107 | 6567192 | 5648523.5 | 7674927.5 | 7535332.5 | 9262592 | 9401211 | 9772804 | 6129455 | 7625851 | 10691659 | 9618479 | 7132907 | 4 | 8 | 2 | 4 | 4 | 8 | 1.240213495 | 1.426632573 | 1.170639599 | 1.346600879 |
| Q9JJ28 | Flii | Protein flightless-1 homolog | 1089.44 | 20.38 | 144803 | 20 | 20183130 | 23525424 | 19545208 | 18099438 | 19328310 | 18914104 | 19354852 | 17686782 | 19642244 | 23543438 | 18201118 | 24241612 | 23044394 | 20244052 | 18924570 | 20116080 | 19735368 | 16952194 | 17119102 | 20030038 | 10 | 9 | 9 | 12 | 10 | 12 | 1.039473345 | 1.055620809 | 0.933168185 | 0.947664276 |
| Q8BTM8 | Flna | Filamin-A | 5224.061 | 41.9 | 281222 | 81 | 177941858 | 191113602 | 82794962 | 106899738 | 121273986 | 169158434 | 162421330 | 125754258 | 116163794 | 140355986 | 232483986 | 232977810 | 182502610 | 178620466 | 181363410 | 211277282 | 208845538 | 217285538 | 221223218 | 220296962 | 3 | 0 | 3 | 0 | 3 | 0 | 1.482224253 | 1.411981388 | 1.586603276 | 1.511413871 |
| Q80X90 | Flnb | Filamin-B | 409.11 | 3.42 | 277825 | 7 | 2099209.5 | 1296679 | 412703.06 | 1205954 | 894698.88 | 2061300.38 | 1625605.12 | 1129517 | 831458.12 | 2071396.5 | 2810910.5 | 2904485.75 | 2327471.75 | 1832218.38 | 2087464.75 | 1346720 | 2662957.5 | 2162883.25 | 2336921.5 | 2090743.12 | 2 | 2 | 2 | 3 | 2 | 3 | 2.024379132 | 1.549698365 | 1.793837686 | 1.373214772 |
| O08917 | Flot1 | Flotillin-1 | 333.29 | 17.99 | 47513 | 6 | 1468992.62 | 570003 | 425506.53 | 135190.97 | 2 | 1007027.25 | 2 | 2 | 349074.38 | 400111.72 | 7496442 | 4268402 | 5295726.5 | 5660567 | 5834044.5 | 1121097.62 | 1176207.75 | 495300.97 | 397656.06 | 2 | 0 | 11.5 | 0 | 7 | 0 | 11.5 | 10.98405031 | 16.25948064 | 1.227168669 | 1.816554426 |
| Q60634 | Flot2 | Flotillin-2 | 115.78 | 5.14 | 47038 | 2 | 2 | 2 | 2 | 2 | 2 | 2 | 2 | 2 | 2 | 146300.62 | 1343533.12 | 950762.62 | 865028.81 | 2 | 848529.69 | 373918.66 | 2 | 2 | 436526.18 | 279612.69 | 2.5 | 5 | 3 | 6 | 3 | 6 | 400785.624 | 27.39316549 | 109006.147 | 7.450425477 |
| Q9JL26 | Fmnl1 | Formin-like protein 1 | 1648.84 | 32.27 | 122060 | 30 | 27389408 | 26417914 | 35360242 | 34733774 | 25977560 | 26362626 | 23601688 | 32817160 | 30350640 | 35740262 | 24797728 | 24818318 | 25431280 | 25391254 | 25537180 | 24318860 | 21293684 | 27001908 | 23546434 | 25718740 | 0 | 2 | 5 | 4 | 5 | 4 | 0.840516989 | 0.846199701 | 0.813187364 | 0.818685301 |
| Q6ZPF4 | Fmnl3 | Formin-like protein 3 | 403.8 | 8.07 | 117169 | 7 | 876172.19 | 849739.44 | 838626.19 | 1381725 | 982193.19 | 1743744.5 | 1108333.5 | 969357.44 | 799533.69 | 1027266.31 | 1363481 | 622024.88 | 364932.84 | 1685317.5 | 822204.94 | 768280 | 2474789.25 | 1295713.75 | 1476643.75 | 1072564.5 | 9 | 7 | 9 | 9 | 9 | 9 | 0.985696362 | 0.860084749 | 1.438176832 | 1.254903647 |
| P35922 | Fmr1 | Synaptic functional regulator FMR1 | 156.42 | 6.03 | 68989 | 3 | 1937220.38 | 2218149 | 2005645.5 | 3007861.5 | 1934500.12 | 2441329 | 2192000.75 | 1954781.12 | 1715698 | 2069097.12 | 221751.55 | 273700.25 | 535028.88 | 524237.44 | 2 | 2098673.75 | 2006420.38 | 1607761 | 1635922.88 | 1557622.38 | 0 | 6 | 0 | 5 | 0 | 6 | 0.140022282 | 0.149882793 | 0.802134413 | 0.858621528 |
| Q80TY0 | Fnbp1 | Formin-binding protein 1 | 690.35 | 18.02 | 71344 | 12 | 23542834 | 19007664 | 24146198 | 21655404 | 19724946 | 16144865 | 20264750 | 22647414 | 16896724 | 18615546 | 10696158 | 15608129 | 10555638 | 7688807 | 12067890 | 23309314 | 11560294 | 18975616 | 16343285 | 16263637 | 0 | 3 | 0 | 10 | 0 | 10 | 0.52385427 | 0.598678668 | 0.799912185 | 0.914167144 |
| Q6NWW9 | Fndc3b | Fibronectin type III domain-containing protein 3B | 48.38 | 1.41 | 132764 | 1 | 2 | 418807.81 | 2 | 2 | 2 | 2 | 2 | 2 | 2 | 2 | 2 | 2 | 2 | 601510.44 | 2 | 2 | 2 | 2 | 2 | 2 | 12 | 10 | 10 | 12.5 | 12 | 12.5 | 1.43623623 | 60151.844 | 2.387684E-05 | 1 |
| Q8K385 | FRRS1 | Ferric-chelate reductase 1 | 209.89 | 8.45 | 66047 | 4 | 19110252 | 14441198 | 10392865 | 12435464 | 12870888 | 15825893 | 15784801 | 10422126 | 12082153 | 13871428 | 5279406 | 5748655 | 4866836 | 6404150 | 5327183 | 14310594 | 12359366 | 8705210 | 8323484 | 9933136 | 0 | 4 | 0 | 5 | 0 | 5 | 0.398930887 | 0.406349352 | 0.774458822 | 0.788860555 |
| P09528 | Fth1 | Ferritin heavy chain | 641.35 | 56.04 | 21067 | 11 | 21131550 | 18141730 | 19321128 | 34893818 | 29652728 | 27886138 | 25057054 | 55739530 | 54762290 | 55680038 | 171254306 | 172532530 | 283515874 | 253427186 | 282415842 | 384739938 | 554799298 | 602124482 | 627704770 | 567048514 | 0 | 0 | 0 | 0 | 0 | 0 | 9.445645013 | 5.308136783 | 22.22182721 | 12.48792414 |
| P29391 | Ftl1 | Ferritin light chain 1 | 1017.49 | 73.22 | 20802 | 15 | 35454822 | 27618676 | 45772886 | 53905598 | 52433102 | 40681886 | 38502082 | 74889490 | 81044098 | 82448042 | 105100170 | 116605522 | 150582738 | 154706914 | 153572834 | 551825986 | 664956482 | 658574466 | 716652034 | 665532546 | 0 | 0 | 0 | 0 | 0 | 0 | 3.162710748 | 2.143079043 | 15.13832396 | 10.25785392 |
| Q9DBE9 | Ftsj3 | pre-rRNA processing protein FTSJ3 | 425.06 | 13.6 | 95532 | 7 | 612300.38 | 1046182.12 | 2 | 966833.31 | 431477.72 | 2 | 2 | 614672.19 | 2 | 2 | 2 | 791690.56 | 2 | 2 | 521866.91 | 466736.88 | 2 | 406095.78 | 501090.94 | 2 | 6.5 | 6 | 10 | 9 | 10 | 9 | 0.429719115 | 2.136986829 | 0.449466635 | 2.235190954 |
| Q32PX7 | Fubp1 | Far upstream element-binding protein 1 | 637.6 | 20.19 | 67197 | 12 | 6748806.5 | 6964945.5 | 6967515 | 9365283 | 5449606.5 | 6731300.5 | 6042149 | 6249227.5 | 7416667 | 8898999 | 7332153.5 | 6596858 | 9215208 | 9276888 | 9044679 | 7257362 | 11252148 | 8585447 | 10588044 | 9576878 | 8 | 2 | 5 | 3 | 8 | 3 | 1.1681768 | 1.173393628 | 1.331408346 | 1.337354131 |
| Q99LJ1 | Fuca1 | Tissue alpha-L-fucosidase | 88.52 | 4.87 | 52281 | 2 | 1142104.62 | 1136052.12 | 1263163.75 | 1478844.62 | 1618179.75 | 1292101.62 | 1367919.75 | 1741055.12 | 1895272.75 | 1377902.75 | 540281.25 | 810616.12 | 1355626.38 | 587712.31 | 1017279.88 | 1075387.5 | 1152969.5 | 1603368 | 1199404.38 | 1250965.62 | 3 | 10 | 1 | 3 | 3 | 10 | 0.649486586 | 0.561815789 | 0.946334536 | 0.818593787 |
| Q9D6K8 | Fundc2 | FUN14 domain-containing protein 2 | 122.84 | 14.57 | 16564 | 2 | 1039575.5 | 906022.25 | 1510879.88 | 938554.5 | 372287.62 | 336519.12 | 562409.06 | 2 | 685354.88 | 925555.5 | 191655.72 | 598328.88 | 2 | 150046.14 | 396364.03 | 668863.69 | 634516.44 | 645586.62 | 2 | 520178.38 | 2 | 4 | 7.5 | 11.5 | 7.5 | 11.5 | 0.280324551 | 0.532462815 | 0.517931932 | 0.983786448 |
| Q8R2K1 | Fuom | Fucose mutarotase | 76.51 | 7.84 | 16805 | 1 | 2 | 149430.55 | 2 | 49925.97 | 148134.98 | 2 | 2 | 2 | 175643.7 | 177203.09 | 784228.19 | 413253.25 | 558491.44 | 611284.88 | 404531.41 | 2 | 2 | 132026.81 | 97612.95 | 2 | 0 | 9 | 0 | 10.5 | 0 | 10.5 | 7.976475005 | 7.855369856 | 0.660859666 | 0.650825972 |
| P56959 | Fus | RNA-binding protein FUS | 667.37 | 23.55 | 52673 | 10 | 12219785 | 18840610 | 22267106 | 18303006 | 17635450 | 15639112 | 16182859 | 19088326 | 14912034 | 13945400 | 15812593 | 17902200 | 16577300 | 18670390 | 18125100 | 18290428 | 16609238 | 18848508 | 13932382 | 18819232 | 9 | 11 | 6 | 9 | 9 | 11 | 0.975596811 | 1.091764576 | 0.96901205 | 1.084395744 |
| Q61584 | Fxr1 | Fragile X mental retardation syndrome-related protein 1 | 774.25 | 23.49 | 76222 | 13 | 15849502 | 15865476 | 17701092 | 17020126 | 15243648 | 19453762 | 16238296 | 14914077 | 17298454 | 19423096 | 17542432 | 16028062 | 18353336 | 15176286 | 15347895 | 17244926 | 18801716 | 20267610 | 18280394 | 17840426 | 12 | 1 | 9 | 9 | 12 | 9 | 1.009404609 | 0.944122256 | 1.131675423 | 1.058485313 |
| Q9WVR4 | Fxr2 | Fragile X mental retardation syndrome-related protein 2 | 119.4 | 3.27 | 73743 | 2 | 1128777.38 | 2 | 735837.5 | 975027.75 | 760828.81 | 1207213.5 | 1498179.12 | 2 | 1076786.12 | 1005349.06 | 1081343.88 | 843567.88 | 783511 | 2 | 914416.19 | 971703.38 | 2 | 523110.16 | 953769.69 | 1240982.88 | 11.5 | 12.5 | 6.5 | 7.5 | 11.5 | 12.5 | 1.00621238 | 0.75672447 | 1.024745265 | 0.770662171 |
| P97808 | Fxyd5 | FXYD domain-containing ion transport regulator 5 | 93.06 | 9.55 | 19454 | 2 | 3190806 | 3763312.75 | 2501074.75 | 3133643.75 | 2262994.5 | 3605343.5 | 3542784.5 | 3411531.75 | 2792344.5 | 3790476.5 | 2227453.5 | 1729037 | 2900714.75 | 2950984.25 | 2530836.25 | 2855389.5 | 3815942.25 | 3983055.5 | 4191708.75 | 4337768 | 6 | 3 | 2 | 4 | 6 | 4 | 0.830808345 | 0.719792306 | 1.291683364 | 1.119083304 |
| O35601 | Fyb1 | FYN-binding protein 1 | 660.75 | 19.41 | 90055 | 13 | 14018317 | 15207384 | 19936664 | 22236778 | 20926844 | 18552562 | 19190034 | 26405952 | 24305064 | 24768982 | 25670928 | 23980002 | 26321990 | 26831058 | 25725852 | 17826972 | 21196780 | 29763582 | 27689308 | 27412306 | 0 | 4 | 6 | 8 | 6 | 8 | 1.392130582 | 1.135195949 | 1.341864323 | 1.094206939 |
| P39688 | Fyn | Tyrosine-protein kinase Fyn | 183.21 | 5.77 | 60675 | 4 | 2 | 2 | 2 | 2 | 2 | 400727.16 | 2 | 2 | 2 | 2 | 2 | 2 | 2 | 516691.41 | 281045.72 | 2 | 749199.56 | 2 | 431872.84 | 2 | 7.5 | 7.5 | 10 | 9 | 10 | 9 | 79774.313 | 1.990699119 | 118107.84 | 2.947279196 |
| P97855 | G3bp1 | Ras GTPase-activating protein-binding protein 1 | 642.19 | 25.59 | 51829 | 12 | 3532843 | 4693111.5 | 5255233 | 5955000.5 | 4125861.75 | 4126449 | 3817857 | 3923463 | 4757718.5 | 4407665.5 | 6351378.5 | 5628655 | 4784394 | 5555381 | 4840932 | 4297503 | 5006097.5 | 5509110 | 4984539 | 5528299 | 6 | 9 | 0 | 2 | 6 | 9 | 1.152732499 | 1.291329954 | 1.074844878 | 1.204077605 |
| P97379 | G3bp2 | Ras GTPase-activating protein-binding protein 2 | 194.77 | 8.71 | 54088 | 4 | 534006.31 | 814831.19 | 415050.56 | 476439.59 | 2 | 658833.12 | 574788.38 | 805040.5 | 609257.12 | 662947.38 | 1217675.38 | 774022.75 | 427684.97 | 568499.62 | 367012.28 | 483265.56 | 569465.06 | 621564.19 | 458361.12 | 994581.88 | 9 | 7 | 9 | 7 | 9 | 7 | 1.497500602 | 1.01329818 | 1.395882883 | 0.944537573 |
| Q6NSQ9 | G6pc3 | Glucose-6-phosphatase 3 | 118.74 | 9.54 | 38782 | 2 | 2 | 91540.34 | 258280.91 | 175652.52 | 302683.22 | 209327.58 | 376627.06 | 2 | 391203 | 390881.25 | 239649.08 | 392972.56 | 440876.72 | 272935.88 | 342615.88 | 74305.98 | 71865.61 | 86434.6 | 139672.12 | 55573.12 | 3 | 6 | 9 | 5 | 9 | 6 | 2.039523981 | 1.234648856 | 0.516629579 | 0.312747545 |
| Q00612 | G6pdx | Glucose-6-phosphate 1-dehydrogenase X | 1459.48 | 53.01 | 59263 | 27 | 77125362 | 81088738 | 74658970 | 74655490 | 76736322 | 74286426 | 86203354 | 89308994 | 82921066 | 81973754 | 150968338 | 156430050 | 119277306 | 100537866 | 119809418 | 107510306 | 97650178 | 102922778 | 105539194 | 110128682 | 0 | 0 | 0 | 0 | 0 | 0 | 1.683794196 | 1.560243484 | 1.362995065 | 1.262983431 |
| P70699 | Gaa | Lysosomal alpha-glucosidase | 890.68 | 20.99 | 106248 | 15 | 7947486 | 5898959 | 7483647.5 | 10927076 | 10831213 | 7626767.5 | 6155746 | 9448248 | 8858814 | 8887092 | 6615255.5 | 4312702.5 | 5445064 | 5713855 | 6156191 | 9879978 | 14970707 | 12796220 | 11582851 | 9029516 | 2 | 4 | 2 | 1 | 2 | 4 | 0.655468296 | 0.689247558 | 1.352087732 | 1.42176696 |
| P60517 | Gabarap | Gamma-aminobutyric acid receptor-associated protein | 183.62 | 21.37 | 13918 | 3 | 5569862.5 | 5121677 | 5286218.5 | 4774505 | 4648064.5 | 5060018.5 | 5348069.5 | 5577156.5 | 5010179 | 6055656.5 | 8922369 | 7807453 | 7833266 | 6261737 | 8233382 | 3973591.5 | 4969096.5 | 5477909 | 7414601 | 4963376.5 | 0 | 12 | 0 | 8 | 0 | 12 | 1.537704858 | 1.443868674 | 1.055048385 | 0.990665604 |
| Q00422 | Gabpa | GA-binding protein alpha chain | 104.11 | 4.41 | 51344 | 2 | 1007518.56 | 1096740.88 | 1565505.12 | 1110874.38 | 1175517.12 | 1001371 | 962455.56 | 806369.88 | 1071268.25 | 1233196.62 | 2 | 955026.38 | 695978.38 | 977074.94 | 827821.81 | 1177315.62 | 830858.19 | 1097263.5 | 699014.56 | 909273.62 | 0 | 6 | 4 | 10 | 4 | 10 | 0.580223801 | 0.681011658 | 0.79140396 | 0.928874895 |
| P18645 | Gale | UDP-glucose 4-epimerase | 68.95 | 3.17 | 38225 | 1 | 2 | 2 | 133931.59 | 2 | 236987.56 | 2 | 2 | 2 | 444622.16 | 446517.44 | 1038062.62 | 890474.75 | 773074.5 | 565671.81 | 2 | 2 | 462403.09 | 2 | 504162.53 | 2 | 3.5 | 10.5 | 3.5 | 10.5 | 3.5 | 10.5 | 8.808477074 | 3.666388164 | 2.605840073 | 1.08463939 |
| Q9R0N0 | Galk1 | Galactokinase | 217.63 | 10.46 | 42295 | 4 | 10940886 | 10992705 | 11593474 | 10197334 | 10297636 | 9270073 | 11020098 | 12402017 | 10719239 | 10747313 | 13167884 | 13685344 | 13082364 | 13585244 | 13489448 | 10090397 | 12080085 | 12447714 | 12229473 | 15741454 | 0 | 5 | 0 | 6 | 0 | 6 | 1.240425023 | 1.237293999 | 1.158585066 | 1.155660619 |
| Q68FH4 | Galk2 | N-acetylgalactosamine kinase | 102.56 | 5.02 | 50503 | 2 | 3759021.5 | 3568158.5 | 3209688.75 | 5135862 | 4346549 | 3148736.25 | 2143766 | 5152401 | 2373793.75 | 3920226.25 | 2747632 | 2595828 | 3484054.75 | 3340957.5 | 3392048.5 | 2383223 | 3221991 | 3881563.5 | 3029992.25 | 1986153 | 3 | 4 | 12 | 10 | 12 | 10 | 0.777276752 | 0.929601057 | 0.724447779 | 0.866419096 |
| Q571E4 | Galns | N-acetylgalactosamine-6-sulfatase | 314.87 | 13.46 | 57673 | 6 | 3667255.5 | 3763997.25 | 4582793.5 | 4868908.5 | 7300380 | 2631194.5 | 4249218.5 | 4443266 | 5696569 | 3422548.25 | 879383.75 | 899451.25 | 421022.31 | 1468553.5 | 1383808 | 3808645.75 | 3649448.5 | 5239998 | 5606179.5 | 4310703 | 0 | 12 | 0 | 10 | 0 | 12 | 0.20891324 | 0.247139322 | 0.935147075 | 1.106256428 |
| Q6P6V1 | Galnt11 | Polypeptide N-acetylgalactosaminyltransferase 11 | 40.34 | 1.15 | 69039 | 1 | 4871399.5 | 4628804 | 6119983 | 6543670 | 5589983.5 | 5645836 | 4830788 | 4855836.5 | 4994786.5 | 4605390.5 | 2111794.5 | 1950528.5 | 2592086 | 3194829.25 | 2603771.5 | 3667474 | 4564209.5 | 4307746.5 | 4141163.25 | 4216927 | 0 | 0 | 0 | 0 | 0 | 0 | 0.448695018 | 0.499466202 | 0.752959599 | 0.83815923 |

| | | | | | | | | | | | | | | | | | | | | | | | | | | | | | | | | | | | | |
|---|---|---|---|---|---|---|---|---|---|---|---|---|---|---|---|---|---|---|---|---|---|---|---|---|---|---|---|---|---|---|---|---|---|---|---|---|
| Q6PB93 | Galnt2 | Polypeptide N-acetylgalactosaminyltransferase 2 | 300.38 | 14.04 | 64514 | 7 | 2789403.5 | 3601219.5 | 3871872.25 | 2740865 | 3696900 | 2953581.75 | 1160750.75 | 3212852.25 | 4626271 | 4072674 | 2930065.75 | 1183471.88 | 2515019 | 524482.75 | 2913797 | 1973708.5 | 2021517.25 | 3022688.5 | 2669925 | 3413067.25 | 4 | 4 | 4 | 8 | 4 | 8 | 0.60279518 | 0.628151434 | 0.784473194 | 0.817471636 |
| Q80VA0 | Galnt7 | N-acetylgalactosaminyltransferase 7 | 398.49 | 15.68 | 75419 | 8 | 1794975.75 | 3216685 | 6382163.5 | 5983846 | 6342949.5 | 2224420.25 | 2008436.25 | 6751366 | 6394791.5 | 5229270.5 | 10799148 | 9374280 | 15640026 | 15476401 | 15987120 | 1218411.88 | 1388120.38 | 6676885 | 7350081 | 6648293 | 0 | 10 | 0 | 12 | 0 | 12 | 2.836223324 | 2.975766472 | 0.981500125 | 1.029790264 |
| Q8BHN3 | Ganab | Neutral alpha-glucosidase AB | 1763.41 | 37.5 | 106911 | 32 | 43316250 | 47558126 | 48293802 | 39643114 | 43920106 | 37777278 | 43530970 | 51039322 | 48575054 | 44223650 | 77357578 | 86386914 | 84247130 | 82613770 | 90453282 | 39537282 | 35388310 | 44553694 | 44031062 | 39572930 | 0 | 6 | 0 | 7 | 0 | 7 | 1.890432502 | 1.870156084 | 0.911785585 | 0.902005947 |
| P16858 | Gapdh | Glyceraldehyde-3-phosphate dehydrogenase | 2487.43 | 85.89 | 35810 | 36 | 1263662850 | 1457022850 | 1177803906 | 1572669442 | 1057915906 | 1390415490 | 1353537410 | 1808562050 | 1145998210 | 1629853314 | 1517367810 | 1772384898 | 1699390338 | 1658613634 | 1805277058 | 1485524738 | 1469444994 | 1381968514 | 1523136514 | 1792500866 | 1 | 5 | 6 | 10 | 6 | 10 | 1.294675555 | 1.153467661 | 1.172076547 | 1.044240303 |
| Q6PAR5 | Gapvd1 | GTPase-activating protein and VPS9 domain-containing protein 1 | 163.33 | 3.36 | 162402 | 3 | 327836.56 | 1344278.25 | 1178924.38 | 1367427.75 | 1092252.75 | 1292598 | 441322.12 | 1400458 | 365298.41 | 1067009.88 | 389514.47 | 1036784.88 | 1403676.5 | 1018529.31 | 352372.81 | 1100758.25 | 2145988.5 | 1939824.38 | 1913511.5 | 1499624 | 9 | 3 | 10 | 2 | 10 | 3 | 0.791018584 | 0.919896308 | 1.619310966 | 1.883139296 |
| Q6AYA1 | Gar1 | H/ACA ribonucleoprotein complex subunit 1 | 164.31 | 14.16 | 23011 | 4 | 5186009.5 | 4950697 | 5769710 | 4512727 | 5766387 | 4027274.5 | 5915924 | 5009789.5 | 5566668 | 4669386 | 4522902 | 5602970.5 | 4788622 | 3976767 | 4145466.5 | 6004444 | 4003516 | 3904423.5 | 3562600.25 | 5411262.5 | 5 | 8 | 8 | 8 | 8 | 8 | 0.879750288 | 0.914553559 | 0.874003536 | 0.908579463 |
| Q9CZD3 | Gars | Glycine--tRNA ligase | 1369.81 | 32.24 | 81878 | 22 | 6380883 | 7186009 | 5970740 | 4144894 | 7662263 | 4526399.5 | 8420981 | 8818800 | 12161376 | 11452732 | 14122017 | 15854628 | 14825674 | 8861426 | 15453353 | 7226901 | 7395229 | 11421728 | 11333434 | 8445279 | 0 | 2 | 2 | 10 | 2 | 10 | 2.205058646 | 1.523064315 | 1.461888003 | 1.009746137 |
| Q64737 | Gart | Trifunctional purine biosynthetic protein adenosine-3 | 1584.33 | 34.06 | 107503 | 25 | 26603370 | 30402736 | 22863802 | 21259324 | 19100118 | 24516794 | 25640046 | 22414796 | 20132084 | 21923018 | 33120782 | 31242790 | 27177104 | 25144052 | 35005498 | 29713622 | 25283986 | 23015848 | 20744890 | 22076252 | 3 | 12 | 1 | 10 | 3 | 12 | 1.261673843 | 1.323340685 | 1.005034112 | 1.054157172 |
| O55148 | Gas7 | Growth arrest-specific protein 7 | 162.18 | 6.4 | 48275 | 4 | 3211542.5 | 2274289 | 5179481 | 3175095.5 | 3353446.25 | 2679248.5 | 2790475.5 | 2735770.25 | 2021708.12 | 1282789.25 | 1223111 | 1444036.62 | 761215.31 | 1325439.62 | 806602.31 | 2665401.5 | 2425076.5 | 1989926 | 1275721.88 | 1899649.88 | 0 | 2 | 2 | 6 | 2 | 6 | 0.323394905 | 0.483093737 | 0.596479161 | 0.891032426 |
| Q8VHR5 | Gatad2b | Transcriptional repressor p66-beta | 121.93 | 4.04 | 65411 | 2 | 608214.06 | 2 | 812617.69 | 778492.62 | 705307.06 | 1078024.62 | 686002.38 | 735521.44 | 757510.56 | 1012065 | 1005809.12 | 510274.5 | 502941.78 | 929691.75 | 331655.09 | 861194.19 | 1101727 | 1054769.38 | 1003001.38 | 1157685.5 | 12 | 0 | 6 | 5 | 12 | 5 | 1.129358426 | 0.768394696 | 1.782798957 | 1.212983612 |
| Q8BFQ8 | Gatd1 | Glutamine amidotransferase-like class 1 domain-containing protein 1 | 51.2 | 4.55 | 23277 | 1 | 252254.8 | 497706.56 | 559466.75 | 276284.44 | 569847.25 | 206312.72 | 326711.41 | 219345.25 | 700138.31 | 297423.84 | 126807.19 | 564441.44 | 266145.06 | 132759.88 | 446337.06 | 386754.62 | 2 | 228141.45 | 281428.16 | 207830.3 | 7 | 4 | 10 | 9 | 10 | 9 | 0.712803528 | 0.878028999 | 0.512236557 | 0.630971276 |
| P50442 | Gatm | Glycine amidinotransferase, mitochondrial | 824.55 | 33.57 | 48242 | 14 | 25437352 | 26092072 | 53985642 | 47772170 | 39542050 | 26654592 | 23759390 | 22077488 | 36202118 | 41397822 | 26460366 | 22881048 | 39924930 | 47173594 | 42137286 | 26657012 | 30455038 | 39494194 | 37778098 | 38508006 | 11 | 10 | 8 | 7 | 11 | 10 | 0.926089743 | 1.189789769 | 0.89660835 | 1.151913677 |
| P17439 | Gba | Glucosylceramidase | 306.74 | 17.09 | 57622 | 6 | 3658209.75 | 4319497 | 2655343.25 | 2841338.75 | 3636878 | 4863366 | 4027177.75 | 4388757.5 | 3940661.25 | 5280667.5 | 6105018 | 5758624 | 6775150 | 4815694 | 5934506 | 3995267.25 | 3145922.5 | 4770864 | 2119991 | 3912580 | 0 | 10 | 2 | 4 | 2 | 10 | 1.717522871 | 1.30614085 | 1.048702297 | 0.797516547 |
| Q01514 | Gbp1 | Guanylate-binding protein 1 | 130.65 | 5.26 | 67712 | 2 | 1807094.5 | 1588703.12 | 828684.75 | 1484069.38 | 1501579.62 | 1256166.5 | 1462324.75 | 1318223.5 | 1329822 | 1039228.88 | 518835.59 | 644644.38 | 715289.25 | 227331.14 | 669539.44 | 1358009 | 1400298.5 | 1077670.38 | 1122079.75 | 998651.44 | 0 | 5 | 0 | 10 | 0 | 10 | 0.384963832 | 0.433303365 | 0.826158188 | 0.929898066 |
| Q61107 | Gbp4 | Guanylate-binding protein 4 | 345.59 | 11.77 | 70801 | 6 | 3626807.75 | 3090258.5 | 3096523.5 | 5244451 | 4698836.5 | 1602165.38 | 3824537.75 | 4509062.5 | 5213511 | 5706552 | 1625176.75 | 1747337.75 | 3185457 | 820131 | 2509833.25 | 2035055.25 | 1287214.75 | 3444341 | 2073505 | 5095386 | 2 | 6 | 4 | 6 | 4 | 6 | 0.5004807 | 0.474108985 | 0.705349425 | 0.66818261 |
| O88986 | Gcat | 2-amino-3-ketobutyrate coenzyme A ligase, mitochondrial | 259.46 | 13.46 | 44931 | 3 | 579542.12 | 675976.81 | 500960.62 | 713589.75 | 312979.94 | 692680 | 288069.56 | 596076.25 | 462179.91 | 758542.31 | 1978093.12 | 1488005 | 1720661.75 | 2476862 | 1706374 | 365969.59 | 409034.56 | 537653.12 | 651583.81 | 656130.06 | 0 | 10 | 0 | 10 | 0 | 10 | 3.36680923 | 3.349360143 | 0.941546812 | 0.936667078 |
| P19468 | Gclc | Glutamate--cysteine ligase catalytic subunit | 53.69 | 1.41 | 72619 | 1 | 1570925.5 | 1646779.12 | 1315105.75 | 1706666.88 | 1192811.5 | 2065112.5 | 2009473.75 | 952832.5 | 1579832.12 | 1670522.25 | 1718312.5 | 1954028.88 | 1396741 | 2 | 1791251.25 | 1412300.5 | 1601507 | 1617985.25 | 2007279.75 | 1658163.12 | 8 | 8 | 10 | 10 | 10 | 10 | 0.92304482 | 0.828765844 | 1.11637692 | 1.002351176 |
| O09172 | Gclm | Glutamate--cysteine ligase regulatory subunit | 282.64 | 22.99 | 30535 | 5 | 2476578.25 | 2534911.5 | 2105851.75 | 2268303 | 3340085 | 3564871.75 | 2651398 | 2589788.75 | 4122152.5 | 2757284.5 | 13735158 | 15253114 | 12957263 | 17591150 | 16191247 | 4133617.25 | 3622444.25 | 6385710.5 | 5451630.5 | 4619630.5 | 0 | 0 | 0 | 1 | 0 | 1 | 5.950773353 | 4.827895427 | 1.9026833 | 1.543657515 |
| E9PVA8 | Gcn1 | eIF-2-alpha kinase activator GCN1 | 751.16 | 7.94 | 293021 | 16 | 3086507.5 | 1795483.25 | 5747501 | 4949021.5 | 6276371 | 4591625 | 4239513.5 | 4693791 | 6864027.5 | 7056868.5 | 5865056.5 | 7351483.5 | 8907562 | 6408196 | 6247750.5 | 1972917.25 | 8823621 | 4838933 | 4635507.5 | 4991057.5 | 2 | 12 | 6 | 12 | 6 | 12 | 1.591408497 | 1.267225447 | 1.155898881 | 0.920432736 |
| Q66H63 | Gdap2 | Ganglioside-induced differentiation-associated-protein 2 | 40.56 | 1.61 | 56130 | 1 | 680279.62 | 646396.44 | 665741.56 | 746857.69 | 836027.19 | 658013.19 | 867614.62 | 771976.44 | 864318.5 | 846694.38 | 265529.03 | 423426.72 | 515671.12 | 604176.25 | 509361 | 635774 | 462448.59 | 535463.44 | 2 | 880435.12 | 0 | 5 | 0 | 5 | 0 | 5 | 0.648382653 | 0.578295219 | 0.70319173 | 0.627179665 |
| P50396 | Gdi1 | Rab GDP dissociation inhibitor alpha | 471.9 | 24.83 | 50522 | 9 | 2208615.5 | 2390015 | 1741772 | 1852468.88 | 2705944.5 | 1490535 | 1582307.25 | 1408183.75 | 2686133 | 1877494.62 | 1250464.75 | 1722105 | 1539269.12 | 730087.19 | 1398362.75 | 2047385.25 | 1778796.25 | 1830386.12 | 1812277 | 1961523.25 | 0 | 7 | 5 | 8 | 5 | 8 | 0.609266996 | 0.734167287 | 0.865265362 | 1.042645552 |
| P50399 | Gdi2 | Rab GDP dissociation inhibitor beta | 1654.71 | 59.1 | 50537 | 28 | 62925746 | 67069786 | 57090730 | 43177562 | 71170962 | 35996554 | 66814918 | 62977614 | 68946882 | 43927114 | 33525008 | 55095934 | 36555642 | 25192590 | 39639566 | 61405142 | 30273944 | 43940078 | 43398842 | 41537826 | 1 | 4 | 4 | 6 | 4 | 6 | 0.630347753 | 0.681858317 | 0.731686727 | 0.791478478 |
| Q9D1H7 | Get4 | Golgi to ER traffic protein 4 homolog | 48.35 | 3.98 | 36525 | 1 | 2 | 278664.69 | 350391.75 | 2 | 919436.12 | 2 | 2 | 2 | 2 | 215368.61 | 564871.56 | 403922.91 | 424930.72 | 467637.03 | 375781.97 | 2 | 2 | 2 | 2 | 2 | 5 | 5 | 0 | 10 | 5 | 10 | 1.444720155 | 10.38712695 | 6.457877E-06 | 4.64303E-05 |
| Q8K0D5 | Gfm1 | Elongation factor G, mitochondrial | 479.66 | 13.72 | 83550 | 10 | 6783559 | 5565749.5 | 7860052 | 7621619 | 5322677 | 8394830 | 7076931 | 7332725 | 6110775 | 7584247 | 7766182 | 6306488 | 8621216 | 6573432.5 | 9893722 | 6491156 | 6115806 | 8798062 | 8082771 | 8010691.5 | 7 | 6 | 9 | 10 | 9 | 10 | 1.181198234 | 1.072919682 | 1.131051307 | 1.027369643 |
| Q5BJP6 | Gfm2 | Ribosome-releasing factor 2, mitochondrial | 89.17 | 2.7 | 85914 | 2 | 350334.75 | 326665.47 | 2 | 422326.72 | 434501.06 | 368654.44 | 541189.62 | 446972.91 | 530317.81 | 530660.94 | 972851.5 | 1102773 | 1053843.12 | 1084255.5 | 712535.94 | 592290.94 | 574670.31 | 595633.31 | 486128 | 804274 | 0 | 0 | 0 | 3 | 0 | 3 | 3.211737324 | 2.037500116 | 1.990439984 | 1.262718986 |
| P47856 | Gfpt1 | Glutamine--fructose-6-phosphate aminotransferase [isomerizing] 1 | 377.01 | 14.2 | 78539 | 8 | 6658558 | 7787280.5 | 6633611.5 | 6179008 | 6642843 | 7966783 | 8024434 | 8520079 | 7322791.5 | 7682186.5 | 8655480 | 9113173 | 8436498 | 8987235 | 8954507 | 6537959.5 | 6277290 | 9139792 | 7883323 | 5519488 | 0 | 12 | 1 | 7 | 1 | 12 | 1.302218254 | 1.117182582 | 1.042964472 | 0.894766862 |
| Q8R0H9 | Gga1 | ADP-ribosylation factor-binding protein GGA1 | 75.2 | 4.72 | 69972 | 1 | 2 | 116602.98 | 2 | 174341.97 | 261285.97 | 122936.16 | 124143.37 | 187945.08 | 218774.09 | 153778.8 | 2 | 2 | 117147.34 | 2 | 151018.98 | 178257.03 | 208847.44 | 183779.91 | 2 | 220933.8 | 9 | 8 | 2 | 10 | 9 | 10 | 0.485612753 | 0.332070074 | 1.433846632 | 0.980488164 |
| Q99J23 | Ghdc | GH3 domain-containing protein | 86.49 | 2.44 | 58508 | 1 | 598072.88 | 381341.06 | 269760.62 | 375788.34 | 306579.22 | 578016 | 444276.69 | 425887.22 | 358958.75 | 420696.97 | 381835.22 | 351408.25 | 361245.31 | 355597.16 | 245871.31 | 475864.94 | 631814.31 | 468758.66 | 539241.62 | 380770.16 | 10 | 5 | 2 | 7 | 10 | 7 | 0.878032756 | 0.7612578 | 1.292464536 | 1.120571759 |
| Q91VC9 | Ghitm | Growth hormone-inducible transmembrane protein | 313.28 | 24.86 | 37275 | 6 | 2576312 | 3687830.75 | 6075269 | 6524067.5 | 6100314 | 2906640.25 | 4822983 | 6980171 | 6177216.5 | 7413621 | 9384260 | 10679017 | 14894054 | 15683354 | 16308080 | 4028501.5 | 2992811.75 | 7750532 | 6651835.5 | 8924991 | 0 | 7 | 0 | 10 | 0 | 10 | 2.681834621 | 2.365627933 | 1.215707543 | 1.072367289 |
| Q6Y7W8 | Gigyf2 | GRB10-interacting GYF protein 2 | 74.73 | 1.7 | 149193 | 1 | 353118.47 | 2 | 466333.62 | 255507.7 | 427056.81 | 425104.56 | 307235.38 | 150967.44 | 363542.38 | 318572.91 | 2 | 531037.19 | 432182.41 | 2 | 388608.34 | 326223.69 | 2 | 310881.25 | 384660.62 | 379373.81 | 12 | 10.5 | 11 | 12 | 12 | 12 | 0.90001012 | 0.86355715 | 0.932838894 | 0.895056266 |
| Q9JLQ2 | Git2 | ARF GTPase-activating protein GIT2 | 76.98 | 2.68 | 78766 | 2 | 1589794.62 | 1771317.75 | 2012959.5 | 2139891.5 | 2158053.75 | 1529414.12 | 918338.38 | 1677075.5 | 787419.31 | 1678205.62 | 366920.91 | 2 | 1031434.25 | 692321.31 | 613285.94 | 1801838.25 | 1463904.5 | 1519093.75 | 1964832.75 | 2 | 0 | 4 | 2 | 11 | 2 | 11 | 0.279565718 | 0.410285065 | 0.697855594 | 1.024158934 |
| Q64516 | Gk | Glycerol kinase | 390.84 | 16.46 | 61227 | 7 | 2637665.75 | 3741703.75 | 2944075.75 | 4035903.75 | 3156509 | 2989530.25 | 3259632.75 | 3307568 | 4728331 | 3371385.75 | 2489381 | 3393635.5 | 2446395.5 | 2601788.5 | 3267848.5 | 3009299.5 | 841371.38 | 1552840.62 | 2567885 | 3352754 | 6 | 5 | 6 | 4 | 6 | 5 | 0.859722153 | 0.804184919 | 0.685653176 | 0.641360633 |
| P51569 | Gla | Alpha-galactosidase A | 523.65 | 24.11 | 47643 | 8 | 12717137 | 12254195 | 6878198.5 | 16050657 | 14430025 | 12027748 | 11342404 | 11550094 | 13878942 | 13755253 | 7479890 | 5911825.5 | 6431722 | 6694278.5 | 7049930 | 14994524 | 28215452 | 9662445 | 19055954 | 9255071 | 2 | 9 | 0 | 10 | 2 | 10 | 0.538545348 | 0.536614914 | 1.302473435 | 1.297804675 |
| P23780 | Glb1 | Beta-galactosidase | 833.69 | 23.34 | 73121 | 14 | 42264458 | 35933210 | 44264378 | 35252802 | 47633602 | 36129026 | 37455510 | 26782392 | 36770010 | 27906636 | 5706196 | 7500823 | 8964626 | 6712162 | 5864396 | 39855654 | 24691772 | 32939704 | 31199270 | 30851970 | 0 | 2 | 0 | 11 | 0 | 11 | 0.169215804 | 0.210539569 | 0.776915385 | 0.966643936 |
| Q61543 | Glg1 | Golgi apparatus protein 1 | 1673.07 | 28.94 | 133734 | 28 | 25615172 | 26871744 | 44065086 | 41590838 | 36088326 | 30504582 | 30847212 | 39377182 | 36388778 | 41076322 | 12825995 | 13107742 | 19986176 | 21584276 | 24333706 | 25931756 | 25789120 | 43788518 | 42409470 | 44149798 | 0 | 10 | 0 | 10 | 0 | 10 | 0.527103716 | 0.515381303 | 1.044983318 | 1.02174363 |
| Q9CWG1 | Glipr1 | Glioma pathogenesis-related protein 1 | 115.94 | 9.41 | 29129 | 2 | 2 | 854614.31 | 989603.81 | 1219983.88 | 1598749.75 | 911999.88 | 865316.19 | 1456372.25 | 1135355.25 | 1156446 | 245185.66 | 898991.5 | 1047552.06 | 1257052.75 | 2 | 2 | 1056889.38 | 1492866 | 930717.62 | 1237875.88 | 10.5 | 11.5 | 7 | 12 | 10.5 | 12 | 0.739613591 | 0.624158987 | 1.011880266 | 0.853924493 |
| Q9CYL5 | Glipr2 | Golgi-associated plant pathogenesis-related protein 1 | 46.81 | 8.44 | 17090 | 1 | 2 | 2 | 2 | 2 | 272615.31 | 2 | 288410.5 | 408759.5 | 276013.22 | 2 | 599081.25 | 2 | 484963.88 | 289800.84 | 641339.31 | 2 | 2 | 313288.19 | 351099.69 | 491465.69 | 3 | 6 | 5 | 10 | 5 | 10 | 7.391837771 | 2.070708738 | 4.239760606 | 1.187703194 |
| Q8BZM1 | Glmn | Glomulin | 53.86 | 1.68 | 67756 | 1 | 298990.91 | 170742.17 | 361162.25 | 352767.59 | 398408.16 | 245417.22 | 256398.94 | 203556.58 | 515639.81 | 384634.94 | 410407.34 | 223702.34 | 315808.56 | 265678.16 | 224841.98 | 414664.84 | 364404.19 | 359118.5 | 297278.97 | 324083.97 | 10 | 10 | 12 | 9 | 12 | 10 | 0.910476399 | 0.897107484 | 1.112181679 | 1.095851039 |
| Q9JHJ3 | Glmp | Glycosylated lysosomal membrane protein | 146.03 | 8.17 | 43804 | 2 | 2 | 230614.86 | 194429.7 | 156101.52 | 219160.42 | 2 | 122646.09 | 2 | 2 | 2 | 2707503.5 | 1630094.25 | 2475595.5 | 1920004.12 | 1679913.75 | 2 | 2 | 2 | 2 | 234816.42 | 0 | 7 | 0 | 12 | 0 | 12 | 13.01137139 | 84.8981972 | 0.293417376 | 1.914525802 |
| Q9CPU0 | Glo1 | Lactoylglutathione lyase | 350.59 | 37.5 | 20810 | 6 | 9899079 | 10759783 | 13349924 | 10598170 | 12377976 | 8869105 | 8081246 | 6699795 | 12244632 | 9692845 | 7672570.5 | 7357579 | 8489167 | 3420820 | 9107245 | 9720097 | 6839863.5 | 11619019 | 6128675.5 | 8774547 | 0 | 3 | 7 | 11 | 7 | 11 | 0.632577424 | 0.790727376 | 0.756027962 | 0.94504164 |
| Q9CPV4 | Glod4 | Glyoxalase domain-containing protein 4 | 463.85 | 36.24 | 33317 | 10 | 19489580 | 18360552 | 19892076 | 20743120 | 18179010 | 16820710 | 17473012 | 16405674 | 16784588 | 19289066 | 10183757 | 12609820 | 10299401 | 10698300 | 11985039 | 21871454 | 18815360 | 19644470 | 19712764 | 21433614 | 0 | 7 | 0 | 1 | 0 | 7 | 0.577010283 | 0.642783871 | 1.049794206 | 1.169460587 |
| Q9QUH0 | Glrx | Glutaredoxin-1 | 446.83 | 78.5 | 11871 | 7 | 16590774 | 18393680 | 20626622 | 18929708 | 28195360 | 12548680 | 18326058 | 17105410 | 23574208 | 17299522 | 8301836 | 13792263 | 12888739 | 11593634 | 12197627 | 23797268 | 14343936 | 17662294 | 15705932 | 19274464 | 0 | 8 | 2 | 11 | 2 | 11 | 0.572087843 | 0.661469148 | 0.88366071 | 1.021721235 |
| Q9CQM9 | Glrx3 | Glutaredoxin-3 | 342.37 | 29.08 | 37778 | 6 | 11594562 | 4945688 | 6471969.5 | 2758655.75 | 4905984 | 1602599.5 | 5837573 | 6789633.5 | 5288878 | 5910969 | 3940652 | 5208554 | 2673197.25 | 1079334.12 | 5632294 | 6084788 | 2202287 | 6527204 | 2291895.5 | 7006759 | 7 | 11 | 5 | 10 | 7 | 11 | 0.604169782 | 0.728835402 | 0.786030059 | 0.948221098 |
| Q80Y14 | Glrx5 | Glutaredoxin-related protein 5, mitochondrial | 114.37 | 20.39 | 16292 | 2 | 607386.69 | 894774 | 599715.06 | 1012731.44 | 618272 | 2 | 731798.69 | 776045.5 | 972144.94 | 783121.56 | 537032.38 | 748820.81 | 979582.31 | 943588.12 | 628393.94 | 1179602.25 | 1014617.25 | 994084.25 | 762950.62 | 956111.25 | 11 | 4 | 12 | 4 | 12 | 4 | 1.028004756 | 1.175999092 | 1.314632853 | 1.503890943 |
| D3Z7P3 | Gls | Glutaminase kidney isoform, mitochondrial | 871.07 | 29.38 | 73964 | 14 | 20602780 | 21176534 | 16943936 | 20668332 | 15664067 | 21888312 | 19613980 | 13120274 | 15428939 | 13470227 | 22598866 | 21569402 | 21549394 | 18837506 | 21779066 | 20492754 | 27388032 | 20966730 | 22435588 | 23570970 | 3 | 4 | 5 | 2 | 5 | 4 | 1.118652443 | 1.27313253 | 1.208282466 | 1.375139994 |
| B0BNM9 | GLTP | Glycolipid transfer protein | 236.33 | 22.01 | 23704 | 5 | 1678254.62 | 1315361.25 | 2348158.75 | 2098829 | 2578301 | 1020392.38 | 1325934.75 | 2097967 | 2620231.5 | 1752848.75 | 2 | 561012.25 | 733115 | 289417.22 | 765984.38 | 2016055.75 | 1166587.5 | 1465446.75 | 1547395.5 | 1663426.88 | 0 | 5 | 0 | 10 | 0 | 10 | 0.234509753 | 0.266466042 | 0.784408344 | 0.89129848 |
| P26443 | Glud1 | Glutamate dehydrogenase 1, mitochondrial | 1661.16 | 51.25 | 61337 | 27 | 113297498 | 115152546 | 117885554 | 129583242 | 125059050 | 110059090 | 115457642 | 119671210 | 116117354 | 113547410 | 80317898 | 112701482 | 127279098 | 118532202 | 139003218 | 113212306 | 87241202 | 101562778 | 109331082 | 118409170 | 12 | 3 | 10 | 5 | 12 | 5 | 0.961489445 | 1.00518601 | 0.881490895 | 0.921551786 |
| Q5RKH0 | Glyr1 | Putative oxidoreductase GLYR1 | 595.04 | 18.84 | 60421 | 8 | 4198848 | 2858047 | 4229403 | 4660015.5 | 5381122 | 5284244 | 3580306.25 | 3609398.5 | 3732485.5 | 5391298.5 | 2569613.5 | 2599059.75 | 3703618 | 2161565 | 3011373.75 | 3537101.5 | 4416767 | 3767762.5 | 3845067.5 | 2951375.25 | 2 | 7 | 2 | 9 | 2 | 9 | 0.65855222 | 0.65031039 | 0.868274751 | 0.857408227 |
| Q8K0C9 | Gmds | GDP-mannose 4,6 dehydratase | 71.34 | 5.11 | 41985 | 1 | 2 | 251705.66 | 288102.16 | 2 | 258358.08 | 2 | 381830.34 | 301469.34 | 315615.59 | 2 | 2 | 2 | 2 | 2 | 230006.81 | 2 | 2 | 2 | 2 | 2 | 6 | 5 | 6 | 5 | 6 | 5 | 0.288177755 | 0.230263663 | 1.252866E-05 | 1.001082E-05 |
| Q9ERL7 | Gmfg | Glia maturation factor gamma | 263.55 | 30.28 | 16748 | 4 | 10874212 | 10541139 | 16520236 | 18279986 | 14080868 | 12778224 | 11267765 | 17283380 | 16281825 | 18123528 | 6446971.5 | 7371342.5 | 13192544 | 14811910 | 9033983 | 8759798 | 10148967 | 18296728 | 15654344 | 17357282 | 5 | 12 | 4 | 11 | 5 | 12 | 0.723461249 | 0.671511688 | 0.998871607 | 0.927145662 |
| Q6PGG2 | Gmip | GEM-interacting protein | 193.14 | 4.53 | 107545 | 3 | 958428 | 743113.38 | 800678.31 | 538967 | 1085566.25 | 610231.38 | 797447.75 | 616476.06 | 758741.81 | 632059.88 | 1106777.25 | 756363.38 | 753855.25 | 749634.38 | 511358.31 | 824624.88 | 677384.88 | 265491.59 | 691822.94 | 828333.25 | 11 | 8 | 11 | 9 | 11 | 9 | 0.939719103 | 1.135589323 | 0.79666934 | 0.962723002 |
| Q5XIC1 | Gmppa | Mannose-1-phosphate guanyltransferase alpha | 89.11 | 6.19 | 46191 | 2 | 1567843 | 1306906.38 | 906230.62 | 1348682.12 | 1013347.06 | 1930939.5 | 1351354.75 | 1150240 | 821716.75 | 1227926.75 | 1617891.25 | 1821874.75 | 1590380 | 1317844.5 | 1448600 | 1355999.38 | 215209.17 | 1022992.94 | 2349720 | 1252099.38 | 3 | 12 | 6 | 12 | 6 | 12 | 1.269180994 | 1.202773327 | 1.008629596 | 0.955854824 |
| Q8BTZ7 | Gmppb | Mannose-1-phosphate guanyltransferase beta | 114.9 | 5 | 39917 | 1 | 462113.59 | 543651.25 | 325736.97 | 2 | 783192 | 2 | 567422.69 | 261592.55 | 2 | 484389.69 | 396611.81 | 689346.25 | 716082.06 | 1025248.81 | 761616.69 | 431084.28 | 2 | 592998.44 | 1015113.69 | 755797.56 | 6 | 9.5 | 2 | 6 | 6 | 9.5 | 1.697126179 | 2.732511968 | 1.321701191 | 2.128047028 |
| Q9DCZ1 | Gmpr | GMP reductase 1 | 121.23 | 8.99 | 37482 | 2 | 2918934 | 925410.56 | 2536706.25 | 2904885.25 | 2185961.5 | 2898652.5 | 2445964 | 2662434.5 | 2997920.5 | 2542904.5 | 1896629.5 | 1981924.5 | 2663020.5 | 2159299.75 | 2222918.25 | 2690206 | 2530750.25 | 2959218 | 3255844 | 3198101.5 | 8 | 5 | 3 | 7 | 8 | 7 | 0.952221936 | 0.80631034 | 1.275649445 | 1.080178159 |
| Q3THK7 | Gmps | GMP synthase [glutamine-hydrolyzing] | 853.6 | 26.41 | 76723 | 15 | 15077493 | 13083123 | 14558206 | 14290489 | 14767446 | 14605928 | 13368783 | 13763161 | 15033546 | 15498302 | 15630660 | 17478024 | 18972180 | 13587330 | 18087274 | 14151812 | 10152675 | 13423085 | 11755078 | 15063229 | 4 | 6 | 4 | 7 | 4 | 7 | 1.16688844 | 1.158928912 | 0.899258781 | 0.893124797 |
| P27601 | Gna13 | Guanine nucleotide-binding protein subunit alpha-13 | 337.86 | 22.55 | 44055 | 6 | 1519471.25 | 1368496.75 | 824845.38 | 1392012.25 | 1282727 | 986958.75 | 1416425.75 | 953664.88 | 1728929.5 | 2162668.5 | 1075150.75 | 1618496.5 | 3200567.25 | 1321075.5 | 2427483.5 | 1264214.88 | 1531027 | 1848905.62 | 1498560.12 | 1712408.25 | 7 | 5 | 8 | 10 | 8 | 10 | 1.509619421 | 1.330285913 | 1.229753604 | 1.083666436 |
| P08752 | Gnai2 | Guanine nucleotide-binding protein G(i) subunit alpha-2 | 1671.55 | 65.35 | 40489 | 22 | 50103158 | 47340502 | 47325442 | 52269762 | 49910698 | 47641506 | 49348982 | 45360682 | 46421414 | 43080854 | 31885220 | 35984118 | 32009802 | 34230586 | 34773162 | 44648494 | 52370350 | 46960338 | 50966458 | 43271574 | 0 | 9 | 0 | 10 | 0 | 10 | 0.683876038 | 0.728403639 | 0.964639144 | 1.027447408 |
| Q9DC51 | Gnai3 | Guanine nucleotide-binding protein G(k) subunit alpha | 1149.71 | 47.18 | 40538 | 16 | 5752976 | 4572239 | 4726301 | 4667543.5 | 5612450.5 | 7063245.5 | 6263146 | 5585737.5 | 5988781 | 5238725.5 | 7110890 | 6606762 | 6539927 | 6349014.5 | 6912081 | 7069470 | 9114574 | 6565600 | 7064013 | 4818314 | 0 | 2 | 4 | 6 | 4 | 6 | 1.323200808 | 1.112112802 | 1.367149886 | 1.149050757 |
| P21279 | Gnaq | Guanine nucleotide-binding protein G(q) subunit alpha | 397.39 | 23.96 | 42158 | 7 | 961926.5 | 1821455.12 | 2415721.5 | 2226604.5 | 4617763 | 974881.88 | 1866045.88 | 3308722.5 | 4202277.5 | 1841703.5 | 2383337 | 6089609 | 3326210.75 | 6426948.5 | 2601075.25 | 2153612.5 | 5755372.5 | 2596017.5 | 1688970 | 1439778.75 | 4 | 12 | 5 | 12 | 5 | 12 | 1.729333774 | 1.708037586 | 1.132045046 | 1.118104276 |
| P63094 | Gnas | Guanine nucleotide-binding protein G(s) subunit alpha isoforms short | 505.01 | 24.62 | 45664 | 8 | 5026894 | 4488970 | 5923516.5 | 5784612.5 | 5686334 | 4814387 | 5760905 | 4146134 | 6679722.5 | 5987157 | 6813710 | 7304083.5 | 8324615 | 5446114.5 | 7597258 | 4672091 | 3425669.5 | 5298360.5 | 5138440 | 5067967 | 3 | 7 | 3 | 7 | 3 | 7 | 1.318667774 | 1.29565449 | 0.877080684 | 0.861773942 |
| P54311 | Gnb1 | Guanine nucleotide-binding protein G(I)/G(S)/G(T) subunit beta-1 | 696.57 | 41.18 | 37377 | 11 | 6536760 | 4148493 | 2986982.25 | 3847832.5 | 2392784.25 | 5679358.5 | 3909425.25 | 2205288.75 | 2614775.25 | 2927658.75 | 7069462.5 | 6126240 | 5490734 | 4156672.75 | 4269139 | 3311884 | 3182917 | 1977472.75 | 2497550.75 | 3045747.75 | 4 | 7 | 3 | 10 | 4 | 10 | 1.36154521 | 1.563881873 | 0.703845549 | 0.808442707 |
| P54313 | Gnb2 | Guanine nucleotide-binding protein G(I)/G(S)/G(T) subunit beta-2 | 1142.47 | 62.65 | 37331 | 18 | 22970378 | 25894944 | 21083106 | 22519498 | 19943470 | 22279910 | 27288682 | 22806836 | 17883462 | 20248970 | 29955782 | 26426594 | 26723160 | 22789936 | 26281214 | 20625876 | 18901542 | 19438148 | 22578084 | 16152432 | 2 | 4 | 5 | 7 | 5 | 7 | 1.175829949 | 1.196084025 | 0.869094109 | 0.884064554 |
| Q9DAS9 | Gng12 | Guanine nucleotide-binding protein G(I)/G(S)/G(O) subunit gamma-12 | 100.42 | 25 | 7997 | 2 | 2840046 | 3039325 | 2952033 | 3701495.75 | 3212946.5 | 2871276.5 | 3004466.5 | 4034091.5 | 3178378.75 | 4413520.5 | 2491961.5 | 2189657.5 | 3300172.5 | 3986711 | 3462711.75 | 3297062.5 | 1531590.25 | 3919242.25 | 2715584 | 4073161.5 | 12 | 11 | 9 | 10 | 12 | 11 | 0.980018095 | 0.881696321 | 0.986713591 | 0.887720081 |
| P63213 | Gng2 | Guanine nucleotide-binding protein G(I)/G(S)/G(O) subunit gamma-2 | 108.69 | 26.76 | 7850 | 2 | 4900839 | 3963083.5 | 5035450 | 6184619 | 4561058 | 4827567 | 4254266 | 3851059.75 | 4289304 | 5563404 | 2989544 | 2787218.75 | 3226758.5 | 3337359 | 3400853 | 3883504 | 5128770 | 5697879.5 | 6212776 | 4825493 | 0 | 10 | 0 | 7 | 0 | 10 | 0.638738147 | 0.690863209 | 1.044770573 | 1.130030443 |
| P63219 | Gng5 | Guanine nucleotide-binding protein G(I)/G(S)/G(O) subunit gamma-5 | 56.23 | 16.18 | 7318 | 1 | 2 | 338255.41 | 157235.94 | 352873.91 | 113089.93 | 2 | 2 | 79445.01 | 91612.26 | 115008.9 | 2 | 294221.81 | 97930.44 | 258838.72 | 489347.38 | 364063.84 | 459801.03 | 117524.04 | 224626.3 | 127341.12 | 12.5 | 8 | 5 | 0 | 12.5 | 8 | 1.186054212 | 3.986226002 | 1.345204283 | 4.521115676 |
| P36916 | Gnl1 | Guanine nucleotide-binding protein-like 1 | 196.89 | 7.91 | 68771 | 3 | 2308742.75 | 1689827.5 | 1553717.25 | 2496493.75 | 2000936.75 | 2073888 | 1247195.25 | 1845201.25 | 1625822.62 | 2758535.25 | 1468655.5 | 1464360.62 | 1685672 | 1334930.12 | 1486147.88 | 1700411.25 | 2140499 | 1821978.5 | 2544159.75 | 1693564.88 | 1 | 11 | 6 | 11 | 6 | 11 | 0.740296008 | 0.778980704 | 0.985163303 | 1.036643714 |
| Q99LH1 | Gnl2 | Nucleolar GTP-binding protein 2 | 168.62 | 5.49 | 83345 | 3 | 436607.03 | 335824.34 | 2 | 413059.66 | 226759.77 | 855812.06 | 2 | 2 | 279102.22 | 316151.31 | 1512346.75 | 1021990.88 | 802155.12 | 978481 | 791115.19 | 2 | 2 | 519372.44 | 230893.27 | 356939.25 | 0 | 11 | 2 | 12 | 2 | 12 | 3.615562979 | 3.518844978 | 0.784001958 | 0.763029539 |
| Q8CI11 | Gnl3 | Guanine nucleotide-binding protein-like 3 | 720.79 | 27.14 | 60786 | 12 | 5920561 | 5999730.5 | 4526905.5 | 4673007 | 6393163.5 | 5907068 | 5196668.5 | 4184427 | 6392547 | 4708152 | 6943035.5 | 6490127.5 | 6221106.5 | 5209271.5 | 8117401 | 4386107.5 | 2537562.5 | 3061264.75 | 2797507 | 2582216.25 | 4 | 0 | 3 | 1 | 4 | 1 | 1.198724293 | 1.249805368 | 0.558443382 | 0.582240254 |
| O88958 | Gnpda1 | Glucosamine-6-phosphate isomerase 1 | 210.4 | 16.96 | 32549 | 4 | 5967354 | 4723525 | 4245984 | 4904127 | 4249126.5 | 4199582.5 | 5208835.5 | 2965362 | 4408621 | 4468334.5 | 3674084.5 | 2324225.5 | 1999416 | 1554132.5 | 649077.31 | 6403557 | 7249584 | 4332663 | 4230696 | 4012467.5 | 0 | 12 | 1 | 10 | 1 | 12 | 0.423449003 | 0.480027424 | 1.088785415 | 1.234261633 |
| Q9JK38 | Gnpnat1 | Glucosamine 6-phosphate N-acetyltransferase | 90.69 | 5.98 | 20791 | 1 | 1921383.75 | 1915372 | 2558620.5 | 1831839 | 3542964.25 | 1017773.88 | 2775182.5 | 2507924.25 | 3371237.75 | 1836101.75 | 843019.12 | 1975685 | 2170528 | 1050499.88 | 1762365.25 | 2404618.25 | 778604.81 | 1808680.38 | 679116.94 | 1117691.12 | 6 | 3 | 6 | 4 | 6 | 4 | 0.662869861 | 0.677958638 | 0.57677213 | 0.589901081 |
| Q8BFR4 | Gns | N-acetylglucosamine-6-sulfatase | 768.17 | 25 | 61175 | 14 | 34371678 | 26735750 | 25553570 | 29391822 | 28688966 | 22658670 | 22628404 | 19831766 | 25202052 | 23195590 | 2423842.25 | 4221129 | 5897701 | 4668385.5 | 4903818 | 24064126 | 21555644 | 18135118 | 19287190 | 19508728 | 0 | 0 | 0 | 5 | 0 | 5 | 0.152788468 | 0.19481643 | 0.70850864 | 0.903400142 |
| Q6AYQ1 | Golga7 | Golgin subfamily A member 7 | 76.83 | 10.95 | 15778 | 1 | 669100.5 | 883763.12 | 615696.19 | 340600.81 | 655986.12 | 527976.62 | 787668.44 | 2 | 582809.12 | 588582.94 | 2 | 754486.5 | 1083706.5 | 854188.75 | 463857.88 | 2 | 529254.56 | 574356.06 | 713793.56 | 2 | 11 | 6 | 9.5 | 9 | 11 | 9 | 0.99718651 | 1.269075989 | 0.574193972 | 0.730751746 |
| Q9CRA5 | Golph3 | Golgi phosphoprotein 3 | 236.69 | 10.4 | 33752 | 3 | 895677.62 | 760683.06 | 768271.62 | 819828.75 | 1714877 | 781294.69 | 859255.81 | 937214.31 | 972951.12 | 1097744.5 | 393232.97 | 691221.38 | 812692.75 | 637722.25 | 752718.12 | 944415.5 | 607882.44 | 702092.62 | 1037057.88 | 1145830.5 | 2 | 12 | 1 | 12 | 2 | 12 | 0.662908525 | 0.707242219 | 0.894732098 | 0.954569584 |
| Q9CR60 | Golt1b | Vesicle transport protein GOT1B | 205.95 | 17.39 | 15422 | 3 | 837335.12 | 947490.5 | 612597.81 | 914738.88 | 779631.69 | 1269546.5 | 1572276.5 | 1458348.62 | 2004970.5 | 1876188.12 | 3400907 | 3160662 | 4960405 | 4752864.5 | 3736450.75 | 1451852.12 | 1523484.75 | 2039682.25 | 2843197.5 | 1856451 | 0 | 0 | 0 | 9 | 0 | 9 | 4.890590594 | 2.445970113 | 2.374182967 | 1.187419079 |
| Q99JX3 | Gorasp2 | Golgi reassembly-stacking protein 2 | 244.22 | 15.52 | 47038 | 4 | 1650941.75 | 1424784.38 | 1956983.5 | 2177194 | 1722735.25 | 1433776.5 | 1595799.38 | 1531710 | 2649507.75 | 2697534.5 | 2326780 | 1762114.12 | 1335447.38 | 1226327.62 | 2246038.5 | 2685863 | 2511430.5 | 1355499 | 2172482.5 | 2835072.5 | 12 | 6 | 9 | 10 | 12 | 10 | 0.995977531 | 0.897901998 | 1.294169355 | 1.166730386 |
| P05201 | Got1 | Aspartate aminotransferase, cytoplasmic | 1401.39 | 65.86 | 46248 | 23 | 113522818 | 117045410 | 126015354 | 141350386 | 134842066 | 109237498 | 105238154 | 119877010 | 116997346 | 125425698 | 70286122 | 76833890 | 76493218 | 81340842 | 81645114 | 131531642 | 147978738 | 141992338 | 148730754 | 151981010 | 0 | 2 | 0 | 0 | 0 | 2 | 0.610957377 | 0.670276473 | 1.141342976 | 1.25215829 |
| P05202 | Got2 | Aspartate aminotransferase, mitochondrial | 1545.57 | 56.28 | 47411 | 26 | 396806242 | 380090466 | 383207074 | 409902914 | 417266818 | 453874786 | 394694914 | 418113346 | 415929090 | 442183714 | 377579874 | 391480738 | 478232354 | 541814274 | 492282114 | 365633346 | 433598914 | 449360354 | 463537890 | 451701858 | 8 | 5 | 10 | 9 | 10 | 9 | 1.147999678 | 1.073698141 | 1.088844765 | 1.018371888 |
| Q9WTK3 | Gpaa1 | Glycosylphosphatidylinositol anchor attachment 1 protein | 91.37 | 3.06 | 67949 | 2 | 998150.25 | 801489.44 | 1806090.62 | 842337.19 | 1293991 | 1784338.25 | 761747.94 | 2178971.75 | 2048846.38 | 2668100.75 | 3157840.75 | 3569428.5 | 4631606 | 3921310 | 4640166 | 1079046.25 | 1462610.38 | 2607028 | 2639096.5 | 2310962 | 0 | 3 | 0 | 11 | 0 | 11 | 3.469200331 | 2.109758584 | 1.758732192 | 1.069554936 |
| Q3ULJ0 | Gpd1l | Glycerol-3-phosphate dehydrogenase 1-like protein | 252.36 | 17.66 | 38226 | 5 | 1532150.5 | 1723690.25 | 1112994.25 | 983500.12 | 1323901.25 | 1226615.5 | 1790738 | 742968.06 | 928917.62 | 685422.62 | 361246.75 | 1262629.62 | 467217.5 | 664568.12 | 621616.88 | 1656395.25 | 990101.56 | 1116075.38 | 828185.5 | 820885.5 | 2 | 7 | 4 | 11 | 4 | 11 | 0.505865683 | 0.62837049 | 0.810582923 | 1.006880692 |
| Q64521 | Gpd2 | Glycerol-3-phosphate dehydrogenase, mitochondrial | 869.14 | 24.07 | 80954 | 13 | 7528557 | 8048480 | 6865208.5 | 6315002.5 | 8066200 | 8260476.5 | 8982288 | 9698333 | 8182471.5 | 7088192 | 10341992 | 10060308 | 11104671 | 7217416.5 | 10265784 | 8130115 | 7000793.5 | 6796087 | 6994010.5 | 7318030 | 3 | 12 | 4 | 2 | 4 | 12 | 1.33040696 | 1.160581088 | 0.984129351 | 0.858505666 |
| P06745 | Gpi | Glucose-6-phosphate isomerase | 1566.22 | 50 | 62767 | 25 | 95635858 | 115971010 | 123780794 | 109734914 | 120243714 | 94177906 | 101394498 | 127361282 | 109968298 | 104739970 | 97667098 | 106099738 | 90005986 | 91010874 | 99050354 | 118657778 | 106306338 | 118459698 | 103322138 | 120754122 | 3 | 12 | 5 | 8 | 5 | 12 | 0.855788643 | 0.899918703 | 1.003774162 | 1.055535324 |
| Q99P91 | Gpnmb | Transmembrane glycoprotein NMB | 168.77 | 4.7 | 63676 | 3 | 34177054 | 29611870 | 7935134.5 | 10534111 | 11133614 | 24805202 | 27124248 | 15861661 | 15724259 | 14278674 | 14915914 | 12240026 | 6077382 | 4291061.5 | 6737862.5 | 28083108 | 37366606 | 17717244 | 13532877 | 16026271 | 6 | 8 | 1 | 9 | 6 | 9 | 0.473941543 | 0.452606766 | 1.207023806 | 1.152688869 |
| Q8CIM5 | Gpr84 | G-protein coupled receptor 84 | 576.73 | 26.77 | 43717 | 9 | 4737317.5 | 5633174.5 | 4121213 | 5519103 | 4899098 | 6839831 | 4338420.5 | 5987096.5 | 5189006 | 5859604.5 | 21072842 | 18636366 | 21370696 | 20801552 | 22792746 | 4651388 | 6195971 | 5983155.5 | 6814149.5 | 5941281 | 0 | 4 | 0 | 10 | 0 | 10 | 4.202111481 | 3.710014743 | 1.187718051 | 1.048627934 |
| Q8BHL4 | Gprc5a | Retinoic acid-induced protein 3 | 67.55 | 3.93 | 40101 | 1 | 2 | 2 | 2 | 2 | 2 | 2 | 2 | 2 | 2 | 2 | 915188 | 664877.88 | 520275.44 | 369602.59 | 595242.12 | 2 | 2 | 2 | 2 | 2 | 0 | 12.5 | 0 | 12.5 | 0 | 12.5 | 306518.603 | 306518.603 | 1 | 1 |
| P97834 | Gps1 | COP9 signalosome complex subunit 1 | 187.73 | 7.64 | 53428 | 4 | 4449191.5 | 4769402 | 5380594 | 4969481.5 | 7536473 | 3560208.5 | 4798125 | 4698583.5 | 6944052 | 4565327 | 3645425.5 | 4262933 | 3871404.5 | 4424618 | 4703242.5 | 4718142.5 | 4636307.5 | 4552542 | 5268358 | 5162446 | 1 | 9 | 7 | 11 | 7 | 11 | 0.771352664 | 0.851069429 | 0.897903283 | 0.990698638 |
| P11352 | Gpx1 | Glutathione peroxidase 1 | 248.75 | 31.34 | 22329 | 5 | 3736602 | 3598004 | 6533950 | 7874368.5 | 5863651 | 3154780 | 3609154 | 5625780 | 4729931.5 | 8089506 | 912262.44 | 1916766.5 | 3012058.25 | 3849631 | 4108609.25 | 2746387.5 | 2312938 | 7702090 | 6109026 | 6029176.5 | 4 | 10 | 4 | 12 | 4 | 12 | 0.499856545 | 0.54739357 | 0.901945191 | 0.987721384 |
| P23764 | Gpx3 | Glutathione peroxidase 3 | 40.1 | 7.52 | 25424 | 1 | 2 | 2 | 2 | 2 | 2 | 2 | 2 | 2 | 2 | 2 | 1075901 | 934652.62 | 769936.75 | 342222.84 | 682758.56 | 2 | 2 | 2 | 2 | 2 | 0 | 12.5 | 0 | 12.5 | 0 | 12.5 | 380547.177 | 380547.177 | 1 | 1 |
| Q60631 | Grb2 | Growth factor receptor-bound protein 2 | 368.79 | 40.55 | 25238 | 7 | 7810937 | 11231547 | 17118800 | 18171862 | 14599754 | 7070851 | 14445533 | 19293238 | 19361236 | 17730244 | 7535205 | 12243289 | 6084945.5 | 11265763 | 13249627 | 14931679 | 12592599 | 15776056 | 17779360 | 17431024 | 6 | 9 | 4 | 10 | 6 | 10 | 0.730838678 | 0.646702398 | 1.138944075 | 1.007825512 |
| O35127 | Grcc10 | Protein C10 | 212.83 | 53.17 | 13194 | 4 | 2830727 | 2124692.25 | 2840743 | 2351709.75 | 3036198 | 2253090 | 1385207.12 | 2971057.5 | 2330686.5 | 2094438.62 | 300331.84 | 871908.38 | 1049285.5 | 216741.56 | 738508.38 | 2947734 | 1376036 | 3159710.25 | 2796134.5 | 3133875 | 0 | 9 | 0 | 7 | 0 | 9 | 0.240955612 | 0.287895373 | 1.017401284 | 1.215597841 |
| Q91Z53 | Grhpr | Glyoxylate reductase/hydroxypyruvate reductase | 208.71 | 13.72 | 35329 | 4 | 3998305.25 | 3800857.25 | 3502499.5 | 3286905.75 | 2101520.25 | 4578127 | 3842619.5 | 4242800.5 | 2118093 | 4206276 | 3896613.75 | 3579385.5 | 3526946.25 | 4461975.5 | 4205869.5 | 3571596.25 | 4466917.5 | 5467272 | 5113549 | 4312383.5 | 5 | 2 | 10 | 6 | 10 | 6 | 1.178591179 | 1.035963636 | 1.373972279 | 1.207700637 |
| P26817 | Grk2 | Beta-adrenergic receptor kinase 1 | 77.28 | 2.32 | 79785 | 2 | 796434.62 | 230835.19 | 1241284.75 | 729656.88 | 2199289 | 626229.06 | 729989.88 | 1165076.25 | 1607492.5 | 913466.75 | 318284.44 | 629825.38 | 498495.19 | 2 | 715886 | 958661.5 | 747198.75 | 439065.12 | 528776.69 | 707763.75 | 4 | 8 | 2 | 6 | 4 | 8 | 0.416064036 | 0.428874234 | 0.650594617 | 0.670625779 |
| P28798 | Grn | Granulins | 877.83 | 35.65 | 63458 | 16 | 32750454 | 32672644 | 36089554 | 50561286 | 40879382 | 39148214 | 28006742 | 29346302 | 25807506 | 48105634 | 16646476 | 13557626 | 20974908 | 26247752 | 17241556 | 28665162 | 23522250 | 32188202 | 31137702 | 28323992 | 0 | 0 | 1 | 10 | 1 | 10 | 0.490628086 | 0.555518308 | 0.745451325 | 0.844044339 |
| P97576 | Grpel1 | GrpE protein homolog 1, mitochondrial | 295.2 | 29.95 | 24297 | 6 | 1735018.38 | 2640217.5 | 1412237 | 466946.25 | 3196520 | 819662 | 2921576.75 | 2171001.5 | 3650719 | 1971295.88 | 6761640 | 7588640 | 6456072.5 | 6640221 | 7442133 | 1896635.38 | 331680.31 | 3010323 | 810692.25 | 1004236.94 | 0 | 9 | 0 | 6 | 0 | 9 | 3.691559751 | 3.0247906 | 0.74633513 | 0.611532154 |
| Q8C5Q4 | Grsf1 | G-rich sequence factor 1 | 140.75 | 7.1 | 53076 | 2 | 512287.38 | 460815.47 | 754002.75 | 518116.41 | 461454.19 | 540634.38 | 501686.78 | 955179.06 | 520193.75 | 765084 | 671290.94 | 701163.12 | 548132.06 | 840572.5 | 295982.69 | 2 | 537484.12 | 838165.12 | 2 | 929937.69 | 8 | 11 | 12 | 10 | 12 | 11 | 1.129481727 | 0.931266549 | 0.851816309 | 0.702329232 |
| Q5XI13 | Grwd1 | Glutamate-rich WD repeat-containing protein 1 | 134.73 | 5.62 | 49169 | 2 | 1693953.38 | 1693639.88 | 1935843.38 | 1965931.5 | 867787.06 | 1651901.38 | 1995974.25 | 1744796 | 2388617.75 | 1311665.12 | 1594165 | 2516848 | 460272.72 | 525323 | 1534498.75 | 2626277.5 | 2523363.75 | 1564935.12 | 1929545.88 | 1415519.88 | 7 | 10 | 7 | 10 | 7 | 10 | 0.812919125 | 0.729257742 | 1.233229218 | 1.10631172 |
| Q9D8T2 | Gsdmdc1 | Gasdermin-D | 428.78 | 18.48 | 53238 | 7 | 11444688 | 13632972 | 9301002 | 10504442 | 13663193 | 7496365 | 11434229 | 11263532 | 13909362 | 11235216 | 6007067 | 9754973 | 7128110 | 5183546 | 7426006 | 11613425 | 5277176 | 8636320 | 7202401 | 9340203 | 1 | 4 | 1 | 6 | 1 | 6 | 0.606352644 | 0.641498616 | 0.718568503 | 0.760218833 |
| Q2NL51 | Gsk3a | Glycogen synthase kinase-3 alpha | 176.17 | 6.53 | 51661 | 3 | 1965488.75 | 1964893 | 2245977 | 2170706.25 | 1974878.62 | 2252921 | 1456064.12 | 2582451.5 | 2347536 | 1776279.75 | 1822748.75 | 2635793.5 | 2802297.75 | 3040403 | 3108643.5 | 516002.91 | 2155114 | 2707450.5 | 1431427 | 2579636.75 | 5 | 12 | 3 | 11 | 5 | 12 | 1.299162928 | 1.287523914 | 0.909676656 | 0.901526994 |
| P13020 | Gsn | Gelsolin | 1466.53 | 35.51 | 85942 | 23 | 8294296 | 9339826 | 12328717 | 11658818 | 11647412 | 9077644 | 9424622 | 16252744 | 13393470 | 14475614 | 6819413.5 | 8005125 | 8884850 | 6625611 | 9994110 | 7612856 | 6150248 | 13387661 | 12479958 | 11701228 | 3 | 11 | 2 | 6 | 3 | 11 | 0.757083055 | 0.643987113 | 0.96363522 | 0.819683731 |
| Q8R050 | Gspt1 | Eukaryotic peptide chain release factor GTP-binding subunit ERF3A | 248.55 | 10.53 | 68625 | 5 | 6012879.5 | 7505538.5 | 5765619.5 | 9179966 | 5185937 | 6935850.5 | 5619667 | 6001038.5 | 4420308.5 | 8029842 | 12633895 | 10799559 | 8682382 | 10345242 | 8190195 | 6150920 | 9236923 | 6795569.5 | 6910118.5 | 7122107 | 2 | 8 | 0 | 7 | 2 | 8 | 1.505241087 | 1.63355863 | 1.076246717 | 1.167993705 |
| P47791 | Gsr | Glutathione reductase, mitochondrial | 566.92 | 17.4 | 53663 | 7 | 3867530 | 5218031.5 | 4319667 | 4161616.75 | 4477000.5 | 6024772.5 | 5336604 | 3577719.25 | 4582224 | 4882817.5 | 6568453 | 7794508.5 | 7674800 | 8705547 | 7990556 | 3867552.5 | 3493894.5 | 6115025 | 5559929.5 | 6028315 | 0 | 9 | 0 | 10 | 0 | 10 | 1.757128268 | 1.587184341 | 1.137039189 | 1.02706833 |
| P46413 | Gss | Glutathione synthetase | 109.24 | 6.33 | 52345 | 2 | 2 | 567246.56 | 505675 | 2 | 383269.34 | 2 | 558727.06 | 2 | 349759.44 | 2 | 2959294.75 | 624753.31 | 2 | 2104725 | 195506.12 | 340885.44 | 4723193.5 | 2 | 63158.21 | 2 | 7 | 11 | 5.5 | 11 | 7 | 11 | 4.040861 | 6.476972765 | 3.520985515 | 5.643680218 |
| P00502 | Gsta1 | Glutathione S-transferase alpha-1 | 52.71 | 4.05 | 25607 | 1 | 2 | 2 | 109434.78 | 2 | 2 | 2 | 2 | 2 | 2 | 2 | 2 | 701278.75 | 2 | 2 | 401080.5 | 2 | 2 | 2 | 2 | 2 | 9 | 10 | 7.5 | 12.5 | 9 | 12.5 | 10.07252603 | 110236.525 | 9.137195E-05 | 1 |
| Q9DCM2 | Gstk1 | Glutathione S-transferase kappa 1 | 79.14 | 13.72 | 25704 | 2 | 1520163.5 | 1437268.75 | 1309692.88 | 679000.31 | 1472444.38 | 733103.25 | 1192299.12 | 1367459.88 | 1321582.12 | 1324225.5 | 219745.61 | 609774.75 | 967223.62 | 495028.44 | 1125010.5 | 1617395.62 | 190076 | 990873.38 | 469045.38 | 1113458.75 | 2 | 7 | 2 | 7 | 2 | 7 | 0.532327764 | 0.575344815 | 0.6825273 | 0.737681876 |
| P10649 | Gstm1 | Glutathione S-transferase Mu 1 | 381.51 | 33.94 | 25970 | 6 | 4878611 | 4387165.5 | 6626978 | 6160112 | 5766978 | 4419559 | 4380431 | 7082468.5 | 6451270 | 6970090 | 3908367 | 3623562.75 | 4291468 | 4691938 | 4830668 | 5762904 | 5095115.5 | 7811414 | 7530384 | 7803221.5 | 2 | 6 | 4 | 6 | 4 | 6 | 0.767294143 | 0.728437618 | 1.222258413 | 1.160362053 |
| P48774 | Gstm5 | Glutathione S-transferase Mu 5 | 385.52 | 28.57 | 26635 | 6 | 6732169.5 | 5669743 | 7112906 | 6233777 | 9932112 | 4723030 | 8137636 | 4022443 | 10034688 | 5599247.5 | 1826832.38 | 3154184 | 3607431.75 | 1216378 | 2951521.5 | 6732161.5 | 3495233 | 5409493 | 5282760 | 5224766 | 0 | 2 | 0 | 9 | 0 | 9 | 0.357513865 | 0.392297265 | 0.732732486 | 0.804021826 |
| O09131 | Gsto1 | Glutathione S-transferase omega-1 | 179.66 | 20.83 | 27498 | 4 | 6389714.5 | 5537287.5 | 1859238.75 | 1752513.12 | 2151074.75 | 6295822.5 | 5941819 | 1370236.5 | 1331644.25 | 1581312.88 | 5644732.5 | 2721196.75 | 3614751 | 3321967.5 | 3962320.5 | 2432520 | 1430321.75 | 1670656.5 | 2010452.88 | 2900621.5 | 9 | 8 | 10 | 11 | 10 | 11 | 1.089042108 | 1.166101356 | 0.590428141 | 0.632206093 |
| Q9D902 | Gtf2e2 | General transcription factor IIE subunit 2 | 58.55 | 4.11 | 33047 | 1 | 2 | 493009.44 | 386220.06 | 2 | 2 | 2 | 2 | 2 | 2 | 2 | 2 | 319281.44 | 2 | 2 | 2 | 434069.72 | 554025.69 | 2 | 305408.28 | 833553.75 | 9 | 6.5 | 10 | 2.5 | 10 | 6.5 | 0.363144391 | 31928.944 | 2.419214693 | 212705.944 |
| Q01750 | Gtf2f2 | General transcription factor IIF subunit 2 | 78.92 | 8.43 | 28349 | 2 | 1333242.25 | 1784199.12 | 1534733.25 | 744327 | 2419774.25 | 925616.38 | 2219493.25 | 2249330 | 2719279.25 | 1991245.5 | 852406.88 | 1877755.5 | 722741.81 | 515966.28 | 1570481.25 | 1563946.25 | 2 | 1125516.25 | 1408031 | 1409183 | 8 | 8 | 2 | 4 | 8 | 8 | 0.7086945 | 0.548181222 | 0.704514348 | 0.544947839 |
| Q9DBA9 | Gtf2h1 | General transcription factor IIH subunit 1 | 81.27 | 2.38 | 61851 | 1 | 211696.97 | 2 | 2 | 192252.75 | 170567.78 | 353954.16 | 190181.2 | 171176.59 | 2 | 147593.89 | 111285.29 | 202966.14 | 2 | 2 | 2 | 246981.03 | 2 | 2 | 2 | 205939.69 | 9 | 12 | 6.5 | 9.5 | 9 | 12 | 0.546989852 | 0.364184233 | 0.788354692 | 0.524884233 |
| D2XV59 | Gtpbp1 | GTP-binding protein 1 | 116.55 | 3.74 | 72489 | 2 | 2191963 | 1774940.5 | 2265729.5 | 2107927 | 2051034 | 1923741.62 | 2187311.5 | 2250741.75 | 2014060.38 | 1786190.75 | 1310647.88 | 1776831.25 | 1574105.75 | 876033 | 1553477.38 | 2358591.5 | 1299925.75 | 1944763.5 | 2167051.25 | 1186078.25 | 1 | 9 | 0 | 10 | 1 | 10 | 0.682387636 | 0.697801925 | 0.861889932 | 0.881358956 |
| Q99ME9 | Gtpbp4 | Nucleolar GTP-binding protein 1 | 619.95 | 22.87 | 74113 | 12 | 739726.38 | 473180.94 | 744325.56 | 2 | 751126.81 | 1230259.75 | 1076656.5 | 320153.53 | 159952.28 | 56230.48 | 4754575.5 | 2967008.5 | 2081422 | 1541569.5 | 910232.94 | 1758115.88 | 1612311 | 58145.13 | 2 | 2400529 | 0 | 8.5 | 2 | 9 | 2 | 9 | 4.524804972 | 4.310137164 | 2.152261654 | 2.0501531 |
| P12265 | Gusb | Beta-glucuronidase | 1200.82 | 39.2 | 74195 | 22 | 85045962 | 82584178 | 146249298 | 143765138 | 155878242 | 70060706 | 76722298 | 78322954 | 90865146 | 78130858 | 23007158 | 26999788 | 19859798 | 23542568 | 23673456 | 83071410 | 70680890 | 68439378 | 67650346 | 77332322 | 0 | 1 | 0 | 7 | 0 | 7 | 0.19083686 | 0.297087503 | 0.598468933 | 0.931673479 |
| Q9Z1E4 | Gys1 | Glycogen [starch] synthase, muscle | 95.5 | 2.57 | 83927 | 2 | 1513803.25 | 1433700.25 | 1114302 | 2 | 1364868.75 | 650580.5 | 1426672.75 | 692771.5 | 1493387.75 | 1127368.88 | 2 | 1165333.5 | 1166827.5 | 2 | 900129.81 | 1521140.75 | 2 | 420106.5 | 2 | 548776.12 | 6 | 8 | 8 | 5 | 8 | 8 | 0.59563067 | 0.599596716 | 0.458849442 | 0.461904721 |
| P10922 | H1f0 | Histone H1.0 | 413.84 | 28.35 | 20861 | 7 | 20894814 | 25386976 | 24833822 | 27617280 | 18044110 | 29353184 | 25204082 | 25838930 | 20350140 | 32344114 | 22351542 | 17115322 | 24746322 | 29858886 | 23969060 | 15662832 | 24816274 | 27106628 | 25053628 | 32486970 | 11 | 11 | 7 | 10 | 11 | 11 | 1.010825162 | 0.886924133 | 1.071498068 | 0.940160109 |
| P01900 | H2-D1 | H-2 class I histocompatibility antigen, D-D alpha chain | 933.12 | 52.05 | 41110 | 17 | 1487373.12 | 3204119.75 | 3367785.5 | 3337204 | 2618029.5 | 1769629.38 | 2554930.5 | 4189572.25 | 3576301.5 | 3958085.5 | 9019624 | 5585289 | 6368448 | 4615714 | 5902007.5 | 2772704.5 | 2786644.5 | 2555594.5 | 3767217.75 | 3963066.5 | 0 | 10 | 0 | 12 | 0 | 12 | 2.247033845 | 1.962242263 | 1.130630014 | 0.987332702 |
| P01902 | H2-K1 | H-2 class I histocompatibility antigen, K-D alpha chain | 1052.88 | 46.2 | 41490 | 19 | 28126020 | 27930064 | 29021526 | 30811482 | 33994242 | 26288954 | 35469082 | 37679242 | 37000442 | 32977470 | 26112974 | 25991176 | 24484522 | 21060914 | 23928656 | 32555414 | 23674954 | 10862026 | 22173152 | 22935142 | 0 | 4 | 0 | 1 | 0 | 4 | 0.811152506 | 0.717634835 | 0.748586818 | 0.662282337 |
| P01897 | H2-L | H-2 class I histocompatibility antigen, L-D alpha chain | 728.66 | 35.36 | 40711 | 12 | 5161332 | 5778816 | 8072738.5 | 8655872 | 8638708 | 5326137 | 6379131 | 9087495 | 8770654 | 10312344 | 7441307 | 7780787 | 16506488 | 10222446 | 11784644 | 6209852 | 8442472 | 8335797 | 8749935 | 10389534 | 6 | 7 | 7 | 12 | 7 | 12 | 1.48001712 | 1.347577342 | 1.160301009 | 1.056471123 |
| P14430 | H2-Q8 | H-2 class I histocompatibility antigen, Q8 alpha chain | 165.67 | 9.51 | 37411 | 3 | 1216790.5 | 807824 | 1095987.62 | 1508230.5 | 1204716.62 | 1210696.5 | 1146037.38 | 1410782.5 | 1553298.75 | 1602475.75 | 1644485.75 | 1334859.5 | 2262272.75 | 1432180.62 | 1650988.12 | 1318081.62 | 1003594.69 | 1228991.38 | 1032187.81 | 1207827.62 | 2 | 12 | 5 | 5 | 5 | 12 | 1.427053479 | 1.202432035 | 0.992651794 | 0.836406157 |
| Q9QZQ8 | H2afy | Core histone macro-H2A.1 | 591.89 | 39.52 | 39735 | 9 | 12152362 | 12422561 | 14022576 | 9076564 | 10097530 | 11339568 | 10689366 | 11672237 | 11306751 | 12315130 | 10611465 | 10189519 | 9095787 | 7780166.5 | 9675806 | 8608321 | 6325368.5 | 11665353 | 7902335 | 11678567 | 6 | 4 | 0 | 7 | 6 | 7 | 0.819654454 | 0.82606808 | 0.799353836 | 0.805608614 |
| P0C0S6 | H2afz | Histone H2A.Z | 549.98 | 56.25 | 13553 | 8 | 16813934 | 11819048 | 27578864 | 12585324 | 27329600 | 7224854 | 20417908 | 24799534 | 27170944 | 11975736 | 5479104 | 13164898 | 4863453.5 | 3262388.25 | 13167663 | 26493662 | 12516676 | 12188558 | 12548880 | 18186318 | 4 | 9 | 4 | 12 | 4 | 12 | 0.415467062 | 0.43605146 | 0.852354594 | 0.894584671 |
| P02301 | H3f3c | Histone H3.3C | 441.36 | 33.09 | 15315 | 8 | 1884889218 | 1358088962 | 1919138306 | 2201927682 | 1533722754 | 1958693250 | 1519249538 | 1643702914 | 1524756738 | 2015931138 | 617554690 | 716589954 | 907555074 | 1032108418 | 938035202 | 1477674242 | 2025817730 | 1718867842 | 1748238850 | 1727472386 | 0 | 11 | 0 | 11 | 0 | 11 | 0.473359594 | 0.486225022 | 0.97755663 | 1.004125617 |
| Q9D3B1 | Hacd2 | Very-long-chain (3R)-3-hydroxyacyl-CoA dehydratase 2 | 48.35 | 3.94 | 28402 | 1 | 1482477.5 | 687002.81 | 572244.38 | 2422366 | 1470149.5 | 2623912.5 | 1696528.38 | 2018779.25 | 1737423.12 | 2153048 | 1671223 | 1581024.38 | 2469854.75 | 3652611.75 | 2474397.25 | 1749166 | 2941869.5 | 2392849 | 3196787.25 | 436611.72 | 2 | 7 | 12 | 9 | 12 | 9 | 1.786053985 | 1.158305841 | 1.615450023 | 1.047664412 |
| Q8K2C9 | Hacd3 | Very-long-chain (3R)-3-hydroxyacyl-CoA dehydratase 3 | 82.18 | 4.97 | 43131 | 2 | 1510929 | 175560.22 | 232013.08 | 1562414.62 | 372875.81 | 2 | 417287.03 | 357978.34 | 548972.88 | 887509.81 | 2875743.75 | 2242437.75 | 2737799.5 | 3237478.25 | 2953928.5 | 364393.75 | 1573279.12 | 547192.25 | 493942.44 | 495456.19 | 0 | 9 | 0 | 9 | 0 | 9 | 3.645081283 | 6.351254603 | 0.901518061 | 1.57082114 |
| Q9QXE0 | Hacl1 | 2-hydroxyacyl-CoA lyase 1 | 83.41 | 3.61 | 63660 | 2 | 883671.56 | 505213.53 | 1353213.75 | 1226554.62 | 1074169.25 | 1082947.62 | 591958.44 | 926444.88 | 972567.25 | 1119172.25 | 82597.95 | 211692.42 | 300034.09 | 943228.75 | 657006 | 1061380.25 | 418947.97 | 1051176 | 417299.84 | 1204110.5 | 3 | 7 | 3 | 11 | 3 | 11 | 0.435184684 | 0.467614941 | 0.823529757 | 0.884899751 |
| Q61425 | Hadh | Hydroxyacyl-coenzyme A dehydrogenase, mitochondrial | 273.26 | 19.11 | 34464 | 5 | 8126321 | 7285312.5 | 7343973 | 5944639 | 9603936 | 4821613 | 8864243 | 29103152 | 9550411 | 4565170.5 | 6355870.5 | 7596669 | 5966863 | 3777246.5 | 6580134.5 | 5111931 | 5177687 | 4138442.5 | 5137643.5 | 5514235.5 | 6 | 0 | 8 | 8 | 8 | 8 | 0.790430243 | 0.532062243 | 0.654757223 | 0.440736674 |
| Q8BMS1 | Hadha | Trifunctional enzyme subunit alpha, mitochondrial | 1276.75 | 28.57 | 82670 | 19 | 18769038 | 13916048 | 19519222 | 19101232 | 18721064 | 17068610 | 16815420 | 15782466 | 15145501 | 18298116 | 12225115 | 9658771 | 12278860 | 10257934 | 13894661 | 16628005 | 19154420 | 17473894 | 15402034 | 13025852 | 0 | 7 | 0 | 12 | 0 | 12 | 0.647756756 | 0.701663599 | 0.90733407 | 0.982843147 |
| Q99JY0 | Hadhb | Trifunctional enzyme subunit beta, mitochondrial | 634.39 | 27.16 | 51386 | 13 | 8528782 | 12196548 | 14580718 | 10857109 | 12967173 | 11115342 | 9028592 | 8946837 | 11359228 | 12341138 | 8119577 | 7226818.5 | 11018466 | 9293246 | 10626496 | 7422196.5 | 4674442 | 11919316 | 9459889 | 11989730 | 4 | 5 | 6 | 10 | 6 | 10 | 0.782755711 | 0.87674951 | 0.768904444 | 0.861234974 |
| O35952 | Hagh | Hydroxyacylglutathione hydrolase, mitochondrial | 95.27 | 6.15 | 34109 | 2 | 2058079.88 | 3244310.75 | 2023365.38 | 639471.12 | 4090580.5 | 419975.44 | 3185438 | 2520431 | 4042854.5 | 442917.22 | 2 | 1412091.38 | 334741.03 | 2 | 310937.88 | 2123825.25 | 2 | 336291.09 | 207979.56 | 682853.5 | 1 | 4 | 2 | 4 | 2 | 4 | 0.170687386 | 0.193917143 | 0.27795329 | 0.315781437 |
| P35492 | Hal | Histidine ammonia-lyase | 384.11 | 13.85 | 72258 | 5 | 125296.52 | 331159.09 | 2 | 2 | 2 | 187163.41 | 79028.74 | 2 | 2 | 2 | 2 | 153887.03 | 2 | 2 | 2 | 327705.94 | 2 | 2 | 2 | 2 | 10 | 10 | 10 | 11 | 10 | 11 | 0.337147805 | 0.578122087 | 0.717944144 | 1.231090224 |
| Q61035 | Hars | Histidine--tRNA ligase, cytoplasmic | 501.97 | 20.04 | 57432 | 10 | 8195298.5 | 8427862 | 9082135 | 8202191.5 | 10430024 | 7761820 | 6932820 | 8614113 | 8706713 | 8212096 | 7331415 | 7572221.5 | 7208220.5 | 6486548 | 7574814.5 | 9002691 | 8001197 | 9223863 | 10011732 | 9345822 | 0 | 10 | 4 | 3 | 4 | 10 | 0.815860401 | 0.899214809 | 1.028143077 | 1.133185874 |
| Q99KK9 | Hars2 | Probable histidine--tRNA ligase, mitochondrial | 177.01 | 7.33 | 56985 | 4 | 495567.41 | 723605.94 | 2 | 1172792 | 1175641.62 | 645177.38 | 787683.25 | 638583.81 | 794762.44 | 1024020.94 | 231236 | 199972.72 | 121326.41 | 217666.75 | 443696.22 | 447109 | 473885.19 | 489683.28 | 1175450.25 | 1149325.88 | 5 | 10 | 0 | 10 | 5 | 10 | 0.340255367 | 0.312037792 | 1.047046812 | 0.960214613 |
| Q8BY71 | Hat1 | Histone acetyltransferase type B catalytic subunit | 186.53 | 9.38 | 49278 | 3 | 2780696.75 | 2782843.75 | 2714967.5 | 2497496 | 1783431.25 | 2627052.25 | 2559159.75 | 2298581.75 | 2353853.25 | 1518116.25 | 2622905 | 3065340.75 | 2286710.25 | 1917012.5 | 1989245.75 | 3243627 | 2468638 | 2115085.5 | 2118878.5 | 1951031.75 | 10 | 9 | 12 | 11 | 12 | 11 | 0.945999085 | 1.046179619 | 0.94727673 | 1.047592566 |
| Q61191 | Hcfc1 | Host cell factor 1 | 527.19 | 8.7 | 210437 | 10 | 4379677.5 | 3908491.5 | 4112907.25 | 4237219.5 | 4337439 | 4437194 | 4425588.5 | 3861110 | 3521111 | 3897759 | 3397782.5 | 3646822.75 | 2666300.75 | 2677242.5 | 3413631.5 | 4909865 | 5375938.5 | 3680031 | 4844359 | 4383987.5 | 0 | 5 | 1 | 6 | 1 | 6 | 0.753336185 | 0.784489218 | 1.105762505 | 1.151489573 |
| P08103 | Hck | Tyrosine-protein kinase HCK | 534.38 | 22.71 | 59129 | 11 | 3192324.5 | 3391359 | 4944170.5 | 2946018.5 | 2983472.25 | 3623149 | 1658407 | 2526531.25 | 2826319.5 | 4905184 | 3762606.5 | 3314935 | 3971553.75 | 4503944.5 | 3932940 | 3250785.5 | 2826909 | 2943183.5 | 2673510.75 | 3982397.25 | 6 | 7 | 6 | 10 | 6 | 10 | 1.116205244 | 1.253957074 | 0.898005179 | 1.008828756 |
| P49710 | Hcls1 | Hematopoietic lineage cell-specific protein | 451.32 | 18.31 | 54240 | 8 | 12014772 | 13816907 | 12499445 | 10183442 | 9893546 | 18547882 | 18216578 | 16460850 | 13372322 | 15995775 | 26479334 | 29366766 | 22933244 | 16679930 | 18042794 | 12454834 | 16105158 | 22432142 | 15169818 | 14916163 | 0 | 2 | 4 | 9 | 4 | 9 | 1.943258635 | 1.374226735 | 1.38813107 | 0.981653596 |
| O09106 | Hdac1 | Histone deacetylase 1 | 255.22 | 14.11 | 55075 | 5 | 7375572.5 | 5336325 | 5731158.5 | 4107336.75 | 6546624 | 4256158 | 5672282.5 | 2596953 | 4829946 | 2568816.5 | 2909645 | 4931517 | 2187275 | 1347213.12 | 3478782.5 | 5465096 | 2834799 | 3623264 | 3489976 | 2148584.5 | 1 | 2 | 8 | 10 | 8 | 10 | 0.510513939 | 0.745548902 | 0.60355739 | 0.881428528 |

| | | | | | | | | | | | | | | | | | | | | | | | | | | | | | | | | | | | | |
|---|---|---|---|---|---|---|---|---|---|---|---|---|---|---|---|---|---|---|---|---|---|---|---|---|---|---|---|---|---|---|---|---|---|---|---|---|
| P70288 | Hdac2 | Histone deacetylase 2 | 246.36 | 12.09 | 55302 | 4 | 2242362 | 1926343 | 2494636.25 | 2854895.75 | 2449081 | 2621280.5 | 2361100.75 | 2269399.25 | 2332764 | 2541165.25 | 1723292.5 | 1615550.75 | 2481558.5 | 1603741.75 | 1848817 | 2109370 | 2440860.75 | 2422634.5 | 2265601.5 | 2210398 | 3 | 8 | 3 | 6 | 3 | 8 | 0.774857031 | 0.764735483 | 0.956677574 | 0.944180999 |
| O88895 | Hdac3 | Histone deacetylase 3 | 41.57 | 4.01 | 48364 | 1 | 440094.06 | 388544.34 | 423147.91 | 558941.38 | 466359.94 | 510164.78 | 485861.22 | 2 | 426612.16 | 2 | 2 | 2 | 371941.31 | 357183.38 | 399400.06 | 614985.56 | 2 | 427655.41 | 626872.19 | 244282.42 | 1 | 12 | 8 | 9 | 8 | 12 | 0.49560181 | 0.793262552 | 0.840458468 | 1.345241716 |
| P51859 | Hdgf | Hepatoma-derived growth factor | 579.17 | 45.57 | 26269 | 11 | 23452478 | 23666196 | 22536396 | 22212260 | 15063241 | 25706892 | 27468902 | 28295384 | 19171326 | 22117322 | 17242266 | 17957466 | 15456733 | 15638646 | 14266881 | 30613244 | 27651012 | 31272702 | 33811758 | 22608278 | 4 | 2 | 0 | 4 | 4 | 4 | 0.753404113 | 0.656256486 | 1.364969743 | 1.188963839 |
| Q3UMU9 | Hdgfl2 | Hepatoma-derived growth factor-related protein 2 | 161.69 | 4.63 | 74291 | 3 | 2783818.75 | 2820569.75 | 2045406.75 | 3762674.5 | 2677877.5 | 3293042.5 | 3003814 | 1115159.25 | 2285499.5 | 2884768.25 | 1798470 | 2 | 98147.8 | 245907.25 | 2 | 3688232.75 | 3108678.5 | 2508590.25 | 3406937.75 | 2337900 | 0 | 11 | 1 | 7 | 1 | 11 | 0.152056512 | 0.170281416 | 1.068131181 | 1.196153246 |
| Q3UGR5 | Hdhd2 | Haloacid dehalogenase-like hydrolase domain-containing protein 2 | 45.79 | 3.09 | 28730 | 1 | 739995.38 | 589639.62 | 547140.94 | 229147.39 | 1230991.38 | 361222.84 | 666331.75 | 648347.38 | 1228357.38 | 297468.06 | 2 | 267936.94 | 2 | 2 | 2 | 610764.56 | 94038.52 | 138035.58 | 2 | 258944.02 | 1 | 4 | 0 | 2 | 1 | 4 | 0.08029721 | 0.083687618 | 0.330180654 | 0.34412195 |
| Q8VDJ3 | Hdlbp | Vigilin | 1289.04 | 21.61 | 141743 | 22 | 20479294 | 20021328 | 17110580 | 17320186 | 15769030 | 19638628 | 19100124 | 20516252 | 15206290 | 18303590 | 18403114 | 16535029 | 14352896 | 14015347 | 12338368 | 19305086 | 19117682 | 18201614 | 16998852 | 17977636 | 4 | 12 | 3 | 9 | 4 | 12 | 0.834006675 | 0.815446004 | 1.00992776 | 0.987451997 |
| Q8BQM4 | Heatr3 | HEAT repeat-containing protein 3 | 148.87 | 6.04 | 74305 | 3 | 1283336.5 | 1126250.5 | 755939.81 | 1002260.31 | 1108846.62 | 1341896.88 | 1711612 | 1094354.62 | 1375081.88 | 974704.12 | 1279773 | 1246054.25 | 1229241.75 | 1020067.12 | 1088303.88 | 1498106.25 | 1043133.75 | 1056936.38 | 1158782 | 1062526.88 | 9 | 10 | 8 | 9 | 9 | 10 | 1.11120845 | 0.902394012 | 1.102878378 | 0.895629298 |
| Q69ZR2 | Hectd1 | E3 ubiquitin-protein ligase HECTD1 | 103.36 | 1.11 | 290086 | 2 | 630236.75 | 274460.47 | 116132.22 | 206269.56 | 160482.44 | 371697.06 | 283039.25 | 160387.36 | 710290.25 | 128325.97 | 289830.72 | 383274.19 | 197592.52 | 234953.91 | 397208.44 | 143646.58 | 163344.69 | 2 | 2 | 67495.87 | 8 | 3 | 10 | 3 | 10 | 3 | 1.083078612 | 0.908764304 | 0.269887683 | 0.226451053 |
| Q6NVF4 | Helb | DNA helicase B | 52.15 | 0.74 | 121471 | 1 | 1072002.5 | 664743.25 | 1101356.62 | 1265751.5 | 1205928.5 | 1226576.38 | 866887 | 1177811.88 | 1014742.75 | 1304429.62 | 852117.06 | 1016990.56 | 906603.75 | 768177.94 | 1065854.88 | 764295.06 | 2370859.5 | 1135513.75 | 958287.44 | 876951.88 | 5 | 11 | 5 | 9 | 5 | 11 | 0.868160664 | 0.824575149 | 1.149935573 | 1.092203708 |
| Q60848 | Hells | Lymphocyte-specific helicase | 180.68 | 4.87 | 95126 | 3 | 1430834.75 | 665003.06 | 1706228.62 | 627069.38 | 2098615 | 2 | 1143350.75 | 799017.75 | 2096970.12 | 2 | 351503.38 | 615685.5 | 495600 | 2 | 295992 | 1185172.5 | 804784.62 | 644398.44 | 640603.69 | 2 | 0 | 6 | 9 | 12 | 9 | 12 | 0.269431682 | 0.435431347 | 0.501698264 | 0.810765899 |
| E9QAM5 | Helz2 | Helicase with zinc finger domain 2 | 45.01 | 0.37 | 331563 | 1 | 164677.91 | 2 | 91465.23 | 2 | 2 | 216154.5 | 187524.53 | 137161.48 | 2 | 180864.16 | 387556.03 | 350597.19 | 321807.91 | 262155.53 | 2 | 2 | 73391.45 | 185616.94 | 2 | 136731.84 | 3.5 | 10 | 4.5 | 6 | 4.5 | 10 | 5.161519028 | 1.831933547 | 1.544975829 | 0.548344981 |
| Q4U2R1 | Herc2 | E3 ubiquitin-protein ligase HERC2 | 54.45 | 0.23 | 527456 | 1 | 2 | 2 | 2 | 285692.94 | 282300.97 | 2 | 206199 | 188737.19 | 121040.45 | 273036.12 | 2 | 188938.59 | 315730.28 | 199479.69 | 83192.73 | 221203.78 | 327394 | 280455.03 | 2 | 255789.95 | 9.5 | 9.5 | 12.5 | 6.5 | 12.5 | 9.5 | 1.386167984 | 0.997881573 | 1.90993826 | 1.374935952 |
| P29416 | Hexa | Beta-hexosaminidase subunit alpha | 650.35 | 31.82 | 60613 | 12 | 18390730 | 17412714 | 40341382 | 27808360 | 41761094 | 14617442 | 19282376 | 26860122 | 27241192 | 24376154 | 6862828.5 | 11301007 | 12451912 | 10431678 | 9951231 | 18990322 | 10419905 | 20485608 | 21849054 | 20666060 | 0 | 8 | 0 | 7 | 0 | 8 | 0.349990794 | 0.45381641 | 0.634192812 | 0.822327645 |
| P20060 | Hexb | Beta-hexosaminidase subunit beta | 798.43 | 33.58 | 61116 | 17 | 31394074 | 31968944 | 61447410 | 45582190 | 62474394 | 24922284 | 33777674 | 37064490 | 46123190 | 32141602 | 10747997 | 12170861 | 15381739 | 11126127 | 14927591 | 25674974 | 21994412 | 30479198 | 32334876 | 28761408 | 0 | 2 | 0 | 5 | 0 | 5 | 0.276356511 | 0.369790243 | 0.597958753 | 0.800123405 |
| Q99L13 | Hibadh | 3-hydroxyisobutyrate dehydrogenase, mitochondrial | 547.92 | 45.07 | 35440 | 10 | 7730235.5 | 7816420.5 | 4946936 | 7270576.5 | 5628029 | 8854423 | 7549463 | 5830091.5 | 8184196.5 | 8137403.5 | 3881021 | 4640570 | 8516468 | 7009161 | 6977092 | 7964389.5 | 6685807 | 7357341.5 | 7667471 | 7329318.5 | 9 | 9 | 6 | 7 | 9 | 9 | 0.929088659 | 0.804664695 | 1.108172875 | 0.959765873 |
| Q8QZS1 | Hibch | 3-hydroxyisobutyryl-CoA hydrolase, mitochondrial | 389.9 | 21.3 | 43038 | 7 | 2704171 | 3453617 | 2910523.5 | 2499818 | 4276334.5 | 2855802.5 | 1640376.62 | 2747243 | 7103248 | 2175878 | 3289915 | 2664352 | 4775891 | 3381175 | 7127442.5 | 3184988.5 | 4075276 | 4436125.5 | 3249766.5 | 5050788.5 | 8 | 5 | 6 | 5 | 8 | 5 | 1.340454022 | 1.285441891 | 1.26207772 | 1.210282146 |
| Q9JLZ6 | Hic2 | Hypermethylated in cancer 2 protein | 47.63 | 1.62 | 66766 | 1 | 2 | 2 | 2 | 2 | 2 | 2 | 2 | 2 | 2 | 2 | 38468358 | 10930912 | 4758361 | 3845766.25 | 5379237 | 2 | 2 | 2 | 2 | 2 | 0 | 12.5 | 0 | 12.5 | 0 | 12.5 | 6338263.425 | 6338263.425 | 1 | 1 |
| Q8VH49 | Higd1a | HIG1 domain family member 1A, mitochondrial | 68.95 | 23.66 | 10301 | 1 | 577207.06 | 452386.66 | 482847.44 | 381128.44 | 377430.38 | 528527.88 | 503848.69 | 456497.84 | 505544.22 | 417457.69 | 2 | 373046.38 | 301787.62 | 436911.66 | 321891.28 | 473667.81 | 553154.69 | 626210.88 | 708117.38 | 387562.12 | 2 | 6 | 1 | 8 | 2 | 8 | 0.631280913 | 0.594408149 | 1.210353547 | 1.139657476 |
| P62959 | Hint1 | Histidine triad nucleotide-binding protein 1 | 218.44 | 42.06 | 13777 | 4 | 22125452 | 24803650 | 22354702 | 28414556 | 29091474 | 27252274 | 23187548 | 27717966 | 26067282 | 30155664 | 29916792 | 32315754 | 29243100 | 30064152 | 30751768 | 25894136 | 32449836 | 29612064 | 33462100 | 30526592 | 0 | 2 | 3 | 5 | 3 | 5 | 1.201133886 | 1.133284225 | 1.198398351 | 1.130703215 |
| Q9D0S9 | Hint2 | Histidine triad nucleotide-binding protein 2, mitochondrial | 37.97 | 7.98 | 17320 | 1 | 2 | 2 | 609161.81 | 971882.44 | 2 | 727908.94 | 2 | 650907.88 | 2 | 1049154.75 | 1030310.69 | 924418.44 | 1463714.25 | 1999049.25 | 1878374 | 2 | 691554.19 | 2 | 1287734 | 1373827.62 | 1 | 8 | 2 | 10 | 2 | 10 | 4.614569733 | 3.004917644 | 2.120817988 | 1.381035234 |
| Q8BLH7 | Hirp3 | HIRA-interacting protein 3 | 238.33 | 7.15 | 65215 | 4 | 2250259.5 | 2780248.25 | 2561791.25 | 2690298 | 2165592 | 2892866.75 | 2533879.75 | 1973267.38 | 756051.94 | 1853524.12 | 1473181 | 1430782.62 | 1592355.88 | 1702746.5 | 1110692.75 | 2579077 | 3188360.5 | 1902503.5 | 2208331.5 | 1569962.62 | 0 | 9 | 5 | 10 | 5 | 10 | 0.587214634 | 0.730275545 | 0.919670734 | 1.143726685 |
| P43275 | Hist1h1a | Histone H1.1 | 467.63 | 30.52 | 21785 | 7 | 10607071 | 7703118 | 12702602 | 9104466 | 7305950.5 | 14258250 | 10732127 | 10083209 | 6351289.5 | 10127272 | 7363134.5 | 6432036.5 | 6004747 | 9395768 | 5747916 | 5886061 | 13891404 | 8581303 | 10939766 | 8286547.5 | 4 | 12 | 3 | 10 | 4 | 12 | 0.736846026 | 0.677830191 | 1.003413392 | 0.923047512 |
| P43276 | Hist1h1b | Histone H1.5 | 778.19 | 37.67 | 22576 | 13 | 60207118 | 51768430 | 71274498 | 61031674 | 39095106 | 87583794 | 63127590 | 65127666 | 39802454 | 74140890 | 48215614 | 40305854 | 40563050 | 54194410 | 39211482 | 40147458 | 56176770 | 50116274 | 62154274 | 56235570 | 6 | 10 | 4 | 5 | 6 | 10 | 0.785139749 | 0.674658241 | 0.934551882 | 0.803045738 |
| P15864 | Hist1h1c | Histone H1.2 | 875.89 | 44.81 | 21267 | 13 | 32299800 | 25940726 | 50457262 | 44882902 | 29103660 | 44393618 | 33803166 | 41144130 | 24282954 | 48932190 | 18905598 | 14478212 | 20307532 | 28068560 | 18994380 | 22161572 | 38665734 | 34190898 | 39975618 | 34267218 | 1 | 12 | 1 | 8 | 1 | 12 | 0.551521146 | 0.523246493 | 0.926521839 | 0.879022149 |
| P43277 | Hist1h1d | Histone H1.3 | 882.47 | 41.18 | 22100 | 13 | 13118786 | 10964394 | 14939886 | 14891720 | 10408818 | 16772242 | 13754269 | 15114581 | 8831848 | 18142716 | 8136131.5 | 6684999 | 7556270 | 13132106 | 7765462.5 | 7608400.5 | 14999577 | 13126991 | 14401588 | 11006492 | 3 | 12 | 1 | 6 | 3 | 12 | 0.672769657 | 0.595945439 | 0.950553836 | 0.842009174 |
| P43274 | Hist1h1e | Histone H1.4 | 815.51 | 37.44 | 21977 | 12 | 25638356 | 24667444 | 34378074 | 29586482 | 21601474 | 35624114 | 27939804 | 33471868 | 19815118 | 34123090 | 17638468 | 17636206 | 17817926 | 24469886 | 17673866 | 19342680 | 29234046 | 28164300 | 29372002 | 27790486 | 1 | 12 | 1 | 6 | 1 | 12 | 0.700927867 | 0.63081296 | 0.985513436 | 0.88693099 |
| C0HKE1 | Hist1h2ab | Histone H2A type 1-B | 705.33 | 63.85 | 14135 | 9 | 9369727 | 7353176 | 8310492.5 | 21733916 | 3487914 | 28546268 | 6436209.5 | 5859032.5 | 5071870 | 13329752 | 24032666 | 3947014.75 | 10651450 | 29584748 | 5331437 | 7281843 | 59136934 | 18498186 | 14395320 | 13327219 | 9 | 7 | 12 | 6 | 12 | 7 | 1.463475987 | 1.24144881 | 2.241349051 | 1.901309033 |
| P10853 | Hist1h2bf | Histone H2B type 1-F/J/L | 1476.61 | 69.05 | 13936 | 21 | 7107562 | 2401588 | 7842684 | 10514171 | 2028339.38 | 17805834 | 2635951 | 4845540.5 | 1959792.12 | 5663646 | 12076098 | 4525456 | 7327556.5 | 14932581 | 3861387.75 | 5456912.5 | 61040694 | 16836758 | 9881230 | 10541165 | 8 | 4 | 9 | 5 | 9 | 5 | 1.429135849 | 1.298149133 | 3.470782238 | 3.152669464 |
| Q64478 | Hist1h2bh | Histone H2B type 1-H | 1477.79 | 69.05 | 13920 | 21 | 5480771 | 1583941.38 | 3327452.25 | 4037991.25 | 1737384.5 | 12563836 | 1885096.12 | 3683460.75 | 3144258.5 | 7091640 | 6059683.5 | 2875650.5 | 5087356.5 | 12575671 | 3125047.25 | 1995420.38 | 41430918 | 11557392 | 6177551.5 | 9275898 | 7 | 3 | 9 | 3 | 9 | 3 | 1.838462008 | 1.047768735 | 4.356703507 | 2.482954612 |
| P62804 | Hist1h4a | Histone H4 | 1276.1 | 65.05 | 11367 | 17 | 1869148290 | 1393522562 | 2368216322 | 2004370946 | 2118236418 | 1667366146 | 1820156674 | 2001113986 | 2066976514 | 1943279874 | 652726914 | 772401666 | 772940994 | 744262914 | 781346306 | 1502400002 | 1477421954 | 1507468034 | 1534740098 | 1533133954 | 0 | 5 | 0 | 5 | 0 | 5 | 0.381778939 | 0.392011861 | 0.774610988 | 0.795373091 |
| Q64522 | Hist2h2ab | Histone H2A type 2-B | 592.52 | 57.69 | 14013 | 7 | 2 | 1179681.12 | 987526.69 | 1016337.38 | 737369.5 | 847785.31 | 2 | 911520.19 | 630339.25 | 709092.62 | 2 | 440680.91 | 750812.56 | 593361.06 | 2 | 2 | 601571.81 | 2 | 2 | 1053628.5 | 5 | 6.5 | 6 | 7.5 | 6 | 7.5 | 0.455214602 | 0.575995047 | 0.422147789 | 0.534154736 |
| Q64524 | Hist2h2be | Histone H2B type 2-E | 811.3 | 42.06 | 13993 | 10 | 2 | 2 | 2 | 118342.77 | 2 | 2 | 167013.02 | 2 | 2 | 2 | 2 | 164922.73 | 2 | 2 | 2 | 646601.31 | 2 | 2 | 2 | 2 | 12 | 12 | 12 | 12 | 12 | 12 | 1.393575471 | 0.987484869 | 5.46349897 | 3.871424746 |
| P17710 | Hk1 | Hexokinase-1 | 1778.07 | 30.08 | 108303 | 28 | 47207126 | 49684014 | 50801102 | 54767726 | 48574598 | 49842558 | 43376034 | 46496154 | 46610750 | 45964134 | 23278056 | 21883030 | 31176500 | 32114470 | 31288544 | 44647010 | 40027282 | 48033654 | 46117506 | 44767790 | 0 | 1 | 0 | 8 | 0 | 8 | 0.556658799 | 0.601579158 | 0.89068707 | 0.962562306 |
| O08528 | Hk2 | Hexokinase-2 | 2031.6 | 42.09 | 102535 | 34 | 29434372 | 33259694 | 19197308 | 18115562 | 22962250 | 38629534 | 35195810 | 28614788 | 28867730 | 23850294 | 47154190 | 52867698 | 41636078 | 37123538 | 50293458 | 42736986 | 35367162 | 27197916 | 26433598 | 30796050 | 0 | 5 | 1 | 11 | 1 | 11 | 1.862864751 | 1.476396523 | 1.321727152 | 1.047522839 |
| Q3TRM8 | Hk3 | Hexokinase-3 | 2205.37 | 45.34 | 100101 | 31 | 69005834 | 73097810 | 49343098 | 67001110 | 70139610 | 74696466 | 67914618 | 64077450 | 65527166 | 64659506 | 67456930 | 63003658 | 66717742 | 68092418 | 59135202 | 77777242 | 78632770 | 72473090 | 72917066 | 80681922 | 7 | 2 | 10 | 2 | 10 | 2 | 0.98727428 | 0.962985534 | 1.164019125 | 1.135382133 |
| Q9D8V0 | Hm13 | Minor histocompatibility antigen H13 | 285.19 | 10.05 | 41748 | 5 | 5183118.5 | 7912295 | 8565135 | 5042682 | 7404219 | 9168869 | 11075718 | 9742800 | 11309677 | 10520672 | 17253918 | 19022720 | 19568800 | 13901836 | 19108150 | 7859244 | 7124779.5 | 10448673 | 9782531 | 10970510 | 0 | 5 | 0 | 7 | 0 | 7 | 2.603635059 | 1.71476855 | 1.353331063 | 0.891311375 |
| P22907 | Hmbs | Porphobilinogen deaminase | 195.89 | 13.85 | 39344 | 3 | 1661123.75 | 1888452.25 | 1658063.62 | 1760285.25 | 1721962.25 | 1487860.88 | 1377829.25 | 1564031.38 | 1611368.62 | 1240569.38 | 1379725.88 | 1116876 | 1406910.12 | 1137799.25 | 990228.12 | 1304303.38 | 1511543.5 | 1274538 | 1413697 | 1855293.25 | 0 | 4 | 4 | 12 | 4 | 12 | 0.694087194 | 0.828319336 | 0.846886612 | 1.01067279 |
| P17095 | Hmga1 | High mobility group protein HMG-I/HMG-Y | 125.73 | 23.36 | 11614 | 2 | 7508813.5 | 5652874 | 4122210.25 | 5767655 | 4118377.5 | 9820769 | 6472627.5 | 5294170 | 4766062 | 7232697 | 7648793.5 | 5821932 | 6028630.5 | 7142351 | 7132211 | 7178966 | 12849916 | 8027079.5 | 7948020 | 8073681 | 4 | 1 | 11 | 5 | 11 | 5 | 1.243062374 | 1.005585383 | 1.622295755 | 1.3123693 |
| P63158 | Hmgb1 | High mobility group protein B1 | 497.76 | 38.6 | 24894 | 8 | 23205570 | 28924132 | 36192366 | 42151882 | 30326520 | 26836270 | 24562834 | 37268022 | 28494522 | 32143542 | 18169290 | 18034210 | 19538952 | 21277812 | 21095964 | 32348298 | 26859146 | 36595802 | 33785598 | 41563262 | 0 | 10 | 0 | 6 | 0 | 10 | 0.610173764 | 0.657152159 | 1.064375658 | 1.146323889 |
| P30681 | Hmgb2 | High mobility group protein B2 | 603.57 | 33.33 | 24162 | 8 | 21961562 | 23457366 | 41478190 | 43007074 | 29077222 | 28563130 | 23595634 | 28329708 | 24294538 | 28383470 | 13630131 | 14304459 | 14980980 | 19119014 | 16910132 | 25979574 | 26381422 | 25905746 | 25673096 | 30006830 | 0 | 11 | 0 | 8 | 0 | 11 | 0.496565693 | 0.592827234 | 0.84253036 | 1.005858742 |
| P38060 | Hmgcl | Hydroxymethylglutaryl-CoA lyase, mitochondrial | 657.01 | 40.62 | 34239 | 10 | 9731430 | 34839118 | 13907799 | 13000718 | 11349875 | 12251256 | 11695107 | 8167647 | 44270046 | 13433311 | 11743790 | 13436908 | 14580310 | 14709682 | 16038312 | 10492196 | 9221245 | 11394106 | 10638087 | 12894660 | 8 | 6 | 7 | 7 | 8 | 8 | 0.851260465 | 0.785026375 | 0.659676364 | 0.608348873 |
| Q8JZK9 | Hmgcs1 | Hydroxymethylglutaryl-CoA synthase, cytoplasmic | 784.4 | 32.12 | 57569 | 12 | 8394162 | 8463691 | 5522955 | 3325473.25 | 3218565 | 6594870 | 10298355 | 4819198.5 | 4418605 | 2988930.5 | 6564970.5 | 11045832 | 5842491 | 1800239.25 | 6650051 | 8637104 | 2736239.5 | 3010839.75 | 2327384.25 | 2838557.5 | 11 | 5 | 10 | 5 | 11 | 5 | 1.102981965 | 1.095591644 | 0.675893826 | 0.671365128 |
| P09602 | Hmgn2 | Non-histone chromosomal protein HMG-17 | 104.93 | 23.33 | 9423 | 2 | 2518673.75 | 2129229.5 | 2180268 | 1833932.25 | 1424810.75 | 2392793 | 1770286.5 | 1843877.38 | 1476029.12 | 2019599 | 2688387.5 | 1555313 | 709729.38 | 166354.34 | 1584221.12 | 1885337.88 | 3473647.25 | 1902768.5 | 2578519 | 1801057.75 | 7 | 10 | 7 | 7 | 7 | 10 | 0.664624004 | 0.705492804 | 1.154102245 | 1.22506985 |
| Q9JL35 | Hmgn5 | High mobility group nucleosome-binding domain-containing protein 5 | 255.24 | 13.55 | 45344 | 5 | 5569288 | 7274098 | 6529054.5 | 6875091.5 | 7456881 | 7318162 | 7319152 | 8842843 | 8387609 | 7571119 | 5536198.5 | 6127594 | 5318245 | 5621212.5 | 4993160 | 4739956 | 3600630 | 6937649 | 8604616 | 7278166.5 | 2 | 12 | 0 | 4 | 2 | 12 | 0.818777351 | 0.699725918 | 0.924538205 | 0.790108988 |
| P14901 | Hmox1 | Heme oxygenase 1 | 559.13 | 51.21 | 32929 | 10 | 523223.28 | 333599.88 | 281882.22 | 486132.62 | 260096.28 | 396796.94 | 492680.75 | 417891.44 | 255481.94 | 966386.88 | 4228262 | 4485262.5 | 9708963 | 9171823 | 7049210 | 1636047.5 | 2272107 | 1785306.5 | 2238398.5 | 1883088.38 | 0 | 1 | 0 | 0 | 0 | 1 | 18.37916625 | 13.69721678 | 5.207050444 | 3.880594896 |
| O70252 | Hmox2 | Heme oxygenase 2 | 720.46 | 46.03 | 35739 | 12 | 10364086 | 20039620 | 10775005 | 8409046 | 10243355 | 10662440 | 10578475 | 8669683 | 9413459 | 9274947 | 9946129 | 8091479 | 9177474 | 9034180 | 9913444 | 11294910 | 10409231 | 10552550 | 11184656 | 11264631 | 4 | 7 | 8 | 4 | 8 | 7 | 0.771550193 | 0.949869384 | 0.914339984 | 1.125660476 |
| Q9CX86 | Hnrnpa0 | Heterogeneous nuclear ribonucleoprotein A0 | 528.66 | 26.23 | 30530 | 8 | 5272880.5 | 7238682 | 9156738 | 7120239 | 5903201 | 6627331 | 5318454 | 7068708.5 | 6364843 | 7199068.5 | 10312660 | 9423261 | 8580416 | 10091398 | 8916231 | 2886863 | 5639788 | 6742153 | 3516319.5 | 6413643 | 2 | 5 | 0 | 6 | 2 | 6 | 1.364127753 | 1.452617647 | 0.726362129 | 0.773480669 |
| P04256 | Hnrnpa1 | Heterogeneous nuclear ribonucleoprotein A1 | 1398.39 | 53.12 | 34212 | 20 | 33517458 | 49451342 | 37017462 | 37578526 | 31037856 | 34853178 | 37642942 | 37499326 | 38780042 | 34458954 | 54197278 | 59644046 | 43467818 | 44470490 | 43285542 | 34269854 | 36620906 | 36975678 | 32237254 | 39187346 | 3 | 11 | 0 | 8 | 3 | 11 | 1.29937295 | 1.337440556 | 0.950628444 | 0.978478915 |
| O88569 | Hnrnpa2b1 | Heterogeneous nuclear ribonucleoproteins A2/B1 | 1527.69 | 60.62 | 37403 | 22 | 56672546 | 74076274 | 58325170 | 56168138 | 69865986 | 53445986 | 64849546 | 62147842 | 61585314 | 55124782 | 87595378 | 98292418 | 60995546 | 61767338 | 65929854 | 59097090 | 54790214 | 64614674 | 62040938 | 62731662 | 6 | 12 | 5 | 11 | 6 | 12 | 1.188736555 | 1.260562544 | 0.962446108 | 1.020599147 |
| Q8URK4 | Hnrnpa3 | Heterogeneous nuclear ribonucleoprotein A3 | 1147.11 | 35.09 | 39652 | 16 | 18569928 | 37536770 | 39661902 | 21641438 | 54703614 | 15348994 | 33242286 | 44343198 | 54934378 | 22108526 | 12461609 | 43073770 | 23615706 | 17418580 | 30162124 | 20058974 | 12516090 | 20821074 | 16282827 | 24884230 | 8 | 4 | 8 | 5 | 8 | 5 | 0.736326186 | 0.745580309 | 0.549422977 | 0.556328106 |
| Q99020 | Hnrnpab | Heterogeneous nuclear ribonucleoprotein A/B | 510.07 | 26.32 | 30831 | 9 | 16071982 | 17270760 | 17912872 | 15058463 | 19816642 | 10655267 | 15431721 | 13103334 | 17450032 | 14468080 | 13599191 | 14894973 | 8801528 | 6211235.5 | 9949263 | 23144590 | 11936647 | 12014533 | 12359325 | 16778120 | 0 | 7 | 5 | 12 | 5 | 12 | 0.620640245 | 0.75175598 | 0.88508741 | 1.072069946 |
| Q9Z204 | Hnrnpc | Heterogeneous nuclear ribonucleoproteins C1/C2 | 1093.61 | 48.56 | 34385 | 17 | 2094205.25 | 2434430 | 2110374 | 838120.12 | 1424170.12 | 306363.81 | 2712307 | 2222432 | 1936538.88 | 943623.56 | 448227.62 | 1628213.62 | 489045.31 | 460520.78 | 793514.56 | 2426816.75 | 490531.88 | 467506.5 | 572944.44 | 753191.06 | 2 | 4 | 6 | 8 | 6 | 8 | 0.42909711 | 0.470311186 | 0.529247514 | 0.580080872 |
| Q60668 | Hnrnpd | Heterogeneous nuclear ribonucleoprotein D0 | 386.65 | 20.56 | 38354 | 7 | 563462.12 | 1063531.38 | 2611195 | 1216766.75 | 4072988 | 168132.3 | 2166759 | 2541813 | 2611810.5 | 546526.94 | 1370817.12 | 1619652.75 | 236198.52 | 102152.12 | 1281884.25 | 2510624.5 | 2 | 693786.12 | 122084.8 | 1327126.88 | 9 | 7 | 7 | 7 | 9 | 7 | 0.483913961 | 0.573824618 | 0.488418558 | 0.579166164 |
| Q794E4 | Hnrnpf | Heterogeneous nuclear ribonucleoprotein F | 975.83 | 47.71 | 45730 | 14 | 11060157 | 15802776 | 18965024 | 15372899 | 15677346 | 13567862 | 11582927 | 15081910 | 15051018 | 15933645 | 39117578 | 30143516 | 30999506 | 32429128 | 32591700 | 13179307 | 13028478 | 20546906 | 16936200 | 21042528 | 0 | 9 | 0 | 8 | 0 | 9 | 2.149912767 | 2.320802447 | 1.102177429 | 1.189785982 |
| O35737 | Hnrnph1 | Heterogeneous nuclear ribonucleoprotein H | 760.14 | 29.18 | 49199 | 12 | 3248334 | 4536220.5 | 4506784 | 4811557.5 | 2916209.25 | 3550374.5 | 3666929.5 | 2868950.5 | 2707650 | 2779467.75 | 4764916 | 3405631 | 1858702.75 | 3291126.75 | 3875979.25 | 4277149.5 | 4631974 | 4405696 | 2997181 | 9207018 | 10 | 11 | 9 | 2 | 10 | 11 | 0.85899722 | 1.10421529 | 1.27473322 | 1.638631511 |
| P70333 | Hnrnph2 | Heterogeneous nuclear ribonucleoprotein H2 | 642.25 | 28.29 | 49280 | 9 | 1677949 | 1299787.25 | 1654852.5 | 1523884.5 | 1672233 | 1328095.5 | 2054938.75 | 941564.75 | 1567379.5 | 1236722.38 | 1738686.88 | 2504463.75 | 2075018.88 | 982578.62 | 1994477.62 | 2025447.25 | 1721949 | 1790852.25 | 2204002 | 1256044.5 | 5 | 5 | 6 | 6 | 6 | 6 | 1.187325907 | 1.303915805 | 1.14939745 | 1.26226295 |
| P61979 | Hnrnpk | Heterogeneous nuclear ribonucleoprotein K | 1543.29 | 47.3 | 50976 | 23 | 122938394 | 122905290 | 139021010 | 119676002 | 130210754 | 108051386 | 131205338 | 136107954 | 144796210 | 125407282 | 126999138 | 129130138 | 113495162 | 96755346 | 124216426 | 141181810 | 135657122 | 117558962 | 125955890 | 125344002 | 9 | 10 | 6 | 11 | 9 | 11 | 0.930436961 | 0.914847165 | 1.017245074 | 1.000200778 |
| F1LQ48 | Hnrnpl | Heterogeneous nuclear ribonucleoprotein L | 1243.93 | 49.92 | 67903 | 19 | 30522246 | 37728666 | 38688674 | 37828566 | 33826794 | 31980084 | 33419606 | 37835326 | 39261094 | 33497320 | 40502062 | 42592106 | 34548546 | 31333480 | 38539838 | 35177722 | 33649730 | 35514226 | 31873650 | 38190494 | 8 | 10 | 8 | 12 | 8 | 12 | 1.049951503 | 1.065471774 | 0.976543995 | 0.990979163 |
| Q921F4 | Hnrnpll | Heterogeneous nuclear ribonucleoprotein L-like | 360.7 | 19.12 | 64125 | 8 | 3936193 | 3755297 | 4827116 | 4196004 | 4640017 | 4873489 | 4133701.75 | 3693744.5 | 4168832.25 | 3809439.25 | 1870949.88 | 3955621.75 | 4099997.5 | 3393113.25 | 2606064 | 5749154 | 2712510.5 | 4590488.5 | 2760747.25 | 3629418.75 | 4 | 8 | 4 | 9 | 4 | 9 | 0.745774973 | 0.770133331 | 0.910449946 | 0.940186886 |
| Q9D0E1 | Hnrnpm | Heterogeneous nuclear ribonucleoprotein M | 2480 | 49.66 | 77649 | 36 | 167336434 | 193341746 | 178978978 | 195731986 | 160736706 | 191903234 | 179516322 | 164910978 | 151263746 | 161281266 | 130172210 | 132983026 | 114726362 | 129031898 | 126715898 | 178981762 | 193186466 | 170282930 | 188495170 | 182766178 | 0 | 11 | 0 | 6 | 0 | 11 | 0.707076349 | 0.746433793 | 1.019625208 | 1.076379818 |
| Q6IMY8 | Hnrnpu | Heterogeneous nuclear ribonucleoprotein U | 2350.89 | 45.36 | 87732 | 35 | 101909426 | 110872058 | 132315826 | 134402754 | 123022706 | 89282778 | 114492530 | 125439818 | 116803170 | 120934722 | 187348850 | 169775058 | 164332562 | 161619522 | 171480514 | 106669650 | 107430194 | 131694650 | 130245258 | 136360066 | 0 | 12 | 0 | 8 | 0 | 12 | 1.418297446 | 1.507279226 | 1.016392821 | 1.080159729 |
| Q8VDM6 | Hnrnpul1 | Heterogeneous nuclear ribonucleoprotein U-like protein 1 | 259.65 | 7.33 | 96002 | 5 | 2442172.5 | 3447036 | 6323572 | 5300485.5 | 3286817.75 | 3567880 | 2959786.75 | 4784262 | 4513808.5 | 5079496.5 | 3858290 | 4840137.5 | 4731360 | 4112679.5 | 4173477.25 | 3358682.5 | 4357831 | 5128614.5 | 5279125 | 5364800.5 | 10 | 10 | 12 | 7 | 12 | 10 | 1.044031578 | 1.038780265 | 1.129276871 | 1.123596788 |
| Q00PI9 | Hnrnpul2 | Heterogeneous nuclear ribonucleoprotein U-like protein 2 | 826.53 | 21.21 | 84940 | 16 | 21956422 | 26279338 | 24787354 | 24778046 | 24879008 | 23416574 | 24467100 | 23254850 | 22053578 | 17778620 | 21607766 | 18754402 | 19596036 | 21650036 | 15870035 | 24787406 | 19865692 | 19737938 | 21662556 | 26410438 | 0 | 8 | 4 | 12 | 4 | 12 | 0.794572396 | 0.878414353 | 0.916725432 | 1.013456775 |
| Q3TEA8 | Hp1bp3 | Heterochromatin protein 1-binding protein 3 | 753.51 | 27.44 | 60867 | 14 | 10764145 | 10801052 | 9942685 | 11520960 | 9277487 | 13437523 | 9359696 | 8447070 | 9447561 | 9877402 | 5420073.5 | 4704245.5 | 7196396 | 7484029 | 6610166 | 9351240 | 10871964 | 11433897 | 10946260 | 10033310 | 0 | 10 | 0 | 8 | 0 | 10 | 0.600594815 | 0.621225523 | 1.006315526 | 1.040882926 |
| Q6P747 | Hp1bp3 | Heterochromatin protein 1-binding protein 3 | 516.42 | 19.53 | 60807 | 9 | 2 | 2 | 2 | 2 | 2 | 2 | 2 | 2 | 2 | 2 | 2 | 2 | 2 | 2 | 2 | 2 | 2 | 254483.53 | 2 | 2 | 12.5 | 10 | 12.5 | 10 | 12.5 | 10 | 1 | 1 | 25449.153 | 25449.153 |
| P62748 | Hpcal1 | Hippocalcin-like protein 1 | 82.86 | 9.84 | 22338 | 2 | 2153310 | 2180834.25 | 1861607.5 | 1367227.5 | 2451932 | 1308597.75 | 2028282.38 | 1288971.25 | 1603568.25 | 1699804.62 | 903877.31 | 1861790.75 | 829316.25 | 503672 | 1033438.75 | 2065273.25 | 602596.06 | 880255.06 | 1197047.5 | 1596693 | 2 | 3 | 4 | 7 | 4 | 7 | 0.512445386 | 0.647237977 | 0.633242244 | 0.799808994 |
| Q8CFE2 | Hpf1 | Histone PARylation factor 1 | 127.41 | 7.8 | 39291 | 2 | 1726564.38 | 1639607.75 | 2618235 | 1101046.5 | 2210294.5 | 1541445.38 | 1812904.25 | 1984333.5 | 1991817.75 | 1452340.25 | 350053.06 | 695089.31 | 332974.34 | 1380569.75 | 706184.25 | 1305231.62 | 1432923 | 607230.62 | 684501.12 | 1680828.12 | 1 | 4 | 0 | 2 | 1 | 4 | 0.372737155 | 0.394504541 | 0.614336189 | 0.650212658 |
| P00493 | Hprt1 | Hypoxanthine-guanine phosphoribosyltransferase | 500.06 | 26.15 | 24570 | 7 | 33790090 | 30398842 | 36526894 | 30844400 | 32075374 | 30106702 | 30306606 | 30455542 | 31334932 | 33212414 | 18558818 | 25044932 | 21425262 | 16412856 | 20107148 | 32734766 | 26674158 | 29056866 | 29366844 | 30421430 | 0 | 4 | 0 | 6 | 0 | 6 | 0.620580216 | 0.653400473 | 0.906001286 | 0.953916437 |
| P59438 | Hps5 | Hermansky-Pudlak syndrome 5 protein homolog | 103.66 | 2.22 | 126337 | 2 | 491528.81 | 266816.91 | 454415.25 | 538369.38 | 321859.94 | 176471.25 | 551637.06 | 545591.62 | 557272.81 | 492974.81 | 2 | 154307.19 | 413289.91 | 2 | 160046.09 | 641954.94 | 234109.66 | 415994.5 | 243006.14 | 308233.81 | 2 | 1 | 8 | 9 | 2 | 9 | 0.351013313 | 0.313054384 | 0.889198111 | 0.793039341 |
| Q3TC93 | Hs1bp3 | HCLS1-binding protein 3 | 43.86 | 2.78 | 43691 | 1 | 686142.38 | 465007.38 | 542651.06 | 640296.56 | 539179.88 | 450064.75 | 340827.25 | 530438.69 | 545059.19 | 636363.5 | 380157.66 | 314721.44 | 304503.88 | 419223.75 | 381951.66 | 512483.22 | 625210.94 | 622367.94 | 579337.31 | 550981.12 | 0 | 12 | 3 | 7 | 3 | 12 | 0.626656681 | 0.719431009 | 1.00595253 | 1.154880282 |
| Q8R3H7 | Hs2st1 | Heparan sulfate 2-O-sulfotransferase 1 | 51.41 | 2.53 | 41813 | 1 | 1103578.38 | 1037478.12 | 1056754.38 | 981248.81 | 1264614.38 | 964091.69 | 1039957.12 | 814184.62 | 1310473.25 | 1118813.62 | 876110.12 | 1004325.5 | 1596332.12 | 1075461.62 | 1194450.38 | 1011297.62 | 1284661.38 | 1147285.62 | 1354728.88 | 981987.19 | 12 | 9 | 10 | 8 | [illegible] | [illegible] | 1.055660142 | 1.095121012 | 1.061759138 | 1.101447991 |
| O70351 | Hsd17b10 | 3-hydroxyacyl-CoA dehydrogenase type-2 | 279.27 | 24.14 | 27246 | 4 | 6497609 | 7073322.5 | 6282428 | 8207786.5 | 6482150 | 7458190 | 6365093.5 | 6231168 | 6313991.5 | 7411593 | 5897380 | 5781495 | 7347602 | 7590852.5 | 6338507.5 | 6479698.5 | 7327937.5 | 7212920 | 7317460 | 6813079 | 9 | 9 | 10 | 10 | 10 | 10 | 0.954044368 | 0.975601003 | 1.017595281 | 1.040587849 |
| O08756 | Hsd17b10 | 3-hydroxyacyl-CoA dehydrogenase type-2 | 629.69 | 50.96 | 27419 | 9 | 11768356 | 13214598 | 11173652 | 11351054 | 10677146 | 13572930 | 11158710 | 11951658 | 10489891 | 13065420 | 10903678 | 12284620 | 13791030 | 14618737 | 10540514 | 11259039 | 13924786 | 10758901 | 14469350 | 11284636 | 10 | 10 | 10 | 10 | 10 | 10 | 1.067951984 | 1.031540735 | 1.060357785 | 1.024205456 |
| Q9EQ06 | Hsd17b11 | Estradiol 17-beta-dehydrogenase 11 | 213.43 | 12.08 | 32881 | 3 | 1812642.75 | 2447585.25 | 2639663.75 | 2097696.5 | 3681367 | 1703346 | 2315954.75 | 3168983.25 | 3170639.25 | 2857470.5 | 2941382 | 3323046.5 | 4491005.5 | 2990805 | 3880440.5 | 2251091.5 | 1884802 | 2843581 | 3048951 | 2226320.5 | 3 | 12 | 4 | 8 | 4 | 12 | 1.390231226 | 1.33369812 | 0.966542255 | 0.927238264 |
| O70503 | Hsd17b12 | Very-long-chain 3-oxoacyl-CoA reductase | 1211.37 | 67.95 | 34742 | 19 | 24645048 | 24625552 | 20024756 | 20379068 | 24375654 | 26629998 | 28338590 | 23588066 | 27183254 | 28153398 | 73031874 | 56624130 | 65838058 | 55050814 | 64825014 | 30608466 | 37458042 | 27887806 | 32889588 | 26303100 | 0 | 0 | 0 | 6 | 0 | 6 | 2.76518785 | 2.355382053 | 1.360341042 | 1.158736061 |
| P51660 | Hsd17b4 | Peroxisomal multifunctional enzyme type 2 | 881.73 | 20.68 | 79482 | 14 | 16972594 | 12812410 | 16793030 | 13023959 | 10466274 | 17170266 | 12934219 | 10029324 | 8144513.5 | 14538211 | 17642472 | 14308450 | 13541020 | 16856404 | 18547100 | 13494369 | 19726740 | 11502725 | 14879096 | 13025683 | 5 | 10 | 5 | 8 | 5 | 10 | 1.154523288 | 1.287805001 | 1.036540735 | 1.15620218 |
| O88736 | Hsd17b7 | 3-keto-steroid reductase | 415.23 | 25.75 | 37317 | 6 | 3117709 | 2385004 | 1333830.75 | 2008091.25 | 1919802.88 | 3794356.75 | 3870693.25 | 2546889.5 | 2239990 | 1838859 | 3015316.75 | 4053680 | 5286140.5 | 5069249 | 4978338.5 | 2876278.25 | 2185816 | 2787950.25 | 2273333.75 | 2676787.5 | 1 | 7 | 2 | 12 | 2 | 12 | 2.08117925 | 1.567633917 | 1.18911604 | 0.895693457 |
| P50171 | Hsd17b8 | (3R)-3-hydroxyacyl-CoA dehydrogenase | 292.19 | 22.78 | 26588 | 5 | 10116503 | 9828969 | 7802038.5 | 12108356 | 12419764 | 11199268 | 10035508 | 10091034 | 10866341 | 10083024 | 5070232 | 4981269.5 | 8139845 | 8076867 | 7564361.5 | 10077686 | 10199456 | 9792618 | 9912318 | 9850846 | 2 | 10 | 0 | 4 | 2 | 10 | 0.647195924 | 0.647201564 | 0.953272558 | 0.953280864 |
| Q4V8B7 | Hsdl1 | Inactive hydroxysteroid dehydrogenase-like protein 1 | 48.71 | 3.33 | 36908 | 1 | 574492.56 | 177739.92 | 474369.03 | 695152.5 | 929593.44 | 304940.22 | 643029.5 | 681247 | 743962.25 | 459293.59 | 64735.39 | 86131.27 | 467198.81 | 343434.94 | 95138.56 | 128474.2 | 428338.56 | 589711.25 | 794991.75 | 414141.81 | 2 | 9 | 3 | 9 | 3 | 9 | 0.370575312 | 0.373044733 | 0.826155918 | 0.831661215 |
| P07901 | Hsp90aa1 | Heat shock protein HSP 90-alpha | 2844.78 | 54.71 | 84788 | 45 | 89224850 | 120213658 | 146816274 | 131314578 | 133045242 | 91868690 | 96453882 | 138049378 | 138592354 | 126546010 | 128666434 | 139199874 | 119517098 | 102369626 | 120045458 | 102612218 | 79447074 | 128179546 | 124910122 | 123311202 | 9 | 7 | 11 | 9 | 11 | 9 | 0.982571934 | 1.030917764 | 0.899850181 | 0.944125823 |
| P11499 | Hsp90ab1 | Heat shock protein HSP 90-beta | 3974.02 | 73.62 | 83281 | 61 | 277141026 | 339814402 | 398408578 | 332775266 | 335200162 | 250017890 | 257515954 | 342290402 | 336572098 | 331610658 | 445201762 | 424404354 | 340753730 | 301181506 | 330782466 | 275469282 | 254223186 | 326748002 | 300549698 | 309572642 | 9 | 3 | 7 | 9 | 9 | 9 | 1.094445826 | 1.213646456 | 0.871222274 | 0.966110702 |
| P08113 | Hsp90b1 | Endoplasmin | 3416.67 | 59.98 | 92476 | 55 | 197288546 | 210405858 | 231678802 | 244539202 | 222799890 | 195361922 | 178001922 | 246966114 | 241121570 | 246667842 | 317766562 | 274095010 | 400029954 | 341421730 | 412321122 | 169938674 | 216094610 | 236078146 | 241949154 | 229261138 | 0 | 12 | 0 | 9 | 0 | 12 | 1.577315424 | 1.575312575 | 0.987900581 | 0.986646161 |
| Q99M31 | Hspa14 | Heat shock 70 kDa protein 14 | 276.43 | 11.2 | 54650 | 5 | 3400396.5 | 3513560 | 3857700.25 | 3167164.25 | 2864660.5 | 2774791.25 | 2855637.25 | 3950441 | 2885469.75 | 2740383.25 | 3292483 | 4874741 | 2083303.75 | 3477415.25 | 3288667.75 | 3266137.5 | 3133584.25 | 3120287 | 3303737.5 | 2734782.5 | 12 | 6 | 8 | 9 | 12 | 9 | 1.012683636 | 1.119018957 | 0.925911023 | 1.023134916 |
| Q61316 | Hspa4 | Heat shock 70 kDa protein 4 | 2016.81 | 42.33 | 94133 | 33 | 17570934 | 15673127 | 16665926 | 17431208 | 15477390 | 16731241 | 16995972 | 16069450 | 14895732 | 15951484 | 7164988 | 16827150 | 14625352 | 15983943 | 14966002 | 22349546 | 27430482 | 20621572 | 23909980 | 17387614 | 5 | 2 | 7 | 0 | 7 | 2 | 0.83997787 | 0.86264991 | 1.348721353 | 1.385092029 |
| P48722 | Hspa4l | Heat shock 70 kDa protein 4L | 136.15 | 4.77 | 94382 | 3 | 2 | 2 | 350136.75 | 2 | 535679.06 | 2 | 295923 | 504377.28 | 2 | 2 | 2 | 589060.69 | 2 | 551476.31 | 2 | 322845.5 | 2 | 2 | 2 | 2 | 10.5 | 9 | 10.5 | 10 | 10.5 | 10 | 1.287553532 | 1.425133138 | 0.36446777 | 0.403412429 |
| P20029 | Hspa5 | Endoplasmic reticulum chaperone BiP | 3356.69 | 62.84 | 72347 | 47 | 422764930 | 379570690 | 324665346 | 340867426 | 316957378 | 392718754 | 402470914 | 347378370 | 362981058 | 393046594 | 455719554 | 494055074 | 421180962 | 415443618 | 506556994 | 434602914 | 392781634 | 338534594 | 341922946 | 355795202 | 2 | 8 | 0 | 9 | 2 | 9 | 1.284694697 | 1.207711686 | 1.044156422 | 0.981587233 |
| P63017 | Hspa8 | Heat shock cognate 71 kDa protein | 3844.221 | 75.08 | 70871 | 54 | 1227405442 | 1234723586 | 1213958018 | 1219160962 | 1150131586 | 1202867202 | 1159717634 | 1061806274 | 1097164290 | 1142324866 | 935175170 | 1060865986 | 878373186 | 769258562 | 839118978 | 1235759234 | 1270444674 | 1119273986 | 1061670786 | 1116368002 | 0 | 10 | 0 | 11 | 0 | 11 | 0.74152364 | 0.7914701 | 0.959992105 | 1.024653843 |
| P38647 | Hspa9 | Stress-70 protein, mitochondrial | 2894.79 | 69.51 | 73461 | 41 | 121863890 | 138234162 | 96252466 | 109482770 | 92253346 | 130259778 | 126977170 | 106917298 | 88876530 | 110469658 | 231566034 | 220905202 | 220763714 | 230550002 | 226099922 | 141427714 | 170057570 | 140447042 | 137563682 | 144059698 | 0 | 1 | 0 | 0 | 0 | 1 | 2.024568956 | 2.005118019 | 1.314411888 | 1.301783746 |
| Q99P31 | Hspbp1 | Hsp70-binding protein 1 | 402.45 | 24.93 | 39167 | 6 | 7060711 | 6794048.5 | 6324013 | 6333823 | 4630153 | 8064256.5 | 7606995 | 5906632 | 5759795 | 5191443 | 6056506.5 | 7314337 | 7059743.5 | 4556591.5 | 6049336 | 5798327.5 | 5048168.5 | 5785981 | 5260233.5 | 4831200 | 11 | 5 | 12 | 5 | 12 | 5 | 0.996588805 | 0.954114746 | 0.858110211 | 0.821538033 |
| P63038 | Hspd1 | 60 kDa heat shock protein, mitochondrial | 4175.81 | 76.44 | 60955 | 54 | 405753922 | 420495266 | 382470690 | 430740898 | 326575970 | 425424674 | 362507586 | 404076610 | 339642530 | 453447074 | 916978498 | 866707522 | 918442114 | 920899522 | 978072386 | 431340546 | 524064386 | 484915778 | 470334530 | 492476642 | 0 | 0 | 0 | 1 | 0 | 1 | 2.340291987 | 2.317819545 | 1.222322974 | 1.210585728 |
| Q64433 | Hspe1 | 10 kDa heat shock protein, mitochondrial | 474.69 | 67.65 | 10963 | 7 | 46814846 | 31449030 | 30110614 | 41931030 | 25769508 | 46977722 | 42066230 | 36473850 | 31469470 | 44223418 | 50963494 | 44858726 | 48755798 | 56888150 | 59396762 | 47254426 | 74640634 | 50619538 | 51834294 | 47250902 | 1 | 0 | 0 | 1 | 0 | 1 | 1.481544163 | 1.296466555 | 1.542523077 | 1.349827855 |
| Q61699 | Hsph1 | Heat shock protein 105 kDa | 770.61 | 20.05 | 96407 | 13 | 11991290 | 11422438 | 8341631.5 | 9525722 | 9428292 | 11394152 | 13075355 | 13311584 | 12758421 | 9872232 | 14364920 | 15689804 | 15478738 | 13196316 | 15050697 | 11865986 | 8940429 | 12852045 | 11250729 | 10716385 | 0 | 9 | 1 | 7 | 1 | 9 | 1.454967197 | 1.221293578 | 1.096948555 | 0.920774179 |
| Q9Z2G9 | Htatip2 | Oxidoreductase HTATIP2 | 346.75 | 31.4 | 26870 | 7 | 4148426 | 3817786 | 4853100.5 | 5939706 | 4709699 | 4484955 | 4283836.5 | 7609679 | 6649794.5 | 9112034 | 2192111.5 | 6835140 | 9538908 | 8115372.5 | 9728563 | 3449188.5 | 3095581.5 | 8583502 | 7317315.5 | 7954892 | 5 | 10 | 8 | 11 | 8 | 11 | 1.551430963 | 1.132848671 | 1.295361773 | 0.945867974 |
| Q9JIY5 | Htra2 | Serine protease HTRA2, mitochondrial | 106.52 | 8.52 | 49348 | 2 | 2 | 1194580.5 | 1536067.62 | 1948775 | 1571739 | 1266280.25 | 1196915.62 | 1432833.75 | 1289950.5 | 1453628.25 | 1727895.12 | 843370.19 | 1531591.88 | 1633930 | 1464699 | 1120214.12 | 1280456.38 | 1774128.75 | 1784050.12 | 1216401.25 | 12 | 12 | 5 | 12 | 12 | 12 | 1.152023215 | 1.084625145 | 1.147826306 | 1.080673772 |
| P42859 | Htt | Huntingtin | 196.88 | 1.19 | 344690 | 3 | 288175.75 | 301949.81 | 291571.62 | 345650.31 | 616362 | 219792.77 | 779292.94 | 594103.75 | 981694.5 | 781164.62 | 312368.66 | 693094.12 | 610145.88 | 384978.28 | 679354.56 | 169189.34 | 2 | 723682.38 | 321148.09 | 416143.28 | 4 | 2 | 4 | 4 | 8 | 12 | 1.45355953 | 0.798540735 | 0.884176764 | 0.48573942 |
| Q7TMY8 | Huwe1 | E3 ubiquitin-protein ligase HUWE1 | 641.27 | 3.04 | 482635 | 12 | 5756171.5 | 4604966 | 5531840 | 5123331 | 5371638 | 5723900 | 5033468 | 4752730.5 | 4381026 | 4870014 | 5189085.5 | 5416213.5 | 4746944 | 5188798.5 | 3556262.25 | 4889614 | 4858302 | 4621653 | 5286672 | 2905321.5 | 8 | 5 | 12 | 10 | 12 | 10 | 0.913193596 | 0.973190459 | 0.854995007 | 0.91116822 |
| Q8R1F5 | Hyi | Putative hydroxypyruvate isomerase | 244.93 | 15.16 | 30449 | 4 | 4508822.5 | 3902198.5 | 5443941 | 6720676 | 5942311.5 | 1907126.5 | 4512956 | 5629331 | 4487224 | 6187359 | 1718843.62 | 2387598.5 | 3082677.25 | 2162378 | 1595607.62 | 2330688 | 7135121 | 5992240.5 | 5055508.5 | 7354827 | 0 | 9 | 3 | 7 | 3 | 9 | 0.412816683 | 0.481742065 | 1.050925336 | 1.226385728 |
| Q9JKR6 | Hyou1 | Hypoxia up-regulated protein 1 | 3040.19 | 52.85 | 111181 | 48 | 47324006 | 48168646 | 35421122 | 33866322 | 40613994 | 41902814 | 42494498 | 51366006 | 50924766 | 59127586 | 115752090 | 114542186 | 119983154 | 105200922 | 134944610 | 32314322 | 41307738 | 53938778 | 52005582 | 53097274 | 0 | 7 | 0 | 12 | 0 | 12 | 2.874585934 | 2.401893101 | 1.132767228 | 0.946496592 |
| P03975 | Iap | IgE-binding protein | 707.99 | 25.31 | 62747 | 11 | 10904472 | 13062561 | 13126207 | 12636507 | 10105517 | 16410198 | 17925098 | 12452111 | 10685703 | 13514713 | 13062716 | 13777400 | 9151308 | 9583375 | 9284160 | 16152140 | 13243808 | 14397614 | 15403137 | 13142806 | 9 | 0 | 5 | 12 | 9 | 12 | 0.916833241 | 0.772793934 | 1.208977786 | 1.01904104 |
| Q8BU30 | Iars | Isoleucine--tRNA ligase, cytoplasmic | 928.84 | 16.56 | 144271 | 18 | 16918918 | 15203795 | 16640322 | 16250181 | 18413708 | 14458544 | 13998497 | 17748718 | 18171026 | 18509632 | 21845364 | 23638150 | 25183278 | 18052888 | 26025708 | 15359781 | 11256623 | 17130926 | 19382788 | 18983754 | 1 | 10 | 2 | 11 | 2 | 11 | 1.375399961 | 1.384369022 | 0.984261052 | 0.990679474 |
| Q8BIJ6 | Iars2 | Isoleucine--tRNA ligase, mitochondrial | 742.37 | 15.91 | 112804 | 13 | 14376774 | 16359546 | 14293188 | 17262170 | 15771938 | 12347232 | 11866860 | 17529840 | 14928774 | 13097612 | 7529638 | 11814743 | 18794886 | 17215332 | 17474372 | 14062054 | 12074779 | 15868722 | 16464583 | 17065024 | 11 | 11 | 12 | 9 | 12 | 11 | 0.932943857 | 1.043838886 | 0.96760134 | 1.082626024 |
| P13597 | Icam1 | Intercellular adhesion molecule 1 | 433.7 | 15.83 | 58844 | 8 | 2101873 | 2443097 | 2638901.25 | 770396.62 | 210194.59 | 3593100 | 3619036.25 | 2738806.5 | 3275944 | 2702585.5 | 4390429.5 | 3499687.5 | 3796883.5 | 4778943 | 3754785 | 2525925.5 | 765915.62 | 1361945.12 | 5303840 | 858726.75 | 0 | 11 | 2 | 5 | 2 | 11 | 2.476676033 | 1.269390987 | 1.324809856 | 0.67901515 |
| P35330 | Icam2 | Intercellular adhesion molecule 2 | 46 | 2.53 | 31390 | 1 | 366375.19 | 2 | 2 | 2 | 183630.92 | 2 | 2 | 2 | 2 | 2 | 2 | 2 | 530602.62 | 2 | 2 | 2 | 498958.84 | 2 | 2 | 2 | 11 | 11 | 10 | 10 | 11 | 11 | 0.96472534 | 53061.062 | 0.907192462 | 49896.684 |
| Q9JHR7 | Ide | Insulin-degrading enzyme | 276.29 | 5.89 | 117772 | 5 | 5368529.5 | 4827248 | 6301815 | 5256706.5 | 7933194 | 3483256.75 | 5202752 | 5180275 | 6840883 | 4475015 | 2126869.5 | 4978679 | 5308843 | 4925804 | 5734039.5 | 5775904 | 3996378.75 | 4483852.5 | 5931625.5 | 6611597.5 | 7 | 10 | 12 | 10 | 12 | 10 | 0.772237573 | 0.916292132 | 0.902715438 | 1.064219078 |
| O88844 | Idh1 | Isocitrate dehydrogenase [NADP] cytoplasmic | 1190.05 | 49.28 | 46674 | 18 | 63547102 | 57337034 | 38121766 | 49629010 | 42696378 | 55477554 | 52441790 | 38631350 | 39434662 | 42982874 | 49497522 | 41061598 | 34580462 | 27627108 | 34567162 | 65967090 | 65663618 | 44915342 | 40087698 | 47429170 | 3 | 10 | 5 | 7 | 5 | 10 | 0.745366214 | 0.818165262 | 1.050656757 | 1.153273177 |
| P54071 | Idh2 | Isocitrate dehydrogenase [NADP], mitochondrial | 1051.17 | 40.49 | 50906 | 18 | 38082038 | 40262122 | 43873622 | 40174962 | 47431150 | 31461612 | 32558562 | 34173254 | 35502062 | 31663588 | 17785820 | 23827426 | 25200788 | 22935422 | 24628980 | 42051914 | 34490554 | 30764826 | 31560086 | 32867800 | 0 | 3 | 0 | 12 | 0 | 12 | 0.545116354 | 0.69169735 | 0.818472943 | 1.038559129 |
| Q9D6R2 | Idh3a | Isocitrate dehydrogenase [NAD] subunit alpha, mitochondrial | 722.19 | 41.53 | 39639 | 13 | 24547176 | 28547432 | 29279290 | 28223896 | 31241236 | 25028598 | 24697762 | 29315538 | 30073450 | 28402092 | 45602310 | 39415898 | 43027398 | 39866254 | 48514402 | 25204506 | 29061532 | 34023446 | 33614218 | 32081326 | 0 | 6 | 0 | 5 | 0 | 6 | 1.525858306 | 1.573809562 | 1.085632269 | 1.119749088 |
| Q68FX0 | Idh3b | Isocitrate dehydrogenase [NAD] subunit beta, mitochondrial | 430.71 | 23.38 | 42354 | 8 | 4113576.5 | 3771308 | 5010667 | 5055859 | 10340922 | 3620184.5 | 3847152.5 | 7148232 | 4135711.25 | 5547320 | 8685390 | 7633763 | 8078491.5 | 7104016 | 10954709 | 6197521.5 | 6254940 | 7892608.5 | 6358638.5 | 6319202 | 4 | 5 | 1 | 4 | 4 | 5 | 1.500631646 | 1.747276348 | 1.167203535 | 1.359045795 |
| P41565 | Idh3g | Isocitrate dehydrogenase [NAD] subunit gamma 1, mitochondrial | 308.85 | 16.54 | 42851 | 7 | 6981381 | 7797810 | 7493047 | 6276695.5 | 6115718.5 | 6791439 | 5479540 | 5468802.5 | 5329397 | 6460638 | 8854416 | 9117726 | 9936011 | 10380129 | 9971880 | 5652533 | 7237330 | 7224916.5 | 5883807 | 7413595 | 0 | 9 | 0 | 4 | 0 | 9 | 1.392200966 | 1.634285875 | 0.963868943 | 1.13147271 |
| P58044 | Idi1 | Isopentenyl-diphosphate Delta-isomerase 1 | 484.01 | 33.04 | 26289 | 8 | 10275524 | 9552742 | 11921270 | 8745653 | 10637929 | 9962784 | 11162172 | 9922140 | 10515463 | 7977631 | 8200821 | 9795442 | 10521084 | 7444468 | 9036308 | 11271856 | 7345498 | 10406190 | 8682548 | 10380313 | 6 | 10 | 7 | 12 | 7 | 12 | 0.880019149 | 0.908315511 | 0.940416053 | 0.970654432 |
| Q9R002 | Ifi202 | Interferon-activable protein 202 | 585.14 | 24.72 | 50466 | 10 | 11856098 | 9811349 | 1345686.5 | 8339616 | 6363246 | 12011938 | 11313004 | 4506352.5 | 4542426 | 5600737.5 | 14152666 | 8192698.5 | 8421738 | 4456950 | 10981706 | 7897588 | 9224174 | 3492592.5 | 2918823.75 | 6604646 | 10 | 9 | 11 | 9 | 11 | 9 | 1.225097147 | 1.216758867 | 0.79907275 | 0.793634086 |
| O35368 | Ifi203 | Interferon-activable protein 203 | 235.24 | 15.2 | 46300 | 5 | 946978.12 | 559788.94 | 1277691.88 | 1056229.5 | 667989.56 | 721464.69 | 637529.75 | 949004.19 | 861182.31 | 936256.62 | 482493.75 | 2 | 728980.94 | 611717.44 | 934276.25 | 831542.94 | 1156267.75 | 922997.12 | 861329.06 | 2 | 3 | 12 | 4 | 9 | 4 | 12 | 0.562091935 | 0.63345854 | 1.017211514 | 1.146362864 |
| P0DOV2 | Ifi204 | Interferon-activable protein 204 | 945.78 | 31.99 | 69438 | 16 | 18930172 | 20552184 | 15202269 | 22115312 | 17688026 | 19186030 | 18232448 | 16486861 | 17210452 | 21211674 | 19965578 | 9397359 | 15387926 | 12464453 | 16033430 | 17865870 | 22356708 | 18805356 | 19470842 | 20488848 | 5 | 10 | 4 | 7 | 5 | 10 | 0.775217749 | 0.793581141 | 1.047621526 | 1.072136271 |
| Q9ESY9 | Ifi30 | Gamma-interferon-inducible lysosomal thiol reductase | 322.3 | 28.23 | 27784 | 6 | 4818800 | 4628118 | 3605326.5 | 3213951 | 8805051 | 3610780.5 | 4337029.5 | 4440209.5 | 5575459.5 | 2720234.5 | 739986.69 | 933777.38 | 1160654.5 | 621738.62 | 257705.12 | 2571911.5 | 3272537.75 | 2392246.75 | 1288238.88 | 1255300.62 | 0 | 1 | 0 | 1 | 0 | 1 | 0.148132336 | 0.17955491 | 0.429983996 | 0.521194357 |
| Q9D8C4 | Ifi35 | Interferon-induced 35 kDa protein homolog | 221.87 | 17.48 | 31876 | 3 | 1327738.75 | 988724.12 | 709578.44 | 750648.06 | 1068472 | 1367260.88 | 1154122.75 | 1011433.75 | 855641.19 | 946389.81 | 1188191.75 | 660073.31 | 1079545.25 | 786960.88 | 904333 | 1018920.06 | 733164.12 | 791963.44 | 1214495.5 | 649693.94 | 12 | 10 | 8 | 7 | 12 | 10 | 0.953343725 | 0.865836076 | 0.909822547 | 0.826309718 |
| Q8BV66 | Ifi44 | Interferon-induced protein 44 | 209.75 | 10.43 | 47852 | 4 | 8697622 | 10107518 | 6048320 | 8683038 | 11484607 | 8148303 | 7329725 | 10206774 | 11158256 | 7700019 | 4681844 | 4418052 | 7154064 | 4582376.5 | 6315248 | 6996473.5 | 6921698 | 4333685 | 5613357.5 | 8851222 | 2 | 5 | 0 | 3 | 2 | 5 | 0.603085697 | 0.609557901 | 0.726691093 | 0.734489806 |
| Q9BDB7 | Ifi44l | Interferon-induced protein 44-like | 383.31 | 19.24 | 49680 | 7 | 15056296 | 14016147 | 7780047.5 | 17854734 | 17541170 | 15945078 | 14687775 | 14851980 | 15055606 | 15542335 | 6896270 | 7779831 | 9234666 | 6036042.5 | 7611214 | 14772263 | 14426800 | 10638963 | 12537147 | 14224228 | 1 | 8 | 4 | 7 | 4 | 8 | 0.519845787 | 0.493646873 | 0.921811501 | 0.875354532 |
| Q8R5F7 | Ifih1 | Interferon-induced helicase C domain-containing protein 1 | 142.46 | 2.93 | 115971 | 3 | 1517989.25 | 1312245.25 | 975211.94 | 1486401 | 1258316.38 | 1202869 | 1280269 | 1996255.25 | 1814069 | 1221278 | 750676.75 | 1463375.25 | 1078926.25 | 506115.47 | 1423174.25 | 1104612.5 | 713715.56 | 820324.94 | 711690.62 | 1814224.75 | 7 | 6 | 6 | 4 | 7 | 6 | 0.79727288 | 0.694936591 | 0.788464001 | 0.687258401 |
| Q64282 | Ifit1 | Interferon-induced protein with tetratricopeptide repeats 1 | 185.2 | 6.91 | 53737 | 3 | 539449.25 | 450480.34 | 2 | 2 | 361077.62 | 2 | 333321.66 | 301532.53 | 423545.03 | 2 | 2 | 368293.56 | 1011494.19 | 441820.16 | 653871.62 | 550936.94 | 340743.56 | 962449.44 | 809888.38 | 2 | 8 | 7 | 5 | 5 | 8 | 7 | 1.832317535 | 2.338883219 | 1.971871366 | 2.517018344 |
| Q99J93 | Ifitm2 | Interferon-induced transmembrane protein 2 | 360.5 | 17.36 | 15743 | 4 | 5355933 | 4821066.5 | 1661589.75 | 4493923.5 | 3759330.25 | 7847195 | 6170795 | 5659796 | 4448472.5 | 6747109 | 4776134.5 | 4546637 | 6464732 | 7641643 | 5867012 | 5749551.5 | 6029808.5 | 5704659.5 | 7240453 | 6420979.5 | 4 | 0 | 11 | 12 | 11 | 12 | 1.458112056 | 0.948913607 | 1.55015406 | 1.008812919 |
| Q9CQW9 | Ifitm3 | Interferon-induced transmembrane protein 3 | 583.48 | 48.91 | 14954 | 6 | 35017494 | 31990272 | 9693378 | 28663176 | 23757432 | 46438882 | 42645270 | 37102474 | 31138636 | 42098354 | 40664406 | 34887610 | 54112154 | 58168450 | 56413882 | 35860642 | 42258166 | 31548276 | 36912518 | 40663122 | 1 | 2 | 7 | 8 | 7 | 8 | 1.891598419 | 1.224762177 | 1.450125336 | 0.938919511 |
| P19182 | Ifrd1 | Interferon-related developmental regulator 1 | 138.58 | 5.35 | 49935 | 2 | 279081.94 | 436668.25 | 2 | 2 | 2 | 187954.36 | 151626.27 | 2 | 2 | 2 | 1547453.25 | 1664108.88 | 1046001.31 | 739880.38 | 1190069.88 | 478172.88 | 491798.84 | 219654.34 | 2 | 167304.45 | 0 | 7.5 | 0 | 4.5 | 0 | 7.5 | 8.644722584 | 18.22072235 | 1.896026229 | 3.996307246 |
| O08836 | Igbp1 | Immunoglobulin-binding protein 1 | 77.79 | 3.53 | 39135 | 1 | 2 | 2 | 368519.69 | 79671.27 | 2 | 2 | 83318.71 | 297220.81 | 237472.59 | 2 | 2 | 2 | 85723.42 | 28784.47 | 259288.05 | 95896.47 | 173209.03 | 73905.38 | 107408.74 | 119476.09 | 11 | 6 | 11 | 7 | 11 | 11 | 0.8340082 | 0.604838505 | 1.271529619 | 0.922137305 |
| Q9CPN8 | Igf2bp3 | Insulin-like growth factor 2 mRNA-binding protein 3 | 502.78 | 17.79 | 63575 | 8 | 5277695.5 | 5748032 | 5982107 | 6969217.5 | 6427466.5 | 6667367 | 6481129.5 | 5427870 | 5968730 | 6746466.5 | 9382402 | 8146840.5 | 8962466 | 5893266 | 8180835.5 | 6501302.5 | 5537251 | 7349937 | 7437216 | 6801160 | 3 | 6 | 4 | 6 | 4 | 6 | 1.334203336 | 1.296381712 | 1.105982537 | 1.074630462 |
| Q8R366 | Igsf8 | Immunoglobulin superfamily member 8 | 47.53 | 1.15 | 65011 | 1 | 3490126 | 3602930.25 | 3416122.75 | 4520469 | 3528644 | 3821155.75 | 3363287.25 | 4309301 | 3609741.75 | 3864288.5 | 2881014.25 | 3541666.75 | 4538075 | 4938769 | 4669505.5 | 3733321 | 3420988.5 | 3765251.25 | 4371907.5 | 4486015 | 7 | 8 | 9 | 10 | 9 | 10 | 1.108347174 | 1.084419829 | 1.065695229 | 1.042688867 |
| Q66HG8 | Ik | Protein Red | 95.98 | 5.57 | 65588 | 2 | 2125162.75 | 2231188.5 | 2464802.25 | 1728169.88 | 2650804.5 | 1292077.75 | 2794753.5 | 1277337.38 | 2181089.5 | 1979706 | 401953.78 | 1063687.5 | 896554.88 | 2 | 542224.12 | 1898005.5 | 764302.25 | 844316.06 | 491318.97 | 985567.5 | 0 | 1 | 0 | 2 | 0 | 2 | 0.259320457 | 0.304927372 | 0.444951195 | 0.523205149 |
| O88351 | Ikbkb | Inhibitor of nuclear factor kappa-B kinase subunit beta | 63.35 | 1.32 | 86690 | 1 | 57400.62 | 37499.38 | 141815.45 | 2 | 167647.66 | 82962.15 | 53548.86 | 205655.19 | 146495.52 | 301990.47 | 35513.59 | 40158.68 | 133493.59 | 2 | 225307.34 | 78226.91 | 2 | 80356.1 | 61270.77 | 208715.45 | 11.5 | 10.5 | 6 | 7 | 11.5 | 10.5 | 1.074462631 | 0.549514952 | 1.059862039 | 0.542047737 |
| Q9R0T8 | Ikbke | Inhibitor of nuclear factor kappa-B kinase subunit epsilon | 174.69 | 4.74 | 80953 | 3 | 1122414.25 | 1269769.5 | 1471137.88 | 669111.19 | 1539089.5 | 927571.56 | 1172691.75 | 1152665.5 | 1215218.5 | 776249.75 | 908365.38 | 873131.62 | 991690.25 | 766585.25 | 1346511.25 | 1236711.62 | 622656.19 | 1025924.12 | 898592.12 | 947251.62 | 7 | 5 | 9 | 10 | 9 | 10 | 0.80478725 | 0.931715065 | 0.779233843 | 0.902131478 |
| P25085 | Il1rn | Interleukin-1 receptor antagonist protein | 177.78 | 19.1 | 20274 | 3 | 2 | 265306.59 | 2 | 2 | 176092.11 | 2 | 2 | 198697.7 | 2 | 2 | 190745.28 | 305789.59 | 2 | 459590.75 | 983281.5 | 325131.38 | 2 | 341015.44 | 2 | 495355.44 | 4.5 | 7 | 4 | 6 | 4.5 | 7 | 4.393721046 | 9.760208791 | 2.631386254 | 5.845359544 |
| P16872 | Il7r | Interleukin-7 receptor subunit alpha | 97.07 | 5.88 | 51605 | 2 | 2 | 2 | 2 | 863705.56 | 2 | 2 | 770374.94 | 2 | 2 | 284662.88 | 2626473 | 986062.5 | 2314582.75 | 2159247.75 | 2444250 | 900625.25 | 1168003.25 | 723302.62 | 2 | 2 | 0 | 7 | 0 | 8 | 0 | 8 | 12.19225503 | 9.981211965 | 3.232478045 | 2.646274086 |
| Q9CXY6 | Ilf2 | Interleukin enhancer-binding factor 2 | 615.98 | 29.74 | 43062 | 10 | 9174005 | 7101038 | 8172763 | 7954904 | 9127248 | 6370014.5 | 8067554 | 7681375 | 8274450.5 | 6909723 | 8185029 | 8743952 | 7847800.5 | 6469826.5 | 7812184 | 10236401 | 9400297 | 7218012 | 7903397 | 7840367 | 8 | 12 | 9 | 7 | 9 | 12 | 0.940496785 | 1.047065102 | 1.0257288 | 1.141954813 |
| Q9Z1X4 | Ilf3 | Interleukin enhancer-binding factor 3 | 531.31 | 12.92 | 96021 | 10 | 3588323.5 | 4670560 | 2536462 | 2821331.25 | 2109882 | 3275674.5 | 2373587.5 | 2260454.5 | 1622073.5 | 3227912.25 | 2655365.25 | 2734457 | 1920687.5 | 1718178 | 2040801.12 | 2736384 | 3770679.75 | 1836089.25 | 2440529.75 | 2585478.5 | 4 | 9 | 9 | 10 | 9 | 10 | 0.703872287 | 0.867535045 | 0.850100868 | 1.047764359 |
| O55222 | Ilk | Integrin-linked protein kinase | 193.16 | 12.61 | 51373 | 5 | 3816587.5 | 5388295 | 3391852.75 | 3280344.75 | 4635111.5 | 5361398 | 4118796 | 3962755.25 | 5048388.5 | 3730336 | 4438423 | 3422191.25 | 3642662.5 | 3805947.5 | 4127954.75 | 4434244.5 | 4585510.5 | 4240618 | 3876236.5 | 4599609.5 | 12 | 10 | 7 | 12 | 12 | 12 | 0.947591534 | 0.874694644 | 1.05967317 | 0.978153997 |
| Q8R0F6 | Ilkap | Integrin-linked kinase-associated serine/threonine phosphatase 2C | 39.31 | 2.3 | 42774 | 1 | 789999.69 | 1087405 | 924436.38 | 943242.56 | 716806.19 | 1210215.5 | 1020943.31 | 684341 | 645690 | 758947.31 | 730512.69 | 668117.56 | 757977.69 | 568719.69 | 617644.62 | 925835.44 | 918283 | 993960.56 | 869362.31 | 846802.94 | 2 | 12 | 5 | 10 | 5 | 12 | 0.74922788 | 0.773811608 | 1.020698501 | 1.054189745 |
| Q8BU33 | Ilvbl | Acetolactate synthase-like protein | 124.9 | 7.28 | 68156 | 2 | 267629.69 | 447548.25 | 354553.59 | 381821.25 | 376890.62 | 265550.22 | 494440 | 576395.5 | 405009.72 | 830548.38 | 497658.19 | 822885 | 1288282.25 | 1380780.25 | 1160162 | 194066.44 | 661153.38 | 837520 | 1191780 | 797544.69 | 0 | 5 | 3 | 7 | 3 | 7 | 2.816476403 | 2.002286228 | 2.013770024 | 1.431627115 |
| Q8CAQ8 | Immt | MICOS complex subunit Mic60 | 2412.86 | 52.31 | 83900 | 36 | 62158350 | 65210714 | 59718474 | 66243846 | 56002790 | 69393970 | 72384346 | 76716050 | 63691034 | 76434706 | 74451978 | 75716754 | 77340354 | 70987426 | 85995450 | 65303298 | 66624202 | 79062746 | 77161610 | 82175578 | 0 | 1 | 7 | 8 | 7 | 8 | 1.242966327 | 1.072142793 | 1.197175951 | 1.032645487 |
| Q5PQR5 | Imp4 | U3 small nucleolar ribonucleoprotein protein IMP4 | 68.3 | 7.56 | 33656 | 2 | 900686.06 | 1422592.5 | 1242228.25 | 1562573.25 | 907246.88 | 1497650.88 | 1030756.31 | 772394.69 | 806886.19 | 1274013.62 | 2106856 | 1760823.75 | 1817512.88 | 2215013.5 | 1872117.88 | 1085036.75 | 1774708.75 | 1425595.88 | 1799022.25 | 1590846.5 | 0 | 4 | 0 | 3 | 0 | 4 | 1.61918718 | 1.815842753 | 1.271714061 | 1.426167887 |
| O55023 | Impa1 | Inositol monophosphatase 1 | 413.89 | 29.24 | 30436 | 7 | 9422377 | 10857228 | 14211266 | 16229520 | 11689397 | 10009847 | 9075987 | 12203611 | 12017610 | 14046848 | 7801058 | 8605827 | 8603751 | 10989196 | 9759351 | 9166444 | 11110482 | 13781328 | 15856460 | 13639834 | 3 | 12 | 3 | 9 | 3 | 12 | 0.733205231 | 0.797839042 | 1.018342636 | 1.108111997 |
| Q8CIN7 | Impa2 | Inositol monophosphatase 2 | 44.04 | 3.45 | 31796 | 1 | 2 | 2 | 2 | 2 | 2 | 2 | 2 | 2 | 2 | 2 | 2 | 2 | 2 | 581917.94 | 600837.62 | 2 | 2 | 2 | 2 | 2 | 7.5 | 12.5 | 7.5 | 12.5 | 7.5 | 12.5 | 118276.156 | 118276.156 | 1 | 1 |
| D3ZLZ7 | Impdh1 | Inosine-5'-monophosphate dehydrogenase 1 | 161.51 | 7.2 | 55293 | 3 | 1065312.5 | 2 | 1215957.38 | 1348535.25 | 1452748.12 | 1161148.88 | 991805 | 1501126.62 | 1211456.12 | 1228220.62 | 672440.62 | 2 | 1026668.56 | 1358373.12 | 477331.97 | 942880.25 | 1352220.5 | 1818499.38 | 1986413.38 | 1635430.25 | 7.5 | 5 | 5 | 6 | 7.5 | 6 | 0.695480147 | 0.580071724 | 1.521959601 | 1.269404647 |
| P24547 | Impdh2 | Inosine-5'-monophosphate dehydrogenase 2 | 1096.96 | 34.82 | 55815 | 17 | 46818662 | 44192842 | 57220642 | 55969838 | 47685234 | 44620610 | 41569466 | 42685438 | 43475630 | 43758310 | 36532262 | 37770430 | 33612442 | 33456942 | 35215622 | 46446602 | 55559318 | 51905486 | 46766378 | 45682334 | 0 | 9 | 0 | 0 | 0 | 9 | 0.70105859 | 0.817121578 | 0.978057243 | 1.139978439 |
| Q9WU62 | Incenp | Inner centromere protein | 149.72 | 2.95 | 101209 | 2 | 378871.59 | 532821.38 | 767677.81 | 382922.94 | 464617.97 | 470475.53 | 417217.88 | 458378.56 | 304881.25 | 398269.38 | 295504.75 | 725572.44 | 532537.25 | 619604.12 | 503925.09 | 690731.5 | 526206.06 | 824912.88 | 339174.53 | 288084.31 | 11 | 12 | 5 | 9 | 11 | 12 | 1.059452794 | 1.306419151 | 1.056273272 | 1.30249846 |
| Q0GNC1 | Inf2 | Inverted formin-2 | 49.76 | 0.71 | 138560 | 1 | 512047.59 | 633266.38 | 427922.09 | 419402.81 | 393267.75 | 565732.19 | 487297.09 | 487301.28 | 384322.03 | 320235.88 | 448557.47 | 418825.47 | 452330 | 308804.72 | 2 | 659975.38 | 276735.94 | 360650.88 | 507952.69 | 303537.16 | 7 | 8 | 6 | 10 | 7 | 10 | 0.682558004 | 0.725434551 | 0.883878704 | 0.939401702 |

| | | | | | | | | | | | | | | | | | | | | | | | | | | | | | | | | | | | | |
|---|---|---|---|---|---|---|---|---|---|---|---|---|---|---|---|---|---|---|---|---|---|---|---|---|---|---|---|---|---|---|---|---|---|---|---|---|
| P49442 | Inpp1 | Inositol polyphosphate 1-phosphatase | 59.77 | 3.03 | 43346 | 1 | 1059431.88 | 1319670.38 | 1438122.5 | 588917.5 | 1159355 | 483260.56 | 1192589.38 | 1539606.38 | 1324068.12 | 649219.25 | 258631.16 | 849523.62 | 481497.44 | 2 | 574198.94 | 975895.75 | 2 | 322871.94 | 2 | 592539.62 | 1 | 2 | 3 | 3 | 3 | 3 | 0.388797812 | 0.417028339 | 0.339827911 | 0.364502743 |
| Q9ES52 | Inpp5d | Phosphatidylinositol 3,4,5-trisphosphate 5-phosphatase 1 | 1415.42 | 28.13 | 133542 | 27 | 38693458 | 37958290 | 31860174 | 36696110 | 31288398 | 38899194 | 32195852 | 30773270 | 33202056 | 34134746 | 25947798 | 25220196 | 20213274 | 21812996 | 22761554 | 36890786 | 39475466 | 35998058 | 34038246 | 36317478 | 0 | 11 | 0 | 5 | 0 | 11 | 0.656986762 | 0.685297344 | 1.035261926 | 1.079872974 |
| Q6P549 | Inppl1 | Phosphatidylinositol 3,4,5-trisphosphate 5-phosphatase 2 | 526.24 | 12.41 | 138973 | 12 | 4396439 | 3927493 | 4761229 | 5658810 | 8078044.5 | 2495282 | 4052363 | 4094670 | 5376734.5 | 3798890.75 | 1310382.38 | 1031670.69 | 1286032.75 | 1388325.25 | 1473479.25 | 4019580.5 | 2992418.75 | 2099189 | 1016628.75 | 2970157.5 | 0 | 1 | 0 | 4 | 0 | 4 | 0.241961322 | 0.327475522 | 0.488329242 | 0.660915026 |
| Q3MHC2 | Ints11 | Integrator complex subunit 11 | 64.75 | 1.67 | 67841 | 1 | 957632.62 | 1208014.88 | 963010.25 | 935597.69 | 931777.5 | 1129076.25 | 1073220.75 | 915894.88 | 831667 | 1061304.25 | 834574 | 1142986.62 | 1262930.62 | 971523.5 | 966960.75 | 962010.69 | 1282541.25 | 1084601.75 | 851395.25 | 1095640.62 | 8 | 9 | 10 | 9 | 10 | 9 | 1.036617563 | 1.033487706 | 1.056075815 | 1.052887209 |
| Q8CIM8 | Ints4 | Integrator complex subunit 4 | 56.44 | 1.04 | 108193 | 1 | 391283.94 | 392142.91 | 270949.97 | 326079.28 | 460756.44 | 241821.92 | 332866.94 | 306268.09 | 507914.84 | 283477.97 | 168540.81 | 237824.05 | 314668.84 | 2 | 314656.75 | 364084.94 | 2 | 222599.23 | 2 | 195095.78 | 2 | 2 | 6 | 4 | 6 | 4 | 0.562505646 | 0.619303733 | 0.424602773 | 0.467476343 |
| Q7TQK1 | Ints7 | Integrator complex subunit 7 | 46.02 | 1.35 | 106861 | 1 | 286159.81 | 395398.41 | 289148.59 | 338059.12 | 304475.25 | 2 | 296016.53 | 221928.84 | 300768.66 | 179972.17 | 2 | 303133.47 | 300577.12 | 2 | 253769.95 | 320166.75 | 2 | 90035.94 | 2 | 286112.44 | 4 | 3 | 12 | 10 | 12 | 10 | 0.531529043 | 0.858610866 | 0.431627421 | 0.697233761 |
| Q8VI75 | Ipo4 | Importin-4 | 263.07 | 5.55 | 119275 | 5 | 2296382 | 2007029 | 2253928.75 | 1806802.12 | 2635760 | 1856008 | 2008396.25 | 2055196.75 | 2777005 | 1642562.12 | 1812234 | 2665389.25 | 3445093.25 | 3156119.75 | 1950643.75 | 1903976 | 2827742 | 3216952 | 2196548 | 2788259.75 | 8 | 7 | 8 | 4 | 8 | 7 | 1.184508749 | 1.260205836 | 1.175781194 | 1.250920538 |
| Q8BKC5 | Ipo5 | Importin-5 | 1069.33 | 22.24 | 123591 | 18 | 17828386 | 15182303 | 18312112 | 12260944 | 23921556 | 16877476 | 19174064 | 14796914 | 17932792 | 10972478 | 14175439 | 25313092 | 23087884 | 19971424 | 18165768 | 20689300 | 17637916 | 12719256 | 14283506 | 13563400 | 8 | 9 | 5 | 11 | 8 | 11 | 1.150942924 | 1.262807577 | 0.901583985 | 0.989212466 |
| Q9EPL8 | Ipo7 | Importin-7 | 731.22 | 15.9 | 119486 | 12 | 8521028 | 7533574 | 6504879 | 5132260 | 8371765 | 7159742 | 10393826 | 8150662.5 | 9201285 | 6202659 | 6657465.5 | 8890540 | 8611296 | 5884468 | 7204751 | 8872543 | 6172364.5 | 6745607.5 | 7758500.5 | 6280825.5 | 10 | 12 | 9 | 7 | 10 | 12 | 1.032859104 | 0.906109818 | 0.993520735 | 0.871598932 |
| Q91YE6 | Ipo9 | Importin-9 | 960.54 | 20.46 | 116052 | 17 | 19476406 | 19567476 | 24585030 | 21887518 | 26464994 | 19757014 | 19773444 | 23340508 | 23017398 | 17601586 | 12420822 | 13855881 | 13840517 | 12214764 | 13538037 | 19448054 | 17855322 | 14532344 | 18771570 | 14701470 | 0 | 0 | 0 | 3 | 0 | 3 | 0.588222748 | 0.636487127 | 0.761811709 | 0.82431927 |
| Q9JKF1 | Iqgap1 | Ras GTPase-activating-like protein IQGAP1 | 6644.04 | 60.35 | 188742 | 102 | 374904002 | 375309954 | 392626946 | 363138082 | 411415170 | 331715074 | 376954690 | 376935202 | 397851842 | 388952322 | 282574434 | 360428866 | 378815394 | 322801570 | 364470850 | 324172866 | 301952354 | 320598626 | 346848162 | 342613826 | 4 | 0 | 5 | 2 | 5 | 2 | 0.891361388 | 0.912776533 | 0.853338282 | 0.873839914 |
| Q5DU25 | Iqsec2 | IQ motif and SEC7 domain-containing protein 2 | 51.54 | 0.54 | 161787 | 1 | 1397428.62 | 851577.81 | 775944.5 | 969356.69 | 748551 | 983643.62 | 1302571.62 | 2 | 1036249 | 766583.56 | 2 | 630366.94 | 558182.56 | 2 | 599698.25 | 760612.12 | 2 | 819784.75 | 2 | 704801.31 | 0 | 3 | 4 | 5 | 4 | 5 | 0.377040914 | 0.437326968 | 0.481819587 | 0.558858975 |
| Q8K4B2 | Irak3 | Interleukin-1 receptor-associated kinase 3 | 48.92 | 1.48 | 68455 | 1 | 2 | 327552.78 | 819705.81 | 795685.81 | 628927 | 412525.03 | 395077.25 | 858291.75 | 997734 | 996676.62 | 863202.19 | 775590.06 | 965108.19 | 809449.44 | 1248902.25 | 459358.28 | 527480.38 | 1585308.62 | 1135047.12 | 1343563.12 | 3 | 6 | 10 | 6 | 10 | 6 | 1.812784459 | 1.273733357 | 1.963843757 | 1.379873536 |
| E9Q1P8 | Irf2bp2 | Interferon regulatory factor 2-binding protein 2 | 116.97 | 6.49 | 59292 | 2 | 1029933.75 | 2 | 2208728 | 2531955.75 | 923402.31 | 1127660.12 | 571151.19 | 1343429.25 | 1327524.25 | 2 | 428141.88 | 2 | 929948.12 | 1107923.75 | 761631.62 | 1114819.25 | 1585666 | 1788596.88 | 2488432.5 | 1545960.62 | 7.5 | 9 | 7.5 | 3 | 7.5 | 9 | 0.482168637 | 0.73863149 | 1.273296606 | 1.950556087 |
| Q5EIC4 | Irf2bpl | Interferon regulatory factor 2-binding protein-like | 100.55 | 4.73 | 81496 | 2 | 595500.38 | 905248.44 | 447754.97 | 175633.69 | 613851.19 | 346456.69 | 774858.94 | 48748.89 | 503302.56 | 2 | 2 | 641633.56 | 2 | 2 | 333688.19 | 970928.88 | 2 | 145496.92 | 295505.41 | 335197.19 | 5 | 7 | 7.5 | 11.5 | 7.5 | 11.5 | 0.356220521 | 0.582852738 | 0.638107242 | 1.044079528 |
| P56477 | Irf5 | Interferon regulatory factor 5 | 212.11 | 7.85 | 56005 | 3 | 1721532.62 | 1347115.75 | 2253555 | 1534578.5 | 2426103 | 1203454 | 1765871.88 | 1806874.62 | 2363468 | 1326406.38 | 1213134.12 | 1837745 | 1396020.5 | 855678 | 1542059.75 | 820852.25 | 987180.75 | 980601.62 | 1558109.38 | 1203661.88 | 6 | 2 | 9 | 3 | 9 | 3 | 0.737339466 | 0.808478246 | 0.597918207 | 0.655605574 |
| P23611 | Irf8 | Interferon regulatory factor 8 | 53.91 | 2.83 | 48237 | 1 | 691769.81 | 829680.19 | 2 | 617626.44 | 543777.81 | 832175.75 | 774707.31 | 521435.84 | 573172.38 | 2 | 338170.94 | 608172.69 | 718042.5 | 2 | 329093.75 | 740648.31 | 590965.69 | 619937.88 | 2 | 859544.88 | 8.5 | 10.5 | 8.5 | 10.5 | 8.5 | 10.5 | 0.743044612 | 0.737918504 | 1.047800738 | 1.040572183 |
| Q60766 | Irgm1 | Immunity-related GTPase family M protein 1 | 473.03 | 23.47 | 46552 | 8 | 3127852.25 | 2593361.5 | 2521695.5 | 4086220 | 5824142 | 2613915 | 3481571 | 4260079 | 5493077 | 5146353 | 5105899.5 | 6035521 | 8364039.5 | 3028783.5 | 8008781.5 | 2724807.75 | 1311771 | 3074645 | 3087886.5 | 5014273 | 4 | 10 | 6 | 6 | 6 | 10 | 1.682508049 | 1.454776484 | 0.838051888 | 0.724619522 |
| Q64339 | Isg15 | Ubiquitin-like protein ISG15 | 150.08 | 25.47 | 17898 | 3 | 1380852.88 | 778908.81 | 438809.38 | 736325 | 903511.31 | 2 | 2 | 94205.6 | 549166.06 | 1225377.75 | 188773.66 | 199159.14 | 1180019.38 | 106462.57 | 173543.25 | 602630.75 | 801479.62 | 76010.14 | 90441.87 | 809534.5 | 4 | 7 | 9 | 9 | 9 | 9 | 0.436002921 | 0.988872042 | 0.561554534 | 1.273628113 |
| Q9JL16 | Isg20 | Interferon-stimulated gene 20 kDa protein | 43.75 | 2.67 | 32503 | 1 | 289973.44 | 261718.09 | 297002.19 | 301518.16 | 240103.88 | 290711.06 | 262436.28 | 229108.72 | 386360.19 | 319806.16 | 184280.02 | 111135 | 90112.27 | 2 | 128588.41 | 169233.66 | 2 | 182507.41 | 147685.86 | 167307.08 | 0 | 0 | 0 | 0 | 0 | 0 | 0.369784847 | 0.345411152 | 0.479557255 | 0.447948113 |
| Q3U1G5 | Isg20l2 | Interferon-stimulated 20 kDa exonuclease-like 2 | 83.92 | 3.26 | 41020 | 1 | 58682.61 | 91342.64 | 38952.41 | 2 | 32463.26 | 81331.81 | 77586.49 | 2 | 2 | 2 | 83476.6 | 84572.61 | 2 | 27217.25 | 2 | 78919.83 | 109894.94 | 2 | 44795.25 | 2 | 10 | 12 | 9 | 10 | 10 | 12 | 0.881809452 | 1.228701086 | 1.054962696 | 1.46997042 |
| Q6I7R3 | Isoc1 | Isochorismatase domain-containing protein 1 | 236.84 | 19.87 | 32002 | 4 | 1541349 | 1976793.62 | 825156.31 | 1576510.5 | 1198735.88 | 748701 | 1935646.12 | 1564863.25 | 1686211.38 | 1955528.75 | 2335085.25 | 2379544.5 | 2759821.5 | 2884490 | 1971250 | 591498 | 928845.38 | 866484 | 650003.69 | 1589160.88 | 1 | 6 | 0 | 4 | 1 | 6 | 1.732122325 | 1.562573641 | 0.649850742 | 0.586240143 |
| P85094 | Isoc2a | Isochorismatase domain-containing protein 2A | 197.92 | 33.01 | 22417 | 3 | 1222176 | 1245107.38 | 1093432 | 1459844.25 | 1188624 | 1033551.25 | 1118069.5 | 1327226.25 | 1443185.88 | 1365190 | 591339.19 | 1053749 | 1341855.62 | 1002786.19 | 1262717 | 1176047.5 | 991112.5 | 1591871.75 | 1179929.12 | 1226213.5 | 8 | 10 | 6 | 11 | 8 | 11 | 0.845915874 | 0.835416065 | 0.992912231 | 0.980587851 |
| Q568Z6 | Ist1 | IST1 homolog | 85.84 | 4.37 | 39942 | 2 | 4596576.5 | 4440334 | 1835135.62 | 4992276 | 4407486 | 3979396.75 | 3976587.5 | 4614698 | 3883703.5 | 4710108.5 | 678293.94 | 2925153.5 | 3085536 | 2248736.25 | 3346931.5 | 4300312 | 1272741.12 | 3486259.5 | 4491164 | 4434493 | 4 | 7 | 0 | 9 | 4 | 9 | 0.605996817 | 0.580436794 | 0.887191192 | 0.849770819 |
| Q6AYK3 | Isyna1 | Inositol-3-phosphate synthase 1 | 564.16 | 21.9 | 60884 | 11 | 9958532 | 12066777 | 7869540.5 | 7765192.5 | 7465034 | 8445617 | 9595111 | 6350789.5 | 6520540.5 | 5259525 | 10786804 | 11426651 | 6674582 | 5088024.5 | 5621605.5 | 11435035 | 11162329 | 7324825 | 7071795 | 7011622 | 8 | 8 | 11 | 6 | 11 | 8 | 0.877509148 | 1.094717558 | 0.975191842 | 1.21657949 |
| Q00651 | Itga4 | Integrin alpha-4 | 142.15 | 3.56 | 115696 | 3 | 2 | 2 | 2 | 2 | 2 | 92279.51 | 2 | 147396.58 | 2 | 2 | 129643.13 | 2 | 2 | 207159.25 | 2 | 2 | 102029.98 | 2 | 197409.41 | 2 | 7.5 | 7.5 | 11.5 | 11.5 | 11.5 | 11.5 | 33680.838 | 1.405229652 | 29944.539 | 1.249344037 |
| P11688 | Itga5 | Integrin alpha-5 | 1108.36 | 24.79 | 115043 | 20 | 11842461 | 11361794 | 18609662 | 15657124 | 13577981 | 15707700 | 12934647 | 19186016 | 17769616 | 23509374 | 35551322 | 29716364 | 37055202 | 33330228 | 29294412 | 11561038 | 14106472 | 24209886 | 18605284 | 22018326 | 0 | 7 | 0 | 12 | 0 | 12 | 2.321601668 | 1.851110177 | 1.273782572 | 1.015640157 |
| Q61739 | Itga6 | Integrin alpha-6 | 40.67 | 0.73 | 122159 | 1 | 2 | 2 | 108714.55 | 2 | 68595.85 | 2 | 2 | 2 | 2 | 85600.09 | 2 | 275851.62 | 627054.94 | 649313.75 | 410644.28 | 2 | 2 | 2 | 2 | 144804.95 | 3.5 | 11 | 3 | 12 | 3.5 | 12 | 11.06985361 | 22.9285175 | 0.816692365 | 1.6915802 |
| P24063 | Itgal | Integrin alpha-L | 76.97 | 1.89 | 128328 | 2 | 856702.25 | 977231.31 | 783113.81 | 1136945.62 | 733153.62 | 1021454.69 | 761024.62 | 744346.38 | 2 | 820970.44 | 1278579.12 | 1364140 | 1183441.88 | 1285109.75 | 1264563.75 | 712654.81 | 1073370.62 | 1000419.81 | 795815.19 | 965764 | 0 | 12 | 0 | 8 | 0 | 12 | 1.420910671 | 1.904485949 | 1.013567156 | 1.358512148 |
| P05555 | Itgam | Integrin alpha-M | 1456.28 | 27.93 | 127481 | 22 | 46727342 | 57519530 | 54105570 | 48778522 | 46165426 | 53299618 | 52724342 | 36114754 | 51656490 | 41754538 | 41694810 | 45241870 | 50168674 | 38972046 | 43008194 | 37767218 | 35145010 | 33982422 | 28642822 | 27613392 | 3 | 0 | 8 | 1 | 8 | 1 | 0.864937688 | 0.930103307 | 0.644110498 | 0.692638687 |
| P09055 | Itgb1 | Integrin beta-1 | 1512.72 | 41.48 | 88231 | 26 | 41431430 | 43414342 | 58204826 | 51590462 | 61205722 | 43682486 | 42196174 | 48714094 | 53679982 | 49977194 | 34736694 | 36740058 | 48913274 | 43384470 | 48488938 | 37749274 | 33516008 | 43610462 | 41615478 | 45042798 | 5 | 5 | 6 | 3 | 6 | 5 | 0.829650591 | 0.890927582 | 0.787713718 | 0.845893302 |
| P11835 | Itgb2 | Integrin beta-2 | 2152.3 | 50.58 | 85026 | 32 | 57889934 | 60608126 | 68087306 | 73967066 | 63443658 | 60340942 | 64724178 | 55744202 | 53610658 | 58985786 | 42107182 | 45060026 | 53500018 | 55665018 | 54271026 | 54387646 | 53489918 | 52105238 | 49769862 | 47157346 | 0 | 0 | 2 | 1 | 2 | 1 | 0.773476217 | 0.854118422 | 0.792941699 | 0.875613365 |
| Q61703 | Itih2 | Inter-alpha-trypsin inhibitor heavy chain H2 | 78.81 | 1.69 | 105928 | 1 | 2 | 2 | 2 | 2 | 2 | 2 | 2 | 2 | 2 | 2 | 422157.69 | 262719.94 | 429381.56 | 513188.09 | 2 | 2 | 2 | 2 | 2 | 2 | 2.5 | 12.5 | 2.5 | 12.5 | 2.5 | 12.5 | 162744.928 | 162744.928 | 1 | 1 |
| O89051 | Itm2b | Integral membrane protein 2B | 360.12 | 31.2 | 30260 | 6 | 2361586.5 | 1945130.25 | 2188844.25 | 1419264 | 4057121.75 | 1740583.75 | 2751589.75 | 2622307 | 5049737 | 1918818 | 3220586.25 | 4862778.5 | 5273106 | 4319535.5 | 4223757.5 | 1768867.75 | 2 | 1095932 | 1665588.25 | 1271955.12 | 1 | 2 | 4 | 1 | 4 | 2 | 1.829256695 | 1.555045697 | 0.484661788 | 0.41200955 |
| Q91VK4 | Itm2c | Integral membrane protein 2C | 278.93 | 27.51 | 30482 | 5 | 750624.88 | 564020.69 | 328609.22 | 335555.47 | 740555 | 747145.12 | 695626 | 2 | 836358.62 | 263746.09 | 4748231.5 | 4340974 | 4261480.5 | 3937329.75 | 3955711 | 279467.75 | 407745.34 | 715998.19 | 2 | 764819.5 | 0 | 10 | 0 | 11.5 | 0 | 11.5 | 7.812016673 | 8.3542066 | 0.797256923 | 0.852590225 |
| Q9D892 | Itpa | Inosine triphosphate pyrophosphatase | 172.31 | 18.69 | 21897 | 3 | 6838660 | 5301901 | 7308243 | 9128029 | 9305616 | 5304724 | 6490282 | 7884006.5 | 8690226 | 7706751.5 | 2299797 | 3972023.25 | 4754557.5 | 3661656.5 | 3792391.75 | 7939065 | 6728717 | 7481885 | 8567852 | 7958636 | 0 | 12 | 0 | 9 | 0 | 12 | 0.487836095 | 0.512263863 | 1.020951813 | 1.072074668 |
| P11881 | Itpr1 | Inositol 1,4,5-trisphosphate receptor type 1 | 93.61 | 0.73 | 313167 | 2 | 330378.31 | 213152.36 | 226356.89 | 252296.06 | 296544.53 | 294254.16 | 318547 | 277317.75 | 225733 | 233852.44 | 155560.88 | 198080.47 | 2 | 206433.48 | 200314.98 | 2 | 2 | 110051.66 | 256834.36 | 167474.98 | 0 | 3 | 0 | 2 | 0 | 3 | 0.576609978 | 0.563376572 | 0.405212401 | 0.395912631 |
| P29995 | Itpr2 | Inositol 1,4,5-trisphosphate receptor type 2 | 79.06 | 0.74 | 307058 | 2 | 164675.09 | 185838.31 | 238587.45 | 231848.25 | 200027.3 | 119436.69 | 178348.72 | 244732.27 | 245750.3 | 212125.98 | 104741.24 | 267557.44 | 293144.62 | 206228.53 | 267429 | 2 | 2 | 151731.89 | 150227.38 | 201045.78 | 7 | 3 | 8 | 4 | 8 | 4 | 1.115697513 | 1.138652247 | 0.492674512 | 0.502810963 |
| Q9JHI5 | Ivd | Isovaleryl-CoA dehydrogenase, mitochondrial | 327.35 | 16.27 | 46325 | 7 | 7954733.5 | 7108030 | 7522451 | 8028482.5 | 8340546 | 7793459 | 7752770 | 6864505.5 | 7690445.5 | 8770396 | 8429215 | 6255998 | 6522889 | 8534583 | 7602623.5 | 7552245.5 | 10415766 | 8625345 | 9932914 | 9108460 | 12 | 3 | 9 | 5 | 12 | 5 | 0.95869681 | 0.960735641 | 1.17149576 | 1.173987144 |
| Q4KM64 | Jagn1 | Protein jagunal homolog 1 | 58.3 | 6.56 | 21102 | 1 | 2 | 2 | 2 | 2 | 2 | 2 | 2 | 2 | 2 | 2 | 677231.81 | 2 | 2 | 897573.44 | 493652.78 | 2 | 2 | 474391.91 | 2 | 2 | 5 | 10 | 5 | 10 | 5 | 10 | 206846.203 | 206846.203 | 47439.991 | 47439.991 |
| P52332 | Jak1 | Tyrosine-protein kinase JAK1 | 185.84 | 4.94 | 133367 | 3 | 920911.88 | 698609.69 | 1819136.75 | 895803.5 | 1774013.75 | 859161.25 | 1075368.75 | 2388282.5 | 1007940.5 | 748217.25 | 973775.62 | 769555.81 | 710491.62 | 748696.25 | 793740.81 | 1006836.69 | 704758 | 2094483.5 | 1052367.25 | 859197.06 | 7 | 12 | 5 | 11 | 7 | 12 | 0.654215616 | 0.657390964 | 0.936017904 | 0.940561027 |
| Q5BK20 | Jpt2 | Jupiter microtubule associated homolog 2 | 106.61 | 14.21 | 20070 | 2 | 1423872.38 | 945373.94 | 1275721.25 | 1368273.62 | 994401.38 | 1851251.75 | 1522901.25 | 1406579.5 | 1187856.5 | 1903906.75 | 2148752.25 | 2442462.5 | 1225073.38 | 2100165.5 | 1611039 | 1295488.75 | 1669351.25 | 1507249.12 | 1855999.25 | 1777350.62 | 3 | 2 | 6 | 12 | 6 | 12 | 1.585895386 | 1.210225186 | 1.349187954 | 1.029589503 |
| P09450 | Junb | Transcription factor jun-B | 53.79 | 4.07 | 35765 | 1 | 2 | 155699.38 | 2 | 2 | 2 | 2 | 2 | 2 | 2 | 2 | 440897.38 | 2 | 544555.31 | 795540.38 | 779976.88 | 2 | 2 | 2 | 2 | 2 | 3 | 10 | 2.5 | 12.5 | 3 | 12.5 | 16.44733827 | 256097.195 | 6.422303E-05 | 1 |
| P15066 | Jund | Transcription factor jun-D | 52.07 | 4.11 | 34905 | 1 | 2 | 2 | 2 | 2 | 2 | 2 | 2 | 2 | 2 | 488542.91 | 2 | 423875.91 | 2 | 2 | 2 | 2 | 2 | 2 | 2 | 2 | 10 | 12.5 | 12 | 10 | 12 | 12.5 | 42388.391 | 0.867635084 | 1 | 2.04687E-05 |
| Q02257 | Jup | Junction plakoglobin | 282.83 | 7.52 | 81801 | 5 | 2 | 3172692.25 | 2 | 2 | 324052.12 | 1130165.12 | 207011.45 | 1122637.62 | 440790.44 | 2 | 2 | 104572.15 | 240020.61 | 264330.78 | 1916451 | 425275.97 | 62110.26 | 800740.62 | 268623.5 | 713819.88 | 10.5 | 7 | 10.5 | 11 | 10.5 | 11 | 0.722206698 | 0.870637374 | 0.649337239 | 0.782791505 |
| Q99MN1 | Kars | Lysine--tRNA ligase | 582.03 | 19.33 | 67840 | 12 | 12846359 | 13768192 | 14812287 | 11741192 | 13898995 | 11986400 | 13348433 | 14859184 | 14313759 | 13778295 | 16617088 | 18498170 | 14495334 | 11915820 | 15225846 | 13845564 | 12206336 | 12720423 | 13193183 | 13794606 | 5 | 10 | 6 | 9 | 6 | 10 | 1.144411251 | 1.12398117 | 0.98051333 | 0.963009162 |
| Q5SVQ0 | Kat7 | Histone acetyltransferase KAT7 | 230.68 | 7.99 | 70641 | 4 | 1040964.75 | 2 | 13446040 | 1267955 | 1165461.25 | 1008862 | 714240.06 | 310453.03 | 1575951.12 | 2 | 1457631.12 | 1064517.12 | 991500.38 | 619630.88 | 273450.78 | 528875.06 | 1053449.62 | 1126110.25 | 955314.5 | 168121 | 9 | 7 | 11 | 11 | 11 | 11 | 0.260438541 | 1.220867227 | 0.226464222 | 1.061604575 |
| Q8BHI4 | Kbtbd3 | Kelch repeat and BTB domain-containing protein 3 | 38.41 | 1.15 | 69551 | 1 | 2 | 2 | 8081787.5 | 3707634 | 266872.09 | 4872034.5 | 2 | 2 | 2 | 2 | 2 | 2 | 2 | 2 | 2 | 2 | 4078737.75 | 2 | 2 | 2 | 5 | 8 | 10 | 12 | 10 | 12 | 8.29442E-07 | 2.052527E-06 | 0.338308317 | 0.83717368 |
| P62482 | Kcnab2 | Voltage-gated potassium channel subunit beta-2 | 376.36 | 31.61 | 41021 | 7 | 9748467 | 10016642 | 7334363.5 | 7986253.5 | 8699198 | 10059875 | 10894465 | 8956674 | 8866016 | 10199262 | 12526044 | 12972302 | 11365093 | 10543452 | 11654158 | 10799396 | 11341114 | 10300543 | 10759586 | 9639148 | 0 | 2 | 1 | 6 | 1 | 6 | 1.348890065 | 1.205910995 | 1.206803214 | 1.078885004 |
| Q6WVG3 | Kctd12 | BTB/POZ domain-containing protein KCTD12 | 361.19 | 24.16 | 35892 | 6 | 681132.56 | 1327032.25 | 738368.94 | 4154678 | 4938768.5 | 799873.94 | 934992.94 | 2818986.5 | 3991157 | 3050283 | 7159216 | 5970121 | 1767244.62 | 1864299 | 4064487.75 | 4453968.5 | 5870222.5 | 3410911.75 | 994118.75 | 4194201.75 | 6 | 7 | 6 | 4 | 6 | 7 | 1.75890229 | 1.796019099 | 1.598264765 | 1.63199176 |
| Q9JHP7 | Kdelc1 | KDEL motif-containing protein 1 | 223.61 | 6.37 | 57985 | 4 | 691918.75 | 496134.69 | 243040.03 | 2 | 797195.38 | 585918 | 772248.75 | 479590.34 | 763722.5 | 175244.22 | 336314.19 | 638945.56 | 657477.88 | 397527.56 | 562777.81 | 246260.53 | 147939.12 | 188024.17 | 2 | 2 | 12 | 5 | 10 | 2 | 12 | 5 | 1.163691445 | 0.933849809 | 0.26128897 | 0.209681574 |
| Q566E5 | Kdelc2 | KDEL motif-containing protein 2 | 118.9 | 3.94 | 58701 | 2 | 2 | 2 | 229625.2 | 220294.27 | 2 | 257863.22 | 2 | 239611.95 | 2 | 222352.22 | 2 | 2 | 321408.03 | 2 | 2 | 2 | 2 | 2 | 126443.26 | 2 | 11 | 9 | 9 | 6 | 11 | 9 | 0.714376161 | 0.446515718 | 0.281049348 | 0.175667888 |
| Q569A6 | Kdelr1 | ER lumen protein-retaining receptor 1 | 116.48 | 10.38 | 24546 | 2 | 177391.19 | 128838.37 | 182111.78 | 423254.69 | 252681.19 | 422402.44 | 321545.25 | 222622.94 | 389383.69 | 570983.88 | 637631.81 | 357808.31 | 1323578 | 1547044.12 | 1105377.12 | 343187.38 | 1107004.75 | 1066359.5 | 1304985.38 | 905430.25 | 1 | 1 | 3 | 3 | 3 | 3 | 4.269979069 | 2.579968242 | 4.0600015 | 2.453097489 |
| P59997 | Kdm2a | Lysine-specific demethylase 2A | 56.95 | 0.86 | 132680 | 1 | 394187.16 | 378353.81 | 476644.09 | 567995.38 | 520363.97 | 516943.25 | 436660.91 | 2 | 350570.59 | 476068.66 | 146290.67 | 310147.97 | 264789.38 | 348998.78 | 311131.16 | 467952.81 | 552363.69 | 350871.56 | 526579.94 | 545058.81 | 0 | 11 | 5 | 5 | 5 | 11 | 0.59094405 | 0.775936819 | 1.045039743 | 1.372185428 |
| Q60749 | Khdrbs1 | KH domain-containing, RNA-binding, signal transduction-associated protein 1 | 306.69 | 23.48 | 48371 | 7 | 10043184 | 10142487 | 12424910 | 9717746 | 8472517 | 9451732 | 10767751 | 11640836 | 9190091 | 10868157 | 9023275 | 10287188 | 9420337 | 8293171 | 9941180 | 11072443 | 8675297 | 12075962 | 11227906 | 13131325 | 8 | 7 | 5 | 7 | 8 | 7 | 0.924495487 | 0.90459259 | 1.105944874 | 1.082135664 |
| Q3U0V1 | Khsrp | Far upstream element-binding protein 2 | 1218.73 | 34.09 | 76775 | 20 | 49038038 | 50065806 | 49674678 | 58683110 | 52704554 | 53206626 | 49638854 | 46294770 | 45225606 | 49750806 | 36379470 | 34166870 | 31685970 | 36121614 | 30526322 | 57075542 | 71344698 | 54589390 | 57041538 | 50320250 | 0 | 5 | 0 | 1 | 0 | 5 | 0.649124502 | 0.69180139 | 1.116099761 | 1.189478078 |
| Q9EQG6 | Kidins220 | Kinase D-interacting substrate of 220 kDa | 131.24 | 2.16 | 195716 | 3 | 196339.48 | 176005.66 | 94377.62 | 882159.31 | 201447.47 | 198711.97 | 252331.3 | 2 | 272423.16 | 237836.84 | 2 | 1183584.25 | 1245977.5 | 1225350.88 | 998675.75 | 241093.23 | 344337.47 | 682989.75 | 267584.31 | 1063273.75 | 5 | 4 | 4.5 | 3 | 5 | 4 | 3.001678198 | 4.840908008 | 1.676597422 | 2.703905399 |
| Q6P9P6 | Kif11 | Kinesin-like protein KIF11 | 55.8 | 1.43 | 118027 | 1 | 2 | 175401.94 | 176720.22 | 2 | 250081.56 | 2 | 139531.36 | 2 | 242201.75 | 2 | 2 | 151950.41 | 2 | 2 | 2 | 172810.48 | 2 | 2 | 2 | 2 | 6 | 6 | 10 | 10 | 10 | 10 | 0.25233554 | 0.398068749 | 0.286974866 | 0.452713582 |
| Q6P9L6 | Kif15 | Kinesin-like protein KIF15 | 117.51 | 1.8 | 160120 | 2 | 2 | 226998.3 | 195330.97 | 2 | 216375.25 | 2 | 2 | 2 | 106662.23 | 155039.91 | 2 | 2 | 2 | 2 | 2 | 202352.25 | 2 | 178307.47 | 2 | 2 | 5 | 8 | 7.5 | 10.5 | 7.5 | 10.5 | 1.565659E-05 | 3.82105E-05 | 0.595992864 | 1.454542912 |
| F1M5N7 | Kif21b | Kinesin-like protein KIF21B | 40.75 | 0.49 | 182563 | 1 | 6733384.5 | 4206716 | 682251.06 | 7773954.5 | 4472198.5 | 8669874 | 5181100 | 4881913 | 2024174.62 | 663443.75 | 2116489.75 | 2171783.25 | 2293462.5 | 2935891 | 2114741 | 4338996.5 | 6872140 | 6657561.5 | 4291220 | 5560767 | 5 | 11 | 10 | 9 | 10 | 11 | 0.48735217 | 0.543048229 | 1.16139178 | 1.294119094 |
| P33174 | Kif4 | Chromosome-associated kinesin KIF4 | 99.36 | 1.62 | 139519 | 2 | 136897.03 | 678491.94 | 720219.94 | 910966.5 | 841799.75 | 421023.69 | 91950.51 | 719713.12 | 911181.06 | 538523.38 | 392604.91 | 711047.69 | 510418.62 | 620548.81 | 283253.44 | 538779.81 | 49478.34 | 440415.81 | 2 | 375068.88 | 6 | 3 | 10 | 6 | 10 | 6 | 0.765689238 | 0.938667315 | 0.426880989 | 0.523318354 |
| Q2PQA9 | Kif5b | Kinesin-1 heavy chain | 634.49 | 12.25 | 109531 | 9 | 4062622.25 | 4825406 | 4382943.5 | 5144260.5 | 4047935 | 4152071.25 | 4551116 | 3452826.5 | 4122998 | 4767190.5 | 4087643 | 5049827 | 3381017.75 | 3428278.75 | 3370014 | 4213868.5 | 3787950.5 | 5671155 | 5967589 | 5342108 | 6 | 8 | 6 | 6 | 6 | 8 | 0.859931295 | 0.917827372 | 1.112161554 | 1.187039386 |
| P70188 | Kifap3 | Kinesin-associated protein 3 | 75.2 | 2.52 | 91291 | 1 | 198499.31 | 179123.69 | 276813.28 | 321219.25 | 316118.97 | 324011.88 | 158522.8 | 303480.44 | 411689.22 | 311406.94 | 88750.02 | 196382.48 | 223778.09 | 321311.41 | 167411.95 | 207248.14 | 288721.44 | 370876.16 | 590049.75 | 251016.45 | 8 | 8 | 6 | 12 | 8 | 12 | 0.772297293 | 0.661073814 | 1.322144027 | 1.131733599 |
| O88447 | Klc1 | Kinesin light chain 1 | 139.11 | 5.18 | 61450 | 3 | 1234729 | 1329136.75 | 1832212.75 | 1471942.75 | 2459430.75 | 1552733.25 | 1545023.12 | 1926138.25 | 2372504 | 1808694.75 | 869947.56 | 514727.22 | 1023695.62 | 1434415.62 | 1109551.75 | 2773553.25 | 2654447 | 1926038.25 | 2834221.75 | 3933038 | 2 | 1 | 0 | 2 | 2 | 2 | 0.594700248 | 0.537999732 | 1.695752584 | 1.534074418 |
| Q9DBS5 | Klc4 | Kinesin light chain 4 | 111.24 | 4.04 | 68613 | 2 | 2 | 506637.66 | 2 | 473443.34 | 2 | 592276.62 | 2 | 458084.06 | 368718.56 | 2 | 2 | 2 | 2 | 2 | 2 | 433792.09 | 2 | 567470.44 | 639219.25 | 548769.06 | 7.5 | 5.5 | 5 | 8 | 7.5 | 8 | 1.020318E-05 | 7.046803E-06 | 2.233733169 | 1.542723343 |
| Q921I2 | Klhdc4 | Kelch domain-containing protein 4 | 57.22 | 2.57 | 64860 | 1 | 711672.81 | 1127711.75 | 1251904.88 | 1313718.38 | 660949.94 | 692486.31 | 794199.25 | 753414.5 | 663563.38 | 599258.31 | 2 | 723639.44 | 2 | 2 | 293572.75 | 1350565.5 | 2 | 989340.06 | 2 | 859778.12 | 2 | 9 | 3 | 10 | 3 | 10 | 0.200794842 | 0.290391354 | 0.631605677 | 0.913433958 |
| Q8C3Y4 | Kntc1 | Kinetochore-associated protein 1 | 40.56 | 0.59 | 250358 | 1 | 2 | 2 | 226789.06 | 341455.78 | 370607.16 | 2 | 30868.52 | 265875.5 | 243533.75 | 215387.59 | 2 | 2 | 224117.75 | 199224.09 | 209565.14 | 2 | 2 | 213255.89 | 244469.08 | 178319.72 | 8 | 9 | 8 | 9 | 8 | 9 | 0.674129984 | 0.837552359 | 0.67747204 | 0.841704596 |
| P52293 | Kpna2 | Importin subunit alpha-1 | 1024.73 | 38.94 | 57928 | 17 | 22200578 | 19943930 | 18613204 | 14678310 | 15951986 | 21973154 | 17812658 | 12809629 | 13428085 | 10944268 | 12243832 | 13568039 | 9762665 | 9236812 | 10206116 | 21722640 | 20739590 | 14108494 | 13122830 | 10767671 | 0 | 8 | 4 | 12 | 4 | 12 | 0.602020606 | 0.714811496 | 0.880435265 | 1.045388218 |
| O35344 | Kpna3 | Importin subunit alpha-4 | 257.33 | 9.6 | 57773 | 5 | 2647977.5 | 2783474.25 | 3592929 | 1348126.12 | 3422469.5 | 2913742 | 3189269 | 3751135.5 | 3670768.25 | 3702509.25 | 3061087 | 3251535.25 | 3725950.5 | 2772714.25 | 3326541.5 | 2984272.75 | 1966333.25 | 3461621.5 | 3139854.5 | 1579036.25 | 9 | 12 | 9 | 4 | 9 | 12 | 1.169833718 | 0.936752268 | 0.951876821 | 0.762221807 |
| O35345 | Kpna6 | Importin subunit alpha-7 | 201.59 | 8.58 | 59964 | 4 | 877408.94 | 436694.09 | 336505.53 | 422007.56 | 282008.06 | 992727.44 | 818428.62 | 151965.16 | 2 | 484646.28 | 1214997.38 | 1113050.5 | 951893.94 | 1053630.88 | 841277.25 | 535373.62 | 648376.38 | 151039.14 | 303494.16 | 2 | 1 | 9 | 2 | 9.5 | 2 | 9.5 | 2.197739238 | 2.114108355 | 0.695773582 | 0.669297211 |
| P70168 | Kpnb1 | Importin subunit beta-1 | 1089.07 | 26.14 | 97184 | 17 | 36525286 | 41772882 | 42326666 | 39070974 | 40922258 | 43487946 | 40615470 | 38205262 | 43298882 | 39054342 | 46304814 | 43665322 | 40873002 | 38839354 | 38599338 | 40089066 | 46993502 | 41664970 | 42512650 | 41438790 | 11 | 7 | 10 | 9 | 11 | 9 | 1.038200767 | 1.017687356 | 1.060218465 | 1.039270015 |
| Q8BGA5 | Krr1 | KRR1 small subunit processome component homolog | 124.48 | 6.58 | 43538 | 3 | 6275591 | 7595902 | 6304253 | 7586612.5 | 6556413.5 | 5938185.5 | 7734956.5 | 6953644.5 | 5881749.5 | 6923790 | 6597206 | 8562543 | 9416835 | 6524481.5 | 8287656.5 | 8708950 | 5280160.5 | 8262376 | 7168474 | 9182037 | 5 | 7 | 6 | 6 | 6 | 7 | 1.147731102 | 1.178162776 | 1.124807074 | 1.154630925 |
| P0C7H8 | KRTAP2-3 | Keratin-associated protein 2-3 Homo sapiens OX=9606 | 145.59 | 35.94 | 13480 | 3 | 2 | 2 | 242852.27 | 2 | 2 | 2 | 2 | 2 | 2 | 156200.89 | 2 | 355386.84 | 2 | 2 | 2 | 2 | 2 | 442802.62 | 175659.17 | 2 | 12 | 10 | 12 | 9 | 12 | 10 | 1.463371675 | 2.275125571 | 2.546599285 | 3.959235547 |
| Q61595 | Ktn1 | Kinectin | 126.94 | 2.41 | 152592 | 2 | 277452.62 | 354237.5 | 470942.06 | 560805.31 | 423850.56 | 355981.19 | 412423.44 | 398571.62 | 472190.44 | 229052.86 | 633446 | 660767.44 | 472053.5 | 440415.44 | 267487.19 | 530268.12 | 470379.25 | 402853.78 | 355768.5 | 460436.44 | 8 | 11 | 6 | 8 | 8 | 11 | 1.185351284 | 1.324346258 | 1.063440233 | 1.188139847 |
| Q58FK9 | Kyat3 | Kynurenine--oxoglutarate transaminase 3 | 40.2 | 2.64 | 51044 | 1 | 2 | 2 | 826640.88 | 909721.56 | 871484.75 | 1168012.75 | 1483824.75 | 2 | 2 | 596012.94 | 2 | 2 | 2 | 2 | 2 | 1020602.88 | 2 | 2 | 2 | 2 | 5 | 9 | 5 | 7 | 5 | 9 | 3.834575E-06 | 3.078956E-06 | 0.391360858 | 0.314241571 |
| Q9EP89 | Lactb | Serine beta-lactamase-like protein LACTB, mitochondrial | 189.55 | 5.99 | 60705 | 3 | 1807807.38 | 1984280.25 | 1597271.25 | 1171047.75 | 2270608 | 1530411.38 | 1861333.88 | 1994968.75 | 1863822.88 | 1675037.75 | 395289.97 | 213890.73 | 2 | 143068.91 | 861726.12 | 1599788.88 | 2 | 921395.12 | 2 | 128094.3 | 0 | 2 | 0 | 1 | 0 | 2 | 0.182762434 | 0.180826198 | 0.299997499 | 0.296819242 |
| P02468 | Lamc1 | Laminin subunit gamma-1 | 132.82 | 1.93 | 177298 | 2 | 2 | 2 | 152717.78 | 2 | 2 | 242443.89 | 2 | 364137.22 | 2 | 389740.38 | 2 | 220980.95 | 2 | 382589 | 2 | 2 | 2 | 341075.28 | 374451.12 | 2 | 9 | 9 | 9 | 10 | 9 | 10 | 3.952024013 | 0.605801975 | 4.685079363 | 0.718171328 |
| P11438 | Lamp1 | Lysosome-associated membrane glycoprotein 1 | 650.84 | 17.98 | 43865 | 8 | 74331226 | 62572362 | 52140682 | 51874898 | 73828626 | 62319906 | 72577098 | 79394050 | 93211778 | 79187994 | 45423382 | 53944802 | 65761122 | 51756866 | 70880122 | 39514306 | 70479338 | 63548010 | 67927218 | 75876026 | 8 | 11 | 2 | 5 | 8 | 11 | 0.914275809 | 0.744176677 | 1.008251381 | 0.820668288 |
| P17047 | Lamp2 | Lysosome-associated membrane glycoprotein 2 | 507.2 | 19.28 | 45681 | 9 | 157081170 | 137547010 | 71488938 | 107170114 | 168836066 | 119275178 | 156146098 | 78395298 | 151156914 | 75984778 | 56481570 | 83011986 | 121031274 | 45781762 | 87051602 | 152674546 | 51581634 | 110279234 | 93187634 | 100482042 | 4 | 7 | 7 | 10 | 7 | 10 | 0.612589818 | 0.677085114 | 0.791444714 | 0.874770392 |
| Q6P791 | Lamtor1 | Ragulator complex protein LAMTOR1 | 151.54 | 16.15 | 17721 | 2 | 2058218.25 | 1895710.62 | 1750621.88 | 637387.25 | 1167043.25 | 2030561.75 | 1864637.62 | 1568915.38 | 1784541.88 | 1708941 | 1589965.38 | 2 | 1668948.88 | 1568703.5 | 1613477 | 1332992.5 | 2461616.5 | 1562254.75 | 1959117.5 | 1290898 | 8 | 10 | 3 | 9 | 8 | 10 | 0.857785703 | 0.719065203 | 1.146211312 | 0.960846826 |
| Q9JHS3 | Lamtor2 | Ragulator complex protein LAMTOR2 | 354.98 | 75.2 | 13480 | 5 | 2430122.75 | 1701781.88 | 1790673.12 | 1979958.5 | 2027068.75 | 2618831 | 2761324.5 | 1674837.5 | 1361867 | 597836.06 | 1500191.75 | 1454392.5 | 2138292.25 | 2240773.5 | 987918.75 | 2210643.5 | 2027906 | 1589974.5 | 2225127 | 1140249.5 | 8 | 12 | 11 | 12 | 11 | 12 | 0.838056373 | 0.923111405 | 0.925907979 | 1.019879144 |
| Q9D1L9 | Lamtor5 | Ragulator complex protein LAMTOR5 | 54.13 | 21.98 | 9642 | 1 | 403871.72 | 277150.12 | 833928.75 | 420557.78 | 848130.06 | 2 | 2 | 429125.91 | 2 | 2 | 2 | 2 | 300533.81 | 602591.44 | 552283.75 | 720479.19 | 2 | 987170.44 | 622620.75 | 1169893.5 | 7 | 9 | 7 | 3 | 7 | 9 | 0.522845562 | 3.391512454 | 1.257406796 | 8.156348866 |
| O89112 | Lancl1 | LanC-like protein 1 | 59.54 | 3.01 | 45341 | 1 | 483709.09 | 297335.81 | 2 | 2 | 656428.06 | 2 | 573104.44 | 2 | 639374.25 | 600430.19 | 2 | 2 | 668446.81 | 626278.69 | 804141.44 | 420171.94 | 2 | 2 | 800374.69 | 2 | 9 | 11 | 9 | 10 | 9 | 11 | 1.460107535 | 1.157734033 | 0.8490937 | 0.673254983 |
| Q9CPY7 | Lap3 | Cytosol aminopeptidase | 622.06 | 26.78 | 56141 | 12 | 8397478 | 8118254.5 | 8317390 | 9704926 | 7873057.5 | 9417272 | 8772117 | 7966034 | 8441125 | 9579080 | 7253780.5 | 9125605 | 9330653 | 10211206 | 7550415.5 | 8514058 | 10474010 | 9784454 | 9519751 | 8888319 | 12 | 3 | 11 | 6 | 12 | 6 | 1.025006516 | 0.984064335 | 1.112458421 | 1.068023119 |
| Q6ZQ58 | Larp1 | La-related protein 1 | 277.35 | 5.69 | 121145 | 5 | 2294953.25 | 2625546 | 2870345.5 | 5316165 | 2320418.5 | 2776398.5 | 2736962.5 | 2723592.5 | 3483347.75 | 2434261.5 | 2949238.25 | 3477564.5 | 3402420 | 2722662.5 | 3900618.5 | 2786696 | 3048197.5 | 3214397.75 | 3146815.25 | 2715963.5 | 6 | 7 | 7 | 8 | 7 | 8 | 1.066445002 | 1.162346308 | 0.966594675 | 1.053516824 |
| Q8BWW4 | Larp4 | La-related protein 4 | 210.68 | 4.31 | 79763 | 3 | 2780900 | 4053761.25 | 2962635 | 3630299 | 2553752.5 | 4192984.75 | 4243206.5 | 3423211.25 | 1734528 | 2117261.5 | 5308124 | 4260656.5 | 4020580 | 3980024.75 | 3018352.5 | 3868441.5 | 4264698 | 3839415.75 | 4223057 | 4146186.75 | 4 | 2 | 7 | 7 | 7 | 7 | 1.28823539 | 1.310386745 | 1.272846278 | 1.294733016 |
| Q6A0A2 | Larp4b | La-related protein 4B | 289.5 | 6.75 | 81627 | 4 | 1021299.5 | 1972223.5 | 1362598 | 899060 | 1807379.12 | 2886783.25 | 2451982.75 | 1145353.25 | 2204609.25 | 553350.38 | 922216.94 | 204993.75 | 1221017.5 | 1512381.12 | 2 | 342352.12 | 1167078.25 | 2181429.75 | 2247568.25 | 2404773.25 | 6 | 8 | 5 | 10 | 6 | 10 | 0.546630577 | 0.417721095 | 1.181328226 | 0.902740793 |
| Q05CL8 | Larp7 | La-related protein 7 | 104.83 | 3.51 | 64802 | 2 | 2196744 | 1754356.25 | 2015552.12 | 2366926.75 | 1449956.5 | 2198811.25 | 1628422.75 | 1426479 | 1410519.5 | 1648539.25 | 1126642.62 | 1410522.88 | 1101205.12 | 1219446.25 | 1282347.88 | 1712274.25 | 1988598 | 1516400.88 | 1808161 | 1540384.5 | 0 | 7 | 1 | 9 | 1 | 9 | 0.627601819 | 0.738642289 | 0.875534057 | 1.030440735 |
| Q8BMJ2 | Lars | Leucine--tRNA ligase, cytoplasmic | 1553.93 | 23.01 | 134192 | 26 | 28805952 | 34325366 | 28222182 | 27971648 | 28316818 | 31343252 | 31594788 | 26374090 | 27425136 | 27230342 | 47512778 | 51184230 | 42921110 | 32073174 | 43096006 | 32615382 | 25850194 | 31438176 | 29499446 | 30783816 | 1 | 9 | 0 | 10 | 1 | 10 | 1.468331152 | 1.50580607 | 1.017237971 | 1.043200037 |
| A2BE28 | Las1l | Ribosomal biogenesis protein LAS1L | 155.84 | 3.99 | 89414 | 2 | 1717041.75 | 1158472.38 | 843209.12 | 372360.28 | 1029906.88 | 576772.06 | 822305.94 | 1169892.25 | 1034128.25 | 575473.25 | 547580.75 | 1380931.62 | 1097741.88 | 584831 | 1144993 | 1346149.75 | 1096405.88 | 884425.62 | 440099.34 | 500208.44 | 12 | 11 | 10 | 12 | 12 | 12 | 0.928741878 | 1.138206673 | 0.833293697 | 1.021231484 |
| Q61792 | Lasp1 | LIM and SH3 domain protein 1 | 694.52 | 44.49 | 29994 | 13 | 40668454 | 53759694 | 30460392 | 38461922 | 33992234 | 49109230 | 43939358 | 50720022 | 43300358 | 52377866 | 37850350 | 40106578 | 37798158 | 40125866 | 36980098 | 38978646 | 39563194 | 59394058 | 53033914 | 61842046 | 12 | 5 | 0 | 10 | 12 | 10 | 0.977290034 | 0.805444143 | 1.28108039 | 1.055816249 |
| Q3U9G9 | Lbr | Lamin-B receptor | 873.65 | 25.4 | 71440 | 14 | 5259683.5 | 4856474.5 | 7822537 | 8392977 | 7031607.5 | 6342602.5 | 6187232 | 7406558 | 7967984.5 | 9603967 | 16012562 | 13508586 | 20155752 | 19196358 | 21768552 | 5297935 | 6453005 | 9279695 | 9859510 | 7567412 | 0 | 8 | 0 | 11 | 0 | 11 | 2.716813555 | 2.416577229 | 1.15269115 | 1.025306716 |
| Q8BYR1 | Lcmt2 | tRNA wybutosine-synthesizing protein 4 | 52.12 | 2.04 | 75295 | 2 | 22302478 | 22041454 | 21297048 | 23016696 | 18353986 | 25093674 | 14945009 | 20615702 | 16699947 | 22241666 | 26546082 | 21469974 | 23440078 | 18289988 | 23052522 | 20228358 | 17784372 | 23179840 | 25899206 | 22557756 | 8 | 10 | 7 | 8 | 8 | 10 | 1.05407805 | 1.132562013 | 1.024650304 | 1.100943152 |
| Q61233 | Lcp1 | Plastin-2 | 4410.49 | 82.3 | 70149 | 63 | 793658882 | 588158786 | 818221442 | 540277890 | 904508930 | 733736130 | 806224322 | 817528962 | 673548866 | 577219906 | 520396514 | 613904258 | 376070914 | 318837890 | 503683554 | 742758594 | 797554818 | 534501314 | 526085122 | 742386754 | 2 | 7 | 1 | 9 | 2 | 9 | 0.640056116 | 0.646542739 | 0.91726921 | 0.926565237 |
| Q60787 | Lcp2 | Lymphocyte cytosolic protein 2 | 265.97 | 9.19 | 60238 | 4 | 2112590.75 | 2041280.12 | 840686.12 | 245124.47 | 1353339.5 | 1635441.75 | 1200299.5 | 959365.81 | 1247842.25 | 533298.19 | 3326787.25 | 4513237 | 2486341 | 2264823 | 2817614.75 | 846759.25 | 1865509.38 | 213854.94 | 1110590.88 | 572904.75 | 0 | 8 | 0 | 9 | 0 | 9 | 2.337138482 | 2.763292519 | 0.699166471 | 0.826652547 |
| P06151 | Ldha | L-lactate dehydrogenase A chain | 1981.46 | 76.2 | 36499 | 27 | 741478338 | 758384898 | 625314434 | 694115586 | 740172546 | 689171074 | 765536514 | 539678466 | 678527042 | 612393154 | 491122018 | 547123842 | 588786626 | 484956546 | 602463042 | 450054146 | 656085570 | 672363010 | 460264258 | 443298530 | 0 | 2 | 3 | 4 | 3 | 4 | 0.76260097 | 0.826240194 | 0.753502257 | 0.816382191 |
| P35951 | Ldlr | Low-density lipoprotein receptor | 316.43 | 7.19 | 94947 | 5 | 661465.25 | 1033295.38 | 936469.56 | 1295552 | 781878.75 | 1051792.62 | 893630.88 | 2 | 2 | 256080.97 | 1722112.5 | 4132536.75 | 3733809.75 | 1971964.5 | 4487035 | 412995.34 | 455623.84 | 377740.19 | 1707570.5 | 599742.38 | 0 | 5 | 0 | 8 | 0 | 8 | 3.408072636 | 7.289301276 | 0.754709735 | 1.614198763 |
| Q6DVA0 | Lemd2 | LEM domain-containing protein 2 | 113.19 | 5.09 | 57507 | 2 | 739807.38 | 966512.31 | 1166310.5 | 666374.25 | 1473873.25 | 481684.5 | 1043519.88 | 988048.69 | 1434808.75 | 834660.19 | 445990.47 | 954363.62 | 474460.44 | 875565.62 | 472043.56 | 956728.38 | 829850.38 | 683156 | 807044.06 | 840066.94 | 4 | 9 | 4 | 7 | 4 | 9 | 0.642829111 | 0.67376354 | 0.821253981 | 0.860774628 |
| Q9Z2I0 | Letm1 | Mitochondrial proton/calcium exchanger protein | 263.78 | 9.21 | 82989 | 4 | 500250.19 | 876857.75 | 2472207.25 | 1211460.38 | 2335306 | 1013646.69 | 891427.62 | 2409222.5 | 2147029.75 | 2889788.75 | 2695883.5 | 3068204 | 3903284.5 | 2489792.75 | 2188538.25 | 124285.63 | 424826.62 | 1269918.62 | 1835252.75 | 1862872.25 | 2 | 9 | 4 | 6 | 4 | 9 | 1.939635585 | 1.53411679 | 0.745956601 | 0.589999769 |
| P16045 | Lgals1 | Galectin-1 | 781.69 | 79.26 | 14866 | 10 | 281825570 | 248817506 | 225968786 | 257049282 | 282939010 | 265381122 | 275253538 | 220794546 | 237212802 | 261775538 | 99424202 | 121197754 | 107703250 | 109459338 | 107044658 | 290200930 | 287853922 | 230620130 | 260718962 | 244264818 | 0 | 10 | 0 | 10 | 0 | 10 | 0.420198317 | 0.432260883 | 1.013156414 | 1.042240935 |
| P16110 | Lgals3 | Galectin-3 | 430.33 | 21.59 | 27515 | 7 | 47853130 | 61230842 | 25886782 | 31648020 | 20372544 | 60890106 | 67134522 | 45795982 | 31408114 | 37851074 | 54942358 | 48347106 | 27178570 | 30006792 | 34761046 | 62014574 | 64364898 | 52578442 | 48663110 | 54971894 | 10 | 3 | 7 | 8 | 10 | 8 | 1.044090571 | 0.80317605 | 1.511262239 | 1.162552052 |
| Q07797 | Lgals3bp | Galectin-3-binding protein | 1553.83 | 42.29 | 64491 | 21 | 50600058 | 47346874 | 33928138 | 39147942 | 33747850 | 41281026 | 34474322 | 30951064 | 34037650 | 37295994 | 30774034 | 32591680 | 49370922 | 42413154 | 51517270 | 53785714 | 62299454 | 38081138 | 49556010 | 53247534 | 12 | 4 | 9 | 1 | 12 | 4 | 1.009260097 | 1.160789682 | 1.254914139 | 1.443326046 |
| Q9JL15 | Lgals8 | Galectin-8 | 64.58 | 4.43 | 36162 | 1 | 2 | 2 | 2 | 2 | 2 | 2 | 2 | 2 | 2 | 493358.12 | 2 | 2 | 207461.83 | 2 | 2 | 2 | 2 | 2 | 2 | 2 | 10 | 12.5 | 12 | 10 | 12 | 12.5 | 20746.983 | 0.420519005 | 1 | 2.026892E-05 |
| O08573 | Lgals9 | Galectin-9 | 153.25 | 11.61 | 40036 | 3 | 1869279.12 | 2123704 | 1868761.62 | 1857493.38 | 1824544.38 | 1437355.38 | 1733110.12 | 1339437.5 | 1406830.75 | 1991098.88 | 2152989.5 | 2122951.25 | 1829442.25 | 1839309.38 | 1669148.75 | 1306399.38 | 2146667.25 | 1410690.62 | 2539072 | 1697757.38 | 11 | 10 | 4 | 10 | 11 | 10 | 1.007340761 | 1.215736546 | 0.953561822 | 1.150831973 |
| O89017 | Lgmn | Legumain | 160.48 | 4.83 | 49373 | 2 | 1776030.75 | 2211846.5 | 1369836.5 | 189032.61 | 3228589.5 | 651365.44 | 1865417 | 1625744 | 2739611.5 | 703187.81 | 366197.44 | 2 | 2 | 2 | 2 | 1485656 | 616930.5 | 417006.81 | 317674.44 | 708792.06 | 1 | 6 | 0 | 4 | 1 | 6 | 0.041731216 | 0.048278143 | 0.404093914 | 0.467489456 |
| P37913 | Lig1 | DNA ligase 1 | 537.95 | 16.38 | 102290 | 11 | 8266347 | 6896158 | 8434862 | 7272140.5 | 5771376 | 7861887.5 | 6763263 | 6344585 | 4718601 | 4825132.5 | 6417197.5 | 6597925.5 | 4231790.5 | 5524123.5 | 5577091 | 7042865 | 5687491.5 | 5551770.5 | 4442994 | 5619220.5 | 2 | 2 | 10 | 10 | 10 | 10 | 0.773674794 | 0.929036551 | 0.773571453 | 0.928912458 |
| P97386 | Lig3 | DNA ligase 3 | 56.44 | 0.99 | 113072 | 1 | 883968.88 | 774607.19 | 1095353.5 | 621403.69 | 926050.5 | 863298.38 | 831029.5 | 707437.56 | 777002.94 | 741963.25 | 557919.75 | 460552.78 | 452626.53 | 635358.38 | 822875.12 | 673641.31 | 787258.69 | 796097.31 | 1080106.88 | 969107.69 | 3 | 12 | 3 | 9 | 3 | 12 | 0.681020975 | 0.747139268 | 1.001122457 | 1.098318448 |
| Q64281 | Lilrb4 | Leukocyte immunoglobulin-like receptor subfamily B member 4 | 500.66 | 28.36 | 37544 | 6 | 3365518.5 | 1745999.75 | 2115142 | 2420224 | 3628811 | 4115080 | 4221960.5 | 4682811 | 4617899 | 4979627 | 8733989 | 7878697 | 8406379 | 7809151 | 7303720.5 | 3501911.75 | 3558118.5 | 4233616 | 3625183.5 | 3347723.5 | 0 | 5 | 0 | 2 | 0 | 5 | 3.022963072 | 1.77438505 | 1.375939482 | 0.807633566 |
| Q9ERG0 | Lima1 | LIM domain and actin-binding protein 1 | 331.74 | 11.69 | 84060 | 7 | 1883585.38 | 4203929.5 | 2054253.25 | 2438088 | 1091149 | 2920460 | 4229631.5 | 2574031 | 2619679 | 3560270.75 | 4395186 | 4165840.75 | 1490866.25 | 3643461 | 3069880 | 3222910 | 2503357.75 | 4191560 | 3012079.25 | 4095693 | 7 | 5 | 9 | 11 | 9 | 11 | 1.436485874 | 1.054147248 | 1.458794663 | 1.070518276 |
| Q9Z0M5 | Lipa | Lysosomal acid lipase/cholesteryl ester hydrolase | 366.45 | 11.34 | 45325 | 7 | 12357164 | 11581532 | 9505586 | 14203772 | 12515728 | 19473484 | 21954418 | 28999422 | 30067432 | 28500100 | 8395092 | 9856939 | 10709936 | 8424222 | 12104490 | 18646134 | 17713950 | 21487026 | 20507386 | 23552954 | 4 | 0 | 0 | 4 | 4 | 4 | 0.822599201 | 0.383663973 | 1.693833842 | 0.790011735 |
| P15304 | Lipe | Hormone-sensitive lipase | 64.15 | 1.5 | 116812 | 1 | 150339.36 | 2 | 254839.17 | 216381.53 | 193696.98 | 194322.08 | 164848.53 | 185698.78 | 190369.94 | 237233.33 | 255467.16 | 265122.56 | 192973.23 | 389269.31 | 313292.03 | 2 | 260893.17 | 2 | 2 | 190920.08 | 3 | 8.5 | 2 | 8 | 3 | 8.5 | 1.737023719 | 1.456209874 | 0.5542033 | 0.464608691 |
| Q9D945 | Llph | Protein LLP homolog | 61.44 | 15.38 | 15391 | 1 | 202180.92 | 214688.22 | 119660.26 | 376084.81 | 156020.62 | 152453.47 | 277811.84 | 175173.23 | 63583.95 | 69881.49 | 74713.06 | 302010.53 | 53126.07 | 759829.19 | 423511.81 | 52190.16 | 315842.28 | 435162.62 | 228938.3 | 364214.22 | 11 | 8 | 8 | 6 | 11 | 8 | 1.509580836 | 2.183220965 | 1.30666486 | 1.889755121 |
| Q9D0F3 | Lman1 | Protein ERGIC-53 | 549.78 | 26.69 | 57789 | 9 | 6423150.5 | 8275196 | 10211947 | 9198566 | 9584530 | 8135166.5 | 7710888.5 | 10351762 | 11277188 | 8645153 | 12454653 | 12536926 | 15139685 | 15051736 | 13324501 | 8285251 | 9182452 | 9929818 | 12054200 | 10191274 | 0 | 8 | 0 | 9 | 0 | 9 | 1.567914547 | 1.485413406 | 1.136167177 | 1.076383888 |
| Q9DBH5 | Lman2 | Vesicular integral-membrane protein VIP36 | 350.07 | 26.82 | 40430 | 7 | 763744.06 | 3320360 | 4822938 | 2338897 | 998194.81 | 1500784 | 1657162.75 | 3605305 | 3722894.5 | 4155203 | 6796239.5 | 3749628 | 7523245.5 | 4496914 | 3858208 | 3564578.5 | 2688383 | 4438215 | 3379571.25 | 3622453.5 | 3 | 6 | 2 | 11 | 3 | 11 | 2.158113859 | 1.804767754 | 1.44503494 | 1.208440626 |
| P59481 | Lman2l | VIP36-like protein | 39.95 | 2.31 | 39881 | 1 | 870673.62 | 648827.31 | 711227.06 | 599312.31 | 681605.75 | 674717.44 | 809237.75 | 690506.06 | 667313.56 | 597732.5 | 677495.44 | 500271.16 | 691035.06 | 539929.69 | 847492.38 | 2 | 2 | 817646.94 | 2 | 2 | 9 | 4 | 12 | 5 | 12 | 5 | 0.927264218 | 0.946712257 | 0.232840932 | 0.237724437 |
| Q6AZ61 | Lmbrd1 | Probable lysosomal cobalamin transporter | 90.91 | 3.35 | 61018 | 1 | 692589.5 | 1797537.38 | 2 | 2363739.75 | 1217265.75 | 885208.5 | 1792280.75 | 1989791.38 | 2 | 1890250.12 | 514906.31 | 571788.69 | 1033670.25 | 1031524 | 960649.5 | 1759195.5 | 478682.84 | 279620.16 | 2262457.5 | 306722.41 | 8 | 10 | 8 | 10 | 8 | 10 | 0.677392147 | 0.627147268 | 0.837846454 | 0.775700039 |
| Q8C3X8 | Lmf2 | Lipase maturation factor 2 | 48.11 | 2.28 | 79997 | 1 | 2 | 2 | 42523.42 | 2 | 127035.21 | 2 | 2 | 42908.84 | 81217.44 | 2 | 2 | 2 | 132647.81 | 96181.34 | 224018.12 | 2 | 2 | 2 | 2 | 2 | 8 | 7.5 | 7 | 7.5 | 8 | 7.5 | 2.670670587 | 3.648134635 | 5.897456E-05 | 8.055922E-05 |
| P48678 | Lmna | Prelamin-A/C | 2967.92 | 58.5 | 74238 | 45 | 266299218 | 283091618 | 242070498 | 279841890 | 252677442 | 278470594 | 298436578 | 262843362 | 246851122 | 288641698 | 160051522 | 189044690 | 163468402 | 154405938 | 185538146 | 277829698 | 286973794 | 279760898 | 300310402 | 279428194 | 0 | 6 | 0 | 9 | 0 | 9 | 0.643898147 | 0.619896614 | 1.075773252 | 1.035673419 |
| P14733 | Lmnb1 | Lamin-B1 | 2621.11 | 65.14 | 66786 | 42 | 25704828 | 27902932 | 31220452 | 38985594 | 23395612 | 27907342 | 23859294 | 23860248 | 25589872 | 34151574 | 33814874 | 26021242 | 26758590 | 27017892 | 31063174 | 28440624 | 33109632 | 23052902 | 21337696 | 28819272 | 12 | 10 | 8 | 12 | 12 | 12 | 0.982788832 | 1.068756422 | 0.915431416 | 0.995507044 |
| P21619 | Lmnb2 | Lamin-B2 | 796.48 | 22.48 | 67318 | 14 | 858275.12 | 1150409.38 | 1735833.5 | 938599.56 | 1474576.25 | 2103602 | 1427345.5 | 1486824 | 1963065.12 | 1065539.5 | 3010602 | 2010957.38 | 1450553.88 | 895245 | 1076324.25 | 1246732.25 | 595589.44 | 1375979.38 | 1435933.88 | 1546699.62 | 9 | 12 | 12 | 7 | 12 | 12 | 1.371241047 | 1.049377059 | 1.007022233 | 0.770649355 |
| Q8C129 | Lnpep | Leucyl-cystinyl aminopeptidase | 1020.69 | 18.93 | 117304 | 18 | 16275149 | 16070031 | 19183966 | 18364332 | 21289944 | 16752599 | 18835510 | 19976424 | 23534044 | 20569438 | 13005185 | 21053844 | 18760902 | 17266674 | 19200566 | 16327579 | 10063839 | 11465616 | 11469761 | 16396055 | 12 | 4 | 8 | 0 | 12 | 4 | 0.979203994 | 0.895845784 | 0.720776305 | 0.659417668 |
| Q8CGK3 | Lonp1 | Lon protease homolog, mitochondrial | 1570.86 | 33.4 | 105843 | 28 | 44156310 | 49725922 | 35161510 | 38901070 | 39886910 | 42212314 | 41384254 | 37038146 | 43888258 | 44531918 | 47421566 | 55039978 | 60719670 | 60006042 | 58199370 | 44219094 | 39078658 | 48603442 | 51987834 | 52342594 | 1 | 5 | 0 | 5 | 1 | 5 | 1.353915674 | 1.345993992 | 1.136648533 | 1.129998069 |
| Q3MIB4 | Lonp2 | Lon protease homolog 2, peroxisomal | 56.49 | 1.41 | 94393 | 1 | 247568.95 | 290642.47 | 203404.61 | 2 | 157344.47 | 381008.62 | 300769.94 | 297640.72 | 340262.12 | 291193.69 | 2 | 270069.25 | 307690.22 | 288161.59 | 286067.97 | 266288.12 | 2 | 286246.72 | 333922.06 | 227536.2 | 7.5 | 8.5 | 3 | 3 | 7.5 | 8.5 | 1.28146728 | 0.715133679 | 1.239200857 | 0.691546543 |
| Q3TFD2 | Lpcat1 | Lysophosphatidylcholine acyltransferase 1 | 55.14 | 4.49 | 59744 | 1 | 2 | 230844.2 | 166301.97 | 277734.56 | 784673.81 | 383892.91 | 2 | 260707.5 | 499249.53 | 378286.53 | 235316.75 | 169087.66 | 249319.06 | 328080.56 | 2 | 259948.52 | 358721.12 | 287766.31 | 265340.12 | 417691.16 | 12.5 | 7 | 5.5 | 11 | 12.5 | 11 | 0.672674201 | 0.645017552 | 1.08900696 | 1.044233006 |
| Q8BYI6 | Lpcat2 | Lysophosphatidylcholine acyltransferase 2 | 359.16 | 22.43 | 60254 | 8 | 843402.56 | 1191794.75 | 1317191.88 | 1394441.75 | 1462377.62 | 1156857 | 1303445.25 | 881019 | 1013884.5 | 810930.75 | 279867.06 | 389445.62 | 191247.09 | 261039.7 | 325689.62 | 910859.44 | 987808.19 | 895796 | 1039600.38 | 1189402.38 | 0 | 5 | 0 | 12 | 0 | 12 | 0.233087531 | 0.280149216 | 0.809034894 | 0.972383597 |
| Q91V01 | Lpcat3 | Lysophospholipid acyltransferase 5 | 104.58 | 4.52 | 56147 | 2 | 625699.56 | 389162.66 | 886404.31 | 640202.94 | 901033.75 | 776814.12 | 1040807.25 | 1087535.25 | 1330276.75 | 1051003.12 | 1962820 | 3066924 | 4068997.25 | 2985422.5 | 4746060 | 725733.44 | 743691.69 | 1234934.88 | 1207846 | 1247278.12 | 0 | 4 | 0 | 12 | 0 | 12 | 4.888949312 | 3.183661391 | 1.49875942 | 0.975985267 |
| P11152 | Lpl | Lipoprotein lipase | 602.75 | 23.21 | 53109 | 10 | 1234617.5 | 358826.94 | 2388391.25 | 2271716.25 | 966526 | 788914.62 | 1749874.38 | 2935389.5 | 1297370.25 | 1868017.88 | 5054586 | 4731905 | 4413182.5 | 3101498.5 | 3686105.25 | 1149003 | 898067.5 | 1699250.75 | 3285494.25 | 1694468.88 | 0 | 11 | 0 | 11 | 0 | 11 | 2.906793725 | 2.42920486 | 1.208613598 | 1.01003728 |
| Q99N69 | Lpxn | Leupaxin | 561.62 | 25.39 | 43478 | 8 | 10556964 | 10671476 | 14942762 | 16373843 | 13352557 | 11490328 | 12524564 | 14518967 | 16175677 | 13825476 | 11145565 | 10580267 | 14288972 | 11966161 | 14069134 | 10245665 | 9712528 | 14620652 | 13031305 | 13719937 | 11 | 8 | 7 | 8 | 11 | 8 | 0.941613915 | 0.905378101 | 0.93068769 | 0.894872346 |
| P62046 | Lrch1 | Leucine-rich repeat and calponin homology domain-containing protein 1 | 48.74 | 1.83 | 79076 | 1 | 449406.31 | 469554 | 339486.06 | 339637.66 | 440189.88 | 279198.12 | 483311.78 | 410006.12 | 454344.69 | 355623.78 | 2 | 314966.44 | 281951.62 | 2 | 221185.06 | 388665.56 | 2 | 378953.34 | 188063.11 | 425474.28 | 0 | 6 | 2 | 7 | 2 | 7 | 0.401372512 | 0.412667602 | 0.67761172 | 0.696680502 |
| Q921G6 | Lrch4 | Leucine-rich repeat and calponin homology domain-containing protein 4 | 185.76 | 6.47 | 73163 | 3 | 875104.88 | 1658548.75 | 2167042.75 | 1566686.75 | 1400699.75 | 1188341.75 | 1029955.12 | 43055.02 | 1134825.62 | 1337789.5 | 1499030.38 | 1345778.12 | 1505332.75 | 1286205.88 | 1168952.62 | 1240794.62 | 982317.94 | 1381688.88 | 1055508.62 | 1529630.75 | 7 | 6 | 3 | 8 | 7 | 8 | 0.887483854 | 1.437546932 | 0.807234469 | 1.307558924 |
| Q60664 | Lrmp | Lymphoid-restricted membrane protein | 222.7 | 10.95 | 59588 | 4 | 3542387.75 | 3652770.5 | 4513666 | 5143267.5 | 1922471.75 | 3226276 | 4281714.5 | 4072407 | 2746777 | 4612182.5 | 1537139.12 | 1656537.25 | 3254918.75 | 3645840 | 2575940 | 3390828 | 4939502.5 | 4096736 | 3821086.5 | 4481724 | 4 | 11 | 4 | 9 | 4 | 11 | 0.674869225 | 0.668997111 | 1.104146949 | 1.09453964 |
| Q91ZX7 | Lrp1 | Prolow-density lipoprotein receptor-related protein 1 | 1651.3 | 7.59 | 504742 | 31 | 55042450 | 56414042 | 37370282 | 40267122 | 33406290 | 53472694 | 47020950 | 25449728 | 23394752 | 29010002 | 18868120 | 14801657 | 12911179 | 10554529 | 12921956 | 41103598 | 43104802 | 27239996 | 29539502 | 27315244 | 0 | 6 | 0 | 12 | 0 | 12 | 0.31486464 | 0.39281288 | 0.756417983 | 0.943677659 |
| Q7TQH7 | Lrp10 | Low-density lipoprotein receptor-related protein 10 | 88.79 | 2.66 | 76461 | 2 | 643770.94 | 767532.75 | 369381.12 | 454843.12 | 731465 | 1047694.75 | 613504.06 | 428188.19 | 513527.25 | 509320.94 | 1562225.5 | 1720724.62 | 1935361.75 | 1287719.38 | 1798435.38 | 1042345.31 | 682084.62 | 531487.31 | 2 | 673811.25 | 0 | 12 | 0 | 10 | 0 | 12 | 2.798950596 | 2.668328749 | 0.987441008 | 0.941358963 |
| P55302 | Lrpap1 | Alpha-2-macroglobulin receptor-associated protein | 548.04 | 33.06 | 42215 | 12 | 11698658 | 14241222 | 4360264 | 6382005 | 6197257 | 13502597 | 13741930 | 7512176.5 | 8754972 | 7718279 | 12950037 | 9400683 | 10809411 | 8916358 | 10585305 | 16464282 | 22213356 | 16670499 | 7217315 | 15994474 | 9 | 2 | 10 | 5 | 10 | 5 | 1.228137209 | 1.027949264 | 1.832113206 | 1.533476396 |
| Q6PB66 | Lrpprc | Leucine-rich PPR motif-containing protein, mitochondrial | 1667.6 | 27.01 | 156615 | 31 | 17457150 | 16036028 | 20971786 | 18079480 | 21687670 | 12404342 | 17736290 | 17397146 | 18906058 | 17697080 | 7343867 | 16626798 | 14112890 | 5255509 | 14695869 | 20346728 | 10736539 | 12408926 | 12514459 | 16211937 | 1 | 4 | 3 | 8 | 3 | 8 | 0.615872133 | 0.68973498 | 0.766390415 | 0.858305239 |
| Q8K1T1 | Lrrc25 | Leucine-rich repeat-containing protein 25 | 140.17 | 13.8 | 32673 | 3 | 115615.92 | 148002.09 | 176173.91 | 807427.88 | 418809.91 | 169263.53 | 140684.17 | 67775.84 | 152311.83 | 2 | 400673.91 | 538426.19 | 912009.62 | 933329.75 | 414571.59 | 117930.78 | 947031.5 | 80851.88 | 213686.41 | 560878.69 | 5 | 12 | 0 | 6 | 5 | 12 | 1.920140464 | 6.035444369 | 1.152668076 | 3.623101631 |
| Q9CRC8 | Lrrc40 | Leucine-rich repeat-containing protein 40 | 199.63 | 7.81 | 68076 | 4 | 1195944.25 | 955617.81 | 1720020.75 | 1144632.25 | 2027153.25 | 678584.31 | 935918.88 | 1676259.62 | 2157526.25 | 672171.88 | 767033.06 | 1547892.62 | 1153690.5 | 500796.47 | 1074755.62 | 1230695.62 | 688865.25 | 1261948.88 | 1058182 | 650382.62 | 6 | 7 | 11 | 11 | 11 | 11 | 0.716158526 | 0.824148429 | 0.694280656 | 0.798971584 |
| Q505F5 | Lrrc47 | Leucine-rich repeat-containing protein 47 | 67.08 | 3.1 | 63590 | 1 | 786676.25 | 966873.81 | 568232.44 | 430225.47 | 555073.5 | 700614.5 | 833156.88 | 598316.38 | 805457.25 | 610071.94 | 770663.19 | 592294.12 | 968056.25 | 492556.12 | 2 | 494647.75 | 571384.69 | 707045.38 | 2 | 706227.38 | 12 | 10 | 8 | 6 | 12 | 10 | 0.853795622 | 0.795906582 | 0.749696439 | 0.69886553 |
| Q922Q8 | Lrrc59 | Leucine-rich repeat-containing protein 59 | 815.66 | 43 | 34877 | 13 | 47843262 | 59636346 | 57088050 | 45758150 | 66942878 | 36082354 | 59707826 | 61203918 | 74747770 | 48043018 | 23095856 | 48425406 | 40964298 | 19970368 | 36422530 | 64977946 | 27198254 | 47622930 | 46597598 | 55797622 | 2 | 8 | 4 | 8 | 4 | 8 | 0.609078726 | 0.603601075 | 0.873500551 | 0.865644865 |
| Q498T9 | Lrrc8c | Volume-regulated anion channel subunit LRRC8C | 173.53 | 3.86 | 92463 | 2 | 2 | 47823.95 | 138879.39 | 2 | 184531.45 | 2 | 81050.48 | 168383.41 | 278093.28 | 172097.62 | 2 | 146105.67 | 2 | 2 | 2 | 2 | 2 | 2 | 2 | 2 | 8 | 5 | 4 | 2.5 | 8 | 5 | 0.393584059 | 0.208845162 | 2.693684E-05 | 1.429333E-05 |
| Q3UZ39 | Lrrfip1 | Leucine-rich repeat flightless-interacting protein 1 | 819.01 | 25.51 | 79249 | 13 | 19315006 | 22981114 | 17006002 | 22322280 | 18633760 | 20521002 | 20178786 | 18780440 | 17121060 | 18106256 | 10886804 | 12055202 | 9525321 | 11502478 | 9276387 | 23092580 | 23360224 | 19555990 | 18568614 | 20385104 | 0 | 8 | 0 | 6 | 0 | 8 | 0.531090845 | 0.562217008 | 1.046922364 | 1.108280371 |
| Q8BUI3 | LRWD1 | Leucine-rich repeat and WD repeat-containing protein 1 | 49.64 | 1.7 | 71594 | 1 | 2 | 2 | 2 | 2 | 2 | 2 | 2 | 2 | 2 | 2 | 340239.28 | 337358.19 | 463514.47 | 340439.41 | 290818.41 | 2 | 2 | 2 | 2 | 2 | 0 | 12.5 | 0 | 12.5 | 0 | 12.5 | 177236.976 | 177236.976 | 1 | 1 |
| Q3UM18 | Lsg1 | Large subunit GTPase 1 homolog | 80.37 | 3.42 | 73157 | 2 | 2 | 2 | 291915.66 | 132555.47 | 155094.14 | 2 | 293026.81 | 2 | 195055.33 | 264454.69 | 189398.03 | 2 | 440356.53 | 157529.41 | 121403.81 | 185634.94 | 250618.86 | 233698.22 | 274941.38 | 235359.66 | 9 | 5 | 12 | 10 | 12 | 10 | 1.567870878 | 1.207495652 | 2.0364314 | 1.568357507 |
| Q9D0R8 | Lsm12 | Protein LSM12 homolog | 89.82 | 6.67 | 21701 | 1 | [illegible] | [illegible] | [illegible] | [illegible] | [illegible] | [illegible] | [illegible] | [illegible] | [illegible] | [illegible] | [illegible] | [illegible] | [illegible] | [illegible] | [illegible] | [illegible] | [illegible] | [illegible] | [illegible] | [illegible] | [illegible] | [illegible] | [illegible] | [illegible] | [illegible] | [illegible] | [illegible] | [illegible] | [illegible] | [illegible] |
| Q8K2F8 | Lsm14a | Protein LSM14 homolog A | 102.1 | 3.9 | 50546 | 2 | [illegible] | [illegible] | [illegible] | [illegible] | [illegible] | [illegible] | [illegible] | [illegible] | [illegible] | [illegible] | [illegible] | [illegible] | [illegible] | [illegible] | [illegible] | [illegible] | [illegible] | [illegible] | [illegible] | [illegible] | 8.5 | 6.5 | 8.5 | 5.5 | 8.5 | 6.5 | 1.779071438 | 1.945742004 | 0.382768475 | 0.418627764 |

| | | | | | | | | | | | | | | | | | | | | | | | | | | | | | | | | | | | | |
|---|---|---|---|---|---|---|---|---|---|---|---|---|---|---|---|---|---|---|---|---|---|---|---|---|---|---|---|---|---|---|---|---|---|---|---|---|
| O35900 | Lsm2 | U6 snRNA-associated Sm-like protein LSm2 | 129.05 | 28.42 | 10835 | 2 | 1347522.75 | 1028049.81 | 1150266.12 | 1316515.38 | 1113439.38 | 1073503.5 | 1141942.5 | 1027156.19 | 1466969.5 | 1023165.06 | 490101.25 | 603094.69 | 537555.5 | 1153655.75 | 584550.62 | 1810117 | 1706473.25 | 627126 | 959613.94 | 1209629.75 | 3 | 12 | 4 | 11 | 4 | 12 | 0.56566062 | 0.58767007 | 1.059969592 | 1.101212251 |
| P62311 | Lsm3 | U6 snRNA-associated Sm-like protein LSm3 | 71.5 | 19.61 | 11845 | 2 | 1271525.12 | 1211671.12 | 1460377.38 | 685369.44 | 1166270 | 1281448.62 | 878357.19 | 915957.44 | 1392405.5 | 1559572.5 | 4632845 | 2379421 | 2587180.75 | 1660958.12 | 2369353 | 803724.88 | 1493493.88 | 1100757.75 | 1271699.5 | 878678.62 | 0 | 12 | 0 | 9 | 0 | 12 | 2.351899357 | 2.26117169 | 0.957403045 | 0.920469941 |
| Q9QXA5 | Lsm4 | U6 snRNA-associated Sm-like protein LSm4 | 48.89 | 7.3 | 15076 | 1 | 1302819.12 | 1966410.38 | 551054.62 | 927543.06 | 1360628.62 | 514828.69 | 1195887.88 | 1707297 | 2 | 660654.31 | 479938.06 | 1579269.5 | 745638.31 | 2 | 1153014.12 | 1173696.25 | 266056.94 | 568794.88 | 855754.5 | 2 | 7 | 4 | 11.5 | 9.5 | 11.5 | 9.5 | 0.647931674 | 0.970380567 | 0.468908127 | 0.702264379 |
| P62322 | Lsm5 | U6 snRNA-associated Sm-like protein LSm5 | 82.79 | 20.88 | 9937 | 1 | 702172.12 | 416537.5 | 604306.19 | 657274.06 | 594387.69 | 621122.75 | 813132.69 | 545057.38 | 858982.5 | 748382.62 | 251735.98 | 238104.47 | 420210.38 | 533256.38 | 2 | 667376.62 | 983892.69 | 2 | 606108 | 403443.97 | 2 | 12 | 0 | 8 | 2 | 12 | 0.48519854 | 0.402408366 | 0.894491328 | 0.741862895 |
| P62313 | Lsm6 | U6 snRNA-associated Sm-like protein LSm6 | 143.81 | 32.5 | 9128 | 3 | 3950095.5 | 4138081 | 5761501.5 | 4072052 | 3530871.5 | 2032799.88 | 3495343.5 | 3227219.5 | 3332043 | 3153708.5 | 1757737.12 | 2143479.5 | 2654418 | 2660743 | 1955671.12 | 5264429.5 | 2717783.5 | 2589992.5 | 3407829 | 3080764.25 | 0 | 4 | 3 | 12 | 3 | 12 | 0.520778272 | 0.733020464 | 0.79527878 | 1.119393131 |
| Q9CQQ8 | Lsm7 | U6 snRNA-associated Sm-like protein LSm7 | 37.69 | 17.48 | 11636 | 1 | 562216 | 2 | 505640.62 | 268186.22 | 2 | 319582 | 349373.19 | 318479.59 | 214040.83 | 257117.45 | 102817.67 | 233644.88 | 2 | 2 | 175817.11 | 247609.33 | 358076.5 | 343126.84 | 622454.12 | 457852.31 | 8 | 9 | 1 | 5 | 8 | 9 | 0.38343241 | 0.351217673 | 1.518748474 | 1.391148193 |
| Q6ZWM4 | Lsm8 | U6 snRNA-associated Sm-like protein LSm8 | 312.6 | 57.29 | 10403 | 4 | 6349568.5 | 5542050 | 7352303 | 7652241 | 6416824 | 6346976.5 | 5074212 | 5597020 | 5463961 | 7083427.5 | 3680483 | 3276212.25 | 2656489 | 5096984 | 2873963.5 | 5121939 | 8497572 | 6933305 | 5948470 | 5959661.5 | 0 | 10 | 1 | 9 | 1 | 10 | 0.527846152 | 0.594749761 | 0.974423218 | 1.097929715 |
| P19973 | Lsp1 | Lymphocyte-specific protein 1 | 728.35 | 44.85 | 36714 | 11 | 36370706 | 43541574 | 41407422 | 45629378 | 17568022 | 41364274 | 34843542 | 35837870 | 27110236 | 42423594 | 17993328 | 16763562 | 9692979 | 12850372 | 13593377 | 34673566 | 45380594 | 32174402 | 29162176 | 28298390 | 1 | 8 | 0 | 9 | 1 | 9 | 0.384211638 | 0.390427398 | 0.919639026 | 0.934516909 |
| Q99KG5 | Lsr | Lipolysis-stimulated lipoprotein receptor | 43.56 | 1.85 | 66108 | 1 | 7163.25 | 100147.45 | 2 | 2 | 2 | 110459.59 | 54202.57 | 2 | 2 | 2 | 132897.11 | 145660.48 | 285682.34 | 187141.3 | 187936.56 | 2 | 2 | 2 | 2 | 2 | 0 | 7.5 | 0 | 7.5 | 0 | 7.5 | 8.752764388 | 5.704307317 | 9.318214E-05 | 6.072819E-05 |
| Q8BLN5 | Lss | Lanosterol synthase | 484.66 | 12.82 | 83141 | 8 | 1722283.62 | 1928754.12 | 1291033.25 | 1290513.75 | 1740296 | 1914380 | 2430977.25 | 1279173.75 | 1924085.5 | 1368363.75 | 2890036 | 3658391.5 | 3882200.75 | 2859282.5 | 3776549.5 | 1096789.38 | 1211127.75 | 1349989.38 | 2098516.25 | 1961507.75 | 0 | 12 | 0 | 9 | 0 | 12 | 2.140563845 | 1.913928233 | 0.968022822 | 0.865531861 |
| P24527 | Lta4h | Leukotriene A-4 hydrolase | 510.55 | 18.99 | 69051 | 9 | 12878679 | 13382945 | 10651675 | 11690592 | 9698045 | 13348327 | 13736008 | 14264084 | 13678302 | 10537676 | 9265850 | 10142074 | 6974557 | 5406179.5 | 6807797.5 | 14645636 | 13526471 | 10522030 | 11694178 | 12217612 | 1 | 8 | 0 | 9 | 1 | 9 | 0.662009886 | 0.588680134 | 1.073822437 | 0.954876882 |
| Q6A009 | Ltn1 | E3 ubiquitin-protein ligase listerin | 89.46 | 1.3 | 198921 | 2 | 2 | 563050 | 186832.3 | 2 | 639409.38 | 2 | 232614.38 | 218471.84 | 219907.34 | 2 | 2 | 190923.84 | 2 | 389792.56 | 186327.84 | 252880.83 | 598220.75 | 2 | 322174.84 | 2 | 10 | 12 | 11 | 8 | 11 | 12 | 0.552113025 | 1.143146094 | 0.844514553 | 1.748561381 |
| Q6NSQ7 | Ltv1 | Protein LTV1 homolog | 223.57 | 10.21 | 54023 | 4 | 1500906.75 | 932489.56 | 869835.06 | 1195999.38 | 1141448 | 1711098.5 | 1609991.62 | 1295601.5 | 1648111.25 | 1245633.25 | 1119932.25 | 1232254.62 | 769113.75 | 1618500.38 | 564034 | 2522826.5 | 1985709 | 1887747 | 538190.56 | 1526645.38 | 11 | 5 | 3 | 8 | 11 | 8 | 0.940283118 | 0.706195341 | 1.500017784 | 1.126581507 |
| Q9CYI4 | Luc7l | Putative RNA-binding protein Luc7-like 1 | 350.08 | 19.41 | 43934 | 7 | 2 | 324675.28 | 2 | 2 | 2 | 2 | 2 | 40474.13 | 2 | 2 | 245011.45 | 2 | 2 | 2 | 2 | 2 | 2 | 2 | 2 | 2 | 12 | 10 | 12 | 10 | 12 | 10 | 0.754641415 | 6.05253355 | 3.079925E-05 | 0.000247023 |
| Q7TNC4 | Luc7l2 | Putative RNA-binding protein Luc7-like 2 | 578.43 | 25.77 | 46583 | 10 | 5125886.5 | 5753026.5 | 5992173.5 | 6881297 | 7304687 | 5237472.5 | 5134608 | 6575725 | 5708022 | 4164100.5 | 3343101.5 | 3488291.5 | 3801143 | 3651439.25 | 3840224 | 4154419.25 | 3974542.5 | 3526939 | 3960842.25 | 4335961 | 0 | 0 | 0 | 1 | 0 | 1 | 0.583577232 | 0.675773598 | 0.642452868 | 0.743950692 |
| Q5SUF2 | Luc7l3 | Luc7-like protein 3 | 557.42 | 24.54 | 51450 | 8 | 3663755 | 6100642 | 9637632 | 6435240 | 7909449 | 3542740 | 7145931 | 6944723.5 | 7271001.5 | 5342010.5 | 1751229.25 | 2869263.75 | 3606497.5 | 2521425.75 | 2488817.25 | 6718200 | 5582930.5 | 5227468 | 5736927 | 5857631.5 | 0 | 7 | 1 | 9 | 1 | 9 | 0.39225247 | 0.437646485 | 0.862992277 | 0.962863374 |
| P05533 | Ly6a | Lymphocyte antigen 6A-2/6E-1 | 44.56 | 15.67 | 14377 | 1 | 165062.11 | 2 | 274153.53 | 338741.47 | 373601.19 | 2 | 266379.06 | 250510.67 | 2 | 285940.34 | 239072.95 | 2 | 444104.41 | 441597.34 | 142153.39 | 2 | 2 | 2 | 2 | 2 | 11.5 | 2.5 | 10 | 5 | 11.5 | 5 | 1.100185626 | 1.578072154 | 8.68387E-06 | 1.245587E-05 |
| O88188 | Ly86 | Lymphocyte antigen 86 | 202.81 | 23.46 | 17811 | 4 | 4048141.5 | 1878042 | 2938628.5 | 1806452.75 | 1220633.25 | 4715146 | 4238420.5 | 894840.75 | 3914739.75 | 4896422 | 3272552.5 | 661350.06 | 2399597.75 | 3463140 | 1062563.38 | 2862801 | 7291829 | 4068717.25 | 5396881.5 | 3817374.5 | 11 | 3 | 4 | 11 | 11 | 11 | 0.913159841 | 0.581964336 | 1.970888352 | 1.256063484 |
| Q08288 | Lyar | Cell growth-regulating nucleolar protein | 352.27 | 13.66 | 43736 | 5 | 559331.75 | 829485.62 | 559237.25 | 1021670.44 | 481315.31 | 2106969.25 | 705999.25 | 522487.25 | 652801.69 | 799514.12 | 1337708.75 | 992289.5 | 1681047.75 | 1757438.75 | 734412.06 | 891386.94 | 1932032 | 1632002 | 615684.62 | 947255.5 | 3 | 4 | 6 | 8 | 6 | 8 | 1.884329394 | 1.358230385 | 1.743926589 | 1.257027614 |
| P25911 | Lyn | Tyrosine-protein kinase Lyn | 807.41 | 34.38 | 58812 | 13 | 10438770 | 10859124 | 11431478 | 13313429 | 12213364 | 11442752 | 11664464 | 8860715 | 10340933 | 11165372 | 10669762 | 10926994 | 13504778 | 13330106 | 12859995 | 10520618 | 11661304 | 8805261 | 10005491 | 10716443 | 8 | 5 | 6 | 9 | 8 | 9 | 1.052105558 | 1.146190008 | 0.887616221 | 0.966991225 |
| P97823 | Lypla1 | Acyl-protein thioesterase 1 | 182.94 | 20.87 | 24688 | 3 | 4441639.5 | 5078306.5 | 6104217 | 7928864 | 6317696.5 | 4594427 | 4740965 | 7755241 | 6751038 | 7513006 | 4790592 | 4403272 | 4868356.5 | 4684057.5 | 4769635 | 4893936.5 | 4775366 | 7486665 | 7153792 | 6502509.5 | 4 | 11 | 7 | 12 | 7 | 12 | 0.787256224 | 0.749996978 | 1.031520679 | 0.98270089 |
| Q9QYL8 | Lypla2 | Acyl-protein thioesterase 2 | 45.21 | 10.82 | 24807 | 1 | 2 | 2 | 2 | 346380.12 | 485172.41 | 264143.22 | 278571.94 | 332939.78 | 227982.55 | 2 | 219801.78 | 134336.8 | 160087.8 | 2 | 2 | 188350.47 | 273347.94 | 2 | 149083.38 | 655850.69 | 12 | 9.5 | 4 | 10.5 | 12 | 10.5 | 0.618393488 | 0.46594054 | 1.523205444 | 1.147688617 |
| P08905 | Lyz2 | Lysozyme C-2 | 264.15 | 31.76 | 16689 | 3 | 20470956 | 16608903 | 6974619 | 9836258 | 8465318 | 14404666 | 13436068 | 6093932 | 5431677 | 7088721 | 7855338.5 | 6276877 | 4867928 | 3033242 | 3661375.75 | 22494642 | 31207344 | 11635266 | 13177206 | 13070076 | 1 | 6 | 5 | 6 | 5 | 6 | 0.412065222 | 0.553110017 | 1.468735241 | 1.971465027 |
| P24668 | M6pr | Cation-dependent mannose-6-phosphate receptor | 584.73 | 35.25 | 31172 | 7 | 38738766 | 30927802 | 37242278 | 41585246 | 34390102 | 38322018 | 37298138 | 41678822 | 33556674 | 46593146 | 40585546 | 38222122 | 46095826 | 43852214 | 48336782 | 40434602 | 46875978 | 44470134 | 48184390 | 48458382 | 3 | 1 | 7 | 3 | 7 | 3 | 1.187048947 | 1.099487524 | 1.249006166 | 1.156874533 |
| Q9QXZ0 | Macf1 | Microtubule-actin cross-linking factor 1 | 371.23 | 1.28 | 831878 | 7 | 1229951.12 | 1429466.25 | 1710885.88 | 1218920 | 2114938.75 | 1192169.25 | 1601373.5 | 1289931.75 | 1732872.25 | 1213097 | 332722.19 | 1270185.25 | 763601.06 | 670582.5 | 544943.06 | 1774378.62 | 913640.12 | 329098.94 | 729521.31 | 1288483.38 | 2 | 6 | 2 | 7 | 2 | 7 | 0.464947915 | 0.509575748 | 0.653558735 | 0.7162903 |
| Q9WTX8 | Mad1l1 | Mitotic spindle assembly checkpoint protein MAD1 | 50.39 | 1.53 | 83541 | 1 | 2 | 32015224 | 2 | 2 | 2 | 2 | 2 | 2 | 2 | 2 | 419607.25 | 2 | 2 | 350870.41 | 333239.88 | 2 | 2 | 2 | 2 | 485402.28 | 9 | 12 | 5 | 10 | 9 | 12 | 0.034474888 | 110372.154 | 0.015161854 | 48541.028 |
| P61327 | Magoh | Protein mago nashi homolog | 272.01 | 45.89 | 17164 | 6 | 1589067.62 | 2127166.5 | 2267745 | 1887214.38 | 4770497.5 | 1489649.5 | 1855024.25 | 2601051 | 1495881.75 | 1922296.75 | 2001320.88 | 1759782.5 | 1151663.12 | 2281696 | 1411576.75 | 2578357.5 | 3155982.5 | 2583534.75 | 2331362 | 1728661.62 | 7 | 8 | 10 | 6 | 10 | 8 | 0.680766462 | 0.919065375 | 0.979133122 | 1.3218738 |
| O35777 | Magt1 | Magnesium transporter protein 1 | 45.34 | 2.69 | 37993 | 1 | 1050560.25 | 1469569.25 | 1625542.75 | 1296910.5 | 1202125.75 | 1125679.62 | 1058852.25 | 1409673 | 1126956.5 | 1801102.25 | 2155995.75 | 2375173.25 | 2 | 3272225.25 | 2684306.75 | 1062893.5 | 1370715.38 | 1622895.25 | 1783739.25 | 1382518.25 | 5 | 9 | 5 | 10 | 5 | 10 | 1.578354114 | 1.607985143 | 1.086994505 | 1.107401057 |
| Q6AY04 | Maip1 | m-AAA protease-interacting protein 1, mitochondrial | 127.28 | 8.59 | 33090 | 2 | 1334924 | 734294 | 469748.62 | 1273225.75 | 971925.62 | 1215032.38 | 1267970.5 | 538976.12 | 1202621.5 | 764693.44 | 1811880.5 | 1495442.75 | 2207069.25 | 1790084 | 1752787.5 | 1043336.88 | 1319363.5 | 1610985 | 1590970.25 | 1301967.5 | 0 | 4 | 0 | 3 | 0 | 4 | 1.89319411 | 1.815339827 | 1.435295522 | 1.376271515 |
| Q8BGS0 | Mak16 | Protein MAK16 homolog | 223.55 | 13.18 | 35185 | 4 | 1424731.88 | 1936496.88 | 1396251.12 | 881491.81 | 1112592 | 1063426.38 | 1079805.5 | 598091.81 | 857213.81 | 644972.12 | 2982412.5 | 2330930.5 | 1487874.38 | 1645897.38 | 1720151.38 | 777187.94 | 489765.47 | 972741.38 | 700232.06 | 488268.94 | 3 | 1 | 0 | 7 | 3 | 7 | 1.505912794 | 2.395956897 | 0.50776323 | 0.807868038 |
| P27046 | Man2a1 | Alpha-mannosidase 2 | 257.01 | 4.78 | 131631 | 5 | 926179.12 | 989275.38 | 391163.22 | 747260.12 | 848456.81 | 1009570.12 | 369294 | 529731.12 | 1395621.75 | 348776.81 | 1054078.75 | 1628515.25 | 1965493.5 | 2551499 | 2764690.5 | 700605.5 | 467985.38 | 861396.31 | 605676.75 | 1073040.25 | 0 | 11 | 1 | 10 | 1 | 11 | 2.553414275 | 2.727701591 | 0.950380868 | 1.015250612 |
| O09159 | Man2b1 | Lysosomal alpha-mannosidase | 675.96 | 14.91 | 114648 | 11 | 9953328 | 10759316 | 17569328 | 14181924 | 13296807 | 9543019 | 10855899 | 5991148.5 | 10765459 | 7942538.5 | 1263374.12 | 891009.44 | 1647786.75 | 2407018 | 1530656.75 | 9685209 | 7562207.5 | 6201755 | 9077272 | 7346064 | 0 | 0 | 0 | 8 | 0 | 8 | 0.117697116 | 0.171622557 | 0.60632727 | 0.884129028 |
| O54782 | Man2b2 | Epididymis-specific alpha-mannosidase | 135.64 | 2.36 | 115610 | 2 | 2632042 | 2228309.75 | 1619328.38 | 2151122.25 | 1474109.62 | 2010066 | 1584338.88 | 784067.56 | 1058784.75 | 1115840.5 | 724718.75 | 794864.62 | 969962.88 | 917494.44 | 800072.12 | 2486890.25 | 3483886.75 | 2040584.75 | 1793583.62 | 1624799.12 | 0 | 10 | 4 | 2 | 4 | 10 | 0.41634334 | 0.642003677 | 1.131107771 | 1.744174287 |
| P21139 | Man2c1 | Alpha-mannosidase 2C1 | 172.26 | 2.88 | 115971 | 2 | 1209001.38 | 1142318.5 | 1326973.25 | 1332167.25 | 1157969.5 | 931813.38 | 726692.69 | 1079227.88 | 1177299 | 1081630.62 | 554613.88 | 695828.88 | 921475.62 | 672888.56 | 761186.38 | 1278839.75 | 465483.03 | 1092417 | 1488780.5 | 1074520.12 | 0 | 8 | 2 | 9 | 2 | 9 | 0.584588524 | 0.721680231 | 0.875431918 | 1.080729235 |
| Q8K2I4 | Manba | Beta-mannosidase | 360.3 | 8.42 | 100831 | 6 | 2916362 | 4043994.5 | 6909605 | 5968031.5 | 6008320 | 1980069.62 | 2449130.75 | 2539066.25 | 2351094 | 3337013.25 | 4470871 | 1923647.25 | 1019560.69 | 2221145 | 876807.5 | 2381421.25 | 918927.25 | 2117669.75 | 1444611.5 | 1333584.25 | 2 | 0 | 6 | 3 | 6 | 3 | 0.406712998 | 0.830572133 | 0.31711347 | 0.647595756 |
| P0C5H9 | Manf | Mesencephalic astrocyte-derived neurotrophic factor | 373.89 | 33.52 | 20388 | 7 | 8962832 | 9601597 | 8523604 | 8483622 | 8697764 | 9809799 | 10501872 | 8796728 | 7521761 | 10415698 | 3883957 | 4898595.5 | 4732790.5 | 4397946.5 | 5379945.5 | 9384418 | 4852431 | 4609467 | 7919918 | 5716704 | 0 | 4 | 0 | 3 | 0 | 4 | 0.526169883 | 0.495117657 | 0.733755688 | 0.690452664 |
| Q8C052 | Map1s | Microtubule-associated protein 1S | 247.59 | 5.65 | 102939 | 5 | 3933269 | 4453431 | 4633019 | 3066923 | 3972475.5 | 3122835.5 | 4316991 | 3927701 | 3487806.75 | 3969207.25 | 2794642.75 | 3797296.5 | 2532643.25 | 2105480 | 2158375.25 | 4179727.5 | 2481982.5 | 4303846 | 3894264 | 4249454.5 | 1 | 10 | 2 | 11 | 2 | 11 | 0.667448992 | 0.711222515 | 0.952647817 | 1.015125627 |
| P31938 | Map2k1 | Dual specificity mitogen-activated protein kinase kinase 1 | 777.88 | 36.64 | 43474 | 14 | 26277690 | 26832960 | 23659244 | 22292620 | 27986626 | 20050372 | 25471042 | 20644578 | 25291824 | 18917658 | 13263176 | 23341218 | 18462452 | 15374852 | 19079240 | 30169198 | 19944066 | 17414920 | 18115524 | 22321942 | 1 | 6 | 4 | 9 | 4 | 9 | 0.704616639 | 0.811058243 | 0.849794418 | 0.978167034 |
| P36506 | Map2k2 | Dual specificity mitogen-activated protein kinase kinase 2 | 310.53 | 18.25 | 44282 | 6 | 3131385.5 | 2672968.25 | 3061244.5 | 3131463 | 3480479.5 | 2096821.12 | 3245312 | 2605355 | 3480977.5 | 2338011 | 784853.12 | 2150770 | 631884.62 | 1256460.88 | 2665592.75 | 4301720.5 | 1914802.38 | 2353139.5 | 1892920.25 | 2587122 | 0 | 5 | 4 | 9 | 4 | 9 | 0.483898669 | 0.544043445 | 0.843138121 | 0.947933519 |
| O09110 | Map2k3 | Dual specificity mitogen-activated protein kinase kinase 3 | 103.15 | 9.22 | 39296 | 2 | 411683.09 | 535137.5 | 609577.12 | 303522 | 753193.12 | 398573.03 | 805872.62 | 650013.88 | 1021020.75 | 226099.3 | 292495.06 | 312937.78 | 173232.17 | 68698.38 | 195380.8 | 426883.84 | 121114.55 | 230076.11 | 331314.81 | 177788.94 | 1 | 3 | 2 | 4 | 2 | 4 | 0.399042926 | 0.336197787 | 0.49258426 | 0.415007327 |
| Q9ESL4 | Map3k20 | Mitogen-activated protein kinase kinase kinase 20 | 223.11 | 5.11 | 91720 | 4 | 1023696.75 | 933503.19 | 2086999 | 1406998.5 | 2110515.5 | 677438.69 | 1208474.75 | 1903018.25 | 2321404.5 | 982177.88 | 463697.25 | 332279.19 | 615450.69 | 559357.56 | 644191.19 | 1119904.38 | 405452.62 | 893746.38 | 593496.69 | 778828.62 | 0 | 2 | 0 | 4 | 0 | 4 | 0.345817925 | 0.368695198 | 0.501398125 | 0.534567666 |
| P27546 | Map4 | Microtubule-associated protein 4 | 708.59 | 16.44 | 117429 | 14 | 24224730 | 23051850 | 20288632 | 26458848 | 19159012 | 27661254 | 22494268 | 22347686 | 24595780 | 20352644 | 14372079 | 16436251 | 16404054 | 12931126 | 12013713 | 21399442 | 24214434 | 26685362 | 30654050 | 31587690 | 0 | 5 | 0 | 7 | 0 | 7 | 0.637526635 | 0.614356921 | 1.1887023 | 1.145501137 |
| P63085 | Mapk1 | Mitogen-activated protein kinase 1 | 341.17 | 20.95 | 41276 | 7 | 3126496.75 | 4000077.5 | 5801612 | 6230949 | 5150668 | 3035029.25 | 2917486.5 | 6023009 | 5394224.5 | 5342078 | 5061589 | 6047416.5 | 7226628.5 | 7640202.5 | 6819541 | 3726414 | 3012426.75 | 6624569.5 | 5784490 | 5913428.5 | 4 | 12 | 3 | 9 | 4 | 12 | 1.349059767 | 1.443977939 | 1.030914504 | 1.10344837 |
| P47811 | Mapk14 | Mitogen-activated protein kinase 14 | 319.61 | 20.28 | 41287 | 6 | 7234879.5 | 4803639 | 6947161 | 9730536 | 9242203 | 5154368 | 6525478 | 7376451 | 4228930.5 | 3840694.75 | 3118528.75 | 3276744 | 3732317.5 | 3999716.75 | 2870739.5 | 6544243 | 5396068 | 5669191 | 4723058.5 | 6532907.5 | 0 | 4 | 1 | 9 | 1 | 9 | 0.447807026 | 0.626634787 | 0.760449701 | 1.064128539 |
| P21708 | Mapk3 | Mitogen-activated protein kinase 3 | 394.97 | 25.26 | 43081 | 9 | 22680142 | 20679622 | 20476836 | 26614924 | 24110434 | 21011560 | 18520278 | 25640478 | 26307282 | 25922306 | 9022878 | 13135374 | 16628484 | 12307230 | 14553613 | 19800010 | 21250588 | 26608302 | 24474420 | 21779054 | 0 | 12 | 0 | 12 | 0 | 12 | 0.573031224 | 0.559169628 | 0.994329846 | 0.970277058 |
| P49186 | Mapk9 | Mitogen-activated protein kinase 9 | 97.37 | 3.07 | 48017 | 1 | 333159.75 | 297943.88 | 366283.31 | 254926.92 | 2 | 259918.69 | 335130.12 | 366384.72 | 397279.12 | 249448.47 | 2 | 250541.62 | 215967.8 | 279447.09 | 278627.47 | 401749.5 | 2 | 323864.41 | 132858.66 | 282049.59 | 6.5 | 11.5 | 5 | 9 | 6.5 | 11.5 | 0.818153002 | 0.637116497 | 0.910732026 | 0.709210132 |
| P49138 | Mapkapk2 | MAP kinase-activated protein kinase 2 | 82.25 | 5.18 | 44050 | 2 | 2074698.5 | 1636873 | 2728220 | 3192011 | 1851495.88 | 2273296.25 | 1593919.5 | 1945089.62 | 2279999 | 2295940.5 | 1574436.75 | 1317869.12 | 1067353.75 | 1408248.5 | 1333513.38 | 1950090.12 | 1822964.5 | 2017713.38 | 1613537 | 2055966.25 | 0 | 7 | 0 | 8 | 0 | 8 | 0.583579846 | 0.645096605 | 0.823828741 | 0.910670798 |
| Q61166 | Mapre1 | Microtubule-associated protein RP/EB family member 1 | 466.15 | 34.33 | 30016 | 7 | 25363538 | 22543502 | 24576258 | 22981514 | 21585326 | 24318858 | 22723282 | 25458536 | 20997186 | 22767938 | 11034099 | 13912940 | 12429939 | 12900727 | 9075289 | 30847968 | 28192894 | 24124650 | 23718524 | 23780502 | 0 | 6 | 0 | 6 | 0 | 6 | 0.507073251 | 0.510494006 | 1.11631255 | 1.12384328 |
| Q3B8Q0 | Mapre2 | Microtubule-associated protein RP/EB family member 2 | 81.81 | 3.68 | 36988 | 1 | 152967.75 | 79864.3 | 188613.55 | 2 | 371195.59 | 2 | 226702.67 | 193319.78 | 422616.78 | 74640.17 | 2 | 181927.2 | 2 | 2 | 2 | 137742.95 | 2 | 2 | 2 | 2 | 5 | 4 | 4 | 4 | 5 | 4 | 0.229529759 | 0.198341752 | 0.173786833 | 0.150173055 |
| Q922Q1 | Marc2 | Mitochondrial amidoxime reducing component 2 | 327.36 | 18.64 | 38194 | 6 | 8562592 | 8565058 | 8431714 | 7960052.5 | 8969418 | 9125462 | 8918437 | 6576704.5 | 8178594 | 9961843 | 10239514 | 9550035 | 13196220 | 10145089 | 13102534 | 7166568 | 6507231.5 | 7959597 | 8808870 | 8663130 | 0 | 8 | 1 | 6 | 1 | 8 | 1.323486339 | 1.315061358 | 0.920368774 | 0.914509938 |
| Q3KNM2 | March5 | E3 ubiquitin-protein ligase MARCH5 | 207.26 | 17.99 | 31232 | 4 | 1078629.88 | 1003967.44 | 970215 | 593966.62 | 1146369.88 | 1078736.12 | 838399.5 | 714066.56 | 1316340.12 | 1289057.12 | 582306.69 | 884668.62 | 1300189.75 | 1467569.5 | 1452474.38 | 741624.31 | 2 | 507526.31 | 335326.78 | 644340.19 | 9 | 2 | 9 | 1 | 9 | 2 | 1.186528763 | 1.086050027 | 0.465001124 | 0.425623465 |
| P26645 | Marcks | Myristoylated alanine-rich C-kinase substrate | 344.4 | 34.63 | 29661 | 7 | 12340544 | 12049810 | 12312248 | 14965269 | 12410288 | 13128232 | 10726036 | 12224280 | 11899050 | 15501013 | 4689887 | 5571308 | 6577917 | 7146906.5 | 6715054 | 12027066 | 11280507 | 12942946 | 14005588 | 15665061 | 0 | 12 | 0 | 10 | 0 | 12 | 0.479119141 | 0.483644365 | 1.028761891 | 1.038478425 |
| P28667 | Marcksl1 | MARCKS-related protein | 235.32 | 34.5 | 20165 | 4 | 12677612 | 12099748 | 8857716 | 10766777 | 7996091 | 13870767 | 11690253 | 8998612 | 8261307.5 | 10189743 | 6593127 | 7158455 | 6134539 | 7129016 | 5710903.5 | 11752624 | 13398182 | 10650684 | 9622999 | 10731034 | 0 | 11 | 0 | 9 | 0 | 11 | 0.624567264 | 0.617348032 | 1.071712337 | 1.059324656 |
| Q68FL6 | Mars | Methionine--tRNA ligase, cytoplasmic | 1008.35 | 24.83 | 101431 | 20 | 19082300 | 21104194 | 21305934 | 22471806 | 21906924 | 19830814 | 20959400 | 19159894 | 20666870 | 20406876 | 17946640 | 24897210 | 22859640 | 17894662 | 25523470 | 22518204 | 22456006 | 23029682 | 23750310 | 24122402 | 10 | 1 | 10 | 0 | 10 | 1 | 1.030702073 | 1.080156989 | 1.094505871 | 1.147022207 |
| Q3THS6 | Mat2a | S-adenosylmethionine synthase isoform type-2 | 360.08 | 15.7 | 43689 | 7 | 521236.09 | 887538.25 | 1400192.38 | 1745892.88 | 1338138.62 | 620640.88 | 695398.56 | 981596.5 | 1285041.38 | 1702137.38 | 1294007.62 | 1688598.5 | 1427671.12 | 1339756 | 1786183.62 | 536552.56 | 471967.84 | 1220357.75 | 1028842.88 | 1082396.5 | 7 | 7 | 4 | 9 | 7 | 9 | 1.278842548 | 1.426013453 | 0.736487161 | 0.821243085 |
| Q5U2R0 | Mat2b | Methionine adenosyltransferase 2 subunit beta | 324.91 | 20.96 | 37375 | 5 | 10457013 | 8519630 | 10139007 | 11731535 | 11906564 | 10571152 | 10092884 | 9016191 | 8631058 | 5970990 | 7475934.5 | 8434391 | 10527513 | 8094077 | 8829116 | 9327360 | 10393228 | 11055322 | 12381740 | 12128556 | 4 | 9 | 9 | 3 | 9 | 9 | 0.821951659 | 0.979196112 | 1.048005252 | 1.24849516 |
| Q8K310 | Matr3 | Matrin-3 | 675.36 | 17.85 | 94630 | 12 | 8762599 | 10119635 | 8771387 | 10458982 | 9944787 | 9635422 | 10605158 | 10069346 | 9002812 | 8925623 | 12143424 | 11012082 | 10020478 | 10596551 | 11541998 | 8924048 | 8164483 | 10896987 | 10055058 | 11798447 | 2 | 10 | 3 | 12 | 3 | 12 | 1.151009928 | 1.1466918 | 1.037073029 | 1.033182346 |
| Q8VCF0 | Mavs | Mitochondrial antiviral-signaling protein | 388.64 | 13.52 | 53399 | 5 | 3344885.5 | 3782574.5 | 4113240.25 | 4912994 | 3441134 | 3941460.5 | 3713501.5 | 4357599 | 3531399.75 | 5914565 | 3464698.25 | 4496007 | 4055855.25 | 3945890.25 | 4563507.5 | 3917051.25 | 4524501.5 | 4340922.5 | 6603556.5 | 4674166 | 9 | 5 | 11 | 7 | 11 | 7 | 1.047519171 | 0.956540933 | 1.227885105 | 1.12124188 |
| Q9Z2E1 | Mbd2 | Methyl-CpG-binding domain protein 2 | 58.26 | 3.38 | 43501 | 1 | 215366.77 | 2 | 2 | 165420.25 | 310895.44 | 300564.03 | 365184.44 | 2 | 323650.44 | 355029.88 | 2 | 2 | 2 | 683928.75 | 252171.75 | 2 | 2 | 176223.31 | 2 | 310164.72 | 12 | 10 | 7.5 | 4.5 | 12 | 10 | 1.353368259 | 0.696284634 | 0.703200161 | 0.361784358 |
| Q9Z2D8 | Mbd3 | Methyl-CpG-binding domain protein 3 | 266.58 | 12.63 | 32168 | 4 | 1940188.25 | 2232216.75 | 2022750.75 | 1926635.38 | 2051323 | 1306859.12 | 2600331 | 1344306.88 | 1752982.5 | 1677560.25 | 762122.44 | 1052624.5 | 1028047.44 | 470699.38 | 653521.75 | 2272112.5 | 1077435.38 | 1782861.25 | 1031583.75 | 1096759.38 | 0 | 5 | 0 | 8 | 0 | 8 | 0.389950949 | 0.456922063 | 0.713719729 | 0.836295671 |
| Q9JKP5 | Mbnl1 | Muscleblind-like protein 1 | 265.29 | 19.35 | 36976 | 5 | 2135167.25 | 2357869.25 | 1403321.38 | 1340723.88 | 839660.31 | 1367623.25 | 545423.81 | 1420712.88 | 1745744.38 | 2062133.75 | 2292476 | 1941620.25 | 1985153.25 | 2298949.75 | 2024614.75 | 1876515.25 | 811614.94 | 1634043.75 | 1457768.5 | 1269794.75 | 8 | 10 | 3 | 12 | 8 | 12 | 1.305330034 | 1.476245911 | 0.872844165 | 0.987131681 |
| F2Z3T4 | Mbnl2 | Muscleblind-like protein 2 | 162.24 | 8.85 | 40156 | 4 | 2 | 2 | 2 | 2 | 2 | 2 | 2 | 2 | 2 | 2 | 1115091.75 | 675604 | 506332.88 | 410095.5 | 2 | 2 | 163488.03 | 2 | 2 | 143535.88 | 2.5 | 7.5 | 2.5 | 7.5 | 2.5 | 7.5 | 270712.613 | 270712.613 | 30702.991 | 30702.991 |
| Q8CHK3 | Mboat7 | Lysophospholipid acyltransferase 7 | 113 | 7.19 | 53436 | 2 | 1499618.25 | 1305189.62 | 1296571.5 | 1496388.88 | 2348207.5 | 1421986 | 1611132.5 | 1659606.62 | 2110623.25 | 1271817.5 | 2203171.5 | 1837423.75 | 1825665 | 1885314 | 2213072.5 | 2095104 | 2225420.5 | 1948577.62 | 2268523.5 | 1848976.5 | 5 | 5 | 3 | 3 | 5 | 5 | 1.254049479 | 1.233986634 | 1.307152507 | 1.286240096 |
| Q8R3F5 | Mcat | Malonyl-CoA-acyl carrier protein transacylase, mitochondrial | 40.83 | 3.94 | 41928 | 1 | 414077.91 | 2 | 691886.5 | 2 | 2 | 419129.66 | 2 | 2 | 2 | 2 | 2 | 423679.59 | 2 | 2 | 2 | 477968.88 | 2 | 442659.03 | 2 | 993711.25 | 10 | 9 | 12 | 6 | 12 | 9 | 0.383091253 | 1.010855455 | 1.730917159 | 4.567337519 |
| Q3ULD5 | Mccc2 | Methylcrotonoyl-CoA carboxylase beta chain, mitochondrial | 211.47 | 7.64 | 61379 | 3 | 792911.62 | 217213.89 | 251025.22 | 297524.62 | 350888.03 | 323363.47 | 311960.91 | 382635 | 589695 | 679120.06 | 467276.53 | 1045751 | 1235960.75 | 2 | 527876.62 | 774427 | 2 | 1512659.75 | 891670.44 | 913891.44 | 7 | 6 | 9 | 5 | 9 | 6 | 1.716029399 | 1.432964635 | 2.143238959 | 1.789704554 |
| P97287 | Mcl1 | Induced myeloid leukemia cell differentiation protein Mcl-1 homolog | 85.46 | 10.27 | 35217 | 2 | 188849.47 | 629096.38 | 231262.31 | 270913.34 | 409582.25 | 2 | 290190.94 | 219409.2 | 218303.42 | 756750.25 | 1041959.38 | 965125.81 | 1235917 | 1231122.88 | 1327856.12 | 2 | 2 | 896149.62 | 983151.38 | 1059280.38 | 0 | 10 | 0 | 9 | 0 | 10 | 3.354320756 | 3.90796382 | 1.698895189 | 1.979304132 |
| P97310 | Mcm2 | DNA replication licensing factor MCM2 | 1236.2 | 27.77 | 102078 | 19 | 11357868 | 14088541 | 22577042 | 16883616 | 15756351 | 11174961 | 11917558 | 18378818 | 17896304 | 15585136 | 20371984 | 21390254 | 21857538 | 20399974 | 18696358 | 11051785 | 9107887 | 12000589 | 10569614 | 11698295 | 5 | 2 | 0 | 3 | 5 | 3 | 1.273391465 | 1.370410972 | 0.674756554 | 0.726166157 |
| P25206 | Mcm3 | DNA replication licensing factor MCM3 | 1799.12 | 39.29 | 91546 | 27 | 54164006 | 60851942 | 66252262 | 60250966 | 49907858 | 54236674 | 52595670 | 57907114 | 51514802 | 53644718 | 69823474 | 64819026 | 64087230 | 63686546 | 65313694 | 46892130 | 50817306 | 45727882 | 49117522 | 41931402 | 4 | 1 | 0 | 0 | 4 | 1 | 1.124569555 | 1.214269029 | 0.804613899 | 0.868792627 |
| P49717 | Mcm4 | DNA replication licensing factor MCM4 | 823.1 | 23.43 | 96736 | 15 | 7513412.5 | 7577932 | 13372276 | 8609271 | 9802424 | 8576464 | 8042229 | 9391006 | 8865856 | 11103449 | 13865468 | 12985963 | 10329315 | 7463710 | 11657063 | 7404898 | 5140945 | 5711311.5 | 4927006.5 | 5384682 | 8 | 0 | 6 | 0 | 8 | 0 | 1.201090988 | 1.224504972 | 0.609464549 | 0.621345408 |
| P49718 | Mcm5 | DNA replication licensing factor MCM5 | 1423.74 | 42.97 | 82343 | 27 | 48241690 | 51845922 | 57230942 | 56657434 | 36504370 | 49953598 | 43545294 | 46156450 | 47658934 | 50634042 | 56936102 | 46879646 | 50028406 | 47186694 | 47953718 | 42607458 | 43167114 | 38538834 | 30731018 | 34186754 | 9 | 3 | 9 | 0 | 9 | 3 | 0.994028306 | 1.046380862 | 0.755473122 | 0.795261675 |
| P97311 | Mcm6 | DNA replication licensing factor MCM6 | 1290.84 | 28.38 | 92867 | 24 | 37789354 | 43780594 | 49054098 | 48599262 | 45923758 | 38590738 | 36866722 | 45899366 | 45316218 | 45477046 | 36577270 | 40009406 | 43314302 | 43449998 | 43148658 | 33311242 | 36269778 | 38589930 | 39959430 | 41951278 | 4 | 3 | 8 | 5 | 8 | 5 | 0.91717666 | 0.973365762 | 0.844255541 | 0.895977268 |
| Q61881 | Mcm7 | DNA replication licensing factor MCM7 | 972.11 | 33.66 | 81211 | 19 | 23858834 | 20655602 | 26618666 | 24918838 | 21428142 | 25774700 | 20231168 | 22700802 | 20291620 | 23536122 | 23931026 | 21826168 | 18654340 | 18879382 | 19868020 | 21404418 | 28331040 | 18841392 | 14965177 | 16449568 | 5 | 6 | 6 | 7 | 6 | 7 | 0.878097242 | 0.916687919 | 0.851136578 | 0.888542386 |
| Q8R3C0 | Mcmbp | Mini-chromosome maintenance complex-binding protein | 271.74 | 12.62 | 72891 | 6 | 3830269.5 | 3374760 | 4359650.5 | 3110883.5 | 4486759 | 2543865 | 3143353 | 2272199 | 2651695 | 2540890.5 | 1803347.88 | 3423129.75 | 3197089.25 | 1577126 | 1512112.25 | 2777775 | 3348105 | 2493730 | 3580766 | 2824275.5 | 3 | 3 | 10 | 6 | 10 | 6 | 0.600804267 | 0.875365187 | 0.784072573 | 1.142385085 |
| Q4G009 | Mcts1 | Malignant T-cell-amplified sequence 1 | 149.22 | 17.03 | 20550 | 3 | 291451.03 | 350097.59 | 561658 | 637270.38 | 736276.12 | 290433.38 | 394638.44 | 446927.62 | 715319.38 | 318958.09 | 274354.94 | 290652 | 274397.56 | 2 | 266392.16 | 811012 | 389487.69 | 360055.78 | 165291.84 | 593023.81 | 0 | 12 | 1 | 12 | 1 | 12 | 0.429144201 | 0.510460438 | 0.899919787 | 1.070440768 |
| P14152 | Mdh1 | Malate dehydrogenase, cytoplasmic | 636.47 | 45.51 | 36511 | 11 | 22036694 | 49910426 | 46172882 | 39982674 | 51026278 | 37292602 | 47256646 | 41479070 | 51109874 | 40001402 | 21705984 | 43116634 | 31204270 | 24709584 | 28738710 | 25527710 | 28274910 | 33593774 | 31366234 | 36460742 | 5 | 5 | 3 | 0 | 5 | 5 | 0.714751253 | 0.688382893 | 0.742237586 | 0.71485521 |
| P08249 | Mdh2 | Malate dehydrogenase, mitochondrial | 1660.27 | 70.41 | 35611 | 26 | 156491234 | 179266498 | 162613058 | 189021090 | 172215122 | 189525922 | 165182690 | 196283874 | 182708578 | 199294098 | 241602178 | 214831170 | 227767010 | 283692354 | 295479394 | 189399058 | 181600914 | 226280642 | 229399794 | 224283810 | 0 | 1 | 0 | 7 | 0 | 7 | 1.469708952 | 1.354103598 | 1.222610118 | 1.12644123 |
| Q9D967 | Mdp1 | Magnesium-dependent phosphatase 1 | 45.11 | 10.98 | 18582 | 1 | 2 | 2 | 140490.3 | 478400.06 | 217462.11 | 290149.84 | 151159.05 | 296112.25 | 531357.06 | 2 | 2 | 2 | 2 | 150706.8 | 281296.31 | 190155.7 | 2 | 596294.94 | 2 | 2 | 10 | 11 | 4.5 | 8.5 | 10 | 11 | 0.516537057 | 0.340491686 | 0.940336649 | 0.619852548 |
| P06801 | Me1 | NADP-dependent malic enzyme | 304.36 | 11.71 | 63954 | 4 | 2988664.25 | 2455303 | 1136488.25 | 1034470.75 | 2135805.5 | 2863664.25 | 3542626 | 2342633.5 | 2572697 | 1667487 | 2565789 | 2910383.5 | 1844046.62 | 1286841.5 | 1277493.12 | 2720108.75 | 1699384.12 | 3287859.5 | 2480004.25 | 2824789 | 11 | 6 | 7 | 12 | 11 | 12 | 1.013724302 | 0.760987893 | 1.334478884 | 1.00177363 |
| Q99KE1 | Me2 | NAD-dependent malic enzyme, mitochondrial | 696.62 | 23.77 | 65799 | 10 | 4310530 | 5014003 | 6893372.5 | 5633564 | 7648637 | 4372517 | 4975544 | 6186280.5 | 6408223 | 7270447.5 | 5413979 | 6128506 | 8614955 | 7298739.5 | 9188428 | 2692426 | 4301348 | 6466821 | 3970947.25 | 5945152 | 6 | 6 | 6 | 6 | 6 | 6 | 1.2421856 | 1.254393333 | 0.792427453 | 0.800215132 |
| Q9DCS3 | Mecr | Enoyl-[acyl-carrier-protein] reductase, mitochondrial | 259.85 | 16.89 | 40343 | 3 | 1114500.12 | 983929.75 | 1021684 | 1123858 | 1152034.12 | 1646451.75 | 1142494.38 | 1443912.62 | 1103186.38 | 1205162.75 | 1402333.5 | 1190058.5 | 870490.5 | 1133127.75 | 1067167.38 | 1184852 | 1083557.25 | 1615096.62 | 1260978.5 | 960860.56 | 9 | 8 | 6 | 9 | 9 | 9 | 1.049512851 | 0.865769401 | 1.131456292 | 0.933366596 |
| A0A096 | Mef2c | Myocyte-specific enhancer factor 2C | 50.12 | 2.11 | 51217 | 1 | 653169.5 | 992310.56 | 1435162.75 | 1802649.5 | 1306426 | 1082852.25 | 871693.19 | 912251.06 | 882648.12 | 1335805 | 2 | 2 | 2 | 529752.94 | 361485.06 | 1183822.25 | 810601.25 | 739613.56 | 901207 | 926765.5 | 0 | 6 | 0 | 9 | 0 | 9 | 0.143987813 | 0.17526062 | 0.737030238 | 0.897106317 |
| Q91VH6 | Memo1 | Protein MEMO1 | 177.21 | 13.47 | 33692 | 4 | 4246485.5 | 5025661.5 | 4376833 | 2429172.25 | 3485846 | 4284441 | 4324059 | 4111959.5 | 2673125.5 | 3841405.25 | 4333500 | 3171920.25 | 3027536.5 | 4979998 | 3511601.25 | 2994766.25 | 4098180.5 | 5040922 | 5216172 | 5538381.5 | 11 | 7 | 12 | 7 | 12 | 7 | 0.972426789 | 0.98905982 | 1.169925593 | 1.189936774 |
| Q9ERE7 | Mesd | LRP chaperone MESD | 283.64 | 30.36 | 25207 | 6 | 12319837 | 12693095 | 13006123 | 13808067 | 10417456 | 14769034 | 11950604 | 12293943 | 9816110 | 13201995 | 15334771 | 13636763 | 16111319 | 18345794 | 9191180 | 12385916 | 15302816 | 14572082 | 15081028 | 14080949 | 6 | 3 | 6 | 4 | 6 | 4 | 1.166685185 | 1.170689235 | 1.147454016 | 1.151392064 |
| Q8BP48 | Metap1 | Methionine aminopeptidase 1 | 236.34 | 13.21 | 43221 | 4 | 6089651 | 6351746 | 6006763 | 7191936.5 | 6441245 | 6110400.5 | 5551956 | 6350584 | 5957317.5 | 5847226 | 5951033 | 7379858.5 | 6975546.5 | 4825710.5 | 6878409.5 | 7024499.5 | 5431972 | 6523473.5 | 7118978 | 6507045 | 12 | 9 | 8 | 5 | 12 | 9 | 0.997793624 | 1.073549935 | 1.01635301 | 1.09351842 |
| O08663 | Metap2 | Methionine aminopeptidase 2 | 224.63 | 9.41 | 52922 | 3 | 602323.19 | 1507991.5 | 504710.69 | 2 | 1034725.19 | 974621.56 | 1725758 | 649487.56 | 1453928.88 | 2 | 187802.45 | 1086006.62 | 2 | 2 | 419189.31 | 1231950.5 | 2 | 329976.5 | 654081.62 | 1074623.75 | 7 | 12.5 | 6 | 9.5 | 7 | 12.5 | 0.463867714 | 0.352429969 | 0.901604782 | 0.685006815 |
| Q9Z120 | Mettl1 | tRNA (guanine-N(7)-)-methyltransferase | 57.62 | 5.6 | 30603 | 1 | 193458.27 | 179367.48 | 172177.12 | 188189.12 | 231476.55 | 109255.9 | 249320.41 | 252363.08 | 179252.75 | 240041.62 | 300996 | 202649.58 | 134346.86 | 47811.61 | 100463.99 | 250313.84 | 304945.56 | 168037.31 | 197131.88 | 182164.89 | 9 | 9 | 8 | 11 | 9 | 11 | 0.815065494 | 0.763193821 | 1.142976509 | 1.070236215 |
| Q91YR5 | Mettl13 | Methyltransferase-like protein 13 | 88.49 | 3.01 | 78757 | 2 | 808707.69 | 1009014.75 | 826023.44 | 1015986.81 | 877510.88 | 825582.69 | 1184860.62 | 799331.94 | 1180821.62 | 1118007.62 | 1151585 | 1201740.62 | 1202171.25 | 997995.62 | 1235742.25 | 927362.75 | 738316.94 | 875536.25 | 961111.62 | 996950.94 | 2 | 11 | 5 | 8 | 5 | 11 | 1.275936513 | 1.133232129 | 0.99163257 | 0.88072555 |
| Q9CQG2 | Mettl16 | U6 small nuclear RNA (adenine-(43)-N(6))-methyltransferase | 67.43 | 2.35 | 62341 | 1 | 372208.03 | 444391.91 | 371230.88 | 296252.56 | 462055.62 | 193416.31 | 421629.94 | 372699.78 | 592393.12 | 290923.56 | 277091.94 | 471576.97 | 330922.53 | 713740.62 | 415329.88 | 580605.19 | 1031156.69 | 207956.98 | 2 | 2 | 11 | 10 | 10 | 10 | 11 | 10 | 1.134894239 | 1.180431809 | 0.935042595 | 0.972561128 |
| Q9DCS2 | Mettl26 | Methyltransferase-like 26 | 192.22 | 20.59 | 22687 | 3 | 2208872 | 2476992.5 | 2182219.25 | 2531833.75 | 2459904.75 | 1884708.12 | 1782221.12 | 2311878.5 | 1085675.5 | 1703384 | 2145717.25 | 1802137.62 | 2113472.5 | 872256.31 | 1944645.5 | 2265350 | 2546903.25 | 2516875.25 | 2713488.25 | 2746460 | 0 | 4 | 10 | 1 | 10 | 4 | 0.748597154 | 1.012587091 | 1.078353156 | 1.458630292 |
| P21956 | Mfge8 | Lactadherin | 784.51 | 33.91 | 51241 | 15 | 12139065 | 11993242 | 9373173 | 10318242 | 10711266 | 12112047 | 11978200 | 12052257 | 12826883 | 12419535 | 7007830 | 15075400 | 21460946 | 19788294 | 19408952 | 9057247 | 12702488 | 11917430 | 12900406 | 10138141 | 5 | 11 | 5 | 9 | 5 | 11 | 1.517217204 | 1.347823342 | 1.039987613 | 0.923875353 |
| Q811U4 | Mfn1 | Mitofusin-1 | 49.41 | 1.89 | 83726 | 1 | 198409.69 | 353020.41 | 351699.16 | 2 | 433478.84 | 2 | 2 | 308302.12 | 332856.38 | 299174.94 | 2 | 2 | 2 | 2 | 2 | 2 | 2 | 245072.36 | 222092.91 | 2 | 2.5 | 5.5 | 5 | 7 | 5 | 7 | 7.481613E-06 | 1.063448E-05 | 0.349519482 | 0.496812368 |
| Q9DC37 | Mfsd1 | Major facilitator superfamily domain-containing protein 1 | 131.8 | 8.19 | 51396 | 3 | 66925.58 | 71070.46 | 277539.81 | 325388.09 | 520446.53 | 706534.88 | 341025.62 | 1224867.25 | 871317.44 | 389695.88 | 2041294.75 | 1586709 | 4156942.75 | 4212782.5 | 3551675.25 | 964719.25 | 2 | 1569121.25 | 1326493.75 | 644430 | 0 | 5 | 0 | 9 | 0 | 9 | 12.32738884 | 4.400640606 | 3.571326868 | 1.274894971 |
| Q921Y4 | Mfsd5 | Molybdate-anion transporter | 56.79 | 2.67 | 49679 | 1 | 2 | 2 | 2 | 153985.73 | 2 | 2 | 2 | 2 | 2 | 195324.52 | 2 | 260299.33 | 537950.88 | 307055.69 | 813703 | 112958.91 | 130504.43 | 208081.17 | 2 | 247338.53 | 3 | 5 | 3 | 5 | 3 | 5 | 12.46161711 | 9.824328791 | 4.538399323 | 3.577924659 |
| P27808 | Mgat1 | Alpha-1,3-mannosyl-glycoprotein 2-beta-N-acetylglucosaminyltransferase | 42.3 | 1.79 | 51690 | 1 | 2 | 2 | 2 | 2 | 2 | 2 | 2 | 505149.5 | 215165.3 | 2 | 1040889.12 | 845396.75 | 737526.44 | 881087.56 | 682514.25 | 2 | 2 | 548608.31 | 2 | 2 | 0 | 10 | 0 | 11 | 0 | 11 | 418741.412 | 5.813262813 | 54861.631 | 0.76162775 |
| Q09326 | Mgat2 | Alpha-1,6-mannosyl-glycoprotein 2-beta-N-acetylglucosaminyltransferase | 80.61 | 4.52 | 51110 | 2 | 209828.31 | 2 | 2 | 2 | 2 | 198351.94 | 250603 | 301783.44 | 397797.12 | 199961.23 | 2 | 443379.88 | 2 | 2 | 399499.47 | 254369.73 | 2 | 381951.03 | 2 | 2 | 9 | 9 | 10 | 7 | 10 | 9 | 4.016870817 | 0.625055539 | 3.032491183 | 0.471878608 |
| Q08834 | Mgat5 | Alpha-1,6-mannosylglycoprotein 6-beta-N-acetylglucosaminyltransferase A | 84.68 | 1.62 | 84562 | 1 | 2 | 2 | 2 | 2 | 76106.23 | 2 | 2 | 2 | 2 | 2 | 153136.39 | 190159.77 | 222033.16 | 2 | 203658.31 | 2 | 2 | 2 | 2 | 2 | 3 | 10 | 2.5 | 12.5 | 3 | 12.5 | 10.10309938 | 76898.963 | 0.000131381 | 1 |
| O35678 | Mgll | Monoglyceride lipase | 46.75 | 2.31 | 33388 | 1 | 2254833.75 | 2698437.75 | 2329699.75 | 2347762 | 1876888 | 2274749 | 2475376 | 2519352 | 2221599.75 | 2425861 | 2768398.5 | 2977746.75 | 2727332 | 2750064.75 | 2562721 | 2928965.75 | 2 | 2427600 | 2690053.75 | 2661585 | 1 | 8 | 0 | 7 | 1 | 8 | 1.198011535 | 1.156862886 | 0.930531712 | 0.898570314 |
| Q9CPU4 | Mgst3 | Microsomal glutathione S-transferase 3 | 256.37 | 23.53 | 16958 | 4 | 3716720.5 | 3163702 | 6959056 | 6691518 | 9663062 | 3191056.75 | 4599241 | 6860160.5 | 6508568 | 5455014 | 2226958.25 | 2409157.5 | 1923441.88 | 3594643.5 | 3118553.75 | 3405266.75 | 3678471.5 | 5036874.5 | 2126275 | 5669032.5 | 1 | 6 | 1 | 7 | 1 | 7 | 0.439581677 | 0.498712512 | 0.659597326 | 0.748323819 |
| Q8BI84 | Mia3 | Transport and Golgi organization protein 1 homolog | 168.28 | 2.18 | 213675 | 3 | 985867.56 | 572670.06 | 507443.19 | 430198.62 | 545346.94 | 728134.38 | 792354.5 | 607216.81 | 448509.75 | 414125.44 | 935228.94 | 1337484.25 | 716950 | 950402.88 | 1141008 | 742003.38 | 1347671 | 448666.66 | 591598.31 | 534720.38 | 3 | 9 | 2 | 10 | 3 | 10 | 1.67056716 | 1.69916216 | 1.204875212 | 1.225498991 |
| P30904 | Mif | Macrophage migration inhibitory factor | 220.99 | 41.74 | 12477 | 4 | 17232154 | 17757322 | 28513126 | 20857658 | 41574978 | 18563244 | 20976512 | 31811134 | 39293986 | 17463574 | 12320615 | 15969944 | 16496787 | 20589474 | 18774240 | 29691590 | 22420934 | 21775508 | 21239306 | 24062388 | 4 | 9 | 4 | 10 | 4 | 10 | 0.668209004 | 0.65687361 | 0.94643666 | 0.930381454 |
| Q5BJQ2 | Mindy1 | Ubiquitin carboxyl-terminal hydrolase MINDY-1 | 57.41 | 2.07 | 52729 | 1 | 2016416.25 | 2212364.25 | 2165413.75 | 2779813.5 | 1658755.88 | 2913990 | 2865222.5 | 2582488.5 | 2246486.5 | 3475449.25 | 2975171.75 | 3140141.5 | 2883333.75 | 2575998.5 | 2336722.25 | 2911418 | 2804470 | 3132453 | 3436198 | 3054314.75 | 2 | 0 | 12 | 8 | 12 | 8 | 1.284193787 | 0.987768145 | 1.415968655 | 1.089125914 |
| Q6PDI6 | Mindy2 | Ubiquitin carboxyl-terminal hydrolase MINDY-2 | 53.7 | 2.5 | 65637 | 1 | 528071.5 | 2 | 411900.22 | 769175.12 | 2 | 620789.94 | 560266 | 652916.69 | 526981.81 | 2 | 417432.81 | 446361.06 | 385032.34 | 2 | 534827.31 | 2 | 634793 | 756179.38 | 561147.25 | 746942.81 | 12 | 8 | 5.5 | 7.5 | 12 | 8 | 1.043591635 | 0.755480063 | 1.579184456 | 1.143208064 |
| F1LP90 | Mink1 | Misshapen-like kinase 1 | 41.82 | 0.52 | 150382 | 1 | 360801.78 | 444664.25 | 367880.47 | 373048.31 | 862124.25 | 351657.44 | 461389.62 | 60002.16 | 620547.5 | 371584.41 | 2 | 2 | 354931.69 | 2 | 373807.56 | 480689.97 | 2 | 319357.31 | 2 | 248693.84 | 3 | 4 | 5 | 6 | 5 | 6 | 0.30256985 | 0.390710177 | 0.435431522 | 0.562275215 |
| Q9Z2L6 | Minpp1 | Multiple inositol polyphosphate phosphatase 1 | 58.11 | 2.29 | 54537 | 1 | 2 | 207437.12 | 294283.69 | 2 | 280041.03 | 230221.94 | 2 | 264269.97 | 449771.25 | 2 | 2 | 199523.55 | 365408.03 | 471652.28 | 2 | 2 | 211579.95 | 240346.08 | 2 | 2 | 11 | 9 | 12 | 8 | 12 | 9 | 1.325956964 | 1.097769682 | 0.578091299 | 0.478606108 |
| Q7TQ72 | Mis12 | Protein MIS12 homolog | 43.76 | 7.77 | 24161 | 1 | 2 | 2 | 303883.12 | 2 | 2 | 2 | 2 | 212204.17 | 2 | 241004.27 | 2 | 2 | 78559.59 | 2 | 2 | 2 | 330340.38 | 151045.67 | 191455.28 | 88409.26 | 12 | 6 | 9 | 9.5 | 12 | 9.5 | 0.258538617 | 0.173356326 | 2.505017554 | 1.679674174 |
| E9PVX6 | Mki67 | Proliferation marker protein Ki-67 | 2137.01 | 14.73 | 350864 | 38 | 18237548 | 21623272 | 20936230 | 20104740 | 13855315 | 24192430 | 20341564 | 18846838 | 13036574 | 17339558 | 14189655 | 12522389 | 7815218 | 8020240.5 | 9065882 | 19992314 | 20396592 | 17253004 | 13625207 | 15613260 | 1 | 7 | 1 | 10 | 1 | 10 | 0.544691446 | 0.550501875 | 0.916874539 | 0.926655187 |
| Q6ZQI3 | Mlec | Malectin | 329.39 | 28.52 | 32342 | 7 | 7347936 | 10400309 | 9692894 | 9708710 | 14184276 | 6081530.5 | 10256436 | 9442073 | 12863759 | 11240364 | 4933768 | 8231612.5 | 7148915 | 6348613 | 8988706 | 10080299 | 6448740 | 7348225.5 | 7589777 | 8601836 | 2 | 6 | 4 | 6 | 4 | 6 | 0.69450126 | 0.714688044 | 0.780550511 | 0.803238453 |
| P97679 | Mlh1 | DNA mismatch repair protein Mlh1 | 57.8 | 1.59 | 84540 | 1 | 665936.25 | 459908.78 | 691015.5 | 2 | 2 | 535016.62 | 426957.38 | 2 | 459299.88 | 482822.91 | 2 | 2 | 2 | 2 | 383626.38 | 604999.44 | 2 | 482963.09 | 2 | 594742.88 | 6 | 11 | 3 | 10 | 6 | 11 | 0.21115189 | 0.201478191 | 0.926161187 | 0.883730098 |
| Q9D2Y4 | Mlkl | Mixed lineage kinase domain-like protein | 120.64 | 5.3 | 54317 | 2 | 2367858.25 | 1408879.25 | 1728854.75 | 2239227.5 | 1661638.5 | 2493080 | 2111188.75 | 2524071.5 | 2342583.75 | 2740276 | 4045111.5 | 3493493.5 | 2694530.5 | 2943524.5 | 3147656.25 | 2513992.75 | 2944931.5 | 2757735 | 3209169 | 2584593 | 0 | 0 | 1 | 3 | 1 | 3 | 1.73543706 | 1.336831454 | 1.489447024 | 1.147341887 |
| P34960 | Mmp12 | Macrophage metalloelastase | 826.32 | 32.35 | 54971 | 16 | 8923396 | 13056070 | 2899567.5 | 3306608.25 | 4041001.5 | 9698140 | 14138530 | 8938428 | 11708752 | 9626907 | 71537586 | 110430474 | 55559586 | 39557898 | 59303914 | 13179429 | 12258858 | 13332498 | 11676975 | 8454364 | 0 | 4 | 0 | 10 | 0 | 10 | 10.43824066 | 6.216683644 | 1.827746177 | 1.088547403 |
| Q9D071 | Mms19 | MMS19 nucleotide excision repair protein homolog | 402.67 | 7.47 | 113089 | 8 | 3899649.25 | 2728397 | 2547440 | 3025475 | 3855509.75 | 2996352.25 | 3834221 | 2866670.5 | 3441679.25 | 4264386 | 3454098.5 | 3666902.75 | 4113587.5 | 5067635.5 | 5253003 | 1740205.5 | 2306669.25 | 2674776.75 | 4493419.5 | 5727415.5 | 4 | 11 | 5 | 10 | 5 | 11 | 1.342463562 | 1.238570622 | 1.05518121 | 0.973520984 |
| D0QMC3 | Mndal | Myeloid cell nuclear differentiation antigen-like protein | 604.6 | 22.49 | 60526 | 10 | 6893806.5 | 6762796 | 6899946 | 8467844 | 7044165 | 7065211 | 6683493.5 | 7077912.5 | 7274583 | 7961776 | 1991161.12 | 2951066.75 | 3462431.5 | 910768.69 | 4816349 | 4785538.5 | 5024533 | 6334546.5 | 5169587.5 | 4217970 | 0 | 0 | 0 | 0 | 0 | 0 | 0.391803223 | 0.391863862 | 0.70787903 | 0.707988589 |
| Q3T1J9 | Mob1a | MOB kinase activator 1A | 187.75 | 21.76 | 25080 | 4 | 11806558 | 13315284 | 13929395 | 12904683 | 11735843 | 11862706 | 10700680 | 11835682 | 10991974 | 11792921 | 5910976 | 8045097.5 | 8506417 | 9261611 | 9258878 | 11638975 | 10315143 | 11980184 | 11727154 | 11619262 | 0 | 2 | 0 | 11 | 0 | 11 | 0.64345808 | 0.716686591 | 0.899342636 | 1.001691995 |
| Q8VI63 | Mob2 | MOB kinase activator 2 | 112.72 | 7.66 | 26851 | 2 | 1728930.88 | 1570227 | 1545294.75 | 618804.25 | 1666562.62 | 2 | 967527.81 | 1484742.88 | 1210190.12 | 1041374.06 | 433636.78 | 1083169.62 | 904803.81 | 2 | 1130663 | 1385707.5 | 2 | 717353.88 | 566497.44 | 604884.31 | 3 | 2 | 8.5 | 7.5 | 8.5 | 7.5 | 0.49822793 | 0.755186735 | 0.459260593 | 0.69612217 |
| A2BDX3 | Mocs3 | Adenylyltransferase and sulfurtransferase MOCS3 | 92.49 | 5.65 | 49375 | 3 | 103970.86 | 191436.5 | 2 | 2 | 2 | 738083.88 | 283095.75 | 2 | 219736.11 | 216858.47 | 1537186.75 | 600257 | 355427.22 | 272997.09 | 658285.38 | 409255.19 | 2 | 2 | 2 | 2 | 0 | 11 | 5 | 6 | 5 | 11 | 11.59105817 | 2.34888827 | 1.385391609 | 0.280744868 |
| Q80UM7 | Mogs | Mannosyl-oligosaccharide glucosidase | 728.44 | 20.62 | 91831 | 11 | 359721.44 | 2 | 463798.38 | 273751.12 | 343742.66 | 501687.34 | 349571.78 | 2 | 502441.91 | 939755.25 | 2235877 | 2079272.5 | 2779748.25 | 1000823.44 | 2416330.75 | 2 | 2 | 547104.31 | 2 | 433724.72 | 0 | 10.5 | 0 | 7.5 | 0 | 10.5 | 7.294891145 | 4.583493858 | 0.680655386 | 0.427666393 |
| Q80TL7 | Mon2 | Protein MON2 homolog | 92 | 1.05 | 189078 | 2 | 704991.69 | 540509.31 | 623480 | 890354.88 | 1005115.69 | 623182.38 | 475511.12 | 880638.88 | 799492.44 | 519943.06 | 2 | 421588.03 | 521498.16 | 2 | 552537.62 | 435533.94 | 344451.91 | 637955.69 | 604572.81 | 685249.81 | 1 | 5 | 4 | 8 | 4 | 8 | 0.397302976 | 0.453389831 | 0.719298445 | 0.820841071 |
| P60762 | Morf4l1 | Mortality factor 4-like protein 1 | 44.26 | 2.76 | 41493 | 1 | 109287.52 | 200330.77 | 137303.36 | 46677.11 | 127905.89 | 210735.53 | 181387.36 | 99646.94 | 2 | 60756.46 | 2 | 2 | 32021.2 | 2 | 2 | 295872.78 | 245505.05 | 69804.91 | 162816.8 | 59273.48 | 0 | 9 | 3 | 9 | 3 | 9 | 0.051534932 | 0.057968435 | 1.340734973 | 1.508109241 |
| Q9CWP6 | Mospd2 | Motile sperm domain-containing protein 2 | 194.4 | 8.11 | 59855 | 4 | 1783912.38 | 2405379.25 | 2052271.38 | 2025955.5 | 2402423.25 | 1553619.75 | 2187250 | 2238720 | 2250888.5 | 2118214.5 | 1945549.5 | 2666123.5 | 2777637 | 1834400.5 | 2374622 | 1874793.62 | 1883672.5 | 2299452.5 | 1979774 | 2342532.5 | 10 | 9 | 8 | 12 | 10 | 12 | 1.087009916 | 1.120753392 | 0.972847402 | 1.003046991 |
| P23249 | Mov10 | Putative helicase MOV-10 | 497.67 | 13.05 | 113583 | 10 | 7510070.5 | 6280248.5 | 6082475 | 6991612.5 | 7052520.5 | 6807734 | 5728600.5 | 6984189.5 | 6828921 | 7558099 | 5072836 | 4444203.5 | 6795630.5 | 5830138 | 3863139 | 6466199 | 5703308.5 | 7710373 | 5850296.5 | 7136244 | 2 | 11 | 2 | 11 | 2 | 11 | 0.766754223 | 0.766966401 | 0.969027088 | 0.96929524 |
| P63030 | Mpc1 | Mitochondrial pyruvate carrier 1 | 70.82 | 14.68 | 12455 | 2 | 2209863.75 | 1931558.75 | 3057470.5 | 3244938 | 2667824 | 2426439 | 1989744.12 | 3168308 | 2642439.25 | 3412142 | 1981376.12 | 1973830.12 | 2 | 3823268 | 4073084 | 1890498.25 | 2794880.5 | 3346465.5 | 3906557.75 | 3756470.25 | 12 | 7 | 10 | 8 | 12 | 8 | 0.903895064 | 0.868941811 | 1.197016872 | 1.150728717 |
| P38718 | Mpc2 | Mitochondrial pyruvate carrier 2 | 188.91 | 29.13 | 14258 | 4 | 5655039 | 3098155.75 | 4030487 | 4954494 | 4268535 | 5854528.5 | 5913731 | 4752773.5 | 5082556.5 | 5962475 | 5056989 | 2406865 | 4702274 | 5879093.5 | 4546548.5 | 4850857.5 | 6735686.5 | 6606043.5 | 6623689.5 | 4775514 | 10 | 4 | 4 | 8 | 10 | 8 | 1.026585493 | 0.819550067 | 1.344671239 | 1.073486242 |
| Q9R0Q9 | Mpdu1 | Mannose-P-dolichol utilization defect 1 protein | 196.53 | 12.55 | 26498 | 3 | 894725.25 | 1111344 | 956907.5 | 530876.25 | 1356653.62 | 1045957.88 | 1299502.5 | 1623486.5 | 1918304.75 | 1632645 | 1621241.88 | 2978616.5 | 3488632.5 | 2346680.75 | 4233256 | 1071924.38 | 578161.38 | 1909798.5 | 1647555.62 | 1802990.5 | 0 | 6 | 3 | 12 | 3 | 12 | 3.024102177 | 1.950615594 | 1.44529859 | 0.932250897 |
| A1L314 | Mpeg1 | Macrophage-expressed gene 1 protein | 1840.74 | 54.98 | 78390 | 26 | 16524213 | 16162328 | 24126432 | 22175300 | 26341332 | 26594852 | 27589238 | 38793926 | 41408134 | 39330802 | 230335586 | 217198898 | 270221090 | 233340978 | 271181218 | 24820612 | 31884024 | 56792498 | 54802182 | 51363146 | 0 | 1 | 0 | 8 | 0 | 8 | 11.60431362 | 7.036030485 | 2.085476937 | 1.264484896 |
| Q810V0 | Mphosph10 | U3 small nucleolar ribonucleoprotein protein MPP10 | 49.77 | 2.64 | 78735 | 1 | 2 | 2 | 80314.07 | 2 | 2 | 2 | 2 | 131115.11 | 2 | 2 | 107843.17 | 85482.25 | 2 | 2 | 2 | 2 | 2 | 161283.62 | 2 | 212389.98 | 9 | 9 | 11 | 9 | 11 | 9 | 2.406952659 | 1.474426743 | 4.65226556 | 2.849837836 |
| Q924M7 | Mpi | Mannose-6-phosphate isomerase | 174.24 | 7.09 | 46575 | 3 | 1262522.5 | 1569284.25 | 1591880 | 1711504.12 | 2037902.25 | 1248247.25 | 1126844.38 | 2306488.25 | 1260217.25 | 1311299.38 | 2 | 765651.62 | 1233381.38 | 459818.5 | 937681.81 | 2859453 | 1428225.75 | 1349508.38 | 1054772.25 | 1580521.75 | 0 | 9 | 1 | 8 | 1 | 9 | 0.415575261 | 0.468287621 | 1.012160391 | 1.140544748 |
| P70290 | Mpp1 | 55 kDa erythrocyte membrane protein | 277.96 | 15.45 | 52227 | 5 | 2040964.38 | 1814750.75 | 1716996 | 865196.25 | 2472083.25 | 1001144.88 | 1941169.5 | 2251189 | 2709654 | 1550445.5 | 2826788.5 | 3714317.5 | 2161789.5 | 1654725.5 | 2795943.5 | 1730907 | 710723.75 | 1598291.62 | 2292917 | 567606.38 | 5 | 7 | 5 | 8 | 5 | 8 | 1.476271418 | 1.391381113 | 0.774461617 | 0.729927609 |
| Q9JLB0 | Mpp6 | MAGUK p55 subfamily member 6 | 254.66 | 10.13 | 62631 | 5 | 2945363.75 | 3530438.5 | 4075186.25 | 1401917.75 | 3926396.5 | 970821.12 | 4097352.5 | 3017014.75 | 5125639 | 3839219.75 | 2345382.5 | 4422723 | 1961109.75 | 2054361.25 | 3881422 | 2108987 | 2 | 3129645.5 | 1798301.5 | 806611.94 | 11 | 4 | 10 | 4 | 11 | 4 | 0.92352912 | 0.860114837 | 0.493947881 | 0.460030866 |
| Q99J99 | Mpst | 3-mercaptopyruvate sulfurtransferase | 106.01 | 8.75 | 33097 | 2 | 2 | 2 | 427705.16 | 2 | 310507.25 | 268953.66 | 132962.69 | 465116.34 | 93027.92 | 351283.06 | 1029353.69 | 459574.16 | 904809.12 | 692614.19 | 675486.81 | 144186.08 | 603379.19 | 282284.91 | 184493.89 | 229563.42 | 0 | 8 | 1 | 11 | 1 | 11 | 5.095833318 | 2.868689617 | 1.955935358 | 1.101090067 |
| Q61216 | Mre11 | Double-strand break repair protein MRE11 | 125.38 | 3.82 | 80223 | 2 | 277325.38 | 271916.56 | 286955.75 | 2 | 2 | 2 | 262724.78 | 2 | 2 | 237909.05 | 2 | 316771.53 | 2 | 2 | 2 | 2 | 2 | 2 | 2 | 170656.16 | 9 | 6 | 11 | 9 | 11 | 9 | 0.378831487 | 0.632749356 | 0.204094493 | 0.340892094 |
| Q9DAT2 | Mrgbp | MRG/MORF4L-binding protein | 65.29 | 6.37 | 22436 | 1 | 634469.69 | 660099.06 | 449156.53 | 401324.41 | 389612.97 | 753026.75 | 629838.69 | 2 | 513509.47 | 380911.69 | 2 | 335809.16 | 146461.84 | 305455.56 | 262794.94 | 564248.25 | 759842.75 | 338792.25 | 586772.5 | 333663.91 | 0 | 11 | 4.5 | 12 | 4.5 | 12 | 0.414462846 | 0.461304509 | 1.019196637 | 1.13438396 |
| Q9CQT1 | Mri1 | Methylthioribose-1-phosphate isomerase | 405.92 | 18.97 | 39411 | 7 | 3068831 | 2861088.5 | 3355616 | 3666644.5 | 3198806 | 4109542.25 | 2558197.5 | 3471665.25 | 3077973.25 | 3644750 | 5734005.5 | 3244468.5 | 3980867.25 | 5378106 | 3870607.5 | 1906868.62 | 5158204 | 5062013.5 | 5017508.5 | 4274164 | 2 | 5 | 5 | 5 | 5 | 5 | 1.3750278 | 1.317037471 | 1.326157958 | 1.270228663 |
| Q99N96 | Mrpl1 | 39S ribosomal protein L1, mitochondrial | 56.11 | 3.57 | 37597 | 1 | 2 | 991315.88 | 1161973.75 | 1110172.25 | 2 | 1254633.25 | 1197891.75 | 941777.62 | 2 | 1114983 | 895158.12 | 990213.62 | 1101125.5 | 759777.44 | 1310468.62 | 1310435.25 | 856433.38 | 1220884.25 | 810295.44 | 1192325.25 | 11 | 6 | 11 | 11 | 11 | 11 | 1.549500894 | 1.121406246 | 1.65173278 | 1.195393602 |
| Q3TBW2 | Mrpl10 | 39S ribosomal protein L10, mitochondrial | 58.14 | 3.05 | 29396 | 1 | 1500845.5 | 1352037.12 | 985123.31 | 1232454.62 | 1148303 | 1574088.88 | 1035979.44 | 1289162.25 | 965777.44 | 978925.88 | 969761.62 | 828084.94 | 1017788.75 | 1050746.88 | 1009605.19 | 1495722.88 | 1849781.12 | 1521415.88 | 1290509.5 | 1599549.38 | 3 | 3 | 8 | 3 | 8 | 3 | 0.784076664 | 0.834367307 | 1.247350651 | 1.327355666 |
| Q9CQF0 | Mrpl11 | 39S ribosomal protein L11, mitochondrial | 89.78 | 15.1 | 20680 | 2 | 1159855.38 | 1746464.5 | 2084219.38 | 1818164.38 | 1893918.5 | 2384154.5 | 1945621.75 | 1578426 | 1151826.88 | 1630715.12 | 535180 | 1075968.88 | 1101514.25 | 1577147.38 | 1482784.75 | 1752114 | 1868868.75 | 2103840.25 | 2007099.25 | 953417.94 | 2 | 11 | 2 | 11 | 2 | 11 | 0.663316776 | 0.66422335 | 0.998014167 | 0.999378182 |
| Q9DB15 | Mrpl12 | 39S ribosomal protein L12, mitochondrial | 51.6 | 6.97 | 21708 | 1 | 2 | 2 | 2 | 2 | 2 | 2 | 2 | 2 | 2 | 513417.78 | 1694530.38 | 1024080.44 | 907024.12 | 1202353.5 | 1152627.75 | 2 | 2 | 2 | 2 | 2 | 0 | 12.5 | 0 | 10 | 0 | 12.5 | 598061.619 | 11.64845324 | 1 | 1.947701E-05 |
| Q7M0E7 | Mrpl14 | 39S ribosomal protein L14, mitochondrial | 69.4 | 6.9 | 15913 | 1 | 2 | 2 | 2 | 2 | 2 | 2 | 2 | 2 | 2 | 2 | 718515.69 | 419700.66 | 289668.5 | 2 | 264955.31 | 2 | 2 | 2 | 2 | 2 | 2.5 | 12.5 | 2.5 | 12.5 | 2.5 | 12.5 | 169284.216 | 169284.216 | 1 | 1 |
| Q9CPR5 | Mrpl15 | 39S ribosomal protein L15, mitochondrial | 283.96 | 23.39 | 33542 | 5 | 1927839.88 | 3177078.75 | 2198457.75 | 1772618.12 | 2217582.5 | 2200819.75 | 2327372.25 | 1148367 | 1964178.25 | 1463587.12 | 1784919.12 | 2530076.25 | 2320614.5 | 1836002.88 | 1576287.38 | 1685108 | 954373.69 | 1133273 | 111953.42 | 543764.06 | [illegible] | [illegible] | [illegible] | [illegible] | [illegible] | [illegible] | 0.889700414 | 1.103640393 | 0.392123078 | 0.486414147 |
| Q5M8I8 | Mrpl16 | 39S ribosomal protein L16, mitochondrial | 40.73 | 3.98 | 28895 | 1 | 766324.81 | 1205346.5 | 956467.25 | 733775.94 | 541500.06 | 1029186.31 | 677045.81 | 771846.31 | 811870.5 | 739554.19 | 853536.81 | 1072108 | 1137162.38 | 1430380.75 | 1184080.88 | 678825.31 | 892603.75 | 857931.25 | 796685.06 | 783537.5 | 5 | 12 | 1 | 10 | 5 | 12 | 1.35063262 | 1.408925282 | 0.953887087 | 0.9950564 |
| Q9D338 | Mrpl19 | 39S ribosomal protein L19, mitochondrial | 313.41 | 18.49 | 33578 | 7 | 5482248 | 4535654.5 | 5267154 | 5069981.5 | 3406188 | 4742736.5 | 3947077.75 | 3851559.75 | 4083688.75 | 4411571 | 4033813.25 | 3965195.5 | 4078571.5 | 4287491 | 4675547.5 | 5757807.5 | 5024416 | 3917273.75 | 4979649.5 | 4438270 | 6 | 11 | 12 | 5 | 12 | 11 | 0.885502236 | 1.000189431 | 1.01499042 | 1.146448478 |
| Q9CQL4 | Mrpl20 | 39S ribosomal protein L20, mitochondrial | 45.54 | 7.38 | 17595 | 1 | 2 | 197798.73 | 145365.58 | 2 | 145747.77 | 2 | 2 | 2 | 256114.42 | 2 | 437219.59 | 265626.47 | 359458.47 | 134686.91 | 689950.81 | 2 | 2 | 2 | 2 | 2 | 3 | 5 | 1 | 10 | 3 | 10 | 3.857861819 | 7.367345077 | 2.044504E-05 | 3.904383E-05 |

| | | | | | | | | | | | | | | | | | | | | | | | | | | | | | | | | | | | | |
|---|---|---|---|---|---|---|---|---|---|---|---|---|---|---|---|---|---|---|---|---|---|---|---|---|---|---|---|---|---|---|---|---|---|---|---|---|
| Q9D1N9 | Mrpl21 | 39S ribosomal protein L21, mitochondrial | 177.53 | 17.7 | 23366 | 3 | 408183.66 | 897826.69 | 717081 | 302972.16 | 311384.34 | 710793.88 | 511730.81 | 206778.38 | 437579.5 | 479329.47 | 4073321.5 | 2806251 | 2887193.75 | 2492870.75 | 2551461 | 306327.84 | 420573.09 | 480779.12 | 358503.91 | 350933.59 | 0 | 11 | 0 | 7 | 0 | 11 | 5.615693216 | 6.312770435 | 0.726883586 | 0.817111803 |
| Q8BU88 | Mrpl22 | 39S ribosomal protein L22, mitochondrial | 89.09 | 12.14 | 23805 | 2 | 601198.19 | 859353.94 | 2 | 2 | 2 | 2 | 668839.5 | 2 | 2 | 2 | 578092.69 | 710789.81 | 669371.75 | 928745.62 | 838921.44 | 327145.22 | 2 | 2 | 2 | 2 | 5 | 9 | 1 | 12 | 5 | 12 | 2.551025689 | 5.570658947 | 0.22399192 | 0.489129764 |
| O35972 | Mrpl23 | 39S ribosomal protein L23, mitochondrial | 47.08 | 13.01 | 17122 | 1 | 2 | 2 | 2 | 2 | 2 | 160500.58 | 2 | 2 | 2 | 2 | 296768.81 | 2 | 2 | 380262.59 | 333383.97 | 2 | 764116.25 | 2 | 2 | 2 | 5 | 10 | 6 | 12 | 6 | 12 | 101041.937 | 6.295111264 | 76412.425 | 4.76064426 |
| Q9D1B9 | Mrpl28 | 39S ribosomal protein L28, mitochondrial | 44.19 | 6.61 | 30170 | 1 | 2 | 2 | 2 | 2 | 2 | 2 | 2 | 2 | 2 | 2 | 176487.12 | 167479.36 | 2 | 2 | 2 | 2 | 2 | 2 | 2 | 2 | 7.5 | 12.5 | 7.5 | 12.5 | 7.5 | 12.5 | 34397.248 | 34397.248 | 1 | 1 |
| P18665 | Mrpl3 | 39S ribosomal protein L3, mitochondrial | 58.25 | 2.02 | 38253 | 1 | 485876.78 | 2 | 470934.88 | 502678.47 | 294553 | 488074.41 | 302611.22 | 2 | 433764.16 | 641439.94 | 866696.38 | 585337 | 812248 | 759058.5 | 878671.25 | 422229.16 | 2 | 581130.75 | 453622.38 | 537530.75 | 0 | 10.5 | 1 | 11.5 | 1 | 11.5 | 2.224578526 | 2.091231269 | 1.137094483 | 1.068933962 |
| Q9JKF7 | Mrpl39 | 39S ribosomal protein L39, mitochondrial | 243.91 | 15.77 | 38549 | 4 | 2460460.75 | 2346654.75 | 2275420.25 | 1638269.5 | 2921566.75 | 2564582.5 | 1839895.5 | 2098618.25 | 2597379.75 | 1957933.88 | 2812433.5 | 2087672.5 | 2845595.25 | 3043019 | 2851413.25 | 2499360.75 | 1701395.5 | 2287375.25 | 2252377.75 | 2601900 | 7 | 12 | 3 | 11 | 7 | 12 | 1.171594027 | 1.233462464 | 0.974235255 | 1.025681755 |
| Q9CY73 | Mrpl44 | 39S ribosomal protein L44, mitochondrial | 222.87 | 17.42 | 37527 | 4 | 1546855.88 | 1363853 | 1949733 | 2149149 | 1133177.75 | 1416111.88 | 1365736 | 1962573 | 2177886.25 | 1159615.25 | 381563.78 | 1059377.5 | 1085884.62 | 961589.62 | 1111026.75 | 1268143.75 | 796855.38 | 1416728.88 | 1459449.5 | 1112039.25 | 0 | 5 | 0 | 7 | 0 | 7 | 0.564849927 | 0.569102505 | 0.743385577 | 0.748982293 |
| Q9DCA2 | Mrps11 | 28S ribosomal protein S11, mitochondrial | 47.41 | 8.9 | 20208 | 1 | 1039372.81 | 1034159.44 | 1481002.75 | 2 | 1091703 | 961168.56 | 1161697.5 | 1307068.38 | 950987.62 | 1057743.75 | 2 | 934747.31 | 1258125.25 | 718750.62 | 707268 | 1570917.88 | 1560262.38 | 977719.31 | 1318895.62 | 1296340.5 | 7.5 | 6 | 4 | 4 | 7.5 | 6 | 0.778886407 | 0.66540091 | 1.447220912 | 1.236357578 |
| Q9CPX7 | Mrps16 | 28S ribosomal protein S16, mitochondrial | 41.88 | 5.93 | 15192 | 1 | 2167735.75 | 1350638 | 2008585.75 | 1460851.62 | 1727561.5 | 1868033.75 | 1886984.75 | 1523566.5 | 2042648.12 | 1603775.88 | 893685.06 | 1413385.12 | 2248836.25 | 1086720.75 | 1811201.62 | 1749886.12 | 1174133.38 | 1799005.25 | 1323099 | 1163266.62 | 9 | 6 | 7 | 4 | 9 | 6 | 0.855250731 | 0.835162049 | 0.827203917 | 0.807774017 |
| Q9CQE3 | Mrps17 | 28S ribosomal protein S17, mitochondrial | 48.21 | 7.5 | 13382 | 1 | 1218650.38 | 996930.5 | 1328337.38 | 1336853.75 | 1420929.25 | 1583339.88 | 961908.06 | 792171.5 | 1236732.12 | 923480.31 | 1119390.25 | 1117841.62 | 1283205.12 | 1144472 | 1307824.25 | 693069 | 1115364.88 | 1128473 | 1018235.06 | 1246307.25 | 7 | 5 | 8 | 12 | 8 | 12 | 0.947796951 | 1.086419277 | 0.825403962 | 0.946125407 |
| Q924T2 | Mrps2 | 28S ribosomal protein S2, mitochondrial | 179.98 | 15.81 | 32313 | 4 | 1259922.62 | 1109058.5 | 1778441.5 | 1642246.12 | 2151082.75 | 1827869.75 | 1106011.12 | 1157782.5 | 1124635.5 | 1780382.25 | 569720.38 | 746277.12 | 1266550.75 | 1157752.12 | 73204.35 | 1875214.75 | 1774523 | 963547.25 | 426749.09 | 1651700.75 | 3 | 10 | 5 | 11 | 5 | 11 | 0.480244814 | 0.545044808 | 0.842708004 | 0.956415581 |
| P58059 | Mrps21 | 28S ribosomal protein S21, mitochondrial | 88.08 | 14.94 | 10561 | 1 | 857617.88 | 1786306 | 2 | 743022.06 | 846326.81 | 1015742.06 | 739743.81 | 715770.62 | 871399.25 | 893198.94 | 1013337.38 | 996648.12 | 1090350.38 | 2 | 949920.62 | 564955.12 | 2 | 1059166.25 | 2 | 2 | 8.5 | 6.5 | 8 | 5 | 8.5 | 6.5 | 0.956767217 | 0.956184479 | 0.383657444 | 0.38342377 |
| Q9CXW2 | Mrps22 | 28S ribosomal protein S22, mitochondrial | 102.97 | 5.57 | 41192 | 2 | 1432670.5 | 2 | 980578.94 | 663971.75 | 1040604.81 | 1562391 | 1356095.62 | 823844.62 | 923120.38 | 1332231.62 | 2272933.75 | 1908351.38 | 2098252.5 | 2580128.25 | 993974.69 | 289212.09 | 1689097.25 | 1369809.75 | 1347517.88 | 1105788.5 | 2 | 7 | 3 | 11 | 3 | 11 | 2.392921844 | 1.642907799 | 1.408855705 | 0.967277737 |
| Q8VE22 | Mrps23 | 28S ribosomal protein S23, mitochondrial | 89.29 | 13.56 | 20348 | 2 | 2 | 2 | 2 | 384104.03 | 2 | 2 | 2 | 2 | 2 | 2 | 2 | 2 | 2 | 2 | 2 | 2 | 2 | 2 | 208820 | 238051.56 | 10 | 11 | 12.5 | 7.5 | 12.5 | 11 | 2.603407E-05 | 1 | 1.163404229 | 44687.756 |
| Q9CQV5 | Mrps24 | 28S ribosomal protein S24, mitochondrial | 48.83 | 7.19 | 18901 | 1 | 576283.25 | 413658.75 | 211829.38 | 241552.62 | 278290.66 | 549568.75 | 542355.44 | 242784.16 | 232469.8 | 177256.91 | 675419.62 | 420964.31 | 242161.16 | 331519.38 | 341535.56 | 274909.28 | 2 | 2 | 2 | 139422.3 | 8 | 2 | 9 | 3 | 9 | 3 | 1.168438023 | 1.153152718 | 0.240668013 | 0.237519636 |
| Q4QR80 | Mrps25 | 28S ribosomal protein S25, mitochondrial | 90.33 | 7.6 | 19775 | 2 | 743944.38 | 462770.78 | 688619.12 | 676542 | 606813.25 | 565961.38 | 666697.62 | 539147.88 | 572212.19 | 678821.69 | 796881.38 | 901585.31 | 1085784.12 | 2 | 697514.88 | 549405 | 828032.12 | 717224.44 | 316381 | 579440.69 | 6 | 11 | 5 | 11 | 6 | 11 | 1.09534689 | 1.151819751 | 0.940791235 | 0.989295662 |
| Q8BK72 | Mrps27 | 28S ribosomal protein S27, mitochondrial | 107.52 | 5.54 | 47779 | 2 | 411305.19 | 143172.17 | 2 | 487775.94 | 913429.19 | 159153.48 | 317404.69 | 624823.75 | 660977.81 | 667672.75 | 503114.06 | 254827.39 | 471942.25 | 276060.09 | 1227178.88 | 310968.97 | 704456.38 | 361261.12 | 895738.19 | 344265.19 | 9 | 11 | 11 | 10 | 11 | 11 | 1.397527405 | 1.124726806 | 1.33799182 | 1.076812706 |
| Q9CY16 | Mrps28 | 28S ribosomal protein S28, mitochondrial | 178.65 | 19.35 | 20520 | 3 | 2092063.62 | 1388579.5 | 2030206.5 | 1824642.62 | 1946840.25 | 2055561.88 | 1628473.75 | 26410.91 | 1587176.38 | 2125883.75 | 1713453.38 | 1799485.5 | 2220242.25 | 2196858 | 2279620.5 | 1293389.62 | 1954137.38 | 2117970.75 | 2131811.75 | 1828313.62 | 8 | 10 | 4 | 9 | 8 | 10 | 1.099902383 | 1.37531494 | 1.004663766 | 1.256228833 |
| Q9D0G0 | Mrps30 | 28S ribosomal protein S30, mitochondrial | 97.82 | 5.66 | 49939 | 2 | 497237.22 | 623024.69 | 405072.41 | 368091.28 | 782877.38 | 2 | 542207.31 | 248143.45 | 778093.5 | 272299.41 | 274946.38 | 687222.62 | 558615.06 | 2 | 638871 | 561033.69 | 283225.31 | 319913.75 | 309849.91 | 190398.3 | 11 | 3 | 9.5 | 11 | 11 | 11 | 0.806955369 | 1.173251197 | 0.621910513 | 0.904210173 |
| Q9JIK9 | Mrps34 | 28S ribosomal protein S34, mitochondrial | 128.94 | 10.09 | 25827 | 3 | 1581247.5 | 2026168.88 | 1500049 | 1047059.31 | 1686311.12 | 1502554 | 1234802.75 | 1964035.62 | 2181164.25 | 2037064.25 | 3358026.25 | 3326065.5 | 4793346 | 4078818 | 3598151 | 1744342.25 | 800204.69 | 1914582 | 1969660.5 | 1035142.38 | 0 | 12 | 0 | 7 | 0 | 12 | 2.442903692 | 2.147446291 | 0.951930636 | 0.836799224 |
| Q8BJZ4 | Mrps35 | 28S ribosomal protein S35, mitochondrial | 67.06 | 4.69 | 35975 | 1 | 2 | 2 | 2 | 2 | 2 | 2 | 2 | 2 | 212916.36 | 2 | 2 | 2 | 2 | 2 | 2 | 2 | 2 | 2 | 2 | 2 | 12.5 | 12.5 | 10 | 10 | 12.5 | 12.5 | 1 | 4.696503E-05 | 1 | 4.696503E-05 |
| Q5I0K8 | Mrps7 | 28S ribosomal protein S7, mitochondrial | 92.78 | 4.96 | 28197 | 1 | 649761.69 | 593133.25 | 627844.31 | 844344.81 | 648851.56 | 708398.19 | 697814.94 | 759827.19 | 598319 | 2 | 2 | 598600.69 | 561840.69 | 2 | 364971.62 | 654105.94 | 601667.69 | 596227.75 | 410087.72 | 806711.31 | 1 | 10 | 5 | 11 | 5 | 11 | 0.453462008 | 0.551815347 | 0.912264905 | 1.110129992 |
| Q9D0I8 | Mrto4 | mRNA turnover protein 4 homolog | 413.3 | 30.54 | 27546 | 9 | 8547096 | 8747208 | 5420332 | 7913051 | 5997416.5 | 8521706 | 6975509.5 | 2803638.75 | 3584787.75 | 5305250 | 7432733.5 | 6925385 | 5063920.5 | 2951439 | 4423258.5 | 4386269 | 8608096 | 5286272.5 | 6032620.5 | 6693707.5 | 4 | 8 | 12 | 10 | 12 | 10 | 0.731649441 | 0.985504135 | 0.846604174 | 1.140343814 |
| P43247 | Msh2 | DNA mismatch repair protein Msh2 | 506.51 | 11.76 | 104151 | 9 | 6151122 | 5423134 | 7880307.5 | 8092073 | 6044809 | 4218181.5 | 6574281.5 | 5102674 | 5259288.5 | 5883686 | 5677365 | 5917701 | 6043555 | 4750814 | 5303222.5 | 6843381 | 6899958.5 | 5406289.5 | 4866287.5 | 6278633.5 | 3 | 9 | 10 | 7 | 10 | 9 | 0.824396125 | 1.024208274 | 0.901853122 | 1.120438829 |
| P54276 | Msh6 | DNA mismatch repair protein Msh6 | 271.13 | 5.08 | 151084 | 5 | 919247.56 | 2215151.5 | 476737.12 | 3154464 | 2239421.25 | 2151199.75 | 2483877 | 1714798.75 | 1410273.62 | 1581373.38 | 1257041.25 | 1247529.25 | 1730045 | 970037.88 | 1684675.62 | 2145818.75 | 2020358.5 | 2442156.5 | 2118253.5 | 1705268.88 | 10 | 12 | 5 | 10 | 10 | 12 | 0.765054148 | 0.737495307 | 1.158448785 | 1.11671905 |
| P26041 | Msn | Moesin | 3630.031 | 69.67 | 67767 | 56 | 212082258 | 202561842 | 231815666 | 179409346 | 211765602 | 210067010 | 230542722 | 246007042 | 273241986 | 287832066 | 297085922 | 334973762 | 285691010 | 298421474 | 316696130 | 192055986 | 232557810 | 306368034 | 291327682 | 292335138 | 0 | 4 | 1 | 8 | 1 | 8 | 1.477271604 | 1.228564213 | 1.266962865 | 1.053662191 |
| P30204 | Msr1 | Macrophage scavenger receptor types I and II | 355.25 | 13.97 | 50170 | 6 | 17479082 | 18735816 | 15746289 | 14879701 | 14074824 | 15840150 | 18270878 | 18315762 | 18400448 | 17014930 | 14053031 | 14185316 | 17897474 | 12365195 | 19839926 | 16276514 | 11617932 | 18640778 | 19608644 | 17661556 | 10 | 9 | 7 | 12 | 10 | 12 | 0.968179604 | 0.891837528 | 1.035712619 | 0.954045488 |
| Q9D6Y7 | Msra | Mitochondrial peptide methionine sulfoxide reductase | 70.49 | 6.87 | 25988 | 1 | 533003.75 | 703331.75 | 1049644.62 | 1168381.62 | 774463.56 | 696883.44 | 1009367.25 | 1343919 | 1204822.12 | 1179339.62 | 2 | 723431.62 | 2 | 2 | 498094.72 | 686820.5 | 2 | 1426450 | 1126839.25 | 983007.5 | 2 | 12 | 1 | 8 | 2 | 12 | 0.288858549 | 0.224780611 | 0.998650677 | 0.777118456 |
| Q2YDW2 | Msto1 | Protein misato homolog 1 | 125.65 | 5.22 | 61230 | 3 | 1978422.25 | 1808277.12 | 978986 | 444342.72 | 1286586.62 | 2067710.12 | 2323445.5 | 1829373.12 | 1366246.75 | 2264531.5 | 1360887.25 | 1848439 | 746664.38 | 2183162.5 | 2 | 2191344.5 | 1712569.25 | 1873215.62 | 1311086.25 | 2179789.75 | 12 | 5 | 5 | 9 | 12 | 9 | 0.944977562 | 0.623181791 | 1.426589968 | 0.940789418 |
| P02802 | Mt1 | Metallothionein-1 | 148.29 | 21.31 | 6018 | 2 | 1606813 | 3580264.5 | 2 | 462210.78 | 100200.93 | 2717617.5 | 2668435.5 | 97178.15 | 591791.06 | 603975.81 | 1681653.75 | 1490950.25 | 1987506.25 | 2531407.25 | 1374914.88 | 1026647.69 | 907057.56 | 873627.75 | 2 | 752606.81 | 7 | 12.5 | 10 | 12 | 10 | 12.5 | 1.576910382 | 1.35745397 | 0.619175103 | 0.53300537 |
| P02798 | Mt2 | Metallothionein-2 | 133.78 | 21.31 | 6115 | 2 | 939969.25 | 1768297.12 | 2 | 2 | 2 | 1488730.5 | 1874345.25 | 2 | 2 | 2 | 1749643.62 | 1398071.62 | 1884817.75 | 2990068.75 | 1444632.38 | 642856.31 | 801594.69 | 690459.5 | 2 | 2 | 3 | 12 | 5 | 12 | 5 | 12 | 3.495672823 | 2.815047276 | 0.788293867 | 0.634808981 |
| Q62599 | Mta1 | Metastasis-associated protein MTA1 | 141.87 | 5.69 | 79412 | 3 | 2 | 514574.94 | 2 | 2 | 333790.84 | 2 | 505429.88 | 568968.75 | 129462.03 | 604590.56 | 510808.31 | 471083.44 | 2 | 2 | 2 | 2 | 2 | 519700.34 | 2 | 2 | 12.5 | 11 | 6.5 | 5 | 12.5 | 11 | 1.157390867 | 0.542948935 | 0.612595035 | 0.287377265 |
| Q9R190 | Mta2 | Metastasis-associated protein MTA2 | 596.9 | 18.41 | 75030 | 12 | 9114256 | 10587944 | 10342166 | 8767436 | 8651210 | 10482642 | 8843768 | 9357658 | 8717441 | 10269779 | 10567941 | 11545380 | 11980942 | 11624290 | 12756540 | 8480835 | 9086683 | 11329690 | 11254962 | 11159037 | 1 | 8 | 0 | 8 | 1 | 8 | 1.232013952 | 1.22663128 | 1.081077766 | 1.076354534 |
| Q9CQ65 | Mtap | S-methyl-5'-thioadenosine phosphorylase | 575.17 | 39.22 | 31062 | 9 | 16582704 | 14896417 | 29206642 | 30538352 | 27405458 | 19915270 | 19103738 | 28390358 | 29362328 | 27730930 | 15487729 | 15512173 | 20228844 | 19793230 | 18796050 | 19423846 | 23424246 | 29001650 | 29829654 | 27098454 | 8 | 12 | 3 | 11 | 8 | 12 | 0.757130147 | 0.721414723 | 1.085545929 | 1.034338441 |
| Q791T5 | Mtch1 | Mitochondrial carrier homolog 1 | 348.46 | 17.99 | 41565 | 7 | 6694270 | 7974040 | 5023342 | 8308722.5 | 4891427 | 9478280 | 6723402 | 5860644 | 6209628 | 8838671 | 6676204.5 | 2320244 | 8938193 | 10047419 | 6131309 | 7871844 | 5213332 | 9214749 | 9389104 | 8460928 | 11 | 5 | 12 | 11 | 12 | 11 | 1.037138981 | 0.919234572 | 1.220667618 | 1.08189924 |
| Q791V5 | Mtch2 | Mitochondrial carrier homolog 2 | 493.73 | 40.26 | 33499 | 9 | 21300666 | 25145198 | 17748042 | 18043398 | 24022332 | 22520314 | 22030334 | 19314234 | 27092348 | 25325352 | 28126458 | 30675712 | 28062624 | 30267702 | 30451634 | 19836344 | 26478974 | 28391534 | 25928412 | 24475570 | 0 | 4 | 0 | 8 | 0 | 8 | 1.388901144 | 1.269185182 | 1.177406951 | 1.075920674 |
| P00397 | Mtco1 | Cytochrome c oxidase subunit 1 | 78.88 | 3.31 | 56910 | 1 | 2 | 2 | 2 | 2 | 2 | 2 | 2 | 556596.75 | 1129433.25 | 1025417.81 | 2 | 2 | 659826.69 | 796806.44 | 597491.12 | 1655145.62 | 2 | 2 | 2 | 741973.75 | 5 | 7.5 | 11 | 11 | 11 | 11 | 205412.825 | 0.757575053 | 239712.537 | 0.884074488 |
| P00405 | Mtco2 | Cytochrome c oxidase subunit 2 | 482.86 | 35.24 | 25976 | 9 | 52670198 | 50879042 | 59135594 | 51120298 | 40771090 | 46469650 | 41628434 | 43951378 | 46546890 | 40509962 | 51321706 | 44968258 | 61718258 | 58499442 | 26737208 | 46141898 | 43590818 | 56079870 | 51043490 | 48057946 | 12 | 9 | 7 | 5 | 12 | 9 | 0.955489362 | 1.110168245 | 0.962045945 | 1.117786236 |
| Q80WJ7 | Mtdh | Protein LYRIC | 386.73 | 15.37 | 63846 | 7 | 11371255 | 10902990 | 7999061 | 8488142 | 10393247 | 10891966 | 12708406 | 8751735 | 10321967 | 9233527 | 9009449 | 8815541 | 9553426 | 8293723 | 9898026 | 10971341 | 12410008 | 10431958 | 11642792 | 9530461 | 9 | 6 | 6 | 8 | 9 | 8 | 0.927076549 | 0.877909287 | 1.118643092 | 1.059316149 |
| Q922D8 | Mthfd1 | C-1-tetrahydrofolate synthase, cytoplasmic | 1185.92 | 25.45 | 101200 | 21 | 30355680 | 29947660 | 31270516 | 27828498 | 26935012 | 29494424 | 33881274 | 27157516 | 27633614 | 28065208 | 29885166 | 29194298 | 26422708 | 22218728 | 25452508 | 31828876 | 33651638 | 29269500 | 27601046 | 25752816 | 4 | 12 | 7 | 12 | 7 | 12 | 0.910043768 | 0.910699267 | 1.01207149 | 1.012800478 |
| Q3V3R1 | Mthfd1l | Monofunctional C1-tetrahydrofolate synthase, mitochondrial | 1816.75 | 34.9 | 105729 | 29 | 34114478 | 37408666 | 32894768 | 34055546 | 32804156 | 35110234 | 38487974 | 31404156 | 32721862 | 35592746 | 33676058 | 40818354 | 38502518 | 32615748 | 39216446 | 38757366 | 28154334 | 34349830 | 31709534 | 34335622 | 8 | 12 | 7 | 10 | 8 | 12 | 1.079120147 | 1.066422531 | 0.976815838 | 0.965322 |
| P18155 | Mthfd2 | Bifunctional methylenetetrahydrofolate dehydrogenase/cyclohydrolase, mitochondrial | 716.33 | 46.86 | 37863 | 13 | 2876670.25 | 3662915.5 | 2877633.5 | 3169769.75 | 3125291.25 | 2438498.5 | 3754124.5 | 3133455.5 | 3870431.5 | 3133534.25 | 8668416 | 9149901 | 8635173 | 9915655 | 10729822 | 3790952.75 | 2160352 | 3966437 | 4080383.25 | 4658974 | 0 | 5 | 0 | 6 | 0 | 6 | 2.997589545 | 2.884191021 | 1.187421476 | 1.142501436 |
| Q8VEL2 | Mtmr14 | Myotubularin-related protein 14 | 43.01 | 1.39 | 72443 | 1 | 595282.62 | 378947.41 | 420663.53 | 585486.75 | 523047.97 | 616517.06 | 467709.44 | 523714.72 | 576818.38 | 427866.47 | 261782.48 | 181290.53 | 249701.34 | 2 | 200625.33 | 555671.06 | 430959.16 | 452093.88 | 489862.62 | 352665.31 | 0 | 9 | 0 | 7 | 0 | 9 | 0.35687129 | 0.341955433 | 0.911251202 | 0.873164383 |
| P03911 | Mtnd4 | NADH-ubiquinone oxidoreductase chain 4 | 100.65 | 6.32 | 51882 | 2 | 727358.81 | 662712.5 | 853325.75 | 877518 | 984943.25 | 969015.81 | 1122449.38 | 926702.75 | 1108312.25 | 990572.19 | 507344.78 | 803674.12 | 1489407.88 | 1037393.12 | 1208384.88 | 678634.75 | 512274.75 | 1143686.88 | 853945.31 | 1161952.25 | 8 | 11 | 12 | 10 | 12 | 11 | 1.229025553 | 0.986154607 | 1.059582093 | 0.850195311 |
| P03921 | Mtnd5 | NADH-ubiquinone oxidoreductase chain 5 | 180.86 | 6.59 | 68475 | 3 | 788216.38 | 808854.81 | 1166704.62 | 771250.69 | 900871.44 | 843731 | 574271.25 | 946179.5 | 802878.06 | 1055978.62 | 1517394.38 | 1689039.75 | 1978260.12 | 2174311.5 | 1775105.5 | 218060.91 | 700296.88 | 948975.75 | 649295.81 | 948953.06 | 0 | 8 | 0 | 10 | 0 | 10 | 2.059134672 | 2.162924018 | 0.781258374 | 0.820637195 |
| P42346 | Mtor | Serine/threonine-protein kinase mTOR | 59.33 | 0.59 | 288794 | 1 | 2 | 2 | 237756.03 | 472569.97 | 360499.91 | 2 | 2 | 2 | 405184.84 | 367488.94 | 2 | 2 | 273702.81 | 2 | 368114.56 | 2 | 2 | 2 | 481423.22 | 268935.81 | 10 | 11 | 11.5 | 12.5 | 11.5 | 12.5 | 0.599370044 | 0.830646002 | 0.700732229 | 0.971120313 |
| P62774 | Mtpn | Myotrophin | 253.8 | 43.22 | 12861 | 4 | 16567104 | 19833034 | 21114356 | 16547071 | 20729234 | 10546610 | 15400051 | 19306180 | 17974042 | 16453684 | 10315752 | 16498966 | 9747111 | 9267369 | 13473707 | 18261988 | 6372816 | 15768122 | 11765191 | 21072496 | 0 | 6 | 4 | 12 | 4 | 12 | 0.62561879 | 0.74425807 | 0.772655297 | 0.919177859 |
| Q9CZU3 | Mtrex | Exosome RNA helicase MTR4 | 331.33 | 6.83 | 117636 | 6 | 3652209 | 4046258 | 3657100.75 | 3074822.25 | 4169899.5 | 3525175 | 2461441.5 | 3653002.75 | 4500285 | 2825856.25 | 1827750.75 | 1645807.38 | 2378964.5 | 2351225 | 2418617 | 3915123 | 2359599.75 | 4000069.5 | 4574835 | 4057627.5 | 0 | 10 | 0 | 8 | 0 | 10 | 0.571085984 | 0.626106011 | 1.016503251 | 1.114436028 |
| Q8R1S4 | Mtss1 | Metastasis suppressor protein 1 | 508.1 | 18.97 | 82408 | 9 | 8323215.5 | 10525482 | 7526187.5 | 10848206 | 9491664 | 14054256 | 10488198 | 13188461 | 11780069 | 14805666 | 7232143.5 | 7342560 | 8209255 | 8412784 | 8045704.5 | 8120439 | 11035094 | 14044684 | 11076789 | 13812344 | 4 | 4 | 0 | 8 | 4 | 8 | 0.840043943 | 0.610144449 | 1.243490413 | 0.903177482 |
| P47802 | Mtx1 | Metaxin-1 | 225.04 | 20.5 | 35624 | 5 | 2760950.5 | 2822076 | 3361507.25 | 3036697 | 2223730.5 | 2670589.75 | 2658944.25 | 2695401 | 2462921.5 | 3662985.75 | 3765784.5 | 3066282.5 | 3837142.25 | 2922647.75 | 2742889 | 3026310 | 1648021.88 | 4277584.5 | 4219654.5 | 3315837.25 | 7 | 8 | 3 | 7 | 7 | 8 | 1.149932458 | 1.154330301 | 1.160679557 | 1.165118502 |
| O88441 | Mtx2 | Metaxin-2 | 147.63 | 22.81 | 29758 | 3 | 686742.75 | 1263067.25 | 1338268.25 | 1741145.25 | 1456249.12 | 1196182.75 | 1268689.25 | 1488211.88 | 1242029.62 | 885831.5 | 1125329.5 | 1331897.38 | 1278211.5 | 1602057.75 | 1443614.75 | 1826575.75 | 798432.69 | 1632698 | 863107.19 | 2730898.5 | 12 | 9 | 7 | 10 | 12 | 10 | 1.04558469 | 1.115140966 | 1.210661518 | 1.291199333 |
| P16332 | Mut | Methylmalonyl-CoA mutase, mitochondrial | 406.29 | 11.63 | 82844 | 7 | 6037114 | 5239626 | 4553507 | 4032908 | 4901558.5 | 4381038 | 5748188 | 4429100 | 4533128 | 5000619 | 4270117 | 4326331 | 4553665.5 | 4008043.25 | 5429440.5 | 5823469 | 4398341.5 | 6224209 | 4807756 | 4783542 | 8 | 11 | 7 | 8 | 8 | 11 | 0.912087969 | 0.93755308 | 1.051387794 | 1.080742097 |
| Q78HU3 | Mvb12a | Multivesicular body subunit 12A | 102.32 | 11.81 | 28705 | 2 | 721360.75 | 728513.31 | 2 | 1869140.25 | 563720.12 | 583130.12 | 903357.06 | 2 | 2 | 1614220.75 | 1479633.5 | 1078146.62 | 3742932.25 | 1969339.38 | 2 | 501763.66 | 2 | 1228005.12 | 1019851.44 | 964242.44 | 6.5 | 11.5 | 6 | 10 | 6.5 | 11.5 | 2.129954968 | 2.667146751 | 0.956507022 | 1.19774579 |
| Q99JF5 | Mvd | Diphosphomevalonate decarboxylase | 388.02 | 21.2 | 44072 | 6 | 6914874 | 7017275.5 | 4724896.5 | 2260423 | 4654891.5 | 5216577 | 4897138 | 3397496.75 | 3704801.75 | 3804400.25 | 6122064 | 5479815.5 | 4703689.5 | 3940254.25 | 5836676 | 5425253.5 | 2815924.5 | 3132969.5 | 5238613 | 6213709 | 12 | 11 | 4 | 10 | 12 | 11 | 1.019948833 | 1.240817596 | 0.892622701 | 1.085919134 |
| Q9R008 | Mvk | Mevalonate kinase | 102.11 | 9.37 | 41877 | 2 | 181997.05 | 181643.23 | 2 | 106314.59 | 2 | 224314.23 | 2 | 2 | 2 | 2 | 2 | 386842.84 | 330708.75 | 408854.09 | 349536 | 158941.56 | 63724.43 | 2 | 148398.45 | 2 | 4 | 10 | 3 | 9 | 4 | 10 | 3.140580536 | 6.579569399 | 0.789576415 | 1.654175959 |
| Q9EQK5 | Mvp | Major vault protein | 1734.31 | 46.69 | 95924 | 29 | 20907910 | 25496224 | 30619960 | 32573696 | 24674756 | 26903444 | 23731840 | 31245976 | 29695352 | 36566502 | 38150878 | 36390578 | 40536630 | 43395038 | 43257378 | 19514982 | 27271498 | 32315004 | 30476170 | 35211850 | 0 | 10 | 1 | 12 | 1 | 12 | 1.502395747 | 1.361727161 | 1.078325453 | 0.977362363 |
| O35682 | Myadm | Myeloid-associated differentiation marker | 362.94 | 25.94 | 35285 | 5 | 11371526 | 11736671 | 12941848 | 18019108 | 20658002 | 17107512 | 17793042 | 17270484 | 17420076 | 20921466 | 20739502 | 17532592 | 27593568 | 20255142 | 34417770 | 15411862 | 20322880 | 25326150 | 26727200 | 25257402 | 3 | 3 | 4 | 6 | 4 | 6 | 1.613049152 | 1.331732827 | 1.512776634 | 1.24894787 |
| Q7TPV4 | Mybbp1a | Myb-binding protein 1A | 1961.03 | 29.39 | 152037 | 30 | 16669950 | 16689174 | 10543772 | 15530982 | 18256520 | 17692290 | 16958520 | 15515895 | 15362147 | 17942284 | 15470809 | 19055898 | 24014454 | 18311890 | 22762764 | 22897156 | 19462508 | 17410934 | 19226136 | 20861750 | 4 | 1 | 4 | 2 | 4 | 2 | 1.282215275 | 1.193416309 | 1.285338814 | 1.196323529 |
| P70402 | Mybph | Myosin-binding protein H | 46.16 | 1.66 | 52641 | 1 | 330678.78 | 309196.12 | 2 | 518573.69 | 230246.08 | 2 | 2 | 536965.06 | 606969.44 | 2 | 1463187.25 | 710325.44 | 556520.56 | 1088478.12 | 536905.75 | 409521.53 | 245297.27 | 600999.75 | 2 | 678561.94 | 0 | 8.5 | 1 | 5 | 1 | 8.5 | 3.136334388 | 8.111053318 | 1.392948173 | 3.602382753 |
| P22366 | Myd88 | Myeloid differentiation primary response protein MyD88 | 56.07 | 5.07 | 33753 | 1 | 406363.78 | 432087.38 | 427746.03 | 174346.62 | 69578.01 | 156431.3 | 430432.09 | 393068.19 | 606969.44 | 141632.77 | 46118.91 | 411710.34 | 127861.98 | 2 | 2 | 397319.44 | 2 | 2 | 2 | 136042.77 | 4 | 3 | 3 | 3 | 4 | 3 | 0.387846346 | 0.338839329 | 0.353195486 | 0.30856684 |
| Q9CPT4 | Mydgf | Myeloid-derived growth factor | 212.84 | 24.1 | 17982 | 4 | 5492793 | 5809749 | 6599282 | 6841732 | 7356390.5 | 6141945 | 5782829.5 | 6167854 | 7519851 | 5625124 | 4247361.5 | 4604795.5 | 6137601 | 5300976 | 5746564 | 5399161.5 | 6084460 | 7122709 | 6740387.5 | 6790018 | 3 | 12 | 3 | 11 | 3 | 12 | 0.811132131 | 0.833524185 | 1.001146092 | 1.028783658 |
| Q8C854 | Myef2 | Myelin expression factor 2 | 734.56 | 32.15 | 63295 | 14 | 13424451 | 15853581 | 13219232 | 14956232 | 13216245 | 17597486 | 16162215 | 13839559 | 14167080 | 14077000 | 14543462 | 14893474 | 14377119 | 15352826 | 14081140 | 16512534 | 17421714 | 15674640 | 15046261 | 15700504 | 9 | 3 | 11 | 8 | 11 | 8 | 1.036483507 | 0.965780529 | 1.13705883 | 1.059495178 |
| Q9JK81 | Myg1 | UPF0160 protein MYG1, mitochondrial | 369.88 | 19.47 | 42723 | 7 | 8160355 | 8206388 | 8268758 | 8292585.5 | 8954494 | 8508474 | 8506411 | 7575559.5 | 9184911 | 9312169 | 9020308 | 7586060.5 | 6620637 | 9071594 | 5879706 | 9504076 | 11756131 | 9371294 | 11114906 | 9337223 | 10 | 0 | 7 | 0 | 10 | 0 | 0.911555712 | 0.886064028 | 1.219686786 | 1.185578206 |
| Q61879 | Myh10 | Myosin-10 | 1148.12 | 10.98 | 228996 | 19 | 1608561.5 | 2316205.25 | 4848067.5 | 3926900 | 2938985.5 | 1769290.38 | 2459663.25 | 3561705.5 | 3345990.5 | 1544438.5 | 2 | 245584.28 | 695842.69 | 262588.81 | 612966.62 | 2145310.5 | 1269062.25 | 3043863.5 | 1823206 | 2124555.75 | 0 | 6 | 0 | 9 | 0 | 9 | 0.116184984 | 0.143283004 | 0.66538682 | 0.820576113 |
| A2AQP0 | Myh7b | Myosin-7B | 254.24 | 1.6 | 221497 | 4 | 2 | 2 | 2 | 2 | 373939.31 | 2 | 908053 | 16453760 | 2 | 2 | 2 | 451685.69 | 2 | 2 | 2 | 360413.69 | 2 | 444529.84 | 546972.38 | 2 | 12 | 7 | 9 | 12 | 12 | 12 | 1.207907312 | 0.026016496 | 3.615268445 | 0.077867412 |
| Q8VDD5 | Myh9 | Myosin-9 | 8403.48 | 61.17 | 226372 | 124 | 349854338 | 412488162 | 393635746 | 388007842 | 376828866 | 353531650 | 334326722 | 375231234 | 367851554 | 368932642 | 274189794 | 286024194 | 297944898 | 278640930 | 292438338 | 351493762 | 376599330 | 340724418 | 332274530 | 335277474 | 0 | 2 | 0 | 8 | 0 | 8 | 0.744079044 | 0.794076869 | 0.903975425 | 0.964717366 |
| Q60605 | Myl6 | Myosin light polypeptide 6 | 909.01 | 70.2 | 16930 | 13 | 94692122 | 92834690 | 115431330 | 116010706 | 126114410 | 88539938 | 98057562 | 77220050 | 93434250 | 91204386 | 41602066 | 49834198 | 47655006 | 52274834 | 49949530 | 125750546 | 124708362 | 102111746 | 102061178 | 104704714 | 0 | 11 | 0 | 0 | 0 | 11 | 0.442713348 | 0.538103033 | 1.026148827 | 1.247249037 |
| Q9JMH9 | Myo18a | Unconventional myosin-XVIIIa | 306.18 | 3.32 | 232755 | 6 | 1833032.38 | 2722957.5 | 2463028 | 1346114.75 | 2194757.5 | 1891221.75 | 2797802.25 | 7783204.5 | 1223726.88 | 2521355.75 | 1026634.75 | 1864904.88 | 1798452.38 | 1005748.88 | 1339092.75 | 1686164.5 | 1482938 | 661897.38 | 1747538.5 | 532859.75 | 3 | 3 | 3 | 3 | 3 | 3 | 0.666184359 | 0.433785452 | 0.578736905 | 0.376844107 |
| Q9WTI7 | Myo1c | Unconventional myosin-Ic | 1593.76 | 35.94 | 121944 | 28 | 5602371.5 | 5251931.5 | 21738910 | 15716824 | 17583818 | 5712148 | 5532782.5 | 16206963 | 18111658 | 14866722 | 17205094 | 20711856 | 25022342 | 18637328 | 24146550 | 4861390.5 | 5368830 | 11516353 | 12278796 | 9881501 | 4 | 7 | 1 | 6 | 4 | 7 | 1.604446575 | 1.749506727 | 0.666327239 | 0.726570773 |
| E9Q634 | Myo1e | Unconventional myosin-Ie | 1213 | 23.49 | 126818 | 23 | 13590545 | 17948514 | 20626254 | 14940812 | 16070648 | 21725522 | 15311711 | 18986062 | 16528284 | 22190964 | 19910644 | 21627506 | 24023130 | 17640432 | 22440310 | 17725520 | 10703938 | 18823208 | 13507526 | 19902370 | 3 | 11 | 7 | 7 | 7 | 11 | 1.270090413 | 1.115043133 | 0.969772679 | 0.851386921 |
| P70248 | Myo1f | Unconventional myosin-If | 448.65 | 7.83 | 125947 | 8 | 2182253 | 2024930.62 | 2623955.25 | 2901526.75 | 2638791.25 | 2270142.25 | 3140347 | 2342848 | 2946771.75 | 2436571.25 | 1112882.12 | 1110496.62 | 1596375 | 883068.88 | 1191938.38 | 2395669.5 | 2596636.75 | 2083452.62 | 992465.81 | 1733047.75 | 0 | 5 | 0 | 5 | 0 | 5 | 0.476480746 | 0.448725316 | 0.792248846 | 0.746099642 |
| Q5SUA5 | Myo1g | Unconventional myosin-Ig | 282.84 | 10.16 | 117227 | 7 | 1583138.88 | 1254330.75 | 2072999.88 | 2948060 | 2891201.75 | 1256227.25 | 2115472.75 | 1288081.75 | 2651459 | 1790297.5 | 3872099.5 | 4988930 | 5222631 | 5679516 | 3881525.5 | 523252 | 981102.88 | 1533186.25 | 1422024.62 | 1591274 | 0 | 4 | 0 | 6 | 0 | 6 | 2.19956215 | 2.59787976 | 0.562882886 | 0.664815066 |
| Q99104 | Myo5a | Unconventional myosin-Va | 56.18 | 0.86 | 215538 | 1 | 158264.8 | 232549.02 | 199760.5 | 2 | 169530.36 | 220468.92 | 205396.69 | 222155.73 | 249565.78 | 200735.02 | 2 | 2 | 2 | 2 | 2 | 237158.28 | 2 | 2 | 2 | 2 | 2.5 | 7 | 0 | 4 | 2.5 | 7 | 1.315605E-05 | 9.104797E-06 | 0.312017097 | 0.215935081 |
| Q63358 | Myo9b | Unconventional myosin-IXb | 50.77 | 0.66 | 225037 | 1 | 172377.67 | 347436.66 | 220151.8 | 311679.94 | 242535.41 | 220394.45 | 298765.94 | 253691.62 | 316316.22 | 276799.53 | 419232.78 | 292581.97 | 2 | 330602.47 | 296521.06 | 2 | 362890.94 | 386930.16 | 256437.53 | 336108.34 | 10 | 8 | 9 | 8 | 10 | 8 | 1.03458464 | 0.980213676 | 1.037233951 | 0.982723758 |
| Q69ZN7 | Myof | Myoferlin | 1054.14 | 13.48 | 233324 | 21 | 8321451.5 | 10958068 | 3596574 | 5368744.5 | 5078450 | 11108537 | 8338027 | 5623081 | 5964350.5 | 5187147.5 | 21512890 | 18692120 | 10017039 | 8825912 | 10773846 | 6775693 | 9485934 | 2872337 | 7864877.5 | 6405382.5 | 3 | 12 | 3 | 12 | 3 | 12 | 2.095285645 | 1.927653332 | 1.002428812 | 0.92222998 |
| Q80UM3 | Naa15 | N-alpha-acetyltransferase 15, NatA auxiliary subunit | 650.22 | 15.49 | 100961 | 12 | 5904237.5 | 5733097 | 9550415 | 12172425 | 11439087 | 7809004 | 6677889.5 | 10439255 | 13126032 | 10973332 | 10806585 | 10176072 | 12938154 | 10700097 | 9512256 | 6826063 | 11017882 | 11359023 | 11970497 | 9750755 | 9 | 10 | 11 | 10 | 11 | 10 | 1.208349472 | 1.104183541 | 1.136720077 | 1.038728968 |
| P61600 | Naa20 | N-alpha-acetyltransferase 20 | 85.87 | 12.36 | 20368 | 2 | 1433044.88 | 1367009.62 | 981901.38 | 635507.5 | 1075894.75 | 636209.88 | 1424462.12 | 432440.81 | 530880.88 | 602358.69 | 716232.69 | 1138079 | 649683.31 | 476170.75 | 612220.56 | 1151809.75 | 635081.75 | 675674.38 | 594379 | 609631.75 | 5 | 4 | 9 | 9 | 9 | 9 | 0.653950867 | 0.990633544 | 0.667456325 | 1.011092207 |
| Q6QI44 | Naa25 | N-alpha-acetyltransferase 25, NatB auxiliary subunit | 163.99 | 4.23 | 111258 | 3 | 1215751.5 | 1391943.25 | 812320.12 | 756202.31 | 2350930.25 | 587926.69 | 1220128.75 | 717674.25 | 1472604.5 | 677782.94 | 247140.41 | 889141.5 | 734568.81 | 404272.88 | 592002.88 | 1168359.75 | 551486.25 | 502220.66 | 106242.71 | 723780.38 | 2 | 2 | 7 | 6 | 7 | 6 | 0.439261792 | 0.613142571 | 0.467599328 | 0.652697455 |
| Q6DKG0 | Naa35 | N-alpha-acetyltransferase 35, NatC auxiliary subunit | 69.24 | 1.52 | 83210 | 1 | 229327.67 | 143763.89 | 229506.78 | 77257.49 | 257895.64 | 219965.06 | 215769.48 | 198086.42 | 270206.34 | 139424.58 | 105055.81 | 207255.56 | 216150.69 | 2 | 110674.71 | 249553.28 | 59454.45 | 167736.69 | 167001.31 | 2 | 6 | 8 | 5 | 6 | 6 | 8 | 0.681565202 | 0.612523474 | 0.686480108 | 0.616940505 |
| Q8VE10 | Naa40 | N-alpha-acetyltransferase 40 | 69.81 | 5.06 | 27229 | 1 | 233354.12 | 270026.41 | 212490.78 | 199728.66 | 158512.53 | 236911.14 | 267705.16 | 2 | 2 | 2 | 332484.62 | 471946.84 | 2 | 239665.23 | 241900.88 | 185940.53 | 2 | 2 | 2 | 2 | 7 | 1 | 5.5 | 9 | 7 | 9 | 1.197267111 | 2.548439833 | 0.1731183 | 0.368490513 |
| Q6PGB6 | Naa50 | N-alpha-acetyltransferase 50 | 306.88 | 34.91 | 19414 | 6 | 1255665.5 | 1596945 | 2280746.25 | 1634449.5 | 4214901 | 1113858.75 | 1192840.38 | 4354943 | 5155985 | 1686218.38 | 1107584.62 | 3504959 | 1851093.12 | 1506079.88 | 2294125.75 | 1588068.75 | 1495361.5 | 1116327.62 | 2655624 | 2122248.5 | 12 | 9 | 11 | 11 | 12 | 11 | 0.934545749 | 0.760068113 | 0.817433276 | 0.664820281 |
| Q9D7V9 | Naaa | N-acylethanolamine-hydrolyzing acid amidase | 59.3 | 4.7 | 40075 | 1 | 363123.12 | 292916.97 | 290651.59 | 203011.05 | 299853.59 | 311291.69 | 292648.16 | 365120.25 | 346967.34 | 451450.06 | 151250.78 | 133950.78 | 55566.71 | 2 | 106855.7 | 224956.39 | 436417.72 | 350475.59 | 339014.47 | 269554.12 | 0 | 10 | 0 | 9 | 0 | 10 | 0.308802055 | 0.253256955 | 1.117871909 | 0.916797125 |
| P70670 | Naca | Nascent polypeptide-associated complex subunit alpha, muscle-specific form | 543.15 | 3.2 | 220499 | 6 | 22080008 | 24246138 | 20593990 | 12143432 | 13942752 | 18883408 | 21786302 | 15413847 | 15978224 | 15325328 | 31278538 | 33329350 | 23388974 | 20894036 | 25667296 | 17586226 | 13222388 | 15178704 | 11405309 | 15506980 | 3 | 7 | 1 | 5 | 3 | 7 | 1.446763983 | 1.539794548 | 0.783813476 | 0.834214655 |
| Q8VBW6 | Nae1 | NEDD8-activating enzyme E1 regulatory subunit | 271.06 | 14.42 | 60274 | 5 | 3761397 | 5073812 | 4865255 | 5910473 | 5463202 | 4910622 | 3844256.5 | 4906931.5 | 5193037 | 5288710 | 4480383 | 6077305.5 | 5428234.5 | 5036432 | 3997026.75 | 5818143 | 4362585 | 5965639 | 3943287 | 6858270 | 12 | 9 | 10 | 8 | 12 | 9 | 0.997816186 | 1.036275713 | 1.074729784 | 1.116153846 |
| Q9QWR8 | Naga | Alpha-N-acetylgalactosaminidase | 531.69 | 27.95 | 47235 | 9 | 12359932 | 10609127 | 15761688 | 18266666 | 17437730 | 9943582 | 10366070 | 11385565 | 11921500 | 11914598 | 5098747.5 | 3398108.25 | 4276914.5 | 3416956 | 4154580 | 12747640 | 14981079 | 11717961 | 12931074 | 10080004 | 0 | 7 | 0 | 6 | 0 | 7 | 0.27332931 | 0.366375373 | 0.839089649 | 1.124730398 |
| Q9QZ08 | Nagk | N-acetyl-D-glucosamine kinase | 166.73 | 8.16 | 37268 | 3 | 3053377 | 3186105.5 | 2681118.75 | 1833943.88 | 4354012 | 2084491.62 | 2752237.75 | 1615376.38 | 3273012.25 | 2140421.75 | 829849.25 | 846692.56 | 1001813.44 | 1178787.5 | 912425.94 | 2571273.5 | 1997661.62 | 2150252.25 | 1693553.12 | 1690950.62 | 0 | 3 | 0 | 9 | 0 | 9 | 0.315686577 | 0.401968119 | 0.668739644 | 0.851515508 |
| Q8BJ48 | Nagpa | N-acetylglucosamine-1-phosphodiester alpha-N-acetylglucosaminidase | 104.53 | 3.09 | 56044 | 1 | 865868.75 | 763555.88 | 1116115.5 | 1194460.75 | 805849.44 | 1024406.12 | 924592.88 | 881313.94 | 1183488.88 | 957340.31 | 896137.94 | 773687 | 614811.69 | 1085210.88 | 822741.75 | 905174.88 | 1452354.25 | 631199.62 | 791581.81 | 1145485 | 9 | 12 | 5 | 10 | 9 | 12 | 0.883422143 | 0.843385514 | 1.037916333 | 0.990878038 |
| Q99KQ4 | Nampt | Nicotinamide phosphoribosyltransferase | 553.8 | 19.55 | 55447 | 9 | 24966870 | 23684338 | 17995112 | 21239972 | 23078814 | 14537018 | 18200334 | 20695948 | 22370062 | 19502956 | 6452214.5 | 12706066 | 11620621 | 7654008 | 11872511 | 27006566 | 12264037 | 15879142 | 15084174 | 21438036 | 0 | 7 | 0 | 11 | 0 | 11 | 0.4533445 | 0.527828811 | 0.826133172 | 0.961866505 |
| Q99J77 | Nans | Sialic acid synthase | 380.1 | 22.01 | 40024 | 6 | 5023453 | 5849230 | 7233714 | 5653359.5 | 5449539.5 | 5348715.5 | 4095853.75 | 5291568 | 4731855 | 6461919.5 | 4707305 | 4225174 | 4371772 | 4462641.5 | 4033709 | 5362727.5 | 6075030.5 | 6917489.5 | 5628015 | 6723912.5 | 0 | 10 | 4 | 3 | 4 | 10 | 0.746358334 | 0.840751087 | 1.0512809 | 1.184237544 |
| P28656 | Nap1l1 | Nucleosome assembly protein 1-like 1 | 420.71 | 24.55 | 45345 | 8 | 7161326 | 4196499 | 6550218 | 5977755 | 8907280 | 5667870.5 | 5914420 | 5771962 | 8805358 | 5554591 | 5560163 | 6618328 | 4706846 | 1232585.88 | 3136786.25 | 7767025 | 4751360 | 6794243 | 5200236 | 5689238 | 5 | 10 | 5 | 10 | 5 | 10 | 0.648146207 | 0.670195311 | 0.920990155 | 0.95232106 |
| Q78ZA7 | Nap1l4 | Nucleosome assembly protein 1-like 4 | 435.36 | 26.67 | 42679 | 8 | 7568224 | 6897492 | 8505788 | 8670282 | 8370651.5 | 6841997.5 | 6219358.5 | 7725769 | 6749182.5 | 7896476 | 6212867 | 7293709.5 | 5995839.5 | 6157236 | 5890278.5 | 8044053.5 | 8434061 | 7017325.5 | 8169645 | 7585750 | 1 | 10 | 3 | 4 | 3 | 10 | 0.788503087 | 0.890416371 | 0.980965856 | 1.107754772 |
| P54921 | Napa | Alpha-soluble NSF attachment protein | 651.47 | 40 | 33193 | 10 | 14403603 | 14394208 | 15317982 | 14994797 | 14840627 | 12426560 | 14625787 | 15146587 | 13576537 | 12389954 | 11278156 | 13709608 | 12978920 | 13956221 | 12916790 | 14068450 | 14616984 | 14343371 | 14297228 | 14292561 | 0 | 2 | 10 | 10 | 10 | 10 | 0.876790101 | 0.951210896 | 0.968457274 | 1.050658659 |
| Q9CWZ7 | Napg | Gamma-soluble NSF attachment protein | 443.68 | 19.23 | 34732 | 7 | 11088988 | 13770494 | 14294186 | 13985626 | 17354466 | 9085534 | 13721403 | 14470126 | 16321228 | 10887309 | 5186496 | 9488301 | 9067052 | 7356326 | 9901191 | 11600356 | 6723215 | 12266312 | 11349709 | 9564898 | 0 | 3 | 2 | 7 | 2 | 7 | 0.581602769 | 0.635791029 | 0.73062481 | 0.798697539 |
| O09043 | Napsa | Napsin-A | 229.4 | 17.42 | 45544 | 4 | 6080154.5 | 4585954 | 6229807 | 8346768.5 | 7231691.5 | 4864598.5 | 5135305 | 5948137.5 | 5553156 | 7747148 | 83108.88 | 2738259.5 | 4933388.5 | 4539107 | 3345277.5 | 5065528.5 | 919437.88 | 6425516 | 7583912 | 2 | 1 | 8 | 1 | 9 | 1 | 9 | 0.481584053 | 0.534701754 | 0.615697641 | 0.683607786 |
| Q8BP47 | Nars | Asparagine--tRNA ligase, cytoplasmic | 1188.9 | 38.64 | 64279 | 20 | 31804814 | 33833638 | 28036132 | 29310378 | 37304910 | 31173156 | 39378294 | 37759178 | 37910234 | 37735682 | 58012282 | 56377894 | 57184398 | 48501910 | 58461622 | 43794998 | 37515638 | 43010070 | 42454946 | 39100374 | 0 | 0 | 0 | 5 | 0 | 5 | 1.737714944 | 1.514151657 | 1.284398218 | 1.119155761 |
| Q99MD9 | Nasp | Nuclear autoantigenic sperm protein | 579.86 | 18.37 | 83954 | 11 | 13208880 | 12452438 | 17121542 | 13823068 | 13185591 | 13209753 | 11816861 | 9962407 | 12527017 | 12088650 | 10472987 | 10609888 | 9106457 | 8618131 | 8650723 | 14604533 | 11288573 | 12418712 | 9078682 | 10498238 | 0 | 4 | 2 | 10 | 2 | 10 | 0.679999328 | 0.79621566 | 0.829452329 | 0.971211157 |
| Q8K224 | Nat10 | RNA cytidine acetyltransferase | 662.64 | 13.18 | 115419 | 11 | 5094473 | 4581250 | 4835921 | 4890791 | 4062015 | 5944862 | 4778394 | 4161183.5 | 4432411.5 | 4774608.5 | 8406592 | 6691413 | 7915289.5 | 7058765 | 6917014.5 | 3866981 | 5908590 | 5118360 | 5307598 | 4477703.5 | 0 | 9 | 0 | 11 | 0 | 11 | 1.576387855 | 1.535360446 | 1.05177119 | 1.024397567 |
| Q6ZQA0 | Nbeal2 | Neurobeachin-like protein 2 | 210.6 | 2.08 | 301924 | 5 | 1696352.62 | 1764908 | 1555963.12 | 956059.38 | 846611.69 | 1267465.25 | 1419487.38 | 937771.31 | 1204597.75 | 737433.25 | 1190752.12 | 1109768.75 | 109576.76 | 1280502.25 | 1005553.56 | 1261480.88 | 597184.38 | 457065.41 | 674779.06 | 340131.69 | 8 | 2 | 10 | 3 | 10 | 3 | 0.688596169 | 0.843607001 | 0.488371377 | 0.59830933 |
| Q8K2Z4 | Ncapd2 | Condensin complex subunit 1 | 113.69 | 1.94 | 155665 | 2 | 650825.5 | 1005955.62 | 2 | 562503.19 | 915220.88 | 806841.94 | 711347.06 | 2 | 1247619 | 519850.62 | 2 | 2 | 2 | 2 | 2 | 601882.75 | 608341.19 | 663055.38 | 358341.25 | 511931.34 | 2.5 | 9 | 2.5 | 8 | 2.5 | 9 | 3.190294E-06 | 3.043528E-06 | 0.875273765 | 0.835007698 |
| Q8C156 | Ncaph | Condensin complex subunit 2 | 310.22 | 10.26 | 82303 | 6 | 5521893.5 | 4810283 | 5856218.5 | 6426872.5 | 6521466.5 | 4906911.5 | 4885790 | 6116669 | 5394795 | 6201698 | 1666832 | 2458803.25 | 2517309.5 | 2168312 | 1927631 | 5288410.5 | 5877861.5 | 7445426.5 | 5279768.5 | 7581697 | 0 | 10 | 0 | 8 | 0 | 10 | 0.368568685 | 0.390421764 | 1.080188466 | 1.144234719 |
| Q3UYV9 | Ncbp1 | Nuclear cap-binding protein subunit 1 | 191.73 | 5.82 | 91927 | 4 | 762120.88 | 504226.12 | 581980.62 | 641390.88 | 766964.25 | 439349.56 | 458360.06 | 545609.88 | 699079.06 | 564937 | 1022877.56 | 978124 | 1607543.38 | 743616.88 | 1067806.12 | 631671.75 | 729634.38 | 534665.69 | 586075.88 | 461346.72 | 2 | 8 | 0 | 8 | 2 | 8 | 1.66426034 | 2.00195647 | 0.903801397 | 1.087192317 |
| Q8BLF1 | Nceh1 | Neutral cholesterol ester hydrolase 1 | 1041.51 | 43.87 | 45740 | 15 | 26176840 | 25915586 | 21717722 | 30599496 | 29800744 | 37316874 | 42046442 | 40541622 | 36308186 | 37626134 | 24738324 | 24444648 | 38843466 | 35782718 | 31673582 | 29965782 | 27721510 | 37738206 | 34190530 | 39442198 | 8 | 3 | 3 | 6 | 8 | 6 | 1.158500026 | 0.802122024 | 1.259650825 | 0.872156795 |
| Q09014 | Ncf1 | Neutrophil cytosol factor 1 | 618.15 | 35.64 | 44667 | 10 | 19092254 | 22366118 | 19613442 | 19779664 | 16892904 | 17447576 | 18480754 | 15885284 | 12198444 | 14429438 | 12480584 | 14664516 | 10638219 | 10931924 | 9954414 | 22929914 | 20823540 | 15071233 | 15207780 | 17223608 | 0 | 10 | 3 | 8 | 3 | 10 | 0.600235592 | 0.747941587 | 0.933619643 | 1.163364796 |
| O70145 | Ncf2 | Neutrophil cytosol factor 2 | 379.19 | 18.29 | 59485 | 9 | 28999916 | 29687588 | 32051138 | 29952018 | 31044792 | 27173200 | 30758052 | 33324726 | 31263748 | 30734950 | 31415010 | 40913958 | 28603830 | 31758302 | 27354930 | 28819208 | 29795978 | 27799992 | 26555076 | 29292606 | 12 | 3 | 10 | 4 | 12 | 4 | 1.05477018 | 1.044314172 | 0.937571663 | 0.928277451 |
| P97369 | Ncf4 | Neutrophil cytosol factor 4 | 823.81 | 46.02 | 38707 | 14 | 35711882 | 34510726 | 33950262 | 32121430 | 34958962 | 24507624 | 36125118 | 33306356 | 30093440 | 26108464 | 24584192 | 27866014 | 22597810 | 15624558 | 26441276 | 36867446 | 31540982 | 18041720 | 20453264 | 27587354 | 0 | 5 | 5 | 10 | 5 | 10 | 0.68386347 | 0.780025765 | 0.78533258 | 0.895763077 |
| Q8K1X4 | Nckap1l | Nck-associated protein 1-like | 1031.8 | 20.63 | 128905 | 18 | 22851172 | 18089718 | 16759047 | 24889264 | 24243830 | 23257604 | 21144028 | 17757226 | 21447312 | 20323660 | 13849821 | 10856500 | 11017968 | 13959618 | 12483787 | 18723194 | 25142020 | 22453040 | 25918880 | 19159376 | 0 | 9 | 0 | 9 | 0 | 9 | 0.581914539 | 0.598169881 | 1.042715993 | 1.071843474 |
| P09405 | Ncl | Nucleolin | 3103.61 | 41.44 | 76723 | 44 | 404668802 | 458011170 | 371939618 | 315591970 | 406140034 | 341727362 | 425879074 | 347482018 | 385869538 | 290736130 | 470698018 | 579638658 | 376851682 | 318124226 | 389706946 | 364136354 | 321098754 | 290560738 | 292442050 | 328998082 | 10 | 3 | 7 | 6 | 10 | 6 | 1.091327109 | 1.191620547 | 0.816436055 | 0.891466885 |
| Q5XIA1 | Ncln | Nicalin | 350.35 | 14.21 | 62992 | 6 | 2361897.5 | 2007178.88 | 2690032 | 2914078.5 | 1989877.75 | 2851215 | 2358659.5 | 2406724.75 | 3158818.75 | 3388624 | 5428977 | 4475739.5 | 6876282 | 6385164 | 7403516.5 | 2375040 | 3738691.5 | 3569224.75 | 3569798.5 | 1988474.88 | 0 | 7 | 0 | 9 | 0 | 9 | 2.555338448 | 2.158259556 | 1.274023848 | 1.076050864 |
| Q91W39 | Ncoa5 | Nuclear receptor coactivator 5 | 45.01 | 2.07 | 65319 | 1 | 574253.75 | 451995.03 | 464504.5 | 331776 | 357867.09 | 605743.12 | 2 | 2 | 325383.25 | 527700.94 | 500491.16 | 476488.12 | 2 | 2 | 2 | 365260.25 | 471663.5 | 440066.44 | 2 | 530090.94 | 8 | 12 | 9 | 11 | 9 | 12 | 0.448076915 | 0.66970408 | 0.828786525 | 1.238719732 |
| P57716 | Ncstn | Nicastrin | 589.04 | 20.62 | 78492 | 11 | 45249702 | 39668602 | 33640378 | 48082610 | 45624214 | 45350878 | 48387974 | 45081746 | 50824898 | 46634502 | 37163574 | 41026630 | 45909194 | 46009026 | 52760210 | 45944998 | 50204118 | 47750718 | 50548614 | 56285862 | 9 | 2 | 9 | 7 | 9 | 7 | 1.049952195 | 0.943239529 | 1.181229653 | 1.061174505 |
| Q8VCB1 | Ndc1 | Nucleoporin NDC1 | 267.23 | 9.51 | 75409 | 5 | 695860.31 | 278639.16 | 588172.81 | 969604.38 | 330837.53 | 537742.06 | 1176861.5 | 1423892.75 | 1452201.12 | 769035.06 | 1845347.5 | 1755499.12 | 2789331.5 | 2344166.25 | 2448270 | 2 | 951978 | 1666394.88 | 460800.28 | 1427610.88 | 0 | 9 | 0 | 11 | 0 | 11 | 3.905752138 | 2.086412781 | 1.574085328 | 0.840860257 |
| Q78PB6 | Ndel1 | Nuclear distribution protein nudE-like 1 | 76.7 | 3.48 | 38366 | 1 | 624331.12 | 577392 | 561622.31 | 651317.56 | 579239.88 | 689109.62 | 548463.81 | 331798.59 | 470465 | 471035.06 | 175582.05 | 274071.72 | 414518.75 | 193059.81 | 2 | 724063.69 | 529171.31 | 467108.91 | 652681.5 | 286680.97 | 0 | 10 | 1 | 12 | 1 | 12 | 0.353129135 | 0.421062601 | 0.888374305 | 1.059275939 |
| Q62433 | Ndrg1 | Protein NDRG1 | 323.93 | 15.99 | 43009 | 4 | 3071850 | 5728749 | 6027499 | 6960406 | 4318586 | 4991685 | 4104167.25 | 5916715 | 4102748.75 | 5511566.5 | 3235001 | 2504239.25 | 4395794 | 5339496.5 | 4190713 | 2352695 | 4493521 | 5422143 | 4517039 | 4768272.5 | 6 | 8 | 7 | 9 | 7 | 9 | 0.753252996 | 0.798527534 | 0.825586862 | 0.875209053 |
| Q99LC3 | Ndufa10 | NADH dehydrogenase [ubiquinone] 1 alpha subcomplex subunit 10, mitochondrial | 460.68 | 27.61 | 40603 | 8 | 2801661.75 | 1632344.12 | 3675778 | 1776944.62 | 3486108.5 | 2136291.5 | 3109806 | 1975757.12 | 1470623.25 | 2452492.25 | 1831973.12 | 3160523.75 | 1881261.5 | 1179273.25 | 3272904 | 3377414.5 | 874533.06 | 2869785.75 | 596124.75 | 1651726.5 | 10 | 7 | 12 | 10 | 12 | 10 | 0.846935892 | 1.016237415 | 0.700642995 | 0.840700734 |
| Q9D8B4 | Ndufa11 | NADH dehydrogenase [ubiquinone] 1 alpha subcomplex subunit 11 | 117.59 | 9.93 | 14982 | 2 | 918465.94 | 487139.06 | 767966.75 | 594368.5 | 902823 | 571151.81 | 844511.81 | 667961.88 | 742479.5 | 893742.69 | 285570.56 | 750268.94 | 872550.88 | 549197 | 572909.19 | 858839.12 | 586432.12 | 795118.25 | 447912.16 | 680589.81 | 7 | 9 | 8 | 10 | 8 | 10 | 0.825576689 | 0.814682972 | 0.917763209 | 0.905653065 |
| Q7TMF3 | Ndufa12 | NADH dehydrogenase [ubiquinone] 1 alpha subcomplex subunit 12 | 103.11 | 6.21 | 17086 | 2 | 1837924.25 | 1523145.75 | 2131985 | 2050716.88 | 1470383.25 | 1900441 | 1433164.5 | 1437625.62 | 1369537.25 | 1576082.5 | 1105276.62 | 1062570.5 | 1294760 | 1794740.75 | 1143760.75 | 1207388.12 | 2241427.25 | 1462901 | 2 | 1409752.88 | 2 | 5 | 4 | 9 | 4 | 9 | 0.710117424 | 0.829497515 | 0.701282722 | 0.819177584 |
| Q9ERS2 | Ndufa13 | NADH dehydrogenase [ubiquinone] 1 alpha subcomplex subunit 13 | 209.49 | 30.56 | 16860 | 4 | 6764583.5 | 5480025.5 | 7088484 | 6463165 | 7985830 | 6041443.5 | 5039351 | 5477610 | 7086066 | 6632238.5 | 3950464.5 | 3709132.75 | 4660833.5 | 3384528.75 | 4019556.5 | 6159757 | 7048975 | 6297673 | 6248064 | 5379534 | 0 | 6 | 0 | 11 | 0 | 11 | 0.583874981 | 0.651474901 | 0.921612749 | 1.028315297 |
| Q9CQ75 | Ndufa2 | NADH dehydrogenase [ubiquinone] 1 alpha subcomplex subunit 2 | 180.68 | 33.33 | 10916 | 3 | 2597328.25 | 2199301.25 | 1944487.12 | 1853403.88 | 1700685.5 | 2592928.5 | 2477623.75 | 1609839 | 1501352.12 | 1535967.62 | 998798.88 | 1008313.38 | 1211659 | 1260986.38 | 1361669.25 | 1798207.25 | 2512981.25 | 1978317.62 | 1853699.88 | 2262556 | 0 | 11 | 0 | 9 | 0 | 11 | 0.56739291 | 0.601111403 | 1.01073859 | 1.070803815 |
| Q62425 | Ndufa4 | Cytochrome c oxidase subunit NDUFA4 | 247.26 | 57.32 | 9327 | 4 | 3777880.5 | 3809582.5 | 6069860.5 | 4735100 | 4200119 | 3982151.75 | 3245223 | 3853514.5 | 3556331 | 4375992 | 3985138.25 | 3607706.5 | 3684530.75 | 3628135.75 | 4419723.5 | 4534269 | 4668633 | 4613494.5 | 4415943.5 | 3940632.5 | 5 | 11 | 10 | 2 | 10 | 11 | 0.85538114 | 1.016410825 | 0.981428828 | 1.166187607 |
| Q9CPP6 | Ndufa5 | NADH dehydrogenase [ubiquinone] 1 alpha subcomplex subunit 5 | 151.12 | 23.28 | 13360 | 2 | 823004.94 | 857750.94 | 1356184.75 | 1084038.62 | 1334671.25 | 642652.56 | 1211387.12 | 756424.12 | 1310209.75 | 721022.62 | 883719.81 | 1414217.25 | 1309258.62 | 1103351.38 | 1115773.88 | 726796.81 | 652087.81 | 1034873.69 | 762595.38 | 873943.38 | 9 | 4 | 7 | 12 | 9 | 12 | 1.067942483 | 1.255213768 | 0.742404058 | 0.872589873 |
| Q9DCJ5 | Ndufa8 | NADH dehydrogenase [ubiquinone] 1 alpha subcomplex subunit 8 | 163.1 | 20.93 | 19992 | 3 | 2837565.75 | 2871971.5 | 2770359.75 | 2268351.75 | 2057943 | 2840368 | 2745397.5 | 2101966 | 2404797.5 | 2034400 | 1159601 | 1176545.62 | 1555857.5 | 1699162.5 | 2070968.75 | 2672819.75 | 1592797.75 | 2164964.75 | 1755581 | 1804521.88 | 1 | 3 | 1 | 5 | 1 | 5 | 0.598314903 | 0.631828171 | 0.780144896 | 0.823842964 |
| Q9DC69 | Ndufa9 | NADH dehydrogenase [ubiquinone] 1 alpha subcomplex subunit 9, mitochondrial | 423.58 | 27.59 | 42525 | 7 | 5430020 | 4816393.5 | 5250783 | 6420175 | 5530324 | 4921741 | 4382376 | 5353719 | 4713723 | 4926963.5 | 6626831 | 8204391 | 7099078 | 7285989 | 7952753 | 4518747.5 | 6346910 | 6364841 | 5466418.5 | 5393559 | 0 | 12 | 0 | 4 | 0 | 12 | 1.354177148 | 1.529683214 | 1.023418378 | 1.156056958 |
| O08776 | Ndufaf3 | NADH dehydrogenase [ubiquinone] 1 alpha subcomplex assembly factor 3 | 49.13 | 5.95 | 20696 | 1 | 2 | 2 | 2 | 2 | 2 | 2 | 2 | 2 | 2 | 2 | 490493.22 | 2 | 673697.75 | 608096 | 641214.25 | 2 | 2 | 2 | 2 | 2 | 2.5 | 12.5 | 2.5 | 12.5 | 2.5 | 12.5 | 241350.322 | 241350.322 | 1 | 1 |
| Q9DCS9 | Ndufb10 | NADH dehydrogenase [ubiquinone] 1 beta subcomplex subunit 10 | 154 | 21.02 | 21024 | 3 | 453797.56 | 581249.94 | 1724883.38 | 1560712.75 | 1358830.12 | 1314558 | 1068626.25 | 1297383.25 | 1091304.75 | 1436429.5 | 1693377.5 | 1694319.25 | 1720879.88 | 1718913.12 | 1670265.75 | 1244151.62 | 1896555.38 | 1462527.25 | 1591063.75 | 1309588 | 5 | 9 | 0 | 5 | 5 | 9 | 1.496222339 | 1.36877295 | 1.321229102 | 1.208685773 |
| Q9CQC7 | Ndufb4 | NADH dehydrogenase [ubiquinone] 1 beta subcomplex subunit 4 | 40.36 | 19.38 | 15081 | 1 | 2 | 2 | 2 | 2 | 2 | 2 | 2 | 331557.88 | 303888 | 2 | 549060.06 | 496550.09 | 2 | 2 | 504872.38 | 2 | 2 | 2 | 2 | 171431.03 | 5 | 10 | 7 | 9 | 7 | 10 | 155048.653 | 2.439974731 | 17143.903 | 0.269790735 |
| Q9CQH3 | Ndufb5 | NADH dehydrogenase [ubiquinone] 1 beta subcomplex subunit 5, mitochondrial | 108.25 | 12.7 | 21710 | 2 | 2120298 | 2030765.62 | 2577926 | 1723672.62 | 1763306.12 | 2382378.5 | 1412097.12 | 1562223.62 | 2165743 | 1833793.38 | 3164803.5 | 1771798.38 | 2177072.5 | 2 | 2 | 2389614.75 | 2600266.25 | 2555679.75 | 2523489.75 | 1757853.5 | 11 | 7 | 11 | 3 | 11 | 7 | 0.696329328 | 0.760314155 | 1.157688002 | 1.264066499 |
| Q91VD9 | Ndufs1 | NADH-ubiquinone oxidoreductase 75 kDa subunit, mitochondrial | 944.3 | 30.26 | 79777 | 18 | 20146806 | 19998182 | 26282816 | 24871234 | 21253196 | 20981858 | 20163288 | 22764546 | 20151290 | 21299490 | 20112368 | 21837064 | 20630704 | 21249114 | 24067600 | 20220126 | 21412238 | 18756854 | 22988490 | 21798084 | 11 | 11 | 11 | 10 | 11 | 11 | 0.958638013 | 1.024073336 | 0.934462056 | 0.99824716 |
| Q91WD5 | Ndufs2 | NADH dehydrogenase [ubiquinone] iron-sulfur protein 2, mitochondrial | 383.61 | 17.06 | 52626 | 6 | 9296235 | 5752037 | 8333236.5 | 8463455 | 8743907 | 7863288.5 | 6392308 | 7029388 | 8820329 | 5630111.5 | 6208146 | 6068435 | 6297680 | 4608767 | 7597978 | 7938045 | 9370727 | 6632652 | 9544523 | 7452202 | 4 | 12 | 6 | 6 | 6 | 12 | 0.758360743 | 0.86135833 | 1.008605278 | 1.145590097 |
| Q9DCT2 | Ndufs3 | NADH dehydrogenase [ubiquinone] iron-sulfur protein 3, mitochondrial | 483.92 | 31.18 | 30149 | 8 | 3719808 | 4586732.5 | 5253568 | 3450449 | 4917838 | 4062306 | 4079880.25 | 5219186.5 | 4220454.5 | 4354091 | 6600296 | 5391325.5 | 7021944 | 3827313.25 | 4706475 | 2964833 | 3140421.75 | 5565839 | 2970358.75 | 2776380.25 | 5 | 5 | 6 | 5 | 6 | 5 | 1.256241194 | 1.255810376 | 0.794304934 | 0.794032534 |
| Q9DC70 | Ndufs7 | NADH dehydrogenase [ubiquinone] iron-sulfur protein 7, mitochondrial | 416.74 | 21.88 | 24683 | 6 | 8331902.5 | 7781588.5 | 11133887 | 10638237 | 8532388 | 8098390 | 6908769.5 | 8169922.5 | 7579060.5 | 9680331 | 4990470 | 6568617 | 7359119 | 9084813 | 7251642 | 7209298.5 | 9233642 | 11329677 | 9709663 | 9693783 | 3 | 11 | 6 | 5 | 6 | 11 | 0.759504044 | 0.871853007 | 1.016331174 | 1.166671062 |
| Q8K3J1 | Ndufs8 | NADH dehydrogenase [ubiquinone] iron-sulfur protein 8, mitochondrial | 162.56 | 17.92 | 24038 | 3 | 167977 | 271683.91 | 287570.25 | 43458.88 | 1634890.5 | 2 | 297183.09 | 964502.88 | 546904.12 | 2 | 2 | 704049.38 | 2 | 2 | 2 | 637198.44 | 557344.19 | 2 | 120637.07 | 2 | 4 | 9 | 8 | 12 | 8 | 12 | 0.292676702 | 0.389284353 | 0.546721957 | 0.727185667 |
| Q91YT0 | Ndufv1 | NADH dehydrogenase [ubiquinone] flavoprotein 1, mitochondrial | 204.16 | 8.62 | 50834 | 4 | 7827861 | 7604368 | 7900273 | 6610103.5 | 7431114 | 6941906 | 6923320.5 | 4786882.5 | 5838122.5 | 6151260.5 | 4656382.5 | 5538700 | 4299929.5 | 5207251 | 4972477 | 7635911 | 6832311 | 6635896.5 | 6983816 | 6690886 | 0 | 7 | 3 | 6 | 3 | 7 | 0.660216332 | 0.805272145 | 0.930568885 | 1.135023729 |
| P69682 | Necap1 | Adaptin ear-binding coat-associated protein 1 | 48.08 | 6.14 | 29792 | 1 | 718526.69 | 2 | 961814.06 | 971884.69 | 566137.69 | 979877.25 | 904641.94 | 2 | 2 | 1168781.38 | 920190.06 | 648644.56 | 2 | 1329704 | 911978.94 | 576371.44 | 2 | 1483644 | 1216216.88 | 1192619.38 | 11.5 | 7.5 | 11 | 7 | 11.5 | 7.5 | 1.183992309 | 1.247998512 | 1.388547762 | 1.463612161 |
| P29595 | Nedd8 | NEDD8 | 177.96 | 34.57 | 8972 | 3 | 1317815.62 | 1238607.38 | 3618282 | 2334638.5 | 3393926 | 1210369.5 | 2545057 | 3402637.75 | 2771530 | 2756943 | 292929.28 | 2685313 | 1685628.38 | 607370.12 | 2140543 | 3876942 | 1149853.88 | 3458624.5 | 2459325 | 2597851.25 | 7 | 10 | 4 | 12 | 7 | 12 | 0.622667896 | 0.584224334 | 1.137720744 | 1.067477781 |
| D3ZBE5 | Nek7 | Serine/threonine-protein kinase Nek7 | 49.19 | 2.65 | 34529 | 1 | 1380538 | 1679725.75 | 1866630.5 | 1962443.88 | 2 | 1385129.62 | 1363561.88 | 2 | 2 | 1946077.12 | 1064100.88 | 1369176.62 | 1434598.75 | 1508242 | 1518856.5 | 1149711.75 | 1206535.25 | 1803839.5 | 1995219.38 | 1615255.12 | 8 | 12 | 8 | 8 | 8 | 12 | 1.000817875 | 1.468649349 | 1.12791078 | 1.655151725 |
| Q8K1R7 | Nek9 | Serine/threonine-protein kinase Nek9 | 198.6 | 4.37 | 107143 | 4 | 3600253.5 | 4028356.25 | 3994219.5 | 3823853 | 3206381.5 | 3409509.5 | 3869755.5 | 5381937 | 4744208.5 | 4607966 | 1187979.88 | 2737968.75 | 2714143.5 | 2382375.75 | 3240947.5 | 3634014 | 3293495.75 | 4088582.5 | 4185709.5 | 3190629.25 | 1 | 12 | 0 | 5 | 1 | 12 | 0.657447781 | 0.557089249 | 0.986027349 | 0.835511581 |
| Q922L6 | Nelfcd | Negative elongation factor D | 86.03 | 4.91 | 66276 | 2 | 932372.62 | 625843.88 | 578636.44 | 565198.44 | 527419.5 | 502168.06 | 399793.25 | 189567.44 | 560186.94 | 567681.25 | 683473.06 | 581470.62 | 544116.12 | 2 | 247163.27 | 534911.62 | 179309.33 | 463383.38 | 133889.77 | 256798.31 | 8 | 1 | 11 | 6 | 11 | 6 | 0.63670649 | 0.926479186 | 0.485618997 | 0.706629978 |
| Q8CCP0 | Nemf | Nuclear export mediator factor Nemf | 75.21 | 1.6 | 121188 | 2 | 1356336.38 | 2 | 1206994.75 | 1305170.25 | 1086612 | 1514420.62 | 2 | 1273825 | 1143388.5 | 1215643.5 | 1198478.75 | 1225668.88 | 1181751.62 | 1042938.62 | 1011424.25 | 1516294 | 1067053.5 | 936327.06 | 1468325 | 1272704.25 | 9 | 10 | 9 | 11 | 9 | 11 | 1.142306826 | 1.099660896 | 1.263482952 | 1.216313135 |
| Q6P5H2 | Nes | Nestin | 3779.04 | 38.84 | 207124 | 56 | 82630850 | 83543066 | 87475674 | 79315674 | 75873674 | 86683522 | 85050642 | 76554850 | 71168058 | 79394706 | 55011118 | 56721710 | 52528002 | 60304046 | 50969522 | 86807266 | 82363346 | 77629242 | 74059122 | 73155674 | 0 | 7 | 0 | 12 | 0 | 12 | 0.67394363 | 0.690819029 | 0.963740518 | 0.987872367 |
| Q60591 | Nfatc2 | Nuclear factor of activated T-cells, cytoplasmic 2 | 183.25 | 7.66 | 100020 | 4 | 2152759.5 | 2687238.25 | 2218399 | 2679699.25 | 2736416.5 | 3473474.25 | 3619936.25 | 2593115.5 | 2325227.25 | 2895045.75 | 1854546.25 | 1555784.25 | 1119287.25 | 1128603 | 1014302.88 | 2416785.75 | 2595109.5 | 1963661.75 | 2453731 | 2780668 | 0 | 11 | 0 | 6 | 0 | 11 | 0.534892536 | 0.447616127 | 0.978792237 | 0.819086378 |
| P09414 | Nfia | Nuclear factor 1 A-type | 58.84 | 1.96 | 55976 | 1 | 100567.18 | 106806.77 | 195057.75 | 108548.04 | 112440.68 | 2 | 68628.5 | 115352.87 | 125519.3 | 63382.67 | 2 | 2 | 2 | 78369.04 | 2 | 49373.26 | 2 | 51491.75 | 98625 | 2 | 0 | 0 | 5 | 6 | 5 | 6 | 0.125721002 | 0.210190725 | 0.319999159 | 0.535000947 |
| P25799 | Nfkb1 | Nuclear factor NF-kappa-B p105 subunit | 430.05 | 9.89 | 105615 | 9 | 3455001.5 | 4925719.5 | 6023282 | 6753021 | 5330443 | 3964123.5 | 3308183.75 | 6573390 | 5697915 | 3591291 | 2459091.75 | 3067309 | 4022647.5 | 2748447.5 | 3117617.25 | 2374237.25 | 1974315 | 3060679 | 4765138.5 | 3321037.25 | 1 | 1 | 3 | 4 | 3 | 4 | 0.58197762 | 0.666314133 | 0.585009016 | 0.66978482 |
| Q9WTK5 | Nfkb2 | Nuclear factor NF-kappa-B p100 subunit | 252.14 | 8.01 | 96832 | 5 | 1678799.38 | 1782291.12 | 1936682.5 | 2538617.75 | 1928344.5 | 1305949 | 2637831 | 2756370.5 | 2611060.5 | 3478273.25 | 1760668.12 | 2017624.25 | 1853927.5 | 1374200.25 | 2030626.75 | 1494737.25 | 1617424.25 | 2554958.5 | 1328760.12 | 2118392.5 | 11 | 9 | 5 | 5 | 11 | 9 | 0.91609624 | 0.706599789 | 0.923924706 | 0.712638011 |

| | | | | | | | | | | | | | | | | | | | | | | | | | | | | | | | | | | | | |
|---|---|---|---|---|---|---|---|---|---|---|---|---|---|---|---|---|---|---|---|---|---|---|---|---|---|---|---|---|---|---|---|---|---|---|---|---|
| O54910 | Nfkbie | NF-kappa-B inhibitor epsilon | 68.07 | 3.3 | 39200 | 1 | 710709.75 | 412876.19 | 581061.31 | 709230.81 | 630205.94 | 733427.19 | 584341.88 | 559367.69 | 929797.75 | 601614.56 | 294136.62 | 183295.42 | 280224.16 | 278984.47 | 234574.44 | 432566.53 | 608283.62 | 443356.38 | 2 | 483614.44 | 0 | 5 | 0 | 3 | 0 | 5 | 0.41760185 | 0.372949042 | 0.646441744 | 0.57731983 |
| Q9QZ23 | Nfu1 | NFU1 iron-sulfur cluster scaffold homolog, mitochondrial | 232.38 | 20 | 28568 | 4 | 3650157.5 | 3010367.5 | 4161302 | 4676266 | 4672414 | 6135233.5 | 5129845.5 | 4691113 | 4105628.25 | 6768579.5 | 4967165.5 | 5272822 | 7961043 | 7968693 | 6626280.5 | 3751522.5 | 2836053.5 | 5139491.5 | 8492115 | 8647118 | 0 | 8 | 6 | 12 | 6 | 12 | 1.625938505 | 1.222344963 | 1.431114275 | 1.075880373 |
| Q8BZW8 | Nhlrc2 | NHL repeat-containing protein 2 | 153.95 | 4.69 | 78430 | 3 | 1826595.5 | 1889379.25 | 1962065 | 2243295.75 | 1882059.38 | 1813269 | 1826638.5 | 2039242.25 | 1496183 | 2392662.5 | 1661667 | 849205.62 | 1265052.62 | 1454385.5 | 1462582 | 2239470 | 2434680 | 2351787 | 2814327.75 | 2125345.75 | 0 | 2 | 1 | 3 | 1 | 3 | 0.682711736 | 0.699508368 | 1.220557842 | 1.250587003 |
| Q8CCH2 | Nhlrc3 | NHL repeat-containing protein 3 | 230.25 | 10.09 | 38195 | 3 | 3263604.5 | 3181009.25 | 675987 | 3109822 | 4117308.5 | 2105077.75 | 3536181 | 3183657 | 2573584 | 1062182.12 | 322726.72 | 1843430.5 | 1804321.25 | 2 | 511184.06 | 4979161.5 | 2388706 | 4857589.5 | 3366518.25 | 4392374.5 | 2 | 5 | 2 | 4 | 2 | 5 | 0.312360502 | 0.359664469 | 1.392857826 | 1.60379263 |
| Q9CRB2 | Nhp2 | H/ACA ribonucleoprotein complex subunit 2 | 192.02 | 18.95 | 17247 | 3 | 2 | 438552 | 2 | 156738.73 | 2 | 225079.64 | 481514.38 | 3138326.75 | 2592455 | 2778211.25 | 599622.69 | 1120493 | 1952721.75 | 1987410.5 | 711390.12 | 3540101.75 | 580403.12 | 2 | 2671292.25 | 2913688.75 | 0 | 3.5 | 10 | 11 | 10 | 11 | 10.7032976 | 0.691397959 | 16.30361361 | 1.053160026 |
| Q9EQ80 | Nif3l1 | NIF3-like protein 1 | 174.32 | 8.78 | 41746 | 3 | 7087108 | 6809518 | 7576722.5 | 7784971.5 | 6580910.5 | 8809040 | 7063845 | 7775607 | 7808562 | 9324528 | 6691021 | 7061022 | 7145302 | 8379003.5 | 7170472 | 8198311 | 8442656 | 9703591 | 11107448 | 9630275 | 11 | 0 | 5 | 4 | 11 | 4 | 1.01695321 | 0.893707863 | 1.313707921 | 1.154498641 |
| Q91VE6 | Nifk | MKI67 FHA domain-interacting nucleolar phosphoprotein | 463.93 | 29.97 | 36265 | 8 | 5459895.5 | 4455114 | 5130856.5 | 3020759.5 | 3620166 | 5815124 | 5426969.5 | 2517076 | 3762796.5 | 3196165 | 6516215.5 | 6685845.5 | 5618609.5 | 5606259 | 7317639.5 | 5651648 | 4636420.5 | 5463489.5 | 3992782.5 | 4215554 | 0 | 8 | 2 | 8 | 2 | 8 | 1.463774344 | 1.532212003 | 1.10481509 | 1.156469881 |
| Q9CXK8 | Nip7 | 60S ribosome subunit biogenesis protein NIP7 homolog | 286.22 | 50.56 | 20452 | 6 | 2190946.5 | 2829305 | 2718762.5 | 2803662 | 2756269 | 2486735 | 3184731.5 | 3131901.5 | 2572290.5 | 2125389 | 2228655.5 | 3974549.25 | 3486735.5 | 2123346 | 4980735 | 3575501.75 | 1718268.38 | 2032862.88 | 2447262.25 | 3634772.5 | 9 | 11 | 9 | 11 | 9 | 11 | 1.262808535 | 1.243905056 | 1.008250486 | 0.993157587 |
| Q6KCD5 | Nipbl | Nipped-B-like protein | 60.23 | 0.39 | 315450 | 1 | 345534.06 | 345123.16 | 316052.91 | 380630.84 | 359529.22 | 440057.06 | 331563.75 | 278631.75 | 336517.62 | 379026.72 | 304127.16 | 246521.72 | 386853.41 | 2 | 2 | 331244.34 | 2 | 403154.16 | 2 | 2 | 5 | 6 | 5 | 5 | 5 | 6 | 0.536677708 | 0.530925323 | 0.420411605 | 0.415905419 |
| O55125 | Nipsnap1 | Protein NipSnap homolog 1 | 99.46 | 9.15 | 33363 | 2 | 1049040.12 | 1453029.62 | 2 | 1628134.38 | 1570484.62 | 2 | 1121792.25 | 2439983 | 1855869.88 | 2453106.5 | 3379245.5 | 3505783 | 2880152 | 632153.94 | 3563725.5 | 2 | 2 | 2027852.38 | 1887936.5 | 1986634.75 | 4 | 9 | 4 | 10 | 4 | 10 | 2.449011984 | 1.773789474 | 1.035388148 | 0.749918992 |
| O55126 | Nipsnap2 | Protein NipSnap homolog 2 | 45.14 | 6.05 | 32933 | 1 | 2 | 314710.03 | 637949.94 | 408583.53 | 2 | 2 | 357181.06 | 255347.89 | 574732.06 | 277415.62 | 1056749.25 | 2 | 961227.19 | 2 | 2 | 2 | 2 | 274568.72 | 153549.66 | 2 | 12 | 7 | 11.5 | 4.5 | 12 | 7 | 1.482450796 | 1.37776465 | 0.314508846 | 0.292299192 |
| Q9CQE1 | Nipsnap3b | Protein NipSnap homolog 3B | 47.48 | 4.45 | 28308 | 1 | 318844.84 | 316624.91 | 320364.41 | 196607.66 | 451262.72 | 150790.58 | 343560.62 | 304094.28 | 450273.53 | 2 | 226206.2 | 2 | 250300.44 | 2 | 235409.38 | 418752.47 | 2 | 170869.62 | 267646.59 | 260945.3 | 3 | 6 | 7 | 10.5 | 7 | 10.5 | 0.443922183 | 0.570119358 | 0.697270571 | 0.895489041 |
| Q80TM9 | Nisch | Nischarin | 430.32 | 5.9 | 175012 | 7 | 2888684 | 3563432.75 | 1734345.75 | 2077359.75 | 3055282.5 | 2884099.5 | 2405558 | 1327257.25 | 2242132.75 | 2514575 | 3549032.5 | 3455348 | 2963497.5 | 2871976 | 3357281 | 3359441.5 | 2960934.25 | 3549464.5 | 3813567.5 | 3697778.75 | 8 | 4 | 1 | 0 | 8 | 4 | 1.21608286 | 1.424096413 | 1.30498159 | 1.528201459 |
| Q8VDK1 | Nit1 | Deaminated glutathione amidase | 92.23 | 5.57 | 35705 | 2 | 1453100.62 | 1793978.12 | 1165273.75 | 608631.81 | 1501805.5 | 593284.62 | 1475366.25 | 1434075.88 | 1432678 | 1180252 | 729199.69 | 1360792.5 | 450124.47 | 373706.66 | 862305.25 | 654782.75 | 560388.44 | 584645 | 190088.25 | 715878.5 | 4 | 2 | 4 | 2 | 4 | 2 | 0.578913116 | 0.617452667 | 0.414819889 | 0.442435384 |
| Q9JHW2 | Nit2 | Omega-amidase NIT2 | 239.2 | 14.86 | 30502 | 3 | 2907915 | 1580953.75 | 2244510.25 | 1947634.5 | 2931020 | 1229578.12 | 904008.31 | 3494207.25 | 3354790.75 | 1701621.12 | 1677937.25 | 1393603 | 1375600 | 2 | 1256064.62 | 3095384.25 | 819708.38 | 2147716.5 | 2500249.25 | 2457042 | 1 | 12 | 8 | 12 | 8 | 12 | 0.491146264 | 0.533797936 | 0.949024164 | 1.031438447 |
| Q8VEJ4 | Nle1 | Notchless protein homolog 1 | 147.95 | 7.63 | 53134 | 3 | 1363441.75 | 1152372.62 | 1019031.12 | 1030047 | 1322283.88 | 1427123.25 | 969060 | 1111331.75 | 950005 | 1354343.25 | 1812520.5 | 1369894.12 | 2004249.25 | 1369809.25 | 1627184.88 | 1197985.5 | 666700.62 | 1234409.25 | 1000239.62 | 1203859.5 | 0 | 9 | 2 | 11 | 2 | 11 | 1.390082017 | 1.408095416 | 0.900804419 | 0.912477507 |
| Q3TL44 | Nlrx1 | NLR family member X1 | 147.88 | 3.28 | 107831 | 3 | 588661.62 | 2 | 686266.31 | 1309574.88 | 1261394.75 | 699580.62 | 2 | 952137.88 | 931891.44 | 1428011 | 588832.69 | 683587.38 | 1604621 | 1054347.62 | 1206998.75 | 475730.75 | 669718.75 | 993324.12 | 1505068.38 | 978523.31 | 10 | 11 | 10 | 10 | 10 | 11 | 1.336069068 | 1.280874977 | 1.201894443 | 1.152243214 |
| P15532 | Nme1 | Nucleoside diphosphate kinase A | 479.88 | 56.58 | 17208 | 9 | 5162010.5 | 4433155 | 6081308 | 4752376 | 4726269.5 | 4044831.5 | 3773797.75 | 5375431 | 4355706 | 5493887 | 4830645 | 5393593 | 4300331 | 2777744.25 | 4567142.5 | 5009792 | 2685147.25 | 5255014 | 5535974.5 | 5378221.5 | 8 | 10 | 12 | 10 | 12 | 10 | 0.869383911 | 0.949044646 | 0.948679641 | 1.03560616 |
| Q01768 | Nme2 | Nucleoside diphosphate kinase B | 648.71 | 78.95 | 17363 | 12 | 16472193 | 16064017 | 14718082 | 13261750 | 17060940 | 13417161 | 14523548 | 13789482 | 12520544 | 10738678 | 16225724 | 18258074 | 10697939 | 8128738 | 13476601 | 19112562 | 12981878 | 15970995 | 12838337 | 14749983 | 9 | 9 | 12 | 6 | 12 | 9 | 0.860913563 | 1.027660859 | 0.975208793 | 1.164093527 |
| Q9WV85 | Nme3 | Nucleoside diphosphate kinase 3 | 40.44 | 10.06 | 19099 | 1 | 822249.19 | 524122.94 | 894998.38 | 1100658.75 | 636907.69 | 584892.62 | 664396.25 | 737392.44 | 633397.5 | 747633.44 | 500719.25 | 474790.16 | 2 | 612435.44 | 2 | 713585.81 | 1064367.12 | 564412.19 | 2 | 2 | 1 | 7 | 1 | 8 | 1 | 8 | 0.399088719 | 0.471521535 | 0.588692193 | 0.695537191 |
| O35309 | Nmi | N-myc-interactor | 45.35 | 5.1 | 35236 | 1 | 517745.31 | 2 | 2 | 757032.75 | 2 | 620735.31 | 2 | 763456 | 559731.12 | 826241.81 | 451389.72 | 568101.81 | 558798 | 469005.75 | 797822.88 | 2 | 689181.31 | 2 | 621854.44 | 543048.25 | 6 | 10 | 9 | 8 | 9 | 10 | 2.231843219 | 1.027056831 | 1.454432996 | 0.66930568 |
| Q8K2T1 | Nmral1 | NmrA-like family domain-containing protein 1 | 202.99 | 11.97 | 34376 | 4 | 5982230.5 | 6827606 | 9352849 | 8491133 | 11117667 | 7360319 | 6306315 | 8417797 | 9027412 | 9043260 | 3947668.5 | 4330879 | 4432504 | 4191282 | 6360666.5 | 7350851 | 3658344 | 7506822 | 7709449.5 | 6711734 | 1 | 7 | 1 | 6 | 1 | 7 | 0.556911006 | 0.57932861 | 0.788509197 | 0.820249434 |
| O70310 | Nmt1 | Glycylpeptide N-tetradecanoyltransferase 1 | 608.25 | 22.38 | 56888 | 11 | 23101998 | 23401698 | 27108654 | 31118298 | 24459014 | 27666790 | 23694986 | 21315858 | 27052930 | 26941786 | 20570810 | 18619410 | 21566356 | 24779246 | 20032614 | 24957422 | 31009470 | 30378344 | 32791592 | 31328242 | 3 | 4 | 3 | 3 | 3 | 4 | 0.817158543 | 0.833397628 | 1.164683518 | 1.187828836 |
| Q61941 | Nnt | NAD(P) transhydrogenase, mitochondrial | 384.86 | 10.22 | 113838 | 8 | 4981756.5 | 5732983 | 5207198.5 | 5326657 | 4993245.5 | 6315677 | 4399176.5 | 5003456 | 4678532.5 | 5204133 | 5881561 | 4837920.5 | 6885334 | 5855049.5 | 6504626 | 5663979 | 4992451 | 5020432 | 5572387 | 4172616.5 | 5 | 11 | 5 | 12 | 5 | 12 | 1.141859352 | 1.170443352 | 0.968753144 | 0.993003802 |
| Q8BHY2 | Noc4l | Nucleolar complex protein 4 homolog | 141.46 | 8.33 | 58676 | 3 | 482483.81 | 367644.16 | 375306.72 | 810527.62 | 430970.44 | 381965.22 | 549821.88 | 390075.38 | 671868.62 | 643146.81 | 2145436.25 | 1800153.62 | 2222172 | 1652804.5 | 1924756.38 | 636618.75 | 701632.12 | 1130868.12 | 778111.75 | 712514.75 | 0 | 4 | 0 | 2 | 0 | 4 | 3.950380386 | 3.695780799 | 1.605129078 | 1.501679496 |
| Q5RJG1 | Nol10 | Nucleolar protein 10 | 73.64 | 3.2 | 80077 | 2 | 441869.38 | 2 | 2 | 2 | 206570.38 | 445058.28 | 390393.88 | 191787.09 | 400325.22 | 217536.81 | 1124332.75 | 1291210.12 | 1116471 | 2 | 1159155.75 | 336545.41 | 646277.75 | 2 | 2 | 2 | 3.5 | 11.5 | 5 | 7 | 5 | 11.5 | 7.234485765 | 2.851600492 | 1.515669036 | 0.597427752 |
| Q8BJW5 | Nol11 | Nucleolar protein 11 | 153.78 | 4.7 | 80818 | 3 | 464722.16 | 2 | 696716.06 | 129107.3 | 2 | 2 | 311365 | 330025.44 | 666895.94 | 520841.44 | 1091020.88 | 901809.25 | 1120244 | 951453.5 | 1077342.12 | 393257.16 | 2 | 901448.44 | 414319.69 | 460637.38 | 0 | 10 | 0 | 10.5 | 0 | 10.5 | 3.984248315 | 2.811101593 | 1.681194434 | 1.186173144 |
| Q9D7Z3 | Nol7 | Nucleolar protein 7 | 50.24 | 7.87 | 28970 | 1 | 2 | 159659.67 | 2 | 2 | 2 | 2 | 2 | 2 | 2 | 64883.62 | 2 | 223887.06 | 68414.18 | 80628.8 | 2 | 2 | 2 | 2 | 2 | 2 | 8 | 10 | 6 | 10 | 8 | 10 | 2.335689122 | 5.747029277 | 6.263009E-05 | 0.000154103 |
| Q3UHX0 | Nol8 | Nucleolar protein 8 | 40.56 | 1.05 | 128635 | 1 | 412848.03 | 509613.88 | 2 | 2 | 338055.72 | 535945.25 | 496577.22 | 347288.5 | 286609.44 | 2 | 2 | 2 | 2 | 295754.19 | 180260.78 | 2 | 2 | 2 | 385637.62 | 298102.75 | 7 | 8 | 4.5 | 6.5 | 7 | 8 | 0.377638081 | 0.285654446 | 0.542431287 | 0.410307954 |
| Q3TZX8 | Nol9 | Polynucleotide 5'-hydroxyl-kinase NOL9 | 48.08 | 0.98 | 80840 | 1 | 417960.38 | 499525.66 | 489977.59 | 161880.64 | 923280.44 | 76429.84 | 843024.62 | 101762.52 | 921998.44 | 233584.03 | 155674.92 | 340435.28 | 2 | 2 | 103087.65 | 274066.75 | 97853.95 | 92435.7 | 349777.41 | 151989.78 | 1 | 2 | 7 | 10 | 7 | 10 | 0.240389918 | 0.275267366 | 0.38759288 | 0.443827561 |
| E9Q5C9 | Nolc1 | Nucleolar and coiled-body phosphoprotein 1 | 352.13 | 10.54 | 73698 | 7 | 5308930 | 3487162.25 | 2647536.5 | 3311617 | 2123570.75 | 4847180 | 5219453 | 2683978 | 2714760 | 3755307 | 5031945.5 | 4436280 | 2759676.25 | 2814023.75 | 2286804.75 | 10113803 | 7991895 | 2553613.25 | 3346953.5 | 2649692.5 | 12 | 9 | 11 | 12 | 12 | 12 | 1.026655527 | 0.901567065 | 1.579255112 | 1.386837512 |
| Q6GQT9 | Nomo1 | Nodal modulator 1 | 1444.12 | 29.98 | 133420 | 26 | 13885431 | 16211102 | 14992131 | 13344158 | 14377059 | 15075999 | 14709042 | 14809197 | 15125735 | 17485566 | 30640490 | 26349218 | 34732982 | 36688214 | 37390762 | 13504927 | 14120330 | 16785052 | 16430040 | 14640522 | 0 | 9 | 0 | 8 | 0 | 9 | 2.277186334 | 2.147535891 | 1.036684444 | 0.977661344 |
| Q5FVM4 | Nono | Non-POU domain-containing octamer-binding protein | 1012.25 | 39.29 | 54925 | 16 | 19095936 | 18283762 | 17843710 | 18360582 | 16657644 | 20318412 | 18016844 | 16189392 | 14237996 | 19340924 | 20743684 | 19883662 | 15625191 | 18210672 | 14889860 | 14436357 | 23840308 | 15414043 | 18361536 | 18027104 | 12 | 11 | 11 | 12 | 12 | 12 | 0.990153492 | 1.014182184 | 0.99820165 | 1.022425653 |
| Q9CQS2 | Nop10 | H/ACA ribonucleoprotein complex subunit 3 | 44.94 | 17.19 | 7706 | 1 | 2 | 2 | 2 | 2 | 2 | 2 | 2 | 2 | 2 | 2 | 2 | 2 | 267803.69 | 405120.25 | 661964.25 | 2 | 2 | 2 | 2 | 2 | 5 | 12.5 | 5 | 12.5 | 5 | 12.5 | 133489.219 | 133489.219 | 1 | 1 |
| Q8R3N1 | Nop14 | Nucleolar protein 14 | 134.69 | 5 | 98769 | 3 | 256864.2 | 578251.88 | 276486.41 | 2 | 651420.38 | 68524.44 | 286210.22 | 244126.3 | 360816.47 | 2 | 172482.86 | 375209.69 | 190176.48 | 2 | 222611.28 | 336289.72 | 109494.95 | 2 | 2 | 2 | 6.5 | 5.5 | 11.5 | 7.5 | 11.5 | 7.5 | 0.544792264 | 1.000836613 | 0.252855577 | 0.464520397 |
| Q1RP77 | Nop16 | Nucleolar protein 16 | 64.55 | 7.87 | 21117 | 1 | 2 | 552282.94 | 2 | 2 | 621762.75 | 676106.56 | 512587.31 | 2 | 2 | 2 | 606349.88 | 834809.12 | 631090.88 | 2 | 2 | 233157.31 | 889083.12 | 653852.25 | 2 | 698343.31 | 8 | 5.5 | 9 | 6.5 | 9 | 6.5 | 1.765044842 | 1.743294445 | 2.107605663 | 2.081633937 |
| Q922K7 | Nop2 | Probable 28S rRNA (cytosine-C(5))-methyltransferase | 895.93 | 24.84 | 86752 | 16 | 8843372 | 7258692 | 8443427 | 7876211 | 8192503.5 | 8632370 | 8133005 | 6747371 | 3752513.75 | 5162488.5 | 18339070 | 16182421 | 13792965 | 12385329 | 13906157 | 9195034 | 7181103 | 10268502 | 7478798.5 | 9958964 | 0 | 9 | 0 | 4 | 0 | 9 | 1.836942052 | 2.300682163 | 1.085393668 | 1.359403717 |
| Q9D6Z1 | Nop56 | Nucleolar protein 56 | 1456.35 | 50.52 | 64464 | 22 | 24714460 | 26917606 | 25666888 | 26463252 | 29232222 | 22555780 | 24241564 | 30136682 | 30168178 | 29537040 | 45194046 | 44934298 | 35367206 | 30939950 | 41319714 | 20329466 | 16356184 | 26771932 | 28669706 | 28750968 | 0 | 11 | 0 | 6 | 0 | 11 | 1.486943603 | 1.447279773 | 0.908897146 | 0.884652553 |
| Q6DFW4 | Nop58 | Nucleolar protein 58 | 850.82 | 29.85 | 60343 | 13 | 12179462 | 13447040 | 14403050 | 12834634 | 14660234 | 11739022 | 13425088 | 12108186 | 12913006 | 11137818 | 8394555 | 8959250 | 9740966 | 8547786 | 9009165 | 12226457 | 12551720 | 9143782 | 11093843 | 12081182 | 0 | 2 | 0 | 8 | 0 | 8 | 0.661267761 | 0.728138457 | 0.845575334 | 0.931084133 |
| Q8BMC4 | Nop9 | Nucleolar protein 9 | 186.68 | 6.6 | 70046 | 3 | 487666.84 | 378638.81 | 315864.78 | 240340.75 | 313415.75 | 226374.36 | 273314.62 | 520693.72 | 473362.56 | 424334.16 | 1369486.5 | 1198407.5 | 1109365.88 | 745497.94 | 565546.31 | 260704.33 | 118071.19 | 2 | 322541.28 | 2 | 0 | 4 | 0 | 3 | 0 | 4 | 2.873568031 | 2.60067653 | 0.404003641 | 0.365636997 |
| O35604 | Npc1 | NPC intracellular cholesterol transporter 1 | 278.93 | 3.84 | 142885 | 5 | 4667573.5 | 4213440.5 | 2370926.75 | 2474595.25 | 2860279.75 | 3712655 | 3394186.75 | 3107550.75 | 3145275.5 | 2257213 | 3786359.5 | 4727541 | 3459669.5 | 2627343 | 3364697 | 3427101 | 3207194.25 | 2790475.5 | 1997139.75 | 2753112.25 | 9 | 10 | 7 | 9 | 9 | 10 | 1.083125916 | 1.150396805 | 0.85459578 | 0.907673097 |
| Q9Z0J0 | Npc2 | NPC intracellular cholesterol transporter 2 | 252.08 | 41.61 | 16442 | 5 | 48277046 | 38412902 | 24695770 | 34918790 | 26873782 | 32027882 | 32829162 | 28393774 | 25873526 | 34777866 | 4590716 | 5648711 | 5558537.5 | 3658984 | 5036400 | 37588486 | 20821178 | 17530012 | 16897220 | 21071624 | 0 | 3 | 0 | 5 | 0 | 5 | 0.14143429 | 0.159148777 | 0.657752886 | 0.7401357 |
| Q6NSR8 | Npepl1 | Probable aminopeptidase NPEPL1 | 219.72 | 9.73 | 55940 | 4 | 1649239.25 | 1470620.75 | 785400.62 | 1221960.12 | 4293510.5 | 1411413.38 | 1347784.75 | 635121.25 | 2277790.25 | 804763.12 | 661980.44 | 2 | 366421.66 | 411752.09 | 298041 | 1972356 | 676263.31 | 2 | 345110 | 267907.12 | 0 | 4 | 1 | 5 | 1 | 5 | 0.184507672 | 0.268369822 | 0.346219242 | 0.503582293 |
| Q11011 | Npepps | Puromycin-sensitive aminopeptidase | 809.73 | 19.24 | 103325 | 14 | 21286894 | 17981266 | 18187018 | 19278726 | 17584040 | 17894802 | 16894598 | 15645107 | 18984584 | 18839842 | 11399520 | 13621685 | 12588147 | 11649918 | 12595574 | 19956640 | 15884341 | 18945772 | 19566078 | 16276044 | 0 | 11 | 0 | 9 | 0 | 11 | 0.655812048 | 0.700833807 | 0.960886881 | 1.026852149 |
| P60670 | Nploc4 | Nuclear protein localization protein 4 homolog | 316.95 | 11.51 | 68017 | 6 | 476110.19 | 1580945.25 | 1198603.38 | 375732.16 | 3969971.25 | 791848.94 | 1068773.62 | 1880249.38 | 2880817 | 773202 | 2003679 | 2551461.75 | 404501.84 | 1497951.75 | 1792848.12 | 594921.5 | 2 | 249794.28 | 634805.12 | 728498.62 | 9 | 6 | 11 | 0 | 11 | 6 | 1.085389988 | 1.115694948 | 0.290477082 | 0.298587435 |
| P13084 | Npm1 | Nucleophosmin | 1204.26 | 56.16 | 32560 | 18 | 210048738 | 109695762 | 175554482 | 190482178 | 145573122 | 208761794 | 218002818 | 160255202 | 161457186 | 176601714 | 218416514 | 221602178 | 166599778 | 90448050 | 178580994 | 226903554 | 232483730 | 186228018 | 202490946 | 194736402 | 10 | 4 | 10 | 6 | 10 | 6 | 1.053278407 | 0.946565412 | 1.254390183 | 1.12730153 |
| Q9CPP0 | Npm3 | Nucleoplasmin-3 | 104.02 | 9.14 | 19023 | 1 | 2 | 400735.5 | 2 | 2 | 141995.56 | 2 | 188897 | 217897.2 | 223805.11 | 113580.45 | 1558095.12 | 1011967.69 | 914635.75 | 1020619.31 | 1013653.88 | 182435.44 | 2 | 245548.44 | 2 | 86724.54 | 0 | 11 | 0 | 9 | 0 | 11 | 10.16877629 | 7.416161006 | 0.948364241 | 0.691648798 |
| P97300 | Nptn | Neuroplastin | 155.46 | 8.56 | 44373 | 3 | 6338515 | 6769829 | 6435087 | 7021597 | 5524188 | 8346771 | 6180451 | 7044575.5 | 7618995 | 9859714 | 10582578 | 9619518 | 11919438 | 11167702 | 13534597 | 6187024 | 7767924.5 | 8093410 | 6846569.5 | 8159578 | 0 | 5 | 1 | 11 | 1 | 11 | 1.770807769 | 1.455136901 | 1.154733914 | 0.948886694 |
| Q9JI75 | Nqo2 | Ribosyldihydronicotinamide dehydrogenase [quinone] | 60.32 | 6.93 | 26248 | 1 | 977400 | 738401.69 | 801079.12 | 842381.69 | 926530.69 | 961878.19 | 1129733.75 | 745140.38 | 939220.12 | 1088188.75 | 223243.05 | 752070 | 483346.5 | 568974.88 | 458673.41 | 1033163.88 | 784627.12 | 1059206.62 | 1163624.62 | 996258.38 | 1 | 4 | 1 | 10 | 1 | 10 | 0.580127815 | 0.511148324 | 1.175250507 | 1.035508575 |
| P08556 | Nras | GTPase NRas | 252.29 | 31.75 | 21199 | 4 | 1505825.88 | 1992710.25 | 1827825.75 | 1627603.38 | 1625730.5 | 1632257.12 | 1799925.75 | 2363048 | 1597107.38 | 1580743.5 | 3801214.5 | 3752262 | 3992486.25 | 4618747 | 4688465.5 | 1851091.5 | 1779815.25 | 1842034.38 | 2 | 1354475.5 | 0 | 11 | 0 | 11 | 0 | 11 | 2.43052619 | 2.323970274 | 0.795764654 | 0.760877792 |
| P47245 | Nrdc | Nardilysin | 297.27 | 8.27 | 132976 | 6 | 943921.69 | 594128.62 | 593431.44 | 464533.5 | 1064315.5 | 291507.94 | 473506.91 | 574166.25 | 1668962.12 | 1179810.25 | 550765.75 | 1284924.25 | 1348820.38 | 1509215.12 | 1590468.88 | 676697.38 | 439176.25 | 503238.19 | 1341703.38 | 940791.75 | 4 | 12 | 7 | 12 | 7 | 12 | 1.716837851 | 1.50054064 | 1.065916502 | 0.931626146 |
| P97333 | Nrp1 | Neuropilin-1 | 428.15 | 9.75 | 103000 | 7 | 845147.62 | 362743.38 | 199757.8 | 415883.88 | 113204.8 | 524876.94 | 545478.56 | 647858.81 | 504211.88 | 493869.06 | 2404788.5 | 2124034.5 | 2040700.38 | 1283572.12 | 1551111 | 952371.5 | 448657.5 | 404888.09 | 851657.06 | 618461.25 | 0 | 4 | 0 | 11 | 0 | 11 | 4.855695001 | 3.462144441 | 1.691522694 | 1.206067492 |
| O35375 | Nrp2 | Neuropilin-2 | 209.45 | 4.19 | 104631 | 4 | 896569.38 | 774227.12 | 1403797 | 625501.81 | 1244303.25 | 1058647.12 | 449272.44 | 774195.94 | 1192028.25 | 556621.25 | 2209377.25 | 2222062 | 2043333 | 2260328.5 | 2114699.5 | 1614882.5 | 425412 | 1159787.25 | 1809376.25 | 1144157.5 | 0 | 9 | 0 | 7 | 0 | 9 | 2.194361987 | 2.691747162 | 1.244562999 | 1.526661936 |
| Q9CR47 | Nsa2 | Ribosome biogenesis protein NSA2 homolog | 55.39 | 6.15 | 30036 | 1 | 417179.56 | 346423.69 | 115346.54 | 351423.19 | 183370.22 | 496158.34 | 423927.38 | 180139.34 | 287259.91 | 173415.61 | 821668.81 | 438538.53 | 567615.25 | 366355.47 | 465856.91 | 230125.67 | 360902.34 | 459852.41 | 353558.91 | 206157.2 | 1 | 8 | 4 | 11 | 4 | 11 | 1.881554564 | 1.704166815 | 1.139242636 | 1.031837998 |
| Q9R1J0 | Nsdhl | Sterol-4-alpha-carboxylate 3-dehydrogenase, decarboxylating | 617.62 | 39.23 | 40686 | 10 | 9952038 | 12261537 | 9401702 | 7892803.5 | 10114061 | 10859824 | 12022310 | 8755560 | 8154216.5 | 9058501 | 11487179 | 11372173 | 13230300 | 15575352 | 16589838 | 13966344 | 12220518 | 12129452 | 11411731 | 12357093 | 2 | 3 | 2 | 1 | 2 | 3 | 1.375491664 | 1.397221434 | 1.251157974 | 1.270923542 |
| P46460 | Nsf | Vesicle-fusing ATPase | 1027.51 | 24.87 | 82613 | 17 | 22363066 | 20965358 | 32809002 | 35373558 | 28531128 | 24349472 | 24603634 | 29078188 | 29350170 | 39861566 | 23855516 | 26181454 | 27173030 | 27869786 | 26478818 | 25872238 | 20194660 | 25951438 | 26732756 | 27903646 | 10 | 8 | 8 | 8 | 10 | 8 | 0.939421736 | 0.893479331 | 0.904404655 | 0.86017476 |
| O35987 | Nsfl1c | NSFL1 cofactor p47 | 436.23 | 28.92 | 40680 | 7 | 4069191.75 | 1169476.88 | 2370348.25 | 1507143.38 | 33549.73 | 3645067.5 | 1990828.75 | 2737113.75 | 1706000.62 | 2918994.5 | 5266786 | 5403880 | 1341750.5 | 2971637.75 | 1768975.12 | 3362079.5 | 3067137.25 | 3354818 | 1371432.12 | 2894198.25 | 6 | 7 | 10 | 10 | 10 | 10 | 1.830990205 | 1.28889235 | 1.535531196 | 1.080909339 |
| Q1HFZ0 | Nsun2 | tRNA (cytosine(34)-C(5))-methyltransferase | 564 | 16.91 | 85452 | 11 | 9735427 | 10858598 | 5841960 | 8799233 | 9504158 | 7408167.5 | 9909605 | 10658448 | 8211530.5 | 9153281 | 9005513 | 9682138 | 10952991 | 6571714.5 | 10659663 | 11050059 | 6021399 | 7424478.5 | 7859928 | 9335637 | 10 | 10 | 10 | 10 | 10 | 10 | 1.047668155 | 1.033766049 | 0.931874899 | 0.91950932 |
| Q9JM14 | Nt5c | 5'(3')-deoxyribonucleotidase, cytosolic type | 295.03 | 38.5 | 23076 | 4 | 11966435 | 11955821 | 8428012 | 9047225 | 4137669.25 | 11616930 | 11192078 | 9704240 | 12608730 | 8915430 | 6105775 | 8636198 | 5152836 | 3368965.5 | 7504954 | 9386367 | 568634.38 | 5845670.5 | 5879917 | 6442842 | 5 | 6 | 0 | 1 | 5 | 6 | 0.675713602 | 0.569396824 | 0.617620087 | 0.520443743 |
| Q3V1L4 | Nt5c2 | Cytosolic purine 5'-nucleotidase | 443.3 | 20.18 | 64809 | 9 | 5484728 | 4252709 | 7509813 | 8759181 | 8108484 | 5417427 | 5520437.5 | 7738427.5 | 8250337 | 7627017 | 4781837.5 | 5615700 | 6172783 | 5370319.5 | 5145547 | 4336530 | 4743261.5 | 6463247 | 6653321 | 6469496.5 | 7 | 8 | 4 | 6 | 7 | 8 | 0.79396906 | 0.783887958 | 0.840273411 | 0.829604378 |
| Q3UFY7 | Nt5c3b | 7-methylguanosine phosphate-specific 5'-nucleotidase | 214.92 | 23.33 | 34425 | 4 | 2394774 | 2455495 | 2476081.5 | 2248555.5 | 3643989.25 | 2435327.75 | 2206629.25 | 2283397.25 | 2797931.5 | 2348677 | 1041307 | 1315714.88 | 1330885.25 | 1288954.12 | 1175893.62 | 2798423 | 2691859.25 | 2287566.25 | 1942872.62 | 1908851.75 | 0 | 9 | 0 | 11 | 0 | 11 | 0.465451519 | 0.509673116 | 0.879768895 | 0.963353939 |
| Q8C5P5 | Nt5dc1 | 5'-nucleotidase domain-containing protein 1 | 48.76 | 2.57 | 53094 | 1 | 307384.69 | 2 | 447487.03 | 326081.56 | 344528.72 | 278940.25 | 389657.06 | 326804.66 | 485621.44 | 2 | 2 | 337390.47 | 304237.91 | 281859.31 | 204223.23 | 388284.03 | 607608.12 | 442999.69 | 2 | 458091.62 | 6.5 | 6.5 | 8.5 | 8.5 | 8.5 | 8.5 | 0.791108788 | 0.761440629 | 1.330765873 | 1.280859496 |
| Q5BJX0 | Ntmt1 | N-terminal Xaa-Pro-Lys N-methyltransferase 1 | 57.15 | 3.14 | 25464 | 1 | 1203289.12 | 1214441.75 | 996393.44 | 1105680 | 1200367.75 | 1211365.75 | 1256591.5 | 791514.12 | 1184055 | 1283295.88 | 1164055.5 | 1424642.88 | 1615731.5 | 891145.31 | 940089.62 | 1342556.25 | 1135871.25 | 1215075.75 | 1317743.75 | 1345202.12 | 12 | 3 | 12 | 6 | 12 | 6 | 1.055154416 | 1.053929133 | 1.111233902 | 1.109943498 |
| Q9R061 | Nubp2 | Cytosolic Fe-S cluster assembly factor NUBP2 | 130.26 | 11.27 | 29518 | 2 | 2140320.5 | 1817808 | 2179952.5 | 2284982.75 | 1962481.12 | 2234874 | 1902162.5 | 2007064.25 | 2291218 | 1911755.62 | 1176419.5 | 1742809.25 | 1881105.88 | 1402613 | 1499760.5 | 1871171 | 2362125 | 2120427 | 2524588 | 1928037.75 | 1 | 11 | 0 | 10 | 1 | 11 | 0.74167588 | 0.744433436 | 1.040518228 | 1.044386883 |
| Q02819 | Nucb1 | Nucleobindin-1 | 177.61 | 8.5 | 53409 | 3 | 500953.97 | 421302.78 | 410834.5 | 717871.44 | 307794.59 | 604045 | 340336.19 | 405246.59 | 523062.5 | 503780.25 | 1203602.12 | 916126.06 | 1552629.75 | 1038532.56 | 1416777 | 348868.31 | 541470.06 | 596151.06 | 2 | 333736.91 | 0 | 10 | 0 | 9 | 0 | 10 | 2.597837235 | 2.578474007 | 0.771689548 | 0.765937687 |
| P81117 | Nucb2 | Nucleobindin-2 | 56.92 | 2.14 | 50304 | 1 | 2227646.75 | 1822346.75 | 1089691.38 | 1209046.5 | 817584.75 | 1545599 | 1720806.62 | 1099201.62 | 1140844.25 | 1260870.88 | 1127426.5 | 1418793 | 1286738.38 | 734427.69 | 2160205 | 1595717.38 | 1164195.12 | 1413226.62 | 999641.12 | 1079182.5 | 12 | 10 | 12 | 9 | 12 | 10 | 0.938779486 | 0.994128874 | 0.872409565 | 0.923845858 |
| Q80XU3 | Nucks1 | Nuclear ubiquitous casein and cyclin-dependent kinase substrate 1 | 145.31 | 11.11 | 26313 | 3 | 11606529 | 12516090 | 13132566 | 20097178 | 10580274 | 13741405 | 10044944 | 13475112 | 11671298 | 17122720 | 8395231 | 6302413.5 | 10457486 | 10489042 | 9489332 | 8067953 | 12490258 | 12249391 | 15531588 | 17033678 | 0 | 12 | 2 | 12 | 2 | 12 | 0.66438617 | 0.68326663 | 0.96231901 | 0.989666096 |
| O35685 | Nudc | Nuclear migration protein nudC | 847.85 | 41.57 | 38358 | 15 | 50607930 | 54291106 | 49094110 | 40203170 | 36052010 | 41775374 | 45680494 | 41383290 | 36090058 | 43495706 | 45988742 | 48320894 | 33299152 | 27606982 | 34469374 | 50288682 | 44753630 | 38617374 | 37071350 | 36877258 | 4 | 8 | 10 | 12 | 10 | 12 | 0.823828548 | 0.910088593 | 0.901671242 | 0.996081908 |
| Q6PIP5 | Nudcd1 | NudC domain-containing protein 1 | 243.73 | 9.11 | 66705 | 5 | 1765560.5 | 1426470.5 | 3041721.5 | 1468120.38 | 1995309.25 | 1615441 | 964541.31 | 1758209.5 | 1400803.75 | 2467015 | 3585880 | 3324700 | 3189778 | 2404625.25 | 3653680.25 | 900858.31 | 1111801.75 | 991375.44 | 3238579.5 | 2223381.5 | 1 | 9 | 1 | 11 | 1 | 11 | 1.666325669 | 1.96912536 | 0.873036763 | 1.031682379 |
| Q5M823 | Nudcd2 | NudC domain-containing protein 2 | 167.75 | 23.57 | 17674 | 3 | 3900363.5 | 4417447.5 | 4734645.5 | 4770043 | 4248829 | 3374329.25 | 967296.94 | 4847373 | 3555804 | 4063488 | 2 | 3378470.75 | 2759369.75 | 3346697.75 | 2693478.5 | 4002302.75 | 900761.94 | 4500749.5 | 4866219 | 3595285.25 | 0 | 9 | 5 | 10 | 5 | 10 | 0.551757396 | 0.724524499 | 0.809435573 | 1.062887252 |
| P56380 | Nudt2 | Bis(5'-nucleosyl)-tetraphosphatase [asymmetrical] | 111.69 | 23.13 | 16989 | 2 | 895758.31 | 716768.12 | 461510.59 | 594085.12 | 577468.75 | 302124.22 | 575965.25 | 464966.84 | 2 | 768278.62 | 274677.91 | 2 | 341203.97 | 396533.16 | 2 | 737703.25 | 491126.34 | 1033468.5 | 1158861.88 | 857927.31 | 0 | 6 | 6 | 3 | 6 | 6 | 0.31193674 | 0.479515621 | 1.318430888 | 2.026719288 |
| Q4KM65 | Nudt21 | Cleavage and polyadenylation specificity factor subunit 5 | 271.43 | 22.47 | 26240 | 5 | 3214062.75 | 2601822.75 | 3005140.5 | 3183400.25 | 2101936.25 | 3716612.5 | 2884948.25 | 2387654.5 | 2554357.75 | 3553367 | 3305185 | 3270167 | 3577469.5 | 3626952.25 | 4055434 | 2965940 | 3731129.5 | 3546442.75 | 2431739.5 | 3795760.5 | 0 | 7 | 6 | 8 | 6 | 8 | 1.26433783 | 1.181378991 | 1.167630014 | 1.091016607 |
| Q9JKX6 | Nudt5 | ADP-sugar pyrophosphatase | 111.46 | 10.09 | 23984 | 2 | 3888450.25 | 4000031 | 2488764.5 | 3098096 | 2550928.5 | 4710536.5 | 4442207.5 | 2970413.25 | 2822938.25 | 3269442.75 | 3656091.75 | 3064425.5 | 2781900.5 | 2421858.5 | 2996478.75 | 4692768.5 | 4430115 | 3336599.5 | 3661888.75 | 3585319.25 | 9 | 6 | 7 | 9 | 9 | 9 | 0.931018557 | 0.819122378 | 1.229649238 | 1.08186158 |
| Q5F2E7 | Nufip2 | Nuclear fragile X mental retardation-interacting protein 2 | 64.57 | 1.88 | 75657 | 1 | 396452.91 | 409327.31 | 336493.75 | 307297.03 | 339079.69 | 636237.69 | 2 | 2 | 455439.81 | 538344.38 | 586940.19 | 708275 | 547975.56 | 493213.88 | 566262.12 | 357064.12 | 2 | 682179.06 | 468984.03 | 641711.75 | 0 | 7 | 5 | 9 | 5 | 9 | 1.622824829 | 1.780748874 | 1.201990401 | 1.318961243 |
| E9Q7G0 | Numa1 | Nuclear mitotic apparatus protein 1 | 2466.11 | 22.21 | 235630 | 40 | 19486082 | 22440948 | 25189754 | 26811114 | 22685102 | 22959490 | 18977908 | 21892120 | 20031708 | 22809720 | 15317192 | 13045566 | 12584389 | 11650330 | 11715564 | 14569204 | 15813670 | 16976060 | 15772472 | 19687302 | 0 | 1 | 0 | 1 | 0 | 1 | 0.551508331 | 0.602910571 | 0.710201333 | 0.77639424 |
| P52590 | Nup107 | Nuclear pore complex protein Nup107 | 129.55 | 2.48 | 107209 | 3 | 264618.94 | 548049.44 | 569377 | 305287.28 | 1070499.25 | 252910.52 | 188552.56 | 752960.56 | 1164661.75 | 309773.25 | 116946.26 | 617373.44 | 308475.75 | 2 | 364207.03 | 649242.69 | 297036.75 | 269348.94 | 316324.56 | 233808.28 | 8 | 8 | 8 | 11 | 8 | 11 | 0.510185003 | 0.527193332 | 0.640271517 | 0.661616615 |
| Q8R0G9 | Nup133 | Nuclear pore complex protein Nup133 | 531.3 | 10.56 | 128620 | 9 | 1443204.5 | 1142987.12 | 2469211.75 | 1808632.75 | 1608601 | 847872.31 | 1560110.62 | 2010400.38 | 1064700.25 | 1273808.25 | 2490051 | 2477286 | 2317309 | 1384915.75 | 2410531 | 1217906 | 1635828.38 | 2035910.38 | 1238685.62 | 768161.88 | 6 | 9 | 2 | 12 | 6 | 12 | 1.307750184 | 1.63982095 | 0.81397234 | 1.020660454 |
| P49791 | Nup153 | Nuclear pore complex protein Nup153 | 435.1 | 7.77 | 152824 | 8 | 1757687.25 | 2034453.25 | 1882636.88 | 1160425.25 | 1419020.25 | 2564808.5 | 1563136 | 1382084.75 | 1919600 | 1360963.25 | 793534.44 | 1660221.5 | 597002.25 | 750862.81 | 1931923.75 | 1493589.12 | 1713479.75 | 1482494.75 | 1673338.75 | 1438272.5 | 6 | 10 | 7 | 12 | 7 | 12 | 0.694619549 | 0.652236439 | 0.945113156 | 0.887445854 |
| Q99P88 | Nup155 | Nuclear pore complex protein Nup155 | 697.24 | 10.93 | 155118 | 13 | 6751792.5 | 5428751.5 | 5891329 | 5829459.5 | 8097353 | 5789014 | 5927092 | 5530573.5 | 6307282 | 8383405.5 | 8842990 | 9840584 | 8150875.5 | 7469547.5 | 7434104 | 5494674 | 5760496.5 | 7030861 | 5558933 | 7446974.5 | 2 | 11 | 3 | 10 | 3 | 11 | 1.304369237 | 1.306873575 | 0.977913265 | 0.97979082 |
| Q9Z0W3 | Nup160 | Nuclear pore complex protein Nup160 | 244.25 | 5.85 | 158232 | 5 | 1152248.38 | 902275 | 1150246.75 | 754167.75 | 1526117 | 852467.5 | 1145140.12 | 1247186 | 1605093.5 | 846823.12 | 912232.5 | 1361285.62 | 1325151.75 | 589118.19 | 894479.75 | 1014101.25 | 760336.62 | 708825.75 | 1065314.75 | 878872.38 | 11 | 6 | 12 | 6 | 12 | 6 | 0.926566447 | 0.892140831 | 0.807184403 | 0.777194304 |
| Q6ZQH8 | Nup188 | Nucleoporin NUP188 homolog | 101.02 | 1.48 | 196696 | 2 | 279351.78 | 2 | 471287.81 | 531176.81 | 537960.5 | 2 | 388604.56 | 500383.41 | 193774.14 | 199249.75 | 435774.62 | 540895.75 | 796859.25 | 370850.94 | 300376.91 | 495052.38 | 174333.81 | 211811.12 | 189815.28 | 610779.81 | 9 | 11 | 5 | 11 | 9 | 11 | 1.343436541 | 1.906966489 | 0.92417403 | 1.311836363 |
| Q9QY81 | Nup210 | Nuclear pore membrane glycoprotein 210 | 436.71 | 5.94 | 204101 | 8 | 1332610.5 | 1035335.38 | 3737213.75 | 2513665.75 | 2436120.25 | 1700809 | 1854578.5 | 2634206.75 | 2992661.5 | 1525478.25 | 2285171.5 | 1913516.25 | 3517606 | 4103474 | 1905868 | 1127992.38 | 1445616.75 | 3209810 | 2350412.5 | 2158621.75 | 10 | 11 | 6 | 11 | 10 | 11 | 1.241583289 | 1.281843175 | 0.931027047 | 0.96121676 |
| Q80U93 | Nup214 | Nuclear pore complex protein Nup214 | 321.78 | 4.22 | 212979 | 6 | 786712.75 | 680023.25 | 927063.31 | 973475.12 | 2 | 611872.69 | 977563.25 | 729538.56 | 715454.88 | 1007867 | 1642213.38 | 841907.62 | 1045734.88 | 222622.94 | 2 | 2 | 2 | 738442.94 | 1858535 | 1395957.5 | 10.5 | 12 | 12 | 12 | 12 | 12 | 1.114396426 | 0.928304228 | 1.185806845 | 0.987789876 |
| Q8R4R6 | Nup35 | Nucleoporin NUP35 | 135.95 | 8.62 | 34786 | 2 | 302388.53 | 917476.81 | 775799.94 | 200763.34 | 205802.42 | 929096.88 | 1094714 | 552278.25 | 375180.56 | 355140.91 | 728136.19 | 1370821.88 | 650536.25 | 1210397.75 | 1223798.62 | 595289.88 | 2 | 914963.88 | 866250.75 | 799076.38 | 4 | 10 | 4 | 12 | 4 | 12 | 2.157865169 | 1.567769801 | 1.321930671 | 0.960432104 |
| Q9CWU9 | Nup37 | Nucleoporin Nup37 | 88.65 | 4.91 | 36732 | 1 | 269038.53 | 293740.59 | 254736.52 | 447784.03 | 322749.34 | 387073.19 | 2 | 2 | 447276.91 | 426401.31 | 422034.75 | 2 | 366765.47 | 632939.25 | 393301.47 | 381940.06 | 620359.31 | 460073.78 | 2 | 435246.56 | 8 | 7 | 11 | 8 | 11 | 8 | 1.142938869 | 1.439647156 | 1.194939009 | 1.505146593 |
| P59235 | Nup43 | Nucleoporin Nup43 | 66.94 | 3.68 | 41990 | 1 | 664322.62 | 863178 | 559149 | 627271 | 515326.72 | 668557.69 | 1110671.75 | 719592.31 | 328418.31 | 2 | 696637.88 | 1018209.12 | 903352.81 | 2 | 783771.88 | 700510.69 | 750286.94 | 2 | 771754.31 | 599108.88 | 7 | 11 | 9.5 | 11.5 | 9.5 | 11.5 | 1.053488114 | 1.203283489 | 0.873783431 | 0.998026614 |
| Q9JIH2 | Nup50 | Nuclear pore complex protein Nup50 | 212.49 | 12.45 | 49485 | 3 | 1528621.5 | 1525921.62 | 1210807.38 | 1137139 | 1043149.75 | 2146894.5 | 1037532.62 | 310899.41 | 1229889.38 | 1951226.25 | 1644181.62 | 1679623.25 | 1043651.88 | 1096700.75 | 1153684 | 1788879.75 | 1060204.62 | 522886.5 | 1370203.62 | 1102540.62 | 11 | 10 | 12 | 11 | 12 | 11 | 1.026716086 | 0.991222771 | 0.906770436 | 0.875423612 |
| P70582 | Nup54 | Nuclear pore complex protein Nup54 | 48.21 | 1.37 | 55745 | 1 | 2 | 2 | 2 | 2 | 2 | 2 | 2 | 2 | 2 | 2 | 455080.44 | 449445.91 | 2 | 2 | 2 | 2 | 2 | 2 | 2 | 2 | 7.5 | 12.5 | 7.5 | 12.5 | 7.5 | 12.5 | 90453.235 | 90453.235 | 1 | 1 |
| Q8R332 | Nup58 | Nucleoporin p58/p45 | 72.44 | 2.21 | 59445 | 1 | 2 | 2 | 2 | 2 | 2 | 2 | 2 | 2 | 2 | 2 | 2 | 187776.58 | 2 | 2 | 2 | 2 | 2 | 2 | 2 | 2 | 10 | 12.5 | 10 | 12.5 | 10 | 12.5 | 18778.458 | 18778.458 | 1 | 1 |
| O08658 | Nup88 | Nuclear pore complex protein Nup88 | 136.47 | 5.26 | 83585 | 2 | 773365.19 | 599666.12 | 1155609.25 | 735809.06 | 880991.06 | 878873.5 | 658673.75 | 780702.38 | 829677.69 | 1061303.62 | 840508.88 | 905784.12 | 575733.62 | 783126.94 | 674284.5 | 755438.38 | 835299.88 | 1170968.5 | 1095757.38 | 2 | 11 | 11 | 10 | 11 | 11 | 11 | 0.911709599 | 0.897892778 | 0.930532225 | 0.91643015 |
| Q8BJ71 | Nup93 | Nuclear pore complex protein Nup93 | 807.17 | 21.73 | 93281 | 15 | 18377648 | 19610594 | 18792608 | 18058746 | 15350537 | 16603451 | 18744664 | 15823840 | 15599971 | 17880754 | 16731226 | 19641012 | 18550106 | 13833926 | 17760068 | 16066949 | 15296228 | 15932976 | 14454636 | 13729195 | 10 | 2 | 10 | 4 | 10 | 4 | 0.95926611 | 1.022015346 | 0.836898466 | 0.891643171 |
| P49793 | Nup98 | Nuclear pore complex protein Nup98-Nup96 | 226.3 | 4.35 | 197283 | 5 | 1432063.25 | 801046.88 | 851889.62 | 1143876 | 1559316 | 1785424 | 1426701 | 1626138.62 | 1596025.5 | 1645104.75 | 1659691.38 | 1608383.25 | 1656565.75 | 1601710.38 | 1556518.12 | 1303273.38 | 659616.75 | 2026962.5 | 1230404.5 | 1225099.75 | 1 | 11 | 12 | 5 | 12 | 11 | 1.396441104 | 1.000430108 | 1.11353548 | 0.797752527 |
| Q9DBY8 | Nvl | Nuclear valosin-containing protein-like | 50.84 | 1.05 | 94476 | 1 | 506069 | 479720.62 | 512455.62 | 429886.25 | 470051.75 | 364655.75 | 518744.88 | 2 | 408759.62 | 393650.88 | 2 | 453393.28 | 2 | 300903.59 | 406606.19 | 2 | 2 | 344031.16 | 2 | 330678.12 | 1 | 0 | 9 | 3.5 | 9 | 3.5 | 0.484077714 | 0.688633301 | 0.28134434 | 0.400231359 |
| Q99JX7 | Nxf1 | Nuclear RNA export factor 1 | 416.65 | 16.67 | 70300 | 7 | 2071560.88 | 2690585.5 | 2026344.38 | 1546525.25 | 1660468.38 | 2194747.5 | 1634368.75 | 1408116 | 1583196.5 | 2175942 | 2801078 | 2562443.5 | 2712958.5 | 953370 | 2120350.5 | 1597022.38 | 3372273.5 | 1971531.25 | 2699200.5 | 1291000.88 | 7 | 12 | 7 | 10 | 7 | 12 | 1.115523777 | 1.239410959 | 1.093596677 | 1.215048692 |
| P11928 | Oas1a | 2'-5'-oligoadenylate synthase 1A | 631.77 | 37.87 | 42429 | 10 | 3303631 | 3524393.25 | 3408435.5 | 3377048.25 | 4853134.5 | 3590256.75 | 5291204 | 4240534 | 5202379.5 | 5104196 | 10773100 | 10577076 | 12851542 | 5960533.5 | 10445943 | 3179996.5 | 2571737.25 | 3008893.5 | 3019295.75 | 3160463.75 | 0 | 0 | 0 | 0 | 0 | 0 | 2.740519534 | 2.160105971 | 0.809047273 | 0.637699467 |
| E9Q9A9 | Oas2 | 2'-5'-oligoadenylate synthase 2 | 112.78 | 4.18 | 85051 | 2 | 284571.09 | 271470.75 | 220109.28 | 2 | 83518.27 | 371825.19 | 351894.5 | 390447.16 | 447291.75 | 362611.12 | 638630.06 | 698067.88 | 883911.31 | 535685.81 | 875197.31 | 181552.08 | 280604.59 | 263616.75 | 285253.91 | 409073.22 | 0 | 6 | 0 | 4 | 0 | 6 | 4.224279663 | 1.887401653 | 1.651910912 | 0.738071253 |
| Q8VI93 | Oas3 | 2'-5'-oligoadenylate synthase 3 | 927.19 | 19.33 | 126333 | 17 | 11612216 | 11760700 | 12480764 | 14431957 | 12397756 | 12092327 | 11576338 | 11575610 | 16958602 | 12467671 | 9233440 | 9120158 | 11011727 | 7660579 | 8836050 | 12478716 | 8774175 | 12345639 | 14349392 | 14873975 | 0 | 11 | 0 | 10 | 0 | 11 | 0.731644409 | 0.70916291 | 1.00220958 | 0.971414329 |
| Q8VI94 | Oasl1 | 2'-5'-oligoadenylate synthase-like protein 1 | 846.77 | 42.86 | 59088 | 14 | 1779162.75 | 1644981.5 | 787050.81 | 2298811.75 | 2646937.25 | 1232091.12 | 2124029.75 | 1673425 | 3238600 | 1647785.88 | 4303419 | 3784448.25 | 7181445.5 | 2712969.5 | 6314237 | 860258.38 | 1354482.38 | 1408830 | 1838586.75 | 693515.62 | 0 | 6 | 1 | 5 | 1 | 6 | 2.653343636 | 2.450250754 | 0.672240989 | 0.620786154 |
| P29758 | Oat | Ornithine aminotransferase, mitochondrial | 1469.72 | 56.95 | 48355 | 22 | 47140514 | 51812810 | 67476250 | 55835770 | 60915078 | 39131574 | 47746034 | 70522890 | 63422630 | 67415578 | 53483274 | 53627326 | 75202898 | 70499082 | 68666666 | 43384234 | 42213018 | 69650210 | 61480222 | 60382934 | 6 | 12 | 8 | 10 | 8 | 12 | 1.13524531 | 1.115322957 | 0.978565594 | 0.961392805 |
| Q4PJX1 | Odr4 | Protein odr-4 homolog | 106.92 | 6.94 | 49998 | 2 | 301044.28 | 315203.06 | 2 | 416859.5 | 472236.91 | 289300.12 | 386600.44 | 2 | 506023.19 | 2 | 2 | 397563.06 | 2 | 448690.19 | 2 | 552151.81 | 2 | 2 | 2 | 479233.88 | 8.5 | 11.5 | 11 | 12 | 11 | 12 | 0.562169355 | 0.715999138 | 0.68515269 | 0.872635142 |
| Q8VIJ5 | Oga | Protein O-GlcNAcase | 51.03 | 1.42 | 102918 | 1 | 441766.81 | 503477.69 | 440238.34 | 632116.88 | 446588.88 | 391368.88 | 314422.81 | 567311.62 | 455505.19 | 486277.09 | 497636.59 | 508030.16 | 728950.5 | 2 | 539310.19 | 448056.34 | 642341.69 | 607235 | 449581.97 | 477942.84 | 9 | 7 | 8 | 8 | 9 | 8 | 0.922790342 | 1.026657743 | 1.065323425 | 1.18523406 |
| Q60597 | Ogdh | 2-oxoglutarate dehydrogenase, mitochondrial | 899.55 | 18.96 | 116449 | 15 | 16460358 | 18108878 | 17075046 | 15700284 | 19487298 | 16409782 | 20262176 | 21279712 | 24133466 | 19943186 | 18205374 | 23252296 | 22778520 | 24820282 | 26415936 | 17354158 | 13808630 | 17271410 | 16594280 | 18583412 | 1 | 12 | 6 | 4 | 6 | 12 | 1.329839101 | 1.131768177 | 0.962917138 | 0.819496865 |
| Q99PG2 | Ogfr | Opioid growth factor receptor | 449.42 | 27.01 | 70679 | 8 | 12223976 | 12846410 | 9481649 | 11222715 | 9249974 | 11736069 | 10407592 | 9355401 | 9094966 | 11035387 | 10545671 | 8096280.5 | 6345851.5 | 5285070 | 6827851.5 | 14814524 | 14180413 | 10158920 | 10835556 | 11241413 | 2 | 8 | 3 | 6 | 3 | 8 | 0.674255531 | 0.718596647 | 1.112787517 | 1.185967844 |
| Q8CGY8 | Ogt | UDP-N-acetylglucosamine--peptide N-acetylglucosaminyltransferase 110 kDa subunit | 564.02 | 10.52 | 116952 | 9 | 4903246.5 | 6481749 | 10862972 | 8923585 | 8787079 | 6723616 | 7380382 | 6055889 | 7341055.5 | 8781219 | 4038127.5 | 5846657.5 | 6227049.5 | 3723606.5 | 4272579 | 5409177.5 | 6388400 | 7946733 | 8438842 | 6555290.5 | 2 | 8 | 1 | 10 | 2 | 10 | 0.603324466 | 0.66445931 | 0.869360178 | 0.957452411 |
| A0JPJ7 | Ola1 | Obg-like ATPase 1 | 423.65 | 21.46 | 44535 | 7 | 16964240 | 17444504 | 14822880 | 14542475 | 12675916 | 14965535 | 16174430 | 13813447 | 15002076 | 15149801 | 13282431 | 15557288 | 12323597 | 11675438 | 10692564 | 16317019 | 13414496 | 10300871 | 14140500 | 11545147 | 4 | 5 | 4 | 6 | 4 | 6 | 0.831017731 | 0.845896725 | 0.85962093 | 0.875012051 |
| Q5RKH6 | Os9 | Protein OS-9 | 61.35 | 3.9 | 75365 | 1 | 2 | 2 | 2 | 347917 | 2 | 268120 | 172662.64 | 2 | 2 | 2 | 2 | 2 | 2 | 2 | 2 | 2 | 2 | 296173.81 | 2 | 2 | 10 | 12 | 7.5 | 11 | 10 | 12 | 2.874183E-05 | 2.268661E-05 | 0.851280621 | 0.671936123 |
| Q3B7Z2 | Osbp | Oxysterol-binding protein 1 | 41.86 | 2.11 | 88797 | 1 | 1077185.38 | 1825117 | 1188061.25 | 1463105.75 | 1103051.5 | 1818394.62 | 1166489.38 | 2 | 1095719.88 | 1045629 | 1160171.12 | 857702.75 | 750624.56 | 1083346.12 | 932686.5 | 1165765.5 | 2005669.12 | 1617770.88 | 1536918.38 | 2 | 3 | 10 | 8 | 8.5 | 8 | 10 | 0.718773536 | 0.933342143 | 0.950365212 | 1.234068674 |
| Q8BX94 | Osbpl2 | Oxysterol-binding protein-related protein 2 | 96.84 | 6.82 | 55384 | 2 | 1239089.12 | 1160142.25 | 597554.5 | 590068.56 | 832064.62 | 783313.31 | 1144709.62 | 641019.19 | 1118895.12 | 257861.62 | 248333.5 | 370578.09 | 501892.12 | 2 | 45503.41 | 1040189.31 | 442366.25 | 642248.5 | 2 | 810059.69 | 0 | 7 | 2 | 9 | 2 | 9 | 0.263935389 | 0.295582507 | 0.664159202 | 0.743795073 |
| B9EJ86 | Osbpl8 | Oxysterol-binding protein-related protein 8 | 1190.42 | 26.1 | 101269 | 18 | 18057228 | 19067434 | 14192709 | 14005068 | 17049184 | 17167398 | 21524196 | 18747644 | 21728194 | 20761762 | 25207212 | 29354882 | 32024490 | 23710978 | 29522204 | 18197060 | 13415282 | 19141312 | 17938698 | 17556794 | 0 | 10 | 0 | 5 | 0 | 10 | 1.697426382 | 1.399188369 | 1.047073529 | 0.863102588 |
| A2A8Z1 | Osbpl9 | Oxysterol-binding protein-related protein 9 | 52.35 | 1.36 | 83109 | 1 | 1319765.25 | 1187311 | 1001321.88 | 1124327.38 | 1007284.69 | 1169712.12 | 1256703.25 | 1030091.62 | 1269390.5 | 1143731.88 | 1293103.25 | 967219.88 | 1090826.75 | 879443 | 1040652.19 | 1077693.75 | 1412491.62 | 755606.75 | 1008768.69 | 967678.88 | 8 | 9 | 7 | 6 | 8 | 9 | 0.93461623 | 0.898054159 | 0.925927349 | 0.889705186 |
| Q8BWU5 | Osgep | Probable tRNA N6-adenosine threonylcarbamoyltransferase | 90.27 | 7.46 | 36301 | 2 | 2147302.5 | 1410814 | 980758.5 | 1914001 | 1605290 | 1634419.12 | 1528404.88 | 979316.44 | 1733378.62 | 1733171 | 491793.88 | 1464403.75 | 1147061.25 | 2 | 1361440 | 1985545.5 | 1835981.75 | 2174308 | 1805086 | 1848179.5 | 4 | 7 | 3 | 0 | 4 | 7 | 0.554059184 | 0.58678969 | 1.19743137 | 1.268168459 |
| Q62422 | Ostf1 | Osteoclast-stimulating factor 1 | 226.09 | 24.65 | 23783 | 4 | 10791940 | 11326515 | 13422198 | 13041645 | 14544777 | 9333125 | 10519160 | 13770844 | 12994330 | 12864358 | 6615299.5 | 7737310.5 | 5831431.5 | 7569145 | 8680595 | 11726163 | 10168663 | 12744627 | 13129828 | 14365744 | 0 | 11 | 0 | 11 | 0 | 11 | 0.577149844 | 0.612519646 | 0.984284873 | 1.044605362 |
| B2RYG6 | Otub1 | Ubiquitin thioesterase OTUB1 | 575.62 | 32.47 | 31270 | 8 | 8595271 | 10289466 | 12230402 | 7348640 | 20404588 | 6200067.5 | 11351350 | 12137798 | 17574936 | 6795789 | 4120724.75 | 6214604 | 4160615.5 | 3441791.75 | 4329821 | 8567423 | 6626194.5 | 7898143.5 | 9136252 | 8307079 | 0 | 5 | 1 | 9 | 1 | 9 | 0.378260144 | 0.411904948 | 0.688571708 | 0.749817547 |
| B2RRE7 | Otud4 | OTU domain-containing protein 4 | 53.34 | 0.72 | 123055 | 1 | 785472.25 | 586863.94 | 1028371.12 | 854736.19 | 709026.44 | 637968.5 | 646698.19 | 731684.75 | 868945.38 | 764413.19 | 2 | 305291.53 | 506463.06 | 263428.81 | 511795.75 | 1216086.75 | 457646.59 | 644749.94 | 611065.12 | 863767.19 | 0 | 11 | 0 | 10 | 0 | 11 | 0.400300967 | 0.434823903 | 0.956827936 | 1.039347121 |
| Q3UCV8 | Otulin | Ubiquitin thioesterase otulin | 83.21 | 5.4 | 40320 | 2 | 595004.19 | 757599 | 2784771 | 1494178.5 | 2963685.5 | 2 | 1017430.88 | 2928645.75 | 3069196.75 | 2329529 | 1129293.62 | 2283874.5 | 1694214.38 | 1172474.38 | 2215538 | 849519.44 | 496611.97 | 1275046.38 | 1270115.25 | 1775571.75 | 12 | 9 | 10 | 8 | 12 | 9 | 0.988383881 | 0.909103555 | 0.659302821 | 0.606418771 |
| Q9D7E3 | Ovca2 | Esterase OVCA2 | 124.9 | 10.67 | 24246 | 2 | 716943.06 | 544077.12 | 950117 | 968451.88 | 808651.88 | 934359.5 | 523190.09 | 711085.81 | 503404.72 | 777657.06 | 1156646.25 | 784171.44 | 1117860.5 | 663384.5 | 884708.62 | 329143.56 | 793449.62 | 678366.5 | 774875.69 | 988483.38 | 9 | 10 | 5 | 11 | 9 | 11 | 1.155088516 | 1.335413246 | 0.893706976 | 1.03322656 |
| Q9D0K2 | Oxct1 | Succinyl-CoA:3-ketoacid coenzyme A transferase 1, mitochondrial | 1318.39 | 46.92 | 55989 | 17 | 50067230 | 58845398 | 67399506 | 67760330 | 58034822 | 54887298 | 50623390 | 64081102 | 58853890 | 64465694 | 43597702 | 42002494 | 56519286 | 61944310 | 56878834 | 47722706 | 52045430 | 61214082 | 62869058 | 62866982 | 5 | 10 | 7 | 10 | 7 | 10 | 0.863741585 | 0.890858632 | 0.94906105 | 0.97885669 |
| Q6P9R2 | Oxsr1 | Serine/threonine-protein kinase OSR1 | 84.35 | 3.8 | 58214 | 2 | 739846.31 | 899719.56 | 864014.62 | 503643.66 | 1333672.75 | 1056420.25 | 824048.88 | 157491.7 | 688381.56 | 501616.5 | 2539005.25 | 1556091.62 | 799448.56 | 571733.75 | 1090789.25 | 378975.88 | 2 | 2 | 128985.12 | 346824.44 | 8 | 0 | 5 | 2 | 8 | 2 | 1.510533095 | 2.031335793 | 0.196915398 | 0.264808032 |
| P51577 | P2rx4 | P2X purinoceptor 4 | 50.61 | 2.06 | 43501 | 1 | 386390.38 | 356502.78 | 2 | 491816.22 | 445528.69 | 555257.94 | 446537.91 | 572563.75 | 802422.62 | 381417.38 | 2 | 220975.92 | 568998.81 | 386747.03 | 2 | 337038.28 | 2 | 598989.25 | 617489.06 | 264068.78 | 10 | 12.5 | 4 | 8 | 10 | 12.5 | 0.700331923 | 0.426628211 | 1.081742664 | 0.658976011 |
| P54001 | P4ha1 | Prolyl 4-hydroxylase subunit alpha-1 | 604.7 | 25.47 | 60898 | 12 | 6783454 | 7883781.5 | 6200231.5 | 4259383.5 | 4676605 | 7030251 | 7911910 | 7997062.5 | 6016769 | 7840461.5 | 9760440 | 6631236 | 9944107 | 9463838 | 11139459 | 7841185.5 | 7333151 | 8202862 | 8174027.5 | 8727136 | 2 | 2 | 4 | 5 | 4 | 5 | 1.574954287 | 1.275641397 | 1.351466175 | 1.094626183 |
| Q60716 | P4ha2 | Prolyl 4-hydroxylase subunit alpha-2 | 103.52 | 4.1 | 61002 | 2 | 133820.75 | 2 | 2 | 2 | 2 | 606676.12 | 44082.44 | 2 | 2 | 2 | 558800.94 | 666648.06 | 106678.48 | 499590.25 | 643076.75 | 77765.08 | 248937.81 | 341136 | 2 | 337363.53 | 1 | 4 | 3 | 7.5 | 3 | 7.5 | 18.49224834 | 3.802902973 | 7.511124628 | 1.544651448 |
| P09103 | P4hb | Protein disulfide-isomerase | 2224.86 | 59.53 | 57058 | 33 | 114519482 | 150233922 | 132855250 | 122891650 | 143822258 | 131357498 | 123295634 | 152910850 | 151538914 | 161825010 | 182502898 | 181850354 | 220581986 | 169958370 | 211269858 | 116546122 | 98607234 | 156125186 | 153432226 | 162141378 | 0 | 9 | 0 | 12 | 0 | 12 | 1.454358953 | 1.340166552 | 1.033913622 | 0.952733471 |
| P50580 | Pa2g4 | Proliferation-associated protein 2G4 | 1077.62 | 56.85 | 43699 | 19 | 43673270 | 48364138 | 56196254 | 46070370 | 52572266 | 39048214 | 44578074 | 52446130 | 48101746 | 46540934 | 62017650 | 65036494 | 48717034 | 41500602 | 58209294 | 51417530 | 36120918 | 44677886 | 42955926 | 51813066 | 7 | 7 | 5 | 11 | 7 | 11 | 1.115866838 | 1.194031411 | 0.919429398 | 0.983833862 |
| P29341 | Pabpc1 | Polyadenylate-binding protein 1 | 1650.55 | 38.21 | 70671 | 27 | 71764874 | 73229554 | 71573130 | 72963914 | 59216166 | 77559842 | 72697714 | 63396762 | 58850238 | 67621882 | 123755802 | 107136274 | 102696002 | 77130242 | 102711738 | 71345338 | 76383450 | 72626754 | 71635618 | 71347490 | 0 | 12 | 1 | 9 | 1 | 12 | 1.472210854 | 1.509527048 | 1.041838311 | 1.068245833 |
| Q8CCS6 | Pabpn1 | Polyadenylate-binding protein 2 | 244.14 | 16.89 | 32297 | 4 | 4768577.5 | 5721077 | 7236725.5 | 7407595.5 | 5324043.5 | 7191275 | 7207758 | 5231805 | 4890943.5 | 6822913 | 6623069 | 5524298 | 4475106.5 | 5262973 | 4948419.5 | 6300692.5 | 7228545.5 | 7613383 | 7605356.5 | 6280941.5 | 7 | 6 | 7 | 6 | 7 | 6 | 0.881011533 | 0.856089569 | 1.150072137 | 1.117539014 |
| Q9QY17 | Pacsin2 | Protein kinase C and casein kinase substrate in neurons 2 protein | 112.87 | 4.1 | 55978 | 2 | 1900056.88 | 2284084.75 | 2416087 | 2735326.25 | 3706764.5 | 1412920.25 | 2687535.5 | 3064914.5 | 3263755.75 | 1859532 | 1028806.88 | 2542706.5 | 1533250.5 | 1037301.75 | 1837797.25 | 3313848.75 | 2136332.75 | 2094559.5 | 2339056.25 | 3241445.75 | 3 | 12 | 4 | 10 | 4 | 12 | 0.611843848 | 0.649368131 | 1.006358042 | 1.068077816 |
| P63004 | Pafah1b1 | Platelet-activating factor acetylhydrolase IB subunit alpha | 611.61 | 26.83 | 46670 | 12 | 22615398 | 27050030 | 21590626 | 20046706 | 26072654 | 21012262 | 30352936 | 24552222 | 27240390 | 22228242 | 20436296 | 26717920 | 19580954 | 19086600 | 22528222 | 22637610 | 15578638 | 21982564 | 21425958 | 23917260 | 7 | 9 | 5 | 6 | 7 | 9 | 0.923106367 | 0.86413114 | 0.899183453 | 0.841736607 |
| O35264 | Pafah1b2 | Platelet-activating factor acetylhydrolase IB subunit beta | 91.81 | 12.23 | 25581 | 2 | 2 | 2 | 234532.31 | 2 | 730701.81 | 2 | 95532.77 | 199484.05 | 379391.56 | 2 | 1654062.38 | 1318293.5 | 619150.81 | 406501.09 | 973710.5 | 168516.06 | 2 | 163807.89 | 824858.06 | 88024.42 | 2 | 9.5 | 0 | 11 | 2 | 11 | 5.15075801 | 7.37192618 | 1.290050428 | 1.846360575 |
| O35263 | Pafah1b3 | Platelet-activating factor acetylhydrolase IB subunit gamma | 66.02 | 3.88 | 25863 | 1 | 2 | 2 | 2 | 2 | 2 | 2 | 2 | 2 | 2 | 2 | 413379.94 | 671812.25 | 756690.94 | 574144.12 | 618646.88 | 2 | 2 | 2 | 2 | 2 | 0 | 12.5 | 0 | 12.5 | 0 | 12.5 | 303467.413 | 303467.413 | 1 | 1 |
| Q9DCL9 | Paics | Multifunctional protein ADE2 | 941.85 | 33.65 | 47006 | 16 | 21599990 | 25329978 | 35616930 | 26412454 | 24926272 | 19765264 | 20783570 | 27306412 | 24724570 | 25355338 | 19768326 | 20947298 | 25108446 | 24540136 | 20200526 | 23518234 | 21178644 | 24707818 | 23369598 | 23745708 | 3 | 4 | 9 | 10 | 9 | 10 | 0.825814816 | 0.937504453 | 0.870295096 | 0.988000592 |
| Q9DCE5 | Pak1ip1 | p21-activated protein kinase-interacting protein 1 | 202.72 | 13.61 | 42116 | 4 | 1907105.88 | 2016600.75 | 2231103.25 | 1124648 | 1485698 | 2437194 | 1045348.69 | 1950239 | 1896721.75 | 1929147.75 | 5736728 | 3492175 | 3144513 | 2827822 | 4486520 | 1369812.75 | 2432455.5 | 1828401.38 | 2121435.5 | 1377698.12 | 0 | 12 | 0 | 11 | 0 | 12 | 2.246138947 | 2.126417509 | 1.041601926 | 0.986083509 |
| Q8CIN4 | Pak2 | Serine/threonine-protein kinase PAK 2 | 517.42 | 26.72 | 57930 | 10 | 6675372.5 | 5027040 | 8720476 | 5088353.5 | 4264764 | 7471475 | 6626016 | 4561763 | 3953846.5 | 7156757.5 | 7198319.5 | 7690433 | 6980808.5 | 7594891 | 5473699 | 4274235 | 6607245 | 5601391.5 | 4993460 | 3959370.75 | 6 | 8 | 6 | 8 | 6 | 8 | 1.173365931 | 1.173608252 | 0.854234858 | 0.854411272 |
| Q8K4K6 | Pank1 | Pantothenate kinase 1 | 58 | 1.64 | 60091 | 1 | 1695302.62 | 924502.94 | 1168101.62 | 1171353.75 | 112944.06 | 192098.38 | 1073594.62 | 934182.31 | 2 | 2 | 1128621.88 | 989238.56 | 261287.89 | 1278981.75 | 175516.52 | 1275888.75 | 2149131.5 | 1333395.25 | 1291431.38 | 2 | 10 | 8 | 6 | 4 | 10 | 8 | 0.755814603 | 1.742662237 | 1.192745343 | 2.750082176 |
| Q80YV4 | Pank4 | Pantothenate kinase 4 | 72.2 | 1.34 | 91522 | 1 | 526074.69 | 372414.56 | 2 | 2 | 543736.12 | 2 | 550251.44 | 2 | 2 | 306316.25 | 2 | 2 | 2 | 2 | 109117.6 | 480586.44 | 2 | 312611.25 | 175937.03 | 2 | 6 | 9 | 9 | 11 | 9 | 11 | 0.075664525 | 0.127397796 | 0.67197267 | 1.13141313 |
| Q60967 | Papss1 | Bifunctional 3'-phosphoadenosine 5'-phosphosulfate synthase 1 | 37.44 | 3.21 | 70794 | 1 | 1169747.5 | 1370697.5 | 1508403.38 | 671089.88 | 1829409 | 503624.75 | 1666178.88 | 1688353.5 | 2117691.5 | 2 | 1156556.12 | 877087.69 | 383241.09 | 386306.53 | 479412.69 | 1190845.38 | 935853.81 | 1698126.88 | 1547700.5 | 1608543.88 | 2 | 10 | 7 | 12 | 7 | 12 | 0.501210883 | 0.549311608 | 1.065918507 | 1.168213679 |
| Q99LX0 | Park7 | Protein/nucleic acid deglycase DJ-1 | 722.07 | 66.14 | 20021 | 11 | 78694658 | 64264302 | 66941346 | 68440810 | 50164094 | 79454594 | 58696734 | 68228786 | 52849882 | 70853074 | 37760494 | 37969918 | 31671824 | 45796206 | 34576822 | 82006202 | 106898666 | 74054738 | 74932866 | 76516146 | 0 | 3 | 0 | 3 | 0 | 3 | 0.571605132 | 0.568872751 | 1.261497856 | 1.255467655 |
| Q5XJY4 | Parl | Presenilins-associated rhomboid-like protein, mitochondrial | 73.21 | 5.31 | 41964 | 1 | 247625.06 | 402403.19 | 323123.53 | 2 | 476489.19 | 556509.94 | 335469.06 | 490533.28 | 410605.81 | 574282.56 | 352757.59 | 710369.88 | 2 | 599278.5 | 458450.91 | 278851.19 | 2 | 436690.75 | 501966.56 | 600110.62 | 7.5 | 8.5 | 12 | 10 | 12 | 10 | 1.463021533 | 0.895859719 | 1.253840537 | 0.767770812 |
| P11103 | Parp1 | Poly [ADP-ribose] polymerase 1 | 851.19 | 17.37 | 113100 | 12 | 3566856 | 3406748.25 | 3686721.5 | 3498917.25 | 3322321.75 | 1694144.5 | 3213946.75 | 2753788.25 | 932581.5 | 3603813 | 1925451.5 | 2510681.75 | 2023688.62 | 1415286.25 | 2137305.5 | 3465690.25 | 2004933.25 | 2668830.25 | 2232173.5 | 2578023.75 | 0 | 2 | 9 | 12 | 9 | 12 | 0.572741271 | 0.820805765 | 0.740760406 | 1.061596993 |
| Q8BZ20 | Parp12 | Poly [ADP-ribose] polymerase 12 | 39.27 | 1.41 | 79917 | 1 | 997237.25 | 943314.12 | 1128986.62 | 967471.5 | 648146.69 | 910874.69 | 756951.12 | 842226.81 | 571179.25 | 640400.5 | 568933 | 859919.25 | 579979.5 | 424915.5 | 2 | 2 | 535448.38 | 720009.88 | 758446.69 | 625398.5 | 1 | 2 | 5 | 6 | 5 | 6 | 0.519459578 | 0.653946712 | 0.563333504 | 0.709179518 |
| Q2EMV9 | Parp14 | Poly [ADP-ribose] polymerase 14 | 305.67 | 4.95 | 203802 | 7 | 2574537.75 | 2469461.5 | 2792454.25 | 3196602.25 | 3465875.25 | 3046859.75 | 3101513.25 | 3046613.5 | 2921148.5 | 2739483 | 2730727 | 1868576.25 | 2056346.25 | 1251089.75 | 1731579 | 2384689.75 | 2042739.75 | 2743162.5 | 4118711.5 | 2952412 | 2 | 10 | 0 | 8 | 2 | 10 | 0.664760612 | 0.648799548 | 0.982259692 | 0.958675398 |
| Q8CAS9 | Parp9 | Poly [ADP-ribose] polymerase 9 | 226.61 | 6.47 | 96659 | 4 | 2867777.75 | 969906.94 | 2 | 3074087 | 3140556 | 3069752 | 2646412.5 | 1751162.38 | 2502295 | 2788115.25 | 2 | 1411346.25 | 2 | 1909249 | 2 | 1235273.38 | 3236266 | 1070194.12 | 1655945.75 | 2757184 | 5.5 | 12 | 1 | 8 | 5.5 | 12 | 0.33033151 | 0.260281366 | 0.990304094 | 0.780300076 |
| Q9D0B6 | Pbdc1 | Protein PBDC1 | 270.14 | 29.29 | 22223 | 5 | 4998629 | 4162354.25 | 5973530 | 4239568 | 3444856.5 | 3305390.25 | 4477362.5 | 4795944.5 | 6557477 | 3937811.5 | 876102.75 | 3426222 | 2267201.25 | 584249.31 | 2455201 | 4301589 | 996218.75 | 4003159.75 | 3221182 | 4239598 | 0 | 7 | 1 | 6 | 1 | 7 | 0.421096565 | 0.41644198 | 0.734554241 | 0.726434855 |
| Q3TVI8 | Pbxip1 | Pre-B-cell leukemia transcription factor-interacting protein 1 | 344.86 | 9.9 | 81164 | 5 | 3501984.75 | 2783742.25 | 3356594.5 | 2899014 | 3353887.5 | 2710739.75 | 2844593.75 | 2783999.75 | 2808963.75 | 1889654.25 | 1452942 | 1704197.88 | 1975134.12 | 777136.5 | 1732772.88 | 3003101.25 | 1965261.75 | 2145500.75 | 2639429.5 | 1934912.62 | 0 | 2 | 1 | 9 | 1 | 9 | 0.480784911 | 0.586149099 | 0.73532821 | 0.896475654 |
| P52873 | Pc | Pyruvate carboxylase, mitochondrial | 262.55 | 7.22 | 129777 | 6 | 2515787.5 | 4950023 | 4467024.5 | 4393428 | 4390722.5 | 2594406.75 | 2381570.75 | 10260166 | 4867294 | 5645472.5 | 2872156 | 2995819 | 2938909 | 3327074 | 6690495 | 4937054.5 | 2136799 | 4991092 | 4362983 | 4854858 | 9 | 11 | 12 | 10 | 12 | 11 | 0.908648268 | 0.731077665 | 1.027310971 | 0.826550969 |

| | | | | | | | | | | | | | | | | | | | | | | | | | | | | | | | | | | | | |
|---|---|---|---|---|---|---|---|---|---|---|---|---|---|---|---|---|---|---|---|---|---|---|---|---|---|---|---|---|---|---|---|---|---|---|---|---|
| P60335 | Pcbp1 | Poly(rC)-binding protein 1 | 945.39 | 54.49 | 37498 | 13 | 45694862 | 57047054 | 42060306 | 28009404 | 47301782 | 39952702 | 51177298 | 40361566 | 47817650 | 26143302 | 63648918 | 52773410 | 31367120 | 34753930 | 42094018 | 34346298 | 27052316 | 18823846 | 21236138 | 23164248 | 12 | 1 | 10 | 2 | 12 | 2 | 1.020552987 | 1.093378646 | 0.566175623 | 0.60657736 |
| Q61990 | Pcbp2 | Poly(rC)-binding protein 2 | 812.3 | 40.88 | 38222 | 11 | 31153232 | 35309106 | 32906330 | 21097998 | 34878770 | 22450778 | 34426682 | 31367742 | 33260546 | 22850534 | 23098526 | 34003106 | 23195664 | 21254674 | 22601510 | 30455592 | 27361742 | 23095686 | 23261528 | 21370938 | 7 | 5 | 9 | 8 | 9 | 8 | 0.799209061 | 0.860049028 | 0.808169775 | 0.869691878 |
| Q91ZA3 | Pcca | Propionyl-CoA carboxylase alpha chain, mitochondrial | 387.37 | 12.02 | 79922 | 6 | 3082533 | 2103787.5 | 2535264.25 | 1921553.62 | 2608584.5 | 3839286 | 1811785.38 | 1893266.75 | 1857442.5 | 1948503.62 | 1830923 | 1124381 | 3242117 | 3529788 | 3597630.5 | 2780355 | 1676942.25 | 1233867.5 | 1735662.5 | 2398341.5 | 10 | 6 | 12 | 8 | 12 | 8 | 1.087589039 | 1.173965269 | 0.801941805 | 0.865631955 |
| Q99MN9 | Pccb | Propionyl-CoA carboxylase beta chain, mitochondrial | 351.2 | 15.71 | 58409 | 6 | 3424139 | 4664593.5 | 5148153.5 | 5345749 | 5074187 | 4795442 | 3878091 | 3512016.25 | 3675988.5 | 3686890.25 | 3607374.25 | 2650944.5 | 4386202 | 3820007.5 | 2726125.75 | 4965854 | 819785.06 | 4876477.5 | 4180616.25 | 3685008 | 3 | 6 | 8 | 9 | 8 | 9 | 0.726667935 | 0.879388051 | 0.783188072 | 0.947786738 |
| Q8BFV2 | Pcid2 | PCI domain-containing protein 2 | 46.15 | 2.51 | 46132 | 1 | 2 | 2 | 250021.14 | 125836.33 | 229244.45 | 2 | 2 | 187216.22 | 299071.44 | 177385.98 | 334583.22 | 366549.41 | 541063 | 346771.69 | 374150.72 | 2 | 2 | 195327.44 | 118237.72 | 221104.06 | 0 | 10 | 0 | 12 | 0 | 12 | 3.244255221 | 2.957939098 | 0.883602692 | 0.805621868 |
| Q8BH04 | Pck2 | Phosphoenolpyruvate carboxykinase [GTP], mitochondrial | 1435.94 | 43.28 | 70528 | 22 | 35227866 | 37885250 | 23587362 | 26004496 | 27810300 | 34771418 | 32445158 | 26792240 | 28357660 | 26213346 | 38652506 | 42818298 | 46889062 | 38998514 | 45055418 | 37405098 | 34916502 | 29689336 | 28191242 | 32629382 | 0 | 9 | 0 | 6 | 0 | 9 | 1.411244137 | 1.42962749 | 1.081827483 | 1.095919741 |
| P17918 | Pcna | Proliferating cell nuclear antigen | 1156.64 | 72.41 | 28785 | 19 | 100384410 | 105890242 | 116633322 | 102681370 | 73523386 | 104045114 | 93340018 | 77542690 | 77869914 | 80154242 | 101204810 | 93706082 | 76311786 | 82986674 | 76749434 | 95331818 | 105468770 | 83725834 | 79866562 | 81471466 | 6 | 7 | 11 | 8 | 11 | 8 | 0.863449798 | 0.995396275 | 0.893314122 | 1.029824259 |
| Q6P8I4 | Pcnp | PEST proteolytic signal-containing nuclear protein | 88.43 | 16.29 | 18963 | 2 | 1451602.5 | 1580737.75 | 1505183.5 | 1716405.88 | 908318.88 | 1435079.38 | 1113560.38 | 1438360.5 | 1164638.5 | 1792343.5 | 1392300.25 | 1485280.25 | 1129674.5 | 1705758.25 | 1224396.25 | 1416856.5 | 1961836.38 | 1646919.12 | 1313062.38 | 1727905.62 | 9 | 9 | 12 | 8 | 12 | 9 | 0.968607762 | 0.99905346 | 1.126263629 | 1.161664834 |
| Q9CQF9 | Pcyox1 | Prenylcysteine oxidase | 403.39 | 21.39 | 56495 | 6 | 5211319 | 5150488.5 | 4901415 | 4319359 | 6967477.5 | 4027441.5 | 5547110 | 5354530.5 | 5960214 | 4337847.5 | 3164005.25 | 5432766 | 5736112.5 | 4372206 | 4839387.5 | 6201409.5 | 2368332.5 | 4526478.5 | 3322134 | 5668533 | 10 | 9 | 11 | 11 | 11 | 11 | 0.886795666 | 0.933299375 | 0.831895986 | 0.875520746 |
| Q8C7K6 | Pcyox1l | Prenylcysteine oxidase-like | 211.61 | 8.48 | 54875 | 4 | 3149152.25 | 3321524 | 1944739.75 | 1842294 | 2226463.75 | 2852605 | 4575130 | 4067655.5 | 3573061 | 3237276.25 | 2617371.75 | 3086431 | 4764610.5 | 3000648 | 5135178 | 2126055.5 | 2600865.75 | 2670356.5 | 1825583.75 | 2651932.75 | 6 | 11 | 12 | 0 | 12 | 11 | 1.490225915 | 1.016307 | 0.951187839 | 0.648692825 |
| O88637 | Pcyt2 | Ethanolamine-phosphate cytidylyltransferase | 147.39 | 10.4 | 45220 | 3 | 3953830.5 | 3750783 | 2708158.5 | 3498189 | 3081090.5 | 4574638 | 4489028 | 3095838 | 2337241.25 | 3321406 | 4240132 | 4096346 | 4200560.5 | 4485653.5 | 4254816.5 | 2824072.75 | 4163368.75 | 3680794 | 4253516.5 | 3055468.5 | 0 | 10 | 10 | 11 | 10 | 11 | 1.252203626 | 1.194147934 | 1.057978226 | 1.008927371 |
| Q3UHX2 | Pdap1 | 28 kDa heat- and acid-stable phosphoprotein | 214.37 | 13.26 | 20605 | 3 | 3459554 | 4165920 | 3314518.25 | 3264785 | 1532367 | 3557703 | 3699228 | 3542589.5 | 2758749 | 2349914 | 3594211.25 | 3369708 | 2851120 | 3203569.5 | 2798266.75 | 4367928.5 | 4303552 | 3970523 | 2588067.25 | 2955913.5 | 10 | 9 | 12 | 7 | 12 | 9 | 1.005066437 | 0.994260313 | 1.155608919 | 1.143184214 |
| Q6NS46 | Pdcd11 | Protein RRP5 homolog | 975.9 | 13 | 207779 | 20 | 6132162.5 | 7580388.5 | 4713025 | 5932954 | 5827317 | 6561284.5 | 5904934.5 | 5506210 | 5704765.5 | 4411420 | 10134853 | 9605872 | 9063067 | 10414823 | 10559770 | 5956619 | 6887934 | 6022601 | 6155115 | 5236416 | 0 | 10 | 0 | 7 | 0 | 10 | 1.649063715 | 1.772190828 | 1.002412985 | 1.077258011 |
| P56812 | Pdcd5 | Programmed cell death protein 5 | 316.41 | 42.86 | 14275 | 5 | 4469920 | 2829261.75 | 3439697.75 | 3793674.25 | 2974762.75 | 3814479.75 | 3250450.75 | 2171153.5 | 2590284.75 | 2622509.75 | 2052369.25 | 1916812.62 | 1453890.12 | 1571807.5 | 1823255.5 | 3463496.25 | 6266344 | 3849766.5 | 2494609 | 3203517.5 | 0 | 11 | 0 | 7 | 0 | 11 | 0.503682845 | 0.610298923 | 1.101124393 | 1.334202738 |
| Q9WU78 | Pdcd6ip | Programmed cell death 6-interacting protein | 1755.69 | 40.85 | 96024 | 27 | 46083634 | 41155586 | 44027574 | 38414790 | 47158686 | 37520918 | 45152774 | 46343350 | 47855510 | 38534326 | 49308090 | 63877674 | 56980014 | 47983106 | 62419478 | 49543442 | 41024222 | 39126866 | 39965446 | 42625254 | 0 | 10 | 0 | 12 | 0 | 12 | 1.293894174 | 1.302504194 | 0.97899357 | 0.985508132 |
| Q4KLJ8 | Pdcl3 | Phosducin-like protein 3 | 105.8 | 7.92 | 27699 | 2 | 141526.94 | 526577.25 | 1039847.62 | 654519.38 | 394746.47 | 278266.03 | 402187.5 | 449809.91 | 341457.78 | 653479.88 | 2 | 1456605.25 | 2 | 2 | 1468712.88 | 2 | 2 | 389295.12 | 582775.12 | 223192.97 | 10 | 5 | 10 | 6 | 10 | 6 | 1.060969604 | 1.376492855 | 0.433504843 | 0.562425462 |
| Q3TIU4 | Pde12 | 2',5'-phosphodiesterase 12 | 127.7 | 4.28 | 67514 | 2 | 1445062.25 | 1744023.75 | 732138.25 | 1665781.25 | 1764370.88 | 1486836 | 1708437.5 | 1876610.5 | 1625518.12 | 1580653.5 | 1019784.25 | 1647181.75 | 2125589 | 1245095.25 | 712949.38 | 1675749 | 836852.88 | 1469904.75 | 1534643 | 1697220.25 | 9 | 11 | 8 | 7 | 9 | 11 | 0.91827697 | 0.815481309 | 0.98136315 | 0.871505364 |
| P26284 | Pdha1 | Pyruvate dehydrogenase E1 component subunit alpha, somatic form, mitochondrial | 93.64 | 4.62 | 43227 | 2 | 6381967.5 | 6951795 | 5911163 | 6415036 | 5637487 | 7366689 | 6154435.5 | 5844258 | 5766943 | 6327423.5 | 4788980.5 | 5111127 | 6528399.5 | 8013801 | 6705987 | 4346289.5 | 6840798 | 7683027 | 7166072.5 | 6649793 | 12 | 7 | 12 | 8 | 12 | 8 | 0.995234324 | 0.990099921 | 1.044365645 | 1.038977774 |
| Q9D051 | Pdhb | Pyruvate dehydrogenase E1 component subunit beta, mitochondrial | 629.52 | 35.1 | 38937 | 10 | 9083238 | 10891227 | 5407464 | 8502919 | 7695178 | 10261159 | 11035968 | 8108928 | 7599641.5 | 8778769 | 7426099.5 | 8458100 | 9237314 | 8045785.5 | 9266578 | 8565924 | 8365759.5 | 7814793.5 | 9753374 | 7665459 | 12 | 12 | 9 | 9 | 12 | 12 | 1.020535124 | 0.926818224 | 1.014076085 | 0.920952326 |
| Q8BKZ9 | Pdhx | Pyruvate dehydrogenase protein X component, mitochondrial | 187.61 | 8.38 | 53999 | 4 | 5692571 | 5348055 | 5277889 | 7223269 | 5903152 | 7323416.5 | 6971629 | 6188923 | 5638475 | 7269680 | 4317165 | 5184994.5 | 6790754 | 5833134 | 7813929 | 6164333 | 4762955.5 | 6583406 | 6126254 | 7443747.5 | 12 | 8 | 8 | 9 | 12 | 9 | 1.016812416 | 0.896617925 | 1.055553186 | 0.93077926 |
| P27773 | Pdia3 | Protein disulfide-isomerase A3 | 2351.18 | 62.77 | 56678 | 36 | 233723186 | 270934146 | 260318658 | 239506866 | 260261010 | 220365842 | 242542418 | 263304578 | 251976130 | 243703170 | 252108402 | 302901378 | 294965122 | 274586050 | 314917730 | 244305410 | 205182530 | 248841170 | 236884642 | 257126386 | 3 | 7 | 1 | 11 | 3 | 11 | 1.138158263 | 1.178073446 | 0.94275226 | 0.975814559 |
| P08003 | Pdia4 | Protein disulfide-isomerase A4 | 2446.24 | 58.15 | 71982 | 40 | 70805106 | 80151162 | 83073042 | 77375466 | 73120162 | 69255962 | 68167754 | 83828586 | 82211426 | 81315226 | 112541330 | 125114514 | 121760498 | 126864722 | 131873794 | 63672930 | 65396930 | 77751346 | 84652154 | 81420026 | 0 | 12 | 0 | 10 | 0 | 12 | 1.607580671 | 1.60651941 | 0.969750851 | 0.96911066 |
| Q922R8 | Pdia6 | Protein disulfide-isomerase A6 | 1543.07 | 53.64 | 48100 | 21 | 78070866 | 80727586 | 70448626 | 70088938 | 63396150 | 82686786 | 73509626 | 61895118 | 65695890 | 69008538 | 95330018 | 84271194 | 89697122 | 76661930 | 72146322 | 77683914 | 92538650 | 71113866 | 71020402 | 63566150 | 4 | 10 | 3 | 9 | 4 | 10 | 1.152659249 | 1.185122949 | 1.036365168 | 1.06555354 |
| Q63065 | Pdk1 | [Pyruvate dehydrogenase (acetyl-transferring)] kinase isozyme 1, mitochondrial | 138.89 | 8.53 | 49081 | 3 | 1415448.75 | 1283489.38 | 1333163.25 | 1457126.75 | 1267866 | 1456968.75 | 1704635.5 | 1387054.25 | 1403062 | 1648937.5 | 2 | 337343.09 | 85595.18 | 484188.22 | 2 | 1400480.25 | 925892.44 | 1134142 | 1345652.25 | 470228.28 | 0 | 6 | 0 | 1 | 0 | 6 | 0.134248609 | 0.119348942 | 0.780867503 | 0.694202426 |
| Q922H2 | Pdk3 | [Pyruvate dehydrogenase (acetyl-transferring)] kinase isozyme 3, mitochondrial | 295.7 | 15.66 | 47923 | 5 | 832475.88 | 720131.12 | 741111.75 | 907957.31 | 952218.56 | 1023594.12 | 213663.59 | 1029792 | 917969.69 | 1185566.12 | 962426.94 | 905889.62 | 915973.19 | 954669 | 967925.31 | 851148.12 | 1114498.75 | 873596.38 | 1108991.25 | 880255.44 | 3 | 6 | 8 | 11 | 8 | 11 | 1.133125534 | 1.076945878 | 1.162400682 | 1.104769582 |
| Q8CI51 | Pdlim5 | PDZ and LIM domain protein 5 | 373.91 | 13.87 | 63299 | 7 | 5665323.5 | 6637987.5 | 6780425 | 7549711 | 8856531 | 7623103 | 7095839.5 | 8481202 | 8763612 | 7475861 | 4790414.5 | 5140974.5 | 5858525.5 | 3951268.25 | 5615782 | 5344975.5 | 5908886 | 7996815.5 | 8926034 | 9355953 | 1 | 10 | 0 | 12 | 1 | 12 | 0.714482403 | 0.642931305 | 1.0575567 | 0.951648783 |
| Q3TJD7 | Pdlim7 | PDZ and LIM domain protein 7 | 40.9 | 2.63 | 50119 | 1 | 685236 | 373015.81 | 2 | 287052.34 | 866985.19 | 502613.44 | 419912 | 2 | 2 | 2 | 2 | 709011.56 | 158792.72 | 628064.75 | 2 | 2 | 874888.25 | 352747.75 | 396105.06 | 345755.62 | 9 | 12.5 | 9 | 9.5 | 9 | 12.5 | 0.676164573 | 1.621487318 | 0.890252854 | 2.134885159 |
| A4L9P7 | Pds5a | Sister chromatid cohesion protein PDS5 homolog A | 151.01 | 1.95 | 150286 | 2 | 154597.8 | 2 | 2 | 2 | 2 | 2 | 219235.47 | 230401.52 | 2 | 263194.97 | 883707.19 | 859203.75 | 731730.94 | 559910.12 | 802242.31 | 2 | 2 | 2 | 2 | 118210.87 | 0 | 12 | 0 | 6 | 0 | 12 | 24.81662596 | 5.382436529 | 0.764647057 | 0.165843022 |
| Q99K01 | Pdxdc1 | Pyridoxal-dependent decarboxylase domain-containing protein 1 | 341.93 | 9.53 | 87335 | 7 | 7307984 | 5494989.5 | 7492156 | 8137889 | 7915961 | 4630663.5 | 7529723 | 6926441 | 8510703 | 9669261 | 3390983 | 4150239 | 6654345 | 5864476 | 5153743 | 5874688 | 4576788.5 | 7612195 | 9993522 | 7505694 | 2 | 12 | 3 | 11 | 3 | 12 | 0.693658704 | 0.676575176 | 0.978373753 | 0.954278221 |
| Q8K183 | Pdxk | Pyridoxal kinase | 515.84 | 25.32 | 35015 | 8 | 9186884 | 6987678 | 9905902 | 8348140 | 10052938 | 10111604 | 8496023 | 9903068 | 9829451 | 8060013 | 4807802 | 6246352 | 2989353 | 6247271 | 5532470.5 | 7867077 | 5164282.5 | 5147282 | 7701528.5 | 9083980 | 0 | 4 | 0 | 2 | 0 | 4 | 0.580538519 | 0.556533621 | 0.786037274 | 0.753535133 |
| Q62048 | Pea15 | Astrocytic phosphoprotein PEA-15 | 57.07 | 7.69 | 15054 | 1 | 2 | 2 | 287190.75 | 232733.36 | 2 | 2 | 2 | 286723.44 | 2 | 2 | 717405.38 | 410440.09 | 392152.5 | 437535.34 | 384581.88 | 351387.66 | 448471.88 | 2 | 425206.34 | 324166.53 | 0 | 3.5 | 0 | 3 | 0 | 3.5 | 4.504673118 | 8.168323606 | 2.979697425 | 5.403085235 |
| P70296 | Pebp1 | Phosphatidylethanolamine-binding protein 1 | 226.84 | 22.46 | 20830 | 4 | 14913318 | 19017374 | 17186178 | 11304594 | 18450446 | 10975424 | 16349182 | 20548624 | 19800590 | 13790762 | 6281577.5 | 13072753 | 8578844 | 7377075.5 | 10106960 | 16111036 | 9218788 | 12548082 | 13648611 | 15926708 | 1 | 6 | 1 | 6 | 1 | 6 | 0.561594378 | 0.557508661 | 0.834074835 | 0.828006765 |
| Q99MZ7 | Pecr | Peroxisomal trans-2-enoyl-CoA reductase | 156.07 | 11.88 | 32410 | 3 | 1325850.5 | 1847321.38 | 1110055.62 | 1406459.12 | 1514281.38 | 1772324 | 1808689.62 | 1355087.62 | 2095742 | 1327678.75 | 2186788.5 | 2905169 | 3032293.5 | 3177936.75 | 2775546 | 1698797.62 | 1374983.75 | 1691746.38 | 2 | 1832638.12 | 0 | 11 | 0 | 10 | 0 | 11 | 1.954163837 | 1.684035734 | 0.915907437 | 0.789299661 |
| Q641Z8 | Pef1 | Peflin | 64.78 | 4.59 | 30012 | 1 | 2 | 2 | 2 | 2 | 2 | 290626 | 434748.44 | 2 | 2 | 2 | 737141.38 | 409857.72 | 268230.44 | 2 | 190527.92 | 2 | 2 | 2 | 2 | 2 | 2.5 | 12.5 | 8.5 | 7.5 | 8.5 | 12.5 | 160575.946 | 2.213679018 | 1 | 1.378587E-05 |
| Q9DBD5 | Pelp1 | Proline-, glutamic acid- and leucine-rich protein 1 | 589.87 | 9.8 | 118069 | 10 | 8466155 | 7871384.5 | 8230362.5 | 8394781 | 8288428.5 | 9322766 | 9220849 | 9037279 | 9094262 | 9944895 | 10076641 | 12322360 | 14452496 | 12724513 | 13143623 | 9965409 | 8891672 | 9831016 | 9582173 | 8922564 | 0 | 0 | 0 | 12 | 0 | 12 | 1.520434983 | 1.345336001 | 1.144037876 | 1.012286194 |
| Q11136 | Pepd | Xaa-Pro dipeptidase | 541.28 | 22.52 | 55029 | 10 | 19124546 | 19551924 | 20142282 | 21426330 | 17979892 | 20776438 | 18723772 | 17727702 | 17098994 | 18245608 | 8415322 | 11312149 | 9745407 | 9913810 | 7866493.5 | 21504780 | 20972542 | 18800478 | 17508138 | 18063210 | 0 | 11 | 0 | 8 | 0 | 11 | 0.481070949 | 0.51044505 | 0.985993114 | 1.046197665 |
| Q3B8N8 | Pes1 | Pescadillo homolog | 221.39 | 7.34 | 67604 | 5 | 2506856 | 2773894.75 | 1981942.12 | 1866994.75 | 2830186.25 | 1865900 | 2821750.5 | 2217475.75 | 2260857.5 | 2193180 | 2713723 | 2578059 | 2169956 | 1571310 | 2101714 | 2761150.75 | 1525931.75 | 1404104.88 | 1801738.5 | 2111315.75 | 10 | 5 | 10 | 5 | 10 | 5 | 0.931009986 | 0.980244871 | 0.803038705 | 0.845506046 |
| Q9Z210 | Pex11b | Peroxisomal membrane protein 11B | 54.72 | 4.63 | 28710 | 1 | 2 | 2 | 2 | 2 | 344777.78 | 332900.16 | 2 | 2 | 2 | 2 | 724620.06 | 161781.72 | 591086.62 | 923490.31 | 825582.06 | 358937.56 | 2 | 576193.69 | 2 | 2 | 1 | 9 | 1 | 9 | 1 | 9 | 9.358160798 | 9.692044707 | 2.71222685 | 2.808994679 |
| Q642G4 | Pex14 | Peroxisomal membrane protein PEX14 | 89.67 | 5.05 | 40936 | 2 | 11449656 | 9093012 | 183558.89 | 92167.52 | 167829.48 | 547344.44 | 11442589 | 186067.16 | 161071.84 | 115977.95 | 1432861.5 | 1346922.12 | 1190978.38 | 1126703.38 | 923192.38 | 391246.56 | 186285.61 | 135680.14 | 2 | 326642.22 | 10 | 10 | 5 | 10 | 10 | 10 | 0.286886187 | 0.483468513 | 0.049549482 | 0.083502154 |
| Q9Z126 | Pf4 | Platelet factor 4 | 42.54 | 11.43 | 11243 | 1 | 2 | 2 | 2 | 2 | 82851.27 | 2 | 2 | 2 | 2 | 2 | 465011.41 | 596885.56 | 435310.44 | 702816.19 | 765936.56 | 145406.72 | 171053.88 | 2 | 109087.7 | 183300.95 | 0 | 3 | 0 | 2.5 | 0 | 3 | 35.79515195 | 296596.016 | 7.3480161 | 60885.125 |
| Q5SUR0 | Pfas | Phosphoribosylformylglycinamidine synthase | 730.95 | 11.44 | 144629 | 13 | 28995282 | 21501994 | 23725600 | 23631096 | 20755394 | 27021230 | 24758644 | 19991874 | 18016396 | 24448922 | 14556991 | 18565342 | 16842270 | 13096330 | 13409008 | 25599420 | 25251280 | 22341134 | 27152226 | 22907224 | 0 | 9 | 1 | 8 | 1 | 9 | 0.644720932 | 0.669396928 | 1.039136184 | 1.078907996 |
| Q9CWM4 | Pfdn1 | Prefoldin subunit 1 | 102.38 | 14.75 | 14255 | 2 | 1026337.62 | 874228.75 | 769224.44 | 1162761 | 620966.75 | 1017767.12 | 885211.31 | 689704.31 | 733995.19 | 995674.88 | 929976.5 | 694433.06 | 854454.12 | 875374.06 | 1007660.94 | 1069790.88 | 1340700.5 | 1346995.75 | 1308532.25 | 1048482.75 | 12 | 2 | 12 | 0 | 12 | 2 | 0.979427529 | 1.009149154 | 1.37295984 | 1.414623562 |
| B0BN18 | Pfdn2 | Prefoldin subunit 2 | 254.13 | 30.52 | 16580 | 4 | 9902071 | 8016953 | 9319836 | 7725765.5 | 9146822 | 6017976 | 8535558 | 9421464 | 8693417 | 8053180 | 3614220 | 5860170 | 5442381 | 1195074.88 | 4245354 | 11936823 | 6738616 | 7656454 | 7174653 | 7789167 | 0 | 6 | 0 | 9 | 0 | 9 | 0.461494715 | 0.499911653 | 0.936167715 | 1.014098613 |
| Q9WU28 | Pfdn5 | Prefoldin subunit 5 | 200.71 | 25.97 | 17356 | 3 | 5020825 | 4666618.5 | 4694435.5 | 5519291.5 | 5624607 | 4763807 | 4119919 | 4529040.5 | 4999391.5 | 5160469 | 3498742.5 | 2819087.5 | 3375099 | 4541273.5 | 2944034.75 | 4818647 | 5521528 | 4763253 | 3204471.25 | 3952763 | 0 | 8 | 2 | 10 | 2 | 10 | 0.672976063 | 0.728736651 | 0.872085571 | 0.944343719 |
| P12382 | Pfkl | ATP-dependent 6-phosphofructokinase, liver type | 524.97 | 13.97 | 85360 | 8 | 7147738 | 6853006.5 | 6941655.5 | 6060000 | 5786239.5 | 6733992.5 | 6137818 | 5296620 | 7605873 | 4666666.5 | 5844731 | 5687493.5 | 6056799.5 | 5518017.5 | 5145961.5 | 4488412 | 6771813.5 | 4256197 | 6150151.5 | 3999054.75 | 2 | 4 | 9 | 7 | 9 | 7 | 0.861670488 | 0.928124268 | 0.782759796 | 0.843127822 |
| P47857 | Pfkm | ATP-dependent 6-phosphofructokinase, muscle type | 177.73 | 5.26 | 85269 | 4 | 2017228.5 | 1863521.5 | 2287508.5 | 2307422.5 | 1766084.25 | 1333542 | 1994168.75 | 1898901.75 | 1938431.75 | 1922248.5 | 1636753.12 | 1019036.12 | 1305465.25 | 1388077.12 | 1773281.75 | 1981965 | 1973366.5 | 2195677 | 1975740 | 1937440.75 | 1 | 11 | 3 | 5 | 3 | 11 | 0.695447824 | 0.783799263 | 0.982661583 | 1.107501379 |
| Q9WUA3 | Pfkp | ATP-dependent 6-phosphofructokinase, platelet type | 2160.22 | 49.87 | 85455 | 31 | 52052906 | 54524506 | 53471202 | 55589866 | 57866402 | 62908750 | 62467594 | 65599658 | 66590114 | 67205882 | 43797434 | 48899562 | 60544678 | 40617486 | 51388550 | 54898078 | 50661634 | 64782966 | 49809742 | 62301834 | 5 | 12 | 0 | 2 | 5 | 12 | 0.896684945 | 0.755138102 | 1.032721069 | 0.869700146 |
| P62962 | Pfn1 | Profilin-1 | 989.64 | 95 | 14957 | 15 | 162198946 | 196126130 | 149374898 | 195677154 | 191199698 | 152114322 | 160290882 | 184939138 | 179733442 | 197231890 | 261492562 | 240977426 | 227605970 | 233037346 | 227395154 | 169898258 | 169066690 | 101699730 | 186274690 | 191303538 | 0 | 9 | 0 | 12 | 0 | 12 | 1.33080628 | 1.361655365 | 0.914670358 | 0.935873101 |
| P25113 | Pgam1 | Phosphoglycerate mutase 1 | 921.69 | 62.2 | 28832 | 13 | 57115262 | 64741810 | 63224042 | 74708242 | 68251122 | 61152098 | 58887798 | 87021602 | 87932458 | 80788282 | 73548426 | 85249738 | 83424618 | 83238578 | 77153458 | 116938746 | 107535178 | 115754066 | 75592394 | 116819578 | 1 | 0 | 12 | 3 | 12 | 3 | 1.227332738 | 1.071404599 | 1.623701944 | 1.417416546 |
| Q562B5 | Pgam5 | Serine/threonine-protein phosphatase PGAM5, mitochondrial | 51.85 | 3.47 | 32060 | 1 | 456048.94 | 492526.69 | 639917.94 | 2 | 2 | 353353.56 | 479121.28 | 1022045.88 | 636768.06 | 738382.19 | 2 | 692166.19 | 796857.19 | 903644.06 | 928331.25 | 543169.5 | 561577.88 | 638720.12 | 830647.12 | 690566 | 4 | 3 | 10 | 11 | 10 | 11 | 2.090655191 | 1.028278336 | 2.055200261 | 1.010840005 |
| Q3UUQ7 | Pgap1 | GPI inositol-deacylase | 92.57 | 2.06 | 104578 | 2 | 2 | 284920.38 | 273569.88 | 286997.88 | 342829.69 | 374254.53 | 424334.88 | 405882.62 | 333264.22 | 2 | 430502.91 | 263827.16 | 289948.34 | 2 | 326597.91 | 395050.06 | 327307.84 | 269943.53 | 205384.23 | 318674.16 | 10.5 | 10 | 8.5 | 7 | 10.5 | 10 | 1.103135946 | 0.852471687 | 1.276053619 | 0.986097484 |
| Q9DCD0 | Pgd | 6-phosphogluconate dehydrogenase, decarboxylating | 1968.88 | 56.11 | 53247 | 28 | 298455074 | 325586754 | 299283042 | 261390402 | 284757730 | 310415618 | 332291074 | 307572770 | 310897538 | 277785634 | 290922530 | 396299714 | 300211906 | 275886306 | 271395362 | 374424898 | 287359650 | 329205122 | 342584546 | 312688098 | 12 | 4 | 7 | 6 | 12 | 6 | 1.044398785 | 0.997240468 | 1.120307969 | 1.069722083 |
| P09411 | Pgk1 | Phosphoglycerate kinase 1 | 3084.831 | 82.97 | 44550 | 45 | 768082050 | 937493058 | 855738754 | 790932738 | 632498562 | 838469954 | 879535554 | 829182402 | 972537154 | 1017636546 | 746589058 | 781059010 | 883768322 | 981040386 | 872042946 | 851427138 | 943809666 | 961683394 | 817959554 | 1016700994 | 9 | 4 | 9 | 12 | 9 | 12 | 1.070206387 | 0.939863315 | 1.152289685 | 1.011949485 |
| Q9CQ60 | Pgls | 6-phosphogluconolactonase | 387.64 | 40.47 | 27254 | 7 | 13461286 | 13437276 | 13949707 | 14905296 | 13462751 | 14817032 | 12143585 | 14065135 | 14920909 | 14593287 | 10584723 | 9786587 | 10765428 | 11833380 | 11567682 | 13407732 | 14583483 | 16726064 | 19172080 | 16711540 | 0 | 6 | 0 | 7 | 0 | 7 | 0.787932718 | 0.77314772 | 1.164478315 | 1.142627707 |
| Q9D0F9 | Pgm1 | Phosphoglucomutase-1 | 1344.41 | 44.31 | 61418 | 23 | 35471650 | 34365374 | 28280304 | 31819486 | 32800462 | 36823222 | 33374102 | 33687850 | 35192290 | 33712866 | 19961366 | 21603780 | 23641106 | 22946542 | 23550148 | 35623742 | 36547914 | 36394966 | 39613918 | 35433310 | 0 | 1 | 0 | 4 | 0 | 4 | 0.686400465 | 0.646465239 | 1.12828391 | 1.062639616 |
| Q7TSV4 | Pgm2 | Phosphoglucomutase-2 | 65.15 | 4.35 | 68748 | 2 | 1805791.75 | 1206048.75 | 1778420.12 | 1387840.38 | 1983409 | 658120.38 | 1359052.25 | 1249144.12 | 2031258.5 | 1084019.5 | 2 | 676990.19 | 837592.56 | 1010925.44 | 851403.75 | 1845686.5 | 1282355.25 | 596209.5 | 1797446.25 | [illegible] | 0 | 8 | 4 | 12 | 4 | 12 | 0.413760927 | 0.529164585 | 0.781624172 | 0.999629991 |
| D3ZDK7 | Pgp | Glycerol-3-phosphate phosphatase | 138.05 | 9.66 | 34601 | 2 | 4246172.5 | 3649662.25 | 3143320.75 | 2589302.5 | 3506447.75 | 2265705.25 | 3599063.75 | 2350552.5 | 2671271.25 | 2417777.75 | 654937.69 | 1913880.5 | 2 | 821033.94 | 1157900 | 4803182 | 2189380.75 | 1884600 | 2090667.88 | 2687291.25 | 0 | 6 | 0 | 9 | 0 | 9 | 0.265408762 | 0.341824074 | 0.796918412 | 1.026363621 |
| O55022 | Pgrmc1 | Membrane-associated progesterone receptor component 1 | 176.4 | 11.79 | 21694 | 3 | 3555214.5 | 2858885.5 | 2329040.75 | 2962519.5 | 2864697 | 3479569 | 3485587.75 | 3442237 | 3450965 | 2926587 | 4154098.5 | 3446211.5 | 3292091 | 3151655.5 | 3781198.5 | 4363704.5 | 4617854 | 3242961 | 4676843 | 3460813.25 | 3 | 2 | 11 | 6 | 11 | 6 | 1.22339176 | 1.061978708 | 1.397506966 | 1.213121332 |
| Q80UU9 | Pgrmc2 | Membrane-associated progesterone receptor component 2 | 414.19 | 30.41 | 23334 | 4 | 2236147.5 | 3077430.75 | 2020058.5 | 3089826.75 | 3439016.75 | 3901849.5 | 4155729.5 | 3621790 | 3541671.5 | 4001262.5 | 4916554 | 5468922.5 | 6076857.5 | 6186110 | 7183534 | 3419730.5 | 4300598.5 | 3852797.75 | 2780236 | 3270899 | 0 | 5 | 0 | 7 | 0 | 7 | 2.151994265 | 1.551946091 | 1.271364246 | 0.916865255 |
| Q8BHF7 | Pgs1 | CDP-diacylglycerol--glycerol-3-phosphate 3-phosphatidyltransferase, mitochondrial | 231.45 | 8.32 | 62489 | 4 | 650251.88 | 655954.88 | 561986.88 | 473843.69 | 499483.09 | 697367.62 | 567186.56 | 884309.56 | 646220.75 | 670437 | 1176801.12 | 1719612.5 | 1817931.38 | 1148640.5 | 1928867.75 | 532782.69 | 401099.5 | 1002155.75 | 1002348.88 | 402113.06 | 0 | 12 | 0 | 10 | 0 | 12 | 2.742142268 | 2.248392709 | 1.175602982 | 0.963924157 |
| Q63068 | Phax | Phosphorylated adapter RNA export protein | 99.57 | 4.94 | 43166 | 2 | 1638308.75 | 1850208 | 1888565.75 | 1824083.75 | 2158349.5 | 1727069.38 | 1547262.5 | 1709620 | 1612579 | 1634208.25 | 1623109.25 | 3072302.25 | 1616010.38 | 1200266.5 | 2503891.75 | 1569894.88 | 1518565.25 | 1772751.25 | 1787157.38 | 2576173.75 | 10 | 7 | 11 | 9 | 11 | 9 | 1.070095975 | 1.216850634 | 0.985579036 | 1.120742908 |
| P67778 | Phb | Prohibitin | 1752.75 | 80.88 | 29820 | 25 | 215902386 | 216467602 | 194506034 | 206260498 | 199560674 | 227240322 | 230726114 | 205789042 | 213743362 | 234699170 | 300793474 | 278884802 | 369353890 | 357096834 | 342199362 | 234440322 | 267370482 | 246789874 | 258797250 | 232440418 | 0 | 0 | 0 | 2 | 0 | 2 | 1.596139092 | 1.482045775 | 1.200582662 | 1.114764039 |
| O35129 | Phb2 | Prohibitin-2 | 1315.11 | 66.89 | 33296 | 21 | 18276052 | 22907158 | 19847916 | 20130556 | 14606617 | 20138358 | 17098080 | 22412610 | 15861471 | 26331250 | 156815106 | 98144122 | 119790498 | 99242730 | 93891698 | 18760020 | 17426810 | 25251846 | 22835156 | 25259078 | 0 | 8 | 0 | 9 | 0 | 9 | 5.92977175 | 5.576141887 | 1.14372826 | 1.075520497 |
| A6H5X4 | Phf11 | PHD finger protein 11 | 41.66 | 2.97 | 37728 | 1 | 1458327 | 2180210.75 | 1417203.38 | 2178104 | 1992728.62 | 1956647.88 | 2049855.25 | 1892231.75 | 1916421.12 | 2205813.75 | 1004646.88 | 1272097.25 | 970532 | 736571.38 | 875836.25 | 2 | 1583367.38 | 1011739 | 1779375.88 | 1585156.25 | 0 | 6 | 0 | 0 | 0 | 6 | 0.526705134 | 0.484951445 | 0.6459213 | 0.594716944 |
| P83870 | Phf5a | PHD finger-like domain-containing protein 5A | 169.02 | 25.45 | 12405 | 3 | 2 | 321757.5 | 2 | 2 | 2 | 319843.84 | 486234.88 | 2 | 2 | 2 | 501436.94 | 440353.25 | 2 | 2 | 2 | 2 | 2 | 2 | 2 | 2 | 9 | 10 | 11.5 | 7.5 | 11.5 | 10 | 2.926964482 | 1.168358817 | 3.107853E-05 | 1.240564E-05 |
| Q2TBE6 | Pi4k2a | Phosphatidylinositol 4-kinase type 2-alpha | 281.46 | 16.49 | 54258 | 6 | 14134694 | 8447061 | 9438110 | 16713823 | 4895660 | 5931340.5 | 9265174 | 9742576 | 9049483 | 23320520 | 5037141 | 4518768.5 | 8721169 | 6223526 | 5178231 | 5289799.5 | 7479837 | 7515172 | 10073169 | 12384067 | 5 | 9 | 2 | 10 | 5 | 10 | 0.553406607 | 0.517873058 | 0.796989822 | 0.745816098 |
| Q5XIL2 | Pi4k2b | Phosphatidylinositol 4-kinase type 2-beta | 62.77 | 1.89 | 54464 | 1 | 201828.95 | 192872.03 | 155043.92 | 128603.2 | 2 | 153298.23 | 211923.09 | 2 | 172433.75 | 2 | 236760.44 | 386366.03 | 399228.41 | 150951.09 | 356458.03 | 160620.44 | 49255.98 | 96434.89 | 2 | 36620.59 | 3 | 6.5 | 3 | 10 | 3 | 10 | 2.255124603 | 2.845230529 | 0.505541165 | 0.637827797 |
| E9Q3L2 | Pi4ka | Phosphatidylinositol 4-kinase alpha | 54.04 | 0.57 | 237042 | 1 | 2 | 2 | 2 | 2 | 2 | 398141.41 | 2 | 593519.56 | 413704.09 | 2 | 2 | 2 | 2 | 2 | 2 | 422106.47 | 2 | 2 | 657323.19 | 355529.69 | 12.5 | 5 | 5 | 12 | 12.5 | 12 | 1 | 7.115569E-06 | 143496.335 | 1.02105802 |
| O55012 | Picalm | Phosphatidylinositol-binding clathrin assembly protein | 872.65 | 24.53 | 69286 | 14 | 34177938 | 38398362 | 30245708 | 30998548 | 30196022 | 34991334 | 33874894 | 26959778 | 27198158 | 26328678 | 15855726 | 16030653 | 16552048 | 16273869 | 14930689 | 36329390 | 37078830 | 29904988 | 29679896 | 31670222 | 0 | 11 | 0 | 6 | 0 | 11 | 0.485578872 | 0.533253897 | 1.003943187 | 1.10251217 |
| E2JF22 | Piezo1 | Piezo-type mechanosensitive ion channel component 1 | 81.58 | 0.71 | 292002 | 2 | 105303.11 | 255191.12 | 316827.97 | 284994.16 | 303752.88 | 283949.47 | 285851.25 | 2 | 396604.06 | 413044.81 | 565501.81 | 401417.19 | 726700.75 | 493278.94 | 421860.88 | 2 | 123512.69 | 319327.56 | 387086.53 | 378066.16 | 0 | 9 | 1 | 10.5 | 1 | 10.5 | 2.060518878 | 1.891157029 | 0.954130234 | 0.875706657 |
| Q9CXY9 | Pigk | GPI-anchor transamidase | 210.43 | 10.13 | 44895 | 4 | 2335566 | 1454087.5 | 1906400.5 | 606371.25 | 2657124 | 3237278 | 2313315.5 | 1848794.75 | 2841323.25 | 3388000 | 3978282.25 | 3122620 | 4777194 | 2168885.25 | 2540831 | 1952409.25 | 695661.06 | 1359033.62 | 2750935.75 | 2487256 | 3 | 11 | 9 | 5 | 9 | 11 | 1.851411498 | 1.21712258 | 1.031892947 | 0.678369021 |
| Q9R1S3 | Pign | GPI ethanolamine phosphate transferase 1 | 266.29 | 3.76 | 105045 | 4 | 963525.62 | 908864.75 | 1026610.81 | 1068950.62 | 848070.75 | 474491.41 | 1015214.75 | 875895.12 | 932954.88 | 1311912.12 | 1623856 | 1851832.75 | 2037209.5 | 1056396 | 2454058.75 | 955597.81 | 576100.31 | 949141.38 | 1009777 | 1132076.25 | 1 | 12 | 1 | 11 | 1 | 12 | 1.873611036 | 1.957144579 | 0.959856957 | 1.00265146 |
| Q6PD26 | Pigs | GPI transamidase component PIG-S | 537.41 | 18.74 | 61711 | 8 | 6209113 | 6569209 | 7305560 | 7186453 | 3837122.5 | 6280324 | 6494984 | 6616174 | 6727003 | 7794197.5 | 7650928 | 10429734 | 11891579 | 8394614 | 10508785 | 6268288 | 3845039 | 6453690 | 7967339 | 7165303.5 | 0 | 12 | 1 | 10 | 1 | 12 | 1.571187231 | 1.44122011 | 1.0190373 | 0.934743499 |
| Q8BXQ2 | Pigt | GPI transamidase component PIG-T | 271.42 | 9.28 | 65705 | 5 | 2942434.75 | 2497266 | 2644310.5 | 4130881 | 4292071.5 | 4188008.75 | 2991306.25 | 4149029 | 4220635 | 4976553 | 6951334 | 6286161 | 8831532 | 9410306 | 8544537 | 2474526.75 | 4586751.5 | 4679590 | 4758880 | 4076328.5 | 0 | 7 | 0 | 12 | 0 | 12 | 2.424665772 | 1.949955304 | 1.246508871 | 1.002462531 |
| Q8CHJ1 | Pigu | Phosphatidylinositol glycan anchor biosynthesis class U protein | 116.17 | 6.21 | 49953 | 2 | 2 | 2 | 2 | 2 | 296775.66 | 2 | 200657.27 | 638889.69 | 541916.31 | 2 | 565129.56 | 989453.44 | 1372125.88 | 964058.12 | 1158465.5 | 2 | 2 | 372500.97 | 509778.47 | 191905.53 | 0 | 7 | 1 | 10 | 1 | 10 | 17.01317552 | 3.654978015 | 3.619434338 | 0.77757106 |
| Q9CQJ2 | Pih1d1 | PIH1 domain-containing protein 1 | 104.37 | 6.9 | 32209 | 2 | 2560342.75 | 2702895 | 1605586.25 | 2234435 | 1616305 | 2196987.75 | 2121588.75 | 1153266.5 | 2336226.5 | 2447206.75 | 1596459 | 2515263.25 | 1735492.5 | 653339.19 | 1645947 | 2619429.25 | 1948265.75 | 1758723.75 | 2 | 1310361.62 | 7 | 8 | 8 | 8 | 8 | 8 | 0.759965698 | 0.794371672 | 0.71241539 | 0.744668616 |
| Q9EQ32 | Pik3ap1 | Phosphoinositide 3-kinase adapter protein 1 | 496.07 | 12.08 | 90928 | 7 | 6059862 | 4418039 | 6732431.5 | 8317002.5 | 6404953.5 | 5592814.5 | 5154366.5 | 4355602.5 | 5159243 | 5644535 | 4363215 | 4029316.25 | 3035976.5 | 3029184.5 | 4195602 | 4906447 | 5975102 | 5712171.5 | 7062495 | 5914852.5 | 0 | 8 | 1 | 4 | 1 | 8 | 0.584151501 | 0.720022001 | 0.926055394 | 1.141450902 |
| P26450 | Pik3r1 | Phosphatidylinositol 3-kinase regulatory subunit alpha | 54.75 | 1.66 | 83517 | 1 | 524096.72 | 472003.22 | 439129.81 | 626613.06 | 568276 | 574040.56 | 436526.25 | 516559.47 | 539121.19 | 629508.12 | 717832.69 | 661966.69 | 704044.44 | 470272.94 | 568379.81 | 156168.22 | 473059.09 | 658941.5 | 653174 | 555072.56 | 5 | 10 | 6 | 11 | 6 | 11 | 1.187207421 | 1.158301065 | 0.949164487 | 0.926054046 |
| P0C0R5 | Pik3r4 | Phosphoinositide 3-kinase regulatory subunit 4 | 181.25 | 2.8 | 152447 | 3 | 147177.28 | 353601.09 | 153198.69 | 79002.98 | 1623440.88 | 2 | 521628.75 | 2 | 837525.25 | 2 | 62135.96 | 479261.88 | 141326.27 | 322631.41 | 279366.88 | 492268.72 | 2 | 2 | 90819.21 | 2 | 11 | 5 | 10 | 10.5 | 11 | 10.5 | 0.545200728 | 0.945232644 | 0.24744897 | 0.429010514 |
| Q2YFS1 | Pilrb2 | Paired immunoglobulin-like type 2 receptor beta-2 | 61.98 | 4 | 25513 | 1 | 1101868.25 | 1176262.5 | 1215418.88 | 1387010.25 | 1134763.25 | 1059051.38 | 971244 | 1140584.25 | 957777.25 | 682217.5 | 1116559.75 | 1113472 | 958727.31 | 1022001.75 | 1316322.62 | 1470758.88 | 1420015.12 | 869280.56 | 796258.62 | 1102560.88 | 6 | 11 | 7 | 9 | 7 | 11 | 0.918834003 | 1.148872948 | 0.940743155 | 1.176267267 |
| Q9QUR7 | Pin1 | Peptidyl-prolyl cis-trans isomerase NIMA-interacting 1 | 99.42 | 18.18 | 18370 | 2 | 4954013.5 | 5675564.5 | 3293099.5 | 5725231.5 | 4869648 | 5687853 | 5594234 | 5397477 | 5068953 | 4392456.5 | 4793596 | 4051810.25 | 3620737.5 | 4376696 | 3423699.75 | 6155299 | 6188597 | 4718935 | 4660450 | 5375308 | 5 | 10 | 1 | 11 | 5 | 11 | 0.826613333 | 0.775278683 | 1.105272805 | 1.036632741 |
| Q9CZX5 | Pinx1 | PIN2/TERF1-interacting telomerase inhibitor 1 | 49.69 | 5.72 | 37221 | 1 | 434186.31 | 374690.06 | 283453.12 | 462345.75 | 2 | 462069.16 | 258316.38 | 437336.34 | 180248.7 | 2 | 402078.19 | 362321.53 | 2 | 2 | 2 | 343084.97 | 419179 | 2 | 307813.97 | 2 | 6.5 | 8 | 7.5 | 10 | 7.5 | 10 | 0.491681296 | 0.571316432 | 0.688298454 | 0.799778677 |
| O70172 | Pip4k2a | Phosphatidylinositol 5-phosphate 4-kinase type-2 alpha | 89.4 | 4.94 | 46152 | 2 | 1657854.62 | 1832224.12 | 1959764.5 | 2341474 | 1413628.38 | 1500180.25 | 1602028.38 | 2063010.38 | 1783427.38 | 2130366.75 | 784693.56 | 1253266.38 | 1203658.62 | 1436902.62 | 1040269.25 | 1600702.25 | 1756655.75 | 1959212.25 | 1893907 | 1930600.12 | 1 | 12 | 0 | 12 | 1 | 12 | 0.621273679 | 0.629891194 | 0.993061529 | 1.006836011 |
| O88377 | Pip4k2b | Phosphatidylinositol 5-phosphate 4-kinase type-2 beta | 127.31 | 5.53 | 47264 | 2 | 819356.75 | 899310.19 | 654602.19 | 832414.31 | 792651.31 | 485738.56 | 663067 | 730967.56 | 810076.06 | 806737.56 | 2 | 671010.69 | 558062.56 | 641535.62 | 850560.44 | 762366.88 | 856227.88 | 717305.5 | 1059488.5 | 813251.69 | 5 | 12 | 9 | 5 | 9 | 12 | 0.68057616 | 0.778236467 | 1.052598322 | 1.203642513 |
| Q4V888 | Pip4p2 | Type 2 phosphatidylinositol 4,5-bisphosphate 4-phosphatase | 49.23 | 3.89 | 28024 | 1 | 550695 | 647970.12 | 832968.31 | 793424.19 | 712161.88 | 675304.5 | 2 | 1017928.62 | 2 | 1066499.5 | 1069712.5 | 111189.23 | 1561643.25 | 1919158.12 | 1756438.62 | 768608.19 | 687737.06 | 1077277.88 | 1349595.5 | 1300067.5 | 5 | 5 | 3 | 4 | 5 | 5 | 1.814459555 | 2.325635596 | 1.465356088 | 1.878181451 |
| D3ZSI8 | Pip5k1a | Phosphatidylinositol 4-phosphate 5-kinase type-1 alpha | 64.07 | 1.83 | 60605 | 1 | 2 | 341267.72 | 264850.38 | 2 | 287595.25 | 399540.88 | 388713.47 | 325673.47 | 2 | 2 | 565857 | 2 | 572651.62 | 603787.88 | 766744.75 | 2 | 507374.28 | 2 | 540450.19 | 2 | 4 | 12 | 4 | 12 | 4 | 12 | 2.807423678 | 2.252420844 | 1.17244056 | 0.940659429 |
| P16446 | Pitpna | Phosphatidylinositol transfer protein alpha isoform | 187.84 | 18.82 | 31907 | 4 | 4334530 | 6674632 | 8077067.5 | 6962338 | 7447116 | 6006457 | 5015879 | 5946294.5 | 5919177.5 | 5316310.5 | 2788739.75 | 3326276.5 | 2094219.75 | 2520374.5 | 2649711 | 5610138 | 6759085.5 | 4754622 | 4716331.5 | 6703671 | 0 | 7 | 0 | 12 | 0 | 12 | 0.399434199 | 0.474374744 | 0.852164966 | 1.012045386 |
| Q8K411 | Pitrm1 | Presequence protease, mitochondrial | 214.48 | 6.85 | 117372 | 5 | 5257413 | 4584484 | 5023033.5 | 6147054 | 4546210 | 4981721 | 5459766 | 3926837.75 | 5618652 | 5169587 | 2540932.25 | 4880542 | 4523107.5 | 2932450.25 | 5068099.5 | 4844412.5 | 3755518.5 | 4555907.5 | 4237047.5 | 4871047 | 5 | 5 | 4 | 4 | 5 | 5 | 0.780381083 | 0.792840059 | 0.871107425 | 0.88501487 |
| P52480 | Pkm | Pyruvate kinase PKM | 4233.3 | 76.46 | 57845 | 56 | 337436226 | 351386146 | 318649762 | 252415826 | 329705762 | 329690498 | 344643714 | 438878754 | 385213602 | 239953474 | 363863298 | 332355202 | 297494018 | 242701602 | 316238562 | 364194434 | 248642642 | 307578562 | 245159890 | 358778050 | 10 | 11 | 8 | 9 | 10 | 11 | 0.976760703 | 0.893160669 | 0.958957976 | 0.876881661 |
| P70268 | Pkn1 | Serine/threonine-protein kinase N1 | 268.76 | 6.66 | 104411 | 6 | 2794838.25 | 3707283 | 3453166 | 3298202.5 | 3197688 | 2347292.5 | 1829337.25 | 1852593.62 | 2811700 | 2822884.25 | 13831608 | 9798076 | 9535423 | 6215663 | 9375427 | 3693195.5 | 2468044.5 | 2159943 | 1750645.38 | 2247562 | 0 | 4 | 0 | 12 | 0 | 12 | 2.963690366 | 4.180126987 | 0.748845497 | 1.056206582 |
| Q68FH0 | Pkp4 | Plakophilin-4 | 41.65 | 0.67 | 131551 | 1 | 7062563.5 | 6936964.5 | 7689407.5 | 10521566 | 5855743.5 | 8480594 | 8160033.5 | 6514535.5 | 5543420 | 6851786.5 | 3092083.25 | 2310216.25 | 2580117.5 | 3805593.75 | 2900856.5 | 5550161 | 8042114.5 | 4957205.5 | 6392400.5 | 5366923.5 | 0 | 5 | 0 | 5 | 0 | 5 | 0.385876444 | 0.413184658 | 0.796212103 | 0.852559493 |
| Q8VEB4 | Pla2g15 | Group XV phospholipase A2 | 129.49 | 8.74 | 47307 | 3 | 4583690 | 4575479 | 5407930.5 | 5392345 | 4804198 | 3885929 | 3200262.75 | 3834957.25 | 3377880.5 | 4688599.5 | 1551756.38 | 987242.62 | 901778.94 | 987062.81 | 957817.5 | 3266837.5 | 3807730.25 | 3586422 | 4223966.5 | 2784121.25 | 0 | 0 | 0 | 9 | 0 | 9 | 0.217482475 | 0.283640377 | 0.713508826 | 0.930557338 |
| P47713 | Pla2g4a | Cytosolic phospholipase A2 | 747.37 | 25.8 | 85222 | 15 | 20304466 | 23770996 | 28871294 | 27812798 | 26023960 | 22527170 | 20861248 | 25426322 | 23689294 | 27097000 | 35088102 | 34042922 | 36816182 | 34356030 | 36528722 | 15799849 | 17825812 | 25708868 | 20152282 | 27661306 | 0 | 5 | 0 | 9 | 0 | 9 | 1.394755141 | 1.478515294 | 0.845126575 | 0.895879521 |
| P27612 | Plaa | Phospholipase A-2-activating protein | 446.93 | 13.48 | 87221 | 9 | 4687275 | 5492631 | 6861951 | 7975981.5 | 7839168 | 5989341.5 | 6159220.5 | 6894608.5 | 6250044 | 7307260 | 7831535 | 5802418 | 6758734 | 7832660 | 5903843 | 7495396.5 | 7272310.5 | 9415640 | 8515547 | 9100257 | 12 | 4 | 12 | 1 | 12 | 4 | 1.038718789 | 1.046892431 | 1.272153353 | 1.282163884 |
| Q3TCN2 | Plbd2 | Putative phospholipase B-like 2 | 401.85 | 11.95 | 66289 | 7 | 15179914 | 12461482 | 8577093 | 11910513 | 13111266 | 10340157 | 9010975 | 8689947 | 9978718 | 10127012 | 1463256 | 654220.38 | 2757886 | 1020292.62 | 2518007.25 | 10568254 | 10977196 | 7184815.5 | 8642018 | 7822277 | 0 | 3 | 0 | 10 | 0 | 10 | 0.137387744 | 0.174750153 | 0.737987634 | 0.938682364 |
| P51432 | Plcb3 | 1-phosphatidylinositol 4,5-bisphosphate phosphodiesterase beta-3 | 90.36 | 1.46 | 139487 | 2 | 377906.88 | 539324.44 | 71637.41 | 560897.5 | 929555.31 | 671609.69 | 2 | 300658.56 | 844578.88 | 509266.03 | 575614.88 | 560455.38 | 515098.25 | 2 | 79993.91 | 344301.22 | 373038.81 | 570606 | 329090.44 | 2 | 10 | 7 | 10.5 | 9.5 | 10.5 | 9.5 | 0.698241189 | 0.744229886 | 0.652210068 | 0.695166988 |
| P24135 | Plcg2 | 1-phosphatidylinositol 4,5-bisphosphate phosphodiesterase gamma-2 | 572.5 | 10.2 | 147735 | 12 | 8884860 | 8867481 | 11872694 | 12902016 | 10641579 | 9961583 | 9455840 | 10208348 | 11092273 | 12054423 | 8135882 | 8688266 | 8352625 | 9221494 | 7277491 | 8102764.5 | 7765962.5 | 10854340 | 11368832 | 9631962 | 2 | 8 | 0 | 8 | 2 | 8 | 0.783841111 | 0.789725407 | 0.897594333 | 0.904332576 |
| O35405 | Pld3 | 5'-3' exonuclease PLD3 | 392.14 | 20.7 | 54389 | 7 | 3214267.25 | 3539842 | 2436896.75 | 2722223 | 3665813.75 | 4079438 | 4483354.5 | 5423772.5 | 6757396 | 6038630.5 | 6374901.5 | 7407418 | 9429739 | 7659524.5 | 9040842 | 6292829 | 5525868.5 | 7797509 | 9898626 | 8771491 | 0 | 0 | 1 | 3 | 1 | 3 | 2.561930514 | 1.490237605 | 2.457553016 | 1.429522737 |
| Q8BG07 | Pld4 | 5'-3' exonuclease PLD4 | 1200.8 | 35.19 | 56154 | 17 | 120049650 | 105438362 | 133711450 | 144558450 | 136347266 | 108612434 | 107793666 | 121716666 | 124928306 | 114563858 | 33612342 | 31923874 | 45000322 | 43251086 | 47578250 | 130859050 | 92868234 | 136091410 | 116522178 | 137899298 | 0 | 10 | 0 | 7 | 0 | 10 | 0.314582479 | 0.348616117 | 0.959592566 | 1.06340771 |
| Q9QXS1 | Plec | Plectin | 10197.982 | 43.42 | 534188 | 176 | 283135394 | 289198786 | 241092738 | 283008674 | 241869634 | 291916386 | 282417794 | 237270754 | 234553794 | 256923426 | 288529474 | 262455442 | 199927042 | 220777426 | 220709730 | 280970050 | 301676866 | 245441794 | 257952994 | 239472850 | 6 | 11 | 7 | 10 | 7 | 11 | 0.890976954 | 0.915060582 | 0.990442635 | 1.017214878 |
| Q9JHK5 | Plek | Pleckstrin | 611.54 | 42.86 | 39901 | 10 | 5096691.5 | 4976140.5 | 4315840.5 | 2013684.25 | 5440159 | 4564926 | 4514470 | 5465476 | 6743922.5 | 5002198.5 | 11522178 | 10172676 | 11221171 | 9560889 | 9548890 | 5431285.5 | 6585702.5 | 6063172 | 6508705 | 5187562 | 0 | 2 | 0 | 7 | 0 | 7 | 2.381863637 | 1.978848574 | 1.363232484 | 1.13257141 |
| Q9ERS5 | Plekha2 | Pleckstrin homology domain-containing family A member 2 | 132.53 | 8.47 | 47380 | 3 | 2160473.5 | 1825008.5 | 1935697 | 2790444.25 | 2953172 | 3507258.5 | 2037350.88 | 2906534 | 2679759.5 | 3281551 | 2497631.5 | 2792465.5 | 3029217 | 2701758.5 | 2285126 | 2108408 | 2471202.75 | 3374016 | 3071162.75 | 2690811.75 | 7 | 7 | 9 | 11 | 9 | 11 | 1.140714279 | 0.923243093 | 1.175811573 | 0.951649273 |
| Q9QZC7 | Plekhb2 | Pleckstrin homology domain-containing family B member 2 | 45.78 | 3.17 | 24574 | 1 | 934162 | 778547.69 | 740760.69 | 984255.5 | 4010127.25 | 1095018.12 | 4222335 | 1375667.62 | 1076331.75 | 944226.69 | 1388310.5 | 1408626.38 | 2133456.5 | 1891817.38 | 2307627 | 4302740.5 | 4907538 | 1388066.5 | 1291564.62 | 1319999.5 | 5 | 3 | 5 | 5 | 5 | 5 | 1.225834828 | 1.047771251 | 1.773653278 | 1.516014125 |
| Q8K124 | Plekho2 | Pleckstrin homology domain-containing family O member 2 | 627.21 | 33.74 | 53872 | 10 | 8577738 | 9347304 | 12693868 | 10578472 | 12283757 | 4981820.5 | 10098154 | 17708800 | 18567624 | 16547193 | 8464597 | 11097413 | 8788309 | 6610923 | 9299608 | 8033445.5 | 6803798 | 9434458 | 16902352 | 7960570.5 | 5 | 7 | 6 | 7 | 6 | 7 | 0.82759737 | 0.651818984 | 0.918728077 | 0.723593891 |
| P20918 | Plg | Plasminogen | 284.73 | 7.76 | 90808 | 6 | 326230.09 | 470106.72 | 4163355 | 3624761.25 | 3971794.5 | 857762.12 | 2183987.5 | 10145757 | 9366997 | 11139141 | 14473096 | 14243628 | 31927990 | 34670938 | 31982884 | 1445346 | 1021498.12 | 9385100 | 8204556.5 | 5765996 | 0 | 6 | 0 | 9 | 0 | 9 | 10.13826268 | 3.778117133 | 2.056545675 | 0.766390722 |
| P43883 | Plin2 | Perilipin-2 | 128.49 | 7.06 | 46646 | 2 | 1024077.44 | 969294.88 | 520466.09 | 1776630.75 | 2165953.5 | 1477168.5 | 1184535.12 | 2508993 | 2292908.75 | 2810969 | 2 | 2 | 2 | 2 | 265069.78 | 469597.19 | 849910.56 | 2204371 | 1001002.19 | 1673552.25 | 0 | 11 | 0 | 4 | 0 | 11 | 0.041056448 | 0.025799393 | 0.960041422 | 0.603278829 |
| Q9DBG5 | Plin3 | Perilipin-3 | 866.94 | 46.45 | 47262 | 13 | 30482700 | 27510904 | 25634202 | 32248348 | 26769156 | 29204616 | 26996374 | 22872160 | 25499482 | 28311760 | 13355199 | 12459760 | 12355926 | 13648157 | 12718288 | 30113760 | 28521308 | 27478300 | 28015446 | 27981612 | 0 | 11 | 0 | 7 | 0 | 11 | 0.45243219 | 0.485665239 | 0.996250252 | 1.069429027 |
| Q63321 | Plod1 | Procollagen-lysine,2-oxoglutarate 5-dioxygenase 1 | 148.92 | 4.4 | 83612 | 3 | 797069.25 | 820683.44 | 789825.5 | 683662 | 508471.25 | 625569.31 | 522607.38 | 660714.81 | 838491.44 | 824516.88 | 973754.69 | 650959.5 | 1044753.5 | 1117535.62 | 1225728.88 | 485274.69 | 898788.12 | 482460.66 | 765427.62 | 804725.56 | 4 | 11 | 3 | 11 | 4 | 11 | 1.392537228 | 1.44380093 | 0.954708928 | 0.989854785 |
| Q9R0E1 | Plod3 | Multifunctional procollagen lysine hydroxylase and glycosyltransferase LH3 | 566.03 | 16.73 | 84922 | 9 | 1540980.75 | 678887.75 | 2109299.5 | 1516739.75 | 533720.06 | 1367653.25 | 476450.5 | 825457.81 | 1440722.75 | 1521858.88 | 3359996.75 | 2470266.5 | 3705967.75 | 1699541.25 | 2976689 | 740764.31 | 1082651.12 | 2631754.75 | 2241530.25 | 2085699.5 | 1 | 7 | 0 | 7 | 1 | 7 | 2.227788466 | 2.523455241 | 1.376632022 | 1.559335342 |
| Q6P742 | Plp2 | Proteolipid protein 2 | 78.35 | 9.93 | 16557 | 1 | 2 | 2 | 2 | 603335.75 | 2 | 574282.62 | 773979.31 | 2 | 2 | 2 | 719333 | 2 | 1579937.62 | 1494082.88 | 1215933.62 | 2 | 2 | 2 | 898829.88 | 748789.75 | 3 | 9 | 4.5 | 11.5 | 4.5 | 11.5 | 8.302545804 | 3.715351384 | 2.730824062 | 1.222031314 |
| Q9Z2Y8 | Plpbp | Pyridoxal phosphate homeostasis protein | 47.28 | 6.57 | 30049 | 1 | 225191.78 | 2 | 197970.09 | 244412.12 | 88358.5 | 184496.12 | 156861.16 | 145925 | 190755.42 | 277854.5 | 2 | 2 | 204363.7 | 2 | 144915.77 | 47669.98 | 1062542.88 | 340296.31 | 151309.83 | 165802.75 | 6.5 | 10 | 4 | 12 | 6.5 | 12 | 0.462057856 | 0.365402574 | 2.338326632 | 1.849185243 |
| Q922V4 | Plrg1 | Pleiotropic regulator 1 | 239.59 | 8.19 | 56938 | 3 | 1099183.75 | 1709131.88 | 1050345.62 | 945956.5 | 1307281.38 | 1517700.25 | 862576.19 | 2 | 776571.38 | 2 | 463060.88 | 1118765.88 | 469766.84 | 749254.88 | 921252.38 | 1269877.12 | 1044692.69 | 1412947.25 | 1353554.5 | 571781.31 | 3 | 12 | 11 | 7 | 11 | 12 | 0.608992521 | 1.179054663 | 0.924893024 | 1.790661454 |
| Q99K51 | Pls3 | Plastin-3 | 1679.24 | 49.68 | 70742 | 26 | 46552794 | 49407098 | 31107656 | 32716232 | 51821034 | 41262362 | 46925406 | 38586450 | 46238054 | 37804018 | 26564152 | 34741274 | 23556636 | 22482366 | 24452356 | 44935010 | 39437606 | 36093254 | 39998378 | 30868706 | 2 | 8 | 0 | 7 | 2 | 8 | 0.622843977 | 0.625173624 | 0.904199438 | 0.907581449 |
| Q9JJ00 | Plscr1 | Phospholipid scramblase 1 | 162.67 | 12.2 | 35914 | 3 | 2734007.25 | 2200552.25 | 2033838.12 | 2582331 | 2311344.5 | 3207438.75 | 2792996 | 2148899.25 | 2478296.5 | 2861763.25 | 4098703.75 | 3502901.5 | 3833390.25 | 3879944.5 | 4522373.5 | 2032689.25 | 1771751.88 | 2018416.38 | 2451088.5 | 2115863.25 | 0 | 4 | 0 | 1 | 0 | 4 | 1.672331076 | 1.470585993 | 0.875884776 | 0.770220623 |
| P70206 | Plxna1 | Plexin-A1 | 1223.66 | 14.94 | 211099 | 23 | 5802266.5 | 6671794.5 | 7855508.5 | 6443910 | 7088993.5 | 8365015.5 | 9139928 | 8274161 | 12718988 | 11241703 | 17555850 | 20203850 | 19241566 | 12703895 | 16273512 | 10199227 | 5724312.5 | 7951007.5 | 8000643 | 9183523 | 0 | 5 | 1 | 6 | 1 | 6 | 2.539054752 | 1.728569089 | 1.212513717 | 0.825470081 |
| B2RXS4 | Plxnb2 | Plexin-B2 | 1359.45 | 17.21 | 206230 | 25 | 25534834 | 23443634 | 32864784 | 26860762 | 28308316 | 21641954 | 23580258 | 27020278 | 27237696 | 23928962 | 19569594 | 21943484 | 22958758 | 20594562 | 25729034 | 23891692 | 21547014 | 24184530 | 21458946 | 22241724 | 2 | 2 | 5 | 6 | 5 | 6 | 0.808653002 | 0.897789457 | 0.827107356 | 0.918278003 |
| Q9CPV5 | Pmf1 | Polyamine-modulated factor 1 | 71.58 | 5.94 | 23121 | 1 | 344803.06 | 429379.69 | 394110.03 | 2 | 508126.56 | 2 | 426630.81 | 306877.38 | 389179.22 | 2 | 2 | 2 | 2 | 2 | 100287.27 | 357906.84 | 2 | 2 | 2 | 2 | 3 | 4 | 6 | 7 | 6 | 7 | 0.059827006 | 0.089334673 | 0.213499334 | 0.318800729 |
| Q60953 | Pml | Protein PML | 369.02 | 9.38 | 98242 | 7 | 2434977.5 | 2824862.25 | 1907620.75 | 2021042 | 2257741 | 1782985.25 | 2631081 | 1491793.25 | 2163519.5 | 2475898.5 | 1911347.88 | 359467.41 | 597221.38 | 199063.78 | 1059082.25 | 2196858.25 | 1459712.75 | 1234477 | 865845.25 | 1666983.38 | 1 | 2 | 2 | 4 | 2 | 4 | 0.360483568 | 0.391282515 | 0.648586292 | 0.70400012 |
| Q03346 | Pmpcb | Mitochondrial-processing peptidase subunit beta | 172.88 | 7.98 | 54265 | 3 | 1403334 | 2 | 1731594.12 | 2273662.75 | 2 | 1613646.62 | 1761746.38 | 2109969 | 2 | 1699691.75 | 1171864.75 | 2 | 221789.14 | 2 | 1504408.25 | 1334741.25 | 2 | 2 | 1852035.62 | 2200768.25 | 9 | 12 | 4 | 11 | 9 | 12 | 0.535826071 | 0.403346368 | 0.996108832 | 0.749827045 |
| Q9D1G2 | Pmvk | Phosphomevalonate kinase | 260.4 | 31.77 | 21916 | 5 | 3636894 | 2541403.25 | 2030934 | 2444370.25 | 2144110 | 3805029.25 | 2841825.5 | 1888259.75 | 2786276.75 | 2813759.5 | 5370741 | 4009593.5 | 4215927 | 3455477.5 | 4213495 | 3885400 | 3226406 | 3750922 | 3528626 | 2805751 | 1 | 3 | 1 | 6 | 1 | 6 | 1.66164349 | 1.504422158 | 1.343764078 | 1.216619851 |
| Q9JLV6 | Pnkp | Bifunctional polynucleotide phosphatase/kinase | 219.71 | 9 | 57223 | 4 | 3289695.5 | 3588072.75 | 2826474.5 | 2814974.5 | 2932938.5 | 2851758.5 | 2716602.5 | 4310539 | 2365091 | 2494353.75 | 2300085 | 1560814.75 | 1986790.75 | 2193547.5 | 2693038 | 2373733 | 3041876.25 | 2744351.75 | 3106984.5 | 3450823 | 0 | 10 | 2 | 9 | 2 | 10 | 0.694678217 | 0.728323036 | 0.952473476 | 0.998603897 |
| O35691 | Pnn | Pinin | 205.47 | 5.1 | 82436 | 4 | 2 | 823435.75 | 780828.19 | 689113.94 | 842637 | 749221.81 | 552581.12 | 406521 | 720993.5 | 1111089.75 | 1438037 | 812780.12 | 1307611 | 1075053.25 | 659047.75 | 344207.91 | 1075959.12 | 2 | 518094.47 | 370822.94 | 6 | 8.5 | 5 | 5 | 6 | 8.5 | 1.68765964 | 1.494892777 | 0.736311866 | 0.652209286 |
| Q9CPS7 | Pno1 | RNA-binding protein PNO1 | 203.2 | 21.37 | 27454 | 4 | 2 | 1239292.5 | 482020 | 1158044.62 | 801052.25 | 1377266.75 | 1354982 | 2351657.75 | 946814.81 | 277599.94 | 1677762.75 | 1702677.75 | 1664910.75 | 783242.38 | 1811064.88 | 1757619.5 | 100004.04 | 1158817.88 | 1203099.25 | 1109276.5 | 3 | 8 | 8 | 10 | 8 | 10 | 2.075762121 | 1.21104462 | 1.447886292 | 0.84472825 |
| P23492 | Pnp | Purine nucleoside phosphorylase | 598.39 | 31.83 | 32277 | 8 | 33232470 | 28558646 | 21936766 | 24280106 | 34489858 | 27551758 | 32179432 | 23696864 | 30479366 | 26921534 | 12566172 | 13037157 | 11674018 | 9223774 | 11664254 | 30929402 | 21531506 | 13419056 | 16808962 | 27898828 | 0 | 5 | 0 | 7 | 0 | 7 | 0.40818424 | 0.413021423 | 0.776066145 | 0.785262908 |
| Q8K1R3 | Pnpt1 | Polyribonucleotide nucleotidyltransferase 1, mitochondrial | 629.51 | 15.84 | 85683 | 11 | 11703812 | 12161811 | 9874756 | 12148300 | 10172642 | 12287917 | 11828411 | 11477675 | 12821094 | 11341771 | 7676404 | 10030360 | 11361609 | 5541631 | 12477436 | 10885593 | 5198780 | 9639358 | 9117209 | 11283243 | 8 | 4 | 5 | 0 | 8 | 4 | 0.839927407 | 0.787983734 | 0.822745204 | 0.771864131 |
| Q91ZW2 | Pofut1 | GDP-fucose protein O-fucosyltransferase 1 | 160.89 | 7.38 | 44688 | 3 | 1658382 | 1274473.38 | 1267832.75 | 1824938 | 930477.06 | 1861231 | 1222118.62 | 964218.06 | 1246718.62 | 1832912.5 | 2317807.5 | 1488719.38 | 1790652.88 | 1961104.25 | 1950898.38 | 2248326.5 | 2377462 | 1709674.25 | 1395890.75 | 1466309.38 | 3 | 5 | 4 | 6 | 4 | 6 | 1.367027218 | 1.334210348 | 1.322243594 | 1.2905018 |
| Q8VHI3 | Pofut2 | GDP-fucose protein O-fucosyltransferase 2 | 176.23 | 9.79 | 49429 | 3 | 224675.31 | 752456.38 | 495950.53 | 266797.56 | 896859.69 | 273511.84 | 240135.22 | 222160.39 | 393940.94 | 813835.5 | 3736976.5 | 593263.06 | 961626.75 | 1859086.75 | 3320988.5 | 500259.84 | 813555.38 | 951140 | 786294.38 | 4617949.5 | 2 | 4 | 1 | 3 | 2 | 4 | 3.97154959 | 5.387954497 | 2.908591913 | 3.94590588 |
| P33611 | Pola2 | DNA polymerase alpha subunit B | 89.24 | 3.5 | 66214 | 2 | 105468.44 | 222482.59 | 2 | 2 | 134266.83 | 2 | 2 | 117935.12 | 194364.91 | 112813.83 | 279290.62 | 925878 | 792196.88 | 2 | 976449.38 | 70031.88 | 2 | 2 | 2 | 2 | 4 | 6 | 4 | 6 | 4 | 6 | 6.433743484 | 6.995276275 | 0.151528705 | 0.164754028 |
| P52431 | Pold1 | DNA polymerase delta catalytic subunit | 386.65 | 8.24 | 123790 | 9 | 6121028 | 6144467.5 | 6871847 | 7219649 | 6500632.5 | 5602079.5 | 6274964 | 5504089 | 6514986 | 5761421.5 | 6187513.5 | 5802157 | 5736734.5 | 6146066.5 | 4152580.25 | 4586546 | 5478202.5 | 4940065.5 | 20320286 | 5890061.5 | 4 | 5 | 11 | 8 | 11 | 8 | 0.852923868 | 0.944955372 | 1.254356112 | 1.389702635 |
| O35654 | Pold2 | DNA polymerase delta subunit 2 | 195.47 | 5.97 | 51355 | 2 | 2045291.12 | 2232080.25 | 2495036 | 2252533.75 | 994885.94 | 2332045.75 | 2151803.75 | 1829730.62 | 2073569.75 | 2652392.75 | 1570398 | 1642892.38 | 1993027.5 | 1500540.25 | 1456470.5 | 1528449.75 | 1169493.88 | 1686549.25 | 1758834.62 | 2032533.25 | 5 | 5 | 1 | 1 | 5 | 5 | 0.814717518 | 0.739462577 | 0.81596825 | 0.740597779 |
| Q9EQ28 | Pold3 | DNA polymerase delta subunit 3 | 79.98 | 5.84 | 50836 | 2 | 712987.12 | 730129.62 | 685763.94 | 791646 | 2 | 885333.12 | 572116.88 | 664230.31 | 2 | 627795.44 | 457080.94 | 2 | 490151.66 | 629989.94 | 2 | 2 | 553650 | 2 | 861877 | 781480.75 | 4 | 11 | 6 | 10 | 6 | 11 | 0.540048297 | 0.573645864 | 0.752265083 | 0.799065113 |
| Q91VA6 | Poldip2 | Polymerase delta-interacting protein 2 | 169.32 | 9.78 | 41870 | 3 | 917589 | 1903049.5 | 1627892.75 | 1193300.12 | 1410304.25 | 1643022.5 | 1585765 | 1636734.88 | 1657868.62 | 1703751.38 | 1646655.38 | 1611164.62 | 1093020.12 | 1388500.12 | 1836376 | 1196013 | 704310.56 | 2099052.5 | 785429.38 | 1055891.75 | 11 | 8 | 9 | 5 | 11 | 8 | 1.074244264 | 0.920819878 | 0.828216799 | 0.709930243 |
| Q8BG81 | Poldip3 | Polymerase delta-interacting protein 3 | 55.34 | 3.33 | 46132 | 1 | 2 | 2 | 2 | 452033.59 | 2 | 595802.31 | 2 | 2 | 2 | 561545.44 | 491862.81 | 290539.03 | 860184.25 | 661886.25 | 318089.09 | 2 | 860556.88 | 2 | 396349.38 | 425390.25 | 2 | 8 | 6 | 11 | 6 | 11 | 5.801593234 | 2.265998127 | 3.72156135 | 1.453575028 |

| | | | | | | | | | | | | | | | | | | | | | | | | | | | | | | | | | | | | |
|---|---|---|---|---|---|---|---|---|---|---|---|---|---|---|---|---|---|---|---|---|---|---|---|---|---|---|---|---|---|---|---|---|---|---|---|---|
| O35134 | Polr1a | DNA-directed RNA polymerase I subunit RPA1 | 210.68 | 4.89 | 194110 | 5 | 1067365.62 | 624039.44 | 747117.25 | 727234.5 | 757911.25 | 709639.31 | 922194 | 760081.38 | 647730 | 599176.75 | 2983033.75 | 2481751 | 3783009 | 2676764 | 2080231.5 | 1238834 | 634122.44 | 621126.25 | 665948.19 | 628923.88 | 0 | 8 | 0 | 10 | 0 | 10 | 3.56931041 | 3.848715712 | 0.965666489 | 1.041258776 |
| P52432 | Polr1c | DNA-directed RNA polymerases I and III subunit RPAC1 | 77.61 | 6.94 | 39107 | 2 | 463302.22 | 2 | 1495342.38 | 373903 | 1387992.25 | 256417.94 | 1167531.75 | 2 | 612997.19 | 294847.69 | 1630705.12 | 586441.69 | 466385.41 | 1315186.5 | 925527.38 | 367262.84 | 284355.59 | 1099666.62 | 2 | 267678 | 8 | 6.5 | 5 | 11.5 | 8 | 11.5 | 1.323529286 | 2.111782033 | 0.542653498 | 0.86584099 |
| Q8K202 | Polr1e | DNA-directed RNA polymerase I subunit RPA49 | 50.55 | 2.49 | 54034 | 1 | 477435.28 | 2 | 460464.78 | 218875.02 | 441425.44 | 2 | 464957.97 | 2 | 2 | 393331.31 | 454601.53 | 254315.3 | 133646.89 | 369987.62 | 269795.34 | 655829.12 | 515054.53 | 2 | 436908.16 | 355175.62 | 10 | 10.5 | 9 | 6.5 | 10 | 10.5 | 0.927508661 | 1.727082409 | 1.228235724 | 2.28705607 |
| P08775 | Polr2a | DNA-directed RNA polymerase II subunit RPB1 | 461.58 | 5.63 | 217176 | 8 | 3836853.5 | 3972113.5 | 3834308.75 | 3999451.25 | 3167988.25 | 4937316.5 | 3347063.75 | 3216390 | 3020126.5 | 3675897.5 | 4101650 | 3320818.5 | 3884468.75 | 3260403 | 3442556 | 3308162.25 | 3828906.25 | 3873516.5 | 3129778 | 3113307.5 | 11 | 5 | 10 | 12 | 11 | 12 | 0.957427509 | 0.98972907 | 0.917225649 | 0.948170885 |
| Q8CFI7 | Polr2b | DNA-directed RNA polymerase II subunit RPB2 | 264.39 | 4.43 | 133911 | 6 | 1606812.88 | 1211215.25 | 289978.16 | 714416.44 | 1234469 | 301079.06 | 267083.47 | 443392.25 | 267658.25 | 399868.97 | 1677612.75 | 1891453.5 | 1661590.25 | 953473.06 | 1785325.25 | 873458 | 411680.97 | 455025.03 | 950150.38 | 717783.75 | 3 | 8 | 0 | 1 | 3 | 8 | 1.575959153 | 4.746316624 | 0.673951176 | 2.029738947 |
| P97760 | Polr2c | DNA-directed RNA polymerase II subunit RPB3 | 176.97 | 9.09 | 31443 | 2 | 554964.12 | 2 | 295332.69 | 220228.97 | 2 | 507808.72 | 226971.39 | 2 | 700156.5 | 274138 | 511248.12 | 233687.05 | 217715.52 | 358872.25 | 620498.38 | 291559.56 | 962344.88 | 519910.81 | 428651.03 | 595178.12 | 7 | 4 | 11 | 6 | 11 | 6 | 1.814075009 | 1.136298577 | 2.61332702 | 1.636933291 |
| B0BNE2 | Polr2e | DNA-directed RNA polymerases I, II, and III subunit RPABC1 | 59.63 | 6.67 | 24570 | 1 | 2 | 2 | 2 | 2 | 2 | 2 | 2 | 2 | 2 | 2 | 939475.38 | 411923.03 | 2 | 2 | 2 | 2 | 2 | 2 | 2 | 2 | 7.5 | 12.5 | 7.5 | 12.5 | 7.5 | 12.5 | 135140.441 | 135140.441 | 1 | 1 |
| P62488 | Polr2g | DNA-directed RNA polymerase II subunit RPB7 | 43.44 | 6.4 | 19294 | 1 | 1095990.88 | 882914.06 | 1167766 | 1022358.31 | 1081986.25 | 1133683.25 | 1256815.25 | 993077.94 | 923100.06 | 748197.25 | 418286.31 | 737084.56 | 564765.06 | 360489.62 | 437070.12 | 1160868.5 | 980921.44 | 929795.81 | 364012.09 | 692128.44 | 0 | 6 | 0 | 8 | 0 | 8 | 0.479468337 | 0.498072908 | 0.786081527 | 0.816583457 |
| Q923G2 | Polr2h | DNA-directed RNA polymerases I, II, and III subunit RPABC3 | 155.02 | 25.33 | 17143 | 3 | 505627.34 | 616863.06 | 2 | 2 | 2 | 2 | 2 | 2 | 309091.12 | 2 | 661360.5 | 2 | 346915.59 | 779768.5 | 479084.97 | 460267.31 | 543145 | 641730.69 | 391480.19 | 2 | 7.5 | 8.5 | 3 | 3 | 7.5 | 8.5 | 2.019722789 | 7.334642557 | 1.814371244 | 6.588906465 |
| Q62086 | Pon2 | Serum paraoxonase/arylesterase 2 | 77.35 | 4.52 | 39617 | 2 | 712906.88 | 2 | 734898.06 | 508484.38 | 591521.44 | 532611.12 | 645105.56 | 665099.06 | 750807.62 | 926493.94 | 3027654.25 | 2254242.5 | 3370617.25 | 3372546.5 | 3097976.75 | 845998.94 | 111451.83 | 791788.44 | 2 | 2 | 0 | 12 | 0 | 8 | 0 | 12 | 5.935694132 | 4.296174235 | 0.686566626 | 0.496927534 |
| P37040 | Por | NADPH--cytochrome P450 reductase | 912.4 | 32.6 | 77044 | 17 | 9732950 | 13376572 | 8151045.5 | 8627471 | 9473038 | 11539862 | 10462630 | 12195338 | 10641674 | 12806978 | 43183954 | 38509494 | 40126274 | 29400900 | 36408986 | 11534504 | 12671956 | 17554934 | 16991208 | 17745150 | 0 | 2 | 0 | 4 | 0 | 4 | 3.801165236 | 3.254831891 | 1.549758584 | 1.327015099 |
| P25425 | Pou2f1 | POU domain, class 2, transcription factor 1 | 171.44 | 4.68 | 79547 | 3 | 1145465.75 | 1638671.75 | 2441183 | 3935758.75 | 2431494 | 2214825.75 | 1242814.25 | 3298106.5 | 2313416 | 1627814.12 | 922307.38 | 497965.03 | 551714.69 | 437041.09 | 458682.25 | 1866592.62 | 872508.12 | 2491786.5 | 1403506.75 | 1860725.25 | 0 | 9 | 0 | 9 | 0 | 9 | 0.247374796 | 0.268086072 | 0.732807036 | 0.794160775 |
| Q9D819 | Ppa1 | Inorganic pyrophosphatase | 572.01 | 34.26 | 32667 | 8 | 19859294 | 21408738 | 14730285 | 16873374 | 23661910 | 16446080 | 20247070 | 18240042 | 20675414 | 14896984 | 8390907 | 15457038 | 9025947 | 6734176.5 | 9072174 | 21951644 | 13143150 | 13726807 | 14114551 | 16397048 | 1 | 5 | 1 | 6 | 1 | 6 | 0.504282882 | 0.537870009 | 0.821819544 | 0.876555802 |
| Q91VM9 | Ppa2 | Inorganic pyrophosphatase 2, mitochondrial | 215.72 | 16.97 | 38115 | 4 | 2582159 | 2153097.25 | 3007511.5 | 3259447.5 | 2874577.75 | 1543056.25 | 2299858.5 | 2828092.5 | 2810034.5 | 2849147.5 | 1433408.12 | 1560441.62 | 1880457.38 | 2003084.5 | 2234763 | 3155189.75 | 2452079.5 | 3191201 | 3059573.75 | 3203869.5 | 1 | 8 | 4 | 3 | 4 | 8 | 0.656647009 | 0.739011741 | 1.085403054 | 1.221547634 |
| Q91YU8 | Ppan | Suppressor of SWI4 1 homolog | 453.3 | 17.23 | 52756 | 8 | 341523.56 | 1759523.25 | 2095264 | 2150489 | 560643.12 | 1759530 | 1951098.5 | 1071427.75 | 2176006.75 | 1417728.62 | 5327990.5 | 4297904.5 | 4366677 | 4073607.5 | 3498687 | 1282525 | 2440819.5 | 1888576.62 | 1762428 | 2527584.5 | 0 | 7 | 0 | 8 | 0 | 8 | 3.121975341 | 2.574666071 | 1.43351653 | 1.182208688 |
| P35433 | Ppat | Amidophosphoribosyltransferase | 44.22 | 1.35 | 57437 | 1 | 2 | 1858454.25 | 1726975.5 | 2424481.5 | 1305613 | 2 | 1752391.62 | 1573035 | 1561889.62 | 1816592.12 | 1481712.25 | 1768407 | 1514191.88 | 1500761.75 | 1937698 | 2650655.75 | 2784098.5 | 2121447.75 | 2428061 | 2280356.25 | 12 | 2 | 12 | 0 | 12 | 2 | 1.121282407 | 1.223580036 | 1.676519068 | 1.829472441 |
| O35711 | Ppfibp2 | Liprin-beta-2 | 403.96 | 10.66 | 98753 | 6 | 1461666.88 | 318578.5 | 561350.12 | 1233049.5 | 1011400.44 | 1207446.62 | 948648.12 | 1742436 | 2065936.5 | 946039.25 | 1880681 | 3156266 | 1253677 | 1281304.25 | 1052216.62 | 2025599.25 | 2067609.25 | 1812770.62 | 1050735.62 | 1645382.5 | 4 | 2 | 8 | 7 | 8 | 7 | 1.880518844 | 1.247975801 | 1.875711297 | 1.24478535 |
| P17742 | Ppia | Peptidyl-prolyl cis-trans isomerase A | 1005.57 | 79.27 | 17971 | 15 | 60414122 | 82944626 | 85645386 | 75539042 | 80923210 | 66253962 | 72046066 | 111711378 | 96904882 | 88969818 | 96424138 | 115568890 | 95211194 | 83686434 | 105514514 | 54515922 | 32693102 | 91447386 | 83766642 | 96970218 | 1 | 11 | 8 | 9 | 8 | 11 | 1.287804042 | 1.138841461 | 0.93235956 | 0.824511874 |
| P24369 | Ppib | Peptidyl-prolyl cis-trans isomerase B | 799.67 | 55.09 | 23713 | 13 | 28149052 | 29411892 | 31258822 | 30730284 | 37851538 | 25219246 | 26443502 | 30111170 | 34181350 | 28220744 | 22658036 | 27981586 | 29597520 | 25948638 | 27200688 | 23868524 | 24335698 | 28109446 | 27080592 | 21983754 | 2 | 0 | 8 | 4 | 8 | 4 | 0.847427715 | 0.925164083 | 0.796548596 | 0.869617714 |
| Q9CR16 | Ppid | Peptidyl-prolyl cis-trans isomerase D | 594.93 | 29.73 | 40743 | 11 | 15131596 | 16743578 | 23337780 | 22240990 | 27823846 | 11924011 | 13117473 | 20427660 | 26051258 | 19870618 | 13933746 | 18928814 | 17492670 | 14680901 | 17438682 | 18206866 | 13113141 | 20246156 | 20571966 | 22427174 | 6 | 9 | 10 | 11 | 10 | 11 | 0.783401827 | 0.902438916 | 0.898245518 | 1.034732986 |
| A2AR02 | Ppig | Peptidyl-prolyl cis-trans isomerase G | 39.03 | 1.6 | 88325 | 1 | 336760.19 | 476171.5 | 102410.55 | 2 | 315526.75 | 465094.59 | 355928.41 | 393838.59 | 391220.84 | 319597.91 | 674373.75 | 576294.69 | 540002.69 | 288214.12 | 527491.56 | 2 | 593619.75 | 2 | 2 | 207379.86 | 3 | 8.5 | 5 | 5 | 5 | 8.5 | 2.117506084 | 1.353483626 | 0.650763253 | 0.4159598 |
| Q9D868 | Ppih | Peptidyl-prolyl cis-trans isomerase H | 166.29 | 24.47 | 20464 | 3 | 2 | 600860.81 | 536468.56 | 325508.56 | 362739.38 | 2 | 419119.88 | 2 | 306852.75 | 331095.62 | 542289.94 | 822027.94 | 807622.38 | 708777.56 | 521427.62 | 455297.78 | 309527.88 | 540649.5 | 502837.78 | 522769.38 | 3 | 11 | 0 | 2 | 3 | 11 | 1.86359772 | 3.218460649 | 1.276900054 | 2.205225158 |
| Q812D3 | Ppil3 | Peptidyl-prolyl cis-trans isomerase-like 3 | 49.68 | 11.8 | 18113 | 1 | 578561.69 | 72705.58 | 484649.5 | 155809.81 | 407429.03 | 2 | 2 | 2 | 434102.72 | 95141.75 | 109789.09 | 330552.25 | 2 | 125859.97 | 2 | 798560 | 2 | 2 | 2 | 2 | 4 | 5 | 10 | 11 | 10 | 11 | 0.333227461 | 1.06982486 | 0.469979321 | 1.508865925 |
| Q6ZQB6 | Ppip5k2 | Inositol hexakisphosphate and diphosphoinositol-pentakisphosphate kinase 2 | 57.88 | 1.33 | 128427 | 1 | 403675.62 | 477484.31 | 560667.38 | 516269.31 | 832887.75 | 2 | 305248.06 | 679657.44 | 2 | 320772.84 | 2 | 2 | 641238.19 | 2 | 197909.02 | 466877.84 | 595237.62 | 642184.06 | 413754.5 | 855412.56 | 4 | 10 | 9 | 4 | 9 | 10 | 0.300665679 | 0.642693237 | 1.065382742 | 2.277327715 |
| P36993 | Ppm1b | Protein phosphatase 1B | 124.79 | 7.69 | 42795 | 2 | 1956610.88 | 1951816.75 | 1879053.25 | 2169385.25 | 1780466.88 | 508927.22 | 1905909 | 2147769.75 | 1950343.12 | 2055357 | 1278307.5 | 970128.56 | 355513.81 | 1540401.88 | 1286992.38 | 1764125.5 | 1769740.62 | 2097859.5 | 2615445 | 1858417.88 | 0 | 10 | 4 | 12 | 4 | 12 | 0.557785599 | 0.633887734 | 1.037818927 | 1.179414973 |
| Q61074 | Ppm1g | Protein phosphatase 1G | 537.85 | 19.37 | 58728 | 10 | 6832508.5 | 10310494 | 9218889 | 7125540 | 10079645 | 10064858 | 7845178 | 8283570.5 | 7996332 | 7329203.5 | 9125973 | 5997953.5 | 9175492 | 13776054 | 4459126 | 8938914 | 7861962 | 6244655 | 5335740 | 9256158 | 9 | 7 | 12 | 10 | 12 | 10 | 0.976301417 | 1.02445755 | 0.863896135 | 0.906507871 |
| Q4FZT2 | Ppme1 | Protein phosphatase methylesterase 1 | 229.32 | 11.66 | 42316 | 4 | 1980937.5 | 2491648.25 | 2819859 | 2268020.5 | 3223547 | 1614171.25 | 2320737 | 3746679.75 | 2655092.75 | 3125136 | 759547.5 | 1875478.25 | 1955847.75 | 2176065 | 2288043.5 | 2437209.5 | 1576131.25 | 2255197.25 | 2468899 | 2947643 | 3 | 9 | 4 | 8 | 4 | 9 | 0.708305172 | 0.6726419 | 0.914038548 | 0.86801657 |
| P51175 | Ppox | Protoporphyrinogen oxidase | 89.06 | 3.98 | 50871 | 2 | 2043131.25 | 1888937.5 | 2063313.88 | 1946111.62 | 1943963.38 | 1577596.75 | 2130830.75 | 1241662.5 | 1896260.5 | 1851780.88 | 940597.56 | 1573155.38 | 1361577.12 | 1501942.38 | 1208410 | 1302271.12 | 1630511.88 | 2247110.5 | 1771737.12 | 2156650.25 | 0 | 10 | 3 | 10 | 3 | 10 | 0.666199046 | 0.757137614 | 0.921381813 | 1.047153747 |
| P62137 | Ppp1ca | Serine/threonine-protein phosphatase PP1-alpha catalytic subunit | 1158.55 | 53.33 | 37540 | 18 | 741704.44 | 1332430.75 | 848783.69 | 1239544.25 | 813496.5 | 1087988.62 | 803158.75 | 856565.69 | 921053.19 | 941258.75 | 2273509.75 | 1924399.12 | 2088640.12 | 34488.61 | 2116970.5 | 758802.44 | 808151.81 | 1053581.62 | 721681.94 | 689296 | 5 | 5 | 5 | 5 | 5 | 5 | 1.695754935 | 1.830360595 | 0.810198255 | 0.874510184 |
| P62141 | Ppp1cb | Serine/threonine-protein phosphatase PP1-beta catalytic subunit | 986.65 | 40.67 | 37187 | 14 | 4340321 | 4548061.5 | 4967435.5 | 5913662.5 | 4833284 | 5494049.5 | 4387050.5 | 3642833.75 | 4569457 | 5020518 | 3915892 | 3429915.25 | 4348444.5 | 3390948.25 | 4252891 | 5736224.5 | 6953079 | 5735692.5 | 7088994.5 | 4974467 | 1 | 3 | 3 | 2 | 3 | 3 | 0.78601293 | 0.836643045 | 1.239228929 | 1.31905243 |
| P63087 | Ppp1cc | Serine/threonine-protein phosphatase PP1-gamma catalytic subunit | 1068.97 | 50.46 | 36984 | 16 | 5326580 | 4732472.5 | 5007804.5 | 5784666.5 | 4578759 | 4957448.5 | 4294018 | 4476670 | 4697473 | 4480961.5 | 4004312.25 | 3487311.75 | 3894091.5 | 3707992.5 | 3830465.5 | 4470837.5 | 5756461.5 | 4873448.5 | 5241667.5 | 5279371 | 0 | 12 | 0 | 5 | 0 | 12 | 0.744158996 | 0.826146065 | 1.00753053 | 1.118534328 |
| O55000 | Ppp1r10 | Serine/threonine-protein phosphatase 1 regulatory subunit 10 | 58.16 | 2.06 | 92828 | 1 | 414771.41 | 2 | 191117.48 | 278994.78 | 2 | 317301.75 | 339947 | 2 | 2 | 2 | 2 | 2 | 2 | 2 | 2 | 277300.97 | 464862.44 | 333845.16 | 2 | 2 | 5 | 11 | 7.5 | 10 | 7.5 | 11 | 1.130087E-05 | 1.52148E-05 | 1.215987754 | 1.637131675 |
| Q9DBR7 | Ppp1r12a | Protein phosphatase 1 regulatory subunit 12A | 229.22 | 4.08 | 114996 | 4 | 3584634.25 | 3873217.25 | 2841830 | 4624534 | 3462829.5 | 3490026.5 | 3593079 | 3834324 | 4277257 | 3581768.5 | 2121407.5 | 2687858.5 | 2863510.25 | 3191500.25 | 2883431 | 3817744 | 3938627 | 4350850 | 5360328 | 4581767 | 3 | 5 | 0 | 3 | 3 | 5 | 0.747684443 | 0.732178012 | 1.199176703 | 1.174306652 |
| Q3UMT1 | Ppp1r12c | Protein phosphatase 1 regulatory subunit 12C | 64.94 | 1.66 | 84685 | 1 | 446182.47 | 574410.12 | 576717.94 | 643908.38 | 592722.19 | 2 | 495331.84 | 559140.62 | 535713.12 | 531522.31 | 279569.25 | 181910.62 | 271368.28 | 495293.81 | 2 | 572833.44 | 818142.12 | 636004 | 845027.06 | 623059.25 | 1 | 6 | 4.5 | 0 | 4.5 | 6 | 0.433369614 | 0.578846319 | 1.233288112 | 1.647287354 |
| Q62084 | Ppp1r14b | Protein phosphatase 1 regulatory subunit 14B | 49.38 | 9.52 | 15957 | 1 | 363866.16 | 363441.72 | 252024.84 | 375220.84 | 224198.52 | 292519.59 | 413348.44 | 272527.94 | 2 | 213915.3 | 2 | 158247.72 | 2 | 120174.29 | 2 | 686074.81 | 513999.94 | 331000.38 | 2 | 397384.25 | 0 | 8 | 3.5 | 6.5 | 3.5 | 8 | 0.176359552 | 0.233519174 | 1.221509954 | 1.617411656 |
| Q8BQ30 | Ppp1r18 | Phostensin | 218.71 | 7.74 | 65630 | 4 | 6649142 | 7633970 | 4705077 | 6798220.5 | 5589067 | 7198436.5 | 7195103 | 5437298 | 6068147.5 | 5055375 | 5664272 | 6314678 | 5829582.5 | 5327209.5 | 5526519.5 | 5288731 | 6560639 | 4502467 | 6351841 | 5105201.5 | 8 | 6 | 10 | 8 | 10 | 8 | 0.913524341 | 0.925952321 | 0.886325328 | 0.898383281 |
| Q3TDD9 | Ppp1r21 | Protein phosphatase 1 regulatory subunit 21 | 38.38 | 2.44 | 88338 | 1 | 332353.56 | 410977.62 | 411663.75 | 2 | 362356.84 | 264588.03 | 277054.69 | 376216.56 | 496499.81 | 285051.91 | 173175.36 | 304574.25 | 2 | 2 | 2 | 352462.34 | 2 | 298134.12 | 2 | 340419.91 | 3.5 | 6 | 3 | 9 | 3.5 | 9 | 0.314861056 | 0.281130115 | 0.653124136 | 0.583155205 |
| Q3UM45 | Ppp1r7 | Protein phosphatase 1 regulatory subunit 7 | 572.52 | 32.69 | 41292 | 10 | 5339011.5 | 5528157.5 | 7671765.5 | 7453816 | 7205143 | 4339640.5 | 6305383.5 | 6055248 | 6328605 | 7153590.5 | 6531187.5 | 4942763 | 5723880.5 | 5635498.5 | 4781093.5 | 6594397 | 5999378.5 | 6843235 | 5344800 | 6239897 | 6 | 9 | 8 | 12 | 8 | 12 | 0.831812506 | 0.914916019 | 0.93444807 | 1.027805546 |
| P63330 | Ppp2ca | Serine/threonine-protein phosphatase 2A catalytic subunit alpha isoform | 637.91 | 38.51 | 35608 | 11 | 23138350 | 25381098 | 24838048 | 24533148 | 21146674 | 17411428 | 24356320 | 20208754 | 22535868 | 22095528 | 12216564 | 21076332 | 15946948 | 11688910 | 18275374 | 25415486 | 20144654 | 21381590 | 21718822 | 24369830 | 0 | 9 | 3 | 10 | 3 | 10 | 0.665372249 | 0.742948032 | 0.949537371 | 1.060243979 |
| Q76MZ3 | Ppp2r1a | Serine/threonine-protein phosphatase 2A 65 kDa regulatory subunit A alpha isoform | 1517.76 | 47.2 | 65323 | 23 | 42804154 | 43828714 | 41020398 | 40088370 | 55477306 | 32251726 | 41078418 | 44277814 | 50947702 | 37609866 | 28830176 | 43283094 | 31712730 | 26994012 | 35978498 | 46030698 | 38412690 | 37431990 | 36017302 | 40395550 | 3 | 5 | 4 | 10 | 4 | 10 | 0.747241737 | 0.809051412 | 0.88831274 | 0.961791401 |
| Q4QQT4 | Ppp2r1b | Serine/threonine-protein phosphatase 2A 65 kDa regulatory subunit A beta isoform | 667.67 | 16.47 | 66006 | 8 | 557970.25 | 622150.69 | 613742.75 | 676283.12 | 456541.81 | 500322.38 | 511853.06 | 200453.05 | 407766.59 | 312497.34 | 681239.31 | 417083.94 | 654562.38 | 441284.75 | 665526.44 | 668464.5 | 2 | 618728.06 | 399857.06 | 615165.38 | 12 | 10 | 4 | 8 | 12 | 10 | 0.977110035 | 1.479490938 | 0.78662861 | 1.191073531 |
| P36876 | Ppp2r2a | Serine/threonine-protein phosphatase 2A 55 kDa regulatory subunit B alpha isoform | 321.04 | 13.87 | 51678 | 6 | 5397344.5 | 4939425 | 4263179.5 | 4056803.75 | 8048544.5 | 2970792 | 3225681 | 6656432 | 8273366 | 5486381 | 8025889.5 | 8759724 | 5674030 | 6571477 | 6935064 | 4969190 | 3254291.75 | 6357979.5 | 3135160 | 5369935.5 | 4 | 10 | 6 | 10 | 6 | 10 | 1.346780909 | 1.351469388 | 0.864493532 | 0.867503049 |
| Q6PD03 | Ppp2r5a | Serine/threonine-protein phosphatase 2A 56 kDa regulatory subunit alpha isoform | 101.07 | 8.44 | 56347 | 2 | 410762.59 | 240170.7 | 234407.61 | 291654.38 | 190610.19 | 592781.88 | 405702.41 | 406715.38 | 258070.44 | 226384.14 | 239769.47 | 206327.61 | 301216.28 | 486190.38 | 155837.89 | 266659.34 | 923826.75 | 621807.69 | 720907.5 | 220288.75 | 12 | 6 | 7 | 8 | 12 | 8 | 1.01589359 | 0.735235893 | 2.013365763 | 1.457139596 |
| Q61151 | Ppp2r5e | Serine/threonine-protein phosphatase 2A 56 kDa regulatory subunit epsilon isoform | 84.89 | 6.21 | 54713 | 2 | 581839.62 | 797561.44 | 818752.25 | 944509.75 | 929895.5 | 1139035.75 | 680427 | 997967.38 | 1044034.38 | 726762.12 | 332185.12 | 574856.12 | 588169.06 | 109137.74 | 415826.06 | 622146.94 | 850091.12 | 820175.19 | 947812.19 | 906005.12 | 1 | 10 | 0 | 8 | 1 | 10 | 0.496045439 | 0.440295187 | 1.018089857 | 0.903667341 |
| P63328 | Ppp3ca | Serine/threonine-protein phosphatase 2B catalytic subunit alpha isoform | 494.34 | 19.58 | 58644 | 9 | 10145184 | 10125261 | 14584545 | 12828923 | 15963079 | 9182245 | 9899852 | 13537429 | 12571545 | 12558725 | 4451651 | 5725251.5 | 5823734.5 | 6028640.5 | 5843319 | 11747729 | 8290227.5 | 10599961 | 10274011 | 13018799 | 0 | 9 | 0 | 10 | 0 | 10 | 0.437924804 | 0.482644069 | 0.84734134 | 0.93386871 |
| P97470 | Ppp4c | Serine/threonine-protein phosphatase 4 catalytic subunit | 220.62 | 14.66 | 35080 | 4 | 2 | 2 | 554230.5 | 372331.28 | 420811.62 | 450032.5 | 404844.09 | 321022.62 | 652601.75 | 382095.28 | 2 | 2 | 208714.84 | 2 | 2 | 2 | 327423.62 | 344902.09 | 413072.66 | 2 | 6 | 9 | 0 | 5 | 6 | 9 | 0.15491045 | 0.09441925 | 0.805566703 | 0.490999826 |
| Q8K2V1 | Ppp4r1 | Serine/threonine-protein phosphatase 4 regulatory subunit 1 | 115.15 | 2 | 106298 | 2 | 1535717.25 | 1636101.75 | 1569247.25 | 1429561.88 | 1083929.88 | 1795421.62 | 1586270.5 | 909595.31 | 1343942.75 | 1307976.25 | 691919.31 | 1538466 | 1223546.38 | 810967.06 | 1345031.88 | 1426490 | 1250302.12 | 1440622.75 | 1415199 | 1307113.88 | 5 | 6 | 7 | 11 | 7 | 11 | 0.77329737 | 0.807974052 | 0.942817983 | 0.985096413 |
| Q0VGB7 | Ppp4r2 | Serine/threonine-protein phosphatase 4 regulatory subunit 2 | 49.85 | 3.36 | 46479 | 1 | 483250.91 | 665609.81 | 683585.88 | 439391.28 | 541398.94 | 369798.75 | 359981.81 | 342479.91 | 646307.94 | 499066.53 | 2 | 412915.75 | 2 | 2 | 262294.09 | 662916.5 | 2 | 382037.91 | 318734.66 | 503627.66 | 0 | 5 | 3 | 12 | 3 | 12 | 0.240013864 | 0.30447565 | 0.663761656 | 0.842031615 |
| Q6P2K6 | Ppp4r3a | Serine/threonine-protein phosphatase 4 regulatory subunit 3A | 77.93 | 2.07 | 93842 | 1 | 2 | 104664.87 | 2 | 2 | 345565.47 | 2 | 145354.77 | 101992.4 | 397059.53 | 2 | 2 | 2 | 2 | 2 | 2 | 125729.03 | 2 | 2 | 2 | 2 | 7.5 | 10 | 5 | 7 | 7.5 | 10 | 2.221056E-05 | 1.551805E-05 | 0.27926895 | 0.195119401 |
| Q60676 | Ppp5c | Serine/threonine-protein phosphatase 5 | 100.28 | 5.21 | 56877 | 2 | 4514620.5 | 732276.19 | 3673207.5 | 605468.44 | 5431588 | 3237836.5 | 4446231 | 6493878 | 5003645 | 2 | 1687259.75 | 2941549 | 396954.44 | 963481.19 | 511116.81 | 5294989.5 | 1458619.75 | 2963278.25 | 2943605 | 611454.19 | 6 | 11 | 5 | 8 | 6 | 11 | 0.434598608 | 0.338885376 | 0.887330625 | 0.691910575 |
| Q64620 | Ppp6c | Serine/threonine-protein phosphatase 6 catalytic subunit | 266.88 | 18.36 | 35159 | 4 | 3603943.5 | 2962975 | 2956527.5 | 2935550.5 | 2792483 | 2913884 | 3057016 | 3150995 | 2587766 | 3193395.5 | 2420825 | 2471460 | 3026381.75 | 2713718.5 | 2621797.25 | 2289637.5 | 3692026 | 3838656 | 3333948 | 3917251.25 | 4 | 6 | 4 | 5 | 4 | 6 | 0.86904241 | 0.889360011 | 1.119335259 | 1.145504531 |
| Q7TSI3 | Ppp6r1 | Serine/threonine-protein phosphatase 6 regulatory subunit 1 | 43.76 | 1.29 | 94527 | 1 | 1446421.38 | 1622224.88 | 1234334.62 | 1847331.25 | 1625511.5 | 1591780.62 | 1785651.62 | 1449110.62 | 1269267.25 | 1498151.5 | 739432.94 | 735131.44 | 811089.5 | 955392.75 | 832180.88 | 1706759 | 2081608.12 | 1625487.88 | 1573111.75 | 1881065.5 | 0 | 6 | 0 | 4 | 0 | 6 | 0.523832291 | 0.536377153 | 1.140462113 | 1.167774174 |
| O88531 | Ppt1 | Palmitoyl-protein thioesterase 1 | 342.59 | 23.53 | 34490 | 6 | 8257738 | 6877490.5 | 8660177 | 15343746 | 14606335 | 8156881 | 7772314 | 9315938 | 10480905 | 10223054 | 3202669.75 | 1628197 | 1424178.75 | 1311020.75 | 1351921.25 | 9633528 | 12755555 | 9233552 | 10548312 | 9008489 | 0 | 10 | 0 | 8 | 0 | 10 | 0.16592998 | 0.194084085 | 0.952255516 | 1.113829105 |
| Q8C6U2 | Pqlc3 | PQ-loop repeat-containing protein 3 | 84.82 | 9.9 | 22739 | 2 | 105594.67 | 2 | 52605.56 | 1010065.69 | 645871 | 2 | 1198348.12 | 373870.06 | 1156774.75 | 1001751.88 | 252773.06 | 1240675.25 | 433173.72 | 803945 | 1329177.88 | 1034985.12 | 86797.18 | 80156.59 | 1208973.12 | 728215.62 | 5 | 7 | 10 | 12 | 10 | 12 | 2.237835739 | 1.088185588 | 1.730367832 | 0.84142071 |
| Q9JIG8 | Praf2 | PRA1 family protein 2 | 45.9 | 6.18 | 19478 | 1 | 2 | 2 | 179009.64 | 2 | 2 | 472012.31 | 2 | 2 | 414179.53 | 2 | 523551.38 | 562800.44 | 895616.12 | 587722.25 | 2 | 2 | 496151.81 | 2 | 2 | 461277.16 | 3 | 9 | 3.5 | 11.5 | 3.5 | 11.5 | 14.3544077 | 2.899682299 | 5.348271656 | 1.080385132 |
| P35700 | Prdx1 | Peroxiredoxin-1 | 1336.8 | 81.91 | 22176 | 22 | 82072738 | 74367090 | 95829290 | 78433474 | 94566354 | 73929258 | 91716154 | 104656386 | 101995874 | 99042450 | 79154522 | 96238754 | 89230146 | 69741634 | 88543362 | 102240738 | 82392306 | 119665258 | 105817586 | 110481466 | 12 | 2 | 5 | 5 | 12 | 5 | 0.994449329 | 0.897246804 | 1.224160285 | 1.104504645 |
| Q61171 | Prdx2 | Peroxiredoxin-2 | 705.1 | 45.45 | 21779 | 10 | 173070546 | 149379778 | 137551250 | 149690610 | 127327018 | 178182018 | 132331386 | 120464218 | 120588674 | 141047858 | 87309658 | 78720674 | 78481938 | 77933194 | 69200850 | 159986754 | 193406914 | 147484274 | 145347842 | 133668658 | 0 | 11 | 0 | 5 | 0 | 11 | 0.53139228 | 0.565461032 | 1.058173844 | 1.126015744 |
| P20108 | Prdx3 | Thioredoxin-dependent peroxide reductase, mitochondrial | 375.49 | 22.96 | 28127 | 7 | 22190338 | 21758742 | 23528866 | 22686238 | 29910876 | 21879406 | 24545814 | 24741882 | 25290274 | 24410962 | 17283366 | 23104258 | 23865946 | 24033058 | 28028534 | 29341426 | 26308136 | 24658146 | 28309670 | 28773042 | 10 | 5 | 8 | 2 | 10 | 5 | 0.968687103 | 0.962329456 | 1.144204467 | 1.136694872 |
| O08807 | Prdx4 | Peroxiredoxin-4 | 761.68 | 50.73 | 31053 | 14 | 17809650 | 27130034 | 32972084 | 32929092 | 32432794 | 24642612 | 26978198 | 36414346 | 31016678 | 32159366 | 20179018 | 31496546 | 41374938 | 24846004 | 41271122 | 26933916 | 23864756 | 31558434 | 31172170 | 31435322 | 11 | 8 | 11 | 10 | 11 | 10 | 1.11093438 | 1.052617981 | 1.011802198 | 0.958689555 |
| P99029 | Prdx5 | Peroxiredoxin-5, mitochondrial | 890.82 | 57.62 | 21897 | 14 | 119736026 | 122183962 | 61644398 | 74211554 | 68228282 | 106144890 | 100462514 | 69477618 | 73488418 | 76216530 | 86558082 | 106636530 | 80979858 | 67003698 | 83355490 | 132448778 | 113995778 | 101893018 | 97160098 | 100718482 | 12 | 8 | 11 | 4 | 12 | 8 | 0.951860178 | 0.997049456 | 1.224688304 | 1.282830016 |
| O08709 | Prdx6 | Peroxiredoxin-6 | 1072.95 | 81.25 | 24871 | 18 | 38099198 | 43332390 | 49689234 | 37493570 | 51388870 | 36171226 | 46098766 | 54894234 | 46825866 | 50382958 | 45106722 | 54125858 | 40435090 | 33686710 | 41198866 | 49009922 | 43690746 | 54765430 | 53459078 | 50535498 | 12 | 5 | 7 | 9 | 12 | 9 | 0.975227567 | 0.915434799 | 1.142986116 | 1.072907802 |
| Q9WUQ2 | Preb | Prolactin regulatory element-binding protein | 229.56 | 12.23 | 45437 | 4 | 1590216.5 | 1210432.5 | 1278671.25 | 1484600.75 | 843390.62 | 3305622.5 | 1227843.5 | 1098627.5 | 1471673.62 | 1823473.38 | 1936050.88 | 2607689.25 | 3466481.75 | 2988726 | 2000235.5 | 1599594.5 | 1727058.25 | 1688256.75 | 1464865.75 | 1616380.75 | 0 | 2 | 4 | 11 | 4 | 11 | 2.028804614 | 1.456125594 | 1.263580809 | 0.906904659 |
| Q9QUR6 | Prep | Prolyl endopeptidase | 840.97 | 22.25 | 80752 | 13 | 16068877 | 17057010 | 14600828 | 15037026 | 13581396 | 17116758 | 13890605 | 13635054 | 11521336 | 15977440 | 16397764 | 16790694 | 17143728 | 15594117 | 16657670 | 18102828 | 18334078 | 18567564 | 15839656 | 14526788 | 5 | 6 | 5 | 4 | 5 | 6 | 1.08171884 | 1.144754745 | 1.118223339 | 1.183386502 |
| Q69ZK0 | Prex1 | Phosphatidylinositol 3,4,5-trisphosphate-dependent Rac exchanger 1 protein | 1024.21 | 13.33 | 184935 | 19 | 12587764 | 12240188 | 18171052 | 19168848 | 22936888 | 13095770 | 11099807 | 16270355 | 20796302 | 17229232 | 7371416.5 | 7588642.5 | 7955791.5 | 6147153.5 | 7983837 | 11453774 | 10371119 | 13508807 | 15818183 | 16322212 | 0 | 6 | 0 | 8 | 0 | 8 | 0.435308785 | 0.471985591 | 0.792835922 | 0.859636065 |
| O89044 | Prim2 | DNA primase large subunit | 143.89 | 4.93 | 58603 | 2 | 551853.31 | 970609.88 | 809258.44 | 476590.53 | 1075688.5 | 557341.12 | 669469.12 | 592665.12 | 1209464.25 | 368183.78 | 270478.09 | 842869 | 2 | 143357.5 | 263835.72 | 557292.19 | 2 | 387330.41 | 67135.01 | 342822.53 | 3 | 2 | 4 | 2 | 4 | 2 | 0.391488685 | 0.447597021 | 0.348759503 | 0.398743874 |
| P54645 | Prkaa1 | 5'-AMP-activated protein kinase catalytic subunit alpha-1 | 42.03 | 1.25 | 63973 | 1 | 589041 | 146182.06 | 547439.88 | 375942.22 | 2 | 2 | 2 | 2 | 2 | 548415.81 | 1069189.88 | 917240.62 | 750215.12 | 539105.81 | 741260.44 | 2 | 2 | 2 | 2 | 426210.28 | 2 | 5 | 1 | 12 | 2 | 12 | 2.421918804 | 7.324648924 | 0.256973616 | 0.77716954 |
| P05132 | Prkaca | cAMP-dependent protein kinase catalytic subunit alpha | 225.89 | 15.38 | 40571 | 4 | 2858580.25 | 3108406 | 3374145.25 | 2908140.25 | 4036980 | 2035747.5 | 3072890 | 2762375.25 | 5191796.5 | 2621666.5 | 1397399 | 1933271.5 | 2109965.25 | 1455798.5 | 1640237.12 | 2617849 | 1684755.62 | 2940111 | 2927366.5 | 2550385.5 | 0 | 4 | 1 | 8 | 1 | 8 | 0.524164277 | 0.544275212 | 0.781055569 | 0.811022812 |
| P68181 | Prkacb | cAMP-dependent protein kinase catalytic subunit beta | 136.21 | 8.55 | 40708 | 3 | 402108.97 | 546788.88 | 636712 | 855679.88 | 625964.19 | 513231.5 | 332859.84 | 660096 | 1311903.5 | 881299.19 | 2073610.62 | 1525540.88 | 2234799.75 | 2453665.5 | 2143883 | 2 | 324822.72 | 1078763.38 | 780548.12 | 561861.69 | 0 | 11 | 0 | 9 | 0 | 11 | 3.400924743 | 2.819789118 | 0.895262662 | 0.742283968 |
| O54950 | Prkag1 | 5'-AMP-activated protein kinase subunit gamma-1 | 38.39 | 3.33 | 37520 | 1 | 2 | 308510.56 | 360454.88 | 310627.31 | 562511.38 | 331210.06 | 430219.91 | 412701.34 | 2 | 235969.42 | 58234.48 | 242199.83 | 562560.06 | 449060.47 | 300693.34 | 2 | 244793.66 | 217574.72 | 115833.73 | 253343.3 | 12 | 4.5 | 10 | 6.5 | 12 | 6.5 | 1.045808812 | 1.143709707 | 0.539228393 | 0.589706971 |
| Q9DBC7 | Prkar1a | cAMP-dependent protein kinase type I-alpha regulatory subunit | 477.98 | 25.46 | 43185 | 8 | 2710606.25 | 2956105.5 | 5157968.5 | 3313074.25 | 4516981.5 | 1407185.25 | 2769253 | 4050832.5 | 3698436.5 | 2919209 | 5640021 | 8269927 | 7735366.5 | 5483762 | 8858641 | 4535994 | 2990896 | 4158960.25 | 5566362 | 4679008 | 0 | 7 | 0 | 2 | 0 | 7 | 1.929146438 | 2.424245236 | 1.1756382 | 1.477355607 |
| P12367 | Prkar2a | cAMP-dependent protein kinase type II-alpha regulatory subunit | 100.01 | 6.73 | 45389 | 2 | 390108.91 | 546053.44 | 447749.03 | 222442.55 | 332044.34 | 936674.75 | 310210.28 | 388834.69 | 362905.09 | 307938.47 | 1158343 | 999147.12 | 636438.38 | 640155.06 | 1093134.12 | 313327.09 | 878818.44 | 385468.62 | 280626.38 | 464386.09 | 0 | 12 | 2 | 12 | 2 | 12 | 2.335545667 | 1.962754597 | 1.198219507 | 1.006964188 |
| P68403 | Prkcb | Protein kinase C beta type | 181.48 | 4.92 | 76751 | 3 | 644298.44 | 684640.75 | 681193.94 | 753202 | 719319.25 | 623077.19 | 236853.73 | 753848 | 939245.44 | 634780.31 | 901138.69 | 905935 | 1114601.25 | 469217.78 | 1061116.12 | 2 | 2 | 746749.75 | 483070.03 | 418916.97 | 5 | 4 | 6 | 5 | 6 | 5 | 1.278337829 | 1.396575167 | 0.473414979 | 0.517202564 |
| P28867 | Prkcd | Protein kinase C delta type | 420.85 | 13.06 | 77547 | 8 | 6811553.5 | 7633023 | 8560850 | 11733033 | 7476933 | 7156835.5 | 7454685 | 9699483 | 10144338 | 11158954 | 8838436 | 8518388 | 10471136 | 12324897 | 10844338 | 5787746.5 | 5168747.5 | 8258316 | 7782506 | 9236412 | 5 | 10 | 8 | 6 | 8 | 10 | 1.208023708 | 1.118009046 | 0.858306079 | 0.794350271 |
| P23298 | Prkch | Protein kinase C eta type | 67.3 | 1.46 | 77919 | 1 | 179823.39 | 159095.2 | 172126.92 | 292217.19 | 278111.72 | 180224.48 | 192252.2 | 142421.42 | 230204.97 | 177740.75 | 2 | 2 | 63347.53 | 2 | 2 | 2 | 2 | 2 | 2 | 144311.05 | 0 | 0 | 0 | 1 | 0 | 1 | 0.058587968 | 0.068652494 | 0.133458909 | 0.156385129 |
| O08795 | Prkcsh | Glucosidase 2 subunit beta | 670.06 | 17.85 | 58793 | 12 | 19468390 | 19573470 | 99787650 | 15690333 | 114337546 | 19053212 | 18818344 | 20786966 | 16790726 | 21968626 | 32663222 | 30295416 | 33689918 | 32289056 | 34402234 | 24689758 | 22567300 | 20641248 | 23267554 | 22069058 | 10 | 10 | 0 | 2 | 10 | 10 | 0.607533409 | 1.67669278 | 0.421170935 | 1.162362853 |
| Q5XIS9 | Prkd2 | Serine/threonine-protein kinase D2 | 68.09 | 1.26 | 96497 | 1 | 69883.27 | 2 | 152440.36 | 2 | 181648.52 | 2 | 73100.15 | 176213.3 | 190110.25 | 2 | 2 | 29272.81 | 2 | 2 | 2 | 2 | 2 | 2 | 2 | 2 | 6 | 5 | 6 | 5 | 6 | 5 | 0.072481531 | 0.066633965 | 2.475394E-05 | 2.275687E-05 |
| Q63009 | Prmt1 | Protein arginine N-methyltransferase 1 | 674.29 | 36.83 | 40522 | 12 | 20197488 | 18622888 | 15182651 | 15349796 | 19059974 | 20149542 | 21556534 | 13359271 | 16756300 | 13873950 | 18015200 | 16415740 | 16256903 | 9861142 | 13833509 | 21110786 | 15862765 | 12616341 | 9772347 | 10075563 | 6 | 7 | 8 | 6 | 8 | 7 | 0.841309138 | 0.867985014 | 0.785381804 | 0.81028436 |
| Q8CIG8 | Prmt5 | Protein arginine N-methyltransferase 5 | 155.39 | 5.34 | 72680 | 4 | 1700641.75 | 1444847.62 | 763948.88 | 1555895.5 | 2066688 | 748573.25 | 996421.62 | 1860536 | 2425524.5 | 812367.75 | 1218772.75 | 2952052 | 2166243.5 | 2 | 2285615.75 | 1897668 | 98737.35 | 843368.88 | 646488.06 | 542314.94 | 9 | 5 | 9 | 6 | 9 | 6 | 1.144803651 | 1.259996035 | 0.53486001 | 0.588678671 |
| Q64695 | Procr | Endothelial protein C receptor | 115.52 | 12.4 | 27189 | 2 | 233846.89 | 270578.75 | 705526.38 | 116987.84 | 704017.38 | 158755.39 | 389496.19 | 1022577.75 | 949646.62 | 1000441.5 | 1579242.5 | 1841332.25 | 1306309.5 | 1487158 | 1677352.25 | 1171111.75 | 818465.31 | 1078614.62 | 1180580.38 | 1255977.12 | 0 | 0 | 0 | 3 | 0 | 3 | 3.885554233 | 2.24128927 | 2.710421013 | 1.563441705 |
| Q99KP6 | Prpf19 | Pre-mRNA-processing factor 19 | 647.5 | 27.38 | 55239 | 11 | 17296906 | 21672258 | 21221408 | 20424166 | 17696278 | 18393522 | 18875198 | 18104980 | 17934056 | 17862196 | 12324100 | 20609950 | 16227917 | 13882515 | 20635476 | 18695578 | 17526546 | 18480058 | 15552445 | 18216924 | 6 | 7 | 10 | 11 | 10 | 11 | 0.851175803 | 0.917845805 | 0.899914929 | 0.970402518 |
| Q922U1 | Prpf3 | U4/U6 small nuclear ribonucleoprotein Prp3 | 195.56 | 5.86 | 77455 | 4 | 709889.38 | 973031.62 | 1013144.5 | 984755.62 | 910943.19 | 2077303 | 1679864.5 | 1977088.5 | 1427422.38 | 2090995.62 | 2861223 | 1791072.5 | 780805.69 | 1359676.75 | 1266485.25 | 873720.88 | 1393408.25 | 1930288 | 1983419.62 | 1693927.75 | 4 | 4 | 7 | 7 | 7 | 7 | 1.755156111 | 0.871019901 | 1.714975763 | 0.851079861 |
| Q8CCF0 | Prpf31 | U4/U6 small nuclear ribonucleoprotein Prp31 | 360.13 | 16.63 | 55430 | 6 | 4910360 | 4937345 | 3945177 | 5673816 | 5759788.5 | 6234981 | 5127047 | 4774661.5 | 4449476 | 4601052.5 | 5474759 | 5429037 | 3627930.5 | 4626277.5 | 3377445.75 | 4871247 | 6709604 | 4966211.5 | 6349193 | 4050588.5 | 7 | 10 | 10 | 9 | 10 | 10 | 0.893324948 | 0.8947177 | 1.068196477 | 1.069861864 |
| Q4FK66 | Prpf38a | Pre-mRNA-splicing factor 38A | 123.39 | 7.05 | 37437 | 2 | 627606.25 | 803540.38 | 644271.75 | 470452.09 | 508239.97 | 569244.44 | 545125.69 | 777755.62 | 639800.56 | 530060.5 | 518593.28 | 424101.97 | 369074 | 580062.62 | 728454.44 | 2 | 2 | 428394 | 2 | 351444.94 | 8 | 0 | 7 | 0 | 8 | 0 | 0.857954014 | 0.855747093 | 0.255342744 | 0.254685924 |
| Q6AXY7 | Prpf38b | Pre-mRNA-splicing factor 38B | 189.5 | 4.98 | 63932 | 3 | 280456.97 | 92532.07 | 490331.38 | 331498.59 | 456297.06 | 259717.08 | 491123.41 | 477784.38 | 460281.47 | 368827.69 | 329019.19 | 636051.31 | 320859.16 | 48105.83 | 287928.5 | 72597.09 | 111122.44 | 214468.52 | 323045.06 | 243352.02 | 11 | 5 | 8 | 1 | 11 | 5 | 0.982344015 | 0.7882282 | 0.584201891 | 0.468760839 |
| Q9DAW6 | Prpf4 | U4/U6 small nuclear ribonucleoprotein Prp4 | 110.77 | 5.57 | 58370 | 2 | 2 | 457556.84 | 2 | 685353.94 | 720503.12 | 1229324.25 | 937317.25 | 2 | 2 | 378816.84 | 1185854.25 | 878711.25 | 1708596.5 | 1704726.38 | 1628989.38 | 242615.11 | 417191.41 | 340568.97 | 329947.28 | 373932.03 | 0 | 10 | 3 | 11 | 3 | 11 | 3.813893684 | 2.791979142 | 0.914585397 | 0.669526621 |
| Q9R1C7 | Prpf40a | Pre-mRNA-processing factor 40 homolog A | 431.57 | 10.49 | 108481 | 9 | 2087894.75 | 4104043.5 | 5109804.5 | 3978483.5 | 3535390.75 | 1763031.75 | 1649801.88 | 2842203.5 | 3655568.5 | 2119387.25 | 1477393.25 | 1706884 | 1678887 | 399794.97 | 2577646.75 | 1463864.62 | 1709662.25 | 475992.94 | 1663198.5 | 2311249 | 1 | 1 | 5 | 5 | 5 | 5 | 0.416707354 | 0.65175483 | 0.405193585 | 0.633746619 |
| Q5RKH1 | Prpf4b | Serine/threonine-protein kinase PRP4 homolog | 43.11 | 1.09 | 117007 | 1 | 141083.72 | 192241.77 | 361962.12 | 2 | 415987.72 | 2 | 2 | 2 | 310608.34 | 139402.78 | 2 | 2 | 2 | 2 | 299826.66 | 154083.22 | 2 | 2 | 2 | 178465.38 | 5 | 5.5 | 10 | 12.5 | 10 | 12.5 | 0.269810831 | 0.666273896 | 0.299254372 | 0.738982108 |
| A1A5S1 | Prpf6 | Pre-mRNA-processing factor 6 | 167.76 | 2.23 | 106764 | 3 | 1529940.25 | 1516255.5 | 1059113.12 | 1425819.5 | 1840514.75 | 2305095.25 | 1311521.12 | 1058917.62 | 1613594.88 | 941827.25 | 2156097.25 | 1769660.75 | 2100752.5 | 1847850.12 | 1897764.75 | 1506419.5 | 1119232 | 1774341.25 | 382220.03 | 947858 | 1 | 7 | 5 | 10 | 5 | 10 | 1.325637339 | 1.351429217 | 0.777312559 | 0.792436116 |
| Q99PV0 | Prpf8 | Pre-mRNA-processing-splicing factor 8 | 926.08 | 8.39 | 273616 | 18 | 8017853 | 9022387 | 10229157 | 9108549 | 12537309 | 6305693 | 8894039 | 12124536 | 11768285 | 11618768 | 6334724 | 7203721.5 | 8372107 | 8487225 | 8784629 | 9301047 | 8472266 | 7868096 | 7258710 | 6086425 | 3 | 4 | 5 | 5 | 5 | 5 | 0.801026316 | 0.772656001 | 0.797022197 | 0.768793698 |
| P60892 | Prps1 | Ribose-phosphate pyrophosphokinase 1 | 521.28 | 29.87 | 34834 | 10 | 1908584.25 | 1872750.62 | 1363867 | 1354321.25 | 1080350.62 | 2218772.25 | 2201345.5 | 1545751.12 | 1090540.88 | 1616451.88 | 2717379.75 | 2741849.5 | 1994902 | 1949452.5 | 2065803.75 | 1042386.56 | 1337658.12 | 907363.25 | 1047229.88 | 558474.12 | 0 | 1 | 6 | 1 | 6 | 1 | 1.513137012 | 1.322445577 | 0.645540031 | 0.564186556 |
| P09330 | Prps2 | Ribose-phosphate pyrophosphokinase 2 | 514.2 | 25.79 | 34813 | 9 | 1523141.75 | 1392618.75 | 1703070.62 | 1190029.12 | 910314.62 | 1478963.25 | 1330535.25 | 1191226.5 | 862879 | 1116321 | 731215.94 | 2 | 2 | 2 | 819162 | 1521057.12 | 973621.69 | 1188639.12 | 1081408.38 | 741813.56 | 0 | 6 | 0 | 9 | 0 | 9 | 0.230740228 | 0.25926478 | 0.819526204 | 0.920837614 |
| Q9D0M1 | Prpsap1 | Phosphoribosyl pyrophosphate synthase-associated protein 1 | 647.95 | 42.98 | 39432 | 13 | 9572516 | 9369766 | 9462825 | 7472326 | 9801881 | 8650699 | 11490563 | 8933024 | 10279339 | 10464273 | 8257498.5 | 10147719 | 7708239.5 | 6242252.5 | 7150238.5 | 11148732 | 12165901 | 9298408 | 9761584 | 11193783 | 7 | 5 | 2 | 8 | 7 | 8 | 0.864854231 | 0.793007124 | 1.172706052 | 1.075284389 |
| Q8R574 | Prpsap2 | Phosphoribosyl pyrophosphate synthase-associated protein 2 | 405.44 | 28.73 | 40881 | 7 | 3234208.75 | 2798035 | 3101815.5 | 2074682 | 3263993.5 | 1767067.38 | 2881516.5 | 3608874.75 | 3455931 | 2657259.75 | 1650128.5 | 2230309 | 1384820.38 | 1244193.88 | 1826214.5 | 2809825.5 | 1617899.38 | 2076396.5 | 1772131.25 | 2163369.5 | 1 | 4 | 2 | 5 | 2 | 5 | 0.575956542 | 0.580047988 | 0.721330302 | 0.726454446 |
| Q3T1I4 | Prrc1 | Protein PRRC1 | 131.13 | 6.32 | 46295 | 2 | 1851173.75 | 930587.12 | 1117869.75 | 2090236.88 | 722956.31 | 1344016 | 1185284 | 461024.78 | 1599534.5 | 966332.81 | 1029324.81 | 2 | 888938.88 | 1885466 | 650195.62 | 1744117 | 2503289.5 | 1244841.12 | 1834080.88 | 2113145.5 | 7 | 6 | 9 | 2 | 9 | 6 | 0.663495339 | 0.80161507 | 1.406185276 | 1.698910665 |
| Q3TLH4 | Prrc2c | Protein PRRC2C | 350.99 | 2.64 | 310892 | 6 | 2474803.75 | 3277973.75 | 1478096.75 | 2782684.25 | 2022148.75 | 3001663.25 | 3256020.75 | 3912211.25 | 2600189 | 3234440.5 | 5232972.5 | 5156903.5 | 1847673.88 | 4108240 | 4485934.5 | 3052977.25 | 2860275.75 | 3688260 | 3759927.5 | 4668373 | 4 | 2 | 5 | 9 | 5 | 9 | 1.730826776 | 1.301614681 | 1.498026923 | 1.12654476 |
| Q8BIW1 | Prune1 | Exopolyphosphatase PRUNE1 | 199.37 | 11.01 | 50239 | 4 | 518169.72 | 204134.19 | 297003.25 | 432005.25 | 290897.91 | 491538.25 | 241802.34 | 148216.84 | 136023.17 | 410617.19 | 365946.72 | 178574.67 | 218985.41 | 361571.25 | 99068.01 | 433242.38 | 752564.25 | 684863.25 | 521839.56 | 453871 | 7 | 2 | 10 | 2 | 10 | 2 | 0.702639656 | 0.857126421 | 1.633775445 | 1.992987568 |
| Q61207 | Psap | Prosaposin | 587.79 | 17.06 | 61422 | 9 | 47805758 | 34913794 | 39723790 | 43951550 | 39073166 | 37368754 | 35162606 | 36529214 | 31290032 | 42193594 | 8348112 | 19671462 | 19434842 | 17199734 | 21339280 | 49969498 | 41935718 | 38279466 | 46950258 | 41313006 | 0 | 9 | 0 | 3 | 0 | 9 | 0.418524567 | 0.471082784 | 1.063172291 | 1.196685219 |
| Q99K85 | Psat1 | Phosphoserine aminotransferase | 1227.63 | 57.3 | 40473 | 20 | 52068278 | 46781450 | 40696466 | 37414430 | 36672946 | 51666058 | 56660414 | 45210554 | 44934514 | 44688190 | 94959266 | 87231602 | 83649170 | 65489202 | 74872690 | 59107458 | 59837818 | 52279634 | 52299282 | 48358058 | 0 | 1 | 0 | 4 | 0 | 4 | 1.9013956 | 1.670514809 | 1.272656961 | 1.118122026 |
| Q8BLR5 | Psd4 | PH and SEC7 domain-containing protein 4 | 76.46 | 1.49 | 112733 | 1 | 321672.28 | 2 | 217242.08 | 184646.84 | 274593.56 | 222986.3 | 395027.88 | 191755.39 | 2 | 2 | 2792147.5 | 362196.56 | 192346.47 | 2 | 220541 | 337326.28 | 2 | 2 | 405404.91 | 2 | 9.5 | 11.5 | 9 | 12 | 9.5 | 12 | 3.57382094 | 4.405223463 | 0.744108761 | 0.917215895 |
| O88777 | Psen2 | Presenilin-2 | 108.64 | 3.35 | 50051 | 1 | 2 | 2 | 2 | 2 | 511886.31 | 2 | 2 | 2 | 2 | 619878.5 | 2 | 2 | 2 | 524266.72 | 991497 | 2 | 2 | 2 | 444076.25 | 646667.06 | 9 | 10 | 10 | 10 | 10 | 10 | 2.961098982 | 2.445237507 | 2.1308096 | 1.7595952 |
| Q99JF8 | Psip1 | PC4 and SFRS1-interacting protein | 517.83 | 20.64 | 59697 | 8 | 3894550.75 | 4661254 | 4191207.75 | 4374920.5 | 3691052 | 5217006.5 | 4319226.5 | 3373615.5 | 3933526 | 4886145 | 3846356.25 | 3552853.75 | 3326459.5 | 3634891 | 3788686 | 2797348.75 | 4761857 | 4878037 | 4588276 | 5079272.5 | 2 | 6 | 4 | 12 | 4 | 12 | 0.872015547 | 0.835234599 | 1.062067322 | 1.017270136 |
| Q9R1P4 | Psma1 | Proteasome subunit alpha type-1 | 494.49 | 38.02 | 29547 | 10 | 34519946 | 31288942 | 36623458 | 39450122 | 37002942 | 30724540 | 31446946 | 33633002 | 32242506 | 36052066 | 29698738 | 29954486 | 27189802 | 22763434 | 31313980 | 41139434 | 24950610 | 34268282 | 34400990 | 34921230 | 1 | 9 | 1 | 8 | 1 | 9 | 0.787769332 | 0.858752268 | 0.948543238 | 1.034012907 |
| P17220 | Psma2 | Proteasome subunit alpha type-2 | 564.08 | 41.45 | 25927 | 9 | 10585845 | 9413731 | 9470277 | 8579442 | 5310389 | 9291446 | 9595182 | 8315244.5 | 7208283 | 10155468 | 16390215 | 13612725 | 9652332 | 9694049 | 12833323 | 8917876 | 10126152 | 9861511 | 10180118 | 11549227 | 2 | 6 | 2 | 5 | 2 | 6 | 1.43411202 | 1.39530515 | 1.167787201 | 1.136187043 |
| O70435 | Psma3 | Proteasome subunit alpha type-3 | 621.82 | 42.35 | 28405 | 10 | 55527578 | 48947946 | 53756230 | 59217982 | 56685734 | 57383914 | 48925678 | 47753194 | 54238354 | 50825762 | 25722154 | 29998250 | 28002414 | 30842340 | 28809934 | 63156806 | 71585738 | 49384850 | 61910538 | 62176678 | 0 | 4 | 0 | 3 | 0 | 4 | 0.523008175 | 0.553300684 | 1.12431496 | 1.189435013 |
| P21670 | Psma4 | Proteasome subunit alpha type-4 | 532.88 | 42.15 | 29498 | 11 | 34637294 | 39758130 | 39268918 | 37490934 | 43698718 | 35293918 | 36303214 | 40721954 | 43006610 | 35717946 | 25081612 | 28813530 | 24741352 | 25480368 | 30297114 | 40128098 | 38799038 | 38497346 | 38588522 | 41773586 | 0 | 11 | 0 | 9 | 0 | 11 | 0.689818942 | 0.703577332 | 1.015050223 | 1.035295328 |
| Q9Z2U1 | Psma5 | Proteasome subunit alpha type-5 | 473.69 | 35.27 | 26411 | 7 | 42563850 | 43813890 | 42247862 | 40708374 | 40158350 | 34532306 | 45288506 | 43918946 | 47255726 | 41435626 | 23457532 | 32946344 | 26612800 | 22477588 | 29353862 | 47086058 | 40044934 | 37144546 | 38535022 | 46169234 | 0 | 10 | 0 | 11 | 0 | 11 | 0.643690051 | 0.634785206 | 0.997553457 | 0.983753246 |
| Q9QUM9 | Psma6 | Proteasome subunit alpha type-6 | 569.7 | 43.9 | 27372 | 9 | 38794286 | 44936838 | 36369986 | 39271418 | 33541046 | 48469886 | 40517038 | 38238906 | 34563398 | 39577570 | 30901338 | 29292222 | 26580762 | 32847798 | 27754360 | 44003914 | 54353530 | 45519670 | 48778306 | 43975778 | 0 | 2 | 0 | 3 | 0 | 3 | 0.763950804 | 0.731880734 | 1.226617667 | 1.175125196 |
| Q9Z2U0 | Psma7 | Proteasome subunit alpha type-7 | 576.33 | 48.79 | 27855 | 10 | 21899462 | 22943680 | 23577004 | 19222746 | 34782590 | 21530004 | 23335184 | 23488042 | 37326618 | 19056022 | 22120728 | 24462668 | 22423940 | 18566966 | 24518312 | 23128086 | 19134298 | 21912476 | 26752924 | 21284470 | 12 | 10 | 12 | 10 | 12 | 10 | 0.915598715 | 0.898639774 | 0.916575963 | 0.899598921 |
| O09061 | Psmb1 | Proteasome subunit beta type-1 | 686.8 | 42.5 | 26372 | 10 | 30704826 | 32197196 | 30691250 | 30795814 | 32116516 | 31011662 | 33044142 | 30466014 | 30323474 | 32932442 | 19310314 | 25698122 | 21726730 | 17989260 | 22893718 | 33663106 | 30663978 | 34819570 | 35796082 | 34771150 | 0 | 5 | 0 | 3 | 0 | 5 | 0.687631258 | 0.682087017 | 1.08439496 | 1.075651689 |
| O35955 | Psmb10 | Proteasome subunit beta type-10 | 341.26 | 26.37 | 29063 | 5 | 12975611 | 12312603 | 10610130 | 15013296 | 12905351 | 12426722 | 9759401 | 13977025 | 12908440 | 13805317 | 10627549 | 8004410.5 | 8696787 | 7796473 | 9808742 | 9218034 | 11566576 | 11675385 | 15095800 | 14541102 | 1 | 11 | 2 | 12 | 2 | 12 | 0.704106552 | 0.714633799 | 0.973046457 | 0.987594682 |
| Q9R1P3 | Psmb2 | Proteasome subunit beta type-2 | 592.88 | 53.73 | 22906 | 10 | 24206446 | 24879762 | 23856666 | 23652522 | 22847502 | 20190262 | 22465826 | 23015590 | 22945968 | 24338064 | 21634956 | 23036882 | 20625522 | 9865072 | 19957822 | 20796992 | 21297840 | 23841526 | 16898832 | 26075374 | 1 | 7 | 6 | 11 | 6 | 11 | 0.796365925 | 0.842102219 | 0.911821178 | 0.964188211 |
| Q9R1P1 | Psmb3 | Proteasome subunit beta type-3 | 516.89 | 46.34 | 22965 | 7 | 27129268 | 24944502 | 25328920 | 25717506 | 24780906 | 24789682 | 22705706 | 22623968 | 25809868 | 23534084 | 16821848 | 14959572 | 16350118 | 14727893 | 15896821 | 24703178 | 23940184 | 26651106 | 26542308 | 26521974 | 0 | 12 | 0 | 4 | 0 | 12 | 0.615758979 | 0.659250554 | 1.00357814 | 1.074461708 |
| P99026 | Psmb4 | Proteasome subunit beta type-4 | 487.82 | 39.39 | 29116 | 7 | 20464140 | 25938466 | 25432194 | 26045580 | 24070534 | 24614136 | 20878742 | 23342914 | 25439566 | 26495772 | 22365788 | 21122790 | 19414126 | 18317766 | 18548646 | 23447842 | 22723018 | 25891202 | 26693088 | 24114718 | 2 | 12 | 2 | 11 | 2 | 12 | 0.818108801 | 0.826100708 | 1.007535442 | 1.017377812 |
| Q60692 | Psmb6 | Proteasome subunit beta type-6 | 171.85 | 12.61 | 25379 | 3 | 4689926 | 4987213 | 5375022 | 3604144 | 3616380 | 4484542 | 5613446.5 | 3007370.25 | 4166580.5 | 5315704 | 5705619 | 4755651 | 2875437.5 | 5380860 | 2969666.5 | 6018036 | 5338362 | 5935906 | 6972756.5 | 5346987.5 | 12 | 2 | 12 | 2 | 12 | 2 | 0.973714395 | 0.960137087 | 1.329523046 | 1.310984403 |
| P70195 | Psmb7 | Proteasome subunit beta type-7 | 146.3 | 10.47 | 29891 | 3 | 10512972 | 6508728 | 11099656 | 10194054 | 9118612 | 12339120 | 6609360.5 | 11881779 | 11022334 | 11316144 | 7901286 | 7165113.5 | 9324732 | 10693786 | 9322815 | 13211356 | 10120152 | 14776816 | 15587466 | 15787860 | 10 | 3 | 5 | 4 | 10 | 4 | 0.936200023 | 0.835222625 | 1.464848374 | 1.306851606 |
| P28063 | Psmb8 | Proteasome subunit beta type-8 | 541.64 | 35.51 | 30260 | 8 | 27969680 | 26381824 | 30800556 | 37793706 | 31573004 | 25829130 | 23334344 | 31318560 | 29709702 | 33224496 | 18300172 | 19963792 | 18408878 | 21407786 | 18903370 | 30420206 | 37176170 | 32637282 | 40535330 | 39873350 | 0 | 5 | 0 | 3 | 0 | 5 | 0.62765189 | 0.676241431 | 1.169064043 | 1.259566895 |
| O35522 | Psmb9 | Proteasome subunit beta type-9 bactrianus (Southwestern Asian house mouse) | 150.38 | 16.89 | 23427 | 3 | 16448648 | 15556475 | 14568322 | 21421180 | 15825280 | 15741578 | 15080285 | 15964824 | 15358306 | 18860432 | 7486548 | 8274746 | 8604166 | 8435180 | 8388597 | 16359024 | 19813396 | 16680886 | 16943994 | 17107386 | 0 | 6 | 0 | 4 | 0 | 6 | 0.491401619 | 0.508475043 | 1.036802487 | 1.072825505 |
| P62192 | Psmc1 | 26S proteasome regulatory subunit 4 | 879.95 | 37.27 | 49185 | 15 | 22715296 | 21359824 | 22409186 | 24681490 | 20494524 | 23904400 | 17986380 | 24827388 | 20429402 | 25615522 | 29614318 | 27341712 | 26399240 | 27098434 | 29712988 | 27664566 | 30281016 | 29915560 | 33298300 | 30705396 | 0 | 0 | 0 | 0 | 0 | 0 | 1.255295453 | 1.243019232 | 1.360060924 | 1.346760144 |
| P46471 | Psmc2 | 26S proteasome regulatory subunit 7 | 1266.46 | 47.34 | 48648 | 19 | 22361660 | 24271934 | 24121212 | 25824980 | 18893120 | 26245738 | 23074908 | 22529782 | 23846304 | 24151868 | 30083154 | 30943400 | 28654206 | 26913780 | 28784956 | 21279928 | 27419438 | 25741208 | 24392818 | 23348070 | 0 | 9 | 0 | 10 | 0 | 10 | 1.25899227 | 1.213026235 | 1.058096364 | 1.019465075 |
| O88685 | Psmc3 | 26S proteasome regulatory subunit 6A | 1226.11 | 50.9 | 49549 | 21 | 24161872 | 23793706 | 24623166 | 29012164 | 26585898 | 25354674 | 22940182 | 24869124 | 31100152 | 31999758 | 34103602 | 33767090 | 36029594 | 29907334 | 37536086 | 24886516 | 24608644 | 33260398 | 31505084 | 31483154 | 0 | 5 | 2 | 9 | 2 | 9 | 1.336776218 | 1.257440295 | 1.137052799 | 1.069570199 |
| P54775 | Psmc4 | 26S proteasome regulatory subunit 6B | 735.1 | 31.58 | 47408 | 12 | 24949404 | 26542094 | 27155168 | 22961684 | 21889486 | 25399606 | 23240448 | 28287206 | 28359804 | 27359132 | 29891976 | 32132454 | 27540392 | 26825146 | 28475932 | 23640106 | 23700210 | 28023468 | 26815266 | 29202728 | 1 | 7 | 5 | 12 | 5 | 12 | 1.173023793 | 1.092122536 | 1.063838706 | 0.99046774 |
| P62196 | Psmc5 | 26S proteasome regulatory subunit 8 | 812.52 | 32.76 | 45626 | 12 | 13112931 | 15307546 | 11222798 | 17231260 | 15514786 | 14393889 | 13872406 | 13183716 | 13753204 | 13698878 | 25314816 | 24029862 | 23996252 | 18986224 | 21943556 | 14151786 | 17091702 | 17790078 | 17508130 | 14550083 | 0 | 7 | 0 | 1 | 0 | 7 | 1.578557561 | 1.658450491 | 1.120217428 | 1.176913145 |
| P62334 | Psmc6 | 26S proteasome regulatory subunit 10B | 792.65 | 31.62 | 44173 | 13 | 20524440 | 21762804 | 20749974 | 22369258 | 23945754 | 17097438 | 22000520 | 25098494 | 27026154 | 23936512 | 23154216 | 30671644 | 27042656 | 21073628 | 30932800 | 21965392 | 17163644 | 22519302 | 19682986 | 21490424 | 4 | 9 | 7 | 6 | 7 | 9 | 1.215109596 | 1.153837806 | 0.940280303 | 0.892866755 |
| Q3TXS7 | Psmd1 | 26S proteasome non-ATPase regulatory subunit 1 | 1482.98 | 32.63 | 105730 | 23 | 55468586 | 54076154 | 57915794 | 61918614 | 59274762 | 59990202 | 53253818 | 59053586 | 62481918 | 60933998 | 51311802 | 56715930 | 56389742 | 52839334 | 58954954 | 60841706 | 58061674 | 66613226 | 71476898 | 69953074 | 7 | 3 | 3 | 6 | 7 | 6 | 0.956895966 | 0.934051849 | 1.132659447 | 1.105619303 |
| Q9Z2X2 | Psmd10 | 26S proteasome non-ATPase regulatory subunit 10 | 65.72 | 3.46 | 25084 | 1 | 2 | 2 | 2 | 2 | 499568.69 | 243766.62 | 338128.34 | 2 | 698670.25 | 2 | 1386957.38 | 2 | 1177710.38 | 1045132.75 | 966720.19 | 443989.03 | 2 | 1041188.69 | 760876.88 | 947804.88 | 3 | 4 | 4 | 5 | 4 | 5 | 9.160801117 | 3.573819099 | 6.393135516 | 2.494095169 |
| Q8BG32 | Psmd11 | 26S proteasome non-ATPase regulatory subunit 11 | 1178.5 | 43.36 | 47437 | 19 | 43115542 | 45971598 | 49639946 | 42768986 | 47344018 | 39988054 | 43694806 | 49444098 | 49294858 | 43117618 | 50376494 | 64643422 | 59851398 | 49765850 | 56213430 | 46826298 | 39495294 | 51290866 | 44728070 | 51810014 | 0 | 10 | 0 | 9 | 0 | 10 | 1.227278813 | 1.24523942 | 1.023205951 | 1.038180055 |
| Q9D8W5 | Psmd12 | 26S proteasome non-ATPase regulatory subunit 12 | 955.69 | 41.45 | 52895 | 17 | 21468450 | 20612268 | 29491250 | 32018038 | 35728310 | 27536166 | 24854764 | 36505706 | 37525834 | 37853278 | 31932082 | 35890094 | 44411050 | 35585162 | 44290626 | 28113742 | 33182398 | 38393738 | 39400170 | 32045832 | 3 | 5 | 9 | 9 | 9 | 9 | 1.378921448 | 1.169430158 | 1.228380337 | 1.041759859 |
| Q9WVJ2 | Psmd13 | 26S proteasome non-ATPase regulatory subunit 13 | 632.09 | 30.59 | 42809 | 13 | 34339210 | 29492540 | 33017972 | 37445674 | 36356310 | 27715364 | 32831232 | 33829290 | 33070110 | 35053626 | 34619530 | 39985766 | 36327830 | 30756688 | 42907478 | 37272298 | 36716010 | 31928936 | 34929970 | 37427390 | 8 | 9 | 5 | 5 | 8 | 9 | 1.081719582 | 1.135985978 | 1.044669334 | 1.097077038 |
| O35593 | Psmd14 | 26S proteasome non-ATPase regulatory subunit 14 | 395.13 | 38.06 | 34577 | 6 | 8660083 | 9181106 | 7990924 | 10538366 | 8660145 | 8050112.5 | 9277148 | 8632354 | 7554669.5 | 8781094 | 12913041 | 11234559 | 11654591 | 13505668 | 11727562 | 9221145 | 10776824 | 10833144 | 11693914 | 9840708 | 0 | 2 | 0 | 1 | 0 | 2 | 1.35542028 | 1.443075435 | 1.162891613 | 1.238095921 |
| Q8VDM4 | Psmd2 | 26S proteasome non-ATPase regulatory subunit 2 | 1719 | 38.22 | 100203 | 31 | 50373054 | 56123102 | 45277794 | 49593886 | 51604238 | 49133034 | 50821934 | 50905102 | 50466018 | 48561538 | 52697726 | 51951054 | 53410050 | 47224442 | 53366362 | 50167990 | 49809618 | 48339306 | 54424330 | 51184662 | 8 | 12 | 5 | 11 | 8 | 12 | 1.022443426 | 1.035063793 | 1.003770503 | 1.016160384 |

| | | | | | | | | | | | | | | | | | | | | | | | | | | | | | | | | | | | | |
|---|---|---|---|---|---|---|---|---|---|---|---|---|---|---|---|---|---|---|---|---|---|---|---|---|---|---|---|---|---|---|---|---|---|---|---|---|
| P14685 | Psmd3 | 26S proteasome non-ATPase regulatory subunit 3 | 512.69 | 18.3 | 60718 | 9 | 15576631 | 16294128 | 33620286 | 29181134 | 23333286 | 21534354 | 20317490 | 32958640 | 37608602 | 35508378 | 26971254 | 41848658 | 57041410 | 53707942 | 57721422 | 16098984 | 16004489 | 32796614 | 34211194 | 31111488 | 2 | 10 | 3 | 7 | 3 | 10 | 2.010844888 | 1.604101629 | 1.10353168 | 0.880315024 |
| O35226 | Psmd4 | 26S proteasome non-ATPase regulatory subunit 4 | 262.79 | 14.63 | 40704 | 5 | 8589708 | 7933462 | 4877478.5 | 6392847 | 7057219 | 7014531.5 | 8462793 | 8950104 | 7308380 | 7332716.5 | 7716910 | 7803515.5 | 7702145 | 8451500 | 6894663.5 | 6517361.5 | 5772600 | 8975346 | 8249782 | 7971222.5 | 10 | 9 | 12 | 11 | 12 | 11 | 1.106684169 | 0.987207324 | 1.075625351 | 0.959501594 |
| Q8BJY1 | Psmd5 | 26S proteasome non-ATPase regulatory subunit 5 | 443.46 | 21.03 | 55972 | 8 | 8620776 | 9456506 | 8177330.5 | 8312016.5 | 7845940 | 9359244 | 7318988.5 | 8203254.5 | 9288543 | 8444428 | 9050752 | 10442376 | 9981055 | 7157187.5 | 9691174 | 8426022 | 9621523 | 7660228.5 | 6721214.5 | 7059148.5 | 6 | 8 | 7 | 8 | 7 | 8 | 1.092189075 | 1.087014752 | 0.931047975 | 0.92663707 |
| Q99JI4 | Psmd6 | 26S proteasome non-ATPase regulatory subunit 6 | 807.35 | 44.73 | 45536 | 14 | 10805017 | 8953385 | 12447206 | 12421972 | 14626454 | 8586283 | 8110821 | 13934345 | 14621918 | 12815959 | 11715856 | 14228815 | 13647904 | 13187670 | 13689256 | 12453567 | 8748414 | 13215045 | 11071256 | 13703584 | 7 | 11 | 10 | 12 | 10 | 12 | 1.121771743 | 1.144657698 | 0.998950822 | 1.019331032 |
| P26516 | Psmd7 | 26S proteasome non-ATPase regulatory subunit 7 | 392.95 | 19.63 | 36540 | 5 | 7096587 | 6687528 | 6197373 | 5849048 | 6022976.5 | 7960283.5 | 7322485.5 | 5274593.5 | 6012081.5 | 6647295.5 | 8920821 | 9329314 | 8413363 | 5281501 | 6806095 | 7407720.5 | 6985313 | 5777008.5 | 6531085.5 | 5910337 | 6 | 12 | 6 | 11 | 6 | 12 | 1.216540688 | 1.166613418 | 1.023794927 | 0.981778013 |
| Q9CX56 | Psmd8 | 26S proteasome non-ATPase regulatory subunit 8 | 335.79 | 17.85 | 39930 | 7 | 17946492 | 13347273 | 14431803 | 13214330 | 19030170 | 11617676 | 17735372 | 16679696 | 19196130 | 15577289 | 5276102 | 6258011 | 10147525 | 3735441 | 7022559.5 | 18271310 | 11114215 | 15466667 | 22895178 | 15523344 | 0 | 10 | 0 | 11 | 0 | 11 | 0.416052459 | 0.401450054 | 1.067983088 | 1.030499542 |
| P97371 | Psme1 | Proteasome activator complex subunit 1 | 716.2 | 37.35 | 28673 | 11 | 50554214 | 46743382 | 47803094 | 52192054 | 61949566 | 35859638 | 49624254 | 48466754 | 55925502 | 42779722 | 7912880 | 16947016 | 17394870 | 9961257 | 18667182 | 51871158 | 24889188 | 35946666 | 38442590 | 43944934 | 0 | 3 | 0 | 8 | 0 | 8 | 0.273424523 | 0.304669747 | 0.752556695 | 0.83855411 |
| P97372 | Psme2 | Proteasome activator complex subunit 2 | 760.54 | 39.33 | 27057 | 9 | 55244570 | 59556870 | 59887874 | 70152786 | 80794386 | 50051582 | 58843426 | 68389258 | 71268506 | 60632126 | 24003782 | 31503716 | 30136716 | 21603628 | 31579852 | 65642118 | 50552778 | 55514486 | 59057146 | 61408438 | 0 | 8 | 0 | 10 | 0 | 10 | 0.426327208 | 0.449011885 | 0.897242719 | 0.944984596 |
| P61290 | Psme3 | Proteasome activator complex subunit 3 | 370.7 | 22.44 | 29506 | 5 | 6409303 | 5355056.5 | 6078221 | 5434244 | 6116694 | 5005415.5 | 5570712.5 | 4727530.5 | 5048589 | 5478206 | 2903544.5 | 5144976 | 3303461.75 | 2504178.25 | 3931916.5 | 7044005.5 | 4316214.5 | 4379919 | 3747592.25 | 4702297 | 0 | 5 | 3 | 5 | 3 | 5 | 0.605170048 | 0.688647491 | 0.822971508 | 0.936492588 |
| Q8BHL8 | Psmf1 | Proteasome inhibitor PI31 subunit | 68.29 | 4.43 | 29664 | 1 | 2231417.5 | 1663058.25 | 1195666.38 | 1451504.12 | 1016401.5 | 1948559.62 | 1657584.88 | 1635742.75 | 1335141.12 | 1313600.5 | 1759093.38 | 1049944.88 | 737799.5 | 896411.94 | 866807.19 | 2028474.75 | 2687988.75 | 1768189 | 1448874.38 | 1409857.75 | 5 | 8 | 4 | 7 | 5 | 8 | 0.702569905 | 0.672957375 | 1.236216671 | 1.184111531 |
| Q9JK23 | Psmg1 | Proteasome assembly chaperone 1 | 225.19 | 17.65 | 33104 | 4 | 4545716.5 | 4343812 | 4475860.5 | 3820979.5 | 3874593.75 | 4087179.5 | 4801963.5 | 4811014 | 5930463 | 4930594 | 3207107 | 5694478.5 | 4962985 | 2274685.25 | 4132872.5 | 4714127 | 2270118.75 | 3093371.5 | 2453976.75 | 3276294 | 12 | 5 | 9 | 1 | 12 | 5 | 0.962545206 | 0.82537159 | 0.750577671 | 0.643611834 |
| Q9EST4 | Psmg2 | Proteasome assembly chaperone 2 | 90.96 | 6.06 | 29524 | 2 | 1919413.62 | 2000728.12 | 1987396.62 | 2330912.25 | 2446069.75 | 1923297.12 | 2085505.25 | 2237558.5 | 2273858 | 1753810 | 945688.56 | 1382125.88 | 1630402.88 | 732623.88 | 1213266.25 | 2364676 | 1031825.62 | 1483787.62 | 1451543.12 | 1967789.25 | 0 | 5 | 0 | 7 | 0 | 7 | 0.552585165 | 0.574663311 | 0.776789349 | 0.807825413 |
| Q9CZH3 | Psmg3 | Proteasome assembly chaperone 3 | 43.49 | 19.67 | 13331 | 1 | 1682673.62 | 1382856.88 | 1371002.88 | 1632936.12 | 2 | 1207703.62 | 1487559.25 | 2 | 1608918.88 | 354035.66 | 1718875.25 | 2 | 792670.12 | 1213012.25 | 693161.38 | 2 | 2 | 1370486.12 | 1045989.5 | 1303218.38 | 8.5 | 4 | 12.5 | 9 | 12.5 | 9 | 0.727859254 | 0.948371172 | 0.612853689 | 0.798523571 |
| Q4KLH4 | Pspc1 | Paraspeckle component 1 | 290.98 | 10.73 | 58760 | 5 | 879982.38 | 1698759.5 | 2453202.5 | 1852635.5 | 2579720.5 | 1197167.75 | 1972825.75 | 2272910.25 | 2181744.5 | 1777676.25 | 759675.44 | 1446199.75 | 1164660.62 | 724556.94 | 1010657.88 | 2001392.5 | 762806.38 | 1491196.75 | 1212497.12 | 1669640.75 | 3 | 6 | 1 | 6 | 3 | 6 | 0.539474702 | 0.543030676 | 0.754153314 | 0.759124353 |
| P97814 | Pstpip1 | Proline-serine-threonine phosphatase-interacting protein 1 | 216.4 | 9.16 | 47590 | 3 | 789463.19 | 3461181.75 | 677371.25 | 3508670.5 | 797336.94 | 3441423.75 | 783989.19 | 774602 | 735802.38 | 792531.5 | 762325.94 | 524735.88 | 2200807.5 | 2578268.5 | 3380515.25 | 3050237.25 | 2727893 | 3330552.5 | 820869.94 | 1317145.25 | 10 | 10 | 12 | 5 | 12 | 10 | 1.023026738 | 1.44702027 | 1.217962872 | 1.722747704 |
| P17225 | Ptbp1 | Polypyrimidine tract-binding protein 1 | 1655.29 | 60.34 | 56478 | 21 | 589310.81 | 1566003.25 | 587812.19 | 526294.94 | 410493.84 | 976441.38 | 572608.88 | 603848.88 | 384870.66 | 477762.06 | 1183280.25 | 432687.31 | 400345.75 | 544764.44 | 657708.44 | 579289.25 | 711397.62 | 484435.94 | 879135.38 | 704377 | 11 | 10 | 12 | 8 | 12 | 10 | 0.874690357 | 1.067402481 | 0.91269368 | 1.113778712 |
| Q8BHD7 | Ptbp3 | Polypyrimidine tract-binding protein 3 | 435.6 | 18.36 | 56701 | 7 | 1362937.12 | 746414.75 | 1561152 | 1616961.75 | 653038 | 1184182.38 | 2 | 536017.56 | 1761629.12 | 2329616 | 1942655.62 | 1884788.88 | 2250588.5 | 1523249.5 | 1623601.75 | 769615.75 | 1240091.75 | 2039106.12 | 599447.31 | 1848843 | 2 | 11 | 7 | 10 | 7 | 11 | 1.552879156 | 1.587364413 | 1.093695812 | 1.117983845 |
| Q14C51 | Ptcd3 | Pentatricopeptide repeat domain-containing protein 3, mitochondrial | 149.68 | 4.23 | 77796 | 3 | 1376886.25 | 1263766.5 | 2118793 | 2207341 | 2830322.5 | 1583386.25 | 1648261.25 | 2285489.5 | 3279645 | 2641429 | 1464733 | 2897579 | 3536529.75 | 3558791 | 3904402 | 1279738 | 1477762 | 2250956 | 2044238.5 | 2037095 | 3 | 11 | 6 | 6 | 6 | 11 | 1.568017091 | 1.343045232 | 0.927803219 | 0.794686293 |
| Q60866 | Pter | Phosphotriesterase-related protein | 148.07 | 10.32 | 39218 | 3 | 2 | 2 | 298667.09 | 299202.38 | 2 | 2 | 407480.59 | 2 | 2 | 2 | 761132.62 | 2 | 987077.06 | 2 | 2 | 2 | 696426.12 | 2 | 2 | 2 | 10.5 | 11 | 9 | 12 | 10.5 | 12 | 2.924046508 | 4.290219954 | 1.164848125 | 1.70908864 |
| P83868 | Ptges3 | Prostaglandin E synthase 3 | 88.26 | 18.75 | 18721 | 2 | 1530239.38 | 1414814.12 | 1462029.75 | 1222458.88 | 2238902.5 | 2 | 1904321.5 | 2013273.62 | 1961896.75 | 1419707 | 1443952.12 | 2755946 | 2289757 | 1448117.5 | 2095585.12 | 1598748.12 | 788037.81 | 1112927.75 | 2 | 135931.55 | 7 | 4 | 6 | 5.5 | 7 | 5.5 | 1.275138634 | 1.374583043 | 0.462054116 | 0.498088393 |
| P43252 | Ptgir | Prostacyclin receptor | 58.46 | 2.65 | 44463 | 1 | 2 | 2 | 2 | 2 | 741393.19 | 2 | 2 | 2 | 2 | 1021421.88 | 228513.16 | 1863760.25 | 2316463 | 1683491.38 | 1663605.38 | 2 | 2 | 908284.81 | 2 | 2 | 1 | 12 | 1 | 12 | 1 | 12 | 10.46104764 | 7.593113656 | 1.225102984 | 0.889236577 |
| Q91YR9 | Ptgr1 | Prostaglandin reductase 1 | 474.21 | 26.75 | 35560 | 7 | 15522848 | 13454757 | 14277792 | 14353274 | 12750000 | 14116422 | 16073166 | 10728833 | 11450650 | 11707313 | 11186127 | 11641085 | 6815450 | 6370235 | 8119334 | 15212488 | 10781982 | 10865785 | 10497802 | 10625268 | 0 | 4 | 3 | 6 | 3 | 6 | 0.627246512 | 0.688744093 | 0.824110578 | 0.904909444 |
| P22437 | Ptgs1 | Prostaglandin G/H synthase 1 | 478.21 | 13.62 | 69042 | 8 | 786628.25 | 467204 | 1391590.25 | 1330005.75 | 767452.62 | 759531.12 | 1077425.38 | 307817.06 | 1197112.25 | 320062.97 | 1852247 | 1676550.75 | 1451391.25 | 1205037.88 | 1311884.62 | 703962 | 436246.16 | 292730.44 | 177268.7 | 343457.06 | 4 | 1 | 0 | 6 | 4 | 6 | 1.580708372 | 2.047301028 | 0.411915124 | 0.53350401 |
| Q05769 | Ptgs2 | Prostaglandin G/H synthase 2 | 972.47 | 34.6 | 69013 | 16 | 3143319 | 2768988.5 | 2693109.5 | 2463459 | 2382702 | 2378090.75 | 2631982.5 | 4180545 | 4386155.5 | 4023008.5 | 10844819 | 12129170 | 17741340 | 17629320 | 19329358 | 1970288.5 | 2564236.25 | 4586106 | 3715207.5 | 3373953 | 0 | 8 | 0 | 10 | 0 | 10 | 5.774341642 | 4.413350455 | 1.20504756 | 0.921022262 |
| Q9QVP9 | Ptk2b | Protein-tyrosine kinase 2-beta | 1119.4 | 21.21 | 115794 | 20 | 12181882 | 12754386 | 15289981 | 14125902 | 16734696 | 13231979 | 15490453 | 19165012 | 20788088 | 15657005 | 11150609 | 14299393 | 13275522 | 12347302 | 11684730 | 11654069 | 9862581 | 15755158 | 15956968 | 15540395 | 6 | 12 | 2 | 8 | 6 | 12 | 0.882829365 | 0.74416777 | 0.967396556 | 0.815452415 |
| P26350 | Ptma | Prothymosin alpha | 690.84 | 38.74 | 12254 | 7 | 69578482 | 83228450 | 82588546 | 98298322 | 82713546 | 74604402 | 85774898 | 62579010 | 69117098 | 76325722 | 37938678 | 49746886 | 34317674 | 43055990 | 33806214 | 73485778 | 84848610 | 62528190 | 61495162 | 75114082 | 0 | 6 | 0 | 9 | 0 | 9 | 0.477574289 | 0.539806819 | 0.858466656 | 0.970333131 |
| P04550 | Ptms | Parathymosin | 222.78 | 22.55 | 11559 | 3 | 40839666 | 36883378 | 29901414 | 56078398 | 31743398 | 41980234 | 29979250 | 41436078 | 29411958 | 37609434 | 12787156 | 11005493 | 10181458 | 11568436 | 10122685 | 45860466 | 53404394 | 28774614 | 29299780 | 39408214 | 0 | 11 | 0 | 12 | 0 | 12 | 0.284810923 | 0.308536569 | 1.006657656 | 1.090515407 |
| P58389 | Ptpa | Serine/threonine-protein phosphatase 2A activator | 202.12 | 12.07 | 36710 | 3 | 3938742.5 | 3713651 | 4093979.25 | 2912235 | 4870607.5 | 2161377.25 | 3375384.75 | 3650901.5 | 3542191 | 2467944.75 | 837075.12 | 2324851 | 1489363.75 | 433269.94 | 1290736.75 | 3603244 | 904473.88 | 1734356.88 | 1888132.75 | 2440945 | 0 | 1 | 1 | 5 | 1 | 5 | 0.326449193 | 0.419488141 | 0.541299401 | 0.695571269 |
| P35821 | Ptpn1 | Tyrosine-protein phosphatase non-receptor type 1 | 267.68 | 11.11 | 49593 | 5 | 5119223.5 | 4060743.5 | 4995889.5 | 4095963.75 | 4165967 | 4567977 | 5326920.5 | 2994029.75 | 4549479.5 | 2855115 | 4714942.5 | 6042202 | 4512932.5 | 3157120 | 4658859 | 4839504 | 4294902 | 4311001 | 1655170 | 4274174 | 11 | 12 | 8 | 10 | 11 | 12 | 1.028891831 | 1.137607178 | 0.863487597 | 0.954725909 |
| P35831 | Ptpn12 | Tyrosine-protein phosphatase non-receptor type 12 | 88.64 | 2.58 | 86526 | 2 | 1597359.88 | 1734961.5 | 1233972.62 | 1314856.88 | 68071.05 | 1603187.75 | 1496194.75 | 1430500.5 | 1162909.12 | 120277.95 | 1745934.25 | 1734952.62 | 1915776.12 | 1867414.88 | 1645269.88 | 1524814.75 | 59299.74 | 1272033.75 | 1940852.5 | 1358502.5 | 2 | 12 | 0 | 12 | 2 | 12 | 1.497565203 | 1.532640695 | 1.034673662 | 1.058907456 |
| O88902 | Ptpn23 | Tyrosine-protein phosphatase non-receptor type 23 | 136.54 | 3 | 163454 | 3 | 654283.19 | 1369009 | 600017.81 | 943320.38 | 618157.12 | 1083491.25 | 1560118.62 | 789826.81 | 1020125.5 | 967767.62 | 2590779.5 | 1828863.88 | 2033796.75 | 2137492.25 | 2953757.5 | 1377971.5 | 2282630.75 | 1130235.5 | 1083242.25 | 1381260.75 | 0 | 2 | 0 | 5 | 0 | 5 | 2.758727864 | 2.12949411 | 1.733741737 | 1.338295403 |
| P29351 | Ptpn6 | Tyrosine-protein phosphatase non-receptor type 6 | 2475.16 | 62.86 | 67559 | 38 | 78698506 | 88988306 | 153955650 | 111864866 | 130765298 | 76364514 | 107054194 | 143812546 | 137764354 | 130518522 | 102587810 | 110137698 | 106480746 | 108974026 | 114324562 | 91195778 | 68063938 | 119715394 | 107847274 | 115917810 | 11 | 10 | 8 | 7 | 11 | 10 | 0.961423285 | 0.910985676 | 0.890952655 | 0.844212032 |
| P49445 | Ptpn7 | Tyrosine-protein phosphatase non-receptor type 7 | 42.79 | 3.62 | 40314 | 1 | 398157.69 | 306346.66 | 130608.91 | 2 | 186062.22 | 217234.67 | 442715.03 | 222743.36 | 201168.89 | 197727.88 | 2 | 2 | 151879.05 | 2 | 337893.69 | 88860.22 | 2 | 2 | 2 | 2 | 7.5 | 3 | 4 | 0 | 7.5 | 3 | 0.479621564 | 0.382164971 | 0.087025245 | 0.06934217 |
| O35239 | Ptpn9 | Tyrosine-protein phosphatase non-receptor type 9 | 41.72 | 1.85 | 67970 | 1 | 200062.72 | 197405.41 | 211414.22 | 187762.52 | 284821.97 | 2 | 176809.06 | 216479.41 | 208875.12 | 2 | 2 | 166338.62 | 158155.69 | 2 | 156748.3 | 136807.47 | 2 | 2 | 2 | 114409.54 | 0 | 0 | 8 | 7 | 8 | 7 | 0.444994328 | 0.799190488 | 0.232298394 | 0.417197827 |
| P18052 | Ptpra | Receptor-type tyrosine-protein phosphatase alpha | 425.71 | 9.41 | 93698 | 8 | 4112161.25 | 3548977.25 | 3554041.75 | 3226486 | 2064478.25 | 5909423 | 4420819.5 | 2276735.5 | 2722222.75 | 2393950 | 7241782 | 4108408.25 | 4666610 | 4702493.5 | 6191638 | 3291798.25 | 2812134.5 | 4208926.5 | 4622688.5 | 2935391 | 1 | 11 | 4 | 9 | 4 | 11 | 1.630358425 | 1.518405623 | 1.082684012 | 1.008338698 |
| P06800 | Ptprc | Receptor-type tyrosine-protein phosphatase C | 2259.24 | 29.23 | 144837 | 40 | 91229538 | 92812330 | 117699322 | 124862818 | 103101458 | 102408978 | 99722210 | 106085930 | 115456482 | 118813210 | 109699242 | 116111770 | 122750130 | 108875706 | 117174642 | 86216010 | 83516202 | 96049058 | 86384962 | 99639066 | 9 | 4 | 6 | 0 | 9 | 4 | 1.084775459 | 1.05921744 | 0.852936824 | 0.832841075 |
| B2GV87 | Ptpre | Receptor-type tyrosine-protein phosphatase epsilon | 50.67 | 2 | 80734 | 1 | 591901.56 | 491960.59 | 306642.19 | 288741.69 | 392512.06 | 564622.19 | 420994.34 | 2 | 374646.16 | 346786.16 | 519523.25 | 457960.38 | 472604.78 | 410880.91 | 442316.31 | 384470.19 | 466310.16 | 571864.69 | 596186.5 | 551027.25 | 9 | 7 | 6 | 4 | 9 | 7 | 1.111754138 | 1.349277691 | 1.240424161 | 1.50543775 |
| Q8R2Y8 | Ptrh2 | Peptidyl-tRNA hydrolase 2, mitochondrial | 226.42 | 27.62 | 19527 | 3 | 100086.88 | 530173.25 | 1231998 | 297622.38 | 931332.5 | 166564.58 | 2 | 504484.69 | 375733.44 | 501903.62 | 2 | 675449.06 | 730380.25 | 367589.09 | 381822.31 | 429193.75 | 252502.7 | 206031.09 | 2 | 496895.91 | 10 | 6 | 9.5 | 10.5 | 10 | 10.5 | 0.697215851 | 1.391656842 | 0.447923015 | 0.894063333 |
| Q3UEB3 | Puf60 | Poly(U)-binding-splicing factor PUF60 | 775.99 | 36.7 | 60249 | 13 | 13664203 | 14686804 | 13551204 | 11025070 | 11878971 | 10581844 | 14159367 | 9059887 | 12229378 | 9267876 | 8976399 | 9452505 | 8900783 | 6323087 | 7762405.5 | 14558116 | 14652739 | 12726757 | 11323936 | 10693201 | 0 | 11 | 2 | 5 | 2 | 11 | 0.639061483 | 0.748940574 | 0.986860789 | 1.156539873 |
| Q8BKS9 | Pum3 | Pumilio homolog 3 | 94 | 4.64 | 72800 | 2 | 580776.75 | 662025.69 | 641102.56 | 940811.31 | 885422.88 | 685543.06 | 604456.81 | 667301.75 | 665940 | 449364.19 | 2 | 403623.62 | 361934.12 | 376858.03 | 331562.38 | 743141.81 | 843267.88 | 691726.94 | 694792.12 | 663709.31 | 0 | 10 | 0 | 3 | 0 | 10 | 0.397284327 | 0.479716645 | 0.980189118 | 1.183568048 |
| P42669 | Pura | Transcriptional activator protein Pur-alpha | 134.37 | 12.77 | 34884 | 3 | 1083241.5 | 743648.94 | 1035008.88 | 1696576.5 | 795160.25 | 634095.94 | 1018748.81 | 161868.7 | 826597.69 | 1717417.75 | 2672070 | 2539794 | 2125801.5 | 1342810.62 | 2668061.25 | 894749.38 | 406294.16 | 79114.3 | 707323.88 | 843116.81 | 1 | 4 | 1 | 9 | 1 | 9 | 2.119781252 | 2.603634605 | 0.547403389 | 0.672351643 |
| O35295 | Purb | Transcriptional activator protein Pur-beta | 266.61 | 24.38 | 33901 | 4 | 2022799.25 | 3296494.25 | 2306953.5 | 2287144.75 | 3296305 | 2555949.25 | 2913621.5 | 3368375.5 | 4166682.5 | 3419599.5 | 2541566.25 | 1994911.12 | 2200332.5 | 1775318.62 | 2153009 | 2683698 | 2229227 | 2659958.5 | 3117017.25 | 4055163.5 | 5 | 10 | 0 | 8 | 5 | 10 | 0.807371864 | 0.64935395 | 1.116230337 | 0.897762989 |
| Q4KM92 | Pus1 | tRNA pseudouridine synthase A | 43.33 | 2.36 | 47515 | 1 | 2 | 2 | 2 | 2 | 2 | 2 | 2 | 2 | 2 | 2 | 772273.25 | 597903.56 | 497151.59 | 2 | 537549.75 | 2 | 2 | 2 | 2 | 2 | 2.5 | 12.5 | 2.5 | 12.5 | 2.5 | 12.5 | 240488.015 | 240488.015 | 1 | 1 |
| Q8BU03 | Pwp2 | Periodic tryptophan protein 2 homolog | 127.92 | 3.26 | 102910 | 3 | 699478.69 | 1888993.12 | 474707.72 | 927675 | 289635.34 | 481527.59 | 846134.06 | 1444760.12 | 1018405.62 | 859638.38 | 3445445 | 2205845.5 | 2232691.5 | 2107989 | 3077145 | 902355.56 | 1556375.62 | 859852.62 | 375902.41 | 801214.94 | 0 | 11 | 0 | 12 | 0 | 12 | 3.053182322 | 2.810281087 | 1.050277255 | 0.966720619 |
| Q66H76 | Pxn | Paxillin | 120.77 | 3.92 | 64019 | 3 | 2518412 | 3231852.75 | 3516242.25 | 3604868 | 2590649 | 2614897 | 2535192.25 | 4073718 | 2117609.5 | 2276496 | 1765667.5 | 1769480.5 | 1596168.5 | 1999467.5 | 1988295.5 | 3093190.75 | 3660646.5 | 3337575.5 | 4356351 | 3254593 | 0 | 7 | 0 | 4 | 0 | 7 | 0.589772691 | 0.669638561 | 1.144892593 | 1.29993172 |
| Q9EPB4 | Pycard | Apoptosis-associated speck-like protein containing a CARD | 328.2 | 43.01 | 21459 | 6 | 8257800 | 8798768 | 10893768 | 8100220 | 10649589 | 5771371.5 | 7889947 | 8020748 | 10379690 | 8949953 | 2545522.25 | 4555039 | 4273481.5 | 3007083 | 4744737 | 7015427 | 3755338.5 | 6547450 | 4273185 | 7990366 | 0 | 0 | 0 | 4 | 0 | 4 | 0.409546111 | 0.466351269 | 0.633440571 | 0.721300498 |
| Q922Q4 | Pycr2 | Pyrroline-5-carboxylate reductase 2 | 856.09 | 45.94 | 33659 | 13 | 35769242 | 30337686 | 38065162 | 44802874 | 32124470 | 36117762 | 34698094 | 40950022 | 38085174 | 41444098 | 39932966 | 36335546 | 45174070 | 52004066 | 43107382 | 34120298 | 37581290 | 43669502 | 50748186 | 39795738 | 4 | 7 | 5 | 10 | 5 | 10 | 1.195774196 | 1.132041403 | 1.137027375 | 1.076425691 |
| P53534 | Pygb | Glycogen phosphorylase, brain form | 871.84 | 19.09 | 96174 | 15 | 18244850 | 17799186 | 18200130 | 16737296 | 23066866 | 13948239 | 17028260 | 22343636 | 22001712 | 16215926 | 7046817.5 | 8231821 | 8822479 | 5368501 | 8261356 | 16399435 | 11798240 | 11463429 | 10741095 | 10378699 | 0 | 0 | 0 | 2 | 0 | 2 | 0.401187084 | 0.412190217 | 0.646273031 | 0.663997998 |
| Q61838 | Pzp | Pregnancy zone protein | 121.74 | 1.61 | 165853 | 3 | 2762633.75 | 2529627.25 | 3738349.25 | 3290433.5 | 2583552.75 | 2923163.5 | 3623689 | 3432464.5 | 3965246.25 | 4719979.5 | 4635071.5 | 6804871 | 8201470.5 | 4946939 | 7872005.5 | 2620860.75 | 1834968.75 | 3838348.25 | 4045025.5 | 3070384.75 | 0 | 10 | 1 | 8 | 1 | 10 | 2.177875631 | 1.739145605 | 1.033881595 | 0.825607582 |
| Q8BML9 | Qars | Glutamine--tRNA ligase | 466.91 | 10.97 | 87677 | 8 | 9134832 | 9949953 | 9600886 | 8470994 | 9108808 | 7796321.5 | 8261620 | 9488416 | 8579497 | 8119739.5 | 9627052 | 10562320 | 12416507 | 6031224.5 | 9596333 | 8530659 | 7505828 | 9506116 | 9800709 | 10827121 | 8 | 12 | 5 | 7 | 8 | 12 | 1.042536332 | 1.141738864 | 0.997945768 | 1.092905286 |
| Q8BVI4 | Qdpr | Dihydropteridine reductase | 281 | 21.58 | 25570 | 4 | 12435898 | 14598934 | 13728056 | 12619882 | 11702171 | 13734372 | 13142331 | 15176777 | 14078325 | 14070203 | 5986159 | 8877218 | 8905944 | 9490798 | 8574603 | 13635406 | 10599172 | 16800260 | 14480734 | 15114327 | 0 | 8 | 0 | 11 | 0 | 11 | 0.642771144 | 0.595919165 | 1.085195714 | 1.006095139 |
| Q91XU1 | Qki | Protein quaking | 303.05 | 27.27 | 37643 | 7 | 5061885 | 4405991.5 | 6675113 | 6601609 | 6459991.5 | 6396417.5 | 5761862 | 5075800.5 | 6198036 | 5415473 | 6259436.5 | 5954313 | 6697351.5 | 8035994.5 | 6864313 | 5133785.5 | 4374312 | 6849428 | 6678429 | 6729037.5 | 6 | 8 | 3 | 9 | 6 | 9 | 1.157742961 | 1.172070515 | 1.019188833 | 1.031801722 |
| P61027 | Rab10 | Ras-related protein Rab-10 | 840.32 | 49.5 | 22541 | 15 | 27415796 | 26846240 | 24438256 | 20186822 | 27459814 | 20709464 | 29458474 | 26351492 | 26316594 | 24731964 | 15835780 | 22042450 | 21809578 | 10851045 | 12866319 | 29932622 | 19510298 | 21180980 | 12055697 | 20943280 | 2 | 7 | 2 | 7 | 2 | 7 | 0.660128215 | 0.653809575 | 0.820145599 | 0.812295299 |
| O35509 | Rab11b | Ras-related protein Rab-11B | 458.53 | 33.49 | 24489 | 8 | 47063222 | 27587908 | 49455914 | 35168842 | 57719694 | 34304226 | 49383438 | 44200158 | 59388214 | 33112596 | 32090512 | 23535516 | 25739658 | 28005316 | 36423618 | 45522150 | 46258442 | 34294670 | 37134062 | 31333646 | 4 | 8 | 2 | 9 | 4 | 9 | 0.671878294 | 0.661534212 | 0.896529644 | 0.882726882 |
| Q9D620 | Rab11fip1 | Rab11 family-interacting protein 1 | 125.37 | 6.2 | 70684 | 3 | 2 | 2 | 2 | 2 | 134348.48 | 99859.26 | 2 | 107080.02 | 2 | 2 | 149498.36 | 353637.44 | 2 | 562949.31 | 380481.5 | 2 | 2 | 234778.64 | 2 | 77814.74 | 3 | 10 | 3.5 | 12.5 | 3.5 | 12.5 | 10.76664564 | 6.990101973 | 2.326641633 | 1.510541241 |
| P35283 | Rab12 | Ras-related protein Rab-12 | 225.03 | 16.46 | 27329 | 4 | 1789760.25 | 1823857.5 | 1956910.12 | 2283628.5 | 1111306.12 | 2066268.75 | 1951885.5 | 2076195.25 | 2472990 | 1994667.75 | 2206791 | 923846.56 | 2041783.12 | 2211209.75 | 2148102 | 1174205 | 2323449.5 | 2159298.5 | 2705226.5 | 1700241.5 | 9 | 9 | 11 | 12 | 11 | 12 | 1.063161264 | 0.902454638 | 1.122353812 | 0.952699687 |
| Q91V41 | Rab14 | Ras-related protein Rab-14 | 819.96 | 55.35 | 23897 | 12 | 33951214 | 29001582 | 30542042 | 29548914 | 25610644 | 33451598 | 30019318 | 31561936 | 32439342 | 32963508 | 29221190 | 23350526 | 29704788 | 27778422 | 30628138 | 35356142 | 39071226 | 35948770 | 39369298 | 35218194 | 10 | 0 | 1 | 0 | 10 | 0 | 0.946376749 | 0.876881282 | 1.244252676 | 1.152883228 |
| P35293 | Rab18 | Ras-related protein Rab-18 | 372.69 | 37.38 | 23035 | 6 | 2290905 | 1535950.75 | 4180238 | 3269490.75 | 1572070.5 | 3549479.25 | 2586163.25 | 3838429.75 | 2067514.5 | 6012491 | 6968783 | 5984985 | 8249985 | 5189242 | 8127000.5 | 3323547 | 5128111 | 5226470 | 6120125.5 | 5291493.5 | 0 | 1 | 2 | 6 | 2 | 6 | 2.686662184 | 1.912033169 | 1.952713883 | 1.389699731 |
| P62821 | Rab1A | Ras-related protein Rab-1A | 1229.34 | 73.17 | 22678 | 19 | 24975582 | 27055422 | 23795682 | 18739630 | 25150482 | 16386293 | 28820616 | 26058816 | 28173304 | 20210276 | 18927840 | 27581530 | 21203534 | 15690180 | 21451458 | 24476954 | 20180678 | 12369655 | 18947442 | 21976260 | 8 | 5 | 8 | 6 | 8 | 6 | 0.875854882 | 0.876348943 | 0.818189182 | 0.818650714 |
| Q9D1G1 | Rab1b | Ras-related protein Rab-1B | 1120.51 | 71.14 | 22187 | 16 | 11542018 | 13488312 | 13843508 | 9740438 | 14180080 | 9676611 | 12323594 | 7487611 | 14972215 | 10310312 | 9826411 | 10031225 | 9421074 | 8395501 | 8038670.5 | 11057081 | 9378104 | 9953696 | 11551486 | 9988770 | 2 | 5 | 7 | 11 | 7 | 11 | 0.727977551 | 0.834628359 | 0.826971408 | 0.948125101 |
| P35295 | Rab20 | Ras-related protein Rab-20 | 42.54 | 4.72 | 25989 | 1 | 297932.09 | 242537.81 | 399136.53 | 340009.69 | 313905.53 | 313369.16 | 245208.33 | 474409.16 | 486812.16 | 503799.5 | 2 | 544268.19 | 477246.12 | 751885.44 | 663417.25 | 309432.06 | 425166.47 | 409183.69 | 502777.31 | 388604.69 | 5 | 4 | 7 | 11 | 7 | 11 | 1.529203572 | 1.204200946 | 1.277148773 | 1.005715517 |
| P35282 | Rab21 | Ras-related protein Rab-21 | 277.01 | 20.72 | 24106 | 5 | 1653953.62 | 10204515 | 1625158.88 | 2597720.5 | 2263994.75 | 2108031 | 9668230 | 2797307.75 | 1651265.25 | 3435621.75 | 3896422.75 | 3914474 | 4583716.5 | 4011467 | 4648642 | 2141869.5 | 1249864.12 | 2333199.5 | 2052885.38 | 2478415.75 | 5 | 10 | 5 | 7 | 5 | 10 | 1.147687592 | 1.070917303 | 0.559064739 | 0.521668184 |
| P35285 | Rab22a | Ras-related protein Rab-22A | 168.91 | 18.04 | 21802 | 2 | 1003017.25 | 757222.88 | 544402.94 | 598353.81 | 657933.44 | 472076.69 | 815376.81 | 629339.19 | 733749.81 | 510489.81 | 607803.31 | 1074431.5 | 939508.25 | 640405.38 | 1009263.94 | 727056.75 | 562197.44 | 254603.67 | 725767.81 | 577850 | 7 | 8 | 5 | 10 | 7 | 10 | 1.199521472 | 1.351271345 | 0.799643749 | 0.90080562 |
| P53994 | Rab2a | Ras-related protein Rab-2A | 692.25 | 59.91 | 23548 | 11 | 11793756 | 9023292 | 10855704 | 11572804 | 11140478 | 12519998 | 10695813 | 10887173 | 11905967 | 11003566 | 11405360 | 12944691 | 16387132 | 14624554 | 9590740 | 10829392 | 13166058 | 15669616 | 12576531 | 12933240 | 6 | 4 | 7 | 4 | 7 | 4 | 1.194285963 | 1.139266961 | 1.198374513 | 1.143167157 |
| Q921E2 | Rab31 | Ras-related protein Rab-31 | 229.49 | 19.07 | 21331 | 3 | 3397679.5 | 3834778 | 3536063.5 | 4649832 | 4444728.5 | 3590516.5 | 3465393.75 | 3326834.5 | 3685449 | 4558727 | 2153244 | 1645682.62 | 2911668.75 | 2702058.25 | 2978499.25 | 3561139 | 4083012 | 4079562 | 4295465.5 | 4601897 | 0 | 10 | 0 | 6 | 0 | 10 | 0.623828325 | 0.665228195 | 1.038160947 | 1.107057671 |
| Q9CZE3 | Rab32 | Ras-related protein Rab-32 | 97.79 | 9.87 | 25068 | 2 | 872238.69 | 689550.25 | 2 | 1024873.38 | 1232055.38 | 872181.75 | 918080.88 | 1099709.25 | 2 | 903327.19 | 824089.31 | 923304.44 | 1133248.5 | 1194591.5 | 976866.69 | 916863.62 | 1234585.5 | 1151276.5 | 2 | 709841.19 | 9 | 10.5 | 6 | 10.5 | 9 | 10.5 | 1.322982789 | 1.331847999 | 1.050762854 | 1.057803938 |
| O35963 | Rab33b | Ras-related protein Rab-33B | 197.4 | 10.48 | 25767 | 3 | 1007412.69 | 1013447.44 | 609945.25 | 988597.5 | 1108526.25 | 905141.88 | 764386.06 | 2 | 804077 | 1190426.5 | 2 | 483798.62 | 1014474.44 | 1318673 | 423266.09 | 761673.75 | 946073 | 1028292.25 | 1185749.12 | 1187936.62 | 9 | 9 | 11.5 | 8 | 11.5 | 9 | 0.685334755 | 0.884329852 | 1.080753243 | 1.394562802 |
| Q5U316 | Rab35 | Ras-related protein Rab-35 | 501.61 | 41.79 | 23025 | 8 | 5412203.5 | 5516776 | 5562216.5 | 4508181.5 | 5592273 | 4556175 | 5965445 | 3190366.5 | 4781147.5 | 4251460.5 | 3433535 | 4851491 | 4064424 | 3977654 | 5022846 | 5348817 | 1380366.75 | 4207829.5 | 4296479 | 4635896 | 2 | 2 | 11 | 10 | 11 | 10 | 0.802881717 | 0.938682376 | 0.747204024 | 0.873587271 |
| P35276 | Rab3d | Ras-related protein Rab-3D | 432.08 | 29.68 | 24416 | 6 | 314832.25 | 2 | 552400.75 | 525182.12 | 381955.47 | 581652.12 | 438051.59 | 454592.19 | 405942.78 | 600732.19 | 378225.69 | 508389.56 | 497607.97 | 444713.91 | 534276.44 | 389031.81 | 370289.84 | 513806.59 | 451590.16 | 518670.16 | 10 | 11 | 11 | 8 | 11 | 11 | 1.331858699 | 0.952535799 | 1.264327781 | 0.90423817 |
| Q80UJ7 | Rab3gap1 | Rab3 GTPase-activating protein catalytic subunit | 207.3 | 6.63 | 110198 | 4 | 2592334.75 | 2188668.75 | 2181525 | 2489567 | 2231633.25 | 3805708.5 | 2556747.5 | 1558627.75 | 2441962 | 1073593 | 1824209.25 | 1040272.19 | 2131923 | 2250571.75 | 1753658.38 | 2212876.5 | 4354276 | 2864533.5 | 2057078.62 | 3291477 | 3 | 8 | 8 | 8 | 8 | 8 | 0.770356345 | 0.786999989 | 1.265027795 | 1.292358878 |
| Q8BMG7 | Rab3gap2 | Rab3 GTPase-activating protein non-catalytic subunit | 552.41 | 7.98 | 152535 | 10 | 7582298 | 7592893.5 | 8719330 | 8808677 | 7446921 | 9575254 | 7838345 | 9391872 | 9333258 | 10039412 | 9122802 | 8720006 | 9865941 | 9699339 | 9652968 | 7132629.5 | 6275008.5 | 10890000 | 11204505 | 10382941 | 1 | 10 | 11 | 10 | 11 | 10 | 1.17212742 | 1.019119761 | 1.142838043 | 0.993653772 |
| Q53B90 | Rab43 | Ras-related protein Rab-43 | 171.58 | 11.9 | 23229 | 2 | 2 | 205771.19 | 271396.97 | 192541.28 | 273206.75 | 73890.59 | 554711 | 227007.59 | 291913.16 | 178219.77 | 64896.35 | 2 | 213653 | 580239.94 | 2 | 186104.06 | 2 | 255346.5 | 2 | 263328.34 | 10 | 8 | 7 | 8 | 10 | 8 | 0.910782398 | 0.647783067 | 0.74744862 | 0.531613875 |
| P51146 | Rab4b | Ras-related protein Rab-4B | 366.45 | 34.27 | 23629 | 6 | 783016.06 | 945966.06 | 988573.31 | 754866.06 | 929258.12 | 283613.88 | 2 | 1109758.25 | 916155.75 | 839536.38 | 770456.62 | 1526323.12 | 1641301.88 | 1006182.06 | 1468072.25 | 848250.5 | 1032498.75 | 914041.5 | 555608.62 | 769018.88 | 4 | 10 | 4 | 11 | 4 | 11 | 1.456792974 | 2.036265801 | 0.93587417 | 1.308139591 |
| Q9CQD1 | Rab5a | Ras-related protein Rab-5A | 543.41 | 59.07 | 23599 | 9 | 5185264.5 | 4166141.5 | 3922643 | 5023185 | 3739090.75 | 4991647.5 | 3796136.75 | 2842799.5 | 3787580.5 | 5418166 | 7975968 | 6574168 | 6063421.5 | 6814707 | 7925080.5 | 3464983.75 | 6135598 | 5481009.5 | 4676521.5 | 5150700.5 | 0 | 8 | 0 | 7 | 0 | 8 | 1.604321292 | 1.696716484 | 1.130352431 | 1.195451068 |
| P61021 | Rab5b | Ras-related protein Rab-5B | 417.59 | 36.74 | 23707 | 6 | 6130193 | 5116326.5 | 4333350.5 | 5980961 | 4430825 | 4854616 | 4575215.5 | 4598084 | 5220267 | 5485281 | 3643185 | 3485026 | 4211917.5 | 4415592 | 3747807 | 4937451.5 | 6101923.5 | 5376425 | 5814321 | 4306203.5 | 1 | 12 | 0 | 8 | 1 | 12 | 0.750376486 | 0.788548175 | 1.020955514 | 1.072891571 |
| P35278 | Rab5c | Ras-related protein Rab-5C | 657.32 | 58.8 | 23413 | 9 | 14647106 | 14468749 | 14864166 | 16169928 | 14013728 | 14822537 | 11434511 | 12849464 | 13491496 | 15774730 | 20073904 | 12238395 | 17516902 | 15808689 | 17567568 | 12882463 | 18280846 | 15838464 | 16233360 | 15082586 | 6 | 7 | 4 | 4 | 6 | 7 | 1.121916568 | 1.216939096 | 1.056011813 | 1.145452432 |
| P35279 | Rab6a | Ras-related protein Rab-6A | 622.61 | 54.33 | 23590 | 10 | 8421500 | 6369542 | 6314879 | 8947371 | 8544034 | 10626456 | 8107574.5 | 7381280.5 | 9346704 | 9507824 | 14078669 | 9253778 | 13666823 | 12652200 | 10621708 | 8864234 | 12937435 | 10966666 | 10985799 | 10797669 | 0 | 1 | 4 | 3 | 4 | 3 | 1.561589474 | 1.340302286 | 1.413357055 | 1.213075346 |
| P09527 | Rab7a | Ras-related protein Rab-7a | 1346.7 | 79.23 | 23504 | 20 | 142025346 | 111623994 | 105559346 | 121953130 | 109165570 | 129989386 | 120343162 | 113408250 | 118454506 | 125129354 | 106986530 | 99218930 | 116968714 | 106786330 | 116374722 | 147085586 | 160508914 | 126807570 | 125795186 | 126541226 | 8 | 3 | 2 | 3 | 8 | 3 | 0.925478368 | 0.899576888 | 1.163318014 | 1.130760085 |
| Q8VEA8 | Rab7b | Ras-related protein Rab-7b | 148.53 | 18.09 | 22502 | 3 | 894017.31 | 526404.19 | 311083.16 | 827742.75 | 916846.94 | 2054835.75 | 653724.88 | 669418.62 | 2228475 | 2299131.25 | 1999164.75 | 1867297.12 | 2707813.5 | 319382.72 | 2903718 | 1006106.75 | 1140042.75 | 2373899.5 | 499680.31 | 1156626 | 4 | 4 | 11 | 11 | 11 | 11 | 2.818501198 | 1.239297974 | 1.77680888 | 0.781264754 |
| P35280 | Rab8a | Ras-related protein Rab-8A | 850.43 | 58.94 | 23668 | 14 | 3282443.5 | 2611452.25 | 3475116.5 | 3077307.75 | 2988160.75 | 2977015.25 | 2784492.25 | 3350539.75 | 3111124.5 | 2691150.75 | 3710744.5 | 3307350 | 3681922.75 | 3228539.5 | 2212151.75 | 3300996 | 3450660.75 | 4052820.25 | 3423931.5 | 2915847.75 | 8 | 7 | 7 | 4 | 8 | 7 | 1.045756496 | 1.082228744 | 1.110776354 | 1.149516262 |
| P61028 | Rab8b | Ras-related protein Rab-8B | 1037.28 | 66.67 | 23603 | 17 | 21683084 | 23391298 | 16911848 | 24809490 | 23378730 | 22275992 | 20404370 | 23634962 | 24434238 | 24865848 | 7470786 | 12518126 | 16382096 | 8425282 | 15693077 | 23650304 | 24370948 | 22649418 | 25149214 | 25766090 | 0 | 5 | 0 | 7 | 0 | 7 | 0.549032621 | 0.523194676 | 1.103576864 | 1.051641593 |
| Q99P75 | Rab9a | Ras-related protein Rab-9A | 168.64 | 18.91 | 22896 | 3 | 695159.56 | 553067.88 | 521908.62 | 527343.88 | 598694.69 | 717365.12 | 712913.31 | 418755.31 | 590447.75 | 600672.5 | 1004375.25 | 789936.06 | 956561.12 | 334041.19 | 952151.12 | 846064.62 | 131960.19 | 820444.38 | 742883.31 | 637669.56 | 5 | 6 | 5 | 7 | 5 | 7 | 1.393930013 | 1.327914557 | 1.097662422 | 1.045677972 |
| O35550 | Rabep1 | Rab GTPase-binding effector protein 1 | 42.4 | 2.2 | 99428 | 1 | 238820.69 | 135367.34 | 411909.31 | 182074.67 | 514256.62 | 269755.94 | 163018.92 | 517876.34 | 174357.36 | 583180.38 | 521927.53 | 611093.38 | 468434.53 | 156939.8 | 369405.84 | 285275.66 | 2 | 598910.81 | 791943.81 | 508594.25 | 7 | 8 | 10 | 9 | 10 | 9 | 1.435348075 | 1.245647381 | 1.473748203 | 1.278972413 |
| A2AWA9 | Rabgap1 | Rab GTPase-activating protein 1 | 51.3 | 0.94 | 120798 | 1 | 2 | 306175.25 | 611046.19 | 601011.31 | 2 | 2 | 476058.47 | 468231.59 | 439527.22 | 474253.19 | 2 | 2 | 465653.16 | 39412.98 | 2 | 2 | 2 | 2 | 1025220.12 | 605461 | 8 | 12 | 4.5 | 11.5 | 8 | 12 | 0.332670211 | 0.271825856 | 1.074066426 | 0.877622992 |
| Q9JM13 | Rabgef1 | Rab5 GDP/GTP exchange factor | 39.75 | 1.43 | 56869 | 1 | 105332.98 | 280560.41 | 224280.27 | 2 | 579116.69 | 2 | 266132.72 | 399723.75 | 588708.88 | 133151.39 | 2 | 2 | 2 | 2 | 2 | 250354 | 2 | 2 | 2 | 2 | 2.5 | 5 | 2.5 | 4 | 2.5 | 5 | 8.408361E-06 | 7.206071E-06 | 0.21051342 | 0.180412639 |
| Q9JHK4 | Rabggta | Geranylgeranyl transferase type-2 subunit alpha | 167.85 | 9.17 | 64989 | 3 | 2206910 | 2030088.88 | 2207134.75 | 2426978 | 1919438.88 | 2074360 | 2050275.75 | 2379156.75 | 2684473.25 | 2563966 | 2684985 | 2534151.5 | 2650485 | 2032869.62 | 2142777.5 | 2301784.5 | 2857025.25 | 3109935 | 3660910.5 | 2415502.5 | 6 | 2 | 11 | 5 | 11 | 5 | 1.116279342 | 1.024934572 | 1.329418526 | 1.220632647 |
| P63001 | Rac1 | Ras-related C3 botulinum toxin substrate 1 | 590.61 | 52.6 | 21450 | 9 | 13113981 | 10963706 | 10547920 | 13947131 | 9313760 | 12354465 | 11564474 | 10449930 | 10728226 | 10724269 | 10238501 | 9267045 | 10156602 | 9058434 | 8874276 | 10864550 | 11911400 | 11017179 | 11705663 | 9874280 | 2 | 12 | 0 | 11 | 2 | 12 | 0.822210008 | 0.852628001 | 0.956580099 | 0.991969168 |
| Q05144 | Rac2 | Ras-related C3 botulinum toxin substrate 2 | 654.62 | 59.9 | 21441 | 10 | 76165706 | 68310834 | 77909298 | 94352282 | 74845786 | 82883874 | 84958714 | 85086098 | 87872690 | 92522450 | 93844834 | 77540514 | 99035434 | 75871666 | 80632674 | 77043906 | 83716146 | 95391546 | 64008558 | 81309266 | 7 | 9 | 10 | 6 | 10 | 9 | 1.090251963 | 0.985233436 | 1.02524495 | 0.926488224 |
| Q9WVM1 | Racgap1 | Rac GTPase-activating protein 1 | 82.63 | 3.34 | 70158 | 2 | 937990.19 | 794472.12 | 2 | 656987.25 | 903442.88 | 874501.12 | 2 | 939008.69 | 97342.98 | 2 | 2 | 2 | 2 | 2 | 2 | 1075373.5 | 2 | 2 | 2 | 2 | 2.5 | 7 | 5 | 9 | 5 | 9 | 3.036842E-06 | 5.233255E-06 | 0.326576366 | 0.562774513 |
| P63245 | Rack1 | Receptor of activated protein C kinase 1 | 1253.07 | 78.86 | 35077 | 20 | 115807394 | 121646634 | 132533306 | 101612298 | 128778722 | 85001386 | 123900170 | 126613370 | 125019570 | 112594514 | 115728258 | 172612978 | 122261890 | 101623338 | 143378018 | 134605298 | 72671202 | 102911314 | 86770730 | 96910114 | 10 | 6 | 10 | 8 | 10 | 8 | 1.091985542 | 1.143903852 | 0.822595709 | 0.861705915 |
| Q9QXK2 | Rad18 | E3 ubiquitin-protein ligase RAD18 | 44.77 | 4.72 | 57412 | 1 | 2 | 2 | 178049.52 | 2 | 269390.81 | 2 | 2 | 2 | 213598.83 | 2 | 403497.66 | 284273.66 | 2 | 319732.03 | 310895.22 | 2 | 2 | 2 | 518815 | 375100.66 | 3.5 | 10.5 | 3 | 9 | 3.5 | 10.5 | 2.946499908 | 6.172089956 | 1.997829907 | 4.184892683 |
| Q61550 | Rad21 | Double-strand-break repair protein rad21 homolog | 134.17 | 3.31 | 72083 | 2 | 961052.38 | 752739.94 | 1009241.69 | 1341840 | 836929.19 | 1410942.75 | 1269813.88 | 1386463.75 | 1052901.38 | 837816.06 | 223998.83 | 269190.53 | 214231.56 | 102923.28 | 150253.69 | 1244446.12 | 2158829.5 | 1088005.62 | 1444630.38 | 749445.25 | 0 | 7 | 0 | 11 | 0 | 11 | 0.195968269 | 0.161229929 | 1.363856646 | 1.122092421 |
| P54726 | Rad23a | UV excision repair protein RAD23 homolog A | 95.14 | 8.26 | 39706 | 1 | 336630.66 | 198972.77 | 280384.28 | 267828.47 | 499901.44 | 243656.89 | 233451.12 | 221963.41 | 206715.16 | 169940.16 | 2 | 2 | 2 | 2 | 92826.03 | 2 | 258244.94 | 216810.12 | 524221.03 | 2 | 0 | 7 | 0 | 12 | 0 | 12 | 0.058617792 | 0.086298896 | 0.630971126 | 0.92893488 |
| P54728 | Rad23b | UV excision repair protein RAD23 homolog B | 116.1 | 9.62 | 43513 | 2 | 707063.06 | 585838.12 | 701974.5 | 648744.56 | 472769.34 | 959283.25 | 619632.31 | 558480.69 | 643650.19 | 1043111.19 | 936187.31 | 725856.06 | 685664.38 | 2 | 538974.5 | 857121.75 | 1446533.5 | 691996.56 | 1382123.5 | 576679.5 | 11 | 6 | 9 | 8 | 11 | 8 | 0.926291202 | 0.754854933 | 1.589805986 | 1.295567623 |
| P70388 | Rad50 | DNA repair protein RAD50 | 315.27 | 5.26 | 153488 | 6 | 7319728 | 3714137.25 | 3224942 | 4863187 | 7808362 | 7030018 | 5683403.5 | 3695340.75 | 6651536 | 3554266.5 | 5113723 | 4189117.5 | 7124342 | 5445181.5 | 5077151.5 | 4055067.5 | 6780610 | 4951042 | 3723984.5 | 4682427 | 11 | 12 | 12 | 12 | 12 | 12 | 1.000711437 | 1.012585242 | 0.898359115 | 0.909018473 |
| Q3SWS8 | Rae1 | mRNA export factor | 92.37 | 5.98 | 40919 | 2 | 1258850.25 | 1348194.38 | 1323115.75 | 1741920.38 | 1404101.88 | 1469847.88 | 1315198.12 | 2 | 1135655.5 | 1358354.12 | 2055619.25 | 1147067.12 | 1092064.38 | 1359044.12 | 1196235.88 | 1464390.25 | 1639245.88 | 1323084.75 | 1559758.38 | 1534997.75 | 8 | 8 | 11 | 3 | 11 | 8 | 0.968040411 | 1.297585903 | 1.062928615 | 1.424776457 |
| O08602 | Raet1a | Retinoic acid early-inducible protein 1-alpha | 40.04 | 4.74 | 28616 | 1 | 560110.94 | 445150.34 | 622163.81 | 2 | 2 | 583702.81 | 573360.62 | 446167.75 | 2 | 2 | 429832.34 | 509150.56 | 642781.25 | 357791.03 | 2 | 2 | 2 | 2 | 2 | 2 | 12 | 5 | 12 | 5 | 12 | 5 | 1.191792129 | 1.209777083 | 6.144661E-06 | 6.237388E-06 |
| P11345 | Raf1 | RAF proto-oncogene serine/threonine-protein kinase | 112.48 | 3.55 | 72928 | 2 | 1107744.25 | 1033716.19 | 792661.06 | 1033771.12 | 677559 | 1028159.25 | 753688 | 958455.44 | 874359.38 | 989522.75 | 530784.38 | 453395.84 | 2 | 537213.94 | 705361.25 | 1129470 | 1261313.5 | 916124.31 | 1063250.38 | 1172414.25 | 1 | 4 | 0 | 3 | 1 | 4 | 0.479341427 | 0.483637712 | 1.193118106 | 1.203811892 |
| P36860 | Ralb | Ras-related protein Ral-B | 394.11 | 26.21 | 23317 | 6 | 2876426.5 | 3106311.5 | 1831598.62 | 3407573.75 | 2503485.25 | 4779119 | 2815179.5 | 1899333.5 | 2165545.75 | 3367487.5 | 3233460 | 2144081.5 | 2520642 | 2615732 | 2390245.5 | 1745448.88 | 1785921 | 2416476.75 | 2949739 | 2376625.75 | 10 | 5 | 10 | 7 | 10 | 7 | 0.94016678 | 0.858750813 | 0.821412489 | 0.750280331 |
| A3KGS3 | Ralgapa2 | Ral GTPase-activating protein subunit alpha-2 | 195.33 | 2.46 | 210288 | 4 | 1117176.25 | 1001787.81 | 682387.88 | 964930.25 | 1767191.12 | 960193.81 | 1121670.25 | 1012967.25 | 1503715.25 | 1026786.5 | 349743.91 | 231863.47 | 700349.62 | 2 | 102621.63 | 1106796.38 | 963941.5 | 595607.38 | 318182.34 | 723244.88 | 1 | 5 | 0 | 4 | 1 | 5 | 0.250219085 | 0.246133094 | 0.670062413 | 0.659120525 |
| Q64012 | Raly | RNA-binding protein Raly | 778.64 | 36.54 | 33188 | 14 | 34756134 | 36133214 | 31583078 | 23312906 | 26017706 | 30593746 | 36845214 | 24730608 | 25359786 | 25242710 | 25575100 | 27553204 | 18217910 | 19723284 | 19853994 | 31012242 | 25285730 | 20988366 | 19162474 | 20849282 | 3 | 3 | 6 | 6 | 6 | 6 | 0.730706667 | 0.776927145 | 0.772699253 | 0.821575949 |
| P62827 | Ran | GTP-binding nuclear protein Ran | 793.22 | 58.33 | 24423 | 13 | 38566218 | 42194822 | 56018026 | 45926046 | 52256634 | 35639902 | 40762794 | 39750642 | 50228138 | 47857074 | 71477210 | 66251958 | 60614574 | 54259090 | 47922050 | 47628582 | 50911470 | 58134030 | 36712922 | 52233958 | 3 | 11 | 1 | 6 | 3 | 11 | 1.279037491 | 1.402758196 | 1.045365751 | 1.146483497 |
| P34022 | Ranbp1 | Ran-specific GTPase-activating protein | 305.31 | 20.69 | 23596 | 5 | 2101473.75 | 7794823.5 | 7079320 | 7149089 | 4844706.5 | 7305336 | 4976903 | 7877269 | 6209366.5 | 10820165 | 15888141 | 16604723 | 14984134 | 16294051 | 12675964 | 6431241.5 | 6141592.5 | 8219074 | 8593937 | 8272734 | 0 | 6 | 0 | 10 | 0 | 10 | 2.638887217 | 2.055632897 | 1.299942782 | 1.01262575 |
| Q6VN19 | Ranbp10 | Ran-binding protein 10 | 116.63 | 3.87 | 67188 | 2 | 2 | 2 | 2 | 2 | 2 | 2 | 2 | 732948.19 | 2 | 2 | 774862 | 744891.94 | 919803.19 | 2 | 846279.94 | 626096 | 2 | 2 | 555923.19 | 2 | 2.5 | 7.5 | 3 | 11 | 3 | 11 | 328583.907 | 4.482995184 | 118202.519 | 1.612681912 |
| Q9ERU9 | Ranbp2 | E3 SUMO-protein ligase RanBP2 | 1746.88 | 13.07 | 341121 | 29 | 6661445.5 | 7327400 | 9076059 | 8100810.5 | 8197001 | 6710492 | 6668599 | 8071434 | 6571912.5 | 5776262 | 7876919.5 | 6777831 | 7058007.5 | 5716218 | 7712835 | 6909958 | 4371010 | 7116499 | 6581919 | 6898456 | 6 | 3 | 9 | 11 | 9 | 11 | 0.892768959 | 1.039738556 | 0.80984864 | 0.94316771 |
| Q9CT10 | Ranbp3 | Ran-binding protein 3 | 405.67 | 14.87 | 52573 | 6 | 6303359 | 7377877 | 7607623 | 6568408 | 5222802.5 | 6576528 | 6867960 | 4077848 | 6318379.5 | 7655495 | 5013996.5 | 6650734 | 5492839 | 5349747 | 6145224 | 6577849 | 5309669 | 7059426.5 | 6159538 | 5988457 | 6 | 9 | 7 | 10 | 7 | 10 | 0.866157204 | 0.9097139 | 0.93999015 | 0.987259705 |
| P46061 | Rangap1 | Ran GTPase-activating protein 1 | 1294.67 | 39.39 | 63531 | 21 | 37230578 | 42114390 | 34928138 | 35216194 | 38420434 | 37296586 | 37738882 | 38428122 | 37520322 | 31811338 | 31271288 | 32811794 | 25313130 | 28053954 | 26883960 | 37733762 | 32375484 | 29420776 | 33397182 | 32738398 | 0 | 3 | 1 | 6 | 1 | 6 | 0.768103509 | 0.789594511 | 0.881623312 | 0.906290519 |
| P62835 | Rap1a | Ras-related protein Rap-1A | 1044.21 | 83.15 | 20987 | 15 | 630975.31 | 2033489.25 | 1526672.75 | 1517399 | 570172.69 | 1802396.25 | 1420271.75 | 711507.94 | 586019.31 | 1702379.75 | 2413593 | 2814916.5 | 1888930.5 | 1325040 | 1969310.25 | 1005176.88 | 793713.81 | 457086.75 | 510436.09 | 454884.34 | 5 | 4 | 3 | 4 | 5 | 4 | 1.65826928 | 1.673228567 | 0.51305099 | 0.517679236 |
| Q99JI6 | Rap1b | Ras-related protein Rap-1b | 1218.61 | 83.15 | 20825 | 16 | 13629994 | 12422928 | 10544779 | 13016406 | 10829974 | 14608352 | 12932520 | 9714775 | 12313395 | 11269606 | 14003465 | 17803998 | 15466155 | 9798287 | 14401966 | 11017321 | 8555235 | 7833452.5 | 6050888 | 7311885 | 5 | 2 | 6 | 1 | 6 | 2 | 1.182479241 | 1.174810311 | 0.674487573 | 0.670113207 |
| P61226 | Rap2b | Ras-related protein Rap-2b | 494.63 | 54.1 | 20504 | 7 | 3106385 | 3161117.5 | 1269210.25 | 1730480.12 | 1295620.12 | 2300642.5 | 2773323.75 | 1836353 | 2285430.75 | 1333396 | 1128291 | 2729920 | 2345406.75 | 1030096.38 | 2412573 | 2773929 | 1332114.75 | 1863858.25 | 1765864.25 | 1291717.38 | 9 | 12 | 12 | 8 | 12 | 12 | 0.913230892 | 0.916150952 | 0.854647681 | 0.857380421 |
| Q8BU31 | Rap2c | Ras-related protein Rap-2c | 477.11 | 54.1 | 20745 | 7 | 940171.94 | 616582.12 | 368383.62 | 697397.06 | 330723.78 | 764207.19 | 440332.53 | 552841.69 | 772406.5 | 406905.28 | 419181 | 745389.75 | 657573.75 | 2 | 682328.75 | 191178.83 | 498686.81 | 354620.62 | 590930.75 | 311583.22 | 12 | 5 | 10 | 5 | 12 | 5 | 0.848037933 | 0.852821554 | 0.659271857 | 0.662990685 |
| F1M386 | Rapgef2 | Rap guanine nucleotide exchange factor 2 | 50.13 | 0.94 | 166388 | 1 | 227724.2 | 320922.09 | 226344 | 286096.66 | 2 | 359699.53 | 342798.25 | 243542.75 | 2 | 305996.47 | 136640.62 | 2 | 2 | 2 | 2 | 2 | 498508.75 | 291287.78 | 300880.03 | 2 | 3 | 11 | 3 | 10 | 3 | 11 | 0.128781494 | 0.109140865 | 1.027887964 | 0.871123471 |
| Q9D0I9 | Rars | Arginine--tRNA ligase, cytoplasmic | 1695.8 | 44.85 | 75674 | 29 | 28191718 | 28314748 | 20498282 | 24726384 | 26574872 | 27343870 | 25145194 | 22467102 | 27191068 | 22871632 | 40809410 | 37054918 | 39739254 | 24907196 | 30945126 | 37023322 | 36738834 | 28191300 | 31265586 | 28915310 | 3 | 2 | 3 | 0 | 3 | 2 | 1.35189234 | 1.387437829 | 1.263653663 | 1.296879081 |
| Q3U186 | Rars2 | Probable arginine--tRNA ligase, mitochondrial | 271.78 | 7.09 | 65336 | 4 | 1176231.62 | 1167882.88 | 841843.25 | 1164175.25 | 1289613.38 | 1103845.38 | 1499025.25 | 364759.34 | 1336745.25 | 728786.62 | 1002466.38 | 446584.69 | 961098 | 592080.56 | 1165748.5 | 1399918.25 | 505706.03 | 1260531 | 1169695 | 1057677.5 | 4 | 12 | 9 | 12 | 9 | 12 | 0.739036448 | 0.82810334 | 0.956342257 | 1.071598322 |
| P50904 | Rasa1 | Ras GTPase-activating protein 1 | 129.57 | 2.41 | 115440 | 3 | 186861.61 | 2 | 142903.52 | 113062.61 | 416065.47 | 2 | 225916.41 | 2 | 305908.41 | 108011.97 | 2 | 2 | 2 | 2 | 2 | 2 | 2 | 2 | 239433.47 | 106627.7 | 2.5 | 6.5 | 5 | 9 | 5 | 9 | 1.164286E-05 | 1.562889E-05 | 0.402921295 | 0.540864502 |
| Q60790 | Rasa3 | Ras GTPase-activating protein 3 | 340.98 | 7.19 | 95987 | 7 | 5219907 | 4639854.5 | 6111745.5 | 5446853.5 | 5240098.5 | 4133152.25 | 4502272 | 4666450 | 4321257.5 | 4103102.5 | 2680542.5 | 2886384.5 | 3532714.25 | 2523634.75 | 2833600.25 | 4680458.5 | 2771065.5 | 3260402.25 | 2530082.75 | 3775586 | 0 | 1 | 0 | 5 | 0 | 5 | 0.542299772 | 0.665411046 | 0.638356291 | 0.783274027 |
| Q6PFQ7 | Rasa4 | Ras GTPase-activating protein 4 | 243.16 | 5.86 | 90060 | 5 | 3411020.5 | 3501443.5 | 221308.16 | 4615899 | 2 | 3818974.5 | 2691590.25 | 206985.19 | 3683763.5 | 2116614.25 | 1033460.31 | 3747171.25 | 4150538.5 | 1695130.62 | 4231861 | 3152882 | 1598275.5 | 2213604.5 | 160293.84 | 2053811.12 | 9 | 9 | 9 | 7 | 9 | 9 | 1.264559574 | 1.186950592 | 0.781201897 | 0.733257707 |
| Q8C2K5 | Rasal3 | RAS protein activator like-3 | 130.31 | 3.07 | 114782 | 3 | 4657640 | 4276998.5 | 5056808 | 5588317.5 | 4006845.25 | 4773707 | 4464023 | 4253322.5 | 4035253.5 | 5228095.5 | 2635268 | 2536821.5 | 3666655.5 | 2539957.5 | 3538142.5 | 4277430 | 4619481.5 | 3123532.5 | 4323948 | 4819902 | 0 | 9 | 0 | 11 | 0 | 11 | 0.632428546 | 0.655558662 | 0.897301252 | 0.930118685 |
| Q8CB96 | Rassf4 | Ras association domain-containing protein 4 | 133.97 | 11.8 | 36934 | 3 | 1225388.25 | 1299489.12 | 812363.75 | 1053161.25 | 946938.75 | 970067.25 | 1314631.75 | 1288030.62 | 1264276.12 | 1248885.5 | 1263175.25 | 1532279.25 | 757604.75 | 863517.25 | 1292491.5 | 1093483 | 1440507.88 | 1134544.88 | 1103146.75 | 1278099.25 | 11 | 7 | 11 | 11 | 11 | 11 | 1.069646453 | 0.938082489 | 1.133482313 | 0.99406669 |
| Q5XI28 | Raver1 | Ribonucleoprotein PTB-binding 1 | 121.06 | 3.61 | 79289 | 2 | 866208 | 931865.38 | 1522117 | 1021599.75 | 668701.12 | 516891.16 | 2 | 1070190.62 | 992400.5 | 866580.19 | 732992.38 | 828401 | 1301089.75 | 1349310 | 988206 | 595811.75 | 698089.56 | 1011431.5 | 670695.31 | 1242270.5 | 12 | 9 | 8 | 10 | 12 | 10 | 1.037822216 | 1.508967673 | 0.841893222 | 1.224091792 |
| Q60972 | Rbbp4 | Histone-binding protein RBBP4 | 446.59 | 12.71 | 47656 | 6 | 2009659 | 1864863.62 | 3444418.5 | 3549477.5 | 2774216 | 2703170.5 | 2480310.25 | 2407554.25 | 1268967.5 | 1973601.75 | 2226215 | 3084254.75 | 599091 | 2131011 | 993655.75 | 1342316 | 2111165.5 | 2 | 1405943 | 2336324 | 7 | 4 | 9 | 6 | 9 | 6 | 0.662205487 | 0.833907838 | 0.527445812 | 0.664206513 |
| Q8BX09 | Rbbp5 | Retinoblastoma-binding protein 5 | 126.17 | 5.58 | 59098 | 3 | 1684333.5 | 1009119 | 857399.69 | 916669.06 | 1612119.12 | 668782.81 | 1671422.75 | 1511831.5 | 1065759.38 | 775974.62 | 1150396.25 | 1199094.38 | 657353.94 | 1364966.88 | 1015951.69 | 527341.56 | 503551.06 | 649598.81 | 651264.75 | 1189235.5 | 12 | 3 | 11 | 3 | 12 | 3 | 0.886197672 | 0.946255668 | 0.57914473 | 0.618393617 |
| Q60973 | Rbbp7 | Histone-binding protein RBBP7 | 263.33 | 8.94 | 47790 | 4 | 731137.25 | 1148787 | 907747.88 | 832848 | 416173.34 | 899097.19 | 2 | 639404.31 | 490231.22 | 851908.19 | 1339490.75 | 1320345.12 | 1109718.62 | 980034.44 | 1047714.5 | 2 | 2 | 585846.62 | 671019.69 | 573915.56 | 3 | 3 | 0 | 8 | 3 | 8 | 1.436151512 | 2.012503323 | 0.453536015 | 0.635547663 |
| Q8C2Q3 | Rbm14 | RNA-binding protein 14 | 549.84 | 13.9 | 69449 | 9 | 5088647 | 4102328.5 | 4774362.5 | 4570536.5 | 3599181.5 | 4149221.5 | 4528677 | 3247756 | 4294441.5 | 5959583 | 13543489 | 11296799 | 10492866 | 13063353 | 12803233 | 4456884 | 6128627 | 6509943 | 5831572.5 | 4187065 | 0 | 6 | 0 | 6 | 0 | 6 | 2.764833303 | 2.759270772 | 1.224938916 | 1.222474478 |
| Q0VBL3 | Rbm15 | RNA-binding protein 15 | 263.52 | 6.76 | 105722 | 5 | 1441874.88 | 781518.69 | 856807 | 688502.19 | 1597246.5 | 1083244.5 | 2382436.25 | 963765.12 | 1348816.5 | 845748.12 | 1844185.38 | 1388112 | 1122265.88 | 704857.19 | 1034046.12 | 1133708.62 | 1525164.88 | 980768.69 | 3346380.75 | 992283.88 | 10 | 7 | 12 | 9 | 12 | 9 | 1.135580356 | 0.919905936 | 1.486839781 | 1.204452625 |
| Q8JZX4 | Rbm17 | Splicing factor 45 | 267.26 | 14.81 | 45304 | 6 | 2466122.25 | 1838096.25 | 2005115 | 2601102 | 2024824.88 | 3345537.5 | 2034896.25 | 2205144.5 | 2106821.75 | 2438935.25 | 3030911 | 2913391 | 2369567.75 | 2872693.25 | 2604453.5 | 2723989.5 | 2023503.88 | 2072477.75 | 1592729.25 | 2093667.12 | 2 | 12 | 6 | 6 | 6 | 12 | 1.261151177 | 1.136809446 | 0.960778906 | 0.866052028 |
| Q8R3C6 | Rbm19 | Probable RNA-binding protein 19 | 228.87 | 4.73 | 106083 | 3 | 718369.81 | 684633.38 | 867500.12 | 1119924.75 | 641335.25 | 848870.31 | 995414.5 | 1164909.25 | 909235.94 | 712165.12 | 1059758.88 | 1060689.88 | 637574.12 | 485055.69 | 475543.66 | 1040784 | 1035235.12 | 474866.38 | 546496.12 | 559654.62 | 8 | 8 | 8 | 8 | 8 | 8 | 0.922331482 | 0.803054928 | 0.907056283 | 0.789755128 |
| Q8BHS3 | Rbm22 | Pre-mRNA-splicing factor RBM22 | 53.59 | 1.67 | 46896 | 1 | 2 | 562019.12 | 330884.5 | 2 | 2 | 2 | 2 | 2 | 2 | 2 | 1408072.75 | 1103865.5 | 1157915.38 | 1343928.88 | 1474358.62 | 2 | 2 | 228606.14 | 2 | 2 | 0 | 9 | 0 | 10 | 0 | 10 | 7.266290994 | 648814.113 | 0.256032789 | 22861.414 |
| [illegible] | Rbm25 | RNA-binding protein 25 | [illegible] | [illegible] | [illegible] | [illegible] | [illegible] | [illegible] | [illegible] | [illegible] | [illegible] | [illegible] | [illegible] | [illegible] | [illegible] | [illegible] | [illegible] | [illegible] | [illegible] | [illegible] | [illegible] | [illegible] | [illegible] | [illegible] | [illegible] | [illegible] | [illegible] | [illegible] | [illegible] | [illegible] | [illegible] | [illegible] | [illegible] | [illegible] | [illegible] | [illegible] |
| [illegible] | Rbm28 | RNA-binding protein 28 | [illegible] | [illegible] | [illegible] | [illegible] | [illegible] | [illegible] | [illegible] | [illegible] | [illegible] | [illegible] | [illegible] | [illegible] | [illegible] | [illegible] | [illegible] | [illegible] | [illegible] | [illegible] | [illegible] | [illegible] | [illegible] | [illegible] | [illegible] | [illegible] | [illegible] | [illegible] | [illegible] | [illegible] | [illegible] | [illegible] | [illegible] | [illegible] | [illegible] | [illegible] |
| [illegible] | Rbm3 | RNA-binding protein 3 | [illegible] | [illegible] | [illegible] | [illegible] | [illegible] | [illegible] | [illegible] | [illegible] | [illegible] | [illegible] | [illegible] | [illegible] | [illegible] | [illegible] | [illegible] | [illegible] | [illegible] | [illegible] | [illegible] | [illegible] | [illegible] | [illegible] | [illegible] | [illegible] | [illegible] | [illegible] | [illegible] | [illegible] | [illegible] | [illegible] | [illegible] | [illegible] | [illegible] | [illegible] |
| [illegible] | Rbm39 | RNA-binding protein 39 | [illegible] | [illegible] | [illegible] | [illegible] | [illegible] | [illegible] | [illegible] | [illegible] | [illegible] | [illegible] | [illegible] | [illegible] | [illegible] | [illegible] | [illegible] | [illegible] | [illegible] | [illegible] | [illegible] | [illegible] | [illegible] | [illegible] | [illegible] | [illegible] | [illegible] | [illegible] | [illegible] | [illegible] | [illegible] | [illegible] | [illegible] | [illegible] | [illegible] | [illegible] |
| [illegible] | Rbm8a | RNA-binding protein 8A | [illegible] | [illegible] | [illegible] | [illegible] | [illegible] | [illegible] | [illegible] | [illegible] | [illegible] | [illegible] | [illegible] | [illegible] | [illegible] | [illegible] | [illegible] | [illegible] | [illegible] | [illegible] | [illegible] | [illegible] | [illegible] | [illegible] | [illegible] | [illegible] | [illegible] | [illegible] | [illegible] | [illegible] | [illegible] | [illegible] | [illegible] | [illegible] | [illegible] | [illegible] |
| [illegible] | Rbmxrt | RNA-binding motif protein, X chromosome retrogene-like | [illegible] | [illegible] | [illegible] | [illegible] | [illegible] | [illegible] | [illegible] | [illegible] | [illegible] | [illegible] | [illegible] | [illegible] | [illegible] | [illegible] | [illegible] | [illegible] | [illegible] | [illegible] | [illegible] | [illegible] | [illegible] | [illegible] | [illegible] | [illegible] | [illegible] | [illegible] | [illegible] | [illegible] | [illegible] | [illegible] | [illegible] | [illegible] | [illegible] | [illegible] |
| P31266 | Rbpj | Recombining binding protein suppressor of hairless | [illegible] | [illegible] | [illegible] | [illegible] | [illegible] | [illegible] | [illegible] | [illegible] | [illegible] | [illegible] | [illegible] | [illegible] | [illegible] | [illegible] | [illegible] | [illegible] | [illegible] | [illegible] | [illegible] | [illegible] | [illegible] | [illegible] | [illegible] | [illegible] | [illegible] | [illegible] | [illegible] | [illegible] | [illegible] | [illegible] | [illegible] | [illegible] | [illegible] | [illegible] |
| P62878 | Rbx1 | E3 ubiquitin-protein ligase RBX1 | [illegible] | [illegible] | [illegible] | [illegible] | [illegible] | [illegible] | [illegible] | [illegible] | [illegible] | [illegible] | [illegible] | [illegible] | [illegible] | [illegible] | [illegible] | [illegible] | [illegible] | [illegible] | [illegible] | [illegible] | [illegible] | [illegible] | [illegible] | [illegible] | [illegible] | [illegible] | [illegible] | [illegible] | [illegible] | [illegible] | [illegible] | [illegible] | [illegible] | [illegible] |
| [illegible] | Rcc1 | Regulator of chromosome condensation | [illegible] | [illegible] | [illegible] | [illegible] | [illegible] | [illegible] | [illegible] | [illegible] | [illegible] | [illegible] | [illegible] | [illegible] | [illegible] | [illegible] | [illegible] | [illegible] | [illegible] | [illegible] | [illegible] | [illegible] | [illegible] | [illegible] | [illegible] | [illegible] | [illegible] | [illegible] | [illegible] | [illegible] | [illegible] | [illegible] | [illegible] | [illegible] | [illegible] | [illegible] |
| [illegible] | Rcc2 | Protein RCC2 | [illegible] | [illegible] | [illegible] | [illegible] | [illegible] | [illegible] | [illegible] | [illegible] | [illegible] | [illegible] | [illegible] | [illegible] | [illegible] | [illegible] | [illegible] | [illegible] | [illegible] | [illegible] | [illegible] | [illegible] | [illegible] | [illegible] | [illegible] | [illegible] | [illegible] | [illegible] | [illegible] | [illegible] | [illegible] | [illegible] | [illegible] | [illegible] | [illegible] | [illegible] |

| | | | | | | | | | | | | | | | | | | | | | | | | | | | | | | | | | | | | |
|---|---|---|---|---|---|---|---|---|---|---|---|---|---|---|---|---|---|---|---|---|---|---|---|---|---|---|---|---|---|---|---|---|---|---|---|---|
| Q9JJT0 | Rcl1 | RNA 3'-terminal phosphate cyclase-like protein | 172.11 | 10.19 | 40841 | 3 | 1629813.5 | 1537623 | 1668742.5 | 538479.88 | 1671939.38 | 1393452.38 | 1412228.62 | 1802545.38 | 2130538.25 | 1047163.38 | 971140.56 | 1854286.38 | 1955794 | 750805.94 | 1513890.25 | 1928820.5 | 2 | 917900.94 | 365722.12 | 451820.22 | 12 | 6 | 11 | 4 | 12 | 6 | 0.999903339 | 0.90495534 | 0.520004922 | 0.470626722 |
| Q8BP92 | Rcn2 | Reticulocalbin-2 | 512.31 | 33.12 | 37271 | 8 | 2 | 542250.5 | 359442.06 | 218010.08 | 2 | 2 | 2 | 2 | 221502.11 | 2 | 2637103.5 | 1087060 | 1112063.25 | 1160075.62 | 249372.33 | 2 | 777871.5 | 2 | 2 | 2 | 2 | 9 | 0 | 12 | 2 | 12 | 5.577956294 | 28.19589002 | 0.694717234 | 3.511711046 |
| Q3UZA1 | Rcsd1 | CapZ-interacting protein | 185.78 | 13.59 | 44114 | 4 | 255294.98 | 861631.25 | 1039746.5 | 1168173.38 | 1769150.12 | 669385 | 262865.72 | 833844.12 | 1398014.62 | 687481.25 | 2 | 620932.25 | 318611.72 | 750197.19 | 479048.91 | 561647.62 | 457824.31 | 820336.5 | 651918.25 | 1353715.5 | 4 | 8 | 6 | 10 | 6 | 10 | 0.425754549 | 0.563089963 | 0.754896943 | 0.998403639 |
| Q9QYF1 | Rdh11 | Retinol dehydrogenase 11 | 470.53 | 26.9 | 35148 | 7 | 17384592 | 19125722 | 15907852 | 19009422 | 15508748 | 19923836 | 17336732 | 17727470 | 14947081 | 15625086 | 22761130 | 22865766 | 29008802 | 28993562 | 26978962 | 18373678 | 20837184 | 19972518 | 21413542 | 21018688 | 0 | 2 | 0 | 1 | 0 | 2 | 1.502343301 | 1.526506651 | 1.168850847 | 1.18765038 |
| Q8CEE7 | Rdh13 | Retinol dehydrogenase 13 | 90.18 | 5.99 | 36464 | 2 | 2573899.75 | 3381271.5 | 1008053.38 | 2823606 | 2248344.5 | 4006794 | 3420210 | 1552175.38 | 974165.88 | 2000315.75 | 2403962.5 | 2246080.5 | 2314497.5 | 2108593 | 1916437.25 | 3549590.25 | 4191457.5 | 2658973.75 | 3591912.75 | 3078943.5 | 7 | 3 | 11 | 6 | 11 | 6 | 0.913120967 | 0.919347699 | 1.4184154 | 1.428087825 |
| P26043 | Rdx | Radixin | 1179.02 | 29.16 | 68543 | 20 | 1153317.75 | 2116427 | 1200595.12 | 848674.38 | 1462502.75 | 1627727.88 | 1483415.75 | 1565809.75 | 2094775.62 | 1227480 | 2814882.5 | 2762675.75 | 2853505.5 | 2071505.88 | 3328681 | 1642778.12 | 2019690 | 1646469.75 | 1807440.12 | 1389065.5 | 1 | 6 | 1 | 8 | 1 | 8 | 2.039551126 | 1.729077291 | 1.254209565 | 1.063285569 |
| P15307 | Rel | Proto-oncogene c-Rel | 51.21 | 1.87 | 64960 | 1 | 592823.38 | 639449.75 | 567581 | 544314 | 423812.78 | 2 | 550020.5 | 514869.59 | 476349.28 | 777114.69 | 1084187.88 | 968475.38 | 1136052.5 | 694820.12 | 984168.19 | 2 | 533138.94 | 457044.5 | 439686.78 | 522171.94 | 0 | 4 | 1 | 8.5 | 1 | 8.5 | 1.758575737 | 2.099636097 | 0.70522313 | 0.841994978 |
| Q8VE91 | Retreg1 | Reticulophagy regulator 1 | 171.66 | 7.29 | 52967 | 3 | 1186824.12 | 1068873.12 | 1807634.62 | 1565975 | 1589422.75 | 1197228.5 | 1511456.5 | 2073828 | 1955992.5 | 2314447.5 | 3484747 | 3356419.25 | 2414921.5 | 2023530.62 | 3823275.5 | 1633857.38 | 1585548.75 | 2385658 | 2147367.75 | 2514557.25 | 0 | 3 | 2 | 7 | 2 | 7 | 2.092181684 | 1.668283694 | 1.42227091 | 1.134103881 |
| Q9CQV4 | Retreg3 | Reticulophagy regulator 3 | 65.12 | 3.22 | 51638 | 1 | 420080.84 | 338088.62 | 272827.5 | 263110.88 | 272936.69 | 415051.12 | 356811.31 | 212285.64 | 273873.41 | 225248.31 | 266299.38 | 325465.72 | 319155.28 | 2 | 247011.94 | 93062.52 | 521134 | 283961.69 | 371991.88 | 2 | 7 | 12 | 10 | 12 | 10 | 12 | 0.738928791 | 0.780663321 | 0.810539883 | 0.856318991 |
| Q64FW2 | Retsat | All-trans-retinol 13,14-reductase | 196.13 | 8.21 | 67334 | 3 | 1610648.25 | 925256.5 | 911101.06 | 1092285.75 | 1156480.12 | 811141.81 | 1155866.5 | 830872.88 | 957370.62 | 617292.62 | 1969051.5 | 1857133.25 | 1658812.12 | 1611660.25 | 1776039 | 1460272.5 | 1337133.75 | 961630.38 | 1033971.88 | 917851.88 | 0 | 12 | 0 | 4 | 0 | 12 | 1.557768924 | 2.029183754 | 1.002649107 | 1.306072581 |
| Q5U1X1 | Rexo2 | Oligoribonuclease, mitochondrial | 256.5 | 17.72 | 26752 | 4 | 7211751.5 | 7610854 | 5521529.5 | 7278375.5 | 8969052 | 8595078 | 8427576 | 7971791.5 | 8708906 | 9022101 | 4494310 | 4600114 | 5490307 | 6030524.5 | 5808641.5 | 9227414 | 6663167 | 6729352 | 9206980 | 9070110 | 2 | 8 | 0 | 10 | 2 | 10 | 0.722130874 | 0.618457979 | 1.117662658 | 0.957205146 |
| P35601 | Rfc1 | Replication factor C subunit 1 | 119.12 | 2.03 | 125985 | 2 | 2 | 297774.78 | 355255.88 | 2 | 465204.28 | 2 | 310305.78 | 278648.59 | 482878.59 | 218871.03 | 340286.06 | 259697.53 | 249447.33 | 2 | 193244.23 | 310442.78 | 2 | 2 | 2 | 270605.47 | 10 | 8 | 9.5 | 7.5 | 10 | 8 | 0.932427867 | 0.807834749 | 0.519615468 | 0.450183275 |
| Q9WUK4 | Rfc2 | Replication factor C subunit 2 | 131.57 | 10.6 | 38725 | 3 | 4096969.5 | 3338042.5 | 4081475.5 | 3800093.25 | 3273991.25 | 4161914.5 | 3250334.5 | 3697608 | 3056406.5 | 3820469.75 | 3936291.25 | 2899428.5 | 3177548.25 | 3371672.5 | 3451018.5 | 3620268.5 | 4269470 | 3036582.25 | 4138577.5 | 3250100.5 | 7 | 12 | 9 | 12 | 9 | 12 | 0.905618127 | 0.936020942 | 0.98517672 | 1.018250424 |
| Q8R323 | Rfc3 | Replication factor C subunit 3 | 215.53 | 12.92 | 40526 | 3 | 412341.03 | 791530.19 | 747369.69 | 1242881.25 | 1136529.75 | 700992.5 | 567473.75 | 502239.41 | 990187.88 | 514259.22 | 373292.41 | 571511.88 | 602068.38 | 2 | 460947.16 | 418910.56 | 544097.19 | 424096 | 426902.88 | 411777.12 | 3 | 4 | 6 | 2 | 6 | 4 | 0.463630389 | 0.613046773 | 0.513960437 | 0.679596927 |
| Q99J62 | Rfc4 | Replication factor C subunit 4 | 240.63 | 15.93 | 39867 | 5 | 6456764 | 5550526 | 7077640.5 | 7530501 | 4911740.5 | 5867971 | 5392639 | 5309192 | 3601785 | 6310186 | 2079287.25 | 2055400.5 | 2318835.25 | 4224212.5 | 1678667.25 | 5871756 | 6340987.5 | 4200538 | 6029969.5 | 4489544 | 0 | 6 | 1 | 10 | 1 | 10 | 0.391928675 | 0.466600282 | 0.854272467 | 1.017031413 |
| Q9D0F6 | Rfc5 | Replication factor C subunit 5 | 250.05 | 17.11 | 38096 | 5 | 3989776.5 | 4724125.5 | 5502698.5 | 4916962 | 2061044 | 5688896 | 5882171 | 3400533 | 4134248.75 | 4151947.75 | 4384236 | 1972659.75 | 4246529 | 2257383 | 3344745.75 | 4648827 | 5571094 | 3671808.5 | 3983295.25 | 3101327.5 | 6 | 10 | 6 | 8 | 6 | 10 | 0.764607425 | 0.6967794 | 0.989702368 | 0.90190626 |
| Q8CFV9 | Rfk | Riboflavin kinase | 83.77 | 13.55 | 17437 | 1 | 903566.25 | 765695.25 | 541264.56 | 374322.31 | 594807.94 | 913338.19 | 1009974.94 | 545286.31 | 611705.06 | 371593.94 | 257626.73 | 598834.62 | 349275.03 | 233687.75 | 263791.94 | 1082747.88 | 570518.38 | 527057.69 | 200588.95 | 515248.28 | 3 | 9 | 2 | 9 | 3 | 9 | 0.535660431 | 0.493414305 | 0.910840952 | 0.839005327 |
| Q6A0D4 | Rftn1 | Raftlin | 163.68 | 7.22 | 61537 | 3 | 1823941.75 | 2308586.75 | 1440984.75 | 1065025 | 1323627 | 2185164.25 | 1799810.75 | 1749477.12 | 1405428.5 | 1714795.75 | 2471732.5 | 1853051 | 2326756.5 | 1643560 | 1226160.5 | 2071629.88 | 2805706.75 | 1863128.25 | 2205068 | 2183334.5 | 7 | 4 | 10 | 3 | 10 | 4 | 1.195812973 | 1.075280462 | 1.397718715 | 1.256835023 |
| O70521 | Rgs19 | Regulator of G-protein signaling 19 | 71.67 | 6.02 | 24738 | 1 | 585786.75 | 612925.94 | 350976.91 | 419235.62 | 476826.56 | 2 | 452303.84 | 301221.25 | 375504.94 | 306868.47 | 354447.97 | 196648 | 76479.82 | 276624.59 | 504680 | 585880.12 | 616058.81 | 363779.31 | 2 | 310367.53 | 4 | 10 | 11 | 8.5 | 11 | 10 | 0.576052072 | 0.981182457 | 0.767080202 | 1.306558337 |
| Q4V8F3 | Rhbdd1 | Rhomboid-related protein 4 | 72.01 | 4.43 | 35879 | 1 | 129295.73 | 130312.66 | 2 | 2 | 2 | 2 | 225774.77 | 2 | 160857.44 | 172236.27 | 410505 | 365722.44 | 417276.16 | 157276.44 | 464485.12 | 119417.41 | 2 | 214276.78 | 2 | 192008.09 | 0 | 9 | 3 | 12 | 3 | 12 | 6.992159256 | 3.248084715 | 2.024950466 | 0.940655156 |
| Q80WQ6 | Rhbdf2 | Inactive rhomboid protein 2 | 86.25 | 2.06 | 93434 | 2 | 895977.88 | 919255.88 | 1267540.88 | 1212610.38 | 940070.5 | 156853.86 | 791225.75 | 898898.25 | 792611.06 | 969389.06 | 1652333.12 | 1739037 | 1952946.12 | 2113357.25 | 2034659.12 | 988986.94 | 1066861 | 1351861.12 | 1357105.62 | 2 | 0 | 9 | 0 | 5 | 0 | 9 | 1.813086287 | 2.630199647 | 0.910105465 | 1.320267596 |
| Q921J2 | Rheb | GTP-binding protein Rheb | 189.45 | 24.46 | 20451 | 5 | 3899164.75 | 3763344 | 3129479.5 | 4462906 | 3290964 | 3681183 | 3731764 | 3813813.5 | 4075623.75 | 4524380.5 | 5425113.5 | 5515014.5 | 7112128.5 | 6554250 | 6154244 | 3122322 | 3344105 | 4334649 | 4473933 | 4910975 | 0 | 9 | 0 | 12 | 0 | 12 | 1.658631813 | 1.551476042 | 1.088436228 | 1.018117895 |
| P61589 | Rhoa | Transforming protein RhoA | 690.79 | 70.98 | 21782 | 11 | 6201595 | 5453976 | 3850716 | 4003925.75 | 4183012.5 | 5600068.5 | 5911258.5 | 4565100.5 | 3987442.5 | 3741532.75 | 5818484 | 5466621.5 | 2977455.25 | 2356635.75 | 3203296 | 5107155 | 6300080.5 | 2772068.25 | 3767106.25 | 2651325.25 | 8 | 8 | 7 | 9 | 8 | 9 | 0.836631243 | 0.832688811 | 0.869351261 | 0.865254643 |
| P62746 | Rhob | Rho-related GTP-binding protein RhoB | 271.99 | 25 | 22123 | 4 | 1140287.5 | 1083259.25 | 1184648 | 1122590.5 | 685750.19 | 1361095.38 | 276501.56 | 1070388.12 | 978140.94 | 525675.5 | 2657647 | 2650178.5 | 1085570.75 | 806448.44 | 1928366.5 | 975822.88 | 309955.44 | 1138145.38 | 1480108.25 | 1041656.88 | 7 | 10 | 4 | 10 | 7 | 10 | 1.74986086 | 2.167293779 | 0.948079216 | 1.17424547 |
| Q62159 | Rhoc | Rho-related GTP-binding protein RhoC | 681.5 | 65.8 | 22006 | 10 | 5657037.5 | 4961224 | 4148133 | 6285029.5 | 5093144 | 7321600.5 | 6023679 | 5114305 | 4865852 | 6714911 | 6185593 | 5593749 | 6089082.5 | 6098007 | 6316691.5 | 2948077.5 | 4329709.5 | 3103206.25 | 2273900.75 | 2078614.5 | 5 | 1 | 11 | 0 | 11 | 1 | 1.158295023 | 1.008081648 | 0.563539948 | 0.490457326 |
| P84096 | Rhog | Rho-related GTP-binding protein RhoG | 690.18 | 50.79 | 21309 | 9 | 22132714 | 22872542 | 19689422 | 24677382 | 23269396 | 25634974 | 22727712 | 23324510 | 22907718 | 29888978 | 18589670 | 21193078 | 28929270 | 27978902 | 29687042 | 24056514 | 25920608 | 27139068 | 32507822 | 30625500 | 9 | 1 | 12 | 4 | 12 | 4 | 1.121948939 | 1.015215382 | 1.245096761 | 1.126647872 |
| Q8BG51 | Rhot1 | Mitochondrial Rho GTPase 1 | 215.18 | 5.39 | 72242 | 4 | 579322.12 | 676448.38 | 989089.19 | 913747.25 | 975509.25 | 667556.5 | 878517.38 | 1223866.88 | 882654.75 | 918014.88 | 2080680 | 888078.31 | 2372307.5 | 2457268.25 | 1881751 | 862302.38 | 1011658.75 | 1621951.75 | 1514847.75 | 1985894.25 | 3 | 3 | 2 | 5 | 3 | 5 | 2.341512579 | 2.117897662 | 1.692418538 | 1.530792232 |
| Q7TSA0 | Rhot2 | Mitochondrial Rho GTPase 2 | 96.62 | 3.22 | 69153 | 2 | 988826.31 | 445198.53 | 422630.44 | 466660.47 | 326684.19 | 582696.12 | 546678.81 | 1341134.75 | 309175.38 | 514630.78 | 2 | 2 | 2 | 934246.75 | 439306.66 | 245737.02 | 1172766.88 | 549138.19 | 2 | 2 | 6 | 9 | 5 | 7 | 6 | 9 | 0.518324317 | 0.416948306 | 0.742507975 | 0.597285198 |
| Q3TIR3 | Ric8a | Synembryn-A | 45.75 | 1.7 | 59847 | 1 | 2 | 2 | 2 | 2 | 115840.2 | 2 | 2 | 77479.71 | 172667.02 | 2 | 144205.75 | 244526.84 | 264455.59 | 2 | 261266.83 | 2 | 2 | 2 | 2 | 2 | 3 | 10 | 4.5 | 7.5 | 4.5 | 10 | 7.893579788 | 3.655594764 | 8.631986E-05 | 3.997558E-05 |
| P52760 | Rida | 2-iminobutanoate/2-iminopropanoate deaminase | 88.74 | 19.26 | 14255 | 2 | 2567178.5 | 597736.25 | 1965444.75 | 697412.31 | 2246829.25 | 1911788.88 | 513523.62 | 1581344.75 | 1516173.75 | 3484611.5 | 1073937.75 | 2 | 411183.38 | 455432.5 | 2 | 3233029 | 4970979 | 766693.69 | 672785.12 | 729623.56 | 2 | 10 | 1 | 12 | 2 | 12 | 0.240328608 | 0.215439358 | 1.284659179 | 1.151615497 |
| O35730 | Ring1 | E3 ubiquitin-protein ligase RING1 | 42.37 | 3.45 | 42631 | 1 | 776934.5 | 496668.62 | 938117.94 | 607374.12 | 809074.62 | 567016.44 | 666305.31 | 607404.5 | 656575.5 | 824806.44 | 2 | 645312.69 | 552256.19 | 399771.31 | 482950.84 | 815403.94 | 429451.28 | 560019.88 | 887272.44 | 706151.44 | 3 | 11 | 2 | 12 | 3 | 12 | 0.573372567 | 0.626196653 | 0.936642761 | 1.02293447 |
| Q9DBU3 | Riok3 | Serine/threonine-protein kinase RIO3 | 44.22 | 1.73 | 58704 | 1 | 2 | 635738.5 | 340251.41 | 565695.12 | 2 | 614227.12 | 685513.81 | 395232.12 | 523267.5 | 556423.12 | 749453.69 | 766342.12 | 726983.81 | 766251.31 | 646078.81 | 2 | 555743.5 | 2 | 720733 | 2 | 0 | 11 | 1 | 7 | 1 | 11 | 2.370847602 | 1.317316322 | 0.827976638 | 0.460049452 |
| Q8CD15 | Riox2 | Ribosomal oxygenase 2 | 52.86 | 4.3 | 53517 | 1 | 2 | 2 | 2 | 2 | 2 | 2 | 2 | 40586.36 | 2 | 2 | 2 | 2 | 2 | 135523.58 | 147280.61 | 2 | 2 | 2 | 2 | 2 | 7.5 | 12.5 | 9 | 10 | 9 | 12.5 | 28281.019 | 6.966736019 | 1 | 0.00024634 |
| Q9QZL0 | Ripk3 | Receptor-interacting serine/threonine-protein kinase 3 | 294.76 | 17.08 | 53322 | 6 | 694077.25 | 4725680.5 | 4865145 | 2222153.75 | 3859112 | 1675472.38 | 4130496.25 | 4649263.5 | 5332154 | 3915735 | 4289625 | 4486154 | 4724883.5 | 4532626 | 4824726.5 | 3262610.25 | 435400.84 | 4058287 | 3172546.5 | 3011559.75 | 9 | 9 | 8 | 5 | 9 | 9 | 1.396662573 | 1.160121528 | 0.851781792 | 0.707522643 |
| P13832 | Rlc-a | Myosin regulatory light chain RLC-A | 531.43 | 57.56 | 19895 | 8 | 266602 | 158144.84 | 223961.97 | 2 | 900876.5 | 2 | 2 | 196898.64 | 377588.16 | 2 | 2 | 199231.52 | 2 | 2 | 2 | 2 | 2 | 2 | 2 | 2 | 4 | 2.5 | 10 | 7.5 | 10 | 7.5 | 0.128575859 | 0.346809429 | 6.453331E-06 | 1.740666E-05 |
| Q8VC42 | Rmc1 | Regulator of MON1-CCZ1 complex | 43.84 | 2.28 | 74922 | 1 | 2 | 2 | 2 | 2 | 134496.16 | 2 | 185890.16 | 2 | 217600.19 | 2 | 2 | 2 | 2 | 2 | 2 | 2 | 2 | 2 | 2 | 2 | 10 | 10 | 7.5 | 7.5 | 10 | 10 | 7.434714E-05 | 2.478337E-05 | 7.434714E-05 | 2.478337E-05 |
| Q9DCV4 | Rmdn1 | Regulator of microtubule dynamics protein 1 | 39.45 | 3.61 | 35000 | 1 | 700219.62 | 427547.53 | 505417 | 488123.66 | 757882.62 | 146030.14 | 896615.12 | 574986.88 | 857559.75 | 493565.09 | 264359.75 | 165078.72 | 2 | 2 | 331216.19 | 748155.88 | 462503.5 | 2 | 294469.59 | 413602.69 | 0 | 5 | 3 | 6 | 3 | 6 | 0.264191855 | 0.256221262 | 0.666414295 | 0.646308766 |
| Q8K3C0 | Rnasek | Ribonuclease kappa | 47.44 | 9.18 | 10992 | 1 | 1469449 | 1658866.75 | 1503862.88 | 2251671.25 | 1755666 | 2006461 | 1054254.12 | 1959435.62 | 1311566.62 | 2644926.75 | 2300367.75 | 691109.81 | 2 | 4392557 | 3132228.5 | 1427579.75 | 854175.62 | 1682789.5 | 2407647.5 | 1748587.75 | 10 | 11 | 11 | 10 | 11 | 11 | 1.217228512 | 1.17151409 | 0.939957775 | 0.904656575 |
| C0HKG5 | Rnaset2a | Ribonuclease T2-A | 52.03 | 6.18 | 29609 | 1 | 2 | 671696.75 | 2 | 2 | 615379.69 | 2 | 569366.12 | 711055.75 | 745889.31 | 211664.5 | 2 | 293094.66 | 2 | 2 | 2 | 400213.78 | 2 | 2 | 2 | 2 | 9 | 9 | 4 | 4 | 9 | 9 | 0.227726407 | 0.130967642 | 0.310952716 | 0.178831891 |
| Q3U2C5 | Rnf149 | E3 ubiquitin-protein ligase RNF149 | 415.94 | 24.11 | 42553 | 7 | 1440000.75 | 1070581.88 | 447641.69 | 319087.47 | 409728.66 | 1192406.88 | 896527.88 | 615830.19 | 468418.28 | 460726.19 | 7997166.5 | 7602215.5 | 5897767.5 | 4531718 | 5693195.5 | 1093578.62 | 1689244.75 | 1025040.88 | 576208.75 | 1380439.12 | 0 | 6 | 0 | 5 | 0 | 6 | 8.603665577 | 8.72945892 | 1.563452367 | 1.58631145 |
| E9Q555 | Rnf213 | E3 ubiquitin-protein ligase RNF213 | 4060.3 | 17.49 | 584784 | 74 | 30747120 | 23916422 | 30173840 | 31645024 | 39179866 | 24718102 | 28126580 | 36374690 | 43883082 | 37697182 | 35411214 | 45491990 | 61310834 | 47371474 | 53929762 | 26974586 | 26101934 | 29141486 | 30224426 | 33080786 | 1 | 9 | 3 | 8 | 3 | 9 | 1.564382113 | 1.425736493 | 0.934865052 | 0.852011289 |
| Q91VI7 | Rnh1 | Ribonuclease inhibitor | 2519.29 | 82.68 | 49816 | 30 | 164062866 | 170076418 | 145901682 | 130123730 | 166484354 | 130097794 | 195301586 | 198912434 | 192759714 | 154316194 | 97392970 | 150362018 | 110252682 | 59717918 | 113266282 | 181184802 | 80336826 | 121376458 | 130056002 | 141794962 | 2 | 6 | 1 | 3 | 2 | 6 | 0.683696027 | 0.609363498 | 0.84304365 | 0.751386591 |
| Q5U2U7 | Rnmt | mRNA cap guanine-N7 methyltransferase | 61.31 | 4.12 | 52817 | 1 | 2 | 2 | 255359.3 | 2 | 2 | 219393.69 | 2 | 2 | 269953.59 | 2 | 2 | 2 | 388037.72 | 414251.12 | 458778.03 | 2 | 2 | 234856.61 | 348979.44 | 2 | 6 | 10 | 7 | 11.5 | 7 | 11.5 | 4.938262926 | 2.57701526 | 2.286283522 | 1.193089071 |
| Q8VCT3 | Rnpep | Aminopeptidase B | 1046.63 | 37.54 | 72416 | 19 | 33920130 | 36046778 | 28082374 | 25419314 | 23577646 | 31102312 | 31619046 | 21730044 | 25029722 | 23384902 | 32063606 | 28532352 | 21683224 | 21349656 | 21109826 | 26507624 | 26394390 | 21986784 | 23653090 | 21546754 | 6 | 5 | 8 | 9 | 8 | 9 | 0.848295491 | 0.938830397 | 0.816672636 | 0.903832572 |
| P60603 | Romo1 | Reactive oxygen species modulator 1 | 161.57 | 46.84 | 8183 | 2 | 749457.25 | 1691864.75 | 1594785.62 | 1237379.62 | 1104837.12 | 2095323.75 | 1578353.75 | 1854438.12 | 1341738.75 | 1674165.25 | 1711448.12 | 1197301.88 | 2199521.25 | 1838059 | 1340091.12 | 1621870 | 1963347.25 | 2356766.5 | 1894710.5 | 2250959 | 5 | 1 | 11 | 5 | 11 | 5 | 1.299153336 | 0.969850461 | 1.581552251 | 1.180668315 |
| Q9EPK2 | Rp2 | Protein XRP2 | 118 | 5.19 | 39377 | 2 | 385329.16 | 456477.72 | 796243.19 | 383561.16 | 939498.69 | 370106.75 | 505488.41 | 904989.5 | 1052128.75 | 533342.69 | 2 | 508399.72 | 379512.72 | 2 | 570623.81 | 488233.19 | 2 | 487231.22 | 396421.69 | 451131.09 | 6 | 10 | 6 | 4 | 6 | 10 | 0.492565386 | 0.43330836 | 0.615654008 | 0.541589069 |
| Q8VEE4 | Rpa1 | Replication protein A 70 kDa DNA-binding subunit | 751.98 | 26.81 | 69037 | 15 | 13808226 | 12899457 | 17949512 | 14143418 | 15592011 | 12758764 | 14812416 | 16261344 | 15890834 | 16602550 | 13892492 | 15970343 | 17607018 | 15822848 | 15430017 | 13760681 | 8442480 | 13770460 | 15063253 | 15769153 | 8 | 9 | 12 | 6 | 12 | 9 | 1.058205959 | 1.031402312 | 0.8980195 | 0.875273269 |
| Q62193 | Rpa2 | Replication protein A 32 kDa subunit | 162.07 | 13.33 | 29718 | 3 | 2468875.25 | 421731.03 | 5391161 | 341404.84 | 444754.16 | 2213536.75 | 1519551.12 | 3550867.5 | 1761336.88 | 227071.72 | 2706263 | 1178008.62 | 1423651.75 | 2474188.5 | 1987386.5 | 3170832 | 2291266 | 2410317.5 | 237238.98 | 1092084 | 8 | 12 | 12 | 11 | 12 | 12 | 1.077368526 | 1.053614634 | 1.014767676 | 0.992394012 |
| Q9CQ71 | Rpa3 | Replication protein A 14 kDa subunit | 47.09 | 8.26 | 13584 | 1 | 1079400.12 | 1024493.25 | 1353243.75 | 1207658.25 | 875332.94 | 919185.75 | 906045.12 | 797955.44 | 838085.75 | 940839.31 | 2 | 646331.31 | 647940.69 | 686944.12 | 616709.5 | 1161057 | 995168.88 | 826258.81 | 857253.75 | 906895.75 | 0 | 5 | 0 | 9 | 0 | 9 | 0.468929143 | 0.590154906 | 0.856773332 | 1.078263086 |
| Q80TE0 | Rpap1 | RNA polymerase II-associated protein 1 | 69.92 | 0.92 | 155270 | 1 | 279815.59 | 2 | 305078.94 | 357011.12 | 287583.88 | 317186.53 | 199250.38 | 2 | 295536.25 | 2 | 332159 | 362089.28 | 418115.06 | 2 | 2 | 255027.86 | 531709.75 | 281753.66 | 281489.78 | 305830.44 | 10 | 11 | 8 | 7 | 10 | 11 | 0.904737701 | 1.369949051 | 1.346744934 | 2.039234072 |
| Q9JJ80 | Rpf2 | Ribosome production factor 2 homolog | 383.38 | 27.45 | 35364 | 7 | 1331525.38 | 1792584.12 | 1049024.5 | 305521.16 | 1892840.25 | 617229 | 1309405.5 | 1354114.62 | 1266479.75 | 505557.62 | 1063925.88 | 3597980.5 | 1095627.25 | 936968.56 | 947553.38 | 1353520.25 | 1902389.88 | 825400.5 | 734263.25 | 1338761.38 | 11 | 12 | 12 | 8 | 12 | 12 | 1.19941318 | 1.512443794 | 0.965916926 | 1.218008177 |
| Q6PDV7 | Rpl10 | 60S ribosomal protein L10 | 440.6 | 35.05 | 24604 | 7 | 10528438 | 19621512 | 15798494 | 14418362 | 11473873 | 15972617 | 13846737 | 18235274 | 16085418 | 17440242 | 28678498 | 35552602 | 27524156 | 28955280 | 28666562 | 14842060 | 13990552 | 20632100 | 17011770 | 18986706 | 0 | 7 | 0 | 10 | 0 | 10 | 2.079282937 | 1.831044014 | 1.189621106 | 1.047596057 |
| P53026 | Rpl10a | 60S ribosomal protein L10a | 597.36 | 35.48 | 24916 | 10 | 73255234 | 71943346 | 70235506 | 73094170 | 70391402 | 89076674 | 69322386 | 77690482 | 65772018 | 82858218 | 58103302 | 64667670 | 68736994 | 85660738 | 68640370 | 72041874 | 88038514 | 85554986 | 93712178 | 89292354 | 5 | 2 | 6 | 5 | 6 | 5 | 0.963472093 | 0.898859621 | 1.194250291 | 1.114161347 |
| P62914 | Rpl11 | 60S ribosomal protein L11 | 649.34 | 50.56 | 20252 | 10 | 49207562 | 54388898 | 46708286 | 44787654 | 39316834 | 59886058 | 53580078 | 54638482 | 48889946 | 57198098 | 65878654 | 71958298 | 60365138 | 70038082 | 64330466 | 51102662 | 50463122 | 62853050 | 63986090 | 64153026 | 0 | 2 | 0 | 8 | 0 | 8 | 1.418760824 | 1.212908601 | 1.248064955 | 1.066979502 |
| P35979 | Rpl12 | 60S ribosomal protein L12 | 545.95 | 51.52 | 17805 | 8 | 47545050 | 56155174 | 55583698 | 54144578 | 49251906 | 65734794 | 58799462 | 64942922 | 62652994 | 71353530 | 67314522 | 74888066 | 72307498 | 74565538 | 67996042 | 52242090 | 51962826 | 76264610 | 74852746 | 62948998 | 0 | 6 | 2 | 12 | 2 | 12 | 1.359338793 | 1.103832013 | 1.211629275 | 0.98388657 |
| P47963 | Rpl13 | 60S ribosomal protein L13 | 460.27 | 34.12 | 24305 | 8 | 57494618 | 68127730 | 49548914 | 45284914 | 34602234 | 67290018 | 63004238 | 56261994 | 47211222 | 62256126 | 120013106 | 102786634 | 60507906 | 87883970 | 81449962 | 64292902 | 60553058 | 45975134 | 62415426 | 37741178 | 1 | 10 | 3 | 9 | 3 | 10 | 1.774658511 | 1.529072618 | 1.062414284 | 0.915392218 |
| P19253 | Rpl13a | 60S ribosomal protein L13a | 466.92 | 34.98 | 23464 | 9 | 69195514 | 86454522 | 70369106 | 61459890 | 56470870 | 85132674 | 68568242 | 82118202 | 69768322 | 83383514 | 73985242 | 82018026 | 87607186 | 84805210 | 96254034 | 38956106 | 26570942 | 70462450 | 55259094 | 64014454 | 3 | 6 | 7 | 2 | 7 | 6 | 1.234684748 | 1.091777403 | 0.742151821 | 0.656252204 |
| Q9CR57 | Rpl14 | 60S ribosomal protein L14 | 449.14 | 29.49 | 23564 | 7 | 51253658 | 52529658 | 43972842 | 41134402 | 44244302 | 52459346 | 51559242 | 51445850 | 47447734 | 53643034 | 70782994 | 71747514 | 49572430 | 68243986 | 55095122 | 50974418 | 49518982 | 56680902 | 39914230 | 54912634 | 2 | 9 | 4 | 12 | 4 | 12 | 1.353045372 | 1.229528923 | 1.080924422 | 0.982249279 |
| P61314 | Rpl15 | 60S ribosomal protein L15 | 717.1 | 47.55 | 24146 | 12 | 47592318 | 62604774 | 70160354 | 52638130 | 55058354 | 54288130 | 54766918 | 73084754 | 68448202 | 68353186 | 94986746 | 106756218 | 85142002 | 86707466 | 84388634 | 34729338 | 27784160 | 39717230 | 32489202 | 45706570 | 0 | 0 | 0 | 0 | 0 | 0 | 1.589914312 | 1.435942049 | 0.626363612 | 0.565704605 |
| P24049 | Rpl17 | 60S ribosomal protein L17 | 330.08 | 36.96 | 21397 | 6 | 37614534 | 51284962 | 46788778 | 43459782 | 32340592 | 41805066 | 45428810 | 46171330 | 38956706 | 44100354 | 26594804 | 40630702 | 34042966 | 33967202 | 35748754 | 50933050 | 42695018 | 42672486 | 43429802 | 53975210 | 5 | 10 | 1 | 9 | 5 | 10 | 0.808480406 | 0.789904084 | 1.105050168 | 1.079659611 |
| P35980 | Rpl18 | 60S ribosomal protein L18 | 658.82 | 45.74 | 21645 | 11 | 66130734 | 79426778 | 66929874 | 58445394 | 44431390 | 92232066 | 63898330 | 66171134 | 65068586 | 90215866 | 217900930 | 188183042 | 173846962 | 171976002 | 153798226 | 49074826 | 45590370 | 90411938 | 81261666 | 86217538 | 0 | 8 | 0 | 10 | 0 | 10 | 2.871934253 | 2.398672634 | 1.117934032 | 0.933711406 |
| P62717 | Rpl18a | 60S ribosomal protein L18a | 571.54 | 51.14 | 20732 | 12 | 33967998 | 39493154 | 41743158 | 31028280 | 35983934 | 40482562 | 38604798 | 39760434 | 37629782 | 37961578 | 51993170 | 62921022 | 61984386 | 58550138 | 59570154 | 40726466 | 37222378 | 46270250 | 35836242 | 44961682 | 0 | 6 | 0 | 10 | 0 | 10 | 1.619056623 | 1.517281185 | 1.125128575 | 1.054401924 |
| P84099 | Rpl19 | 60S ribosomal protein L19 | 315.85 | 14.29 | 23466 | 4 | 2932831.25 | 4805663.5 | 4594362.5 | 5031310.5 | 4198651.5 | 4386003 | 3857302 | 5382448.5 | 4102763.25 | 5255082.5 | 6678651.5 | 6500682 | 4718349 | 5934667 | 4680558 | 3807777 | 3767795 | 4097301.75 | 4363548.5 | 4931568.5 | 4 | 9 | 4 | 6 | 4 | 9 | 1.322318161 | 1.240576256 | 0.972414159 | 0.912302313 |
| O09167 | Rpl21 | 60S ribosomal protein L21 | 303.88 | 38.12 | 18562 | 6 | 38344534 | 39227422 | 42229910 | 41982594 | 35936282 | 35935538 | 43926834 | 41244518 | 34839654 | 46160114 | 37975914 | 36074238 | 34238458 | 35705746 | 37217510 | 53003118 | 58739866 | 51015458 | 53441762 | 54538458 | 3 | 0 | 7 | 0 | 7 | 0 | 0.916504076 | 0.896615024 | 1.36929823 | 1.339583093 |
| P67984 | Rpl22 | 60S ribosomal protein L22 | 426.57 | 56.25 | 14759 | 7 | 26171456 | 4369093 | 19429210 | 24898026 | 19040844 | 27591256 | 22743604 | 18406340 | 20145584 | 22627728 | 27777980 | 23541890 | 19081468 | 21485718 | 18219436 | 18686950 | 29212894 | 22487546 | 22804848 | 19020596 | 11 | 12 | 12 | 12 | 12 | 12 | 1.172485353 | 0.987373661 | 1.194915049 | 1.006262163 |
| Q9D7S7 | Rpl22l1 | 60S ribosomal protein L22-like 1 | 334.03 | 50.82 | 14467 | 5 | 7764551.5 | 7720018.5 | 13056604 | 12927331 | 12154583 | 6347811 | 6761531.5 | 12465460 | 12551930 | 14294729 | 3383156.5 | 2665428.75 | 4057711 | 3555043 | 3602887 | 5413088.5 | 6767973.5 | 7582463.5 | 7505820 | 7097291.5 | 0 | 0 | 0 | 8 | 0 | 8 | 0.321955092 | 0.329335081 | 0.640892539 | 0.655583343 |
| P62830 | Rpl23 | 60S ribosomal protein L23 | 474.17 | 67.86 | 14865 | 8 | 60085382 | 64951198 | 59085922 | 66386874 | 50137894 | 72040490 | 56214030 | 53466090 | 47504130 | 62393830 | 120180258 | 84940842 | 80969018 | 91672450 | 92185498 | 63318970 | 71536106 | 55455534 | 60225630 | 63629694 | 0 | 10 | 0 | 8 | 0 | 10 | 1.563121016 | 1.611516256 | 1.044965198 | 1.077317998 |
| P62751 | Rpl23a | 60S ribosomal protein L23a | 398.22 | 37.18 | 17695 | 8 | 13074343 | 11750140 | 11062038 | 11624124 | 10018829 | 14539240 | 12914109 | 13109376 | 10052779 | 13293364 | 21104400 | 19033714 | 13120010 | 13691022 | 13507884 | 13034986 | 15597614 | 16075098 | 11739028 | 13709102 | 0 | 3 | 4 | 8 | 4 | 8 | 1.398535818 | 1.258933737 | 1.219476264 | 1.097747937 |
| P83732 | Rpl24 | 60S ribosomal protein L24 | 310.24 | 26.75 | 17779 | 5 | 18581420 | 23483404 | 14974894 | 14746763 | 10709080 | 25206022 | 21202960 | 13305952 | 16940472 | 22941020 | 57307642 | 50541802 | 36416354 | 38102042 | 35962134 | 15786142 | 18317374 | 21522118 | 15510011 | 18318230 | 0 | 9 | 0 | 9 | 0 | 9 | 2.646566329 | 2.192146674 | 1.084347738 | 0.898163504 |
| P61255 | Rpl26 | 60S ribosomal protein L26 | 540.15 | 46.9 | 17258 | 11 | 30649744 | 36092766 | 28029440 | 22090888 | 16435398 | 38577186 | 33020174 | 34170542 | 25847226 | 30254586 | 47288362 | 44684826 | 38590826 | 35387318 | 37688770 | 30045584 | 32627954 | 34334762 | 29825760 | 34689230 | 1 | 7 | 2 | 12 | 2 | 12 | 1.527702902 | 1.258049434 | 1.211743642 | 0.997859859 |
| P61354 | Rpl27 | 60S ribosomal protein L27 | 284.49 | 42.65 | 15798 | 5 | 24660006 | 27295986 | 24421514 | 20859728 | 16697353 | 28249224 | 26376850 | 28144434 | 23617054 | 30782118 | 59127082 | 52115650 | 48279594 | 39976346 | 41816674 | 26166294 | 20042938 | 31124874 | 21480350 | 28751348 | 0 | 8 | 0 | 10 | 0 | 10 | 2.118016595 | 1.759246985 | 1.119640729 | 0.929985431 |
| P14115 | Rpl27a | 60S ribosomal protein L27a | 337.53 | 32.43 | 16605 | 6 | 61261942 | 65413842 | 63601414 | 55034730 | 54850222 | 59794706 | 68245474 | 53551498 | 50403722 | 54411162 | 68832978 | 62496970 | 51773478 | 50278622 | 46837058 | 24157912 | 23149892 | 15184820 | 12620028 | 13894153 | 8 | 0 | 10 | 0 | 10 | 0 | 0.933559098 | 0.978396249 | 0.296529076 | 0.31077083 |
| P41105 | Rpl28 | 60S ribosomal protein L28 | 379.06 | 40.88 | 15733 | 7 | 38160010 | 48705542 | 39380130 | 31052262 | 22922522 | 47155322 | 46042230 | 45894298 | 30262412 | 44075134 | 56918850 | 66560590 | 54140282 | 65680586 | 56142658 | 37155226 | 48837874 | 56112394 | 47490134 | 46672238 | 0 | 5 | 0 | 5 | 0 | 5 | 1.661536964 | 1.403007138 | 1.310993536 | 1.107007143 |
| P47915 | Rpl29 | 60S ribosomal protein L29 | 95.98 | 11.88 | 17587 | 2 | 38707466 | 43819890 | 45158330 | 49810546 | 6586432 | 40824594 | 36131098 | 44362238 | 37447570 | 45243742 | 31242586 | 35148966 | 32799324 | 39393994 | 34224954 | 52324550 | 58511962 | 56768114 | 53750122 | 58263698 | 6 | 0 | 2 | 0 | 6 | 0 | 0.938762077 | 0.847068605 | 1.518983048 | 1.370616563 |
| P27659 | Rpl3 | 60S ribosomal protein L3 | 1071.69 | 46.15 | 46110 | 18 | 60903566 | 83046514 | 68413602 | 59743898 | 51943142 | 76359530 | 66186314 | 84800106 | 69915914 | 88865842 | 137650482 | 114641554 | 118726570 | 116060122 | 126282690 | 37496106 | 35695854 | 69466378 | 53174178 | 70514010 | 0 | 9 | 0 | 3 | 0 | 9 | 1.89279448 | 1.588493673 | 0.821928507 | 0.689788694 |
| P62889 | Rpl30 | 60S ribosomal protein L30 | 461.99 | 50.43 | 12784 | 7 | 22865492 | 23192966 | 18168254 | 14812558 | 17972404 | 30706280 | 22071526 | 18318900 | 17170188 | 20302838 | 50727006 | 49045626 | 43146486 | 45831402 | 41534918 | 18088366 | 24748842 | 32699872 | 33652170 | 32232914 | 0 | 3 | 0 | 5 | 0 | 5 | 2.373790993 | 2.121083232 | 1.457785008 | 1.302592918 |
| P62900 | Rpl31 | 60S ribosomal protein L31 | 215.49 | 31.2 | 14463 | 4 | 12382484 | 13173400 | 9016778 | 9520594 | 6105968.5 | 17994486 | 16508778 | 12917846 | 8372157.5 | 15218908 | 28241364 | 27672740 | 23389516 | 20331160 | 20381924 | 11065463 | 18983612 | 13329313 | 11186331 | 13017694 | 0 | 5 | 0 | 11 | 0 | 11 | 2.39080793 | 1.690086287 | 1.346284005 | 0.951701768 |
| P62911 | Rpl32 | 60S ribosomal protein L32 | 362.73 | 40 | 15860 | 6 | 38568698 | 47736594 | 31130412 | 29451414 | 23360220 | 43458122 | 39642962 | 36934162 | 34013938 | 40259558 | 37636338 | 34217218 | 50151118 | 55928574 | 44770402 | 30586644 | 27026006 | 42730574 | 33938594 | 33038176 | 5 | 12 | 7 | 4 | 7 | 12 | 1.308118251 | 1.146132941 | 0.982746608 | 0.861052325 |
| P11250 | Rpl34 | 60S ribosomal protein L34 | 200.48 | 21.37 | 13507 | 4 | 9418797 | 19888446 | 16166396 | 9918688 | 9086686 | 16699977 | 13727236 | 26812662 | 19946138 | 23668262 | 40902906 | 33866758 | 53146242 | 60538870 | 58190038 | 9795318 | 7735543 | 29501414 | 21636354 | 26323290 | 0 | 8 | 0 | 12 | 0 | 12 | 3.825195246 | 2.445556363 | 1.473222287 | 0.941873004 |
| P17078 | Rpl35 | 60S ribosomal protein L35 | 181.46 | 21.95 | 14553 | 4 | 1121194.88 | 1613077 | 841653.06 | 648526.88 | 517271.66 | 1719203.62 | 1417661.62 | 2322194.75 | 861991.88 | 770731 | 3363037.5 | 4435335.5 | 2794574 | 1924032.88 | 2439354.75 | 785092.69 | 699684.38 | 266629.34 | 580719.25 | 294968.41 | 0 | 5 | 1 | 1 | 1 | 5 | 3.154197982 | 2.108966801 | 0.554037805 | 0.370441977 |
| O55142 | Rpl35a | 60S ribosomal protein L35a | 157.15 | 27.27 | 12554 | 4 | 9759211 | 15400194 | 9233290 | 7045611.5 | 7984857.5 | 17627738 | 20876836 | 26175630 | 14490232 | 20712130 | 56736386 | 53604822 | 45209298 | 43600994 | 55554366 | 6846346 | 6555198 | 18914630 | 19017804 | 17726820 | 0 | 10 | 0 | 6 | 0 | 10 | 5.153572645 | 2.55005329 | 1.397336642 | 0.691419942 |
| P39032 | Rpl36 | 60S ribosomal protein L36 | 114.52 | 19.05 | 12268 | 2 | 1187076.75 | 1351084.12 | 724337.75 | 618222.5 | 259250.17 | 2 | 856009.31 | 2 | 2 | 1133793.62 | 5625721.5 | 3448654.5 | 2651835.5 | 2354726.25 | 2584549.25 | 515020.03 | 545382.06 | 2 | 534504.81 | 395464.47 | 0 | 4 | 0 | 11.5 | 0 | 11.5 | 4.025507868 | 8.37542075 | 0.480769849 | 1.000283665 |
| P47964 | Rpl36 | 60S ribosomal protein L36 | 87.35 | 18.1 | 12216 | 2 | 2 | 2 | 2 | 2 | 2 | 2 | 3067728.25 | 2 | 2 | 2 | 4058658.75 | 4245923 | 2526637 | 1771514.5 | 1990539.38 | 2 | 1324170.75 | 1058940.88 | 2 | 2 | 0 | 7.5 | 3 | 11 | 3 | 11 | 1459327.263 | 4.757016719 | 238311.763 | 0.776832634 |
| P83882 | Rpl36a | 60S ribosomal protein L36a | 131.46 | 17.92 | 12441 | 3 | 6477954 | 8548935 | 12781729 | 13852250 | 8563354 | 7733672.5 | 8461798 | 16015631 | 8521872 | 8946752 | 18051964 | 16259811 | 10641787 | 13927320 | 12388990 | 12158961 | 8377294.5 | 12794609 | 15762999 | 13522917 | 4 | 8 | 3 | 8 | 4 | 8 | 1.419033868 | 1.434586671 | 1.246744658 | 1.260409148 |
| P61928 | Rpl37 | 60S ribosomal protein L37 | 48.17 | 7.22 | 11078 | 1 | 3452314 | 4585279 | 4669056.5 | 4746459 | 3557778.25 | 3006567.75 | 3633208 | 4620881.5 | 3459302 | 4068130 | 2623986.5 | 2380897 | 2471176.5 | 3337402.75 | 2530356 | 4309645 | 3585380.75 | 3071689.25 | 3344069.75 | 5172138 | 0 | 9 | 1 | 12 | 1 | 12 | 0.635090699 | 0.710227558 | 0.92727751 | 1.036982659 |
| P61514 | Rpl37a | 60S ribosomal protein L37a | 123.76 | 19.57 | 10275 | 1 | 531654.81 | 658775 | 1222192.88 | 2047246.25 | 709240.62 | 573045.56 | 633002.75 | 1757756.12 | 886809.94 | 2289959 | 1815273.62 | 861951.81 | 2177465.5 | 4403283.5 | 2883112.25 | 1226776.25 | 1858185.62 | 2251851.25 | 3041730.5 | 2750179.75 | 3 | 2 | 5 | 4 | 5 | 4 | 2.3487772 | 1.977191045 | 2.152928515 | 1.812326423 |
| P63174 | Rpl38 | 60S ribosomal protein L38 | 210.24 | 48.57 | 8218 | 4 | 224857.86 | 402180.69 | 1136925.12 | 1048542.69 | 1349236.25 | 769805.62 | 709168.5 | 1710797.5 | 1938542.25 | 1553965.5 | 10208006 | 5299055 | 4971177 | 3641907 | 4062600 | 2 | 673339.88 | 1835169.25 | 1255803.25 | 1227221.25 | 0 | 10 | 0 | 8 | 0 | 10 | 6.771861607 | 4.217534682 | 1.199385954 | 0.746980986 |
| P62892 | Rpl39 | 60S ribosomal protein L39 | 45.07 | 19.61 | 6407 | 1 | 7477343.5 | 4970857.5 | 6605486.5 | 5956417.5 | 4692699.5 | 6355581.5 | 5753667 | 6768819 | 4517022.5 | 7163695.5 | 7437683.5 | 5819069.5 | 5001640 | 4628952.5 | 3377543.75 | 6063533.5 | 7161184.5 | 4159952.75 | 3709189.25 | 4147637.75 | 8 | 7 | 9 | 6 | 9 | 7 | 0.884256207 | 0.85948734 | 0.849801834 | 0.825998067 |
| Q9D8E6 | Rpl4 | 60S ribosomal protein L4 | 1404.29 | 45.58 | 47154 | 22 | 119786658 | 137520850 | 138266946 | 122117658 | 119121866 | 149028082 | 131906010 | 143372242 | 136161922 | 119666978 | 179218850 | 190535714 | 175851858 | 158785986 | 171850658 | 129887706 | 113461530 | 145477522 | 137004834 | 159035522 | 0 | 9 | 0 | 12 | 0 | 12 | 1.375979637 | 1.288336528 | 1.07545867 | 1.006957263 |
| P09895 | Rpl5 | 60S ribosomal protein L5 | 723.45 | 39.73 | 34459 | 14 | 31035334 | 38214966 | 34511214 | 29019112 | 29498174 | 33252704 | 32885818 | 34534822 | 27726854 | 31003046 | 39617890 | 46156338 | 38935110 | 33234358 | 33860166 | 32940480 | 30669598 | 31634226 | 28138722 | 34308306 | 4 | 11 | 3 | 11 | 4 | 11 | 1.181940352 | 1.203261974 | 0.971730947 | 0.989260495 |
| P47911 | Rpl6 | 60S ribosomal protein L6 | 746.49 | 41.55 | 33510 | 13 | 65477570 | 72599906 | 64250318 | 59205750 | 52766282 | 88694394 | 68742018 | 68403410 | 62912738 | 86232802 | 131684330 | 106170858 | 74571682 | 79514162 | 71467938 | 63165190 | 79678522 | 91297746 | 71065010 | 80512690 | 1 | 4 | 6 | 10 | 6 | 10 | 1.474416884 | 1.235805493 | 1.227233126 | 1.028624573 |
| P14148 | Rpl7 | 60S ribosomal protein L7 | 1339.69 | 51.48 | 31420 | 25 | 119151890 | 126349314 | 177330370 | 144068994 | 132946218 | 124753690 | 111746586 | 151655858 | 145821234 | 172552594 | 177549570 | 188805154 | 165974274 | 153630146 | 158017234 | 90795930 | 95392570 | 150649458 | 129371034 | 140207490 | 3 | 9 | 3 | 7 | 3 | 9 | 1.205944494 | 1.194537279 | 0.866498917 | 0.85830257 |
| P12970 | Rpl7a | 60S ribosomal protein L7a | 1040.68 | 54.14 | 29977 | 18 | 39124486 | 48596426 | 33825094 | 28209090 | 24239524 | 55066674 | 44661142 | 41411074 | 32895968 | 45595086 | 162444658 | 133649706 | 105265634 | 107677834 | 101414394 | 34670294 | 45752530 | 46871902 | 42848122 | 44195314 | 0 | 6 | 0 | 12 | 0 | 12 | 3.508454606 | 2.779458096 | 1.231866606 | 0.975905917 |
| Q9D8M4 | Rpl7l1 | 60S ribosomal protein L7-like 1 | 280.39 | 29.67 | 28544 | 7 | 2596336.75 | 2431704 | 2980346.75 | 2519771.5 | 2667291.5 | 3160950.5 | 2830177.5 | 2033455.38 | 1356665 | 1845360.5 | 3582913.25 | 3228058.75 | 3062702 | 3407623.75 | 2784824.5 | 2491296.25 | 3496652.5 | 2214886 | 2309514.25 | 2688494.5 | 1 | 10 | 3 | 8 | 3 | 10 | 1.217550113 | 1.431075263 | 1.000408701 | 1.175853158 |
| P62918 | Rpl8 | 60S ribosomal protein L8 | 518.98 | 36.96 | 28025 | 10 | 104141098 | 133724586 | 110534506 | 105760946 | 83239890 | 143571346 | 116170970 | 143862434 | 113287530 | 143882802 | 160788210 | 155422706 | 146592578 | 172592450 | 153507586 | 96432314 | 99652474 | 165842434 | 151572626 | 173456946 | 0 | 8 | 0 | 10 | 0 | 10 | 1.467997811 | 1.193906295 | 1.278294534 | 1.039622729 |
| P51410 | Rpl9 | 60S ribosomal protein L9 | 1018.54 | 74.48 | 21881 | 15 | 76408610 | 79425650 | 87899578 | 81198906 | 74264906 | 79156290 | 76102370 | 80313418 | 81164802 | 85569050 | 91404658 | 102349058 | 104989802 | 95383490 | 101243074 | 65567018 | 56496358 | 85309458 | 79399162 | 76813250 | 0 | 8 | 0 | 7 | 0 | 8 | 1.240914324 | 1.231326821 | 0.910790046 | 0.903753136 |
| P14869 | Rplp0 | 60S acidic ribosomal protein P0 | 1093.11 | 56.47 | 34216 | 17 | 137619442 | 142038194 | 139902658 | 116522106 | 138853378 | 112637146 | 138884242 | 124318482 | 117367914 | 102136338 | 106872466 | 133554146 | 106897066 | 80483226 | 115530954 | 158288130 | 102556970 | 93766506 | 81183490 | 112170434 | 1 | 5 | 8 | 7 | 8 | 7 | 0.805021568 | 0.912645037 | 0.81187803 | 0.920418141 |
| P47955 | Rplp1 | 60S acidic ribosomal protein P1 | 222.13 | 66.67 | 11475 | 3 | 21010508 | 17846756 | 18974596 | 18821982 | 33063202 | 16097444 | 24837326 | 20325490 | 27478460 | 17633676 | 10935700 | 16095510 | 9570694 | 11259321 | 10582081 | 28479230 | 38681610 | 18035336 | 7630153 | 22779256 | 0 | 11 | 0 | 10 | 0 | 11 | 0.532672991 | 0.549421732 | 1.053670248 | 1.086800611 |
| P99027 | Rplp2 | 60S acidic ribosomal protein P2 | 498.87 | 71.3 | 11651 | 6 | 15746718 | 9963271 | 9627175 | 8751212 | 8858731 | 10396751 | 10412105 | 10097364 | 8121642.5 | 10364215 | 5841279 | 7653945.5 | 6029774 | 5125963.5 | 7164217.5 | 12105275 | 14878255 | 10268505 | 10550704 | 10291777 | 0 | 5 | 0 | 6 | 0 | 6 | 0.600886079 | 0.644135277 | 1.097217946 | 1.176190979 |
| Q91YQ5 | Rpn1 | Dolichyl-diphosphooligosaccharide--protein glycosyltransferase subunit 1 | 1934.48 | 60.2 | 68528 | 32 | 44076746 | 49187770 | 54164002 | 43592598 | 52538506 | 43553838 | 46487062 | 54449558 | 55251610 | 38940658 | 87041738 | 92271426 | 97006218 | 93561594 | 100863514 | 44778466 | 45878306 | 53981466 | 47961970 | 53504362 | 0 | 11 | 0 | 12 | 0 | 12 | 1.932769012 | 1.972260406 | 1.010448973 | 1.031095019 |
| Q9DBG6 | Rpn2 | Dolichyl-diphosphooligosaccharide--protein glycosyltransferase subunit 2 | 1163.91 | 31.54 | 69063 | 16 | 18775494 | 13843042 | 18711782 | 19092820 | 21979078 | 18053264 | 17055230 | 17980770 | 22922778 | 20778968 | 59039218 | 30686852 | 32196334 | 27173100 | 35940026 | 15949060 | 24794678 | 25914940 | 25330744 | 19842254 | 0 | 5 | 0 | 7 | 0 | 7 | 2.002501001 | 1.911701614 | 1.210270498 | 1.155393213 |
| Q9CQH8 | Rpp14 | Ribonuclease P protein subunit p14 | 61.88 | 13.11 | 13565 | 1 | 2 | 2 | 2 | 2 | 2 | 2 | 2 | 2 | 2 | 2 | 122617.98 | 157664.42 | 204098.47 | 242716.3 | 131538.2 | 2 | 2 | 2 | 2 | 2 | 0 | 12.5 | 0 | 12.5 | 0 | 12.5 | 85863.537 | 85863.537 | 1 | 1 |
| O88796 | Rpp30 | Ribonuclease P protein subunit p30 | 251.25 | 23.51 | 29473 | 5 | 750166.94 | 2068819.5 | 2106339.25 | 1597380.25 | 1362529.88 | 354796.06 | 2056958 | 1320686.75 | 1930795.75 | 2076528.38 | 2271721.25 | 2642791.75 | 2134418.75 | 1836972.25 | 2479721.75 | 1419229.5 | 506628.62 | 1996361.25 | 2295742 | 1878285.75 | 2 | 12 | 3 | 12 | 3 | 12 | 1.441380576 | 1.468471696 | 1.026760303 | 1.046058528 |
| Q9CSU0 | Rprd1b | Regulation of nuclear pre-mRNA domain-containing protein 1B | 151.68 | 11.04 | 36884 | 3 | 206546.02 | 742127 | 209432.84 | 202060.88 | 2 | 173849.34 | 209260.69 | 352022.66 | 447803.81 | 2 | 629211.75 | 970842.81 | 500226.81 | 206637.27 | 324758.78 | 491069.19 | 110621.34 | 2 | 2 | 2 | 5 | 6.5 | 5 | 7.5 | 5 | 7.5 | 1.934816867 | 2.224695045 | 0.442369033 | 0.508645657 |
| Q6NXI6 | Rprd2 | Regulation of nuclear pre-mRNA domain-containing protein 2 | 54.66 | 0.95 | 156586 | 1 | 244108.56 | 210210.88 | 2 | 147413.12 | 2 | 247246.48 | 403335.44 | 233124.81 | 351894.94 | 216515.64 | 2 | 200853.11 | 514613.03 | 2 | 2 | 2 | 2 | 200471.5 | 2 | 68193.62 | 11 | 8 | 5 | 0 | 11 | 8 | 1.189012248 | 0.492709601 | 0.44649293 | 0.185020258 |
| P63325 | Rps10 | 40S ribosomal protein S10 | 348.69 | 24.24 | 18916 | 5 | 7829103 | 8679740 | 7061984 | 7032239 | 3822406.25 | 9502366 | 9222597 | 6733453.5 | 2889025 | 7618783 | 13789864 | 10730584 | 5890281 | 7654103 | 6812145 | 10777984 | 13390899 | 10459331 | 10745734 | 8948775 | 10 | 0 | 9 | 2 | 10 | 2 | 1.303598007 | 1.247753347 | 1.577980473 | 1.51038158 |
| P62281 | Rps11 | 40S ribosomal protein S11 | 547.41 | 50.63 | 18431 | 12 | 112600698 | 131488018 | 117845770 | 122627258 | 106653970 | 134574930 | 117821850 | 147051554 | 131069210 | 138592082 | 157729810 | 163061570 | 161982578 | 187301890 | 163159106 | 113914594 | 137601714 | 169374866 | 159010082 | 169764946 | 0 | 3 | 0 | 7 | 0 | 7 | 1.410102834 | 1.245946735 | 1.26800791 | 1.120393689 |
| P63323 | Rps12 | 40S ribosomal protein S12 | 405.46 | 49.24 | 14525 | 6 | 14446056 | 18686352 | 16991748 | 12376538 | 13472290 | 14341465 | 16893730 | 12593742 | 14547456 | 10797275 | 22751702 | 27575998 | 21609772 | 15906435 | 20968696 | 21577948 | 11802368 | 12812913 | 13992302 | 12528391 | 2 | 9 | 1 | 11 | 2 | 11 | 1.432253905 | 1.573035031 | 0.957102356 | 1.051179215 |
| P62278 | Rps13 | 40S ribosomal protein S13 | 613.29 | 54.97 | 17222 | 11 | 49151486 | 44094458 | 56417326 | 44395258 | 51261946 | 40749730 | 44498926 | 47178762 | 42707690 | 47869218 | 57423894 | 58372766 | 41186058 | 30900404 | 41155566 | 46095466 | 31024836 | 39608558 | 28946896 | 31898196 | 10 | 2 | 12 | 3 | 12 | 3 | 0.933630546 | 1.027059394 | 0.72384481 | 0.796280302 |
| P13471 | Rps14 | 40S ribosomal protein S14 | 468.14 | 39.74 | 16259 | 8 | 17645522 | 31860406 | 19319550 | 14823905 | 13597129 | 29420234 | 21776538 | 30690020 | 16782084 | 24653142 | 70370954 | 63243558 | 56363666 | 61370130 | 60286494 | 19024394 | 23726732 | 26668310 | 25383668 | 25606376 | 0 | 6 | 0 | 12 | 0 | 12 | 3.204585908 | 2.527000507 | 1.238188162 | 0.97638266 |
| P62843 | Rps15 | 40S ribosomal protein S15 | 142.51 | 28.28 | 17040 | 2 | 3134937 | 2761665 | 2576118.5 | 2426432.5 | 2679518 | 3351933.75 | 2768912.5 | 2336597.5 | 2581188 | 1550124.62 | 618450 | 1720631.25 | 2353061 | 758509.44 | 2240902.5 | 2808228.5 | 1225052.75 | 2550723.25 | 1480589.88 | 3263151 | 0 | 10 | 4 | 10 | 4 | 10 | 0.566443814 | 0.610986023 | 0.834230786 | 0.899830376 |
| P62245 | Rps15a | 40S ribosomal protein S15a | 542.84 | 71.54 | 14840 | 10 | 112554538 | 102753242 | 116387394 | 121793546 | 96654738 | 116203466 | 105127418 | 110444370 | 91688954 | 109722290 | 103377586 | 111319482 | 114781226 | 120455442 | 98988098 | 106169866 | 131059394 | 128828018 | 119659546 | 117367506 | 12 | 5 | 10 | 3 | 12 | 5 | 0.997779445 | 1.02951188 | 1.096231031 | 1.131094527 |
| P14131 | Rps16 | 40S ribosomal protein S16 | 769.18 | 64.38 | 16445 | 14 | 87129538 | 78053866 | 85512258 | 84388994 | 86635434 | 85250138 | 98477426 | 105833554 | 90691770 | 86924906 | 170714802 | 151698722 | 135426146 | 140187378 | 145977714 | 105846314 | 128481410 | 108636194 | 109695826 | 97306370 | 0 | 0 | 0 | 2 | 0 | 2 | 1.764214652 | 1.59255164 | 1.30410224 | 1.17720945 |
| P63276 | Rps17 | 40S ribosomal protein S17 | 673.51 | 66.67 | 15524 | 10 | 7666385 | 9606818 | 10011876 | 6619614 | 9509126 | 5866135 | 9619298 | 9125800 | 12875750 | 8991390 | 14263983 | 20198710 | 19690400 | 19744370 | 20699806 | 5482000.5 | 7975365 | 13627371 | 8279384.5 | 15141538 | 0 | 11 | 0 | 12 | 0 | 12 | 2.178966771 | 2.035296481 | 1.163354438 | 1.086648601 |
| P62270 | Rps18 | 40S ribosomal protein S18 | 921.28 | 67.11 | 17719 | 17 | 66810106 | 82964610 | 81242186 | 74724898 | 67565954 | 70254218 | 66919146 | 83780890 | 74357986 | 83271602 | 83863634 | 80887882 | 85065426 | 70957450 | 83233922 | 57324542 | 92354514 | 88773874 | 72038826 | 75216282 | 5 | 10 | 7 | 10 | 7 | 10 | 1.082239278 | 1.067156781 | 1.033217322 | 1.018818014 |
| P17074 | Rps19 | 40S ribosomal protein S19 | 638.56 | 54.48 | 16085 | 11 | 38710946 | 39534102 | 36789310 | 34843242 | 26225338 | 38875498 | 35388278 | 36034854 | 31809362 | 39631314 | 60782074 | 57711394 | 49952042 | 47571414 | 50652074 | 36655762 | 40941726 | 45946374 | 38067226 | 43469410 | 0 | 5 | 0 | 4 | 0 | 5 | 1.514279097 | 1.467316036 | 1.164548987 | 1.128432272 |
| Q8C6B9 | Rps19bp1 | Active regulator of SIRT1 | 136.37 | 22.38 | 15978 | 2 | 2 | 564864.94 | 312439.66 | 431924.91 | 323228.41 | 2 | 2 | 319665.69 | 420211.38 | 455671.03 | 534340.69 | 431877.28 | 323514.47 | 385974.34 | 413639.72 | 432151.88 | 2 | 468478.06 | 635825.94 | 469484.69 | 9 | 7.5 | 7 | 5 | 9 | 7.5 | 1.279876139 | 1.747599707 | 1.228785188 | 1.677837854 |
| P25444 | Rps2 | 40S ribosomal protein S2 | 960.17 | 52.9 | 31231 | 17 | 132219490 | 146763362 | 111191138 | 121924210 | 104001450 | 148599090 | 134877026 | 134017578 | 119291602 | 137493746 | 222510818 | 228394226 | 184462754 | 228779138 | 210035410 | 143669634 | 135663154 | 155764146 | 121340786 | 162900850 | 0 | 5 | 0 | 7 | 0 | 7 | 1.743520461 | 1.593082803 | 1.167568542 | 1.066826232 |
| P60867 | Rps20 | 40S ribosomal protein S20 | 183.56 | 25.21 | 13373 | 3 | 12978826 | 13550240 | 10523446 | 11035351 | 8864144 | 21405204 | 17623458 | 11536492 | 9663226 | 12542327 | 23840140 | 21187356 | 16854374 | 16037971 | 16796298 | 12482341 | 13328561 | 9099111 | 7238545.5 | 10551166 | 0 | 10 | 7 | 6 | 7 | 10 | 1.663086939 | 1.301569586 | 0.925335687 | 0.724188711 |
| P05765 | Rps21 | 40S ribosomal protein S21 | 291.21 | 59.04 | 9127 | 5 | 25574940 | 31173774 | 23938218 | 24560128 | 19897434 | 34245690 | 27175730 | 3968527 | 18238444 | 23850518 | 23558860 | 23357190 | 21335602 | 27879806 | 18749858 | 23737486 | 31225124 | 25769726 | 30471940 | 26991898 | 7 | 7 | 12 | 9 | 12 | 9 | 0.917989376 | 1.068873113 | 1.104292882 | 1.285798072 |
| P62267 | Rps23 | 40S ribosomal protein S23 | 366.08 | 48.95 | 15808 | 6 | 1471789.25 | 2779051 | 6946694 | 4369206 | 5683720 | 2600594.75 | 2432724.75 | 10628139 | 2929394 | 3791393 | 8013984.5 | 7275402.5 | 8931206 | 14134983 | 10678276 | 5100226 | 7907163 | 9637856 | 5354051 | 11940118 | 0 | 4 | 3 | 4 | 3 | 4 | 2.307425412 | 2.190747662 | 1.879461128 | 1.784423909 |
| P62849 | Rps24 | 40S ribosomal protein S24 | 317.63 | 35.34 | 15423 | 5 | 7779609 | 10424034 | 4968428 | 3457600.25 | 3098356.25 | 8805184 | 8207040 | 5509281 | 3866838 | 4560566 | 27407248 | 21248128 | 18351492 | 15650499 | 14951270 | 5673178 | 5671896 | 2529750.5 | 3168696 | 2950088.25 | 0 | 7 | 0 | 6 | 0 | 7 | 3.283387605 | 3.153863582 | 0.672550802 | 0.646019824 |
| P62852 | Rps25 | 40S ribosomal protein S25 | 309.13 | 27.2 | 13742 | 5 | 74740466 | 66851378 | 65518294 | 79753714 | 64779122 | 74375794 | 67572954 | 57744978 | 53960682 | 65548346 | 44985978 | 43677914 | 38681578 | 41912702 | 42932830 | 68210834 | 65849002 | 64611654 | 62019826 | 59876426 | 0 | 5 | 0 | 12 | 0 | 12 | 0.60342739 | 0.664753043 | 0.911628458 | 1.004276241 |
| P62855 | Rps26 | 40S ribosomal protein S26 | 210.31 | 20.87 | 13015 | 3 | 61373914 | 66496694 | 70983330 | 68631466 | 66796042 | 68861090 | 69673994 | 67996146 | 62220346 | 73096314 | 48215274 | 54967702 | 59984586 | 64170738 | 63038242 | 52319314 | 66505890 | 68442042 | 78930522 | 67013566 | 2 | 12 | 2 | 9 | 2 | 12 | 0.868658867 | 0.849432015 | 0.996798769 | 0.974735676 |
| P62982 | Rps27a | Ubiquitin-40S ribosomal protein S27a | 830.26 | 64.74 | 17951 | 13 | 346528642 | 299085602 | 202035842 | 289455746 | 200119106 | 299098754 | 277193634 | 218380098 | 184806434 | 286665282 | 373035330 | 252350722 | 313437730 | 406304482 | 306882498 | 276546626 | 538717058 | 331257986 | 367960002 | 291660290 | 5 | 6 | 3 | 4 | 5 | 6 | 1.235402298 | 1.304757199 | 1.350664283 | 1.426489936 |
| P24051 | Rps27l | 40S ribosomal protein S27-like | 186.84 | 26.19 | 9477 | 3 | 15804327 | 17299142 | 16003991 | 13967390 | 13454693 | 15400417 | 16189034 | 17624378 | 12734804 | 14237708 | 19105952 | 18597552 | 17340744 | 17313114 | 16873208 | 17237974 | 14104560 | 15183184 | 15680335 | 15768366 | 1 | 12 | 3 | 12 | 3 | 12 | 1.165962405 | 1.171214798 | 1.018879977 | 1.023469798 |
| P62858 | Rps28 | 40S ribosomal protein S28 | 220.66 | 46.38 | 7841 | 3 | 5482068 | 7701256.5 | 5106754 | 5134743.5 | 3735691.5 | 7543302.5 | 6270473.5 | 5097110.5 | 4620315 | 5530636 | 14258162 | 10677707 | 6977953.5 | 7657022 | 6328913.5 | 6556024 | 8254472 | 6097289.5 | 6922970.5 | 5960269.5 | 3 | 4 | 2 | 6 | 3 | 6 | 1.689944411 | 1.579382515 | 1.24412322 | 1.162728458 |
| P62908 | Rps3 | 40S ribosomal protein S3 | 1486.45 | 81.07 | 26674 | 25 | 348559362 | 331974050 | 299133090 | 341544738 | 289411170 | 390739586 | 329705858 | 296465602 | 287629346 | 320028482 | 395873090 | 334577474 | 328282050 | 338376258 | 316178146 | 346338690 | 424142178 | 350467906 | 356783074 | 347006082 | 10 | 2 | 7 | 4 | 10 | 4 | 1.063742195 | 1.054610269 | 1.132939613 | 1.123213647 |
| P49242 | Rps3a | 40S ribosomal protein S3a | 1212.84 | 52.27 | 29945 | 20 | 217755970 | 251055218 | 222922210 | 225058418 | 195240274 | 231895618 | 210259810 | 186803442 | 194513842 | 205554690 | 166661410 | 218984354 | 202986962 | 173267906 | 208131810 | 225696754 | 175608802 | 215110882 | 197981634 | 230576386 | 4 | 10 | 9 | 11 | 9 | 11 | 0.87230616 | 0.942669204 | 0.939698114 | 1.015497212 |
| P62702 | Rps4x | 40S ribosomal protein S4, X isoform | 1445.9 | 69.58 | 29598 | 25 | 134926738 | 161825138 | 137239554 | 107868186 | 98122490 | 142683410 | 138275330 | 132895514 | 117951090 | 133460978 | 228055442 | 230582226 | 196979362 | 176075506 | 195666578 | 130702506 | 115150122 | 131593218 | 126216538 | 132910570 | 0 | 10 | 0 | 5 | 0 | 10 | 1.605293499 | 1.544282463 | 0.994673051 | 0.956869351 |
| P97461 | Rps5 | 40S ribosomal protein S5 | 692.29 | 40.2 | 22889 | 10 | 94344818 | 90517682 | 83234082 | 91066194 | 72602634 | 93147394 | 89804258 | 87222946 | 72275434 | 87465842 | 89710218 | 91123282 | 74220290 | 75110162 | 73544722 | 98653010 | 114924018 | 89348282 | 103033394 | 92164610 | 9 | 4 | 10 | 3 | 10 | 4 | 0.935018565 | 0.939041097 | 1.153689718 | 1.158652993 |
| P62754 | Rps6 | 40S ribosomal protein S6 | 758.24 | 36.95 | 28681 | 10 | 8559045 | 18397554 | 16334593 | 14672886 | 7308984.5 | 13321723 | 11575501 | 16554768 | 9740905 | 14747013 | 61741774 | 44267190 | 25870918 | 42601434 | 25540082 | 9013057 | 19855524 | 17451118 | 13821894 | 12681002 | 0 | 10 | 0 | 10 | 0 | 10 | 3.064378939 | 3.033389005 | 1.115660767 | 1.104378138 |
| P18653 | Rps6ka1 | Ribosomal protein S6 kinase alpha-1 | 136.18 | 3.45 | 81595 | 2 | 925991.44 | 1106768.38 | 866476.44 | 712228.81 | 1567237.25 | 2 | 1004296.69 | 924545.31 | 1387298.12 | 707959.62 | 2 | 582952 | 494748.88 | 2 | 398986.31 | 696317.88 | 2 | 2 | 2 | 266189.59 | 0 | 0 | 4 | 3.5 | 4 | 3.5 | 0.285146954 | 0.366961694 | 0.185859972 | 0.239187161 |
| P62082 | Rps7 | 40S ribosomal protein S7 | 342.07 | 37.63 | 22127 | 7 | 38168246 | 34546778 | 37466070 | 41988334 | 27235076 | 44975918 | 33352782 | 37863226 | 31111702 | 40932986 | 61280718 | 38836270 | 41130162 | 39727938 | 43164934 | 37758870 | 59948266 | 40140538 | 39742238 | 37704502 | 3 | 6 | 6 | 10 | 6 | 10 | 1.249355601 | 1.190735518 | 1.200050217 | 1.143743555 |
| P62242 | Rps8 | 40S ribosomal protein S8 | 811.89 | 55.29 | 24205 | 12 | 101469178 | 110569450 | 99636418 | 95703322 | 88469770 | 104153346 | 100354914 | 87081946 | 83625466 | 85556050 | 94070922 | 87682602 | 61028106 | 59951138 | 61165186 | 37705414 | 39662934 | 22164404 | 19233892 | 17156836 | 1 | 0 | 6 | 0 | 6 | 0 | 0.733889927 | 0.789757567 | 0.274123203 | 0.294990933 |
| P29314 | Rps9 | 40S ribosomal protein S9 | 869.05 | 52.58 | 22591 | 17 | 59167042 | 78397690 | 114224442 | 78293546 | 135403106 | 44239842 | 68772026 | 124574402 | 144147730 | 97838778 | 115823450 | 132945074 | 106560402 | 87536650 | 103615722 | 46598418 | 35863366 | 88285442 | 77260298 | 81862538 | 8 | 7 | 10 | 7 | 10 | 7 | 1.174002016 | 1.139516926 | 0.708657586 | 0.687841506 |
| P14206 | Rpsa | 40S ribosomal protein SA | 1004.15 | 55.93 | 32838 | 14 | 39552950 | 39643370 | 40869158 | 34886698 | 30784336 | 34651310 | 31271908 | 33407126 | 30563100 | 45537182 | 166420482 | 117667602 | 89669994 | 88936682 | 99037282 | 34641590 | 42945042 | 43497398 | 37912674 | 36192738 | 0 | 10 | 0 | 6 | 0 | 10 | 3.024349041 | 3.202018113 | 1.050894301 | 1.112630368 |
| Q8K4Q0 | Rptor | Regulatory-associated protein of mTOR | 69.31 | 1.42 | 149471 | 1 | 2 | 86198.84 | 158243.36 | 142620.06 | 153469.33 | 97977.07 | 88113.13 | 118015.31 | 115864.62 | 2 | 64775.38 | 212346.19 | 250037.33 | 278298.88 | 246871.92 | 152621.03 | 112144.02 | 176399.91 | 126119.21 | 194086.7 | 4 | 8 | 4 | 2 | 4 | 8 | 1.946834978 | 2.505713177 | 1.408554221 | 1.812908085 |
| Q63486 | Rraga | Ras-related GTP-binding protein A | 429.75 | 34.5 | 36566 | 7 | 5441339 | 6623591.5 | 5997288 | 8470086 | 7383029 | 5988673 | 6408659 | 6470891 | 6177374.5 | 8123683 | 9010431 | 7269356 | 8928903 | 8155160.5 | 8093535 | 5414666 | 6750714.5 | 7361039.5 | 8678438 | 8175548.5 | 4 | 10 | 2 | 7 | 4 | 10 | 1.222378825 | 1.249872921 | 1.072683142 | 1.09681024 |
| Q99K70 | Rragc | Ras-related GTP-binding protein C | 343.81 | 17.59 | 44121 | 6 | 7221371 | 8656698 | 8959086 | 3672536 | 13839950 | 2503904.25 | 9961906 | 10234323 | 14051282 | 4308855.5 | 3558735 | 9837712 | 7315492 | 4751244 | 7766280 | 8603456 | 2517215.5 | 3209616 | 3516225 | 2905702.5 | 9 | 2 | 9 | 6 | 9 | 6 | 0.784645683 | 0.809285043 | 0.490021037 | 0.505408625 |
| P10833 | Rras | Ras-related protein R-Ras | 544.04 | 39.91 | 23764 | 9 | 9405102 | 8291536.5 | 7386057 | 7902766.5 | 11071432 | 9557019 | 8961949 | 9005553 | 8846654 | 8630120 | 8268202 | 8143514 | 9435614 | 8684972 | 9617284 | 8966024 | 6808417 | 8682965 | 8134239.5 | 7341599 | 10 | 8 | 10 | 4 | 10 | 8 | 1.002103916 | 0.981073678 | 0.90640172 | 0.887379897 |
| Q99PL5 | Rrbp1 | Ribosome-binding protein 1 | 1944.78 | 20.93 | 172879 | 32 | 84019658 | 87183882 | 61799698 | 68681394 | 62309134 | 92410674 | 90738066 | 66249690 | 61493098 | 64416754 | 57677234 | 59390538 | 37609762 | 41816170 | 43344074 | 105951170 | 107665218 | 73603698 | 82960746 | 78521610 | 0 | 6 | 0 | 6 | 0 | 6 | 0.658906279 | 0.639042061 | 1.232720129 | 1.195556995 |
| P07742 | Rrm1 | Ribonucleoside-diphosphate reductase large subunit | 170.33 | 6.31 | 90210 | 4 | 2540730 | 651897.12 | 3157018 | 2340451.5 | 2392762 | 1963065.25 | 2413208.75 | 2157141.25 | 1986422.5 | 2708402.25 | 193121.7 | 1212229 | 1626405.75 | 1488615.62 | 1122542.5 | 2438012 | 2225532.75 | 1745919.25 | 605482.25 | 1474614.12 | 4 | 6 | 0 | 7 | 4 | 7 | 0.509156957 | 0.502564478 | 0.766008181 | 0.756090035 |
| P11157 | Rrm2 | Ribonucleoside-diphosphate reductase subunit M2 | 263.34 | 9.74 | 45096 | 3 | 3181308.5 | 3103195.75 | 2425905.75 | 2758342.5 | 2154076.5 | 3279557.5 | 2895177.5 | 2024701.5 | 1959319 | 2306456.25 | 1798755.25 | 1784488.25 | 854542.75 | 1584438.75 | 567508.19 | 3182356.5 | 3783539 | 2343587 | 2119742 | 1920030.25 | 0 | 11 | 0 | 11 | 0 | 11 | 0.483727219 | 0.52864992 | 0.979917956 | 1.070920817 |
| Q6P5B0 | Rrp12 | RRP12-like protein | 74.93 | 1.78 | 143131 | 1 | 169517.47 | 2 | 2 | 2 | 124676.45 | 2 | 2 | 2 | 2 | 2 | 302619.78 | 237375.28 | 281383.62 | 412541.53 | 329021.41 | 2 | 2 | 2 | 2 | 2 | 0 | 7.5 | 0 | 12.5 | 0 | 12.5 | 5.312515449 | 156294.162 | 3.399049E-05 | 1 |
| Q5M947 | Rrp15 | RRP15-like protein | 119.05 | 6.79 | 30926 | 2 | 320898.09 | 706679.69 | 2 | 2 | 2 | 2 | 2 | 2 | 2 | 575549.31 | 797622.31 | 2 | 331406.12 | 2 | 2 | 902635.06 | 917645.5 | 2 | 2 | 2 | 11.5 | 10.5 | 10 | 9 | 11.5 | 10.5 | 1.098727376 | 1.961636852 | 1.77142399 | 3.162650406 |
| Q91YK2 | Rrp1b | Ribosomal RNA processing protein 1 homolog B | 164.75 | 4.83 | 80582 | 3 | 885470.25 | 1338587.62 | 1133953.75 | 735808.88 | 585837.38 | 1334884.75 | 993108.81 | 716538.62 | 1019275.75 | 569168.12 | 893004.62 | 1041727.81 | 568975.75 | 670000.06 | 517617.81 | 1598231.25 | 1339775 | 1239611.12 | 893089.19 | 854082.56 | 7 | 6 | 7 | 7 | 7 | 7 | 0.788802546 | 0.796750514 | 1.266073134 | 1.278830077 |
| Q91WM3 | Rrp9 | U3 small nucleolar RNA-interacting protein 2 | 95.55 | 5.05 | 52107 | 2 | 363805.66 | 473543.38 | 425381.44 | 427391.78 | 384977.81 | 458874.5 | 427053.69 | 2 | 514519.56 | 540498.12 | 542539.5 | 1005943.44 | 591390.12 | 672489.19 | 785815.5 | 469372.88 | 418737.62 | 576220.69 | 2 | 645149.75 | 0 | 9 | 0 | 10.5 | 0 | 10.5 | 1.733977943 | 1.853825033 | 1.016569259 | 1.086831322 |
| Q9CYH6 | Rrs1 | Ribosome biogenesis regulatory protein homolog | 276.92 | 13.97 | 41552 | 5 | 1131972.5 | 1476569 | 1476071.62 | 879592.5 | 1231039.25 | 1155563 | 1835926.75 | 1079656.75 | 1034957.88 | 898773.06 | 4734407.5 | 3921502.5 | 2105275.5 | 2219920.5 | 3252690.75 | 1139418.5 | 1636476.5 | 1117142.75 | 1253305.62 | 1634018 | 0 | 9 | 0 | 7 | 0 | 9 | 2.62036402 | 2.703435151 | 1.094446065 | 1.129142341 |
| Q8BVY0 | Rsl1d1 | Ribosomal L1 domain-containing protein 1 | 1072.2 | 30.53 | 50421 | 15 | 12525552 | 17197862 | 14468086 | 17039706 | 7808994.5 | 19277084 | 12216742 | 17503144 | 13973891 | 23560136 | 31268438 | 24233662 | 23961430 | 29965902 | 25035312 | 15650673 | 17715812 | 22835416 | 24177676 | 18588116 | 0 | 2 | 0 | 8 | 0 | 8 | 1.947629686 | 1.553948858 | 1.433479223 | 1.143725329 |
| Q01730 | Rsu1 | Ras suppressor protein 1 | 181.97 | 20.58 | 31550 | 4 | 5233442.5 | 4247136 | 4905571 | 2809467 | 5192786.5 | 2003213.62 | 4444730 | 3923886 | 4382350.5 | 3227243.75 | 874575 | 3193500.25 | 1272742 | 520025.88 | 1753797.75 | 3940799.25 | 1845003.75 | 2034958.38 | 1811618.75 | 2378603.5 | 1 | 1 | 1 | 5 | 1 | 5 | 0.340115411 | 0.423472631 | 0.536482376 | 0.667966214 |
| Q9D7H3 | RtcA | RNA 3'-terminal phosphate cyclase | 315.33 | 19.95 | 39254 | 5 | 2550319 | 2651029.5 | 2489997.5 | 2769123.5 | 2473752.75 | 3836092.5 | 2951871.5 | 1303655.5 | 3458873.75 | 3266991.75 | 4561338.5 | 3728036.5 | 5439276 | 1581357.12 | 3837093.5 | 3102336.25 | 2694428.75 | 4227264 | 4530229 | 2721738.25 | 5 | 2 | 5 | 11 | 5 | 11 | 1.480344256 | 1.292196457 | 1.335681104 | 1.165919604 |
| Q6AYT3 | Rtcb | tRNA-splicing ligase RtcB homolog | 669.83 | 29.5 | 55249 | 14 | 20345266 | 18515768 | 13131262 | 19635250 | 16184950 | 21249302 | 20994016 | 18971988 | 16670135 | 17449726 | 18045398 | 16980720 | 13381151 | 14369938 | 14867545 | 23082606 | 25329648 | 20639528 | 20043030 | 21229274 | 7 | 1 | 3 | 5 | 7 | 5 | 0.884210739 | 0.814439775 | 1.256359755 | 1.157223399 |
| Q3T1J8 | Rtfdc1 | Protein RTF2 homolog | 49.34 | 3.27 | 33812 | 1 | 2 | 2 | 340331 | 2 | 390358.12 | 2 | 265881.47 | 2 | 401211.5 | 2 | 2 | 234428.39 | 2 | 2 | 2 | 2 | 2 | 2 | 2 | 2 | 9 | 7.5 | 9 | 7.5 | 9 | 7.5 | 0.32084023 | 0.351426701 | 1.36856E-05 | 1.499028E-05 |

| | | | | | | | | | | | | | | | | | | | | | | | | | | | | | | | | | | | | |
|---|---|---|---|---|---|---|---|---|---|---|---|---|---|---|---|---|---|---|---|---|---|---|---|---|---|---|---|---|---|---|---|---|---|---|---|---|
| Q9ES97 | Rtn3 | Reticulon-3 | 222.47 | 7.37 | 103879 | 4 | 21774568 | 18691292 | 21911046 | 26600386 | 23987002 | 26728948 | 18778148 | 24104686 | 21210296 | 31676760 | 16004878 | 12916179 | 20616366 | 23634562 | 20293514 | 23161982 | 30989366 | 33139572 | 32652858 | 33644494 | 5 | 2 | 4 | 4 | 5 | 4 | 0.82738975 | 0.762990903 | 1.359617863 | 1.253793705 |
| Q99P72 | Rtn4 | Reticulon-4 | 1181.88 | 22.2 | 126613 | 17 | 59608790 | 55170834 | 52711738 | 59402866 | 61431926 | 70111346 | 65731314 | 63068474 | 62195446 | 68134098 | 47164086 | 49820166 | 55193914 | 55655506 | 64511938 | 57462874 | 67680338 | 63227642 | 67245930 | 66565998 | 9 | 3 | 2 | 11 | 9 | 11 | 0.944574768 | 0.827193078 | 1.117424755 | 0.978563111 |
| Q9CQE8 | RTRAF | RNA transcription, translation and transport factor protein | 482.15 | 37.7 | 28152 | 8 | 2412755.75 | 2993737.25 | 3249002.25 | 1596407.12 | 4875344.5 | 1458948.25 | 3239405.5 | 2705338 | 4595946 | 1878315.5 | 382856.06 | 1719776.75 | 719565.44 | 922486.31 | 1208036.88 | 2219269.5 | 1610793.38 | 1330839.25 | 1849604.62 | 2122728.75 | 1 | 4 | 1 | 6 | 1 | 6 | 0.32740402 | 0.356876936 | 0.60376059 | 0.65811113 |
| Q8BIJ7 | Rufy1 | RUN and FYVE domain-containing protein 1 | 236.32 | 7.58 | 80376 | 5 | 3310324 | 3152040.5 | 3095710.75 | 5668589.5 | 4988930 | 2895603.5 | 4066053.5 | 4339435 | 3715427 | 2969100 | 1132675.5 | 2196545.75 | 1643968.25 | 2146838.5 | 1942653.88 | 2738147 | 2267570.5 | 4409835 | 3501089.5 | 4780345 | 0 | 9 | 0 | 12 | 0 | 12 | 0.448301521 | 0.503884903 | 0.875412632 | 0.983952067 |
| Q03347 | Runx1 | Runt-related transcription factor 1 | 45.77 | 1.55 | 48610 | 1 | 2 | 820748.06 | 485842.75 | 91999.77 | 386583.09 | 2 | 1287830.38 | 736021.88 | 1525080.62 | 2 | 526785.81 | 382767.88 | 1019598.56 | 2 | 1284044.12 | 334424.59 | 1554380 | 484602.78 | 474638.69 | 1165674.25 | 8.5 | 7 | 11 | 11 | 11 | 11 | 1.799933992 | 0.905397441 | 2.248361535 | 1.13096413 |
| P60122 | Ruvbl1 | RuvB-like 1 | 891.48 | 39.69 | 50214 | 16 | 21169430 | 22514924 | 21319002 | 24075628 | 20066322 | 23467716 | 20196014 | 20059222 | 21600058 | 22857650 | 26000134 | 25986380 | 24600252 | 24747890 | 27922516 | 22149760 | 19734388 | 14346254 | 23154966 | 23509356 | 0 | 11 | 0 | 12 | 0 | 12 | 1.184266889 | 1.194826987 | 0.942731555 | 0.951137884 |
| Q9WTM5 | Ruvbl2 | RuvB-like 2 | 1178.63 | 52.27 | 51113 | 18 | 23127862 | 26879614 | 24735270 | 21821776 | 21499636 | 23595598 | 24770486 | 20748286 | 25418052 | 27539686 | 38637762 | 38122930 | 31110866 | 26699648 | 34679290 | 24790096 | 25157960 | 24061224 | 24927234 | 24285848 | 1 | 7 | 1 | 12 | 1 | 12 | 1.433546801 | 1.38647967 | 1.043689839 | 1.009422742 |
| Q8CCI5 | Rybp | RING1 and YY1-binding protein | 177.34 | 28.51 | 24777 | 3 | 374475.91 | 1092868.5 | 1676324 | 1977373.75 | 1106544.25 | 1478197.25 | 1205570.88 | 980146 | 1220637.62 | 1227461.62 | 2 | 305352.28 | 914365.69 | 1362424.75 | 642484.12 | 783378.88 | 1038432.62 | 1312756.5 | 675202.81 | 1177622.88 | 5 | 9 | 4 | 6 | 5 | 9 | 0.517797527 | 0.527588643 | 0.800854996 | 0.815998492 |
| P56565 | S100a1 | Protein S100-A1 | 51.04 | 19.15 | 10505 | 1 | 676743.19 | 390121.16 | 607398.56 | 2 | 808424.31 | 2 | 488628.06 | 566254.94 | 627059.31 | 416537.12 | 2 | 204616.44 | 2 | 2 | 228255.8 | 615383 | 189799.42 | 345913.88 | 307329.41 | 430541.66 | 3.5 | 8 | 3.5 | 9 | 3.5 | 9 | 0.174358609 | 0.206281663 | 0.760855348 | 0.900159202 |
| P05943 | S100a10 | Protein S100-A10 | 98.91 | 17.89 | 11075 | 1 | 11001008 | 9393460 | 7963365.5 | 8564046 | 10712490 | 9116927 | 10527626 | 6492688.5 | 9769004 | 9487778 | 2356027.25 | 2953880.75 | 3011131.25 | 1526155.5 | 1747811.62 | 9678669 | 3971003.75 | 7388694.5 | 7632011.5 | 6368850.5 | 0 | 3 | 0 | 5 | 0 | 5 | 0.243416812 | 0.255430241 | 0.735587132 | 0.771890803 |
| P50543 | S100a11 | Protein S100-A11 | 403.69 | 54.08 | 11083 | 6 | 10671228 | 10670198 | 12432641 | 15443588 | 10546433 | 10812181 | 12204094 | 13666889 | 11308174 | 12880784 | 5095480.5 | 6751019 | 8899674 | 9728116 | 8814536 | 11772969 | 11930135 | 13466477 | 14600459 | 13965316 | 0 | 7 | 0 | 7 | 0 | 7 | 0.657398562 | 0.645432165 | 1.099913982 | 1.079892631 |
| P07091 | S100a4 | Protein S100-A4 | 493.93 | 51.49 | 11721 | 9 | 88327370 | 116157114 | 163944466 | 143027602 | 114054770 | 81118722 | 86242946 | 133616802 | 115081586 | 131295946 | 57573170 | 76519338 | 76355466 | 65711698 | 75551826 | 75056370 | 70227930 | 90545482 | 77844002 | 110398274 | 0 | 2 | 0 | 4 | 0 | 4 | 0.562278388 | 0.642564431 | 0.677960643 | 0.774764607 |
| P14069 | S100a6 | Protein S100-A6 | 387.42 | 78.65 | 10051 | 6 | 119124594 | 98337914 | 106512554 | 164490178 | 67167138 | 105661602 | 110420474 | 124099138 | 121879874 | 149827522 | 39358382 | 12969644 | 22147996 | 21893664 | 24985826 | 106061162 | 69931898 | 54505910 | 121893946 | 106547890 | 0 | 10 | 0 | 5 | 0 | 10 | 0.21840972 | 0.198329418 | 0.825979234 | 0.750039792 |
| Q9EP69 | Sacm1l | Phosphatidylinositide phosphatase SAC1 | 370.79 | 11.75 | 66944 | 6 | 7586786.5 | 8751889 | 6932019.5 | 8775891 | 8888231 | 9471690 | 9048156 | 9396998 | 6992678.5 | 8723046 | 9730185 | 8877567 | 11690291 | 10376453 | 10160118 | 7032398 | 7315409 | 10126378 | 9834976 | 10451977 | 1 | 8 | 3 | 8 | 3 | 8 | 1.241842952 | 1.165061232 | 1.093473509 | 1.025865301 |
| Q9R1T2 | Sae1 | SUMO-activating enzyme subunit 1 | 519.05 | 22.57 | 38620 | 8 | 7198050 | 7096838.5 | 9900424 | 8190355.5 | 8891356 | 9487016 | 6953776 | 7704534 | 8103181.5 | 9218402 | 8748524 | 7432218 | 8139818 | 2351886.25 | 7638378 | 7420605.5 | 5611648 | 8598028 | 10681371 | 8934089 | 9 | 11 | 8 | 12 | 9 | 12 | 0.831232994 | 0.827426607 | 0.999242133 | 0.994666398 |
| D3YXK2 | Safb | Scaffold attachment factor B1 | 440.98 | 10.99 | 105104 | 8 | 2358497.25 | 3100693 | 2214554.5 | 1799454.25 | 2012680.75 | 3266018.75 | 2613414 | 1513298.75 | 865839.12 | 1684394 | 1579759.5 | 1724517.88 | 867551.81 | 1395130.38 | 753799.38 | 2692943 | 3069591.75 | 1209091.38 | 2216057 | 2414951.5 | 0 | 10 | 7 | 10 | 7 | 10 | 0.550306906 | 0.635701643 | 1.010165079 | 1.16691903 |
| Q80YR5 | Safb2 | Scaffold attachment factor B2 | 373.01 | 9.49 | 111838 | 7 | 1296563 | 2270550.75 | 1875448 | 1655078.75 | 869381 | 1769176.5 | 1783822.75 | 1647474.75 | 2042319.25 | 1516406.75 | 1437138.12 | 1476064 | 1551873.5 | 584199.5 | 1197897 | 2112390.5 | 1916743 | 1799332.62 | 2300885.25 | 1691418.25 | 7 | 6 | 1 | 5 | 7 | 6 | 0.78412894 | 0.713212636 | 1.232677685 | 1.121194815 |
| Q69Z37 | Samd9l | Sterile alpha motif domain-containing protein 9-like | 104.82 | 1.86 | 180203 | 2 | 2 | 2 | 2 | 2 | 77222.45 | 2 | 2 | 2 | 85804.77 | 39294.83 | 2 | 39770.59 | 133210.17 | 2 | 62791.36 | 2 | 2 | 88987.3 | 2 | 2 | 8 | 12 | 9 | 11 | 9 | 12 | 3.052890667 | 1.884616836 | 1.152334345 | 0.711361442 |
| Q60710 | Samhd1 | Deoxynucleoside triphosphate triphosphohydrolase SAMHD1 | 555.56 | 26.14 | 75893 | 11 | 7444854 | 5846682.5 | 6033026.5 | 8582454 | 13852412 | 7235161.5 | 6962161 | 9855420 | 11346352 | 9843856 | 2625795.5 | 1700432.5 | 1853306.12 | 1299092 | 2552601.25 | 8640591 | 4774947 | 6330687.5 | 6968790 | 7195451 | 0 | 10 | 0 | 4 | 0 | 10 | 0.240214668 | 0.221719124 | 0.812043347 | 0.749519342 |
| Q8BGH2 | Samm50 | Sorting and assembly machinery component 50 homolog | 733.11 | 31.56 | 51864 | 13 | 15020742 | 15960356 | 12920978 | 14849282 | 14670690 | 17489704 | 16224836 | 16038938 | 18184382 | 19100916 | 32387718 | 26557496 | 29954280 | 26219598 | 27886112 | 15113241 | 16554093 | 22016088 | 14802404 | 18027764 | 0 | 4 | 0 | 10 | 0 | 10 | 1.947714725 | 1.643005687 | 1.178305323 | 0.993966069 |
| O55128 | Sap18 | Histone deacetylase complex subunit SAP18 | 289.15 | 39.87 | 17595 | 5 | 605760.81 | 1271467.25 | 121106.09 | 199374.56 | 2 | 2 | 164574.25 | 2 | 77595.45 | 169751.56 | 461447.88 | 992187.69 | 102184.52 | 2 | 132723.91 | 407055.25 | 894276.12 | 2 | 254974.8 | 543965.69 | 10.5 | 11.5 | 8 | 4 | 10.5 | 11.5 | 0.768320413 | 4.09915624 | 0.955664388 | 5.098677027 |
| P36536 | Sar1a | GTP-binding protein SAR1a | 218.81 | 28.79 | 22371 | 4 | 2 | 2 | 2 | 1487042.38 | 78065 | 2205663 | 2 | 2 | 2 | 492340.78 | 1618179.12 | 2 | 1123737.25 | 2 | 477562.19 | 2 | 899985.38 | 1766347.25 | 1198863.25 | 1116290.5 | 9 | 6.5 | 11 | 7.5 | 11 | 7.5 | 2.057028329 | 1.193280537 | 3.18282908 | 1.84635668 |
| Q5HZY2 | Sar1b | GTP-binding protein SAR1b | 153.04 | 18.18 | 22410 | 3 | 2563497 | 2448004.75 | 2489419.5 | 2663990 | 3523273.75 | 2622371 | 2774789 | 3087905.5 | 3510372 | 2640752.5 | 1683966.75 | 2310315.75 | 1939122.75 | 1454351.75 | 2269699 | 2378401 | 2811529.5 | 2846717.25 | 2760782.25 | 2742306.25 | 0 | 9 | 0 | 10 | 0 | 10 | 0.705532253 | 0.659834014 | 0.989154972 | 0.925086122 |
| Q498U4 | Sarnp | SAP domain-containing ribonucleoprotein | 413.96 | 34.29 | 23605 | 6 | 6865031 | 7213109.5 | 6653731.5 | 6881309.5 | 3694967.25 | 5459687 | 6026840 | 5596412 | 5678481 | 6988612.5 | 8723149 | 12616195 | 8230035.5 | 6659188.5 | 9608730 | 3840273 | 5871609 | 7792794.5 | 5392251 | 8470118 | 3 | 12 | 1 | 12 | 3 | 12 | 1.464069254 | 1.540747829 | 1.001881196 | 1.054353319 |
| P26638 | Sars | Serine--tRNA ligase, cytoplasmic | 1470.6 | 48.63 | 58389 | 24 | 77054826 | 81006650 | 59921162 | 73771042 | 69507762 | 88254218 | 82434162 | 92004146 | 90575170 | 96642722 | 118495394 | 119375442 | 122594314 | 106728194 | 118132714 | 87444866 | 100045874 | 94118394 | 102344306 | 101557522 | 0 | 0 | 0 | 5 | 0 | 5 | 1.620228427 | 1.300983562 | 1.343932414 | 1.079128072 |
| Q5XIW8 | Sart1 | U4/U6.U5 tri-snRNP-associated protein 1 | 208.15 | 6.33 | 91011 | 4 | 3237630.5 | 2773134.5 | 3224954 | 1402528.75 | 2436161.5 | 2150430.5 | 2275486.5 | 2911863.75 | 1824526 | 2263786 | 1542622.62 | 2696085 | 1301621.5 | 862361.88 | 1383761.75 | 2717102.5 | 1707680 | 1099355 | 2156516.5 | 1714879.5 | 3 | 5 | 4 | 6 | 4 | 6 | 0.595549107 | 0.681462414 | 0.718620117 | 0.822287523 |
| Q9JLI8 | Sart3 | Squamous cell carcinoma antigen recognized by T-cells 3 | 141.57 | 4.26 | 109619 | 3 | 1183100.62 | 1326538.75 | 350120.09 | 1476966.5 | 1218214.5 | 1041937.75 | 1028392.19 | 1450672.75 | 1318730 | 1310574.5 | 893452 | 1153336.25 | 1088245.62 | 876239.69 | 1039569.12 | 1104418.5 | 943187.06 | 1459900.25 | 963898.75 | 1302844.38 | 5 | 10 | 5 | 9 | 5 | 10 | 0.909252352 | 0.821234212 | 1.039479898 | 0.938855371 |
| P70122 | Sbds | Ribosome maturation protein SBDS | 40.97 | 3.6 | 28781 | 1 | 2 | 1324472.5 | 1260116.62 | 1115398.38 | 2 | 998933.38 | 1225124.12 | 2 | 1041147.94 | 1241952.25 | 1080448.38 | 684199.38 | 1073480.88 | 2 | 2 | 1836788.75 | 2 | 2 | 2 | 2 | 8 | 9 | 8 | 7 | 8 | 9 | 0.767064638 | 0.629694272 | 0.496432695 | 0.40752866 |
| Q6ZPE2 | Sbf1 | Myotubularin-related protein 5 | 62.11 | 0.54 | 208693 | 1 | 603692.5 | 610769.12 | 614214.81 | 581255.62 | 623971.81 | 822218.62 | 705932.44 | 682210.25 | 829181.25 | 640015.38 | 539204.5 | 518415.03 | 715339.56 | 861355.5 | 475254.97 | 636797.12 | 593921.06 | 663124.62 | 676732.25 | 711932.06 | 10 | 4 | 8 | 5 | 10 | 5 | 1.024940045 | 0.845093245 | 1.081941703 | 0.89209279 |
| Q63623 | Scaf8 | Protein SCAF8 | 75.3 | 0.95 | 139504 | 1 | 990841.06 | 982865.5 | 1003612.62 | 945580.38 | 1120100.5 | 869558.06 | 927541.56 | 920176.62 | 1256072.25 | 828604.81 | 667548.56 | 621888.12 | 784115.81 | 2 | 800262.06 | 894869.69 | 498700.78 | 762440.12 | 632900.06 | 618342.56 | 0 | 0 | 0 | 2 | 0 | 2 | 0.569862486 | 0.598468242 | 0.675640129 | 0.709555674 |
| Q9ERN0 | Scamp2 | Secretory carrier-associated membrane protein 2 | 89.49 | 6.08 | 36465 | 1 | 2 | 2 | 2 | 2 | 2 | 2 | 2 | 2 | 2 | 2 | 611043.38 | 290807.75 | 2 | 432939.22 | 2 | 2 | 2 | 2 | 2 | 2 | 5 | 12.5 | 5 | 12.5 | 5 | 12.5 | 133479.435 | 133479.435 | 1 | 1 |
| O35609 | Scamp3 | Secretory carrier-associated membrane protein 3 | 338.71 | 17.19 | 38458 | 4 | 289626.97 | 687346.19 | 1696406.62 | 1303280.88 | 2 | 1808945.75 | 1691631.25 | 1727114.75 | 1212598.25 | 2141305.5 | 4791401 | 5599834 | 6499896.5 | 5779287 | 5510377 | 824772.5 | 1974613.88 | 3148148 | 1375415.12 | 2016337.88 | 0 | 3 | 0 | 11 | 0 | 11 | 7.086544148 | 3.283864347 | 2.34852392 | 1.088292658 |
| O35114 | Scarb2 | Lysosome membrane protein 2 | 297.19 | 13.18 | 54044 | 4 | 2282379 | 1792893.75 | 918774.62 | 1486002.75 | 2014082.5 | 2113843 | 1531629.25 | 2852010.25 | 3571978 | 2003659.5 | 2721037.5 | 3379849.5 | 3244550.5 | 4271344 | 2965426.25 | 2866306 | 2094200 | 3597808.25 | 2096494.25 | 2855979.5 | 0 | 2 | 5 | 8 | 5 | 8 | 1.952195532 | 1.373481565 | 1.590602432 | 1.119080072 |
| Q8R127 | Sccpdh | Saccharopine dehydrogenase-like oxidoreductase | 270.94 | 20.05 | 47129 | 6 | 2725873.75 | 2120487 | 1642000.75 | 639801.88 | 1641921.62 | 1635942.75 | 2517243 | 1750709.25 | 2082845.5 | 771968.75 | 2353159.75 | 2224423.5 | 2730510.75 | 2541627.5 | 2369183.25 | 819529.62 | 1949240.75 | 2248493 | 2480847 | 1883097.5 | 4 | 10 | 3 | 10 | 4 | 10 | 1.393248156 | 1.395057696 | 1.069682662 | 1.071071958 |
| Q8BRF7 | Scfd1 | Sec1 family domain-containing protein 1 | 354.79 | 18.31 | 72323 | 7 | 2608326.75 | 2644161.75 | 2916612.25 | 2024707 | 2498712.75 | 2583842.5 | 2125898 | 1344516.75 | 2160425 | 2470964 | 3094923.5 | 2945792.25 | 2106827 | 1058899.62 | 2070743.62 | 2578827.75 | 2060298.12 | 2056306.5 | 2224588 | 1908325.75 | 12 | 5 | 12 | 10 | 12 | 10 | 0.888490666 | 1.05535835 | 0.853128117 | 1.013354351 |
| Q9JLI6 | Scly | Selenocysteine lyase | 402.04 | 18.06 | 47174 | 7 | 7425405 | 9750254 | 10050072 | 10042924 | 9260524 | 7577197.5 | 8526412 | 7397674.5 | 7318078 | 7392763 | 2229914.25 | 4899934 | 4693641.5 | 4191719 | 4401672.5 | 8648386 | 7943784 | 8279063.5 | 8696999 | 6899012.5 | 0 | 4 | 0 | 7 | 0 | 7 | 0.438797367 | 0.534303739 | 0.869717581 | 1.059015823 |
| P32020 | Scp2 | Non-specific lipid-transfer protein | 358.07 | 9.14 | 59126 | 6 | 6592078.5 | 7252175.5 | 8965610 | 9652925 | 7954333 | 4571237 | 5893385 | 7148682.5 | 6628815 | 6797775 | 4333670 | 6134082 | 7986403.5 | 7354191.5 | 7591911 | 8141533 | 6990909.5 | 7229456.5 | 7670108 | 7644862 | 7 | 9 | 8 | 1 | 8 | 9 | 0.826388826 | 1.076042897 | 0.93220069 | 1.213820782 |
| Q920A5 | Scpep1 | Retinoid-inducible serine carboxypeptidase | 460.22 | 16.81 | 50965 | 7 | 22047296 | 19227024 | 14002990 | 16087729 | 16378486 | 17345966 | 18864862 | 15306315 | 17257936 | 16338235 | 12910629 | 14662654 | 14695024 | 11396414 | 15659937 | 22309040 | 21942278 | 20055088 | 20606080 | 20603684 | 3 | 4 | 1 | 0 | 3 | 4 | 0.790082892 | 0.814498399 | 1.202552211 | 1.239714036 |
| Q5M9F8 | Scyl1 | N-terminal kinase-like protein | 41.2 | 0.87 | 89126 | 1 | 1132899.5 | 876070.62 | 1381842.25 | 1388516.25 | 1147327.88 | 1230261.62 | 792726.88 | 1356292 | 1134871.25 | 1138138.12 | 1439083.12 | 1178870.38 | 1503201.62 | 1392559.62 | 1376603.12 | 1025555.31 | 1865056.5 | 1850907.75 | 1279979.75 | 1285944.12 | 4 | 8 | 2 | 6 | 4 | 8 | 1.162597809 | 1.219031228 | 1.232979072 | 1.292828853 |
| Q8CFE4 | Scyl2 | SCY1-like protein 2 | 43.38 | 1.18 | 103317 | 1 | 173464.62 | 280603.22 | 181580.8 | 2 | 1123364.88 | 2 | 255197.89 | 275600.19 | 962996.44 | 2 | 2 | 112434.88 | 2 | 55492.62 | 2 | 214666.56 | 2 | 2 | 2 | 2 | 3.5 | 5 | 7 | 6 | 7 | 6 | 0.095470164 | 0.112420449 | 0.122042448 | 0.143710519 |
| Q9DBQ7 | Scyl3 | Protein-associating with the carboxyl-terminal domain of ezrin | 93.94 | 3.95 | 81333 | 2 | 1181338.5 | 1011650.69 | 1088263 | 1181949.25 | 1443407.25 | 1151600 | 1309330.5 | 1396113.75 | 1152293.12 | 954630.5 | 619290.94 | 966185.5 | 792917.94 | 377741.94 | 783893.88 | 1026389.81 | 748272.69 | 1033415.62 | 1244954.88 | 1020811.25 | 0 | 7 | 1 | 6 | 1 | 7 | 0.599333795 | 0.59356963 | 0.859011408 | 0.850749763 |
| O08992 | Sdcbp | Syntenin-1 | 702.34 | 53.18 | 32379 | 11 | 3832622.5 | 5880057 | 3926586 | 4341193 | 3925483 | 5630713 | 6097584 | 6470009 | 7865225.5 | 7833768.5 | 40593762 | 35881806 | 48236338 | 44669734 | 35225198 | 4435854 | 6549850 | 9149730 | 8121668.5 | 9681491 | 0 | 1 | 0 | 7 | 0 | 7 | 9.34024397 | 6.036080691 | 1.731886005 | 1.119221693 |
| Q9ESP1 | Sdf2l1 | Stromal cell-derived factor 2-like protein 1 | 207.62 | 19.46 | 23648 | 3 | 314453.12 | 2 | 3097009.25 | 3082743.5 | 2398559 | 2115213.75 | 321910.03 | 2078074.62 | 2369989.25 | 2549111.75 | 366600.91 | 3146461 | 3990298.5 | 5287316 | 5021196.5 | 1006644.38 | 2 | 3942794.25 | 450094.31 | 3897467.75 | 3 | 10.5 | 4 | 12 | 4 | 12 | 2.002961864 | 1.887991058 | 1.045456698 | 0.985447069 |
| Q8K2B3 | Sdha | Succinate dehydrogenase [ubiquinone] flavoprotein subunit, mitochondrial | 984.54 | 33.43 | 72585 | 17 | 40076370 | 46117394 | 47329858 | 53380286 | 52451206 | 42231566 | 42119998 | 46766386 | 40213342 | 52871598 | 35841634 | 38840218 | 57873922 | 57148030 | 51783158 | 37875290 | 42728714 | 50647066 | 58727902 | 61411858 | 12 | 11 | 11 | 8 | 12 | 11 | 1.008906632 | 1.07709121 | 1.050283931 | 1.121264895 |
| Q9CQA3 | Sdhb | Succinate dehydrogenase [ubiquinone] iron-sulfur subunit, mitochondrial | 581.69 | 34.04 | 31814 | 9 | 22873316 | 22867406 | 25181834 | 22721846 | 35227602 | 19885726 | 20474962 | 29806240 | 29286202 | 20524770 | 6601999 | 13779550 | 16132749 | 11327063 | 17380944 | 25931516 | 13290488 | 20188190 | 20664492 | 22878700 | 0 | 7 | 0 | 10 | 0 | 10 | 0.506101426 | 0.543619325 | 0.798880927 | 0.858102917 |
| Q9CZB0 | Sdhc | Succinate dehydrogenase cytochrome b560 subunit, mitochondrial | 63.75 | 7.69 | 18382 | 1 | 1435003.62 | 1041456.12 | 786725.88 | 890318.38 | 872385.69 | 1541062.12 | 1727094.25 | 2 | 1247998.88 | 743896.25 | 747015.06 | 1074835.5 | 1610400.38 | 1191169.38 | 1417648.88 | 1096556.5 | 854332.62 | 441389 | 695988.94 | 687975.88 | 8 | 5 | 12 | 7 | 12 | 7 | 1.20199001 | 1.148480562 | 0.751358102 | 0.717909607 |
| Q5M8N4 | Sdr39u1 | Epimerase family protein SDR39U1 | 44.24 | 6.17 | 32995 | 1 | 85859.92 | 92982.28 | 154940.16 | 90359.4 | 105739.34 | 102072.14 | 52013.35 | 2 | 132811.81 | 202154.62 | 176586.92 | 117888.69 | 183798.55 | 279900.09 | 176999.59 | 2 | 71612.83 | 96631.11 | 101838.05 | 127366.42 | 1 | 10 | 5 | 9.5 | 5 | 10 | 1.764874875 | 1.912210089 | 0.750074705 | 0.812692412 |
| P42667 | Sec11a | Signal peptidase complex catalytic subunit SEC11A | 225.82 | 16.2 | 20599 | 4 | 1007853.81 | 2732271.25 | 2168367 | 292805.44 | 498653.12 | 1139665.88 | 1190433.12 | 1049093.5 | 316108.44 | 1153780.75 | 3085148.5 | 4047475 | 4111522.5 | 3519881 | 4213876 | 243213.81 | 2 | 979051.88 | 509392.97 | 928253.25 | 0 | 6 | 0 | 3 | 0 | 6 | 2.832543712 | 3.913710722 | 0.397005002 | 0.548539719 |
| Q5XFW8 | Sec13 | Protein SEC13 homolog | 112.3 | 6.83 | 35548 | 2 | 2740121.5 | 3413309.5 | 2534112.25 | 1913116.12 | 1903830.88 | 2584389.5 | 2232038.5 | 1961668.25 | 1642189 | 1892469.62 | 3588792.5 | 3437884 | 2584127 | 1374033.88 | 3140804 | 2115541 | 1020524.19 | 1638621.75 | 1484886 | 1516709.5 | 8 | 2 | 6 | 3 | 8 | 3 | 1.129645519 | 1.369725312 | 0.621879204 | 0.754045116 |
| O08547 | Sec22b | Vesicle-trafficking protein SEC22b | 630.95 | 43.72 | 24741 | 10 | 5436720.5 | 6338159 | 5188988.5 | 6332689 | 5591379 | 7792231.5 | 5212002.5 | 6442542.5 | 5928865 | 7721434 | 7649776.5 | 6259790 | 7505622.5 | 8343102.5 | 7364479 | 5532310.5 | 8272267.5 | 7918247 | 7921520 | 7444177 | 2 | 3 | 9 | 6 | 9 | 6 | 1.285061366 | 1.121632952 | 1.283875802 | 1.120598163 |
| Q9D662 | Sec23b | Protein transport protein Sec23B | 544.07 | 16.69 | 86437 | 9 | 7376193 | 7728683 | 9570496 | 9197736 | 8096919 | 7386000.5 | 6636415 | 8259694 | 8756874 | 8709012 | 7043156.5 | 6889831.5 | 7373043.5 | 6180845 | 6309096.5 | 7064165 | 8278637 | 8338973 | 7718718.5 | 8584532 | 0 | 10 | 3 | 12 | 3 | 12 | 0.805240678 | 0.850256034 | 0.952704307 | 1.00596332 |
| Q6NZC7 | Sec23ip | SEC23-interacting protein | 56.25 | 0.8 | 110780 | 1 | 1932839.12 | 2259049.75 | 1825957.5 | 2102788.5 | 1936469.88 | 2334753.25 | 1870426.88 | 1872502 | 1991176.75 | 2529631.25 | 1490542.38 | 1760109 | 1783496.5 | 1481181.25 | 1370724.38 | 2423845.25 | 1971338.88 | 2451424.25 | 2152265 | 2741245 | 0 | 3 | 0 | 7 | 0 | 7 | 0.78412761 | 0.744073298 | 1.167345739 | 1.107716121 |
| Q3UPL0 | Sec31a | Protein transport protein Sec31A | 657.72 | 12.11 | 133569 | 10 | 7319392.5 | 9112508 | 6225977 | 8906831 | 10244437 | 4715937.5 | 7109971 | 9624596 | 10674874 | 9383669 | 6926425.5 | 7322314 | 7388233.5 | 3339849 | 3604037 | 8491790 | 7718602.5 | 8801693 | 9591577 | 9099617 | 5 | 12 | 5 | 11 | 5 | 12 | 0.683603041 | 0.688545286 | 1.045304298 | 1.052861536 |
| P61620 | Sec61a1 | Protein transport protein Sec61 subunit alpha isoform 1 | 409.6 | 24.16 | 52265 | 7 | 8621367 | 6291027.5 | 16741497 | 8259011 | 13128453 | 9086017 | 15083565 | 24726082 | 15644648 | 13308394 | 34289962 | 34015670 | 46941578 | 40086746 | 49122062 | 18931488 | 17729576 | 16487887 | 17550012 | 21226606 | 0 | 1 | 0 | 5 | 0 | 5 | 3.85465296 | 2.626325195 | 1.733092379 | 1.18082334 |
| Q9CQS8 | Sec61b | Protein transport protein Sec61 subunit beta | 161.01 | 37.5 | 9958 | 3 | 14511030 | 13737012 | 13301174 | 17340382 | 12870361 | 14551708 | 14236868 | 16459350 | 13303834 | 15610830 | 10089208 | 9109832 | 9717178 | 9010318 | 11199416 | 17623090 | 20536134 | 17465326 | 16914198 | 19503906 | 0 | 1 | 0 | 0 | 0 | 1 | 0.684587236 | 0.662408797 | 1.282646413 | 1.241092767 |
| P60060 | Sec61g | Protein transport protein Sec61 subunit gamma | 68.72 | 17.65 | 7741 | 1 | 663210.62 | 722639.88 | 473569.62 | 2 | 2 | 2 | 2 | 2 | 2 | 504709.19 | 656061.25 | 548155.75 | 1447934.75 | 1062748.62 | 2 | 435058.94 | 831238.56 | 853020.81 | 583645.56 | 547806.44 | 8 | 7 | 3 | 1 | 8 | 7 | 1.997877907 | 7.360364267 | 1.748267259 | 6.440775893 |
| Q8BU14 | Sec62 | Translocation protein SEC62 | 101.25 | 4.77 | 45581 | 2 | 3073701.5 | 3409986.5 | 3042914.5 | 3635743 | 2150073.25 | 4024070 | 3083394.5 | 2898358.5 | 2390697 | 3324120 | 2150099.5 | 1909699.38 | 1834859.75 | 1987701.12 | 1842400.25 | 1724340.88 | 2841435 | 1794644.25 | 2377395.75 | 2391615.5 | 1 | 3 | 0 | 2 | 1 | 3 | 0.635089737 | 0.618598225 | 0.726823865 | 0.707950273 |
| Q8R2U0 | Seh1l | Nucleoporin SEH1 | 101.18 | 5.28 | 39775 | 2 | 960620.5 | 585121.44 | 994536.62 | 909010 | 706294.19 | 1016299.88 | 868918.38 | 954560.94 | 1199068.5 | 1192927.75 | 1351225.75 | 393264.25 | 1268357.25 | 1347125.88 | 1348571 | 670258.38 | 736002.94 | 1131216.25 | 1664827.5 | 1059512.5 | 5 | 7 | 5 | 11 | 5 | 11 | 1.373704838 | 1.091129423 | 1.266204498 | 1.005742242 |
| Q9Z2G6 | Sel1l | Protein sel-1 homolog 1 | 479.7 | 14.81 | 88340 | 7 | 1449648.25 | 2004069.38 | 1703608.5 | 1904361.62 | 1583826.75 | 1011187.62 | 1792948.75 | 2054909.75 | 2034347.12 | 2669072.5 | 2215766.5 | 3716867.75 | 3618209.75 | 2573970.5 | 3259437.75 | 2106538.75 | 1209293 | 1531699.88 | 1695045.5 | 922644.81 | 0 | 8 | 2 | 7 | 2 | 8 | 1.779449014 | 1.608816457 | 0.863479171 | 0.780679575 |
| Q923V8 | Selenof | Selenoprotein F | 144.31 | 17.28 | 17822 | 2 | 491594.06 | 532515.38 | 669065 | 845846.44 | 634432.25 | 902992.62 | 567793.69 | 851722.31 | 471901.03 | 875641.88 | 532241.44 | 445070.03 | 2 | 1589918.75 | 632996.5 | 392883.25 | 370041.88 | 519696.31 | 531933.31 | 371564.88 | 8 | 2 | 8 | 2 | 8 | 2 | 1.008437367 | 0.871984683 | 0.688877239 | 0.59566456 |
| Q9EP97 | Senp3 | Sentrin-specific protease 3 | 70.87 | 2.11 | 64403 | 1 | 103757.62 | 2 | 2 | 2 | 2 | 2 | 2 | 2 | 170578.7 | 89210.13 | 400270.19 | 473290.22 | 319120.59 | 207233.36 | 341449.38 | 2 | 2 | 2 | 2 | 2 | 0 | 10 | 0 | 7.5 | 0 | 10 | 16.78170226 | 6.702842162 | 9.637103E-05 | 3.849191E-05 |
| P42208 | Sept2 | Septin-2 | 689.31 | 39.06 | 41526 | 10 | 17090746 | 16211686 | 15121541 | 14958838 | 15363959 | 17069030 | 16037154 | 16985324 | 17666864 | 17704028 | 16559614 | 16653101 | 11707141 | 11848974 | 11644210 | 18107826 | 16725038 | 19182964 | 18820134 | 14488566 | 8 | 6 | 2 | 9 | 8 | 9 | 0.868772649 | 0.800504549 | 1.108928379 | 1.021788857 |
| O55131 | Sept7 | Septin-7 | 664.46 | 34.86 | 50550 | 12 | 6697754 | 6178711 | 5724120 | 4337728 | 5392747.5 | 5390248 | 5919212.5 | 4480593 | 5413140.5 | 5249857.5 | 6798666 | 7255116.5 | 7059636 | 6577122.5 | 8478074 | 4678953.5 | 6410497 | 5623928.5 | 4981192.5 | 6443041.5 | 1 | 12 | 0 | 9 | 1 | 12 | 1.276641762 | 1.367275719 | 0.993171893 | 1.063681179 |
| Q8CHH9 | Sept8 | Septin-8 | 357.8 | 17.95 | 49812 | 7 | 4316638 | 5652763 | 3866300.25 | 3465091.5 | 4572104 | 4650090 | 5290636 | 4318094 | 3914149.75 | 5718270.5 | 5260338 | 5807344 | 2573192.5 | 4538788 | 5525942 | 4222789 | 4080808 | 5781644 | 5528089 | 5686586.5 | 9 | 7 | 11 | 10 | 11 | 10 | 1.083788982 | 0.992229966 | 1.156678824 | 1.058962039 |
| Q80UG5 | Sept9 | Septin-9 | 355.09 | 13.89 | 65575 | 7 | 8021211 | 8537528 | 7617742 | 8445325 | 7922355 | 8852651 | 6907943 | 6791401 | 8124623.5 | 7802961.5 | 8040037 | 6756646 | 7180407 | 6769498 | 6040958 | 6010277.5 | 8481496 | 7696427 | 6404903 | 6989098 | 3 | 5 | 5 | 8 | 5 | 8 | 0.858016176 | 0.904052123 | 0.877615928 | 0.924703479 |
| Q6AXS5 | Serbp1 | Plasminogen activator inhibitor 1 RNA-binding protein | 647.37 | 27.52 | 44754 | 10 | 22412848 | 29288102 | 25523634 | 22451074 | 17854296 | 26850720 | 25072404 | 25975926 | 23870214 | 27002392 | 37386042 | 34262174 | 24075520 | 25351548 | 27860358 | 25124008 | 31900352 | 29701316 | 29506310 | 24560294 | 5 | 4 | 7 | 7 | 7 | 7 | 1.267214331 | 1.156587145 | 1.197926786 | 1.093348369 |
| Q9EPB5 | Serhl | Serine hydrolase-like protein | 51.93 | 2.89 | 35311 | 1 | 1257119.38 | 1631392 | 1314560.25 | 1567836.62 | 1645631.25 | 1576269 | 1072955.75 | 1382099.62 | 2 | 2 | 1180611.38 | 1154063.5 | 1070616.62 | 1572765 | 1447647.88 | 1229949 | 1658540 | 1503478.62 | 1640093.5 | 408116.75 | 5 | 11 | 9 | 6 | 9 | 11 | 0.866401963 | 1.593942192 | 0.868353478 | 1.597532446 |
| Q9QZI9 | Serinc3 | Serine incorporator 3 | 137.98 | 6.57 | 52623 | 3 | 1077126.75 | 617930.88 | 590843.75 | 478443.31 | 781988 | 746361.44 | 1212246.62 | 1011099.06 | 238469.72 | 518441 | 1611046.5 | 2132591.5 | 2428575 | 1548024.25 | 1568451.62 | 1559047.62 | 673105.81 | 761067.38 | 2 | 505818.62 | 0 | 12 | 0 | 11 | 0 | 12 | 2.619237867 | 2.492525198 | 0.986664742 | 0.938932185 |
| Q4G075 | Serpinb1a | Leukocyte elastase inhibitor A | 76.5 | 2.64 | 42727 | 1 | 387030.69 | 545643.56 | 259102.84 | 287193.97 | 314463.31 | 581456.38 | 571179.94 | 699026.12 | 795223.5 | 671208.62 | 2292143.25 | 1203094.38 | 2249718.25 | 1771185.5 | 1215161.38 | 601279 | 386916.78 | 1041289.94 | 1056485.75 | 1368197 | 0 | 2 | 0 | 8 | 0 | 8 | 4.868481895 | 2.631420715 | 2.483597139 | 1.342387442 |
| Q60854 | Serpinb6 | Serpin B6 | 1039.49 | 51.85 | 42599 | 15 | 106981778 | 98591338 | 99536754 | 106529506 | 105132810 | 90846442 | 95201674 | 87321202 | 89142626 | 93028650 | 46619314 | 48816950 | 43516294 | 40650730 | 46861850 | 101960290 | 85956554 | 91821602 | 99014770 | 96654858 | 0 | 3 | 0 | 7 | 0 | 7 | 0.438230122 | 0.497134923 | 0.919956776 | 1.043612974 |
| P32261 | Serpinc1 | Antithrombin-III | 592.06 | 16.56 | 52004 | 9 | 18538054 | 14376824 | 5103202.5 | 8564908 | 11437041 | 24077028 | 30826878 | 22985050 | 25600706 | 19469042 | 53209658 | 82062754 | 90721018 | 58913722 | 62685310 | 23230312 | 28633794 | 27377490 | 23029742 | 18607772 | 0 | 0 | 0 | 12 | 0 | 12 | 5.990904606 | 2.82690408 | 2.083403112 | 0.983087053 |
| Q64268 | Serpind1 | Heparin cofactor 2 | 91.62 | 2.92 | 54552 | 2 | 2 | 2 | 314710.41 | 255790.31 | 413022.44 | 138487.03 | 343577.31 | 574945.69 | 789393.62 | 2 | 1096599.12 | 1728344.62 | 2421166.5 | 1441650.12 | 2288313.5 | 2 | 2 | 515004.19 | 468467.06 | 332526.38 | 0 | 9 | 0 | 9 | 0 | 9 | 9.126411781 | 4.861376946 | 1.338042998 | 0.712737003 |
| Q63945 | Set | Protein SET | 542.6 | 41.18 | 33406 | 9 | 21827170 | 20215858 | 16366938 | 16894680 | 21247058 | 19225722 | 17991874 | 14619464 | 19207608 | 16592527 | 31194516 | 32905262 | 17365424 | 16777540 | 17213094 | 20411890 | 25696612 | 17119312 | 17112628 | 18273002 | 10 | 11 | 9 | 8 | 10 | 11 | 1.195792837 | 1.317429614 | 1.02135374 | 1.125246466 |
| Q91WC0 | Setd3 | Histone-lysine N-methyltransferase setd3 | 48.06 | 1.68 | 67176 | 1 | 156476.73 | 327107.09 | 258126.47 | 147648.72 | 166855.7 | 2 | 275623.84 | 229835.75 | 221492.92 | 248657.05 | 2 | 334227.22 | 135105.41 | 2 | 2 | 191948.33 | 2 | 2 | 2 | 75760.02 | 5 | 3 | 7.5 | 3.5 | 7.5 | 3.5 | 0.444359111 | 0.481071206 | 0.253465841 | 0.274406701 |
| Q8VHL1 | Setd7 | Histone-lysine N-methyltransferase SETD7 | 75.02 | 4.92 | 40506 | 1 | 378866.72 | 435084.53 | 413309.97 | 363087.31 | 431491.31 | 370060.44 | 394743.91 | 306899.38 | 431652.78 | 244623.55 | 2 | 2 | 303458.09 | 225321.92 | 2 | 457695.25 | 386273 | 560355.81 | 356516.47 | 262709.88 | 0 | 12 | 1 | 9 | 1 | 12 | 0.261537041 | 0.30251261 | 1.000846046 | 1.15765074 |
| Q64213 | Sf1 | Splicing factor 1 | 394.03 | 9.95 | 70403 | 6 | 1234353.5 | 2332551.5 | 3675655 | 4228454 | 2506023.5 | 3465056.75 | 3420105.75 | 2467733.5 | 2321710.25 | 1931790 | 5295259 | 4129318.25 | 2074429.25 | 2013093.5 | 1890214.25 | 4496012 | 4420448.5 | 3209468.25 | 4212367 | 2400121.25 | 12 | 6 | 12 | 5 | 12 | 6 | 1.101972736 | 1.131990717 | 1.340657274 | 1.377177076 |
| Q8K4Z5 | Sf3a1 | Splicing factor 3A subunit 1 | 926.52 | 29.08 | 88545 | 18 | 7592403 | 9370178 | 9686491 | 7098193.5 | 7261149 | 5910342.5 | 8540280 | 9411626 | 7771951 | 8306208 | 12519729 | 12565173 | 8225003.5 | 7883453.5 | 8158357.5 | 5844157.5 | 8150373 | 8033212.5 | 8466384 | 7394134.5 | 6 | 11 | 9 | 8 | 9 | 11 | 1.203453616 | 1.235633775 | 0.923914323 | 0.948619803 |
| Q62203 | Sf3a2 | Splicing factor 3A subunit 2 | 209.23 | 13.47 | 49911 | 4 | 807075.88 | 711547.75 | 810309.88 | 214294.73 | 509417.78 | 457040.78 | 2 | 484539.81 | 140169.11 | 513887.19 | 1010707.19 | 782341.56 | 221127.62 | 456445.75 | 811730.25 | 286550.97 | 237842.16 | 711342.62 | 2 | 824558.88 | 10 | 9 | 6 | 10.5 | 10 | 10.5 | 1.075248276 | 2.057077194 | 0.674921565 | 1.291204823 |
| Q9D554 | Sf3a3 | Splicing factor 3A subunit 3 | 350.51 | 16.77 | 58842 | 6 | 1763693.25 | 2939448.25 | 2171996.5 | 2307528.25 | 5740935 | 1444022 | 1920641.5 | 2525262.5 | 3673293.75 | 1510420.5 | 2430001 | 2409676.75 | 1897069.25 | 2372694.5 | 2114179 | 2950980.75 | 2577504.75 | 2209258.75 | 1881850.38 | 2457934.5 | 11 | 12 | 11 | 9 | 11 | 12 | 0.752071857 | 1.013543898 | 0.809290528 | 1.090655725 |
| Q99NB9 | Sf3b1 | Splicing factor 3B subunit 1 | 713.23 | 14.88 | 145816 | 16 | 8557603 | 9345948 | 6949399 | 7948405 | 7905751 | 9387026 | 9092406 | 10738769 | 7765459.5 | 6947940.5 | 4408444 | 9930924 | 7267625 | 5871514 | 7877961 | 9373455 | 8429539 | 7289597 | 7314254.5 | 6955003.5 | 7 | 11 | 7 | 8 | 7 | 11 | 0.868557642 | 0.804807182 | 0.966952772 | 0.895980299 |
| Q921M3 | Sf3b3 | Splicing factor 3B subunit 3 | 1843.07 | 30.07 | 135550 | 27 | 23377764 | 23822430 | 20626898 | 21264542 | 21852732 | 28609580 | 23651334 | 20244020 | 18568900 | 21799660 | 28950900 | 24272998 | 24309110 | 23157900 | 21350664 | 21219476 | 28254392 | 22178834 | 22665644 | 23326412 | 5 | 10 | 7 | 10 | 7 | 10 | 1.100024962 | 1.081224366 | 1.060394162 | 1.042270898 |
| Q6AYL5 | Sf3b4 | Splicing factor 3B subunit 4 | 283.83 | 16.27 | 44356 | 5 | 2360113 | 2732745.5 | 2192282.25 | 1966953.25 | 1656481.25 | 2523014.5 | 1639327.38 | 2074980.38 | 1646344.5 | 2681475.75 | 3035288 | 2661603.25 | 2098841.75 | 2335352 | 503519.28 | 1802466.12 | 1018179.62 | 1578152.5 | 1533225.62 | 2285858.5 | 11 | 4 | 10 | 5 | 11 | 5 | 0.974884807 | 1.006574617 | 0.753341493 | 0.777829769 |
| P59708 | Sf3b6 | Splicing factor 3B subunit 6 | 112.71 | 18.4 | 14585 | 2 | 1521540.12 | 603998.12 | 1151458.5 | 2 | 895604.5 | 2 | 2 | 616785.94 | 1048266.44 | 881235.56 | 592570.19 | 669559 | 642441.44 | 2 | 637394.69 | 492116.22 | 435484.91 | 634239.38 | 956035.31 | 1041110.81 | 7.5 | 10 | 12 | 10 | 12 | 10 | 0.609204176 | 0.998301601 | 0.852941539 | 1.397713504 |
| Q8VIJ6 | Sfpq | Splicing factor, proline- and glutamine-rich | 1424.8 | 31.33 | 75442 | 22 | 53683502 | 56954818 | 54665158 | 44815674 | 51501558 | 51961398 | 55199194 | 47283370 | 48711982 | 47624998 | 54963158 | 62130142 | 55136538 | 47369234 | 59211786 | 53026862 | 46517958 | 48644262 | 41690858 | 51653382 | 6 | 6 | 6 | 9 | 6 | 9 | 1.065706373 | 1.111770519 | 0.923219427 | 0.96312471 |
| Q8BP27 | Sfr1 | Swi5-dependent recombination DNA repair protein 1 homolog | 65.27 | 3.76 | 35183 | 1 | 1312765.88 | 1612694.38 | 1292757.12 | 1458227.5 | 990601.75 | 1130347.5 | 1376673.75 | 1234842.62 | 838600.44 | 1215851.25 | 615109.19 | 329179.59 | 585893.19 | 567439.88 | 623369.69 | 1363618 | 1296308.62 | 1203165.12 | 1536355.75 | 1257274.88 | 0 | 11 | 0 | 6 | 0 | 11 | 0.408125508 | 0.469434723 | 0.998451449 | 1.148440298 |
| Q99JR1 | Sfxn1 | Sideroflexin-1 | 483.52 | 29.5 | 35649 | 8 | 6229848.5 | 8289035.5 | 9225031 | 9054769 | 9778035 | 8935171 | 7884046 | 7921555.5 | 9897513 | 12318944 | 11817776 | 10529592 | 11586878 | 17203718 | 12501369 | 9078886 | 11294480 | 10587108 | 13280445 | 9904075 | 0 | 2 | 3 | 5 | 3 | 5 | 1.494697912 | 1.355261664 | 1.271704238 | 1.153070455 |
| Q91V61 | Sfxn3 | Sideroflexin-3 | 317.52 | 27.1 | 35406 | 5 | 5388601 | 4688643 | 4078988 | 5003397 | 5175066 | 6469363.5 | 2315676.75 | 2326551.25 | 1779398.38 | 5322748 | 5407045 | 2106785.25 | 4781817 | 3987501 | 4339212 | 2485966.5 | 5768948 | 5407893 | 3598336.5 | 5576126 | 8 | 10 | 11 | 7 | 11 | 10 | 0.847446835 | 1.132242068 | 0.93846543 | 1.253848614 |
| Q8R0X7 | Sgpl1 | Sphingosine-1-phosphate lyase 1 | 478.87 | 19.19 | 63677 | 9 | 8090855 | 9528189 | 8730407 | 8180365 | 9072290 | 8932219 | 8776986 | 8291352.5 | 9125721 | 9289983 | 9977245 | 9836623 | 13593261 | 11121256 | 13829608 | 7439228.5 | 6665845 | 9569956 | 12551461 | 9404609 | 0 | 11 | 0 | 10 | 0 | 11 | 1.338421429 | 1.313888 | 1.0465343 | 1.027351199 |
| Q9JJU8 | Sh3bgrl | SH3 domain-binding glutamic acid-rich-like protein | 374.06 | 50.88 | 12811 | 6 | 18866464 | 19760272 | 18795056 | 19020624 | 23148000 | 19019854 | 21931450 | 24001816 | 24158814 | 20272888 | 13675312 | 15154845 | 11057924 | 11294087 | 10625626 | 22101468 | 19115978 | 20325136 | 23793260 | 18102208 | 0 | 9 | 0 | 9 | 0 | 9 | 0.6206199 | 0.565049089 | 1.038634581 | 0.945634395 |
| Q8BG73 | Sh3bgrl2 | SH3 domain-binding glutamic acid-rich-like protein 2 | 127.64 | 32.71 | 12255 | 2 | 3736095.25 | 6846200 | 1790566.75 | 2358104.5 | 2982749 | 3429238 | 4808621 | 3049467.5 | 3921219.5 | 2768475.25 | 1626488.25 | 3922293.5 | 465400.66 | 1372206.62 | 1490734.5 | 4517214.5 | 3635836.75 | 3099161.25 | 2922479.75 | 3349616.5 | 4 | 10 | 4 | 12 | 4 | 12 | 0.501144073 | 0.493803918 | 0.989307339 | 0.974817157 |
| Q91VW3 | Sh3bgrl3 | SH3 domain-binding glutamic acid-rich-like protein 3 | 109.88 | 31.18 | 10477 | 2 | 13821302 | 14168911 | 12134226 | 12008428 | 13059073 | 13300386 | 13863918 | 12608497 | 13517594 | 13337844 | 11464357 | 11573538 | 9181436 | 9144313 | 7553110 | 13221516 | 17893010 | 14644429 | 15952626 | 15471372 | 0 | 2 | 0 | 4 | 0 | 4 | 0.750349721 | 0.734174499 | 1.18393398 | 1.158412021 |
| P55194 | Sh3bp1 | SH3 domain-binding protein 1 | 192.84 | 8.53 | 74172 | 5 | 3190609.5 | 3809920 | 4490802.5 | 4387910.5 | 3666784 | 3547409.5 | 4282484.5 | 3895506.25 | 3468490.25 | 1560622.88 | 844693.75 | 1724855.5 | 2606129 | 1226067.38 | 1137558.5 | 3966598.25 | 1698651.75 | 2320277 | 2257081.25 | 2907223.5 | 0 | 3 | 2 | 8 | 2 | 8 | 0.385720552 | 0.449986458 | 0.672762403 | 0.784853099 |
| O35964 | Sh3gl1 | Endophilin-A2 | 78.61 | 4.35 | 41492 | 2 | 1384779.62 | 3759137 | 119754.23 | 1238054.38 | 2959341.5 | 579323.5 | 1329700.88 | 1282584.5 | 1386485.5 | 3519271.5 | 366949.62 | 2866758 | 2111280.25 | 186663.2 | 687892.5 | 4177322.25 | 3269312.75 | 800273.06 | 2857573.5 | 2035265.88 | 9 | 9 | 9 | 7 | 9 | 9 | 0.657382909 | 0.76809467 | 1.388823038 | 1.62271875 |
| Q6AYE2 | Sh3glb1 | Endophilin-B1 | 241.1 | 9.32 | 40787 | 3 | 2665214 | 2020140.5 | 3000006.5 | 4654448.5 | 4677355.5 | 1177232.62 | 1990481.5 | 3083532.5 | 3286194 | 3761170.5 | 559197.56 | 680699.88 | 1143869.25 | 281448.28 | 475682.72 | 2409213.25 | 2296083 | 3231830 | 3674788.75 | 3883393.25 | 0 | 11 | 0 | 9 | 0 | 11 | 0.184572324 | 0.236182385 | 0.910569313 | 1.165182447 |
| A2AAY5 | Sh3pxd2b | SH3 and PX domain-containing protein 2B | 119.87 | 2.75 | 101517 | 3 | 4576874 | 4816674.5 | 5494536 | 5309791 | 5257893 | 5907214 | 4185331.75 | 4097920.5 | 5079915 | 5497254.5 | 3760530 | 3752843.25 | 3529200.75 | 2788497 | 3702362.25 | 4593471 | 5079869.5 | 4366471 | 4576754 | 4856949 | 0 | 5 | 0 | 10 | 0 | 10 | 0.688780354 | 0.707917115 | 0.922129477 | 0.947749504 |
| P50431 | Shmt1 | Serine hydroxymethyltransferase, cytosolic | 249.77 | 5.86 | 52601 | 3 | 446557.66 | 2 | 535281.5 | 415730.38 | 2 | 571669.06 | 404969.81 | 2 | 162112.84 | 2 | 698596.25 | 424896.19 | 261675.58 | 465953.41 | 2 | 410110.28 | 721313 | 2 | 473515.25 | 2 | 10 | 12 | 8 | 10 | 10 | 12 | 1.324526672 | 1.62556676 | 1.148377874 | 1.409382641 |
| Q9CZN7 | Shmt2 | Serine hydroxymethyltransferase, mitochondrial | 2163.04 | 62.5 | 55759 | 34 | 113253394 | 109288442 | 107143194 | 126275066 | 98402882 | 123445994 | 102799906 | 115512834 | 106383762 | 136037330 | 176867714 | 164704290 | 184667970 | 210115906 | 200868290 | 110115274 | 134129266 | 143371474 | 144863922 | 137707794 | 0 | 2 | 0 | 4 | 0 | 4 | 1.690632685 | 1.604341897 | 1.208933058 | 1.147228473 |
| Q8K2Q9 | Shtn1 | Shootin-1 | 469.61 | 12.84 | 71343 | 9 | 7977429.5 | 7276175 | 6536327 | 6379734 | 6639198.5 | 6335016 | 7400578 | 6604102 | 5431539.5 | 6573212.5 | 5379158 | 7878235.5 | 2830949.25 | 1546203.12 | 2755671 | 7818663 | 6229875 | 6055072 | 5863111.5 | 7413393 | 4 | 8 | 5 | 12 | 5 | 12 | 0.585776567 | 0.630408559 | 0.958954435 | 1.032019916 |
| P70665 | Siae | Sialate O-acetylesterase | 113.6 | 4.99 | 60775 | 2 | 1721404.12 | 1097780.12 | 324347.06 | 837222.62 | 1968767.5 | 761982.5 | 1443061.5 | 287876.59 | 1286508.88 | 1257775.12 | 2 | 420774.91 | 2 | 2 | 2 | 798645.38 | 119568.66 | 94193.4 | 811969.88 | 635668.25 | 1 | 3 | 1 | 5 | 1 | 5 | 0.070725506 | 0.083535005 | 0.413486295 | 0.488375155 |
| Q9EPK6 | Sil1 | Nucleotide exchange factor SIL1 | 244.25 | 9.25 | 52430 | 4 | 1497068.25 | 2517387 | 2459088 | 2005829.5 | 2166620 | 2116765.5 | 2371273.25 | 2413807 | 3401428.5 | 2133487.75 | 1206036.75 | 1640443.5 | 2941061.5 | 1920445.75 | 2319848.25 | 2438253.5 | 2324777.25 | 3068811.5 | 2907117.5 | 2734025 | 10 | 4 | 6 | 7 | 10 | 7 | 0.941935241 | 0.806305994 | 1.265545174 | 1.083319336 |
| Q8C8M1 | Sinhcaf | SIN3-HDAC complex-associated factor | 60.32 | 5.88 | 24829 | 1 | 111809.68 | 2 | 136999.59 | 116955.89 | 96719.53 | 125379.56 | 2 | 119714.19 | 2 | 2 | 2 | 87577.48 | 203492.38 | 2 | 2 | 56609.27 | 2 | 2 | 2 | 2 | 7.5 | 3 | 12.5 | 9 | 12.5 | 9 | 0.629371323 | 1.187581219 | 0.122419242 | 0.230996849 |
| P97797 | Sirpa | Tyrosine-protein phosphatase non-receptor type substrate 1 | 347.79 | 15.98 | 56425 | 6 | 14675151 | 9433290 | 17798348 | 19245050 | 12877282 | 12444094 | 12067628 | 14062358 | 12707352 | 14926931 | 11313716 | 8923925 | 9676290 | 11177919 | 11147156 | 14281373 | 14509298 | 13808406 | 11948080 | 14928646 | 4 | 10 | 0 | 9 | 4 | 10 | 0.705654819 | 0.789009177 | 0.93849288 | 1.049350865 |
| P97710 | Sirpa | Tyrosine-protein phosphatase non-receptor type substrate 1 | 115.09 | 4.52 | 55691 | 1 | 3004251 | 2840580.25 | 2833474.5 | 3310780.75 | 2 | 3535186.5 | 2738166.75 | 2370096.5 | 2288529.5 | 2875641.75 | 842845.62 | 1988843.12 | 2197786.5 | 2283478 | 2116495.25 | 842536.56 | 1066708.5 | 2599202 | 162172.5 | 2107699.5 | 5 | 5 | 0 | 2 | 5 | 5 | 0.786502534 | 0.682916231 | 0.565374012 | 0.490911437 |
| Q5RJQ4 | Sirt2 | NAD-dependent protein deacetylase sirtuin-2 | 109.25 | 7.14 | 39319 | 2 | 2 | 915988.75 | 842974.94 | 472963.06 | 1362017.62 | 1231519 | 2 | 1148902.25 | 1327891.38 | 1273240.88 | 2 | 2 | 389209.31 | 1223188.62 | 841574.69 | 2 | 2 | 852479.69 | 2 | 2 | 8 | 5 | 5 | 3 | 8 | 5 | 0.682808358 | 0.492612521 | 0.237201005 | 0.171128815 |
| Q8R216 | Sirt4 | NAD-dependent protein lipoamidase sirtuin-4, mitochondrial | 47.77 | 3.6 | 37525 | 1 | 247708.95 | 220979.75 | 236209.08 | 340827.22 | 222727.58 | 330338.91 | 256206.77 | 144016.11 | 238919.61 | 269433.56 | 2 | 2 | 267320 | 2 | 201603.42 | 257148.61 | 2 | 332192 | 2 | 2 | 4 | 8 | 4 | 8 | 4 | 8 | 0.369686205 | 0.378500087 | 0.464618559 | 0.475695773 |
| Q3UND0 | Skap2 | Src kinase-associated phosphoprotein 2 | 431.72 | 29.89 | 40712 | 8 | 13963265 | 11612911 | 16303496 | 17918418 | 16991520 | 14008840 | 15100614 | 16365465 | 18337558 | 16508020 | 9154706 | 10048664 | 9674024 | 10207499 | 10739722 | 15923850 | 14268601 | 16875846 | 16008804 | 15401702 | 0 | 11 | 0 | 11 | 0 | 11 | 0.648845788 | 0.620322544 | 1.021997677 | 0.977070685 |
| Q6PEC4 | Skp1 | S-phase kinase-associated protein 1 | 357.08 | 30.06 | 18672 | 5 | 7714227 | 8018556 | 9415634 | 9755121 | 8676100 | 7041177.5 | 8738194 | 8569774 | 9685045 | 8540013 | 6860478 | 6549053 | 7013263.5 | 20096202 | 5289640.5 | 9893174 | 10753308 | 8727770 | 17146248 | 16563612 | 5 | 2 | 5 | 2 | 5 | 2 | 1.051147717 | 1.075971674 | 1.44755934 | 1.481744973 |
| Q8CI08 | Slain2 | SLAIN motif-containing protein 2 | 100.94 | 5.34 | 62378 | 2 | 2 | 2 | 1009006.88 | 2 | 499313.09 | 1330878 | 730303.56 | 1520810 | 1553184 | 2221552.75 | 1000865.94 | 826454.94 | 830245.44 | 1963593.62 | 1879365.75 | 944722.75 | 349700.16 | 2068916 | 1893080.75 | 1887804 | 3 | 3 | 11 | 12 | 11 | 12 | 4.309761828 | 0.883616387 | 4.736524997 | 0.971114245 |
| P41251 | Slc11a1 | Natural resistance-associated macrophage protein 1 | 61.13 | 2.55 | 59741 | 1 | 425383.78 | 434529.16 | 2 | 2 | 326375.72 | 2 | 311587.44 | 537489.12 | 2 | 2 | 790142.94 | 933358.75 | 941690.69 | 583074.94 | 1195092.38 | 2 | 2 | 613744.06 | 942486.62 | 2 | 0 | 12 | 0 | 10.5 | 0 | 12 | 3.745584753 | 5.233130333 | 1.311848865 | 1.832844947 |
| Q63632 | Slc12a4 | Solute carrier family 12 member 4 | 253.29 | 4.52 | 120628 | 6 | 441647.84 | 386516.81 | 356887 | 110253.45 | 522900.12 | 272370.91 | 605334.69 | 490248.28 | 437769.94 | 2 | 2 | 436707.56 | 2 | 272530.34 | 467162.44 | 371132.47 | 2 | 2 | 2 | 2 | 8 | 2 | 8 | 4 | 8 | 4 | 0.647014059 | 0.651485584 | 0.204124631 | 0.20553534 |
| Q924N4 | Slc12a6 | Solute carrier family 12 member 6 | 236.24 | 3.91 | 127527 | 6 | 2 | 2 | 2 | 2 | 2 | 393485.03 | 2 | 482910.72 | 2 | 2 | 410261.47 | 563117.81 | 629829.31 | 941283.25 | 880737.69 | 2 | 2 | 449662.5 | 2 | 2 | 0 | 10 | 1 | 10 | 1 | 10 | 342522.953 | 3.908286959 | 44967.05 | 0.513087177 |
| Q5RK27 | Slc12a7 | Solute carrier family 12 member 7 | 51.97 | 0.65 | 119378 | 1 | 1018795.5 | 939524.94 | 888938.38 | 1099566.88 | 1232809.75 | 1333430.75 | 1476397 | 1279761.62 | 1288075.5 | 1262896.88 | 1727469.38 | 2004153.12 | 2578650 | 2331914.75 | 2294464.75 | 1059226.12 | 1546978.62 | 1349643.25 | 1560873.12 | 1750682.88 | 0 | 2 | 0 | 6 | 0 | 6 | 2.111471378 | 1.646946811 | 1.403072487 | 1.094395966 |
| Q99MR3 | Slc12a9 | Solute carrier family 12 member 9 | 248.7 | 5.58 | 96330 | 5 | 2100301.5 | 2051488.75 | 1955887.12 | 2216476.5 | 2240842.75 | 2099289.75 | 2045757 | 1975483.88 | 1888544.12 | 1794772.88 | 1338324.25 | 1089027.5 | 1410522.75 | 1406231.5 | 1553677.5 | 1691158.12 | 2369065.5 | 1590572.5 | 1687382 | 1961738.25 | 0 | 6 | 0 | 7 | 0 | 7 | 0.643425052 | 0.693379146 | 0.880257392 | 0.948598624 |
| Q8BPX9 | Slc15a3 | Solute carrier family 15 member 3 | 686.54 | 19.9 | 64051 | 10 | 16337541 | 15653552 | 7273231 | 13383588 | 9678633 | 21308606 | 18243898 | 13165218 | 11726833 | 15117490 | 45517138 | 34738442 | 49076998 | 50358462 | 52886698 | 14389028 | 15866988 | 15696881 | 18914514 | 20250774 | 0 | 4 | 0 | 9 | 0 | 9 | 3.731600043 | 2.923224736 | 1.365681172 | 1.069834102 |
| P53986 | Slc16a1 | Monocarboxylate transporter 1 | 308.93 | 11.56 | 53267 | 5 | 1753786.12 | 1571419.25 | 1593456.25 | 1967701 | 1440187.62 | 1833914.38 | 1861837.38 | 2252977 | 1714045.88 | 2368603 | 5249233 | 3753128.75 | 4730612.5 | 5150874 | 4936248.5 | 2235543.5 | 2474423.25 | 2531022.5 | 2921630.25 | 2711864.25 | 0 | 0 | 0 | 2 | 0 | 2 | 2.86074017 | 2.37455887 | 1.54619661 | 1.283421302 |
| Q3U9N9 | Slc16a10 | Monocarboxylate transporter 10 | 66.2 | 2.34 | 55320 | 1 | 2 | 2 | 2 | 530905.5 | 2 | 364630.09 | 2 | 2 | 2 | 2 | 487970.97 | 483232.72 | 602327.75 | 259279.33 | 676193.06 | 2 | 2 | 2 | 2 | 2 | 3 | 10 | 1 | 10 | 3 | 10 | 4.725824131 | 6.88080565 | 1.883546E-05 | 2.742445E-05 |
| P51912 | Slc1a5 | Neutral amino acid transporter B(0) | 922.38 | 30.02 | 58483 | 12 | 1092364 | 2690801 | 3025113.5 | 4033137.5 | 2239726.25 | 2677644.25 | 3116386.75 | 4434341.5 | 3646883.5 | 4249711 | 31546544 | 35856170 | 34753970 | 29187606 | 33871846 | 1293785.38 | 2617361.75 | 4687115 | 4614672 | 6186011.5 | 0 | 7 | 0 | 10 | 0 | 10 | 12.6301001 | 9.115389617 | 1.482970314 | 1.070288604 |
| Q61609 | Slc20a1 | Sodium-dependent phosphate transporter 1 | 61.15 | 1.47 | 74153 | 1 | 2 | 2 | 2 | 2 | 2 | 2 | 2 | 2 | 2 | 2 | 1769839.75 | 1187912.75 | 1417739.62 | 2 | 1068840.12 | 2 | 2 | 2 | 2 | 2 | 2.5 | 12.5 | 2.5 | 12.5 | 2.5 | 12.5 | 544433.424 | 544433.424 | 1 | 1 |
| Q9EPR4 | Slc23a2 | Solute carrier family 23 member 2 | 297.67 | 8.02 | 70049 | 5 | 1027715.25 | 861613.62 | 542322.19 | 396800.47 | 756733.06 | 864292.25 | 693639.06 | 470775.25 | 910536.88 | 528150.56 | 1156607 | 913100.75 | 1442371.25 | 333761.88 | 1359789.5 | 877451.56 | 445937.97 | 405265.06 | 327745.47 | 472447.31 | 6 | 7 | 5 | 5 | 6 | 7 | 1.451983921 | 1.501309162 | 0.705360437 | 0.729322185 |
| P32089 | Slc25a1 | Tricarboxylate transport protein, mitochondrial | 345.63 | 23.15 | 33835 | 7 | 6715794 | 8123958 | 7897652.5 | 7537323.5 | 8056010.5 | 6914926 | 6647434 | 8102028 | 7939634 | 7607597.5 | 9627372 | 9127762 | 13492733 | 12708465 | 6493620 | 5458284 | 5218107 | 9524316 | 3939660.5 | 8957593 | 5 | 10 | 5 | 10 | 5 | 10 | 1.34226352 | 1.382631358 | 0.863483507 | 0.889452299 |
| Q9QZD8 | Slc25a10 | Mitochondrial dicarboxylate carrier | 621.61 | 37.28 | 31715 | 9 | 10468875 | 12346649 | 14357291 | 19095500 | 13901232 | 12591567 | 8682461 | 14253406 | 7453462 | 13598113 | 6719080.5 | 6848351 | 11173330 | 10720918 | 8438949 | 11076610 | 11175018 | 13420392 | 12837986 | 13057450 | 2 | 8 | 5 | 12 | 5 | 12 | 0.625636482 | 0.77591724 | 0.877409911 | 1.088167804 |
| Q9CR62 | Slc25a11 | Mitochondrial 2-oxoglutarate/malate carrier protein | 824.16 | 51.27 | 34155 | 15 | 40990838 | 34279970 | 40765342 | 48517298 | 47246262 | 38406062 | 35319746 | 41036366 | 35057978 | 44828770 | 35184742 | 34025690 | 45032310 | 35586478 | 43782222 | 38180486 | 40120198 | 45971418 | 45834034 | 43574566 | 8 | 11 | 12 | 6 | 12 | 11 | 0.914125152 | 0.994669994 | 1.008880994 | 1.097774906 |
| Q8BH59 | Slc25a12 | Calcium-binding mitochondrial carrier protein Aralar1 | 1287.36 | 38.4 | 74570 | 22 | 10385789 | 10417374 | 12246940 | 10677751 | 9999886 | 12710475 | 12794154 | 11233565 | 12156012 | 13957124 | 8842633 | 11672392 | 11149695 | 9055088 | 9726832 | 9036491 | 8872389 | 10680648 | 10868957 | 11487091 | 8 | 12 | 1 | 1 | 8 | 12 | 0.938930988 | 0.802634407 | 0.948217364 | 0.81057276 |
| Q9QXX4 | Slc25a13 | Calcium-binding mitochondrial carrier protein Aralar2 | 1603.81 | 47.93 | 74467 | 28 | 28052970 | 29168192 | 42127186 | 41563970 | 41078670 | 27656632 | 30199164 | 40654346 | 40154678 | 42388354 | 24200506 | 30224540 | 42109838 | 34204522 | 41140254 | 25192810 | 21923060 | 37044090 | 33738458 | 32695784 | 11 | 6 | 12 | 6 | 12 | 6 | 0.944440502 | 0.949332487 | 0.827481644 | 0.83176781 |
| O70579 | Slc25a17 | Peroxisomal membrane protein PMP34 | 126.54 | 5.21 | 34413 | 2 | 2 | 2 | 2 | 2 | 2 | 2 | 2 | 2 | 2 | 2 | 550915.38 | 178131.73 | 258723.31 | 520870.75 | 2 | 2 | 2 | 2 | 2 | 2 | 2.5 | 12.5 | 2.5 | 12.5 | 2.5 | 12.5 | 150864.317 | 150864.317 | 1 | 1 |
| Q9D6M3 | Slc25a22 | Mitochondrial glutamate carrier 1 | 168.34 | 8.67 | 34670 | 3 | 849905.69 | 875144.38 | 1489792.5 | 2268682.5 | 1950351.62 | 1831790.25 | 1430428 | 1971744.75 | 1108165.5 | 2315186 | 1589950.12 | 1391970.5 | 2273507.5 | 2143743.75 | 2451138.5 | 2056274.25 | 875498.56 | 1665801.75 | 2195476.25 | 1328623.62 | 6 | 10 | 9 | 11 | 9 | 11 | 1.325057003 | 1.137802071 | 1.092522081 | 0.938128611 |
| Q8BMD8 | Slc25a24 | Calcium-binding mitochondrial carrier protein SCaMC-1 | 590.52 | 28 | 52902 | 10 | 10309415 | 11024610 | 10142066 | 13881845 | 9989448 | 12489211 | 11892545 | 13636157 | 10635635 | 15574492 | 10775625 | 10570809 | 14259062 | 11672733 | 16076398 | 10387177 | 9501452 | 10637625 | 10912346 | 13676156 | 5 | 12 | 11 | 6 | 11 | 12 | 1.144672474 | 0.986401375 | 0.995796947 | 0.858110507 |
| P16036 | Slc25a3 | Phosphate carrier protein, mitochondrial | 885.03 | 41.29 | 39445 | 14 | 43734926 | 48152374 | 62615658 | 63512710 | 65593422 | 51883262 | 52563958 | 66100750 | 63655398 | 71751218 | 61837946 | 67780338 | 95112138 | 98172258 | 95705666 | 51756286 | 48365274 | 76813874 | 82918018 | 79935186 | 3 | 6 | 4 | 10 | 4 | 10 | 1.476004687 | 1.368204188 | 1.198087967 | 1.11058521 |
| P48962 | Slc25a4 | ADP/ATP translocase 1 | 1787.46 | 74.5 | 32904 | 27 | 14947885 | 17230060 | 15403444 | 14904180 | 14671976 | 14905865 | 16623267 | 16690955 | 16055364 | 16295180 | 14901348 | 17058606 | 19568936 | 15064500 | 19548840 | 15660254 | 11122692 | 15030301 | 14073492 | 14909741 | 7 | 9 | 9 | 3 | 9 | 9 | 1.116445968 | 1.069151736 | 0.917557447 | 0.878688414 |
| Q8CFJ7 | Slc25a45 | Solute carrier family 25 member 45 | 111.53 | 8.68 | 31690 | 2 | 586012 | 299800.09 | 294023.12 | 554124.12 | 611583.31 | 801961.19 | 948889.12 | 574163.62 | 605812 | 579075 | 395038.19 | 693326.88 | 555764.88 | 130483.44 | 598071.06 | 796160 | 572747.19 | 765343.5 | 625795.06 | 712576.12 | 11 | 2 | 5 | 12 | 11 | 12 | 1.011571655 | 0.675997556 | 1.480519608 | 0.989378885 |
| P51881 | Slc25a5 | ADP/ATP translocase 2 | 2069.79 | 81.21 | 32931 | 29 | 136445954 | 119045322 | 98111826 | 111667826 | 107200954 | 129107666 | 130604322 | 118307490 | 112127714 | 122774722 | 169108754 | 164528994 | 193597986 | 185790258 | 176077282 | 132594658 | 111761234 | 108288658 | 82816778 | 101264122 | 0 | 10 | 0 | 5 | 0 | 10 | 1.553095099 | 1.450597953 | 0.937557751 | 0.875683244 |
| Q5HZI9 | Slc25a51 | Solute carrier family 25 member 51 | 60.11 | 6.38 | 33689 | 1 | 2 | 384858.66 | 2 | 220366.67 | 95621.55 | 2 | 2 | 24886.86 | 189449.86 | 189591.02 | 2 | 2 | 2 | 153327.34 | 165447.05 | 2 | 2 | 43131.16 | 363642.69 | 163686.89 | 9 | 11 | 9 | 12 | 9 | 12 | 0.45484767 | 0.789193714 | 0.813960225 | 1.412280055 |
| Q91VE0 | Slc27a4 | Long-chain fatty acid transport protein 4 | 156.29 | 5.44 | 72319 | 3 | 503544.16 | 2004745.88 | 1253422.25 | 1128619.88 | 1596266 | 715051.19 | 543567.12 | 2 | 1524229.75 | 426487.72 | 2291631 | 2498608.5 | 2030176.5 | 828116.62 | 2199066.25 | 1077444.5 | 576496.56 | 638931.5 | 361299.22 | 670078.56 | 4 | 4 | 1 | 11 | 4 | 11 | 1.518145353 | 3.068420822 | 0.51247977 | 1.035805692 |
| P11167 | Slc2a1 | Solute carrier family 2, facilitated glucose transporter member 1 | 473.83 | 13.82 | 53963 | 7 | 3092552.75 | 3433677 | 561326.44 | 936228.94 | 1531889.62 | 3129515.5 | 4705844 | 3448770.5 | 4108368.75 | 2028025.62 | 20963490 | 21958016 | 32201954 | 21074078 | 23149682 | 3006910.25 | 2648212.25 | 4745789.5 | 2745733 | 3565960 | 0 | 6 | 0 | 11 | 0 | 11 | 12.48966955 | 6.85095451 | 1.74897173 | 0.959362913 |
| P32037 | Slc2a3 | Solute carrier family 2, facilitated glucose transporter member 3 | 52.41 | 1.83 | 53479 | 1 | 2 | 2 | 136959.09 | 2 | 2 | 2 | 284874.81 | 102861.85 | 2 | 126622.92 | 1171268.62 | 1095802.5 | 1190946.12 | 1028640.81 | 681002.62 | 110935.09 | 2 | 100500.05 | 144059.31 | 161354.94 | 0 | 5 | 0 | 11 | 0 | 11 | 37.72921415 | 10.04670795 | 3.77354436 | 1.004836676 |
| Q60738 | Slc30a1 | Zinc transporter 1 | 177.97 | 8.15 | 54716 | 3 | 441014.28 | 646542 | 347457.31 | 623168.25 | 3829908 | 456879.41 | 664247.19 | 1147027.88 | 716529.88 | 1173772.88 | 1334396.88 | 1591366.75 | 1829477.5 | 1645201.62 | 1534198.88 | 528622.88 | 567665.44 | 780867.06 | 826258.69 | 603975.56 | 5 | 11 | 0 | 9 | 5 | 11 | 1.347574824 | 1.908073396 | 0.561708418 | 0.795340541 |
| Q99J27 | Slc33a1 | Acetyl-coenzyme A transporter 1 | 218.24 | 8.36 | 61075 | 4 | 775929.12 | 1430168.5 | 566479.12 | 113103.79 | 1423356.5 | 867867.38 | 783350.69 | 903027.75 | 649925.5 | 1117740.5 | 434493.19 | 1364551.25 | 2358453.75 | 1893495.38 | 2479712 | 773142.88 | 681820.69 | 650043.5 | 1131394.38 | 2 | 6 | 9 | 5 | 8 | 6 | 9 | 1.979724359 | 1.973826845 | 0.751073483 | 0.748836067 |
| Q61420 | Slc35a1 | CMP-sialic acid transporter | 49.74 | 3.57 | 36453 | 1 | 658202.5 | 701569.94 | 876091.62 | 800750.06 | 771266.94 | 819186.25 | 2 | 802064.31 | 2 | 1052502.12 | 516254.28 | 715846.25 | 950129.38 | 931961.62 | 991948.25 | 801811.31 | 840910.94 | 781562.12 | 867023.75 | 879859 | 8 | 5 | 9 | 9 | 9 | 9 | 1.07832669 | 1.535719316 | 1.095403731 | 1.560039906 |
| P97858 | Slc35b1 | Solute carrier family 35 member B1 | 68.19 | 3.11 | 35883 | 1 | 841177.56 | 990431.88 | 867423.44 | 1350912.88 | 1183026.62 | 1361369.62 | 1310151.88 | 1366369.38 | 2049040.25 | 1356156.62 | 1610329.12 | 1475943 | 2965527.5 | 1568622.5 | 1774108.5 | 1061722.5 | 735469.44 | 1787086.62 | 1232221.25 | 1744720.5 | 0 | 8 | 4 | 8 | 4 | 8 | 1.795257062 | 1.262181898 | 1.253822844 | 0.881518602 |
| Q91ZN5 | Slc35b2 | Adenosine 3'-phospho 5'-phosphosulfate transporter 1 | 65.68 | 2.32 | 47370 | 1 | 242179.84 | 516026.78 | 405062.41 | 2 | 2 | 2 | 2 | 195755.38 | 2 | 261278.17 | 1322608.88 | 1439141.25 | 1604640.75 | 1910172.5 | 1650316.62 | 264819.88 | 2 | 364685.72 | 366459.38 | 249601.11 | 0 | 12 | 0 | 4.5 | 0 | 12 | 6.814290193 | 17.34396947 | 1.070744406 | 2.725296071 |
| P0C6B1 | Slc35e1 | Solute carrier family 35 member E1 | 52.29 | 2.93 | 44384 | 1 | 246792.55 | 234380.05 | 189341.36 | 435592.25 | 187102.45 | 250620.34 | 260734.75 | 171044.92 | 360484.22 | 297138.16 | 232430.11 | 209972.89 | 383070.16 | 381190 | 2 | 337656.09 | 402699.09 | 389979.06 | 2 | 237627.81 | 12 | 10 | 12 | 10 | 12 | 10 | 0.933078472 | 0.900481342 | 1.057806132 | 1.020851637 |

| | | | | | | | | | | | | | | | | | | | | | | | | | | | | | | | | | | | | |
|---|---|---|---|---|---|---|---|---|---|---|---|---|---|---|---|---|---|---|---|---|---|---|---|---|---|---|---|---|---|---|---|---|---|---|---|---|
| Q9WU81 | Slc37a2 | Glucose-6-phosphate exchanger SLC37A2 | 290.42 | 11.18 | 55073 | 5 | 8284388 | 8088368 | 5438423.5 | 9965466 | 9921874 | 13788850 | 13205134 | 11421332 | 11904541 | 13245604 | 16187891 | 15760414 | 17926422 | 18540176 | 16922654 | 15151308 | 14198181 | 13826449 | 16089322 | 17467638 | 0 | 0 | 0 | 0 | 0 | 0 | 2.046536856 | 1.342514561 | 1.840182791 | 1.207147668 |
| Q8K2P7 | Slc38a1 | Sodium-coupled neutral amino acid transporter 1 | 103.41 | 7.42 | 53795 | 2 | 2 | 2 | 2 | 2 | 2 | 251625.44 | 164979.72 | 2 | 172143.39 | 2 | 1321254 | 1296056.12 | 1966036.62 | 1627361.5 | 1767368.38 | 127891.23 | 245056.58 | 302188.34 | 205133.95 | 120290.04 | 0 | 0 | 0 | 8 | 0 | 8 | 797807.662 | 13.55081455 | 100056.014 | 1.699457845 |
| Q8CFE6 | Slc38a2 | Sodium-coupled neutral amino acid transporter 2 | 369.19 | 15.67 | 55503 | 6 | 2612774.5 | 2595230.5 | 172236.84 | 718003.38 | 1404909 | 2045983.62 | 4463505 | 321326.78 | 1863998.88 | 647917.88 | 3853018.75 | 10905270 | 4345602.5 | 2327168.25 | 9106990 | 3345153.5 | 332711.03 | 2 | 2 | 850471 | 2 | 8 | 3 | 7 | 3 | 8 | 4.070028231 | 3.268642296 | 0.603524784 | 0.484691143 |
| Q6P6S2 | Slc39a11 | Zinc transporter ZIP11 | 92.52 | 6.57 | 34868 | 1 | 365221.38 | 344759.03 | 265149.03 | 304560.44 | 267653.91 | 348006.41 | 407202.19 | 507917.09 | 438557.38 | 455154.31 | 288443.69 | 369328.09 | 647489.81 | 414370.56 | 489306.38 | 280032.31 | 519215.81 | 431734.38 | 499361.62 | 421972.44 | 3 | 3 | 12 | 12 | 12 | 12 | 1.427568033 | 1.024156272 | 1.39097502 | 0.997903959 |
| Q78IQ7 | Slc39a4 | Zinc transporter ZIP4 | 155.25 | 4.09 | 71063 | 2 | 2 | 2 | 2 | 2 | 212244.03 | 490096.81 | 2 | 2 | 2 | 637962.69 | 1122110 | 834824.88 | 803855.31 | 439034.25 | 899065.19 | 520224.38 | 2 | 621530.69 | 675542.12 | 557467.5 | 0 | 3 | 2 | 6.5 | 2 | 6.5 | 19.31142722 | 3.633556411 | 11.18842863 | 2.105167377 |
| Q31125 | Slc39a7 | Zinc transporter SLC39A7 | 140.34 | 8.19 | 50657 | 2 | 1324677.25 | 1669090 | 1386560 | 1472587.62 | 1343896.62 | 1535726.5 | 2235623.5 | 1678386.5 | 1192941.88 | 1713843.12 | 1849698.5 | 1982342.75 | 1884691.62 | 1529736.12 | 2167158 | 2487957.25 | 1903563.25 | 1502018.38 | 2135818.25 | 1880676.12 | 1 | 1 | 8 | 7 | 8 | 7 | 1.308027451 | 1.126500661 | 1.377003311 | 1.185904117 |
| P10852 | Slc3a2 | Amino acid transporter heavy chain SLC3A2 | 1538.43 | 43.35 | 58337 | 23 | 19888242 | 22148812 | 20073252 | 21779348 | 18065094 | 23246186 | 27277712 | 22806582 | 22682570 | 25025252 | 74960418 | 88608218 | 90566370 | 74906730 | 89635922 | 27318094 | 29236768 | 27323670 | 25414426 | 28803020 | 0 | 0 | 0 | 1 | 0 | 1 | 4.10650476 | 3.459050987 | 1.35448305 | 1.140927919 |
| B4F795 | Slc44a2 | Choline transporter-like protein 2 | 125.32 | 5.25 | 79897 | 3 | 11193577 | 15308189 | 15545989 | 13261794 | 15009054 | 8955410 | 12573645 | 16853618 | 15880833 | 12128422 | 6206593 | 9371565 | 9315797 | 6793429 | 9750167 | 11089294 | 8715246 | 11471693 | 12463449 | 11802536 | 0 | 3 | 3 | 5 | 3 | 5 | 0.58928291 | 0.624135377 | 0.789865208 | 0.836580887 |
| Q8BTY2 | Slc4a7 | Sodium bicarbonate cotransporter 3 | 326.62 | 5.03 | 116514 | 5 | 2 | 2 | 2 | 2 | 2 | 140841.45 | 421306.94 | 196736 | 369459.84 | 299633.56 | 655779.62 | 1772453.38 | 1444893.12 | 237210.3 | 1943857 | 406224.03 | 2 | 2 | 2 | 314551.47 | 0 | 7.5 | 3 | 7 | 3 | 7.5 | 605419.342 | 4.239697187 | 72078.15 | 0.5047568 |
| Q9CXK4 | Slc50a1 | Sugar transporter SWEET1 | 61.71 | 9.05 | 24649 | 1 | 2 | 2 | 2 | 1186288 | 1521149 | 2 | 2 | 1489781.12 | 1826221.25 | 1549834.38 | 2 | 249207.94 | 1735358 | 2281148.25 | 2323128 | 330953.91 | 2 | 1689443.25 | 1438615.25 | 1949933.75 | 5.5 | 6.5 | 8 | 11 | 8 | 11 | 2.433604028 | 1.354101897 | 1.997806846 | 1.111616355 |
| Q09143 | Slc7a1 | High affinity cationic amino acid transporter 1 | 252.8 | 11.74 | 67092 | 4 | 2 | 2 | 2 | 2 | 132812.67 | 2 | 2 | 2 | 2 | 130933.38 | 4413334 | 4058870.75 | 5414258.5 | 3760409.75 | 4264950 | 2 | 294749.12 | 2 | 94392.73 | 662648.31 | 0 | 7 | 0 | 7 | 0 | 7 | 164.9729895 | 167.3407062 | 7.918904189 | 8.03255747 |
| Q9WTR6 | Slc7a11 | Cystine/glutamate transporter | 182.95 | 9.96 | 55456 | 4 | 2 | 2 | 2 | 2 | 2 | 2 | 2 | 2 | 2 | 2 | 4133776.5 | 4614765.5 | 2624050.25 | 2364724 | 2970044.5 | 2 | 2 | 2 | 407776.38 | 2 | 0 | 10 | 0 | 10 | 0 | 10 | 1670736.075 | 1670736.075 | 40778.438 | 40778.438 |
| Q9Z127 | Slc7a5 | Large neutral amino acids transporter small subunit 1 | 325.93 | 12.7 | 55872 | 5 | 2 | 2 | 2 | 2 | 2 | 2 | 2 | 2 | 2 | 2 | 256247.91 | 779812.25 | 1133775.88 | 986283.56 | 1245036 | 2 | 2 | 2 | 2 | 2 | 0 | 12.5 | 0 | 12.5 | 0 | 12.5 | 440115.56 | 440115.56 | 1 | 1 |
| P70441 | Slc9a3r1 | Na(+)/H(+) exchange regulatory cofactor NHE-RF1 | 41.91 | 3.94 | 38600 | 1 | 1085570.5 | 2 | 635252.31 | 2 | 350144.47 | 334706.38 | 2 | 2 | 222892.86 | 2 | 438319.12 | 567632.62 | 2 | 2 | 313676.66 | 1260194.38 | 2 | 668077.81 | 2 | 2 | 10 | 12 | 8 | 10.5 | 10 | 12 | 0.637204587 | 2.366606885 | 0.93109847 | 3.458142162 |
| B1ARD6 | Slfn9 | Schlafen family member 9 | 151.73 | 3.85 | 103992 | 3 | 2271834.5 | 2426148.75 | 2726213.25 | 1914708.12 | 2006276 | 1143569.38 | 1881881.62 | 1823455.75 | 1888983.5 | 1557205.25 | 1691341.5 | 1848053.25 | 1163455.25 | 878350 | 1502683.38 | 1810381.62 | 642126.88 | 1114413.25 | 942001.19 | 1114537.5 | 0 | 0 | 7 | 2 | 7 | 2 | 0.624395822 | 0.853984548 | 0.495669538 | 0.677925943 |
| O08815 | Slk | STE20-like serine/threonine-protein kinase | 183.61 | 3.57 | 137888 | 4 | 3353757.75 | 3206636.75 | 1326854.75 | 1616287.5 | 1412683.12 | 2589408 | 3205904.5 | 2516103 | 3610585.25 | 3090205.75 | 2553396.5 | 1684455.25 | 2516874.5 | 1903709 | 392765.06 | 2090857.25 | 2142359 | 1844386.25 | 2004460.25 | 3062743.5 | 12 | 10 | 2 | 2 | 12 | 10 | 0.829151521 | 0.602922716 | 1.020940067 | 0.742382957 |
| Q3URD3 | Slmap | Sarcolemmal membrane-associated protein | 231.64 | 5.44 | 96933 | 4 | 730023.25 | 769954.62 | 1193905.25 | 522448 | 807375.88 | 349430.88 | 697521.88 | 456420.38 | 1033504.38 | 565168.81 | 370504.72 | 1058225.75 | 476702.03 | 607242.06 | 1308872.38 | 378837.56 | 389223.75 | 394877.16 | 288113.22 | 641139.62 | 10 | 1 | 9 | 6 | 10 | 6 | 0.949757758 | 1.231943863 | 0.519966118 | 0.67445521 |
| Q8CH25 | Sltm | SAFB-like transcription modulator | 100.94 | 2.42 | 116919 | 2 | 583122.69 | 2 | 627551 | 473011.62 | 2 | 1080283.75 | 674671.62 | 690659.06 | 973873.94 | 2 | 982828.88 | 1234435.88 | 670646.5 | 1790835 | 2 | 2 | 545903.69 | 655530 | 816369.38 | 2 | 4 | 10 | 9.5 | 6 | 9.5 | 10 | 2.778866761 | 1.368258937 | 1.198443833 | 0.590089999 |
| O54835 | Smad9 | Mothers against decapentaplegic homolog 9 | 48.62 | 3 | 48992 | 1 | 662835.94 | 627516.25 | 583921.12 | 676483.75 | 479214.62 | 728819 | 2 | 398323.84 | 496492.78 | 449646.19 | 265135.94 | 255265.61 | 2 | 260381.84 | 366772.97 | 465305.72 | 652066.81 | 531065.88 | 269777.5 | 400759.56 | 0 | 4 | 4.5 | 11 | 4.5 | 11 | 0.378735672 | 0.55349796 | 0.765345592 | 1.118503631 |
| Q9R0P4 | Smap | Small acidic protein | 90.55 | 8.84 | 20046 | 1 | 562305.44 | 739130.94 | 608453.06 | 456924.94 | 556191.38 | 515419.59 | 692143.81 | 635906.25 | 611140.62 | 723324.94 | 2 | 551844.94 | 2 | 2 | 359122.09 | 602423.44 | 2 | 222984.88 | 535403.25 | 491198.34 | 1 | 5 | 1 | 2 | 1 | 5 | 0.311656256 | 0.286655633 | 0.633598447 | 0.582772079 |
| Q3TKT4 | Smarca4 | Transcription activator BRG1 | 97.15 | 1.49 | 181427 | 2 | 2 | 114915.91 | 541838 | 251983 | 555687.81 | 616035.62 | 195631.06 | 519485.94 | 582447.75 | 2 | 422905.09 | 624087.12 | 408537 | 553175.5 | 361722.66 | 58803.27 | 405638.12 | 2 | 2 | 2 | 7 | 5.5 | 11 | 4.5 | 11 | 5.5 | 1.618672575 | 1.238725143 | 0.317153043 | 0.242708411 |
| Q91ZW3 | Smarca5 | SWI/SNF-related matrix-associated actin-dependent regulator of chromatin subfamily A member 5 | 461.51 | 7.99 | 121627 | 9 | 6787082 | 8142618.5 | 8106559.5 | 8262912 | 9132291 | 7800210 | 7277738 | 7491777.5 | 7536482.5 | 6979846 | 7097229 | 5473063 | 7352491.5 | 6731863 | 5459363 | 6025054.5 | 4882014.5 | 5225097.5 | 7143520 | 5203866 | 2 | 1 | 3 | 1 | 3 | 1 | 0.794282648 | 0.865932232 | 0.704390848 | 0.767931592 |
| B4F769 | Smarcal1 | SWI/SNF-related matrix-associated actin-dependent regulator of chromatin subfamily A-like protein 1 | 75.11 | 1.76 | 101212 | 2 | 689316.5 | 797524.12 | 1305054.25 | 1613860.25 | 1488575.88 | 746774.25 | 480024.22 | 1161558.38 | 1149046 | 1626379.25 | 127988.93 | 579675.12 | 1032949.38 | 565510.81 | 802130.88 | 537377.75 | 605887.19 | 1017867.88 | 1062089.5 | 1435899.75 | 4 | 7 | 6 | 10 | 6 | 10 | 0.527329585 | 0.601933827 | 0.7904412 | 0.902269302 |
| Q9Z0H3 | Smarcb1 | SWI/SNF-related matrix-associated actin-dependent regulator of chromatin subfamily B member 1 | 144.33 | 7.79 | 44141 | 2 | 1488132.62 | 1715032.5 | 1559418.25 | 1895814.25 | 1319306.62 | 1893164 | 1357167.12 | 1608657.12 | 1618796 | 1758259.5 | 1336243.88 | 1298207.62 | 1274361.38 | 1698039.38 | 883076.19 | 1532301.62 | 1119267.5 | 1501320.75 | 1361394.38 | 1366406 | 4 | 6 | 3 | 4 | 4 | 6 | 0.813508279 | 0.787991013 | 0.862490015 | 0.835436341 |
| P97496 | Smarcc1 | SWI/SNF complex subunit SMARCC1 | 492.98 | 9.69 | 122890 | 9 | 2348759.5 | 1942556 | 1786859.5 | 2068306.62 | 1707181.25 | 2488814.5 | 1459893 | 1736827.38 | 1379013.5 | 1607387.25 | 2316447.25 | 1982225.75 | 1723529.62 | 3271086.75 | 1642581.88 | 2532588.75 | 3211519.25 | 2129521 | 2400148 | 1680010.62 | 12 | 6 | 6 | 4 | 12 | 6 | 1.10982803 | 1.261064625 | 1.213131378 | 1.378445151 |
| Q6PDG5 | Smarcc2 | SWI/SNF complex subunit SMARCC2 | 456.53 | 7.34 | 132604 | 8 | 2 | 748183.62 | 2 | 344740.34 | 1197671.62 | 386388.28 | 528039.88 | 2 | 950975.56 | 805363.12 | 2 | 980429.62 | 2 | 2 | 2 | 534054.12 | 2 | 2 | 2 | 1530208.75 | 8 | 11 | 7 | 9.5 | 8 | 11 | 0.42802663 | 0.367099393 | 0.901191499 | 0.772911844 |
| Q61466 | Smarcd1 | SWI/SNF-related matrix-associated actin-dependent regulator of chromatin subfamily D member 1 | 108.84 | 5.83 | 58245 | 2 | 315970.09 | 171934.92 | 433715.28 | 622457.88 | 515849.06 | 2 | 218719.53 | 2 | 175322.06 | 2 | 974517.88 | 2 | 102547.34 | 227920.38 | 157549.3 | 68004 | 2 | 98750.6 | 99552.29 | 356660.41 | 6 | 2 | 7.5 | 9.5 | 7.5 | 9.5 | 0.70999445 | 3.71157428 | 0.302422965 | 1.580949398 |
| O54941 | Smarce1 | SWI/SNF-related matrix-associated actin-dependent regulator of chromatin subfamily E member 1 | 295.93 | 15.82 | 46638 | 5 | 2085180.75 | 2293400.75 | 2577778.5 | 2412403.75 | 2306473 | 2575538 | 2261587 | 2542401.5 | 1876429.12 | 2045170.75 | 782059.31 | 351291.19 | 868636.56 | 424600.12 | 1028748 | 2224819 | 2025830.88 | 1507150.88 | 1624470 | 2032158.38 | 0 | 1 | 0 | 4 | 0 | 4 | 0.295954185 | 0.305751397 | 0.806358735 | 0.833052285 |
| Q9CU62 | Smc1a | Structural maintenance of chromosomes protein 1A | 547.84 | 11.03 | 143235 | 13 | 4418352.5 | 4640420.5 | 5209664.5 | 3706750.5 | 4042251 | 2695981 | 4153632.5 | 3908496.75 | 4280454.5 | 3915259.5 | 4888279.5 | 3429005 | 2839242 | 3731776.25 | 2850777.75 | 2947660 | 3185510.5 | 2823271 | 3084082.75 | 1853840.88 | 5 | 0 | 9 | 4 | 9 | 4 | 0.80568319 | 0.935910361 | 0.63106182 | 0.733063942 |
| Q8CG48 | Smc2 | Structural maintenance of chromosomes protein 2 | 1124.25 | 21.07 | 134239 | 21 | 11892427 | 10943507 | 14211970 | 9454951 | 7219781 | 8773906 | 9710824 | 9499144 | 6891039 | 9713547 | 8488712 | 11760622 | 8487167 | 2745544.25 | 7819888 | 10881706 | 5986109.5 | 7376451.5 | 5845497 | 6037436 | 6 | 3 | 8 | 6 | 8 | 6 | 0.731571199 | 0.881437333 | 0.67247631 | 0.81023655 |
| P97690 | Smc3 | Structural maintenance of chromosomes protein 3 | 500.25 | 10.16 | 138448 | 9 | 2329267.25 | 2809433.5 | 3606026.25 | 3010853 | 2307765.25 | 2776988 | 2880509 | 1307065.5 | 1956221.75 | 2633183 | 3205330.5 | 1545135.62 | 470959.38 | 1284611.5 | 806805.94 | 1324254.5 | 2098115 | 2559514 | 2870973.25 | 678296.25 | 4 | 5 | 6 | 9 | 6 | 9 | 0.519993132 | 0.632929173 | 0.677730144 | 0.824924703 |
| Q8CG47 | Smc4 | Structural maintenance of chromosomes protein 4 | 1006.57 | 16.64 | 146895 | 20 | 15776996 | 14905833 | 19238018 | 18377084 | 16628891 | 16274200 | 14682889 | 14707581 | 15214558 | 17012034 | 11344594 | 13815537 | 11965476 | 12212104 | 11445322 | 15223131 | 12435067 | 15887319 | 11517246 | 11711581 | 0 | 3 | 0 | 6 | 0 | 6 | 0.715710674 | 0.780357532 | 0.786257421 | 0.857276443 |
| Q6P5D8 | Smchd1 | Structural maintenance of chromosomes flexible hinge domain-containing protein 1 | 414.65 | 4.48 | 225648 | 8 | 67104846 | 92187898 | 127087618 | 134103490 | 111719626 | 96436498 | 84626298 | 125653634 | 118114410 | 124203978 | 52074698 | 60931918 | 83537474 | 102096458 | 79721882 | 83575730 | 72080954 | 137908642 | 113950130 | 136546002 | 4 | 10 | 2 | 12 | 4 | 12 | 0.710935658 | 0.689141048 | 1.022280914 | 0.990941631 |
| Q8BKX6 | Smg1 | Serine/threonine-protein kinase SMG1 | 44.87 | 0.46 | 409769 | 1 | 2 | 2 | 2 | 221419.84 | 2 | 292494.88 | 2 | 2 | 2 | 279945.47 | 309397.59 | 2 | 274468.56 | 329826.19 | 223281.39 | 2 | 354525.81 | 333771.78 | 244623.42 | 166618.38 | 3 | 4 | 7.5 | 7.5 | 7.5 | 7.5 | 5.134746064 | 1.986169935 | 4.965687196 | 1.920776314 |
| Q6ZPR5 | Smpd4 | Sphingomyelin phosphodiesterase 4 | 41.1 | 0.97 | 93276 | 1 | 1221498.12 | 478604.72 | 747962.62 | 810437.62 | 571676.19 | 943050.69 | 1136309.25 | 281380.72 | 890128.25 | 772620.19 | 488557.12 | 415460.94 | 791539 | 536642.69 | 469120.06 | 1170846.12 | 707491.69 | 510354.97 | 445136.66 | 497002.19 | 5 | 8 | 6 | 9 | 6 | 9 | 0.705272422 | 0.671387381 | 0.869628128 | 0.827846565 |
| P58242 | Smpdl3b | Acid sphingomyelinase-like phosphodiesterase 3b | 386.91 | 20.83 | 51600 | 7 | 1463865 | 2 | 3295985 | 2519254 | 2141971.25 | 1802970.5 | 2734977.75 | 2186380.5 | 1849952.5 | 1599829 | 2971756.75 | 2043719.38 | 4238402 | 2881039.75 | 4197585 | 906992.38 | 450373.09 | 1865889.5 | 1888849.75 | 398684.31 | 5 | 7 | 2 | 6 | 5 | 7 | 1.733613094 | 1.605300363 | 0.584942558 | 0.541648252 |
| P97355 | Sms | Spermine synthase | 243.69 | 17.21 | 41313 | 5 | 3176047.5 | 3187345.25 | 5210159 | 4005633.75 | 5406645 | 2594035 | 3447213.75 | 3724127.5 | 4999467.5 | 4134271.75 | 2909311 | 2121658 | 1419095.38 | 1816747.5 | 1916777 | 2957715 | 3772278.5 | 3778734.5 | 3555714 | 3498675 | 0 | 8 | 1 | 11 | 1 | 11 | 0.485260227 | 0.538839444 | 0.836903595 | 0.92930894 |
| Q3UKJ7 | Smu1 | WD40 repeat-containing protein SMU1 | 295.68 | 16.37 | 57544 | 6 | 1000514.5 | 239832.06 | 696038.19 | 2726092.25 | 2594501.25 | 1393835.5 | 3694634 | 1160656.38 | 2 | 415417.5 | 3349215 | 1005544 | 2195736.75 | 3357715.5 | 3189960.5 | 1875002.25 | 2683724.25 | 2908130 | 3170775 | 2893182.25 | 4 | 3 | 7 | 5 | 7 | 5 | 1.804907125 | 1.965351123 | 1.864524501 | 2.03026808 |
| O09044 | Snap23 | Synaptosomal-associated protein 23 | 129.35 | 18.57 | 23261 | 2 | 2 | 2 | 1019384.88 | 1003348.38 | 693380.44 | 649713.25 | 362969.5 | 809870.38 | 845520.31 | 1154038.25 | 1153278.12 | 1353400.75 | 1113679.62 | 1220323.5 | 1015030.56 | 556527.69 | 644453.12 | 1026501.81 | 2 | 944368.38 | 1 | 12 | 3 | 10 | 3 | 12 | 2.155912665 | 1.532062123 | 1.167789231 | 0.82986926 |
| Q78PY7 | Snd1 | Staphylococcal nuclease domain-containing protein 1 | 2517.89 | 53.52 | 102088 | 39 | 71533346 | 70860674 | 62891774 | 66915826 | 53659766 | 79586450 | 65520950 | 65120950 | 62073150 | 71505618 | 93469730 | 82128786 | 72985490 | 65013222 | 69768034 | 70199178 | 74506306 | 74614106 | 72117050 | 66315898 | 5 | 5 | 7 | 7 | 7 | 7 | 1.176467291 | 1.115059119 | 1.097867232 | 1.040561755 |
| F1LNJ2 | Snrnp200 | U5 small nuclear ribonucleoprotein 200 kDa helicase | 1536.78 | 17.39 | 244875 | 28 | 13779742 | 12999439 | 10647068 | 16712211 | 12713440 | 14560439 | 12067499 | 10890421 | 13727565 | 12203743 | 11290642 | 10673719 | 11349829 | 8996344 | 11196346 | 13506712 | 18247368 | 14653594 | 14627803 | 12118727 | 4 | 8 | 3 | 5 | 4 | 8 | 0.800379346 | 0.843296467 | 1.094272624 | 1.152948588 |
| Q6PE01 | Snrnp40 | U5 small nuclear ribonucleoprotein 40 kDa protein | 177.57 | 12.29 | 39276 | 4 | 2 | 2 | 1009125.12 | 881010.31 | 756917.94 | 889579.06 | 2 | 827932.5 | 814988.81 | 1232936.25 | 2019779.25 | 1498711 | 1485365.5 | 1250833.38 | 1949984.88 | 781509.06 | 840304.12 | 1036728.06 | 759379 | 753730.31 | 0 | 9 | 0 | 10 | 0 | 10 | 3.099545217 | 2.178942439 | 1.57595774 | 1.107879047 |
| Q62376 | Snrnp70 | U1 small nuclear ribonucleoprotein 70 kDa | 523.92 | 23.21 | 51992 | 10 | 15061200 | 23722930 | 22027248 | 23325986 | 17034648 | 16335981 | 20248828 | 24318458 | 14511913 | 23108676 | 19284014 | 17584504 | 17837614 | 15476033 | 12664371 | 9595069 | 7143686 | 16311627 | 13795266 | 9226812 | 7 | 1 | 7 | 1 | 7 | 1 | 0.818868127 | 0.840877929 | 0.55422897 | 0.569125715 |
| Q62189 | Snrpa | U1 small nuclear ribonucleoprotein A | 314.43 | 23.34 | 31835 | 6 | 7552845 | 4747318 | 6512862.5 | 7738000.5 | 13026633 | 7387676 | 4474387.5 | 6225780 | 5283202.5 | 9486997 | 5267264.5 | 4817631.5 | 5364472 | 5439635.5 | 5949348 | 6095791 | 9885601 | 8341667 | 7525287 | 7868508 | 5 | 10 | 8 | 6 | 8 | 10 | 0.678118721 | 0.816797017 | 1.003517009 | 1.208740703 |
| P57784 | Snrpa1 | U2 small nuclear ribonucleoprotein A' | 840.84 | 50.59 | 28357 | 13 | 17520504 | 14251923 | 16313620 | 14983261 | 11892303 | 16682445 | 13602146 | 12600880 | 11334971 | 14528315 | 9955409 | 8868412 | 9746193 | 8234028.5 | 10349696 | 12782103 | 18016530 | 14226346 | 14730334 | 16221990 | 0 | 12 | 0 | 7 | 0 | 12 | 0.629038489 | 0.685884961 | 1.013549495 | 1.105144388 |
| P17136 | Snrpb | Small nuclear ribonucleoprotein-associated protein B | 338.03 | 19.48 | 23656 | 6 | 15090585 | 13530329 | 15971527 | 17942868 | 13379040 | 20241172 | 13761482 | 14220442 | 15501166 | 14815506 | 22406536 | 23853284 | 38591414 | 38106618 | 34259586 | 13813062 | 18725580 | 15067281 | 15945593 | 17658434 | 0 | 9 | 0 | 9 | 0 | 9 | 2.070984472 | 2.001755824 | 1.069757576 | 1.033997834 |
| Q9CQI7 | Snrpb2 | U2 small nuclear ribonucleoprotein B'' | 187.31 | 20 | 25323 | 4 | 1578493.12 | 1433000.75 | 221146.67 | 2187524.25 | 548119.81 | 1998013.12 | 2 | 2 | 1432012 | 92148.82 | 1147144.38 | 2 | 2 | 2 | 226666.83 | 1886180.12 | 2661857.25 | 1945311.62 | 2 | 253080.7 | 3 | 11 | 9 | 8 | 9 | 11 | 0.230186277 | 0.390047645 | 1.130380359 | 1.915414782 |
| D3ZCL3 | Snrpc | U1 small nuclear ribonucleoprotein C | 115.07 | 13.21 | 17364 | 2 | 4499091.5 | 5410393 | 5943541.5 | 4372082.5 | 6755793 | 2838375.5 | 5505882.5 | 7584885.5 | 6568210 | 4802548.5 | 853303.06 | 2686158 | 1953502.88 | 1461114.38 | 2645405 | 6224008 | 1153085.62 | 3162350.25 | 2222208 | 5683057.5 | 0 | 7 | 0 | 7 | 0 | 7 | 0.355788087 | 0.351630688 | 0.683620945 | 0.675632805 |
| P62315 | Snrpd1 | Small nuclear ribonucleoprotein Sm D1 | 100.48 | 20.17 | 13282 | 2 | 2445829.75 | 2596576 | 1694342.62 | 2189160 | 1595066 | 3239067 | 1891299.25 | 2017864.62 | 1935478.75 | 3187325.75 | 2 | 6021884.5 | 7633901.5 | 5850211 | 6009020 | 1905429.5 | 3485992.5 | 3161345.5 | 2952780 | 2797427.25 | 5 | 3 | 5 | 10 | 5 | 10 | 2.425157414 | 2.079288196 | 1.359472445 | 1.165588259 |
| P62317 | Snrpd2 | Small nuclear ribonucleoprotein Sm D2 | 302.68 | 44.07 | 13527 | 6 | 912534.25 | 1225675.5 | 360644.03 | 994331.88 | 175024.23 | 515330.38 | 1074006.25 | 1019221.25 | 705900.62 | 920810.25 | 8044850.5 | 8167752 | 6049217.5 | 6107753 | 7684966 | 1173560.62 | 2060604.5 | 1610716.5 | 904615 | 1347343.25 | 0 | 4 | 0 | 3 | 0 | 4 | 9.828919304 | 8.512928253 | 1.934687513 | 1.675652784 |
| P62320 | Snrpd3 | Small nuclear ribonucleoprotein Sm D3 | 156.21 | 30.95 | 13916 | 3 | 17761068 | 17989994 | 19162858 | 24995466 | 15405266 | 22061778 | 19434404 | 19632798 | 18839780 | 23083578 | 36306738 | 22050618 | 30800628 | 34589718 | 27684276 | 16976084 | 23509180 | 23309940 | 26262592 | 24172646 | 1 | 7 | 2 | 5 | 2 | 7 | 1.588758652 | 1.46946669 | 1.198456267 | 1.108470164 |
| P62305 | Snrpe | Small nuclear ribonucleoprotein E | 117.95 | 25 | 10804 | 2 | 23474350 | 15399955 | 18612938 | 22316754 | 20984680 | 21704308 | 17777164 | 15130419 | 20895604 | 20702192 | 13849982 | 12760876 | 18704840 | 15329359 | 14893430 | 12113276 | 28132700 | 21060276 | 7065839 | 20080882 | 2 | 10 | 3 | 11 | 3 | 11 | 0.749473941 | 0.785144296 | 0.877608236 | 0.919376996 |
| P62307 | Snrpf | Small nuclear ribonucleoprotein F | 168.12 | 40.7 | 9725 | 3 | 517004.69 | 1012267.56 | 411603.72 | 296207 | 2 | 328566.91 | 556382.06 | 441216.5 | 2 | 629587.12 | 2530098.25 | 1238572.5 | 1211369.5 | 858502.12 | 1797688.5 | 345010.34 | 423670.25 | 254081.73 | 2 | 284998.62 | 1 | 7.5 | 0 | 6.5 | 1 | 7.5 | 3.413473772 | 3.90449339 | 0.58458349 | 0.668674356 |
| P62309 | Snrpg | Small nuclear ribonucleoprotein G | 150 | 17.11 | 8496 | 3 | 2 | 2 | 2 | 2 | 2 | 2 | 216023.05 | 263144.88 | 2 | 323453.34 | 1526589.38 | 563779.25 | 621266 | 118485.49 | 2 | 2 | 2 | 2 | 2 | 2 | 2.5 | 12.5 | 7 | 5 | 7 | 12.5 | 283012.212 | 3.526081505 | 1 | 1.245911E-05 |
| Q61235 | Sntb2 | Beta-2-syntrophin | 80.74 | 1.73 | 56382 | 1 | 751266.94 | 721832.38 | 354319.94 | 383212 | 548151.06 | 813307.19 | 778506.69 | 622657.25 | 824675.81 | 701036.06 | 619271 | 579983.38 | 505674.69 | 406886.22 | 480691.78 | 612852.5 | 682464.56 | 441933.59 | 761215.75 | 381485.81 | 12 | 11 | 0 | 3 | 12 | 11 | 0.939728753 | 0.693149792 | 1.043921512 | 0.770003021 |
| P55770 | Snu13 | NHP2-like protein 1 | 272.97 | 43.75 | 14174 | 6 | 14393259 | 17299510 | 11749541 | 15508244 | 12719277 | 17584850 | 16245432 | 12160483 | 10669095 | 13455071 | 15632916 | 13823916 | 12451756 | 12276650 | 9447075 | 15026598 | 13850274 | 13039300 | 14844858 | 13300856 | 8 | 12 | 10 | 12 | 10 | 12 | 0.887853538 | 0.907542974 | 0.977564549 | 0.999243456 |
| Q9CSN1 | Snw1 | SNW domain-containing protein 1 | 373.7 | 15.86 | 61475 | 7 | 1938379 | 4205845 | 3649853.75 | 2390391 | 2259855.25 | 5318064 | 3939412.75 | 2662758.25 | 3576042.25 | 4563269 | 5296478 | 4536913.5 | 3077093 | 2785943.75 | 3241025.75 | 3975121 | 2610273.25 | 4267644.5 | 3395543 | 3126066.5 | 6 | 7 | 10 | 8 | 10 | 8 | 1.31106544 | 0.944061933 | 1.202870294 | 0.866153603 |
| Q99N27 | Snx1 | Sorting nexin-1 | 554.12 | 19.73 | 59044 | 9 | 11846404 | 13135452 | 9969574 | 12690623 | 9599338 | 11676498 | 11070362 | 10588197 | 9759390 | 12295280 | 12118136 | 12007211 | 8446994 | 7887957.5 | 9095801 | 11832306 | 13433174 | 11205600 | 10619724 | 12003817 | 6 | 11 | 8 | 7 | 8 | 11 | 0.865738911 | 0.894680335 | 1.032375698 | 1.066887746 |
| O70493 | Snx12 | Sorting nexin-12 | 187.6 | 26.06 | 19116 | 4 | 3277677.75 | 3786535.75 | 4023411.75 | 4478184.5 | 5031220 | 3719509 | 3473516.5 | 3945523.5 | 3859011.75 | 4230772.5 | 2562999.25 | 2764646.25 | 3221425 | 2879050.75 | 3251033.5 | 3400517.5 | 3644481.25 | 4985979 | 3776848.5 | 4150284 | 0 | 10 | 0 | 12 | 0 | 12 | 0.712683087 | 0.763412749 | 0.968980018 | 1.037953212 |
| Q8BVL3 | Snx17 | Sorting nexin-17 | 49.35 | 3.62 | 52797 | 1 | 2 | 2 | 2 | 2 | 2 | 2 | 2 | 2 | 2 | 541783.94 | 1184046.5 | 1496173.75 | 1648145.88 | 1562972.62 | 1601976.12 | 2 | 2 | 2 | 2 | 818645 | 0 | 10 | 0 | 12 | 0 | 12 | 749331.487 | 13.83061341 | 81865.3 | 1.511009928 |
| Q91ZR2 | Snx18 | Sorting nexin-18 | 251.79 | 8.63 | 67904 | 5 | 5196432 | 5758152 | 5921530 | 6030658 | 4855884 | 5723148 | 4911123.5 | 6615733 | 6218401 | 6418817 | 6453263 | 6162818 | 8991630 | 8825816 | 8424361 | 3959154 | 4457405 | 6355241 | 6186657 | 6458674.5 | 0 | 10 | 4 | 9 | 4 | 10 | 1.399645913 | 1.300150524 | 0.987554343 | 0.917352942 |
| Q9CWK8 | Snx2 | Sorting nexin-2 | 705.45 | 27.55 | 58471 | 13 | 28008800 | 28245520 | 30118582 | 28565694 | 27457274 | 27159664 | 22949344 | 25627796 | 22638730 | 27853322 | 18316154 | 18631186 | 13212924 | 13219069 | 15585130 | 25983278 | 26836168 | 25588860 | 21557936 | 23151830 | 0 | 0 | 0 | 10 | 0 | 10 | 0.554541807 | 0.625565861 | 0.864618279 | 0.975355999 |
| Q3UHD6 | Snx27 | Sorting nexin-27 | 75.48 | 3.71 | 60989 | 2 | 1955912.88 | 1196651 | 2140806.75 | 2348554 | 2666683.75 | 2225524.75 | 1546214.88 | 2003280.5 | 1531538.38 | 2089889.38 | 1702430.12 | 1243629.75 | 1391420.38 | 2220989.75 | 976486.5 | 1478602.12 | 2288197.5 | 2397319.5 | 1595844.75 | 2825645 | 6 | 11 | 6 | 8 | 6 | 11 | 0.730938282 | 0.80189414 | 1.026870794 | 1.12655431 |
| Q5U211 | Snx3 | Sorting nexin-3 | 385.9 | 43.83 | 18762 | 8 | 8640204 | 9870248 | 12874294 | 13758889 | 13111206 | 7320338 | 8975867 | 10010031 | 10096814 | 12877094 | 8322998 | 7134324.5 | 8233186 | 11147242 | 10715566 | 9425280 | 11402415 | 14404332 | 11161117 | 12566633 | 4 | 12 | 10 | 6 | 10 | 12 | 0.781966197 | 0.924374663 | 1.0121009 | 1.19642055 |
| Q91YJ2 | Snx4 | Sorting nexin-4 | 65.94 | 2 | 51778 | 1 | 929865.44 | 947126.81 | 666955.56 | 1160018.25 | 866250.75 | 888115.69 | 615093.62 | 1112215.12 | 1041578.5 | 919043 | 876416.69 | 887829.81 | 1344899 | 960181.12 | 830631 | 653338.25 | 798353.75 | 708583.56 | 865996.25 | 1124756 | 11 | 7 | 11 | 9 | 11 | 9 | 1.072149927 | 1.070784187 | 0.908278093 | 0.907121098 |
| Q9D8U8 | Snx5 | Sorting nexin-5 | 1077.28 | 44.55 | 46797 | 19 | 52047294 | 48658690 | 57072854 | 47560386 | 65534822 | 41640470 | 48681466 | 48379626 | 57577694 | 44404850 | 22220066 | 22047642 | 18501490 | 18091088 | 17174900 | 49635094 | 38683350 | 38202426 | 38331730 | 40232758 | 0 | 2 | 0 | 4 | 0 | 4 | 0.361921666 | 0.407318903 | 0.757124431 | 0.852093482 |
| Q6P8X1 | Snx6 | Sorting nexin-6 | 248.09 | 10.34 | 46649 | 4 | 3510285 | 3409826 | 3952335.5 | 1829702 | 4776025.5 | 1141655.5 | 3120143 | 4353509.5 | 4146828 | 1828254 | 582066.25 | 1983521.25 | 1178477.38 | 499392.5 | 1855999.12 | 4917313 | 178726.98 | 1268898.75 | 1114658.5 | 2326046.25 | 2 | 6 | 5 | 8 | 5 | 8 | 0.348975614 | 0.418046159 | 0.561022177 | 0.672061763 |
| Q91VH2 | Snx9 | Sorting nexin-9 | 47.26 | 2.18 | 66546 | 1 | 1840820.88 | 839676.94 | 2 | 1329550.88 | 1521785.75 | 1186620 | 1641743.25 | 1466348.75 | 1653452.25 | 1533025.62 | 762958.62 | 1051057 | 1029008.44 | 2 | 1263276.25 | 1285676.38 | 2 | 1793998.75 | 1160883.38 | 653549.81 | 7.5 | 9.5 | 1 | 6 | 7.5 | 9.5 | 0.742303636 | 0.548883584 | 0.884717103 | 0.654188759 |
| Q61263 | Soat1 | Sterol O-acyltransferase 1 | 317.58 | 13.15 | 63799 | 6 | 3936999.75 | 3715354.25 | 4343658.5 | 4242494 | 7031317 | 2396802 | 3604695 | 4959330.5 | 7797527 | 4608763 | 4220646 | 6401208 | 8143720 | 7151065.5 | 7613762 | 3400892 | 2331613.75 | 4357750.5 | 4181356.25 | 3987286.25 | 4 | 8 | 6 | 7 | 6 | 8 | 1.440939228 | 1.434939568 | 0.784659959 | 0.781392859 |
| P08228 | Sod1 | Superoxide dismutase [Cu-Zn] | 303.98 | 41.56 | 15943 | 5 | 4445996.5 | 3917995.75 | 12267304 | 13643726 | 9986887 | 5378816.5 | 3109375 | 9691028 | 4807349 | 8332123 | 4255234 | 5658031 | 9143916 | 14160305 | 10941188 | 16195545 | 18213628 | 21294936 | 16866326 | 21270154 | 12 | 0 | 7 | 0 | 12 | 0 | 0.997667628 | 1.409978255 | 2.120120677 | 2.996312569 |
| P09671 | Sod2 | Superoxide dismutase [Mn], mitochondrial | 436 | 40.99 | 24603 | 7 | 39840390 | 41133922 | 26598114 | 32391068 | 35334958 | 40511462 | 44791502 | 42865002 | 40751490 | 38829334 | 49554490 | 73128018 | 79063234 | 60683918 | 90468634 | 47739830 | 30503558 | 45923254 | 50417366 | 56798022 | 0 | 4 | 0 | 5 | 0 | 5 | 2.01312841 | 1.698677975 | 1.319931964 | 1.113758737 |
| Q9QX47 | Son | Protein SON | 397.64 | 5.56 | 265651 | 8 | 604928.81 | 2462437.5 | 2464862 | 1439938.5 | 1864746.62 | 2525749.25 | 1705175.5 | 1021683.12 | 1201321.62 | 78385.77 | 1816082 | 1845499.75 | 1208354.75 | 1728702.75 | 2656369 | 419051.97 | 222893.17 | 1165735.75 | 863545.12 | 676510.5 | 12 | 3 | 5 | 6 | 12 | 6 | 1.047312314 | 1.416803672 | 0.378835499 | 0.512488509 |
| O35892 | Sp100 | Nuclear autoantigen Sp-100 | 295.51 | 12.24 | 54727 | 6 | 1447781.12 | 2332736.25 | 927597 | 2583610.5 | 1333819.25 | 2409581.25 | 1732866.5 | 2025773.75 | 1575940.25 | 2522889.5 | 780577.12 | 871127.38 | 1048141.56 | 1163209.25 | 765167.62 | 1129312.5 | 1682078.38 | 1722726 | 2039552.25 | 2305335.5 | 2 | 12 | 0 | 8 | 2 | 12 | 0.536571707 | 0.450784049 | 1.029384872 | 0.864805718 |
| Q8BVK9 | Sp110 | Sp110 nuclear body protein | 515.7 | 25.62 | 50140 | 10 | 8079860 | 9576880 | 11339911 | 12294665 | 11197758 | 10633997 | 10523113 | 11723016 | 11555364 | 10963564 | 5905158.5 | 7225355.5 | 10802161 | 7776167 | 9369299 | 7931567.5 | 6555060 | 10599959 | 8725991 | 11156973 | 3 | 5 | 2 | 4 | 3 | 5 | 0.782603652 | 0.741495351 | 0.856741167 | 0.811738599 |
| Q7TNE3 | Spag7 | Sperm-associated antigen 7 | 87 | 15.86 | 25921 | 2 | 1305485.75 | 2 | 377539.84 | 1775567.75 | 1289422.88 | 1341820.38 | 605546.69 | 1317980.12 | 998753.19 | 704408.75 | 455112.78 | 925473.75 | 394852.12 | 2 | 485827.38 | 1204031 | 1285739.5 | 284587.28 | 740191.62 | 351415.03 | 8.5 | 8 | 2 | 8 | 8.5 | 8 | 0.476255129 | 0.455120031 | 0.814226958 | 0.778093454 |
| Q58A65 | Spag9 | C-Jun-amino-terminal kinase-interacting protein 4 | 293.53 | 7.27 | 146219 | 7 | 4042364.5 | 3117856 | 5380525 | 5774084 | 5362477.5 | 3367363.75 | 4741961 | 4428129 | 5617291 | 5317781 | 1397446.75 | 2674842.25 | 2263180 | 3344001 | 2385621 | 3546482 | 3403521.25 | 4645272 | 4881703.5 | 6166646 | 1 | 11 | 0 | 12 | 1 | 12 | 0.509563482 | 0.514009064 | 0.956342913 | 0.964686331 |
| Q9CYN2 | Spcs2 | Signal peptidase complex subunit 2 | 414.12 | 42.04 | 24978 | 7 | 1722744.88 | 2605666.25 | 2344027.75 | 4973374 | 1782055.62 | 2501666.5 | 3095051.5 | 4277617 | 4776263 | 4640303.5 | 9324331 | 6856048 | 9855954 | 9661505 | 12678596 | 1873690.88 | 1027797.12 | 4065473.5 | 2068306.62 | 3801496 | 0 | 12 | 0 | 4 | 0 | 12 | 3.60268899 | 2.507733192 | 0.955979284 | 0.665431013 |
| Q6ZWQ7 | Spcs3 | Signal peptidase complex subunit 3 | 257.02 | 23.89 | 20313 | 5 | 3235333 | 3445539.25 | 5148136.5 | 4595276 | 4170507.25 | 3416268.25 | 4225007 | 5648856 | 3542826 | 6034589 | 10854318 | 8906214 | 12945522 | 12575963 | 12378706 | 3635967.5 | 3007613.5 | 6410599 | 5245986 | 1979988.62 | 0 | 12 | 0 | 10 | 0 | 12 | 2.799772049 | 2.52150897 | 0.984722478 | 0.886853115 |
| Q62407 | Speg | Striated muscle-specific serine/threonine-protein kinase | 60.48 | 0.34 | 354343 | 1 | 2 | 2 | 320797.25 | 2 | 904784.5 | 601805.81 | 2 | 272997.97 | 403250.75 | 2 | 522652.03 | 959602.25 | 545598.62 | 726107.12 | 819560.94 | 556606 | 644869.88 | 2 | 508775.28 | 2 | 4 | 10 | 2 | 10 | 4 | 10 | 2.915760997 | 2.796054231 | 1.395457127 | 1.338166539 |
| Q5XIC4 | Spg21 | Maspardin | 42.1 | 7.28 | 29576 | 1 | 412565.12 | 402515.66 | 391503.66 | 520339.09 | 438782.56 | 469151.94 | 436683.12 | 379057.72 | 2 | 367824.75 | 492082.69 | 207853.08 | 368737.22 | 175753 | 342675.53 | 494537.47 | 332254.34 | 523294.12 | 122164.26 | 544435.31 | 4 | 11 | 10 | 8 | 10 | 11 | 0.73283329 | 0.960296948 | 0.93119076 | 1.220222466 |
| Q9JIA7 | Sphk2 | Sphingosine kinase 2 | 269.84 | 8.91 | 65618 | 4 | 2695946.5 | 3268200.25 | 2825148.5 | 3535600.5 | 3259539.5 | 2693454.25 | 2734366 | 2320909.75 | 3804473 | 3294433.25 | 4322568.5 | 5356724 | 6931207 | 4950845 | 4531275 | 2618167.25 | 2838938 | 3611277.75 | 3451589.25 | 4419155.5 | 0 | 9 | 0 | 8 | 0 | 9 | 1.674274305 | 1.757358482 | 1.086925992 | 1.1408636 |
| P17433 | Spi1 | Transcription factor PU.1 | 131 | 8.09 | 31349 | 2 | 1267179.25 | 1272033.5 | 1803587.38 | 1334020.12 | 1835273.62 | 1011703.81 | 1577422.5 | 1584240.12 | 1698268.75 | 763863.5 | 705330.38 | 837873.06 | 1122656.25 | 496517.72 | 775307.12 | 854688.19 | 447602.41 | 1127073.25 | 1090755.75 | 998143 | 0 | 0 | 4 | 6 | 4 | 6 | 0.524179356 | 0.593427069 | 0.601465141 | 0.680922839 |
| P15702 | Spn | Leukosialin | 249.72 | 14.43 | 40038 | 4 | 8833089 | 11937924 | 7804647.5 | 8503242 | 8507665 | 13681885 | 11808580 | 10691202 | 9921858 | 10451851 | 13127055 | 12057570 | 10561999 | 9281626 | 12783704 | 7583025.5 | 9361346 | 8072104.5 | 9212802 | 9896493 | 2 | 12 | 11 | 0 | 11 | 12 | 1.268179579 | 1.022218542 | 0.96795555 | 0.780222397 |
| Q2YDU8 | Spns1 | Protein spinster homolog 1 | 53.89 | 4.17 | 56796 | 1 | 1544778.75 | 101528.74 | 2640876.5 | 2 | 103638.51 | 84203.62 | 2025633.62 | 119043.83 | 140391.97 | 78895.88 | 350135.81 | 293219.56 | 392647.53 | 367167.06 | 281999.97 | 3884064 | 166047.33 | 154375.58 | 2 | 75623.34 | 10 | 12.5 | 5 | 12 | 10 | 12.5 | 0.383793506 | 0.688338912 | 0.974785544 | 1.748291229 |
| Q3UHX9 | Spout1 | Putative methyltransferase C9orf114 homolog | 40.69 | 3.12 | 42958 | 1 | 233345.39 | 212540.66 | 2 | 346753.5 | 278303.34 | 507501.94 | 256100.83 | 2 | 2 | 302659.34 | 318333.72 | 234455.09 | 259908.97 | 359361.62 | 152172.8 | 416299.06 | 490741.47 | 318469 | 389621 | 389901.97 | 9 | 1 | 10 | 5 | 10 | 5 | 1.236508258 | 1.241934061 | 1.872208849 | 1.880424109 |
| P08721 | Spp1 | Osteopontin | 37.96 | 2.84 | 34963 | 1 | 1246595.25 | 1519479.38 | 506590.41 | 2 | 2 | 1710957.38 | 2458041 | 1291590.12 | 1703393.88 | 972375.75 | 505818.69 | 4224851 | 541153.69 | 2332170.25 | 4580380 | 1659389.25 | 1780588.75 | 1450554.5 | 2428940.5 | 1835320.12 | 5 | 1 | 11 | 9 | 11 | 9 | 3.723069299 | 1.497521795 | 2.797347672 | 1.125170866 |
| Q9JJF9 | Sppl2a | Signal peptide peptidase-like 2A | 41.24 | 1.53 | 58129 | 1 | 2 | 2 | 33412.34 | 2 | 2 | 33080.26 | 33682.33 | 113363.77 | 121146.61 | 204463.77 | 46032.05 | 35513.35 | 363713.94 | 203035.56 | 731963.5 | 52937.93 | 57320.18 | 167385.64 | 117643.42 | 185016.06 | 0 | 0 | 7 | 10 | 7 | 10 | 41.29995087 | 2.72920334 | 17.36377398 | 1.147441315 |
| Q64105 | Spr | Sepiapterin reductase | 646.75 | 47.13 | 27883 | 9 | 8596762 | 9521466 | 19387886 | 19838100 | 16915356 | 10012386 | 8632079 | 29663884 | 14790500 | 16886172 | 16210028 | 12070529 | 12621936 | 12403702 | 15322001 | 11995612 | 13624233 | 20681542 | 22953934 | 18263652 | 10 | 8 | 12 | 9 | 12 | 9 | 0.924166353 | 0.858013102 | 1.178554804 | 1.094192036 |
| Q4FZT8 | Spryd4 | SPRY domain-containing protein 4 | 46.52 | 3.38 | 23250 | 1 | 1641478.5 | 1091024.75 | 1147326.75 | 1131429.38 | 1640098.12 | 1726478.5 | 1417202 | 975941.94 | 1172939.62 | 1158785.12 | 1030590.38 | 868021.94 | 1081206.12 | 1133404.38 | 1551450.12 | 1717987.12 | 1159352.38 | 1612807.12 | 1652298 | 1390545.62 | 5 | 6 | 7 | 8 | 7 | 8 | 0.851656664 | 0.87806047 | 1.132549294 | 1.16766158 |
| O35704 | Sptlc1 | Serine palmitoyltransferase 1 | 126.36 | 6.55 | 52535 | 2 | 1334312.5 | 1189607.25 | 2 | 1351342 | 2 | 2 | 2 | 782693.44 | 466320.38 | 2153002.5 | 689958 | 596859.75 | 1025669.5 | 564098.31 | 149698.64 | 2 | 1838779.75 | 1200005.88 | 1422905.88 | 1314777.88 | 10 | 8 | 10 | 8 | 10 | 8 | 0.780923012 | 0.889555004 | 1.490600068 | 1.69795323 |
| P97363 | Sptlc2 | Serine palmitoyltransferase 2 | 94.69 | 4.64 | 62982 | 2 | 2 | 636194.88 | 784218.5 | 772791.12 | 325828.53 | 199649.91 | 280170.47 | 286343.78 | 416251.84 | 426494.09 | 493504.84 | 1518366 | 1776914.25 | 1085160.25 | 1651489.12 | 2 | 258981.05 | 726510.19 | 941528.5 | 903433.25 | 3 | 10.5 | 0 | 9 | 3 | 10.5 | 2.590450066 | 4.055810515 | 1.123626689 | 1.759237516 |
| P52019 | Sqle | Squalene monooxygenase | 209.32 | 10.14 | 63770 | 4 | 2 | 2 | 1744045.88 | 2 | 214764.5 | 2 | 2 | 1971132.38 | 355855.59 | 2 | 466247 | 445134.16 | 711531.94 | 652596.12 | 875228.94 | 2 | 2872722.25 | 294015.12 | 2 | 408057.19 | 5 | 9 | 5 | 10 | 5 | 10 | 1.608490817 | 1.353994983 | 1.824978899 | 1.536230264 |
| Q9R112 | Sqor | Sulfide:quinone oxidoreductase, mitochondrial | 912.84 | 31.11 | 50282 | 12 | 16160938 | 17841290 | 31438418 | 34465622 | 30591056 | 22223632 | 23337364 | 34282298 | 32039598 | 38688178 | 23796132 | 29805502 | 38293962 | 37668458 | 34698726 | 21636856 | 23002258 | 35057302 | 38254990 | 34027010 | 6 | 7 | 9 | 12 | 9 | 12 | 1.258744432 | 1.090931877 | 1.164609444 | 1.009346722 |
| Q64337 | Sqstm1 | Sequestosome-1 | 1253.64 | 70.14 | 48163 | 18 | 1451793.62 | 2246445.75 | 1834648 | 1120818.62 | 4345621.5 | 2448268 | 2032139.75 | 5285084 | 3792636.5 | 2957902 | 25558482 | 28205200 | 29292420 | 30737276 | 26396636 | 961176.19 | 2662543.25 | 4325963.5 | 4868603.5 | 3270165.5 | 0 | 8 | 0 | 12 | 0 | 12 | 12.74532594 | 8.488118021 | 1.462675964 | 0.974111315 |
| Q497V5 | Srbd1 | S1 RNA-binding domain-containing protein 1 | 117.71 | 2.14 | 110281 | 2 | 1322633 | 1283518.38 | 765956.5 | 1112442.62 | 872685.5 | 505730.78 | 1248252 | 868273 | 759636.56 | 935639.38 | 524848.12 | 1089177.12 | 1428847 | 833902.56 | 1044002.19 | 539162.38 | 2 | 1110706.12 | 545034.94 | 539515 | 10 | 2 | 9 | 7 | 10 | 7 | 0.918529068 | 1.139719939 | 0.510416274 | 0.633329554 |
| Q8BZX4 | Srek1 | Splicing regulatory glutamine/lysine-rich protein 1 | 68.59 | 2.02 | 56763 | 1 | 2 | 2 | 149068.36 | 127582.22 | 2 | 2 | 2 | 36581.3 | 2 | 72787.01 | 271561.5 | 118809.24 | 2 | 2 | 2 | 2 | 147143.78 | 2 | 2 | 74474.85 | 12.5 | 11.5 | 10.5 | 10.5 | 12.5 | 11.5 | 1.411051709 | 3.569181282 | 0.801082085 | 2.026295114 |
| D4A208 | Srgap2 | SLIT-ROBO Rho GTPase-activating protein 2 | 578.37 | 10.64 | 120878 | 11 | 9433355 | 11535967 | 9897790 | 10723176 | 8647508 | 7102847 | 9296492 | 10668539 | 11349170 | 6327836 | 5936505.5 | 6142075.5 | 3648795.25 | 5335513.5 | 5361621.5 | 7810840 | 4298244 | 9135708 | 10281437 | 10134199 | 0 | 7 | 0 | 10 | 0 | 10 | 0.525988665 | 0.590559387 | 0.829264644 | 0.931065728 |
| Q812A2 | Srgap3 | SLIT-ROBO Rho GTPase-activating protein 3 | 57.24 | 1.27 | 124420 | 1 | 2 | 217286.08 | 2 | 351501.03 | 383985.47 | 213831.41 | 2 | 316652.12 | 557336.88 | 251876.2 | 2 | 2 | 2 | 2 | 2 | 2 | 2 | 190923.53 | 2 | 480819.91 | 5 | 10 | 2.5 | 6.5 | 5 | 10 | 1.049564E-05 | 7.464365E-06 | 0.70504403 | 0.50141833 |
| Q6P069 | Sri | Sorcin | 59.31 | 5.56 | 21627 | 1 | 4549208 | 3679921 | 6511501 | 7057636 | 5594882.5 | 3575417.25 | 4071559.5 | 5233924.5 | 4959482.5 | 5972833.5 | 2905182.25 | 2743201.75 | 3901619 | 5129380 | 4151781.25 | 4603750.5 | 6399663 | 6280801 | 7400320 | 7075008 | 4 | 7 | 6 | 3 | 6 | 7 | 0.687440666 | 0.790786228 | 1.159397303 | 1.333693896 |
| Q64674 | Srm | Spermidine synthase | 414.73 | 36.75 | 33995 | 8 | 13714213 | 12298902 | 10338538 | 8934702 | 8277694 | 12858428 | 9748980 | 10285272 | 8398522 | 9336026 | 19217998 | 17457266 | 12088638 | 11657164 | 12772805 | 12271424 | 12995239 | 12574114 | 10747001 | 10356995 | 5 | 8 | 3 | 4 | 5 | 8 | 1.366473827 | 1.445741232 | 1.100454019 | 1.164289955 |
| P16254 | Srp14 | Signal recognition particle 14 kDa protein | 187.04 | 28.18 | 12510 | 3 | 244074.97 | 334826.31 | 2 | 2 | 722679.31 | 594999.25 | 2 | 552726.31 | 2 | 2 | 1372225.5 | 1408954.5 | 785962.25 | 1617634.5 | 1300888.88 | 2 | 674579.69 | 794429.12 | 582132.88 | 2 | 0 | 10 | 0 | 8 | 0 | 10 | 4.982899828 | 5.650855876 | 1.575883508 | 1.787130163 |
| Q9D7A6 | Srp19 | Signal recognition particle 19 kDa protein | 51.46 | 8.33 | 16191 | 1 | 2 | 2 | 2 | 2 | 2 | 258494.36 | 371691.5 | 2 | 2 | 2 | 1169767.38 | 1155451 | 1088914.38 | 767659.38 | 1043021.19 | 2 | 380564.72 | 519635.16 | 2 | 450782.34 | 0 | 5 | 0 | 7 | 0 | 7 | 522481.333 | 8.290829605 | 135098.622 | 2.143769708 |
| P14576 | Srp54 | Signal recognition particle 54 kDa protein | 106.3 | 6.15 | 55721 | 2 | 1726393 | 1707138.38 | 1962641.75 | 1618209.75 | 1892016.38 | 1441793.5 | 1579971.25 | 2 | 1782899.62 | 2 | 1733300.25 | 2119204.5 | 2277148 | 2105068.25 | 2540877.75 | 1806713.75 | 2 | 2115559.5 | 1904813.75 | 2315737.25 | 2 | 8 | 1 | 4 | 2 | 8 | 1.209871513 | 2.242735173 | 0.914266923 | 1.694773837 |
| Q8BMA6 | Srp68 | Signal recognition particle subunit SRP68 | 592.58 | 17.76 | 70574 | 10 | 3538575 | 5459216 | 3329790.25 | 2213799 | 5688544.5 | 4127733.5 | 3622973.5 | 4373182.5 | 5546809.5 | 4439063.5 | 6254830 | 5197185.5 | 5419242 | 5185843.5 | 4474042.5 | 1896241.88 | 2758291.25 | 4673435.5 | 3819788 | 3779511.75 | 8 | 10 | 4 | 6 | 8 | 10 | 1.311480088 | 1.199974152 | 0.836744011 | 0.765601547 |
| P49962 | Srp9 | Signal recognition particle 9 kDa protein | 66.87 | 12.79 | 10194 | 1 | 2 | 612658.25 | 683686.31 | 605986.69 | 689663.88 | 450960.78 | 770303.38 | 589586 | 417303.12 | 586694.5 | 2 | 633251.12 | 799958.88 | 395321.5 | 704316.25 | 2 | 734129.06 | 586507.12 | 528889.44 | 767944.94 | 10.5 | 12.5 | 12 | 12 | 12 | 12.5 | 0.977180769 | 0.899817663 | 1.009828495 | 0.929880677 |
| O70551 | Srpk1 | SRSF protein kinase 1 | 266.49 | 8.49 | 73088 | 4 | 1740105.5 | 2606641.5 | 2000471.75 | 1608278.75 | 1415783.75 | 1368655.5 | 1974083.75 | 688658.56 | 1985034.75 | 1656587.25 | 1080898.5 | 1517807.62 | 1581940.75 | 2066292 | 1867932.25 | 2702282.5 | 2764105.75 | 1847100.38 | 2896680.25 | 1774687.75 | 9 | 4 | 12 | 4 | 12 | 4 | 0.865929738 | 1.057585061 | 1.278892001 | 1.561947828 |
| Q9DBG7 | Srpra | Signal recognition particle receptor subunit alpha | 890.55 | 25.16 | 69623 | 13 | 8883435 | 10801511 | 10122705 | 10574733 | 9839620 | 10135145 | 9217604 | 7991960 | 10003137 | 11679127 | 14866605 | 17255778 | 17384604 | 11201151 | 17821756 | 8897067 | 5895018.5 | 10815798 | 10091625 | 12821886 | 0 | 12 | 1 | 12 | 1 | 12 | 1.563655126 | 1.601769173 | 0.966138159 | 0.989687748 |
| P47758 | Srprb | Signal recognition particle receptor subunit beta | 577.44 | 47.21 | 29579 | 10 | 1644052.25 | 3644542 | 2173715.25 | 1191985.88 | 1946561 | 1123986.75 | 2326468.5 | 1861481.88 | 3526959.25 | 3132839.5 | 8178715 | 7087527 | 6688063.5 | 4698027 | 6462670 | 2598112 | 2181117.5 | 1770034.25 | 2166121.5 | 1211288.12 | 0 | 11 | 0 | 9 | 0 | 11 | 3.123804466 | 2.766098654 | 0.936402967 | 0.829175774 |
| Q52KI8 | Srrm1 | Serine/arginine repetitive matrix protein 1 | 471.81 | 10.25 | 106862 | 6 | 8501666 | 8367737 | 5465718 | 6422313 | 7979181 | 9026008 | 7528744.5 | 7740896 | 6283068 | 6295670.5 | 4239778 | 4102588.5 | 3496546.25 | 3361611.5 | 3310263 | 9675304 | 7939394.5 | 8023116.5 | 6684109 | 8120188.5 | 0 | 10 | 0 | 6 | 0 | 10 | 0.503878413 | 0.501995796 | 1.100866601 | 1.096753486 |
| Q8BTI8 | Srrm2 | Serine/arginine repetitive matrix protein 2 | 603.32 | 6.03 | 294840 | 12 | 3638587.75 | 3782651 | 3040603 | 3756826.5 | 2246795 | 3402126.5 | 1990826 | 5004844.5 | 3833181 | 5430869.5 | 6518241 | 4401726.5 | 2354939.5 | 7140228.5 | 4867360.5 | 3199006.5 | 3012468 | 4607823 | 4639623 | 5166274 | 4 | 7 | 8 | 12 | 8 | 12 | 1.535486467 | 1.285865736 | 1.252633721 | 1.048995752 |
| Q99MR6 | Srrt | Serrate RNA effector molecule homolog | 630.18 | 18.29 | 100452 | 13 | 21995720 | 23455960 | 19229110 | 19756030 | 17280948 | 18112560 | 23224250 | 16981704 | 17988418 | 16172708 | 12476360 | 15949953 | 12719538 | 9912660 | 8687871 | 20492866 | 16400138 | 14768377 | 13165956 | 18199110 | 0 | 4 | 0 | 9 | 0 | 9 | 0.587374096 | 0.646049033 | 0.816243304 | 0.89778082 |
| Q6PDM2 | Srsf1 | Serine/arginine-rich splicing factor 1 | 678.64 | 43.95 | 27745 | 13 | 15907760 | 17264516 | 14064191 | 14594075 | 11815843 | 14174410 | 14533153 | 16057024 | 12581223 | 15604316 | 23469638 | 20748142 | 16556724 | 13756276 | 19264574 | 13768264 | 14594574 | 12225312 | 12645730 | 14690257 | 5 | 9 | 4 | 8 | 5 | 9 | 1.27359074 | 1.285746292 | 0.922301034 | 0.931103765 |
| Q9R0U0 | Srsf10 | Serine/arginine-rich splicing factor 10 | 260.63 | 19.08 | 31301 | 4 | 4095790.75 | 4472141.5 | 3417730 | 2484113.25 | 4796650.5 | 3177282.5 | 4385638 | 3396305 | 4919169 | 2251103.75 | 2091603.75 | 3286911.75 | 1475725.25 | 2 | 2095273 | 3532369 | 3229088 | 3175951 | 2614245.5 | 2795175.75 | 1 | 6 | 2 | 8 | 2 | 8 | 0.46451354 | 0.493643874 | 0.796558181 | 0.846511527 |
| Q62093 | Srsf2 | Serine/arginine-rich splicing factor 2 | 438.25 | 25.79 | 25476 | 6 | 14947994 | 28941844 | 31387342 | 33428616 | 29273396 | 33300920 | 30474002 | 28718134 | 24113236 | 29661070 | 25552486 | 21521948 | 20232016 | 21393772 | 19811762 | 30600030 | 34160210 | 29225562 | 30376608 | 28061872 | 5 | 11 | 1 | 10 | 5 | 11 | 0.786437306 | 0.741874213 | 1.104690351 | 1.042093601 |
| P84104 | Srsf3 | Serine/arginine-rich splicing factor 3 | 346.26 | 31.71 | 19330 | 6 | 12184573 | 15034282 | 26518374 | 25783702 | 20921014 | 14937576 | 13847682 | 22115274 | 19281516 | 22071534 | 11625829 | 11755492 | 19931674 | 18327930 | 18822078 | 15959072 | 16269203 | 22331548 | 22619292 | 22324894 | 6 | 12 | 7 | 6 | 7 | 12 | 0.801089654 | 0.872193808 | 0.990661909 | 1.078592363 |
| Q8VE97 | Srsf4 | Serine/arginine-rich splicing factor 4 | 259.94 | 10.22 | 55979 | 5 | 3537528.25 | 4868679.5 | 4742036 | 4323918.5 | 4666186.5 | 2645047.25 | 4120089.75 | 4388367.5 | 2900566.75 | 2536712.75 | 1416730.5 | 2549506.25 | 1225663 | 768305.38 | 863210.06 | 4682822.5 | 3670940 | 2106839 | 3316167.25 | 3213644.5 | 0 | 4 | 1 | 11 | 1 | 11 | 0.308216989 | 0.411277441 | 0.767465245 | 1.024087424 |
| O35326 | Srsf5 | Serine/arginine-rich splicing factor 5 | 418.67 | 25.65 | 30891 | 8 | 10937538 | 10982293 | 10987580 | 10292766 | 9157683 | 8857001 | 11269048 | 8214608 | 8681382 | 6688126 | 4166753 | 6373755 | 4360477.5 | 2824101.25 | 5229903.5 | 11962478 | 8997805 | 8260227 | 7920289 | 9187980 | 0 | 6 | 0 | 9 | 0 | 9 | 0.438424914 | 0.52516366 | 0.884848598 | 1.059908582 |
| G3V6S8 | Srsf6 | Serine/arginine-rich splicing factor 6 | 609.87 | 25.96 | 39025 | 10 | 14658255 | 17243126 | 9144687 | 8273567.5 | 7156091 | 9171984 | 16665326 | 7871629 | 14452997 | 7785860 | 18532228 | 17716734 | 10021170 | 7169104 | 8976417 | 8520730 | 16887218 | 13952871 | 7447790.5 | 7861654 | 9 | 11 | 10 | 11 | 10 | 11 | 1.105176628 | 1.115605215 | 0.968031168 | 0.977165633 |
| Q8BL97 | Srsf7 | Serine/arginine-rich splicing factor 7 | 358.95 | 23.22 | 30818 | 7 | 13855693 | 16376365 | 16275690 | 18174838 | 13899296 | 20434752 | 17082410 | 18086314 | 14864874 | 17854044 | 17138854 | 10114629 | 16049271 | 19054992 | 18746338 | 14901592 | 17872740 | 19857122 | 19992620 | 20088172 | 9 | 4 | 11 | 9 | 11 | 9 | 1.032096482 | 0.918273162 | 1.179817073 | 1.049702593 |
| Q9D975 | Srxn1 | Sulfiredoxin-1 | 45.05 | 6.62 | 14149 | 1 | 884547 | 1015408.5 | 419838.62 | 2 | 540690 | 1012313.19 | 1355964.38 | 424432.44 | 1356292.12 | 1661937.12 | 2668304.5 | 2834787.5 | 2518493 | 2501508.75 | 2235425.25 | 1430154.5 | 1663177.5 | 1931213.62 | 1946348.88 | 1901633 | 0 | 0 | 0 | 1 | 0 | 1 | 4.46026251 | 2.195603576 | 3.101755131 | 1.526866332 |
| P32067 | Ssb | Lupus La protein homolog | 904.95 | 29.4 | 47756 | 16 | 6337511 | 9612328 | 7216263 | 5035573 | 5421868.5 | 7294489 | 5495446 | 6151656 | 6590092.5 | 7342951.5 | 21787622 | 20509834 | 14319389 | 13757959 | 14239020 | 5670681.5 | 5397544 | 6107091 | 5579438 | 5849388.5 | 0 | 9 | 0 | 4 | 0 | 9 | 2.516505258 | 2.573833109 | 0.850717682 | 0.87009766 |
| Q9CYR0 | Ssbp1 | Single-stranded DNA-binding protein, mitochondrial | 266.79 | 36.18 | 17319 | 5 | 8055811.5 | 7497129 | 7120608 | 9846342 | 8257693 | 7805785.5 | 7630332 | 9821935 | 10083657 | 11084735 | 8307833 | 8087119 | 9858164 | 8537546 | 10510160 | 10211013 | 13767776 | 10558659 | 9443554 | 9924926 | 4 | 1 | 12 | 7 | 12 | 7 | 1.110924633 | 0.975754712 | 1.321950037 | 1.161103948 |
| Q7TPJ0 | Ssr1 | Translocon-associated protein subunit alpha | 296.52 | 15.99 | 35629 | 6 | 5429944.5 | 5059912 | 5510637.5 | 4019153.75 | 6217843.5 | 3664652.5 | 5438331 | 5794526 | 7370894.5 | 4657428.5 | 8908013 | 15967832 | 12839976 | 6555656 | 13749827 | 5586816.5 | 2573361.5 | 3653790 | 4448721.5 | 3942394.75 | 0 | 5 | 1 | 5 | 1 | 5 | 2.21138917 | 2.154856456 | 0.770084459 | 0.750397755 |
| Q08013 | Ssr3 | Translocon-associated protein subunit gamma | 93.1 | 7.57 | 21064 | 1 | 1990162.5 | 1654031.38 | 1374648.12 | 1178728.25 | 1448017 | 2074515.88 | 1456043.62 | 1468482.38 | 1474651.38 | 1440502.75 | 3474867.75 | 2769901.25 | 2115703 | 2559082 | 2402459.5 | 1356450.12 | 1848921.75 | 1651666.62 | 1671163.62 | 1477542 | 0 | 10 | 0 | 9 | 0 | 10 | 1.742444768 | 1.683305984 | 1.047106501 | 1.011567581 |
| Q07984 | Ssr4 | Translocon-associated protein subunit delta | 210.41 | 24.86 | 18980 | 3 | 1604991.25 | 2103111.5 | 1522137.38 | 1401177.25 | 1476040.88 | 1438376.25 | 1062950.5 | 1479739 | 1950872.38 | 2112323.5 | 10892649 | 5730854 | 6593353 | 5531906 | 6863768 | 1248981.88 | 1865891.5 | 1890804.75 | 3059338 | 1067931.88 | 0 | 12 | 0 | 12 | 0 | 12 | 4.392564088 | 4.427072569 | 1.126487207 | 1.135337018 |
| Q04931 | Ssrp1 | FACT complex subunit SSRP1 | 222.1 | 9.87 | 80914 | 4 | 1213887.38 | 1386102.38 | 582564.75 | 712948 | 1366580.75 | 1836165 | 1582354.25 | 1340123 | 1094968.38 | 1617337.25 | 940903.06 | 678691.12 | 504982.66 | 607014.19 | 742832.12 | 1263210.25 | 536042.44 | 1162496.25 | 574433.12 | 156094.97 | 6 | 5 | 0 | 2 | 6 | 5 | 0.660275214 | 0.465057876 | 0.701675952 | 0.494218048 |
| Q4KLK9 | Ssu72 | RNA polymerase II subunit A C-terminal domain phosphatase SSU72 | 49.25 | 6.19 | 22544 | 1 | 2 | 2 | 2 | 2 | 570810.69 | 481183.53 | 621870.94 | 563600.75 | 396803.56 | 2 | 258031.94 | 355347.75 | 2 | 2 | 2 | 2 | 429466.5 | 599957.62 | 813723 | 338183.62 | 11 | 5 | 3.5 | 12.5 | 11 | 12.5 | 1.07457184 | 0.297260649 | 3.821410858 | 1.057123431 |
| Q99L47 | St13 | Hsc70-interacting protein | 606.59 | 18.87 | 41656 | 9 | 22941456 | 24133922 | 27933810 | 21167742 | 27570188 | 17042504 | 24190704 | 29245124 | 26823918 | 21757036 | 20128512 | 22641602 | 16082786 | 15071298 | 18315630 | 24806658 | 18331928 | 19989994 | 25008376 | 21050166 | 1 | 6 | 4 | 9 | 4 | 9 | 0.745389707 | 0.774738629 | 0.882340727 | 0.917081949 |
| O88829 | St3gal5 | Lactosylceramide alpha-2,3-sialyltransferase | 104.41 | 7.97 | 47360 | 3 | 1508451 | 1490368.25 | 3024865.75 | 1947725.38 | 1905721.5 | 873787.19 | 1701127.38 | 2284464 | 2484890 | 1780576.88 | 616628.75 | 1537065.38 | 1305410.5 | 714953.62 | 1379898.38 | 944010.75 | 307755.94 | 1113385.62 | 1104520.25 | 645942.31 | 2 | 0 | 3 | 3 | 3 | 3 | 0.562304594 | 0.60866309 | 0.41668117 | 0.451033926 |
| O35638 | Stag2 | Cohesin subunit SA-2 | 52.73 | 1.22 | 141281 | 1 | 160922.34 | 120282.05 | 2 | 144687.84 | 150210.31 | 165798.66 | 194188.08 | 174453.09 | 164480 | 197183.12 | 2 | 149381.17 | 217339.84 | 2 | 2 | 181781.84 | 221659.28 | 2 | 2 | 120897.95 | 9.5 | 12 | 5 | 8 | 9.5 | 12 | 0.636563305 | 0.409246516 | 0.910152643 | 0.585137087 |
| P70297 | Stam | Signal transducing adapter molecule 1 | 45.08 | 1.28 | 59771 | 1 | 594976.31 | 494790.69 | 421963.38 | 478133.44 | 386811.88 | 508226.12 | 433856.09 | 387892.28 | 466577.19 | 2 | 511112.38 | 931898.12 | 1072674.12 | 400573.5 | 664809.56 | 471562.31 | 2 | 342161.09 | 399405.34 | 418445.81 | 5 | 4 | 3 | 9.5 | 5 | 9.5 | 1.506754868 | 1.993298458 | 0.686495238 | 0.90817022 |
| O88811 | Stam2 | Signal transducing adapter molecule 2 | 92.88 | 4.78 | 57455 | 2 | 2 | 2 | 2 | 2 | 255056.23 | 290814.09 | 474631.09 | 734466.25 | 279305.66 | 222598.14 | 711042.31 | 541629.94 | 2 | 315834.38 | 375815.62 | 2 | 2 | 2 | 2 | 2 | 3 | 10 | 11 | 0 | 11 | 10 | 7.622880911 | 0.971280576 | 3.920581E-05 | 4.995466E-06 |
| Q8R424 | Stambp | STAM-binding protein | 134.07 | 8.25 | 48512 | 3 | 1578720.12 | 1685489.25 | 1958346.75 | 2110224.75 | 2225203 | 1847553.38 | 1852332.75 | 3455404.25 | 2718708.25 | 2438574.75 | 1317026.5 | 1788807 | 1627179.25 | 1976353.62 | 1728325.38 | 1697981.75 | 1428464 | 2276739.5 | 2418442.75 | 2345869.5 | 8 | 8 | 2 | 6 | 8 | 8 | 0.882789913 | 0.685290677 | 1.063770104 | 0.825781677 |
| Q76N33 | Stambpl1 | AMSH-like protease | 63 | 4.59 | 49640 | 1 | 968191.56 | 1136997.25 | 922286 | 1162122.25 | 964085.81 | 932400.75 | 1407890.12 | 1073707 | 1212732.25 | 1124464.25 | 571817.5 | 886405.06 | 994708.38 | 858282.62 | 2 | 1301703.75 | 1057825.25 | 1290474.75 | 794081.19 | 1297895.12 | 3 | 7 | 1 | 12 | 3 | 12 | 0.642495016 | 0.575743984 | 1.114150832 | 0.998397844 |
| Q9JM90 | Stap1 | Signal-transducing adaptor protein 1 | 528.62 | 51.85 | 34628 | 12 | 10493390 | 12171022 | 17410320 | 17362668 | 15931860 | 18146626 | 15900156 | 23202226 | 22330234 | 25613038 | 31580526 | 26355964 | 32139602 | 23570396 | 30711744 | 12507830 | 11517922 | 25113514 | 17780154 | 22784918 | 0 | 7 | 1 | 8 | 1 | 8 | 1.967557421 | 1.372327247 | 1.222641989 | 0.852765412 |
| P42225 | Stat1 | Signal transducer and activator of transcription 1 | 870.44 | 21.36 | 87197 | 14 | 15190246 | 20221744 | 16674483 | 18256158 | 24827878 | 11540651 | 20518686 | 23817032 | 25944386 | 22072716 | 17075622 | 25652304 | 22784080 | 13681152 | 22380990 | 14362149 | 8133001 | 12770302 | 12087104 | 16204479 | 10 | 1 | 12 | 4 | 12 | 4 | 1.067285959 | 0.97767595 | 0.667822792 | 0.611751965 |
| Q9WVL2 | Stat2 | Signal transducer and activator of transcription 2 | 431.2 | 8.23 | 105417 | 8 | 2994720 | 3259707.25 | 3116399.75 | 4517291 | 4329102.5 | 3896341.25 | 3674515.25 | 3174896 | 3791599 | 3353165 | 1867926.12 | 3047160.75 | 4086024.5 | 2480172.25 | 2460988.25 | 2644342.75 | 2756682.75 | 2260161 | 2868967 | 3869501.5 | 4 | 3 | 5 | 4 | 5 | 4 | 0.765334749 | 0.779310752 | 0.790441934 | 0.804876427 |

| | | | | | | | | | | | | | | | | | | | | | | | | | | | | | | | | | | | | |
|---|---|---|---|---|---|---|---|---|---|---|---|---|---|---|---|---|---|---|---|---|---|---|---|---|---|---|---|---|---|---|---|---|---|---|---|---|
| P42227 | Stat3 | Signal transducer and activator of transcription 3 | 279.55 | 10.39 | 88054 | 5 | 2409887 | 2424468 | 3072996.5 | 2249426 | 2716452.5 | 2033209.5 | 2176323.5 | 6053349 | 2754671.5 | 2042811.75 | 2332530 | 2164906.5 | 2049241.75 | 1160529.25 | 3348259 | 2506687 | 1842355.75 | 2738012.5 | 2307584.5 | 2380756.75 | 6 | 9 | 11 | 12 | 11 | 12 | 0.858795073 | 0.734076918 | 0.914719655 | 0.781879875 |
| P52633 | Stat6 | Signal transducer and transcription activator 6 | 44.19 | 2.15 | 93497 | 1 | 205695.19 | 229973.66 | 2 | 2 | 2 | 261475.75 | 162797.86 | 2 | 2 | 2 | 342154.78 | 266777.97 | 252979.62 | 1522707.5 | 327992.72 | 2 | 215311.94 | 103722.41 | 2 | 2 | 0 | 11.5 | 1 | 11.5 | 1 | 11.5 | 6.226231764 | 6.393454991 | 0.732290032 | 0.751957771 |
| P35173 | Stfa3 | Stefin-3 | 375.91 | 61.17 | 11640 | 8 | 266558.41 | 279679.75 | 2 | 2 | 2 | 2 | 2 | 2 | 157418.36 | 2 | 4211531.5 | 6178260.5 | 114237.37 | 2 | 584903.5 | 2 | 2 | 137406 | 2 | 2 | 5.5 | 9 | 4 | 12 | 5.5 | 12 | 20.30032663 | 70.43886977 | 0.2515615 | 0.87287796 |
| Q60864 | Stip1 | Stress-induced-phosphoprotein 1 | 1674.58 | 53.41 | 62582 | 30 | 60641998 | 68856458 | 77419146 | 71272322 | 67208698 | 57467882 | 70964122 | 76693562 | 68995274 | 60131582 | 45957194 | 54877430 | 45202874 | 41257334 | 44543050 | 72501842 | 47892182 | 63093210 | 58956938 | 63316910 | 0 | 6 | 0 | 9 | 0 | 9 | 0.671218318 | 0.693601203 | 0.885241175 | 0.914761007 |
| O55098 | Stk10 | Serine/threonine-protein kinase 10 | 317.37 | 7.35 | 111906 | 6 | 3226348.25 | 3269965.75 | 1500675.75 | 3328566.5 | 2615094.75 | 2948284 | 3113582.5 | 2289352 | 2339252 | 2056819.25 | 2803016 | 2931004.75 | 1776468.12 | 2001977.38 | 2462995.25 | 3239231 | 1890415.25 | 2486952 | 2114820.25 | 2239043.75 | 7 | 7 | 9 | 10 | 9 | 10 | 0.859031727 | 0.93945158 | 0.858673117 | 0.939059399 |
| Q3TAA7 | Stk11ip | Serine/threonine-protein kinase 11-interacting protein | 126.66 | 2.71 | 118005 | 2 | 965213.62 | 1001463.25 | 321509.12 | 1154242.12 | 905046.38 | 1154486.25 | 847609.5 | 1156196 | 1016858.25 | 878303.06 | 775808.81 | 702731.44 | 861731.25 | 1014576.19 | 503618.62 | 860628.38 | 454392.28 | 987899.25 | 1147854.25 | 1157975.75 | 8 | 11 | 3 | 11 | 8 | 11 | 0.887519023 | 0.763530652 | 1.060098207 | 0.912000142 |
| B0LT89 | Stk24 | Serine/threonine-protein kinase 24 | 219.81 | 15.31 | 47990 | 4 | 868059.12 | 1145763.12 | 1066439 | 644190.94 | 648120.44 | 1333364.25 | 922208.44 | 1023541.88 | 884072.5 | 1033282.5 | 432859.62 | 437901.97 | 798315.12 | 222136.58 | 457158.12 | 1188569 | 1183854.88 | 478888 | 2 | 846363.62 | 2 | 12 | 0 | 8 | 2 | 12 | 0.53706859 | 0.451916706 | 0.845652622 | 0.711574936 |
| Q9JI10 | Stk3 | Serine/threonine-protein kinase 3 | 323.75 | 19.72 | 56855 | 7 | 2033628.25 | 1741370.25 | 1650258 | 376582.19 | 2485376.75 | 528989.5 | 2665760.25 | 1233831.5 | 2247299 | 707906.12 | 913360.62 | 2202276.5 | 643447.75 | 1107219.38 | 2143665 | 2320003 | 207170.92 | 748104.12 | 831279.25 | 474089.66 | 11 | 7 | 11 | 8 | 11 | 8 | 0.845877521 | 0.949373248 | 0.552736559 | 0.620365585 |
| Q9JI11 | Stk4 | Serine/threonine-protein kinase 4 | 247.22 | 13.96 | 55541 | 5 | 190670.11 | 987513.75 | 1110420.5 | 2 | 1655742.5 | 468273.56 | 954659.25 | 815627.62 | 1278122 | 458096.16 | 2 | 623075.94 | 284974.41 | 2 | 243518.3 | 343591.09 | 2 | 415095.66 | 555072.12 | 377386.16 | 7 | 8.5 | 2 | 2 | 7 | 8.5 | 0.291955071 | 0.289719949 | 0.42875189 | 0.425469493 |
| P54227 | Stmn1 | Stathmin | 232.43 | 30.87 | 17274 | 4 | 1617348.12 | 1145098.88 | 2292979.5 | 2209439 | 1383900.5 | 1768516.88 | 1443395 | 1681724.38 | 1550935.38 | 1769040.75 | 2095532.5 | 950760.94 | 818187.75 | 1334130 | 1002633.25 | 1468354.38 | 2324538 | 1527054.88 | 1323993.62 | 1374668.25 | 4 | 11 | 5 | 7 | 5 | 11 | 0.717009159 | 0.754995993 | 0.927139101 | 0.976258527 |
| Q6AYD2 | Stn1 | CST complex subunit STN1 | 61.44 | 1.96 | 46783 | 1 | 738034.69 | 694909.94 | 874280.88 | 1197826.88 | 841897.69 | 1164587.62 | 512084.84 | 978744.06 | 916877.94 | 1417687 | 670599.81 | 691053.88 | 835298.75 | 2 | 664846.25 | 753674.88 | 1170137.12 | 997569.56 | 1141868.62 | 1042972.25 | 2 | 7 | 4 | 11 | 4 | 11 | 0.658346803 | 0.573509283 | 1.174667833 | 1.02329487 |
| P54116 | Stom | Erythrocyte band 7 integral membrane protein | 814.88 | 51.06 | 31375 | 13 | 92341338 | 93551802 | 55041202 | 65966150 | 65776542 | 108484890 | 99222178 | 64689742 | 69707938 | 64106626 | 121065202 | 122221826 | 102101578 | 99342218 | 113663586 | 81509154 | 101558538 | 54415990 | 62695590 | 64967366 | 0 | 10 | 2 | 9 | 2 | 10 | 1.498333299 | 1.374640017 | 0.979793775 | 0.898907961 |
| Q99JB2 | Stoml2 | Stomatin-like protein 2, mitochondrial | 819.92 | 39.09 | 38385 | 13 | 3247057.75 | 4795111.5 | 6000894 | 4381175 | 5679558.5 | 5151733 | 4573503 | 6593433 | 4864398.5 | 7093289 | 13879182 | 12203344 | 14314509 | 13304579 | 14631388 | 3618325.5 | 4639076 | 5955419.5 | 7335541 | 6708059 | 0 | 8 | 0 | 12 | 0 | 12 | 2.834947652 | 2.416612692 | 1.172280919 | 0.999294976 |
| Q5XIG8 | Strap | Serine-threonine kinase receptor-associated protein | 580.18 | 46.29 | 38456 | 10 | 7591131.5 | 11770988 | 12785068 | 8313877.5 | 10723248 | 7296734.5 | 10672625 | 11024789 | 8449587 | 7188077.5 | 8112180 | 13929784 | 3731211.5 | 6371791 | 6865514.5 | 13879658 | 4840809 | 5826592 | 5490896 | 10217425 | 6 | 7 | 7 | 8 | 7 | 8 | 0.762156972 | 0.874051005 | 0.786478857 | 0.901943643 |
| Q8C079 | Strip1 | Striatin-interacting protein 1 | 126.23 | 3.23 | 95585 | 3 | 1562713.75 | 502404.88 | 1208587.75 | 1054526.12 | 1637548 | 987432.75 | 497703.66 | 1129510.25 | 1690999.38 | 1068504.62 | 240533.05 | 613878.38 | 359929.12 | 778762.62 | 758764.31 | 814717.69 | 711580.75 | 452649.91 | 972676.25 | 948182.62 | 3 | 4 | 3 | 4 | 3 | 4 | 0.461275349 | 0.512056259 | 0.653696062 | 0.725660196 |
| P58404 | Strn4 | Striatin-4 | 46.16 | 2.24 | 81645 | 1 | 889791.56 | 2 | 593704.19 | 2 | 531959.88 | 2 | 2 | 2 | 671361.06 | 561512.94 | 548449.75 | 652041.81 | 349479.34 | 2 | 582922.94 | 851570.12 | 729044.31 | 735284.62 | 688369.44 | 2 | 12 | 8 | 9.5 | 3.5 | 12 | 8 | 1.058267706 | 1.730010901 | 1.490613082 | 2.436790677 |
| P46978 | Stt3a | Dolichyl-diphosphooligosaccharide--protein glycosyltransferase subunit STT3A | 742.2 | 23.55 | 80598 | 14 | 6265917 | 6299527.5 | 8810481 | 8555198 | 11637380 | 5810295.5 | 6522062.5 | 11171837 | 13748020 | 10351836 | 20175936 | 19876758 | 24285232 | 21233918 | 26076642 | 8345627 | 7653878 | 11265142 | 10778248 | 9083614 | 0 | 9 | 0 | 12 | 0 | 12 | 2.685891399 | 2.345356827 | 1.133707135 | 0.989968459 |
| Q3TDQ1 | Stt3b | Dolichyl-diphosphooligosaccharide--protein glycosyltransferase subunit STT3B | 171.27 | 5.1 | 93246 | 4 | 2248959.5 | 2001860 | 2059143.38 | 2013612.75 | 1022686.25 | 2201152.75 | 2223372 | 1766951.75 | 1383887.5 | 2808298 | 3212638 | 4031338.25 | 3849412.25 | 3625664.25 | 5197618 | 1275616.5 | 450539.84 | 2204986.5 | 2967601.75 | 2462916.75 | 0 | 10 | 0 | 12 | 0 | 12 | 2.130977176 | 1.918077722 | 1.00164766 | 0.901576086 |
| Q9WUD1 | Stub1 | STIP1 homology and U box-containing protein 1 | 98.02 | 7.57 | 34909 | 2 | 3633414 | 3433135.5 | 3413697.5 | 1360550 | 2743220.25 | 3084917 | 3096468.5 | 2422199 | 2653850 | 2761489 | 2764643.75 | 3266364.5 | 1617037.38 | 2769006 | 1120524.38 | 3617563 | 4039921 | 3427116 | 3571265 | 2775813.25 | 7 | 7 | 11 | 2 | 11 | 7 | 0.791110968 | 0.82300014 | 1.195259026 | 1.243439145 |
| G3V7P1 | Stx12 | Syntaxin-12 | 105.92 | 8.03 | 31187 | 2 | 1719255.62 | 1305995 | 1133618.62 | 1473813.5 | 1893167.75 | 1800021.12 | 2088166.75 | 1544452.25 | 1622441.12 | 1265656.12 | 585853.19 | 782807.81 | 457016.34 | 2 | 764505 | 1801625.25 | 1706609.25 | 1024475.19 | 424019.41 | 1220210.5 | 0 | 8 | 0 | 7 | 0 | 8 | 0.344171645 | 0.311292645 | 0.820762997 | 0.742354834 |
| Q08849 | Stx3 | Syntaxin-3 | 78.75 | 4.84 | 33257 | 1 | 432458.69 | 501518.94 | 225729.06 | 2 | 646878 | 226879.25 | 554609.19 | 227072.75 | 493059.22 | 299282.03 | 139418.73 | 433584.03 | 2 | 2 | 205169.52 | 374707.75 | 2 | 2 | 173964.33 | 248626.97 | 6 | 6 | 3 | 5 | 6 | 6 | 0.430743946 | 0.432103518 | 0.441331188 | 0.442724177 |
| P70452 | Stx4 | Syntaxin-4 | 93.88 | 9.06 | 34165 | 2 | 969219.75 | 2 | 1021763.5 | 1214278.88 | 175930.44 | 517646.75 | 1119727.75 | 2 | 2 | 1244888.38 | 2 | 1023416.88 | 1101597.88 | 1260600.88 | 1526316.75 | 1206732 | 2 | 2 | 2 | 1465786 | 6.5 | 10.5 | 8 | 12 | 8 | 12 | 1.452721601 | 1.704191386 | 0.790408225 | 0.927229889 |
| Q8K1E0 | Stx5 | Syntaxin-5 | 137.32 | 10.42 | 39713 | 2 | 346727.41 | 568969.69 | 395473.34 | 2 | 2 | 2 | 563798.38 | 392773.66 | 880050.5 | 2 | 335988.06 | 700640.88 | 360601.47 | 267024.53 | 396951.94 | 246784.72 | 2 | 206135.14 | 2 | 2 | 9 | 7 | 12 | 7 | 12 | 7 | 1.572031011 | 1.122278719 | 0.345435242 | 0.246607489 |
| O70439 | Stx7 | Syntaxin-7 | 351.45 | 31.03 | 29821 | 6 | 7253678 | 5556575.5 | 7881542 | 10734388 | 8796522 | 6502048 | 6835730 | 8374755 | 7810959.5 | 8618078 | 5882819.5 | 4659002.5 | 6912013 | 8071952.5 | 5756211 | 6762232.5 | 10606674 | 9824234 | 10883201 | 9401004 | 6 | 7 | 5 | 4 | 6 | 7 | 0.7777199 | 0.820154967 | 1.180361811 | 1.244766403 |
| O88983 | Stx8 | Syntaxin-8 | 365.45 | 26.69 | 26925 | 6 | 2183242.25 | 2031838.12 | 1736252.38 | 1043383.5 | 1758451 | 2198775.5 | 2337117.5 | 1311888.5 | 1790352.5 | 1610645.5 | 1845459.75 | 2155532.5 | 2588980.5 | 2864797.75 | 2170086.75 | 2150621.75 | 2225218.5 | 2722657 | 2750273.75 | 2111508.25 | 4 | 2 | 6 | 5 | 6 | 5 | 1.328074389 | 1.256907168 | 1.366394461 | 1.293173791 |
| O08599 | Stxbp1 | Syntaxin-binding protein 1 | 253.54 | 9.6 | 67569 | 5 | 2830521.25 | 2553104 | 2806965.75 | 3315331.75 | 3158927 | 3347387.25 | 1027199.81 | 3041361.5 | 3251000.5 | 3451557.5 | 1113120.25 | 1857823.5 | 2303259 | 1247540.88 | 2527596.5 | 1556264.25 | 3031557 | 3256119 | 3614570 | 4088398.5 | 0 | 8 | 5 | 10 | 5 | 10 | 0.617076907 | 0.640955904 | 1.060147838 | 1.101172329 |
| Q64324 | Stxbp2 | Syntaxin-binding protein 2 | 838.44 | 29.85 | 66357 | 13 | 8264539.5 | 10177083 | 9969377 | 10936564 | 14781492 | 8387572.5 | 9428416 | 9893411 | 12293981 | 8434021 | 6286799 | 7404432 | 7382004.5 | 5837009 | 7525609 | 7825187.5 | 3038385.5 | 7859298 | 8973492 | 9092270 | 0 | 2 | 0 | 4 | 0 | 4 | 0.636180572 | 0.710935195 | 0.679646683 | 0.759508806 |
| Q60770 | Stxbp3 | Syntaxin-binding protein 3 | 107.12 | 3.38 | 67942 | 2 | 2 | 762512.25 | 602319.19 | 862775.88 | 656877.12 | 464605.66 | 2 | 1075808.38 | 591125.94 | 932466.19 | 2 | 865739.25 | 714719.38 | 839812.31 | 941881.75 | 2 | 2 | 1001337.38 | 2 | 702812.94 | 7.5 | 9.5 | 11.5 | 8.5 | 11.5 | 9.5 | 1.165599062 | 1.097306046 | 0.5908006 | 0.556185305 |
| Q8K400 | Stxbp5 | Syntaxin-binding protein 5 | 54.98 | 1.04 | 127651 | 1 | 521611.69 | 2 | 985762.75 | 1300739.88 | 2 | 2 | 2 | 991364.88 | 2 | 1168492.12 | 2 | 636272.25 | 634469 | 666027.38 | 2 | 2 | 2 | 962248.94 | 1412182.25 | 1052503.88 | 11 | 11 | 12 | 10 | 12 | 11 | 0.689704781 | 0.89671087 | 1.220368474 | 1.5866465 |
| P11031 | Sub1 | Activated RNA polymerase II transcriptional coactivator p15 | 196.06 | 33.07 | 14427 | 4 | 5567745.5 | 7847156.5 | 5433774.5 | 6893635 | 9637934 | 6392736.5 | 4532797.5 | 8208163 | 8172471.5 | 6586468.5 | 4462873 | 6141244.5 | 2827539 | 2772189.5 | 3344648 | 10151775 | 10043001 | 3875539.5 | 5310045 | 5722626 | 2 | 12 | 1 | 12 | 2 | 12 | 0.552525674 | 0.576777015 | 0.992163452 | 1.035711281 |
| Q9Z2I9 | Sucla2 | Succinate--CoA ligase [ADP-forming] subunit beta, mitochondrial | 379.02 | 20.3 | 50114 | 8 | 8267958 | 11891756 | 9807541 | 9745442 | 11189640 | 8115380 | 11404510 | 16291266 | 11835889 | 10944860 | 6647734 | 9511638 | 8385688 | 11521565 | 13197789 | 8729625 | 7301525 | 11203993 | 11242852 | 13635176 | 11 | 11 | 9 | 9 | 11 | 11 | 0.967822244 | 0.840805808 | 1.023787395 | 0.889426125 |
| Q9WUM5 | Suclg1 | Succinate--CoA ligase [ADP/GDP-forming] subunit alpha, mitochondrial | 389.17 | 28.61 | 36155 | 7 | 266026.28 | 146935.88 | 629895.81 | 336985.84 | 497807.41 | 778835 | 435396.75 | 421639.06 | 2 | 2 | 4491276.5 | 3400693.25 | 3587354.75 | 3475179 | 2992594.5 | 2 | 955763.31 | 2155392.5 | 570741.94 | 2094940.75 | 0 | 6 | 0 | 5 | 0 | 6 | 9.558270359 | 10.97094832 | 3.076631293 | 3.531346326 |
| Q9Z2I8 | Suclg2 | Succinate--CoA ligase [GDP-forming] subunit beta, mitochondrial | 880.65 | 40.88 | 46840 | 13 | 10280670 | 10705420 | 10269305 | 5408170 | 11952663 | 11235765 | 9224445 | 9855076 | 11100422 | 8714956 | 12401597 | 13851580 | 15877882 | 12241767 | 13992783 | 8732808 | 5630603 | 10992689 | 10710215 | 10440839 | 0 | 12 | 0 | 10 | 0 | 12 | 1.40623022 | 1.363748324 | 0.956617901 | 0.927718691 |
| Q9CX34 | Sugt1 | Protein SGT1 homolog | 227.78 | 15.77 | 38159 | 4 | 1592947.5 | 3544906.75 | 3653119 | 3695952 | 4184516.25 | 992010.12 | 2334251.25 | 4007306.5 | 3961089.25 | 4800442.5 | 19229942 | 2791940.5 | 2922712.5 | 3176703 | 2399710 | 3526220 | 3020503.25 | 4058051.25 | 3675885 | 4152477.25 | 9 | 12 | 12 | 11 | 12 | 12 | 1.830735992 | 1.89629196 | 1.105671441 | 1.145263912 |
| Q8R0F3 | Sumf1 | Formylglycine-generating enzyme | 172.93 | 11.83 | 40659 | 3 | 1668955.88 | 1440478.38 | 995784.12 | 1315132.12 | 1283694 | 1613124.25 | 1272631.25 | 752226.06 | 950690.25 | 1668558.75 | 1305645.75 | 1327368 | 1306579.62 | 1160713.75 | 1431787.75 | 1498957 | 930479.12 | 1475689.75 | 775642.44 | 861578.5 | 11 | 8 | 11 | 9 | 11 | 9 | 0.974351359 | 1.04392747 | 0.826716889 | 0.885750774 |
| P61957 | Sumo2 | Small ubiquitin-related modifier 2 | 78.56 | 12.63 | 10871 | 1 | 4775276.5 | 6700373 | 7066345.5 | 5891485 | 5417384 | 7045077 | 5835696 | 8913092 | 7494279 | 10235150 | 7207287.5 | 8110611.5 | 8296581.5 | 9112198 | 8674276 | 4994726.5 | 5416452 | 8803607 | 9635162 | 9859506 | 0 | 8 | 10 | 11 | 10 | 11 | 1.386926506 | 1.047507693 | 1.296761578 | 0.979408586 |
| Q8BJS4 | Sun2 | SUN domain-containing protein 2 | 276.31 | 13 | 81605 | 6 | 2 | 258487.31 | 1483114.88 | 1498523.25 | 1129878.75 | 337749.91 | 418505.84 | 1130413 | 804252.06 | 2119391.5 | 562468.06 | 439829.06 | 1751986.5 | 2025948.62 | 3073066 | 2 | 380469.69 | 1368341.5 | 480596.91 | 265386.59 | 6 | 9.5 | 8 | 7 | 8 | 9.5 | 1.797090873 | 1.632596333 | 0.570890882 | 0.518635076 |
| Q920B9 | Supt16h | FACT complex subunit SPT16 | 1017.63 | 18.15 | 119825 | 18 | 17750218 | 15314425 | 19065550 | 18994786 | 16594437 | 14050922 | 14326595 | 13621839 | 15304444 | 13320341 | 11515145 | 10904390 | 12699456 | 9708514 | 11218954 | 14280917 | 16959898 | 11255006 | 14666848 | 13371911 | 0 | 2 | 0 | 12 | 0 | 12 | 0.638928775 | 0.793587833 | 0.804093133 | 0.998731864 |
| O55201 | Supt5h | Transcription elongation factor SPT5 | 329.43 | 7.67 | 120664 | 6 | 4276605 | 4065461.25 | 3637201.25 | 4089017.5 | 3288601.75 | 5345375 | 3834117.5 | 3474317.5 | 3370784 | 3484663.5 | 3631732.25 | 2055235 | 1810321.12 | 1841415.62 | 4398005 | 4200435.5 | 5160890 | 3253116.5 | 963337.75 | 2656638 | 6 | 9 | 7 | 8 | 7 | 9 | 0.709654872 | 0.704112342 | 0.838689504 | 0.832139191 |
| Q62383 | Supt6h | Transcription elongation factor SPT6 | 102.72 | 1.33 | 199086 | 2 | 2 | 2 | 2 | 334848.75 | 584775.94 | 319071.69 | 512746.66 | 177594.94 | 568425.88 | 365806.84 | 88105.25 | 69810.42 | 173537.77 | 2 | 2 | 577105.31 | 465375.69 | 2 | 559150.88 | 331658.97 | 12 | 8.5 | 0 | 11 | 12 | 11 | 0.360424509 | 0.170533851 | 2.102249165 | 0.99467333 |
| Q64310 | Surf4 | Surfeit locus protein 4 | 390.08 | 24.54 | 30381 | 6 | 1767345.62 | 2316656.25 | 1433498.25 | 792287.88 | 3037570.5 | 2517544.75 | 7970820 | 3036490.5 | 4339631.5 | 1194346 | 4340165 | 4017280.75 | 5784855 | 5199556 | 4739178 | 2008746.12 | 3897484.5 | 3418967.5 | 2736783.25 | 3025052 | 0 | 4 | 6 | 11 | 6 | 11 | 2.57623956 | 1.263510471 | 1.614042445 | 0.79160322 |
| P70279 | Surf6 | Surfeit locus protein 6 | 63.68 | 2.82 | 41235 | 1 | 2 | 2 | 2 | 2 | 2 | 2 | 2 | 2 | 2 | 2 | 439448.81 | 279553.12 | 2 | 2 | 2 | 2 | 2 | 2 | 2 | 2 | 7.5 | 12.5 | 7.5 | 12.5 | 7.5 | 12.5 | 71900.793 | 71900.793 | 1 | 1 |
| Q6A028 | Swap70 | Switch-associated protein 70 | 448.95 | 18.8 | 68996 | 9 | 4625262.5 | 5905758 | 9236765 | 7586059.5 | 7828983 | 4957702 | 5720579 | 10953964 | 10354843 | 10214384 | 8321118 | 10166607 | 8120107 | 13133908 | 9553778 | 4267708 | 3780981.75 | 8005628.5 | 9044627 | 7420454 | 2 | 10 | 12 | 6 | 12 | 10 | 1.401124378 | 1.168099492 | 0.92429748 | 0.770574999 |
| P48025 | Syk | Tyrosine-protein kinase SYK | 262.77 | 12.24 | 71376 | 5 | 424650.44 | 1579013.25 | 1552581.38 | 1399843.62 | 2373728.5 | 1508722.75 | 1243663.5 | 638870.62 | 783504.25 | 2089194.12 | 877336 | 2074101.38 | 1922007.25 | 728629.5 | 1438969.5 | 1531455.25 | 1821804.75 | 1889443.88 | 1585712 | 1819092.38 | 12 | 7 | 11 | 5 | 12 | 7 | 0.960602897 | 1.124057143 | 1.179771342 | 1.380518846 |
| Q80X82 | Sympk | Symplekin | 278.58 | 5.61 | 142284 | 6 | 1987055.5 | 1715299 | 1010320.06 | 1038999.88 | 1816224.75 | 1609541.62 | 1531388.5 | 1443532.75 | 2274122 | 2987971.25 | 3190075.75 | 3778085 | 3465361.5 | 2437852.25 | 3442991.25 | 1638325.25 | 1884755 | 1326838 | 1325411.88 | 1515635.25 | 0 | 12 | 1 | 7 | 1 | 12 | 2.155732435 | 1.656860079 | 1.016261605 | 0.78108176 |
| Q7TMK9 | Syncrip | Heterogeneous nuclear ribonucleoprotein Q | 912.04 | 28.41 | 69633 | 16 | 27590934 | 28765316 | 26831762 | 27516798 | 27919320 | 27714686 | 28541028 | 22041382 | 25947028 | 25245578 | 32498486 | 37939514 | 31049280 | 26364940 | 30722858 | 28370962 | 27309780 | 26859600 | 24046460 | 24090228 | 5 | 6 | 2 | 12 | 5 | 12 | 1.143921177 | 1.22461536 | 0.942671597 | 1.009169285 |
| Q6ZWR6 | Syne1 | Nesprin-1 | 120.6 | 0.31 | 1009926 | 3 | 2 | 96423.89 | 438364.69 | 2 | 1568945.75 | 2 | 2 | 500783.81 | 1409141.5 | 1296424.62 | 248898.22 | 232644.98 | 679580.75 | 2 | 412342.44 | 276885.38 | 105403.98 | 154308.08 | 407864.72 | 259670.75 | 11 | 10 | 10 | 10 | 11 | 10 | 0.747939213 | 0.490734468 | 0.572377702 | 0.375545849 |
| O55100 | Syngr1 | Synaptogyrin-1 | 66.49 | 5.13 | 25653 | 1 | 1289213 | 1003822.62 | 767296.56 | 980711.88 | 723178.69 | 1422675 | 1250296.38 | 984970.31 | 950670.94 | 1111251 | 1294552.12 | 968464.94 | 1577547.75 | 622994.62 | 1208548.12 | 1325949 | 681229.31 | 1685408.62 | 1634376.25 | 1789278.88 | 9 | 5 | 12 | 6 | 12 | 6 | 1.190563046 | 0.991650836 | 1.493683741 | 1.244127923 |
| O55101 | Syngr2 | Synaptogyrin-2 | 51.91 | 4.46 | 24778 | 1 | 1709030.25 | 1906913 | 1621035.25 | 1618487.88 | 1532192.12 | 1364996.5 | 1477045.25 | 1885007.5 | 1630333.62 | 2016633.25 | 4265865 | 3648800.25 | 4277986.5 | 2728492.75 | 5369453 | 1276229.5 | 2112639.25 | 1555302.62 | 2919791.25 | 2134649 | 0 | 9 | 0 | 8 | 0 | 9 | 2.419101529 | 2.423042565 | 1.192062316 | 1.194004344 |
| Q62910 | Synj1 | Synaptojanin-1 | 110.2 | 1.78 | 172881 | 2 | 1474355.75 | 1432817.88 | 1119810.25 | 1273536.75 | 1869187.5 | 1371752.75 | 1475986.62 | 1224387.62 | 1578217.5 | 1238388.5 | 492486.06 | 326820.38 | 690777.62 | 2 | 318969.66 | 1334767.75 | 885104.69 | 1291559.38 | 1267210.25 | 1246238 | 0 | 6 | 0 | 8 | 0 | 8 | 0.255108812 | 0.265514097 | 0.840324315 | 0.874599158 |
| Q9D6K5 | Synj2bp | Synaptojanin-2-binding protein | 71.4 | 8.28 | 15815 | 1 | 618740.88 | 751674.5 | 589130.69 | 900847.94 | 692452.88 | 969005.19 | 535767.56 | 752219.94 | 642525.25 | 1111032 | 725292.31 | 898978 | 1159158.62 | 1036526.06 | 762794.44 | 640371.94 | 945141.25 | 764107.62 | 867594.44 | 1080466.88 | 4 | 5 | 8 | 11 | 8 | 11 | 1.289880924 | 1.142673573 | 1.209644621 | 1.071594219 |
| O09117 | Sypl1 | Synaptophysin-like protein 1 | 125.51 | 4.21 | 28899 | 2 | 8253654 | 7307984 | 5692099 | 8393402 | 5300985.5 | 9185214 | 7912980 | 6284083.5 | 5742225.5 | 7183404 | 8138993 | 7628890.5 | 8592746 | 9858888 | 9655848 | 9111536 | 11133977 | 7520642 | 8201029 | 7637180 | 4 | 6 | 4 | 6 | 4 | 6 | 1.255442635 | 1.208424531 | 1.247688241 | 1.200960551 |
| B2RYT9 | Taco1 | Translational activator of cytochrome c oxidase 1 | 118.65 | 8.81 | 32754 | 2 | 1950473.38 | 2449153.5 | 1839345.75 | 2496905.5 | 2523794.75 | 456197.75 | 1626878 | 2 | 1769701.62 | 2977902 | 651729.88 | 1545331.25 | 2573037.25 | 2029557 | 2175947.5 | 1287536.5 | 1700813.62 | 2393455.25 | 1985575.88 | 2129455.25 | 9 | 6 | 9 | 8 | 9 | 8 | 0.797145972 | 1.314012819 | 0.843438047 | 1.390320524 |
| Q8C092 | Taf5 | Transcription initiation factor TFIID subunit 5 | 67.13 | 1.37 | 87045 | 1 | 2 | 2 | 182366.03 | 2 | 151470.38 | 2 | 118859.85 | 152093.77 | 167532.38 | 173081.61 | 2 | 136821.81 | 86151.5 | 2 | 2 | 145522.56 | 2 | 2 | 2 | 2 | 10.5 | 9 | 4.5 | 4 | 10.5 | 9 | 0.667917866 | 0.364601685 | 0.435925921 | 0.23796238 |
| Q9WVA4 | Tagln2 | Transgelin-2 | 1591.4 | 83.42 | 22395 | 23 | 271746498 | 303059138 | 336224290 | 351228450 | 324337634 | 310738722 | 314420674 | 319820578 | 310664898 | 313230402 | 136568002 | 155350546 | 127087682 | 130126770 | 137312098 | 367751906 | 351242018 | 312328130 | 334709794 | 326324866 | 0 | 7 | 0 | 3 | 0 | 7 | 0.432652732 | 0.437539624 | 1.066658874 | 1.078706983 |
| Q93092 | Taldo1 | Transaldolase | 1629.57 | 67.95 | 37387 | 27 | 111736498 | 103671690 | 127061218 | 156285314 | 110375042 | 125047282 | 90031530 | 105123410 | 106103394 | 143167522 | 76683154 | 60308898 | 84690282 | 115144058 | 75213858 | 116108610 | 181350546 | 163401074 | 179843282 | 150744578 | 3 | 3 | 3 | 2 | 3 | 3 | 0.676440843 | 0.723546419 | 1.299309506 | 1.389790031 |
| Q3TUH1 | Tamm41 | Phosphatidate cytidylyltransferase, mitochondrial | 45.47 | 3.26 | 37828 | 1 | 2 | 664719.44 | 580041.25 | 735298 | 421582.97 | 2 | 2 | 748295.62 | 600101.44 | 2 | 500943.94 | 251571.97 | 852576.62 | 860893.75 | 876314.44 | 406804.38 | 784567.88 | 673177.69 | 873611.38 | 810485.38 | 7 | 5 | 4 | 3 | 7 | 5 | 1.391672202 | 2.478710424 | 1.477590855 | 2.631740327 |
| Q0VGY8 | Tanc1 | Protein TANC1 | 324.2 | 4.47 | 200805 | 6 | 3377116.5 | 3186368 | 4144158.5 | 3501937 | 3687900.5 | 4097850.5 | 3693836.25 | 3221105.5 | 4101495.5 | 3537083.5 | 1708581.25 | 1498440 | 3373667.75 | 2730449.75 | 2044732.88 | 3536040.75 | 3086901.5 | 4460313.5 | 3702257 | 3301847.25 | 1 | 12 | 1 | 10 | 1 | 12 | 0.634495544 | 0.608849155 | 1.010609287 | 0.969760333 |
| Q53UA7 | Taok3 | Serine/threonine-protein kinase TAO3 | 71.53 | 1.45 | 105474 | 1 | 347816.47 | 464784.03 | 627049.62 | 2 | 287511.34 | 468833.19 | 373780.5 | 601947.69 | 679055.06 | 2 | 504168.84 | 2 | 2 | 452750.19 | 2 | 359743 | 2 | 2 | 2 | 2 | 8.5 | 5 | 6.5 | 3 | 8.5 | 5 | 0.554044277 | 0.450610624 | 0.208290071 | 0.169404726 |
| P21958 | Tap1 | Antigen peptide transporter 1 | 908.43 | 26.52 | 78864 | 16 | 14532402 | 15249153 | 13516872 | 15937900 | 14591451 | 19014616 | 13841585 | 14275234 | 16447614 | 15930579 | 18227734 | 15151523 | 25446886 | 20147402 | 22182980 | 11036463 | 14783282 | 15032196 | 15948184 | 13703052 | 2 | 12 | 4 | 7 | 4 | 12 | 1.370168895 | 1.272255041 | 0.954968156 | 0.886725027 |
| P36371 | Tap2 | Antigen peptide transporter 2 | 826.87 | 21.51 | 77445 | 13 | 18989506 | 18299886 | 10651545 | 22540646 | 19470248 | 19264330 | 20304520 | 20842396 | 21134410 | 20363852 | 19160328 | 17158938 | 24486118 | 19022970 | 25137522 | 16726754 | 11421113 | 18291356 | 23330020 | 22326858 | 8 | 12 | 10 | 10 | 10 | 12 | 1.166912055 | 1.029990999 | 1.023837981 | 0.903704697 |
| Q9R233 | Tapbp | Tapasin | 388.41 | 19.78 | 49736 | 8 | 9727317 | 8397552 | 8588021 | 11483399 | 9566606 | 12305502 | 3546520 | 11835830 | 10184644 | 15558076 | 19232952 | 15875474 | 25049460 | 23214156 | 27107662 | 7931140 | 7166080 | 10750760 | 12770198 | 11950635 | 0 | 11 | 0 | 11 | 0 | 11 | 2.313086424 | 2.067724523 | 1.058746816 | 0.946439671 |
| Q921F2 | Tardbp | TAR DNA-binding protein 43 | 783.04 | 34.06 | 44548 | 11 | 12123194 | 14165007 | 15718791 | 17689606 | 12619078 | 13910926 | 11890182 | 14522097 | 13324718 | 15382147 | 27984660 | 22676334 | 21800302 | 18882880 | 19553854 | 13306463 | 16828668 | 15655214 | 16607968 | 15748365 | 0 | 8 | 0 | 4 | 0 | 8 | 1.533526839 | 1.60651771 | 1.080632614 | 1.132067199 |
| Q9D0R2 | Tars | Threonine--tRNA ligase, cytoplasmic | 1425.85 | 36.29 | 83356 | 25 | 43427090 | 46151354 | 44764730 | 46996278 | 47236006 | 45772214 | 46084098 | 49251734 | 47537162 | 46575022 | 59273970 | 61582594 | 56476550 | 53788234 | 55759810 | 53585938 | 53679594 | 54340378 | 48295926 | 42010406 | 0 | 5 | 0 | 6 | 0 | 6 | 1.255082941 | 1.219627912 | 1.10209663 | 1.070963335 |
| Q3UQ84 | Tars2 | Threonine--tRNA ligase, mitochondrial | 68.02 | 1.38 | 81700 | 1 | 595230.5 | 746916.81 | 700275.69 | 926635.12 | 847855.12 | 722337.25 | 616062.62 | 658601.12 | 616871.75 | 732629.56 | 458517.72 | 405745.81 | 811836.94 | 614155.25 | 710012.75 | 694987.94 | 621910.56 | 792585.56 | 237977.62 | 910796.94 | 6 | 9 | 8 | 10 | 8 | 10 | 0.78604576 | 0.896538595 | 0.853637066 | 0.973631071 |
| Q60949 | Tbc1d1 | TBC1 domain family member 1 | 45.05 | 0.8 | 142026 | 1 | 2 | 2 | 2 | 2 | 2 | 2 | 2 | 2 | 2 | 2 | 280200 | 276184.81 | 158689.73 | 2 | 2 | 2 | 2 | 2 | 2 | 2 | 5 | 12.5 | 5 | 12.5 | 5 | 12.5 | 71507.854 | 71507.854 | 1 | 1 |
| Q9CXF4 | Tbc1d15 | TBC1 domain family member 15 | 327.71 | 11.77 | 76527 | 7 | 3621638.75 | 3877034.5 | 4960185 | 4758842 | 4710793 | 2278784 | 3387401.5 | 4699296.5 | 6263753 | 3616537.5 | 2796822 | 2171313.25 | 4130744.5 | 2556932 | 3459173.75 | 4486463 | 3950920.5 | 3797552 | 3483994.5 | 3581666.5 | 2 | 5 | 7 | 12 | 7 | 12 | 0.689285184 | 0.746574896 | 0.880160633 | 0.953314896 |
| Q8BYH7 | Tbc1d17 | TBC1 domain family member 17 | 70.46 | 1.55 | 72860 | 1 | 920407.62 | 751114.19 | 604610.44 | 683799.94 | 976370 | 529437 | 1036321.56 | 2 | 885718.44 | 516639.38 | 467216.31 | 638067.56 | 603212 | 409441.44 | 342135.12 | 740228.44 | 976877.88 | 665949.69 | 672962.19 | 490113.47 | 1 | 9 | 9 | 11 | 9 | 11 | 0.624970419 | 0.828832316 | 0.900878921 | 1.194740646 |
| Q80XQ2 | Tbc1d5 | TBC1 domain family member 5 | 46.73 | 1.84 | 91837 | 1 | 662448.62 | 884389.56 | 725738.19 | 954803.56 | 2 | 869047.88 | 2 | 2 | 2 | 951087.69 | 596726.19 | 538006.94 | 2 | 668601.12 | 2 | 863705.56 | 830058.94 | 983363.94 | 927173.25 | 831322.75 | 5 | 7 | 12 | 7 | 12 | 7 | 0.558761959 | 0.990768125 | 1.374372335 | 2.436966725 |
| Q5SVR0 | Tbc1d9b | TBC1 domain family member 9B | 126.47 | 1.66 | 141779 | 2 | 2 | 262940.12 | 419029.72 | 666304.62 | 621123.38 | 602292.69 | 481103.75 | 563391.25 | 592592.62 | 873939.12 | 294526 | 381395.12 | 328630.16 | 393490.84 | 175761.81 | 394673.88 | 1074196.62 | 1178879.25 | 336850.22 | 467598.16 | 9 | 8 | 0 | 10 | 9 | 10 | 0.799128698 | 0.505506732 | 1.752918864 | 1.108848034 |
| P48428 | Tbca | Tubulin-specific chaperone A | 212.54 | 33.33 | 12758 | 4 | 4672721 | 4814423 | 5042815.5 | 5478890 | 4346868 | 4928993.5 | 4850359 | 5189918 | 4862399 | 5944287 | 2356882 | 2567843.5 | 433375.59 | 1666626.38 | 1630974.62 | 4685166.5 | 5884220.5 | 4447779 | 6212237 | 4325394.5 | 0 | 12 | 0 | 9 | 0 | 12 | 0.355386865 | 0.335805272 | 1.049231972 | 0.99141995 |
| Q9D1E6 | Tbcb | Tubulin-folding cofactor B | 49.61 | 8.2 | 27386 | 1 | 910396.38 | 1148104.12 | 1020927.5 | 969228.06 | 1014498.38 | 678925.31 | 1023519.62 | 1276668.12 | 1484437.25 | 1096883.25 | 2 | 2 | 1055166.25 | 225330.06 | 654739.62 | 1171033.12 | 610607.62 | 873739.69 | 1101962.75 | 1095556.5 | 4 | 12 | 2 | 9 | 4 | 12 | 0.382220205 | 0.348037597 | 0.958473564 | 0.872755629 |
| Q8VCN9 | Tbcc | Tubulin-specific chaperone C | 161.34 | 9.68 | 38125 | 3 | 4096669 | 3502541.5 | 4166846 | 3510925.5 | 3653037.5 | 4089593.75 | 3425092.5 | 3843232 | 3280742 | 3015101.5 | 2015552.12 | 858024.06 | 2225289.75 | 2972205.5 | 2231574.75 | 3424835.25 | 4434461.5 | 3469486.5 | 3726348.5 | 3871972 | 0 | 11 | 0 | 8 | 0 | 11 | 0.544249106 | 0.583594949 | 0.999845972 | 1.07212865 |
| Q8BYA0 | Tbcd | Tubulin-specific chaperone D | 375.68 | 7.36 | 133321 | 7 | 61629518 | 4066142.5 | 5397410 | 4726422 | 4857177 | 3097525.75 | 4766743.5 | 7214080 | 6295231 | 4497703.5 | 4720924.5 | 4703834 | 8048474 | 4176551 | 8764482 | 4757922.5 | 3329569.5 | 5267652.5 | 5531536 | 4757436 | 11 | 11 | 10 | 11 | 11 | 11 | 0.376989602 | 1.175599394 | 0.293072541 | 0.913913539 |
| Q8BHJ5 | Tbl1xr1 | F-box-like/WD repeat-containing protein TBL1XR1 | 46.76 | 1.36 | 55661 | 1 | 1092834.88 | 963483 | 1517772.25 | 1177737.5 | 1094568 | 873923.88 | 1413439.38 | 1189776.62 | 1400101.12 | 1078490.25 | 2 | 450232.88 | 546666.5 | 632096.19 | 673092.38 | 1296168.88 | 781580.19 | 1461068.5 | 974176.5 | 1370266.5 | 0 | 12 | 0 | 12 | 0 | 12 | 0.393762259 | 0.386533551 | 1.006305584 | 0.987831775 |
| Q8C4J7 | Tbl3 | Transducin beta-like protein 3 | 364.77 | 10.11 | 88266 | 6 | 3903115.5 | 3454141.25 | 3559560.5 | 2739951.5 | 3564379.25 | 2907548.75 | 3330561.25 | 2836116.5 | 2723130 | 2928765.25 | 4870644.5 | 3952600.5 | 4350620 | 3085734 | 4627522 | 5275702 | 3527989.5 | 3012992.5 | 3350734.75 | 3027150.5 | 4 | 10 | 1 | 2 | 4 | 10 | 1.212876226 | 1.418372152 | 1.05652476 | 1.235530275 |
| P29037 | Tbp | TATA-box-binding protein | 118.96 | 8.54 | 34709 | 2 | 2 | 507002.69 | 504958.28 | 264374.59 | 82279.74 | 243533.53 | 211302.11 | 278813.28 | 205131.84 | 2 | 324130.03 | 150660.33 | 371987.91 | 583699.56 | 475218.31 | 136121.2 | 301136.19 | 128605.52 | 160056.5 | 168593.44 | 9 | 11 | 4 | 9 | 9 | 11 | 1.402673247 | 2.029964994 | 0.658399426 | 0.952843286 |
| P62340 | Tbpl1 | TATA box-binding protein-like protein 1 | 120.01 | 15.05 | 20887 | 2 | 2 | 465718.5 | 595138.81 | 629357.44 | 498311.41 | 402033.5 | 558854.94 | 393450.94 | 358601.94 | 721257.25 | 303562.75 | 2 | 359331.84 | 324220.94 | 2 | 386149.06 | 277661.75 | 342922.53 | 107858.62 | 430780.34 | 4 | 5 | 1 | 4 | 4 | 5 | 0.451042645 | 0.405521366 | 0.706124019 | 0.634858766 |
| P10711 | Tcea1 | Transcription elongation factor A protein 1 | 759.49 | 43.52 | 33880 | 11 | 778876.56 | 939366.56 | 772368.25 | 953662.94 | 895706.06 | 966225.19 | 923676.44 | 1019491.75 | 1121519.62 | 1083839.75 | 1248137.62 | 1323004.38 | 984665.94 | 1226688.75 | 890175.38 | 563141.75 | 1259822.62 | 1110589.5 | 1060024.62 | 1260701.88 | 3 | 5 | 8 | 8 | 8 | 8 | 1.307073209 | 1.109080409 | 1.210669156 | 1.027279446 |
| Q8CGF7 | Tcerg1 | Transcription elongation regulator 1 | 73.59 | 1.36 | 123788 | 2 | 1706855.38 | 2215348.5 | 2009227 | 2289965.5 | 1518857.75 | 2108764.75 | 1641574.5 | 1549209.62 | 1606553.88 | 1779895.62 | 1641183.88 | 1150635.62 | 1475551.88 | 1693170.12 | 1205931.38 | 2353019.5 | 2050613.12 | 1867384.62 | 1771781.38 | 2081247.12 | 2 | 10 | 5 | 5 | 5 | 10 | 0.735758306 | 0.825060353 | 1.039402628 | 1.165559249 |
| O08784 | Tcof1 | Treacle protein | 1147.84 | 20.38 | 135001 | 20 | 20892942 | 23440140 | 18648366 | 25059364 | 15162637 | 28178136 | 19686218 | 17102636 | 17530600 | 18334964 | 25584686 | 23910906 | 18504184 | 25166870 | 20723308 | 22376166 | 26680658 | 18309836 | 23858510 | 23502550 | 8 | 8 | 6 | 7 | 8 | 8 | 1.103547944 | 1.129495877 | 1.111665561 | 1.137804364 |
| P11983 | Tcp1 | T-complex protein 1 subunit alpha | 2515.75 | 76.08 | 60449 | 34 | 90973114 | 94899386 | 95042610 | 88726442 | 82915250 | 95490058 | 89679170 | 96641698 | 94252050 | 92220618 | 151331442 | 134397826 | 143738098 | 127757386 | 122795282 | 89502690 | 92142834 | 110300778 | 102088530 | 99428210 | 0 | 5 | 0 | 9 | 0 | 9 | 1.502618082 | 1.452154299 | 1.090389184 | 1.053769657 |
| Q80VL1 | Tdrkh | Tudor and KH domain-containing protein | 41.01 | 2.32 | 62134 | 1 | 2 | 2 | 2 | 2 | 2 | 2 | 2 | 2 | 2 | 2 | 181581.39 | 124069.71 | 160362.06 | 2 | 219937.55 | 2 | 2 | 2 | 2 | 2 | 2.5 | 12.5 | 2.5 | 12.5 | 2.5 | 12.5 | 68595.271 | 68595.271 | 1 | 1 |
| Q64232 | Tecr | Very-long-chain enoyl-CoA reductase | 373.7 | 20.45 | 36123 | 7 | 3047384.75 | 5641587.5 | 8403883 | 6865663 | 6606188 | 3098219 | 5448722 | 7183319 | 10625561 | 5413980.5 | 7728804 | 10586639 | 15910204 | 11813662 | 16889300 | 2921052.5 | 2 | 4221806.5 | 7987675 | 7507977.5 | 1 | 9 | 2 | 9 | 2 | 9 | 2.058865166 | 1.980768089 | 0.74067499 | 0.712579633 |
| Q5SSK3 | Tefm | Transcription elongation factor, mitochondrial | 46.47 | 2.75 | 41874 | 1 | 2 | 296534.62 | 421154.56 | 361732.56 | 389067.38 | 297153.53 | 379115.56 | 552704.88 | 378574.41 | 470914.88 | 2 | 477225.5 | 625790.75 | 744566.5 | 443323.97 | 2 | 236235.03 | 382282.59 | 345312.44 | 300554.38 | 4.5 | 8.5 | 8 | 5 | 8 | 8.5 | 1.56004261 | 1.102212757 | 0.861010613 | 0.608327539 |
| Q5EAN7 | Terf2ip | Telomeric repeat-binding factor 2-interacting protein 1 | 43.87 | 3.56 | 43418 | 1 | 2 | 2 | 2 | 2 | 129402.89 | 2 | 1214735.38 | 81788.77 | 150565.39 | 2 | 2 | 2 | 2 | 2 | 174048.55 | 229536.45 | 2 | 2 | 2 | 2 | 12 | 12 | 8 | 8 | 12 | 12 | 1.344991523 | 0.120280096 | 1.773764557 | 0.158624473 |
| P47226 | Tes | Testin | 453.74 | 18.68 | 47983 | 9 | 10818867 | 12386318 | 12795695 | 10784258 | 11755765 | 10686125 | 13779215 | 15689953 | 13232390 | 14455313 | 8433696 | 7313029 | 8954180 | 7460007 | 10542883 | 14978544 | 7800558.5 | 15889664 | 16578833 | 16903282 | 0 | 5 | 0 | 6 | 0 | 6 | 0.729469359 | 0.629450312 | 1.232486651 | 1.063497866 |
| Q3URQ0 | Tex10 | Testis-expressed protein 10 | 139.3 | 3.66 | 105209 | 3 | 553290 | 448033.88 | 437391.16 | 473350.31 | 133525.14 | 460767.16 | 556541.62 | 519022.81 | 725264.38 | 549603.19 | 1182027.62 | 1790131 | 1743041.25 | 859426.62 | 1452986 | 493846.81 | 380683.28 | 742256 | 98833.7 | 504249.19 | 0 | 11 | 0 | 7 | 0 | 11 | 3.435493333 | 2.499862902 | 1.085197155 | 0.789651979 |
| Q8JZM0 | Tfb1m | Dimethyladenosine transferase 1, mitochondrial | 40.62 | 2.61 | 38962 | 1 | 2 | 475636.06 | 468599.41 | 497829.62 | 459842.59 | 618886.38 | 445425.88 | 487584.75 | 520100.84 | 452144.81 | 307868.06 | 528936.56 | 631013.38 | 552076.56 | 782017.88 | 371310.47 | 584009.12 | 367502.12 | 462756.12 | 499714.41 | 4 | 11 | 7 | 9 | 7 | 11 | 1.473210042 | 1.110045198 | 1.20157769 | 0.905373645 |
| Q62351 | Tfrc | Transferrin receptor protein 1 | 1208.87 | 30.14 | 85731 | 21 | 18225534 | 17123336 | 17031192 | 11489983 | 15566815 | 14231232 | 16261950 | 15696246 | 17438198 | 16183347 | 18365210 | 19177910 | 17272828 | 14655296 | 21044256 | 4678022 | 2313503 | 6171230 | 4403883 | 4090596.25 | 5 | 0 | 5 | 0 | 5 | 0 | 1.139464727 | 1.1341235 | 0.272634571 | 0.2713566 |
| Q8R3N6 | Thoc1 | THO complex subunit 1 | 89.4 | 2.44 | 75436 | 2 | 2553907.5 | 1953997.75 | 2132083.5 | 1702342 | 2727929 | 2305959 | 2362493.5 | 1974791.75 | 2657728.5 | 1651528 | 1727699.5 | 1827820 | 1385355.25 | 1769682.25 | 1443713.75 | 1567458.38 | 2136725 | 1680335.38 | 2111774.75 | 1369010.5 | 3 | 5 | 3 | 5 | 3 | 5 | 0.73659254 | 0.744512229 | 0.80082168 | 0.809431947 |
| B1AZI6 | Thoc2 | THO complex subunit 2 | 289.03 | 3.26 | 182773 | 6 | 2251055 | 2548139 | 2297063 | 1932945.5 | 2152337.5 | 1881885.12 | 2412699.75 | 1948901.5 | 2271278 | 2090517.75 | 1971809 | 2471329 | 1954421.5 | 1750097.25 | 2182414 | 2267132.75 | 1673024.25 | 1491940 | 1169086.25 | 1684287.25 | 8 | 3 | 12 | 3 | 12 | 3 | 0.923850449 | 0.974049595 | 0.740995471 | 0.781258849 |
| Q68FX7 | Thoc5 | THO complex subunit 5 homolog | 109.64 | 3.81 | 78660 | 2 | 901123.69 | 1263710.88 | 1039118.12 | 486357.41 | 1409757.5 | 2 | 1358638.12 | 1310993.62 | 1349717.12 | 537518.88 | 2 | 1087244.25 | 471011.12 | 433490.06 | 593766.75 | 1180150.25 | 174822.8 | 2 | 291980.47 | 528622.81 | 4 | 4 | 6.5 | 5.5 | 6.5 | 5.5 | 0.506956845 | 0.567388213 | 0.426578332 | 0.477428247 |
| Q5U4D9 | Thoc6 | THO complex subunit 6 homolog | 101.61 | 7.04 | 37315 | 2 | 1239103.38 | 1310266.12 | 1318260.12 | 1232637.88 | 831799.31 | 1208322.12 | 531119.44 | 2 | 1023886.62 | 1674085.25 | 2215526 | 1630476.62 | 1859927 | 557558.62 | 694533.69 | 2 | 2 | 1607290.88 | 1841821.12 | 1594776.38 | 10 | 10 | 7 | 11 | 10 | 11 | 1.172950702 | 1.568034826 | 0.85027572 | 1.136673467 |
| Q8C1A5 | Thop1 | Thimet oligopeptidase | 308.61 | 9.17 | 78026 | 6 | 5584359 | 5270105 | 3934341 | 4058996.5 | 5734575 | 3956735.5 | 5067787 | 6504592 | 5260157.5 | 3445239.25 | 1828827 | 3815298 | 2370955 | 2625764.25 | 2223260 | 5681658 | 5168942.5 | 3650092.5 | 5101303 | 4008322 | 0 | 9 | 1 | 12 | 1 | 12 | 0.523305965 | 0.530817565 | 0.960457098 | 0.974243621 |
| Q569Z6 | Thrap3 | Thyroid hormone receptor-associated protein 3 | 707.63 | 14.62 | 108178 | 13 | 14776905 | 15780427 | 13603096 | 14775476 | 12441717 | 21131974 | 14137676 | 43437662 | 14761771 | 14634802 | 15559783 | 15369651 | 13504895 | 12643530 | 14779977 | 15276815 | 16716153 | 17881606 | 17623472 | 17090674 | 11 | 1 | 9 | 10 | 11 | 10 | 1.006727809 | 0.664710949 | 1.18508741 | 0.782476226 |
| Q99J36 | Thumpd1 | THUMP domain-containing protein 1 | 212.05 | 22.29 | 38885 | 5 | 821305.5 | 770122.38 | 1082200.88 | 1327887 | 1455501.75 | 647640.19 | 808882.5 | 730320.88 | 1066600.5 | 1022416.38 | 305965.19 | 870748.12 | 900062.25 | 476438.25 | 3683798.25 | 1036965.38 | 1014947.31 | 1025329.12 | 1357523.75 | 1110220.25 | 9 | 12 | 11 | 4 | 11 | 12 | 1.142934222 | 1.458656599 | 1.016120216 | 1.296811689 |
| P97770 | Thumpd3 | THUMP domain-containing protein 3 | 122.9 | 4.75 | 56431 | 2 | 1535571.25 | 1835977.75 | 1718866.75 | 1614412 | 1678549.25 | 2511968.25 | 1726190.88 | 1716928.12 | 1652624 | 1644381.88 | 1496671.62 | 1687695.88 | 1455282.25 | 1383347.38 | 1923595 | 2153200.25 | 2401306.75 | 1720427.75 | 2028630.5 | 2250965.25 | 8 | 1 | 6 | 6 | 8 | 6 | 0.947898696 | 0.858896686 | 1.258983164 | 1.140772185 |
| P70318 | Tial1 | Nucleolysin TIAR | 178.83 | 8.16 | 43389 | 3 | 1998053.25 | 2353421 | 2570837 | 2475182.5 | 3069492.25 | 1871950 | 2206768.5 | 1030495.88 | 2438760.75 | 2069003.75 | 1632870.5 | 4534189 | 990666.69 | 2433861.5 | 1722032.88 | 2792902 | 3452093.75 | 1927175.75 | 2947032.75 | 2263294.5 | 7 | 11 | 11 | 4 | 11 | 11 | 0.907486426 | 1.176421484 | 1.073434971 | 1.391549146 |
| Q8BZA9 | Tigar | Fructose-2,6-bisphosphatase TIGAR | 188.63 | 19.33 | 29191 | 3 | 2969677 | 2023488.5 | 1571850 | 1225828.25 | 2884427 | 2818207 | 2993921.5 | 3778726.5 | 3617442 | 2470240 | 3544495 | 2909391 | 2375111 | 2446639.5 | 1959277 | 2564986.5 | 3587710.5 | 3984530.5 | 3609015.75 | 3516649.5 | 8 | 2 | 5 | 10 | 8 | 10 | 1.239773099 | 0.844142122 | 1.617091796 | 1.101052525 |
| P62075 | Timm13 | Mitochondrial import inner membrane translocase subunit Tim13 | 91.99 | 14.74 | 10458 | 1 | 825765.62 | 931080.81 | 760500.81 | 718933.75 | 599405.88 | 1357459.12 | 689098.38 | 2 | 528331.56 | 899521.5 | 632144.56 | 1021881.5 | 904212.31 | 770224.25 | 669200 | 1499492.38 | 1122269.62 | 1370500 | 1617780.88 | 1237384.38 | 11 | 0 | 10 | 2 | 11 | 2 | 1.042228617 | 1.150601016 | 1.785189326 | 1.970815826 |
| Q9Z0V8 | Timm17a | Mitochondrial import inner membrane translocase subunit Tim17-A | 238.9 | 49.12 | 18112 | 4 | 1558100 | 1126865.38 | 1889037 | 1088783.12 | 1492365.5 | 2006193.38 | 1938143.38 | 1144090.75 | 1767242.25 | 742960.25 | 1633407.12 | 2409109 | 2296100.5 | 1531515.5 | 1971008.5 | 1145675.38 | 2647929.25 | 1839519.12 | 942907.56 | 1354308.5 | 3 | 12 | 7 | 12 | 7 | 12 | 1.375392444 | 1.295120384 | 1.108339965 | 1.043653895 |
| Q9CQ85 | Timm22 | Mitochondrial import inner membrane translocase subunit Tim22 | 58.2 | 7.73 | 20114 | 1 | 2 | 507968.41 | 2 | 478942.12 | 586042.38 | 2 | 2 | 2 | 331881.5 | 537399.38 | 541699.44 | 2 | 714894.12 | 783891.56 | 757568 | 614105.75 | 477746.44 | 794924.25 | 322359.94 | 655118.62 | 5 | 6 | 3.5 | 3 | 5 | 6 | 1.778850458 | 3.218793685 | 1.820936722 | 3.294947923 |
| Q9WTQ8 | Timm23 | Mitochondrial import inner membrane translocase subunit Tim23 | 213.54 | 22.49 | 21978 | 3 | 1847632.75 | 1751525.25 | 2307177.75 | 2294477 | 2387046.25 | 2239514.5 | 1505880.12 | 2277688 | 2088310.5 | 2608296.75 | 2984624.5 | 2435659.5 | 2740435.75 | 3736102.5 | 3233737.75 | 1600146 | 2482868 | 2842089.75 | 2431995.5 | 2657136.5 | 0 | 5 | 1 | 6 | 1 | 6 | 1.42904812 | 1.41147367 | 1.134718147 | 1.120763371 |
| O35857 | Timm44 | Mitochondrial import inner membrane translocase subunit TIM44 | 775.68 | 25.44 | 51091 | 12 | 6439846 | 6466518.5 | 6092465 | 6893164 | 8389081 | 6047664.5 | 7590714.5 | 15133596 | 8267661.5 | 12554782 | 6518055 | 13095654 | 9183214 | 7621516 | 10394820 | 8384739.5 | 7452130.5 | 6703118 | 6269101 | 6132744.5 | 3 | 12 | 12 | 7 | 12 | 12 | 1.36557152 | 0.943921925 | 1.019274746 | 0.704551733 |
| Q9D880 | Timm50 | Mitochondrial import inner membrane translocase subunit TIM50 | 492.39 | 25.78 | 39776 | 8 | 22266716 | 22571318 | 23017344 | 17028398 | 26085900 | 21977838 | 25395480 | 20585894 | 22458416 | 15344472 | 13055034 | 19934690 | 11990790 | 12937207 | 14904554 | 24423870 | 15623731 | 13176702 | 19356810 | 16248049 | 1 | 5 | 1 | 7 | 1 | 7 | 0.656235808 | 0.68854793 | 0.800481404 | 0.839895974 |
| A2VCX1 | Tiprl | TIP41-like protein | 79.62 | 6.64 | 31225 | 2 | 1036806.94 | 665196.25 | 718172.25 | 881903.56 | 848840.94 | 1044343.38 | 695369.25 | 635424.75 | 892883.12 | 808007.81 | 1008940 | 483195.25 | 742234.62 | 710417.94 | 164565.64 | 111663.29 | 1127149.5 | 1235256.25 | 1130068.75 | 802371.88 | 7 | 8 | 8 | 8 | 8 | 8 | 0.749075746 | 0.762839022 | 1.061574237 | 1.08107926 |
| Q9Z0U1 | Tjp2 | Tight junction protein ZO-2 | 794.43 | 15 | 131280 | 16 | 3978762.75 | 5022813.5 | 11241929 | 10674462 | 9426633 | 5282887 | 7259928.5 | 15312769 | 12609734 | 15641169 | 9899733 | 9663896 | 12702253 | 12705071 | 12315847 | 3785616.5 | 3149424.5 | 13034722 | 8825821 | 11426192 | 4 | 12 | 12 | 7 | 12 | 12 | 1.419937232 | 1.021037006 | 0.996955621 | 0.716882803 |
| P04184 | Tk1 | Thymidine kinase, cytosolic | 164.13 | 13.3 | 25776 | 3 | 2892030.75 | 1987739.88 | 3241999.75 | 3374822.75 | 2555100.75 | 2446995.75 | 2102579.25 | 1693552 | 1756944.38 | 1738435.88 | 722069.44 | 756153.38 | 563158.69 | 696827.5 | 414510.78 | 2882336.25 | 2294389.25 | 1846818 | 1730751.5 | 1416123.62 | 0 | 3 | 0 | 12 | 0 | 12 | 0.224365818 | 0.323737479 | 0.723785951 | 1.044350879 |
| P40142 | Tkt | Transketolase | 2049.05 | 56.18 | 67630 | 30 | 146613682 | 164280802 | 205331442 | 157071186 | 175797906 | 183975298 | 158662450 | 185782114 | 206493330 | 157999794 | 241136482 | 182809506 | 173872626 | 176096690 | 137225074 | 190512146 | 177544050 | 224882418 | 135126050 | 167885122 | 9 | 9 | 11 | 12 | 11 | 12 | 1.07307234 | 1.020413402 | 1.055182008 | 1.003401003 |
| Q8K0P3 | Tldc1 | TLD domain-containing protein 1 | 41.38 | 2.64 | 50845 | 1 | 278263.5 | 240734.64 | 247524.88 | 198591.5 | 232911.95 | 2 | 239026.94 | 283527.78 | 290068.44 | 336447.59 | 120986.15 | 176921.89 | 284280.88 | 130630.73 | 327491.12 | 278453.03 | 2 | 324752.69 | 336498.09 | 350597.44 | 10 | 5 | 10 | 8.5 | 10 | 8.5 | 0.868353743 | 0.905348047 | 1.077023991 | 1.122908232 |
| P26039 | Tln1 | Talin-1 | 7906.86 | 56.87 | 269821 | 114 | 364391874 | 383670498 | 427454530 | 454319330 | 403382498 | 389375746 | 427172450 | 395900578 | 391508738 | 370266338 | 272798882 | 370026722 | 298986882 | 298096002 | 280454658 | 371571490 | 389903138 | 390702082 | 588120002 | 415774850 | 1 | 12 | 0 | 11 | 1 | 12 | 0.747761726 | 0.770106767 | 1.060422831 | 1.092110989 |
| Q6R5N8 | Tlr13 | Toll-like receptor 13 | 334.8 | 7.16 | 114443 | 5 | 3616679.5 | 3277248.5 | 2957877.25 | 3072278.25 | 3419532.25 | 2984219.5 | 2732992.25 | 1702208.5 | 2593311 | 2450427.5 | 2170578.5 | 2693165.25 | 3673427.75 | 3302230 | 3336064.5 | 2332339.5 | 2442633 | 2191242 | 2234356 | 2625356.75 | 11 | 0 | 6 | 7 | 11 | 7 | 0.928525623 | 1.217625989 | 0.723580842 | 0.948870787 |
| Q9QUN7 | Tlr2 | Toll-like receptor 2 | 843 | 22.07 | 89449 | 14 | 2888974.75 | 3942519.5 | 4686806 | 4625219.5 | 3821702 | 3835734.5 | 3117337.25 | 3318860.5 | 4562523.5 | 4257348.5 | 7781609.5 | 5565905 | 5563690 | 4671440 | 5993425.5 | 2761803.25 | 4445320.5 | 3319632.5 | 1377558.75 | 4284962 | 1 | 7 | 0 | 10 | 1 | 10 | 1.481379489 | 1.549150076 | 0.810873889 | 0.847969987 |
| Q99MB1 | Tlr3 | Toll-like receptor 3 | 247.34 | 5.86 | 103671 | 4 | 299594.03 | 235861.94 | 2 | 678723.75 | 567800.19 | 284373.28 | 430598.94 | 2 | 340718.41 | 302025.5 | 394786.81 | 230575.95 | 255282.84 | 343672.53 | 305275.94 | 296062.88 | 518071.62 | 227866.45 | 264597.34 | 577606.12 | 12 | 12 | 12 | 11 | 12 | 12 | 0.858366778 | 1.126591769 | 1.057364499 | 1.387772888 |
| P58681 | Tlr7 | Toll-like receptor 7 | 656.46 | 12.48 | 121837 | 11 | 4559862 | 3675712.75 | 6031433 | 3394438.75 | 6692631 | 3174893 | 3943359.75 | 2068307.88 | 4392243 | 2795030.25 | 1411709 | 1404682 | 1941228.75 | 1770185 | 2178378.25 | 3319142 | 1487698.25 | 1816245.38 | 1915010.25 | 1446163.5 | 0 | 0 | 1 | 3 | 1 | 3 | 0.357483588 | 0.531713163 | 0.409962536 | 0.609769187 |
| P58021 | Tm9sf2 | Transmembrane 9 superfamily member 2 | 539.06 | 11.33 | 75330 | 7 | 4694399 | 2974438.25 | 7628271 | 4693756.5 | 11036714 | 3051993.5 | 7189068 | 7777712 | 9279103 | 3196636.5 | 1366338.75 | 3831423.5 | 4147797.75 | 2 | 4873423 | 5212247 | 3187994.5 | 4067095.5 | 5378844 | 3432295.25 | 5 | 9 | 6 | 9 | 6 | 9 | 0.458269242 | 0.466280114 | 0.685792353 | 0.697780491 |
| Q9ET30 | Tm9sf3 | Transmembrane 9 superfamily member 3 | 351.12 | 16.87 | 67545 | 8 | 8369321.5 | 6508217.5 | 6655307.5 | 8137158 | 8554881 | 6409926 | 8456617 | 7489752 | 9731092 | 7953941.5 | 8512144 | 11143944 | 12879708 | 10080441 | 12893246 | 8932684 | 4969304.5 | 7563403.5 | 6554831 | 6217084.5 | 1 | 8 | 1 | 7 | 1 | 8 | 1.452181799 | 1.386304728 | 0.8956811 | 0.85504924 |
| Q8BH24 | Tm9sf4 | Transmembrane 9 superfamily member 4 | 188.46 | 8.4 | 74693 | 4 | 2 | 2 | 334859.97 | 451891.38 | 452252.06 | 688459.69 | 650141.75 | 676567.81 | 974322.56 | 1097224.12 | 6443011 | 1697644.38 | 5361137 | 5718504 | 2963633.75 | 938133.12 | 2243355.5 | 1233456.62 | 2210930.5 | 1528219.38 | 0 | 0 | 0 | 2 | 0 | 2 | 17.90459843 | 5.428302458 | 6.581151214 | 1.995268391 |
| Q5I0H4 | Tmco1 | Calcium load-activated calcium channel | 230.78 | 24.47 | 21175 | 4 | 2733688 | 4988388 | 3927994.75 | 4257729 | 1717561.25 | 3494565.25 | 3253680.75 | 4113250 | 3575605.25 | 2046692.38 | 14430378 | 13558328 | 13931276 | 12779253 | 16817506 | 3812244.75 | 3258885 | 6359126.5 | 3860515.75 | 5267642.5 | 0 | 9 | 0 | 5 | 0 | 9 | 4.057604324 | 4.33860934 | 1.279883828 | 1.368520803 |
| Q3V009 | Tmed1 | Transmembrane emp24 domain-containing protein 1 | 42.27 | 3.96 | 25263 | 1 | 2 | 2 | 2 | 2 | 2 | 2 | 2 | 2 | 2 | 2 | 1025601.25 | 2 | 2 | 225029.86 | 2 | 2 | 2 | 2 | 2 | 2 | 7.5 | 12.5 | 7.5 | 12.5 | 7.5 | 12.5 | 125063.711 | 125063.711 | 1 | 1 |
| Q63584 | Tmed10 | Transmembrane emp24 domain-containing protein 10 | 753.52 | 45.21 | 24858 | 11 | 31932010 | 36519910 | 38413858 | 36951726 | 37010030 | 33726378 | 36537450 | 41492530 | 42742558 | 39646530 | 37358306 | 47292086 | 50894558 | 37753334 | 57856722 | 30874806 | 35023962 | 47675326 | 41295238 | 38373638 | 2 | 10 | 6 | 11 | 6 | 11 | 1.278317527 | 1.19062801 | 1.068658991 | 0.995351547 |
| Q63524 | Tmed2 | Transmembrane emp24 domain-containing protein 2 | 228.39 | 16.92 | 22733 | 3 | 2 | 2 | 729971.75 | 2 | 358908.47 | 2 | 2 | 301405.44 | 2 | 2 | 824468.12 | 681770.5 | 756637.31 | 2 | 1169182.5 | 512704.5 | 2 | 2 | 2 | 497762.06 | 4.5 | 12.5 | 3 | 9 | 4.5 | 12.5 | 3.151899957 | 11.38655406 | 0.927987279 | 3.352446925 |
| [illegible] | [illegible] | [illegible] | [illegible] | [illegible] | [illegible] | [illegible] | [illegible] | [illegible] | [illegible] | [illegible] | [illegible] | [illegible] | [illegible] | [illegible] | [illegible] | [illegible] | [illegible] | [illegible] | [illegible] | [illegible] | [illegible] | [illegible] | [illegible] | [illegible] | [illegible] | [illegible] | [illegible] | [illegible] | [illegible] | [illegible] | [illegible] | [illegible] | [illegible] | [illegible] | [illegible] | [illegible] |

| | | | | | | |
|---|---|---|---|---|---|---|
| Q6AY41 | Tmem30a | Cell cycle control protein 50A | 41.46 | 2.44 | 37173 | 1 |
| Q9CR67 | Tmem33 | Transmembrane protein 33 | 224.78 | 17 | 28031 | 4 |
| Q9DAV9 | Tmem38b | Trimeric intracellular cation channel type B | 89.35 | 9.59 | 32640 | 2 |
| Q9DBS1 | Tmem43 | Transmembrane protein 43 | 475.03 | 25.75 | 44783 | 8 |
| Q9JHJ0 | Tmod3 | Tropomodulin-3 | 192.69 | 14.49 | 39503 | 4 |
| Q61033 | Tmpo | Lamina-associated polypeptide 2, isoforms alpha/zeta | 737.94 | 25.97 | 75168 | 12 |
| Q61029 | Tmpo | Lamina-associated polypeptide 2, isoforms beta/delta/epsilon/gamma | 870.72 | 42.04 | 50373 | 14 |
| P63312 | Tmsb10 | Thymosin beta-10 | 67.1 | 31.82 | 5026 | 1 |
| P62329 | Tmsb4x | Thymosin beta-4 | 93.01 | 54.55 | 5053 | 2 |
| Q8VBT0 | Tmx1 | Thioredoxin-related transmembrane protein 1 | 284.98 | 17.27 | 31396 | 5 |
| Q5XIK2 | Tmx2 | Thioredoxin-related transmembrane protein 2 | 65.63 | 5.08 | 33866 | 1 |
| Q8BXZ1 | Tmx3 | Protein disulfide-isomerase TMX3 | 482.09 | 23.46 | 51848 | 8 |
| Q8C0L0 | Tmx4 | Thioredoxin-related transmembrane protein 4 | 318.99 | 20.6 | 37131 | 5 |
| P06804 | Tnf | Tumor necrosis factor | 140.04 | 13.62 | 25896 | 3 |
| Q9D8Y7 | Tnfaip8l2 | Tumor necrosis factor alpha-induced protein 8-like protein 2 | 201.03 | 26.63 | 20615 | 4 |
| Q8BFY9 | Tnpo1 | Transportin-1 | 765.23 | 16.93 | 102357 | 14 |
| Q6P2B1 | Tnpo3 | Transportin-3 | 162.76 | 4.77 | 104170 | 4 |
| Q5SSZ5 | Tns3 | Tensin-3 | 110.83 | 2.99 | 155589 | 2 |
| Q9D2E2 | Toe1 | Target of EGR1 protein 1 | 51.37 | 1.57 | 56871 | 1 |
| A2RUW1 | Tollip | Toll-interacting protein | 193.69 | 15.69 | 30315 | 4 |
| O88746 | Tom1 | Target of Myb protein 1 | 258.73 | 12.2 | 54325 | 5 |
| Q62760 | Tomm20 | Mitochondrial import receptor subunit TOM20 homolog | 84.5 | 13.79 | 16284 | 1 |
| Q9CPQ3 | Tomm22 | Mitochondrial import receptor subunit TOM22 homolog | 259.44 | 37.32 | 15537 | 4 |
| Q9CYG7 | Tomm34 | Mitochondrial import receptor subunit TOM34 | 569.13 | 43.69 | 34278 | 11 |
| Q9QYA2 | Tomm40 | Mitochondrial import receptor subunit TOM40 homolog | 615.64 | 35.73 | 37895 | 9 |
| B1AXP6 | Tomm5 | Mitochondrial import receptor subunit TOM5 homolog | 76.26 | 33.33 | 6037 | 2 |
| Q9CZW5 | Tomm70 | Mitochondrial import receptor subunit TOM70 | 637.83 | 21.6 | 67590 | 12 |
| Q04750 | Top1 | DNA topoisomerase 1 | 601.75 | 14.86 | 90876 | 12 |
| Q01320 | Top2a | DNA topoisomerase 2-alpha | 927.92 | 12.83 | 172790 | 18 |
| Q9ER39 | Tor1a | Torsin-1A | 101.79 | 6.91 | 37830 | 2 |
| Q921T2 | Tor1aip1 | Torsin-1A-interacting protein 1 | 456.38 | 17.98 | 66781 | 7 |
| Q9ER41 | Tor1b | Torsin-1B | 160.64 | 13.1 | 37818 | 3 |
| Q9ER38 | Tor3a | Torsin-3A | 617.75 | 42.34 | 43814 | 12 |
| P70399 | Tp53bp1 | TP53-binding protein 1 | 42.47 | 0.76 | 212735 | 1 |
| Q62393 | Tpd52 | Tumor protein D52 | 641.24 | 44.2 | 24313 | 9 |
| Q6PCT3 | Tpd52l2 | Tumor protein D54 | 300.59 | 20.45 | 23992 | 5 |
| P17751 | Tpi1 | Triosephosphate isomerase | 1281.95 | 73.91 | 32192 | 18 |
| P04692 | Tpm1 | Tropomyosin alpha-1 chain | 517.84 | 25.35 | 32681 | 9 |
| Q63610 | Tpm3 | Tropomyosin alpha-3 chain | 1333.24 | 55.24 | 29007 | 22 |
| P21107 | Tpm3 | Tropomyosin alpha-3 chain | 698.72 | 32.28 | 32994 | 12 |
| Q6IRU2 | Tpm4 | Tropomyosin alpha-4 chain | 1213.37 | 47.18 | 28468 | 18 |
| O89023 | Tpp1 | Tripeptidyl-peptidase 1 | 433.52 | 22.78 | 61342 | 7 |
| Q64514 | Tpp2 | Tripeptidyl-peptidase 2 | 1050.67 | 20.29 | 139879 | 20 |
| F6ZDS4 | Tpr | Nucleoprotein TPR | 1488.92 | 12.01 | 273990 | 25 |
| F1MA98 | Tpr | Nucleoprotein TPR | 1169.47 | 9.7 | 267306 | 20 |
| P63028 | Tpt1 | Translationally-controlled tumor protein | 416.63 | 45.93 | 19462 | 6 |
| A2APB8 | Tpx2 | Targeting protein for Xklp2 | 116.35 | 4.03 | 85894 | 3 |
| Q6PFR5 | Tra2a | Transformer-2 protein homolog alpha | 263.49 | 14.95 | 32316 | 5 |
| P62996 | Tra2b | Transformer-2 protein homolog beta | 429.34 | 24.31 | 33666 | 7 |
| Q5XI41 | Tram1 | Translocating chain-associated membrane protein 1 | 201.03 | 6.95 | 43030 | 3 |
| Q9CQN1 | Trap1 | Heat shock protein 75 kDa, mitochondrial | 1214.69 | 35.13 | 80209 | 19 |
| Q8K2L8 | Trappc12 | Trafficking protein particle complex subunit 12 | 39.82 | 1.13 | 87694 | 1 |
| O55013 | Trappc3 | Trafficking protein particle complex subunit 3 | 86.43 | 10.56 | 20302 | 2 |
| Q9CQA1 | Trappc5 | Trafficking protein particle complex subunit 5 | 141.64 | 14.36 | 20795 | 3 |
| Q9D289 | Trappc6b | Trafficking protein particle complex subunit 6B | 51.61 | 6.96 | 17936 | 1 |
| Q99NH8 | Trem2 | Triggering receptor expressed on myeloid cells 2 | 194.24 | 16.3 | 24527 | 3 |
| Q91XB0 | Trex1 | Three-prime repair exonuclease 1 | 221.76 | 10.19 | 33675 | 3 |
| Q8BVW3 | Trim14 | Tripartite motif-containing protein 14 | 181.65 | 12.05 | 49641 | 4 |
| Q61510 | Trim25 | E3 ubiquitin/ISG15 ligase TRIM25 | 165.01 | 6.94 | 71726 | 4 |
| Q62318 | Trim28 | Transcription intermediary factor 1-beta | 1340.44 | 32.49 | 88847 | 20 |
| Q80VI1 | Trim56 | E3 ubiquitin-protein ligase TRIM56 | 69.54 | 2.72 | 79513 | 1 |
| P97531 | Trip10 | Cdc42-interacting protein 4 | 104.88 | 7.68 | 62799 | 2 |
| F1LP64 | Trip12 | E3 ubiquitin-protein ligase TRIP12 | 550.22 | 5.78 | 223928 | 10 |
| Q9D735 | Trir | Telomerase RNA component interacting RNase | 45.97 | 6.94 | 18377 | 1 |
| Q3TX08 | Trmt1 | tRNA (guanine(26)-N(2))-dimethyltransferase | 52.8 | 3.62 | 72350 | 1 |
| A2RSY6 | Trmt1l | TRMT1-like protein | 179.67 | 5.49 | 80861 | 3 |
| Q8BNV1 | Trmt2a | tRNA (uracil-5-)-methyltransferase homolog A | 81.96 | 2.96 | 63301 | 2 |
| Q9D0C4 | Trmt5 | tRNA (guanine(37)-N1)-methyltransferase | 51.59 | 1.8 | 56794 | 1 |
| Q8CE96 | Trmt6 | tRNA (adenine(58)-N(1))-methyltransferase non-catalytic subunit TRM6 | 123.77 | 6.24 | 55518 | 2 |
| Q8K1J6 | Trnt1 | CCA tRNA nucleotidyltransferase 1, mitochondrial | 212.47 | 11.52 | 49895 | 5 |
| O08848 | Trove2 | 60 kDa SS-A/Ro ribonucleoprotein | 40.67 | 2.04 | 60124 | 1 |
| Q9WTR1 | Trpv2 | Transient receptor potential cation channel subfamily V member 2 | 453.96 | 10.71 | 85965 | 7 |
| Q9CZR8 | Tsfm | Elongation factor Ts, mitochondrial | 244.76 | 18.83 | 35334 | 3 |
| Q61187 | Tsg101 | Tumor susceptibility gene 101 protein | 190.2 | 14.07 | 44124 | 4 |
| Q62348 | Tsn | Translin | 525.42 | 33.77 | 26201 | 8 |
| Q9QZE7 | Tsnax | Translin-associated protein X | 341.71 | 27.93 | 32926 | 5 |
| Q9CQ88 | Tspan31 | Tetraspanin-31 | 76.53 | 5.71 | 22694 | 1 |
| P50637 | Tspo | Translocator protein | 278.33 | 40.24 | 18841 | 4 |
| Q5SWD9 | Tsr1 | Pre-rRNA-processing protein TSR1 homolog | 337.28 | 9.71 | 92105 | 6 |
| Q5XIB1 | Tssc4 | Protein TSSC4 | 42.85 | 3.15 | 33939 | 1 |
| P23591 | Tsta3 | GDP-L-fucose synthase | 172.47 | 11.53 | 35878 | 4 |
| Q91Z38 | Ttc1 | Tetratricopeptide repeat protein 1 | 63.63 | 6.16 | 33263 | 1 |
| Q9D6K7 | Ttc33 | Tetratricopeptide repeat protein 33 | 70.89 | 4.96 | 29371 | 1 |
| A3KMP2 | Ttc38 | Tetratricopeptide repeat protein 38 | 57.91 | 1.94 | 52224 | 1 |
| Q8BGB2 | Ttc7a | Tetratricopeptide repeat protein 7A | 61.34 | 2.33 | 96156 | 1 |
| Q3UDE2 | Ttll12 | Tubulin--tyrosine ligase-like protein 12 | 245.17 | 8.29 | 74043 | 5 |
| P68373 | Tuba1c | Tubulin alpha-1C chain | 2305.94 | 80.18 | 49909 | 32 |
| P05214 | Tuba3a | Tubulin alpha-3 chain | 1849.96 | 66.89 | 49960 | 25 |
| P68368 | Tuba4a | Tubulin alpha-4A chain | 1857.84 | 58.71 | 49924 | 25 |
| Q6AY56 | Tuba8 | Tubulin alpha-8 chain | 1236.36 | 37.64 | 50037 | 16 |
| A2AQ07 | Tubb1 | Tubulin beta-1 chain | 463.68 | 8.43 | 50441 | 8 |
| P85108 | Tubb2a | Tubulin beta-2A chain | 2109.57 | 58.2 | 49907 | 31 |
| Q3KRE8 | Tubb2b | Tubulin beta-2B chain | 2111.15 | 58.2 | 49953 | 31 |
| Q4QRB4 | Tubb3 | Tubulin beta-3 chain | 1686.73 | 38.89 | 50419 | 25 |
| Q9D6F9 | Tubb4a | Tubulin beta-4A chain | 1798.94 | 59.91 | 49586 | 27 |
| P68372 | Tubb4b | Tubulin beta-4B chain | 2291.51 | 68.09 | 49831 | 33 |
| P69897 | Tubb5 | Tubulin beta-5 chain | 2527.68 | 68.92 | 49671 | 35 |
| Q922F4 | Tubb6 | Tubulin beta-6 chain | 1617.11 | 46.31 | 50090 | 24 |
| P83887 | Tubg1 | Tubulin gamma-1 chain | 40.67 | 2 | 51101 | 1 |
| Q8BFR5 | Tufm | Elongation factor Tu, mitochondrial | 960.94 | 47.57 | 49508 | 18 |
| Q91YR1 | Twf1 | Twinfilin-1 | 164.4 | 11.71 | 40079 | 3 |
| Q9Z0P5 | Twf2 | Twinfilin-2 | 344.31 | 19.2 | 39471 | 5 |
| P10639 | Txn | Thioredoxin | 384.46 | 55.24 | 11675 | 6 |
| P97493 | Txn2 | Thioredoxin, mitochondrial | 64.89 | 9.04 | 18255 | 1 |
| Q498E0 | Txndc12 | Thioredoxin domain-containing protein 12 | 143.72 | 22.35 | 19019 | 3 |
| Q9CQM5 | Txndc17 | Thioredoxin domain-containing protein 17 | 303.58 | 42.28 | 14015 | 5 |
| Q91W90 | Txndc5 | Thioredoxin domain-containing protein 5 | 510.22 | 25.9 | 46415 | 8 |
| Q8K581 | Txndc9 | Thioredoxin domain-containing protein 9 | 58.03 | 3.98 | 26272 | 1 |
| Q5M7W1 | Txnip | Thioredoxin-interacting protein | 157.4 | 7.11 | 44018 | 3 |
| Q8CDN6 | Txnl1 | Thioredoxin-like protein 1 | 374.41 | 33.91 | 32237 | 7 |
| Q9JMH6 | Txnrd1 | Thioredoxin reductase 1, cytoplasmic | 1007.85 | 35.73 | 67084 | 14 |
| Q9D883 | U2af1 | Splicing factor U2AF 35 kDa subunit | 387.93 | 25.94 | 27815 | 6 |
| P26369 | U2af2 | Splicing factor U2AF 65 kDa subunit | 864.42 | 40.63 | 53517 | 14 |
| Q6NV83 | U2surp | U2 snRNP-associated SURP motif-containing protein | 153.5 | 3.3 | 118261 | 3 |
| Q91YN5 | Uap1 | UDP-N-acetylhexosamine pyrophosphorylase | 135.3 | 4.41 | 58609 | 2 |
| Q3TW96 | Uap1l1 | UDP-N-acetylhexosamine pyrophosphorylase-like protein 1 | 778.2 | 33.14 | 56614 | 15 |
| Q02053 | Uba1 | Ubiquitin-like modifier-activating enzyme 1 | 1794.04 | 31 | 117809 | 24 |
| Q9Z1F9 | Uba2 | SUMO-activating enzyme subunit 2 | 981.28 | 33.7 | 70569 | 14 |
| Q8C878 | Uba3 | NEDD8-activating enzyme E1 catalytic subunit | 334.1 | 18.18 | 51737 | 6 |
| Q8VE47 | Uba5 | Ubiquitin-like modifier-activating enzyme 5 | 130.75 | 7.94 | 44790 | 2 |
| Q8C7R4 | Uba6 | Ubiquitin-like modifier-activating enzyme 6 | 278.14 | 6.27 | 117966 | 6 |
| Q80X50 | Ubap2l | Ubiquitin-associated protein 2-like | 516.49 | 11.2 | 116799 | 9 |
| Q9D1C1 | Ube2c | Ubiquitin-conjugating enzyme E2 C | 52.57 | 9.5 | 19606 | 1 |
| P61078 | Ube2d3 | Ubiquitin-conjugating enzyme E2 D3 | 163.63 | 36.05 | 16687 | 3 |
| P62254 | Ube2g1 | Ubiquitin-conjugating enzyme E2 G1 | 51.57 | 5.88 | 19509 | 1 |
| P60605 | Ube2g2 | Ubiquitin-conjugating enzyme E2 G2 | 50.83 | 14.55 | 18566 | 1 |
| P63280 | Ube2i | SUMO-conjugating enzyme UBC9 | 157.36 | 15.82 | 18007 | 3 |
| Q9JJZ4 | Ube2j1 | Ubiquitin-conjugating enzyme E2 J1 | 147.32 | 11.01 | 34990 | 3 |
| P61087 | Ube2k | Ubiquitin-conjugating enzyme E2 K | 281.61 | 35 | 22407 | 5 |
| P68037 | Ube2l3 | Ubiquitin-conjugating enzyme E2 L3 | 340.42 | 62.34 | 17862 | 6 |
| P61082 | Ube2m | NEDD8-conjugating enzyme Ubc12 | 312.56 | 37.16 | 20900 | 7 |
| P61089 | Ube2n | Ubiquitin-conjugating enzyme E2 N | 261.68 | 43.42 | 17138 | 5 |
| Q6ZPJ3 | Ube2o | (E3-independent) E2 ubiquitin-conjugating enzyme UBE2O | 48.76 | 1.16 | 140834 | 1 |
| Q9CZY3 | Ube2v1 | Ubiquitin-conjugating enzyme E2 variant 1 | 297.36 | 29.25 | 16355 | 4 |
| Q9D2M8 | Ube2v2 | Ubiquitin-conjugating enzyme E2 variant 2 | 301.15 | 34.48 | 16367 | 5 |
| O08759 | Ube3a | Ubiquitin-protein ligase E3A | 228.14 | 7.82 | 99819 | 5 |
| Q9EPV8 | Ubl5 | Ubiquitin-like protein 5 | 46.08 | 12.33 | 8547 | 1 |
| Q8BGR9 | Ublcp1 | Ubiquitin-like domain-containing CTD phosphatase 1 | 67.24 | 6.6 | 36837 | 2 |
| Q8R317 | Ubqln1 | Ubiquilin-1 | 100.52 | 7.9 | 61976 | 3 |
| A2AN08 | Ubr4 | E3 ubiquitin-protein ligase UBR4 | 1173.26 | 5 | 572290 | 22 |
| P25976 | Ubtf | Nucleolar transcription factor 1 | 47.89 | 1.57 | 89509 | 1 |
| Q922Y1 | Ubxn1 | UBX domain-containing protein 1 | 295.92 | 17.85 | 33573 | 5 |
| Q8VCH8 | Ubxn4 | UBX domain-containing protein 4 | 229.49 | 12.45 | 56472 | 5 |
| Q9JKB1 | Uchl3 | Ubiquitin carboxyl-terminal hydrolase isozyme L3 | 202.41 | 15.65 | 26152 | 3 |
| Q9WUP7 | Uchl5 | Ubiquitin carboxyl-terminal hydrolase isozyme L5 | 652.35 | 47.11 | 37617 | 13 |
| Q91YL3 | Uckl1 | Uridine-cytidine kinase-like 1 | 45.09 | 1.46 | 60842 | 1 |
| Q3U1V6 | Uevld | Ubiquitin-conjugating enzyme E2 variant 3 | 48.93 | 4.25 | 51681 | 1 |
| Q6BBI8 | Ufc1 | Ubiquitin-fold modifier-conjugating enzyme 1 | 108.62 | 26.35 | 19492 | 3 |

| | | | | | | | | | | | | | | | | | | | | | | | | | | | | | | | | | | | | |
|---|---|---|---|---|---|---|---|---|---|---|---|---|---|---|---|---|---|---|---|---|---|---|---|---|---|---|---|---|---|---|---|---|---|---|---|---|
| P70362 | Ufd1 | Ubiquitin recognition factor in ER-associated degradation protein 1 | 120.75 | 9.12 | 34481 | 2 | 554061.62 | 1419051.75 | 1399341.5 | 1229009.75 | 951031.56 | 1019379.25 | 1432083.5 | 488804.72 | 1437425.88 | 737853.62 | 1390428.25 | 1543720.5 | 1292340.38 | 1057006.12 | 1349468 | 843395.81 | 858799.19 | 1223481 | 728118.69 | 764990.88 | 9 | 6 | 8 | 10 | 9 | 10 | 1.194591232 | 1.296628354 | 0.795819651 | 0.863795327 |
| Q8CCJ3 | Ufl1 | E3 UFM1-protein ligase 1 | 252.81 | 10.21 | 89520 | 5 | 1440844.38 | 908792.5 | 1095232.75 | 1084736.62 | 1239839.38 | 632074.5 | 1239599.75 | 1572877.38 | 1452905.38 | 982697.31 | 2308764.5 | 1685610.5 | 1631462.75 | 1862654.62 | 1564454.25 | 1896383.5 | 1271037 | 1866804.5 | 1297193.5 | 1173703.25 | 0 | 4 | 1 | 7 | 1 | 7 | 1.569118976 | 1.539576366 | 1.300839323 | 1.276347752 |
| P61961 | Ufm1 | Ubiquitin-fold modifier 1 | 53.23 | 17.65 | 9118 | 1 | 1267837.38 | 1445704.38 | 1349218.62 | 1072879.12 | 2 | 1492547.38 | 1183892.12 | 1193208.12 | 2 | 1413730.12 | 715620.94 | 1433380.38 | 786448.94 | 575942.06 | 912320.69 | 1094595.5 | 1381808.25 | 1509901.25 | 2 | 1178670.5 | 8 | 11.5 | 8 | 10.5 | 8 | 11.5 | 0.861374964 | 0.837288483 | 1.005712237 | 0.977589678 |
| Q99K23 | Ufsp2 | Ufm1-specific protease 2 | 265.14 | 11.71 | 52515 | 5 | 1458187.5 | 1733843.62 | 1584242 | 1971278.25 | 2688134 | 1601678.12 | 2737949.5 | 1696991.12 | 1684853.62 | 1122219.12 | 2734549 | 3357247 | 4470830.5 | 1742195 | 2929767.25 | 1904241.25 | 1529679 | 1960265 | 804756.12 | 1731051 | 2 | 9 | 2 | 12 | 2 | 12 | 1.614571507 | 1.72265041 | 0.840425688 | 0.896683516 |
| O70199 | Ugdh | UDP-glucose 6-dehydrogenase | 138.32 | 6.49 | 54892 | 3 | 1467086.12 | 2 | 1232817.38 | 935081 | 117390.56 | 562456.06 | 1489868.75 | 423676.22 | 715608.12 | 125971.45 | 1037990 | 1356449.25 | 489445.69 | 423844.25 | 766753.62 | 1410334 | 1496037.62 | 515546.31 | 449358.91 | 350215.94 | 12 | 10 | 9 | 11 | 12 | 11 | 1.085840454 | 1.228148853 | 1.125018278 | 1.272461257 |
| Q6P5E4 | Uggt1 | UDP-glucose:glycoprotein glucosyltransferase 1 | 2837.679 | 40.1 | 176434 | 48 | 17573274 | 20983830 | 25060222 | 22836502 | 27461082 | 17572300 | 19367838 | 37924546 | 23408338 | 22899904 | 42935366 | 37941030 | 36196442 | 39393822 | 41769666 | 19283892 | 13183063 | 22117168 | 19600098 | 17753384 | 0 | 5 | 1 | 6 | 1 | 6 | 1.740214042 | 1.6359787 | 0.807072621 | 0.758730585 |
| Q91ZJ5 | Ugp2 | UTP--glucose-1-phosphate uridylyltransferase | 373.5 | 12.8 | 56979 | 6 | 7157190 | 6947708.5 | 5712874 | 6286316 | 5207761.5 | 5538120.5 | 7186109.5 | 6064384.5 | 6635129 | 7811349 | 4308885 | 6530129 | 5282071.5 | 4911969 | 5473974.5 | 7538540 | 8574576 | 6475859.5 | 7261910.5 | 8220578 | 5 | 2 | 2 | 5 | 5 | 5 | 0.846549437 | 0.797561463 | 1.215880378 | 1.145520025 |
| Q6ZQM8 | Ugt1a7c | UDP-glucuronosyltransferase 1-7C | 993.27 | 36.16 | 59758 | 18 | 37914282 | 37325462 | 39810114 | 35764642 | 40193370 | 32468334 | 44044550 | 49294306 | 51068746 | 44349178 | 27234166 | 48273118 | 45260666 | 32349146 | 46107406 | 38553726 | 26733904 | 37813214 | 41014594 | 41310558 | 10 | 10 | 9 | 4 | 10 | 10 | 1.043017243 | 0.90055102 | 0.970776733 | 0.838177875 |
| Q09426 | Ugt8 | 2-hydroxyacylsphingosine 1-beta-galactosyltransferase | 42.84 | 1.29 | 61126 | 1 | 5231538.5 | 6088538.5 | 7125840.5 | 8207048.5 | 6179887.5 | 6730344 | 4833056.5 | 5598135 | 4795786 | 8426761 | 9113146 | 8816571 | 9385500 | 10784755 | 9414617 | 7063549 | 9171870 | 8665436 | 8791171 | 9978919 | 0 | 2 | 0 | 1 | 0 | 2 | 1.447165992 | 1.563798709 | 1.330098981 | 1.437296815 |
| Q8VDF2 | Uhrf1 | E3 ubiquitin-protein ligase UHRF1 | 619.72 | 18.93 | 88304 | 11 | 1521471.25 | 1235918.75 | 2303327 | 1830958.5 | 1954379 | 1734090.5 | 1607617.38 | 1460947 | 1917865.25 | 1733488.88 | 1450707.88 | 1197161.75 | 1037818.88 | 1137527.62 | 1202489.5 | 1240414.25 | 515538.16 | 1731079 | 1126336.25 | 1733958.38 | 1 | 5 | 0 | 5 | 1 | 5 | 0.681174373 | 0.712763095 | 0.717531871 | 0.750806633 |
| O70405 | Ulk1 | Serine/threonine-protein kinase ULK1 | 53 | 1.33 | 112463 | 2 | 13464157 | 10412046 | 10542626 | 15056821 | 9640529 | 12401917 | 8543847 | 10514360 | 8948724 | 14584795 | 15432105 | 10027609 | 11747891 | 10011760 | 12594345 | 10665855 | 14542376 | 13239438 | 12102886 | 13074338 | 12 | 9 | 9 | 7 | 12 | 9 | 1.011799325 | 1.087647712 | 1.076268698 | 1.156949959 |
| P13439 | Umps | Uridine 5'-monophosphate synthase | 432.98 | 17.05 | 52292 | 7 | 8241257 | 6503955.5 | 7150913.5 | 6587577 | 7583852 | 7474301 | 6255790 | 6658105 | 6479353.5 | 6391590.5 | 7632873.5 | 8049397.5 | 7775707 | 4925786.5 | 7470648.5 | 8762540 | 6522905.5 | 7401696 | 6187109 | 6177662 | 10 | 9 | 6 | 12 | 10 | 12 | 0.994090478 | 1.078031873 | 0.97184055 | 1.053903153 |
| Q62885 | Unc119 | Protein unc-119 homolog A | 65.42 | 5.42 | 27048 | 1 | 530441.31 | 614716.12 | 618683.81 | 708742.12 | 576161.56 | 671616.31 | 512196.72 | 508071.16 | 459355.41 | 2 | 2 | 2 | 2 | 308633.47 | 2 | 2 | 2 | 688924 | 703444.06 | 779116.38 | 0 | 12 | 3 | 9 | 3 | 12 | 0.101235583 | 0.143471319 | 0.712256518 | 1.0094117 |
| Q32PZ3 | Unc45a | Protein unc-45 homolog A | 44.59 | 1.17 | 103241 | 1 | 956794.06 | 788032.69 | 590349.62 | 881377.31 | 639722.62 | 986303.44 | 756080.19 | 590910.25 | 723141.06 | 2 | 853364.94 | 558825.62 | 836602.56 | 886549.94 | 705428.38 | 2 | 1049749.62 | 968114 | 698507.06 | 738068.38 | 12 | 11 | 10 | 10.5 | 12 | 11 | 0.995979318 | 1.256617269 | 0.895797083 | 1.130218332 |
| Q8VCW4 | Unc93b1 | Protein unc-93 homolog B1 | 352.79 | 11.04 | 66981 | 5 | 2478440 | 2992381.75 | 5455438 | 6291410 | 5758102.5 | 3708939.75 | 4685693.5 | 5436337.5 | 6490236 | 6201145.5 | 8317285.5 | 7943283.5 | 10902756 | 10745370 | 11697678 | 2913205.25 | 3965492.25 | 8254524.5 | 6852612.5 | 6433489.5 | 0 | 7 | 0 | 10 | 0 | 10 | 2.159073152 | 1.87036099 | 1.236925736 | 1.071523511 |
| Q9EPU0 | Upf1 | Regulator of nonsense transcripts 1 | 631.26 | 12.54 | 123967 | 13 | 3240071.25 | 4802541 | 4240917 | 3027405 | 3745973 | 3704145 | 2448105.25 | 3390813.5 | 3882387.25 | 3364117.5 | 14621800 | 10192607 | 10034338 | 8584375 | 10402870 | 1926275.38 | 2527963.5 | 4716080.5 | 4097874.25 | 4224170 | 0 | 10 | 0 | 9 | 0 | 10 | 2.825011913 | 3.206514211 | 0.917901494 | 1.041859035 |
| Q9CWU6 | Uqcc1 | Ubiquinol-cytochrome-c reductase complex assembly factor 1 | 186.32 | 10.51 | 34300 | 3 | 3518964.25 | 3381768.5 | 2588719.5 | 2011969.38 | 3267102 | 2456047 | 3907423.5 | 2392539.75 | 3218082.25 | 1950307.5 | 892420.44 | 2027782.25 | 1019839.75 | 1122364.25 | 1193738.5 | 3168842.5 | 1166184.75 | 1528653.5 | 982111.69 | 1262096.12 | 1 | 2 | 1 | 3 | 1 | 3 | 0.423613446 | 0.4492937 | 0.548997907 | 0.582279205 |
| Q8R1I1 | Uqcr10 | Cytochrome b-c1 complex subunit 9 | 50.45 | 10.94 | 7446 | 1 | 4273253 | 2348988.75 | 2811913.75 | 3338998 | 2833952.75 | 4186415.75 | 3206350 | 1173279 | 2382683.5 | 3078339 | 3042000 | 1581178.5 | 2709038.5 | 918278.81 | 2483155.25 | 2343565.75 | 4282472 | 3944247.25 | 3980629 | 2744929.75 | 5 | 11 | 7 | 9 | 7 | 11 | 0.687741269 | 0.765209924 | 1.108203114 | 1.233033495 |
| Q9CZ13 | Uqcrc1 | Cytochrome b-c1 complex subunit 1, mitochondrial | 808.89 | 37.5 | 52852 | 15 | 22230196 | 21688912 | 22093690 | 22553350 | 16763938 | 20451570 | 20826100 | 27409688 | 21874630 | 24165412 | 33487584 | 36614250 | 34903210 | 35761834 | 33505876 | 22533500 | 15514598 | 23883202 | 23208680 | 22255374 | 0 | 7 | 0 | 12 | 0 | 12 | 1.654539179 | 1.51901598 | 1.019607579 | 0.936091588 |
| Q9DB77 | Uqcrc2 | Cytochrome b-c1 complex subunit 2, mitochondrial | 1419.99 | 60.04 | 48235 | 22 | 36867706 | 35965642 | 41686002 | 36415238 | 41294194 | 36541434 | 37479622 | 38270654 | 36515050 | 42680914 | 57217626 | 57827102 | 63116582 | 68379978 | 71991418 | 33909246 | 32613004 | 46429490 | 39629014 | 44407670 | 0 | 12 | 0 | 11 | 0 | 12 | 1.657050014 | 1.663463237 | 1.024760298 | 1.028726392 |
| Q9CR68 | Uqcrfs1 | Cytochrome b-c1 complex subunit Rieske, mitochondrial | 567.31 | 50 | 29368 | 10 | 11001169 | 14564724 | 15278864 | 14145642 | 11592536 | 12613836 | 11332484 | 14155510 | 12390126 | 12673427 | 25456502 | 21151276 | 25195038 | 25890698 | 24571972 | 11126492 | 14104117 | 16107096 | 14976332 | 13993696 | 0 | 11 | 0 | 7 | 0 | 11 | 1.836288623 | 1.935640697 | 1.055942232 | 1.113073802 |
| P99028 | Uqcrh | Cytochrome b-c1 complex subunit 6, mitochondrial | 81.57 | 20.22 | 10435 | 1 | 444645.31 | 175214.52 | 359706.06 | 347411.34 | 645973.56 | 2 | 133614.97 | 352083.75 | 247570.38 | 313376.59 | 2 | 213769.53 | 284711.72 | 2 | 545013.62 | 613584.44 | 179960.94 | 769532.12 | 363922.56 | 763866.44 | 6 | 7 | 11 | 3 | 11 | 7 | 0.528902634 | 0.996991519 | 1.363879177 | 2.570938173 |
| Q9CQ69 | Uqcrq | Cytochrome b-c1 complex subunit 8 | 85.95 | 20.73 | 9768 | 2 | 3266451.5 | 3510785 | 3732386.5 | 3716923 | 2590808 | 3555647.25 | 3006931 | 6981310 | 3302641.25 | 5442073.5 | 5118538.5 | 5781453 | 7181998.5 | 6426698.5 | 6403099.5 | 2289120.75 | 3963625.25 | 4036654.25 | 3649315.5 | 3715539.5 | 0 | 9 | 5 | 12 | 5 | 12 | 1.838088679 | 1.386887639 | 1.049764145 | 0.79207545 |
| P70697 | Urod | Uroporphyrinogen decarboxylase | 168.13 | 11.44 | 40692 | 3 | 3959032 | 3692270 | 4103924.5 | 3451816 | 3630446 | 3542656.5 | 2937370 | 2303506.75 | 3369352.75 | 2995431 | 1532966.88 | 2875904 | 2819779 | 1580387 | 2728993.5 | 3169567.5 | 3315289 | 3659382.25 | 2642614 | 3453313.5 | 0 | 3 | 3 | 9 | 3 | 9 | 0.612503646 | 0.761670777 | 0.862119504 | 1.072077264 |
| P41542 | Uso1 | General vesicular transport factor p115 | 332.77 | 7.3 | 107162 | 7 | 4133280.5 | 6151225.5 | 4011338.5 | 4600044 | 4169569.25 | 4723475 | 5047672.5 | 5486432 | 4950103.5 | 5135330 | 7694405.5 | 7110123 | 7725306 | 7290339 | 5878338 | 6774903 | 4088854 | 5634967 | 4791233.5 | 5474425.5 | 1 | 7 | 0 | 10 | 1 | 10 | 1.547704446 | 1.40861355 | 1.160366436 | 1.056085281 |
| P52479 | Usp10 | Ubiquitin carboxyl-terminal hydrolase 10 | 61.92 | 1.52 | 87022 | 1 | 342825.56 | 337677.75 | 306090.97 | 320144.53 | 2 | 515493.31 | 376061.25 | 416176.75 | 374659.5 | 346709.97 | 382514.09 | 337871.97 | 2 | 513550.22 | 2 | 437551.12 | 533017.94 | 281261 | 230254.41 | 190495.42 | 10 | 12 | 7 | 9 | 10 | 12 | 0.94428847 | 0.608121732 | 1.279963002 | 0.824296115 |
| Q9JMA1 | Usp14 | Ubiquitin carboxyl-terminal hydrolase 14 | 998.19 | 32.45 | 56002 | 18 | 36570422 | 38229974 | 34679658 | 42290310 | 37123266 | 40248026 | 38315354 | 35018138 | 38872202 | 36984262 | 30136378 | 30304518 | 30525440 | 27126072 | 29537430 | 40099274 | 33367412 | 40063774 | 41734154 | 40891718 | 0 | 9 | 0 | 7 | 0 | 9 | 0.781550114 | 0.779304321 | 1.038448634 | 1.035464641 |
| Q8R5H1 | Usp15 | Ubiquitin carboxyl-terminal hydrolase 15 | 380.48 | 8.46 | 112325 | 8 | 4812414 | 5641152 | 4847889 | 5492666.5 | 4554627.5 | 4280706 | 4907044 | 4627096 | 4318142 | 4426590.5 | 3344385.25 | 4199675 | 4374966.5 | 3598892.5 | 3277001.5 | 4999270.5 | 4902428.5 | 5055083.5 | 4316238 | 4880080.5 | 0 | 12 | 2 | 6 | 2 | 12 | 0.741453582 | 0.833123755 | 0.952832071 | 1.070636182 |
| Q9WTV6 | Usp18 | Ubl carboxyl-terminal hydrolase 18 | 63.02 | 3.26 | 42472 | 1 | 122777.7 | 89398.98 | 105823.52 | 2 | 94885.78 | 103769.71 | 167254.28 | 115380.99 | 154500.91 | 138184.73 | 374113.53 | 333022.09 | 501318.31 | 2 | 389022.44 | 49222.13 | 75154.16 | 2 | 2 | 2 | 4.5 | 3.5 | 5 | 0 | 5 | 3.5 | 3.869035785 | 2.352378789 | 0.301249482 | 0.183160077 |
| Q3UJD6 | Usp19 | Ubiquitin carboxyl-terminal hydrolase 19 | 271.27 | 4.41 | 150549 | 6 | 2272072.75 | 1758229.62 | 1857747.5 | 2385021.5 | 2378754.5 | 2679601.5 | 1892667.75 | 1366663.75 | 1624497.38 | 2242679.75 | 2295942.5 | 1152203.62 | 1632182.75 | 1821230.75 | 1796778.5 | 2548550 | 1771562.25 | 2375215 | 2879339 | 1719701.62 | 5 | 11 | 10 | 8 | 10 | 11 | 0.816605362 | 0.887032473 | 1.06032224 | 1.15176841 |
| Q3TIX9 | Usp39 | U4/U6.U5 tri-snRNP-associated protein 2 | 428.69 | 20.21 | 65146 | 8 | 6384584 | 6019572.5 | 4885444.5 | 5318307 | 7144127.5 | 5566229 | 5751611 | 5947523 | 7360321.5 | 6054210 | 4269631.5 | 5544140 | 3851634.75 | 3606556 | 4900780.5 | 4521855.5 | 3643144.75 | 5032963.5 | 5126938 | 5391743 | 3 | 4 | 0 | 0 | 3 | 4 | 0.745251287 | 0.722712484 | 0.797143602 | 0.773035408 |
| P35123 | Usp4 | Ubiquitin carboxyl-terminal hydrolase 4 | 347.73 | 8.32 | 108343 | 7 | 2906627.5 | 2793725.5 | 2451677 | 3249423.25 | 2757548.25 | 4257333.5 | 3170231 | 2911414 | 2949848 | 3699529.5 | 2547911.25 | 3079752.5 | 2450708 | 3220789 | 3264814 | 2630746.5 | 3387362 | 1103158.12 | 3725122.5 | 3116911.25 | 11 | 10 | 8 | 9 | 11 | 10 | 1.028601823 | 0.857291591 | 0.986178324 | 0.821933586 |
| Q8BY87 | Usp47 | Ubiquitin carboxyl-terminal hydrolase 47 | 103.39 | 1.82 | 157455 | 2 | 329708.56 | 696934.75 | 708609.12 | 571888.88 | 1214668.25 | 277669.16 | 950425.5 | 691276.19 | 1360077.75 | 664822.88 | 2 | 599707.31 | 241500.61 | 2 | 2 | 321183.03 | 2 | 2 | 430001.72 | 363789.22 | 2 | 2 | 1 | 3 | 2 | 3 | 0.238858435 | 0.213274853 | 0.316592351 | 0.282682867 |
| P56399 | Usp5 | Ubiquitin carboxyl-terminal hydrolase 5 | 1449.79 | 36.95 | 95833 | 25 | 51957214 | 52525038 | 51150610 | 55603746 | 59788110 | 46814170 | 51362414 | 49778382 | 54167258 | 51825414 | 35246270 | 38502378 | 35917062 | 28440520 | 33893134 | 54139702 | 48857490 | 48671754 | 49216422 | 49368970 | 0 | 3 | 0 | 8 | 0 | 8 | 0.634626116 | 0.677302476 | 0.923363521 | 0.985456451 |
| Q4VSI4 | Usp7 | Ubiquitin carboxyl-terminal hydrolase 7 | 40.36 | 1 | 128431 | 1 | 636450.44 | 545429.06 | 2 | 674303.06 | 2 | 708495.25 | 575427.56 | 565415.94 | 440816.03 | 491079.06 | 448734.41 | 535420.94 | 380365.47 | 587351.81 | 350248.72 | 494136.81 | 2 | 564174.38 | 2 | 705724.56 | 11 | 12 | 7 | 8 | 11 | 12 | 1.240242441 | 0.827733834 | 0.950356924 | 0.634265168 |
| Q80U87 | Usp8 | Ubiquitin carboxyl-terminal hydrolase 8 | 40.82 | 0.93 | 122611 | 1 | 767222.44 | 660738.38 | 1007004.38 | 830976.75 | 1052967.25 | 714349.06 | 879253.62 | 623184 | 718333.5 | 361721.88 | 429887.53 | 493372.38 | 436103.44 | 663423.81 | 557217.44 | 663565.56 | 2 | 749677.25 | 2 | 709453.62 | 1 | 3 | 6 | 8 | 6 | 8 | 0.597374124 | 0.782568456 | 0.491489941 | 0.643858696 |
| P70398 | Usp9x | Probable ubiquitin carboxyl-terminal hydrolase FAF-X | 332.72 | 3.13 | 290711 | 7 | 4835860 | 4789264 | 5414444 | 4549858.5 | 4502260 | 4840592 | 4465822 | 4886081.5 | 4357288 | 5089332 | 4594501.5 | 4634738 | 5413293 | 4369747.5 | 4589620 | 4051168 | 4513945.5 | 5269434.5 | 5500075.5 | 4801142 | 10 | 12 | 12 | 11 | 12 | 12 | 0.979669896 | 0.998425681 | 1.001829635 | 1.021009669 |
| Q640M1 | Utp14a | U3 small nucleolar RNA-associated protein 14 homolog A | 237.78 | 7.95 | 87265 | 4 | 424755.06 | 634321.19 | 365195 | 416893.75 | 438898.88 | 808959.12 | 488472.22 | 278924.41 | 217859.06 | 280060.66 | 661061.25 | 884583.62 | 929867.81 | 392150.69 | 780119.56 | 301394.34 | 176636.28 | 634467.25 | 226350.66 | 338252.06 | 4 | 5 | 4 | 11 | 4 | 11 | 1.59985997 | 1.758581723 | 0.735549826 | 0.808523561 |
| Q8C7V3 | Utp15 | U3 small nucleolar RNA-associated protein 15 homolog | 245.67 | 10.04 | 59375 | 5 | 2457419 | 3096592.75 | 3619165.5 | 2954373.75 | 3319631.25 | 3703848.5 | 4004879.5 | 2253348.5 | 2687517.5 | 1621797.75 | 4093532.25 | 4882541 | 4623868 | 4266883 | 3907862.5 | 4688559 | 2877398.5 | 2121874 | 1409428.38 | 2469480.25 | 0 | 7 | 1 | 11 | 1 | 11 | 1.409621923 | 1.525757763 | 0.878266334 | 0.950624884 |
| Q5SSI6 | Utp18 | U3 small nucleolar RNA-associated protein 18 homolog | 436.94 | 16.85 | 61218 | 7 | 2401870 | 1828966 | 2294339.25 | 1114091 | 1388069.75 | 3094868 | 2598916.5 | 939626.94 | 2278153.25 | 1103376.25 | 3978814.5 | 3006803 | 1540704.75 | 1738164.25 | 1686825 | 2580703.5 | 4048447.5 | 1325390 | 478421.5 | 1428818.5 | 9 | 12 | 10 | 12 | 10 | 12 | 1.323902367 | 1.193348176 | 1.092435354 | 0.984706855 |
| Q5XG71 | Utp20 | Small subunit processome component 20 homolog | 39.62 | 0.5 | 317744 | 1 | 292033.97 | 231845.16 | 255537.06 | 2 | 242668.11 | 2 | 197446.11 | 307149.62 | 471890.09 | 2 | 160431.16 | 257268.41 | 398231.94 | 346113.34 | 351637.97 | 209092.03 | 2 | 241826.45 | 383941 | 211810.55 | 5 | 9.5 | 8 | 11 | 8 | 11 | 1.480973593 | 1.550126575 | 1.024054456 | 1.071871932 |
| Q9JI13 | Utp3 | Something about silencing protein 10 | 154.91 | 8.96 | 53399 | 3 | 761862.81 | 876591 | 2 | 696475.25 | 170663.92 | 732079.69 | 2 | 2 | 2 | 2 | 1339718.25 | 1321932.75 | 1052715.25 | 278440.72 | 1757686.75 | 71853.21 | 1017095.31 | 317874.59 | 2 | 2 | 3 | 9 | 1 | 8 | 3 | 9 | 2.295061159 | 7.854924756 | 0.561474269 | 1.921664753 |
| Q80WQ2 | Vac14 | Protein VAC14 homolog | 512.27 | 15.98 | 88048 | 11 | 6418253 | 7154448 | 6626949 | 8635488 | 10394551 | 6428043 | 6578552 | 7092412 | 6090404 | 9585462 | 7888703 | 8295298 | 8368720 | 6166671.5 | 8120360 | 7252658 | 7824170.5 | 8560420 | 7088657.5 | 8524633 | 12 | 11 | 8 | 6 | 12 | 11 | 0.990060168 | 1.085671289 | 1.000531485 | 1.097153832 |
| P63024 | Vamp3 | Vesicle-associated membrane protein 3 | 48.87 | 23.3 | 11480 | 1 | 848162.62 | 658646 | 2 | 983470.69 | 2 | 1153390.12 | 533229.94 | 399417.78 | 470502.72 | 474081.34 | 779611 | 672609.06 | 939780.94 | 973769.62 | 405687.91 | 493268.88 | 1129507.12 | 854994.06 | 511039.06 | 266040.34 | 9 | 10 | 8 | 11 | 9 | 11 | 1.514469665 | 1.244450365 | 1.307019746 | 1.073987309 |
| P70280 | Vamp7 | Vesicle-associated membrane protein 7 | 187.64 | 19.09 | 24967 | 4 | 1635746.62 | 2140641 | 329281.72 | 2114918.75 | 1654199.5 | 2262587.75 | 1984663.75 | 1659870.88 | 1339635.5 | 1621420.62 | 6744768 | 3070343 | 3539290 | 3272297.25 | 3552770.5 | 1372447.25 | 3000810.25 | 1714071.88 | 2 | 2204767.25 | 0 | 11 | 0 | 12 | 0 | 12 | 2.562541341 | 2.275491946 | 1.052993307 | 0.935039662 |
| O70404 | Vamp8 | Vesicle-associated membrane protein 8 | 151.69 | 30.69 | 11451 | 3 | 3721272.5 | 2873398.75 | 2212145.25 | 2 | 2546649.25 | 4111961.5 | 3512886.5 | 4635420 | 3538196.25 | 3936588.75 | 4362770.5 | 3936863.25 | 5550547.5 | 5222436.5 | 5254522.5 | 2943725 | 4564241.5 | 3507635.5 | 4396576.5 | 4301657 | 0 | 2 | 3 | 12 | 3 | 12 | 2.142705716 | 1.232686847 | 1.736371295 | 0.998924883 |
| Q9WV55 | Vapa | Vesicle-associated membrane protein-associated protein A | 640.84 | 61.85 | 27855 | 13 | 26413584 | 31079540 | 23345656 | 24419186 | 23578750 | 29828764 | 30623798 | 22647174 | 21246396 | 23947742 | 22142514 | 27667172 | 25932932 | 27838368 | 28562550 | 33945938 | 31952902 | 30099460 | 30510236 | 35705502 | 10 | 2 | 12 | 2 | 12 | 2 | 1.025666752 | 1.030006593 | 1.259066849 | 1.264394261 |
| Q9QY76 | Vapb | Vesicle-associated membrane protein-associated protein B | 552.73 | 31.69 | 26946 | 10 | 8452380 | 13045572 | 4863369 | 5556996 | 6119141 | 8763073 | 7166344 | 7125603 | 6236399.5 | 12580681 | 7650806.5 | 5776161 | 6750693 | 12468193 | 5686950.5 | 10023198 | 9747719 | 6666794 | 8991320 | 12028282 | 11 | 6 | 8 | 8 | 11 | 8 | 1.007764609 | 0.915473634 | 1.247646806 | 1.133387445 |
| Q9Z1Q9 | Vars | Valine--tRNA ligase | 1888.22 | 30.4 | 140215 | 32 | 50986826 | 55554998 | 61330266 | 55920878 | 45779634 | 57261562 | 51417678 | 57305882 | 53100306 | 53590518 | 60279890 | 58725378 | 48101930 | 48691702 | 52006226 | 53852958 | 57617438 | 53836030 | 55492242 | 57162198 | 12 | 11 | 11 | 8 | 12 | 11 | 0.993443414 | 0.982136965 | 1.031116901 | 1.019381688 |
| Q9CZT5 | Vasn | Vasorin | 54.58 | 1.04 | 72261 | 1 | 1015842.25 | 1813874.88 | 637234.94 | 751756.94 | 507963.62 | 1474410.5 | 1170301.62 | 982576.12 | 1059798.62 | 1505575.5 | 13321597 | 10826836 | 10036818 | 9256282 | 7948387 | 1646626 | 1686999.25 | 2187867.25 | 2246833.25 | 1851863.12 | 0 | 2 | 0 | 0 | 0 | 2 | 10.87232479 | 8.29851799 | 2.035298321 | 1.553481897 |
| P70460 | Vasp | Vasodilator-stimulated phosphoprotein | 576.19 | 33.33 | 39667 | 11 | 13484306 | 15671691 | 10997197 | 13703274 | 12207399 | 14980350 | 12997234 | 15550622 | 14424016 | 15123393 | 16390229 | 17488518 | 14037108 | 15089688 | 13243461 | 16208733 | 16522083 | 15100845 | 15971224 | 15308952 | 5 | 2 | 10 | 3 | 10 | 3 | 1.154171069 | 1.0434261 | 1.197505393 | 1.082602411 |
| Q62465 | Vat1 | Synaptic vesicle membrane protein VAT-1 homolog | 685.73 | 30.05 | 43097 | 10 | 29689820 | 26953332 | 21245174 | 25236126 | 19650526 | 30773062 | 26496514 | 20837246 | 20235510 | 25918066 | 14295416 | 12976245 | 10520504 | 11816617 | 11204814 | 28919982 | 29476540 | 30522274 | 26522550 | 27099740 | 0 | 5 | 0 | 5 | 0 | 5 | 0.495325652 | 0.489404484 | 1.160994596 | 1.147115962 |
| P27870 | Vav1 | Proto-oncogene vav | 1182.78 | 26.15 | 98137 | 21 | 36002466 | 41106910 | 36917594 | 43063690 | 45634882 | 33955034 | 34862950 | 43891186 | 46048802 | 41923774 | 44282850 | 42354018 | 45706754 | 38604950 | 41875074 | 38086910 | 30486918 | 41284286 | 42636742 | 41972110 | 8 | 11 | 10 | 10 | 10 | 11 | 1.049811701 | 1.060503261 | 0.959262282 | 0.969031663 |
| Q9R0C8 | Vav3 | Guanine nucleotide exchange factor VAV3 | 227.49 | 3.78 | 97968 | 4 | 1349489.88 | 1052709.75 | 1696430.88 | 1737423.38 | 1674872.88 | 1716416.62 | 1281913.12 | 1657558.75 | 2102974.5 | 1472897.12 | 743395.19 | 611824.94 | 993360.56 | 1262519.75 | 986387.25 | 1159935.62 | 1447178.5 | 1400253.25 | 1445472.25 | 1408097.75 | 1 | 9 | 0 | 4 | 1 | 9 | 0.612106579 | 0.558506034 | 0.913460826 | 0.833471491 |
| P61759 | Vbp1 | Prefoldin subunit 3 | 295.7 | 31.63 | 22436 | 6 | 3918979.5 | 3944803 | 3984370 | 3781369 | 3446785.75 | 2836780 | 4317837.5 | 3451772 | 2107163.75 | 2605783.25 | 3886825.5 | 3546592 | 1506527.75 | 960939.19 | 2414493.5 | 4842112 | 3579572.25 | 3455285.25 | 4344448.5 | 2899490.5 | 3 | 12 | 9 | 4 | 9 | 12 | 0.645585007 | 0.803910662 | 1.002338044 | 1.248155134 |
| P85972 | Vcl | Vinculin | 1078.48 | 21.76 | 116615 | 19 | 28655644 | 27941326 | 20108586 | 22346392 | 16981428 | 28427046 | 24004922 | 13960240 | 15978833 | 18211654 | 9927106 | 13637514 | 9991975 | 10192090 | 8740704 | 28058218 | 29302430 | 21505680 | 19961664 | 22095792 | 0 | 11 | 0 | 7 | 0 | 11 | 0.452364577 | 0.521853078 | 1.042146563 | 1.202232491 |
| P46462 | Vcp | Transitional endoplasmic reticulum ATPase | 2688.03 | 50.62 | 89349 | 40 | 47629514 | 51799822 | 49372922 | 45948862 | 48753146 | 47381394 | 45716394 | 50656322 | 44724666 | 44804334 | 99144786 | 79581594 | 66948146 | 61632582 | 69792594 | 52829846 | 54300410 | 51747294 | 45710514 | 51466926 | 0 | 7 | 0 | 3 | 0 | 7 | 1.548636943 | 1.616489518 | 1.051542111 | 1.097614782 |
| Q8CDG3 | Vcpip1 | Deubiquitinating protein VCIP135 | 174.94 | 4.02 | 134503 | 4 | 1305174.75 | 1738111.88 | 1826271.25 | 1964256.75 | 2265904.75 | 1824407.5 | 2296907.5 | 2393859.5 | 2485559 | 2554345 | 2077501 | 2114728.5 | 1365264.75 | 1706199.62 | 2520058.5 | 1409874.88 | 2047932.88 | 2784567 | 1753731 | 2918763 | 10 | 8 | 6 | 11 | 10 | 11 | 1.075170779 | 0.846705833 | 1.199473116 | 0.944594947 |
| Q9Z2L0 | Vdac1 | Voltage-dependent anion-selective channel protein 1 | 1450.83 | 81.98 | 30756 | 18 | 60167878 | 59722070 | 61659314 | 65666914 | 56339218 | 68231370 | 65049754 | 64415722 | 60508558 | 77020298 | 116506082 | 94395450 | 116189730 | 109699994 | 123360874 | 59242662 | 69538130 | 75454322 | 78370978 | 66042982 | 0 | 4 | 0 | 9 | 0 | 9 | 1.845304485 | 1.670970116 | 1.148551734 | 1.040042789 |
| Q60930 | Vdac2 | Voltage-dependent anion-selective channel protein 2 | 1125.74 | 64.07 | 31733 | 17 | 196537442 | 209380466 | 186308658 | 205320002 | 191581698 | 236378466 | 223776658 | 234575538 | 217723074 | 228284386 | 254540242 | 236025938 | 282303778 | 278693570 | 276677826 | 202572610 | 193546914 | 244126194 | 243741954 | 244090386 | 0 | 5 | 1 | 10 | 1 | 10 | 1.342840357 | 1.164370094 | 1.140477021 | 0.988901866 |
| Q60931 | Vdac3 | Voltage-dependent anion-selective channel protein 3 | 714.93 | 50.88 | 30753 | 11 | 47109246 | 48709058 | 37230782 | 38317190 | 62646186 | 56217470 | 59861410 | 52504606 | 71175170 | 47098922 | 58731614 | 56570650 | 65294418 | 63417374 | 61519382 | 42614706 | 42892854 | 55931062 | 51198046 | 50366942 | 3 | 9 | 7 | 4 | 7 | 9 | 1.30562892 | 1.065104991 | 1.038421663 | 0.847122853 |
| P20152 | Vim | Vimentin | 4660.771 | 89.7 | 53688 | 70 | 1889461378 | 2305572354 | 1013441986 | 1095957762 | 1028872386 | 1969269762 | 2157559298 | 1184613506 | 1131866882 | 1160073730 | 1329626242 | 1633923586 | 916101826 | 690902658 | 1013319298 | 1782584066 | 1371426306 | 1190947202 | 1240137730 | 1256292866 | 6 | 10 | 6 | 10 | 6 | 10 | 0.761440162 | 0.734393293 | 0.932920063 | 0.899782111 |
| A2AIV2 | Virma | Protein virilizer homolog | 65.28 | 0.66 | 201439 | 1 | 314971.19 | 228099.5 | 2 | 284063.06 | 192868.02 | 380445.28 | 344675.84 | 302680.38 | 344824.47 | 225198.92 | 200124.47 | 192052.2 | 201227.56 | 2 | 129095.71 | 388329.09 | 352603.31 | 222922.09 | 273278.53 | 284839.97 | 6.5 | 6 | 0 | 11 | 6.5 | 11 | 0.708332617 | 0.452178424 | 1.492124867 | 0.952528027 |
| Q6TEK4 | Vkorc1 | Vitamin K epoxide reductase complex subunit 1 | 64.36 | 8.07 | 17783 | 1 | 2 | 202863.5 | 2 | 165812.97 | 169864.92 | 2 | 2 | 317022.69 | 413992.56 | 494239.03 | 1074096.25 | 2 | 2 | 1448982.12 | 1701825.88 | 369664.69 | 2 | 544800.25 | 357661.28 | 179871.12 | 8 | 5 | 8 | 11 | 8 | 11 | 7.845036516 | 3.448177677 | 2.696150347 | 1.185055726 |
| Q6TEK3 | Vkorc1l1 | Vitamin K epoxide reductase complex subunit 1-like protein 1 | 65.58 | 6.25 | 19779 | 1 | 2 | 2 | 95691.94 | 2 | 251327.27 | 229610.09 | 142357.86 | 552914.88 | 464456.78 | 426546.97 | 538409.12 | 772853.69 | 855542.12 | 874415.88 | 948482.5 | 2 | 2 | 597613.31 | 2 | 566688.25 | 0 | 10.5 | 1 | 10 | 1 | 10.5 | 11.4968688 | 2.197110411 | 3.355109446 | 0.64117857 |
| Q78T54 | Vma21 | Vacuolar ATPase assembly integral membrane protein Vma21 | 181.95 | 37.62 | 11365 | 4 | 546356.31 | 475445.41 | 279699.47 | 716482.56 | 1198286.25 | 1649381.5 | 381259.06 | 159911.34 | 1927684.88 | 1264982 | 4473417 | 4535038 | 3490414 | 1160670.88 | 2741137.25 | 854761.44 | 677094.38 | 2896646.25 | 516005.09 | 2 | 1 | 11 | 3 | 11 | 3 | 11 | 5.099284926 | 3.046630241 | 1.537342686 | 0.918504219 |
| Q91ZQ0 | Vmp1 | Vacuole membrane protein 1 | 92.36 | 3.94 | 45901 | 2 | 931438 | 608458.81 | 646860.81 | 1707590.75 | 1750455.75 | 643602.31 | 1360828 | 814626.69 | 2299991.5 | 1117358.12 | 294071.75 | 1822041.12 | 2405583.25 | 2981760 | 255893.98 | 1450547.38 | 2121670.75 | 1852486.25 | 1819718.12 | 1646560.38 | 10 | 4 | 11 | 5 | 11 | 5 | 1.374600417 | 1.244202082 | 1.575073765 | 1.425657983 |
| Q91W86 | Vps11 | Vacuolar protein sorting-associated protein 11 homolog | 123.05 | 2.98 | 107719 | 2 | 465810.88 | 1250757.25 | 957945.88 | 1041641.5 | 999223 | 182182.17 | 362992.5 | 1160486.88 | 1120420 | 1201309.25 | 510731.25 | 561978.31 | 763763.75 | 811255.44 | 434932.47 | 668043.94 | 509809.31 | 954704.12 | 869496.62 | 965292.81 | 4 | 6 | 10 | 10 | 10 | 10 | 0.653746293 | 0.765423912 | 0.84136338 | 0.985091092 |
| Q8BX70 | Vps13c | Vacuolar protein sorting-associated protein 13C | 2107.49 | 13.85 | 420089 | 40 | 36710974 | 35834090 | 36290782 | 32823178 | 40141670 | 34702422 | 36884898 | 33890582 | 38300538 | 30373368 | 20762064 | 21228888 | 22420232 | 27711270 | 26053608 | 37540190 | 33643306 | 29878902 | 30991280 | 31981720 | 0 | 5 | 0 | 7 | 0 | 7 | 0.650030863 | 0.678580736 | 0.902281473 | 0.941910393 |
| Q920Q4 | Vps16 | Vacuolar protein sorting-associated protein 16 homolog | 313.08 | 9.54 | 94928 | 5 | 1092791.38 | 1241800.38 | 1539708.62 | 1619977.25 | 2210453.75 | 773375.88 | 1317035.75 | 1677803.25 | 1659838.25 | 1470818.62 | 510353.84 | 810719.81 | 1010704.94 | 274139.19 | 536890.81 | 1712562.5 | 522165.94 | 1257584.25 | 671575.44 | 1445096 | 0 | 8 | 2 | 8 | 2 | 8 | 0.407906316 | 0.455553995 | 0.727992172 | 0.813029193 |
| Q8R307 | Vps18 | Vacuolar protein sorting-associated protein 18 homolog | 45.88 | 0.72 | 110219 | 1 | 473860.69 | 2 | 504842.91 | 587246.69 | 482856.97 | 544764.88 | 2 | 588634.69 | 575210.06 | 642001.12 | 760648.31 | 540769.31 | 875074.06 | 756779.19 | 895857.88 | 322614.03 | 467896.31 | 734421.94 | 596660.75 | 496444.25 | 1 | 10 | 4 | 12 | 4 | 12 | 1.868953262 | 1.628991738 | 1.277833584 | 1.113768008 |
| P0C0A1 | Vps25 | Vacuolar protein-sorting-associated protein 25 | 61.07 | 8.52 | 20762 | 1 | 609085.38 | 462877.62 | 505933.94 | 372415.81 | 578294.25 | 339562.25 | 575928.5 | 500324.16 | 651941.38 | 691960.62 | 487620.34 | 730509.94 | 611553.94 | 488255.53 | 453801.84 | 536537.44 | 343097.53 | 500212.62 | 409730.84 | 557677.31 | 10 | 9 | 11 | 7 | 11 | 9 | 1.09615357 | 1.004357215 | 0.928280172 | 0.850542217 |
| P40336 | Vps26a | Vacuolar protein sorting-associated protein 26A | 355.17 | 28.75 | 38114 | 7 | 5420480.5 | 6425514 | 6605964 | 7873572.5 | 11506948 | 8673093 | 7671731 | 7160480.5 | 10848586 | 8359787 | 6639582.5 | 6393673 | 6522068 | 3620339.75 | 6771320 | 6823164 | 7502488 | 7080682 | 7201265.5 | 6124029.5 | 9 | 12 | 0 | 2 | 9 | 12 | 0.791568093 | 0.701109926 | 0.918037356 | 0.813126638 |
| Q8C0E2 | Vps26b | Vacuolar protein sorting-associated protein 26B | 86.48 | 5.95 | 39125 | 2 | 769989 | 689666.88 | 759103.19 | 695184.38 | 2 | 679834.5 | 613522.69 | 806408.81 | 716242.94 | 826762 | 803761.94 | 580984.38 | 686022.44 | 956728.06 | 683866.19 | 490257.66 | 724374.44 | 805884.69 | 226341 | 558440.88 | 12 | 11 | 12 | 6 | 12 | 11 | 1.273655624 | 1.018829641 | 0.962714889 | 0.770100211 |
| B5DEN9 | Vps28 | Vacuolar protein sorting-associated protein 28 homolog | 169.07 | 16.23 | 26110 | 3 | 813347.69 | 1366617.62 | 955957.31 | 1061970.25 | 684789.56 | 967268.62 | 1085964.62 | 842370.69 | 1119210.12 | 892973.81 | 1318905.5 | 1010706.19 | 801635.88 | 2 | 2 | 1199996.25 | 1539832.5 | 939660.25 | 1079917.25 | 1030629.56 | 8 | 7 | 8 | 7 | 8 | 7 | 0.641297405 | 0.638016895 | 1.185830922 | 1.179764891 |
| Q9QZ88 | Vps29 | Vacuolar protein sorting-associated protein 29 | 207.44 | 20.88 | 20496 | 4 | 3490242 | 3687587 | 5294479 | 5067793.5 | 6161209 | 3180262 | 3432617.75 | 5746108 | 4771971.5 | 4515376 | 2481991.5 | 3689138.75 | 4250820.5 | 3082923 | 4900413.5 | 5108286.5 | 3399564 | 5942527.5 | 5668257 | 7797946.5 | 6 | 9 | 8 | 6 | 8 | 9 | 0.776551459 | 0.850272669 | 1.177849702 | 1.289667797 |
| Q63615 | Vps33a | Vacuolar protein sorting-associated protein 33A | 115.7 | 6.53 | 67514 | 2 | 453075.5 | 527791.62 | 568094.75 | 491251.5 | 613865.56 | 135939.11 | 117545.12 | 703051 | 694951.19 | 434291 | 204163.66 | 411082.38 | 370113.91 | 183615.72 | 261050.84 | 330095.03 | 2 | 300283.34 | 419353 | 333363.06 | 0 | 0 | 10 | 8 | 10 | 8 | 0.538803309 | 0.685608395 | 0.521121062 | 0.663108353 |
| Q9EQH3 | Vps35 | Vacuolar protein sorting-associated protein 35 | 1306.29 | 28.27 | 91713 | 20 | 30047288 | 24635438 | 29207304 | 29779862 | 37465010 | 25953864 | 30422920 | 32259542 | 38844022 | 33918538 | 25631336 | 30343294 | 31806814 | 24587448 | 31221152 | 28909232 | 25219446 | 28320662 | 29745414 | 31410086 | 12 | 9 | 5 | 5 | 12 | 9 | 0.950078652 | 0.889659449 | 0.950176552 | 0.889751123 |
| P0C0A2 | Vps36 | Vacuolar protein-sorting-associated protein 36 | 243.8 | 12.18 | 43718 | 4 | 5500451.5 | 5713930.5 | 4268811.5 | 5409231.5 | 5234844 | 4412802.5 | 5538306 | 5359284.5 | 5889099 | 5965031.5 | 4034437 | 5344872 | 4717911 | 3152934.75 | 4037304.75 | 5435576 | 2878204 | 5710082 | 5869895 | 5595939.5 | 3 | 9 | 2 | 11 | 3 | 11 | 0.814760222 | 0.783649288 | 0.97559743 | 0.938345062 |
| O08700 | Vps45 | Vacuolar protein sorting-associated protein 45 | 283.87 | 8.42 | 64894 | 5 | 1696794 | 1665274.88 | 1967764 | 2248604.5 | 2452695.75 | 1885417.75 | 1582488.12 | 1499971.5 | 2125026.75 | 1264495.88 | 1460551.62 | 1243699.5 | 1867863.5 | 1541318.5 | 1899828.75 | 1875450 | 1146046.25 | 1948122 | 1123398.38 | 1341887.12 | 4 | 4 | 10 | 8 | 10 | 8 | 0.798839151 | 0.958822345 | 0.741182841 | 0.88961923 |
| P46467 | Vps4b | Vacuolar protein sorting-associated protein 4B | 318.58 | 16.67 | 49419 | 6 | 4427319.5 | 5374847 | 5165953.5 | 3213315.5 | 4034591.75 | 3680319 | 4795621.5 | 5872638 | 3799101.25 | 4982728.5 | 9579873 | 9014684 | 7938859 | 5036418.5 | 6282244.5 | 4045603.5 | 2614145.5 | 4888478.5 | 3280572.5 | 4648136 | 2 | 9 | 1 | 7 | 2 | 9 | 1.703818535 | 1.636463939 | 0.876706523 | 0.842048951 |
| F1LSG8 | Vps50 | Syndetin | 107 | 2.39 | 111188 | 2 | 653030.75 | 1582079.12 | 864747 | 1780181.75 | 1403921 | 1550003.25 | 1722078.88 | 1447936.5 | 1588502.38 | 971969.5 | 1328695 | 1244506.75 | 951969.69 | 730020.25 | 403699.03 | 1327817.88 | 1452747 | 1264991.5 | 1063881.62 | 1468092.5 | 7 | 12 | 2 | 7 | 7 | 12 | 0.741394121 | 0.6399144 | 1.046717499 | 0.903446065 |
| Q3UVL4 | Vps51 | Vacuolar protein sorting-associated protein 51 homolog | 361.29 | 10.1 | 86187 | 7 | 3536787.75 | 3066622.75 | 2965203.25 | 2621920 | 3917567.25 | 2697742.75 | 3327398.75 | 3304563.25 | 3803573.5 | 3063523.75 | 1518954.88 | 2065540.25 | 2593693.75 | 2708926.5 | 2658056.75 | 3604963.75 | 3306237.75 | 3008309 | 3223568 | 3143753.75 | 2 | 10 | 1 | 12 | 2 | 12 | 0.716730801 | 0.712805659 | 1.011095737 | 1.00555852 |
| Q8CCB4 | Vps53 | Vacuolar protein sorting-associated protein 53 homolog | 59.11 | 1.8 | 94423 | 1 | 624702.19 | 2 | 670273.38 | 645026.44 | 975861.5 | 912788.88 | 413961.12 | 2 | 760998.81 | 344601.47 | 2 | 269257.38 | 261334.69 | 177576.06 | 2 | 908277.56 | 2 | 260548.27 | 989631.31 | 495584.44 | 4 | 11.5 | 4 | 11.5 | 4 | 11.5 | 0.242868585 | 0.291147025 | 0.910207817 | 1.09114276 |
| Q80X41 | Vrk1 | Serine/threonine-protein kinase VRK1 | 109.04 | 5 | 49741 | 2 | 978046.62 | 788956.38 | 1815290.25 | 475299.34 | 3667109.5 | 644484 | 2217056.75 | 3112822.5 | 3193335.25 | 881839.81 | 570600.44 | 2 | 537899 | 368466.69 | 543111.44 | 2 | 823664.31 | 680205 | 393043.47 | 444291.09 | 3 | 3 | 0 | 2 | 3 | 3 | 0.261509058 | 0.201012177 | 0.303080409 | 0.23296651 |
| Q8BN21 | Vrk2 | Serine/threonine-protein kinase VRK2 | 324.75 | 16.3 | 58119 | 6 | 1980232.5 | 2254226.5 | 3405887.5 | 1538701.62 | 5666701.5 | 3036892.5 | 3087615.75 | 1096893.88 | 6304819 | 1607044.62 | 1916531.75 | 2433435.25 | 2031280.5 | 1861623.88 | 2756570.5 | 2484215.5 | 1164458 | 2054151.38 | 1191708.38 | 2201807.5 | 10 | 7 | 10 | 8 | 10 | 8 | 0.740915222 | 0.726838612 | 0.612723574 | 0.60108247 |
| Q9CR26 | Vta1 | Vacuolar protein sorting-associated protein VTA1 homolog | 95.89 | 7.77 | 33913 | 2 | 1895736.62 | 1706983.62 | 1914528 | 1920838.25 | 890168.38 | 1113728.62 | 1569134 | 931938.12 | 2223510.75 | 2225500.5 | 891580.06 | 1578245.5 | 2211677.75 | 1086336 | 833766.12 | 1859055 | 828784.81 | 1736346.25 | 1740271.5 | 1979142.62 | 8 | 11 | 7 | 12 | 8 | 12 | 0.792675721 | 0.818670554 | 0.977827925 | 1.009894599 |
| O88384 | Vti1b | Vesicle transport through interaction with t-SNAREs homolog 1B | 77.71 | 8.19 | 26713 | 2 | 1063271 | 465131.25 | 491723.66 | 534842.75 | 510579.44 | 1276905.25 | 1018398.62 | 1026685.81 | 811042.06 | 1325304.25 | 2034035.75 | 1802559.12 | 2075501.62 | 2125032 | 1784635 | 491219.19 | 983944.75 | 1440267.5 | 1229325 | 1229025 | 0 | 5 | 0 | 12 | 0 | 12 | 3.203917593 | 1.799406176 | 1.752959427 | 0.984509098 |
| Q99KC8 | Vwa5a | von Willebrand factor A domain-containing protein 5A | 1672.17 | 37.45 | 87143 | 26 | 97203218 | 98990634 | 115614418 | 107912442 | 115783762 | 82159258 | 97843546 | 93395466 | 108455882 | 92003738 | 35362374 | 44539886 | 46947962 | 35819954 | 48290238 | 85062826 | 67895530 | 79923730 | 77929010 | 78708538 | 0 | 0 | 0 | 1 | 0 | 1 | 0.393947062 | 0.445197639 | 0.727388197 | 0.822017829 |
| P32921 | Wars | Tryptophan--tRNA ligase, cytoplasmic | 554.26 | 23.08 | 54358 | 9 | 9464140 | 13007876 | 8930346 | 8952283 | 7959832 | 10295216 | 11647668 | 9195988 | 8908527 | 9800459 | 17773804 | 18824938 | 17222980 | 15497672 | 17019038 | 11225826 | 11062462 | 11349393 | 10603952 | 8473847 | 0 | 8 | 0 | 9 | 0 | 9 | 1.787009554 | 1.732038957 | 1.091090772 | 1.057527487 |
| P70315 | Was | Wiskott-Aldrich syndrome protein homolog | 165 | 9.81 | 54192 | 3 | 1631680.38 | 2014299.12 | 1864197.88 | 1483118.25 | 1555410.12 | 1559895.25 | 1950293.62 | 1155010 | 1027908.69 | 2057072.38 | 1083570.12 | 1241296.62 | 763381.81 | 314577.84 | 1428877.88 | 2249134.75 | 1662551.75 | 1400666.12 | 1594774.25 | 1444301 | 0 | 10 | 5 | 10 | 5 | 10 | 0.565197167 | 0.623431237 | 0.97692307 | 1.077578577 |
| Q8BH43 | Wasf2 | Wiskott-Aldrich syndrome protein family member 2 | 574.4 | 23.54 | 54074 | 9 | 8905475 | 8737580 | 7916331.5 | 7972427.5 | 7971929 | 8633420 | 7722390 | 7030047 | 7204585 | 8038061 | 6544242 | 7176856 | 6487607 | 7156739 | 4827495 | 6885549 | 8410950 | 8636886 | 5919564.5 | 7909005.5 | 0 | 6 | 2 | 12 | 2 | 12 | 0.775663511 | 0.833398566 | 0.909844565 | 0.977567135 |
| Q6PGL7 | Washc2 | WASH complex subunit 2 | 76.6 | 1.65 | 145311 | 1 | 2 | 2 | 2 | 2 | 495407.91 | 2 | 131958.83 | 2 | 2 | 63881.21 | 2 | 2 | 2 | 142894.05 | 199267.75 | 2 | 2 | 198244.7 | 181994.23 | 167889.14 | 11 | 9 | 10.5 | 7 | 11 | 9 | 0.690667766 | 1.747126467 | 1.106407887 | 2.798790673 |
| Q3UMB9 | Washc4 | WASH complex subunit 4 | 407.14 | 9.55 | 136370 | 7 | 1634553.38 | 1061743.62 | 2137427.5 | 1962022.88 | 3224147.75 | 1136953.12 | 1662380.5 | 1841466 | 2714920 | 1317200.88 | 789322.75 | 1130095.12 | 1237552.25 | 707799.88 | 1287356.75 | 1748264.62 | 1306162.75 | 1746232.12 | 1933238.25 | 1354970.25 | 3 | 8 | 2 | 12 | 3 | 12 | 0.514189688 | 0.594047501 | 0.807280704 | 0.93265792 |
| Q8C2E7 | Washc5 | WASH complex subunit 5 | 297.23 | 5.52 | 134110 | 5 | 1617371.25 | 1367381.5 | 2711810.5 | 3390143.5 | 2901820.5 | 1263408.62 | 1615320.75 | 2898598.75 | 3089311 | 3149421.5 | 1938355 | 2953030 | 3069249.5 | 2795611.75 | 3268608 | 2020784 | 1390340.62 | 3041467 | 3489362 | 2913051.75 | 8 | 9 | 10 | 11 | 10 | 11 | 1.16985631 | 1.167175724 | 1.07227561 | 1.069818618 |
| Q5PQQ2 | Wbp11 | WW domain-binding protein 11 | 44.7 | 1.72 | 69996 | 1 | 241733.67 | 377462.5 | 2 | 574926.19 | 2 | 594088.69 | 411018.16 | 2 | 436189.31 | 507528 | 439600 | 314921.56 | 429410.34 | 396391.34 | 321113.31 | 339827.56 | 193485.14 | 455189 | 2 | 2 | 7 | 11 | 8 | 6 | 8 | 11 | 1.592324409 | 0.975682998 | 0.827806615 | 0.507231338 |
| P97765 | Wbp2 | WW domain-binding protein 2 | 82.39 | 7.66 | 28032 | 2 | 587538.88 | 793644.12 | 712062.56 | 667332.75 | 692643.88 | 736450 | 2 | 707253.81 | 561766.69 | 874039.62 | 1833610.88 | 1718290.12 | 1684907.88 | 1697151.62 | 2362006 | 699766.19 | 1331791.75 | 816563.44 | 782835.56 | 982002.06 | 0 | 3 | 0 | 5 | 0 | 5 | 2.69196883 | 3.228313031 | 1.335841931 | 1.601993257 |
| P59328 | Wdhd1 | WD repeat and HMG-box DNA-binding protein 1 | 90.16 | 1.52 | 124254 | 2 | 794815.12 | 1749766.75 | 1095572.38 | 460122.22 | 2 | 1318797.75 | 1309954.38 | 864865.62 | 530459.56 | 1240876 | 1019538.44 | 1293964.25 | 542149.88 | 1108216.75 | 614082.38 | 401373.28 | 358731.31 | 650367.19 | 2 | 1202560.5 | 10 | 8.5 | 9 | 3 | 10 | 8.5 | 1.116497753 | 0.869514207 | 0.63728215 | 0.496307208 |
| O88342 | Wdr1 | WD repeat-containing protein 1 | 815.55 | 22.94 | 66407 | 13 | 94452602 | 103854826 | 97041266 | 111003442 | 88398122 | 87175874 | 82373018 | 74499474 | 83440130 | 77747098 | 69147250 | 63404738 | 66659022 | 66015102 | 63532254 | 82033626 | 88809962 | 89248370 | 92050570 | 88170490 | 0 | 3 | 0 | 3 | 0 | 3 | 0.664493572 | 0.811277121 | 0.889970265 | 1.08656057 |
| Q8K1X1 | Wdr11 | WD repeat-containing protein 11 | 69.08 | 1.47 | 135937 | 1 | 2 | 106214.77 | 167675.55 | 161729.36 | 319743.97 | 2 | 111559.8 | 187036.83 | 250676.48 | 115191.3 | 2 | 2 | 69734.84 | 49839.43 | 158649.23 | 52264.12 | 2 | 49700.98 | 2 | 202215.97 | 5 | 7 | 6 | 7 | 6 | 7 | 0.368334859 | 0.418723198 | 0.402699103 | 0.457788483 |
| Q9JJA4 | Wdr12 | Ribosome biogenesis protein WDR12 | 306.57 | 19.15 | 47347 | 6 | 2 | 356754.12 | 2 | 1074678.88 | 203723.42 | 2 | 2 | 134528.75 | 126716.2 | 2 | 235150.27 | 620931.12 | 2 | 73831.39 | 440418.16 | 2 | 2 | 119700.2 | 414865.25 | 2 | 11 | 9 | 5.5 | 12.5 | 11 | 12.5 | 0.838041897 | 5.245274477 | 0.326922939 | 2.046199066 |
| Q8C6G8 | Wdr26 | WD repeat-containing protein 26 | 377.34 | 12.32 | 70543 | 6 | 2578093 | 2253415.5 | 3431840.5 | 2126202.25 | 2330961.5 | 1627820.25 | 3257984.75 | 3529572.5 | 2047921.38 | 2027943.88 | 950980.38 | 3654498 | 2315045.75 | 464880.38 | 2495557 | 3279998.5 | 772471.62 | 1633989.25 | 1876081.62 | 1495402 | 10 | 4 | 11 | 6 | 11 | 6 | 0.776773838 | 0.791031101 | 0.712073732 | 0.72514346 |
| Q8BHB4 | Wdr3 | WD repeat-containing protein 3 | 192.81 | 4.88 | 105775 | 3 | 946664.81 | 832429.5 | 1104522 | 888200.38 | 1458225.12 | 909825.69 | 774971.56 | 989083.38 | 1152588.12 | 1088799.25 | 1672426.12 | 1426613.12 | 1282753.12 | 1063075.12 | 1946496.5 | 921609.44 | 1336111 | 1270437.38 | 1246714 | 1050712.88 | 4 | 8 | 2 | 5 | 4 | 8 | 1.413251413 | 1.503756047 | 1.113869623 | 1.185201845 |
| Q6ZQL4 | Wdr43 | WD repeat-containing protein 43 | 707.57 | 23.93 | 75381 | 11 | 6248391.5 | 5780576 | 5086972 | 2776046.5 | 5214896 | 5934959 | 6196405.5 | 5425418 | 5411452 | 5554511 | 7856185 | 5663910 | 7301125 | 5740570 | 5139069.5 | 6318533.5 | 5200360 | 4969212 | 6290249.5 | 4890226 | 7 | 11 | 9 | 10 | 9 | 11 | 1.262636256 | 1.111423846 | 1.102031746 | 0.970053216 |
| Q9Z0H1 | Wdr46 | WD repeat-containing protein 46 | 252.08 | 11.9 | 69048 | 4 | 459916 | 408652.75 | 349996.5 | 325746.91 | 447069.22 | 354702.44 | 273722.44 | 365732.06 | 327849.16 | 288688.69 | 296305.28 | 433723.78 | 424960.88 | 217227.72 | 252882.81 | 314932.88 | 530710.5 | 266815.38 | 2 | 203630.23 | 6 | 5 | 12 | 7 | 12 | 7 | 0.81606692 | 1.008943768 | 0.66089349 | 0.817095205 |
| Q4V7A0 | Wdr61 | WD repeat-containing protein 61 | 194.81 | 9.51 | 33747 | 3 | 1707942.62 | 989921.38 | 1336334 | 923264.12 | 1886369.5 | 507422.47 | 1634028.12 | 711756.44 | 2194133.25 | 1070411.62 | 2 | 940021.25 | 559723.5 | 1039827.75 | 822389.19 | 1595544.25 | 2 | 533218.75 | 2 | 690208.38 | 3 | 3 | 7 | 5 | 7 | 5 | 0.491239977 | 0.549542339 | 0.411900166 | 0.460786156 |
| Q8VCG3 | Wdr74 | WD repeat-containing protein 74 | 57.79 | 3.39 | 42636 | 1 | 815712.62 | 722700.75 | 2 | 613035.06 | 2 | 2 | 676947.38 | 2 | 455087.91 | 475166.91 | 2 | 488489.03 | 533596.06 | 2 | 2 | 2 | 1022006.06 | 2 | 698864.62 | 2 | 7 | 11 | 11 | 12 | 11 | 12 | 0.475070271 | 0.635942725 | 0.799867409 | 1.070725511 |
| Q3U821 | Wdr75 | WD repeat-containing protein 75 | 265.82 | 6.51 | 94037 | 4 | 2768597.25 | 1758667 | 1274314 | 1565066 | 1187015.88 | 998214.5 | 1436145 | 2264347.75 | 2341571.25 | 1234738.25 | 2742927.5 | 2991042.25 | 2621062 | 2684044.5 | 2540631.5 | 2335289.75 | 1267917.5 | 1375730 | 2980809 | 1098942.88 | 4 | 12 | 0 | 11 | 4 | 12 | 1.587590288 | 1.641048974 | 1.059042444 | 1.094703419 |
| Q8BFQ4 | Wdr82 | WD repeat-containing protein 82 | 139.12 | 10.22 | 35079 | 2 | 662135.44 | 1073867.5 | 1630959.62 | 630871.12 | 136562.7 | 380189.5 | 707156.88 | 946699.62 | 986236.88 | 752570.81 | 437256.69 | 806279.62 | 858754.81 | 2 | 1622483.75 | 828897.62 | 2 | 967650 | 2 | 496281.25 | 11 | 7 | 12 | 8 | 12 | 8 | 0.900923987 | 0.987257173 | 0.554574999 | 0.607718469 |
| B2RYI0 | Wdr91 | WD repeat-containing protein 91 | 105.37 | 3.08 | 83146 | 2 | 2 | 2 | 2 | 56281.43 | 2 | 2 | 2 | 347050.97 | 225744.78 | 2 | 390548.62 | 294252.31 | 2 | 2 | 2 | 2 | 2 | 390875.38 | 261046.23 | 2 | 9 | 9 | 11.5 | 11.5 | 11.5 | 11.5 | 12.16581745 | 1.195539172 | 11.58170566 | 1.1381383 |
| Q8BGF3 | Wdr92 | WD repeat-containing protein 92 | 47.07 | 3.36 | 39790 | 1 | 519696.75 | 1120434.88 | 2 | 857835.88 | 642267.44 | 1162377.5 | 925191.62 | 686994.69 | 553839.69 | 826790.94 | 917265.31 | 766143.94 | 370917.09 | 808306.38 | 663710 | 378837.19 | 1044402.06 | 896505.06 | 2 | 874553.44 | 11 | 11.5 | 8 | 10 | 11 | 11.5 | 1.122954343 | 0.848658895 | 1.017216153 | 0.768748562 |
| Q6DID7 | Wls | Protein wntless homolog | 137.34 | 5.36 | 62188 | 3 | 1521271.25 | 1041345.69 | 2173184.5 | 2269878.5 | 2298018 | 2850203 | 2633320.5 | 2671636.5 | 2565973.25 | 3009524 | 3656848 | 3487582.5 | 4777241.5 | 3867260.5 | 4124858 | 2482199.5 | 1893867.75 | 2426248 | 2567370.5 | 2613731 | 0 | 3 | 0 | 2 | 0 | 3 | 2.140416706 | 1.450315898 | 1.28802728 | 0.872748954 |
| P83741 | Wnk1 | Serine/threonine-protein kinase WNK1 | 498.43 | 3.91 | 250934 | 10 | 13381832 | 10494573 | 11477634 | 23655360 | 8640574 | 10878378 | 11813146 | 10369643 | 8441893 | 21715466 | 4353864.5 | 6101208.5 | 16626742 | 18580564 | 9350636 | 8384629.5 | 12718117 | 7553896 | 20015170 | 17210682 | 9 | 11 | 9 | 12 | 9 | 12 | 0.813200842 | 0.870204013 | 0.973873182 | 1.042139048 |
| Q9ER69 | Wtap | Pre-mRNA-splicing regulator WTAP | 56.14 | 3.03 | 44177 | 1 | 500269.69 | 2 | 421669.41 | 514523.5 | 2 | 536073.12 | 446603.06 | 880094.44 | 2 | 463258.31 | 2 | 300681.41 | 2 | 2 | 281285.94 | 486463.91 | 540731.06 | 359591.06 | 440623.62 | 562233.38 | 7 | 7 | 3.5 | 12 | 7 | 12 | 0.405142278 | 0.250200177 | 1.663556278 | 1.027347917 |
| Q91WL8 | Wwox | WW domain-containing oxidoreductase | 83.54 | 4.59 | 46512 | 2 | 2943334.75 | 3098931.5 | 3332114 | 3007047.5 | 2402386.75 | 3079514.25 | 2890151.75 | 2751383.5 | 2308709.25 | 3257819.5 | 4760212 | 3974356 | 5784928.5 | 4524703.5 | 4559092 | 2403216.5 | 2468684.25 | 4502699 | 1597683.5 | 2561968 | 0 | 8 | 0 | 8 | 0 | 8 | 1.596563052 | 1.652014889 | 0.915477616 | 0.94727399 |
| Q9DBH0 | Wwp2 | NEDD4-like E3 ubiquitin-protein ligase WWP2 | 163.64 | 4.83 | 98761 | 3 | 56324.41 | 129730.71 | 102532.84 | 2 | 2 | 116411.34 | 119961.66 | 76825.57 | 29113.76 | 55735.29 | 185378.8 | 257702.28 | 171020.06 | 235191.47 | 2 | 2 | 2 | 98891.66 | 105236.38 | 114567.57 | 4 | 12 | 5 | 9 | 5 | 12 | 2.942890751 | 2.133650768 | 1.104326018 | 0.800656992 |
| Q00519 | Xdh | Xanthine dehydrogenase/oxidase | 771.4 | 13.18 | 146562 | 14 | 20120926 | 20496500 | 13793951 | 17366834 | 14716912 | 13916289 | 20705110 | 15643754 | 16614903 | 14312318 | 4170935 | 8116236.5 | 7206664 | 7903994 | 6768019 | 20349710 | 15579121 | 14499611 | 15275404 | 16632358 | 0 | 11 | 0 | 11 | 0 | 11 | 0.395003178 | 0.420801201 | 0.951917301 | 1.014087899 |
| Q6P1B1 | Xpnpep1 | Xaa-Pro aminopeptidase 1 | 888.17 | 29.53 | 69591 | 16 | 2 | 94193.61 | 2 | 239912.41 | 2 | 130478.02 | 2 | 89831.88 | 2 | 66216.48 | 441961.19 | 2 | 2 | 307072.03 | 2 | 2 | 649446.56 | 71090.1 | 127969.69 | 74393.32 | 10.5 | 8.5 | 12 | 9 | 12 | 9 | 2.241880493 | 2.614170337 | 2.762252223 | 3.220955733 |
| B7ZMP1 | Xpnpep3 | Xaa-Pro aminopeptidase 3 | 91.46 | 3.36 | 56677 | 1 | 506093.94 | 501078.78 | 525338.5 | 307813.22 | 650968.12 | 446897.19 | 490146.38 | 462616.5 | 765919.44 | 386609.25 | 304894.16 | 386890.34 | 375864.66 | 2 | 375027.09 | 494837 | 307061.31 | 427982.81 | 2 | 408060.94 | 3 | 3 | 1 | 6 | 3 | 6 | 0.57908825 | 0.565270983 | 0.657467568 | 0.64178014 |
| Q6P5F9 | Xpo1 | Exportin-1 | 1059.78 | 21.48 | 123093 | 18 | 18333160 | 13175708 | 15895322 | 17035398 | 20464702 | 17864674 | 17573544 | 18227954 | 20810922 | 13394997 | 11056813 | 14376917 | 17233248 | 12332939 | 12570035 | 19184680 | 15051578 | 15405900 | 14956071 | 12941478 | 4 | 7 | 2 | 7 | 4 | 7 | 0.795836724 | 0.768958053 | 0.913260178 | 0.882415635 |
| Q924C1 | Xpo5 | Exportin-5 | 373.65 | 6.81 | 136973 | 6 | 2815128 | 3312891.5 | 2073454.5 | 2033927.12 | 3063127.5 | 1439362 | 3345498 | 1888996.62 | 2494954 | 2242880 | 3102705.25 | 4702594 | 3782850.25 | 971250.5 | 3483782.5 | 3087354 | 1415715.62 | 1620227.75 | 2100270 | 2107605 | 6 | 8 | 6 | 9 | 6 | 9 | 1.206387786 | 1.405855016 | 0.776865822 | 0.905314796 |
| Q60596 | Xrcc1 | DNA repair protein XRCC1 | 85.16 | 2.69 | 68971 | 1 | 365807.78 | 349284.34 | 299007.28 | 351177.31 | 354196.44 | 314226.19 | 270287.41 | 2 | 389933.09 | 2 | 161924.39 | 247665.48 | 317532.53 | 188602.8 | 342435.56 | 266206.22 | 62653.34 | 258445.94 | 357933.88 | 399740.47 | 2 | 9 | 11 | 10 | 11 | 10 | 0.731712944 | 1.291148719 | 0.782204625 | 1.380244135 |
| P27641 | Xrcc5 | X-ray repair cross-complementing protein 5 | 90.53 | 2.6 | 83057 | 2 | 1368449 | 1168766.75 | 1096088.25 | 773614.12 | 1909140 | 977823.44 | 1554098.25 | 1568093 | 1653722.25 | 1293487.12 | 821357.5 | 1505811.75 | 794044.69 | 972466.56 | 1121423.38 | 1136115.88 | 422790.44 | 993479.62 | 811577.81 | 1025208.62 | 9 | 5 | 3 | 3 | 9 | 5 | 0.82568966 | 0.740022431 | 0.694922733 | 0.622822878 |
| P23475 | Xrcc6 | X-ray repair cross-complementing protein 6 | 98.86 | 4.77 | 69484 | 2 | 881231.81 | 879788.25 | 718916.75 | 758275.44 | 806272.56 | 1466116.38 | 968649.88 | 2 | 686150.31 | 423276.5 | 1529740.38 | 1435201.5 | 1084767.5 | 654500.44 | 1144450 | 896176.75 | 811219.5 | 814933.31 | 955044.62 | 377755.25 | 5 | 9 | 6 | 12 | 6 | 12 | 1.446082776 | 1.650208215 | 0.953181829 | 1.0877306 |
| Q9DBR1 | Xrn2 | 5'-3' exoribonuclease 2 | 888.82 | 20.5 | 108687 | 16 | 7825260.5 | 8871738 | 7823233 | 4782985.5 | 8312840 | 4509506 | 7564678 | 3114902.75 | 5817656.5 | 3995659.5 | 7136970.5 | 7789142.5 | 5075806.5 | 4034270.75 | 2342429.5 | 7105889 | 5898220.5 | 4634929.5 | 5028446 | 5176130.5 | 3 | 4 | 11 | 8 | 11 | 8 | 0.701259564 | 1.05504339 | 0.740205586 | 1.113637588 |
| Q91WQ3 | Yars | Tyrosine--tRNA ligase, cytoplasmic | 718.91 | 28.41 | 59105 | 14 | 20234012 | 20456820 | 16717416 | 14047377 | 17313632 | 16761284 | 19375068 | 18257526 | 18334892 | 16982260 | 16422093 | 22274502 | 14412419 | 13971934 | 14856322 | 21574020 | 11856518 | 16924536 | 12308977 | 16148334 | 8 | 8 | 5 | 6 | 8 | 8 | 0.923036564 | 0.913346664 | 0.887834231 | 0.87851388 |
| Q8BYL4 | Yars2 | Tyrosine--tRNA ligase, mitochondrial | 67.19 | 2.54 | 52597 | 1 | 292642.97 | 398422.78 | 383249.75 | 317815.78 | 373712.19 | 423366.12 | 319955.69 | 349734.22 | 416304.97 | 233900.98 | 422311 | 496498.62 | 632156.81 | 578202.81 | 555907.88 | 362938.44 | 391451.41 | 271406.25 | 2 | 347446.19 | 0 | 8 | 1 | 9 | 1 | 9 | 1.520563496 | 1.540260242 | 0.777670452 | 0.787744072 |
| P62960 | Ybx1 | Nuclease-sensitive element-binding protein 1 | 759.75 | 49.07 | 35730 | 9 | 11491526 | 13630891 | 9594266 | 8267771 | 7404750.5 | 12728250 | 13867912 | 10636318 | 9506304 | 11067551 | 14220984 | 14111705 | 9123450 | 7953942 | 8604576 | 11087734 | 13934378 | 12063640 | 10958790 | 10740578 | 10 | 7 | 10 | 10 | 10 | 10 | 1.071948993 | 0.934407224 | 1.166621315 | 1.016932141 |
| Q9JKB3 | Ybx3 | Y-box-binding protein 3 | 449.34 | 26.87 | 38814 | 6 | 1746590.5 | 3383782.5 | 1050931.5 | 634122.56 | 909462.75 | 3301444.25 | 2318392 | 2439463.5 | 1531160.12 | 2356911.25 | 4703031 | 5050664 | 2731809.25 | 2417936.5 | 1922235.5 | 2277510.25 | 2706880.5 | 2579646.75 | 2454643.75 | 2053367.62 | 3 | 5 | 7 | 11 | 7 | 11 | 2.178112137 | 1.408316196 | 1.562747064 | 1.01043558 |
| Q6PEC3 | Yif1b | Protein YIF1B | 77.34 | 5.02 | 28418 | 1 | 2 | 2 | 2 | 2 | 2 | 2 | 2 | 2 | 2 | 2 | 1490649.12 | 947074.12 | 1220666.75 | 1555737.12 | 1131963.12 | 2 | 2 | 514099.47 | 468217.5 | 370149.06 | 0 | 5 | 0 | 5 | 0 | 5 | 634609.023 | 634609.023 | 135247.003 | 135247.003 |
| Q3UDR8 | Yipf3 | Protein YIPF3 | 41.55 | 6.63 | 37998 | 1 | 235145.56 | 256590.23 | 154755.05 | 148951.34 | 206023.95 | 2 | 175691.48 | 394423.16 | 517126.75 | 374200.06 | 1102692.12 | 565297.31 | 1522148.88 | 1258537.75 | 1425271.25 | 265192.12 | 410385.62 | 743035.5 | 1005383.94 | 418522.5 | 0 | 0 | 0 | 5 | 0 | 5 | 5.865347947 | 4.019277865 | 2.838358278 | 1.945008327 |
| Q5M7T4 | Yipf4 | Protein YIPF4 | 41.37 | 4.47 | 27283 | 1 | 636903.94 | 469120.12 | 493692.66 | 645758.81 | 774972.25 | 879537.31 | 977600.81 | 973376.94 | 895919.88 | 1176813.75 | 933535.88 | 1097438 | 1794655.75 | 2 | 2 | 1286831 | 2 | 1193342.25 | 1255987.75 | 1834436.62 | 10 | 5 | 11 | 5 | 11 | 5 | 1.266578305 | 0.78022427 | 1.844295954 | 1.136103831 |
| Q5EGY4 | Ykt6 | Synaptobrevin homolog YKT6 | 276.08 | 27.27 | 22369 | 5 | 4953738 | 4221914 | 3031028.75 | 1538098.5 | 5965042 | 4491326.5 | 4073466.5 | 3293818 | 5710508 | 2849113.5 | 2250526.5 | 2582040.5 | 729111.5 | 655605 | 2165103.5 | 5418693 | 4360353 | 2210635.25 | 3304922 | 3302746.5 | 3 | 12 | 0 | 11 | 3 | 12 | 0.425289854 | 0.410534408 | 0.943557504 | 0.910820746 |
| O88967 | Yme1l1 | ATP-dependent zinc metalloprotease YME1L1 | 111.82 | 4.62 | 80028 | 2 | 2 | 178797.89 | 168358.17 | 2 | 2 | 2 | 2 | 2 | 2 | 2 | 2 | 2 | 439501.78 | 2 | 2 | 2 | 2 | 2 | 2 | 2 | 11 | 7.5 | 10 | 12.5 | 11 | 12.5 | 1.266007524 | 43950.978 | 2.880499E-05 | 1 |
| E9Q5K9 | Ythdc1 | YTH domain-containing protein 1 | 50.08 | 1.22 | 85649 | 1 | 2 | 2 | 2 | 2 | 2 | 2 | 2 | 2 | 2 | 2 | 2 | 731972.56 | 203479.11 | 2 | 2 | 2 | 2 | 2 | 2 | 2 | 7.5 | 12.5 | 7.5 | 12.5 | 7.5 | 12.5 | 93545.767 | 93545.767 | 1 | 1 |

| | | | | | | | | | | | | | | |
|---|---|---|---|---|---|---|---|---|---|---|---|---|---|---|
| Q91YT7 | Ythdf2 | YTH domain-containing family protein 2 | 157.54 | 6.22 | 62280 | 3 | 492057.16 | 624506.31 | 552024.44 | 2 | 416995.41 | 521541.94 | 544007.94 | 388009.5 |
| Q8BYK6 | Ythdf3 | YTH domain-containing family protein 3 | 275.61 | 9.4 | 63961 | 4 | 1296984.5 | 742622.94 | 1412299.5 | 827178.81 | 572006.31 | 1721977.62 | 1121235.62 | 542118.81 |
| Q9CQV8 | Ywhab | 14-3-3 protein beta/alpha | 998 | 69.11 | 28086 | 17 | 13124176 | 13731282 | 13586167 | 14715585 | 12299983 | 11910798 | 10042159 | 10495820 |
| P62259 | Ywhae | 14-3-3 protein epsilon | 1503.85 | 70.59 | 29174 | 24 | 161167762 | 150037426 | 183540978 | 168920994 | 197059554 | 131969858 | 149601826 | 167928322 |
| P61982 | Ywhag | 14-3-3 protein gamma | 965.61 | 59.51 | 28303 | 16 | 65808054 | 70518098 | 63453730 | 73393930 | 70788194 | 64014962 | 63607302 | 66275114 |
| P68510 | Ywhah | 14-3-3 protein eta | 1175.95 | 58.94 | 28212 | 20 | 77621730 | 82781970 | 98546138 | 94003298 | 118651634 | 64174706 | 89997306 | 98789842 |
| P68254 | Ywhaq | 14-3-3 protein theta | 1277.78 | 62.86 | 27778 | 20 | 30573510 | 38028930 | 31267838 | 29141440 | 38228498 | 24833428 | 31985984 | 34165622 |
| P63101 | Ywhaz | 14-3-3 protein zeta/delta | 1197.55 | 74.29 | 27771 | 20 | 119783570 | 196716274 | 205914594 | 206822978 | 251440898 | 161605330 | 165403474 | 197796786 |
| Q6NZF1 | Zc3h11a | Zinc finger CCCH domain-containing protein 11A | 181.23 | 5.81 | 86492 | 4 | 1197310.62 | 1712958.75 | 2 | 818086.69 | 1034430.75 | 1869358.75 | 757492 | 1484297.5 |
| Q8BJ05 | Zc3h14 | Zinc finger CCCH domain-containing protein 14 | 67.74 | 1.9 | 82409 | 1 | 310876.94 | 2 | 2 | 2 | 2 | 361339.53 | 2 | 321011.28 |
| Q3TIV5 | Zc3h15 | Zinc finger CCCH domain-containing protein 15 | 124.96 | 6.34 | 48327 | 2 | 2 | 146090.77 | 227515.52 | 2 | 2 | 2 | 2 | 83032.48 |
| Q6ZPZ3 | Zc3h4 | Zinc finger CCCH domain-containing protein 4 | 40.69 | 1 | 140967 | 1 | 303405 | 277803.75 | 325365.66 | 297280.69 | 226710.42 | 339579.47 | 350608.06 | 2 |
| Q3UPF5 | Zc3hav1 | Zinc finger CCCH-type antiviral protein 1 | 104.86 | 2.96 | 106688 | 2 | 2 | 2 | 587612.5 | 2 | 673782.5 | 1516664.38 | 435937.44 | 2 |
| Q9CYA6 | Zcchc8 | Zinc finger CCHC domain-containing protein 8 | 114.84 | 5.08 | 78026 | 2 | 862905.06 | 889528.19 | 894699.88 | 691171.38 | 809333.81 | 921941.69 | 1260015.25 | 757541.12 |
| Q9D270 | Zdhhc21 | Probable palmitoyltransferase ZDHHC21 | 88.5 | 6.04 | 31331 | 2 | 171134.08 | 183658.73 | 301324.25 | 320345.16 | 186390.03 | 204552.97 | 180098.97 | 433269.47 |
| Q9R0G7 | Zeb2 | Zinc finger E-box-binding homeobox 2 | 47.18 | 0.91 | 136615 | 1 | 2 | 269654.88 | 244839.36 | 2 | 205720 | 2 | 200271.55 | 468549.19 |
| Q91X58 | Zfand2b | AN1-type zinc finger protein 2B | 83.73 | 8.56 | 27895 | 2 | 720935.12 | 655820.38 | 239000.3 | 856556.12 | 965126 | 279772.44 | 338141 | 770406.25 |
| Q9DB43 | Zfpl1 | Zinc finger protein-like 1 | 57.9 | 3.87 | 34155 | 1 | 560931.56 | 312382 | 388487.09 | 429818.97 | 413123.25 | 440843.41 | 517347.25 | 353770.22 |
| O88532 | Zfr | Zinc finger RNA-binding protein | 254.3 | 4.28 | 116859 | 3 | 1293851.62 | 1960891.75 | 1779541.38 | 1755371.12 | 1513254.5 | 2050407.62 | 2028313.12 | 899249.25 |
| Q3TXX3 | Zfyve27 | Protrudin | 97.24 | 3.37 | 46202 | 1 | 264804.31 | 2 | 197241.95 | 183435 | 292416.75 | 155971.44 | 287409.94 | 2 |
| Q80W54 | Zmpste24 | CAAX prenyl protease 1 homolog | 228.03 | 13.68 | 54735 | 5 | 3193989.25 | 2831700.25 | 2784568.75 | 4624603.5 | 2780870.75 | 3525645 | 3710543.25 | 4315759 |
| Q9JMD0 | Znf207 | BUB3-interacting and GLEBS motif-containing protein ZNF207 | 82.77 | 5.05 | 52793 | 2 | 2 | 216012.55 | 2 | 2 | 2 | 2 | 2 | 521500.75 |
| O88291 | Znf326 | DBIRD complex subunit ZNF326 | 76.2 | 2.76 | 65225 | 1 | 271651.12 | 306350.69 | 230680.86 | 2 | 187647.61 | 2 | 448959.56 | 297416.84 |
| Q8R151 | Znfx1 | NFX1-type zinc finger-containing protein 1 | 47.7 | 0.52 | 218828 | 1 | 2 | 252048.17 | 2 | 2 | 818815 | 2 | 363969.28 | 265403.41 |
| O35986 | Zranb2 | Zinc finger Ran-binding domain-containing protein 2 | 133.33 | 8.18 | 37350 | 3 | 5654370 | 6570487.5 | 6087805 | 7326659 | 6541948.5 | 5675426.5 | 4949323.5 | 5546762 |
| O54692 | Zw10 | Centromere/kinetochore protein zw10 homolog | 71.21 | 2.18 | 88063 | 2 | 576424 | 424001.03 | 367169.28 | 2 | 849967.12 | 271336.72 | 457555.56 | 597282 |
| Q8R060 | Zwilch | Protein zwilch homolog | 75.85 | 1.7 | 66839 | 1 | 370313.25 | 306428.97 | 378426.72 | 250683.61 | 499696.19 | 152003.41 | 421387 | 2 |
| Q62523 | Zyx | Zyxin | 282.75 | 13.3 | 60546 | 6 | 6705190 | 8314274 | 8944082 | 10806205 | 8022379.5 | 9275074 | 6780046.5 | 9589633 |
| Q5U2Q3 | | Ester hydrolase C11orf54 homolog | 140.75 | 7.62 | 34993 | 3 | 3892808.5 | 4236515.5 | 4328884.5 | 6043029 | 4995700.5 | 4484480 | 3440668.5 | 2235540 |
| Q6PIU9 | | Uncharacterized protein FLJ45252 homolog | 66.76 | 3.39 | 37705 | 1 | 1244333.12 | 1360104.5 | 1490624.12 | 1551362 | 1096650 | 1212925 | 1391489 | 1482986.62 |
| P10404 | | MLV-related proviral Env polyprotein | 181.57 | 3.43 | 69613 | 3 | 1736666.38 | 2479842 | 2844977.5 | 2408473.5 | 3305649.5 | 1980012.12 | 2119979.5 | 2445384 |
| Q99M08 | | Uncharacterized protein C4orf3 homolog | 99.56 | 21.54 | 7404 | 1 | 489674.47 | 456930.31 | 2 | 680351.75 | 290150.22 | 535083.19 | 457029.41 | 605096.88 |
| Q5XI83 | | UPF0505 protein C16orf62 homolog | 140.47 | 4.49 | 106151 | 3 | 192197.03 | 2 | 466330.81 | 178844.89 | 1569628.75 | 2 | 174074.81 | 313374.53 |

| | | | | | | | | | | | | |
|---|---|---|---|---|---|---|---|---|---|---|---|---|
| Q91YT7 | 2 | 78015.83 | 565409.69 | 760847.5 | 187205.22 | 461315.5 | 2 | 580881.5 | 518519.56 | 363314 | 469503.22 | 313484.78 |
| Q8BYK6 | 1207628.25 | 2156863 | 807113.75 | 1941835.38 | 1281636.5 | 1939145.62 | 1686723.62 | 707515.5 | 1498014.12 | 879894.19 | 1830610.75 | 1757883.75 |
| Q9CQV8 | 9924122 | 10697516 | 13562094 | 12299544 | 10827705 | 12269950 | 12172022 | 13602864 | 12572039 | 12881728 | 11091309 | 13087585 |
| P62259 | 165367250 | 164627426 | 100901802 | 110061482 | 95774914 | 90161330 | 107176594 | 168867858 | 160980210 | 165289586 | 185618498 | 158997074 |
| P61982 | 65476766 | 66460934 | 43439574 | 53476046 | 56987190 | 59511494 | 57989378 | 72701082 | 63393538 | 72227578 | 76992922 | 81781682 |
| P68510 | 109820050 | 97920210 | 31085374 | 53592170 | 50225894 | 42203482 | 54093454 | 80189410 | 46278910 | 76349874 | 79044778 | 97991506 |
| P68254 | 50971466 | 25445520 | 17479840 | 20284504 | 15816199 | 13867849 | 15501930 | 26600342 | 17400452 | 20782536 | 19886744 | 22642332 |
| P63101 | 228187442 | 193380546 | 109198674 | 154572290 | 149939138 | 122986714 | 155438994 | 169852354 | 106413714 | 187680610 | 202101154 | 216874402 |
| Q6NZF1 | 2 | 658897.81 | 1044243.25 | 999779.69 | 649011.94 | 1029868.38 | 803159.56 | 1093351 | 1114947 | 1451074 | 1489881.25 | 1431198.62 |
| Q8BJ05 | 315553.75 | 476323.91 | 2 | 2 | 276148.72 | 2 | 2 | 436114.5 | 461766.94 | 2 | 366511.06 | 519009.94 |
| Q3TIV5 | 132803.47 | 227055.36 | 319012.25 | 297827.34 | 327328.72 | 294162.66 | 368993.19 | 2 | 2 | 55652.3 | 2 | 118858.49 |
| Q6ZPZ3 | 2 | 2 | 231118.36 | 234529.77 | 2 | 2 | 2 | 170385.08 | 2 | 2 | 204557.02 | 259448.2 |
| Q3UPF5 | 502489.78 | 2 | 2 | 1344647.62 | 1204632 | 1523418.88 | 1176842 | 2 | 674534.88 | 2 | 2 | 935434 |
| Q9CYA6 | 1049101.88 | 648034 | 173342.89 | 424998.12 | 361856.53 | 394171.31 | 257171.66 | 1176601.12 | 725557.19 | 816331.06 | 743430.62 | 776412.12 |
| Q9D270 | 295106.06 | 361936.34 | 749243.38 | 651979.19 | 1083678.62 | 1446460 | 1089062.25 | 148355.22 | 157868.84 | 596876.38 | 496641.28 | 488547.06 |
| Q9R0G7 | 225480.72 | 2 | 2 | 261800.83 | 231442.17 | 2 | 287445.41 | 2 | 2 | 2 | 2 | 252100 |
| Q91X58 | 631804.62 | 433814.12 | 92448.66 | 202936.75 | 103927.91 | 274257.38 | 117574.75 | 205704.64 | 2 | 839195.25 | 918703.06 | 191608.36 |
| Q9DB43 | 367739.25 | 457649.41 | 322634.47 | 562387.5 | 656652 | 413394.69 | 522226.72 | 569200.62 | 412315.75 | 582267.81 | 146506.75 | 376831.91 |
| O88532 | 1287674 | 1339879 | 1420984 | 1824267 | 279122.47 | 895833.31 | 1523377.38 | 2457557.5 | 1740226.5 | 798371.56 | 1232658 | 1582066.5 |
| Q3TXX3 | 268714.38 | 122031.69 | 86018.71 | 159169.2 | 2 | 2 | 141837.02 | 192237.45 | 2 | 2 | 2 | 203384.66 |
| Q80W54 | 3886651.25 | 4662634.5 | 5134520.5 | 4776145 | 5731597.5 | 6055413 | 6247878.5 | 3827867.75 | 4479173.5 | 5618494 | 4844236 | 4593169.5 |
| Q9JMD0 | 254913.3 | 448598.19 | 884679.69 | 2 | 616792.25 | 315869.75 | 588370.62 | 2 | 2 | 432687.56 | 474022.31 | 2 |
| O88291 | 209302.72 | 262719.5 | 2 | 253433.56 | 158557.39 | 2 | 2 | 208729.25 | 374094.75 | 2 | 2 | 296348.53 |
| Q8R151 | 844248.12 | 2 | 2 | 2 | 2 | 2 | 2 | 2 | 2 | 2 | 2 | 2 |
| O35986 | 5706232 | 4704939 | 2412629.25 | 2120365.5 | 1935293.25 | 2163655 | 513532.25 | 5631302 | 5246629 | 5557038 | 5967596 | 6137981 |
| O54692 | 735908.44 | 2 | 2 | 337072.94 | 2 | 2 | 2 | 818926.62 | 419542.78 | 2 | 187726.72 | 2 |
| Q8R060 | 2 | 185397.58 | 2 | 257183.2 | 186754.64 | 2 | 2 | 313688.09 | 153107.86 | 124922.35 | 123322.15 | 2 |
| Q62523 | 8567889 | 11262406 | 7275480.5 | 6931583 | 8356129.5 | 11705391 | 8619541 | 6911272.5 | 7875665.5 | 11651428 | 11844924 | 11419700 |
| Q5U2Q3 | 4208480 | 4595630 | 2049414.25 | 2056728 | 2055007.75 | 3259545 | 1439517.88 | 4825769 | 5777268 | 4943061.5 | 5888461 | 5392283 |
| Q6PIU9 | 1154820.25 | 1607568.38 | 710716.06 | 701748.06 | 737789.44 | 1112046.12 | 982075.81 | 1286719.38 | 1339875.12 | 1998855.88 | 2306265.75 | 1731470.75 |
| P10404 | 2694892 | 2655380 | 2447336.5 | 3195508.25 | 3503414 | 1943056 | 3789582 | 1875327.75 | 1445945.5 | 2133931.75 | 1683757 | 1725597.25 |
| Q99M08 | 2 | 602255.25 | 2 | 544195.5 | 339087.19 | 599295.5 | 531055.62 | 622297.62 | 816952.38 | 535370.25 | 671375.94 | 468481.41 |
| Q5XI83 | 1019761.06 | 2 | 176952.97 | 368024.25 | 246754.78 | 109047.62 | 206665.55 | 2 | 2 | 205764 | 2 | 2 |

| | | | | | | | | | | |
|---|---|---|---|---|---|---|---|---|---|---|
| Q91YT7 | 12.5 | 11 | 9.5 | 10 | 12.5 | 11 | 0.946870833 | 1.289376662 | 1.076773527 | 1.466268266 |
| Q8BYK6 | 5 | 6 | 10 | 12 | 10 | 12 | 1.578295109 | 1.134319304 | 1.375755856 | 0.988754522 |
| Q9CQV8 | 2 | 6 | 1 | 1 | 2 | 6 | 0.906223818 | 1.151890653 | 0.937417082 | 1.191540051 |
| P62259 | 0 | 10 | 0 | 8 | 0 | 10 | 0.58564015 | 0.64667038 | 0.975632814 | 1.077304625 |
| P61982 | 0 | 7 | 0 | 5 | 0 | 7 | 0.789051341 | 0.832948017 | 1.067259743 | 1.126633769 |
| P68510 | 0 | 5 | 0 | 6 | 0 | 6 | 0.490241806 | 0.501843527 | 0.805450882 | 0.824512123 |
| P68254 | 0 | 0 | 0 | 2 | 0 | 2 | 0.495995066 | 0.495515657 | 0.641666272 | 0.641046064 |
| P63101 | 4 | 8 | 0 | 12 | 4 | 12 | 0.705772525 | 0.731355805 | 0.900317894 | 0.932953175 |
| Q6NZF1 | 9 | 7 | 12 | 9 | 12 | 9 | 0.95029677 | 0.94885057 | 1.381638391 | 1.379535759 |
| Q8BJ05 | 12 | 3 | 3 | 7.5 | 12 | 7.5 | 0.88829237 | 0.187322624 | 5.736541757 | 1.209718885 |
| Q3TIV5 | 0 | 10.5 | 0 | 8 | 0 | 10.5 | 4.302118006 | 3.629128879 | 0.467106663 | 0.394036211 |
| Q6ZPZ3 | 2 | 1 | 10.5 | 12 | 10.5 | 12 | 0.325503532 | 0.674671827 | 0.443457004 | 0.919154226 |
| Q3UPF5 | 3.5 | 10.5 | 7 | 11 | 7 | 11 | 4.161676184 | 2.138223253 | 1.276338674 | 0.655768712 |
| Q9CYA6 | 0 | 10 | 0 | 10 | 0 | 10 | 0.388544127 | 0.347566905 | 1.021866369 | 0.91409677 |
| Q9D270 | 0 | 10 | 0 | 10 | 0 | 10 | 4.31733562 | 3.403760422 | 1.623842393 | 1.280227194 |
| Q9R0G7 | 11 | 8 | 11 | 8 | 11 | 8 | 1.083966452 | 0.872959458 | 0.350043898 | 0.28190368 |
| Q91X58 | 1 | 7 | 0 | 10 | 1 | 10 | 0.23015556 | 0.322398248 | 0.626982468 | 0.878267068 |
| Q9DB43 | 7 | 12 | 8 | 11 | 8 | 12 | 1.177006187 | 1.159050185 | 0.991628417 | 0.976500475 |
| O88532 | 7 | 9 | 9 | 12 | 9 | 12 | 0.715843469 | 0.781482637 | 0.940740019 | 1.027001045 |
| Q3TXX3 | 4 | 6.5 | 7 | 7.5 | 7 | 7.5 | 0.412654788 | 0.463991446 | 0.421823335 | 0.474300614 |
| Q80W54 | 0 | 3 | 0 | 5 | 0 | 5 | 1.723360601 | 1.390240813 | 1.440757656 | 1.162264064 |
| Q9JMD0 | 3 | 9 | 6 | 10 | 6 | 10 | 11.13650674 | 1.963822382 | 4.197359325 | 0.740166408 |
| O88291 | 5.5 | 12 | 4.5 | 9 | 5.5 | 12 | 0.413513602 | 0.338145716 | 0.882412974 | 0.721582471 |
| Q8R151 | 7.5 | 7.5 | 5 | 5 | 7.5 | 7.5 | 9.338209E-06 | 6.785988E-06 | 9.338209E-06 | 6.785988E-06 |
| O35986 | 0 | 3 | 0 | 7 | 0 | 7 | 0.284186275 | 0.344038833 | 0.886868231 | 1.073651821 |
| O54692 | 3 | 8 | 4 | 9 | 4 | 9 | 0.152005095 | 0.163466097 | 0.643138366 | 0.691630226 |
| Q8R060 | 1 | 2 | 11 | 12 | 11 | 12 | 0.245877516 | 0.585066587 | 0.396025006 | 0.942343171 |
| Q62523 | 12 | 8 | 10 | 8 | 12 | 8 | 1.002243275 | 0.943113343 | 1.161498374 | 1.092972776 |
| Q5U2Q3 | 0 | 7 | 1 | 0 | 1 | 7 | 0.462196942 | 0.572651108 | 1.141716529 | 1.414559849 |
| Q6PIU9 | 1 | 6 | 0 | 6 | 1 | 6 | 0.629442248 | 0.619635924 | 1.284753395 | 1.264737726 |
| P10404 | 8 | 2 | 7 | 2 | 8 | 2 | 1.164633082 | 1.250784928 | 0.693865892 | 0.745193497 |
| Q99M08 | 10.5 | 5 | 9.5 | 5 | 10.5 | 5 | 1.050350331 | 0.91551092 | 1.624570124 | 1.416014872 |
| Q5XI83 | 11 | 5 | 10 | 7 | 11 | 7 | 0.460092883 | 0.734762865 | 0.085488867 | 0.136524704 |
