## Supplemental Tables S2 to S5 for "Beyond the ink: cellular and molecular effects of iron-based pigments on macrophages"

| accession | abund.PR101#1 | abund.PR101#2 | abund.PR101#3 | abund.PR101#4 | abund.PR101#5 | abund.PBk11 #1 | abund.PBk11 #2 | abund.PBk11 #3 | abund.PBk11 #4 | abund.PBk11 #5 | U ctrl vs PR101 | U ctrl vs PBK11 | U ERM vs PR101 | U ERM vs PBK11 | Max U PR101 | Max U PBk11 | ratio PR101 Ctrl | ratio PR101 ERM | ratio PBk11 Ctrl | ratio PBK11 ERM |
|---|---|---|---|---|---|---|---|---|---|---|---|---|---|---|---|---|---|---|---|---|
| P14046 | 804531.81 | 796653.44 | 1528199.62 | 1643296 | 1381671.75 | 2 | 2 | 339412.78 | 583525.38 | 354524.78 | 0 | 8 | 0 | 10 | 0 | 10 | 7.65 | 3.71 | 1.59 | 0.77 |
| Q8BGQ7 | 14376877 | 14293355 | 10615027 | 10255833 | 10941392 | 8147467.5 | 7186804 | 7815803 | 6506542.5 | 4954248 | 0 | 11 | 0 | 9 | 0 | 11 | 1.79 | 1.84 | 1.02 | 1.05 |
| P55096 | 1356213.5 | 1148035.38 | 1504144.5 | 1173876.12 | 1367783.25 | 364442.75 | 2 | 978311.38 | 2 | 646735.25 | 0 | 11 | 0 | 12 | 0 | 12 | 4.04 | 3.07 | 1.23 | 0.93 |
| P61222 | 34438630 | 33029642 | 32882174 | 28954910 | 32443174 | 26514610 | 23552572 | 27166056 | 25107636 | 26451888 | 0 | 6 | 0 | 5 | 0 | 6 | 1.33 | 1.37 | 1.06 | 1.09 |
| Q6P542 | 9530251 | 8395944 | 7432936 | 7366759.5 | 8117239.5 | 5481887.5 | 5415308 | 6962911 | 5784079 | 4388544.5 | 0 | 8 | 0 | 11 | 0 | 11 | 1.63 | 1.39 | 1.12 | 0.95 |
| Q6AYT7 | 9928425 | 10001447 | 13420353 | 11155857 | 13529037 | 3605408.25 | 3115539 | 7362916 | 5457305 | 8016842 | 0 | 9 | 0 | 7 | 0 | 9 | 1.82 | 1.63 | 0.86 | 0.78 |
| Q5SSL4 | 1298322.5 | 1708274.38 | 951102.44 | 1425104.38 | 1746725.12 | 4205417 | 1823019.62 | 2151851.25 | 2548934 | 2576422 | 0 | 10 | 0 | 9 | 0 | 10 | 0.47 | 0.49 | 0.89 | 0.91 |
| Q4KML4 | 3244150 | 3238398.75 | 2356760.75 | 1314134.38 | 2500888 | 7758814 | 5931499.5 | 4284298.5 | 3279311 | 4009400 | 0 | 3 | 0 | 4 | 0 | 4 | 0.30 | 0.36 | 0.60 | 0.72 |
| Q8JZN5 | 13395500 | 14618983 | 18045232 | 17511716 | 12765865 | 19926202 | 19981610 | 22734662 | 21098514 | 22717526 | 0 | 12 | 0 | 3 | 0 | 12 | 0.71 | 0.78 | 1.00 | 1.08 |
| Q9JIX8 | 3362996.25 | 2986162.75 | 2702165.5 | 3615508 | 3506767.5 | 3966059.75 | 4420869 | 3939507.25 | 4204008 | 4560664.5 | 0 | 2 | 0 | 0 | 0 | 2 | 0.57 | 0.52 | 0.75 | 0.68 |
| P54987 | 20002776 | 19841840 | 22009238 | 14388771 | 22568140 | 4476999 | 5382819 | 6600090.5 | 5672641 | 7470627 | 0 | 0 | 0 | 4 | 0 | 4 | 7.11 | 4.67 | 2.13 | 1.40 |
| Q9D358 | 9311773 | 11091374 | 10357466 | 10997143 | 9846320 | 17781652 | 21960460 | 15401542 | 17242272 | 16758012 | 0 | 12 | 0 | 1 | 0 | 12 | 0.62 | 0.71 | 1.07 | 1.23 |
| Q9QUJ7 | 89543818 | 91066066 | 91863090 | 78868394 | 93616482 | 75037770 | 69010842 | 66721334 | 63455134 | 69597050 | 0 | 9 | 0 | 8 | 0 | 9 | 1.41 | 1.37 | 1.09 | 1.06 |
| Q9QXG4 | 7307326.5 | 7458311 | 8089299 | 8152918 | 9004635 | 15379777 | 12087815 | 12756392 | 12197419 | 12498191 | 0 | 12 | 0 | 8 | 0 | 12 | 0.66 | 0.65 | 1.07 | 1.06 |
| P57780 | 27095216 | 40868066 | 27574836 | 29738726 | 34009346 | 57831258 | 46718858 | 50458038 | 46790306 | 54159386 | 0 | 1 | 0 | 6 | 0 | 6 | 0.52 | 0.56 | 0.83 | 0.90 |
| P61161 | 58896066 | 75830002 | 66273258 | 57209086 | 80672202 | 142445106 | 75853762 | 96370498 | 102546354 | 113596122 | 0 | 4 | 0 | 8 | 0 | 8 | 0.50 | 0.55 | 0.78 | 0.87 |
| Q4V7C7 | 91478762 | 96705882 | 84592490 | 75955322 | 86810746 | 121355322 | 116082322 | 121633922 | 119218602 | 129139090 | 0 | 0 | 0 | 12 | 0 | 12 | 0.65 | 0.71 | 0.90 | 1.00 |
| Q9Z0F8 | 841951.88 | 645758.12 | 791699.94 | 1167681 | 867609.25 | 299561.25 | 230305.8 | 442023.25 | 15863.36 | 2 | 0 | 10 | 0 | 9.5 | 0 | 10 | 5.01 | 5.43 | 1.15 | 1.24 |
| Q05910 | 15753664 | 12194551 | 10485853 | 9631999 | 10020924 | 1748590.38 | 1596504.25 | 1306191.38 | 2465615.25 | 1533906.75 | 0 | 5 | 0 | 11 | 0 | 11 | 9.54 | 6.50 | 1.42 | 0.97 |
| Q8JZQ2 | 7953209.5 | 8205233 | 11657622 | 13026175 | 12836860 | 5787243.5 | 8783274 | 8861670 | 9259989 | 8774111 | 0 | 2 | 0 | 2 | 0 | 2 | 1.86 | 1.79 | 1.44 | 1.38 |
| P50247 | 110514234 | 115190322 | 94739314 | 101615874 | 91543746 | 188086002 | 158783122 | 152496530 | 161247826 | 152698050 | 0 | 9 | 0 | 11 | 0 | 11 | 0.60 | 0.62 | 0.95 | 0.98 |
| O70200 | 500938.94 | 367437.47 | 275214.69 | 678180.56 | 2 | 2220180.5 | 1811020 | 1192203.5 | 739106.12 | 986513.06 | 0 | 3 | 0 | 9 | 0 | 9 | 0.16 | 0.22 | 0.61 | 0.84 |
| O08915 | 1518733.12 | 1060656.12 | 2095438 | 1822907.25 | 1215229.75 | 3442565 | 2799465 | 3990173.5 | 3298561.5 | 2734841.5 | 0 | 6 | 0 | 6 | 0 | 6 | 0.53 | 0.55 | 1.13 | 1.15 |
| Q9WTP6 | 9463196 | 8058540.5 | 7168981.5 | 10841718 | 8515407 | 17559652 | 25295778 | 17166932 | 19596966 | 18396718 | 0 | 7 | 0 | 5 | 0 | 7 | 0.50 | 0.53 | 1.11 | 1.18 |
| P06214 | 4085427.75 | 4150240 | 2907144 | 3895298.5 | 3933843.5 | 5965630 | 6557158 | 5666135.5 | 6405606.5 | 5814811 | 0 | 2 | 0 | 8 | 0 | 8 | 0.70 | 0.68 | 1.13 | 1.08 |
| Q61490 | 10319752 | 10759348 | 7847498 | 9404139 | 10426511 | 3304052 | 3045259.5 | 2667592 | 2930240.5 | 2746924.25 | 0 | 6 | 0 | 6 | 0 | 6 | 4.00 | 3.83 | 1.20 | 1.15 |
| Q571I9 | 3246910.5 | 2585368 | 2347460.5 | 2874486.75 | 2713110.5 | 1514888.75 | 3639365.25 | 2033837.25 | 2299912.5 | 2786625 | 0 | 4 | 0 | 5 | 0 | 5 | 1.73 | 1.68 | 1.54 | 1.50 |
| Q9Z110 | 39710534 | 35184546 | 44583054 | 48924290 | 47650102 | 17545054 | 23329384 | 23320368 | 19758636 | 17604710 | 0 | 3 | 0 | 9 | 0 | 9 | 2.48 | 2.34 | 1.17 | 1.10 |
| P05064 | 614135874 | 651428674 | 697288066 | 810012418 | 688416642 | 1032909826 | 1217700098 | 1095962370 | 1275317506 | 1156400898 | 0 | 0 | 0 | 1 | 0 | 1 | 0.77 | 0.71 | 1.29 | 1.18 |
| P05063 | 26245102 | 23378532 | 27989454 | 30687114 | 26776524 | 36423338 | 41220870 | 45593106 | 40814906 | 45788706 | 0 | 8 | 0 | 6 | 0 | 8 | 0.61 | 0.70 | 0.94 | 1.09 |
| O08583 | 6266498.5 | 6644567.5 | 6099895 | 7345934 | 6444590 | 5912227.5 | 5891946.5 | 7305514.5 | 5144697.5 | 7032554 | 0 | 3 | 0 | 2 | 0 | 3 | 1.32 | 1.41 | 1.26 | 1.35 |
| B2RXR6 | 2 | 320308.06 | 606909.69 | 591800.19 | 767775.94 | 841410.06 | 437206.31 | 807747.75 | 1102335.12 | 1567634 | 0 | 3 | 0 | 2 | 0 | 3 | 0.24 | 0.26 | 0.50 | 0.55 |
| Q8BH79 | 777151.5 | 852273.19 | 1374865.12 | 1062282.75 | 961519.62 | 290066.38 | 2 | 479395.94 | 503243.41 | 583772.56 | 0 | 5.5 | 0 | 9.5 | 0 | 9.5 | 5.74 | 3.14 | 2.12 | 1.16 |
| O35381 | 7219836.5 | 15248762 | 7751460.5 | 6406946 | 10426322 | 20829754 | 11469968 | 11301684 | 10228472 | 16302438 | 0 | 4 | 0 | 7 | 0 | 7 | 0.48 | 0.56 | 0.71 | 0.84 |
| P97822 | 6384944.5 | 18182350 | 12450206 | 10589692 | 14407733 | 30720846 | 13666128 | 15249906 | 10109668 | 17604306 | 0 | 4 | 0 | 5 | 0 | 5 | 0.44 | 0.49 | 0.62 | 0.69 |
| O35639 | 53432270 | 53534506 | 67959578 | 76970082 | 70776042 | 32335598 | 42426562 | 39826022 | 33507966 | 34065218 | 0 | 12 | 0 | 9 | 0 | 12 | 1.85 | 1.93 | 1.05 | 1.09 |
| Q07076 | 53988222 | 50497058 | 43796738 | 42667494 | 49499678 | 30753040 | 30077082 | 26561904 | 29410552 | 30364024 | 0 | 7 | 0 | 11 | 0 | 11 | 1.76 | 1.64 | 1.08 | 1.00 |
| O35643 | 6781939.5 | 5704941.5 | 7107218 | 4930555 | 7176565.5 | 11480509 | 10142868 | 12698150 | 11347795 | 13937547 | 0 | 0 | 0 | 4 | 0 | 4 | 0.40 | 0.44 | 0.75 | 0.83 |
| P35585 | 3099140.5 | 2370494.5 | 2248026.75 | 2022577 | 2302721.5 | 3630415.5 | 3943334.75 | 4156437.25 | 3960636 | 3223179.5 | 0 | 10 | 0 | 10 | 0 | 10 | 0.60 | 0.60 | 0.94 | 0.94 |
| Q7TN05 | 775720.44 | 2 | 2 | 718021 | 2 | 980667 | 854427.5 | 2 | 1452535.88 | 2 | 0 | 3 | 0 | 1 | 0 | 3 | 0.20 | 0.19 | 0.45 | 0.43 |
| P62743 | 3498136.75 | 1413719.62 | 1262728.5 | 1105073 | 1488366.12 | 2 | 2 | 2 | 2 | 2 | 0 | 12.5 | 0 | 10 | 0 | 12.5 | 876802.40 | 23.14 | 1.00 | 0.00 |
| O54774 | 2031895.12 | 1865146.25 | 1577748.88 | 1735517.62 | 1667938.25 | 3413284.75 | 1314963.62 | 2922310.25 | 2167795 | 2715637.75 | 0 | 8 | 0 | 9 | 0 | 9 | 0.62 | 0.61 | 0.87 | 0.87 |
| Q8R5A3 | 2036772.75 | 1898100.88 | 1560816.75 | 1382374.12 | 1703166.25 | 281777.69 | 278846.5 | 2 | 2 | 2 | 0 | 10.5 | 0 | 7 | 0 | 10.5 | 8.41 | 6.89 | 0.55 | 0.45 |
| O35841 | 9813522 | 10870504 | 8826902 | 9062649 | 8833780 | 17293072 | 17253454 | 13930846 | 13168368 | 12390238 | 0 | 6 | 0 | 12 | 0 | 12 | 0.59 | 0.63 | 0.92 | 0.99 |
| Q9D7N9 | 3951952.5 | 1831665 | 2141260.75 | 1777084.25 | 1997864.38 | 473584.06 | 596468.75 | 1084666.12 | 726964.25 | 766984.25 | 0 | 9 | 0 | 6 | 0 | 9 | 3.31 | 4.66 | 1.03 | 1.45 |
| Q8VBT6 | 6984240 | 8092972.5 | 8685173 | 5829414 | 7365107.5 | 14895387 | 12503558 | 17470098 | 18724398 | 19805150 | 0 | 8 | 0 | 12 | 0 | 12 | 0.40 | 0.44 | 0.89 | 0.99 |
| Q5XJY5 | 14824155 | 16118553 | 16458550 | 14717467 | 12848068 | 20689190 | 17854228 | 21141150 | 22718240 | 20076190 | 0 | 12 | 0 | 12 | 0 | 12 | 0.72 | 0.74 | 0.99 | 1.01 |
| Q6Y5D8 | 471807.03 | 459191.34 | 668398.56 | 442574.38 | 2 | 716455.75 | 762442.5 | 910481.75 | 845766.19 | 813679.88 | 0 | 9 | 0 | 8 | 0 | 9 | 0.52 | 0.44 | 1.04 | 0.88 |
| Q640N3 | 1026521.44 | 1018189.25 | 1052182.88 | 886350.62 | 899897.5 | 300406.12 | 365594.06 | 164644.23 | 262098.05 | 2 | 0 | 7.5 | 0 | 6 | 0 | 7.5 | 3.15 | 2.58 | 0.71 | 0.58 |
| Q3TBD2 | 3740066.25 | 4884473.5 | 5015516 | 4531720.5 | 5416835.5 | 8506425 | 7069089 | 7603472 | 8723605 | 8062815 | 0 | 2 | 0 | 9 | 0 | 9 | 0.45 | 0.54 | 0.75 | 0.91 |
| Q5XI73 | 103417106 | 105216970 | 93833882 | 96809474 | 92593586 | 132299490 | 156260386 | 161944370 | 147219794 | 142174338 | 0 | 3 | 0 | 11 | 0 | 11 | 0.59 | 0.65 | 0.88 | 0.97 |
| Q9JM76 | 40141946 | 41659018 | 49016626 | 36794566 | 36587090 | 87001578 | 79442010 | 90808738 | 57191478 | 80051026 | 0 | 9 | 0 | 6 | 0 | 9 | 0.57 | 0.57 | 1.09 | 1.11 |
| Q4KLF8 | 39164490 | 41572970 | 41487010 | 42810470 | 44390294 | 78574538 | 89374226 | 83094882 | 91923106 | 86940682 | 0 | 8 | 0 | 3 | 0 | 8 | 0.47 | 0.53 | 0.96 | 1.09 |
| A1L108 | 1114495.12 | 2224783.25 | 1138557.88 | 1863533.12 | 1875632 | 4405968 | 2909724.5 | 3047416.25 | 3344755 | 2055927.75 | 0 | 6 | 0 | 10 | 0 | 10 | 0.39 | 0.44 | 0.74 | 0.84 |
| P29066 | 2 | 237990.5 | 297565.94 | 66171.43 | 175562.75 | 412344.41 | 167970.64 | 305472.34 | 297133.78 | 305457.84 | 0 | 1 | 0 | 2 | 0 | 2 | 0.27 | 0.29 | 0.51 | 0.56 |
| P50428 | 271358.75 | 2 | 2 | 2 | 2 | 663678.12 | 1377756 | 1107358 | 1153772.62 | 592369.38 | 0 | 9 | 0 | 7 | 0 | 9 | 0.06 | 0.07 | 1.09 | 1.35 |
| P50429 | 982859.5 | 2694667.25 | 2482796.75 | 1703945.5 | 3113993.75 | 16842870 | 12119315 | 10708555 | 11000842 | 12631576 | 0 | 0 | 0 | 2 | 0 | 2 | 0.07 | 0.11 | 0.42 | 0.65 |
| Q1AAU6 | 1040991.5 | 1979642.75 | 1939111.75 | 1411923.12 | 1728913.25 | 3362983.5 | 1381564.88 | 3573960.5 | 4817299.5 | 4555063 | 0 | 8 | 0 | 8 | 0 | 8 | 0.38 | 0.37 | 0.83 | 0.82 |
| Q3KRE0 | 2887723.75 | 2205978.5 | 3028997.75 | 1823179.25 | 2942015.75 | 377643.38 | 696432 | 580788.56 | 383538.69 | 409622.25 | 0 | 8 | 0 | 6 | 0 | 8 | 6.69 | 6.52 | 1.27 | 1.24 |
| O08997 | 10027890 | 9524634 | 11068398 | 11784971 | 11218188 | 10836040 | 8510422 | 18714182 | 16028714 | 18859590 | 0 | 6 | 0 | 9 | 0 | 9 | 0.50 | 0.68 | 0.68 | 0.92 |
| O55143 | 51009886 | 48171486 | 46003126 | 44210522 | 49306630 | 42423274 | 40996514 | 37055514 | 43775230 | 37127102 | 0 | 4 | 0 | 3 | 0 | 4 | 1.43 | 1.33 | 1.20 | 1.12 |
| O35143 | 3594067.5 | 2 | 2936256 | 3131063.5 | 3892749.25 | 6122254 | 7265000.5 | 5627148 | 5263701.5 | 6616508 | 0 | 8 | 0 | 5 | 0 | 8 | 0.48 | 0.52 | 1.08 | 1.18 |
| Q9DB20 | 25401152 | 20806226 | 25484574 | 24161616 | 26546278 | 16202130 | 22864852 | 20146762 | 18568622 | 20435636 | 0 | 4 | 0 | 4 | 0 | 4 | 1.42 | 1.46 | 1.14 | 1.17 |
| P15920 | 588741 | 697374.12 | 686893.5 | 842727.62 | 707264.38 | 2 | 2 | 450995.16 | 546834.69 | 404734.31 | 0 | 6 | 0 | 12 | 0 | 12 | 15.83 | 2.44 | 6.30 | 0.97 |
| Q5FVI6 | 20291176 | 23482254 | 22020416 | 18243908 | 21647282 | 39976774 | 34725002 | 34369574 | 41254678 | 41174810 | 0 | 10 | 0 | 10 | 0 | 10 | 0.54 | 0.52 | 0.98 | 0.94 |
| Q64430 | 1551131.25 | 1868661.38 | 2581115.25 | 1878435.5 | 2764956.25 | 1100947.62 | 1291759.25 | 1416808.5 | 1479432.25 | 957584.25 | 0 | 6 | 0 | 3 | 0 | 6 | 2.30 | 2.62 | 1.35 | 1.54 |
| Q99J86 | 404401.5 | 378675.28 | 392320.97 | 238304.66 | 532639.81 | 2 | 2 | 2 | 2 | 2 | 0 | 12.5 | 0 | 12.5 | 0 | 12.5 | 194634.22 | 194634.22 | 1.00 | 1.00 |
| Q9Z277 | 887648.38 | 2420304.5 | 1873248.38 | 1942934.25 | 2522276.25 | 2936937 | 1533712.5 | 2919676 | 2949842.5 | 3029727.5 | 0 | 0 | 0 | 5 | 0 | 5 | 0.46 | 0.56 | 0.63 | 0.77 |
| P24288 | 10205009 | 9245840 | 6986470.5 | 6745172 | 6915704.5 | 6600257 | 6544434.5 | 3549133.25 | 4581167 | 3258155.75 | 0 | 2 | 0 | 6 | 0 | 6 | 3.03 | 2.17 | 1.85 | 1.33 |
| Q9Z0H7 | 2 | 478889.97 | 2 | 2 | 2 | 875730.75 | 407561.66 | 720654.56 | 652777.75 | 713016.88 | 0 | 1 | 0 | 1 | 0 | 1 | 0.10 | 0.10 | 0.69 | 0.68 |
| P53563 | 3178506.5 | 3726258.75 | 3068271.5 | 2736069 | 2172570.5 | 6483247 | 5799423 | 4580708 | 4605383.5 | 4065793 | 0 | 11 | 0 | 11 | 0 | 11 | 0.57 | 0.60 | 0.98 | 1.03 |
| Q8R016 | 2126503.25 | 2848438 | 1940982.75 | 3054732.5 | 2134410.5 | 5263597.5 | 4646043 | 4200817.5 | 2778729 | 4690108.5 | 0 | 8 | 0 | 12 | 0 | 12 | 0.51 | 0.56 | 0.91 | 1.00 |
| Q9CY64 | 27281938 | 28413794 | 28861326 | 25251022 | 23350476 | 31471798 | 28921738 | 30461610 | 31793530 | 30824600 | 0 | 4 | 0 | 2 | 0 | 4 | 0.81 | 0.80 | 0.93 | 0.93 |
| Q923D2 | 14671318 | 6608738 | 6307858 | 8127343 | 6285399 | 1212510.25 | 1520904.88 | 1950822.5 | 1578359 | 1225602.38 | 0 | 0 | 0 | 0 | 0 | 0 | 20.62 | 13.02 | 3.68 | 2.32 |
| Q9ESU6 | 769168.62 | 1204432.5 | 1276054.62 | 562412.44 | 919077.69 | 2484575.25 | 2474942.5 | 2230089.5 | 2290493.25 | 2190506.5 | 0 | 7 | 0 | 5 | 0 | 7 | 0.38 | 0.38 | 0.95 | 0.94 |
| P18572 | 5974050.5 | 5489107.5 | 5946719.5 | 7031816 | 6179538.5 | 5059563 | 6006546.5 | 3882911.25 | 4168914 | 3876563.75 | 0 | 2 | 0 | 8 | 0 | 8 | 1.91 | 1.56 | 1.44 | 1.17 |
| O08901 | 31996902 | 39303550 | 40061906 | 48747586 | 40773638 | 61797510 | 77785898 | 65190994 | 67593570 | 75526626 | 0 | 5 | 0 | 3 | 0 | 5 | 0.65 | 0.68 | 1.13 | 1.18 |
| Q9WVA3 | 17059766 | 21671838 | 16436074 | 23576734 | 22049990 | 30316034 | 36363978 | 27500792 | 29730698 | 29196570 | 0 | 8 | 0 | 5 | 0 | 8 | 0.65 | 0.71 | 0.99 | 1.08 |
| O54825 | 2902069.5 | 3208845.75 | 2921822.5 | 2149537.5 | 2000373.88 | 1623538.38 | 1731947.75 | 1040340.19 | 1017715.81 | 912993 | 0 | 9 | 0 | 9 | 0 | 9 | 2.67 | 2.80 | 1.28 | 1.34 |
| P14106 | 849015.94 | 657943.25 | 2 | 2 | 2 | 2066759.25 | 2418033.75 | 764220.81 | 447096.72 | 2 | 0 | 7 | 0 | 8 | 0 | 8 | 0.16 | 0.19 | 0.60 | 0.73 |
| P01027 | 6384931.5 | 5884645 | 8649081 | 9174553 | 8417449 | 902166.38 | 2294937.5 | 3349028 | 4154984.75 | 3636309.5 | 0 | 9 | 0 | 9 | 0 | 9 | 3.08 | 2.21 | 1.15 | 0.82 |
| P01029 | 105530.12 | 116954.55 | 478942 | 435041.59 | 458600.28 | 2 | 2 | 277277.56 | 288529.41 | 125331.89 | 0 | 5 | 0 | 6 | 0 | 6 | 159506.85 | 17.61 | 69114.29 | 7.63 |
| P30993 | 19034822 | 19478004 | 21360574 | 21672352 | 21959542 | 17688686 | 20563804 | 13774112 | 15895223 | 14815232 | 0 | 6 | 0 | 4 | 0 | 6 | 1.46 | 1.55 | 1.17 | 1.24 |
| P00920 | 288087.12 | 207276.84 | 226640.8 | 176559.75 | 108265.89 | 338823.22 | 354136.59 | 330817.91 | 248126.77 | 247569.97 | 0 | 1 | 0 | 3 | 0 | 3 | 0.46 | 0.53 | 0.70 | 0.80 |
| Q63450 | 2 | 367121.75 | 250972.03 | 140350.34 | 215962.78 | 1612253.12 | 739433.94 | 591837.94 | 723871 | 468101.88 | 0 | 7 | 0 | 7 | 0 | 7 | 0.17 | 0.16 | 0.73 | 0.68 |
| Q6PHZ2 | 10608928 | 10671437 | 11223797 | 8371907 | 12397556 | 17281780 | 15815405 | 19507128 | 21090912 | 19628640 | 0 | 4 | 0 | 5 | 0 | 5 | 0.66 | 0.64 | 1.15 | 1.12 |
| P35564 | 92233210 | 77760794 | 81179882 | 63588282 | 83527874 | 54067514 | 54062926 | 52782790 | 50798246 | 50104354 | 0 | 0 | 0 | 8 | 0 | 8 | 1.76 | 1.58 | 1.16 | 1.04 |
| P40124 | 222119330 | 226758658 | 169648098 | 177702498 | 163722226 | 506864450 | 398874722 | 300150242 | 309237986 | 311801634 | 0 | 7 | 0 | 8 | 0 | 8 | 0.47 | 0.49 | 0.90 | 0.93 |
| P24452 | 6362775 | 4258558.5 | 2413275 | 2430783 | 2980359.25 | 2 | 2 | 2 | 2 | 2 | 0 | 10 | 0 | 12.5 | 0 | 12.5 | 20.90 | 1844575.08 | 0.00 | 1.00 |
| O35350 | 172390.72 | 1760429.38 | 754564.12 | 641610.5 | 1441714.12 | 3422802.5 | 1220637.38 | 3353927.75 | 2379836.5 | 3110247.5 | 0 | 9 | 0 | 12 | 0 | 12 | 0.29 | 0.34 | 0.82 | 0.95 |
| O88456 | 2847223 | 6101771.5 | 2315000.75 | 1051738.62 | 4044087.5 | 11144080 | 8247007.5 | 5794765 | 5712035 | 6976809 | 0 | 3 | 0 | 6 | 0 | 6 | 0.30 | 0.34 | 0.69 | 0.80 |
| P47753 | 11103678 | 9524960 | 8189003 | 7174848 | 9206734 | 16221279 | 17780278 | 13523088 | 12902043 | 13518558 | 0 | 6 | 0 | 7 | 0 | 7 | 0.55 | 0.57 | 0.90 | 0.94 |
| P47754 | 12204783 | 11001484 | 13006926 | 12325875 | 13858605 | 29088270 | 29478204 | 26228878 | 23719384 | 21982158 | 0 | 10 | 0 | 10 | 0 | 10 | 0.46 | 0.46 | 0.97 | 0.96 |
| Q5XI32 | 39567682 | 46893322 | 44563590 | 46664402 | 48374170 | 91263938 | 91504266 | 86511234 | 93787242 | 92017610 | 0 | 11 | 0 | 3 | 0 | 11 | 0.50 | 0.53 | 1.01 | 1.07 |
| Q4AE70 | 228177.22 | 214448.17 | 321026.53 | 345460.75 | 214950.98 | 59143.31 | 392782 | 159819.75 | 2 | 181364.05 | 0 | 7.5 | 0 | 2.5 | 0 | 7.5 | 5.09 | 132406.37 | 3.05 | 79311.11 |
| P51125 | 2620520 | 2997422 | 2243961.75 | 2436643.75 | 1861749.25 | 4736277 | 4154599.25 | 5415365 | 4734693.5 | 4932092.5 | 0 | 11 | 0 | 8 | 0 | 11 | 0.51 | 0.47 | 1.01 | 0.93 |
| P24270 | 3345282 | 5556524 | 4422586 | 3836579.5 | 5190314.5 | 2830377.25 | 1359751.88 | 2928296 | 3977639.75 | 2898963.5 | 0 | 0 | 0 | 5 | 0 | 5 | 3.82 | 2.27 | 2.39 | 1.42 |
| P22682 | 894043.44 | 888159.38 | 853576.31 | 1230390.25 | 790115.19 | 2 | 2 | 552526.38 | 724146.62 | 565990.31 | 0 | 5 | 0 | 10 | 0 | 10 | 465628.46 | 3.03 | 184266.73 | 1.20 |
| Q8VEH6 | 441313.59 | 598301.56 | 585462.88 | 357843.06 | 619200.38 | 167533.69 | 2 | 2 | 95171.94 | 245965.44 | 0 | 11 | 0 | 5 | 0 | 11 | 4.86 | 2.55 | 0.95 | 0.50 |
| P83917 | 4180597.75 | 3956777 | 4060838.5 | 4783398.5 | 4579744 | 9722690 | 11320977 | 7190009.5 | 8184884.5 | 7555478 | 0 | 11 | 0 | 11 | 0 | 11 | 0.49 | 0.52 | 1.00 | 1.07 |
| P23198 | 10257938 | 9489630 | 7177663 | 7514349 | 8341837.5 | 10456609 | 16212969 | 11878497 | 12867423 | 11522276 | 0 | 3 | 0 | 9 | 0 | 9 | 0.54 | 0.65 | 0.80 | 0.96 |
| P80314 | 95542666 | 86454266 | 81077826 | 83109146 | 72384178 | 64904890 | 73764274 | 71461770 | 69159962 | 66055374 | 0 | 3 | 0 | 6 | 0 | 6 | 1.33 | 1.29 | 1.10 | 1.07 |
| P21855 | 5685576.5 | 4950914 | 5794479 | 5462664.5 | 5506759 | 2771005.75 | 2558361.5 | 4043862.25 | 3980384 | 3514743.75 | 0 | 7 | 0 | 3 | 0 | 7 | 2.07 | 2.32 | 1.27 | 1.43 |
| P40237 | 1528681.12 | 1270900.5 | 1804171.62 | 1707588.62 | 1965921.75 | 709602.69 | 2 | 294829.88 | 2 | 2 | 0 | 11.5 | 0 | 8.5 | 0 | 11.5 | 10.62 | 5.73 | 1.29 | 0.70 |
| Q61081 | 5231879.5 | 6011082.5 | 4728049.5 | 4351062 | 5163146 | 9370558 | 10353200 | 8327007.5 | 8593367 | 9637675 | 0 | 11 | 0 | 11 | 0 | 11 | 0.54 | 0.53 | 0.98 | 0.97 |
| P60766 | 19090246 | 18736476 | 18718286 | 11450051 | 16932422 | 26588534 | 37559450 | 24744980 | 25928838 | 24017224 | 0 | 4 | 0 | 4 | 0 | 4 | 0.71 | 0.71 | 1.15 | 1.17 |
| Q4V8C8 | 7694914.5 | 7987646 | 7272199 | 2378364.75 | 2353211.25 | 285681.22 | 1185893.12 | 233703.89 | 996589.5 | 233024.5 | 0 | 3 | 0 | 2 | 0 | 3 | 4.50 | 3.44 | 0.48 | 0.36 |
| Q8VDP6 | 2581199.5 | 2403590.75 | 4043855 | 4379123 | 3077969.5 | 1526571.25 | 1784353 | 2228243.25 | 2356180.5 | 2231270 | 0 | 6 | 0 | 6 | 0 | 6 | 1.93 | 1.85 | 1.19 | 1.13 |
| P11440 | 6934705.5 | 8777903 | 6981411 | 6292311 | 6694706.5 | 13258054 | 10062064 | 8526360 | 7900430 | 7271599 | 0 | 3 | 0 | 5 | 0 | 5 | 0.57 | 0.65 | 0.75 | 0.86 |
| Q04899 | 540974.31 | 690629.25 | 570322.06 | 331825.75 | 594809.94 | 223020.81 | 161912.7 | 385973.56 | 280306.88 | 299368.03 | 0 | 10 | 0 | 12 | 0 | 12 | 2.34 | 2.01 | 1.16 | 0.99 |
| Q641Z4 | 465585.06 | 367857.44 | 434891.38 | 353494.59 | 376377.28 | 2 | 2 | 2 | 2 | 2 | 0 | 7.5 | 0 | 7.5 | 0 | 7.5 | 5.28 | 8.07 | 0.00 | 0.00 |
| Q99L43 | 1371127 | 1584039.75 | 1696750.5 | 1849520.25 | 1681232.75 | 2 | 2 | 164354.97 | 1401031.5 | 680030.88 | 0 | 7 | 0 | 9 | 0 | 9 | 16.56 | 2.74 | 4.54 | 0.75 |
| Q8BGS7 | 1977661.25 | 2127569.5 | 2015721.25 | 2064738.25 | 1956105.62 | 154077.41 | 2 | 242225.23 | 2 | 353603.53 | 0 | 9 | 0 | 2 | 0 | 9 | 9.67 | 4.61 | 0.72 | 0.34 |
| Q924Z4 | 1102118.5 | 1426117.38 | 2419867.5 | 1652745.5 | 2182233 | 183355.59 | 873208.75 | 780294.81 | 150521.55 | 78248.88 | 0 | 11 | 0 | 9 | 0 | 11 | 3.94 | 5.51 | 0.93 | 1.30 |
| Q9D6J1 | 903536.06 | 672312.81 | 1027753.06 | 1234819.38 | 1142286.25 | 2 | 2 | 652073.31 | 795254.31 | 508848.66 | 0 | 9 | 0 | 10 | 0 | 10 | 4.43 | 2.74 | 1.74 | 1.07 |
| Q9D6K9 | 3270555.5 | 4278985 | 4629232.5 | 3096323.25 | 4330059.5 | 747337.31 | 282292.97 | 1584979.62 | 1185369.75 | 1199595.62 | 0 | 7 | 0 | 6 | 0 | 7 | 2.81 | 2.62 | 0.72 | 0.67 |
| Q6PDQ2 | 12740057 | 9912070 | 11242066 | 13268734 | 10669761 | 16715333 | 18694924 | 14284036 | 17150234 | 16006203 | 0 | 11 | 0 | 10 | 0 | 11 | 0.68 | 0.72 | 0.97 | 1.03 |
| Q9D8B3 | 6363682 | 6555359 | 6855648.5 | 7694914 | 7289633.5 | 4237323 | 6028029 | 4288894.5 | 3589109.5 | 4791954 | 0 | 9 | 0 | 11 | 0 | 11 | 1.66 | 1.46 | 1.10 | 0.96 |
| D4A4T9 | 3997395 | 5199590 | 4257780.5 | 3451181.5 | 5023869.5 | 8846882 | 6807567.5 | 6789221.5 | 6791580 | 6978797 | 0 | 8 | 0 | 0 | 0 | 8 | 0.67 | 0.69 | 1.11 | 1.14 |
| B0K020 | 2283674.25 | 1725444.12 | 2860612.75 | 2954805.75 | 3399822.25 | 448586.75 | 556483.12 | 1080794.5 | 1025185.44 | 1637559.12 | 0 | 8 | 0 | 12 | 0 | 12 | 4.45 | 3.02 | 1.60 | 1.08 |
| Q9CQB5 | 9108032 | 8995554 | 9684727 | 9927192 | 6099127 | 2177535.5 | 5158616 | 4744773 | 4363574 | 5079340 | 0 | 3 | 0 | 5 | 0 | 5 | 3.58 | 2.70 | 1.76 | 1.33 |
| A2AGT5 | 7306344.5 | 10917230 | 6760339.5 | 7577599.5 | 6989689 | 15036311 | 9376876 | 13761917 | 12746102 | 11546591 | 0 | 3 | 0 | 11 | 0 | 11 | 0.52 | 0.59 | 0.82 | 0.93 |
| Q04447 | 95051754 | 121946738 | 122833034 | 111181650 | 101794042 | 124975410 | 128055778 | 167563298 | 150542530 | 148797714 | 0 | 6 | 0 | 3 | 0 | 6 | 0.68 | 0.66 | 0.88 | 0.86 |
| Q8VBZ3 | 4435151.5 | 4877026 | 4646717 | 5908635 | 5385962 | 4286360 | 5302041 | 4720796.5 | 1679070.75 | 4269669 | 0 | 5 | 0 | 7 | 0 | 7 | 2.20 | 1.49 | 1.77 | 1.19 |
| O08585 | 2353338.5 | 1684241.12 | 6685222 | 2313229.5 | 5212441.5 | 15590410 | 15659413 | 14860430 | 19435998 | 16973350 | 0 | 11 | 0 | 7 | 0 | 11 | 0.23 | 0.24 | 1.03 | 1.08 |
| Q5I0K3 | 893941.31 | 934923.44 | 1158842.25 | 998002.12 | 917607.75 | 1639086.75 | 1520903.62 | 1656044.88 | 1333862.38 | 508131.22 | 0 | 9 | 0 | 12 | 0 | 12 | 0.70 | 0.67 | 0.95 | 0.91 |
| Q9DBL7 | 3833864.5 | 3114185.25 | 3334404.75 | 3445516.5 | 3371358 | 2565154.25 | 2425627.5 | 2955260.5 | 3116308.75 | 3028344.75 | 0 | 8 | 0 | 5 | 0 | 8 | 1.29 | 1.33 | 1.06 | 1.10 |
| Q8CIE6 | 40265106 | 40837738 | 44116738 | 36684994 | 40788714 | 57514306 | 61179482 | 55639322 | 53523154 | 53331334 | 0 | 7 | 0 | 4 | 0 | 7 | 0.68 | 0.66 | 0.94 | 0.92 |
| P23514 | 9424965 | 15110004 | 15151232 | 9784707 | 13539165 | 20379126 | 17989354 | 16425265 | 19841698 | 17398334 | 0 | 4 | 0 | 3 | 0 | 4 | 0.60 | 0.60 | 0.87 | 0.87 |
| O35142 | 34425702 | 34324630 | 31019382 | 31574358 | 31988566 | 43237810 | 40324462 | 44933562 | 47517874 | 45399338 | 0 | 6 | 0 | 3 | 0 | 6 | 0.78 | 0.81 | 1.05 | 1.09 |
| P61924 | 2040044.75 | 3843425.25 | 3240284 | 2022807.75 | 3490172 | 7463687 | 4741617 | 6226048 | 6188686 | 5135334 | 0 | 5 | 0 | 10 | 0 | 10 | 0.41 | 0.47 | 0.82 | 0.96 |
| Q9WUM4 | 19133772 | 18659226 | 17965376 | 19039818 | 18496726 | 37734350 | 34852418 | 37349514 | 37194498 | 41645930 | 0 | 4 | 0 | 2 | 0 | 4 | 0.54 | 0.57 | 1.09 | 1.14 |
| P19783 | 24113178 | 21657602 | 22750404 | 25271566 | 25269576 | 13697534 | 16728997 | 16813934 | 15888168 | 16324582 | 0 | 8 | 0 | 1 | 0 | 8 | 1.57 | 1.84 | 1.05 | 1.22 |
| Q9CPQ1 | 21348290 | 21548734 | 19244470 | 21701218 | 21927320 | 14318079 | 20776030 | 11594254 | 12461696 | 10714111 | 0 | 11 | 0 | 9 | 0 | 11 | 1.68 | 1.81 | 1.11 | 1.20 |
| Q66H71 | 2 | 494528.78 | 2 | 2 | 461718.19 | 1120445.5 | 2 | 2 | 642677.31 | 704329.62 | 0 | 2 | 0 | 5 | 0 | 5 | 0.17 | 0.21 | 0.44 | 0.55 |
| Q9WVJ3 | 614710.12 | 236829.23 | 614426.19 | 1012373.62 | 417944.97 | 2744713.5 | 3158618 | 3846206.5 | 3411793.5 | 3640248.5 | 0 | 9 | 0 | 0 | 0 | 9 | 0.21 | 0.30 | 1.22 | 1.76 |
| P47941 | 2 | 2 | 2 | 2 | 2 | 1751191.25 | 1001856.62 | 1280819.75 | 1675077 | 1781967.75 | 0 | 9 | 0 | 11 | 0 | 11 | 0.00 | 0.00 | 0.87 | 0.98 |
| Q99KP3 | 342654.72 | 200679.23 | 350701.22 | 2 | 2 | 624801.38 | 943206.56 | 2 | 822363.12 | 703246.31 | 0 | 10 | 0 | 12 | 0 | 12 | 0.23 | 0.26 | 0.81 | 0.90 |
| P09581 | 5957355.5 | 6134714.5 | 6690032.5 | 5705836 | 7150953.5 | 7398196 | 6294002.5 | 6751680 | 7439528 | 8636705 | 0 | 2 | 0 | 5 | 0 | 5 | 0.53 | 0.79 | 0.61 | 0.91 |
| P32577 | 9236026 | 9805139 | 9181255 | 8359036 | 8915433 | 15947652 | 14225793 | 15611836 | 14215982 | 13185240 | 0 | 6 | 0 | 9 | 0 | 9 | 0.56 | 0.66 | 0.90 | 1.06 |
| Q8BIQ5 | 1816623.25 | 1418831.25 | 1615494.25 | 1672684.5 | 1652313.5 | 2 | 171670.06 | 692029.12 | 2 | 2 | 0 | 12 | 0 | 10.5 | 0 | 12 | 18.49 | 14.47 | 1.95 | 1.53 |

| | | | | | | | | | | | | | | | | | | | | | | | | | | | | | | | | | | | | |
|---|---|---|---|---|---|---|---|---|---|---|---|---|---|---|---|---|---|---|---|---|---|---|---|---|---|---|---|---|---|---|---|---|---|---|---|---|
| O88712 | Ctbp1 | C-terminal-binding protein 1 | 432.26 | 19.27 | 47745 | 8 | 4837460 | 4373362 | 4441901 | 4639444 | 4206856.5 | 5290492 | 4579939 | 4116397.5 | 3925816.5 | 4041799 | 2912457 | 2926564.25 | 2902575 | 2998265.75 | 1697815 | 4692173 | 4455718.5 | 3836497 | 4373472.5 | 3559868.75 | 0 | 9 | 0 | 10 | 0 | 10 | 0.60 | 0.61 | 0.93 | 0.95 |
| Q5SUQ9 | Ctc1 | CST complex subunit CTC1 | 68.16 | 0.91 | 134028 | 1 | 377259.66 | 305969.53 | 354404.06 | 386156.78 | 445666.09 | 405283.44 | 550210.38 | 417154.94 | 376980.75 | 452127.88 | 175849.25 | 214894.31 | 265026.94 | 129092.23 | 282463 | 225921.61 | 2 | 334925.38 | 398748.81 | 402998.91 | 0 | 9 | 0 | 2 | 0 | 9 | 0.57 | 0.48 | 0.73 | 0.62 |
| Q61164 | Ctcf | Transcriptional repressor CTCF | 89.16 | 2.45 | 83745 | 1 | 305099.44 | 267622.34 | 321708.66 | 233637.69 | 295764.78 | 310896.09 | 365388.44 | 220003.52 | 231989.31 | 167699.58 | 2 | 147670.44 | 2 | 2 | 2 | 277983.22 | 331633 | 177337.94 | 2 | 2 | 0 | 7 | 0 | 8 | 0 | 8 | 0.10 | 0.11 | 0.55 | 0.61 |
| Q8VCN5 | Cth | Cystathionine gamma-lyase | 942.24 | 40.95 | 43567 | 15 | 38931342 | 39684586 | 22701240 | 39828798 | 34159694 | 48102042 | 41071706 | 38456990 | 35483394 | 40035890 | 105159770 | 82884922 | 94313754 | 95149810 | 99341730 | 46959098 | 56565026 | 57118294 | 58389214 | 59422318 | 0 | 0 | 0 | 1 | 0 | 1 | 2.72 | 2.31 | 1.59 | 1.35 |
| P16675 | Ctsa | Lysosomal protective protein | 1131.48 | 43.46 | 53844 | 18 | 133395418 | 130606898 | 129632746 | 148174402 | 166887522 | 105849642 | 124751426 | 111381714 | 116145970 | 101272954 | 26000130 | 26040210 | 26977068 | 24063008 | 26794096 | 146341250 | 131279202 | 121700338 | 124675034 | 132276658 | 0 | 7 | 0 | 2 | 0 | 7 | 0.18 | 0.23 | 0.93 | 1.17 |
| P10605 | Ctsb | Cathepsin B | 1390.44 | 59.88 | 37280 | 19 | 227687010 | 237342786 | 144279778 | 188717474 | 282802850 | 167335506 | 245120722 | 215620834 | 222223698 | 180320594 | 47628342 | 80215610 | 64645382 | 39798438 | 80146114 | 250517506 | 154833202 | 152005266 | 150147826 | 187809458 | 0 | 8 | 0 | 7 | 0 | 8 | 0.29 | 0.30 | 0.83 | 0.87 |
| P97821 | Ctsc | Dipeptidyl peptidase 1 | 350.48 | 11.04 | 52376 | 5 | 14882918 | 13536377 | 12625771 | 16922566 | 16400908 | 10056369 | 11104187 | 10505408 | 12143344 | 14608572 | 3926110.5 | 3988632.25 | 6805714 | 4995329.5 | 5691353.5 | 12099124 | 14572128 | 7845456.5 | 11764732 | 10175244 | 0 | 2 | 0 | 11 | 0 | 11 | 0.34 | 0.43 | 0.76 | 0.97 |
| P24268 | Ctsd | Cathepsin D | 576.48 | 19.66 | 44681 | 10 | 28492142 | 21037556 | 15338439 | 10309086 | 49814782 | 10294403 | 23875046 | 19274632 | 27578042 | 10057534 | 1218670.5 | 4259972 | 1393277.75 | 545463 | 2177100.25 | 2 | 10114845 | 5529561 | 7100870 | 8432397 | 0 | 1 | 0 | 1 | 0 | 1 | 0.08 | 0.11 | 0.25 | 0.34 |
| P18242 | Ctsd | Cathepsin D | 1781.51 | 54.63 | 44954 | 28 | 214010386 | 141707890 | 175723282 | 232967138 | 266084386 | 211509906 | 212960738 | 234712674 | 231453986 | 235514466 | 101865506 | 93374226 | 94719018 | 93631594 | 112494722 | 277949826 | 294248354 | 217899106 | 254964818 | 258123506 | 0 | 4 | 0 | 3 | 0 | 4 | 0.48 | 0.44 | 1.26 | 1.16 |
| O70370 | Ctss | Cathepsin S | 706.33 | 38.53 | 38475 | 9 | 67497538 | 67125162 | 35839430 | 47299234 | 64095410 | 64323454 | 79080074 | 42865774 | 52315650 | 42232778 | 15772356 | 18940196 | 15141739 | 13974331 | 17016190 | 71862594 | 54928322 | 39866838 | 42796870 | 45202370 | 0 | 10 | 0 | 10 | 0 | 10 | 0.29 | 0.29 | 0.90 | 0.91 |
| Q9WUU7 | Ctsz | Cathepsin Z | 644.7 | 45.1 | 33996 | 11 | 301307426 | 366608706 | 186394642 | 260748866 | 247046274 | 239288658 | 269751266 | 173890210 | 163964898 | 179187298 | 53963922 | 55904946 | 51712094 | 47698950 | 51216690 | 257881410 | 248935074 | 148243970 | 174790066 | 177257506 | 0 | 4 | 0 | 12 | 0 | 12 | 0.19 | 0.25 | 0.74 | 0.98 |
| Q6MGD0 | Cuta | Protein CutA | 64.27 | 7.91 | 18659 | 1 | 3473019 | 2974854.75 | 2806350.5 | 3537871.5 | 2915292 | 2675715.5 | 3425591.75 | 2715197.75 | 2662199 | 2883027.5 | 1393326.5 | 1102284.75 | 1084233.12 | 1089290.12 | 988306.62 | 3349193 | 3400780.75 | 980415.38 | 2987158.25 | 2824246.25 | 0 | 10 | 0 | 10 | 0 | 10 | 0.36 | 0.39 | 0.86 | 0.94 |
| Q9CQX2 | Cyb5b | Cytochrome b5 type B | 316.49 | 58.22 | 16318 | 5 | 1376888.12 | 2354756 | 3105410 | 1008773.75 | 1437789.25 | 1263738 | 1654581.62 | 1468857 | 1813716.25 | 2239677 | 6360090 | 6773005 | 8791580 | 8028418 | 9347346 | 2433243.5 | 2596683 | 3948626 | 2649847.25 | 3138392 | 0 | 3 | 0 | 0 | 0 | 3 | 4.23 | 4.66 | 1.59 | 1.75 |
| Q9DB73 | Cyb5r1 | NADH-cytochrome b5 reductase 1 | 682.64 | 50.49 | 34135 | 12 | 14111291 | 15806638 | 7516598.5 | 11459522 | 10617681 | 12623020 | 14057831 | 12205023 | 12645846 | 15056724 | 21034610 | 26214476 | 33145326 | 27885792 | 32187298 | 10676115 | 12902689 | 12910420 | 14340060 | 19072378 | 0 | 8 | 0 | 10 | 0 | 10 | 2.36 | 2.11 | 1.17 | 1.05 |
| Q9DCN2 | Cyb5r3 | NADH-cytochrome b5 reductase 3 | 972.82 | 46.18 | 34128 | 14 | 43607954 | 42809978 | 37728798 | 43531538 | 36114694 | 45600294 | 43297914 | 41942534 | 40805942 | 51465694 | 76787674 | 67449250 | 71572978 | 52350722 | 76243866 | 41668962 | 44562422 | 49733694 | 52286794 | 50384954 | 0 | 3 | 0 | 8 | 0 | 8 | 1.69 | 1.54 | 1.17 | 1.07 |
| Q8BKE6 | Cyp20a1 | Cytochrome P450 20A1 | 131.96 | 7.58 | 52149 | 3 | 746238.81 | 643629.56 | 580776.12 | 2 | 681437.25 | 392378.03 | 701323.62 | 579586.31 | 519170.19 | 2 | 1175618.75 | 1897439.75 | 751774.25 | 907403.94 | 979668.5 | 745240.69 | 2 | 2 | 2 | 415797.16 | 0 | 6.5 | 0 | 8.5 | 0 | 8.5 | 2.15 | 2.61 | 0.44 | 0.53 |
| P61804 | Dad1 | Dolichyl-diphosphooligosaccharide--protein glycosyltransferase subunit DAD1 | 275.11 | 35.4 | 12497 | 5 | 10470926 | 12811956 | 14223651 | 14010104 | 13973814 | 12306418 | 12763802 | 17467618 | 15727106 | 20385484 | 22323038 | 25797446 | 40525470 | 41786242 | 37469026 | 10679342 | 13241611 | 19977326 | 24555152 | 20216342 | 0 | 7 | 0 | 10 | 0 | 10 | 2.56 | 2.13 | 1.35 | 1.13 |
| P31786 | Dbi | Acyl-CoA-binding protein | 217.95 | 50.57 | 10000 | 3 | 10634191 | 9164357 | 12138297 | 14119228 | 12166713 | 10097446 | 8429456 | 12619304 | 10330353 | 9496982 | 4078808 | 5613093 | 6361463.5 | 4408121 | 6142115.5 | 12271616 | 11459901 | 10699329 | 9096512 | 12448279 | 0 | 12 | 0 | 8 | 0 | 12 | 0.46 | 0.52 | 0.96 | 1.10 |
| O54734 | Ddost | Dolichyl-diphosphooligosaccharide--protein glycosyltransferase 48 kDa subunit | 939.88 | 39.68 | 49028 | 14 | 20915626 | 20031972 | 24709282 | 23269566 | 21556386 | 25729610 | 20606310 | 23082530 | 23075630 | 26319756 | 65929622 | 46711034 | 56523034 | 49633682 | 52499346 | 17429464 | 22019734 | 30279706 | 22128570 | 25773916 | 0 | 9 | 0 | 11 | 0 | 11 | 2.46 | 2.28 | 1.06 | 0.99 |
| Q501J6 | Ddx17 | Probable ATP-dependent RNA helicase DDX17 | 1195.27 | 31.69 | 72399 | 20 | 16069296 | 21299302 | 22525162 | 22054908 | 25131066 | 16138516 | 19960414 | 21083836 | 19944432 | 18820302 | 9851357 | 9546976 | 11767068 | 7322714 | 11627910 | 21681464 | 17088558 | 17356752 | 18575230 | 20090958 | 0 | 6 | 0 | 12 | 0 | 12 | 0.47 | 0.52 | 0.89 | 0.99 |
| Q61655 | Ddx19a | ATP-dependent RNA helicase DDX19A | 430.32 | 17.36 | 53933 | 8 | 6650934.5 | 4279058 | 5378205.5 | 6522997.5 | 3363693.5 | 4094296.5 | 4073897.5 | 6928175.5 | 5466416.5 | 3924953.5 | 9063342 | 8575493 | 6998061.5 | 9036992 | 7657545 | 1560178.75 | 4610726.5 | 4420766.5 | 6975163.5 | 3725118 | 0 | 10 | 0 | 11 | 0 | 11 | 1.58 | 1.69 | 0.81 | 0.87 |
| Q921N6 | Ddx27 | Probable ATP-dependent RNA helicase DDX27 | 100.23 | 2.63 | 85939 | 2 | 2 | 321288.94 | 2 | 2 | 2 | 2 | 2 | 413319.34 | 2 | 2 | 1874121.62 | 1293865.62 | 1140336.38 | 1041999.62 | 1162560 | 2 | 2 | 282114.09 | 2 | 2 | 0 | 12 | 0 | 12 | 0 | 12 | 20.27 | 15.76 | 0.88 | 0.68 |
| Q63413 | Ddx39b | Spliceosome RNA helicase Ddx39b | 1241.82 | 50.93 | 49035 | 20 | 25447358 | 24410644 | 32732670 | 33207510 | 24134258 | 27626986 | 24174278 | 25638444 | 28859106 | 26629166 | 21657576 | 19966526 | 15534338 | 17997040 | 19397272 | 24395200 | 22276408 | 20457640 | 20125906 | 27366250 | 0 | 4 | 0 | 4 | 0 | 4 | 0.68 | 0.71 | 0.82 | 0.86 |
| Q9CWX9 | Ddx47 | Probable ATP-dependent RNA helicase DDX47 | 473.91 | 19.12 | 50639 | 9 | 4210993.5 | 4199856.5 | 3807033.75 | 1444020 | 3295991 | 4044662 | 3522336 | 2000602.25 | 2435649 | 2248510 | 4814973.5 | 4235272.5 | 4894923 | 4578365 | 4310187 | 3254863 | 2041247.75 | 2079435.62 | 3086381.5 | 2711935 | 0 | 5 | 0 | 11 | 0 | 11 | 1.35 | 1.60 | 0.78 | 0.92 |
| O09005 | Degs1 | Sphingolipid delta(4)-desaturase DES1 | 145.14 | 10.53 | 38241 | 2 | 2 | 2 | 164094.52 | 334792.81 | 2 | 2 | 2 | 2 | 2 | 2 | 633214.75 | 422370.75 | 1008304.19 | 1006414.75 | 842824.5 | 2 | 2 | 457219.59 | 264788.66 | 358493.06 | 0 | 8 | 0 | 5 | 0 | 8 | 7.84 | 391312.89 | 2.17 | 108050.53 |
| Q6AXS3 | Dek | Protein DEK | 445.44 | 20.37 | 42892 | 7 | 20493654 | 19553952 | 22402142 | 19361510 | 21256832 | 15014022 | 19694926 | 18151994 | 18670892 | 15232538 | 8541964 | 10528663 | 7752561 | 6916722 | 7525699.5 | 20402430 | 11690834 | 12359950 | 10419368 | 14940667 | 0 | 2 | 0 | 5 | 0 | 5 | 0.40 | 0.48 | 0.68 | 0.80 |
| A6H8H2 | Dennd4c | DENN domain-containing protein 4C | 179.86 | 2.31 | 211458 | 4 | 1806835.75 | 1408024.75 | 1728051.38 | 1760529.25 | 1958200.5 | 1782875.88 | 1582826.62 | 1751823.5 | 2685984.75 | 2315327.25 | 2 | 390661 | 819668 | 2 | 804231.69 | 1718979.25 | 1521341.25 | 1183484.5 | 672851.38 | 1322616.5 | 0 | 2 | 0 | 1 | 0 | 2 | 0.23 | 0.20 | 0.74 | 0.63 |
| Q9CQJ6 | Denr | Density-regulated protein | 140.13 | 13.13 | 22166 | 2 | 3091532.5 | 3653996.25 | 4545085.5 | 3561991.75 | 4084616.5 | 2365028.5 | 3543943.5 | 3723218 | 3941202.5 | 3293574.5 | 1662105.25 | 2201602.5 | 1880427.38 | 1644686.75 | 1970223.25 | 4741275.5 | 3523439.75 | 2841897.75 | 2781883.25 | 3306993.75 | 0 | 7 | 0 | 11 | 0 | 11 | 0.49 | 0.55 | 0.91 | 1.02 |
| Q99JS6 | Derl1 | Derlin-1 | 165.68 | 13.15 | 28835 | 3 | 180752.86 | 2 | 163375.83 | 2 | 39241.09 | 147228.64 | 253868.34 | 330686.69 | 418369.12 | 133970.81 | 1741929.88 | 1817824.25 | 2741966.25 | 1940256.5 | 2248538.5 | 183937.23 | 2 | 375452.97 | 160496.42 | 496785 | 0 | 6 | 0 | 12 | 0 | 12 | 27.36 | 8.17 | 3.17 | 0.95 |
| O88455 | Dhcr7 | 7-dehydrocholesterol reductase | 250.55 | 13.38 | 53919 | 5 | 8420224 | 8326428.5 | 6577549 | 9156462 | 7734344 | 7811316.5 | 7582275.5 | 9103084 | 8379915 | 6764625 | 12156979 | 10105335 | 16013407 | 12918322 | 16240539 | 12110634 | 11161273 | 11089765 | 9033594 | 12011790 | 0 | 1 | 0 | 1 | 0 | 1 | 1.68 | 1.70 | 1.38 | 1.40 |
| Q99J47 | Dhrs7b | Dehydrogenase/reductase SDR family member 7B | 311.79 | 19.81 | 34987 | 5 | 2487142.25 | 2193360.5 | 2090305.75 | 2643748 | 1884103.62 | 2774615 | 2580111.5 | 1696947.75 | 2079898.75 | 2242783.75 | 3696339.75 | 3548176.5 | 3308227.25 | 3226782 | 3673769 | 2843676.75 | 2885475 | 2441491.5 | 3737674.25 | 2831290.25 | 0 | 2 | 0 | 2 | 0 | 2 | 1.54 | 1.53 | 1.30 | 1.30 |
| O35286 | Dhx15 | Pre-mRNA-splicing factor ATP-dependent RNA helicase DHX15 | 1768.96 | 36.86 | 91007 | 28 | 47725702 | 47857806 | 49693378 | 48751798 | 47388102 | 49460418 | 49558526 | 44943334 | 49148714 | 45145742 | 41762806 | 43193062 | 44577294 | 38775022 | 43388310 | 51012074 | 47104654 | 44544570 | 46021050 | 46747242 | 0 | 5 | 0 | 11 | 0 | 11 | 0.88 | 0.89 | 0.98 | 0.99 |
| Q8C9B9 | Dido1 | Death-inducer obliterator 1 | 52.52 | 0.58 | 247176 | 1 | 459719.41 | 415242.25 | 612249.12 | 566930.69 | 456856.34 | 804627.81 | 514245.09 | 504927.03 | 573178.88 | 702817.56 | 311777.41 | 2 | 2 | 2 | 350266.28 | 497476.62 | 2 | 438412.22 | 636770.19 | 628040.44 | 0 | 11 | 0 | 6 | 0 | 11 | 0.26 | 0.21 | 0.88 | 0.71 |
| Q9D0D4 | Dimt1 | Probable dimethyladenosine transferase | 73.1 | 10.22 | 35274 | 2 | 2 | 2 | 2 | 2 | 247011.91 | 2 | 2 | 2 | 236158.92 | 2 | 430642.16 | 587505.25 | 345660.59 | 288312.94 | 479209.59 | 2 | 2 | 2 | 2 | 2 | 0 | 10 | 0 | 10 | 0 | 10 | 8.63 | 9.02 | 0.00 | 0.00 |
| Q9ESX5 | Dkc1 | H/ACA ribonucleoprotein complex subunit DKC1 | 491.3 | 23.18 | 57402 | 8 | 6855922 | 7131872 | 5296310 | 6087363.5 | 6350568.5 | 6325512 | 6275570 | 5156337.5 | 6520654 | 6617723.5 | 11448150 | 10178244 | 9067182 | 7312616.5 | 8735823 | 5460428 | 3884198.75 | 6075846.5 | 4183797.25 | 5407961 | 0 | 3 | 0 | 3 | 0 | 3 | 1.47 | 1.51 | 0.79 | 0.81 |
| Q8BMF4 | Dlat | Dihydrolipoyllysine-residue acetyltransferase component of pyruvate dehydrogenase complex, mitochondrial | 775.5 | 26.32 | 67942 | 13 | 28863490 | 31774864 | 31888394 | 34470678 | 37913546 | 31598538 | 30933024 | 31700982 | 32499396 | 37391942 | 47253438 | 47557834 | 54315390 | 50696094 | 54020226 | 29725828 | 29937930 | 40128374 | 34624230 | 38293722 | 0 | 9 | 0 | 11 | 0 | 11 | 1.54 | 1.55 | 1.05 | 1.05 |
| Q01205 | Dlst | Dihydrolipoyllysine-residue succinyltransferase component of 2-oxoglutarate dehydrogenase complex, mitochondrial | 868.78 | 32.16 | 48925 | 14 | 62412966 | 52688806 | 41609106 | 47722082 | 44143446 | 60143174 | 61715730 | 50255858 | 50464742 | 56512398 | 79403370 | 76885458 | 78721682 | 75284986 | 85600338 | 57553314 | 69854426 | 56178822 | 51460630 | 50691002 | 0 | 6 | 0 | 11 | 0 | 11 | 1.59 | 1.42 | 1.15 | 1.02 |
| Q9QYJ0 | Dnaja2 | DnaJ homolog subfamily A member 2 | 603.23 | 43.93 | 45746 | 12 | 3131704.25 | 4848873.5 | 4247618 | 4079323.75 | 3602772.5 | 6211460.5 | 5959330.5 | 3644132.75 | 5210506.5 | 6984792.5 | 16248890 | 15479597 | 14773827 | 16830914 | 15631852 | 4844743 | 4387009 | 3594779.5 | 4470630 | 6007625 | 0 | 7 | 0 | 6 | 0 | 7 | 3.97 | 2.82 | 1.17 | 0.83 |
| Q5U458 | Dnajc11 | DnaJ homolog subfamily C member 11 | 84.52 | 4.65 | 63233 | 2 | 1037263.44 | 1017516.69 | 986345.06 | 1208397.75 | 619565.38 | 1524811.62 | 871355 | 1082608 | 1067692.62 | 1268481.62 | 2473401.5 | 1916764.5 | 2535218 | 2474392.25 | 2228923.25 | 552010.5 | 1039655.12 | 1202565.5 | 1566417 | 1163570.88 | 0 | 8 | 0 | 12 | 0 | 12 | 2.39 | 2.00 | 1.13 | 0.95 |
| P56542 | Dnase2 | Deoxyribonuclease-2-alpha | 178.03 | 12.18 | 38811 | 4 | 3046905 | 2833234.5 | 5074117.5 | 4271193 | 6219831.5 | 1999076.38 | 2989927.75 | 4601271.5 | 3845534 | 3550079 | 450472.59 | 1073032.38 | 1237456.62 | 866165.62 | 1565803.75 | 3342988.25 | 1418778.38 | 2989654.5 | 1852334.5 | 3520259.5 | 0 | 5 | 0 | 5 | 0 | 5 | 0.24 | 0.31 | 0.61 | 0.77 |
| O88508 | Dnmt3a | DNA (cytosine-5)-methyltransferase 3A | 346.73 | 8.37 | 101627 | 6 | 2578082 | 2728381.5 | 3201360.5 | 2745604 | 2859341.75 | 2098206.75 | 2583196.25 | 3262621.75 | 2573124.5 | 2300572.25 | 913542.56 | 944162.5 | 328877.19 | 965993.62 | 561245.38 | 1596132 | 2574168 | 2037701.25 | 1854950.25 | 2259786.75 | 0 | 4 | 0 | 4 | 0 | 4 | 0.26 | 0.29 | 0.73 | 0.81 |
| Q80VJ3 | Dnph1 | 2'-deoxynucleoside 5'-phosphate N-hydrolase 1 | 51.09 | 4.62 | 18977 | 1 | 723878.5 | 628221.81 | 722847.56 | 677860.31 | 537921.38 | 753576.56 | 830447.19 | 746383.25 | 651398.81 | 637537.94 | 282006.5 | 253685.58 | 2 | 288783.31 | 249066.61 | 861323.12 | 1073947.62 | 815715.06 | 751323.69 | 735453.56 | 0 | 6 | 0 | 6 | 0 | 6 | 0.33 | 0.30 | 1.29 | 1.17 |
| Q8C147 | Dock8 | Dedicator of cytokinesis protein 8 | 789.21 | 10.81 | 238978 | 16 | 7162572 | 8235190 | 8870338 | 8755393 | 7592066 | 7829460 | 7272329 | 7824176.5 | 8026211 | 8169520 | 4926037 | 5455893.5 | 5184860.5 | 5550080.5 | 4467253 | 8000438 | 4979872 | 6214086 | 7611319.5 | 7042958.5 | 0 | 4 | 0 | 4 | 0 | 4 | 0.63 | 0.65 | 0.83 | 0.87 |
| Q61103 | Dpf2 | Zinc finger protein ubi-d4 | 204.23 | 8.7 | 44230 | 3 | 2021057.62 | 2242443.25 | 1816908.12 | 2554518.75 | 2242532 | 2774882 | 1906940 | 2185902.5 | 1436040.62 | 1939905.25 | 655157 | 1038061.62 | 430762.44 | 1187797.25 | 1281222 | 1797644 | 2333036 | 2668887.5 | 2065799.62 | 2109442.25 | 0 | 12 | 0 | 10 | 0 | 12 | 0.42 | 0.45 | 1.01 | 1.07 |
| Q99KK7 | Dpp3 | Dipeptidyl peptidase 3 | 694.81 | 31.17 | 82898 | 13 | 11403524 | 12171415 | 9164446 | 10831242 | 13670032 | 9960280 | 12681797 | 9754570 | 17428356 | 11084307 | 8568424 | 9042664 | 8663324 | 5768870 | 7595527 | 10351509 | 11849008 | 12034576 | 9078905 | 9729000 | 0 | 8 | 0 | 8 | 0 | 8 | 0.69 | 0.65 | 0.93 | 0.87 |
| Q9EPB1 | Dpp7 | Dipeptidyl peptidase 2 | 46.43 | 2 | 55114 | 1 | 1325564 | 1413193.75 | 1048353.75 | 814386.75 | 1181039.25 | 1232362 | 1008559.12 | 758895.88 | 1109484.62 | 789596.88 | 2 | 290522.53 | 275817.16 | 298784.75 | 254544.31 | 933929.88 | 1027867.69 | 952214.38 | 677195.62 | 843438.38 | 0 | 4 | 0 | 9 | 0 | 9 | 0.19 | 0.23 | 0.77 | 0.91 |
| O08553 | Dpysl2 | Dihydropyrimidinase-related protein 2 | 1857.08 | 63.29 | 62278 | 26 | 183940514 | 197865314 | 199506082 | 213494050 | 191185666 | 204487522 | 185658178 | 186302738 | 170798274 | 179177954 | 84239362 | 93429706 | 90996938 | 100954730 | 98888978 | 181593186 | 212221810 | 182453154 | 187842802 | 189860258 | 0 | 4 | 0 | 8 | 0 | 8 | 0.48 | 0.51 | 0.97 | 1.03 |
| A0JPP1 | Drap1 | Dr1-associated corepressor | 151.34 | 11.22 | 22326 | 3 | 3970337.25 | 3065957.5 | 3521112.5 | 2940893.5 | 4329134.5 | 2115163.25 | 3540594 | 3537646.5 | 4123338.75 | 2532914.25 | 525952.25 | 1961701.88 | 1539841.5 | 752996.88 | 1427972.75 | 4055291.25 | 1803777 | 2340476 | 2886541 | 3029603 | 0 | 5 | 0 | 9 | 0 | 9 | 0.35 | 0.39 | 0.79 | 0.89 |
| P32233 | Drg1 | Developmentally-regulated GTP-binding protein 1 | 354.77 | 16.35 | 40512 | 6 | 1374078.5 | 1492018.5 | 1484171.75 | 792938.81 | 1299491.75 | 2174285.5 | 1797074 | 1422477.25 | 916385.44 | 1246717.62 | 3040175.5 | 3152560.5 | 2579766.5 | 2471443.75 | 2619185 | 1313201.88 | 1700365.5 | 1332502.5 | 1173468.5 | 1679211.5 | 0 | 10 | 0 | 11 | 0 | 11 | 2.15 | 1.83 | 1.12 | 0.95 |
| Q3UIR3 | Dtx3l | E3 ubiquitin-protein ligase DTX3L | 251.72 | 8.96 | 83044 | 5 | 2934914.75 | 2278330.5 | 2402400 | 3082405.25 | 3344188.25 | 3569071.75 | 2609245 | 3647822.25 | 3079284.25 | 3088358.5 | 343808.38 | 1032422.75 | 546028.75 | 129778.22 | 701802.62 | 1487467.5 | 3209973 | 2707638.5 | 3046488 | 2762401.5 | 0 | 11 | 0 | 6 | 0 | 11 | 0.20 | 0.17 | 0.94 | 0.83 |
| Q9D7X3 | Dusp3 | Dual specificity protein phosphatase 3 | 475.25 | 50.81 | 20472 | 7 | 9857928 | 11076486 | 11509561 | 10437444 | 10983042 | 10484899 | 11010231 | 10318837 | 9993306 | 9967412 | 8977778 | 9555490 | 8397068 | 9744912 | 8440193 | 10662021 | 9921944 | 11822272 | 11059273 | 12052829 | 0 | 6 | 0 | 9 | 0 | 9 | 0.84 | 0.87 | 1.03 | 1.07 |
| Q9JHU4 | Dync1h1 | Cytoplasmic dynein 1 heavy chain 1 | 6142.521 | 27.07 | 532045 | 108 | 145004402 | 139568482 | 146092610 | 143011314 | 134834354 | 119510082 | 135156370 | 142608082 | 141465986 | 140054114 | 97889538 | 117178562 | 104135938 | 77970714 | 96465922 | 132771786 | 114973050 | 121877570 | 114409282 | 117620170 | 0 | 0 | 0 | 2 | 0 | 2 | 0.70 | 0.73 | 0.85 | 0.89 |
| P42125 | Eci1 | Enoyl-CoA delta isomerase 1, mitochondrial | 465.66 | 47.75 | 32250 | 7 | 3433240.5 | 3336398.25 | 4370804.5 | 2670969.25 | 3691248.5 | 4118672.5 | 4342862.5 | 4328974.5 | 5047957 | 5524563.5 | 7395487 | 6793461 | 8554648 | 9825218 | 9418746 | 2214910 | 2998927.75 | 6200716.5 | 6207767 | 5054152.5 | 0 | 9 | 0 | 11 | 0 | 11 | 2.40 | 1.80 | 1.30 | 0.97 |
| P10126 | Eef1a1 | Elongation factor 1-alpha 1 | 2160.719 | 61.9 | 50114 | 34 | 1030543618 | 1170794114 | 1197046914 | 1041232386 | 1137700610 | 979320002 | 1061729026 | 1169864706 | 1206226946 | 1098271234 | 1806016386 | 2043400578 | 1524526850 | 1392806274 | 1625211906 | 1178606850 | 1028110082 | 1149407106 | 1067687234 | 1210632066 | 0 | 11 | 0 | 10 | 0 | 11 | 1.50 | 1.52 | 1.01 | 1.02 |
| O70251 | Eef1b | Elongation factor 1-beta | 586.81 | 39.56 | 24694 | 9 | 13771842 | 12854370 | 12870794 | 14759090 | 10387106 | 13726210 | 12097874 | 12449892 | 12244898 | 13739346 | 10792082 | 10154066 | 9624926 | 10784266 | 10200162 | 14701659 | 18835906 | 15644208 | 16918427 | 15086946 | 0 | 1 | 0 | 1 | 0 | 1 | 0.81 | 0.81 | 1.26 | 1.26 |
| P57776 | Eef1d | Elongation factor 1-delta | 848.2 | 54.09 | 31293 | 13 | 107040474 | 101984394 | 112340730 | 117554170 | 100273450 | 113836402 | 102780050 | 93984738 | 93946170 | 107623186 | 74352722 | 82442754 | 77204906 | 74214418 | 71849442 | 113664906 | 120213322 | 124013970 | 123229770 | 125278306 | 0 | 1 | 0 | 1 | 0 | 1 | 0.70 | 0.74 | 1.12 | 1.18 |
| P58252 | Eef2 | Elongation factor 2 | 3438.76 | 62.35 | 95314 | 54 | 329060322 | 351599810 | 367255362 | 346052962 | 348820674 | 353570722 | 306345506 | 353023106 | 377328962 | 367936002 | 493306274 | 474916226 | 397677058 | 414912578 | 381094370 | 348080866 | 392176906 | 332216066 | 393574690 | 393441154 | 0 | 7 | 0 | 8 | 0 | 8 | 1.24 | 1.23 | 1.07 | 1.06 |
| Q4FZY0 | Efhd2 | EF-hand domain-containing protein D2 | 1184.57 | 59.83 | 26759 | 18 | 67667330 | 70539330 | 81479018 | 61647334 | 77141626 | 50672538 | 67809018 | 73450810 | 80635314 | 72807418 | 43513966 | 47312902 | 40859390 | 45750734 | 45903554 | 58002854 | 44637410 | 67405514 | 62757134 | 66464482 | 0 | 3 | 0 | 4 | 0 | 4 | 0.62 | 0.65 | 0.83 | 0.87 |
| Q641Z6 | Ehd1 | EH domain-containing protein 1 | 1300.64 | 40.64 | 60603 | 22 | 19867912 | 20930430 | 17494354 | 17740606 | 19019650 | 20654194 | 18432034 | 18865630 | 18848330 | 20025234 | 24294008 | 29272462 | 24268062 | 21376980 | 23749172 | 19046954 | 20437218 | 17560422 | 20522000 | 20101454 | 0 | 9 | 0 | 10 | 0 | 10 | 1.29 | 1.27 | 1.03 | 1.01 |
| Q9EQP2 | Ehd4 | EH domain-containing protein 4 | 897.71 | 33.46 | 61481 | 16 | 25445490 | 22944646 | 25472126 | 28310800 | 25653818 | 20906108 | 23713010 | 23802804 | 27698488 | 26318510 | 14672710 | 19894054 | 18987064 | 15068005 | 20059944 | 21992832 | 18147290 | 19688014 | 22885084 | 21309906 | 0 | 3 | 0 | 3 | 0 | 3 | 0.69 | 0.72 | 0.81 | 0.85 |
| Q99LD9 | Eif2b2 | Translation initiation factor eIF-2B subunit beta | 43.31 | 4.56 | 38898 | 1 | 358795 | 413217.25 | 278451.28 | 369331.28 | 258486.33 | 333236.28 | 355701.22 | 2 | 2 | 2 | 360525.62 | 685621.12 | 554450.31 | 536127.88 | 491320.31 | 507121.31 | 2 | 2 | 364145.09 | 475630.75 | 0 | 8 | 0 | 12 | 0 | 12 | 1.65 | 2.64 | 0.50 | 0.80 |
| Q99L45 | Eif2s2 | Eukaryotic translation initiation factor 2 subunit 2 | 677.22 | 48.64 | 38092 | 13 | 30187998 | 35690918 | 28081482 | 22938526 | 23885110 | 31115042 | 30063770 | 30930912 | 28557302 | 32691598 | 18887052 | 17339320 | 15913156 | 18329932 | 20291966 | 32045076 | 30169030 | 32022004 | 33022038 | 35767238 | 0 | 5 | 0 | 5 | 0 | 5 | 0.65 | 0.59 | 1.16 | 1.06 |
| Q9Z0N1 | Eif2s3x | Eukaryotic translation initiation factor 2 subunit 3, X-linked | 993.83 | 45.13 | 51065 | 15 | 41026146 | 37746286 | 30992512 | 35576362 | 33689946 | 39430634 | 40416566 | 47899294 | 42414654 | 42234882 | 62572882 | 57265502 | 53799214 | 51660934 | 58259138 | 41309214 | 43896446 | 45763762 | 44792794 | 47832466 | 0 | 7 | 0 | 7 | 0 | 7 | 1.58 | 1.34 | 1.25 | 1.05 |
| Q8R1B4 | Eif3c | Eukaryotic translation initiation factor 3 subunit C | 1215.83 | 24.59 | 105531 | 22 | 26428630 | 28545168 | 30785486 | 30230136 | 29459800 | 27935156 | 29476892 | 32486432 | 34040622 | 30967914 | 44159542 | 42834982 | 38036718 | 37864214 | 39112274 | 27397046 | 28968726 | 29393904 | 27916540 | 30374588 | 0 | 10 | 0 | 4 | 0 | 10 | 1.39 | 1.30 | 0.99 | 0.93 |
| Q9Z1D1 | Eif3g | Eukaryotic translation initiation factor 3 subunit G | 412.07 | 21.25 | 35638 | 7 | 13172290 | 12824299 | 13440771 | 16176288 | 12311142 | 12888302 | 12036649 | 13953353 | 11801702 | 15783256 | 11248912 | 10792458 | 9923438 | 8955490 | 10937724 | 12803637 | 11926550 | 14637110 | 17033636 | 15425285 | 0 | 11 | 0 | 9 | 0 | 11 | 0.76 | 0.78 | 1.06 | 1.08 |
| B0BNA7 | Eif3i | Eukaryotic translation initiation factor 3 subunit I | 460.44 | 31.69 | 36461 | 9 | 2621145 | 2257842 | 2700000.5 | 2388992 | 2740422.5 | 2645388 | 2068384.62 | 2512597.75 | 3021868.5 | 1766340.38 | 7253555.5 | 5549140 | 6511433 | 6064594 | 6546310 | 3024335.5 | 3507737 | 3520692.5 | 3247407.5 | 2735716.5 | 0 | 1 | 0 | 1 | 0 | 1 | 2.51 | 2.66 | 1.26 | 1.33 |
| Q8QZY1 | Eif3l | Eukaryotic translation initiation factor 3 subunit L | 966.92 | 28.37 | 66613 | 15 | 5480722.5 | 5829148.5 | 8327198 | 8944528 | 8713883 | 4721745 | 6392368 | 8097364 | 10770226 | 7273625 | 11395667 | 13219535 | 14181032 | 12278080 | 15104551 | 6440975.5 | 6879478.5 | 7448142.5 | 7800854.5 | 8006947 | 0 | 10 | 0 | 12 | 0 | 12 | 1.77 | 1.78 | 0.98 | 0.98 |
| Q62448 | Eif4g2 | Eukaryotic translation initiation factor 4 gamma 2 | 941.63 | 22.08 | 102106 | 20 | 10674454 | 11612718 | 12654184 | 8810106 | 13159489 | 8165306 | 11064697 | 10153783 | 12431849 | 6713764 | 21811288 | 28220834 | 20006296 | 17815760 | 21311028 | 11309470 | 7730185 | 8962351 | 6634666 | 9799681 | 0 | 4 | 0 | 9 | 0 | 9 | 1.92 | 2.25 | 0.78 | 0.92 |
| Q5PPK9 | Eipr1 | EARP and GARP complex-interacting protein 1 | 47.73 | 2.33 | 43157 | 1 | 1313429 | 1378099.88 | 1441632.38 | 1912876.12 | 1297479.12 | 1302592.25 | 1347871.62 | 1246203.88 | 1672854.38 | 1849405.75 | 888687 | 2 | 737996.25 | 2 | 2 | 1206113.5 | 862123 | 1289087.75 | 1354703.5 | 1374582.12 | 0 | 4 | 0 | 7 | 0 | 7 | 0.22 | 0.22 | 0.83 | 0.82 |
| Q8BPU7 | Elmo1 | Engulfment and cell motility protein 1 | 455.51 | 12.65 | 83936 | 9 | 10474585 | 11417404 | 12314479 | 12679575 | 11089144 | 9561596 | 10135867 | 11454115 | 11787528 | 11515365 | 5902627 | 8792404 | 5802504 | 2810125.75 | 8685426 | 10600521 | 8862784 | 8724269 | 8454968 | 6767100 | 0 | 1 | 0 | 2 | 0 | 2 | 0.55 | 0.59 | 0.75 | 0.80 |
| Q9JLJ5 | Elovl1 | Elongation of very long chain fatty acids protein 1 | 161.51 | 12.54 | 32678 | 3 | 232668.17 | 162598.12 | 146736.78 | 63342.02 | 248787.86 | 128353.64 | 2 | 405928.25 | 331378.53 | 60543.55 | 1397396.25 | 1239972.75 | 633588 | 1194960.5 | 978077.25 | 191717.67 | 260532.53 | 335626.31 | 432603.5 | 621376.75 | 0 | 2 | 0 | 5 | 0 | 5 | 6.37 | 5.88 | 2.16 | 1.99 |
| Q7TT37 | Elp1 | Elongator complex protein 1 | 94.95 | 1.5 | 149584 | 2 | 1031274.44 | 1091911.75 | 1102650.5 | 2 | 1134935 | 763406.69 | 1065633.62 | 975952.38 | 953457.75 | 526756.31 | 1702377.88 | 1914105.25 | 1894216.88 | 1295163.5 | 1324139 | 1253002.25 | 771725.19 | 2 | 648942.75 | 732505.38 | 0 | 8.5 | 0 | 9 | 0 | 9 | 1.86 | 1.90 | 0.78 | 0.79 |
| Q8C7X2 | Emc1 | ER membrane protein complex subunit 1 | 1552.84 | 33.3 | 111605 | 26 | 18481804 | 17367878 | 19549058 | 16301832 | 15415819 | 19980334 | 20930014 | 16942430 | 16584822 | 19260798 | 34832042 | 31491022 | 30645086 | 31147820 | 33525678 | 21983420 | 21412606 | 22343084 | 21251526 | 19101662 | 0 | 1 | 0 | 3 | 0 | 3 | 1.86 | 1.73 | 1.22 | 1.13 |
| O35130 | Emg1 | Ribosomal RNA small subunit methyltransferase NEP1 | 233.85 | 24.59 | 26974 | 4 | 426419.81 | 1011391.56 | 737607.38 | 536937.12 | 555231.12 | 736113.19 | 550164 | 923984.62 | 604435.62 | 725618.69 | 2476015.75 | 2024096.12 | 2218142.75 | 2108873.25 | 2135799 | 270581.66 | 404482.84 | 941546.81 | 632047.25 | 370633.56 | 0 | 7 | 0 | 7 | 0 | 7 | 3.36 | 3.10 | 0.80 | 0.74 |
| Q8C522 | Endod1 | Endonuclease domain-containing 1 protein | 56.85 | 2.2 | 55262 | 1 | 2 | 2 | 2 | 2 | 2 | 2 | 2 | 2 | 2 | 2 | 315772.56 | 381703.91 | 737180.94 | 646391.44 | 684715.44 | 2 | 2 | 2 | 2 | 427658.72 | 0 | 10 | 0 | 10 | 0 | 10 | 276576.43 | 276576.43 | 42766.67 | 42766.67 |
| Q8CGC7 | Eprs | Bifunctional glutamate/proline--tRNA ligase | 2976.59 | 37.43 | 170079 | 47 | 69261714 | 79808250 | 73123498 | 78397066 | 72556914 | 75767354 | 73700258 | 73577666 | 83873530 | 83770346 | 154616690 | 150332530 | 132186002 | 144640738 | 126726594 | 79761218 | 94751962 | 91702866 | 86288674 | 89401194 | 0 | 1 | 0 | 2 | 0 | 2 | 1.90 | 1.81 | 1.18 | 1.13 |
| Q9DC16 | Ergic1 | Endoplasmic reticulum-Golgi intermediate compartment protein 1 | 152.45 | 9.66 | 32562 | 3 | 3864363.75 | 4039279.5 | 4870981 | 4051686 | 3267568 | 4070570 | 4009455.5 | 4029218 | 4189291.5 | 4597938 | 6824986 | 7070171 | 8384725 | 7331553 | 8133665 | 4438117 | 4922859 | 5580743 | 5117602 | 5355347.5 | 0 | 1 | 0 | 1 | 0 | 1 | 1.88 | 1.81 | 1.26 | 1.22 |
| Q9CQE7 | Ergic3 | Endoplasmic reticulum-Golgi intermediate compartment protein 3 | 101.81 | 4.44 | 43208 | 2 | 239605.31 | 38680.39 | 73586.16 | 458031.59 | 78549.29 | 40155.32 | 2 | 93574.88 | 88107.98 | 544705.62 | 874711.56 | 763483 | 1259647.88 | 1116063.88 | 1332706.88 | 33148.82 | 28714.62 | 344427.19 | 242481.55 | 55899.18 | 0 | 9 | 0 | 12 | 0 | 12 | 6.02 | 6.97 | 0.79 | 0.92 |
| Q3UVK0 | Ermp1 | Endoplasmic reticulum metallopeptidase 1 | 206.72 | 3.34 | 100148 | 3 | 161155.33 | 2 | 505708.59 | 2 | 2 | 2 | 720145.38 | 147366.53 | 484164.84 | 812815.81 | 1083981.5 | 995245.62 | 1707930.25 | 1775645.5 | 1830113.75 | 865565.75 | 158379.88 | 743712.69 | 597387.44 | 540757.5 | 0 | 2 | 0 | 8 | 0 | 8 | 11.09 | 3.42 | 4.36 | 1.34 |
| Q9R0P3 | Esd | S-formylglutathione hydrolase | 1127.09 | 71.63 | 31320 | 17 | 65296098 | 76205842 | 46468122 | 51129030 | 57718778 | 73324242 | 75570362 | 63368618 | 69002194 | 71205010 | 184859842 | 195729378 | 162199010 | 148743954 | 158067074 | 78615178 | 86860162 | 105089498 | 92808050 | 95206258 | 0 | 0 | 0 | 0 | 0 | 0 | 2.86 | 2.41 | 1.54 | 1.30 |
| Q3U7R1 | Esyt1 | Extended synaptotagmin-1 | 708.63 | 17.67 | 121554 | 12 | 9560872 | 8811495 | 5688600.5 | 6542616.5 | 5472162 | 8356517 | 9430540 | 6008448 | 7311805.5 | 8862323 | 24161692 | 17831466 | 15742915 | 13596386 | 15048117 | 7582002.5 | 8500848 | 8491353 | 9051297 | 7323864 | 0 | 9 | 0 | 11 | 0 | 11 | 2.39 | 2.16 | 1.14 | 1.02 |
| P20934 | Evi2a | Protein EVI2A | 165.69 | 9.42 | 24087 | 2 | 119787.56 | 429897.19 | 103185.62 | 308999.88 | 2 | 455494 | 131082.31 | 646475.25 | 174416.25 | 567103.06 | 1167299.75 | 1300729.25 | 1863121.12 | 2205133.25 | 1681069.88 | 2 | 2 | 2 | 363274.22 | 2 | 0 | 6 | 0 | 2 | 0 | 6 | 8.54 | 4.16 | 0.38 | 0.18 |
| Q8VEG4 | Exd2 | Exonuclease 3'-5' domain-containing protein 2 | 49.04 | 2 | 74332 | 1 | 222700.02 | 182885.28 | 225938 | 234069.62 | 149773.81 | 264727.19 | 165069.14 | 185012.58 | 225833.27 | 197412.7 | 2 | 137431.47 | 2 | 2 | 101095.98 | 199459.58 | 177371.64 | 344801.78 | 2 | 124948.03 | 0 | 8 | 0 | 9 | 0 | 9 | 0.23 | 0.23 | 0.83 | 0.82 |
| O54924 | Exoc8 | Exocyst complex component 8 | 76.12 | 3.35 | 81043 | 2 | 1100021.62 | 819506.5 | 817617.5 | 991021.44 | 900820.56 | 1090983.75 | 828897.94 | 818560.44 | 898674.38 | 933237.56 | 671809.69 | 2 | 586981.44 | 757944.44 | 532290.06 | 858672.44 | 1469939.88 | 2 | 992163.44 | 872366.88 | 0 | 12 | 0 | 12 | 0 | 12 | 0.55 | 0.56 | 0.91 | 0.92 |
| Q9DAA6 | Exosc1 | Exosome complex component CSL4 | 39.81 | 5.64 | 21424 | 1 | 374710.69 | 435638.91 | 350818.19 | 467606.12 | 351473.19 | 443275.66 | 2 | 2 | 396776.75 | 414255.28 | 688142.31 | 631663.06 | 658752.69 | 566467.94 | 874838.88 | 499370.97 | 469931.16 | 411223.62 | 319517.53 | 516584.34 | 0 | 7 | 0 | 5 | 0 | 7 | 1.73 | 2.73 | 1.12 | 1.77 |
| P35505 | Fah | Fumarylacetoacetase | 136.15 | 8.81 | 46176 | 3 | 2168883.25 | 3525974.25 | 3840234.5 | 3910919 | 3219120 | 3889039 | 3035370 | 3492837.25 | 3775399.25 | 4430304.5 | 6703918.5 | 6432458 | 4880465 | 5707814.5 | 5494775 | 3935399 | 3314026 | 5310956.5 | 3996080.5 | 2235877 | 0 | 7 | 0 | 11 | 0 | 11 | 1.75 | 1.57 | 1.13 | 1.01 |
| Q4QQV3 | Fam162a | Protein FAM162A | 67.82 | 15.48 | 17827 | 2 | 139769.56 | 405552.72 | 412854.28 | 505323.53 | 141133.23 | 335212.5 | 292857.47 | 466097.34 | 461709.72 | 749852.25 | 1208003.25 | 1038436.88 | 1530561.25 | 1643258.62 | 1716172.25 | 312297.22 | 298463.75 | 578279 | 392708.16 | 2 | 0 | 11 | 0 | 8 | 0 | 11 | 4.45 | 3.10 | 0.99 | 0.69 |
| Q9D8B6 | Fam210b | Protein FAM210B, mitochondrial | 52.75 | 6.84 | 20344 | 1 | 2 | 2 | 2 | 2 | 2 | 2 | 2 | 2 | 2 | 2 | 541837.56 | 218933.42 | 304704.22 | 247822.86 | 242151.64 | 2 | 2 | 2 | 2 | 2 | 0 | 12.5 | 0 | 12.5 | 0 | 12.5 | 155544.97 | 155544.97 | 1.00 | 1.00 |
| Q921M7 | Fam49b | CYFIP-related Rac1 interactor B | 689.92 | 38.89 | 36776 | 10 | 12861141 | 14445928 | 17839438 | 17254798 | 18573588 | 13732099 | 15099142 | 15022374 | 16201185 | 13642780 | 8373122 | 10964909 | 11621341 | 9739654 | 7291179 | 15004552 | 14686504 | 14977756 | 10381945 | 14345370 | 0 | 7 | 0 | 8 | 0 | 8 | 0.59 | 0.65 | 0.86 | 0.94 |
| Q922J9 | Far1 | Fatty acyl-CoA reductase 1 | 1092.32 | 49.51 | 59435 | 19 | 16641804 | 10686342 | 10341209 | 13125131 | 12728094 | 16481948 | 14269302 | 13616062 | 11013636 | 11615346 | 25758894 | 29708074 | 31270736 | 29192340 | 32006064 | 15893355 | 16773403 | 14739104 | 14015383 | 13724871 | 0 | 4 | 0 | 6 | 0 | 6 | 2.33 | 2.21 | 1.18 | 1.12 |
| Q9WUA2 | Farsb | Phenylalanine--tRNA ligase beta subunit | 656.07 | 23.94 | 65697 | 14 | 21526610 | 23939552 | 25638974 | 24736648 | 22805866 | 25088838 | 19701588 | 20885812 | 22240146 | 25957974 | 30056540 | 29085762 | 28121212 | 30596406 | 28850276 | 21601754 | 28802046 | 28916514 | 28906888 | 27575658 | 0 | 4 | 0 | 3 | 0 | 4 | 1.24 | 1.29 | 1.14 | 1.19 |
| P20491 | Fcer1g | High affinity immunoglobulin epsilon receptor subunit gamma | 172.4 | 25.58 | 9652 | 3 | 7293618 | 10586340 | 7397920 | 8877770 | 3926941.75 | 10143556 | 8692348 | 10305551 | 6647899 | 10310799 | 38111330 | 33805394 | 41004698 | 33926402 | 44153050 | 6364498 | 8049496.5 | 12716928 | 10700599 | 12268295 | 0 | 6 | 0 | 9 | 0 | 9 | 5.02 | 4.14 | 1.32 | 1.09 |
| P39749 | Fen1 | Flap endonuclease 1 | 588.82 | 22.75 | 42315 | 9 | 18350324 | 18601704 | 25782404 | 22303604 | 22462160 | 15569020 | 15663790 | 20306910 | 16889446 | 17730516 | 12328346 | 12408609 | 8817527 | 8912347 | 12664128 | 18775342 | 12994049 | 17653564 | 11721029 | 17579514 | 0 | 2 | 0 | 10 | 0 | 10 | 0.51 | 0.64 | 0.73 | 0.91 |
| P84817 | Fis1 | Mitochondrial fission 1 protein | 421.25 | 44.74 | 16995 | 6 | 2273554.75 | 3469042.5 | 4548336.5 | 3376777.5 | 4943141.5 | 2082440.38 | 3233517.5 | 4000489 | 2830745.75 | 1699995.12 | 385355.56 | 986572.75 | 778236.81 | 999233.12 | 894572.5 | 3490586.5 | 2396232.25 | 2365250.5 | 3037691 | 3024747.5 | 0 | 7 | 0 | 11 | 0 | 11 | 0.22 | 0.29 | 0.77 | 1.03 |
| P26883 | Fkbp1a | Peptidyl-prolyl cis-trans isomerase FKBP1A | 143.76 | 25 | 11923 | 2 | 2 | 2 | 2 | 2 | 2 | 1336983.12 | 1785448.75 | 2 | 2 | 2 | 5570327 | 5588228.5 | 3596534.25 | 4939032.5 | 5138845.5 | 2 | 2 | 1320234.75 | 2 | 2 | 0 | 10 | 0 | 9 | 0 | 10 | 2483296.78 | 7.95 | 132024.28 | 0.42 |
| O08917 | Flot1 | Flotillin-1 | 333.29 | 17.99 | 47513 | 6 | 1468992.62 | 570003 | 425506.53 | 135190.97 | 2 | 1007027.25 | 2 | 2 | 349074.38 | 400111.72 | 7496442 | 4268402 | 5295726.5 | 5660567 | 5834044.5 | 1121097.62 | 1176207.75 | 495300.97 | 397656.06 | 2 | 0 | 11.5 | 0 | 7 | 0 | 11.5 | 10.98 | 16.26 | 1.23 | 1.82 |
| P35922 | Fmr1 | Synaptic functional regulator FMR1 | 156.42 | 6.03 | 68989 | 3 | 1937220.38 | 2218149 | 2005645.5 | 3007861.5 | 1934500.12 | 2441329 | 2192000.75 | 1954781.12 | 1715698 | 2069097.12 | 221751.55 | 273700.25 | 535028.88 | 524237.44 | 2 | 2098673.75 | 2006420.38 | 1607761 | 1635922.88 | 1557622.38 | 0 | 6 | 0 | 6 | 0 | 6 | 0.14 | 0.15 | 0.80 | 0.86 |
| Q80TY0 | Fnbp1 | Formin-binding protein 1 | 690.35 | 18.02 | 71344 | 12 | 23542834 | 19007664 | 24146198 | 21655404 | 19724946 | 16144865 | 20264750 | 22647414 | 16896724 | 18615546 | 10696158 | 15608129 | 10555638 | 7688807 | 12067890 | 23309314 | 11560294 | 18975616 | 16343285 | 16263637 | 0 | 3 | 0 | 10 | 0 | 10 | 0.52 | 0.60 | 0.80 | 0.91 |
| Q8K385 | Frrs1 | Ferric-chelate reductase 1 | 209.81 | 8.45 | 66047 | 4 | 19110252 | 14441198 | 10392865 | 12435464 | 12870888 | 15825893 | 15784801 | 10422126 | 12082153 | 13871428 | 5279406 | 5748655 | 4866836 | 6404150 | 5327183 | 14310594 | 12359366 | 8705210 | 8323484 | 9933136 | 0 | 4 | 0 | 5 | 0 | 5 | 0.40 | 0.41 | 0.77 | 0.79 |
| P09528 | Fth1 | Ferritin heavy chain | 641.35 | 56.04 | 21067 | 11 | 21131550 | 18141730 | 19321128 | 34893818 | 29652728 | 27886138 | 25057054 | 55739530 | 54782290 | 55680038 | 171254306 | 172530250 | 283451874 | 253427186 | 282415842 | 384739938 | 554799298 | 602124482 | 627704770 | 567048514 | 0 | 0 | 0 | 0 | 0 | 0 | 9.45 | 5.31 | 22.22 | 12.49 |
| P29391 | Ftl1 | Ferritin light chain 1 | 1017.49 | 73.22 | 20802 | 15 | 35454822 | 27618676 | 45772886 | 53905598 | 52433102 | 40681886 | 38502082 | 74889490 | 81044098 | 82448042 | 105100170 | 116605522 | 150582738 | 154706914 | 153572834 | 551825986 | 664956482 | 658574466 | 716652034 | 665532546 | 0 | 0 | 0 | 0 | 0 | 0 | 3.16 | 2.14 | 15.14 | 10.26 |
| Q8R2K1 | Fuom | Fucose mutarotase | 76.51 | 7.84 | 16805 | 1 | 2 | 149430.55 | 2 | 49925.97 | 148134.98 | 2 | 2 | 2 | 175643.7 | 177203.09 | 784228.19 | 413253.25 | 558491.44 | 611284.88 | 404531.41 | 2 | 2 | 132026.81 | 97612.95 | 2 | 0 | 9 | 0 | 10.5 | 0 | 10.5 | 7.98 | 7.86 | 0.66 | 0.65 |
| Q00612 | G6pdx | Glucose-6-phosphate 1-dehydrogenase X | 1459.48 | 53.01 | 59263 | 27 | 77125362 | 81088738 | 74658970 | 74655490 | 76736322 | 74286426 | 86203354 | 89308994 | 82921066 | 81973754 | 150968338 | 156430050 | 119277306 | 100537866 | 119809418 | 107510306 | 97650178 | 102922778 | 105539194 | 110128682 | 0 | 0 | 0 | 0 | 0 | 0 | 1.68 | 1.56 | 1.36 | 1.26 |
| P60517 | Gabarap | Gamma-aminobutyric acid receptor-associated protein | 183.62 | 21.37 | 13918 | 3 | 5569862.5 | 5121677 | 5286218.5 | 4774505 | 4648064.5 | 5060018.5 | 5348069.5 | 5577156.5 | 5010179 | 6055656.5 | 8922369 | 7807453 | 7833266 | 6261737 | 8233382 | 3973591.5 | 4969096.5 | 5477909 | 7414601 | 4963376.5 | 0 | 12 | 0 | 8 | 0 | 12 | 1.44 | 1.44 | 1.06 | 0.99 |
| Q9R0N0 | Galk1 | Galactokinase | 217.63 | 10.46 | 42295 | 4 | 10940886 | 10992705 | 11593474 | 10197334 | 10297636 | 9270073 | 11020098 | 12402017 | 10719239 | 10747313 | 13167884 | 13685344 | 13082364 | 13585244 | 13489448 | 10090397 | 12080085 | 12447714 | 12229473 | 15741454 | 0 | 5 | 0 | 6 | 0 | 6 | 1.24 | 1.24 | 1.16 | 1.16 |
| Q571E4 | Galns | N-acetylgalactosamine-6-sulfatase | 314.87 | 13.46 | 57673 | 6 | 3667255.5 | 3763997.25 | 4582793.5 | 4868908.5 | 7300380 | 2631194.5 | 4249218.5 | 4443266 | 5696569 | 3422548.25 | 879383.75 | 899451.25 | 421022.31 | 1468553.5 | 1383808 | 3808645.75 | 3649448.5 | 5239998 | 5606179.5 | 4310703 | 0 | 12 | 0 | 10 | 0 | 12 | 0.21 | 0.25 | 0.94 | 1.11 |
| Q6P6V1 | Galnt11 | Polypeptide N-acetylgalactosaminyltransferase 11 | 40.34 | 1.15 | 69039 | 1 | 4871399.5 | 4628804 | 6119983 | 6543670 | 5589983.5 | 5645836 | 4607888 | 4855838.5 | 4994786.5 | 4605390.5 | 2111794.5 | 1950528.5 | 2592086 | 3194829.25 | 2603715.5 | 3667474 | 4564209.5 | 4307746.5 | 4141163.25 | 4216927 | 0 | 10 | 0 | 6 | 0 | 10 | 0.45 | 0.50 | 0.75 | 0.84 |
| Q80VA0 | Galnt7 | N-acetylgalactosaminyltransferase 7 | 398.49 | 15.68 | 75419 | 8 | 1794975.75 | 3216685 | 6382163.5 | 5983846 | 6342949.5 | 2224420.25 | 2008436.25 | 6751366 | 6394791.5 | 5229070.5 | 10799148 | 9374280 | 15640026 | 15476401 | 15987120 | 1218411.88 | 1388120.38 | 6676885 | 7350081 | 6648293 | 0 | 10 | 0 | 12 | 0 | 12 | 2.84 | 2.98 | 0.98 | 1.03 |
| Q8BHN3 | Ganab | Neutral alpha-glucosidase AB | 1763.41 | 37.5 | 106911 | 32 | 43316250 | 47558126 | 48293802 | 39643114 | 43920106 | 37777278 | 43530970 | 51039322 | 48575054 | 44223650 | 77357578 | 86386914 | 84247130 | 82613770 | 90453282 | 39537282 | 35388310 | 44553694 | 44031062 | 39572930 | 0 | 6 | 0 | 7 | 0 | 7 | 1.89 | 1.87 | 0.91 | 0.90 |
| Q01514 | Gbp1 | Guanylate-binding protein 1 | 130.65 | 5.26 | 67712 | 2 | 1807094.5 | 1588703.12 | 828684.75 | 1484069.38 | 1501579.62 | 1256166.5 | 1462324.75 | 1318223.5 | 1329822 | 1039228.88 | 518835.59 | 644644.38 | 715289.25 | 227331.14 | 669539.44 | 1358009 | 1400298.5 | 1077670.38 | 1122079.75 | 998651.44 | 0 | 5 | 0 | 10 | 0 | 10 | 0.38 | 0.43 | 0.83 | 0.93 |
| O88986 | Gcat | 2-amino-3-ketobutyrate coenzyme A ligase, mitochondrial | 259.46 | 13.46 | 44931 | 3 | 579542.12 | 675976.81 | 500960.62 | 713589.75 | 312979.94 | 692680 | 288069.56 | 596076.25 | 462179.91 | 758542.31 | 1978093.12 | 1488005 | 1720661.75 | 2476862 | 1706374 | 365969.59 | 409034.56 | 537653.12 | 651583.81 | 656130.06 | 0 | 10 | 0 | 10 | 0 | 10 | 3.37 | 3.35 | 0.94 | 0.94 |
| O09172 | Gclm | Glutamate--cysteine ligase regulatory subunit | 282.64 | 22.99 | 30535 | 5 | 2476578.25 | 2534911.5 | 2105851.75 | 2268303 | 3340085 | 3564871.75 | 2651398 | 2589788.75 | 4122152.5 | 2757284.5 | 13735158 | 15253114 | 12957263 | 17591150 | 16191247 | 4133617.25 | 3622444.25 | 6385710.5 | 5451630.5 | 4619630.5 | 0 | 0 | 0 | 1 | 0 | 1 | 5.95 | 4.83 | 1.90 | 1.54 |
| Q66H63 | Gdap2 | Ganglioside-induced differentiation-associated-protein 2 | 40.56 | 1.61 | 56130 | 1 | 680279.62 | 646396.44 | 665741.56 | 746857.69 | 836027.19 | 658013.19 | 867614.62 | 771976.44 | 864318.5 | 846694.38 | 265529.03 | 423426.72 | 515671.12 | 604176.25 | 509361 | 635774 | 462448.59 | 535463.44 | 2 | 880435.12 | 0 | 5 | 0 | 5 | 0 | 5 | 0.65 | 0.58 | 0.70 | 0.63 |
| Q5BJP6 | Gfm2 | Ribosome-releasing factor 2, mitochondrial | 89.17 | 2.7 | 85914 | 2 | 350334.75 | 326665.47 | 2 | 422326.72 | 434501.06 | 368654.44 | 541189.62 | 446972.91 | 530317.81 | 530660.94 | 972851.5 | 1102773 | 1053843.12 | 1084255.5 | 712535.94 | 592290.94 | 574670.31 | 595633.31 | 486128 | 804274 | 0 | 0 | 0 | 3 | 0 | 3 | 3.21 | 2.04 | 1.99 | 1.26 |
| Q91VC9 | Ghitm | Growth hormone-inducible transmembrane protein | 313.28 | 24.86 | 37275 | 6 | 2576312 | 3687830.75 | 6075269 | 6524067.5 | 6100314 | 2906640.25 | 4822983 | 6980171 | 6177216.5 | 7413621 | 9384260 | 10679017 | 14894054 | 15683354 | 16308080 | 4028501.5 | 2992811.75 | 7750532 | 6651835.5 | 8924991 | 0 | 7 | 0 | 10 | 0 | 10 | 2.68 | 2.37 | 1.22 | 1.07 |
| P23780 | Glb1 | Beta-galactosidase | 833.69 | 23.34 | 73121 | 14 | 42264458 | 35933210 | 44264378 | 35252802 | 47633602 | 36129026 | 37455510 | 26782392 | 36770010 | 27906636 | 5706196 | 7500823 | 8964626 | 6712162 | 5864396 | 39855654 | 24691772 | 32939704 | 31199270 | 30851970 | 0 | 2 | 0 | 11 | 0 | 11 | 0.17 | 0.21 | 0.78 | 0.97 |
| Q61543 | Glg1 | Golgi apparatus protein 1 | 1673.07 | 28.94 | 133734 | 28 | 25615172 | 26871744 | 44065086 | 41590838 | 36088326 | 30504582 | 30847212 | 39377182 | 36388778 | 41076322 | 12825995 | 13107742 | 19986176 | 21584276 | 24333706 | 25931756 | 25789120 | 43788518 | 42409470 | 44149798 | 0 | 10 | 0 | 10 | 0 | 10 | 0.53 | 0.52 | 1.04 | 1.02 |
| Q9JHJ3 | Glmp | Glycosylated lysosomal membrane protein | 146.03 | 8.17 | 43804 | 2 | 2 | 230614.86 | 194429.7 | 156101.52 | 219160.42 | 2 | 122646.09 | 2 | 2 | 2 | 2707503.5 | 1630094.25 | 2475595.5 | 1920004.12 | 1679913.75 | 2 | 2 | 2 | 2 | 234816.42 | 0 | 7 | 0 | 12 | 0 | 12 | 13.01 | 84.90 | 0.29 | 1.91 |
| Q9CPV4 | Glod4 | Glyoxalase domain-containing protein 4 | 463.85 | 36.24 | 33317 | 10 | 19489580 | 18360552 | 19892076 | 20743120 | 18179010 | 16820710 | 17473012 | 16405674 | 16784588 | 19289066 | 10183757 | 12609820 | 10299401 | 10698300 | 11985039 | 21871454 | 18815360 | 19644470 | 19712764 | 21433614 | 0 | 7 | 0 | 7 | 0 | 7 | 0.58 | 0.64 | 1.05 | 1.17 |
| B0BNM9 | GLTP | Glycolipid transfer protein | 236.33 | 22.01 | 23704 | 5 | 1678254.62 | 1315361.25 | 2348158.75 | 2098829 | 2578301 | 1020392.38 | 1325934.75 | 2097967 | 2620231.5 | 1752848.75 | 2 | 561012.25 | 733115 | 289417.22 | 765984.38 | 2016055.75 | 1166587.5 | 1465446.75 | 1547395.5 | 1663426.88 | 0 | 5 | 0 | 10 | 0 | 10 | 0.23 | 0.27 | 0.78 | 0.89 |
| P08752 | Gnai2 | Guanine nucleotide-binding protein G(i) subunit alpha-2 | 1671.55 | 65.35 | 40489 | 22 | 50103158 | 47340502 | 47325442 | 52269762 | 49910698 | 47641506 | 49348982 | 45360682 | 46421414 | 43080854 | 31885220 | 35984118 | 32009802 | 34230586 | 34773162 | 44648494 | 52370350 | 46960338 | 50966458 | 43271574 | 0 | 9 | 0 | 10 | 0 | 10 | 0.68 | 0.73 | 0.96 | 1.03 |
| P63213 | Gng2 | Guanine nucleotide-binding protein G(I)/G(S)/G(O) subunit gamma-2 | 108.69 | 26.76 | 7850 | 2 | 4900839 | 3963083.5 | 5035450 | 6184619 | 4561058 | 4827567 | 4254266 | 3851059.75 | 4289304 | 5563404 | 2989544 | 2787218.75 | 3226758.5 | 3337359 | 3400853 | 3883504 | 5128770 | 5697879.5 | 6212776 | 4825493 | 0 | 10 | 0 | 7 | 0 | 10 | 0.64 | 0.69 | 1.04 | 1.13 |
| Q8BFR4 | Gns | N-acetylglucosamine-6-sulfatase | 768.17 | 25 | 61175 | 14 | 34371678 | 26735750 | 25553570 | 29391822 | 28688966 | 22658670 | 22628404 | 19831766 | 25202052 | 23195590 | 2423842.25 | 4221129 | 5897701 | 4668385.5 | 4903818 | 24064126 | 21555644 | 18135118 | 19287190 | 19508728 | 0 | 5 | 0 | 5 | 0 | 5 | 0.15 | 0.19 | 0.71 | 0.90 |
| Q9CR60 | Golt1b | Vesicle transport protein GOT1B | 205.95 | 17.39 | 15422 | 3 | 837335.12 | 947490.5 | 612597.81 | 914738.88 | 779631.69 | 1269546.5 | 1572276.5 | 1458348.62 | 2004970.5 | 1876188.12 | 3400907 | 3160662 | 4960405 | 4752864.5 | 3736450.75 | 1451852.12 | 1523484.75 | 2039682.25 | 2843197.5 | 1856451 | 0 | 9 | 0 | 9 | 0 | 9 | 4.89 | 2.45 | 2.37 | 1.19 |
| P05201 | Got1 | Aspartate aminotransferase, cytoplasmic | 1401.39 | 65.86 | 46248 | 23 | 113522818 | 117045410 | 126015354 | 141350386 | 134842066 | 109237498 | 105238154 | 119877036 | 116997346 | 125425698 | 70286122 | 76833890 | 76493218 | 81340842 | 81645114 | 131531642 | 147978738 | 141992338 | 148730754 | 151981010 | 0 | 2 | 0 | 0 | 0 | 2 | 0.61 | 0.67 | 1.14 | 1.25 |
| Q9WTK3 | Gpaa1 | Glycosylphosphatidylinositol anchor attachment 1 protein | 91.37 | 3.06 | 67949 | 2 | 998150.25 | 801489.44 | 1806090.62 | 842337.19 | 1293991 | 1784338.25 | 761747.94 | 2178971.75 | 2048846.38 | 2668100.75 | 3157840.75 | 3569428.5 | 4631606 | 3921310 | 4640166 | 1079046.25 | 1462610.38 | 2607028 | 2639096.5 | 2310962 | 0 | 3 | 0 | 11 | 0 | 11 | 3.47 | 2.11 | 1.76 | 1.07 |
| Q8CIM5 | Gpr84 | G-protein coupled receptor 84 | 576.73 | 26.77 | 43717 | 9 | 4737317.5 | 5633174.5 | 4121213 | 5519103 | 4899098 | 6839831 | 4338420.5 | 5987096.5 | 5189006 | 5859604.5 | 21072842 | 18636366 | 21370696 | 20801552 | 22792746 | 4651388 | 6195971 | 5983155.5 | 6814149.5 | 5941281 | 0 | 4 | 0 | 10 | 0 | 10 | 4.20 | 3.71 | 1.19 | 1.05 |
| Q8BHL4 | Gprc5a | Retinoic acid-induced protein 3 | 67.55 | 3.93 | 40101 | 1 | 2 | 2 | 2 | 2 | 2 | 2 | 2 | 2 | 2 | 2 | 915188 | 664877.88 | 520275.44 | 369602.59 | 595242.12 | 2 | 2 | 2 | 2 | 2 | 0 | 12.5 | 0 | 12.5 | 0 | 12.5 | 306518.60 | 306518.60 | 1.00 | 1.00 |
| P23764 | Gpx3 | Glutathione peroxidase 3 | 40.1 | 7.52 | 25424 | 1 | 2 | 2 | 2 | 2 | 2 | 2 | 2 | 2 | 2 | 2 | 1075901 | 934652.62 | 769936.75 | 342222.84 | 682758.56 | 2 | 2 | 2 | 2 | 2 | 0 | 12.5 | 0 | 12.5 | 0 | 12.5 | 380547.18 | 380547.18 | 1.00 | 1.00 |
| O35127 | Grcc10 | Protein C10 | 212.83 | 53.17 | 13194 | 4 | 2830727 | 2124692.25 | 2840743 | 2351709.75 | 3036198 | 2253090 | 1385207.12 | 2971057.5 | 2330686.5 | 2094438.62 | 300331.84 | 871908.38 | 1049285.5 | 216741.56 | 738508.38 | 2947734 | 1376036 | 3159710.25 | 2796134.5 | 3133875 | 0 | 9 | 0 | 7 | 0 | 9 | 0.24 | 0.29 | 1.02 | 1.22 |
| P97576 | Grpel1 | GrpE protein homolog 1, mitochondrial | 295.2 | 29.95 | 24297 | 6 | 1735018.38 | 2640217.5 | 1412237 | 466946.25 | 3196520 | 819662 | 2921576.75 | 2171001.5 | 3650719 | 1971295.88 | 6761640 | 7588640 | 6456072.5 | 6640221 | 7442133 | 1896635.38 | 331680.31 | 3010323 | 810692.25 | 1004236.94 | 0 | 9 | 0 | 6 | 0 | 9 | 3.69 | 3.02 | 0.75 | 0.61 |
| P47791 | Gsr | Glutathione reductase, mitochondrial | 566.92 | 17.4 | 53663 | 7 | 3867530 | 5218031.5 | 4319667 | 4161616.75 | 4477000.5 | 6024772.5 | 5336604 | 3577719.25 | 4582224 | 4882817.5 | 6568453 | 7794508.5 | 7674800 | 8705547 | 7990556 | 3867552.5 | 3493894.5 | 6115025 | 5559929.5 | 6028315 | 0 | 9 | 0 | 10 | 0 | 10 | 1.76 | 1.59 | 1.14 | 1.03 |
| P48774 | Gstm5 | Glutathione S-transferase Mu 5 | 385.52 | 28.57 | 26635 | 6 | 6732169.5 | 5669743 | 7112906 | 6233777 | 9932112 | 4723030 | 8137636 | 4022443 | 10034688 | 5599247.5 | 1826832.38 | 3154184 | 3607431.75 | 1216378 | 2951521.5 | 6732161.5 | 3495233 | 5409493 | 5282760 | 5224766 | 0 | 2 | 0 | 9 | 0 | 9 | 0.36 | 0.39 | 0.73 | 0.80 |
| P12265 | Gusb | Beta-glucuronidase | 1200.82 | 39.2 | 74195 | 22 | 85045962 | 82584178 | 146249298 | 143765138 | 155878242 | 70060706 | 76722298 | 78322954 | 90865146 | 78130858 | 23007158 | 26999788 | 19859798 | 23542568 | 23673456 | 83071410 | 70680890 | 68439378 | 67650346 | 77332322 | 0 | 1 | 0 | 7 | 0 | 7 | 0.19 | 0.30 | 0.60 | 0.93 |
| P01900 | H2-D1 | H-2 class I histocompatibility antigen, D-D alpha chain | 933.12 | 52.05 | 41110 | 17 | 1487373.12 | 3204119.75 | 3367785.5 | 3337204 | 2618029.5 | 1769629.38 | 2554930.5 | 4189572.25 | 3576301.5 | 3958085.5 | 9019624 | 5585289 | 6368448 | 4615714 | 5902007.5 | 2772704.5 | 2786644.5 | 2555594.5 | 3767217.75 | 3963066.5 | 0 | 10 | 0 | 12 | 0 | 12 | 2.25 | 1.96 | 1.13 | 0.99 |
| P01902 | H2-K1 | H-2 class I histocompatibility antigen, K-D alpha chain | 1052.88 | 46.2 | 41490 | 19 | 28126020 | 27930064 | 29021526 | 30811482 | 33994242 | 26288954 | 35469082 | 37679242 | 37000442 | 32977470 | 26112974 | 25991176 | 24484522 | 21060914 | 23928656 | 32555414 | 23674954 | 10862026 | 22173152 | 22935142 | 0 | 4 | 0 | 1 | 0 | 4 | 0.81 | 0.72 | 0.75 | 0.66 |
| P02301 | H3f3c | Histone H3.3C | 441.36 | 33.09 | 15315 | 8 | 1884889218 | 1358088962 | 1919138306 | 2201927682 | 1533722754 | 1958693250 | 1519249538 | 1643702914 | 1524756738 | 2015931138 | 617554690 | 716589954 | 907555074 | 1032108418 | 938035202 | 1477674242 | 2025817730 | 1718867842 | 1748238850 | 1727472386 | 0 | 11 | 0 | 11 | 0 | 11 | 0.47 | 0.49 | 0.98 | 1.00 |
| Q8K2C9 | Hacd3 | Very-long-chain (3R)-3-hydroxyacyl-CoA dehydratase 3 | 82.18 | 4.97 | 43131 | 2 | 1510929 | 175560.22 | 232013.08 | 1562414.62 | 372875.81 | 2 | 417287.03 | 357978.34 | 548972.88 | 887509.81 | 2875743.75 | 2242437.75 | 2737799.5 | 3237478.25 | 2953928.5 | 364393.75 | 1573279.12 | 547192.25 | 493942.44 | 495456.19 | 0 | 9 | 0 | 9 | 0 | 9 | 3.65 | 6.35 | 0.90 | 1.57 |
| Q8BMS1 | Hadha | Trifunctional enzyme subunit alpha, mitochondrial | 1276.75 | 28.57 | 82670 | 19 | 18769038 | 13916048 | 19519222 | 19101232 | 18721064 | 17068610 | 16815420 | 15782466 | 15145501 | 18298116 | 12225115 | 9658771 | 12278860 | 10257934 | 13894661 | 16628005 | 19154420 | 17473894 | 15402034 | 13025852 | 0 | 7 | 0 | 12 | 0 | 12 | 0.65 | 0.70 | 0.91 | 0.98 |
| P29416 | Hexa | Beta-hexosaminidase subunit alpha | 650.35 | 31.82 | 60613 | 12 | 18390730 | 17412714 | 40341382 | 27808360 | 41761094 | 14617442 | 19282376 | 26860122 | 27241192 | 24376154 | 6862828.5 | 11301007 | 12451912 | 10431678 | 9951231 | 18990322 | 10419905 | 20485608 | 21849054 | 20666060 | 0 | 8 | 0 | 7 | 0 | 8 | 0.35 | 0.45 | 0.63 | 0.82 |
| P20060 | Hexb | Beta-hexosaminidase subunit beta | 798.43 | 33.58 | 61116 | 17 | 31394074 | 31968944 | 61447410 | 45582190 | 62474394 | 24922284 | 33777674 | 37064490 | 46123190 | 32141602 | 10747997 | 12170861 | 15381739 | 11126127 | 14927591 | 25674974 | 21994412 | 30479198 | 32334876 | 28761408 | 0 | 2 | 0 | 5 | 0 | 5 | 0.28 | 0.37 | 0.60 | 0.80 |
| Q9JLZ6 | Hic2 | Hypermethylated in cancer 2 protein | 47.63 | 1.62 | 66766 | 1 | 2 | 2 | 2 | 2 | 2 | 2 | 2 | 2 | 2 | 2 | 38468358 | 10930912 | 4758361 | 3845766.25 | 5379237 | 2 | 2 | 2 | 2 | 2 | 0 | 12.5 | 0 | 12.5 | 0 | 12.5 | 6338263.43 | 6338263.43 | 1.00 | 1.00 |

| | | | | | | | | | | | | | | | | | | | | | | | | | | | | | | | | | | | | |
|---|---|---|---|---|---|---|---|---|---|---|---|---|---|---|---|---|---|---|---|---|---|---|---|---|---|---|---|---|---|---|---|---|---|---|---|---|
| P62804 | Hist1h4b | Histone H4 | 1276.1 | 65.05 | 11367 | 17 | 1869148290 | 1393522562 | 2368216322 | 2004370946 | 2118236418 | 1667366146 | 1820156674 | 2001113986 | 2066976514 | 1943279874 | 652726914 | 772401666 | 772940994 | 744262914 | 781346306 | 1502400002 | 1477421954 | 1507468034 | 1534740098 | 1533133954 | 0 | 5 | 0 | 0 | 0 | 5 | 0.38 | 0.39 | 0.77 | 0.80 |
| P17710 | Hk1 | Hexokinase-1 | 1778.07 | 30.08 | 108303 | 28 | 47207126 | 49684014 | 50801102 | 54767726 | 48574598 | 49842558 | 43376034 | 46496154 | 46610750 | 45964134 | 23278056 | 21883030 | 31176500 | 32114470 | 31288544 | 44647010 | 40027282 | 48033654 | 46117506 | 44767790 | 0 | 1 | 0 | 8 | 0 | 8 | 0.56 | 0.60 | 0.89 | 0.96 |
| Q9D8V0 | Hm13 | Minor histocompatibility antigen H13 | 285.19 | 10.05 | 41748 | 5 | 5183118.5 | 7912295 | 8585135 | 5042682 | 7404219 | 9168869 | 11075718 | 9742800 | 11309677 | 10520672 | 17253918 | 19022720 | 19568800 | 13901836 | 19108150 | 7859244 | 7124779.5 | 10448673 | 9782531 | 10970510 | 0 | 5 | 0 | 7 | 0 | 7 | 2.60 | 1.71 | 1.35 | 0.89 |
| P63158 | Hmgb1 | High mobility group protein B1 | 497.76 | 38.6 | 24894 | 8 | 23205570 | 28924132 | 36192366 | 42151882 | 30326520 | 26836270 | 24562834 | 37268022 | 28494522 | 32143542 | 18169290 | 18034210 | 19538952 | 21277812 | 21095964 | 32348298 | 26859146 | 36595802 | 33785598 | 41563262 | 0 | 10 | 0 | 6 | 0 | 10 | 0.61 | 0.66 | 1.06 | 1.15 |
| P30681 | Hmgb2 | High mobility group protein B2 | 603.57 | 33.33 | 24162 | 8 | 21961562 | 23457366 | 41478190 | 43007074 | 29077222 | 28563130 | 23595634 | 28329708 | 24294538 | 28383470 | 13630131 | 14304459 | 14980980 | 19119014 | 16910132 | 25979574 | 26381422 | 25905746 | 25673096 | 30006830 | 0 | 11 | 0 | 12 | 0 | 12 | 0.50 | 0.59 | 0.84 | 1.01 |
| P14901 | Hmox1 | Heme oxygenase 1 | 559.13 | 51.21 | 32929 | 10 | 523223.28 | 333599.88 | 281882.22 | 486132.62 | 260096.28 | 396796.94 | 492680.75 | 417891.44 | 255481.94 | 966386.88 | 4228262 | 4485262.5 | 9708963 | 9171823 | 7049210 | 1636047.5 | 2272107 | 1785306.5 | 2238398.5 | 1883088.38 | 0 | 0 | 0 | 0 | 0 | 0 | 18.38 | 13.70 | 5.21 | 3.88 |
| Q794E4 | Hnrnpf | Heterogeneous nuclear ribonucleoprotein F | 975.83 | 47.71 | 45730 | 14 | 11060157 | 15802776 | 18965024 | 15372899 | 15677346 | 13567862 | 11582927 | 15081910 | 15051018 | 15933645 | 39117578 | 30143516 | 30999506 | 32429128 | 32591700 | 13179307 | 13028478 | 20546906 | 16936200 | 21042528 | 0 | 9 | 0 | 8 | 0 | 9 | 2.15 | 2.32 | 1.10 | 1.19 |
| Q9D0E1 | Hnrnpm | Heterogeneous nuclear ribonucleoprotein M | 2480 | 49.66 | 77649 | 36 | 167336434 | 193341746 | 178978978 | 195731986 | 160736706 | 191903234 | 179516322 | 164910978 | 151263746 | 161281266 | 130172210 | 132983026 | 114726362 | 129031898 | 126715898 | 178981762 | 193186466 | 170282930 | 188495170 | 182766178 | 0 | 11 | 0 | 6 | 0 | 11 | 0.71 | 0.75 | 1.02 | 1.08 |
| Q6IMY8 | Hnrnpu | Heterogeneous nuclear ribonucleoprotein U | 2350.89 | 45.36 | 87732 | 35 | 101909426 | 110872058 | 132315826 | 134402754 | 123022706 | 89282778 | 114492530 | 125439818 | 116803170 | 120934722 | 187348850 | 169775058 | 164332562 | 161619522 | 171480514 | 106669650 | 107430194 | 131694650 | 130245258 | 136360066 | 0 | 12 | 0 | 8 | 0 | 12 | 1.42 | 1.51 | 1.02 | 1.08 |
| Q3TEA8 | Hp1bp3 | Heterochromatin protein 1-binding protein 3 | 753.51 | 27.44 | 60867 | 14 | 10764145 | 10801052 | 9942685 | 11520960 | 9277487 | 13437523 | 9359696 | 8447070 | 9447561 | 9877402 | 5420073.5 | 4704245.5 | 7196396 | 7484029 | 6610166 | 9351240 | 10871964 | 11433897 | 10946260 | 10033310 | 0 | 10 | 0 | 8 | 0 | 10 | 0.60 | 0.62 | 1.01 | 1.04 |
| P00493 | Hprt1 | Hypoxanthine-guanine phosphoribosyltransferase | 500.08 | 26.15 | 24570 | 7 | 33790090 | 30398842 | 36526894 | 30844400 | 32075374 | 30106702 | 30306606 | 30455542 | 31334932 | 33212414 | 18558818 | 25044932 | 21425262 | 16412856 | 20107148 | 32734766 | 26674158 | 29056866 | 29366844 | 30421430 | 0 | 4 | 0 | 6 | 0 | 6 | 0.62 | 0.65 | 0.91 | 0.95 |
| O70503 | Hsd17b12 | Very-long-chain 3-oxoacyl-CoA reductase | 1211.37 | 67.95 | 34742 | 19 | 24645048 | 24625552 | 20024756 | 20379068 | 24375654 | 26629998 | 28338590 | 23588066 | 27183254 | 28153398 | 73031874 | 56624130 | 65838058 | 55050814 | 64825014 | 30608466 | 37458042 | 27887806 | 32889588 | 26303100 | 0 | 0 | 0 | 6 | 0 | 6 | 2.77 | 2.36 | 1.36 | 1.16 |
| P08113 | Hsp90b1 | Endoplasmin | 3416.67 | 59.98 | 92476 | 55 | 197288546 | 210405858 | 231678802 | 244539202 | 222799890 | 195361922 | 178001922 | 246966114 | 241121570 | 246667842 | 317766562 | 274095010 | 400029954 | 341421730 | 412321122 | 169938674 | 216094610 | 236078146 | 241949154 | 229261138 | 0 | 12 | 0 | 9 | 0 | 12 | 1.58 | 1.58 | 0.99 | 0.99 |
| P63017 | Hspa8 | Heat shock cognate 71 kDa protein | 3844.221 | 75.08 | 70871 | 54 | 1227405442 | 1234723586 | 1213958018 | 1219160962 | 1150131586 | 1202867202 | 1159717634 | 1061806274 | 1097164290 | 1142324866 | 935175170 | 1060865986 | 878373186 | 769258562 | 839118978 | 1235759234 | 1270444674 | 1119273986 | 1061670786 | 1116368002 | 0 | 10 | 0 | 11 | 0 | 11 | 0.74 | 0.79 | 0.96 | 1.02 |
| P38647 | Hspa9 | Stress-70 protein, mitochondrial | 2894.79 | 69.51 | 73461 | 41 | 121863890 | 138234162 | 96252466 | 109482770 | 92253346 | 130259778 | 126977170 | 106917298 | 88876530 | 110469658 | 231566034 | 220905202 | 220763714 | 230550002 | 226099922 | 141427714 | 170057570 | 140447042 | 137563682 | 144059698 | 0 | 1 | 0 | 0 | 0 | 1 | 2.02 | 2.01 | 1.31 | 1.30 |
| P63038 | Hspd1 | 60 kDa heat shock protein, mitochondrial | 4175.81 | 76.44 | 60955 | 54 | 405753922 | 420495266 | 382470690 | 430740898 | 326575970 | 425424674 | 362507586 | 404076610 | 339642530 | 453447074 | 916978498 | 866707522 | 918442114 | 920899522 | 978072386 | 431340546 | 524064386 | 484915778 | 470334530 | 492476642 | 0 | 0 | 0 | 1 | 0 | 1 | 2.34 | 2.32 | 1.22 | 1.21 |
| Q9JKR6 | Hyou1 | Hypoxia up-regulated protein 1 | 3040.19 | 52.85 | 111181 | 48 | 47324006 | 48168646 | 35421122 | 33866322 | 40613994 | 41902814 | 42494498 | 51366006 | 50924766 | 59127586 | 115752090 | 114542186 | 119983154 | 105200922 | 134944610 | 32314322 | 41307738 | 53938778 | 52005582 | 53097274 | 0 | 7 | 0 | 12 | 0 | 12 | 2.87 | 2.40 | 1.13 | 0.95 |
| P54071 | Idh2 | Isocitrate dehydrogenase [NADP], mitochondrial | 1051.17 | 40.49 | 50906 | 18 | 38082038 | 40262122 | 43873622 | 40174962 | 47431150 | 31461612 | 32558562 | 34173254 | 35502062 | 31663588 | 17785820 | 23827426 | 25200788 | 22935422 | 24628980 | 42051914 | 34490554 | 30764826 | 31560086 | 32867800 | 0 | 3 | 0 | 12 | 0 | 12 | 0.55 | 0.69 | 0.82 | 1.04 |
| Q9D6R2 | Idh3a | Isocitrate dehydrogenase [NAD] subunit alpha, mitochondrial | 722.19 | 41.53 | 39639 | 13 | 24547176 | 28547432 | 29279290 | 28223896 | 31241236 | 25028598 | 24697762 | 29315538 | 30073450 | 28402092 | 45602310 | 39415898 | 43027398 | 39866254 | 48514402 | 25204506 | 29061532 | 34023446 | 33614218 | 32081326 | 0 | 6 | 0 | 5 | 0 | 6 | 1.53 | 1.57 | 1.09 | 1.12 |
| P41565 | Idh3g | Isocitrate dehydrogenase [NAD] subunit gamma 1, mitochondrial | 308.85 | 16.54 | 42851 | 7 | 6981381 | 7797810 | 7493047 | 6276695.5 | 6115718.5 | 6791439 | 5479540 | 5468802.5 | 5329397 | 6460638 | 8854416 | 9117726 | 9936011 | 10380129 | 9971880 | 5652533 | 7237330 | 7224916.5 | 5883807 | 7413595 | 0 | 9 | 0 | 4 | 0 | 9 | 1.39 | 1.63 | 0.96 | 1.13 |
| Q9ESY9 | Ifi30 | Gamma-interferon-inducible lysosomal thiol reductase | 322.3 | 28.23 | 27784 | 6 | 4818800 | 4628118 | 3605326.5 | 3213951 | 8805051 | 3610780.5 | 4337029.5 | 4440209.5 | 5575459.5 | 2720234.5 | 739986.69 | 933777.38 | 1160654.5 | 621738.62 | 257705.12 | 2571911.5 | 3272537.75 | 2392246.5 | 1288238.38 | 1255300.62 | 0 | 1 | 0 | 1 | 0 | 1 | 0.15 | 0.18 | 0.43 | 0.52 |
| P19182 | Ifrd1 | Interferon-related developmental regulator 1 | 138.58 | 5.35 | 49935 | 2 | 279081.94 | 436668.25 | 2 | 2 | 2 | 187954.36 | 151626.27 | 2 | 2 | 2 | 1547453.25 | 1664108.88 | 1046001.31 | 739880.38 | 1190069.88 | 478172.88 | 491758.84 | 219854.34 | 2 | 167304.45 | 0 | 7.5 | 0 | 4.5 | 0 | 7.5 | 8.64 | 18.22 | 1.90 | 4.00 |
| Q66HG8 | Ik | Protein Red | 95.98 | 5.57 | 65588 | 2 | 2125162.75 | 2231188.5 | 2464802.25 | 1728169.88 | 2650804.5 | 1292077.75 | 2794753.5 | 1277337.38 | 2181089.5 | 1979706 | 401953.78 | 1063687.5 | 896554.88 | 2 | 542224.12 | 1898005.5 | 764302.25 | 844316.06 | 491318.97 | 985567.5 | 0 | 1 | 0 | 2 | 0 | 2 | 0.26 | 0.30 | 0.44 | 0.52 |
| P16872 | Il7r | Interleukin-7 receptor subunit alpha | 97.07 | 5.88 | 51605 | 2 | 2 | 2 | 2 | 863705.56 | 2 | 2 | 770374.94 | 2 | 2 | 284662.88 | 2626473 | 986062.5 | 2314582.75 | 2159247.75 | 2444250 | 900625.25 | 1168003.25 | 723302.62 | 2 | 2 | 0 | 7 | 0 | 8 | 0 | 8 | 12.19 | 9.98 | 3.23 | 2.65 |
| Q5PQR5 | Imp4 | U3 small nucleolar ribonucleoprotein protein IMP4 | 68.3 | 7.56 | 33656 | 2 | 900686.06 | 1422592.5 | 1242228.25 | 1562573.25 | 907246.88 | 1497650.88 | 1030756.31 | 772394.69 | 806886.19 | 1274013.62 | 2106856 | 1760823.75 | 1817512.88 | 2215013.5 | 1872117.88 | 1085036.75 | 1774708.75 | 1425595.88 | 1799022.25 | 1590846.5 | 0 | 4 | 0 | 3 | 0 | 4 | 1.62 | 1.82 | 1.27 | 1.43 |
| P24547 | Impdh2 | Inosine-5'-monophosphate dehydrogenase 2 | 1096.96 | 34.82 | 55815 | 17 | 46818662 | 44192842 | 57220642 | 55969838 | 47685234 | 44620610 | 41569466 | 42685438 | 43475630 | 43758310 | 36532262 | 37770430 | 33612442 | 33456942 | 35215622 | 46446602 | 55559318 | 51905486 | 46766378 | 45682334 | 0 | 9 | 0 | 0 | 0 | 9 | 0.70 | 0.82 | 0.98 | 1.14 |
| Q9ES52 | Inpp5d | Phosphatidylinositol 3,4,5-trisphosphate 5-phosphatase 1 | 1415.42 | 28.13 | 133542 | 27 | 38693458 | 37958290 | 31860174 | 36696110 | 31288398 | 38899194 | 32195852 | 30773270 | 33202056 | 34134746 | 25947798 | 25220196 | 20213274 | 21812996 | 22761554 | 36890786 | 39475466 | 35998058 | 34038246 | 36317478 | 0 | 11 | 0 | 5 | 0 | 11 | 0.66 | 0.69 | 1.04 | 1.08 |
| Q6P549 | Inppl1 | Phosphatidylinositol 3,4,5-trisphosphate 5-phosphatase 2 | 526.24 | 12.41 | 138973 | 12 | 4396439 | 3927493 | 4761229 | 5658810 | 8078044.5 | 2495282 | 4052363 | 4094670 | 5376734.5 | 3798890.75 | 1310382.38 | 1031670.69 | 1286032.75 | 1388325.25 | 1473479.25 | 4019580.5 | 2992418.75 | 2099189 | 1016628.75 | 2970157.5 | 0 | 1 | 0 | 4 | 0 | 4 | 0.24 | 0.33 | 0.49 | 0.66 |
| Q91YE6 | Ipo9 | Importin-9 | 960.54 | 20.46 | 116052 | 17 | 19476406 | 19567476 | 24585030 | 21887518 | 26464994 | 19757014 | 19773444 | 23340508 | 23017398 | 17601586 | 12420822 | 13855881 | 13840517 | 12214764 | 13538037 | 19448054 | 17855322 | 14532344 | 18771570 | 14701470 | 0 | 0 | 0 | 3 | 0 | 3 | 0.59 | 0.64 | 0.76 | 0.82 |
| Q9JL16 | Isg20 | Interferon-stimulated gene 20 kDa protein | 43.75 | 2.67 | 32503 | 1 | 289973.44 | 261718.09 | 297002.19 | 301518.16 | 240103.88 | 290711.06 | 262436.28 | 229108.72 | 386360.19 | 319806.16 | 184280.02 | 111135 | 90112.27 | 2 | 128588.41 | 169233.66 | 2 | 182507.41 | 147685.86 | 167307.08 | 0 | 0 | 0 | 0 | 0 | 0 | 0.37 | 0.35 | 0.48 | 0.45 |
| P11688 | Itga5 | Integrin alpha-5 | 1108.36 | 24.79 | 115043 | 20 | 11842461 | 11361794 | 18609662 | 15657124 | 13577981 | 15707700 | 12934647 | 19186016 | 17769616 | 23509374 | 35551322 | 29716364 | 37055202 | 33330228 | 29294412 | 11561038 | 14106472 | 24209886 | 18605284 | 22018326 | 0 | 7 | 0 | 12 | 0 | 12 | 2.32 | 1.85 | 1.27 | 1.02 |
| P24063 | Itgal | Integrin alpha-L | 76.97 | 1.89 | 128328 | 2 | 856702.25 | 977231.31 | 783113.81 | 1136945.62 | 733153.62 | 1021454.69 | 761024.62 | 744346.38 | 2 | 820970.44 | 1278579.12 | 1364140 | 1183441.88 | 1285109.75 | 1264563.75 | 712654.81 | 1073370.62 | 1000419.81 | 795815.19 | 965764 | 0 | 12 | 0 | 8 | 0 | 12 | 1.42 | 1.90 | 1.01 | 1.36 |
| Q91VK4 | Itm2c | Integral membrane protein 2C | 278.93 | 27.51 | 30482 | 5 | 750624.88 | 564020.69 | 328609.22 | 335555.47 | 740555 | 747145.12 | 695626 | 2 | 836358.62 | 263746.09 | 4748231.5 | 4340974 | 4261480.5 | 3937329.75 | 3955711 | 279467.75 | 407745.34 | 715998.19 | 2 | 764819.5 | 0 | 10 | 0 | 11.5 | 0 | 11.5 | 7.81 | 8.35 | 0.80 | 0.85 |
| Q9D892 | Itpa | Inosine triphosphate pyrophosphatase | 172.31 | 18.69 | 21897 | 3 | 6838660 | 5301901 | 7308243 | 9128029 | 9305616 | 5304724 | 6490282 | 7884006.5 | 8690226 | 7706751.5 | 2299797 | 3972023.25 | 4754557.5 | 3661656.5 | 3792391.75 | 7939065 | 6728717 | 7481885 | 8567852 | 7958636 | 0 | 12 | 0 | 9 | 0 | 12 | 0.49 | 0.51 | 1.02 | 1.07 |
| P11881 | Itpr1 | Inositol 1,4,5-trisphosphate receptor type 1 | 93.61 | 0.73 | 313167 | 2 | 330378.31 | 213152.36 | 226356.89 | 252296.06 | 296544.53 | 294254.16 | 318547 | 277317.75 | 225733 | 233852.44 | 155560.88 | 198080.47 | 2 | 206433.48 | 200314.98 | 2 | 2 | 110051.66 | 256834.36 | 167474.98 | 0 | 3 | 0 | 2 | 0 | 3 | 0.58 | 0.56 | 0.41 | 0.40 |
| Q3U0V1 | Khsrp | Far upstream element-binding protein 2 | 1218.73 | 34.09 | 76775 | 20 | 49038038 | 50065806 | 49674678 | 58683110 | 52704554 | 53206626 | 49638854 | 46294770 | 45225606 | 49750806 | 36379470 | 34166870 | 31685970 | 36121614 | 30526322 | 57075542 | 71344698 | 54589390 | 57041538 | 50320250 | 0 | 5 | 0 | 1 | 0 | 5 | 0.65 | 0.69 | 1.12 | 1.19 |
| Q9EP89 | Lactb | Serine beta-lactamase-like protein LACTB, mitochondrial | 189.55 | 5.99 | 60705 | 3 | 1807807.38 | 1984280.25 | 1597271.25 | 1171047.75 | 2270608 | 1530411.38 | 1861333.88 | 1994968.75 | 1863822.88 | 1675037.75 | 395289.97 | 213890.73 | 2 | 143068.91 | 861726.12 | 1599788.88 | 2 | 921395.12 | 2 | 128094.3 | 0 | 2 | 0 | 1 | 0 | 2 | 0.18 | 0.18 | 0.30 | 0.30 |
| Q3U9G9 | Lbr | Lamin-B receptor | 873.65 | 25.4 | 71440 | 14 | 5259683.5 | 4856474.5 | 7822537 | 8392977 | 7031607.5 | 6342602.5 | 6187232 | 7406558 | 7967984.5 | 9603967 | 16012562 | 13508586 | 20155752 | 19196358 | 21768552 | 5297935 | 6453005 | 9279695 | 9859510 | 7567412 | 0 | 8 | 0 | 11 | 0 | 11 | 2.72 | 2.42 | 1.15 | 1.03 |
| Q60787 | Lcp2 | Lymphocyte cytosolic protein 2 | 265.97 | 9.19 | 60238 | 4 | 2112590.75 | 2041280.12 | 840686.12 | 245124.47 | 1353339.5 | 1635441.75 | 1200299.5 | 959365.81 | 1247842.25 | 533298.19 | 3326787.25 | 4513237 | 2486341 | 2264823 | 2817614.75 | 846759.25 | 1865509.38 | 213854.94 | 1110590.88 | 572904.75 | 0 | 8 | 0 | 9 | 0 | 9 | 2.34 | 2.76 | 0.70 | 0.83 |
| P35951 | Ldlr | Low-density lipoprotein receptor | 316.43 | 7.19 | 94947 | 5 | 661465.25 | 1033295.38 | 936469.56 | 1295552 | 781878.75 | 1051792.62 | 893630.88 | 2 | 2 | 256080.97 | 1722112.5 | 4132536.75 | 3733809.75 | 1971964.5 | 4487035 | 412995.34 | 455623.84 | 377740.19 | 1707570.5 | 599742.38 | 0 | 5 | 0 | 8 | 0 | 8 | 3.41 | 7.29 | 0.75 | 1.61 |
| P16045 | Lgals1 | Galectin-1 | 781.69 | 79.26 | 14866 | 10 | 281825570 | 248817506 | 225968786 | 257049282 | 282939010 | 265381122 | 275253538 | 220794546 | 237212802 | 261775538 | 99424202 | 121197754 | 107703250 | 109459338 | 107044658 | 290200930 | 287853922 | 230620130 | 260718962 | 244264818 | 0 | 10 | 0 | 10 | 0 | 10 | 0.42 | 0.43 | 1.01 | 1.04 |
| Q64281 | Lilrb4 | Leukocyte immunoglobulin-like receptor subfamily B member 4 | 500.66 | 28.36 | 37544 | 6 | 3365518.5 | 1745999.75 | 2115142 | 2420224 | 3628811 | 4115080 | 4221960.5 | 4682811 | 4617899 | 4979627 | 8733989 | 7878697 | 8406379 | 7809151 | 7303720.5 | 3501911.75 | 3558118.5 | 4233616 | 3625183.5 | 3347723.5 | 0 | 5 | 0 | 2 | 0 | 5 | 3.02 | 1.77 | 1.38 | 0.81 |
| Q9D0F3 | Lman1 | Protein ERGIC-53 | 549.78 | 26.69 | 57789 | 9 | 6423150.5 | 8275196 | 10211947 | 9198566 | 9584530 | 8135166.5 | 7710888.5 | 10351762 | 11277188 | 8645153 | 12454653 | 12536926 | 15139685 | 15051736 | 13324501 | 8285251 | 9182452 | 9929818 | 12054200 | 10191274 | 0 | 8 | 0 | 9 | 0 | 9 | 1.57 | 1.49 | 1.14 | 1.08 |
| P48678 | Lmna | Prelamin-A/C | 2967.92 | 58.5 | 74238 | 45 | 266299218 | 283091618 | 242070498 | 279841890 | 252677442 | 278470594 | 298436578 | 262843362 | 246851122 | 288641698 | 160051522 | 189044690 | 163468402 | 154405938 | 185538146 | 277829698 | 286973794 | 279760898 | 300310402 | 279428194 | 0 | 6 | 0 | 9 | 0 | 9 | 0.64 | 0.62 | 1.08 | 1.04 |
| Q8BYI6 | Lpcat2 | Lysophosphatidylcholine acyltransferase 2 | 359.16 | 22.43 | 60254 | 8 | 843402.56 | 1191794.75 | 1317191.88 | 1394441.75 | 1462377.62 | 1156857 | 1303445.25 | 881019 | 1013884.5 | 810930.75 | 279867.06 | 389445.62 | 191247.09 | 261039.7 | 325689.62 | 910859.44 | 987808.19 | 895796 | 1039600.38 | 1189402.38 | 0 | 5 | 0 | 12 | 0 | 12 | 0.23 | 0.28 | 0.81 | 0.97 |
| Q91V01 | Lpcat3 | Lysophospholipid acyltransferase 5 | 104.58 | 4.52 | 56147 | 2 | 625699.56 | 389162.66 | 886404.31 | 640202.94 | 901033.75 | 776814.12 | 1040807.25 | 1087535.25 | 1330276.75 | 1051003.12 | 1962820 | 3066924 | 4068997.25 | 2985422.5 | 4746060 | 725733.44 | 743691.69 | 1234934.88 | 1207846 | 1247278.12 | 0 | 4 | 0 | 12 | 0 | 12 | 4.89 | 3.18 | 1.50 | 0.98 |
| P11152 | Lpl | Lipoprotein lipase | 602.75 | 23.21 | 53109 | 10 | 1234617.5 | 358826.94 | 2388391.25 | 2271716.25 | 966526 | 788914.62 | 1749874.38 | 2935389.5 | 1297370.25 | 1868017.88 | 5054586 | 4731905 | 4413182.5 | 3101498.5 | 3686105.25 | 1149003 | 898067.5 | 1699250.75 | 3285494.25 | 1694468.88 | 0 | 11 | 0 | 11 | 0 | 11 | 2.91 | 2.43 | 1.21 | 1.01 |
| Q91ZX7 | Lrp1 | Prolow-density lipoprotein receptor-related protein 1 | 1651.3 | 7.59 | 504742 | 31 | 55042450 | 56414042 | 37370282 | 40267122 | 33406290 | 53472694 | 47020950 | 25449728 | 23394752 | 29010002 | 18868120 | 14801657 | 12911179 | 10554529 | 12921956 | 41103598 | 43104802 | 27239996 | 29539502 | 27315244 | 0 | 6 | 0 | 12 | 0 | 12 | 0.31 | 0.39 | 0.76 | 0.94 |
| Q7TQH7 | Lrp10 | Low-density lipoprotein receptor-related protein 10 | 88.79 | 2.66 | 76461 | 2 | 643770.94 | 767532.75 | 369381.12 | 454843.12 | 731465 | 1047694.75 | 613504.06 | 428188.19 | 513527.25 | 509320.94 | 1562225.5 | 1720724.62 | 1935361.75 | 1287719.38 | 1798435.38 | 1042345.31 | 682084.62 | 531487.31 | 2 | 673811.25 | 0 | 12 | 0 | 10 | 0 | 12 | 2.80 | 2.67 | 0.99 | 0.94 |
| Q3UZ39 | Lrrfip1 | Leucine-rich repeat flightless-interacting protein 1 | 819.01 | 25.51 | 79249 | 13 | 19315006 | 22981114 | 17006002 | 22322280 | 18633760 | 20521002 | 20178786 | 18780440 | 17121060 | 18106256 | 10886804 | 12055202 | 9525321 | 11502478 | 9276387 | 23092580 | 23360224 | 19555990 | 18568614 | 20385104 | 0 | 8 | 0 | 6 | 0 | 8 | 0.53 | 0.56 | 1.05 | 1.11 |
| Q8BUI3 | LRWD1 | Leucine-rich repeat and WD repeat-containing protein 1 | 49.64 | 1.7 | 71594 | 1 | 2 | 2 | 2 | 2 | 2 | 2 | 2 | 2 | 2 | 2 | 340239.28 | 337358.19 | 463514.47 | 340439.41 | 290818.41 | 2 | 2 | 2 | 2 | 2 | 0 | 12.5 | 0 | 12.5 | 0 | 12.5 | 177236.98 | 177236.98 | 1.00 | 1.00 |
| Q9D0R8 | Lsm12 | Protein LSM12 homolog | 89.82 | 6.67 | 21701 | 1 | 2 | 471239.38 | 328720.88 | 2 | 2 | 2 | 2 | 506729.59 | 2 | 2 | 1242698.25 | 1175439.75 | 1239589.5 | 1293901.25 | 1257084.38 | 2 | 2 | 2 | 419952.78 | 2 | 0 | 10 | 0 | 12 | 0 | 12 | 7.76 | 12.25 | 0.52 | 0.83 |
| P62311 | Lsm3 | U6 snRNA-associated Sm-like protein LSm3 | 71.5 | 19.61 | 11845 | 2 | 1271525.12 | 1211671.12 | 1460377.38 | 685369.44 | 1166270 | 1281448.62 | 878357.19 | 915957.44 | 1392405.5 | 1559572.5 | 4632845 | 2379421 | 2587180.75 | 1660958.12 | 2369353 | 803724.88 | 1493493.88 | 1100757.75 | 1271699.5 | 878678.62 | 0 | 12 | 0 | 9 | 0 | 12 | 2.35 | 2.26 | 0.96 | 0.92 |
| Q99KG5 | Lsr | Lipolysis-stimulated lipoprotein receptor | 43.56 | 1.85 | 66108 | 1 | 7163.25 | 100147.45 | 2 | 2 | 2 | 110459.59 | 54202.57 | 2 | 2 | 2 | 132897.11 | 145660.48 | 285682.34 | 187141.3 | 187936.56 | 2 | 2 | 2 | 2 | 2 | 0 | 7.5 | 0 | 7.5 | 0 | 7.5 | 8.75 | 5.70 | 0.00 | 0.00 |
| Q8BLN5 | Lss | Lanosterol synthase | 484.66 | 12.82 | 83141 | 8 | 1722283.62 | 1928754.12 | 1291033.25 | 1290513.75 | 1740296 | 1914380 | 2430977.25 | 1279173.75 | 1924085.5 | 1368363.75 | 2890036 | 3658391.5 | 3882200.75 | 2859282.5 | 3776549.5 | 1096789.38 | 1211127.75 | 1349989.38 | 2098516.25 | 1961507.75 | 0 | 12 | 0 | 9 | 0 | 12 | 2.14 | 1.91 | 0.97 | 0.87 |
| Q7TNC4 | Luc7l2 | Putative RNA-binding protein Luc7-like 2 | 578.43 | 25.77 | 46583 | 10 | 5125886.5 | 5753026.5 | 5992173.5 | 6881297 | 7304687 | 5237472.5 | 5134608 | 6575725 | 5708022 | 4164100.5 | 3343101.5 | 3488291.5 | 3801143 | 3651439.25 | 3840224 | 4154419.25 | 3974542.5 | 3526939 | 3960842.25 | 4335961 | 0 | 0 | 0 | 1 | 0 | 1 | 0.58 | 0.68 | 0.64 | 0.74 |
| Q6AY04 | Maip1 | m-AAA protease-interacting protein 1, mitochondrial | 127.28 | 8.59 | 33090 | 2 | 1334924 | 734294 | 469748.62 | 1273225.75 | 971925.62 | 1215032.38 | 1267970.5 | 538976.12 | 1202621.5 | 764693.44 | 1811880.5 | 1495442.75 | 2207069.25 | 1790084 | 1752787.5 | 1043336.88 | 1319363.5 | 1610985 | 1590970.25 | 1301967.5 | 0 | 4 | 0 | 3 | 0 | 4 | 1.89 | 1.82 | 1.44 | 1.38 |
| O09159 | Man2b1 | Lysosomal alpha-mannosidase | 675.96 | 14.91 | 114648 | 11 | 9953328 | 10759316 | 17569328 | 14181924 | 13296807 | 9543019 | 10855899 | 5991148.5 | 10765459 | 7942538.5 | 1263374.12 | 891009.44 | 1647786.75 | 2407018 | 1530656.75 | 9685209 | 7562207.5 | 6201755 | 9077272 | 7346064 | 0 | 0 | 0 | 8 | 0 | 8 | 0.12 | 0.17 | 0.61 | 0.88 |
| P0C5H9 | Manf | Mesencephalic astrocyte-derived neurotrophic factor | 373.89 | 33.52 | 20388 | 7 | 8962832 | 9601597 | 8523604 | 8483622 | 8697764 | 9809799 | 10501872 | 8796728 | 7521761 | 10415698 | 3883957 | 4898595.5 | 4732790.5 | 4397946.5 | 5379945.5 | 9384418 | 4852431 | 4609467 | 7919918 | 5716704 | 0 | 4 | 0 | 3 | 0 | 4 | 0.53 | 0.50 | 0.73 | 0.69 |
| Q9ESL4 | Map3k20 | Mitogen-activated protein kinase kinase kinase 20 | 223.11 | 5.11 | 91720 | 4 | 1023696.75 | 933503.19 | 2086999 | 1406998.5 | 2110515.5 | 677438.69 | 1208474.75 | 1903018.25 | 2321404.5 | 982177.88 | 463697.25 | 332279.19 | 615450.69 | 559357.56 | 644191.19 | 1119904.38 | 405452.62 | 893746.38 | 593496.69 | 778828.62 | 0 | 2 | 0 | 4 | 0 | 4 | 0.35 | 0.37 | 0.50 | 0.53 |
| P27546 | Map4 | Microtubule-associated protein 4 | 708.59 | 16.44 | 117429 | 14 | 24224730 | 23051850 | 20288632 | 26458848 | 19159012 | 27661254 | 22494268 | 22347686 | 24595780 | 20352644 | 14372079 | 16436251 | 16404054 | 12931126 | 12013713 | 21399442 | 24214434 | 26685362 | 30654050 | 31587690 | 0 | 5 | 0 | 7 | 0 | 7 | 0.64 | 0.61 | 1.19 | 1.15 |
| P21708 | Mapk3 | Mitogen-activated protein kinase 3 | 394.97 | 25.26 | 43081 | 9 | 22680142 | 20679622 | 20476836 | 26614924 | 24110434 | 21011560 | 18520278 | 25640478 | 26307282 | 25922306 | 9022878 | 13135374 | 16628484 | 12307230 | 14553613 | 19800010 | 21250588 | 26608302 | 24474420 | 21779054 | 0 | 12 | 0 | 12 | 0 | 12 | 0.57 | 0.56 | 0.99 | 0.97 |
| P49138 | Mapkapk2 | MAP kinase-activated protein kinase 2 | 82.25 | 5.18 | 44050 | 2 | 2074698.5 | 1636873 | 2728220 | 3192011 | 1851495.88 | 2273296.25 | 1593919.5 | 1945089.62 | 2279999 | 2295940.5 | 1574436.75 | 1317869.12 | 1067353.75 | 1408248.5 | 1333513.38 | 1950090.12 | 1822964.5 | 2017713.38 | 1613537 | 2055966.25 | 0 | 7 | 0 | 8 | 0 | 8 | 0.58 | 0.65 | 0.82 | 0.91 |
| Q61166 | Mapre1 | Microtubule-associated protein RP/EB family member 1 | 466.15 | 34.33 | 30016 | 7 | 25363538 | 22543502 | 24576258 | 22981514 | 21585326 | 24318858 | 22723282 | 25458536 | 20997186 | 22767938 | 11034099 | 13912940 | 12429939 | 12900727 | 9075289 | 30847968 | 28192894 | 24124650 | 23718524 | 23780502 | 0 | 6 | 0 | 6 | 0 | 6 | 0.51 | 0.51 | 1.12 | 1.12 |
| P26645 | Marcks | Myristoylated alanine-rich C-kinase substrate | 344.4 | 34.63 | 29661 | 7 | 12340544 | 12049810 | 12312248 | 14965269 | 12410288 | 13128232 | 10726036 | 12224280 | 11899050 | 15501013 | 4689887 | 5571308 | 6577917 | 7146906.5 | 6715054 | 12027066 | 11280507 | 12942946 | 14005588 | 15665061 | 0 | 12 | 0 | 10 | 0 | 12 | 0.48 | 0.48 | 1.03 | 1.04 |
| P28667 | Marcksl1 | MARCKS-related protein | 235.32 | 34.5 | 20165 | 4 | 12677612 | 12099748 | 8857716 | 10766777 | 7996091 | 13870767 | 11690253 | 8998612 | 8261307.5 | 10189743 | 6593127 | 7158455 | 6134539 | 7129016 | 5710903.5 | 11752624 | 13398182 | 10650684 | 9622999 | 10731034 | 0 | 11 | 0 | 9 | 0 | 11 | 0.62 | 0.62 | 1.07 | 1.06 |
| Q9Z2D8 | Mbd3 | Methyl-CpG-binding domain protein 3 | 266.58 | 12.63 | 32168 | 4 | 1940188.25 | 2232216.75 | 2022750.75 | 1926635.38 | 2051323 | 1306859.12 | 2600331 | 1344306.88 | 1752982.5 | 1677560.25 | 762122.44 | 1052624.5 | 1028047.44 | 470699.38 | 653521.75 | 2272112.5 | 1077435.38 | 1782861.25 | 1031583.75 | 1096759.38 | 0 | 5 | 0 | 8 | 0 | 8 | 0.39 | 0.46 | 0.71 | 0.84 |
| P97287 | Mcl1 | Induced myeloid leukemia cell differentiation protein Mcl-1 homolog | 85.46 | 10.27 | 35217 | 2 | 188849.47 | 629096.38 | 231262.31 | 270913.34 | 409582.25 | 2 | 290190.94 | 219409.2 | 218303.42 | 756750.25 | 1041959.38 | 965125.81 | 1235917 | 1231122.88 | 1327856.12 | 2 | 2 | 896149.62 | 983151.38 | 1059280.38 | 0 | 10 | 0 | 9 | 0 | 10 | 3.35 | 3.91 | 1.70 | 1.98 |
| P08249 | Mdh2 | Malate dehydrogenase, mitochondrial | 1660.27 | 70.41 | 35611 | 26 | 156491234 | 179266498 | 162613058 | 189021090 | 172215122 | 189525922 | 165182690 | 196283874 | 182708578 | 199294098 | 241602178 | 214831170 | 227767010 | 283692354 | 295479394 | 189399058 | 181600914 | 226280642 | 229399794 | 224283810 | 0 | 1 | 0 | 7 | 0 | 7 | 1.47 | 1.35 | 1.22 | 1.13 |
| A0A096 | Mef2c | Myocyte-specific enhancer factor 2C | 50.12 | 2.11 | 51217 | 1 | 653169.5 | 992310.56 | 1435162.75 | 1802649.5 | 1306426 | 1082852.25 | 871693.19 | 912251.06 | 882648.12 | 1335805 | 2 | 2 | 2 | 529752.94 | 361485.06 | 1183822.25 | 810601.25 | 739613.56 | 901207 | 926765.5 | 0 | 6 | 0 | 9 | 0 | 9 | 0.14 | 0.18 | 0.74 | 0.90 |
| Q9DC37 | Mfsd1 | Major facilitator superfamily domain-containing protein 1 | 131.8 | 8.19 | 51396 | 3 | 66925.58 | 71070.46 | 277539.81 | 325388.09 | 520446.53 | 706534.88 | 341025.62 | 1224867.25 | 871317.44 | 389695.88 | 2041294.75 | 1586709 | 4156942.75 | 4212782.5 | 3551675.25 | 964719.25 | 2 | 1569121.25 | 1326493.75 | 644430 | 0 | 5 | 0 | 9 | 0 | 9 | 12.33 | 4.40 | 3.57 | 1.27 |
| P27808 | Mgat1 | Alpha-1,3-mannosyl-glycoprotein 2-beta-N-acetylglucosaminyltransferase | 42.3 | 1.79 | 51690 | 1 | 2 | 2 | 2 | 2 | 2 | 2 | 2 | 505149.5 | 215165.3 | 2 | 1040889.12 | 845396.75 | 737526.44 | 881087.56 | 682514.25 | 2 | 2 | 548608.31 | 2 | 2 | 0 | 10 | 0 | 11 | 0 | 11 | 418741.41 | 5.81 | 54861.63 | 0.76 |
| P34960 | Mmp12 | Macrophage metalloelastase | 826.32 | 32.35 | 54971 | 16 | 8923396 | 13056070 | 2899567.5 | 3306608.25 | 4041001.5 | 9698140 | 14138530 | 8938428 | 11708752 | 9626907 | 71537586 | 110430474 | 55559586 | 39557898 | 59303914 | 13179429 | 12258858 | 13332498 | 11676975 | 8454364 | 0 | 4 | 0 | 10 | 0 | 10 | 10.44 | 6.22 | 1.83 | 1.09 |
| D0QMC3 | Mndal | Myeloid cell nuclear differentiation antigen-like protein | 604.6 | 22.49 | 60526 | 10 | 6893806.5 | 6762796 | 6899946 | 8467844 | 7044165 | 7065211 | 6683493.5 | 7077912.5 | 7274583 | 7961776 | 1991161.12 | 2951066.75 | 3462431.5 | 910768.69 | 4816349 | 4785538.5 | 5024533 | 6334546.5 | 5169587.5 | 4217970 | 0 | 0 | 0 | 0 | 0 | 0 | 0.39 | 0.39 | 0.71 | 0.71 |
| Q3T1J9 | Mob1a | MOB kinase activator 1A | 187.75 | 21.76 | 25080 | 4 | 11806558 | 13315284 | 13929395 | 12904683 | 11735843 | 11862706 | 10700680 | 11835682 | 10991974 | 11792921 | 5910976 | 8045097.5 | 8506417 | 9261611 | 9258878 | 11638975 | 10315143 | 11980184 | 11727154 | 11619262 | 0 | 2 | 0 | 11 | 0 | 11 | 0.64 | 0.72 | 0.90 | 1.00 |
| Q80UM7 | Mogs | Mannosyl-oligosaccharide glucosidase | 728.44 | 20.62 | 91831 | 11 | 359721.44 | 2 | 463798.38 | 273751.12 | 343742.66 | 501687.34 | 349571.78 | 2 | 502441.91 | 939755.25 | 2235877 | 2079272.5 | 2779748.25 | 1000823.44 | 2416330.75 | 2 | 2 | 547104.31 | 2 | 433724.72 | 0 | 10.5 | 0 | 7.5 | 0 | 10.5 | 7.29 | 4.58 | 0.68 | 0.43 |
| A1L314 | Mpeg1 | Macrophage-expressed gene 1 protein | 1840.74 | 54.98 | 78390 | 26 | 16524213 | 16162328 | 24126432 | 22175300 | 26341332 | 26594852 | 27589238 | 38793926 | 41408134 | 39330802 | 230335586 | 217198898 | 270221090 | 233340978 | 271181218 | 24820612 | 31884024 | 56792498 | 54802182 | 51363146 | 0 | 1 | 0 | 8 | 0 | 8 | 11.60 | 7.04 | 2.09 | 1.26 |
| Q9DB15 | Mrpl12 | 39S ribosomal protein L12, mitochondrial | 51.6 | 6.97 | 21708 | 1 | 2 | 2 | 2 | 2 | 2 | 2 | 2 | 2 | 2 | 513417.78 | 1694530.38 | 1024080.44 | 907024.12 | 1202353.5 | 1152627.75 | 2 | 2 | 2 | 2 | 2 | 0 | 12.5 | 0 | 10 | 0 | 12.5 | 598061.62 | 11.65 | 1.00 | 0.00 |
| Q9D1N9 | Mrpl21 | 39S ribosomal protein L21, mitochondrial | 177.53 | 17.7 | 23366 | 3 | 408183.66 | 897826.69 | 717081 | 302972.16 | 311384.34 | 710793.88 | 511730.81 | 206778.38 | 437579.5 | 479329.47 | 4073321.5 | 2806251 | 2887193.75 | 2492870.75 | 2551461 | 306327.84 | 420573.09 | 480779.12 | 358503.91 | 350933.59 | 0 | 11 | 0 | 7 | 0 | 11 | 5.62 | 6.31 | 0.73 | 0.82 |
| Q9CY73 | Mrpl44 | 39S ribosomal protein L44, mitochondrial | 222.87 | 17.42 | 37527 | 4 | 1546855.88 | 1363853 | 1949733 | 2149149 | 1133177.75 | 1416111.88 | 1365736 | 1962573 | 2177886.25 | 1159615.25 | 381563.78 | 1059377.5 | 1085884.62 | 961589.62 | 1111026.75 | 1268143.75 | 796855.38 | 1416728.88 | 1459449.5 | 1112039.25 | 0 | 5 | 0 | 7 | 0 | 7 | 0.56 | 0.57 | 0.74 | 0.75 |
| Q9JIK9 | Mrps34 | 28S ribosomal protein S34, mitochondrial | 128.94 | 10.09 | 25827 | 3 | 1581247.5 | 2026168.88 | 1500049 | 1047059.31 | 1686311.12 | 1502554 | 1234802.75 | 1964035.62 | 2181164.25 | 2037064.25 | 3358026.25 | 3326065.5 | 4793346 | 4078818 | 3598151 | 1744342.25 | 800204.69 | 1914582 | 1969660.5 | 1035142.38 | 0 | 12 | 0 | 7 | 0 | 12 | 2.44 | 2.15 | 0.95 | 0.84 |
| Q791V5 | Mtch2 | Mitochondrial carrier homolog 2 | 493.73 | 40.26 | 33499 | 9 | 21300666 | 25145198 | 17748042 | 18043398 | 24022332 | 22520314 | 22030334 | 19314234 | 27092348 | 25325352 | 28126458 | 30675712 | 28062624 | 30267702 | 30451634 | 19836344 | 26478974 | 28391534 | 25928412 | 24475570 | 0 | 4 | 0 | 8 | 0 | 8 | 1.39 | 1.27 | 1.18 | 1.08 |
| P18155 | Mthfd2 | Bifunctional methylenetetrahydrofolate dehydrogenase/cyclohydrolase, mitochondrial | 716.33 | 46.86 | 37863 | 13 | 2876670.25 | 3662915.5 | 2877633.5 | 3169769.75 | 3125291.25 | 2438498.5 | 3754124.5 | 3133455.5 | 3870431.5 | 3133534.25 | 8668416 | 9149901 | 8635173 | 9915655 | 10729822 | 3790952.75 | 2160352 | 3966437 | 4080383.25 | 4658974 | 0 | 5 | 0 | 6 | 0 | 6 | 3.00 | 2.88 | 1.19 | 1.14 |
| Q8VEL2 | Mtmr14 | Myotubularin-related protein 14 | 43.01 | 1.39 | 72443 | 1 | 595282.62 | 378947.41 | 420663.53 | 585486.75 | 523047.97 | 616517.06 | 467709.44 | 523714.72 | 576818.38 | 427866.47 | 261782.48 | 181290.53 | 249701.34 | 2 | 200625.33 | 555671.06 | 430959.16 | 452093.88 | 489862.62 | 352665.31 | 0 | 9 | 0 | 7 | 0 | 9 | 0.36 | 0.34 | 0.91 | 0.87 |
| P03921 | Mtnd5 | NADH-ubiquinone oxidoreductase chain 5 | 180.86 | 6.59 | 68475 | 3 | 788216.38 | 808854.81 | 1166704.62 | 771250.69 | 900871.44 | 843731 | 574271.25 | 946179.5 | 802878.06 | 1055978.62 | 1517394.38 | 1689039.75 | 1978260.12 | 2174311.5 | 1775105.5 | 218060.91 | 700296.88 | 948975.75 | 649295.81 | 948953.06 | 0 | 8 | 0 | 10 | 0 | 10 | 2.06 | 2.16 | 0.78 | 0.82 |
| Q9CZU3 | Mtrex | Exosome RNA helicase MTR4 | 331.33 | 6.83 | 117636 | 6 | 3652209 | 4046258 | 3657100.75 | 3074822.25 | 4169989.5 | 3525175 | 2461441.5 | 3653002.75 | 4500285 | 2825856.25 | 1827750.75 | 1645807.38 | 2378964.5 | 2351225 | 2418617 | 3915123 | 2359599.75 | 4000069.5 | 4574835 | 4057627.5 | 0 | 10 | 0 | 8 | 0 | 10 | 0.57 | 0.63 | 1.02 | 1.11 |
| Q61879 | Myh10 | Myosin-10 | 1148.12 | 10.98 | 228996 | 19 | 1608561.5 | 2316205.25 | 4848067.5 | 3926900 | 2938985.5 | 1769290.38 | 2459663.25 | 3561705.5 | 3345990.5 | 1544438.5 | 2 | 245584.28 | 695842.69 | 262588.81 | 612966.62 | 2145310.5 | 1269062.25 | 3043663.5 | 1823206 | 2124555.75 | 0 | 6 | 0 | 9 | 0 | 9 | 0.12 | 0.14 | 0.67 | 0.82 |
| Q8VDD5 | Myh9 | Myosin-9 | 8403.48 | 61.17 | 226372 | 124 | 349854338 | 412488162 | 393635746 | 388007842 | 376828866 | 353531650 | 334326722 | 375231234 | 367851554 | 368932642 | 274189794 | 286024194 | 297944898 | 278640930 | 292438338 | 351493762 | 376599330 | 340724418 | 332274530 | 335277474 | 0 | 2 | 0 | 8 | 0 | 8 | 0.74 | 0.79 | 0.90 | 0.96 |
| Q60605 | Myl6 | Myosin light polypeptide 6 | 909.01 | 70.2 | 16930 | 13 | 94692122 | 92834690 | 115431330 | 116010706 | 126114410 | 88539938 | 98057562 | 77220050 | 93434250 | 91204386 | 41602066 | 49834198 | 47655006 | 52274834 | 49949530 | 125750546 | 124708362 | 102111746 | 102061178 | 104704714 | 0 | 11 | 0 | 0 | 0 | 11 | 0.44 | 0.54 | 1.03 | 1.25 |
| P70248 | Myo1f | Unconventional myosin-If | 448.65 | 7.83 | 125947 | 8 | 2182253 | 2024930.62 | 2623955.25 | 2901526.75 | 2638791.25 | 2270142.25 | 3140347 | 2342848 | 2946771.75 | 2436571.25 | 1112882.12 | 1110496.62 | 1596375 | 883068.88 | 1191938.38 | 2395669.5 | 2596636.75 | 2083452.62 | 992465.81 | 1733047.75 | 0 | 5 | 0 | 5 | 0 | 5 | 0.48 | 0.45 | 0.79 | 0.75 |
| Q5SUA5 | Myo1g | Unconventional myosin-Ig | 282.84 | 10.16 | 117227 | 7 | 1583138.88 | 1254330.75 | 2072999.88 | 2948060 | 2891201.75 | 1256227.25 | 2115472.75 | 1288081.75 | 2651459 | 1790297.5 | 3872099.5 | 4988930 | 5222631 | 5679516 | 3881525.5 | 523252 | 981102.88 | 1533186.25 | 1422024.62 | 1591274 | 0 | 4 | 0 | 6 | 0 | 6 | 2.20 | 2.60 | 0.56 | 0.66 |
| Q9D7V9 | Naaa | N-acylethanolamine-hydrolyzing acid amidase | 59.3 | 4.7 | 40075 | 1 | 363123.12 | 292916.97 | 290651.59 | 203011.05 | 299853.59 | 311291.69 | 292648.16 | 365120.25 | 346967.34 | 451450.06 | 151250.78 | 133950.78 | 55566.71 | 2 | 106855.7 | 224956.39 | 436417.72 | 350475.59 | 339014.47 | 269554.12 | 0 | 10 | 0 | 9 | 0 | 10 | 0.31 | 0.25 | 1.12 | 0.92 |
| Q9QWR8 | Naga | Alpha-N-acetylgalactosaminidase | 531.69 | 27.95 | 47235 | 9 | 12359932 | 10609127 | 15761688 | 18266666 | 17437730 | 9943582 | 10366070 | 11385565 | 11921500 | 11914598 | 5098747.5 | 3398108.25 | 4276914.5 | 3416956 | 4154580 | 12747640 | 14981079 | 11717961 | 12931074 | 10080004 | 0 | 7 | 0 | 6 | 0 | 7 | 0.27 | 0.37 | 0.84 | 1.12 |
| Q9QZ08 | Nagk | N-acetyl-D-glucosamine kinase | 166.73 | 8.16 | 37268 | 3 | 3053377 | 3186105.5 | 2681118.75 | 1833943.88 | 4354012 | 2084491.62 | 2752237.75 | 1615376.38 | 3273012.25 | 2140421.75 | 829849.25 | 846692.56 | 1001813.44 | 1178787.5 | 912425.94 | 2571273.5 | 1997661.62 | 2150252.25 | 1693553.12 | 1690950.62 | 0 | 3 | 0 | 9 | 0 | 9 | 0.32 | 0.40 | 0.67 | 0.85 |
| Q99KQ4 | Nampt | Nicotinamide phosphoribosyltransferase | 553.8 | 19.55 | 55447 | 9 | 24966870 | 23684338 | 17995112 | 21239972 | 23078814 | 14537018 | 18200334 | 20695948 | 22370062 | 19502956 | 6452214.5 | 12706066 | 11620621 | 7654008 | 11872511 | 27006566 | 12264037 | 15879142 | 15084174 | 21438036 | 0 | 7 | 0 | 11 | 0 | 11 | 0.45 | 0.53 | 0.83 | 0.96 |
| Q8BP47 | Nars | Asparagine--tRNA ligase, cytoplasmic | 1188.9 | 38.64 | 64279 | 20 | 31804814 | 33833638 | 28036132 | 29310378 | 37304910 | 31173156 | 39378294 | 37759178 | 37910234 | 37735682 | 58012282 | 56377894 | 57184398 | 48501910 | 58461622 | 43794998 | 37515638 | 43010070 | 42454946 | 39100374 | 0 | 0 | 0 | 5 | 0 | 5 | 1.74 | 1.51 | 1.28 | 1.12 |
| Q8K224 | Nat10 | RNA cytidine acetyltransferase | 662.64 | 13.18 | 115419 | 11 | 5094473 | 4581250 | 4835921 | 4890791 | 4062015 | 5944862 | 4778394 | 4161183.5 | 4432411.5 | 4774608.5 | 8406592 | 6691413 | 7915289.5 | 7058765 | 6917014.5 | 3866981 | 5908590 | 5118360 | 5307598 | 4477703.5 | 0 | 9 | 0 | 11 | 0 | 11 | 1.58 | 1.54 | 1.05 | 1.02 |
| Q8C156 | Ncaph | Condensin complex subunit 2 | 310.22 | 10.26 | 82303 | 6 | 5521893.5 | 4810283 | 5856218.5 | 6426872.5 | 6521466.5 | 4906911.5 | 4885790 | 6116669 | 5394795 | 6201698 | 1666832 | 2458803.25 | 2517309.5 | 2168312 | 1927631 | 5288410.5 | 5877861.5 | 7445426.5 | 5279768.5 | 7581697 | 0 | 10 | 0 | 8 | 0 | 10 | 0.37 | 0.39 | 1.08 | 1.14 |
| Q8K1X4 | Nckap1l | Nck-associated protein 1-like | 1031.8 | 20.63 | 128905 | 18 | 22851172 | 18089718 | 16759047 | 24889264 | 24243830 | 23257604 | 21144028 | 17757226 | 21447312 | 20323660 | 13849821 | 10856500 | 11017968 | 13959618 | 12483787 | 18723194 | 25142020 | 22453040 | 25918880 | 19159376 | 0 | 9 | 0 | 9 | 0 | 9 | 0.58 | 0.60 | 1.04 | 1.07 |
| Q5XIA1 | Ncln | Nicalin | 350.35 | 14.21 | 62992 | 6 | 2361897.5 | 2007178.88 | 2690032 | 2914078.5 | 1989877.75 | 2851215 | 2358659.5 | 2406724.75 | 3158818.75 | 3388624 | 5428977 | 4475739.5 | 6876282 | 6385164 | 7403516.5 | 2375040 | 3738691.5 | 3569224.75 | 3569798.5 | 1988474.88 | 0 | 7 | 0 | 9 | 0 | 9 | 2.56 | 2.16 | 1.27 | 1.08 |
| Q8VCB1 | Ndc1 | Nucleoporin NDC1 | 267.23 | 9.51 | 75409 | 5 | 695860.31 | 278639.16 | 588172.81 | 969604.38 | 330837.53 | 537742.06 | 1176861.5 | 1423892.75 | 1452201.12 | 769035.06 | 1845347.5 | 1755499.12 | 2789331.5 | 2344166.25 | 2448270 | 2 | 951978 | 1666394.88 | 460800.28 | 1427610.88 | 0 | 9 | 0 | 11 | 0 | 11 | 3.91 | 2.09 | 1.57 | 0.84 |
| Q9ERS2 | Ndufa13 | NADH dehydrogenase [ubiquinone] 1 alpha subcomplex subunit 13 | 209.49 | 30.56 | 16860 | 4 | 6764583.5 | 5480025.5 | 7088484 | 6463165 | 7985830 | 6041443.5 | 5039351 | 5477610 | 7086066 | 6632238.5 | 3950464.5 | 3709132.75 | 4660833.5 | 3384528.75 | 4019556.5 | 6159757 | 7048975 | 6297673 | 6248064 | 5379534 | 0 | 6 | 0 | 11 | 0 | 11 | 0.58 | 0.65 | 0.92 | 1.03 |
| Q9CQ75 | Ndufa2 | NADH dehydrogenase [ubiquinone] 1 alpha subcomplex subunit 2 | 180.68 | 33.33 | 10916 | 3 | 2597328.25 | 2199301.25 | 1944487.12 | 1853403.88 | 1700685.5 | 2592928.5 | 2477623.75 | 1609839 | 1501352.12 | 1535967.62 | 998798.88 | 1008313.38 | 1211659 | 1260986.38 | 1361669.25 | 1798207.25 | 2512981.25 | 1978317.62 | 1853699.88 | 2262556 | 0 | 11 | 0 | 9 | 0 | 11 | 0.57 | 0.60 | 1.01 | 1.07 |
| Q9DC69 | Ndufa9 | NADH dehydrogenase [ubiquinone] 1 alpha subcomplex subunit 9, mitochondrial | 423.58 | 27.59 | 42525 | 7 | 5430020 | 4816393.5 | 5250783 | 6420175 | 5530324 | 4921741 | 4382376 | 5353719 | 4713723 | 4926963.5 | 6626831 | 8204391 | 7099078 | 7285989 | 7952753 | 4518747.5 | 6346910 | 6364841 | 5466418.5 | 5393559 | 0 | 12 | 0 | 4 | 0 | 12 | 1.35 | 1.53 | 1.02 | 1.16 |
| Q6P5H2 | Nes | Nestin | 3779.04 | 38.84 | 207124 | 56 | 82630850 | 83543066 | 87475674 | 79315674 | 75873674 | 86683522 | 85050642 | 76554850 | 71168058 | 79394706 | 55011118 | 56721710 | 52528002 | 60304046 | 50969522 | 86807266 | 82363346 | 77629242 | 74059122 | 73155674 | 0 | 7 | 0 | 12 | 0 | 12 | 0.67 | 0.69 | 0.96 | 0.99 |
| Q60591 | Nfatc2 | Nuclear factor of activated T-cells, cytoplasmic 2 | 183.25 | 7.66 | 100020 | 4 | 2152759.5 | 2687238.25 | 2218399 | 2679699.25 | 2736416.5 | 3473474.25 | 3619936.25 | 2593115.5 | 2325227.25 | 2895045.75 | 1854546.25 | 1555784.25 | 1119287.25 | 1128603 | 1014302.88 | 2416785.75 | 2595109.5 | 1963661.75 | 2453731 | 2780668 | 0 | 11 | 0 | 6 | 0 | 11 | 0.53 | 0.45 | 0.98 | 0.82 |
| O54910 | Nfkbie | NF-kappa-B inhibitor epsilon | 68.07 | 3.3 | 39200 | 1 | 710709.75 | 412876.19 | 581061.31 | 709230.81 | 630205.94 | 733427.19 | 584341.88 | 559367.69 | 929797.75 | 601614.56 | 294136.62 | 183295.42 | 280224.16 | 278984.47 | 234574.44 | 432566.53 | 608283.62 | 443356.38 | 2 | 483614.44 | 0 | 5 | 0 | 3 | 0 | 5 | 0.42 | 0.37 | 0.65 | 0.58 |
| Q8BHY2 | Noc4l | Nucleolar complex protein 4 homolog | 141.46 | 8.33 | 58676 | 3 | 482483.81 | 367644.16 | 375306.72 | 810527.62 | 430970.44 | 381965.22 | 549821.88 | 390075.38 | 671868.62 | 643146.81 | 2145436.25 | 1800153.62 | 2222172 | 1652804.5 | 1924756.38 | 636618.75 | 701632.12 | 1130868.12 | 778111.75 | 712514.75 | 0 | 4 | 0 | 2 | 0 | 4 | 3.95 | 3.70 | 1.61 | 1.50 |
| Q8BJW5 | Nol11 | Nucleolar protein 11 | 153.78 | 4.7 | 80818 | 3 | 464722.16 | 2 | 696716.06 | 129107.3 | 2 | 2 | 311365 | 330025.44 | 666895.94 | 520841.44 | 1091020.88 | 901809.25 | 1120244 | 951453.5 | 1077342.12 | 393257.16 | 2 | 901448.44 | 414319.69 | 460637.38 | 0 | 10 | 0 | 10.5 | 0 | 10.5 | 3.98 | 2.81 | 1.68 | 1.19 |
| Q6GQT9 | Nomo1 | Nodal modulator 1 | 1444.12 | 29.98 | 133420 | 26 | 13885431 | 16211102 | 14992131 | 13344158 | 14377059 | 15075999 | 14709042 | 14809197 | 15125735 | 17485566 | 30640490 | 26349218 | 34732982 | 36688214 | 37390762 | 13504927 | 14120330 | 16785052 | 16430040 | 14640522 | 0 | 9 | 0 | 8 | 0 | 9 | 2.28 | 2.15 | 1.04 | 0.98 |
| Q922K7 | Nop2 | Probable 28S rRNA (cytosine-C(5))-methyltransferase | 895.93 | 24.84 | 86752 | 16 | 8843372 | 7258692 | 8443427 | 7876211 | 8192503.5 | 8632370 | 8133005 | 6747371 | 3752513.75 | 5162488.5 | 18339070 | 16182421 | 13792965 | 12385329 | 13906157 | 9195034 | 7181103 | 10268502 | 7478798.5 | 9958964 | 0 | 9 | 0 | 4 | 0 | 9 | 1.84 | 2.30 | 1.09 | 1.36 |
| Q9D6Z1 | Nop56 | Nucleolar protein 56 | 1456.35 | 50.52 | 64464 | 22 | 24714460 | 26917606 | 25666888 | 26463252 | 29232222 | 22555780 | 24241564 | 30136682 | 30168178 | 29537040 | 45194046 | 44934298 | 35367206 | 30939950 | 41319714 | 20329466 | 16356184 | 26771932 | 28669706 | 28750968 | 0 | 11 | 0 | 6 | 0 | 11 | 1.49 | 1.45 | 0.91 | 0.88 |
| Q6DFW4 | Nop58 | Nucleolar protein 58 | 850.82 | 29.85 | 60343 | 13 | 12179462 | 13447040 | 14403050 | 12834634 | 14660234 | 11739022 | 13425088 | 12108186 | 12913006 | 11137818 | 8394555 | 8959250 | 9740966 | 8547786 | 9009165 | 12226457 | 12551720 | 9143782 | 11093843 | 12081182 | 0 | 2 | 0 | 8 | 0 | 8 | 0.66 | 0.73 | 0.85 | 0.93 |
| Q8BMC4 | Nop9 | Nucleolar protein 9 | 186.68 | 6.6 | 70046 | 3 | 487666.84 | 378638.81 | 315864.78 | 240340.75 | 313415.75 | 226374.36 | 273314.62 | 520693.72 | 473362.56 | 424334.16 | 1369486.5 | 1198407.5 | 1109365.88 | 745497.94 | 565546.31 | 260704.33 | 118071.19 | 2 | 322541.28 | 2 | 0 | 4 | 0 | 3 | 0 | 4 | 2.87 | 2.60 | 0.40 | 0.37 |
| Q9Z0J0 | Npc2 | NPC intracellular cholesterol transporter 2 | 252.08 | 41.61 | 16442 | 5 | 48277046 | 38412902 | 24695770 | 34918790 | 26873782 | 32027882 | 32829162 | 28393774 | 25873526 | 34777866 | 4590716 | 5648711 | 5558537.5 | 3658984 | 5036400 | 37588486 | 20821178 | 17530012 | 16897220 | 21071624 | 0 | 3 | 0 | 5 | 0 | 5 | 0.14 | 0.16 | 0.66 | 0.74 |
| Q11011 | Npepps | Puromycin-sensitive aminopeptidase | 809.73 | 19.24 | 103325 | 14 | 21286894 | 17981266 | 18187018 | 19278726 | 17584040 | 17894802 | 16894598 | 15645107 | 18984584 | 18839842 | 11399520 | 13621685 | 12588147 | 11649918 | 12595574 | 19956640 | 15884341 | 18945772 | 19566078 | 16276044 | 0 | 11 | 0 | 9 | 0 | 11 | 0.66 | 0.70 | 0.96 | 1.03 |
| Q9CPP0 | Npm3 | Nucleoplasmin-3 | 104.02 | 9.14 | 19023 | 1 | 2 | 400735.5 | 2 | 2 | 141995.56 | 2 | 188897 | 217897.2 | 223805.11 | 113580.45 | 1558095.12 | 1011967.69 | 914635.75 | 1020619.31 | 1013653.88 | 182435.44 | 2 | 245548.44 | 2 | 86724.54 | 0 | 11 | 0 | 9 | 0 | 11 | 10.17 | 7.42 | 0.95 | 0.69 |
| P08556 | Nras | GTPase NRas | 252.29 | 31.75 | 21199 | 4 | 1505825.88 | 1992710.25 | 1827825.75 | 1627603.38 | 1625730.5 | 1632257.12 | 1799925.75 | 2363048 | 1597107.38 | 1580743.5 | 3801214.5 | 3752262 | 3992486.25 | 4618747 | 4688465.5 | 1851091.5 | 1779815.25 | 1842034.38 | 2 | 1354475.5 | 0 | 11 | 0 | 11 | 0 | 11 | 2.43 | 2.32 | 0.80 | 0.76 |
| P97333 | Nrp1 | Neuropilin-1 | 428.15 | 9.75 | 103000 | 7 | 845147.62 | 362743.38 | 199757.8 | 415883.88 | 113204.8 | 524876.94 | 545478.56 | 647858.81 | 504211.88 | 493869.06 | 2404788.5 | 2124034.5 | 2040700.38 | 1283572.12 | 1551111 | 952371.5 | 448657.5 | 404888.09 | 851657.06 | 618461.25 | 0 | 4 | 0 | 11 | 0 | 11 | 4.86 | 3.46 | 1.69 | 1.21 |
| O35375 | Nrp2 | Neuropilin-2 | 209.45 | 4.19 | 104631 | 4 | 896569.38 | 774227.12 | 1403797 | 625501.81 | 1244303.25 | 1058647.12 | 449272.44 | 774195.94 | 1192028.25 | 556621.25 | 2209377.25 | 2222062 | 2043333 | 2260328.5 | 2114699.5 | 1614882.5 | 425412 | 1159787.25 | 1809376.25 | 1144157.5 | 0 | 9 | 0 | 7 | 0 | 9 | 2.19 | 2.69 | 1.24 | 1.53 |
| Q3UFY7 | Nt5c3b | 7-methylguanosine phosphate-specific 5'-nucleotidase | 214.92 | 23.33 | 34425 | 4 | 2394774 | 2455495 | 2476081.5 | 2248555.5 | 3643989.25 | 2435327.75 | 2206629.25 | 2283397.25 | 2797931.5 | 2348677 | 1041307 | 1315714.88 | 1330885.25 | 1288954.12 | 1175893.62 | 2798423 | 2691859.25 | 2287566.25 | 1942872.62 | 1908851.75 | 0 | 9 | 0 | 11 | 0 | 11 | 0.47 | 0.51 | 0.88 | 0.96 |
| Q02819 | Nucb1 | Nucleobindin-1 | 177.61 | 8.5 | 53409 | 3 | 500953.97 | 421302.78 | 410834.5 | 717871.44 | 307794.59 | 604045 | 340336.19 | 405246.59 | 523062.5 | 503780.25 | 1203602.12 | 916126.06 | 1552629.75 | 1038532.56 | 1416777 | 348868.31 | 541470.06 | 596151.06 | 2 | 333736.91 | 0 | 10 | 0 | 9 | 0 | 10 | 2.60 | 2.58 | 0.77 | 0.77 |
| E9Q7G0 | Numa1 | Nuclear mitotic apparatus protein 1 | 2466.11 | 22.21 | 235630 | 40 | 19486082 | 22440948 | 25189754 | 26811114 | 22685102 | 22959490 | 18977908 | 21892120 | 20031708 | 22809720 | 15317192 | 13045566 | 12584389 | 11650330 | 11715564 | 14569204 | 15813670 | 16976060 | 15772472 | 19687302 | 0 | 1 | 0 | 1 | 0 | 1 | 0.55 | 0.60 | 0.71 | 0.78 |
| P11928 | Oas1a | 2'-5'-oligoadenylate synthase 1A | 631.77 | 37.87 | 42429 | 10 | 3303631 | 3524393.25 | 3408435.5 | 3377048.25 | 4853134.5 | 3590256.75 | 5291204 | 4240534 | 5202379.5 | 5104196 | 10773100 | 10577076 | 12851542 | 5960533.5 | 10445943 | 3179996.5 | 2571737.25 | 3008893.5 | 3019295.75 | 3160463.75 | 0 | 0 | 0 | 0 | 0 | 0 | 2.74 | 2.16 | 0.81 | 0.64 |
| E9Q9A9 | Oas2 | 2'-5'-oligoadenylate synthase 2 | 112.78 | 4.18 | 85051 | 2 | 284571.09 | 271470.75 | 220109.28 | 2 | 83518.27 | 371825.19 | 351894.5 | 390447.16 | 447291.75 | 362611.12 | 638630.06 | 698067.88 | 883911.31 | 535685.81 | 875197.31 | 181552.08 | 280604.59 | 263616.75 | 285253.91 | 409073.22 | 0 | 6 | 0 | 4 | 0 | 6 | 4.22 | 1.89 | 1.65 | 0.74 |
| Q8VI93 | Oas3 | 2'-5'-oligoadenylate synthase 3 | 927.19 | 19.33 | 126333 | 17 | 11612216 | 11760700 | 12480764 | 14431957 | 12397756 | 12092327 | 11576338 | 11575610 | 16958602 | 12467671 | 9233440 | 9120158 | 11011727 | 7660579 | 8836050 | 12478716 | 8774175 | 12345639 | 14349392 | 14873975 | 0 | 11 | 0 | 10 | 0 | 11 | 0.73 | 0.71 | 1.00 | 0.97 |
| B9EJ86 | Osbpl8 | Oxysterol-binding protein-related protein 8 | 1190.42 | 26.1 | 101269 | 18 | 18057228 | 19067434 | 14192709 | 14005068 | 17049184 | 17167398 | 21524196 | 18747644 | 21728194 | 20761762 | 25207212 | 29354882 | 32024490 | 23710978 | 29522204 | 18197060 | 13415282 | 19141312 | 17938698 | 17556794 | 0 | 10 | 0 | 5 | 0 | 10 | 1.70 | 1.40 | 1.05 | 0.86 |
| Q62422 | Ostf1 | Osteoclast-stimulating factor 1 | 226.09 | 24.65 | 23783 | 4 | 10791940 | 11326515 | 13422198 | 13041645 | 14544777 | 9333125 | 10519160 | 13770844 | 12994330 | 12864358 | 6615299.5 | 7737310.5 | 5831431.5 | 7569145 | 8680595 | 11726163 | 10168663 | 12744627 | 13129828 | 14365744 | 0 | 11 | 0 | 11 | 0 | 11 | 0.58 | 0.61 | 0.98 | 1.04 |
| B2RRE7 | Otud4 | OTU domain-containing protein 4 | 53.34 | 0.72 | 123055 | 1 | 785472.25 | 586863.94 | 1028371.12 | 854736.19 | 709026.44 | 637968.5 | 646698.19 | 731684.75 | 868945.38 | 764413.19 | 2 | 305291.53 | 506463.06 | 263428.81 | 511795.75 | 1216086.75 | 457646.59 | 644749.94 | 611065.12 | 863767.19 | 0 | 11 | 0 | 10 | 0 | 11 | 0.40 | 0.43 | 0.96 | 1.04 |
| P09103 | P4hb | Protein disulfide-isomerase | 2224.86 | 59.53 | 57058 | 33 | 114519482 | 150233922 | 132855250 | 122891650 | 143822258 | 131357498 | 123295634 | 152910850 | 151538914 | 161825010 | 182502898 | 181850354 | 220581986 | 169958370 | 211269858 | 116546122 | 98607234 | 156125186 | 153432226 | 162141378 | 0 | 9 | 0 | 12 | 0 | 12 | 1.45 | 1.34 | 1.03 | 0.95 |
| O35263 | Pafah1b3 | Platelet-activating factor acetylhydrolase IB subunit gamma | 66.02 | 3.88 | 25863 | 1 | 2 | 2 | 2 | 2 | 2 | 2 | 2 | 2 | 2 | 2 | 413379.94 | 671812.25 | 756690.94 | 574144.12 | 618646.88 | 2 | 2 | 2 | 2 | 2 | 0 | 12.5 | 0 | 12.5 | 0 | 12.5 | 303467.41 | 303467.41 | 1.00 | 1.00 |
| Q9DCE5 | Pak1ip1 | p21-activated protein kinase-interacting protein 1 | 202.72 | 13.61 | 42116 | 4 | 1907105.88 | 2016600.75 | 2231103.25 | 1124648 | 1485698 | 2437194 | 1045348.69 | 1950239 | 1896721.75 | 1929147.75 | 5736728 | 3492175 | 3144513 | 2827822 | 4486520 | 1369812.75 | 2432455.5 | 1828401.38 | 2121435.5 | 1377698.12 | 0 | 12 | 0 | 11 | 0 | 12 | 2.25 | 2.13 | 1.04 | 0.99 |
| Q99LX0 | Park7 | Protein/nucleic acid deglycase DJ-1 | 722.07 | 66.14 | 20021 | 11 | 78694658 | 64264302 | 66941346 | 68440810 | 50164094 | 79454594 | 58696734 | 68228786 | 52849882 | 70853074 | 37760494 | 37969918 | 31671824 | 45796206 | 34576822 | 82006202 | 106898666 | 74054738 | 74932866 | 76516146 | 0 | 3 | 0 | 3 | 0 | 3 | 0.57 | 0.57 | 1.26 | 1.26 |
| Q8BFV2 | Pcid2 | PCI domain-containing protein 2 | 46.15 | 2.51 | 46132 | 1 | 2 | 2 | 250021.14 | 125836.33 | 229244.45 | 2 | 2 | 187216.22 | 299071.44 | 177385.98 | 334583.22 | 366549.41 | 541063 | 346771.69 | 374150.72 | 2 | 2 | 195327.44 | 118237.12 | 221104.06 | 0 | 10 | 0 | 12 | 0 | 12 | 3.24 | 2.96 | 0.88 | 0.81 |
| Q8BH04 | Pck2 | Phosphoenolpyruvate carboxykinase [GTP], mitochondrial | 1435.94 | 43.28 | 70528 | 22 | 35227866 | 37885250 | 23587362 | 26004496 | 27810300 | 34771418 | 32445158 | 26792240 | 28357660 | 26213346 | 38652506 | 42818298 | 46889062 | 38998514 | 45055418 | 37405098 | 34916502 | 29689336 | 28191242 | 32629382 | 0 | 9 | 0 | 6 | 0 | 9 | 1.41 | 1.43 | 1.08 | 1.10 |
| Q6NS46 | Pdcd11 | Protein RRP5 homolog | 975.9 | 13 | 207779 | 20 | 6132162.5 | 7580388.5 | 4713025 | 5932954 | 5827317 | 6561284.5 | 5904934.5 | 5506210 | 5704765.5 | 4411420 | 10134853 | 9605872 | 9063067 | 10414823 | 10559770 | 5956619 | 6887934 | 6022601 | 6155115 | 5236416 | 0 | 10 | 0 | 7 | 0 | 10 | 1.65 | 1.77 | 1.00 | 1.08 |
| P56812 | Pdcd5 | Programmed cell death protein 5 | 316.41 | 42.86 | 14275 | 5 | 4469920 | 2829261.75 | 3439697.75 | 3793674.25 | 2974762.75 | 3814479.75 | 3250450.75 | 2171153.5 | 2590284.75 | 2622509.75 | 2052369.25 | 1916812.62 | 1453890.12 | 1571807.5 | 1823255.5 | 3463496.25 | 6266344 | 3849766.5 | 2494609 | 3203517.5 | 0 | 11 | 0 | 7 | 0 | 11 | 0.50 | 0.61 | 1.10 | 1.33 |
| Q9WU78 | Pdcd6ip | Programmed cell death 6-interacting protein | 1755.69 | 40.85 | 96024 | 27 | 46083634 | 41155586 | 44027574 | 38414790 | 47158686 | 37520918 | 45152774 | 46343350 | 47855510 | 38534326 | 49308090 | 63877674 | 56980014 | 47983106 | 62419478 | 49543442 | 41024222 | 39126866 | 39965446 | 42625254 | 0 | 10 | 0 | 12 | 0 | 12 | 1.29 | 1.30 | 0.98 | 0.99 |
| P08003 | Pdia4 | Protein disulfide-isomerase A4 | 2446.24 | 58.15 | 71982 | 40 | 70805106 | 80151162 | 83073042 | 77375466 | 73120162 | 69255962 | 68167754 | 83828586 | 82211426 | 81315226 | 112541330 | 125114514 | 121760498 | 126864722 | 131873794 | 63672930 | 65396930 | 77751346 | 84652154 | 81420026 | 0 | 12 | 0 | 10 | 0 | 12 | 1.61 | 1.61 | 0.97 | 0.97 |
| Q63065 | Pdk1 | [Pyruvate dehydrogenase (acetyl-transferring)] kinase isozyme 1, mitochondrial | 138.89 | 8.53 | 49081 | 3 | 1415448.75 | 1283489.38 | 1333163.25 | 1457126.75 | 1267866 | 1456968.75 | 1704635.5 | 1387054.25 | 1403062 | 1648937.5 | 2 | 337343.09 | 85595.18 | 484188.22 | 2 | 1400480.25 | 925892.44 | 1134142 | 1345652.25 | 470228.28 | 0 | 6 | 0 | 1 | 0 | 6 | 0.13 | 0.12 | 0.78 | 0.69 |
| A4L9P7 | Pds5a | Sister chromatid cohesion protein PDS5 homolog A | 151.01 | 1.95 | 150286 | 2 | 154597.8 | 2 | 2 | 2 | 2 | 2 | 219235.47 | 230401.52 | 2 | 263194.97 | 883707.19 | 859203.75 | 731730.94 | 559910.12 | 802242.31 | 2 | 2 | 2 | 2 | 118210.87 | 0 | 12 | 0 | 6 | 0 | 12 | 24.82 | 5.38 | 0.76 | 0.17 |
| Q8K183 | Pdxk | Pyridoxal kinase | 515.84 | 25.32 | 35015 | 8 | 9186884 | 6987678 | 9905902 | 8348140 | 10052938 | 10111604 | 8496023 | 9903068 | 9829451 | 8060013 | 4807802 | 6246352 | 2989353 | 6247271 | 5532470.5 | 7867077 | 5164282.5 | 5147282 | 7701528.5 | 9083980 | 0 | 4 | 0 | 2 | 0 | 4 | 0.58 | 0.56 | 0.79 | 0.75 |

| | | | | | | | | | | | | | | | | | | | | | | | | | | | | | | | | | | | | |
|---|---|---|---|---|---|---|---|---|---|---|---|---|---|---|---|---|---|---|---|---|---|---|---|---|---|---|---|---|---|---|---|---|---|---|---|---|
| Q62048 | Pea15 | Astrocytic phosphoprotein PEA-15 | 57.07 | 7.69 | 15054 | 1 | 2 | 2 | 287190.75 | 232733.36 | 2 | 2 | 2 | 286723.44 | 2 | 2 | 717405.38 | 410440.09 | 392152.5 | 437535.34 | 384581.88 | 351387.66 | 448471.88 | 2 | 425206.34 | 324166.53 | 0 | 3.5 | 0 | 3 | 0 | 3.5 | 4.50 | 8.17 | 2.98 | 5.40 |
| Q99MZ7 | Pecr | Peroxisomal trans-2-enoyl-CoA reductase | 156.07 | 11.88 | 32410 | 3 | 1325850.5 | 1847321.38 | 1110055.62 | 1406459.12 | 1514281.38 | 1772324 | 1808689.62 | 1355087.62 | 2095742 | 1327678.75 | 2186788.5 | 2905169 | 3032293.5 | 3177936.75 | 2775546 | 1698797.62 | 1374983.75 | 1691746.38 | 2 | 1832638.12 | 0 | 11 | 0 | 10 | 0 | 11 | 1.95 | 1.68 | 0.92 | 0.79 |
| Q9DBD5 | Pelp1 | Proline-, glutamic acid- and leucine-rich protein 1 | 589.87 | 9.8 | 118069 | 10 | 8466155 | 7871384.5 | 8230362.5 | 8394781 | 8288428.5 | 9322766 | 9220849 | 9037279 | 9094262 | 9944895 | 10076641 | 12322360 | 14452496 | 12724513 | 13143623 | 9965409 | 8891672 | 9831016 | 9582173 | 8922564 | 0 | 0 | 0 | 12 | 0 | 12 | 1.52 | 1.35 | 1.14 | 1.01 |
| Q11136 | Pepd | Xaa-Pro dipeptidase | 541.28 | 22.52 | 55029 | 10 | 19124546 | 19551924 | 20142282 | 21426330 | 17979892 | 20776438 | 18723772 | 17727702 | 17098994 | 18245608 | 8415322 | 11312149 | 9745407 | 9913810 | 7866493.5 | 21504780 | 20972542 | 18800478 | 17508138 | 18063210 | 0 | 11 | 0 | 8 | 0 | 11 | 0.48 | 0.51 | 0.99 | 1.05 |
| Q9Z126 | Pf4 | Platelet factor 4 | 42.54 | 11.43 | 11243 | 1 | 2 | 2 | 2 | 2 | 82851.27 | 2 | 2 | 2 | 2 | 2 | 465011.41 | 596885.56 | 435310.44 | 702816.19 | 765936.56 | 145406.72 | 171053.88 | 2 | 109087.7 | 183300.95 | 0 | 3 | 0 | 2.5 | 0 | 3 | 35.80 | 296596.02 | 7.35 | 60885.13 |
| B0BN18 | Pfdn2 | Prefoldin subunit 2 | 254.13 | 30.52 | 16580 | 4 | 9902071 | 8016953 | 9319836 | 7725765.5 | 9146822 | 6017976 | 8535558 | 9421464 | 8693417 | 8053180 | 3614220 | 5860170 | 5442381 | 1195074.88 | 4245354 | 11936823 | 6738616 | 7656454 | 7174653 | 7789167 | 0 | 6 | 0 | 9 | 0 | 9 | 0.46 | 0.50 | 0.94 | 1.01 |
| P62962 | Pfn1 | Profilin-1 | 989.64 | 95 | 14957 | 15 | 162198946 | 196126130 | 149374898 | 195677154 | 191199698 | 152114322 | 160290882 | 184939138 | 179733442 | 197231890 | 261492562 | 240977426 | 227605970 | 233037346 | 227395154 | 169898258 | 169066690 | 101699730 | 186274690 | 191303538 | 0 | 9 | 0 | 12 | 0 | 12 | 1.33 | 1.36 | 0.91 | 0.94 |
| Q9CQ60 | Pgls | 6-phosphogluconolactonase | 387.64 | 40.47 | 27254 | 7 | 13461286 | 13437276 | 13949707 | 14905296 | 13462751 | 14817032 | 12143585 | 14065135 | 14920909 | 14593287 | 10584723 | 9786587 | 10765428 | 11833380 | 11567682 | 13407732 | 14583483 | 16726064 | 19172080 | 16711540 | 0 | 6 | 0 | 7 | 0 | 7 | 0.79 | 0.77 | 1.16 | 1.14 |
| Q9D0F9 | Pgm1 | Phosphoglucomutase-1 | 1344.41 | 44.31 | 61418 | 23 | 35471650 | 34365374 | 28280304 | 31819486 | 32800462 | 36823222 | 33374102 | 33687850 | 35192290 | 33712866 | 19961366 | 21603780 | 23641106 | 22946542 | 23550148 | 35623742 | 36547914 | 36394966 | 39613918 | 35433310 | 0 | 1 | 0 | 4 | 0 | 4 | 0.69 | 0.65 | 1.13 | 1.06 |
| D3ZDK7 | Pgp | Glycerol-3-phosphate phosphatase | 138.05 | 9.66 | 34601 | 2 | 4246172.5 | 3649662.25 | 3143320.75 | 2589302.5 | 3506447.75 | 2265705.25 | 3599063.75 | 2350552.5 | 2671271.25 | 2417777.75 | 654937.69 | 1913880.5 | 2 | 821033.94 | 1157900 | 4803182 | 2189380.75 | 1884600 | 2090667.88 | 2687291.25 | 0 | 6 | 0 | 9 | 0 | 9 | 0.27 | 0.34 | 0.80 | 1.03 |
| Q80UU9 | Pgrmc2 | Membrane-associated progesterone receptor component 2 | 414.19 | 30.41 | 23334 | 4 | 2236147.5 | 3077430.75 | 2020058.5 | 3089826.75 | 3439016.75 | 3901849.5 | 4155729.5 | 3621790 | 3541671.5 | 4001262.5 | 4916554 | 5468922.5 | 6076857.5 | 6186110 | 7183534 | 3419730.5 | 4300598.5 | 3852797.75 | 2780236 | 3270899 | 0 | 5 | 0 | 7 | 0 | 7 | 2.15 | 1.55 | 1.27 | 0.92 |
| Q8BHF7 | Pgs1 | CDP-diacylglycerol--glycerol-3-phosphate 3-phosphatidyltransferase, mitochondrial | 231.45 | 8.32 | 62489 | 4 | 650251.88 | 655954.88 | 561986.88 | 473843.69 | 499483.09 | 697367.62 | 567186.56 | 884309.56 | 646220.75 | 670437 | 1176801.12 | 1719612.5 | 1817931.38 | 1148640.5 | 1928867.75 | 532782.69 | 401099.5 | 1002155.75 | 1002348.88 | 402113.06 | 0 | 12 | 0 | 10 | 0 | 12 | 2.74 | 2.25 | 1.18 | 0.96 |
| P67778 | Phb | Prohibitin | 1752.75 | 80.88 | 29820 | 25 | 215902386 | 216467602 | 194506034 | 206260498 | 199560674 | 227240322 | 230726114 | 205789042 | 213743362 | 234699170 | 300793474 | 278884802 | 369353890 | 357096834 | 342199362 | 234440322 | 267370482 | 246789874 | 258797250 | 232440418 | 0 | 0 | 0 | 2 | 0 | 2 | 1.60 | 1.48 | 1.20 | 1.11 |
| O35129 | Phb2 | Prohibitin-2 | 1315.11 | 66.89 | 33296 | 21 | 18276052 | 22907158 | 19847916 | 20130556 | 14606617 | 20138358 | 17098080 | 22412610 | 15861471 | 26331250 | 156815106 | 98144122 | 119790498 | 99242730 | 93891698 | 18760020 | 17426810 | 25251846 | 22835156 | 25259078 | 0 | 8 | 0 | 9 | 0 | 9 | 5.93 | 5.58 | 1.14 | 1.08 |
| A6H5X4 | Phf11 | PHD finger protein 11 | 41.66 | 2.97 | 37728 | 1 | 1458327 | 2180210.75 | 1417203.38 | 2178104 | 1992728.62 | 1956647.88 | 2049855.25 | 1892231.75 | 1916421.12 | 2205813.75 | 1004646.88 | 1272097.25 | 970532 | 736571.38 | 875836.25 | 2 | 1583367.38 | 1011739 | 1779375.88 | 1585156.25 | 0 | 6 | 0 | 0 | 0 | 6 | 0.53 | 0.48 | 0.65 | 0.59 |
| O55012 | Picalm | Phosphatidylinositol-binding clathrin assembly protein | 872.65 | 24.53 | 69286 | 14 | 34177938 | 38398362 | 30245708 | 30998548 | 30196022 | 34991334 | 33874894 | 26959778 | 27198158 | 26328678 | 15855726 | 16030653 | 16552048 | 16273869 | 14930689 | 36329390 | 37078830 | 29904988 | 29679896 | 31670222 | 0 | 11 | 0 | 6 | 0 | 11 | 0.49 | 0.53 | 1.00 | 1.10 |
| Q8BXQ2 | Pigt | GPI transamidase component PIG-T | 271.42 | 9.28 | 65705 | 5 | 2942434.75 | 2497266 | 2644310.5 | 4130881 | 4292071.5 | 4188008.75 | 2991306.25 | 4149029 | 4220635 | 4976553 | 6951334 | 6286161 | 8831532 | 9410306 | 8544537 | 2474526.75 | 4586751.5 | 4679590 | 4758880 | 4076328.5 | 0 | 7 | 0 | 12 | 0 | 12 | 2.42 | 1.95 | 1.25 | 1.00 |
| P16446 | Pitpna | Phosphatidylinositol transfer protein alpha isoform | 187.84 | 18.82 | 31907 | 4 | 4334530 | 6674632 | 8077067.5 | 6962338 | 7447116 | 6006457 | 5015879 | 5946294.5 | 5919177.5 | 5316310.5 | 2788739.75 | 3326276.5 | 2094219.75 | 2520374.5 | 2649711 | 5610138 | 6759085.5 | 4754622 | 4716331.5 | 6703671 | 0 | 7 | 0 | 12 | 0 | 12 | 0.40 | 0.47 | 0.85 | 1.01 |
| P70268 | Pkn1 | Serine/threonine-protein kinase N1 | 268.76 | 6.66 | 104411 | 6 | 2794838.25 | 3707283 | 3453166 | 3298202.5 | 3197688 | 2347292.5 | 1829337.25 | 1852593.62 | 2811700 | 2822884.25 | 13831608 | 9798076 | 9535423 | 6215663 | 9375427 | 3693195.5 | 2468044.5 | 2159943 | 1750645.38 | 2247562 | 0 | 4 | 0 | 12 | 0 | 12 | 2.96 | 4.18 | 0.75 | 1.06 |
| Q68FH0 | Pkp4 | Plakophilin-4 | 41.65 | 0.67 | 131551 | 1 | 7062563.5 | 6936964.5 | 7689407.5 | 10521566 | 5855743.5 | 8480594 | 8160033.5 | 6514535.5 | 5543420 | 6851786.5 | 3092083.25 | 2310216.25 | 2580117.5 | 3805593.75 | 2900856.5 | 5550161 | 8042114.5 | 4957205.5 | 6392400.5 | 5366923.5 | 0 | 5 | 0 | 5 | 0 | 5 | 0.39 | 0.41 | 0.80 | 0.85 |
| Q8VEB4 | Pla2g15 | Group XV phospholipase A2 | 129.49 | 8.74 | 47307 | 3 | 4583690 | 4575479 | 5407930.5 | 5392345 | 4804198 | 3885929 | 3200262.75 | 3834957.25 | 3377880.5 | 4688599.5 | 1551756.38 | 987242.62 | 901778.94 | 987062.81 | 957817.5 | 3266837.5 | 3807730.25 | 3586422 | 4223966.5 | 2784121.25 | 0 | 0 | 0 | 9 | 0 | 9 | 0.22 | 0.28 | 0.71 | 0.93 |
| P47713 | Pla2g4a | Cytosolic phospholipase A2 | 747.37 | 25.8 | 85222 | 15 | 20304466 | 23770996 | 28871294 | 27812798 | 26023960 | 22527170 | 20861248 | 25426322 | 23689294 | 27097000 | 35088102 | 34042922 | 36816182 | 34356030 | 36528722 | 15799849 | 17825812 | 25708868 | 20152282 | 27661306 | 0 | 5 | 0 | 9 | 0 | 9 | 1.39 | 1.48 | 0.85 | 0.90 |
| Q3TCN2 | Plbd2 | Putative phospholipase B-like 2 | 401.85 | 11.95 | 66289 | 7 | 15179914 | 12461482 | 8577093 | 11910513 | 13111266 | 10340157 | 9010975 | 8689947 | 9978718 | 10127012 | 1463256 | 654220.38 | 2757886 | 1020292.62 | 2518007.25 | 10568254 | 10977196 | 7184815.5 | 8642018 | 7822277 | 0 | 3 | 0 | 10 | 0 | 10 | 0.14 | 0.17 | 0.74 | 0.94 |
| Q8BG07 | Pld4 | 5'-3' exonuclease PLD4 | 1200.8 | 35.19 | 56154 | 17 | 120049650 | 105438362 | 133711450 | 144558450 | 136347266 | 108612434 | 107793666 | 121716666 | 124928306 | 114563858 | 33612342 | 31923874 | 45000322 | 43251086 | 47578250 | 130859050 | 92868234 | 136091410 | 116522178 | 137899298 | 0 | 10 | 0 | 7 | 0 | 10 | 0.31 | 0.35 | 0.96 | 1.06 |
| Q9JHK5 | Plek | Pleckstrin | 611.54 | 42.86 | 39901 | 10 | 5096691.5 | 4976140.5 | 4315840.5 | 2013684.25 | 5440159 | 4564926 | 4514470 | 5465476 | 6743922.5 | 5002198.5 | 11522178 | 10172676 | 11221171 | 9560889 | 9548980 | 5431285.5 | 6585702.5 | 6063172 | 6508705 | 5187562 | 0 | 2 | 0 | 7 | 0 | 7 | 2.38 | 1.98 | 1.36 | 1.13 |
| P20918 | Plg | Plasminogen | 284.73 | 7.76 | 90808 | 6 | 326230.09 | 470106.72 | 4163355 | 3624761.25 | 3971794.5 | 857762.12 | 2183987.5 | 10145757 | 9366997 | 11139141 | 14473096 | 14243628 | 31927990 | 34670938 | 31982884 | 1445346 | 1021498.12 | 9385100 | 8204556.5 | 5765996 | 0 | 6 | 0 | 9 | 0 | 9 | 10.14 | 3.78 | 2.06 | 0.77 |
| P43883 | Plin2 | Perilipin-2 | 128.49 | 7.06 | 46646 | 2 | 1024077.44 | 969294.88 | 520466.09 | 1776630.75 | 2165953.5 | 1477168.5 | 1184535.12 | 2508993 | 2292908.75 | 2810969 | 2 | 2 | 2 | 2 | 265069.78 | 469597.19 | 849910.56 | 2204371 | 1001002.19 | 1673552.25 | 0 | 11 | 0 | 4 | 0 | 11 | 0.04 | 0.03 | 0.96 | 0.60 |
| Q9DBG5 | Plin3 | Perilipin-3 | 866.94 | 46.45 | 47262 | 13 | 30482700 | 27510904 | 25634202 | 32248348 | 26769156 | 29204616 | 26996374 | 22872160 | 25499482 | 28311760 | 13355199 | 12459760 | 12355926 | 13648157 | 12718288 | 30113760 | 28521308 | 27478300 | 28015446 | 27981612 | 0 | 11 | 0 | 7 | 0 | 11 | 0.45 | 0.49 | 1.00 | 1.07 |
| Q9JJ00 | Plscr1 | Phospholipid scramblase 1 | 162.67 | 12.2 | 35914 | 3 | 2734007.25 | 2200552.25 | 2033838.12 | 2582331 | 2311344.5 | 3207438.75 | 2792996 | 2148899.25 | 2478296.5 | 2861763.25 | 4098703.75 | 3502901.5 | 3833390.25 | 3879944.5 | 4522373.5 | 2032689.25 | 1771751.88 | 2018416.38 | 2451088.5 | 2115863.25 | 0 | 4 | 0 | 1 | 0 | 4 | 1.67 | 1.47 | 0.88 | 0.77 |
| P23492 | Pnp | Purine nucleoside phosphorylase | 598.39 | 31.83 | 32277 | 8 | 33232470 | 28558646 | 21936766 | 24280106 | 34489858 | 27551758 | 32179432 | 23696864 | 30479366 | 26921534 | 12566172 | 13037157 | 11674018 | 9223774 | 11664254 | 30929402 | 21531506 | 13419056 | 16808962 | 27898828 | 0 | 5 | 0 | 7 | 0 | 7 | 0.41 | 0.41 | 0.78 | 0.79 |
| O35134 | Polr1a | DNA-directed RNA polymerase I subunit RPA1 | 210.68 | 4.89 | 194110 | 5 | 1067365.62 | 624039.44 | 747117.25 | 727234.5 | 757911.25 | 709639.31 | 922194 | 760081.38 | 647730 | 599176.75 | 2983033.75 | 2481751 | 3783009 | 2676764 | 2080231.5 | 1238834 | 634122.44 | 621126.25 | 665948.19 | 628923.88 | 0 | 8 | 0 | 10 | 0 | 10 | 3.57 | 3.85 | 0.97 | 1.04 |
| P62488 | Polr2g | DNA-directed RNA polymerase II subunit RPB7 | 43.44 | 6.4 | 19294 | 1 | 1095990.88 | 882914.06 | 1167766 | 1022358.31 | 1081986.25 | 1133683.25 | 1256815.25 | 993077.94 | 923100.06 | 748197.25 | 418286.31 | 737084.56 | 564765.06 | 360489.62 | 437070.12 | 1160868.5 | 980921.44 | 929795.81 | 364012.09 | 692128.44 | 0 | 6 | 0 | 8 | 0 | 8 | 0.48 | 0.50 | 0.79 | 0.82 |
| Q62086 | Pon2 | Serum paraoxonase/arylesterase 2 | 77.35 | 4.52 | 39617 | 2 | 712906.88 | 2 | 734898.06 | 508484.38 | 591521.44 | 532611.12 | 645105.56 | 665099.06 | 750807.62 | 926493.94 | 3027654.25 | 2254242.5 | 3370617.25 | 3372546.5 | 3097976.75 | 845998.94 | 111451.83 | 791788.44 | 2 | 2 | 0 | 12 | 0 | 8 | 0 | 12 | 5.94 | 4.30 | 0.69 | 0.50 |
| P37040 | Por | NADPH--cytochrome P450 reductase | 912.4 | 32.6 | 77044 | 17 | 9732950 | 13376572 | 8151045.5 | 8627471 | 9473038 | 11539862 | 10462630 | 12195338 | 10641674 | 12806978 | 43183954 | 38509494 | 40126274 | 29400900 | 36408986 | 11534504 | 12671956 | 17554934 | 16991208 | 17745150 | 0 | 2 | 0 | 4 | 0 | 4 | 3.80 | 3.25 | 1.55 | 1.33 |
| P25425 | Pou2f1 | POU domain, class 2, transcription factor 1 | 171.44 | 4.68 | 79547 | 3 | 1145465.75 | 1638671.75 | 2441183 | 3935758.75 | 2431494 | 2214825.75 | 1242814.25 | 3298106.5 | 2313416 | 1627814.12 | 922307.38 | 497965.03 | 551714.69 | 437041.09 | 458682.25 | 1866592.62 | 872508.12 | 2491786.5 | 1403506.75 | 1860725.25 | 0 | 9 | 0 | 9 | 0 | 9 | 0.25 | 0.27 | 0.73 | 0.79 |
| Q91YU8 | Ppan | Suppressor of SWI4 1 homolog | 453.3 | 17.23 | 52756 | 8 | 341523.56 | 1759523.25 | 2095264 | 2150489 | 560643.12 | 1759530 | 1951098.5 | 1071427.75 | 2176006.75 | 1417728.62 | 5327990.5 | 4297904.5 | 4366677 | 4073607.5 | 3498687 | 1282525 | 2440819.5 | 1888576.62 | 1762428 | 2527584.5 | 0 | 7 | 0 | 8 | 0 | 8 | 3.12 | 2.57 | 1.43 | 1.18 |
| P63087 | Ppp1cc | Serine/threonine-protein phosphatase PP1-gamma catalytic subunit | 1068.97 | 50.46 | 36984 | 16 | 5326580 | 4732472.5 | 5007804.5 | 5784666.5 | 4578759 | 4957448.5 | 4294018 | 4476670 | 4697473 | 4480961.5 | 4004312.25 | 3487311.75 | 3894091.5 | 3707992.5 | 3830465.5 | 4470837.5 | 5756461.5 | 4873448.5 | 5241667.5 | 5279371 | 0 | 12 | 0 | 5 | 0 | 12 | 0.74 | 0.83 | 1.01 | 1.12 |
| P63328 | Ppp3ca | Serine/threonine-protein phosphatase 2B catalytic subunit alpha isoform | 494.34 | 19.58 | 58644 | 9 | 10145184 | 10125261 | 14584545 | 12828923 | 15963079 | 9182245 | 9899852 | 13537429 | 12571545 | 12558725 | 4451651 | 5725251.5 | 5823734.5 | 6028640.5 | 5843319 | 11747729 | 8290227.5 | 10599961 | 10274011 | 13018799 | 0 | 9 | 0 | 10 | 0 | 10 | 0.44 | 0.48 | 0.85 | 0.93 |
| Q7TSI3 | Ppp6r1 | Serine/threonine-protein phosphatase 6 regulatory subunit 1 | 43.76 | 1.29 | 94527 | 1 | 1446421.38 | 1622224.88 | 1234334.62 | 1847331.25 | 1625511.5 | 1591780.62 | 1785651.62 | 1449110.62 | 1269267.25 | 1498151.5 | 739432.94 | 735131.44 | 811089.5 | 955392.75 | 832180.88 | 1706759 | 2081608.12 | 1625487.88 | 1573111.75 | 1881065.5 | 0 | 6 | 0 | 4 | 0 | 6 | 0.52 | 0.54 | 1.14 | 1.17 |
| O88531 | Ppt1 | Palmitoyl-protein thioesterase 1 | 342.59 | 23.53 | 34490 | 6 | 8257738 | 6877490.5 | 8660177 | 15343746 | 14606335 | 8156881 | 7772314 | 9315938 | 10480905 | 10223054 | 3202669.75 | 1628197 | 1424178.75 | 1311020.75 | 1351921.25 | 9633528 | 12755555 | 9233552 | 10548312 | 9008489 | 0 | 10 | 0 | 8 | 0 | 10 | 0.17 | 0.19 | 0.95 | 1.11 |
| Q61171 | Prdx2 | Peroxiredoxin-2 | 705.1 | 45.45 | 21779 | 10 | 173070546 | 149379778 | 137551250 | 149690610 | 127327018 | 178182018 | 132331386 | 120464218 | 120588674 | 141047858 | 87309658 | 78720674 | 78481938 | 77933194 | 69200850 | 159986754 | 193406914 | 147484274 | 145347842 | 133668658 | 0 | 11 | 0 | 5 | 0 | 11 | 0.53 | 0.57 | 1.06 | 1.13 |
| Q69ZK0 | Prex1 | Phosphatidylinositol 3,4,5-trisphosphate-dependent Rac exchanger 1 protein | 1024.21 | 13.33 | 184935 | 19 | 12587764 | 12240188 | 18171052 | 19168848 | 22936888 | 13095770 | 11099807 | 16270355 | 20796302 | 17229232 | 7371416.5 | 7588642.5 | 7955791.5 | 6147153.5 | 7983837 | 11453774 | 10371119 | 13508807 | 15818183 | 16322212 | 0 | 6 | 0 | 8 | 0 | 8 | 0.44 | 0.47 | 0.79 | 0.86 |
| P68181 | Prkacb | cAMP-dependent protein kinase catalytic subunit beta | 136.21 | 8.55 | 40708 | 3 | 402108.97 | 546788.88 | 636712 | 855679.88 | 625964.19 | 513231.5 | 332859.84 | 660096 | 1311903.5 | 881299.19 | 2073610.62 | 1525540.88 | 2234799.75 | 2453665.5 | 2143883 | 2 | 324822.72 | 1078763.38 | 780548.12 | 561861.69 | 0 | 11 | 0 | 9 | 0 | 11 | 3.40 | 2.82 | 0.90 | 0.74 |
| Q9DBC7 | Prkar1a | cAMP-dependent protein kinase type I-alpha regulatory subunit | 477.98 | 25.46 | 43185 | 8 | 2710606.25 | 2956105.5 | 5157968.5 | 3313074.25 | 4516981.5 | 1407185.25 | 2769253 | 4050832.5 | 3698436.5 | 2919209 | 5640021 | 8269927 | 7735366.5 | 5483762 | 8858641 | 4535994 | 2990896 | 4158960.25 | 5566362 | 4679008 | 0 | 7 | 0 | 2 | 0 | 7 | 1.93 | 2.42 | 1.18 | 1.48 |
| P23298 | Prkch | Protein kinase C eta type | 67.3 | 1.46 | 77919 | 1 | 179823.39 | 159095.2 | 172126.92 | 292217.19 | 278111.72 | 180224.48 | 192252.2 | 142421.42 | 230204.97 | 177740.75 | 2 | 2 | 63347.53 | 2 | 2 | 2 | 2 | 2 | 2 | 144311.05 | 0 | 0 | 0 | 1 | 0 | 1 | 0.06 | 0.07 | 0.13 | 0.16 |
| Q64695 | Procr | Endothelial protein C receptor | 115.52 | 12.4 | 27189 | 2 | 233846.89 | 270578.75 | 705526.38 | 116987.84 | 704017.38 | 158755.39 | 389496.19 | 1022577.75 | 949646.62 | 1000441.5 | 1579242.5 | 1841332.25 | 1306309.5 | 1487158 | 1677352.25 | 1171111.75 | 818465.31 | 1078614.62 | 1180580.38 | 1255977.12 | 0 | 0 | 0 | 3 | 0 | 3 | 3.89 | 2.24 | 2.71 | 1.56 |
| P09330 | Prps2 | Ribose-phosphate pyrophosphokinase 2 | 514.2 | 25.79 | 34813 | 9 | 1523141.75 | 1392618.75 | 1703070.62 | 1190029.12 | 910314.62 | 1478963.25 | 1330535.25 | 1191226.5 | 862879 | 1116321 | 731215.94 | 2 | 2 | 2 | 819162 | 1521057.12 | 973621.69 | 1188639.12 | 1081408.38 | 741813.56 | 0 | 6 | 0 | 9 | 0 | 9 | 0.23 | 0.26 | 0.82 | 0.92 |
| Q61207 | Psap | Prosaposin | 587.79 | 17.06 | 61422 | 9 | 47805758 | 34913794 | 39723790 | 43951550 | 39073166 | 37368754 | 35162606 | 36529214 | 31290032 | 42193594 | 8348112 | 19671462 | 19434842 | 17199734 | 21339280 | 49969498 | 41935718 | 38279466 | 46950258 | 41313006 | 0 | 9 | 0 | 3 | 0 | 9 | 0.42 | 0.47 | 1.06 | 1.20 |
| Q99K85 | Psat1 | Phosphoserine aminotransferase | 1227.63 | 57.3 | 40473 | 20 | 52068278 | 46781450 | 40696466 | 37414430 | 36672946 | 51666058 | 56660414 | 45210554 | 44934514 | 44688190 | 94959266 | 87231602 | 83649170 | 65489202 | 74872690 | 59107458 | 59837818 | 52279634 | 52299282 | 48358058 | 0 | 1 | 0 | 4 | 0 | 4 | 1.90 | 1.67 | 1.27 | 1.12 |
| O70435 | Psma3 | Proteasome subunit alpha type-3 | 621.82 | 42.35 | 28405 | 10 | 55527578 | 48947946 | 53756230 | 59217982 | 56685734 | 57383914 | 48925678 | 47753194 | 54238354 | 50825762 | 25722154 | 29998250 | 28002414 | 30842340 | 28809934 | 63156806 | 71585738 | 49384850 | 61910538 | 62176678 | 0 | 4 | 0 | 3 | 0 | 4 | 0.52 | 0.55 | 1.12 | 1.19 |
| P21670 | Psma4 | Proteasome subunit alpha type-4 | 532.88 | 42.15 | 29498 | 11 | 34637294 | 39758130 | 39268918 | 37490934 | 43698718 | 35293918 | 36303214 | 40721954 | 43006610 | 35717946 | 25081612 | 28813530 | 24741352 | 25480368 | 30297114 | 40128098 | 38799038 | 38497346 | 38588522 | 41773586 | 0 | 11 | 0 | 9 | 0 | 11 | 0.69 | 0.70 | 1.02 | 1.04 |
| Q9Z2U1 | Psma5 | Proteasome subunit alpha type-5 | 473.69 | 35.27 | 26411 | 7 | 42563850 | 43813890 | 42247862 | 40708374 | 40158350 | 34532306 | 45288506 | 43918946 | 47255726 | 41435626 | 23457532 | 32946344 | 26612800 | 22477588 | 29353862 | 47086058 | 40044934 | 37144546 | 38535022 | 46169234 | 0 | 10 | 0 | 11 | 0 | 11 | 0.64 | 0.63 | 1.00 | 0.98 |
| Q9QUM9 | Psma6 | Proteasome subunit alpha type-6 | 569.7 | 43.9 | 27372 | 9 | 38794286 | 44936838 | 36369986 | 39271418 | 33541046 | 48469886 | 40517038 | 38238906 | 34563398 | 39577570 | 30901338 | 29292222 | 26580762 | 32847798 | 27754360 | 44003914 | 54353530 | 45519670 | 48778306 | 43975778 | 0 | 2 | 0 | 3 | 0 | 3 | 0.76 | 0.73 | 1.23 | 1.18 |
| O09061 | Psmb1 | Proteasome subunit beta type-1 | 686.8 | 42.5 | 26372 | 10 | 30704826 | 32197196 | 30691250 | 30795814 | 32116516 | 31011662 | 33044142 | 30466014 | 30323474 | 32932442 | 19310314 | 25698122 | 21726730 | 17989260 | 22893718 | 33663106 | 30663978 | 34819570 | 35796082 | 34771150 | 0 | 5 | 0 | 3 | 0 | 5 | 0.69 | 0.68 | 1.08 | 1.08 |
| Q9R1P1 | Psmb3 | Proteasome subunit beta type-3 | 516.89 | 46.34 | 22965 | 7 | 27129268 | 24944502 | 25328920 | 25717506 | 24780906 | 24789682 | 22705706 | 22623968 | 25809868 | 23534084 | 16821848 | 14959572 | 16350118 | 14727893 | 15896821 | 24703178 | 23940184 | 26651106 | 26542308 | 26521974 | 0 | 12 | 0 | 4 | 0 | 12 | 0.62 | 0.66 | 1.00 | 1.07 |
| P28063 | Psmb8 | Proteasome subunit beta type-8 | 541.64 | 35.51 | 30260 | 8 | 27969680 | 26381824 | 30800556 | 37793706 | 31573004 | 25829130 | 23334344 | 31318560 | 29709702 | 33224496 | 18300172 | 19963792 | 18408878 | 21407786 | 18903370 | 30420206 | 37176170 | 32637282 | 40535330 | 39873350 | 0 | 5 | 0 | 3 | 0 | 5 | 0.63 | 0.68 | 1.17 | 1.26 |
| O35522 | Psmb9 | Proteasome subunit beta type-9 bactrianus (Southwestern Asian house mouse) | 150.38 | 16.89 | 23427 | 3 | 16448648 | 15556475 | 14568322 | 21421180 | 15825280 | 15741578 | 15080285 | 15964824 | 15358306 | 18860432 | 7486548 | 8274746 | 8604166 | 8435180 | 8388597 | 16359024 | 19813396 | 16680886 | 16943994 | 17107386 | 0 | 6 | 0 | 4 | 0 | 6 | 0.49 | 0.51 | 1.04 | 1.07 |
| P62192 | Psmc1 | 26S proteasome regulatory subunit 4 | 879.95 | 37.27 | 49185 | 15 | 22715296 | 21359824 | 22409186 | 24681490 | 20494524 | 23904400 | 17986380 | 24827388 | 20429402 | 25615522 | 29614318 | 27341712 | 26399240 | 27098434 | 29712988 | 27664566 | 30281016 | 29915560 | 33298300 | 30705396 | 0 | 0 | 0 | 0 | 0 | 0 | 1.26 | 1.24 | 1.36 | 1.35 |
| P46471 | Psmc2 | 26S proteasome regulatory subunit 7 | 1266.46 | 47.34 | 48648 | 19 | 22361660 | 24271934 | 24121212 | 25824980 | 18893120 | 26245738 | 23074908 | 22529782 | 23846304 | 24151868 | 30083154 | 30943400 | 28654206 | 26913780 | 28784956 | 21279928 | 27419438 | 25741208 | 24392818 | 23348070 | 0 | 9 | 0 | 10 | 0 | 10 | 1.26 | 1.21 | 1.06 | 1.02 |
| P62196 | Psmc5 | 26S proteasome regulatory subunit 8 | 812.52 | 32.76 | 45626 | 12 | 13112931 | 15307546 | 11222798 | 17231260 | 15514786 | 14393889 | 13872406 | 13183716 | 13753204 | 13698878 | 25314816 | 24029862 | 23996252 | 18986224 | 21943556 | 14151786 | 17091702 | 17790078 | 17508130 | 14550083 | 0 | 7 | 0 | 1 | 0 | 7 | 1.58 | 1.66 | 1.12 | 1.18 |
| Q8BG32 | Psmd11 | 26S proteasome non-ATPase regulatory subunit 11 | 1178.5 | 43.36 | 47437 | 19 | 43115542 | 45971598 | 49639946 | 42768986 | 47344018 | 39988054 | 43694806 | 49444098 | 49294858 | 43117618 | 50376494 | 64643422 | 59851398 | 49765850 | 56213430 | 46826298 | 39495294 | 51290866 | 44728070 | 51810014 | 0 | 10 | 0 | 9 | 0 | 10 | 1.23 | 1.25 | 1.02 | 1.04 |
| O35593 | Psmd14 | 26S proteasome non-ATPase regulatory subunit 14 | 395.13 | 38.06 | 34577 | 6 | 8660083 | 9181106 | 7990924 | 10538366 | 8660145 | 8050112.5 | 9277148 | 8632354 | 7554669.5 | 8781094 | 12913041 | 11234559 | 11654591 | 13505668 | 11727562 | 9221145 | 10776824 | 10833144 | 11693914 | 9840708 | 0 | 2 | 0 | 1 | 0 | 2 | 1.36 | 1.44 | 1.16 | 1.24 |
| Q9CX56 | Psmd8 | 26S proteasome non-ATPase regulatory subunit 8 | 335.79 | 17.85 | 39930 | 7 | 17946492 | 13347273 | 14431803 | 13214330 | 19030170 | 11617676 | 17735372 | 16679696 | 19196130 | 15577289 | 5276102 | 6258011 | 10147525 | 3735441 | 7022559.5 | 18271310 | 11114215 | 15466667 | 22895178 | 15523344 | 0 | 10 | 0 | 11 | 0 | 11 | 0.42 | 0.40 | 1.07 | 1.03 |
| P97371 | Psme1 | Proteasome activator complex subunit 1 | 716.2 | 37.35 | 28673 | 11 | 50554214 | 46743382 | 47803094 | 52192054 | 61949566 | 35859638 | 49624254 | 48466754 | 55925502 | 42779722 | 7912880 | 16947016 | 17394870 | 9961257 | 18667182 | 51871158 | 24889188 | 35946666 | 38442590 | 43944934 | 0 | 3 | 0 | 8 | 0 | 8 | 0.27 | 0.30 | 0.75 | 0.84 |
| P97372 | Psme2 | Proteasome activator complex subunit 2 | 760.54 | 39.33 | 27057 | 9 | 55244570 | 59556870 | 59887874 | 70152786 | 80794386 | 50051582 | 58843426 | 68389258 | 71268506 | 60632126 | 24003782 | 31503716 | 30136716 | 21603628 | 31579852 | 65642118 | 50552778 | 55514486 | 59057146 | 61408438 | 0 | 8 | 0 | 10 | 0 | 10 | 0.43 | 0.45 | 0.90 | 0.94 |
| Q9EST4 | Psmg2 | Proteasome assembly chaperone 2 | 90.96 | 6.06 | 29524 | 2 | 1919413.62 | 2000728.12 | 1987396.62 | 2330912.25 | 2446069.75 | 1923297.12 | 2085505.25 | 2237558.5 | 2273858 | 1753810 | 945688.56 | 1382125.88 | 1630402.88 | 732623.88 | 1213266.25 | 2364676 | 1031825.62 | 1483787.62 | 1451543.12 | 1967789.25 | 0 | 5 | 0 | 7 | 0 | 7 | 0.55 | 0.57 | 0.78 | 0.81 |
| Q05769 | Ptgs2 | Prostaglandin G/H synthase 2 | 972.47 | 34.6 | 69013 | 16 | 3143319 | 2768988.5 | 2693109.5 | 2463459 | 2382702 | 2378090.75 | 2631982.5 | 4180545 | 4386155.5 | 4023008.5 | 10844819 | 12129170 | 17741340 | 17629320 | 19329358 | 1970288.5 | 2564236.25 | 4586106 | 3715207.5 | 3373953 | 0 | 8 | 0 | 10 | 0 | 10 | 5.77 | 4.41 | 1.21 | 0.92 |
| P26350 | Ptma | Prothymosin alpha | 690.84 | 38.74 | 12254 | 7 | 69578482 | 83228450 | 82588546 | 98298322 | 82713546 | 74604402 | 85774898 | 62579010 | 69117098 | 76325722 | 37938678 | 49746886 | 34317674 | 43055990 | 33806214 | 73485778 | 84848610 | 62528190 | 61495162 | 75114082 | 0 | 6 | 0 | 9 | 0 | 9 | 0.48 | 0.54 | 0.86 | 0.97 |
| P04550 | Ptms | Parathymosin | 222.78 | 22.55 | 11559 | 3 | 40839666 | 36883378 | 29901414 | 56078398 | 31743398 | 41980234 | 29979250 | 41436078 | 29411958 | 37609434 | 12787156 | 11005493 | 10181458 | 11568436 | 10122685 | 45860466 | 53404394 | 28774614 | 29299780 | 39408214 | 0 | 11 | 0 | 12 | 0 | 12 | 0.28 | 0.31 | 1.01 | 1.09 |
| O88902 | Ptpn23 | Tyrosine-protein phosphatase non-receptor type 23 | 136.54 | 3 | 163454 | 3 | 654283.19 | 1369009 | 600017.81 | 943320.38 | 618157.12 | 1083491.25 | 1560118.62 | 789826.81 | 1020125.5 | 967767.62 | 2590779.5 | 1828863.88 | 2033796.75 | 2137492.25 | 2953757.5 | 1377971.5 | 2282630.75 | 1130235.5 | 1083242.25 | 1381260.75 | 0 | 2 | 0 | 5 | 0 | 5 | 2.76 | 2.13 | 1.73 | 1.34 |
| Q8BKS9 | Pum3 | Pumilio homolog 3 | 94 | 4.64 | 72800 | 2 | 580776.75 | 662025.69 | 641102.56 | 940811.31 | 885422.88 | 685543.06 | 604456.81 | 667301.75 | 665940 | 449364.19 | 2 | 403623.62 | 361934.12 | 376858.03 | 331562.38 | 743141.81 | 843267.88 | 691726.94 | 694792.12 | 663709.31 | 0 | 10 | 0 | 3 | 0 | 10 | 0.40 | 0.48 | 0.98 | 1.18 |
| Q8BU03 | Pwp2 | Periodic tryptophan protein 2 homolog | 127.92 | 3.26 | 102910 | 3 | 699478.69 | 1888993.12 | 474707.72 | 927675 | 289635.34 | 481527.59 | 846134.06 | 1444760.12 | 1018405.62 | 859638.38 | 3445445 | 2205845.5 | 2232691.5 | 2107989 | 3077145 | 902355.56 | 1556375.62 | 859852.62 | 375902.41 | 801214.94 | 0 | 11 | 0 | 12 | 0 | 12 | 3.05 | 2.81 | 1.05 | 0.97 |
| Q66H76 | Pxn | Paxillin | 120.77 | 3.92 | 64019 | 3 | 2518412 | 3231852.75 | 3516242.25 | 3604868 | 2590649 | 2614897 | 2535192.25 | 4073718 | 2117609.5 | 2276496 | 1765667.5 | 1769480.5 | 1596168.5 | 1999467.5 | 1988295.5 | 3093190.75 | 3660646.5 | 3337575.5 | 4356351 | 3254593 | 0 | 7 | 0 | 4 | 0 | 7 | 0.59 | 0.67 | 1.14 | 1.30 |
| Q9EPB4 | Pycard | Apoptosis-associated speck-like protein containing a CARD | 328.2 | 43.01 | 21459 | 6 | 8257800 | 8798768 | 10893768 | 8100220 | 10649589 | 5771371.5 | 7889947 | 8020748 | 10379690 | 8949953 | 2545522.25 | 4555039 | 4273481.5 | 3007083 | 4744737 | 7015427 | 3755338.5 | 6547450 | 4273185 | 7990366 | 0 | 0 | 0 | 4 | 0 | 4 | 0.41 | 0.47 | 0.63 | 0.72 |
| P53534 | Pygb | Glycogen phosphorylase, brain form | 871.84 | 19.09 | 96174 | 15 | 18244850 | 17799186 | 18200130 | 16737296 | 23066866 | 13948239 | 17028260 | 22343636 | 22001712 | 16215926 | 7046817.5 | 8231821 | 8822479 | 5368501 | 8261356 | 16399435 | 11798240 | 11463429 | 10741095 | 10378699 | 0 | 0 | 0 | 2 | 0 | 2 | 0.40 | 0.41 | 0.65 | 0.66 |
| Q8BVI4 | Qdpr | Dihydropteridine reductase | 281 | 21.58 | 25570 | 4 | 12435898 | 14598934 | 13728056 | 12619882 | 11702171 | 13734372 | 13142331 | 15176777 | 14078325 | 14070203 | 5986159 | 8877218 | 8905944 | 9490798 | 8574603 | 13635406 | 10599172 | 16800260 | 14480734 | 15114327 | 0 | 8 | 0 | 11 | 0 | 11 | 0.64 | 0.60 | 1.09 | 1.01 |
| Q921E2 | Rab31 | Ras-related protein Rab-31 | 229.49 | 19.07 | 21333 | 3 | 3397679.5 | 3834778 | 3536063.5 | 4649832 | 4444728.5 | 3590516.5 | 3465393.75 | 3326834.5 | 3685449 | 4558727 | 2153244 | 1645682.62 | 2911668.75 | 2702058.25 | 2978499.25 | 3561139 | 4083012 | 4079562 | 4295465.5 | 4601897 | 0 | 10 | 0 | 6 | 0 | 10 | 0.62 | 0.67 | 1.04 | 1.11 |
| Q9CQD1 | Rab5a | Ras-related protein Rab-5A | 543.41 | 59.07 | 23599 | 9 | 5185264.5 | 4166141.5 | 3922643 | 5023185 | 3739090.75 | 4991647.5 | 3796136.75 | 2842799.5 | 3787580.5 | 5418166 | 7975968 | 6574168 | 6063421.5 | 6814707 | 7925080.5 | 3464983.75 | 6135598 | 5481009.5 | 4676521.5 | 5150700.5 | 0 | 8 | 0 | 7 | 0 | 8 | 1.60 | 1.70 | 1.13 | 1.20 |
| P61028 | Rab8b | Ras-related protein Rab-8B | 1037.28 | 66.67 | 23603 | 17 | 21683084 | 23391298 | 16911848 | 24809490 | 23378730 | 22275992 | 20404370 | 23634962 | 24434238 | 24865848 | 7470786 | 12518126 | 16382096 | 8425282 | 15693077 | 23650304 | 24370948 | 22649418 | 25149214 | 25766090 | 0 | 5 | 0 | 7 | 0 | 7 | 0.55 | 0.52 | 1.10 | 1.05 |
| Q61550 | Rad21 | Double-strand-break repair protein rad21 homolog | 134.17 | 3.31 | 72083 | 2 | 961052.38 | 752739.94 | 1009241.69 | 1341840 | 836929.19 | 1410942.75 | 1269813.88 | 1386463.75 | 1052901.38 | 837816.06 | 223998.83 | 269190.53 | 214231.56 | 102923.28 | 150253.69 | 1244446.12 | 2158829.5 | 1088005.62 | 1444630.38 | 749445.25 | 0 | 7 | 0 | 11 | 0 | 11 | 0.20 | 0.16 | 1.36 | 1.12 |
| P54726 | Rad23a | UV excision repair protein RAD23 homolog A | 95.14 | 8.26 | 39706 | 1 | 336630.66 | 198972.77 | 280384.28 | 267828.47 | 499901.44 | 243656.89 | 233451.12 | 221963.41 | 206715.16 | 169940.16 | 2 | 2 | 2 | 2 | 92826.03 | 2 | 258244.94 | 216810.12 | 524221.03 | 2 | 0 | 7 | 0 | 12 | 0 | 12 | 0.06 | 0.09 | 0.63 | 0.93 |
| P34022 | Ranbp1 | Ran-specific GTPase-activating protein | 305.31 | 20.69 | 23596 | 5 | 2101473.75 | 7794823.5 | 7079320 | 7149089 | 4844706.5 | 7305336 | 4976903 | 7877269 | 6209366.5 | 10820165 | 15888141 | 16604723 | 14984134 | 16294051 | 12675964 | 6431241.5 | 6141592.5 | 8219074 | 8593937 | 8272734 | 0 | 6 | 0 | 10 | 0 | 10 | 2.64 | 2.06 | 1.30 | 1.01 |
| Q60790 | Rasa3 | Ras GTPase-activating protein 3 | 340.98 | 7.19 | 95987 | 7 | 5219907 | 4639854.5 | 6111745.5 | 5446853.5 | 5240098.5 | 4133152.25 | 4502272 | 4666450 | 4321257.5 | 4103102.5 | 2680542.5 | 2886384.5 | 3532714.25 | 2523634.75 | 2833600.25 | 4680458.5 | 2771065.5 | 3260402.25 | 2530082.75 | 3775586 | 0 | 1 | 0 | 5 | 0 | 5 | 0.54 | 0.67 | 0.64 | 0.78 |
| Q8C2K5 | Rasal3 | RAS protein activator like-3 | 130.31 | 3.07 | 114782 | 3 | 4657640 | 4276998.5 | 5056808 | 5588317.5 | 4006845.25 | 4773707 | 4464023 | 4253322.5 | 4035253.5 | 5228095.5 | 2635268 | 2536821.5 | 3666655.5 | 2539957.5 | 3538142.5 | 4277430 | 4619481.5 | 3123532.5 | 4323948 | 4819902 | 0 | 9 | 0 | 11 | 0 | 11 | 0.63 | 0.66 | 0.90 | 0.93 |
| Q8C2Q3 | Rbm14 | RNA-binding protein 14 | 549.84 | 13.9 | 69449 | 9 | 5088647 | 4102328.5 | 4774362.5 | 4570536.5 | 3599181.5 | 4149221.5 | 4528677 | 3247756 | 4294441.5 | 5959583 | 13543489 | 11296799 | 10492866 | 13063353 | 12803233 | 4456884 | 6128627 | 6509943 | 5831572.5 | 4187065 | 0 | 6 | 0 | 6 | 0 | 6 | 2.76 | 2.76 | 1.22 | 1.22 |
| Q8BHS3 | Rbm22 | Pre-mRNA-splicing factor RBM22 | 53.59 | 1.67 | 46896 | 1 | 2 | 562019.12 | 330884.5 | 2 | 2 | 2 | 2 | 2 | 2 | 2 | 1408072.75 | 1103865.5 | 1157915.38 | 1343928.88 | 1474358.62 | 2 | 2 | 228606.14 | 2 | 2 | 0 | 9 | 0 | 10 | 0 | 10 | 7.27 | 648814.11 | 0.26 | 22861.41 |
| P84586 | Rbmxrtl | RNA-binding motif protein, X chromosome retrogene-like | 675.67 | 33.51 | 42204 | 14 | 3365446.5 | 5001976.5 | 3758197.5 | 5190724 | 4033865.25 | 4857600.5 | 5310631 | 4950059.5 | 5070440 | 5661981 | 13099638 | 12252538 | 13273482 | 13410254 | 17635272 | 5411076.5 | 3490867.75 | 5734795 | 4851748 | 4652391 | 0 | 8 | 0 | 9 | 0 | 9 | 3.26 | 2.70 | 1.13 | 0.93 |
| P31266 | Rbpj | Recombining binding protein suppressor of hairless | 394.17 | 16.54 | 58537 | 7 | 7605564.5 | 4736370 | 7434527 | 6332384.5 | 6457803 | 8492469 | 7414432 | 6681940.5 | 5057985 | 5272308 | 14118880 | 10700679 | 8588044 | 9170090 | 8898610 | 5249852.5 | 7615695 | 4022623.5 | 3454719.5 | 3793122 | 0 | 6 | 0 | 5 | 0 | 6 | 1.58 | 1.56 | 0.74 | 0.73 |
| Q9QYF1 | Rdh11 | Retinol dehydrogenase 11 | 470.53 | 26.9 | 35148 | 7 | 17384592 | 19125722 | 15907852 | 19009422 | 15508748 | 19923836 | 17336732 | 17727470 | 14947081 | 15625086 | 22761130 | 22865766 | 29008802 | 28993562 | 26978962 | 18373678 | 20837184 | 19972518 | 21413542 | 21018688 | 0 | 2 | 0 | 1 | 0 | 2 | 1.50 | 1.53 | 1.17 | 1.19 |
| Q64FW2 | Retsat | All-trans-retinol 13,14-reductase | 196.13 | 8.21 | 67334 | 3 | 1610648.25 | 925256.5 | 911101.06 | 1092285.75 | 1156480.12 | 811141.81 | 1155866.5 | 830872.88 | 957370.62 | 617292.62 | 1969051.5 | 1857133.25 | 1658812.12 | 1611660.25 | 1776039 | 1460272.5 | 1337133.75 | 961630.38 | 1033971.88 | 917851.88 | 0 | 12 | 0 | 4 | 0 | 12 | 1.56 | 2.03 | 1.00 | 1.31 |
| Q80WQ6 | Rhbdf2 | Inactive rhomboid protein 2 | 86.25 | 2.06 | 93434 | 2 | 895977.88 | 919255.88 | 1267540.88 | 1212610.38 | 940070.5 | 156853.86 | 791225.75 | 898898.25 | 792611.06 | 969389.06 | 1652333.12 | 1739037 | 1952946.12 | 2113357.25 | 2034659.12 | 988986.94 | 1066861 | 1351861.12 | 1357105.62 | 2 | 0 | 9 | 0 | 5 | 0 | 9 | 1.81 | 2.63 | 0.91 | 1.32 |
| Q921J2 | Rheb | GTP-binding protein Rheb | 189.45 | 24.46 | 20451 | 5 | 3899164.75 | 3763344 | 3129479.5 | 4462906 | 3290964 | 3681183 | 3731764 | 3813813.5 | 4075623.75 | 4524380.5 | 5425113.5 | 5515014.5 | 7112128.5 | 6554250 | 6154244 | 3122322 | 3344105 | 4334649 | 4473933 | 4910975 | 0 | 9 | 0 | 12 | 0 | 12 | 1.66 | 1.55 | 1.09 | 1.02 |
| Q3U2C5 | Rnf149 | E3 ubiquitin-protein ligase RNF149 | 415.94 | 24.11 | 42553 | 7 | 1440000.75 | 1070581.88 | 447641.69 | 319087.47 | 409728.66 | 1192406.88 | 896527.88 | 615830.19 | 468418.28 | 460726.19 | 7997166.5 | 7602215.5 | 5897767.5 | 4531718 | 5693195.5 | 1093578.62 | 1689244.75 | 1025040.88 | 576208.75 | 1380439.12 | 0 | 6 | 0 | 5 | 0 | 6 | 8.60 | 8.73 | 1.56 | 1.59 |
| Q9CQ71 | Rpa3 | Replication protein A 14 kDa subunit | 47.09 | 8.26 | 13584 | 1 | 1079400.12 | 1024493.25 | 1353243.75 | 1207658.25 | 875332.94 | 919185.75 | 906045.12 | 797955.44 | 838085.75 | 940839.31 | 2 | 646331.31 | 647940.69 | 686944.12 | 616709.5 | 1161057 | 995168.88 | 826258.81 | 857253.75 | 906895.75 | 0 | 5 | 0 | 9 | 0 | 9 | 0.47 | 0.59 | 0.86 | 1.08 |
| Q6PDV7 | Rpl10 | 60S ribosomal protein L10 | 440.6 | 35.05 | 24604 | 7 | 10528438 | 19621512 | 15798494 | 14418362 | 11473873 | 15972617 | 13846737 | 18235274 | 16085418 | 17440242 | 28678498 | 35552602 | 27524156 | 28955280 | 28666562 | 14842060 | 13990552 | 20632100 | 17011770 | 18986706 | 0 | 7 | 0 | 10 | 0 | 10 | 2.08 | 1.83 | 1.19 | 1.05 |
| P62914 | Rpl11 | 60S ribosomal protein L11 | 649.34 | 50.56 | 20252 | 10 | 49207562 | 54388898 | 46708286 | 44787654 | 39316834 | 59886058 | 53580078 | 54638482 | 48889946 | 57198098 | 65878654 | 71958298 | 60365138 | 70038082 | 64330466 | 51102662 | 50463122 | 62853050 | 63986090 | 64153026 | 0 | 2 | 0 | 8 | 0 | 8 | 1.42 | 1.21 | 1.25 | 1.07 |
| P61314 | Rpl15 | 60S ribosomal protein L15 | 717.1 | 47.55 | 24146 | 12 | 47592318 | 62604774 | 70160354 | 52638130 | 55058354 | 54288130 | 54766918 | 73084754 | 68448202 | 68353186 | 94986746 | 106756218 | 85142002 | 86707466 | 84388634 | 34729338 | 27784160 | 39717230 | 32489202 | 45706570 | 0 | 0 | 0 | 0 | 0 | 0 | 1.59 | 1.44 | 0.63 | 0.57 |
| P35980 | Rpl18 | 60S ribosomal protein L18 | 658.82 | 45.74 | 21645 | 11 | 66130734 | 79426778 | 66929874 | 58445394 | 44431390 | 92232066 | 63898330 | 66171134 | 65068586 | 90215866 | 217900930 | 188183042 | 173846962 | 171976002 | 153798226 | 49074826 | 45590370 | 90411938 | 81261666 | 86217538 | 0 | 8 | 0 | 10 | 0 | 10 | 2.87 | 2.40 | 1.12 | 0.93 |
| P62717 | Rpl18a | 60S ribosomal protein L18a | 571.54 | 51.14 | 20732 | 12 | 33967998 | 39493154 | 41743158 | 31028280 | 35983934 | 40482562 | 38604798 | 39760434 | 37629782 | 37961578 | 51993170 | 62921022 | 61984386 | 58550138 | 59570154 | 40726466 | 37222378 | 46270250 | 35836242 | 44961682 | 0 | 6 | 0 | 10 | 0 | 10 | 1.62 | 1.52 | 1.13 | 1.05 |
| Q9D7S7 | Rpl22l1 | 60S ribosomal protein L22-like 1 | 334.03 | 50.82 | 14467 | 5 | 7764551.5 | 7720018.5 | 13056604 | 12927331 | 12154583 | 6347811 | 6761531.5 | 12465460 | 12551930 | 14294729 | 3383156.5 | 2665428.75 | 4057711 | 3555043 | 3602887 | 5413088.5 | 6767973.5 | 7582463.5 | 7505820 | 7097291.5 | 0 | 0 | 0 | 8 | 0 | 8 | 0.32 | 0.33 | 0.64 | 0.66 |
| P62830 | Rpl23 | 60S ribosomal protein L23 | 474.17 | 67.86 | 14865 | 8 | 60085382 | 64951198 | 59085922 | 66386874 | 50137894 | 72040490 | 56214030 | 53466090 | 47504130 | 62393830 | 120180258 | 84940842 | 80969018 | 91672450 | 92185498 | 63318970 | 71536106 | 55455534 | 60225630 | 63629694 | 0 | 10 | 0 | 8 | 0 | 10 | 1.56 | 1.61 | 1.04 | 1.08 |
| P83732 | Rpl24 | 60S ribosomal protein L24 | 310.24 | 26.75 | 17779 | 5 | 18581420 | 23483404 | 14974894 | 14746763 | 10709080 | 25206022 | 21202960 | 13305952 | 16940472 | 22941020 | 57307642 | 50541802 | 36416354 | 38102042 | 35962134 | 15786142 | 18317374 | 21522118 | 15510011 | 18318230 | 0 | 9 | 0 | 9 | 0 | 9 | 2.65 | 2.19 | 1.08 | 0.90 |
| P61354 | Rpl27 | 60S ribosomal protein L27 | 284.49 | 42.65 | 15798 | 5 | 24660006 | 27295986 | 24421514 | 20859728 | 16697353 | 28249224 | 26376850 | 28144434 | 23617054 | 30782118 | 59127082 | 52115650 | 48279594 | 39976346 | 41816674 | 26166294 | 20042938 | 31124874 | 21480350 | 28751348 | 0 | 8 | 0 | 10 | 0 | 10 | 2.12 | 1.76 | 1.12 | 0.93 |
| P41105 | Rpl28 | 60S ribosomal protein L28 | 379.06 | 40.88 | 15733 | 7 | 38160010 | 48705542 | 39380130 | 31052262 | 22922522 | 47155322 | 46042230 | 45894298 | 30262412 | 44075134 | 56918850 | 66560590 | 54140282 | 65680586 | 56142658 | 37155226 | 48837874 | 56112394 | 47490134 | 46672238 | 0 | 5 | 0 | 5 | 0 | 5 | 1.66 | 1.40 | 1.31 | 1.11 |
| P27659 | Rpl3 | 60S ribosomal protein L3 | 1071.69 | 46.15 | 46110 | 18 | 60903566 | 83046514 | 68413602 | 59743898 | 51943142 | 76359530 | 66186314 | 84800106 | 69915914 | 88865842 | 137650482 | 114641554 | 118726570 | 116060122 | 126282690 | 37496106 | 35695854 | 69466378 | 53174178 | 70514010 | 0 | 9 | 0 | 3 | 0 | 9 | 1.89 | 1.59 | 0.82 | 0.69 |
| P62889 | Rpl30 | 60S ribosomal protein L30 | 461.99 | 50.43 | 12784 | 7 | 22865492 | 23192966 | 18168254 | 14812558 | 17972404 | 30706280 | 22071526 | 18318900 | 17170188 | 20302838 | 50727006 | 49045626 | 43146486 | 45831402 | 41534918 | 18088366 | 24748842 | 32699872 | 33652170 | 32232914 | 0 | 3 | 0 | 5 | 0 | 5 | 2.37 | 2.12 | 1.46 | 1.30 |
| P62900 | Rpl31 | 60S ribosomal protein L31 | 215.49 | 31.2 | 14463 | 4 | 12382484 | 13173400 | 9016778 | 9520594 | 6105968.5 | 17994486 | 16508778 | 12917846 | 8372157.5 | 15218908 | 28241364 | 27672740 | 23389516 | 20331160 | 20381924 | 11065463 | 18983612 | 13329313 | 11186331 | 13017694 | 0 | 5 | 0 | 11 | 0 | 11 | 2.39 | 1.69 | 1.35 | 0.95 |
| P11250 | Rpl34 | 60S ribosomal protein L34 | 200.48 | 21.37 | 13507 | 4 | 9418797 | 19888446 | 16166396 | 9918688 | 9086686 | 16699977 | 13727236 | 26812662 | 19946138 | 23668262 | 40902906 | 33866758 | 53146242 | 60538870 | 58190038 | 9795318 | 7735543 | 29501414 | 21636354 | 26323290 | 0 | 8 | 0 | 12 | 0 | 12 | 3.83 | 2.45 | 1.47 | 0.94 |
| O55142 | Rpl35a | 60S ribosomal protein L35a | 157.15 | 27.27 | 12554 | 4 | 9759211 | 15400194 | 9233290 | 7045611.5 | 7984857.5 | 17627738 | 20876836 | 26175630 | 14490232 | 20712130 | 56736386 | 53604822 | 45209298 | 43600994 | 55554366 | 6846346 | 6555198 | 18914630 | 19017804 | 17726820 | 0 | 10 | 0 | 6 | 0 | 10 | 5.15 | 2.55 | 1.40 | 0.69 |
| P39032 | Rpl36 | 60S ribosomal protein L36 | 114.52 | 19.05 | 12268 | 2 | 1187076.75 | 1351084.12 | 724337.75 | 618222.5 | 259250.17 | 2 | 856009.31 | 2 | 2 | 1133793.62 | 5625721.5 | 3448654.5 | 2651835.5 | 2354726.25 | 2584549.25 | 515020.03 | 545382.06 | 2 | 534504.81 | 395464.47 | 0 | 4 | 0 | 11.5 | 0 | 11.5 | 4.03 | 8.38 | 0.48 | 1.00 |
| P63174 | Rpl38 | 60S ribosomal protein L38 | 210.24 | 48.57 | 8218 | 4 | 224857.86 | 402180.69 | 1136925.12 | 1048542.69 | 1349236.25 | 769805.62 | 709168.5 | 1710797.5 | 1938542.25 | 1553965.5 | 10208006 | 5299055 | 4971177 | 3641907 | 4062600 | 2 | 673339.88 | 1835169.25 | 1255803.25 | 1227221.25 | 0 | 10 | 0 | 8 | 0 | 10 | 6.77 | 4.22 | 1.20 | 0.75 |
| Q9D8E6 | Rpl4 | 60S ribosomal protein L4 | 1404.29 | 45.58 | 47154 | 22 | 119786658 | 137520850 | 138266946 | 122117658 | 119121866 | 149028082 | 131906010 | 143372242 | 136161922 | 119666978 | 179218850 | 190535714 | 175851858 | 158785986 | 171850658 | 129887706 | 113461530 | 145477522 | 137004834 | 159035522 | 0 | 9 | 0 | 12 | 0 | 12 | 1.38 | 1.29 | 1.08 | 1.01 |
| P12970 | Rpl7a | 60S ribosomal protein L7a | 1040.68 | 54.14 | 29977 | 18 | 39124486 | 48596426 | 33825094 | 28209090 | 24239524 | 55066674 | 44661142 | 41411074 | 32895968 | 45595086 | 162444658 | 133649706 | 105265634 | 107677834 | 101414394 | 34670294 | 45752530 | 46871902 | 42848122 | 44195314 | 0 | 6 | 0 | 12 | 0 | 12 | 3.51 | 2.78 | 1.23 | 0.98 |
| P62918 | Rpl8 | 60S ribosomal protein L8 | 518.98 | 36.96 | 28025 | 10 | 104141098 | 133724586 | 110534506 | 105760946 | 83239890 | 143571346 | 116170970 | 143862434 | 113287530 | 143882802 | 160788210 | 155422706 | 146592578 | 172592450 | 153507586 | 96432314 | 99652474 | 165842434 | 151572626 | 173456946 | 0 | 8 | 0 | 10 | 0 | 10 | 1.47 | 1.19 | 1.28 | 1.04 |
| P51410 | Rpl9 | 60S ribosomal protein L9 | 1018.54 | 74.48 | 21881 | 15 | 76408610 | 79425650 | 87899578 | 81198906 | 74264906 | 79156290 | 76102370 | 80313418 | 81164802 | 85569050 | 91404658 | 102349058 | 104989802 | 95383490 | 101243074 | 65567018 | 56496358 | 85309458 | 79399162 | 76813250 | 0 | 8 | 0 | 7 | 0 | 8 | 1.24 | 1.23 | 0.91 | 0.90 |
| P47955 | Rplp1 | 60S acidic ribosomal protein P1 | 222.13 | 66.67 | 11475 | 3 | 21010508 | 17846756 | 18974596 | 18821982 | 33063202 | 16097444 | 24837326 | 20325490 | 27478460 | 17633676 | 10935700 | 16095510 | 9570694 | 11259321 | 10582081 | 28479230 | 38681610 | 18035336 | 7630153 | 22779256 | 0 | 11 | 0 | 10 | 0 | 11 | 0.53 | 0.55 | 1.05 | 1.09 |
| P99027 | Rplp2 | 60S acidic ribosomal protein P2 | 498.87 | 71.3 | 11651 | 6 | 15746718 | 9963271 | 9627175 | 8751212 | 8858731 | 10396751 | 10412105 | 10097364 | 8121642.5 | 10364215 | 5841279 | 7653945.5 | 6029774 | 5125963.5 | 7164217.5 | 12105275 | 14878255 | 10268505 | 10550704 | 10291777 | 0 | 5 | 0 | 6 | 0 | 6 | 0.60 | 0.64 | 1.10 | 1.18 |
| Q91YQ5 | Rpn1 | Dolichyl-diphosphooligosaccharide--protein glycosyltransferase subunit 1 | 1934.48 | 60.2 | 68528 | 32 | 44076746 | 49187770 | 54164002 | 43592598 | 52538506 | 43553838 | 46487062 | 54449558 | 55251610 | 38940658 | 87041738 | 92271426 | 97006218 | 93561594 | 100863514 | 44778466 | 45878306 | 53981466 | 47961970 | 53504362 | 0 | 11 | 0 | 12 | 0 | 12 | 1.93 | 1.97 | 1.01 | 1.03 |
| Q9DBG6 | Rpn2 | Dolichyl-diphosphooligosaccharide--protein glycosyltransferase subunit 2 | 1163.91 | 31.54 | 69063 | 16 | 18775494 | 13843042 | 18711782 | 19092820 | 21979078 | 18053264 | 17055230 | 17980770 | 22922778 | 20778968 | 59039218 | 30686852 | 32196334 | 27173100 | 35940026 | 15949060 | 24794678 | 25914940 | 25330744 | 19842254 | 0 | 5 | 0 | 7 | 0 | 7 | 2.00 | 1.91 | 1.21 | 1.16 |
| Q9CQH8 | Rpp14 | Ribonuclease P protein subunit p14 | 61.88 | 13.11 | 13565 | 1 | 2 | 2 | 2 | 2 | 2 | 2 | 2 | 2 | 2 | 2 | 122617.98 | 157664.42 | 204098.47 | 242716.3 | 131538.2 | 2 | 2 | 2 | 2 | 2 | 0 | 12.5 | 0 | 12.5 | 0 | 12.5 | 85863.54 | 85863.54 | 1.00 | 1.00 |
| P62281 | Rps11 | 40S ribosomal protein S11 | 547.41 | 50.63 | 18431 | 12 | 112600698 | 131488018 | 117845770 | 122627258 | 106653970 | 134574930 | 117821850 | 147051554 | 131069210 | 138592082 | 157729810 | 163061570 | 161982578 | 187301890 | 163599106 | 113914594 | 137601714 | 169374866 | 159010082 | 169764946 | 0 | 3 | 0 | 7 | 0 | 7 | 1.41 | 1.25 | 1.27 | 1.12 |
| P13471 | Rps14 | 40S ribosomal protein S14 | 468.14 | 39.74 | 16259 | 8 | 17645522 | 31860406 | 19319550 | 14823905 | 13597129 | 29420234 | 21776538 | 30690020 | 16782084 | 24653142 | 70370954 | 63243558 | 56363666 | 61370130 | 60286494 | 19024394 | 23726732 | 26668310 | 25383668 | 25606376 | 0 | 6 | 0 | 12 | 0 | 12 | 3.20 | 2.53 | 1.24 | 0.98 |
| P14131 | Rps16 | 40S ribosomal protein S16 | 769.18 | 64.38 | 16445 | 14 | 87129538 | 78053866 | 85512258 | 84388994 | 86635434 | 85250138 | 98477426 | 105833554 | 90691770 | 86924906 | 170714802 | 151698722 | 135426146 | 140187378 | 145977714 | 105846314 | 128481410 | 108636194 | 109695826 | 97306370 | 0 | 0 | 0 | 2 | 0 | 2 | 1.76 | 1.59 | 1.30 | 1.18 |
| P63276 | Rps17 | 40S ribosomal protein S17 | 673.51 | 66.67 | 15524 | 10 | 7666385 | 9606818 | 10011876 | 6619614 | 9509126 | 5866135 | 9619298 | 9125800 | 12875750 | 8991390 | 14263983 | 20198710 | 19690400 | 19744370 | 20699806 | 5482000.5 | 7975365 | 13627371 | 8279384.5 | 15141538 | 0 | 11 | 0 | 12 | 0 | 12 | 2.18 | 2.04 | 1.16 | 1.09 |
| P17074 | Rps19 | 40S ribosomal protein S19 | 638.56 | 54.48 | 16085 | 11 | 38710946 | 39534102 | 36789310 | 34843242 | 26225338 | 38875498 | 35388278 | 36034854 | 31809362 | 39631314 | 60782074 | 57711394 | 49952042 | 47571414 | 50652074 | 36655762 | 40941726 | 45946374 | 38067226 | 43469410 | 0 | 5 | 0 | 4 | 0 | 5 | 1.51 | 1.47 | 1.16 | 1.13 |
| P25444 | Rps2 | 40S ribosomal protein S2 | 960.17 | 52.9 | 31231 | 17 | 132219490 | 146763362 | 111191138 | 121924210 | 104001450 | 148599090 | 134877026 | 134017578 | 119291602 | 137493746 | 222510818 | 228394226 | 184462754 | 228779138 | 210035410 | 143669634 | 135663154 | 155764146 | 121340786 | 162900850 | 0 | 5 | 0 | 7 | 0 | 7 | 1.74 | 1.59 | 1.17 | 1.07 |
| P62849 | Rps24 | 40S ribosomal protein S24 | 317.63 | 35.34 | 15423 | 5 | 7779609 | 10424034 | 4968428 | 3457600.25 | 3098356.25 | 8805184 | 8207040 | 5509281 | 3866838 | 4560566 | 27407248 | 21248128 | 18351492 | 15650499 | 14951270 | 5673178 | 5671896 | 2529750.5 | 3168696 | 2950088.25 | 0 | 7 | 0 | 6 | 0 | 7 | 3.28 | 3.15 | 0.67 | 0.65 |
| P62852 | Rps25 | 40S ribosomal protein S25 | 309.13 | 27.2 | 13742 | 5 | 74740466 | 66851378 | 65518294 | 79753714 | 64779122 | 74375794 | 67572954 | 57744978 | 53960682 | 65548346 | 44985978 | 43677914 | 38681578 | 41912702 | 42932830 | 68210834 | 65849002 | 64611654 | 62019826 | 59876426 | 0 | 5 | 0 | 12 | 0 | 12 | 0.60 | 0.66 | 0.91 | 1.00 |
| P62702 | Rps4x | 40S ribosomal protein S4, X isoform | 1445.9 | 69.58 | 29598 | 25 | 134926738 | 161825138 | 137239554 | 107868186 | 98122490 | 142683410 | 138275330 | 132895514 | 117951090 | 133460978 | 228055442 | 230582226 | 196979362 | 176075506 | 195666578 | 130702506 | 115150122 | 131593218 | 126216538 | 132910570 | 0 | 10 | 0 | 5 | 0 | 10 | 1.61 | 1.54 | 0.99 | 0.96 |
| P62754 | Rps6 | 40S ribosomal protein S6 | 758.24 | 36.95 | 28681 | 10 | 8559045 | 18397554 | 16334593 | 14672886 | 7308984.5 | 13321723 | 11575501 | 16554768 | 9740905 | 14747013 | 61741774 | 44267190 | 25870918 | 42601434 | 25540082 | 9013057 | 19855524 | 17451118 | 13821894 | 12681002 | 0 | 10 | 0 | 10 | 0 | 10 | 3.06 | 3.03 | 1.12 | 1.10 |
| P14206 | Rpsa | 40S ribosomal protein SA | 1004.15 | 55.93 | 32838 | 14 | 39552950 | 39643370 | 40869158 | 34886698 | 30784336 | 34651310 | 31271908 | 33407126 | 30563100 | 45537182 | 166420482 | 117667602 | 89669994 | 88936682 | 99037282 | 34641590 | 42945042 | 43497398 | 37912674 | 36192738 | 0 | 10 | 0 | 6 | 0 | 10 | 3.02 | 3.20 | 1.05 | 1.11 |
| Q99PL5 | Rrbp1 | Ribosome-binding protein 1 | 1944.78 | 20.93 | 172879 | 32 | 84019658 | 87183882 | 61799698 | 68681394 | 62309134 | 92410674 | 90738066 | 66249690 | 61493098 | 64416754 | 57677234 | 59390538 | 37609762 | 41816170 | 43344074 | 105951170 | 107665218 | 73603698 | 82960746 | 78521610 | 0 | 6 | 0 | 6 | 0 | 6 | 0.66 | 0.64 | 1.23 | 1.20 |
| P11157 | Rrm2 | Ribonucleoside-diphosphate reductase subunit M2 | 263.34 | 9.74 | 45096 | 3 | 3181308.5 | 3103195.75 | 2425905.75 | 2758342.5 | 2154076.5 | 3279557.5 | 2895177.5 | 2024701.5 | 1959319 | 2306456.25 | 1798755.25 | 1784488.25 | 854542.75 | 1584438.75 | 567508.19 | 3182356.5 | 3783539 | 2343587 | 2119742 | 1920030.25 | 0 | 11 | 0 | 11 | 0 | 11 | 0.48 | 0.53 | 0.98 | 1.07 |
| Q6P5B0 | Rrp12 | RRP12-like protein | 74.93 | 1.78 | 143131 | 1 | 169517.47 | 2 | 2 | 2 | 124676.45 | 2 | 2 | 2 | 2 | 2 | 302619.78 | 237375.28 | 281383.62 | 412541.53 | 329021.41 | 2 | 2 | 2 | 2 | 2 | 0 | 7.5 | 0 | 12.5 | 0 | 12.5 | 5.31 | 156294.16 | 0.00 | 1.00 |
| Q91WM3 | Rrp9 | U3 small nucleolar RNA-interacting protein 2 | 95.55 | 5.05 | 52107 | 2 | 363805.66 | 473543.38 | 425381.44 | 427391.78 | 384977.81 | 458874.5 | 427053.69 | 2 | 514519.56 | 540498.12 | 542539.5 | 1005943.44 | 591390.12 | 672489.19 | 785815.5 | 469372.88 | 418737.62 | 576220.69 | 2 | 645149.75 | 0 | 9 | 0 | 10.5 | 0 | 10.5 | 1.73 | 1.85 | 1.02 | 1.09 |
| Q9CYH6 | Rrs1 | Ribosome biogenesis regulatory protein homolog | 276.92 | 13.97 | 41552 | 5 | 1131972.5 | 1476569 | 1476071.62 | 879592.5 | 1231039.25 | 1155563 | 1835926.75 | 1079656.75 | 1034957.88 | 898773.06 | 4734407.5 | 3921502.5 | 2105275.5 | 2219920.5 | 3252690.75 | 1139418.5 | 1636476.5 | 1117142.75 | 1253305.62 | 1634018 | 0 | 9 | 0 | 7 | 0 | 9 | 2.62 | 2.70 | 1.09 | 1.13 |
| Q8BVY0 | Rsl1d1 | Ribosomal L1 domain-containing protein 1 | 1072.2 | 30.53 | 50421 | 15 | 12525552 | 17197862 | 14468086 | 17039706 | 7808994.5 | 19277084 | 12216742 | 17503144 | 13973891 | 23560136 | 31268438 | 24233662 | 23961430 | 29965902 | 25035312 | 15650673 | 17715812 | 22835416 | 24177676 | 18588116 | 0 | 2 | 0 | 8 | 0 | 8 | 1.95 | 1.55 | 1.43 | 1.14 |
| Q8BIJ7 | Rufy1 | RUN and FYVE domain-containing protein 1 | 236.32 | 7.58 | 80376 | 5 | 3310324 | 3152040.5 | 3095710.75 | 5668589.5 | 4988930 | 2895603.5 | 4066053.5 | 4339435 | 3715427 | 2969100 | 1132675.5 | 2196545.75 | 1643968.25 | 2146838.5 | 1942653.88 | 2738147 | 2267570.5 | 4409835 | 3501089.5 | 4780345 | 0 | 9 | 0 | 12 | 0 | 12 | 0.45 | 0.50 | 0.88 | 0.98 |
| P60122 | Ruvbl1 | RuvB-like 1 | 891.48 | 39.69 | 50214 | 16 | 21169430 | 22514924 | 21319002 | 24075628 | 20066322 | 23467716 | 20196014 | 20059222 | 21600058 | 22857650 | 26000134 | 25986380 | 24600252 | 24747890 | 27922516 | 22149760 | 19734388 | 14346254 | 23154966 | 23509356 | 0 | 11 | 0 | 12 | 0 | 12 | 1.18 | 1.19 | 0.94 | 0.95 |
| P05943 | S100a10 | Protein S100-A10 | 98.91 | 17.89 | 11075 | 1 | 11001008 | 9393460 | 7963365.5 | 8564046 | 10712490 | 9116927 | 10527626 | 6492688.5 | 9769004 | 9487778 | 2356027.25 | 2953880.75 | 3011131.25 | 1526155.5 | 1747811.62 | 9678669 | 3971003.75 | 7388694.5 | 7632011.5 | 6368850.5 | 0 | 3 | 0 | 5 | 0 | 5 | 0.24 | 0.26 | 0.74 | 0.77 |
| P50543 | S100a11 | Protein S100-A11 | 403.69 | 54.08 | 11083 | 6 | 10671228 | 10670198 | 12432641 | 15443588 | 10546433 | 10812181 | 12204094 | 13666889 | 11308174 | 12880784 | 5095480.5 | 6751019 | 8899674 | 9728116 | 8814536 | 11772969 | 11930135 | 13466477 | 14600459 | 13965316 | 0 | 7 | 0 | 7 | 0 | 7 | 0.66 | 0.65 | 1.10 | 1.08 |
| P07091 | S100a4 | Protein S100-A4 | 493.93 | 51.49 | 11721 | 9 | 88327370 | 116157114 | 163944466 | 143027602 | 114054770 | 81118722 | 86242946 | 133616802 | 115081586 | 131295946 | 57573170 | 76519338 | 76355466 | 65711698 | 75551826 | 75056370 | 70227930 | 90545482 | 77844002 | 110398274 | 0 | 2 | 0 | 4 | 0 | 4 | 0.56 | 0.64 | 0.68 | 0.77 |
| P14069 | S100a6 | Protein S100-A6 | 387.42 | 78.65 | 10051 | 6 | 119124594 | 98337914 | 106512554 | 164490178 | 67167138 | 105661602 | 110420474 | 124099138 | 121879874 | 149827522 | 39358382 | 12969644 | 22147996 | 21893664 | 24985826 | 106061162 | 69931898 | 54505910 | 121893946 | 106547890 | 0 | 10 | 0 | 5 | 0 | 10 | 0.22 | 0.20 | 0.83 | 0.75 |
| Q60710 | Samhd1 | Deoxynucleoside triphosphate triphosphohydrolase SAMHD1 | 555.56 | 26.14 | 75893 | 11 | 7444854 | 5846682.5 | 6033026.5 | 8582454 | 13852412 | 7235161.5 | 6962161 | 9855420 | 11346352 | 9843856 | 2625795.5 | 1700432.5 | 1853306.12 | 1299092 | 2552601.25 | 8640591 | 4774947 | 6330687.5 | 6968790 | 7195451 | 0 | 10 | 0 | 4 | 0 | 10 | 0.24 | 0.22 | 0.81 | 0.75 |
| Q8BGH2 | Samm50 | Sorting and assembly machinery component 50 homolog | 733.11 | 31.56 | 51864 | 13 | 15020742 | 15960356 | 12920978 | 14849282 | 14670690 | 17489704 | 16224836 | 16038938 | 18184382 | 19100916 | 32387718 | 26557496 | 29954280 | 26219598 | 27886112 | 15113241 | 16554093 | 22016088 | 14802404 | 18027764 | 0 | 4 | 0 | 10 | 0 | 10 | 1.95 | 1.64 | 1.18 | 0.99 |
| Q5HZY2 | Sar1b | GTP-binding protein SAR1b | 153.04 | 18.18 | 22410 | 3 | 2563497 | 2448004.75 | 2489419.5 | 2663990 | 3523273.75 | 2622371 | 2774789 | 3087905.5 | 3510372 | 2640752.5 | 1683966.75 | 2310315.75 | 1939122.75 | 1454351.75 | 2269699 | 2378401 | 2811529.5 | 2846717.25 | 2760782.25 | 2742306.25 | 0 | 9 | 0 | 10 | 0 | 10 | 0.71 | 0.66 | 0.99 | 0.93 |

| | | | | | | | | | | | | | | | | | | | | | | | | | | | | | | | | | | | | |
|---|---|---|---|---|---|---|---|---|---|---|---|---|---|---|---|---|---|---|---|---|---|---|---|---|---|---|---|---|---|---|---|---|---|---|---|---|
| P26638 | Sars | Serine--tRNA ligase, cytoplasmic | 1470.6 | 48.63 | 58389 | 24 | 77054826 | 81006650 | 59921162 | 73771042 | 69507762 | 88254218 | 82434162 | 92004146 | 90575170 | 96642722 | 118495394 | 119375442 | 122594314 | 106728194 | 118132714 | 87444866 | 100045874 | 94118394 | 102344306 | 101557522 | 0 | 0 | 0 | 5 | 0 | 5 | 1.62 | 1.30 | 1.34 | 1.08 |
| Q63623 | Scaf8 | Protein SCAF8 | 75.3 | 0.95 | 139504 | 1 | 990841.06 | 982865.5 | 1003612.62 | 945580.38 | 1120100.5 | 869558.06 | 927541.56 | 920176.62 | 1256072.25 | 828604.81 | 667548.56 | 621888.12 | 784115.81 | 2 | 800262.06 | 894869.69 | 498700.78 | 762440.12 | 632900.06 | 618342.56 | 0 | 0 | 0 | 2 | 0 | 2 | 0.57 | 0.60 | 0.68 | 0.71 |
| O35609 | Scamp3 | Secretory carrier-associated membrane protein 3 | 338.71 | 17.19 | 38458 | 4 | 289626.97 | 687346.19 | 1696406.62 | 1303280.88 | 2 | 1808945.75 | 1691631.25 | 1727114.75 | 1212598.25 | 2141305.5 | 4791401 | 5599834 | 6499896.5 | 5779287 | 5510377 | 824772.5 | 1974613.88 | 3148148 | 1375415.12 | 2016337.88 | 0 | 3 | 0 | 11 | 0 | 11 | 7.09 | 3.28 | 2.35 | 1.09 |
| Q9JLI6 | Scly | Selenocysteine lyase | 402.04 | 18.06 | 47174 | 7 | 7425405 | 9750254 | 10050072 | 10042924 | 9260524 | 7577197.5 | 8526412 | 7397674.5 | 7318078 | 7392763 | 2229914.25 | 4899934 | 4693641.5 | 4191719 | 4401672.5 | 8648386 | 7943784 | 8279063.5 | 8696999 | 6899012.5 | 0 | 4 | 0 | 7 | 0 | 7 | 0.44 | 0.53 | 0.87 | 1.06 |
| O08992 | Sdcbp | Syntenin-1 | 702.34 | 53.18 | 32379 | 11 | 3832622.5 | 5880057 | 3926586 | 4341193 | 3925483 | 5630713 | 6097584 | 6470009 | 7865225.5 | 7833768.5 | 40593762 | 35881806 | 48236338 | 44669734 | 35225198 | 4435854 | 6549850 | 9149730 | 8121668.5 | 9681491 | 0 | 1 | 0 | 7 | 0 | 7 | 9.34 | 6.04 | 1.73 | 1.12 |
| Q9CQA3 | Sdhb | Succinate dehydrogenase [ubiquinone] iron-sulfur subunit, mitochondrial | 581.69 | 34.04 | 31814 | 9 | 22873316 | 22867406 | 25181834 | 22721846 | 35227602 | 19885726 | 20474962 | 29806240 | 29286202 | 20524770 | 6601999 | 13779550 | 16132749 | 11327063 | 17380944 | 25931516 | 13290488 | 20188190 | 20664492 | 22878700 | 0 | 7 | 0 | 10 | 0 | 10 | 0.51 | 0.54 | 0.80 | 0.86 |
| P42667 | Sec11a | Signal peptidase complex catalytic subunit SEC11A | 225.82 | 16.2 | 20599 | 4 | 1007853.81 | 2732271.25 | 2168367 | 292805.44 | 498653.12 | 1139665.88 | 1190433.12 | 1049093.5 | 316108.44 | 1153780.75 | 3085148.5 | 4047475 | 4111522.5 | 3519881 | 4213876 | 243213.81 | 2 | 979051.88 | 509392.97 | 928253.25 | 0 | 6 | 0 | 3 | 0 | 6 | 2.83 | 3.91 | 0.40 | 0.55 |
| Q6NZC7 | Sec23ip | SEC23-interacting protein | 56.25 | 0.8 | 110780 | 1 | 1932839.12 | 2259049.75 | 1825957.5 | 2102788.5 | 1936469.88 | 2334753.25 | 1870426.88 | 1872502 | 1991176.75 | 2529631.25 | 1490542.38 | 1760109 | 1783496.5 | 1481181.25 | 1370724.38 | 2423845.25 | 1971338.88 | 2451424.25 | 2152265 | 2741245 | 0 | 3 | 0 | 7 | 0 | 7 | 0.78 | 0.74 | 1.17 | 1.11 |
| P61620 | Sec61a1 | Protein transport protein Sec61 subunit alpha isoform 1 | 409.6 | 24.16 | 52265 | 7 | 8621367 | 6291027.5 | 16741497 | 8259011 | 13128453 | 9086017 | 15083565 | 24726082 | 15644648 | 13308394 | 34289962 | 34015670 | 46941578 | 40086746 | 49122062 | 18931488 | 17729576 | 16487887 | 17550012 | 21226606 | 0 | 1 | 0 | 5 | 0 | 5 | 3.85 | 2.63 | 1.73 | 1.18 |
| Q9CQS8 | Sec61b | Protein transport protein Sec61 subunit beta | 161.01 | 37.5 | 9958 | 3 | 14511030 | 13737012 | 13301174 | 17340382 | 12870361 | 14551708 | 14236868 | 16459350 | 13303834 | 15610830 | 10089208 | 9109832 | 9717178 | 9010318 | 11199416 | 17623090 | 20536134 | 17465326 | 16914198 | 19503906 | 0 | 1 | 0 | 0 | 0 | 1 | 0.68 | 0.66 | 1.28 | 1.24 |
| Q9EP97 | Senp3 | Sentrin-specific protease 3 | 70.87 | 2.11 | 64403 | 1 | 103757.62 | 2 | 2 | 2 | 2 | 2 | 2 | 2 | 170578.7 | 89210.13 | 400270.19 | 473290.22 | 319120.59 | 207233.36 | 341449.38 | 2 | 2 | 2 | 2 | 2 | 0 | 10 | 0 | 7.5 | 0 | 10 | 16.78 | 6.70 | 0.00 | 0.00 |
| Q9QZI9 | Serinc3 | Serine incorporator 3 | 137.98 | 6.57 | 52623 | 3 | 1077126.75 | 617930.88 | 590843.75 | 478443.31 | 781988 | 746361.44 | 1212246.62 | 1011099.06 | 238469.72 | 518441 | 1611046.5 | 2132591.5 | 2428575 | 1548024.25 | 1568451.62 | 1559047.62 | 673105.81 | 761067.38 | 2 | 505818.62 | 0 | 12 | 0 | 11 | 0 | 12 | 2.62 | 2.49 | 0.99 | 0.94 |
| Q4G075 | Serpinb1a | Leukocyte elastase inhibitor A | 76.5 | 2.64 | 42727 | 1 | 387030.69 | 545643.56 | 259102.84 | 287193.97 | 314463.31 | 581456.38 | 571179.94 | 699026.12 | 795223.5 | 671208.62 | 2292143.25 | 1203094.38 | 2249718.25 | 1771185.5 | 1215161.38 | 601279 | 386916.78 | 1041289.94 | 1056485.75 | 1368197 | 0 | 2 | 0 | 8 | 0 | 8 | 4.87 | 2.63 | 2.48 | 1.34 |
| Q60854 | Serpinb6 | Serpin B6 | 1039.49 | 51.85 | 42599 | 15 | 106981778 | 98591338 | 99536754 | 106529506 | 105132810 | 90846442 | 95201674 | 87321202 | 89142626 | 93028650 | 46619314 | 48816950 | 43516294 | 40650730 | 46861850 | 101960290 | 85956554 | 91821602 | 99014770 | 96654858 | 0 | 3 | 0 | 7 | 0 | 7 | 0.44 | 0.50 | 0.92 | 1.04 |
| P32261 | Serpinc1 | Antithrombin-III | 592.06 | 16.56 | 52004 | 9 | 18538054 | 14376824 | 5103202.5 | 8564908 | 11437041 | 24077028 | 30826878 | 22985050 | 25600706 | 19469042 | 53209658 | 82062754 | 90721018 | 58913722 | 62685310 | 23230312 | 28633794 | 27377490 | 23029742 | 18607772 | 0 | 0 | 0 | 12 | 0 | 12 | 5.99 | 2.83 | 2.08 | 0.98 |
| Q64268 | Serpind1 | Heparin cofactor 2 | 91.62 | 2.92 | 54552 | 2 | 2 | 2 | 314710.41 | 255790.31 | 413022.44 | 138487.03 | 343577.31 | 574945.69 | 789393.62 | 2 | 1096599.12 | 1728344.62 | 2421166.5 | 1441650.12 | 2288313.5 | 2 | 2 | 515004.19 | 468467.06 | 332526.38 | 0 | 9 | 0 | 9 | 0 | 9 | 9.13 | 4.86 | 1.34 | 0.71 |
| Q8BP27 | Sfr1 | Swi5-dependent recombination DNA repair protein 1 homolog | 65.27 | 3.76 | 35183 | 1 | 1312765.88 | 1612694.38 | 1292757.12 | 1458227.5 | 990601.75 | 1130347.5 | 1376673.75 | 1234842.62 | 838600.44 | 1215851.25 | 615109.19 | 329179.59 | 585893.19 | 567439.88 | 623369.94 | 1363618 | 1296308.62 | 1203165.12 | 1536355.75 | 1257274.88 | 0 | 11 | 0 | 6 | 0 | 11 | 0.41 | 0.47 | 1.00 | 1.15 |
| Q8R0X7 | Sgpl1 | Sphingosine-1-phosphate lyase 1 | 478.87 | 19.19 | 63677 | 9 | 8090855 | 9528189 | 8730407 | 8180365 | 9072290 | 8932219 | 8776986 | 8291352.5 | 9125721 | 9289983 | 9977245 | 9836623 | 13593261 | 11121256 | 13829608 | 7439228.5 | 6665845 | 9569956 | 12551461 | 9404609 | 0 | 11 | 0 | 10 | 0 | 11 | 1.34 | 1.31 | 1.05 | 1.03 |
| Q9JJU8 | Sh3bgrl | SH3 domain-binding glutamic acid-rich-like protein | 374.06 | 50.88 | 12811 | 6 | 18866464 | 19760272 | 18795056 | 19020624 | 23148000 | 19019854 | 21931450 | 24001816 | 24158814 | 20272888 | 13675312 | 15154845 | 11057924 | 11294087 | 10625626 | 22101468 | 19115978 | 20325136 | 23793260 | 18102208 | 0 | 9 | 0 | 9 | 0 | 9 | 0.62 | 0.57 | 1.04 | 0.95 |
| Q91VW3 | Sh3bgrl3 | SH3 domain-binding glutamic acid-rich-like protein 3 | 109.88 | 31.18 | 10477 | 2 | 13821302 | 14168911 | 12134226 | 12008428 | 13059073 | 13300386 | 13863918 | 12608497 | 13517594 | 13337844 | 11464357 | 11573538 | 9181436 | 9144313 | 7553110 | 13221516 | 17893010 | 14644429 | 15952626 | 15471372 | 0 | 2 | 0 | 4 | 0 | 4 | 0.75 | 0.73 | 1.18 | 1.16 |
| Q6AYE2 | Sh3glb1 | Endophilin-B1 | 241.1 | 9.32 | 40787 | 3 | 2665214 | 2020140.5 | 3000006.5 | 4654448.5 | 4677355.5 | 1177232.62 | 1990481.5 | 3083532.5 | 3286194 | 3761170.5 | 559197.56 | 680699.88 | 1143869.25 | 281448.28 | 475682.72 | 2409213.25 | 2296083 | 3231830 | 3674788.75 | 3883393.25 | 0 | 11 | 0 | 9 | 0 | 11 | 0.18 | 0.24 | 0.91 | 1.17 |
| A2AAY5 | Sh3pxd2b | SH3 and PX domain-containing protein 2B | 119.87 | 2.75 | 101517 | 3 | 4576874 | 4816674.5 | 5494536 | 5309791 | 5257893 | 5907214 | 4185331.75 | 4097920.5 | 5079915 | 5497254.5 | 3760530 | 3752843.25 | 3529200.75 | 2788497 | 3702362.25 | 4593471 | 5079869.5 | 4366471 | 4576754 | 4856949 | 0 | 5 | 0 | 10 | 0 | 10 | 0.69 | 0.71 | 0.92 | 0.95 |
| Q9CZN7 | Shmt2 | Serine hydroxymethyltransferase, mitochondrial | 2163.04 | 62.5 | 55759 | 34 | 113253394 | 109288442 | 107143194 | 126275066 | 98402882 | 123445994 | 102799906 | 115512834 | 106383762 | 136037330 | 176867714 | 164704290 | 184667970 | 210115906 | 200868290 | 110115274 | 134129266 | 143371474 | 144863922 | 137707794 | 0 | 2 | 0 | 4 | 0 | 4 | 1.69 | 1.60 | 1.21 | 1.15 |
| Q3UND0 | Skap2 | Src kinase-associated phosphoprotein 2 | 431.72 | 29.89 | 40712 | 8 | 13963265 | 11612911 | 16303496 | 17918418 | 16991520 | 14008840 | 15100614 | 16365465 | 18337558 | 16508020 | 9154706 | 10048664 | 9674024 | 10207499 | 10739722 | 15923850 | 14268601 | 16875846 | 16008804 | 15401702 | 0 | 11 | 0 | 11 | 0 | 11 | 0.65 | 0.62 | 1.02 | 0.98 |
| P41251 | Slc11a1 | Natural resistance-associated macrophage protein 1 | 61.13 | 2.55 | 59741 | 1 | 425383.78 | 434529.16 | 2 | 2 | 326375.72 | 2 | 311587.44 | 537489.12 | 2 | 2 | 790142.94 | 933358.75 | 941690.69 | 583074.94 | 1195092.38 | 2 | 2 | 613744.06 | 942486.62 | 2 | 0 | 12 | 0 | 10.5 | 0 | 12 | 3.75 | 5.23 | 1.31 | 1.83 |
| Q5RK27 | Slc12a7 | Solute carrier family 12 member 7 | 51.97 | 0.65 | 119378 | 1 | 1018795.5 | 939524.94 | 888938.38 | 1099566.88 | 1232809.75 | 1333430.75 | 1476397 | 1279761.62 | 1288075.5 | 1262896.88 | 1727469.38 | 2004153.12 | 2578650 | 2331914.75 | 2294464.75 | 1059226.12 | 1546978.62 | 1349643.25 | 1560873.12 | 1750682.88 | 0 | 2 | 0 | 6 | 0 | 6 | 2.11 | 1.65 | 1.40 | 1.09 |
| Q99MR3 | Slc12a9 | Solute carrier family 12 member 9 | 248.7 | 5.58 | 96330 | 5 | 2100301.5 | 2051488.75 | 1955887.12 | 2216476.5 | 2240842.75 | 2099289.75 | 2045757 | 1975483.88 | 1888544.12 | 1794772.88 | 1338324.25 | 1089027.5 | 1410522.75 | 1406231.5 | 1553677.5 | 1691158.12 | 2369065.5 | 1590572.5 | 1687382 | 1961738.25 | 0 | 6 | 0 | 7 | 0 | 7 | 0.64 | 0.69 | 0.88 | 0.95 |
| Q8BPX9 | Slc15a3 | Solute carrier family 15 member 3 | 686.54 | 19.9 | 64051 | 10 | 16337541 | 15653552 | 7273231 | 13383588 | 9678633 | 21308606 | 18243898 | 13165218 | 11726833 | 15117490 | 45517138 | 34738442 | 49076998 | 50358462 | 52886698 | 14389032 | 15866988 | 15696881 | 18914514 | 20250774 | 0 | 4 | 0 | 9 | 0 | 9 | 3.73 | 2.92 | 1.37 | 1.07 |
| P53986 | Slc16a1 | Monocarboxylate transporter 1 | 308.93 | 11.56 | 53267 | 5 | 1753786.12 | 1571419.25 | 1593456.25 | 1967701 | 1440187.62 | 1833914.38 | 1861837.38 | 2252977 | 1714045.88 | 2368603 | 5249233 | 3753128.75 | 4730612.5 | 5150874 | 4936248.5 | 2235543.5 | 2474423.25 | 2531022.5 | 2921630.25 | 2711864.25 | 0 | 0 | 0 | 2 | 0 | 2 | 2.86 | 2.37 | 1.55 | 1.28 |
| P51912 | Slc1a5 | Neutral amino acid transporter B(0) | 922.38 | 30.02 | 58483 | 12 | 1092364 | 2690801 | 3025113.5 | 4033137.5 | 2239726.25 | 2677644.25 | 3116386.75 | 4434341.5 | 3646883.5 | 4249711 | 31546544 | 35856170 | 34753970 | 29187606 | 33871846 | 1293785.38 | 2617361.75 | 4687115 | 4614672 | 6186011.5 | 0 | 7 | 0 | 10 | 0 | 10 | 12.63 | 9.12 | 1.48 | 1.07 |
| P51881 | Slc25a5 | ADP/ATP translocase 2 | 2069.79 | 81.21 | 32931 | 29 | 136445954 | 119045322 | 98111826 | 111667826 | 107200954 | 129107666 | 130604322 | 118307490 | 112127714 | 122774722 | 169108754 | 164528994 | 193597986 | 185790258 | 176077282 | 132594658 | 111761234 | 108288658 | 82816778 | 101264122 | 0 | 10 | 0 | 5 | 0 | 10 | 1.55 | 1.45 | 0.94 | 0.88 |
| P11167 | Slc2a1 | Solute carrier family 2, facilitated glucose transporter member 1 | 473.83 | 13.82 | 53963 | 7 | 3092552.75 | 3433677 | 561326.44 | 936228.94 | 1531889.62 | 3129515.5 | 4705844 | 3448770.5 | 4108368.75 | 2028025.62 | 20963490 | 21958016 | 32201954 | 21074078 | 23149682 | 3006910.25 | 2648212.25 | 4745789.5 | 2745733 | 3565960 | 0 | 6 | 0 | 11 | 0 | 11 | 12.49 | 6.85 | 1.75 | 0.96 |
| P32037 | Slc2a3 | Solute carrier family 2, facilitated glucose transporter member 3 | 52.41 | 1.83 | 53479 | 1 | 2 | 2 | 136959.09 | 2 | 2 | 2 | 284874.81 | 102861.85 | 2 | 126622.92 | 1171268.62 | 1095802.5 | 1190946.12 | 1028640.81 | 681002.62 | 110935.09 | 2 | 100500.05 | 144059.31 | 161354.94 | 0 | 5 | 0 | 11 | 0 | 11 | 37.73 | 10.05 | 3.77 | 1.00 |
| Q91ZN5 | Slc35b2 | Adenosine 3'-phospho 5'-phosphosulfate transporter 1 | 65.68 | 2.32 | 47370 | 1 | 242179.84 | 516026.78 | 405062.41 | 2 | 2 | 2 | 2 | 195755.38 | 2 | 261278.17 | 1322608.88 | 1439141.25 | 1604640.75 | 1910172.5 | 1650316.62 | 264819.88 | 2 | 364685.72 | 366459.38 | 249601.11 | 0 | 12 | 0 | 4.5 | 0 | 12 | 6.81 | 17.34 | 1.07 | 2.73 |
| Q9WU81 | Slc37a2 | Glucose-6-phosphate exchanger SLC37A2 | 290.42 | 11.18 | 55073 | 5 | 8284388 | 8088368 | 5438423.5 | 9965466 | 9921874 | 13788850 | 13205134 | 11421332 | 11904541 | 13245604 | 16187891 | 15760414 | 17926422 | 18540176 | 16922654 | 15151308 | 14198181 | 13826449 | 16089322 | 17467638 | 0 | 0 | 0 | 0 | 0 | 0 | 2.05 | 1.34 | 1.84 | 1.21 |
| Q8K2P7 | Slc38a1 | Sodium-coupled neutral amino acid transporter 1 | 103.41 | 7.42 | 53795 | 2 | 2 | 2 | 2 | 2 | 2 | 251625.44 | 164979.72 | 2 | 172143.39 | 2 | 1321254 | 1296056.12 | 1966036.62 | 1627361.5 | 1767368.38 | 127891.23 | 245056.58 | 302188.34 | 205133.95 | 120290.04 | 0 | 0 | 0 | 8 | 0 | 8 | 797807.66 | 13.55 | 100056.01 | 1.70 |
| P10852 | Slc3a2 | Amino acid transporter heavy chain SLC3A2 | 1538.43 | 43.35 | 58337 | 23 | 19888242 | 22148812 | 20073252 | 21779348 | 18065094 | 23246186 | 27277712 | 22806582 | 22682570 | 25025252 | 74960418 | 88608218 | 90566370 | 74906730 | 89635922 | 27318094 | 29236768 | 27323670 | 25414426 | 28803020 | 0 | 0 | 0 | 1 | 0 | 1 | 4.11 | 3.46 | 1.35 | 1.14 |
| Q09143 | Slc7a1 | High affinity cationic amino acid transporter 1 | 252.8 | 11.74 | 67092 | 4 | 2 | 2 | 2 | 2 | 132812.67 | 2 | 2 | 2 | 2 | 130933.38 | 4413334 | 4058870.75 | 5414258.5 | 3760409.75 | 4264950 | 2 | 294749.12 | 2 | 94392.73 | 662648.31 | 0 | 7 | 0 | 7 | 0 | 7 | 164.97 | 167.34 | 7.92 | 8.03 |
| Q9WTR6 | Slc7a11 | Cystine/glutamate transporter | 182.95 | 9.96 | 55456 | 4 | 2 | 2 | 2 | 2 | 2 | 2 | 2 | 2 | 2 | 2 | 4133776.5 | 4614765.5 | 2624050.25 | 2364724 | 2970044.5 | 2 | 2 | 2 | 407776.38 | 2 | 0 | 10 | 0 | 10 | 0 | 10 | 1670736.08 | 1670736.08 | 40778.44 | 40778.44 |
| Q9Z127 | Slc7a5 | Large neutral amino acids transporter small subunit 1 | 325.93 | 12.7 | 55872 | 5 | 2 | 2 | 2 | 2 | 2 | 2 | 2 | 2 | 2 | 2 | 256247.91 | 779812.25 | 1133775.88 | 986283.56 | 1245036 | 2 | 2 | 2 | 2 | 2 | 0 | 12.5 | 0 | 12.5 | 0 | 12.5 | 440115.56 | 440115.56 | 1.00 | 1.00 |
| O54941 | Smarce1 | SWI/SNF-related matrix-associated actin-dependent regulator of chromatin subfamily E member 1 | 295.93 | 15.82 | 46638 | 5 | 2085180.75 | 2293400.75 | 2577778.5 | 2412403.75 | 2306473 | 2575538 | 2261587 | 2542401.5 | 1876429.12 | 2045170.75 | 782059.31 | 351291.19 | 868636.56 | 424600.12 | 1028748 | 2224819 | 2025830.88 | 1507150.88 | 1624470 | 2032158.38 | 0 | 1 | 0 | 4 | 0 | 4 | 0.30 | 0.31 | 0.81 | 0.83 |
| Q8CG47 | Smc4 | Structural maintenance of chromosomes protein 4 | 1006.57 | 16.64 | 146895 | 20 | 15776996 | 14905833 | 19238018 | 18377084 | 16628891 | 16274200 | 14682889 | 14707581 | 15214558 | 17012034 | 11344594 | 13815537 | 11965476 | 12212104 | 11445322 | 15223131 | 12435067 | 15887319 | 11517246 | 11711581 | 0 | 3 | 0 | 6 | 0 | 6 | 0.72 | 0.78 | 0.79 | 0.86 |
| Q6PE01 | Snrnp40 | U5 small nuclear ribonucleoprotein 40 kDa protein | 177.57 | 12.29 | 39276 | 4 | 2 | 2 | 1009125.12 | 881010.31 | 756917.94 | 889579.06 | 2 | 827932.5 | 814988.81 | 1232936.25 | 2019779.25 | 1498711 | 1485365.5 | 1250833.38 | 1949984.88 | 781509.06 | 840304.12 | 1036728.06 | 759379 | 753730.31 | 0 | 9 | 0 | 10 | 0 | 10 | 3.10 | 2.18 | 1.58 | 1.11 |
| P57784 | Snrpa1 | U2 small nuclear ribonucleoprotein A' | 840.84 | 50.59 | 28357 | 13 | 17520504 | 14251923 | 16313620 | 14983261 | 11892303 | 16682445 | 13602146 | 12600880 | 11334971 | 14528315 | 9955409 | 8868412 | 9746193 | 8234028.5 | 10349696 | 12782103 | 18016530 | 14226346 | 14730334 | 16221990 | 0 | 12 | 0 | 7 | 0 | 12 | 0.63 | 0.69 | 1.01 | 1.11 |
| P17136 | Snrpb | Small nuclear ribonucleoprotein-associated protein B | 338.03 | 19.48 | 23656 | 6 | 15090585 | 13530329 | 15971527 | 17942868 | 13379040 | 20241172 | 13761482 | 14220442 | 15501166 | 14815506 | 22406536 | 23853284 | 38591414 | 38106618 | 34259586 | 13813062 | 18725580 | 15067281 | 15945593 | 17658434 | 0 | 9 | 0 | 9 | 0 | 9 | 2.07 | 2.00 | 1.07 | 1.03 |
| D3ZCL3 | Snrpc | U1 small nuclear ribonucleoprotein C | 115.07 | 13.21 | 17364 | 2 | 4499091.5 | 5410393 | 5943541.5 | 4372082.5 | 6755793 | 2838375.5 | 5505882.5 | 7584885.5 | 6568210 | 4802548.5 | 853303.06 | 2686158 | 1953502.88 | 1461114.38 | 2645405 | 6224008 | 1153085.62 | 3162350.25 | 2222208 | 5683057.5 | 0 | 7 | 0 | 7 | 0 | 7 | 0.36 | 0.35 | 0.68 | 0.68 |
| P62317 | Snrpd2 | Small nuclear ribonucleoprotein Sm D2 | 302.68 | 44.07 | 13527 | 6 | 912534.25 | 1225675.5 | 360644.03 | 994331.88 | 175024.23 | 515330.38 | 1074006.25 | 1019221.25 | 705900.62 | 920810.25 | 8044850.5 | 8167752 | 6049217.5 | 6107753 | 7684966 | 1173560.62 | 2060604.5 | 1610716.5 | 904615 | 1347343.25 | 0 | 4 | 0 | 3 | 0 | 4 | 9.83 | 8.51 | 1.93 | 1.68 |
| O70493 | Snx12 | Sorting nexin-12 | 187.6 | 26.06 | 19116 | 4 | 3277677.75 | 3786535.75 | 4023411.75 | 4478184.5 | 5031220 | 3719509 | 3473516.5 | 3945523.5 | 3859011.75 | 4230772.5 | 2562999.25 | 2764646.25 | 3221425 | 2879050.75 | 3251033.5 | 3400517.5 | 3644481.25 | 4985979 | 3776848.5 | 4150284 | 0 | 10 | 0 | 12 | 0 | 12 | 0.71 | 0.76 | 0.97 | 1.04 |
| Q8BVL3 | Snx17 | Sorting nexin-17 | 49.35 | 3.62 | 52797 | 1 | 2 | 2 | 2 | 2 | 2 | 2 | 2 | 2 | 2 | 541783.94 | 1184046.5 | 1496173.75 | 1648145.88 | 1562972.62 | 1601976.12 | 2 | 2 | 2 | 2 | 818645 | 0 | 10 | 0 | 12 | 0 | 12 | 749331.49 | 13.83 | 81865.30 | 1.51 |
| Q9CWK8 | Snx2 | Sorting nexin-2 | 705.45 | 27.55 | 58471 | 13 | 28008800 | 28245520 | 30118582 | 28565694 | 27457274 | 27159664 | 22949344 | 25627796 | 22638730 | 27853322 | 18316154 | 18631186 | 13212924 | 13219069 | 15585130 | 25983278 | 26836168 | 25588860 | 21557936 | 23151830 | 0 | 0 | 0 | 10 | 0 | 10 | 0.55 | 0.63 | 0.86 | 0.98 |
| Q9D8U8 | Snx5 | Sorting nexin-5 | 1077.28 | 44.55 | 46797 | 19 | 52047294 | 48658690 | 57072854 | 47560386 | 65534822 | 41640470 | 48681466 | 48379626 | 57577694 | 44404850 | 22220066 | 22047642 | 18501490 | 18091088 | 17174900 | 49635094 | 38683350 | 38202426 | 38331730 | 40232758 | 0 | 2 | 0 | 4 | 0 | 4 | 0.36 | 0.41 | 0.76 | 0.85 |
| P09671 | Sod2 | Superoxide dismutase [Mn], mitochondrial | 436 | 40.99 | 24603 | 7 | 39840390 | 41133922 | 26598114 | 32391068 | 35334958 | 40511462 | 44791502 | 42865002 | 40751490 | 38829334 | 49554490 | 73128018 | 79063234 | 60683918 | 90468634 | 47739830 | 30503558 | 45923254 | 50417366 | 56798022 | 0 | 4 | 0 | 5 | 0 | 5 | 2.01 | 1.70 | 1.32 | 1.11 |
| Q9CYN2 | Spcs2 | Signal peptidase complex subunit 2 | 414.12 | 42.04 | 24978 | 7 | 1722744.88 | 2605666.25 | 2344027.75 | 4973374 | 1782055.62 | 2501666.5 | 3095051.5 | 4277617 | 4776263 | 4640303.5 | 9324331 | 6856048 | 9855954 | 9661505 | 12678596 | 1873690.88 | 1027797.12 | 4065473.5 | 2068306.62 | 3801496 | 0 | 12 | 0 | 4 | 0 | 12 | 3.60 | 2.51 | 0.96 | 0.67 |
| Q6ZWQ7 | Spcs3 | Signal peptidase complex subunit 3 | 257.02 | 23.89 | 20313 | 5 | 3235333 | 3445539.25 | 5148136.5 | 4595276 | 4170507.25 | 3416268.25 | 4225007 | 5648856 | 3542826 | 6034589 | 10854318 | 8906214 | 12945522 | 12575963 | 12378706 | 3635967.5 | 3007613.5 | 6410599 | 5245986 | 1979988.62 | 0 | 12 | 0 | 10 | 0 | 12 | 2.80 | 2.52 | 0.98 | 0.89 |
| Q9JIA7 | Sphk2 | Sphingosine kinase 2 | 269.84 | 8.91 | 65618 | 4 | 2695946.5 | 3268200.25 | 2825148.5 | 3535600.5 | 3259539.5 | 2693454.25 | 2734366 | 2320909.75 | 3804473 | 3294433.25 | 4322568.5 | 5356724 | 6931207 | 4950845 | 4531275 | 2618167.25 | 2838938 | 3611277.75 | 3451589.25 | 4419155.5 | 0 | 9 | 0 | 8 | 0 | 9 | 1.67 | 1.76 | 1.09 | 1.14 |
| Q64337 | Sqstm1 | Sequestosome-1 | 1253.64 | 70.14 | 48163 | 18 | 1451793.62 | 2246445.75 | 1834648 | 1120818.62 | 4345621.5 | 2448268 | 2032139.75 | 5285084 | 3792636.5 | 2957902 | 25558482 | 28205200 | 29292420 | 30737276 | 26396636 | 961176.19 | 2662543.25 | 4325963.5 | 4868603.5 | 3270165.5 | 0 | 8 | 0 | 12 | 0 | 12 | 12.75 | 8.49 | 1.46 | 0.97 |
| D4A208 | Srgap2 | SLIT-ROBO Rho GTPase-activating protein 2 | 578.37 | 10.64 | 120878 | 11 | 9433355 | 11535967 | 9897790 | 10723176 | 8647508 | 7102847 | 9296492 | 10668539 | 11349170 | 6327836 | 5936505.5 | 6142075.5 | 3648795.25 | 5335513.5 | 5361621.5 | 7810840 | 4298244 | 9135708 | 10281437 | 10134199 | 0 | 7 | 0 | 10 | 0 | 10 | 0.53 | 0.59 | 0.83 | 0.93 |
| P16254 | Srp14 | Signal recognition particle 14 kDa protein | 187.04 | 28.18 | 12510 | 3 | 244074.97 | 334826.31 | 2 | 2 | 722679.31 | 594999.25 | 2 | 552726.31 | 2 | 2 | 1372225.5 | 1408954.5 | 785962.25 | 1617634.5 | 1300888.88 | 2 | 674579.69 | 794429.12 | 582132.88 | 2 | 0 | 10 | 0 | 8 | 0 | 10 | 4.98 | 5.65 | 1.58 | 1.79 |
| Q9D7A6 | Srp19 | Signal recognition particle 19 kDa protein | 51.46 | 8.33 | 16191 | 1 | 2 | 2 | 2 | 2 | 2 | 258494.36 | 371691.5 | 2 | 2 | 2 | 1169767.38 | 1155451 | 1088914.38 | 767659.38 | 1043021.19 | 2 | 380564.72 | 519635.16 | 2 | 450782.34 | 0 | 5 | 0 | 7 | 0 | 7 | 522481.33 | 8.29 | 135098.62 | 2.14 |
| P47758 | Srprb | Signal recognition particle receptor subunit beta | 577.44 | 47.21 | 29579 | 10 | 1644052.25 | 3644542 | 2173715.25 | 1191985.88 | 1946561 | 1123986.75 | 2326468.5 | 1861481.88 | 3526959.25 | 3132839.5 | 8178715 | 7087527 | 6688063.5 | 4698027 | 6462670 | 2598112 | 2181117.5 | 1770034.25 | 2166121.5 | 1211288.12 | 0 | 11 | 0 | 9 | 0 | 11 | 3.12 | 2.77 | 0.94 | 0.83 |
| Q52KI8 | Srrm1 | Serine/arginine repetitive matrix protein 1 | 471.81 | 10.25 | 106862 | 6 | 8501666 | 8367737 | 5465718 | 6422313 | 7979181 | 9026008 | 7528744.5 | 7740896 | 6283068 | 6295670.5 | 4239778 | 4102588.5 | 3496546.25 | 3361611.5 | 3310263 | 9675304 | 7939394.5 | 8023116.5 | 6684109 | 8120188.5 | 0 | 10 | 0 | 6 | 0 | 10 | 0.50 | 0.50 | 1.10 | 1.10 |
| Q99MR6 | Srrt | Serrate RNA effector molecule homolog | 630.18 | 18.29 | 100452 | 13 | 21995720 | 23455960 | 19229110 | 19756030 | 17280948 | 18112560 | 23224250 | 16981704 | 17988418 | 16172708 | 12476360 | 15949953 | 12719538 | 9912660 | 8687871 | 20492866 | 16400138 | 14768377 | 13165956 | 18199110 | 0 | 4 | 0 | 9 | 0 | 9 | 0.59 | 0.65 | 0.82 | 0.90 |
| O35326 | Srsf5 | Serine/arginine-rich splicing factor 5 | 418.67 | 25.65 | 30891 | 8 | 10937538 | 10982293 | 10987580 | 10292766 | 9157683 | 8857001 | 11269048 | 8214608 | 8681382 | 6688126 | 4166753 | 6373755 | 4360477.5 | 2824101.25 | 5229903.5 | 11962478 | 8997805 | 8260227 | 7920289 | 9187980 | 0 | 6 | 0 | 9 | 0 | 9 | 0.44 | 0.53 | 0.88 | 1.06 |
| Q9D975 | Srxn1 | Sulfiredoxin-1 | 45.05 | 6.62 | 14149 | 1 | 884547 | 1015408.5 | 419838.62 | 2 | 540690 | 1012313.19 | 1355964.38 | 424432.44 | 1356292.12 | 1661937.12 | 2668304.5 | 2834787.5 | 2518493 | 2501508.75 | 2235425.25 | 1430154.5 | 1663177.5 | 1931213.62 | 1946348.88 | 1901633 | 0 | 0 | 0 | 1 | 0 | 1 | 4.46 | 2.20 | 3.10 | 1.53 |
| P32067 | Ssb | Lupus La protein homolog | 904.95 | 29.4 | 47756 | 16 | 6337511 | 9612328 | 7216263 | 5035573 | 5421868.5 | 7294489 | 5495446 | 6151656 | 6590092.5 | 7342951.5 | 21787622 | 20509834 | 14319389 | 13757959 | 14239020 | 5670681.5 | 5397544 | 6107091 | 5579438 | 5849388.5 | 0 | 9 | 0 | 4 | 0 | 9 | 2.52 | 2.57 | 0.85 | 0.87 |
| Q08013 | Ssr3 | Translocon-associated protein subunit gamma | 93.1 | 7.57 | 21064 | 1 | 1990162.5 | 1654031.38 | 1374648.12 | 1178728.25 | 1448017 | 2074515.88 | 1456043.62 | 1468482.38 | 1474651.38 | 1440502.75 | 3474867.75 | 2769901.25 | 2115703 | 2559082 | 2402459.5 | 1356450.12 | 1848921.75 | 1651666.62 | 1671163.62 | 1477542 | 0 | 10 | 0 | 9 | 0 | 10 | 1.74 | 1.68 | 1.05 | 1.01 |
| Q07984 | Ssr4 | Translocon-associated protein subunit delta | 210.41 | 24.86 | 18980 | 3 | 1604991.25 | 2103111.5 | 1522137.38 | 1401177.25 | 1476040.88 | 1438376.25 | 1062950.5 | 1479739 | 1950872.38 | 2112323.5 | 10892649 | 5730854 | 6593353 | 5531906 | 6863768 | 1248981.88 | 1865891.5 | 1890804.75 | 3059338 | 1067931.88 | 0 | 12 | 0 | 12 | 0 | 12 | 4.39 | 4.43 | 1.13 | 1.14 |
| Q60864 | Stip1 | Stress-induced-phosphoprotein 1 | 1674.58 | 53.41 | 62582 | 30 | 60641998 | 68856458 | 77419146 | 71272322 | 67208698 | 57467882 | 70964122 | 76693562 | 68995274 | 60131582 | 45957194 | 54877430 | 45202874 | 41257334 | 44543050 | 72501842 | 47892182 | 63093210 | 58956938 | 63316910 | 0 | 6 | 0 | 9 | 0 | 9 | 0.67 | 0.69 | 0.89 | 0.91 |
| Q99JB2 | Stoml2 | Stomatin-like protein 2, mitochondrial | 819.92 | 39.09 | 38385 | 13 | 3247057.75 | 4795111.5 | 6000894 | 4381175 | 5679558.5 | 5151733 | 4573503 | 6593433 | 4864398.5 | 7093289 | 13879182 | 12203344 | 14314509 | 13304579 | 14631388 | 3618325.5 | 4639076 | 5955419.5 | 7335541 | 6708059 | 0 | 8 | 0 | 12 | 0 | 12 | 2.83 | 2.42 | 1.17 | 1.00 |
| P46978 | Stt3a | Dolichyl-diphosphooligosaccharide--protein glycosyltransferase subunit STT3A | 742.2 | 23.55 | 80598 | 14 | 6265917 | 6299527.5 | 8810481 | 8555198 | 11637380 | 5810295.5 | 6522062.5 | 11171837 | 13748020 | 10351836 | 20175936 | 19876758 | 24285232 | 21233918 | 26076642 | 8345627 | 7653878 | 11265142 | 10778248 | 9083614 | 0 | 9 | 0 | 12 | 0 | 12 | 2.69 | 2.35 | 1.13 | 0.99 |
| Q3TDQ1 | Stt3b | Dolichyl-diphosphooligosaccharide--protein glycosyltransferase subunit STT3B | 171.27 | 5.1 | 93246 | 4 | 2248959.5 | 2001860 | 2059143.38 | 2013612.75 | 1022686.25 | 2201152.75 | 2223372 | 1766951.75 | 1383887.5 | 2808298 | 3212638 | 4031338.25 | 3849412.25 | 3625664.25 | 5197618 | 1275616.5 | 450539.84 | 2204986.5 | 2967601.75 | 2462916.75 | 0 | 10 | 0 | 12 | 0 | 12 | 2.13 | 1.92 | 1.00 | 0.90 |
| G3V7P1 | Stx12 | Syntaxin-12 | 105.92 | 8.03 | 31187 | 2 | 1719255.62 | 1305995 | 1133618.62 | 1473813.5 | 1893167.75 | 1800021.12 | 2088166.75 | 1544452.25 | 1622441.12 | 1265656.12 | 585853.19 | 782807.81 | 457016.34 | 2 | 764505 | 1801625.25 | 1706609.25 | 1024475.19 | 424019.41 | 1220210.5 | 0 | 8 | 0 | 7 | 0 | 8 | 0.34 | 0.31 | 0.82 | 0.74 |
| Q64324 | Stxbp2 | Syntaxin-binding protein 2 | 838.44 | 29.85 | 66357 | 13 | 8264539.5 | 10177083 | 9969377 | 10936564 | 14781492 | 8387572.5 | 9428416 | 9893411 | 12293981 | 8434021 | 6286799 | 7404432 | 7382004.5 | 5837009 | 7525609 | 7825187.5 | 3038385.5 | 7859298 | 8973492 | 9092270 | 0 | 2 | 0 | 4 | 0 | 4 | 0.64 | 0.71 | 0.68 | 0.76 |
| Q9WUM5 | Suclg1 | Succinate--CoA ligase [ADP/GDP-forming] subunit alpha, mitochondrial | 389.17 | 28.61 | 36155 | 7 | 266026.28 | 146935.88 | 629895.81 | 336985.84 | 497807.41 | 778835 | 435396.75 | 421639.06 | 2 | 2 | 4491276.5 | 3400693.25 | 3587354.75 | 3475179 | 2992594.5 | 2 | 955763.31 | 2155392.5 | 570741.94 | 2094940.75 | 0 | 6 | 0 | 5 | 0 | 6 | 9.56 | 10.97 | 3.08 | 3.53 |
| Q9Z2I8 | Suclg2 | Succinate--CoA ligase [GDP-forming] subunit beta, mitochondrial | 880.65 | 40.88 | 46840 | 13 | 10280670 | 10705420 | 10269305 | 5408170 | 11952663 | 11235765 | 9224445 | 9855076 | 11100422 | 8714956 | 12401597 | 13851580 | 15877882 | 12241767 | 13992783 | 8732808 | 5630603 | 10992689 | 10710215 | 10440839 | 0 | 12 | 0 | 10 | 0 | 12 | 1.41 | 1.36 | 0.96 | 0.93 |
| Q920B9 | Supt16h | FACT complex subunit SPT16 | 1017.63 | 18.15 | 119825 | 18 | 17750218 | 15314425 | 19065550 | 18994786 | 16594437 | 14050922 | 14326595 | 13621839 | 15304444 | 13320341 | 11515145 | 10904390 | 12699456 | 9708514 | 11218954 | 14280917 | 16959898 | 11255006 | 14666848 | 13371911 | 0 | 2 | 0 | 12 | 0 | 12 | 0.64 | 0.79 | 0.80 | 1.00 |
| O55101 | Syngr2 | Synaptogyrin-2 | 51.91 | 4.46 | 24778 | 1 | 1709030.25 | 1906913 | 1621035.25 | 1618487.88 | 1532192.12 | 1364996.5 | 1477045.25 | 1885007.5 | 1630333.62 | 2016633.25 | 4265865 | 3648800.25 | 4277986.5 | 2728492.75 | 5369453 | 1276229.5 | 2112639.25 | 1555302.62 | 2919791.25 | 2134649 | 0 | 9 | 0 | 8 | 0 | 9 | 2.42 | 2.42 | 1.19 | 1.19 |
| Q62910 | Synj1 | Synaptojanin-1 | 110.2 | 1.78 | 172881 | 2 | 1474355.75 | 1432817.88 | 1119810.25 | 1273536.75 | 1869187.5 | 1371752.75 | 1475986.62 | 1224387.62 | 1578217.5 | 1238388.5 | 492486.06 | 326820.38 | 690777.62 | 2 | 318969.66 | 1334767.75 | 885104.69 | 1291559.38 | 1267210.25 | 1246238 | 0 | 6 | 0 | 8 | 0 | 8 | 0.26 | 0.27 | 0.84 | 0.87 |
| Q9WVA4 | Tagln2 | Transgelin-2 | 1591.4 | 83.42 | 22395 | 23 | 271746498 | 303059138 | 336224290 | 351228450 | 324337634 | 310738722 | 314420674 | 319820578 | 310664898 | 313230402 | 136568002 | 155350546 | 127087682 | 130126770 | 137312098 | 367751906 | 351242018 | 312328130 | 334709794 | 326324866 | 0 | 7 | 0 | 3 | 0 | 7 | 0.43 | 0.44 | 1.07 | 1.08 |
| Q9R233 | Tapbp | Tapasin | 388.41 | 19.78 | 49736 | 8 | 9727317 | 8397552 | 8588021 | 11483399 | 9566606 | 12305502 | 3546520 | 11835830 | 10184644 | 15558076 | 19232952 | 15875474 | 25049460 | 23214156 | 27107662 | 7931140 | 7166080 | 10750760 | 12770198 | 11950635 | 0 | 11 | 0 | 11 | 0 | 11 | 2.31 | 2.07 | 1.06 | 0.95 |
| Q921F2 | Tardbp | TAR DNA-binding protein 43 | 783.04 | 34.06 | 44548 | 11 | 12123194 | 14165007 | 15718791 | 17689606 | 12619078 | 13910926 | 11890182 | 14522097 | 13324718 | 15382147 | 27984660 | 22676334 | 21800302 | 18882880 | 19553854 | 13306463 | 16828668 | 15655214 | 16607968 | 15748365 | 0 | 8 | 0 | 4 | 0 | 8 | 1.53 | 1.61 | 1.08 | 1.13 |
| Q9D0R2 | Tars | Threonine--tRNA ligase, cytoplasmic | 1425.85 | 36.29 | 83356 | 25 | 43427090 | 46151354 | 44764730 | 46996278 | 47236006 | 45772214 | 46084098 | 49251734 | 47537162 | 46575022 | 59273970 | 61582594 | 56476550 | 53788234 | 55759810 | 53585938 | 53679594 | 54340378 | 48295926 | 42010406 | 0 | 5 | 0 | 6 | 0 | 6 | 1.26 | 1.22 | 1.10 | 1.07 |
| P48428 | Tbca | Tubulin-specific chaperone A | 212.54 | 33.33 | 12758 | 4 | 4672721 | 4814423 | 5042815.5 | 5478890 | 4346868 | 4928993.5 | 4850359 | 5189918 | 4862399 | 5944287 | 2356882 | 2567843.5 | 433375.59 | 1666626.38 | 1630974.62 | 4685166.5 | 5884220.5 | 4447779 | 6212237 | 4325394.5 | 0 | 12 | 0 | 9 | 0 | 12 | 0.36 | 0.34 | 1.05 | 0.99 |
| Q8VCN9 | Tbcc | Tubulin-specific chaperone C | 161.34 | 9.68 | 38125 | 3 | 4096669 | 3502541.5 | 4166846 | 3510925.5 | 3653037.5 | 4089593.75 | 3425092.5 | 3843232 | 3280742 | 3015101.5 | 2015552.12 | 858024.06 | 2225289.75 | 2972205.5 | 2231574.75 | 3424835.25 | 4434461.5 | 3469486.5 | 3726348.5 | 3871972 | 0 | 11 | 0 | 8 | 0 | 11 | 0.54 | 0.58 | 1.00 | 1.07 |
| Q8BHJ5 | Tbl1xr1 | F-box-like/WD repeat-containing protein TBL1XR1 | 46.76 | 1.36 | 55661 | 1 | 1092834.88 | 963483 | 1517772.25 | 1177737.5 | 1094568 | 873923.88 | 1413439.38 | 1189776.62 | 1400101.12 | 1078490.25 | 2 | 450232.88 | 546666.5 | 632096.19 | 673092.38 | 1296168.88 | 781580.19 | 1461068.5 | 974176.5 | 1370266.5 | 0 | 12 | 0 | 12 | 0 | 12 | 0.39 | 0.39 | 1.01 | 0.99 |
| P11983 | Tcp1 | T-complex protein 1 subunit alpha | 2515.75 | 76.08 | 60449 | 34 | 90973114 | 94899386 | 95042610 | 88726442 | 82915250 | 95490058 | 89679170 | 96641698 | 94252050 | 92220618 | 151331442 | 134397826 | 143738098 | 127757386 | 122795282 | 89502690 | 92142834 | 110300778 | 102088530 | 99428210 | 0 | 5 | 0 | 9 | 0 | 9 | 1.50 | 1.45 | 1.09 | 1.05 |
| P47226 | Tes | Testin | 453.74 | 18.68 | 47983 | 9 | 10818867 | 12386318 | 12795695 | 10784258 | 11755765 | 10686125 | 13779215 | 15689953 | 13232390 | 14455313 | 8433696 | 7313029 | 8954180 | 7460007 | 10542883 | 14978544 | 7800558.5 | 15889664 | 16578833 | 16903282 | 0 | 5 | 0 | 6 | 0 | 6 | 0.73 | 0.63 | 1.23 | 1.06 |
| Q3URQ0 | Tex10 | Testis-expressed protein 10 | 139.3 | 3.66 | 105209 | 3 | 553290 | 448033.88 | 437391.16 | 473350.31 | 133525.14 | 460767.16 | 556541.62 | 519022.81 | 725264.38 | 549603.19 | 1182027.62 | 1790131 | 1743041.25 | 859426.62 | 1452986 | 493846.81 | 380683.28 | 742256 | 98833.7 | 504249.19 | 0 | 11 | 0 | 7 | 0 | 11 | 3.44 | 2.50 | 1.09 | 0.79 |
| P04184 | Tk1 | Thymidine kinase, cytosolic | 164.13 | 13.3 | 25776 | 3 | 2892030.75 | 1987739.88 | 3241999.75 | 3374822.75 | 2555100.75 | 2446995.75 | 2102579.25 | 1693552 | 1756944.38 | 1738435.88 | 722069.44 | 756153.38 | 563158.69 | 696827.5 | 414510.78 | 2882336.25 | 2294389.25 | 1846818 | 1730751.5 | 1416123.62 | 0 | 3 | 0 | 12 | 0 | 12 | 0.22 | 0.32 | 0.72 | 1.04 |
| Q8BH24 | Tm9sf4 | Transmembrane 9 superfamily member 4 | 188.46 | 8.4 | 74693 | 4 | 2 | 2 | 334859.97 | 451891.38 | 452252.06 | 688459.69 | 650141.75 | 676567.81 | 974322.56 | 1097224.12 | 6443011 | 1697644.38 | 5361137 | 5718504 | 2963633.75 | 938133.12 | 2243355.5 | 1233456.62 | 2210930.5 | 1528219.38 | 0 | 0 | 0 | 2 | 0 | 2 | 17.90 | 5.43 | 6.58 | 2.00 |
| Q5I0H4 | Tmco1 | Calcium load-activated calcium channel | 230.78 | 24.47 | 21175 | 4 | 2733688 | 4988388 | 3927994.75 | 4257729 | 1717561.25 | 3494565.25 | 3253680.75 | 4113250 | 3575605.25 | 2046692.38 | 14430378 | 13558328 | 13931276 | 12779253 | 16817506 | 3812244.75 | 3258885 | 6359126.5 | 3860515.75 | 5267642.5 | 0 | 9 | 0 | 5 | 0 | 9 | 4.06 | 4.34 | 1.28 | 1.37 |
| Q8VC04 | Tmem106a | Transmembrane protein 106A | 222.41 | 17.62 | 29109 | 4 | 9620297 | 7243960.5 | 5672861 | 8146288.5 | 8766442 | 6758617 | 6608150.5 | 7364197 | 7345812.5 | 7623131.5 | 13810544 | 9788569 | 16667479 | 19443758 | 15550414 | 6260677.5 | 10893116 | 13576774 | 11895540 | 11176185 | 0 | 4 | 0 | 5 | 0 | 5 | 1.91 | 2.11 | 1.36 | 1.51 |
| Q2TA63 | Tmem147 | Transmembrane protein 147 | 88.6 | 7.59 | 25291 | 1 | 2 | 2 | 2 | 2 | 2 | 2 | 2 | 2 | 2 | 2 | 490748.38 | 347444.12 | 548589.38 | 502947.12 | 593281.06 | 2 | 2 | 2 | 2 | 2 | 0 | 12.5 | 0 | 12.5 | 0 | 12.5 | 248301.01 | 248301.01 | 1.00 | 1.00 |
| Q8BRG8 | Tmem209 | Transmembrane protein 209 | 54.83 | 1.78 | 62982 | 1 | 2 | 2 | 2 | 2 | 2 | 2 | 2 | 261868.62 | 2 | 2 | 431343.62 | 473719.16 | 431002.72 | 481933.38 | 342241.75 | 2 | 283077.53 | 345796.38 | 2 | 2 | 0 | 7.5 | 0 | 9 | 0 | 9 | 216024.06 | 8.25 | 62887.99 | 2.40 |
| Q9JHJ0 | Tmod3 | Tropomodulin-3 | 192.69 | 14.49 | 39503 | 4 | 7974889.5 | 9211874 | 9561117 | 10204860 | 9012192 | 8528276 | 10169114 | 9601069 | 9530439 | 9133718 | 4034687.5 | 5742610 | 4698123.5 | 6810591.5 | 5908413.5 | 9181456 | 7431710 | 11646372 | 8862454 | 11894171 | 0 | 12 | 0 | 12 | 0 | 12 | 0.59 | 0.58 | 1.07 | 1.04 |
| Q9D2E2 | Toe1 | Target of EGR1 protein 1 | 51.37 | 1.57 | 56871 | 1 | 240689.75 | 348054.19 | 2 | 209692.22 | 2 | 422291.28 | 306588.62 | 131209.36 | 198089.73 | 240515.98 | 739350.19 | 539524.94 | 804628.62 | 558759.5 | 627909.5 | 255743.98 | 298254.44 | 238489.62 | 2 | 205131.27 | 0 | 11 | 0 | 10 | 0 | 11 | 4.10 | 2.52 | 1.25 | 0.77 |
| Q9CYG7 | Tomm34 | Mitochondrial import receptor subunit TOM34 | 569.13 | 43.69 | 34278 | 11 | 11272926 | 10393163 | 7823002.5 | 8969852 | 8048139 | 9347286 | 11255762 | 7570602 | 9270702 | 9041349 | 4554240.5 | 5541701.5 | 5770787 | 4719654 | 5222508 | 11937507 | 10024173 | 8946195 | 9347988 | 10358782 | 0 | 9 | 0 | 7 | 0 | 9 | 0.55 | 0.56 | 1.09 | 1.09 |
| Q9QYA2 | Tomm40 | Mitochondrial import receptor subunit TOM40 homolog | 615.64 | 35.73 | 37895 | 9 | 9251899 | 7763679.5 | 9583023 | 5813566.5 | 8914863 | 11950823 | 9052842 | 10105860 | 10013694 | 12979997 | 19852198 | 15459592 | 18983086 | 16809520 | 19347960 | 9875995 | 9366529 | 14624446 | 13078961 | 12573201 | 0 | 1 | 0 | 9 | 0 | 9 | 2.19 | 1.67 | 1.44 | 1.10 |
| Q01320 | Top2a | DNA topoisomerase 2-alpha | 927.92 | 12.83 | 172790 | 18 | 10445980 | 12916295 | 10039114 | 10077986 | 6893747.5 | 12427716 | 10683110 | 9451205 | 6800433.5 | 8486842 | 4750600.5 | 3208588.25 | 2691038.75 | 4423362 | 3203772 | 11646091 | 7310758.5 | 7668849.5 | 4947201 | 6196644.5 | 0 | 6 | 0 | 6 | 0 | 6 | 0.36 | 0.38 | 0.75 | 0.79 |
| Q62393 | Tpd52 | Tumor protein D52 | 641.24 | 44.2 | 24313 | 9 | 21269064 | 15228310 | 22625582 | 27181578 | 21754294 | 18938474 | 16340134 | 20154434 | 22096812 | 23683718 | 3302527.5 | 4750591.5 | 6413511 | 6912796 | 6467339 | 18385800 | 21085200 | 21389488 | 15781546 | 20142348 | 0 | 6 | 0 | 9 | 0 | 9 | 0.26 | 0.28 | 0.90 | 0.96 |
| Q6PCT3 | Tpd52l2 | Tumor protein D54 | 300.59 | 20.45 | 23992 | 5 | 11525836 | 10089134 | 9574889 | 11059420 | 10928764 | 12482674 | 8609182 | 9754232 | 9691154 | 9380944 | 2478254.5 | 5149497 | 4558451.5 | 4748220 | 4460422.5 | 10945002 | 9330399 | 11797052 | 11371377 | 11984581 | 0 | 8 | 0 | 8 | 0 | 8 | 0.40 | 0.43 | 1.04 | 1.11 |
| P17751 | Tpi1 | Triosephosphate isomerase | 1281.95 | 73.91 | 32192 | 18 | 420965602 | 392756386 | 425830978 | 455259010 | 452426626 | 433234690 | 438809730 | 444903810 | 450406722 | 444476130 | 245421186 | 318317026 | 262772386 | 238289714 | 270637122 | 478385858 | 387606402 | 397192098 | 434600450 | 453474594 | 0 | 12 | 0 | 11 | 0 | 12 | 0.62 | 0.60 | 1.00 | 0.97 |
| Q63610 | Tpm3 | Tropomyosin alpha-3 chain | 1333.24 | 55.24 | 29007 | 22 | 99961754 | 123507162 | 133188962 | 149942562 | 128354010 | 105693666 | 106475194 | 127017666 | 116495874 | 124950882 | 60231322 | 65645806 | 63939442 | 75625986 | 64422370 | 107682018 | 111694250 | 113125362 | 123686450 | 118300330 | 0 | 6 | 0 | 12 | 0 | 12 | 0.52 | 0.57 | 0.90 | 0.99 |
| Q6IRU2 | Tpm4 | Tropomyosin alpha-4 chain | 1213.37 | 47.18 | 28468 | 18 | 25153866 | 32230522 | 37013850 | 30255678 | 43154262 | 19190398 | 33364026 | 32973046 | 45577914 | 24853768 | 8483115 | 7784271.5 | 9279722 | 5448241.5 | 10423978 | 26501210 | 17918886 | 29596338 | 22434162 | 22281102 | 0 | 2 | 0 | 6 | 0 | 6 | 0.25 | 0.27 | 0.71 | 0.76 |
| Q5XI41 | Tram1 | Translocating chain-associated membrane protein 1 | 201.03 | 6.95 | 43030 | 3 | 2408784.75 | 3156368.25 | 3551630.5 | 3537265.75 | 4136328 | 3345318 | 3510926 | 4229045.5 | 4435865 | 3933066.25 | 5836675 | 5500307 | 9011670 | 6949177 | 7263579.5 | 2335481 | 1538624.12 | 4616389 | 5792151.5 | 4170847.5 | 0 | 10 | 0 | 12 | 0 | 12 | 2.06 | 1.78 | 1.10 | 0.95 |
| Q9CQN1 | Trap1 | Heat shock protein 75 kDa, mitochondrial | 1214.69 | 35.13 | 80209 | 19 | 13624086 | 15375283 | 15122497 | 14231375 | 14619045 | 12139896 | 13643221 | 15703286 | 14251235 | 15162128 | 20543750 | 21792984 | 24347620 | 20815458 | 23132360 | 15798590 | 12425747 | 15525485 | 13213821 | 15969035 | 0 | 10 | 0 | 9 | 0 | 10 | 1.52 | 1.56 | 1.00 | 1.03 |
| O55013 | Trappc3 | Trafficking protein particle complex subunit 3 | 86.43 | 10.56 | 20302 | 2 | 3493005.75 | 2784441 | 2786345.75 | 3886372 | 2920875.5 | 2894540.25 | 2758802.75 | 3592779 | 2750826.5 | 4583073 | 2419999.25 | 2237663 | 2146241.5 | 2299636 | 2114581.75 | 2843326 | 4144808.5 | 4038111 | 4367495.5 | 4002949.5 | 0 | 3 | 0 | 7 | 0 | 7 | 0.71 | 0.68 | 1.22 | 1.17 |
| Q99NH8 | Trem2 | Triggering receptor expressed on myeloid cells 2 | 194.24 | 16.3 | 24527 | 3 | 1560126.88 | 2236838.75 | 1913290.5 | 2410745.75 | 2005547.38 | 2210698.5 | 1396864.62 | 1849918.25 | 1662521.75 | 2426231 | 3481204.5 | 3109150.75 | 3310466.5 | 3836597.75 | 3688794 | 1869784.38 | 2769195.75 | 3110672.5 | 2797246.25 | 3466022.25 | 0 | 4 | 0 | 2 | 0 | 4 | 1.72 | 1.83 | 1.38 | 1.47 |
| Q8BVW3 | Trim14 | Tripartite motif-containing protein 14 | 181.65 | 12.05 | 49641 | 4 | 2 | 499605.31 | 462687.38 | 511045.81 | 413822.38 | 2 | 473778 | 2 | 2 | 606742.88 | 1495364 | 1014437.19 | 1337591.75 | 687667.12 | 1389748.5 | 355876.75 | 2 | 2 | 625252.5 | 329150.47 | 0 | 8 | 0 | 11 | 0 | 11 | 3.14 | 5.48 | 0.69 | 1.21 |
| A2RSY6 | Trmt1l | TRMT1-like protein | 179.67 | 5.49 | 80861 | 3 | 2479836.25 | 2302663.75 | 2263770.25 | 2397787 | 1996628.25 | 2416480.75 | 2270919.75 | 2213206.25 | 1983192.5 | 2285049 | 3340707 | 3070068 | 3101322.25 | 3324588.5 | 3073118.5 | 2135581.75 | 2575828.25 | 2332297.75 | 2914780 | 764852.69 | 0 | 11 | 0 | 10 | 0 | 11 | 1.39 | 1.42 | 0.94 | 0.96 |
| Q62348 | Tsn | Translin | 525.42 | 33.77 | 26201 | 8 | 19076874 | 20370690 | 26537616 | 23180734 | 25773350 | 17717720 | 18627812 | 23739586 | 23126020 | 22124158 | 12172159 | 14707650 | 14121185 | 11947830 | 12807245 | 18820460 | 17992070 | 24211582 | 20048980 | 21886516 | 0 | 6 | 0 | 12 | 0 | 12 | 0.57 | 0.62 | 0.90 | 0.98 |
| P50637 | Tspo | Translocator protein | 278.33 | 40.24 | 18841 | 4 | 4287615.5 | 6739886.5 | 5024804 | 7337101 | 7697459 | 4806849.5 | 7448837 | 13223339 | 11964723 | 15505280 | 21132006 | 21069352 | 39585806 | 37803186 | 40052866 | 6480883.5 | 7766612.5 | 19523136 | 17983276 | 18807326 | 0 | 3 | 0 | 7 | 0 | 7 | 5.14 | 3.02 | 2.27 | 1.33 |
| P68373 | Tuba1c | Tubulin alpha-1C chain | 2305.94 | 80.18 | 49909 | 32 | 2 | 2 | 413527.72 | 2 | 2 | 2 | 2 | 591094.25 | 416624.31 | 2 | 1484944.88 | 997684.88 | 1194934.12 | 1258070.38 | 1448321.5 | 2 | 2 | 805727.44 | 2 | 2 | 0 | 12 | 0 | 11 | 0 | 12 | 15.44 | 6.34 | 1.95 | 0.80 |
| A2AQ07 | Tubb1 | Tubulin beta-1 chain | 463.68 | 8.43 | 50441 | 8 | 2 | 2 | 2 | 2 | 2 | 2 | 2 | 233981.84 | 182717.28 | 2 | 573589 | 664710.19 | 1315254.12 | 905927.88 | 1055717.12 | 2 | 2 | 122445.7 | 143303.28 | 2 | 0 | 7.5 | 0 | 10.5 | 0 | 10.5 | 451519.83 | 10.84 | 26575.50 | 0.64 |
| P68372 | Tubb4b | Tubulin beta-4B chain | 2291.51 | 68.09 | 49831 | 33 | 11016610 | 11014618 | 12763672 | 12201752 | 11396399 | 10344409 | 10515986 | 10820189 | 9957500 | 11114736 | 3197898 | 7296630 | 6887597.5 | 1426209.38 | 7677148 | 12744088 | 13478899 | 11215009 | 12671318 | 10832722 | 0 | 10 | 0 | 1 | 0 | 10 | 0.45 | 0.50 | 1.04 | 1.16 |
| P69897 | Tubb5 | Tubulin beta-5 chain | 2527.68 | 68.92 | 49671 | 35 | 97629258 | 93015314 | 120647458 | 80243370 | 117016386 | 91340050 | 105993826 | 99439618 | 101362994 | 90368986 | 43552702 | 74479610 | 70365914 | 67344922 | 53489874 | 103073642 | 55799598 | 65936114 | 70952834 | 77053466 | 0 | 3 | 0 | 4 | 0 | 4 | 0.61 | 0.63 | 0.73 | 0.76 |
| Q922F4 | Tubb6 | Tubulin beta-6 chain | 1617.11 | 46.31 | 50090 | 24 | 19082296 | 15870838 | 18061344 | 19007570 | 16211167 | 17591572 | 17070202 | 16608073 | 16639332 | 16997958 | 10307702 | 11359461 | 11204661 | 10334042 | 10674064 | 18231686 | 14192812 | 16495737 | 16328222 | 15975744 | 0 | 8 | 0 | 5 | 0 | 8 | 0.61 | 0.63 | 0.92 | 0.96 |
| Q9Z0P5 | Twf2 | Twinfilin-2 | 344.31 | 19.2 | 39471 | 5 | 8568626 | 7160764 | 6939692 | 8671824 | 7342823.5 | 8324899 | 7461092 | 5954537 | 6013728.5 | 6596521 | 3837115.5 | 2801972.5 | 3591545.25 | 1825475.5 | 3401412 | 8693229 | 6357085 | 6178413 | 6215489.5 | 6022162 | 0 | 5 | 0 | 12 | 0 | 12 | 0.40 | 0.45 | 0.87 | 0.97 |
| Q9CQM5 | Txndc17 | Thioredoxin domain-containing protein 17 | 303.58 | 42.28 | 14015 | 5 | 14281974 | 11760762 | 12871074 | 13277621 | 16649070 | 14787084 | 10781177 | 14383235 | 13442847 | 12984878 | 6089121.5 | 7178804 | 6151846 | 5731685.5 | 6575796 | 12205070 | 12275872 | 13486504 | 14374301 | 12262969 | 0 | 10 | 0 | 9 | 0 | 10 | 0.46 | 0.48 | 0.94 | 0.97 |
| Q9JMH6 | Txnrd1 | Thioredoxin reductase 1, cytoplasmic | 1007.85 | 35.73 | 67084 | 14 | 19807386 | 24552198 | 22245692 | 23546662 | 22307386 | 24420478 | 23366882 | 22487040 | 24123004 | 25967258 | 37900402 | 37285750 | 35741662 | 40957738 | 37186502 | 24680212 | 24894844 | 32046844 | 31508106 | 33738122 | 0 | 0 | 0 | 2 | 0 | 2 | 1.68 | 1.57 | 1.31 | 1.22 |
| Q9JJZ4 | Ube2j1 | Ubiquitin-conjugating enzyme E2 J1 | 147.32 | 11.01 | 34990 | 3 | 2985318 | 3073730.75 | 2877738.25 | 3071225.75 | 4172331 | 3066257 | 3194095.25 | 1965920.38 | 2947142 | 2791589 | 862471.62 | 1818722.25 | 1042621.94 | 1544912 | 1359086.75 | 2959714 | 3065397.5 | 2512270 | 2762480.25 | 3457661.5 | 0 | 7 | 0 | 12 | 0 | 12 | 0.41 | 0.47 | 0.91 | 1.06 |
| P68037 | Ube2l3 | Ubiquitin-conjugating enzyme E2 L3 | 340.42 | 62.34 | 17862 | 6 | 13169140 | 14885278 | 14486531 | 15483211 | 13772558 | 13631515 | 14023117 | 17290274 | 16760224 | 16935024 | 11699139 | 12609283 | 12211300 | 10137944 | 13039888 | 14334794 | 15800081 | 15732680 | 17766614 | 17858352 | 0 | 3 | 0 | 9 | 0 | 9 | 0.83 | 0.76 | 1.14 | 1.04 |
| P61082 | Ube2m | NEDD8-conjugating enzyme Ubc12 | 312.56 | 37.16 | 20900 | 7 | 14929850 | 15979026 | 16769130 | 19614070 | 17308140 | 14770654 | 15003998 | 16251912 | 17159102 | 17832462 | 7851057 | 10764796 | 7626575 | 12322930 | 9811398 | 15846132 | 18000384 | 17916920 | 18118710 | 18954794 | 0 | 8 | 0 | 3 | 0 | 8 | 0.57 | 0.60 | 1.05 | 1.10 |
| Q9CZY3 | Ube2v1 | Ubiquitin-conjugating enzyme E2 variant 1 | 297.36 | 29.25 | 16355 | 4 | 3402311.75 | 4655651.5 | 3767725 | 4151930.5 | 3476824.25 | 3407446 | 3629556.75 | 3424093.75 | 3444209.25 | 3567288.25 | 2328740.5 | 2398093 | 2677703.75 | 3220009.5 | 2655091.5 | 3431545.5 | 2835464 | 3835946.75 | 3257173.75 | 4007043 | 0 | 7 | 0 | 12 | 0 | 12 | 0.68 | 0.76 | 0.89 | 0.99 |
| Q9EPV8 | Ubl5 | Ubiquitin-like protein 5 | 46.08 | 12.33 | 8547 | 1 | 3163037 | 3515567.75 | 4237460.5 | 4350828 | 4281769.5 | 2683395.5 | 3748777.25 | 4645939.5 | 3309751 | 3864105.5 | 959313.31 | 1632661.38 | 1687121.88 | 946106.69 | 1684074.25 | 3400760.25 | 2274436.25 | 2453956.75 | 2882438.5 | 2884175.5 | 0 | 1 | 0 | 4 | 0 | 4 | 0.35 | 0.38 | 0.71 | 0.76 |
| Q8VCH8 | Ubxn4 | UBX domain-containing protein 4 | 229.49 | 12.45 | 56472 | 5 | 13434748 | 10150237 | 13749887 | 18220232 | 11866540 | 11109355 | 9007712 | 8570850 | 7783915 | 10789661 | 2908340.75 | 411428.5 | 2228670.25 | 3016401.25 | 2391121.25 | 16625953 | 21541256 | 8390709 | 10751921 | 10248383 | 0 | 11 | 0 | 8 | 0 | 11 | 0.16 | 0.23 | 1.00 | 1.43 |
| Q09426 | Ugt8 | 2-hydroxyacylsphingosine 1-beta-galactosyltransferase | 42.84 | 1.29 | 61126 | 1 | 5231538.5 | 6088538.5 | 7125840.5 | 8207048.5 | 6179887.5 | 6730344 | 4833056.5 | 5598135 | 4795786 | 8426761 | 9113146 | 8816571 | 9385500 | 10784755 | 9414617 | 7063549 | 9171870 | 8665436 | 8791171 | 9978919 | 0 | 2 | 0 | 1 | 0 | 2 | 1.45 | 1.56 | 1.33 | 1.44 |
| Q8VCW4 | Unc93b1 | Protein unc-93 homolog B1 | 352.79 | 11.04 | 66981 | 5 | 2478440 | 2992381.75 | 5455438 | 6291410 | 5758102.5 | 3708939.75 | 4685693.5 | 5436337.5 | 6490236 | 6201145.5 | 8317285.5 | 7943283.5 | 10902756 | 10745370 | 11697678 | 2913205.25 | 3965492.25 | 8254524.5 | 6852612.5 | 6433489.5 | 0 | 7 | 0 | 10 | 0 | 10 | 2.16 | 1.87 | 1.24 | 1.07 |
| Q9EPU0 | Upf1 | Regulator of nonsense transcripts 1 | 631.26 | 12.54 | 123967 | 13 | 3240071.25 | 4802541 | 4240917 | 3027405 | 3745973 | 3704145 | 2448105.25 | 3390813.5 | 3882387.25 | 3364117.5 | 14621800 | 10192607 | 10034338 | 8584375 | 10402870 | 1926275.38 | 2527963.5 | 4716080.5 | 4097874.25 | 4224170 | 0 | 10 | 0 | 9 | 0 | 10 | 2.83 | 3.21 | 0.92 | 1.04 |
| Q9CZ13 | Uqcrc1 | Cytochrome b-c1 complex subunit 1, mitochondrial | 808.89 | 37.5 | 52852 | 15 | 22230196 | 21688912 | 22093690 | 22553350 | 16763938 | 20451570 | 20826100 | 27409688 | 21874630 | 24165412 | 33487584 | 36614250 | 34903210 | 35761834 | 33505876 | 22533500 | 15514598 | 23883202 | 23208680 | 22255374 | 0 | 7 | 0 | 12 | 0 | 12 | 1.65 | 1.52 | 1.02 | 0.94 |
| Q9DB77 | Uqcrc2 | Cytochrome b-c1 complex subunit 2, mitochondrial | 1419.99 | 60.04 | 48235 | 22 | 36867706 | 35965642 | 41686002 | 36415238 | 41294194 | 36541434 | 37479622 | 38270654 | 36515050 | 42680914 | 57217626 | 57827102 | 63116582 | 68379978 | 71991418 | 33909246 | 32613004 | 46429490 | 39629014 | 44407670 | 0 | 12 | 0 | 11 | 0 | 12 | 1.66 | 1.66 | 1.02 | 1.03 |
| Q9CR68 | Uqcrfs1 | Cytochrome b-c1 complex subunit Rieske, mitochondrial | 567.31 | 50 | 29368 | 10 | 11001169 | 14564724 | 15278864 | 14145642 | 11592536 | 12613836 | 11332484 | 14155510 | 12390126 | 12673427 | 25456502 | 21151276 | 25195038 | 25890698 | 24571972 | 11126492 | 14104117 | 16107096 | 14976332 | 13993696 | 0 | 11 | 0 | 7 | 0 | 11 | 1.84 | 1.94 | 1.06 | 1.11 |
| Q9JMA1 | Usp14 | Ubiquitin carboxyl-terminal hydrolase 14 | 998.19 | 32.45 | 56002 | 18 | 36570422 | 38229974 | 34679658 | 42290310 | 37123266 | 40248026 | 38315354 | 35018138 | 38872202 | 36984262 | 30136378 | 30304518 | 30525440 | 27126072 | 29537430 | 40099274 | 33367412 | 40063774 | 41734154 | 40891718 | 0 | 9 | 0 | 7 | 0 | 9 | 0.78 | 0.78 | 1.04 | 1.04 |
| P56399 | Usp5 | Ubiquitin carboxyl-terminal hydrolase 5 | 1449.79 | 36.95 | 95833 | 25 | 51957214 | 52525038 | 51150610 | 55603746 | 59788110 | 46814170 | 51362414 | 49778382 | 54167258 | 51825414 | 35246270 | 38502378 | 35917062 | 28440520 | 33893134 | 54139702 | 48857490 | 48671754 | 49216422 | 49368970 | 0 | 3 | 0 | 8 | 0 | 8 | 0.63 | 0.68 | 0.92 | 0.99 |
| P70280 | Vamp7 | Vesicle-associated membrane protein 7 | 187.64 | 19.09 | 24967 | 4 | 1635746.62 | 2140641 | 329281.72 | 2114918.75 | 1654199.5 | 2262587.75 | 1984663.75 | 1659870.88 | 1339635.5 | 1621420.62 | 6744768 | 3070343 | 3539290 | 3272297.25 | 3552770.5 | 1372447.25 | 3000810.25 | 1714071.88 | 2 | 2204767.25 | 0 | 11 | 0 | 12 | 0 | 12 | 2.56 | 2.28 | 1.05 | 0.94 |
| Q9CZT5 | Vasn | Vasorin | 54.58 | 1.04 | 72261 | 1 | 1015842.25 | 1813874.88 | 637234.94 | 751756.94 | 507963.62 | 1474410.5 | 1170301.62 | 982576.12 | 1059798.62 | 1505575.5 | 13321597 | 10826836 | 10036818 | 9256282 | 7948387 | 1646626 | 1686999.25 | 2187867.25 | 2246833.25 | 1851863.12 | 0 | 2 | 0 | 0 | 0 | 2 | 10.87 | 8.30 | 2.04 | 1.55 |
| Q62465 | Vat1 | Synaptic vesicle membrane protein VAT-1 homolog | 685.73 | 30.05 | 43097 | 10 | 29689820 | 26953332 | 21245174 | 25236126 | 19650526 | 30773062 | 26496514 | 20837246 | 20235510 | 25918066 | 14295416 | 12976245 | 10520504 | 11816617 | 11204814 | 28919982 | 29476540 | 30522274 | 26522550 | 27099740 | 0 | 5 | 0 | 5 | 0 | 5 | 0.50 | 0.49 | 1.16 | 1.15 |
| P85972 | Vcl | Vinculin | 1078.48 | 21.76 | 116615 | 19 | 28655644 | 27941326 | 20108586 | 22346392 | 16981428 | 28427046 | 24004922 | 13960240 | 15978833 | 18211654 | 9927106 | 13637514 | 9991975 | 10192090 | 8740704 | 28058218 | 29302430 | 21505680 | 19961664 | 22095792 | 0 | 11 | 0 | 7 | 0 | 11 | 0.45 | 0.52 | 1.04 | 1.20 |
| P46462 | Vcp | Transitional endoplasmic reticulum ATPase | 2688.03 | 50.62 | 89349 | 40 | 47629514 | 51799822 | 49372922 | 45948862 | 48753146 | 47381394 | 45716394 | 50656322 | 44724666 | 44804334 | 99144786 | 79581594 | 66948146 | 61632582 | 69792594 | 52829846 | 54300410 | 51747294 | 45710514 | 51466926 | 0 | 7 | 0 | 3 | 0 | 7 | 1.55 | 1.62 | 1.05 | 1.10 |
| Q9Z2L0 | Vdac1 | Voltage-dependent anion-selective channel protein 1 | 1450.83 | 81.98 | 30756 | 18 | 60167878 | 59722070 | 61659314 | 65666914 | 56339218 | 68231370 | 65049754 | 64415722 | 60508558 | 77020298 | 116506082 | 94395450 | 116189730 | 109699994 | 123360874 | 59242662 | 69538130 | 75454322 | 78370978 | 66042982 | 0 | 4 | 0 | 9 | 0 | 9 | 1.85 | 1.67 | 1.15 | 1.04 |
| Q8BX70 | Vps13c | Vacuolar protein sorting-associated protein 13C | 2107.49 | 13.85 | 420089 | 40 | 36710974 | 35834090 | 36290782 | 32823178 | 40141670 | 34702422 | 36884898 | 33890582 | 38300538 | 30373368 | 20762064 | 21228888 | 22420232 | 27711270 | 26053608 | 37540190 | 33643306 | 29878902 | 30991280 | 31981720 | 0 | 5 | 0 | 7 | 0 | 7 | 0.65 | 0.68 | 0.90 | 0.94 |
| O88384 | Vti1b | Vesicle transport through interaction with t-SNAREs homolog 1B | 77.71 | 8.19 | 26713 | 2 | 1063271 | 465131.25 | 491723.66 | 534842.75 | 510579.44 | 1276905.25 | 1018398.62 | 1026685.81 | 811042.06 | 1325304.25 | 2034035.75 | 1802559.12 | 2075501.62 | 2125032 | 1784635 | 491219.19 | 983944.75 | 1440267.5 | 1229325 | 1229025 | 0 | 5 | 0 | 12 | 0 | 12 | 3.20 | 1.80 | 1.75 | 0.98 |
| Q99KC8 | Vwa5a | von Willebrand factor A domain-containing protein 5A | 1672.17 | 37.45 | 87143 | 26 | 97203218 | 98990634 | 115614418 | 107912442 | 115783762 | 82159258 | 97843546 | 93395466 | 108455882 | 92003738 | 35362374 | 44539886 | 46947962 | 35819954 | 48290238 | 85062826 | 67895530 | 79923730 | 77929010 | 78708538 | 0 | 0 | 0 | 1 | 0 | 1 | 0.39 | 0.45 | 0.73 | 0.82 |
| P32921 | Wars | Tryptophan--tRNA ligase, cytoplasmic | 554.26 | 23.08 | 54358 | 9 | 9464140 | 13007876 | 8930346 | 8952283 | 7959832 | 10295216 | 11647668 | 9195988 | 8908527 | 9800459 | 17773804 | 18824938 | 17222980 | 15497672 | 17019038 | 11225826 | 11062462 | 11349393 | 10603952 | 8473847 | 0 | 8 | 0 | 9 | 0 | 9 | 1.79 | 1.73 | 1.09 | 1.06 |
| P97765 | Wbp2 | WW domain-binding protein 2 | 82.39 | 7.66 | 28032 | 2 | 587538.88 | 793644.12 | 712062.56 | 667332.75 | 692643.88 | 736450 | 2 | 707253.81 | 561766.69 | 874039.62 | 1833610.88 | 1718290.12 | 1684907.88 | 1697151.62 | 2362006 | 699766.19 | 1331791.75 | 816563.44 | 782835.56 | 982002.06 | 0 | 3 | 0 | 5 | 0 | 5 | 2.69 | 3.23 | 1.34 | 1.60 |

| | | | | | | | | | | | | | | | | | | | | | | | | | | | | | | | | | | | | |
|---|---|---|---|---|---|---|---|---|---|---|---|---|---|---|---|---|---|---|---|---|---|---|---|---|---|---|---|---|---|---|---|---|---|---|---|---|
| O88342 | Wdr1 | WD repeat-containing protein 1 | 815.55 | 22.94 | 66407 | 13 | 94452602 | 103854826 | 97041266 | 111003442 | 88398122 | 87175874 | 82373018 | 74499474 | 83440130 | 77747098 | 69147250 | 63404738 | 66659022 | 66015102 | 63532254 | 82033626 | 88809962 | 89248370 | 92050570 | 88170490 | 0 | 3 | 0 | 3 | 0 | 3 | 0.66 | 0.81 | 0.89 | 1.09 |
| Q6DID7 | Wls | Protein wntless homolog | 137.34 | 5.36 | 62188 | 3 | 1521271.25 | 1041345.69 | 2173184.5 | 2269878.5 | 2298018 | 2850203 | 2633320.5 | 2671636.5 | 2565973.25 | 3009524 | 3656848 | 3487582.5 | 4777241.5 | 3867260.5 | 4124858 | 2482199.5 | 1893867.75 | 2426248 | 2567370.5 | 2613731 | 0 | 2 | 0 | 2 | 0 | 3 | 2.14 | 1.45 | 1.29 | 0.87 |
| Q91WL8 | Wwox | WW domain-containing oxidoreductase | 83.54 | 4.59 | 46512 | 2 | 2943334.75 | 3098931.5 | 3332114 | 3007047.5 | 2402386.75 | 3079514.25 | 2890151.75 | 2751383.5 | 2308709.25 | 3257819.5 | 4760212 | 3974356 | 5784928.5 | 4524703.5 | 4559092 | 2403216.5 | 2468684.25 | 2502699 | 1597683.5 | 2561968 | 0 | 8 | 0 | 8 | 0 | 8 | 1.60 | 1.65 | 0.92 | 0.95 |
| Q00519 | Xdh | Xanthine dehydrogenase/oxidase | 771.4 | 13.18 | 146562 | 14 | 20120926 | 20496500 | 13793951 | 17366834 | 14716912 | 13916289 | 20705110 | 15643754 | 16614903 | 14312318 | 4170935 | 8116236.5 | 7206664 | 7903994 | 6768019 | 20349710 | 15579121 | 14499611 | 15275404 | 16632358 | 0 | 11 | 0 | 11 | 0 | 11 | 0.40 | 0.42 | 0.95 | 1.01 |
| Q6PEC3 | Yif1b | Protein YIF1B | 77.34 | 5.02 | 28418 | 1 | 2 | 2 | 2 | 2 | 2 | 2 | 2 | 2 | 2 | 2 | 1490649.12 | 947074.12 | 1220666.75 | 1555737.12 | 1131963.12 | 2 | 2 | 2 | 2 | 2 | 0 | 1 | 0 | 1 | 0 | 1 | 634609.02 | 634609.02 | 135247.00 | 135247.00 |
| Q3UDR8 | Yipf3 | Protein YIPF3 | 41.55 | 6.63 | 37998 | 1 | 235145.56 | 256590.23 | 154755.05 | 148951.34 | 206023.95 | 2 | 175691.48 | 394423.16 | 517126.75 | 374200.06 | 1102692.12 | 565297.31 | 1522148.88 | 1258537.75 | 1425271.25 | 265192.12 | 410385.62 | 743035.5 | 1005383.94 | 418522.5 | 0 | 5 | 0 | 5 | 0 | 5 | 5.87 | 4.02 | 2.84 | 1.95 |
| P62259 | Ywhae | 14-3-3 protein epsilon | 1503.85 | 70.59 | 29174 | 24 | 161167762 | 150037426 | 183540978 | 168920994 | 197059554 | 131969858 | 149601826 | 167928322 | 165367250 | 164627426 | 100901802 | 110061482 | 95774914 | 90161330 | 107176594 | 168867858 | 160980210 | 165289586 | 185618498 | 158997074 | 0 | 10 | 0 | 8 | 0 | 10 | 0.59 | 0.65 | 0.98 | 1.08 |
| P61982 | Ywhag | 14-3-3 protein gamma | 965.61 | 59.51 | 28303 | 16 | 65808054 | 70518098 | 63453730 | 73393930 | 70788194 | 64014962 | 63607302 | 66275114 | 65476766 | 66460934 | 43439574 | 53476046 | 56987190 | 59511494 | 57989378 | 72701082 | 63393538 | 72227578 | 76992922 | 81781682 | 0 | 7 | 0 | 5 | 0 | 7 | 0.79 | 0.83 | 1.07 | 1.13 |
| P68510 | Ywhah | 14-3-3 protein eta | 1175.95 | 58.94 | 28212 | 20 | 77621730 | 82781970 | 98546138 | 94003298 | 118651634 | 64174706 | 89997306 | 98789842 | 109820050 | 97920210 | 31085374 | 53592170 | 50225894 | 42203482 | 54093454 | 80189410 | 46278910 | 76349874 | 79044778 | 97991506 | 0 | 5 | 0 | 6 | 0 | 6 | 0.49 | 0.50 | 0.81 | 0.82 |
| P68254 | Ywhaq | 14-3-3 protein theta | 1277.78 | 62.86 | 27778 | 20 | 30573510 | 38028930 | 31267838 | 29141440 | 38228498 | 24833428 | 31985984 | 34165622 | 50971466 | 25445520 | 17479840 | 20284504 | 15816199 | 13867849 | 15501930 | 26600342 | 17400452 | 20782536 | 19886744 | 22642332 | 0 | 2 | 0 | 2 | 0 | 2 | 0.50 | 0.50 | 0.64 | 0.64 |
| Q3TIV5 | Zc3h15 | Zinc finger CCCH domain-containing protein 15 | 124.96 | 6.34 | 48327 | 2 | 2 | 146090.77 | 227515.52 | 2 | 2 | 2 | 2 | 83032.48 | 132803.47 | 227055.36 | 319012.25 | 297827.34 | 327328.72 | 294162.66 | 368993.19 | 2 | 2 | 55652.3 | 2 | 118858.49 | 0 | 10.5 | 0 | 8 | 0 | 10.5 | 4.30 | 3.63 | 0.47 | 0.39 |
| Q9CYA6 | Zcchc8 | Zinc finger CCHC domain-containing protein 8 | 114.84 | 5.08 | 78026 | 2 | 862905.06 | 889528.19 | 894699.88 | 691171.38 | 809333.81 | 921941.69 | 1260015.25 | 757541.12 | 1049101.88 | 648034 | 173342.89 | 424998.12 | 361856.53 | 394171.31 | 257171.66 | 1176601.12 | 725557.19 | 816331.06 | 743430.62 | 776412.12 | 0 | 10 | 0 | 10 | 0 | 10 | 0.39 | 0.35 | 1.02 | 0.91 |
| Q9D270 | Zdhhc21 | Probable palmitoyltransferase ZDHHC21 | 88.5 | 6.04 | 31331 | 2 | 171134.08 | 183658.73 | 301324.25 | 320345.16 | 186390.03 | 204552.97 | 180098.97 | 433269.47 | 295106.06 | 361936.34 | 749243.38 | 651979.19 | 1083678.62 | 1446460 | 1089062.25 | 148355.22 | 157868.84 | 596876.38 | 496641.28 | 488547.06 | 0 | 10 | 0 | 10 | 0 | 10 | 4.32 | 3.40 | 1.62 | 1.28 |
| Q80W54 | Zmpste24 | CAAX prenyl protease 1 homolog | 228.03 | 13.68 | 54735 | 5 | 3193989.25 | 2831700.25 | 2784568.75 | 4624603.5 | 2780870.75 | 3525645 | 3710543.25 | 4315759 | 3886651.25 | 4662634.5 | 5134520.5 | 4776145 | 5731597.5 | 6055413 | 6247878.5 | 3827867.75 | 4479173.5 | 5618494 | 4844236 | 4593169.5 | 0 | 3 | 0 | 5 | 0 | 5 | 1.72 | 1.39 | 1.44 | 1.16 |
| O35986 | Zranb2 | Zinc finger Ran-binding domain-containing protein 2 | 133.33 | 8.18 | 37350 | 3 | 5654370 | 6570487.5 | 6087805 | 7326659 | 6541948.5 | 5675426.5 | 4949323.5 | 5546762 | 5706232 | 4704939 | 2412629.25 | 2120365.5 | 1935293.25 | 2163655 | 513532.25 | 5631302 | 5246629 | 5557038 | 5967596 | 6137981 | 0 | 3 | 0 | 7 | 0 | 7 | 0.28 | 0.34 | 0.89 | 1.07 |
| P11497 | Acaca | Acetyl-CoA carboxylase 1 | 960.71 | 8.61 | 265194 | 17 | 11324967 | 10542662 | 8519150 | 10532117 | 13300456 | 12657131 | 11578942 | 9833489 | 11738617 | 9746258 | 8657329 | 8002260.5 | 7694931 | 7734428.5 | 7352655 | 12502676 | 12914949 | 8288740 | 9242459 | 9698856 | 1 | 10 | 0 | 9 | 1 | 10 | 0.73 | 0.71 | 0.97 | 0.95 |
| Q9QYR9 | Acot2 | Acyl-coenzyme A thioesterase 2, mitochondrial | 369.48 | 22.3 | 49657 | 8 | 7293702 | 8286939 | 6672335 | 9951024 | 6500513 | 9967002 | 6756689 | 6678616 | 7720353.5 | 8637121 | 12033325 | 9562777 | 10658222 | 13313819 | 11936642 | 8458113 | 6291363 | 9791124 | 9576298 | 6315352.5 | 1 | 12 | 1 | 11 | 1 | 12 | 1.49 | 1.45 | 1.04 | 1.02 |
| Q7TPR4 | Actn1 | Alpha-actinin-1 | 1445.12 | 29.48 | 103068 | 24 | 16411922 | 15038022 | 9307904 | 10521349 | 10581996 | 11564938 | 13728450 | 9574411 | 9593676 | 7617270.5 | 4884120.5 | 8370109 | 4183680.5 | 3218683 | 4789386 | 14157026 | 9845708 | 5638205 | 10378459 | 9918144 | 0 | 6 | 1 | 11 | 1 | 11 | 0.41 | 0.49 | 0.81 | 0.96 |
| Q920A7 | Afg3l1 | AFG3-like protein 1 | 254.34 | 8.62 | 87047 | 5 | 3626907 | 4021630.5 | 3526595.25 | 3658533.5 | 3200927.75 | 3270029 | 4235210 | 3223704.25 | 1750377.25 | 2905540.5 | 6135459 | 3967930.75 | 6268462.5 | 6547007.5 | 5385317 | 4012403 | 4594071.5 | 2750356 | 4289658 | 2185402.5 | 1 | 11 | 1 | 9 | 1 | 11 | 1.57 | 1.84 | 0.99 | 1.16 |
| B5DFN2 | Ahcyl1 | S-adenosylhomocysteine hydrolase-like protein 1 | 359.2 | 18.01 | 53753 | 8 | 2557811 | 2881662.75 | 3449470.5 | 3998142.75 | 3023553 | 3471000 | 2682895.25 | 3045911.25 | 3017800.5 | 3185049.25 | 2109430.25 | 1988876.12 | 2462204 | 2648915.75 | 2050080.38 | 2812777.75 | 3285509.75 | 3398386.25 | 4335576.5 | 3533494.5 | 1 | 9 | 0 | 6 | 1 | 9 | 0.71 | 0.73 | 1.09 | 1.13 |
| Q9DB25 | Alg5 | Dolichyl-phosphate beta-glucosyltransferase | 331.67 | 24.07 | 36791 | 6 | 2779985.5 | 2720061.5 | 3054323 | 3024555.25 | 4804590.5 | 3536512.75 | 2812052 | 3054136.5 | 2508318.75 | 4274237 | 4608453 | 5748151.5 | 6561112.5 | 5310270 | 5853950.5 | 5010623 | 4460241.5 | 5822724 | 4409222.5 | 3988750 | 1 | 3 | 0 | 1 | 1 | 3 | 1.71 | 1.74 | 1.45 | 1.46 |
| P48999 | Alox5 | Arachidonate 5-lipoxygenase | 348.54 | 10.09 | 77967 | 6 | 5211182 | 5592098.5 | 4024445 | 2274519 | 4344241.5 | 6652062.5 | 6064063.5 | 2567617.75 | 4628740 | 2405004 | 1702517.25 | 2301356 | 1716003.5 | 1800951.38 | 1540111.5 | 5600932 | 5980421.5 | 2033284.75 | 1795194.12 | 2009383 | 1 | 10 | 0 | 6 | 1 | 10 | 0.42 | 0.41 | 0.81 | 0.78 |
| Q8K298 | Anln | Anillin | 266.91 | 6.78 | 122794 | 5 | 2118728.75 | 1980347.5 | 2003021.25 | 1720598.62 | 1435279.5 | 2017354.75 | 1842732.25 | 1817433.5 | 1193087.75 | 1589734.5 | 940191.38 | 1243722.75 | 924029.62 | 315938.81 | 634309.62 | 2038471.75 | 1392028.75 | 467074.84 | 1302800.88 | 1285070.25 | 0 | 4 | 1 | 8 | 1 | 8 | 0.44 | 0.48 | 0.70 | 0.77 |
| Q9EST5 | Anp32b | Acidic leucine-rich nuclear phosphoprotein 32 family member B | 385.18 | 27.94 | 31079 | 8 | 39030546 | 36995526 | 36545950 | 25332370 | 39812518 | 27436470 | 42139894 | 33950414 | 34211506 | 21965270 | 11975588 | 22429962 | 10864926 | 7448038 | 14780276 | 34219746 | 17061458 | 17385138 | 21568990 | 21096050 | 0 | 1 | 1 | 4 | 1 | 4 | 0.38 | 0.42 | 0.63 | 0.70 |
| P22892 | Ap1g1 | AP-1 complex subunit gamma-1 | 556.66 | 16.67 | 91350 | 10 | 11817118 | 11262903 | 15879660 | 15824366 | 18750032 | 10234695 | 11808577 | 13903822 | 17361028 | 12827280 | 6541864 | 7704800 | 9013904 | 6820035 | 10284966 | 10553962 | 10304549 | 12437823 | 15157330 | 12707895 | 0 | 6 | 1 | 10 | 1 | 10 | 0.55 | 0.61 | 0.83 | 0.92 |
| Q3UIA2 | Arhgap17 | Rho GTPase-activating protein 17 | 459.8 | 13.71 | 92202 | 8 | 3759071.5 | 4572134.5 | 5820784 | 5900824 | 5833046.5 | 4571993 | 3755844 | 4929673.5 | 5082329.5 | 5510133 | 3673514.75 | 4110693.25 | 2436612.75 | 3373478.25 | 3229381.25 | 4536390 | 3661387 | 3899119.5 | 5247094.5 | 3997795.5 | 1 | 5 | 1 | 7 | 1 | 7 | 0.65 | 0.71 | 0.82 | 0.89 |
| Q61210 | Arhgef1 | Rho guanine nucleotide exchange factor 1 | 640.85 | 13.91 | 102805 | 10 | 3345640.5 | 4915514 | 4754303 | 4210843.5 | 4258991 | 3702200.5 | 3019504 | 3578559.75 | 5567374.5 | 2735761 | 5710911 | 7730284.5 | 5326105 | 6046377 | 6065336 | 2426557.75 | 4138239.75 | 4672820 | 3345446.25 | 2955711.5 | 0 | 4 | 1 | 11 | 1 | 11 | 1.44 | 1.66 | 0.82 | 0.94 |
| P29067 | Arrb2 | Beta-arrestin-2 | 500.44 | 20.49 | 46340 | 11 | 9895255 | 7191552 | 6662854.5 | 4177038.75 | 6848940.5 | 7203335 | 9796674 | 6731424 | 6953158.5 | 7675919 | 14506648 | 11737890 | 13602688 | 11407377 | 9561296 | 10253096 | 8429972 | 5677624.5 | 6115687.5 | 4089589.25 | 1 | 11 | 1 | 11 | 1 | 11 | 1.75 | 1.59 | 0.99 | 0.90 |
| E9PZJ8 | Ascc3 | Activating signal cointegrator 1 complex subunit 3 | 38.22 | 0.5 | 250557 | 1 | 490778.41 | 599253 | 390407.34 | 660616.94 | 526985.62 | 583743.81 | 527209.38 | 678654.75 | 508565.44 | 531420.56 | 2 | 2 | 334125.56 | 2 | 466349.5 | 578242.06 | 422132.5 | 618223 | 267297.72 | 488781.44 | 1 | 9 | 0 | 7 | 1 | 9 | 0.30 | 0.28 | 0.89 | 0.84 |
| Q61024 | Asns | Asparagine synthetase [glutamine-hydrolyzing] | 917.48 | 31.91 | 64283 | 16 | 21166886 | 22116378 | 17689506 | 21294528 | 23271476 | 20062274 | 24133718 | 22337614 | 28311134 | 22627548 | 24710390 | 38047358 | 38547094 | 28788770 | 33352748 | 30047160 | 26982828 | 26724758 | 28907746 | 27183986 | 0 | 3 | 1 | 3 | 1 | 3 | 1.55 | 1.39 | 1.33 | 1.19 |
| Q64518 | Atp2a3 | Sarcoplasmic/endoplasmic reticulum calcium ATPase 3 | 929.76 | 21.48 | 113638 | 15 | 2471695 | 3325290 | 6237073 | 6447011.5 | 5239079 | 2757748.5 | 3332570 | 7296791 | 5232757 | 4921322 | 2086031.88 | 1312184.12 | 2259472 | 2117376.5 | 2715179 | 2068664.12 | 1188284.25 | 3657723.5 | 4233920.5 | 3033257.25 | 1 | 5 | 0 | 5 | 1 | 5 | 0.44 | 0.45 | 0.60 | 0.60 |
| P15999 | Atp5f1a | ATP synthase subunit alpha, mitochondrial | 2944.79 | 63.47 | 59754 | 42 | 7766863.5 | 7417389.5 | 7649394 | 7779600.5 | 6668269.5 | 8189972 | 5242943 | 7285566.5 | 6824187 | 7559088 | 4754507.5 | 5047347.5 | 5240527.5 | 6344206.5 | 5071641.5 | 8549246 | 9591793 | 7875812.5 | 9648108 | 9336178 | 0 | 0 | 1 | 1 | 1 | 1 | 0.71 | 0.75 | 1.21 | 1.28 |
| Q9CQQ7 | Atp5pb | ATP synthase F(0) complex subunit B1, mitochondrial | 474.73 | 34.38 | 28949 | 8 | 11709389 | 14118780 | 11899932 | 12754266 | 9931300 | 16283180 | 10441256 | 14817111 | 8848232 | 16445706 | 16346674 | 16710861 | 19404772 | 18597346 | 19045068 | 16827610 | 18605342 | 17558612 | 19664778 | 11344490 | 0 | 4 | 1 | 3 | 1 | 4 | 1.49 | 1.35 | 1.39 | 1.26 |
| Q8BVE3 | Atp6v1h | V-type proton ATPase subunit H | 637.89 | 25.88 | 55855 | 12 | 7183891 | 10154310 | 8809174 | 7921626 | 12365190 | 7347120 | 7116445.5 | 11549696 | 11255460 | 8644788 | 13085035 | 14729671 | 13944304 | 11467183 | 14185712 | 6137229 | 4406123 | 7390519 | 11056899 | 8463675 | 1 | 7 | 1 | 7 | 1 | 7 | 1.45 | 1.47 | 0.81 | 0.82 |
| Q91XV3 | Basp1 | Brain acid soluble protein 1 | 1206.97 | 83.19 | 22087 | 15 | 221095810 | 166081298 | 114296738 | 193591650 | 127235106 | 211086002 | 164883106 | 155059042 | 127267466 | 223221042 | 85773626 | 65896242 | 86146162 | 118788858 | 88783970 | 192481586 | 293953730 | 165916290 | 160184034 | 180322370 | 1 | 10 | 0 | 9 | 1 | 10 | 0.54 | 0.51 | 1.21 | 1.13 |
| O35855 | Bcat2 | Branched-chain-amino-acid aminotransferase, mitochondrial | 340.87 | 15.01 | 44127 | 5 | 20837462 | 20085942 | 18106018 | 23489116 | 17680850 | 22667158 | 19699926 | 20346566 | 18642548 | 19331194 | 13677339 | 13989961 | 17698942 | 17427866 | 17293580 | 16663638 | 19558416 | 19213962 | 22115774 | 21004682 | 1 | 12 | 0 | 11 | 1 | 12 | 0.80 | 0.80 | 0.98 | 0.98 |
| P59017 | Bcl2l13 | Bcl-2-like protein 13 | 90.48 | 9.45 | 46719 | 2 | 822995.94 | 851050.56 | 915134.25 | 775541.25 | 949020.94 | 363038 | 779211.5 | 782276.56 | 1813582.88 | 432619.81 | 391183.88 | 197470.09 | 305857.62 | 296963.59 | 326555.19 | 664107.94 | 484148.91 | 428570.44 | 762784 | 497649.94 | 0 | 1 | 1 | 9 | 1 | 9 | 0.35 | 0.36 | 0.66 | 0.68 |
| Q566Q8 | Bles03 | UPF0696 protein C11orf68 homolog | 95.59 | 7.57 | 27493 | 2 | 4445206 | 4077209.5 | 3795122.25 | 2788341.5 | 4359151 | 1993332 | 3673701 | 3701075 | 4043024 | 3071245.25 | 636979.25 | 2494373.5 | 1510295.88 | 533081.88 | 1955884 | 4284398 | 923915.56 | 1870041.25 | 1390528.62 | 1006317.06 | 0 | 3 | 1 | 5 | 1 | 5 | 0.37 | 0.43 | 0.49 | 0.57 |
| Q91VR8 | Brk1 | Protein BRICK1 | 88.76 | 25.33 | 8761 | 2 | 2839962.75 | 2790175.5 | 3936429.75 | 3163345.5 | 4452210 | 2477315 | 3252593 | 3816226.75 | 4241510 | 3348926 | 1727652.25 | 2463406.75 | 2104739.75 | 2744413.25 | 2148803.5 | 2599547.5 | 2296945.25 | 3564559.5 | 3317075.75 | 3073138.25 | 0 | 8 | 1 | 7 | 1 | 8 | 0.65 | 0.65 | 0.86 | 0.87 |
| Q9CXW3 | Cacybp | Calcyclin-binding protein | 572.84 | 45.41 | 26510 | 10 | 30273778 | 31879846 | 32453310 | 24680994 | 30928280 | 25114858 | 31127102 | 28016542 | 29031882 | 25933426 | 16201527 | 25225594 | 17202650 | 15691935 | 14723398 | 30705798 | 21281410 | 23241872 | 22392810 | 24945556 | 1 | 3 | 1 | 4 | 1 | 4 | 0.59 | 0.64 | 0.82 | 0.88 |
| Q6ZQ38 | Cand1 | Cullin-associated NEDD8-dissociated protein 1 | 1054.47 | 20.65 | 136332 | 18 | 28769238 | 26692090 | 30039876 | 32621932 | 32496030 | 26715000 | 27008500 | 31462830 | 33765678 | 30072296 | 20132430 | 22119792 | 26975474 | 21094742 | 23364316 | 25948690 | 21893548 | 30268794 | 28521350 | 30111600 | 1 | 7 | 1 | 8 | 1 | 8 | 0.75 | 0.76 | 0.91 | 0.92 |
| O89110 | Casp8 | Caspase-8 | 238.4 | 9.79 | 55357 | 4 | 6704277 | 7112024.5 | 7250546.5 | 7527506.5 | 7648271.5 | 6182639.5 | 7790562 | 7713102 | 7300013.5 | 7314465.5 | 3254475.25 | 5075877 | 5196326 | 1477953.38 | 6546232.5 | 8036769 | 4251604 | 6537527.5 | 5975522.5 | 7179114.5 | 0 | 7 | 1 | 5 | 1 | 7 | 0.59 | 0.59 | 0.88 | 0.88 |
| Q8VDP4 | Ccar2 | Cell cycle and apoptosis regulator protein 2 | 250.74 | 8.24 | 103002 | 5 | 2323680 | 2261175.25 | 2173934 | 2850118 | 2670660 | 2255687.75 | 2361709.25 | 1950865.88 | 2554406.75 | 1564996.75 | 462239.41 | 1101663 | 1647431.5 | 179453.59 | 1230882.38 | 2058792.12 | 1810425.75 | 1317677.25 | 1962819.5 | 2876560.5 | 0 | 5 | 1 | 10 | 1 | 10 | 0.38 | 0.43 | 0.82 | 0.94 |
| Q9WU84 | Ccs | Copper chaperone for superoxide dismutase | 122.34 | 8.03 | 28912 | 2 | 3169196 | 3745303.75 | 3702982.5 | 4573268 | 4706114 | 3486547.25 | 3555463.5 | 2670772.25 | 3659285.5 | 5210381 | 1896295.75 | 2059991.38 | 2280457.25 | 2863105.25 | 2 | 3474351 | 2367599.5 | 3356016.5 | 3420140.5 | 3407005.25 | 0 | 4 | 1 | 4 | 1 | 4 | 0.46 | 0.49 | 0.81 | 0.86 |
| P80313 | Cct7 | T-complex protein 1 subunit eta | 2304.58 | 64.71 | 59652 | 33 | 100202282 | 113481114 | 115497986 | 124026946 | 103497242 | 112411538 | 100522530 | 111683706 | 99555314 | 115008602 | 132479162 | 117021098 | 124916234 | 129594778 | 130966026 | 121111114 | 131870674 | 130732914 | 133505162 | 128493874 | 1 | 1 | 0 | 0 | 1 | 1 | 1.14 | 1.18 | 1.16 | 1.20 |
| P40240 | Cd9 | CD9 antigen | 140.35 | 16.37 | 25258 | 2 | 2 | 2 | 542973.62 | 354011.75 | 1492847.75 | 2 | 590755.38 | 1081013.38 | 1219159.5 | 1147816 | 1230283 | 1786222.75 | 2162968 | 1529887.12 | 1717307.25 | 2 | 2 | 1564702.5 | 1345326.38 | 1363522.12 | 1 | 10 | 0 | 9 | 1 | 10 | 3.53 | 2.09 | 1.79 | 1.06 |
| Q9CY57 | Chtop | Chromatin target of PRMT1 protein | 118.72 | 11.65 | 26585 | 2 | 405960.84 | 336235.22 | 401906.53 | 2 | 410551.94 | 1284919.5 | 575524.31 | 2 | 448337 | 889128.62 | 2047352.25 | 1713699.38 | 1087901 | 2159011.75 | 1529772.12 | 416941.78 | 433482.81 | 136987.75 | 885272.38 | 1113177.38 | 0 | 4 | 1 | 10 | 1 | 10 | 5.49 | 2.67 | 1.92 | 0.93 |
| Q8WTY4 | Ciapin1 | Anamorsin | 195.18 | 12.94 | 33429 | 3 | 3957603.75 | 4255422 | 4180778.25 | 4184645.5 | 4409513 | 4698365 | 4326254 | 3408517.5 | 2922341 | 3600711.5 | 2738113.5 | 3124015 | 2138176.5 | 2643184.75 | 2345807.5 | 4416690.5 | 4350587.5 | 3662743.5 | 4235965.5 | 4193105 | 0 | 10 | 1 | 8 | 1 | 10 | 0.62 | 0.69 | 0.99 | 1.10 |
| Q9R0Q8 | Clec4e | C-type lectin domain family 4 member E | 144.93 | 12.15 | 24431 | 2 | 2 | 2 | 293403.88 | 2 | 217834.89 | 2 | 454512.56 | 382112.5 | 434754.38 | 1242368.88 | 1635191.75 | 1565770.75 | 1157819 | 3619881.5 | 3175214 | 305161.75 | 376993.97 | 1157214 | 595455.56 | 1301800 | 0 | 1 | 0 | 10 | 0 | 10 | 21.82 | 4.44 | 7.31 | 1.49 |
| Q99KN9 | Clint1 | Clathrin interactor 1 | 253.82 | 6.66 | 68513 | 4 | 7828398 | 9154773 | 10557484 | 4486338 | 12086314 | 2794233.75 | 9493212 | 9684301 | 10927460 | 5199250.5 | 1080036.38 | 4311833 | 2031879.88 | 890859.81 | 2423061 | 8177334.5 | 2024662 | 2976371 | 2477946.5 | 5088433.5 | 0 | 3 | 1 | 4 | 1 | 4 | 0.24 | 0.28 | 0.47 | 0.54 |
| Q8K297 | Colgalt1 | Procollagen galactosyltransferase 1 | 492.62 | 16.21 | 71061 | 10 | 6316465.5 | 6631513.5 | 10341376 | 6919533.5 | 9945237 | 7080995 | 9312300 | 10230210 | 12401221 | 9881968 | 13057111 | 12143810 | 13827149 | 14545194 | 15242906 | 9368100 | 7388491 | 8904424 | 7925501 | 6700500.5 | 0 | 11 | 1 | 11 | 1 | 11 | 1.71 | 1.41 | 1.00 | 0.82 |
| O89001 | Cpd | Carboxypeptidase D | 302.47 | 4.28 | 152406 | 6 | 5757836 | 7323969 | 4113734.5 | 5498823 | 8541857 | 5424900.5 | 6386209 | 5074758 | 6348866.5 | 7666359.5 | 8072861 | 9626536 | 9687440 | 9924379 | 11156255 | 5105822.5 | 8358155 | 11883878 | 10465033 | 9911125 | 1 | 5 | 0 | 4 | 1 | 5 | 1.55 | 1.57 | 1.46 | 1.48 |
| Q9D0M3 | Cyc1 | Cytochrome c1, heme protein, mitochondrial | 354.02 | 43.69 | 35328 | 7 | 6370051 | 7750000 | 9304479 | 7627395.5 | 9136189 | 6764911 | 6593894.5 | 7329207.5 | 7615947 | 6479375 | 9955296 | 10793203 | 10228164 | 10125768 | 9202940 | 8406211 | 9031330 | 8105640.5 | 8375333 | 9660940 | 1 | 8 | 1 | 5 | 1 | 8 | 1.25 | 1.45 | 1.08 | 1.25 |
| P62897 | Cycs | Cytochrome c, somatic | 493.77 | 58.1 | 11605 | 8 | 49704282 | 48117066 | 35428586 | 43605522 | 38304610 | 52847270 | 47305654 | 41012570 | 36324342 | 43760318 | 26555042 | 30518744 | 33549436 | 35179258 | 37381330 | 48252870 | 48410638 | 46447130 | 47034958 | 55866758 | 1 | 6 | 1 | 6 | 1 | 6 | 0.76 | 0.74 | 1.14 | 1.11 |
| Q80Y44 | Ddx10 | Probable ATP-dependent RNA helicase DDX10 | 105.88 | 3.31 | 100739 | 2 | 708754.69 | 773512.06 | 647299.31 | 818686.25 | 189144.8 | 1016384.88 | 530037.88 | 184747.17 | 182454.56 | 773671.25 | 1213203.12 | 1159950.62 | 1546014.75 | 983782 | 1147964.88 | 611027.62 | 498868.31 | 701127.62 | 510031.47 | 748771 | 0 | 8 | 1 | 12 | 1 | 12 | 1.93 | 2.25 | 0.98 | 1.14 |
| Q9D0R4 | Ddx56 | Probable ATP-dependent RNA helicase DDX56 | 205.22 | 9.52 | 61212 | 4 | 1250722.88 | 1271492.12 | 1179409.88 | 2037641.12 | 1315141.38 | 635658.44 | 1485957.38 | 1181388 | 1514121.5 | 494334.31 | 2984934.25 | 1895738.25 | 2850929.5 | 2076176.5 | 2104065 | 1557556.5 | 894174.12 | 1319310.88 | 1196419.38 | 655754.88 | 1 | 10 | 0 | 10 | 1 | 10 | 1.69 | 2.24 | 0.80 | 1.06 |
| Q9DBB8 | Dhdh | Trans-1,2-dihydrobenzene-1,2-diol dehydrogenase | 115.3 | 10.21 | 36301 | 2 | 3280341.5 | 3367484.75 | 2658273 | 2130175.5 | 2155294.75 | 2473015.5 | 3352970 | 2550494.5 | 1653441.62 | 2335235 | 904253.62 | 2037383.5 | 1237699.25 | 468387.84 | 1401610.88 | 2927108 | 1230199.12 | 2163536 | 1689074.38 | 2085685.62 | 0 | 5 | 1 | 7 | 1 | 7 | 0.45 | 0.49 | 0.74 | 0.82 |
| Q9CXR1 | Dhrs7 | Dehydrogenase/reductase SDR family member 7 | 195.4 | 9.47 | 38167 | 3 | 191769.73 | 344904.69 | 2 | 291593.91 | 839271.38 | 2 | 478300.94 | 482029.91 | 1246240.25 | 2 | 1226951.62 | 1783041.88 | 1788327.62 | 1838871.62 | 1456250 | 2 | 160828.67 | 2 | 42885.94 | 2 | 0 | 3.5 | 1 | 7 | 1 | 7 | 4.85 | 3.67 | 0.12 | 0.09 |
| O70152 | Dpm1 | Dolichol-phosphate mannosyltransferase subunit 1 | 255.06 | 23.08 | 29175 | 4 | 1999628.25 | 211640.92 | 1527133.12 | 1569813.75 | 2789389 | 2682183 | 2383246.25 | 1961016.75 | 2116420.5 | 1911999 | 4913640 | 2999986 | 3349860.25 | 2796365.25 | 2384457.75 | 1853080.62 | 3279293.5 | 1983776.75 | 2296371 | 2368952.5 | 1 | 6 | 1 | 12 | 1 | 12 | 2.03 | 1.49 | 1.45 | 1.07 |
| O35474 | Edil3 | EGF-like repeat and discoidin I-like domain-containing protein 3 | 136.03 | 7.29 | 53712 | 3 | 278383.06 | 318849.03 | 60915.11 | 138268.36 | 208195.23 | 326443.91 | 413373.47 | 373618.25 | 1091285.5 | 2 | 966419.81 | 2406893 | 3626450.25 | 1772868 | 3352026 | 2 | 412693.19 | 383384.06 | 157010.42 | 2 | 0 | 12 | 1 | 8 | 1 | 12 | 12.07 | 5.50 | 0.95 | 0.43 |
| O08810 | Eftud2 | 116 kDa U5 small nuclear ribonucleoprotein component | 1077.28 | 23.58 | 109361 | 20 | 15342416 | 12522402 | 13872018 | 12258745 | 16827270 | 14835546 | 14414613 | 13162887 | 16330368 | 11214880 | 11009032 | 10212536 | 10092388 | 9350003 | 11614514 | 16261616 | 13698714 | 12518574 | 12985200 | 11538068 | 0 | 9 | 1 | 9 | 1 | 9 | 0.74 | 0.75 | 0.95 | 0.96 |
| Q61749 | Eif2b4 | Translation initiation factor eIF-2B subunit delta | 328.67 | 14.5 | 57624 | 6 | 3247852.5 | 5040503.5 | 5460648 | 4645250 | 4113787.5 | 3081380.25 | 2184176 | 5403548 | 5400072 | 6830617 | 7597456.5 | 5931220.5 | 8772362 | 9960244 | 9340261 | 2214191.25 | 2649621.75 | 5818426 | 7585296 | 6346404.5 | 0 | 10 | 1 | 10 | 1 | 10 | 1.85 | 1.82 | 1.09 | 1.07 |
| B0BNG0 | Emc2 | ER membrane protein complex subunit 2 | 215.4 | 17.51 | 34870 | 4 | 3389873.75 | 2336424 | 2317062.25 | 1303853.75 | 2490317.5 | 2473223.75 | 3715608.5 | 3923817.5 | 3470784.5 | 3128738.5 | 3839772.5 | 5214697 | 6114067.5 | 5526436 | 6913969.5 | 3083924.25 | 2043757.5 | 3501314.5 | 2826593 | 3243633.25 | 0 | 7 | 1 | 7 | 1 | 7 | 2.33 | 1.65 | 1.24 | 0.88 |
| Q5U2V8 | Emc3 | ER membrane protein complex subunit 3 | 308.13 | 27.2 | 29980 | 6 | 4587161 | 2964865.5 | 2797672.75 | 3408000 | 3217148.25 | 3201659.75 | 4357719.5 | 2878517.25 | 3319192.5 | 3109040 | 7005619 | 4582609.5 | 6556658 | 6691622.5 | 6354937.5 | 4056032.5 | 6490242.5 | 3744880.75 | 3916222 | 1642932.25 | 1 | 8 | 0 | 8 | 1 | 8 | 1.84 | 1.85 | 1.17 | 1.18 |
| O70378 | Emc8 | ER membrane protein complex subunit 8 | 291.31 | 26.09 | 23348 | 4 | 4785793 | 4529128.5 | 4848187.5 | 5255306 | 4241928 | 6027472 | 4414625.5 | 4325981.5 | 4057900.5 | 5482924 | 7291946.5 | 7445748 | 7710460 | 5920541 | 7935277.5 | 4050450.75 | 6183388 | 6369153 | 6755309.5 | 6744128 | 0 | 5 | 1 | 5 | 1 | 5 | 1.53 | 1.49 | 1.27 | 1.24 |
| Q8BTW3 | Exosc6 | Exosome complex component MTR3 | 111.5 | 11.36 | 28370 | 2 | 784200.94 | 2 | 2 | 1548933.25 | 1261206.75 | 702942.88 | 820891.69 | 2 | 1371478.38 | 1225940.5 | 1481039 | 2101587.5 | 1965669 | 1744897.25 | 2113186.5 | 2 | 2 | 1436280.75 | 1609791.75 | 1646646.12 | 1 | 9 | 0 | 9 | 1 | 9 | 2.62 | 2.28 | 1.31 | 1.14 |
| Q8VE88 | Fam114a2 | Protein FAM114A2 | 99.94 | 7.65 | 54045 | 2 | 1009979.75 | 1037351.75 | 815890.31 | 317230.69 | 1080864.5 | 548480.75 | 1123763.12 | 1016401.25 | 1097146.62 | 893140.25 | 259426.77 | 638725.88 | 219288.16 | 2 | 314136.81 | 1018743.62 | 279006.47 | 356725.5 | 347987.53 | 549989.75 | 1 | 6 | 1 | 4 | 1 | 6 | 0.34 | 0.31 | 0.60 | 0.55 |
| P22509 | Fbl | rRNA 2'-O-methyltransferase fibrillarin | 960.05 | 63.3 | 34222 | 16 | 21404730 | 27398548 | 25358232 | 28009130 | 22077558 | 24209648 | 22411264 | 24153646 | 22849104 | 21489198 | 27786556 | 32750108 | 30876884 | 35032110 | 30618022 | 26485854 | 27545930 | 28453998 | 27057136 | 31108998 | 1 | 5 | 0 | 5 | 1 | 5 | 1.26 | 1.36 | 1.13 | 1.22 |
| Q6P9Q6 | Fkbp15 | FK506-binding protein 15 | 44.07 | 1.56 | 132961 | 1 | 400036.34 | 678246.5 | 399313.22 | 477087.44 | 320379.12 | 396883.75 | 460948.53 | 609284.5 | 408377 | 478115.53 | 310378.62 | 282808.72 | 2 | 321161.12 | 2 | 516595.5 | 656064.25 | 454333.09 | 451608 | 512084.19 | 1 | 7 | 0 | 8 | 1 | 8 | 0.40 | 0.39 | 1.14 | 1.10 |
| P47856 | Gfpt1 | Glutamine--fructose-6-phosphate aminotransferase [isomerizing] 1 | 377.01 | 14.2 | 78539 | 8 | 6658558 | 7787280.5 | 6633611.5 | 6179008 | 6642843 | 7966783 | 8024434 | 8520079 | 7322791.5 | 7682186.5 | 8655480 | 9113173 | 8436498 | 8987235 | 8954507 | 6537959.5 | 6277290 | 9139792 | 7883323 | 5519488 | 0 | 12 | 1 | 7 | 1 | 12 | 1.30 | 1.12 | 1.04 | 0.89 |
| O88958 | Gnpda1 | Glucosamine-6-phosphate isomerase 1 | 210.4 | 16.96 | 32549 | 4 | 5967354 | 4723525 | 4245984 | 4904127 | 4249126.5 | 4199582.5 | 5208835.5 | 2965362 | 4408621 | 4468334.5 | 3674084.5 | 2324225.5 | 1999416 | 1554132.5 | 649077.31 | 6403557 | 7249584 | 4332663 | 4230696 | 4012467.5 | 0 | 12 | 1 | 10 | 1 | 12 | 0.42 | 0.48 | 1.09 | 1.23 |
| P28798 | Grn | Granulins | 877.83 | 35.65 | 63458 | 16 | 32750454 | 32672644 | 36089554 | 50561286 | 40879382 | 39148214 | 28006742 | 29346302 | 25807506 | 48105634 | 16646476 | 13557626 | 20974908 | 26247752 | 17241556 | 28665162 | 23522250 | 32188202 | 31137702 | 28323992 | 0 | 0 | 1 | 10 | 1 | 10 | 0.49 | 0.56 | 0.75 | 0.84 |
| Q9D8T2 | Gsdmdc1 | Gasdermin-D | 428.78 | 18.48 | 53238 | 7 | 11444688 | 13632972 | 9301002 | 10504442 | 13663193 | 7496365 | 11434229 | 11263532 | 13909362 | 11235216 | 6007067 | 9754973 | 7128110 | 5183546 | 7426006 | 11613425 | 5277176 | 8636320 | 7202401 | 9340203 | 1 | 4 | 1 | 6 | 1 | 6 | 0.61 | 0.64 | 0.72 | 0.76 |
| D2XV59 | Gtpbp1 | GTP-binding protein 1 | 116.55 | 3.74 | 72489 | 2 | 2191963 | 1774940.5 | 2265729.5 | 2107927 | 2051034 | 1923741.62 | 2187311.5 | 2250741.75 | 2014060.38 | 1786190.75 | 1310647.88 | 1776831.25 | 1574105.75 | 876033 | 1553477.38 | 2358591.5 | 1299925.75 | 1944763.5 | 2167051.25 | 1186078.25 | 1 | 9 | 0 | 10 | 1 | 10 | 0.68 | 0.70 | 0.86 | 0.88 |
| Q61191 | Hcfc1 | Host cell factor 1 | 527.19 | 8.7 | 210437 | 10 | 4379677.5 | 3908491.5 | 4112907.25 | 4237219.5 | 4337439 | 4437194 | 4425588.5 | 3861110 | 3521111 | 3897759 | 3397782.5 | 3646822.75 | 2666300.75 | 2677242.5 | 3413631.5 | 4909865 | 5375938.5 | 3680031 | 4844359 | 4383987.5 | 0 | 5 | 1 | 6 | 1 | 6 | 0.75 | 0.78 | 1.11 | 1.15 |
| Q3UMU9 | Hdgfl2 | Hepatoma-derived growth factor-related protein 2 | 161.69 | 4.63 | 74291 | 3 | 2783818.75 | 2820569.75 | 2045406.75 | 3762674.5 | 2677877.5 | 3293042.5 | 3003814 | 1115159.25 | 2285499.5 | 2884768.25 | 1798470 | 2 | 98147.8 | 245907.25 | 2 | 3688232.75 | 3108678.5 | 2508590.25 | 3406937.75 | 2337900 | 0 | 11 | 1 | 7 | 1 | 11 | 0.15 | 0.17 | 1.07 | 1.20 |
| Q3UGR5 | Hdhd2 | Haloacid dehalogenase-like hydrolase domain-containing protein 2 | 45.79 | 3.09 | 28730 | 1 | 739995.38 | 589639.62 | 547140.94 | 229147.39 | 1230991.38 | 361222.84 | 668331.75 | 648347.38 | 1228357.38 | 297468.06 | 2 | 267936.94 | 2 | 2 | 2 | 610764.56 | 94038.52 | 138035.58 | 2 | 258944.02 | 1 | 4 | 0 | 2 | 1 | 4 | 0.08 | 0.08 | 0.33 | 0.34 |
| P15864 | Hist1h1c | Histone H1.2 | 875.89 | 44.81 | 21267 | 13 | 32299800 | 25940726 | 50457262 | 44882992 | 29103660 | 44393618 | 33803166 | 41144130 | 24282954 | 48932190 | 18905598 | 14478212 | 20307532 | 28068560 | 18994380 | 22161572 | 38665734 | 34190898 | 39975618 | 34267218 | 1 | 12 | 1 | 8 | 1 | 12 | 0.55 | 0.52 | 0.93 | 0.88 |
| P43274 | Hist1h1e | Histone H1.4 | 815.51 | 37.44 | 21977 | 12 | 25638356 | 24667444 | 34378074 | 29586482 | 21601474 | 35624114 | 27939804 | 33471868 | 19815118 | 34123090 | 17638468 | 17636206 | 17817926 | 22469886 | 17673866 | 19342680 | 29234046 | 28164300 | 29372002 | 27790486 | 1 | 12 | 1 | 7 | 1 | 12 | 0.70 | 0.63 | 0.99 | 0.89 |
| O08528 | Hk2 | Hexokinase-2 | 2031.6 | 42.09 | 102535 | 34 | 29434372 | 33259694 | 19197308 | 18115562 | 22962250 | 38629534 | 35195810 | 28614788 | 28867730 | 23850294 | 47154190 | 52867698 | 41636078 | 37123538 | 50293458 | 42736986 | 35367162 | 27197916 | 26433598 | 30796050 | 0 | 5 | 1 | 11 | 1 | 11 | 1.86 | 1.48 | 1.32 | 1.05 |
| Q8CFE2 | Hpf1 | Histone PARylation factor 1 | 127.41 | 7.8 | 39291 | 2 | 1726564.38 | 1639607.75 | 2618235 | 1101046.5 | 2210294.5 | 1541445.38 | 1812904.25 | 1984333.5 | 1991817.75 | 1452340.25 | 350053.06 | 695089.31 | 332974.34 | 1380569.75 | 706184.25 | 1305231.62 | 1432923 | 607230.62 | 684501.12 | 1680828.12 | 1 | 4 | 0 | 2 | 1 | 4 | 0.37 | 0.39 | 0.61 | 0.65 |
| Q64433 | Hspe1 | 10 kDa heat shock protein, mitochondrial | 474.69 | 67.65 | 10963 | 7 | 46814846 | 31449030 | 30110614 | 41931030 | 25769508 | 46977722 | 42066230 | 36473850 | 31469470 | 44223418 | 50963494 | 44858726 | 48755798 | 56888150 | 59396762 | 47254426 | 74640634 | 50619538 | 51834294 | 47250902 | 1 | 0 | 0 | 0 | 1 | 0 | 1.48 | 1.30 | 1.54 | 1.35 |
| Q61699 | Hsph1 | Heat shock protein 105 kDa | 770.61 | 20.05 | 96407 | 13 | 11991290 | 11422438 | 8341631.5 | 9525722 | 9428292 | 11394152 | 13075355 | 13311584 | 12758421 | 9872232 | 14364920 | 15689804 | 15478738 | 13199316 | 15050697 | 11865986 | 8940429 | 12852045 | 11250729 | 10716385 | 0 | 9 | 1 | 7 | 1 | 9 | 1.45 | 1.22 | 1.10 | 0.92 |
| Q9BDB7 | Ifi44l | Interferon-induced protein 44-like | 383.31 | 19.24 | 49680 | 7 | 15056296 | 14016147 | 7780047.5 | 17854734 | 17541170 | 15945078 | 14687775 | 14851980 | 15055606 | 15542335 | 6896270 | 7779831 | 9234666 | 6036042.5 | 7611214 | 14772263 | 14426800 | 10638963 | 12537147 | 14224228 | 1 | 8 | 0 | 1 | 1 | 8 | 0.52 | 0.49 | 0.92 | 0.88 |
| Q6I7R3 | Isoc1 | Isochorismatase domain-containing protein 1 | 236.84 | 19.87 | 32002 | 4 | 1541349 | 1976793.62 | 825156.31 | 1576510.5 | 1198735.88 | 748701 | 1935646.12 | 1564863.25 | 1686211.38 | 1955528.75 | 2335085.25 | 2379544.5 | 2759821.5 | 2884490 | 1971250 | 591498 | 928845.38 | 866484 | 650003.69 | 1589160.88 | 1 | 6 | 0 | 4 | 1 | 6 | 1.73 | 1.56 | 0.65 | 0.59 |
| P62482 | Kcnab2 | Voltage-gated potassium channel subunit beta-2 | 376.36 | 31.61 | 41021 | 7 | 9748467 | 10016642 | 7334363.5 | 7986253.5 | 8699198 | 10059875 | 10894465 | 8956674 | 8866016 | 10199262 | 12526044 | 12972302 | 11365093 | 10543452 | 11654158 | 10799396 | 11341114 | 10300543 | 10759586 | 9639148 | 0 | 6 | 1 | 6 | 1 | 6 | 1.35 | 1.21 | 1.21 | 1.08 |
| Q05CL8 | Larp7 | La-related protein 7 | 104.83 | 3.51 | 64802 | 2 | 2196744 | 1754356.25 | 2015552.12 | 2366926.75 | 1449956.5 | 2198811.25 | 1628422.75 | 1426479 | 1410519.5 | 1648539.25 | 1126642.62 | 1410522.88 | 1101205.12 | 1219446.25 | 1282347.88 | 1712274.25 | 1988598 | 1516400.88 | 1808161 | 1540384.5 | 0 | 7 | 1 | 9 | 1 | 9 | 0.63 | 0.74 | 0.88 | 1.03 |
| Q8BMJ2 | Lars | Leucine--tRNA ligase, cytoplasmic | 1553.93 | 23.01 | 134192 | 26 | 28805952 | 34325366 | 28222182 | 27971648 | 28316818 | 31343252 | 31594788 | 26374090 | 27425136 | 27230342 | 47512778 | 51184230 | 42921110 | 32073174 | 43096006 | 32615382 | 25850194 | 31438176 | 29499446 | 30783816 | 1 | 9 | 1 | 10 | 1 | 10 | 1.47 | 1.51 | 1.02 | 1.04 |
| O89017 | Lgmn | Legumain | 160.48 | 4.83 | 49373 | 2 | 1776030.75 | 2211846.5 | 1369836.5 | 189032.61 | 3228589.5 | 651365.44 | 1865417 | 1625744 | 2739611.5 | 703187.81 | 366197.44 | 2 | 2 | 2 | 2 | 1485656 | 616930.5 | 417006.81 | 317674.44 | 708792.06 | 1 | 6 | 0 | 4 | 1 | 6 | 0.04 | 0.05 | 0.40 | 0.47 |
| Q8CGK3 | Lonp1 | Lon protease homolog, mitochondrial | 1570.86 | 33.4 | 105843 | 28 | 44156310 | 49725922 | 35161510 | 38901070 | 39886910 | 42212314 | 41384254 | 37038146 | 43888258 | 44531918 | 47421566 | 55039978 | 60719670 | 60006042 | 58199370 | 44219094 | 39078658 | 48603442 | 51987834 | 52342594 | 1 | 5 | 0 | 5 | 1 | 5 | 1.35 | 1.35 | 1.14 | 1.13 |
| Q6ZWM4 | Lsm8 | U6 snRNA-associated Sm-like protein LSm8 | 312.6 | 57.29 | 10403 | 4 | 6349568.5 | 5542050 | 7352303 | 7652241 | 6416824 | 6346976.5 | 5074212 | 5597020 | 5463961 | 7083427.5 | 3680483 | 3276212.25 | 2656489 | 5096984 | 2873963.5 | 5121939 | 8497572 | 6933305 | 5948470 | 5959661.5 | 0 | 10 | 1 | 9 | 1 | 10 | 0.53 | 0.59 | 0.97 | 1.10 |
| P19973 | Lsp1 | Lymphocyte-specific protein 1 | 728.35 | 44.85 | 36714 | 11 | 36370706 | 43541574 | 41407422 | 45629378 | 17568022 | 41364274 | 34843542 | 35837878 | 27110236 | 42423594 | 17993328 | 16763562 | 9692979 | 12850372 | 13593377 | 34673566 | 45380594 | 32174402 | 29162176 | 28298390 | 1 | 8 | 0 | 9 | 1 | 9 | 0.38 | 0.39 | 0.92 | 0.93 |
| P24527 | Lta4h | Leukotriene A-4 hydrolase | 510.55 | 18.99 | 69051 | 9 | 12878679 | 13382945 | 10651675 | 11690592 | 9698045 | 13348327 | 13736008 | 14264084 | 13678302 | 10537676 | 9265850 | 10142074 | 6974557 | 5406179.5 | 6807797.5 | 14645636 | 13526471 | 10522030 | 11694178 | 12217612 | 1 | 8 | 0 | 9 | 1 | 9 | 0.66 | 0.59 | 1.07 | 0.95 |
| Q5SUF2 | Luc7l3 | Luc7-like protein 3 | 557.42 | 24.54 | 51450 | 8 | 3663755 | 6100642 | 9637632 | 6435240 | 7909449 | 3542740 | 7145931 | 6944723.5 | 7271001.5 | 5342010.5 | 1751229.25 | 2869263.75 | 3606497.5 | 2521425.75 | 2488817.25 | 6718200 | 5582930.5 | 5227468 | 5736927 | 5857631.5 | 0 | 7 | 1 | 9 | 1 | 9 | 0.39 | 0.44 | 0.86 | 0.96 |
| P27046 | Man2a1 | Alpha-mannosidase 2 | 257.01 | 4.78 | 131631 | 5 | 926179.12 | 989275.38 | 391163.22 | 747260.12 | 848456.81 | 1009570.12 | 369294 | 529731.12 | 1395621.75 | 348776.81 | 1054078.75 | 1628515.25 | 1965493.5 | 2551499 | 2764690.5 | 700605.5 | 467985.38 | 861396.31 | 605676.75 | 1073040.25 | 0 | 11 | 1 | 10 | 1 | 11 | 2.55 | 2.73 | 0.95 | 1.02 |
| P47811 | Mapk14 | Mitogen-activated protein kinase 14 | 319.61 | 20.28 | 41287 | 6 | 7234879.5 | 4803639 | 6947161 | 9730536 | 9242203 | 5154368 | 6547161 | 7376451 | 4228930.5 | 3840694.75 | 3118528.75 | 3276744 | 3732317.5 | 3999716.75 | 2870739.5 | 6544243 | 5396068 | 5669191 | 4723058.5 | 6532907.5 | 0 | 4 | 1 | 9 | 1 | 9 | 0.45 | 0.63 | 0.76 | 1.06 |
| Q922Q1 | Marc2 | Mitochondrial amidoxime reducing component 2 | 327.36 | 18.64 | 38194 | 6 | 8562592 | 8565058 | 8431714 | 7960052.5 | 8969418 | 9125462 | 8918437 | 6576704.5 | 8176594 | 9961843 | 10239514 | 9550035 | 13196220 | 10145089 | 13102534 | 7166568 | 6507231.5 | 7959597 | 8808870 | 8663130 | 0 | 8 | 1 | 6 | 1 | 8 | 1.32 | 1.32 | 0.92 | 0.91 |
| Q4G009 | Mcts1 | Malignant T-cell-amplified sequence 1 | 149.22 | 17.03 | 20550 | 3 | 291451.03 | 350097.59 | 561658 | 637270.38 | 736276.12 | 290433.38 | 394638.44 | 446927.62 | 715319.38 | 318958.09 | 274354.94 | 290652 | 274397.56 | 2 | 266392.16 | 811012 | 389487.69 | 360055.78 | 165291.84 | 593023.81 | 0 | 12 | 1 | 12 | 1 | 12 | 0.43 | 0.51 | 0.90 | 1.07 |
| O35678 | Mgll | Monoglyceride lipase | 46.75 | 2.31 | 33388 | 1 | 2254833.75 | 2698437.75 | 2329699.75 | 2347762 | 1876888 | 2274749 | 2475376 | 2519352 | 2221599.75 | 2425861 | 2768398.5 | 2977746.75 | 2727332 | 2750064.75 | 2562721 | 2928965.75 | 2 | 2427600 | 2690053.75 | 2661585 | 1 | 8 | 0 | 7 | 1 | 8 | 1.20 | 1.16 | 0.93 | 0.90 |
| Q9CPU4 | Mgst3 | Microsomal glutathione S-transferase 3 | 256.37 | 23.53 | 16958 | 4 | 3716720.5 | 3163702 | 6959056 | 6691518 | 9663062 | 3191056.75 | 4599241 | 6860160.5 | 6508568 | 5455014 | 2226958.25 | 2409157.5 | 1923441.88 | 3594643.5 | 3118553.75 | 3405266.75 | 3678471.5 | 5036874.5 | 2126275 | 5669032.5 | 1 | 6 | 1 | 7 | 1 | 7 | 0.44 | 0.50 | 0.66 | 0.75 |
| E9PVX6 | Mki67 | Proliferation marker protein Ki-67 | 2137.01 | 14.73 | 350864 | 38 | 18237548 | 21623272 | 20936230 | 20104740 | 13855315 | 24192430 | 20341564 | 18846838 | 13036574 | 17339558 | 14189655 | 12522389 | 7815218 | 8020240.5 | 9065882 | 19992314 | 20396592 | 17253004 | 13625207 | 15613260 | 1 | 7 | 1 | 10 | 1 | 10 | 0.54 | 0.55 | 0.92 | 0.93 |
| Q9D2Y4 | Mlkl | Mixed lineage kinase domain-like protein | 120.64 | 5.3 | 54317 | 2 | 2367858.25 | 1408879.25 | 1728854.75 | 2239227.5 | 1661638.5 | 2493080 | 2111188.75 | 2524071.5 | 2342583.75 | 2740276 | 4045111.5 | 3493493.5 | 2694530.5 | 2943524.5 | 3147656.25 | 2513992.75 | 2944931.5 | 2757735 | 3209169 | 2584593 | 0 | 1 | 0 | 3 | 1 | 3 | 1.74 | 1.34 | 1.49 | 1.15 |
| Q924M7 | Mpi | Mannose-6-phosphate isomerase | 174.24 | 7.09 | 46575 | 3 | 1262522.5 | 1569284.25 | 1591880 | 1711504.12 | 2037902.25 | 1248247.25 | 1126844.38 | 2306488.25 | 1260217.25 | 1311299.38 | 2 | 765651.62 | 1233381.38 | 459818.5 | 937681.81 | 2859453 | 1428225.75 | 1349508.38 | 1054772.25 | 1580521.75 | 0 | 9 | 1 | 8 | 1 | 9 | 0.42 | 0.47 | 1.01 | 1.14 |
| Q99J99 | Mpst | 3-mercaptopyruvate sulfurtransferase | 106.01 | 8.75 | 33097 | 2 | 2 | 2 | 427705.16 | 2 | 310507.25 | 268953.66 | 132962.69 | 465116.34 | 93027.92 | 351283.06 | 1029353.69 | 459574.16 | 904809.12 | 692614.19 | 675486.81 | 144186.08 | 603379.19 | 282284.91 | 184493.89 | 229563.42 | 0 | 8 | 1 | 11 | 1 | 11 | 5.10 | 2.87 | 1.96 | 1.10 |
| P18665 | Mrpl3 | 39S ribosomal protein L3, mitochondrial | 58.25 | 2.02 | 38253 | 1 | 485876.78 | 2 | 470934.88 | 502678.47 | 294553 | 488074.41 | 302611.22 | 2 | 433764.16 | 641439.94 | 866696.38 | 585337 | 812248 | 759058.5 | 878671.25 | 422229.16 | 2 | 581130.75 | 453622.38 | 537530.75 | 0 | 10.5 | 1 | 11.5 | 1 | 11.5 | 2.22 | 2.09 | 1.14 | 1.07 |
| P26041 | Msn | Moesin | 3630.031 | 69.67 | 67767 | 56 | 212082258 | 202561842 | 231815666 | 179409346 | 211765652 | 210067010 | 230542722 | 246007042 | 273241986 | 287832066 | 297085922 | 334973762 | 285601010 | 298421474 | 316690318 | 192055986 | 232557810 | 306368034 | 291327682 | 292335138 | 0 | 4 | 1 | 4 | 1 | 4 | 1.40 | 1.31 | 1.35 | 1.26 |
| Q9R190 | Mta2 | Metastasis-associated protein MTA2 | 596.9 | 18.41 | 75030 | 12 | 9114256 | 10587944 | 10342166 | 8767436 | 8651210 | 10482642 | 8843768 | 9357658 | 8717441 | 10269779 | 10567941 | 11545380 | 11980942 | 11624290 | 12756540 | 8480835 | 9086683 | 11329690 | 11254962 | 11159037 | 1 | 8 | 0 | 8 | 1 | 8 | 1.23 | 1.23 | 1.08 | 1.08 |
| Q9EQK5 | Mvp | Major vault protein | 1734.31 | 46.69 | 95924 | 29 | 20907910 | 25496224 | 30619960 | 32573696 | 24674756 | 26903444 | 23731840 | 31245976 | 29695352 | 36566502 | 38150878 | 36390578 | 40536630 | 43395038 | 43257378 | 19514982 | 27271498 | 32315004 | 30476170 | 35211850 | 0 | 10 | 1 | 12 | 1 | 12 | 1.50 | 1.36 | 1.08 | 0.98 |
| P70402 | Mybph | Myosin-binding protein H | 46.16 | 1.66 | 52641 | 1 | 330678.78 | 309196.12 | 2 | 518573.69 | 230246.08 | 2 | 2 | 536965.06 | 2 | 2 | 1463187.25 | 710325.44 | 556520.56 | 1088478.12 | 536905.75 | 409521.53 | 245297.27 | 600999.75 | 2 | 678561.94 | 0 | 8.5 | 1 | 5 | 1 | 8.5 | 3.14 | 8.11 | 1.39 | 3.60 |
| O09043 | Napsa | Napsin-A | 229.4 | 17.42 | 45544 | 4 | 6080154.5 | 4585954 | 6229807 | 8346768.5 | 7231691.5 | 4864598.5 | 5135305 | 5948137.5 | 5553156 | 7747148 | 8310888 | 2738259.5 | 4933388.5 | 4539107 | 3345277.5 | 5065528.5 | 919437.88 | 6425516 | 7583912 | 2 | 1 | 8 | 1 | 5 | 1 | 9 | 0.48 | 0.53 | 0.62 | 0.68 |
| Q78PB6 | Ndel1 | Nuclear distribution protein nudE-like 1 | 76.7 | 3.48 | 38366 | 1 | 624331.12 | 577392 | 561622.31 | 651317.56 | 579239.88 | 689109.62 | 548463.81 | 331798.59 | 470465 | 471035.06 | 175582.05 | 274071.72 | 414518.75 | 193059.81 | 2 | 724063.69 | 529171.31 | 467108.91 | 652681.5 | 286680.97 | 0 | 10 | 1 | 12 | 1 | 12 | 0.35 | 0.42 | 0.89 | 1.06 |
| Q9DCJ5 | Ndufa8 | NADH dehydrogenase [ubiquinone] 1 alpha subcomplex subunit 8 | 163.1 | 20.93 | 19992 | 3 | 2837565.75 | 2871971.5 | 2770359.75 | 2268351.75 | 2057943 | 2840368 | 2745397.5 | 2101966 | 2404797.5 | 2034400 | 1159601 | 1176545.62 | 1555857.5 | 1699162.5 | 2070968.75 | 2672819.75 | 1592797.75 | 2164964.75 | 1755581 | 1804521.88 | 1 | 3 | 1 | 5 | 1 | 5 | 0.60 | 0.63 | 0.78 | 0.82 |
| Q8K1R7 | Nek9 | Serine/threonine-protein kinase Nek9 | 198.6 | 4.37 | 107143 | 4 | 3600253.5 | 4028356.25 | 3994219.5 | 3823853 | 3206381.5 | 3409509.5 | 3869755.5 | 5381937 | 4744208.5 | 4607966 | 1187979.88 | 2737968.75 | 2714143.5 | 2382375.75 | 3240947.5 | 3634014 | 3293495.75 | 4088582.5 | 4185709.5 | 3190629.25 | 1 | 12 | 0 | 5 | 1 | 12 | 0.66 | 0.56 | 0.99 | 0.84 |
| Q8BZW8 | Nhlrc2 | NHL repeat-containing protein 2 | 153.95 | 4.69 | 78430 | 3 | 1826595.5 | 1889379.25 | 1962065 | 2243295.75 | 1882059.38 | 1813269 | 1826638.5 | 2039242.25 | 1496183 | 2392662.5 | 1661667 | 849205.62 | 1265052.62 | 1454385.5 | 1462582 | 2239470 | 2434680 | 2351787 | 2814327.75 | 2125345.75 | 0 | 2 | 1 | 3 | 1 | 3 | 0.68 | 0.70 | 1.22 | 1.25 |
| Q9WV85 | Nme3 | Nucleoside diphosphate kinase 3 | 40.44 | 10.06 | 19099 | 1 | 822249.19 | 524122.94 | 894998.38 | 1100658.75 | 636907.69 | 584892.62 | 664396.25 | 737392.44 | 633397.5 | 747633.44 | 500719.25 | 474790.16 | 2 | 612435.44 | 2 | 713585.81 | 1064367.12 | 564412.19 | 2 | 2 | 1 | 7 | 1 | 8 | 1 | 8 | 0.40 | 0.47 | 0.59 | 0.70 |
| Q8K2T1 | Nmral1 | NmrA-like family domain-containing protein 1 | 202.99 | 11.97 | 34376 | 4 | 5982230.5 | 6827606 | 9352849 | 8491133 | 11117667 | 7360319 | 6306315 | 8417797 | 9027412 | 9043260 | 3947668.5 | 4330879 | 4432504 | 4191282 | 6360666.5 | 7350851 | 3658344 | 7506822 | 7709449.5 | 6711734 | 1 | 7 | 1 | 6 | 1 | 7 | 0.56 | 0.58 | 0.79 | 0.82 |
| Q6NSR8 | Npepl1 | Probable aminopeptidase NPEPL1 | 219.72 | 9.73 | 55940 | 4 | 1649239.25 | 1470620.75 | 785400.62 | 1221960.12 | 4293510.5 | 1411413.38 | 1347784.75 | 635121.25 | 2277790.25 | 804763.12 | 661980.44 | 2 | 366421.66 | 411752.09 | 298041 | 1972356 | 676263.31 | 2 | 345110 | 267907.12 | 0 | 4 | 1 | 5 | 1 | 5 | 0.18 | 0.27 | 0.35 | 0.50 |
| P97300 | Nptn | Neuroplastin | 155.46 | 8.56 | 44373 | 3 | 6338515 | 6769829 | 6435087 | 7021597 | 5524188 | 8346771 | 6180451 | 7044575.5 | 7618995 | 9859714 | 10582578 | 9619518 | 11919438 | 11167702 | 13534597 | 6187024 | 7767924.5 | 8093410 | 6846569.5 | 8159578 | 0 | 5 | 1 | 11 | 1 | 11 | 1.77 | 1.46 | 1.15 | 0.95 |
| Q9JI75 | Nqo2 | Ribosyldihydronicotinamide dehydrogenase [quinone] | 60.32 | 6.93 | 26248 | 1 | 977400 | 738401.69 | 801079.12 | 842381.69 | 926530.69 | 961878.19 | 1129733.75 | 745140.38 | 939220.12 | 1088188.75 | 223243.05 | 752070 | 483346.5 | 568974.88 | 458673.41 | 1033163.88 | 784627.12 | 1059206.62 | 1163624.62 | 996258.38 | 1 | 4 | 1 | 10 | 1 | 10 | 0.58 | 0.51 | 1.18 | 1.04 |
| Q9R061 | Nubp2 | Cytosolic Fe-S cluster assembly factor NUBP2 | 130.26 | 11.27 | 29518 | 2 | 2140320.5 | 1817808 | 2179952.5 | 2284982.75 | 1962481.12 | 2234874 | 1902162.5 | 2007064.25 | 2291218 | 1911755.62 | 1176419.5 | 1742809.25 | 1881105.88 | 1402613 | 1499760.5 | 1871171 | 2362125 | 2120427 | 2524588 | 1928037.75 | 1 | 11 | 0 | 10 | 1 | 11 | 0.74 | 0.74 | 1.04 | 1.04 |
| Q6PIP5 | Nudcd1 | NudC domain-containing protein 1 | 243.73 | 9.11 | 66705 | 5 | 1765560.5 | 1426470.5 | 3041721.5 | 1468120.38 | 1995309.25 | 1615441 | 964541.31 | 1758209.5 | 1400803.75 | 2467015 | 3585880 | 3324700 | 3189778 | 2404625.25 | 3653680.25 | 900858.31 | 1111801.75 | 991375.44 | 3238579.5 | 2223381.5 | 1 | 9 | 1 | 11 | 1 | 11 | 1.67 | 1.97 | 0.87 | 1.03 |
| Q8VI94 | Oasl1 | 2'-5'-oligoadenylate synthase-like protein 1 | 846.77 | 42.86 | 59088 | 14 | 1779162.75 | 1644981.5 | 787050.81 | 2298811.75 | 2646937.25 | 1232091.12 | 2124029.75 | 1673425 | 3238600 | 1647785.88 | 4303419 | 3784448.25 | 7181445.5 | 2712969.5 | 6314237 | 860258.38 | 1354482.38 | 1408830 | 1838586.75 | 693515.62 | 0 | 6 | 1 | 5 | 1 | 6 | 2.65 | 2.45 | 0.67 | 0.62 |
| B2RYG6 | Otub1 | Ubiquitin thioesterase OTUB1 | 575.62 | 32.47 | 31270 | 8 | 8595271 | 10289466 | 12230402 | 7348640 | 20404588 | 6200067.5 | 11351350 | 12137798 | 17574936 | 6795789 | 4120724.75 | 6214604 | 4160615.5 | 3441791.75 | 4329821 | 8567423 | 6626194.5 | 7898143.5 | 9136252 | 8307079 | 0 | 5 | 1 | 9 | 1 | 9 | 0.38 | 0.41 | 0.69 | 0.75 |
| P29341 | Pabpc1 | Polyadenylate-binding protein 1 | 1650.55 | 38.21 | 70671 | 27 | 71764874 | 73229554 | 71573130 | 72963914 | 59216166 | 77559842 | 72697714 | 63396762 | 58850238 | 67621882 | 123755802 | 107136274 | 102696002 | 77130242 | 102711738 | 71345338 | 76383450 | 72626754 | 71635618 | 71347490 | 0 | 12 | 1 | 9 | 1 | 12 | 1.47 | 1.51 | 1.04 | 1.07 |
| Q9D0B6 | Pbdc1 | Protein PBDC1 | 270.14 | 29.29 | 22223 | 5 | 4998629 | 4162354.25 | 5973530 | 4239568 | 3444856.5 | 3305390.25 | 4477362.5 | 4795944.5 | 6557477 | 3937811.5 | 876102.75 | 3426222 | 2267201.25 | 584249.31 | 2455201 | 4301589 | 996218.75 | 4003159.75 | 3221182 | 4239598 | 0 | 7 | 1 | 6 | 1 | 7 | 0.42 | 0.42 | 0.73 | 0.73 |
| Q3TVI8 | Pbxip1 | Pre-B-cell leukemia transcription factor-interacting protein 1 | 344.86 | 9.9 | 81164 | 5 | 3501984.75 | 2783742.25 | 3356594.5 | 2899014 | 3353887.5 | 2710739.75 | 2844593.75 | 2783999.75 | 2808963.75 | 1889654.25 | 1452942 | 1704197.88 | 1975134.12 | 777136.5 | 1732772.88 | 3003101.25 | 1965261.75 | 2145500.75 | 2639429.5 | 1934912.62 | 0 | 2 | 1 | 9 | 1 | 9 | 0.48 | 0.59 | 0.74 | 0.90 |
| Q8CI51 | Pdlim5 | PDZ and LIM domain protein 5 | 373.91 | 13.87 | 63299 | 7 | 5665323.5 | 6637987.5 | 6780425 | 7549711 | 8856531 | 7623103 | 7095839.5 | 8481202 | 8763612 | 7475861 | 4790414.5 | 5140974.5 | 5858525.5 | 3951268.25 | 5615782 | 5344975.5 | 5908886 | 7996815.5 | 8926034 | 9355953 | 1 | 10 | 0 | 12 | 1 | 12 | 0.71 | 0.64 | 1.06 | 0.95 |
| P70296 | Pebp1 | Phosphatidylethanolamine-binding protein 1 | 226.84 | 22.46 | 20830 | 4 | 14913318 | 19017374 | 17186178 | 11304594 | 18450446 | 10975424 | 16349182 | 20548624 | 19800590 | 13790762 | 6281577.5 | 13072753 | 8578844 | 7377075.5 | 10106960 | 16111036 | 9218788 | 12548082 | 13648611 | 15926708 | 1 | 6 | 1 | 6 | 1 | 6 | 0.56 | 0.56 | 0.83 | 0.83 |
| Q9Z210 | Pex11b | Peroxisomal membrane protein 11B | 54.72 | 4.63 | 28710 | 1 | 2 | 2 | 2 | 2 | 344777.78 | 332900.16 | 2 | 2 | 2 | 2 | 724620.06 | 161781.72 | 591086.62 | 923490.31 | 825582.06 | 358937.56 | 2 | 576193.69 | 2 | 2 | 1 | 9 | 1 | 9 | 1 | 9 | 9.36 | 9.69 | 2.71 | 2.81 |
| Q5SUR0 | Pfas | Phosphoribosylformylglycinamidine synthase | 730.95 | 11.44 | 144629 | 13 | 28995282 | 21501994 | 23725600 | 23631096 | 20755394 | 27021230 | 24758644 | 19991874 | 18016396 | 24448922 | 14556991 | 18565342 | 16842270 | 13096330 | 13409008 | 25599420 | 25251280 | 22341134 | 27152226 | 22907224 | 0 | 9 | 1 | 8 | 1 | 9 | 0.64 | 0.67 | 1.04 | 1.08 |
| E2JF22 | Piezo1 | Piezo-type mechanosensitive ion channel component 1 | 81.58 | 0.71 | 292002 | 2 | 105303.11 | 255191.12 | 316827.97 | 284994.16 | 303752.88 | 283949.47 | 285851.25 | 2 | 396604.06 | 413044.81 | 565501.81 | 401417.19 | 726700.75 | 493278.94 | 421860.88 | 2 | 123512.69 | 319327.56 | 387086.53 | 378066.16 | 0 | 9 | 1 | 10.5 | 1 | 10.5 | 2.06 | 1.89 | 0.95 | 0.88 |
| Q9R1S3 | Pign | GPI ethanolamine phosphate transferase 1 | 266.29 | 3.76 | 105045 | 4 | 963525.62 | 908864.75 | 1026610.81 | 1068950.62 | 848070.75 | 474491.41 | 1015214.75 | 875895.12 | 932954.88 | 1311912.12 | 1623856 | 1851832.75 | 2037209.5 | 1056396 | 2454058.75 | 955597.81 | 576100.31 | 949141.38 | 1009777 | 1132076.25 | 1 | 12 | 1 | 11 | 1 | 12 | 1.87 | 1.96 | 0.96 | 1.00 |
| Q6PD26 | Pigs | GPI transamidase component PIG-S | 537.41 | 18.74 | 61711 | 8 | 6209113 | 6569209 | 7305560 | 7186453 | 3837122.5 | 6280324 | 6494984 | 6616174 | 6727003 | 7794197.5 | 7650928 | 10429734 | 11891579 | 8394614 | 10508785 | 6268288 | 3845039 | 6453690 | 7967339 | 7165303.5 | 0 | 12 | 1 | 10 | 1 | 12 | 1.57 | 1.44 | 1.02 | 0.93 |
| Q8CHJ1 | Pigu | Phosphatidylinositol glycan anchor biosynthesis class U protein | 116.17 | 6.21 | 49953 | 2 | 2 | 2 | 2 | 2 | 296775.66 | 2 | 200657.27 | 638889.69 | 541916.31 | 2 | 565129.56 | 989453.44 | 1372125.88 | 964058.12 | 1158465.5 | 2 | 2 | 372500.97 | 509778.47 | 191905.53 | 0 | 7 | 1 | 10 | 1 | 10 | 17.01 | 3.65 | 3.62 | 0.78 |
| Q9EQ32 | Pik3ap1 | Phosphoinositide 3-kinase adapter protein 1 | 496.07 | 12.08 | 90928 | 7 | 6059862 | 4418039 | 6732431.5 | 8317002.5 | 6404953.5 | 5592814.5 | 5154366.5 | 4355602.5 | 5159243 | 5644535 | 4363215 | 4029316.25 | 3035976.5 | 3029184.5 | 4195602 | 4906447 | 5975102 | 5712171.5 | 7062495 | 5914852.5 | 0 | 8 | 1 | 4 | 1 | 8 | 0.58 | 0.72 | 0.93 | 1.14 |
| O70172 | Pip4k2a | Phosphatidylinositol 5-phosphate 4-kinase type-2 alpha | 89.4 | 4.94 | 46152 | 2 | 1657854.62 | 1832224.12 | 1959764.5 | 2341474 | 1413628.38 | 1500180.25 | 1602028.38 | 2063010.38 | 1783427.38 | 2130366.75 | 784693.56 | 1253266.38 | 1203658.62 | 1436902.62 | 1040269.25 | 1600702.25 | 1756655.75 | 1959212.25 | 1893907 | 1930600.12 | 1 | 12 | 0 | 12 | 1 | 12 | 0.62 | 0.63 | 0.99 | 1.01 |
| O35405 | Pld3 | 5'-3' exonuclease PLD3 | 392.14 | 20.7 | 54389 | 7 | 3214267.25 | 3539842 | 2436896.75 | 2722223 | 3665813.75 | 4079438 | 4483354.5 | 5423772.5 | 6757396 | 6038630.5 | 6374901.5 | 7407418 | 9429739 | 7659524.5 | 9040842 | 6292829 | 5525868.5 | 7797509 | 9898626 | 8771491 | 0 | 1 | 1 | 3 | 1 | 3 | 2.56 | 1.49 | 2.46 | 1.43 |
| Q9R0E1 | Plod3 | Multifunctional procollagen lysine hydroxylase and glycosyltransferase LH3 | 566.03 | 16.73 | 84922 | 9 | 1540980.75 | 678887.75 | 2109299.5 | 1516739.75 | 533720.06 | 1367653.25 | 476450.5 | 825457.81 | 1440722.75 | 1521858.88 | 3359996.75 | 2470266.5 | 3705967.75 | 1699541.25 | 2976689 | 740764.31 | 1082651.12 | 2631754.75 | 2241530.25 | 2085699.5 | 1 | 7 | 0 | 7 | 1 | 7 | 2.23 | 2.52 | 1.38 | 1.56 |
| P70206 | Plxna1 | Plexin-A1 | 1223.66 | 14.94 | 211099 | 23 | 5802266.5 | 6671794.5 | 7855508.5 | 6443910 | 7088993.5 | 8365015.5 | 9139928 | 8274161 | 12718988 | 11241703 | 17555850 | 20203850 | 19241566 | 12703895 | 16273512 | 10199227 | 5724312.5 | 7951007.5 | 8000643 | 9183523 | 0 | 5 | 1 | 6 | 1 | 6 | 2.54 | 1.73 | 1.21 | 0.83 |
| Q9D1G2 | Pmvk | Phosphomevalonate kinase | 260.4 | 31.77 | 21916 | 5 | 3636894 | 2541403.25 | 2030934 | 2444370.25 | 2144110 | 3805029.25 | 2841825.5 | 1888259.75 | 2786276.75 | 2813759.5 | 5370741 | 4009593.5 | 4215927 | 3455477.5 | 4213495 | 3885400 | 3226406 | 3750922 | 3528626 | 2805751 | 1 | 3 | 1 | 5 | 1 | 5 | 1.66 | 1.50 | 1.34 | 1.22 |
| Q9D819 | Ppa1 | Inorganic pyrophosphatase | 572.01 | 34.26 | 32667 | 8 | 19859294 | 21408738 | 14730285 | 16873374 | 23661910 | 16446080 | 20247070 | 18240042 | 20675414 | 14896984 | 8390907 | 15457038 | 9025947 | 6734176.5 | 9072174 | 21951644 | 13143150 | 13726807 | 14114551 | 16397048 | 1 | 6 | 1 | 6 | 1 | 6 | 0.50 | 0.54 | 0.82 | 0.88 |
| Q61151 | Ppp2r5e | Serine/threonine-protein phosphatase 2A 56 kDa regulatory subunit epsilon isoform | 84.89 | 6.21 | 54713 | 2 | 581839.62 | 797561.44 | 818752.25 | 944509.75 | 929895.5 | 1139035.75 | 680427 | 997967.38 | 1044034.38 | 726762.12 | 332185.12 | 574856.12 | 588169.06 | 109137.74 | 415826.06 | 622146.94 | 820175.19 | 947812.19 | 906005.12 | 2 | 1 | 10 | 0 | 8 | 1 | 10 | 0.51 | 0.50 | 1.02 | 0.90 |
| P05132 | Prkaca | cAMP-dependent protein kinase catalytic subunit alpha | 225.89 | 15.38 | 40571 | 4 | 2858580.25 | 3108406 | 3374145.25 | 2908140.25 | 4036980 | 2035747.5 | 3072890 | 2762375.25 | 5191796.5 | 2621666.5 | 1397399 | 1933271.5 | 2109965.25 | 1455798.5 | 1640237.12 | 2617849 | 1684755.62 | 2940111 | 2927366.5 | 2550385.5 | 0 | 4 | 1 | 8 | 1 | 8 | 0.52 | 0.54 | 0.78 | 0.81 |
| Q9R1P4 | Psma1 | Proteasome subunit alpha type-1 | 494.49 | 38.02 | 29547 | 10 | 34519946 | 31288942 | 36623458 | 39450122 | 37002942 | 30724540 | 31446946 | 33633002 | 32242506 | 36052066 | 26698738 | 29954486 | 27189802 | 22763434 | 31313980 | 41139434 | 24950610 | 34268282 | 34400990 | 34921230 | 1 | 9 | 1 | 8 | 1 | 9 | 0.79 | 0.86 | 0.95 | 1.03 |
| P43252 | Ptgir | Prostacyclin receptor | 58.46 | 2.65 | 44463 | 1 | 2 | 2 | 2 | 2 | 741393.19 | 2 | 2 | 2 | 2 | 1021421.88 | 228513.16 | 1863760.25 | 2316463 | 1683491.38 | 1663605.38 | 2 | 2 | 908284.81 | 2 | 2 | 1 | 12 | 1 | 12 | 1 | 12 | 10.46 | 7.59 | 1.23 | 0.89 |

| | | | | | | | | | | | | | | | | | | | | | | | | | | | | | | | | | | | | |
|---|---|---|---|---|---|---|---|---|---|---|---|---|---|---|---|---|---|---|---|---|---|---|---|---|---|---|---|---|---|---|---|---|---|---|---|---|
| P58389 | Ptpa | Serine/threonine-protein phosphatase 2A activator | 202.12 | 12.07 | 36710 | 3 | 3938742.5 | 3713651 | 4093979.25 | 2912235 | 4870607.5 | 2161377.25 | 3375384.75 | 3650901.5 | 3542191 | 2467944.75 | 837075.12 | 2324851 | 1489363.75 | 433269.94 | 1290736.75 | 3603244 | 904473.88 | 1734356.88 | 1888132.75 | 2440945 | 0 | 1 | 1 | 5 | 1 | 5 | 0.33 | 0.42 | 0.54 | 0.70 |
| P42669 | Pura | Transcriptional activator protein Pur-alpha | 134.37 | 12.77 | 34884 | 3 | 1083241.5 | 743648.94 | 1035008.88 | 1696576.5 | 795160.25 | 634095.94 | 1018748.81 | 161868.7 | 826597.69 | 1717417.75 | 2672070 | 2539794 | 2125801.5 | 1342810.62 | 2668061.25 | 894749.38 | 406294.16 | 79114.3 | 707323.88 | 843116.81 | 1 | 4 | 1 | 9 | 1 | 9 | 2.12 | 2.60 | 0.55 | 0.67 |
| Q61838 | Pzp | Pregnancy zone protein | 121.74 | 1.61 | 165853 | 3 | 2762633.75 | 2529627.25 | 3738349.25 | 3290433.5 | 2583552.75 | 2923163.5 | 3623689 | 3432464.5 | 3965246.25 | 4719979.5 | 4635071.5 | 6804871 | 8201470.5 | 4946939 | 7872005.5 | 2620860.75 | 1834968.75 | 3838348.25 | 4045025.5 | 3070384.75 | 0 | 10 | 1 | 8 | 1 | 10 | 2.18 | 1.74 | 1.03 | 0.83 |
| P61021 | Rab5b | Ras-related protein Rab-5B | 417.59 | 36.74 | 23707 | 6 | 6130193 | 5116326.5 | 4333350.5 | 5980961 | 4430825 | 4854616 | 4575215.5 | 4598084 | 5220267 | 5485281 | 3643185 | 3485026 | 4211917.5 | 4415592 | 3747807 | 4937451.5 | 6101923.5 | 5376425 | 5814321 | 4306203.5 | 1 | 12 | 0 | 8 | 1 | 12 | 0.75 | 0.79 | 1.02 | 1.07 |
| P11345 | Raf1 | RAF proto-oncogene serine/threonine-protein kinase | 112.48 | 3.55 | 72928 | 2 | 1107744.25 | 1033716.19 | 792661.06 | 1033771.12 | 677559 | 1028159.25 | 753688 | 958455.44 | 874359.38 | 989522.75 | 530784.38 | 453395.84 | 2 | 537213.94 | 705361.25 | 1129470 | 1261313.5 | 916124.31 | 1063250.38 | 1172414.25 | 0 | 4 | 0 | 3 | 1 | 4 | 0.48 | 0.48 | 1.19 | 1.20 |
| A3KGS3 | Ralgapa2 | Ral GTPase-activating protein subunit alpha-2 | 195.33 | 2.46 | 210288 | 4 | 1117176.25 | 1001787.81 | 682387.88 | 964930.25 | 1767191.12 | 960193.81 | 1121670.25 | 1012967.25 | 1503715.25 | 1026786.5 | 349743.91 | 231863.47 | 700349.62 | 2 | 102621.63 | 1106796.38 | 963941.5 | 595607.38 | 318182.34 | 723244.88 | 1 | 5 | 0 | 4 | 1 | 5 | 0.25 | 0.25 | 0.67 | 0.66 |
| P46061 | Rangap1 | Ran GTPase-activating protein 1 | 1294.67 | 39.39 | 63531 | 21 | 37230578 | 42114390 | 34928138 | 35216194 | 38420434 | 37296586 | 37738882 | 38428122 | 37520322 | 31811338 | 31271288 | 32811794 | 25313130 | 28053954 | 26883960 | 37733762 | 32375484 | 29420776 | 33397182 | 32738398 | 0 | 3 | 1 | 6 | 1 | 6 | 0.77 | 0.79 | 0.88 | 0.91 |
| P26043 | Rdx | Radixin | 1179.02 | 29.16 | 68543 | 20 | 1153317.75 | 2116427 | 1200595.12 | 848674.38 | 1462502.75 | 1627727.88 | 1483415.75 | 1565809.75 | 2094775.62 | 1227480 | 2814882.5 | 2762675.75 | 2853505.5 | 2071505.88 | 3328681 | 1642778.12 | 2019690 | 1646469.75 | 1807440.12 | 1389065.5 | 1 | 6 | 1 | 8 | 1 | 8 | 2.04 | 1.73 | 1.25 | 1.06 |
| P15307 | Rel | Proto-oncogene c-Rel | 51.21 | 1.87 | 64960 | 1 | 592823.38 | 639449.75 | 567581 | 544314 | 423812.78 | 2 | 550020.5 | 514869.59 | 476349.28 | 777114.69 | 1084187.88 | 968475.38 | 1136052.5 | 694820.12 | 984168.19 | 2 | 533138.94 | 457044.5 | 439686.78 | 522171.94 | 0 | 4 | 1 | 8.5 | 1 | 8.5 | 1.76 | 2.10 | 0.71 | 0.84 |
| Q99J62 | Rfc4 | Replication factor C subunit 4 | 240.63 | 15.93 | 39867 | 5 | 6456764 | 5550526 | 7077640.5 | 7530501 | 4911740.5 | 5867971 | 5392639 | 5309192 | 3601785 | 6310186 | 2079287.25 | 2055400.5 | 2318835.25 | 4224212.5 | 1678667.25 | 5871756 | 6340987.5 | 4200538 | 6029969.5 | 4489544 | 0 | 6 | 1 | 10 | 1 | 10 | 0.39 | 0.47 | 0.85 | 1.02 |
| Q9DBU3 | Riok3 | Serine/threonine-protein kinase RIO3 | 44.22 | 1.73 | 58704 | 1 | 2 | 635738.5 | 340251.41 | 565695.12 | 2 | 614227.12 | 685513.81 | 395232.12 | 523267.5 | 556423.12 | 749453.69 | 766342.12 | 726983.81 | 766251.31 | 646078.81 | 2 | 555743.5 | 2 | 720733 | 2 | 0 | 11 | 1 | 7 | 1 | 11 | 2.37 | 1.32 | 0.83 | 0.46 |
| P17078 | Rpl35 | 60S ribosomal protein L35 | 181.46 | 21.95 | 14553 | 4 | 1121194.88 | 1613077 | 841653.06 | 648526.88 | 517271.66 | 1719203.62 | 1417661.62 | 2322194.75 | 861991.88 | 770731 | 3363037.5 | 4435335.5 | 2794574 | 1924032.88 | 2439354.75 | 785092.69 | 699684.38 | 266629.34 | 580719.25 | 294968.41 | 0 | 5 | 1 | 1 | 1 | 5 | 3.15 | 2.11 | 0.55 | 0.37 |
| P61928 | Rpl37 | 60S ribosomal protein L37 | 48.17 | 7.22 | 11078 | 1 | 3452314 | 4585279 | 4669056.5 | 4746459 | 3557778.25 | 3006567.75 | 3633208 | 4620881.5 | 3459302 | 4068130 | 2623986.5 | 2380897 | 2471176.5 | 3337402.75 | 2530356 | 4309645 | 3585380.75 | 3071689.25 | 3344069.75 | 5172138 | 0 | 9 | 1 | 12 | 1 | 12 | 0.64 | 0.71 | 0.93 | 1.04 |
| Q01730 | Rsu1 | Ras suppressor protein 1 | 181.97 | 20.58 | 31550 | 4 | 5233442.5 | 4247136 | 4905571 | 2809467 | 5192786.5 | 2003213.62 | 4444730 | 3923886 | 4382350.5 | 3227243.75 | 874575 | 3193500.25 | 1272742 | 520025.88 | 1753797.75 | 3940799.25 | 1845003.75 | 2034958.38 | 1811618.75 | 2378603.5 | 1 | 1 | 1 | 5 | 1 | 5 | 0.34 | 0.42 | 0.54 | 0.67 |
| Q9CQE8 | RTRAF | RNA transcription, translation and transport factor protein | 482.15 | 37.7 | 28152 | 8 | 2412755.75 | 2993737.25 | 3249002.25 | 1596407.12 | 4875344.5 | 1458948.25 | 3239405.5 | 2705338 | 4595946 | 1878315.5 | 382856.06 | 1719776.75 | 719565.44 | 922486.31 | 1208036.88 | 2219269.5 | 1610793.38 | 1330839.25 | 1849604.62 | 2122728.75 | 1 | 4 | 1 | 6 | 1 | 6 | 0.33 | 0.36 | 0.60 | 0.66 |
| Q9WTM5 | Ruvbl2 | RuvB-like 2 | 1178.63 | 52.27 | 51113 | 18 | 23127862 | 26879614 | 24735270 | 21821776 | 21499636 | 23595598 | 24770486 | 20748286 | 25418052 | 27539686 | 38637762 | 38122930 | 31110866 | 26699648 | 34679290 | 24790096 | 25157960 | 24061224 | 24927234 | 24285848 | 1 | 7 | 1 | 12 | 1 | 12 | 1.43 | 1.39 | 1.04 | 1.01 |
| Q9DBQ7 | Scyl3 | Protein-associating with the carboxyl-terminal domain of ezrin | 93.94 | 3.95 | 81333 | 2 | 1181338.5 | 1011650.69 | 1088263 | 1181949.25 | 1443407.25 | 1151600 | 1309330.5 | 1396113.75 | 1152293.12 | 954630.5 | 619290.94 | 966185.5 | 792917.94 | 377741.94 | 783893.88 | 1026389.81 | 748272.69 | 1033415.62 | 1244954.88 | 1020811.25 | 0 | 7 | 1 | 6 | 1 | 7 | 0.60 | 0.59 | 0.86 | 0.85 |
| Q8BU14 | Sec62 | Translocation protein SEC62 | 101.25 | 4.77 | 45581 | 2 | 3073701.5 | 3409986.5 | 3042914.5 | 3635743 | 2150073.25 | 4024070 | 3083394.5 | 2898358.5 | 2390697 | 3324120 | 2150099.5 | 1909699.38 | 1834859.75 | 1987701.12 | 1842400.25 | 1724340.88 | 2841435 | 1794644.25 | 2377395.75 | 2391615.5 | 1 | 3 | 0 | 2 | 1 | 3 | 0.64 | 0.62 | 0.73 | 0.71 |
| O55131 | Sept7 | Septin-7 | 664.46 | 34.86 | 50550 | 12 | 6697754 | 6178711 | 5724120 | 4337728 | 5392747.5 | 5390248 | 5919212.5 | 4480593 | 5413140.5 | 5249857.5 | 6798666 | 7255116.5 | 7059636 | 6577122.5 | 8478074 | 4678953.5 | 6410497 | 5623928.5 | 4981192.5 | 6443041.5 | 1 | 12 | 0 | 9 | 1 | 12 | 1.28 | 1.37 | 0.99 | 1.06 |
| Q8VHL1 | Setd7 | Histone-lysine N-methyltransferase SETD7 | 75.02 | 4.92 | 40506 | 1 | 378866.72 | 435084.53 | 413309.97 | 363087.31 | 431491.31 | 370060.44 | 394743.91 | 306899.38 | 431652.78 | 244623.55 | 2 | 2 | 303458.09 | 225321.92 | 2 | 457695.25 | 386273 | 560355.81 | 356516.47 | 262709.88 | 0 | 12 | 1 | 9 | 1 | 12 | 0.26 | 0.30 | 1.00 | 1.16 |
| P70665 | Siae | Sialate O-acetylesterase | 113.6 | 4.99 | 60775 | 2 | 1721404.12 | 1097780.12 | 324347.06 | 837222.62 | 1968767.5 | 761982.5 | 1443061.5 | 287876.59 | 1286508.88 | 1257775.12 | 2 | 420774.91 | 2 | 2 | 2 | 798645.38 | 119568.66 | 94193.4 | 811969.88 | 635668.25 | 1 | 3 | 1 | 5 | 1 | 5 | 0.07 | 0.08 | 0.41 | 0.49 |
| Q924N4 | Slc12a6 | Solute carrier family 12 member 6 | 236.24 | 3.91 | 127527 | 6 | 2 | 2 | 2 | 2 | 2 | 393485.03 | 2 | 482910.72 | 2 | 2 | 410261.47 | 563117.81 | 629829.31 | 941283.25 | 880737.69 | 2 | 2 | 449662.5 | 2 | 2 | 0 | 10 | 1 | 10 | 1 | 10 | 342522.95 | 3.91 | 44967.05 | 0.51 |
| Q9R0P4 | Smap | Small acidic protein | 90.55 | 8.84 | 20046 | 1 | 562305.44 | 739130.94 | 608453.06 | 456924.94 | 556191.38 | 515419.59 | 692143.81 | 635906.25 | 611140.62 | 723324.94 | 2 | 551844.94 | 2 | 2 | 359122.09 | 602423.44 | 2 | 222984.88 | 535403.25 | 491198.34 | 1 | 5 | 1 | 2 | 1 | 5 | 0.31 | 0.29 | 0.63 | 0.58 |
| P97355 | Sms | Spermine synthase | 243.69 | 17.21 | 41313 | 5 | 3176047.5 | 3187345.25 | 5210159 | 4005633.75 | 5406645 | 2594035 | 3447213.75 | 3724127.5 | 4999467.5 | 4134271.75 | 2909311 | 2121658 | 1419095.38 | 1816747.5 | 1916777 | 2957715 | 3772278.5 | 3778734.5 | 3555714 | 3498675 | 0 | 8 | 1 | 11 | 1 | 11 | 0.49 | 0.54 | 0.84 | 0.93 |
| P62307 | Snrpf | Small nuclear ribonucleoprotein F | 168.12 | 40.7 | 9725 | 3 | 517004.69 | 1012267.56 | 411603.72 | 296207 | 2 | 328566.91 | 556382.06 | 441216.5 | 2 | 629587.12 | 2530098.25 | 1238572.5 | 1211369.5 | 858502.12 | 1797688.5 | 345010.34 | 423670.25 | 254081.73 | 2 | 284998.62 | 1 | 7.5 | 0 | 6.5 | 1 | 7.5 | 3.41 | 3.90 | 0.58 | 0.67 |
| Q58A65 | Spag9 | C-Jun-amino-terminal kinase-interacting protein 4 | 293.53 | 7.27 | 146219 | 7 | 4042364.5 | 3117856 | 5380525 | 5774084 | 5362477.5 | 3367363.75 | 4741961 | 4428129 | 5617291 | 5317781 | 1397446.75 | 2674842.25 | 2263180 | 3344001 | 2385621 | 3546482 | 3403521.25 | 4645272 | 4881703.5 | 6166646 | 1 | 11 | 0 | 12 | 1 | 12 | 0.51 | 0.51 | 0.96 | 0.96 |
| Q9DBG7 | Srpra | Signal recognition particle receptor subunit alpha | 890.55 | 25.16 | 69623 | 13 | 8883435 | 10801511 | 10122705 | 10574733 | 9839620 | 10135145 | 9217604 | 7991960 | 10003137 | 11679127 | 14866605 | 17255778 | 17384604 | 11201151 | 17821756 | 8897067 | 5895018.5 | 10815798 | 10091625 | 12821886 | 0 | 12 | 1 | 12 | 1 | 12 | 1.56 | 1.60 | 0.97 | 0.99 |
| Q8VE97 | Srsf4 | Serine/arginine-rich splicing factor 4 | 259.94 | 10.22 | 55979 | 5 | 3537528.25 | 4868679.5 | 4742036 | 4323918.5 | 4666186.5 | 2645047.25 | 4120089.75 | 4388367.5 | 2900566.75 | 2536712.75 | 1416730.5 | 2549506.25 | 1225663 | 768305.38 | 863210.06 | 4682822.5 | 3670940 | 2106839 | 3316167.25 | 3213644.5 | 0 | 4 | 1 | 11 | 1 | 11 | 0.31 | 0.41 | 0.77 | 1.02 |
| Q7TPJ0 | Ssr1 | Translocon-associated protein subunit alpha | 296.52 | 15.99 | 35629 | 6 | 5429944.5 | 5059912 | 5510637.5 | 4019153.75 | 6217843.5 | 3664652.5 | 5438331 | 5794526 | 7370894.5 | 4657428.5 | 8908013 | 15967832 | 12839976 | 6555656 | 13749827 | 5586816.5 | 2573361.5 | 3653790 | 4448721.5 | 3942394.75 | 0 | 5 | 1 | 5 | 1 | 5 | 2.21 | 2.15 | 0.77 | 0.75 |
| Q9JM90 | Stap1 | Signal-transducing adaptor protein 1 | 528.62 | 51.85 | 34628 | 12 | 10493390 | 12171022 | 17410320 | 17362668 | 15931860 | 18146626 | 15900156 | 23202226 | 22330234 | 25613038 | 31580526 | 26355964 | 32139602 | 23570396 | 30711744 | 12507830 | 11517922 | 25113514 | 17780154 | 22784918 | 0 | 7 | 1 | 8 | 1 | 8 | 1.97 | 1.37 | 1.22 | 0.85 |
| P52633 | Stat6 | Signal transducer and activator of transcription 6 | 44.19 | 2.15 | 93497 | 1 | 205695.19 | 229973.66 | 2 | 2 | 2 | 261475.75 | 162797.86 | 2 | 2 | 2 | 342154.78 | 266777.97 | 252979.62 | 1522707.5 | 327992.72 | 2 | 215311.94 | 103722.41 | 2 | 2 | 0 | 11.5 | 1 | 11.5 | 1 | 11.5 | 6.23 | 6.39 | 0.73 | 0.75 |
| Q80X82 | Sympk | Symplekin | 278.58 | 5.61 | 142284 | 6 | 1987055.5 | 1715299 | 1010320.06 | 1038999.88 | 1816224.75 | 1609541.62 | 1531388.5 | 1443532.75 | 2274122 | 2987971.25 | 3190075.75 | 3778085 | 3465361.5 | 2437852.25 | 3442991.25 | 1638325.25 | 1884755 | 1326838 | 1325411.88 | 1515635.25 | 0 | 12 | 1 | 7 | 1 | 12 | 2.16 | 1.66 | 1.02 | 0.78 |
| Q0VGY8 | Tanc1 | Protein TANC1 | 324.2 | 4.47 | 200805 | 6 | 3377116.5 | 3186368 | 4144158.5 | 3501937 | 3687900.5 | 4097850.5 | 3693836.25 | 3221105.5 | 4101495.5 | 3537083.5 | 1708581.25 | 1498440 | 3373667.75 | 2730449.75 | 2044732.88 | 3536040.75 | 3086901.5 | 4460313.5 | 3702257 | 3301847.25 | 1 | 12 | 1 | 10 | 1 | 12 | 0.63 | 0.61 | 1.01 | 0.97 |
| Q8C1A5 | Thop1 | Thimet oligopeptidase | 308.61 | 9.17 | 78026 | 6 | 5584359 | 5270105 | 3934341 | 4058996.5 | 5734575 | 3956735.5 | 5067787 | 6504592 | 5260157.5 | 3445239.25 | 1828827 | 3815298 | 2370955 | 2625764.25 | 2223260 | 5681658 | 5168942.5 | 3650092.5 | 5101303 | 4008322 | 0 | 9 | 1 | 12 | 1 | 12 | 0.52 | 0.53 | 0.96 | 0.97 |
| Q9WTQ8 | Timm23 | Mitochondrial import inner membrane translocase subunit Tim23 | 213.54 | 22.49 | 21978 | 3 | 1847632.75 | 1751525.25 | 2307177.75 | 2294477 | 2387046.25 | 2239514.5 | 1505880.12 | 2277688 | 2088310.5 | 2608296.75 | 2984624.5 | 2435659.5 | 2740435.75 | 3736102.5 | 3233737.75 | 1600146 | 2482868 | 2842089.75 | 2431995.5 | 2657136.5 | 0 | 5 | 1 | 6 | 1 | 6 | 1.43 | 1.41 | 1.13 | 1.12 |
| Q9D880 | Timm50 | Mitochondrial import inner membrane translocase subunit TIM50 | 492.39 | 25.78 | 39776 | 8 | 22266716 | 22571318 | 23017344 | 17028398 | 26085900 | 21977838 | 25395480 | 20585894 | 22458416 | 15344472 | 13055034 | 19934690 | 11990790 | 12937207 | 14904554 | 24423870 | 15623731 | 13176702 | 19356810 | 16248049 | 1 | 5 | 1 | 7 | 1 | 7 | 0.66 | 0.69 | 0.80 | 0.84 |
| P26039 | Tln1 | Talin-1 | 7906.86 | 56.87 | 269821 | 114 | 364391874 | 383670498 | 427454530 | 454319330 | 403382498 | 389375746 | 427172450 | 395900578 | 391508738 | 370266338 | 272798882 | 370026722 | 298986882 | 298096002 | 280454658 | 371571490 | 389903138 | 390702082 | 588120002 | 415774850 | 1 | 12 | 0 | 11 | 1 | 12 | 0.75 | 0.77 | 1.06 | 1.09 |
| Q9QUN7 | Tlr2 | Toll-like receptor 2 | 843 | 22.07 | 89449 | 14 | 2888974.75 | 3942519.5 | 4686806 | 4625219.5 | 3821702 | 3835734.5 | 3117337.25 | 3318860.5 | 4562523.5 | 4257348.5 | 7781609.5 | 5565905 | 5563690 | 4671440 | 5993425.5 | 2761803.25 | 4445320.5 | 3319632.5 | 1377558.75 | 4284962 | 1 | 7 | 0 | 10 | 1 | 10 | 1.48 | 1.55 | 0.81 | 0.85 |
| P58681 | Tlr7 | Toll-like receptor 7 | 656.46 | 12.48 | 121837 | 11 | 4559862 | 3675712.75 | 6031433 | 3394438.75 | 6692631 | 3174893 | 3943359.75 | 2068307.88 | 4392243 | 2795030.25 | 1411709 | 1404682 | 1941228.75 | 1770185 | 2178378.25 | 3319142 | 1487698.25 | 1816245.38 | 1915010.25 | 1446163.5 | 0 | 0 | 1 | 3 | 1 | 3 | 0.36 | 0.53 | 0.41 | 0.61 |
| Q9ET30 | Tm9sf3 | Transmembrane 9 superfamily member 3 | 351.12 | 16.87 | 67545 | 8 | 8369321.5 | 6508217.5 | 6655307.5 | 8137158 | 8554881 | 6409926 | 8455047 | 7489752 | 9731092 | 7953941.5 | 8512144 | 11143944 | 12879708 | 10080441 | 12893246 | 8932684 | 4969304.5 | 7563403.5 | 6554831 | 6217084.5 | 1 | 8 | 1 | 7 | 1 | 8 | 1.45 | 1.39 | 0.90 | 0.86 |
| Q9CR64 | Tmem167a | Protein kish-A | 109.39 | 25 | 8074 | 2 | 2371256.25 | 2697340.25 | 2536733 | 2193765.5 | 2167969.25 | 2441662 | 1822424 | 2691341.5 | 2974276.5 | 3301230.75 | 3291546.75 | 3668902 | 6576632 | 6070472.5 | 5718173 | 2126870.5 | 2152006 | 3249964.5 | 2851435.75 | 3024925.5 | 0 | 10 | 1 | 12 | 1 | 12 | 2.12 | 1.91 | 1.12 | 1.01 |
| Q9R1Q6 | Tmem176b | Transmembrane protein 176B | 232.86 | 13.31 | 28367 | 4 | 926599.25 | 779227.81 | 1444292.5 | 2153564.5 | 1977967.12 | 989472.19 | 1074480.75 | 2398171.25 | 1493185.5 | 1908020.75 | 2394531.5 | 2916016 | 4412176.5 | 3035488.75 | 2903080.75 | 789012.62 | 667135.69 | 1563275.12 | 1516792.75 | 1443976.25 | 0 | 9 | 1 | 8 | 1 | 9 | 2.15 | 1.99 | 0.82 | 0.76 |
| Q8BM55 | Tmem214 | Transmembrane protein 214 | 303.73 | 10.33 | 76430 | 6 | 2511938.5 | 1815017.5 | 1770423.5 | 1845143.25 | 1873291.88 | 1758638.5 | 2364171 | 1696120.75 | 1930323 | 1763541.75 | 2205714.75 | 2813785.75 | 3386112.25 | 2548880.75 | 3784878.25 | 2219188.5 | 508232.59 | 2575419 | 2631999 | 1725058.12 | 1 | 11 | 1 | 10 | 1 | 11 | 1.50 | 1.55 | 0.98 | 1.02 |
| Q9DAV9 | Tmem38b | Trimeric intracellular cation channel type B | 89.35 | 9.59 | 32640 | 2 | 783306.31 | 1076811.5 | 526068 | 1171907.5 | 741440.31 | 1890396.38 | 1917870.62 | 1300335.25 | 721151.19 | 915288.56 | 365880.03 | 648338.69 | 2 | 2 | 2 | 1503081.62 | 1085745.12 | 804934.88 | 2 | 2 | 1 | 12 | 0 | 6 | 1 | 12 | 0.24 | 0.15 | 0.79 | 0.50 |
| Q8BXZ1 | Tmx3 | Protein disulfide-isomerase TMX3 | 482.09 | 23.46 | 51848 | 8 | 2697409 | 2095425.25 | 3656439.5 | 3165329.5 | 3767248.5 | 2831280.75 | 3411983 | 3776667.75 | 3566646 | 2703025.25 | 4031658.25 | 4440919 | 5134994 | 3663860.5 | 5395752 | 2808078.75 | 2131037.25 | 4062352.5 | 2852462.75 | 4209042 | 1 | 10 | 1 | 12 | 1 | 12 | 1.47 | 1.39 | 1.04 | 0.99 |
| F6ZDS4 | Tpr | Nucleoprotein TPR | 1488.92 | 12.01 | 273990 | 25 | 3043812 | 3128306 | 2892832 | 3059373 | 2259754 | 3035505 | 2275177 | 3528124 | 2993972 | 3530629 | 1920891.5 | 771862.62 | 2744544.25 | 1703949.12 | 1416281.12 | 2229034.25 | 1532764.38 | 2912381.5 | 3956295 | 2547340.5 | 1 | 8 | 1 | 7 | 1 | 8 | 0.59 | 0.56 | 0.92 | 0.86 |
| Q9D6K7 | Ttc33 | Tetratricopeptide repeat protein 33 | 70.89 | 4.96 | 29371 | 1 | 775487.75 | 943554.69 | 965418.88 | 1129877.25 | 953504.12 | 1069394.5 | 1001955.19 | 721082.44 | 839861 | 870492.5 | 808153.81 | 681535.25 | 611224.62 | 459759.19 | 619335.44 | 765483.88 | 1183430.25 | 800745.88 | 715542.56 | 722205.5 | 1 | 8 | 1 | 7 | 1 | 8 | 0.67 | 0.71 | 0.88 | 0.93 |
| Q4QRB4 | Tubb3 | Tubulin beta-3 chain | 1686.73 | 38.89 | 50419 | 25 | 950453.62 | 923385.88 | 737481.88 | 848886.62 | 923146.81 | 1123720 | 1104887.75 | 957678.12 | 1045370.75 | 801635.62 | 275100.69 | 2 | 744041.88 | 395539.47 | 460658.84 | 785347.69 | 1367593.62 | 773293.94 | 1094668.88 | 613333.81 | 1 | 12 | 0 | 8 | 1 | 12 | 0.43 | 0.37 | 1.06 | 0.92 |
| Q498E0 | Txndc12 | Thioredoxin domain-containing protein 12 | 143.72 | 22.35 | 19019 | 3 | 967360.75 | 1294911 | 2 | 1045005.38 | 353950.06 | 684772.44 | 1155173.25 | 450806.84 | 2 | 1849042 | 1983154 | 1455474.75 | 2213132.75 | 2011255.75 | 2633908.5 | 955939.06 | 682766.94 | 1056607.75 | 889939.31 | 1496925.12 | 0 | 10 | 1 | 10 | 1 | 10 | 2.81 | 2.49 | 1.39 | 1.23 |
| Q922Y1 | Ubxn1 | UBX domain-containing protein 1 | 295.92 | 17.85 | 33573 | 5 | 1149114.38 | 628840.19 | 1651762.25 | 1375892.25 | 133887.5 | 584431.38 | 681691.88 | 1882278.38 | 1211611.62 | 1204768.88 | 3283528.5 | 3089872.5 | 2045153.75 | 1896686.5 | 1475845 | 1973369.12 | 3614178.25 | 1953563.88 | 726638.81 | 1880239.25 | 1 | 7 | 1 | 9 | 1 | 9 | 2.39 | 2.12 | 1.40 | 1.24 |
| Q9JKB1 | Uchl3 | Ubiquitin carboxyl-terminal hydrolase isozyme L3 | 202.41 | 15.65 | 26152 | 3 | 4094560.5 | 4466292 | 4289601.5 | 4440666 | 3418284.5 | 4672345 | 4301939.5 | 5010154 | 3837831.75 | 4053465.5 | 2062258.12 | 2827663 | 2807361.25 | 2777464.5 | 3442978.5 | 4877162 | 5787133 | 5031195 | 4499798.5 | 4325064 | 1 | 2 | 0 | 5 | 1 | 5 | 0.67 | 0.64 | 1.18 | 1.12 |
| Q91YL3 | Uckl1 | Uridine-cytidine kinase-like 1 | 45.09 | 1.46 | 60842 | 1 | 149231.95 | 569288.5 | 596598.5 | 651782.75 | 638175.62 | 196874.72 | 297555.97 | 638500.56 | 913490.75 | 833132.25 | 1182787.38 | 983620.69 | 839694.62 | 932796.25 | 1240416 | 164171.27 | 631276.44 | 806736.94 | 867626.94 | 943344.81 | 0 | 6 | 1 | 11 | 1 | 11 | 1.99 | 1.80 | 1.31 | 1.19 |
| Q8CCJ3 | Ufl1 | E3 UFM1-protein ligase 1 | 252.81 | 10.21 | 89520 | 5 | 1440844.38 | 908792.5 | 1095232.75 | 1084736.62 | 1239839.38 | 632074.5 | 1239599.75 | 1572877.38 | 1452905.38 | 982697.31 | 2308764.5 | 1685610.5 | 1631462.75 | 1862654.62 | 1564454.25 | 1896383.5 | 1271037 | 1866804.5 | 1297193.5 | 1173703.25 | 0 | 4 | 1 | 7 | 1 | 7 | 1.57 | 1.54 | 1.30 | 1.28 |
| Q6P5E4 | Uggt1 | UDP-glucose:glycoprotein glucosyltransferase 1 | 2837.679 | 40.1 | 176434 | 48 | 17573274 | 20983830 | 25060222 | 22836502 | 27461082 | 17572300 | 19367838 | 37924546 | 23408338 | 22899904 | 42935366 | 37941030 | 36196442 | 39393822 | 41769666 | 30768892 | 13183063 | 22117168 | 19600098 | 17753384 | 0 | 6 | 1 | 5 | 1 | 6 | 1.74 | 1.64 | 0.81 | 0.76 |
| Q8VDF2 | Uhrf1 | E3 ubiquitin-protein ligase UHRF1 | 619.72 | 18.93 | 88304 | 11 | 1521471.25 | 1235918.75 | 2303327 | 1830958.5 | 1954379 | 1734090.5 | 1607617.38 | 1460947 | 1917865.25 | 1733488.88 | 1450707.88 | 1197161.75 | 1037818.88 | 1137527.62 | 1202489.5 | 1240414.25 | 515538.16 | 1731079 | 1126336.25 | 1733958.38 | 1 | 5 | 0 | 5 | 1 | 5 | 0.68 | 0.71 | 0.72 | 0.75 |
| Q9CWU6 | Uqcc1 | Ubiquinol-cytochrome-c reductase complex assembly factor 1 | 186.32 | 10.51 | 34300 | 3 | 3518964.25 | 3381768.5 | 2588719.5 | 2011969.38 | 3267102 | 2456047 | 3907423.5 | 2392539.75 | 3218082.25 | 1950307.5 | 892420.44 | 2027782.25 | 1019839.75 | 1122364.25 | 1193738.5 | 3168842.5 | 1166184.75 | 1528653.5 | 982111.69 | 1262096.12 | 1 | 2 | 1 | 3 | 1 | 3 | 0.42 | 0.45 | 0.55 | 0.58 |
| P41542 | Uso1 | General vesicular transport factor p115 | 332.77 | 7.3 | 107162 | 7 | 4133280.5 | 6151225.5 | 4011338.5 | 4600044 | 4169569.25 | 4723475 | 5047672.5 | 5486432 | 4950103.5 | 5135330 | 7694405.5 | 7110123 | 7725306 | 7290339 | 5878338 | 6774903 | 4088854 | 5634967 | 4791233.5 | 5474425.5 | 1 | 7 | 0 | 10 | 1 | 10 | 1.55 | 1.41 | 1.16 | 1.06 |
| Q8C7V3 | Utp15 | U3 small nucleolar RNA-associated protein 15 homolog | 245.67 | 10.04 | 59375 | 5 | 2457419 | 3096592.75 | 3619165.5 | 2954373.75 | 3319631.25 | 3703848.5 | 4004879.5 | 2253348.5 | 2687517.5 | 1621797.75 | 4093532.25 | 4882541 | 4623868 | 4266883 | 3907862.5 | 4688559 | 2877398.5 | 2121874 | 1409428.38 | 2469480.25 | 0 | 7 | 1 | 11 | 1 | 11 | 1.41 | 1.53 | 0.88 | 0.95 |
| Q9R0C8 | Vav3 | Guanine nucleotide exchange factor VAV3 | 227.49 | 3.78 | 97968 | 4 | 1349489.88 | 1052709.75 | 1696430.88 | 1737423.38 | 1674872.88 | 1716416.62 | 1281913.12 | 1657558.75 | 2102974.5 | 1472897.12 | 743395.19 | 611824.94 | 993360.56 | 1262519.75 | 986387.25 | 1159935.62 | 1447178.5 | 1400253.25 | 1445472.25 | 1408097.75 | 1 | 9 | 0 | 4 | 1 | 9 | 0.61 | 0.56 | 0.91 | 0.83 |
| Q60930 | Vdac2 | Voltage-dependent anion-selective channel protein 2 | 1125.74 | 64.07 | 31733 | 17 | 196537442 | 209380466 | 186308658 | 205320002 | 191581698 | 236378466 | 223776658 | 234575538 | 217723074 | 228284386 | 254540242 | 236025938 | 282303778 | 278693570 | 276677826 | 202572610 | 193546914 | 244126194 | 243741954 | 244090386 | 0 | 5 | 1 | 10 | 1 | 10 | 1.34 | 1.16 | 1.14 | 0.99 |
| Q6TEK3 | Vkorc1l1 | Vitamin K epoxide reductase complex subunit 1-like protein 1 | 65.58 | 6.25 | 19779 | 1 | 2 | 2 | 95691.94 | 2 | 251327.27 | 229610.09 | 142357.86 | 552914.88 | 464456.78 | 426546.97 | 538409.12 | 772853.69 | 855542.12 | 874415.88 | 948482.5 | 2 | 2 | 597613.31 | 2 | 566688.25 | 0 | 10.5 | 1 | 10 | 1 | 10.5 | 11.50 | 2.20 | 3.36 | 0.64 |
| Q8BYL4 | Yars2 | Tyrosine--tRNA ligase, mitochondrial | 67.19 | 2.54 | 52597 | 1 | 292642.97 | 398422.78 | 383249.75 | 317815.78 | 373712.19 | 423366.12 | 319955.69 | 349734.22 | 416304.97 | 233900.98 | 422311 | 496498.62 | 632156.81 | 578202.81 | 555907.88 | 362938.44 | 391451.41 | 271406.25 | 2 | 347446.19 | 0 | 8 | 1 | 9 | 1 | 9 | 1.52 | 1.54 | 0.78 | 0.79 |
| Q91X58 | Zfand2b | AN1-type zinc finger protein 2B | 83.73 | 8.56 | 27895 | 2 | 720935.12 | 655820.38 | 239000.3 | 856556.12 | 965126 | 279772.44 | 338141 | 770406.25 | 631804.62 | 433814.12 | 92448.66 | 202936.75 | 103927.91 | 274257.38 | 117574.75 | 205704.64 | 2 | 839195.25 | 918703.06 | 191608.36 | 1 | 7 | 0 | 10 | 1 | 10 | 0.23 | 0.32 | 0.63 | 0.88 |
| Q5U2Q3 | | Ester hydrolase C11orf54 homolog | 140.75 | 7.62 | 34993 | 3 | 3892808.5 | 4236515.5 | 4328884.5 | 6043029 | 4995700.5 | 4484480 | 3440668.5 | 2235540 | 4208480 | 4595630 | 2049414.25 | 2056728 | 2055007.75 | 3259545 | 1439517.88 | 4825769 | 5777268 | 4943061.5 | 5888461 | 5392283 | 0 | 7 | 1 | 1 | 1 | 7 | 0.46 | 0.57 | 1.14 | 1.41 |
| Q6PIU9 | | Uncharacterized protein FLJ45252 homolog | 66.76 | 3.39 | 37705 | 1 | 1244333.12 | 1360104.5 | 1490624.12 | 1551362 | 1096650 | 1212925 | 1391489 | 1482986.62 | 1154820.25 | 1607568.38 | 710716.06 | 701748.06 | 737789.44 | 1112046.12 | 982075.81 | 1286719.38 | 1339875.12 | 1998855.88 | 2306265.75 | 1731470.75 | 1 | 6 | 0 | 6 | 1 | 6 | 0.63 | 0.62 | 1.28 | 1.26 |
| P0C1X8 | Aak1 | AP2-associated protein kinase 1 | 92.63 | 2.91 | 103761 | 2 | 751131.5 | 861356.56 | 1313063.88 | 769912.19 | 925448.12 | 891970.88 | 724858.19 | 1145019.75 | 1303018.5 | 656801.88 | 464786.66 | 528626.75 | 398620.97 | 749964.44 | 2 | 2 | 601887.31 | 962370 | 1107653.88 | 1182037.38 | 1 | 12 | 0 | 10 | 2 | 12 | 0.46 | 0.45 | 0.83 | 0.82 |
| Q99LE6 | Abcf2 | ATP-binding cassette sub-family F member 2 | 192.56 | 7.64 | 71782 | 4 | 2303904.5 | 3130276.25 | 3072528.25 | 2674455.25 | 1796052 | 1817489.62 | 1458207.5 | 2525999.25 | 1439237.38 | 2512478 | 3386342.5 | 4271862 | 4888278 | 3023831.5 | 3361912.5 | 2069335.75 | 1333197.25 | 2697136.5 | 2199707.25 | 2128842.25 | 2 | 6 | 0 | 11 | 2 | 11 | 1.46 | 1.94 | 0.80 | 1.07 |
| Q8CBW3 | Abi1 | Abl interactor 1 | 184.31 | 9.15 | 52288 | 4 | 4543707.5 | 5116722.5 | 4458697.5 | 2476582.5 | 7211364 | 2204648.75 | 5048914 | 3960553.5 | 5673222 | 2357758 | 623636.25 | 3057447 | 1031224.56 | 357051.44 | 1646344.62 | 4977649.5 | 1997275.25 | 2138443.75 | 1779962.5 | 2423521.5 | 1 | 3 | 2 | 5 | 2 | 5 | 0.28 | 0.35 | 0.56 | 0.69 |
| P45952 | Acadm | Medium-chain specific acyl-CoA dehydrogenase, mitochondrial | 471.07 | 28.27 | 46481 | 10 | 16555526 | 21097706 | 20319710 | 26387190 | 24250584 | 23797926 | 19384488 | 20730774 | 23363098 | 26513874 | 8270995.5 | 12266707 | 15937865 | 18617272 | 20106558 | 21444492 | 22173790 | 26162712 | 23046294 | 24281606 | 2 | 8 | 1 | 11 | 2 | 11 | 0.69 | 0.66 | 1.08 | 1.03 |
| Q8QZT1 | Acat1 | Acetyl-CoA acetyltransferase, mitochondrial | 850.68 | 43.16 | 44816 | 13 | 26204268 | 27220776 | 37970174 | 36559342 | 30341418 | 32688850 | 28288316 | 32972544 | 34100694 | 35013850 | 16894826 | 17765628 | 24847262 | 28959072 | 24038362 | 25668278 | 25894530 | 36957378 | 31649110 | 32498500 | 2 | 10 | 1 | 11 | 2 | 11 | 0.71 | 0.69 | 0.96 | 0.94 |
| Q8CAY6 | Acat2 | Acetyl-CoA acetyltransferase, cytosolic | 365.41 | 24.94 | 41298 | 6 | 14564278 | 17322250 | 13904134 | 16146889 | 13305429 | 18757684 | 13281943 | 13431818 | 12677805 | 12988482 | 11740502 | 11273730 | 12236470 | 13253706 | 11692324 | 17763148 | 17505086 | 19557784 | 16864940 | 20933670 | 0 | 1 | 2 | 3 | 2 | 3 | 0.90 | 0.85 | 1.23 | 1.30 |
| Q9CZW4 | Acsl3 | Long-chain-fatty-acid--CoA ligase 3 | 1163.46 | 37.08 | 80492 | 21 | 22665738 | 26771378 | 19117168 | 23176984 | 22844330 | 25316566 | 28297000 | 19276960 | 19943004 | 20877052 | 29094542 | 24873816 | 31847766 | 30600920 | 30825496 | 24952532 | 26800570 | 26447298 | 25128348 | 24583282 | 1 | 4 | 2 | 8 | 2 | 8 | 1.29 | 1.29 | 1.12 | 1.12 |
| Q99JW2 | Acy1 | Aminoacylase-1 | 137.59 | 12.75 | 45781 | 3 | 652034.56 | 883274.12 | 813575.31 | 736385.38 | 648893 | 1104219.25 | 474431.75 | 2 | 485066.56 | 1246737 | 1968802.12 | 1610324.62 | 1079691.62 | 1741335.38 | 1370829.62 | 755020.06 | 871366.75 | 1194566.62 | 1128399.75 | 832918.88 | 0 | 4 | 2 | 8 | 2 | 8 | 2.08 | 2.35 | 1.28 | 1.44 |
| Q8CHT0 | Aldh4a1 | Delta-1-pyrroline-5-carboxylate dehydrogenase, mitochondrial | 45.12 | 3.38 | 61841 | 1 | 308800.81 | 163457.16 | 189896.17 | 149361.06 | 163490.69 | 171352.52 | 216010.88 | 161437.25 | 203621.31 | 58791.71 | 2 | 117141.09 | 113651.38 | 2 | 49082.12 | 269982.09 | 61557.95 | 89877.54 | 2 | 29709.31 | 0 | 4 | 2 | 7 | 2 | 7 | 0.29 | 0.35 | 0.46 | 0.56 |
| O08739 | Ampd3 | AMP deaminase 3 | 108.6 | 2.61 | 88652 | 2 | 2 | 275841.75 | 445564.69 | 509218.06 | 767836.69 | 2 | 2 | 512972.81 | 966341.88 | 618622.75 | 748828.62 | 1006023.31 | 1099069.12 | 947782.75 | 1151622.88 | 2 | 2 | 751337 | 472971.81 | 355352.84 | 1 | 10 | 2 | 10 | 2 | 10 | 2.48 | 2.36 | 0.79 | 0.75 |
| Q78IK4 | Apool | MICOS complex subunit Mic27 | 433.13 | 27.92 | 29261 | 6 | 10763256 | 10324312 | 10618502 | 10179094 | 10640308 | 12087951 | 11637010 | 10536372 | 10380883 | 9318317 | 6968096 | 7585226.5 | 9436677 | 6036326 | 9584420 | 9387035 | 11838509 | 9529942 | 13245232 | 9422321 | 0 | 10 | 2 | 12 | 2 | 12 | 0.75 | 0.73 | 1.02 | 0.99 |
| O55043 | Arhgef7 | Rho guanine nucleotide exchange factor 7 | 156.7 | 7.59 | 73140 | 3 | 2288450.25 | 1883114 | 1764299.25 | 1128315.12 | 1674032.75 | 1586991.5 | 2364424.5 | 1161651.12 | 2009553.25 | 1182007.62 | 1058506.62 | 1492338.38 | 1123380.25 | 424721.34 | 507937.38 | 2041622.75 | 1016375.75 | 1334076.5 | 1203312.25 | 1141759.5 | 1 | 7 | 2 | 8 | 2 | 8 | 0.53 | 0.55 | 0.77 | 0.81 |
| Q9WV54 | Asah1 | Acid ceramidase | 914.02 | 39.59 | 44670 | 15 | 10032551 | 9483134 | 12095022 | 11631176 | 10700058 | 9734821 | 9004514 | 8800062 | 10457309 | 10576928 | 7946214.5 | 7091462 | 9203370 | 7335724 | 7416714.5 | 6994265 | 9057073 | 9850942 | 10922684 | 8035628 | 0 | 4 | 2 | 10 | 2 | 10 | 0.72 | 0.80 | 0.83 | 0.92 |
| Q8C0M9 | Asrgl1 | Isoaspartyl peptidase/L-asparaginase | 200.55 | 10.12 | 33950 | 3 | 3246340.75 | 3407937 | 5489696.5 | 6247459.5 | 5462548.5 | 3874943.5 | 3763051 | 5260210 | 5187130.5 | 6155852 | 1714815.25 | 1842438 | 2685574 | 3444235 | 1311502 | 4174999.75 | 5568604 | 5833915 | 7343994 | 6052836.5 | 2 | 6 | 2 | 6 | 2 | 6 | 0.45 | 0.46 | 1.21 | 1.20 |
| P97370 | Atp1b3 | Sodium/potassium-transporting ATPase subunit beta-3 | 353.62 | 41.37 | 31776 | 7 | 18224662 | 14946475 | 16402326 | 18606252 | 14984770 | 18422382 | 15571598 | 14491956 | 14943833 | 18553568 | 20196828 | 18120950 | 21763448 | 23313734 | 23312618 | 13911182 | 18365040 | 16030410 | 13427631 | 16457997 | 2 | 9 | 2 | 9 | 2 | 9 | 1.28 | 1.30 | 0.94 | 0.95 |
| P56480 | Atp5f1b | ATP synthase subunit beta, mitochondrial | 3291.26 | 82.61 | 56300 | 46 | 328098754 | 304828194 | 362188098 | 333184738 | 364119714 | 327507426 | 292508834 | 402535170 | 355526402 | 369485410 | 383475714 | 395858978 | 514923746 | 489951106 | 518279810 | 318385090 | 328095682 | 378541026 | 337576258 | 373592930 | 0 | 10 | 2 | 12 | 2 | 12 | 1.36 | 1.32 | 1.03 | 0.99 |
| Q91VR2 | Atp5f1c | ATP synthase subunit gamma, mitochondrial | 681.95 | 41.61 | 32886 | 11 | 21448328 | 21415392 | 27669524 | 27411542 | 24160046 | 26044114 | 20811692 | 33378534 | 24969740 | 34403098 | 40006862 | 30622354 | 40935542 | 44161346 | 46866194 | 22689466 | 24426624 | 34245358 | 34104162 | 29499408 | 0 | 5 | 2 | 6 | 2 | 6 | 1.66 | 1.45 | 1.19 | 1.04 |
| P35434 | Atp5f1d | ATP synthase subunit delta, mitochondrial | 130.32 | 13.69 | 17595 | 2 | 22896798 | 23222358 | 26522842 | 27974298 | 22762096 | 23202274 | 22157522 | 25487514 | 24270808 | 26327454 | 29590250 | 24443698 | 30642502 | 31920642 | 31453250 | 21308748 | 26389202 | 27720842 | 26426546 | 24269120 | 2 | 12 | 2 | 8 | 2 | 12 | 1.20 | 1.22 | 1.02 | 1.04 |
| O55099 | Aurkb | Aurora kinase B | 54.54 | 2.92 | 39234 | 1 | 313606.66 | 412414.94 | 498848.53 | 195868.14 | 453747.72 | 273312.66 | 227879.3 | 307634.72 | 504960.97 | 300610 | 206467.17 | 2 | 228473.78 | 2 | 2 | 2 | 2 | 317109.22 | 2 | 255875.11 | 2 | 3 | 1 | 5 | 2 | 5 | 0.23 | 0.27 | 0.31 | 0.35 |
| Q6MG49 | Bag6 | Large proline-rich protein BAG6 | 185.4 | 3.66 | 120012 | 3 | 1679902.88 | 2133678.75 | 1889555.75 | 877168.12 | 1769627.62 | 509804.97 | 2349261.5 | 1662198.12 | 2997992.5 | 1219981.25 | 230423.88 | 492289.22 | 817690.31 | 2 | 1082244.5 | 1156720.25 | 996902.25 | 687448.88 | 916249.25 | 1181538.25 | 1 | 4 | 2 | 5 | 2 | 5 | 0.31 | 0.30 | 0.59 | 0.57 |
| O54962 | Banf1 | Barrier-to-autointegration factor | 259.47 | 52.81 | 10103 | 4 | 10598436 | 9011016 | 14254400 | 14565587 | 15305168 | 10615507 | 11217534 | 13989371 | 13986866 | 13936608 | 6724573.5 | 5618558 | 10235035 | 7481617 | 9347490 | 12392250 | 11550010 | 15967121 | 11576564 | 14718053 | 2 | 10 | 0 | 9 | 2 | 10 | 0.62 | 0.62 | 1.04 | 1.02 |
| O08539 | Bin1 | Myc box-dependent-interacting protein 1 | 72.24 | 4.59 | 64470 | 2 | 4157506.5 | 3862174 | 3264043.25 | 3437358 | 3223071.5 | 4200793.5 | 3632246.5 | 2921400.5 | 3057574 | 3228750 | 3082021.5 | 2679215.5 | 2398512.25 | 2617835.5 | 2607420.75 | 4310908.5 | 5144775 | 3176149.75 | 3810881 | 3538203.75 | 0 | 9 | 2 | 6 | 2 | 9 | 0.75 | 0.79 | 1.11 | 1.17 |
| Q80Y55 | Bsdc1 | BSD domain-containing protein 1 | 131.84 | 7.26 | 46953 | 2 | 208326.16 | 713448.81 | 594767.19 | 451121.25 | 432251.94 | 391390.5 | 690462.38 | 320638.03 | 665693.5 | 513951.84 | 222754.75 | 2 | 265128.97 | 151229.44 | 141569.52 | 330918.25 | 156453.83 | 725670.75 | 371800.41 | 509307 | 2 | 10 | 0 | 9 | 2 | 10 | 0.33 | 0.30 | 0.87 | 0.81 |
| Q64152 | Btf3 | Transcription factor BTF3 | 267.65 | 33.82 | 22031 | 4 | 2 | 371407.34 | 1088299.12 | 1240705.38 | 664365.12 | 600144.12 | 329985.41 | 617994.25 | 456508.38 | 542940.56 | 1159616.38 | 1160822.5 | 2085161 | 2547377.5 | 1916867.5 | 1623086 | 1833734.12 | 1485502.75 | 1652312.88 | 1747117.62 | 2 | 0 | 0 | 2 | 2 | 2 | 2.64 | 3.49 | 2.48 | 3.28 |
| O08529 | Capn2 | Calpain-2 catalytic subunit | 516.64 | 19.29 | 79872 | 10 | 6698480.5 | 5953105 | 4311564.5 | 4285090 | 7881370.5 | 5359264 | 6396876.5 | 4802521.5 | 7413379 | 3991678.25 | 4408469 | 3782247 | 3816807 | 2688660 | 2557550.25 | 5292069.5 | 4648910.5 | 3651381.5 | 3251504.5 | 3050047.5 | 2 | 4 | 1 | 3 | 2 | 4 | 0.59 | 0.62 | 0.68 | 0.71 |
| Q61686 | Cbx5 | Chromobox protein homolog 5 | 262.84 | 19.37 | 22186 | 4 | 2025084.25 | 2683737.5 | 5088881 | 3586381 | 3396031.5 | 2932006 | 3068397.25 | 3600424.75 | 2997733 | 3293764.75 | 2204987.5 | 2645630.5 | 1109065.38 | 1504620.75 | 1821255.75 | 2163299.5 | 3095867 | 3661661.5 | 3137389 | 2359425.25 | 2 | 10 | 0 | 11 | 2 | 11 | 0.55 | 0.58 | 0.86 | 0.91 |
| Q3URS9 | Ccdc51 | Mitochondrial potassium channel | 42.33 | 2.71 | 45132 | 1 | 148176.33 | 212491.25 | 2 | 261195.73 | 146049.83 | 250172.66 | 176588.73 | 327104.94 | 239003.53 | 243864.48 | 401227.12 | 349304.19 | 310800.06 | 271989.75 | 357688 | 227905.56 | 284536.88 | 238637.48 | 340868.94 | 293878.47 | 0 | 2 | 2 | 10 | 2 | 10 | 2.20 | 1.37 | 1.80 | 1.13 |
| Q9JLQ0 | Cd2ap | CD2-associated protein | 135.81 | 5.18 | 70450 | 3 | 3164551.75 | 3150614.25 | 3373515.75 | 2522259 | 1638692 | 2793771 | 3527361 | 3103920 | 3304581.5 | 2406082 | 1343720.75 | 1774949.25 | 1466846.75 | 960120.69 | 1743996 | 3146648.25 | 2397158.5 | 1897132.75 | 2306739 | 1984848 | 2 | 6 | 0 | 3 | 2 | 6 | 0.53 | 0.48 | 0.85 | 0.78 |
| P28659 | Celf1 | CUGBP Elav-like family member 1 | 160.88 | 6.58 | 52107 | 3 | 865261.88 | 734913.44 | 745036.25 | 872370.88 | 600001.44 | 906230.38 | 581172.19 | 663452.06 | 734091.81 | 627396.31 | 1389668 | 939876.62 | 979234.5 | 899674.75 | 872242.56 | 880577.31 | 1094080.62 | 772794.38 | 978676.38 | 937467.69 | 1 | 2 | 2 | 3 | 2 | 3 | 1.33 | 1.45 | 1.22 | 1.33 |
| Q499T7 | Cfap20 | Cilia- and flagella-associated protein 20 | 47.94 | 5.49 | 19132 | 1 | 420250.09 | 377187.91 | 495680.75 | 280988.38 | 622377.38 | 104615.73 | 592011.12 | 377426.44 | 554728.19 | 164516.72 | 2 | 303008.06 | 68334.57 | 2 | 2 | 283015 | 2 | 73445.95 | 2 | 127790.95 | 1 | 1 | 2 | 3 | 2 | 3 | 0.17 | 0.21 | 0.22 | 0.27 |
| Q8BIG7 | Comtd1 | Catechol O-methyltransferase domain-containing protein 1 | 372.41 | 41.22 | 28961 | 6 | 5550451 | 4390174 | 2861018.5 | 3596811 | 3763666.25 | 4555779 | 4514774 | 3654933 | 3389992.5 | 4001443 | 8755382 | 5453733 | 6245979.5 | 7598881 | 4628597 | 3880878 | 4534800 | 4036096.25 | 5051660.5 | 4237763 | 2 | 8 | 0 | 8 | 2 | 8 | 1.62 | 1.62 | 1.08 | 1.08 |
| Q9D2V7 | Coro7 | Coronin-7 | 303.11 | 5.97 | 100812 | 5 | 9053420 | 10865616 | 6785693.5 | 13387078 | 9046619 | 9451717 | 7881362.5 | 9787965 | 8731591 | 9752732 | 6007479.5 | 6176702.5 | 6958483.5 | 7741362.5 | 6268989 | 8896002 | 9479314 | 7546859 | 10548104 | 11020179 | 2 | 12 | 0 | 10 | 2 | 12 | 0.67 | 0.73 | 0.97 | 1.04 |
| P18395 | Csde1 | Cold shock domain-containing protein E1 | 767.53 | 22.68 | 88895 | 15 | 13324688 | 13540787 | 13771010 | 11039766 | 10099697 | 13382956 | 11447522 | 13586364 | 10761577 | 13809623 | 19450140 | 18738514 | 17474372 | 13413791 | 19336656 | 13286310 | 14525364 | 15547939 | 13829385 | 13714606 | 2 | 4 | 2 | 4 | 2 | 4 | 1.43 | 1.40 | 1.15 | 1.13 |
| Q61093 | Cybb | Cytochrome b-245 heavy chain | 839.23 | 29.3 | 65305 | 14 | 7644749.5 | 6439553 | 10833598 | 13610290 | 11816560 | 8851723 | 9517807 | 10575317 | 12291670 | 11611803 | 12012609 | 13311517 | 22758782 | 20157186 | 22831006 | 7401418.5 | 5422021.5 | 12068021 | 12187429 | 12963714 | 2 | 12 | 1 | 12 | 2 | 12 | 1.81 | 1.72 | 0.99 | 0.95 |
| Q8CBY8 | Dctn4 | Dynactin subunit 4 | 156.48 | 5.78 | 53057 | 2 | 2073636.5 | 1296095.25 | 2169572 | 2416345.25 | 2020182.25 | 1681754.12 | 1625209.5 | 1824548 | 2325559.75 | 2105989.75 | 1308593.62 | 490002.59 | 627527.62 | 1297798.62 | 1236672.62 | 1325548.75 | 1945475.5 | 2173865.75 | 2504719.25 | 2622946 | 2 | 9 | 0 | 8 | 2 | 9 | 0.50 | 0.52 | 1.06 | 1.11 |
| Q6Q899 | Ddx58 | Antiviral innate immune response receptor RIG-I | 240.08 | 6.8 | 105975 | 5 | 3704273.25 | 3759491 | 3360024.5 | 2085483 | 4515250 | 2445025.5 | 3720996.75 | 5571987.5 | 4484877 | 3936143.5 | 648171.25 | 2362976.75 | 2345742.25 | 539298.12 | 1750258.75 | 2240114.5 | 697283.5 | 1960737.25 | 1394739.5 | 2075040.75 | 2 | 1 | 0 | 0 | 2 | 1 | 0.44 | 0.38 | 0.48 | 0.42 |
| Q99LB2 | Dhrs4 | Dehydrogenase/reductase SDR family member 4 | 317.79 | 21.86 | 29885 | 5 | 5420507 | 5516992 | 4335593.5 | 5652825.5 | 3679009.5 | 6274157.5 | 4908779 | 3334603.5 | 4218530 | 4869705 | 6880136 | 5470969 | 6126124 | 6592027 | 6812585 | 6238955 | 6500336 | 3780289.75 | 5564403.5 | 4349295.5 | 2 | 8 | 2 | 9 | 2 | 9 | 1.30 | 1.35 | 1.07 | 1.12 |
| Q3USZ8 | Dipk2a | Deleted in autism protein 1 homolog | 116.92 | 5.35 | 49463 | 2 | 2 | 2 | 397896.81 | 226714.94 | 370995.53 | 2 | 265037.03 | 510568.91 | 597538.75 | 667352 | 502886.34 | 1327066.25 | 973958.88 | 910389.31 | 2 | 2 | 2 | 2 | 142025.61 | 271269.19 | 0 | 8 | 2 | 9 | 2 | 9 | 4.40 | 3.19 | 0.42 | 0.30 |
| O08749 | Dld | Dihydrolipoyl dehydrogenase, mitochondrial | 1330.55 | 42.63 | 54272 | 19 | 70020714 | 76878370 | 57145674 | 71355106 | 67879138 | 81577594 | 74153938 | 68227010 | 67463050 | 68787338 | 78200698 | 77967290 | 90110986 | 87822738 | 89216170 | 87098290 | 83602866 | 73668490 | 81040898 | 79327906 | 0 | 1 | 2 | 4 | 2 | 4 | 1.23 | 1.18 | 1.18 | 1.12 |
| Q498R3 | Dnajc10 | DnaJ homolog subfamily C member 10 | 228.02 | 8.32 | 90747 | 5 | 1369493 | 1847762.12 | 786174.94 | 1526362.25 | 545242 | 1860527.38 | 1640256 | 1677857.25 | 952675.31 | 2155436.5 | 2913996.5 | 1802018.25 | 3108334.25 | 2899967 | 2655206 | 1248649.5 | 1623875.75 | 1320857.38 | 991799.81 | 1267014.5 | 1 | 12 | 2 | 5 | 2 | 12 | 2.20 | 1.61 | 1.06 | 0.78 |
| Q642C0 | Dnajc8 | DnaJ homolog subfamily C member 8 | 104.9 | 9.09 | 29813 | 3 | 4354400.5 | 5994667 | 5928528 | 4735088 | 4878608.5 | 1629393.38 | 5512871.5 | 5041466.5 | 5530726 | 3367557 | 1519193 | 3160000.75 | 845380.12 | 1230680.5 | 2384441.5 | 3804513.5 | 1538477.75 | 2723140.5 | 2289484 | 4020354.75 | 0 | 0 | 2 | 6 | 2 | 6 | 0.35 | 0.43 | 0.56 | 0.68 |
| Q8BVG4 | Dpp9 | Dipeptidyl peptidase 9 | 93.73 | 2.9 | 98001 | 2 | 944856.62 | 722211.81 | 656356.88 | 555557 | 1018953.38 | 634722.5 | 982295.81 | 803284.56 | 795669.5 | 224261.39 | 101126.98 | 554225.81 | 495323.19 | 106688.02 | 2 | 992497.38 | 857045.81 | 443120.25 | 715922.25 | 499796.56 | 0 | 9 | 2 | 12 | 2 | 12 | 0.32 | 0.37 | 0.90 | 1.02 |
| P70583 | Dut | Deoxyuridine 5'-triphosphate nucleotidohydrolase | 66.86 | 5.85 | 22003 | 1 | 2 | 315644.81 | 2 | 2 | 2 | 2 | 2 | 2 | 2 | 2 | 926151 | 393771.91 | 51107.13 | 151629.69 | 412534.69 | 2 | 2 | 2 | 2 | 2 | 2 | 10 | 0 | 12.5 | 2 | 12.5 | 6.13 | 193519.44 | 0.00 | 1.00 |
| Q3UJB9 | Edc4 | Enhancer of mRNA-decapping protein 4 | 522.49 | 7.75 | 152484 | 9 | 7991692 | 7875347 | 9350608 | 8552192 | 8658630 | 8079244 | 8093085.5 | 8713856 | 8554901 | 7415124.5 | 5799534 | 6969027.5 | 7638182.5 | 7639382 | 7359331 | 7600140 | 9313051 | 9223503 | 8368825.5 | 9317376 | 0 | 11 | 2 | 6 | 2 | 11 | 0.83 | 0.87 | 1.03 | 1.07 |
| Q91VC3 | Eif4a3 | Eukaryotic initiation factor 4A-III | 1330.12 | 50.12 | 46840 | 22 | 47629754 | 45397810 | 49000850 | 41542466 | 50426294 | 43652706 | 49422302 | 52445638 | 52945362 | 45960322 | 38884346 | 45033154 | 40377630 | 31224234 | 44650270 | 48865194 | 37435774 | 35312734 | 41539406 | 43012858 | 2 | 4 | 2 | 6 | 2 | 6 | 0.86 | 0.82 | 0.88 | 0.84 |
| B5DF91 | Elavl1 | ELAV-like protein 1 | 776.42 | 43.25 | 36169 | 15 | 34766586 | 31866640 | 32819010 | 33502792 | 29180594 | 36762278 | 34003106 | 27135548 | 26841338 | 30704490 | 29494540 | 25853546 | 23527778 | 21310742 | 23892150 | 33660830 | 39922698 | 30203866 | 31638644 | 32283148 | 1 | 12 | 2 | 9 | 2 | 12 | 0.77 | 0.80 | 1.03 | 1.08 |
| Q8CB77 | Eloa | Elongin-A | 102.91 | 5.43 | 87161 | 2 | 310114.16 | 339946.28 | 478889.5 | 317594.78 | 499651.5 | 355039.66 | 450818.81 | 450546.59 | 348876.03 | 263342.84 | 102164.05 | 330001.5 | 79621.67 | 216488.08 | 2 | 352338.78 | 146348.56 | 209268.62 | 300729.91 | 341288.5 | 2 | 6 | 1 | 4 | 2 | 6 | 0.37 | 0.39 | 0.69 | 0.72 |
| P62869 | Elob | Elongin-B | 421.8 | 49.15 | 13170 | 7 | 10117124 | 11082114 | 11185229 | 11492752 | 11318186 | 9141868 | 10096282 | 11598962 | 8021507.5 | 8515193 | 7229582 | 7636213 | 8266598 | 5479339 | 8369458 | 10867246 | 11081651 | 10892940 | 12324654 | 12227384 | 2 | 12 | 2 | 3 | 2 | 12 | 0.67 | 0.78 | 1.04 | 1.21 |
| P83940 | Eloc | Elongin-C | 145.55 | 17.86 | 12473 | 1 | 1388173 | 1240285.38 | 1037827 | 848497.56 | 1216629.5 | 1177662.88 | 1549580.5 | 1239994.62 | 1081936.88 | 1011246.12 | 503042.78 | 784307.25 | 891560.31 | 523465.25 | 1030187.38 | 1091573.12 | 1079793.75 | 910376.19 | 1016080.25 | 935421.5 | 2 | 7 | 1 | 4 | 2 | 7 | 0.65 | 0.62 | 0.88 | 0.83 |
| Q6P6T4 | Eml2 | Echinoderm microtubule-associated protein-like 2 | 89.11 | 4.78 | 70711 | 2 | 1969360.75 | 1691764.62 | 2002754.88 | 2089358.88 | 2504038 | 1593188.5 | 1930653 | 2120536.25 | 1854927.5 | 1967721.62 | 1378754.5 | 330893.44 | 1791958.88 | 1088378.88 | 1757108.25 | 2057380.5 | 995826 | 2375247 | 1986140.5 | 2268018.25 | 1 | 12 | 2 | 7 | 2 | 12 | 0.62 | 0.67 | 0.94 | 1.02 |
| Q80TH2 | Erbin | Erbin | 45.36 | 0.86 | 157248 | 1 | 503436.59 | 418326.12 | 650377.25 | 715211.25 | 545994.56 | 417698.62 | 378373.84 | 461853.22 | 506947.28 | 685470.06 | 418373.72 | 232612.19 | 263356.53 | 231893.73 | 299813.72 | 442198.59 | 589745.81 | 447277.34 | 516479.06 | 486257.12 | 1 | 8 | 2 | 10 | 2 | 10 | 0.51 | 0.59 | 0.88 | 0.85 |
| P57759 | Erp29 | Endoplasmic reticulum resident protein 29 | 525.23 | 43.89 | 28823 | 9 | 6102849 | 8488076 | 8270290 | 5082312.5 | 7252702.5 | 4310599.5 | 5652218.5 | 4488450 | 6211205 | 5150833.5 | 12503332 | 12514592 | 8984732 | 7726597.5 | 8493772 | 5623980 | 4721745 | 3698637.5 | 4587794 | 3901948 | 2 | 1 | 0 | 7 | 2 | 7 | 1.43 | 1.95 | 0.64 | 0.87 |
| P08101 | Fcgr2 | Low affinity immunoglobulin gamma Fc region receptor II | 334.24 | 29.7 | 36695 | 6 | 2 | 1566742.75 | 1805839.25 | 651330.38 | 1088704.75 | 2 | 950864.69 | 529951.75 | 508079.88 | 140311.62 | 1274329.38 | 2832895.25 | 2282229.75 | 1992983.12 | 2232350 | 2009672.25 | 837369.56 | 1357900 | 725059.12 | 1491630.75 | 2 | 10 | 0 | 2 | 2 | 10 | 2.08 | 4.99 | 1.26 | 3.02 |
| P24483 | Fdx1 | Adrenodoxin, mitochondrial | 97.76 | 9.04 | 20135 | 2 | 805551.12 | 1089451.38 | 496347.47 | 291768 | 1028971.19 | 357153.78 | 227476.5 | 1235398.5 | 356417.5 | 1115426.12 | 1335928 | 1057570.75 | 1445007.75 | 1547390.5 | 1595789.12 | 2 | 360429 | 1114900.25 | 368067.06 | 1126759.88 | 1 | 12 | 2 | 12 | 2 | 12 | 1.88 | 2.12 | 0.80 | 0.90 |
| Q80X90 | Flnb | Filamin-B | 409.11 | 3.42 | 277825 | 7 | 2099209.5 | 1296679 | 412703.06 | 1205954 | 894698.88 | 2061300.38 | 1625605.12 | 1129517 | 831458.12 | 2071396.5 | 2810910.5 | 2904485.75 | 2327471.75 | 1832218.38 | 2087464.75 | 1346720 | 2662957.5 | 2162883.25 | 2336921.5 | 2090743.12 | 2 | 2 | 2 | 3 | 2 | 3 | 2.02 | 1.55 | 1.79 | 1.37 |
| P70699 | Gaa | Lysosomal alpha-glucosidase | 890.68 | 20.99 | 106248 | 15 | 7947486 | 5898959 | 7483647.5 | 10927076 | 10831213 | 7626767.5 | 6155746 | 9448248 | 8858814 | 8887092 | 6615255.5 | 4312702.5 | 5445064 | 5713855 | 6156191 | 9879978 | 14970707 | 12796220 | 11582851 | 9029516 | 2 | 4 | 2 | 1 | 2 | 4 | 0.66 | 0.69 | 1.35 | 1.42 |
| Q9CZD3 | Gars | Glycine--tRNA ligase | 1369.61 | 32.24 | 81878 | 22 | 6380883 | 7186009 | 5970740 | 4144894 | 7662263 | 4526399.5 | 8420981 | 8818800 | 12161376 | 11452732 | 14122017 | 15854628 | 14825674 | 8861426 | 15453353 | 7226901 | 7395229 | 11421728 | 11333434 | 8445279 | 0 | 2 | 2 | 10 | 2 | 10 | 2.01 | 2.24 | 1.52 | 1.46 |
| O55148 | Gas7 | Growth arrest-specific protein 7 | 162.18 | 6.4 | 48275 | 4 | 3211542.5 | 2274289 | 5179481 | 3175095.5 | 3353446.25 | 2679248.5 | 2790475.5 | 2735770.25 | 2021708.12 | 1282789.25 | 1223111 | 1444036.62 | 761215.31 | 1325439.62 | 806602.31 | 2665401.5 | 2425076.5 | 1989926 | 1275721.88 | 1899649.88 | 0 | 2 | 2 | 6 | 2 | 6 | 0.32 | 0.48 | 0.60 | 0.89 |
| P17439 | Gba | Glucosylceramidase | 306.74 | 17.09 | 57622 | 6 | 3658209.75 | 4319497 | 2655343.25 | 2841338.75 | 3636878 | 4863366 | 4027177.75 | 4388757.5 | 3940661.25 | 5280667.5 | 6105018 | 5758624 | 6775150 | 4815694 | 5934506 | 3995267.25 | 3145922.5 | 4770864 | 2119991 | 3912580 | 0 | 10 | 2 | 4 | 2 | 10 | 1.72 | 1.31 | 1.05 | 0.80 |
| Q9JLQ2 | Git2 | ARF GTPase-activating protein GIT2 | 76.98 | 2.68 | 78766 | 2 | 1589794.62 | 1771317.75 | 2012959.5 | 2139891.5 | 2158053.75 | 1529414.12 | 918338.38 | 1677075.5 | 787419.31 | 1678205.62 | 366920.91 | 2 | 1031434.25 | 692321.31 | 613285.94 | 1801838.25 | 1463904.5 | 1519093.75 | 1964832.75 | 2 | 2 | 4 | 2 | 11 | 2 | 11 | 0.28 | 0.41 | 0.70 | 1.02 |
| P51569 | Gla | Alpha-galactosidase A | 523.65 | 24.11 | 47643 | 8 | 12717137 | 12254195 | 6878198.5 | 16050657 | 14430025 | 12027748 | 11342404 | 11550094 | 13878942 | 13755253 | 7479890 | 5911825.5 | 6431722 | 6694278.5 | 7049930 | 14994524 | 28215452 | 9662445 | 19055954 | 9255071 | 2 | 9 | 0 | 10 | 2 | 10 | 0.54 | 0.54 | 1.30 | 1.30 |
| Q9QUH0 | Glrx | Glutaredoxin-1 | 446.83 | 78.5 | 11871 | 7 | 16590774 | 18393680 | 20626622 | 18929708 | 28195360 | 12548680 | 18326058 | 17105410 | 23574208 | 17299522 | 8301836 | 13792263 | 12888739 | 11593634 | 12197627 | 23797268 | 14343936 | 17662294 | 15705932 | 19274464 | 0 | 8 | 2 | 11 | 2 | 11 | 0.57 | 0.66 | 0.88 | 1.02 |
| Q5RKH0 | Glyr1 | Putative oxidoreductase GLYR1 | 595.04 | 18.84 | 60421 | 8 | 4198848 | 2858047 | 4229403 | 4660015.5 | 5381122 | 5284244 | 3580306.25 | 3609398.5 | 3732485.5 | 5391298.5 | 2569613.5 | 2599059.75 | 3703618 | 2161565 | 3011373.75 | 3537101.5 | 4416767 | 3767762.5 | 3845067.5 | 2951375.25 | 2 | 7 | 2 | 9 | 2 | 9 | 0.66 | 0.65 | 0.87 | 0.86 |
| Q99LH1 | Gnl2 | Nucleolar GTP-binding protein 2 | 168.62 | 5.49 | 83345 | 3 | 436607.03 | 335824.34 | 2 | 413059.66 | 226759.77 | 855812.06 | 2 | 2 | 279102.22 | 316151.31 | 1512346.75 | 1021990.88 | 802155.12 | 978481 | 791115.19 | 2 | 2 | 519372.44 | 230893.27 | 356939.25 | 0 | 11 | 2 | 12 | 2 | 12 | 3.62 | 3.52 | 0.78 | 0.76 |
| Q9CRA5 | Golph3 | Golgi phosphoprotein 3 | 236.69 | 10.4 | 33752 | 3 | 895677.62 | 760683.06 | 768271.62 | 819828.75 | 1714877 | 781294.69 | 859255.81 | 937214.31 | 972951.12 | 1097744.5 | 393232.97 | 691221.38 | 812692.75 | 637722.25 | 752718.12 | 944415.5 | 607882.44 | 702092.62 | 1037057.88 | 1145830.5 | 2 | 12 | 1 | 12 | 2 | 12 | 0.66 | 0.71 | 0.89 | 0.95 |
| Q8R050 | Gspt1 | Eukaryotic peptide chain release factor GTP-binding subunit ERF3A | 248.55 | 10.53 | 68625 | 5 | 6012879.5 | 7505538.5 | 5765619.5 | 9179966 | 5185937 | 6935850.5 | 5619667 | 6001038.5 | 4420308.5 | 8029842 | 12633895 | 10799559 | 8682382 | 10345242 | 8190195 | 6150920 | 9236923 | 6795569.5 | 6910118.5 | 7122107 | 2 | 8 | 0 | 7 | 2 | 8 | 1.51 | 1.63 | 1.08 | 1.17 |
| Q9DCM2 | Gstk1 | Glutathione S-transferase kappa 1 | 79.14 | 13.72 | 25704 | 2 | 1520163.5 | 1437268.75 | 1309692.88 | 679000.31 | 1472444.38 | 733103.25 | 1192299.12 | 1367459.88 | 1321582.12 | 1324225.5 | 219745.61 | 609774.75 | 967223.62 | 495028.44 | 1125010.5 | 1617395.62 | 190076 | 990873.38 | 469045.38 | 1113458.75 | 2 | 7 | 2 | 7 | 2 | 7 | 0.53 | 0.58 | 0.68 | 0.74 |
| Q99ME9 | Gtpbp4 | Nucleolar GTP-binding protein 1 | 619.95 | 22.87 | 74113 | 12 | 739726.38 | 473180.94 | 744325.56 | 2 | 751126.81 | 1230259.75 | 1076656.5 | 320153.53 | 159952.28 | 56230.48 | 4754575.5 | 2967008.5 | 2081422 | 1541569.5 | 910232.94 | 1758115.88 | 1612311 | 58145.13 | 2 | 2400529 | 0 | 8.5 | 2 | 9 | 2 | 9 | 4.52 | 4.31 | 2.15 | 2.05 |
| O35952 | Hagh | Hydroxyacylglutathione hydrolase, mitochondrial | 95.27 | 6.15 | 34109 | 2 | 2058079.88 | 3244310.75 | 2023365.38 | 639471.12 | 4090580.5 | 419975.44 | 3185438 | 2520431 | 4042854.5 | 442917.22 | 2 | 1412091.38 | 334741.03 | 2 | 310937.88 | 2123825.25 | 2 | 336291.09 | 207979.56 | 682853.5 | 1 | 4 | 2 | 4 | 2 | 4 | 0.17 | 0.19 | 0.28 | 0.32 |
| Q8VH49 | Higd1a | HIG1 domain family member 1A, mitochondrial | 68.95 | 23.66 | 10301 | 1 | 577207.06 | 452386.66 | 482847.44 | 381128.44 | 377430.38 | 528527.88 | 503848.69 | 456497.84 | 505544.22 | 417457.69 | 2 | 373046.38 | 301787.62 | 436911.66 | 321891.28 | 473667.81 | 553154.69 | 626210.88 | 708117.38 | 387562.12 | 2 | 6 | 1 | 8 | 2 | 8 | 0.63 | 0.59 | 1.21 | 1.14 |
| Q9D0S9 | Hint2 | Histidine triad nucleotide-binding protein 2, mitochondrial | 37.97 | 7.98 | 17320 | 1 | 2 | 2 | 609161.81 | 971882.44 | 2 | 727908.94 | 2 | 650907.88 | 2 | 1049154.75 | 1030310.69 | 924418.44 | 1463714.25 | 1999049.25 | 1878374 | 2 | 691554.19 | 2 | 1287734 | 1373827.62 | 1 | 8 | 2 | 10 | 2 | 10 | 4.61 | 3.00 | 2.12 | 1.38 |
| Q9JL35 | Hmgn5 | High mobility group nucleosome-binding domain-containing protein 5 | 255.24 | 13.55 | 45344 | 5 | 5569288 | 7274098 | 6529054.5 | 6875091.5 | 7456881 | 7318162 | 7319152 | 8842843 | 8387609 | 7571119 | 5536198.5 | 6127594 | 5318245 | 5621212.5 | 4993160 | 4739956 | 3600630 | 6937649 | 8604616 | 7278166.5 | 2 | 12 | 0 | 4 | 2 | 12 | 0.78 | 0.70 | 0.92 | 0.79 |
| Q9CX86 | Hnrnpa0 | Heterogeneous nuclear ribonucleoprotein A0 | 528.66 | 26.23 | 30530 | 8 | 5272880.5 | 7328682 | 9156738 | 7120239 | 5903201 | 6627331 | 5318454 | 7068708.5 | 6364843 | 7199068.5 | 10312660 | 9423261 | 8580416 | 10091398 | 8916231 | 2886863 | 5639788 | 6742153 | 3516319.5 | 6413643 | 2 | 5 | 0 | 6 | 2 | 6 | 1.36 | 1.45 | 0.73 | 0.77 |
| P59438 | Hps5 | Hermansky-Pudlak syndrome 5 protein homolog | 103.66 | 2.22 | 126337 | 2 | 491528.81 | 266816.91 | 454415.25 | 538369.38 | 321859.94 | 176471.25 | 551637.06 | 545991.62 | 557272.81 | 492974.81 | 2 | 154307.19 | 413289.91 | 2 | 160046.09 | 641954.94 | 234109.66 | 415994.5 | 243006.14 | 308233.81 | 2 | 8 | 1 | 9 | 2 | 9 | 0.35 | 0.31 | 0.89 | 0.79 |
| O88736 | Hsd17b7 | 3-keto-steroid reductase | 415.23 | 25.75 | 37317 | 6 | 3117709 | 2385004 | 1333830.75 | 2008091.25 | 1919802.88 | 3794356.75 | 3870693.25 | 2546889.5 | 2239990 | 1838859 | 3015316.75 | 4053680 | 5286140.5 | 5069249 | 4978338.5 | 2876278.25 | 2185816 | 2787950.25 | 2273333.75 | 2676787.5 | 1 | 7 | 2 | 12 | 2 | 12 | 2.08 | 1.57 | 1.19 | 0.90 |
| P50171 | Hsd17b8 | (3R)-3-hydroxyacyl-CoA dehydrogenase | 292.19 | 22.78 | 26588 | 5 | 10116503 | 9828969 | 7802038.5 | 12108356 | 12419764 | 11199268 | 10035508 | 10091034 | 10866341 | 10083024 | 5070232 | 4981269.5 | 8139845 | 8076867 | 7564361.5 | 10077686 | 10199456 | 9792618 | 9912318 | 9850846 | 2 | 10 | 0 | 4 | 2 | 10 | 0.65 | 0.65 | 0.95 | 0.95 |
| P06761 | Hspa5 | Endoplasmic reticulum chaperone BiP | 3356.69 | 62.84 | 72347 | 47 | 422764930 | 379570690 | 324665346 | 340867426 | 316957378 | 392718754 | 402470914 | 347378370 | 362981058 | 393046594 | 455719554 | 494055074 | 421180962 | 415443618 | 506556994 | 434602914 | 392781634 | 338534594 | 341922946 | 355795202 | 2 | 8 | 0 | 9 | 2 | 9 | 1.28 | 1.21 | 1.04 | 0.98 |
| Q8BU30 | Iars | Isoleucine--tRNA ligase, cytoplasmic | 928.84 | 16.56 | 144271 | 18 | 16918918 | 15203795 | 16640322 | 16250181 | 18413708 | 14458544 | 13998497 | 17748718 | 18171026 | 18509632 | 21845364 | 23638150 | 25183278 | 18052888 | 26025708 | 15359781 | 11256623 | 17130926 | 19382788 | 18983754 | 1 | 10 | 2 | 11 | 2 | 11 | 1.38 | 1.38 | 0.98 | 0.99 |
| P13597 | Icam1 | Intercellular adhesion molecule 1 | 433.7 | 15.83 | 58844 | 8 | 2101873 | 2443097 | 2638901.25 | 770396.62 | 210194.59 | 3593100 | 3619036.25 | 2738806.5 | 3275944 | 2702585.5 | 4390429.5 | 3499687.5 | 3796883.5 | 478943 | 3754785 | 2525925.5 | 765915.62 | 1361945.12 | 5303840 | 858726.75 | 0 | 11 | 2 | 5 | 2 | 11 | 0.68 | 1.27 | 1.32 | 0.68 |
| Q8BV66 | Ifi44 | Interferon-induced protein 44 | 209.75 | 10.43 | 47852 | 4 | 8697622 | 10107518 | 6048320 | 8683038 | 11484607 | 8148303 | 7329725 | 10206774 | 11158256 | 7700019 | 4681844 | 4418052 | 7154064 | 4582376.5 | 6315248 | 6996473.5 | 6921698 | 4333685 | 5613357.5 | 8851222 | 2 | 5 | 0 | 3 | 2 | 5 | 0.60 | 0.61 | 0.73 | 0.73 |

| | | | | | | | | | | | | | | | | | | | | | | | | | | | | | | | | | | | | |
|---|---|---|---|---|---|---|---|---|---|---|---|---|---|---|---|---|---|---|---|---|---|---|---|---|---|---|---|---|---|---|---|---|---|---|---|---|
| P11835 | Itgb2 | Integrin beta-2 | 2152.3 | 50.58 | 85026 | 32 | 57889934 | 60608126 | 68087306 | 73967066 | 63443658 | 60340942 | 64724178 | 55744202 | 53610658 | 58985786 | 42107182 | 45060026 | 53500018 | 55665018 | 54271026 | 54387646 | 53489918 | 52105238 | 49769862 | 47157346 | 0 | 0 | 2 | 1 | 2 | 1 | 0.77 | 0.85 | 0.79 | 0.88 |
| O88447 | Klc1 | Kinesin light chain 1 | 139.11 | 5.18 | 61450 | 3 | 1234729 | 1329136.75 | 1832212.75 | 1471942.75 | 2459430.75 | 1552733.25 | 1545023.12 | 1926138.25 | 2372504 | 1808694.75 | 869947.56 | 514727.22 | 1023695.62 | 1434415.62 | 1109551.75 | 2773553.25 | 2654447 | 1926038.25 | 2834221.75 | 3933038 | 2 | 1 | 0 | 2 | 2 | 2 | 0.59 | 0.54 | 1.70 | 1.53 |
| O35345 | Kpna6 | Importin subunit alpha-7 | 201.59 | 8.58 | 59964 | 4 | 877408.94 | 436694.09 | 336505.53 | 422007.56 | 282008.06 | 992727.44 | 818428.62 | 151965.16 | 2 | 484646.28 | 1214997.38 | 1113050.5 | 951893.94 | 1053630.88 | 841277.25 | 535373.62 | 648376.38 | 151039.14 | 303494.16 | 2 | 1 | 9 | 2 | 9.5 | 2 | 9.5 | 2.20 | 2.11 | 0.70 | 0.67 |
| Q61233 | Lcp1 | Plastin-2 | 4410.49 | 82.3 | 70149 | 63 | 793658882 | 588158786 | 818221442 | 540277890 | 904508930 | 733736130 | 806224322 | 817528962 | 673548866 | 577219906 | 520396514 | 613904258 | 376070914 | 318837890 | 503683554 | 742758594 | 797554818 | 534501314 | 526085122 | 742386754 | 2 | 7 | 1 | 9 | 2 | 9 | 0.64 | 0.65 | 0.92 | 0.93 |
| P62046 | Lrch1 | Leucine-rich repeat and calponin homology domain-containing protein 1 | 48.74 | 1.83 | 79076 | 1 | 449406.31 | 469554 | 339486.06 | 339637.66 | 440189.88 | 279198.12 | 483311.78 | 410006.12 | 454344.69 | 355623.78 | 2 | 314966.44 | 281951.62 | 2 | 221185.06 | 388665.56 | 2 | 378953.34 | 188063.11 | 425474.28 | 0 | 6 | 2 | 7 | 2 | 7 | 0.40 | 0.41 | 0.68 | 0.70 |
| P62322 | Lsm5 | U6 snRNA-associated Sm-like protein LSm5 | 82.79 | 20.88 | 9937 | 1 | 702172.12 | 416537.5 | 604306.19 | 657274.06 | 594387.69 | 621122.75 | 813132.69 | 545057.38 | 858982.5 | 748382.62 | 251735.98 | 238104.47 | 420210.38 | 533256.38 | 2 | 667376.62 | 983892.69 | 2 | 606108 | 403443.97 | 2 | 12 | 0 | 8 | 2 | 12 | 0.49 | 0.40 | 0.89 | 0.74 |
| Q9QXZ0 | Macf1 | Microtubule-actin cross-linking factor 1 | 371.23 | 1.28 | 831878 | 7 | 1229951.12 | 1429466.25 | 1710885.88 | 1218920 | 2114938.75 | 1192169.25 | 1601373.5 | 1289931.75 | 1732872.25 | 1213097 | 332722.19 | 1270185.25 | 763601.06 | 670582.5 | 544943.06 | 1774378.62 | 913640.12 | 329098.94 | 729521.31 | 1288483.38 | 2 | 6 | 2 | 7 | 2 | 7 | 0.46 | 0.51 | 0.65 | 0.72 |
| P21139 | Man2c1 | Alpha-mannosidase 2C1 | 172.26 | 2.88 | 115971 | 2 | 1209001.38 | 1142318.5 | 1326973.25 | 1332167.25 | 1157969.5 | 931813.38 | 726692.69 | 1079227.88 | 1177299 | 1081630.62 | 554613.88 | 695828.88 | 921475.62 | 672888.56 | 761186.38 | 1278839.75 | 465483.03 | 1092417 | 1488780.5 | 1074520.12 | 0 | 8 | 2 | 9 | 2 | 9 | 0.58 | 0.72 | 0.88 | 1.08 |
| Q8C052 | Map1s | Microtubule-associated protein 1S | 247.59 | 5.65 | 102939 | 5 | 3933269 | 4453431 | 4633019 | 3066923 | 3972475.5 | 3122835.5 | 4316991 | 3927701 | 3487806.75 | 3969207.25 | 2794642.75 | 3797296.5 | 2532643.25 | 2105480 | 2158375.25 | 4179727.5 | 2481982.5 | 4303846 | 3894264 | 4249454.5 | 1 | 10 | 2 | 11 | 2 | 11 | 0.67 | 0.71 | 0.95 | 1.02 |
| O09110 | Map2k3 | Dual specificity mitogen-activated protein kinase kinase 3 | 103.15 | 9.22 | 39296 | 2 | 411683.09 | 535137.5 | 609577.12 | 303522 | 753193.12 | 398573.03 | 805872.62 | 650013.88 | 1021020.75 | 226099.3 | 292495.06 | 312937.78 | 173232.17 | 68698.38 | 195380.8 | 426883.84 | 121114.55 | 230076.11 | 331314.81 | 177788.94 | 1 | 3 | 2 | 4 | 2 | 4 | 0.40 | 0.34 | 0.49 | 0.42 |
| P23249 | Mov10 | Putative helicase MOV-10 | 497.67 | 13.05 | 113583 | 10 | 7510070.5 | 6280248.5 | 6082475 | 6991612.5 | 7052520.5 | 6807734 | 5728600.5 | 6984189.5 | 6828921 | 7558099 | 5072836 | 4444203.5 | 6795630.5 | 5830138 | 3863139 | 6466199 | 5703308.5 | 7710373 | 5850296.5 | 7136244 | 2 | 11 | 2 | 11 | 2 | 11 | 0.77 | 0.77 | 0.97 | 0.97 |
| Q9CQF0 | Mrpl11 | 39S ribosomal protein L11, mitochondrial | 89.78 | 15.1 | 20680 | 2 | 1159855.38 | 1746464.5 | 2084219.38 | 1818164.38 | 1893918.5 | 2384154.5 | 1945621.75 | 1578426 | 1151826.88 | 1630715.12 | 535180 | 1075968.88 | 1101514.25 | 1577147.38 | 1482784.75 | 1752114 | 1868868.75 | 2103840.25 | 2007099.25 | 953417.94 | 2 | 11 | 2 | 11 | 2 | 11 | 0.66 | 0.66 | 1.00 | 1.00 |
| Q9D6Y7 | Msra | Mitochondrial peptide methionine sulfoxide reductase | 70.49 | 6.87 | 25988 | 1 | 533003.75 | 703331.75 | 1049644.62 | 1168381.62 | 774463.56 | 696883.44 | 1009367.25 | 1343919 | 1204822.12 | 1179339.62 | 2 | 723431.62 | 2 | 2 | 498094.72 | 686820.5 | 2 | 1426450 | 1126839.25 | 983007.5 | 2 | 12 | 1 | 8 | 2 | 12 | 0.29 | 0.22 | 1.00 | 0.78 |
| Q9CWZ7 | Napg | Gamma-soluble NSF attachment protein | 443.68 | 19.23 | 34732 | 7 | 11088988 | 13770494 | 14294186 | 13985626 | 17354466 | 9085534 | 13721403 | 14470126 | 16321228 | 10887309 | 5186496 | 9488301 | 9067052 | 7356326 | 9901191 | 11600356 | 6723215 | 12266312 | 11349709 | 9564898 | 0 | 3 | 2 | 7 | 2 | 7 | 0.58 | 0.64 | 0.73 | 0.80 |
| Q99MD9 | Nasp | Nuclear autoantigenic sperm protein | 579.86 | 18.37 | 83954 | 11 | 13208880 | 12452438 | 17121542 | 13823068 | 13185591 | 13209753 | 11816861 | 9962407 | 12527017 | 12088650 | 10472987 | 10609888 | 9106457 | 8618131 | 8650723 | 14604533 | 11288573 | 12418712 | 9078682 | 10498238 | 0 | 4 | 2 | 10 | 2 | 10 | 0.68 | 0.80 | 0.83 | 0.97 |
| Q3UYV9 | Ncbp1 | Nuclear cap-binding protein subunit 1 | 191.73 | 5.82 | 91927 | 4 | 762120.88 | 504226.12 | 581980.62 | 641390.88 | 766964.25 | 439349.56 | 458360.06 | 545609.88 | 699079.06 | 564937 | 1022877.56 | 978124 | 1607543.38 | 743616.88 | 1067806.12 | 631671.75 | 729634.38 | 534665.69 | 586075.88 | 461346.72 | 2 | 8 | 0 | 8 | 2 | 8 | 1.66 | 2.00 | 0.90 | 1.09 |
| Q8CCH2 | Nhlrc3 | NHL repeat-containing protein 3 | 230.25 | 10.09 | 38195 | 3 | 3263604.5 | 3181009.25 | 675987 | 3109822 | 4117308.5 | 2105077.75 | 3536181 | 3183657 | 2573584 | 1062182.12 | 322726.72 | 1843430.5 | 1804321.25 | 2 | 511184.06 | 4979161.5 | 2388706 | 4857589.5 | 3366518.25 | 4392374.5 | 2 | 5 | 2 | 4 | 2 | 5 | 0.31 | 0.36 | 1.39 | 1.60 |
| Q91VE6 | Nifk | MKI67 FHA domain-interacting nucleolar phosphoprotein | 463.93 | 29.97 | 36265 | 8 | 5459895.5 | 4455114 | 5130856.5 | 3020759.5 | 3620166 | 5815124 | 5426969.5 | 2517076 | 3762796.5 | 3196165 | 6516215.5 | 6685845.5 | 5618609.5 | 5606259 | 7317639.5 | 5651648 | 4636420.5 | 5463489.5 | 3992782.5 | 4215554 | 0 | 8 | 2 | 8 | 2 | 8 | 1.46 | 1.53 | 1.10 | 1.16 |
| Q8VEJ4 | Nle1 | Notchless protein homolog 1 | 147.95 | 7.63 | 53134 | 3 | 1363441.75 | 1152372.62 | 1019031.12 | 1030047 | 1322283.88 | 1427123.25 | 969060 | 1111331.75 | 950005 | 1354343.25 | 1812520.5 | 1369894.12 | 2004249.25 | 1369809.25 | 1627184.88 | 1197985.5 | 666700.62 | 1234409.25 | 1000239.62 | 1203859.5 | 0 | 9 | 2 | 11 | 2 | 11 | 1.39 | 1.41 | 0.90 | 0.91 |
| Q9R1J0 | Nsdhl | Sterol-4-alpha-carboxylate 3-dehydrogenase, decarboxylating | 617.62 | 39.23 | 40686 | 10 | 9952038 | 12261537 | 9401702 | 7892803.5 | 10114061 | 10859824 | 12022310 | 8755560 | 8154216.5 | 9058501 | 11487179 | 11372173 | 13230300 | 15575352 | 16589838 | 13966344 | 12220518 | 12129452 | 11411731 | 12357093 | 2 | 3 | 2 | 1 | 2 | 3 | 1.38 | 1.40 | 1.25 | 1.27 |
| Q80XU3 | Nucks1 | Nuclear ubiquitous casein and cyclin-dependent kinase substrate 1 | 145.31 | 11.11 | 26313 | 3 | 11606529 | 12516090 | 13132566 | 20097178 | 10580274 | 13741405 | 10044944 | 13475112 | 11671298 | 17122720 | 8395231 | 6302413.5 | 10457486 | 10489042 | 9489332 | 8067953 | 12490258 | 12249391 | 15531588 | 17033678 | 0 | 12 | 2 | 12 | 2 | 12 | 0.66 | 0.68 | 0.96 | 0.99 |
| Q8CGY8 | Ogt | UDP-N-acetylglucosamine--peptide N-acetylglucosaminyltransferase 110 kDa subunit | 564.02 | 10.52 | 116952 | 9 | 4903246.5 | 6481749 | 10862972 | 8923585 | 8787079 | 6723616 | 7380382 | 6055889 | 7341055.5 | 8781219 | 4038127.5 | 5846657.5 | 6227049.5 | 3723606.5 | 4272579 | 5409177.5 | 6388400 | 7946733 | 8438842 | 6555290.5 | 2 | 8 | 1 | 10 | 2 | 10 | 0.60 | 0.66 | 0.87 | 0.96 |
| Q8BX94 | Osbpl2 | Oxysterol-binding protein-related protein 2 | 96.84 | 6.82 | 55384 | 2 | 1239089.12 | 1160142.25 | 597554.5 | 590068.56 | 832064.62 | 783313.31 | 1144709.62 | 641019.19 | 1118895.12 | 257861.62 | 248333.5 | 370578.09 | 501892.12 | 2 | 45503.41 | 1040189.31 | 442366.25 | 642248.5 | 2 | 810059.69 | 0 | 7 | 2 | 9 | 2 | 9 | 0.26 | 0.30 | 0.66 | 0.74 |
| O35264 | Pafah1b2 | Platelet-activating factor acetylhydrolase IB subunit beta | 91.81 | 12.23 | 25581 | 2 | 2 | 2 | 234532.31 | 2 | 730701.81 | 2 | 95532.77 | 199484.05 | 379391.56 | 2 | 1654062.38 | 1318293.5 | 619150.81 | 406501.09 | 973710.5 | 168516.06 | 2 | 163807.89 | 824858.06 | 88024.42 | 2 | 9.5 | 0 | 11 | 2 | 11 | 5.15 | 7.37 | 1.29 | 1.85 |
| Q2EMV9 | Parp14 | Poly [ADP-ribose] polymerase 14 | 305.67 | 4.95 | 203802 | 7 | 2574537.75 | 2469461.5 | 2792454.25 | 3196602.25 | 3465875.25 | 3046859.75 | 3101513.25 | 3046613.5 | 2921148.5 | 2739483 | 2730727 | 1868576.25 | 2056346.25 | 1251089.75 | 1731579 | 2384689.75 | 2042739.75 | 2743162.5 | 4118711.5 | 2952412 | 2 | 10 | 0 | 8 | 2 | 10 | 0.66 | 0.65 | 0.98 | 0.96 |
| Q9WU28 | Pfdn5 | Prefoldin subunit 5 | 200.71 | 25.97 | 17356 | 3 | 5020825 | 4666618.5 | 4694435.5 | 5519291.5 | 5624607 | 4763807 | 4119919 | 4529040.5 | 4999391.5 | 5160469 | 3498742.5 | 2819087.5 | 3375099 | 4541273.5 | 2944034.75 | 4818647 | 5521528 | 4763253 | 3204471.25 | 3952763 | 0 | 8 | 2 | 10 | 2 | 10 | 0.67 | 0.73 | 0.87 | 0.94 |
| P24135 | Plcg2 | 1-phosphatidylinositol 4,5-bisphosphate phosphodiesterase gamma-2 | 572.5 | 10.2 | 147735 | 12 | 8884860 | 8867481 | 11872694 | 12902016 | 10641579 | 9961583 | 9455840 | 10208348 | 11092273 | 12054423 | 8135882 | 8688266 | 8352625 | 9221494 | 7277491 | 8102764.5 | 7765962.5 | 10854340 | 11368832 | 9631962 | 2 | 8 | 0 | 8 | 2 | 8 | 0.78 | 0.79 | 0.90 | 0.90 |
| Q99K51 | Pls3 | Plastin-3 | 1679.24 | 49.68 | 70742 | 26 | 46552794 | 49407098 | 31107656 | 32716232 | 51821034 | 41262362 | 46925406 | 38586450 | 46238054 | 37804018 | 26564152 | 34741274 | 23556636 | 22482366 | 24452356 | 44935010 | 39437606 | 36093254 | 39998378 | 30868706 | 2 | 8 | 0 | 7 | 2 | 8 | 0.62 | 0.63 | 0.90 | 0.91 |
| Q60953 | Pml | Protein PML | 369.02 | 9.38 | 98242 | 7 | 2434977.5 | 2824862.25 | 1907620.75 | 2021042 | 2257741 | 1782985.25 | 2631081 | 1491793.25 | 2163519.5 | 2475898.5 | 1911347.88 | 359467.41 | 597221.38 | 199063.78 | 1059082.25 | 2196858.25 | 1459712.75 | 1234477 | 865845.25 | 1666983.38 | 1 | 2 | 2 | 4 | 2 | 4 | 0.36 | 0.39 | 0.65 | 0.70 |
| Q9JLV6 | Pnkp | Bifunctional polynucleotide phosphatase/kinase | 219.71 | 9 | 57223 | 4 | 3289695.5 | 3588072.75 | 2826474.5 | 2814974.5 | 2932938.5 | 2851758.5 | 2716602.5 | 4310539 | 2365091 | 2494353.75 | 2300085 | 1560814.75 | 1986790.75 | 2193547.5 | 2693038 | 2373733 | 3041876.25 | 2744351.75 | 3106984.5 | 3450823 | 0 | 10 | 2 | 9 | 2 | 10 | 0.69 | 0.73 | 0.95 | 1.00 |
| Q8VHI3 | Pofut2 | GDP-fucose protein O-fucosyltransferase 2 | 176.23 | 9.79 | 49429 | 3 | 224675.31 | 752456.38 | 495950.53 | 266797.56 | 896859.69 | 273511.84 | 240135.22 | 222160.39 | 393940.94 | 813835.5 | 3736976.5 | 593263.06 | 961626.75 | 1859086.75 | 3320988.5 | 500259.84 | 813555.38 | 951140 | 786294.38 | 4617949.5 | 2 | 4 | 1 | 3 | 2 | 4 | 3.97 | 5.39 | 2.91 | 3.95 |
| P54645 | Prkaa1 | 5'-AMP-activated protein kinase catalytic subunit alpha-1 | 42.03 | 1.25 | 63973 | 1 | 589041 | 146182.06 | 547439.88 | 375942.22 | 2 | 2 | 2 | 2 | 2 | 548415.81 | 1069189.88 | 917240.62 | 750215.12 | 539105.81 | 741260.44 | 2 | 2 | 2 | 2 | 426210.28 | 2 | 5 | 1 | 12 | 2 | 12 | 2.42 | 7.32 | 0.26 | 0.78 |
| P12367 | Prkar2a | cAMP-dependent protein kinase type II-alpha regulatory subunit | 100.01 | 6.73 | 45389 | 2 | 390108.91 | 546053.44 | 447749.03 | 222442.55 | 332044.34 | 936674.75 | 310210.28 | 388834.69 | 362905.09 | 307938.47 | 1158343 | 999147.12 | 636438.38 | 640155.06 | 1093134.12 | 313327.09 | 878818.44 | 385468.62 | 280626.38 | 464386.09 | 0 | 12 | 2 | 12 | 2 | 12 | 2.34 | 1.96 | 1.20 | 1.01 |
| Q8R574 | Prpsap2 | Phosphoribosyl pyrophosphate synthase-associated protein 2 | 405.44 | 28.73 | 40881 | 7 | 3234208.75 | 2798035 | 3101815.5 | 2074682 | 3263993.5 | 1767067.38 | 2881516.5 | 3608874.75 | 3455931 | 2657259.75 | 1650128.5 | 2230309 | 1384820.38 | 1244193.88 | 1826214.5 | 2809825.5 | 1617899.38 | 2076396.5 | 1772131.25 | 2163369.5 | 1 | 4 | 2 | 5 | 2 | 5 | 0.58 | 0.58 | 0.72 | 0.73 |
| P17220 | Psma2 | Proteasome subunit alpha type-2 | 564.08 | 41.45 | 25927 | 9 | 10585845 | 9413731 | 9470277 | 8579442 | 5310389 | 9291446 | 9595182 | 8315244.5 | 7208283 | 10155468 | 16390215 | 13612725 | 9652332 | 9694049 | 12833323 | 8917876 | 10126152 | 9861511 | 10180118 | 11549227 | 2 | 6 | 2 | 5 | 2 | 6 | 1.43 | 1.40 | 1.17 | 1.14 |
| O35955 | Psmb10 | Proteasome subunit beta type-10 | 341.26 | 26.37 | 29063 | 5 | 12975611 | 12312603 | 10610130 | 15013296 | 12905351 | 12426722 | 9759401 | 13977025 | 12908440 | 13805317 | 10627549 | 8004410.5 | 8696787 | 7796473 | 9808742 | 9218034 | 11566576 | 11675385 | 15095800 | 14541102 | 1 | 11 | 2 | 12 | 2 | 12 | 0.70 | 0.71 | 0.97 | 0.99 |
| P99026 | Psmb4 | Proteasome subunit beta type-4 | 487.82 | 39.39 | 29116 | 7 | 20464140 | 25938466 | 25432194 | 26045580 | 24070534 | 24614136 | 20878742 | 23342914 | 25439566 | 26495772 | 22365788 | 21122790 | 19414126 | 18317766 | 18548646 | 23447842 | 22723018 | 25891202 | 26693088 | 24114718 | 2 | 12 | 2 | 11 | 2 | 12 | 0.82 | 0.83 | 1.01 | 1.02 |
| O88685 | Psmc3 | 26S proteasome regulatory subunit 6A | 1226.11 | 50.9 | 49549 | 21 | 24161872 | 23793706 | 24623166 | 29012164 | 26585898 | 25354674 | 22940182 | 24869124 | 31100152 | 31999758 | 34103602 | 33767090 | 36029594 | 29907334 | 37536086 | 24886516 | 24608644 | 33260398 | 31505084 | 31483154 | 0 | 5 | 2 | 9 | 2 | 9 | 1.34 | 1.26 | 1.14 | 1.07 |
| P35831 | Ptpn12 | Tyrosine-protein phosphatase non-receptor type 12 | 88.64 | 2.58 | 86526 | 2 | 1597359.88 | 1734961.5 | 1233972.62 | 1314856.88 | 68071.05 | 1603187.75 | 1496194.75 | 1430500.5 | 1162909.12 | 120277.95 | 1745934.25 | 1734952.62 | 1915776.12 | 1867414.88 | 1645269.88 | 1524814.75 | 59299.74 | 1272033.75 | 1940852.5 | 1358502.5 | 2 | 12 | 0 | 12 | 2 | 12 | 1.50 | 1.53 | 1.03 | 1.06 |
| Q3UEB3 | Puf60 | Poly(U)-binding-splicing factor PUF60 | 775.99 | 36.7 | 60249 | 13 | 13664203 | 14686804 | 13551204 | 11025070 | 11878971 | 10581844 | 14159367 | 9059887 | 12229378 | 9267876 | 8976399 | 9452505 | 8900783 | 6323087 | 7762405.5 | 14558116 | 14652739 | 12726757 | 11323936 | 10693201 | 0 | 11 | 2 | 5 | 2 | 11 | 0.64 | 0.75 | 0.99 | 1.16 |
| P61027 | Rab10 | Ras-related protein Rab-10 | 840.32 | 49.5 | 22541 | 15 | 27415796 | 26846240 | 24438256 | 20186822 | 27459814 | 20709464 | 29458474 | 26351492 | 26316594 | 24731964 | 15835780 | 22042450 | 21809578 | 10851045 | 12866319 | 29932622 | 19510298 | 21180980 | 12055697 | 20943280 | 2 | 7 | 2 | 7 | 2 | 7 | 0.66 | 0.65 | 0.82 | 0.81 |
| P35293 | Rab18 | Ras-related protein Rab-18 | 372.69 | 37.38 | 23035 | 6 | 2290905 | 1535950.75 | 4180238 | 3269490.75 | 1572070.5 | 3549479.25 | 2586163.25 | 3838429.75 | 2067514.5 | 6012491 | 6968783 | 5984985 | 8249985 | 5189242 | 8127000.5 | 3323547 | 5128111 | 5226470 | 6120125.5 | 5291493.5 | 0 | 1 | 2 | 6 | 2 | 6 | 2.69 | 1.91 | 1.95 | 1.39 |
| P63001 | Rac1 | Ras-related C3 botulinum toxin substrate 1 | 590.61 | 52.6 | 21450 | 9 | 13113981 | 10963706 | 10547920 | 13947131 | 9313760 | 12354465 | 11564474 | 10449930 | 10728226 | 10724269 | 10238501 | 9267045 | 10156602 | 9058434 | 8874276 | 10864550 | 11911400 | 11017179 | 11705663 | 9874280 | 2 | 12 | 0 | 11 | 2 | 12 | 0.82 | 0.85 | 0.96 | 0.99 |
| Q27W01 | Rbm8a | RNA-binding protein 8A | 169.19 | 22.41 | 19889 | 3 | 4239646 | 5075659 | 4558940 | 4106249 | 4949064.5 | 3836961.75 | 5608400.5 | 4789178 | 4497641.5 | 3809997.5 | 1996881.62 | 3919466 | 3064949.75 | 2322331.25 | 2089426.88 | 4993281 | 4223137 | 3486694.5 | 3662641.25 | 3646119.5 | 0 | 5 | 2 | 6 | 2 | 6 | 0.58 | 0.59 | 0.87 | 0.89 |
| Q8BP92 | Rcn2 | Reticulocalbin-2 | 512.31 | 33.12 | 37271 | 8 | 2 | 542250.5 | 359442.06 | 218010.08 | 2 | 2 | 2 | 2 | 221502.11 | 2 | 2637103.5 | 1087060 | 1112063.25 | 1160075.62 | 249372.33 | 2 | 777871.5 | 2 | 2 | 2 | 2 | 9 | 0 | 12 | 2 | 12 | 5.58 | 28.20 | 0.69 | 3.51 |
| Q8VE91 | Retreg1 | Reticulophagy regulator 1 | 171.66 | 7.29 | 52967 | 3 | 1186824.12 | 1068873.12 | 1807634.62 | 1565975 | 1589422.75 | 1197228.5 | 1511456.5 | 2073828 | 1955992.5 | 2314447.5 | 3484747 | 3356419.25 | 2414921.5 | 2023530.62 | 3823275.5 | 1633857.38 | 1585548.75 | 2385658 | 2147367.75 | 2514557.25 | 0 | 3 | 2 | 7 | 2 | 7 | 2.09 | 1.67 | 1.42 | 1.13 |
| Q5U1X1 | Rexo2 | Oligoribonuclease, mitochondrial | 256.5 | 17.72 | 26752 | 4 | 7211751.5 | 7610854 | 5521529.5 | 7278375.5 | 8969052 | 8595078 | 8427576 | 7971791.5 | 8708906 | 9022101 | 4494310 | 4600114 | 5490307 | 6030524.5 | 5808641.5 | 9227414 | 6663167 | 6729352 | 9206980 | 9070110 | 2 | 8 | 0 | 10 | 2 | 10 | 0.72 | 0.62 | 1.12 | 0.96 |
| P52760 | Rida | 2-iminobutanoate/2-iminopropanoate deaminase | 88.74 | 19.26 | 14255 | 2 | 2567178.5 | 597736.25 | 1965444.75 | 697412.31 | 2246829.25 | 1911788.88 | 513523.62 | 1581344.75 | 1516173.75 | 3484611.5 | 1073937.75 | 2 | 411183.38 | 455432.5 | 2 | 3233029 | 4970979 | 766693.69 | 672785.12 | 729623.56 | 2 | 10 | 1 | 12 | 2 | 12 | 0.24 | 0.22 | 1.28 | 1.15 |
| Q91VI7 | Rnh1 | Ribonuclease inhibitor | 2519.29 | 82.68 | 49816 | 30 | 164062866 | 170076418 | 145901682 | 130123730 | 166484354 | 130097794 | 195301586 | 198912434 | 192759714 | 154316194 | 97392970 | 150362018 | 110252682 | 59717918 | 113266282 | 181184802 | 80336826 | 121376458 | 130056002 | 141794962 | 2 | 6 | 1 | 3 | 2 | 6 | 0.68 | 0.61 | 0.84 | 0.75 |
| P35979 | Rpl12 | 60S ribosomal protein L12 | 545.95 | 51.52 | 17805 | 8 | 47545050 | 56155174 | 55583698 | 54144578 | 49251906 | 65734794 | 58799462 | 64942922 | 62652994 | 71353530 | 67314522 | 74888066 | 72307498 | 74565538 | 67996042 | 52242090 | 51962826 | 76264610 | 74852746 | 62948998 | 0 | 6 | 2 | 12 | 2 | 12 | 1.36 | 1.10 | 1.21 | 0.98 |
| P61255 | Rpl26 | 60S ribosomal protein L26 | 540.15 | 46.9 | 17258 | 11 | 30649744 | 36092766 | 28029440 | 22090888 | 16435398 | 38577186 | 33020174 | 34170542 | 25847226 | 30254586 | 47288362 | 44684826 | 38590826 | 35387318 | 37688770 | 30045584 | 32627954 | 34334762 | 29825760 | 34689230 | 1 | 7 | 2 | 12 | 2 | 12 | 1.53 | 1.26 | 1.21 | 1.00 |
| P63323 | Rps12 | 40S ribosomal protein S12 | 405.46 | 49.24 | 14525 | 6 | 14446056 | 18686352 | 16991748 | 12376538 | 13472290 | 14341465 | 16893730 | 12593742 | 14547456 | 10797275 | 22751702 | 27575998 | 21609772 | 15906435 | 20968696 | 21577948 | 11802368 | 12812913 | 13992302 | 12528391 | 2 | 9 | 1 | 11 | 2 | 11 | 1.43 | 1.57 | 0.96 | 1.05 |
| P62855 | Rps26 | 40S ribosomal protein S26 | 210.31 | 20.87 | 13015 | 3 | 61373914 | 66496694 | 70983330 | 68631466 | 66796042 | 68861090 | 69673994 | 67996146 | 62220346 | 73096314 | 48215274 | 54967702 | 59984586 | 64170738 | 63038242 | 52319314 | 66505890 | 68442042 | 78930522 | 67013566 | 2 | 12 | 2 | 9 | 2 | 12 | 0.87 | 0.85 | 1.00 | 0.97 |
| Q9Z2G6 | Sel1l | Protein sel-1 homolog 1 | 479.7 | 14.81 | 88340 | 7 | 1449648.25 | 2004069.38 | 1703608.5 | 1904361.62 | 1583826.75 | 1011187.62 | 1792948.75 | 2054909.75 | 2034347.12 | 2669072.5 | 2215766.5 | 3716867.75 | 3618209.75 | 2573970.5 | 3259437.75 | 2106538.5 | 1209293 | 1531699.88 | 1695045.5 | 922644.81 | 0 | 8 | 2 | 7 | 2 | 8 | 1.78 | 1.61 | 0.86 | 0.78 |
| P55194 | Sh3bp1 | SH3 domain-binding protein 1 | 192.84 | 8.53 | 74172 | 5 | 3190609.5 | 3809920 | 4490802.5 | 4387910.5 | 3666784 | 3547409.5 | 4282484.5 | 3895506.25 | 3468490.25 | 1560622.88 | 844693.75 | 1724855.5 | 2606129 | 1226067.38 | 1137558.5 | 3966598.25 | 1698651.75 | 2320277 | 2257081.25 | 2907223.5 | 0 | 3 | 2 | 8 | 2 | 8 | 0.39 | 0.45 | 0.67 | 0.78 |
| Q78IQ7 | Slc39a4 | Zinc transporter ZIP4 | 155.25 | 4.09 | 71063 | 2 | 2 | 2 | 2 | 2 | 212244.03 | 490096.81 | 2 | 2 | 2 | 637962.69 | 1122110 | 834824.88 | 803855.31 | 439034.25 | 899065.19 | 520224.38 | 2 | 621530.69 | 675542.12 | 557467.5 | 0 | 3 | 2 | 6.5 | 2 | 6.5 | 19.31 | 3.63 | 11.19 | 2.11 |
| P62320 | Snrpd3 | Small nuclear ribonucleoprotein Sm D3 | 156.21 | 30.95 | 13916 | 3 | 17761068 | 17989994 | 19162858 | 24995466 | 15405266 | 22061778 | 19434404 | 19632798 | 18839780 | 23083578 | 36306738 | 22050618 | 30800628 | 34589718 | 27684276 | 16976084 | 23509180 | 23309940 | 26262592 | 24172646 | 1 | 7 | 2 | 5 | 2 | 7 | 1.59 | 1.47 | 1.20 | 1.11 |
| O35892 | Sp100 | Nuclear autoantigen Sp-100 | 295.51 | 12.24 | 54727 | 6 | 1447781.12 | 2332736.25 | 927597 | 2583610.5 | 1333819.25 | 2409581.25 | 1732866.5 | 2025773.75 | 1575940.25 | 2522889.5 | 780577.12 | 871127.38 | 1048141.56 | 1163209.25 | 765167.62 | 1129312.5 | 1682078.38 | 1722726 | 2039552.25 | 2305335.5 | 2 | 12 | 0 | 8 | 2 | 12 | 0.54 | 0.45 | 1.03 | 0.86 |
| P14576 | Srp54 | Signal recognition particle 54 kDa protein | 106.3 | 6.15 | 55721 | 2 | 1726393 | 1707138.38 | 1962641.75 | 1618209.75 | 1892016.38 | 1441793.5 | 1579971.25 | 2 | 1782899.62 | 2 | 1733300.25 | 2119204.5 | 2277148 | 2105068.25 | 2540877.75 | 1806713.75 | 2 | 2115559.5 | 1904813.75 | 2315737.25 | 2 | 8 | 1 | 4 | 2 | 8 | 1.21 | 2.24 | 0.91 | 1.69 |
| Q9R0U0 | Srsf10 | Serine/arginine-rich splicing factor 10 | 260.63 | 19.08 | 31301 | 4 | 4095790.75 | 4472141.5 | 3417730 | 2484113.25 | 4796650.5 | 3177282.5 | 4385638 | 3396305 | 4919169 | 2251103.75 | 2091603.75 | 3286911.75 | 1475725.25 | 2 | 2095273 | 3532369 | 3229088 | 3175951 | 2614245.5 | 2795175.75 | 1 | 6 | 2 | 8 | 2 | 8 | 0.46 | 0.49 | 0.80 | 0.85 |
| B0LT89 | Stk24 | Serine/threonine-protein kinase 24 | 219.81 | 15.31 | 47990 | 4 | 868059.12 | 1145763.12 | 1066439 | 644190.94 | 648120.44 | 1333364.25 | 922208.44 | 1023541.88 | 884072.5 | 1033282.5 | 432859.62 | 437901.97 | 798315.12 | 222136.58 | 457158.12 | 1188569 | 1183854.88 | 478888 | 2 | 846363.62 | 2 | 12 | 0 | 8 | 2 | 12 | 0.54 | 0.45 | 0.85 | 0.71 |
| P54116 | Stom | Erythrocyte band 7 integral membrane protein | 814.88 | 51.06 | 31375 | 13 | 92341338 | 93551802 | 55041202 | 65966150 | 65776542 | 108484890 | 99222178 | 64689742 | 69707938 | 64106626 | 121065202 | 122221826 | 102101578 | 99342218 | 113663586 | 81509154 | 101558538 | 54415990 | 62695590 | 64967366 | 0 | 10 | 2 | 9 | 2 | 10 | 1.50 | 1.37 | 0.98 | 0.90 |
| P11031 | Sub1 | Activated RNA polymerase II transcriptional coactivator p15 | 196.06 | 33.07 | 14427 | 4 | 5567745.5 | 7847156.5 | 5433774.5 | 6893635 | 9637934 | 6392736.5 | 4532797.5 | 8208163 | 8172471.5 | 6586468.5 | 4462873 | 6141244.5 | 2827539 | 2772189.5 | 3344648 | 10151775 | 10043001 | 3875539.5 | 5310045 | 5722626 | 2 | 12 | 1 | 12 | 2 | 12 | 0.55 | 0.58 | 0.99 | 1.04 |
| Q64232 | Tecr | Very-long-chain enoyl-CoA reductase | 373.7 | 20.45 | 36123 | 7 | 3047384.75 | 5641587.5 | 8403883 | 6865663 | 6606188 | 3098219 | 5448722 | 7183319 | 10625561 | 5413980.5 | 7728804 | 10586639 | 15910204 | 11813662 | 16889300 | 2921052.5 | 2 | 4221806.5 | 7987675 | 7507977.5 | 1 | 9 | 2 | 9 | 2 | 9 | 2.06 | 1.98 | 0.74 | 0.71 |
| Q9DBS1 | Tmem43 | Transmembrane protein 43 | 475.03 | 25.75 | 44783 | 8 | 3122653.5 | 3581449.5 | 4353855.5 | 3039145.75 | 4067336.25 | 3624366.25 | 4095359.5 | 5684562 | 5107968 | 5773514 | 5240789.5 | 6770302.5 | 10151934 | 7720914 | 10049352 | 3680465.75 | 2564547.25 | 6800108 | 5643969 | 5670339 | 0 | 7 | 2 | 12 | 2 | 12 | 2.20 | 1.64 | 1.34 | 1.00 |
| Q61033 | Tmpo | Lamina-associated polypeptide 2, isoforms alpha/zeta | 737.94 | 25.97 | 75168 | 12 | 4216975 | 2238663.25 | 2708178.5 | 2511581.25 | 2316937 | 3364739.5 | 4776040 | 4205402.5 | 2894091 | 2534569.5 | 1660559.5 | 2128492 | 1690574.88 | 2468295.5 | 1994614.25 | 3546784 | 2291515.25 | 4369545.5 | 3453658.5 | 3133056.25 | 2 | 7 | 0 | 12 | 2 | 12 | 0.71 | 0.56 | 1.20 | 0.94 |
| P62329 | Tmsb4x | Thymosin beta-4 | 93.01 | 54.55 | 5053 | 2 | 8261753.5 | 7529783 | 6927429.5 | 12027363 | 4758757 | 11756675 | 6623982 | 7770494.5 | 4789711 | 8331191.5 | 6431981 | 3396202.75 | 4608032.5 | 6017787 | 3234264.75 | 10805646 | 22119038 | 10087836 | 11456546 | 9575092 | 2 | 4 | 2 | 4 | 2 | 4 | 0.60 | 0.60 | 1.62 | 1.63 |
| Q9CZW5 | Tomm70 | Mitochondrial import receptor subunit TOM70 | 637.83 | 21.6 | 67590 | 12 | 19442726 | 23894000 | 21517074 | 19766492 | 20738092 | 18661022 | 24172026 | 22814826 | 20957520 | 18433304 | 14953526 | 15488843 | 15166697 | 14273428 | 18711490 | 23516640 | 18685278 | 19344990 | 19091856 | 21191560 | 0 | 7 | 2 | 12 | 2 | 12 | 0.75 | 0.75 | 0.97 | 0.97 |
| P63028 | Tpt1 | Translationally-controlled tumor protein | 416.63 | 45.93 | 19462 | 6 | 12700666 | 13387536 | 15328356 | 8758859 | 22009476 | 7821847 | 14378943 | 12714519 | 16373450 | 8015683.5 | 5910026.5 | 10803955 | 5481636 | 2639217 | 7485891.5 | 15169318 | 5760480.5 | 7821905 | 7062240.5 | 6405474 | 1 | 3 | 2 | 5 | 2 | 5 | 0.45 | 0.54 | 0.58 | 0.71 |
| A2APB8 | Tpx2 | Targeting protein for Xklp2 | 116.35 | 4.03 | 85894 | 3 | 2538195 | 2828537 | 2030094.75 | 1790090.25 | 1748067.75 | 3204617.5 | 2357157.5 | 1633654.5 | 1414462.75 | 1430975 | 1137939 | 630592.94 | 887002.12 | 1520624.38 | 950536.38 | 2609980.25 | 927241.56 | 1855177.62 | 2166354 | 1853723 | 0 | 11 | 2 | 12 | 2 | 12 | 0.47 | 0.51 | 0.86 | 0.94 |
| Q61510 | Trim25 | E3 ubiquitin/ISG15 ligase TRIM25 | 165.01 | 6.94 | 71726 | 4 | 5902786.5 | 3228285.25 | 5112480 | 3915305.75 | 5566492 | 2911935 | 5270226 | 5180984.5 | 6144201 | 4459854 | 1990821.12 | 4659185 | 1588432.5 | 1261317.62 | 2495482.75 | 4096069.25 | 3294960.25 | 3636313.5 | 4669466 | 4186648.5 | 2 | 8 | 2 | 6 | 2 | 8 | 0.51 | 0.50 | 0.84 | 0.83 |
| Q80VI1 | Trim56 | E3 ubiquitin-protein ligase TRIM56 | 69.54 | 2.72 | 79513 | 1 | 480550.44 | 387312.47 | 269205.44 | 280460.5 | 297999.16 | 400112.38 | 464090.34 | 331981.19 | 602560.38 | 247824.03 | 2 | 192712.23 | 264234.91 | 263507.38 | 209170.81 | 352204.62 | 203347.08 | 262363.19 | 344840.78 | 263446.03 | 0 | 6 | 2 | 6 | 2 | 6 | 0.54 | 0.45 | 0.83 | 0.70 |
| Q8K1J6 | Trnt1 | CCA tRNA nucleotidyltransferase 1, mitochondrial | 212.47 | 11.52 | 49895 | 5 | 38374378 | 43606618 | 56773978 | 76890282 | 61005202 | 45659674 | 38743090 | 60897226 | 68951514 | 76985914 | 24955058 | 33170368 | 38026086 | 41416454 | 42814306 | 38648754 | 38575538 | 57220186 | 63771038 | 64867174 | 2 | 12 | 2 | 8 | 2 | 12 | 0.65 | 0.62 | 0.95 | 0.90 |
| Q9WTR1 | Trpv2 | Transient receptor potential cation channel subfamily V member 2 | 453.96 | 10.71 | 85965 | 7 | 7199856 | 4911928 | 6331832 | 6757846.5 | 6298708 | 7770425.5 | 6756932 | 5097009 | 7055648 | 9462220 | 8470749 | 7972006 | 11748722 | 9570345 | 10221135 | 6140808 | 8154650 | 10254779 | 8057335.5 | 8181157.5 | 0 | 4 | 2 | 7 | 2 | 7 | 1.52 | 1.33 | 1.29 | 1.13 |
| Q3KRE8 | Tubb2b | Tubulin beta-2B chain | 2111.15 | 58.2 | 49953 | 31 | 1807671.38 | 1968283.5 | 2257569.5 | 2286151.25 | 2126188 | 1709265.88 | 1695818.38 | 2487098 | 2050119 | 2434077.5 | 2 | 1272916.62 | 1600081.38 | 1389774.75 | 1763336.12 | 1776069.12 | 2044303.12 | 2495019.75 | 2547721 | 2824471.5 | 0 | 8 | 2 | 6 | 2 | 8 | 0.58 | 0.58 | 1.12 | 1.13 |
| Q91YR1 | Twf1 | Twinfilin-1 | 164.4 | 11.71 | 40079 | 3 | 9463100 | 9279111 | 6825505 | 8495180 | 9009675 | 9120412 | 9601400 | 7738679 | 8248810 | 8633291 | 6679430 | 5992468 | 7119808 | 6881073.5 | 6150404.5 | 9195612 | 10269293 | 8867696 | 8993438 | 9325793 | 2 | 9 | 0 | 6 | 2 | 9 | 0.76 | 0.76 | 1.08 | 1.08 |
| Q5M7W1 | Txnip | Thioredoxin-interacting protein | 157.4 | 7.11 | 44018 | 3 | 1554157.38 | 1729114.75 | 1076746 | 1535635.5 | 1288496.12 | 1651214.25 | 1704915.25 | 1318544.5 | 1528536 | 1028645.12 | 1196712.75 | 1130762.75 | 956625.19 | 589353 | 729589.5 | 909123.12 | 1565760.88 | 1504573.38 | 1195876.5 | 1306505.75 | 2 | 9 | 2 | 7 | 2 | 9 | 0.64 | 0.64 | 0.90 | 0.90 |
| Q99K23 | Ufsp2 | Ufm1-specific protease 2 | 265.14 | 11.71 | 52515 | 5 | 1458187.5 | 1733843.62 | 1584242 | 1971278.25 | 2688134 | 1601678.12 | 2737949.5 | 1696991.12 | 1684853.62 | 1122219.12 | 2734549 | 3357247 | 4470830.5 | 1742195 | 2929767.25 | 1904241.25 | 1529679 | 1960265 | 804756.12 | 1731051 | 2 | 9 | 2 | 12 | 2 | 12 | 1.61 | 1.72 | 0.84 | 0.90 |
| Q8R5H1 | Usp15 | Ubiquitin carboxyl-terminal hydrolase 15 | 380.48 | 8.46 | 112325 | 8 | 4812414 | 5641152 | 4847889 | 5492666.5 | 4554627.5 | 4280706 | 4907044 | 4627096 | 4318142 | 4426590.5 | 3344385.25 | 4199675 | 4374966.5 | 3598892.5 | 3277001.5 | 4999270.5 | 4902428.5 | 5055083.5 | 4316238 | 4880080.5 | 0 | 12 | 2 | 6 | 2 | 12 | 0.74 | 0.83 | 0.95 | 1.07 |
| Q8BY87 | Usp47 | Ubiquitin carboxyl-terminal hydrolase 47 | 103.39 | 1.82 | 157455 | 2 | 329708.56 | 696934.75 | 708609.12 | 571888.88 | 1214668.25 | 277669.16 | 950425.5 | 691276.19 | 1360077.75 | 664822.88 | 2 | 599707.31 | 241500.61 | 2 | 2 | 321183.03 | 2 | 2 | 430001.72 | 363789.22 | 2 | 2 | 1 | 3 | 2 | 3 | 0.24 | 0.21 | 0.32 | 0.28 |
| Q920Q4 | Vps16 | Vacuolar protein sorting-associated protein 16 homolog | 313.08 | 9.54 | 94928 | 5 | 1092791.38 | 1241800.38 | 1539708.62 | 1619977.25 | 2210453.75 | 773375.88 | 1317035.75 | 1677803.25 | 1659838.25 | 1470818.62 | 510353.84 | 810719.81 | 1010704.94 | 274139.19 | 536890.81 | 1712562.5 | 522165.94 | 1257584.25 | 671575.44 | 1445096 | 0 | 8 | 2 | 8 | 2 | 8 | 0.41 | 0.46 | 0.73 | 0.81 |
| P46467 | Vps4b | Vacuolar protein sorting-associated protein 4B | 318.58 | 16.67 | 49419 | 6 | 4427319.5 | 5374847 | 5165953.5 | 3213315.5 | 4034591.75 | 3680319 | 4795621.5 | 5872638 | 3799101.25 | 4982728.5 | 9579873 | 9014684 | 7938859 | 5036418.5 | 6282244.5 | 4045603.5 | 2614145.5 | 4888478.5 | 3280572.5 | 4648136 | 2 | 9 | 1 | 7 | 2 | 9 | 1.70 | 1.64 | 0.88 | 0.84 |
| Q3UVL4 | Vps51 | Vacuolar protein sorting-associated protein 51 homolog | 361.29 | 10.1 | 86187 | 7 | 3536787.75 | 3066622.75 | 2965203.25 | 2621920 | 3917567.25 | 2697742.75 | 3327398.75 | 3304563.25 | 3803573.5 | 3063523.75 | 1518954.88 | 2065540.25 | 2593693.75 | 2708926.5 | 2658056.75 | 3604963.75 | 3306237.75 | 3008309 | 3223568 | 3143753.75 | 2 | 10 | 1 | 12 | 2 | 12 | 0.72 | 0.71 | 1.01 | 1.01 |
| Q8BH43 | Wasf2 | Wiskott-Aldrich syndrome protein family member 2 | 574.4 | 23.54 | 54074 | 9 | 8905475 | 8737580 | 7916331.5 | 7972427.5 | 7971929 | 8633420 | 7722390 | 7030047 | 7204585 | 8038061 | 6544242 | 7176856 | 6487607 | 7156739 | 4827495 | 6885549 | 8410950 | 8636886 | 5919564.5 | 7909005.5 | 0 | 6 | 2 | 12 | 2 | 12 | 0.78 | 0.83 | 0.91 | 0.98 |
| Q9CQV8 | Ywhab | 14-3-3 protein beta/alpha | 998 | 69.11 | 28086 | 17 | 13124176 | 13731282 | 13586167 | 14715585 | 12299983 | 11910798 | 10042159 | 10495820 | 9924122 | 10697516 | 13562094 | 12299544 | 10827705 | 12269950 | 12172022 | 13602864 | 12572039 | 12881728 | 11091309 | 13087585 | 2 | 6 | 1 | 1 | 2 | 6 | 0.91 | 1.15 | 0.94 | 1.19 |
| P15535 | B4galt1 | Beta-1,4-galactosyltransferase 1 | 110.41 | 6.77 | 44411 | 2 | 234257.55 | 257301.67 | 2 | 329867.66 | 415033.53 | 304749.88 | 319913.84 | 252402.5 | 294196.59 | 2 | 2 | 2 | 2 | 2 | 2 | 246571.19 | 2 | 2 | 301713.75 | 324450.59 | 2.5 | 9 | 2.5 | 10 | 2.5 | 10 | 0.00 | 0.00 | 0.71 | 0.75 |
| P51670 | Ccl9 | C-C motif chemokine 9 | 76.53 | 17.21 | 13871 | 2 | 2 | 2 | 2 | 2 | 2 | 2 | 2 | 2 | 2 | 2 | 768454.06 | 473038.34 | 2 | 361407.91 | 524812.62 | 2 | 2 | 2 | 2 | 2 | 2.5 | 12.5 | 2.5 | 12.5 | 2.5 | 12.5 | 212771.49 | 212771.49 | 1.00 | 1.00 |
| P10818 | Cox6a1 | Cytochrome c oxidase subunit 6A1, mitochondrial | 64.06 | 18.92 | 12301 | 1 | 2 | 2 | 2 | 2 | 2 | 2 | 2 | 2 | 2 | 2 | 300907.22 | 202004.3 | 178053.77 | 2 | 159018.69 | 2 | 2 | 2 | 2 | 2 | 2.5 | 12.5 | 2.5 | 12.5 | 2.5 | 12.5 | 83998.60 | 83998.60 | 1.00 | 1.00 |
| Q61703 | Itih2 | Inter-alpha-trypsin inhibitor heavy chain H2 | 78.81 | 1.69 | 105928 | 1 | 2 | 2 | 2 | 2 | 2 | 2 | 2 | 2 | 2 | 2 | 422157.69 | 262719.94 | 429381.56 | 513188.09 | 2 | 2 | 2 | 2 | 2 | 2 | 2.5 | 12.5 | 2.5 | 12.5 | 2.5 | 12.5 | 162744.93 | 162744.93 | 1.00 | 1.00 |
| F2Z3T4 | Mbnl2 | Muscleblind-like protein 2 | 162.24 | 8.85 | 40156 | 4 | 2 | 2 | 2 | 2 | 2 | 2 | 2 | 2 | 2 | 2 | 1115091.75 | 675604 | 506332.88 | 410095.5 | 2 | 2 | 163488.03 | 2 | 2 | 143535.88 | 2.5 | 7.5 | 2.5 | 7.5 | 2.5 | 7.5 | 270712.61 | 270712.61 | 30702.99 | 30702.99 |
| Q7M0E7 | Mrpl14 | 39S ribosomal protein L14, mitochondrial | 69.4 | 6.9 | 15913 | 1 | 2 | 2 | 2 | 2 | 2 | 2 | 2 | 2 | 2 | 2 | 718515.69 | 419700.66 | 289668.5 | 2 | 264955.31 | 2 | 2 | 2 | 2 | 2 | 2.5 | 12.5 | 2.5 | 12.5 | 2.5 | 12.5 | 169284.22 | 169284.22 | 1.00 | 1.00 |
| Q99104 | Myo5a | Unconventional myosin-Va | 56.18 | 0.86 | 215538 | 1 | 158264.8 | 232549.02 | 199760.5 | 2 | 169530.36 | 220468.92 | 205396.69 | 222155.73 | 249565.78 | 200735.02 | 2 | 2 | 2 | 2 | 2 | 237158.28 | 2 | 2 | 2 | 2 | 2.5 | 7 | 0 | 4 | 2.5 | 7 | 0.00 | 0.00 | 0.31 | 0.22 |
| Q8K2Z4 | Ncapd2 | Condensin complex subunit 1 | 113.69 | 1.94 | 155665 | 2 | 650825.5 | 1005955.62 | 2 | 562503.19 | 915220.88 | 806841.94 | 711347.06 | 2 | 1247619 | 519850.62 | 2 | 2 | 2 | 2 | 2 | 601882.75 | 608341.19 | 663055.38 | 358341.25 | 511931.34 | 2.5 | 9 | 2.5 | 8 | 2.5 | 9 | 0.00 | 0.00 | 0.88 | 0.84 |
| O08776 | Ndufaf3 | NADH dehydrogenase [ubiquinone] 1 alpha subcomplex assembly factor 3 | 49.13 | 5.95 | 20696 | 1 | 2 | 2 | 2 | 2 | 2 | 2 | 2 | 2 | 2 | 2 | 490493.22 | 2 | 673697.75 | 608096 | 641214.25 | 2 | 2 | 2 | 2 | 2 | 2.5 | 12.5 | 2.5 | 12.5 | 2.5 | 12.5 | 241350.32 | 241350.32 | 1.00 | 1.00 |
| Q4KM92 | Pus1 | tRNA pseudouridine synthase A | 43.33 | 2.36 | 47515 | 1 | 2 | 2 | 2 | 2 | 2 | 2 | 2 | 2 | 2 | 2 | 772273.25 | 597903.56 | 497151.59 | 2 | 537549.75 | 2 | 2 | 2 | 2 | 2 | 2.5 | 12.5 | 2.5 | 12.5 | 2.5 | 12.5 | 240488.02 | 240488.02 | 1.00 | 1.00 |
| Q9JM13 | Rabgef1 | Rab5 GDP/GTP exchange factor | 39.75 | 1.43 | 56869 | 1 | 105332.98 | 280560.41 | 224280.27 | 2 | 579116.69 | 2 | 266132.72 | 399723.75 | 588708.88 | 133151.39 | 2 | 2 | 2 | 2 | 2 | 250354 | 2 | 2 | 2 | 2 | 2.5 | 5 | 2.5 | 4 | 2.5 | 5 | 0.00 | 0.00 | 0.21 | 0.18 |
| Q61609 | Slc20a1 | Sodium-dependent phosphate transporter 1 | 61.15 | 1.47 | 74153 | 1 | 2 | 2 | 2 | 2 | 2 | 2 | 2 | 2 | 2 | 2 | 1769839.75 | 1187912.75 | 1417739.62 | 2 | 1068840.12 | 2 | 2 | 2 | 2 | 2 | 2.5 | 12.5 | 2.5 | 12.5 | 2.5 | 12.5 | 544433.42 | 544433.42 | 1.00 | 1.00 |
| O70579 | Slc25a17 | Peroxisomal membrane protein PMP34 | 126.54 | 5.21 | 34413 | 2 | 2 | 2 | 2 | 2 | 2 | 2 | 2 | 2 | 2 | 2 | 550915.38 | 178131.73 | 258723.31 | 520870.75 | 2 | 2 | 2 | 2 | 2 | 2 | 2.5 | 12.5 | 2.5 | 12.5 | 2.5 | 12.5 | 150864.32 | 150864.32 | 1.00 | 1.00 |
| Q80VL1 | Tdrkh | Tudor and KH domain-containing protein | 41.01 | 2.32 | 62134 | 1 | 2 | 2 | 2 | 2 | 2 | 2 | 2 | 2 | 2 | 2 | 181581.39 | 124069.71 | 160362.06 | 2 | 219937.55 | 2 | 2 | 2 | 2 | 2 | 2.5 | 12.5 | 2.5 | 12.5 | 2.5 | 12.5 | 68595.27 | 68595.27 | 1.00 | 1.00 |

Supplementary table 3: List of modulated proteins by treatment with PBk11, selected by the Mann Whitney U test (p<0.05)

| accession | gene_name | description | protein_set_score | coverage | MW | #peptides | abund.CTRL #1 | abund.CTRL #2 | abund.CTRL #3 | abund.CTRL #4 | abund.CTRL #5 | abund.ERM #1 | abund.ERM #2 | abund.ERM #3 | abund.ERM #4 | abund.ERM #5 | abund.PR101#1 | abund.PR101#2 | abund.PR101#3 | abund.PR101#4 | abund.PR101#5 | abund.PBk11 #1 | abund.PBk11 #2 | abund.PBk11 #3 | abund.PBk11 #4 | abund.PBk11 #5 | U ctrl vs PR101 | U ctrl vs PBK11 | U ERM vs PR101 | U ERM vs PBK11 | Max U PR101 | Max U PBk11 | ratio PR101 Ctrl | ratio PR101 ERM | ratio PBk11 Ctrl | ratio PBk11 ERM |
|---|---|---|---|---|---|---|---|---|---|---|---|---|---|---|---|---|---|---|---|---|---|---|---|---|---|---|---|---|---|---|---|---|---|---|---|---|
| P60710 | Actb | Actin, cytoplasmic 1 | 4410.089 | 93.33 | 41737 | 59 | 1903567.75 | 3456343 | 1590567.5 | 2177505.25 | 2814027.25 | 1746062.88 | 2369415 | 1691173.88 | 3222573.5 | 2456823.75 | 4577360 | 5946585 | 4084039.5 | 2183052.5 | 3250668.75 | 912575.69 | 878834.81 | 1134181.62 | 966919.25 | 969713.25 | 3 | 0 | 3 | 0 | 3 | 0 | 1.678252195 | 1.74487378 | 0.407152926 | 0.423315678 |
| Q78IK2 | Atp5md | ATP synthase F(0) complex subunit k, mitochondrial | 228.84 | 44.83 | 6382 | 3 | 7492585 | 6021504 | 8170398 | 8883460 | 7988478 | 7433931 | 6588542.5 | 8779794 | 6479979 | 9199603 | 5081172.5 | 5363472 | 6653119 | 6803642.5 | 6982287 | 9853499 | 11115798 | 9575196 | 10097592 | 9661366 | 3 | 0 | 6 | 0 | 6 | 0 | 0.800999911 | 0.802552201 | 1.304671037 | 1.307199411 |
| Q923D2 | Blvrb | Flavin reductase (NADPH) | 485.1 | 52.43 | 22197 | 6 | 2 | 208048.06 | 129867.8 | 1111843.12 | 587203.44 | 1038699.06 | 404462.66 | 954894.12 | 628881.62 | 198019.28 | 14671318 | 6608738 | 6307858 | 8127343 | 6285399 | 1212510.25 | 1520904.88 | 1950822.5 | 1578359 | 1225602.38 | 0 | 0 | 0 | 0 | 0 | 0 | 20.6192389 | 13.02363392 | 3.676156017 | 2.32195332 |
| Q9D8S9 | Bola1 | BolA-like protein 1 | 52.91 | 11.68 | 14379 | 1 | 1291989 | 1512927.5 | 1583565.75 | 1672233.75 | 1496165.12 | 1506708.88 | 1604919.75 | 1232906 | 1290167.88 | 1557782.12 | 1581180.62 | 1359919.88 | 1187520 | 1436074.38 | 1659663.5 | 1763800.38 | 2460316 | 1676388.62 | 1676782.25 | 2172626.25 | 9 | 0 | 12 | 0 | 12 | 0 | 0.955997357 | 1.004431535 | 1.290203372 | 1.355569598 |
| Q64152 | Btf3 | Transcription factor BTF3 | 267.65 | 33.82 | 22031 | 4 | 2 | 371407.34 | 1088299.12 | 1240705.38 | 664365.12 | 600144.12 | 326985.41 | 617994.25 | 456508.38 | 542940.56 | 1159616.38 | 1160822.5 | 2085161 | 2547377.5 | 1916867.5 | 1623086 | 1833734.12 | 1485502.75 | 1652312.88 | 1747117.62 | 2 | 0 | 0 | 0 | 2 | 0 | 2.636085456 | 3.485789504 | 2.479138591 | 3.278253085 |
| O88668 | Creg1 | Protein CREG1 | 232.76 | 17.73 | 24452 | 4 | 2844224.75 | 2328332 | 1808015.5 | 2865231.5 | 2694705.75 | 3335767 | 3588065.75 | 3052793.5 | 4687482 | 4756861 | 3787437.75 | 3041952 | 4271372 | 5075047.5 | 4898390 | 5104928.5 | 6124831 | 5883303 | 5480842.5 | 6140475.5 | 0 | 0 | 9 | 0 | 9 | 0 | 1.680489876 | 1.085126029 | 2.291324806 | 1.4795544 |
| Q62426 | Cstb | Cystatin-B | 161.85 | 41.84 | 11046 | 3 | 8472974 | 9935852 | 7092119.5 | 10921310 | 9204859 | 11780998 | 8575034 | 9470133 | 7088252 | 9584553 | 10517191 | 7793224 | 9727302 | 13101336 | 9661600 | 13334215 | 23149752 | 15108231 | 13749050 | 16594676 | 9 | 0 | 7 | 0 | 9 | 0 | 1.11338737 | 1.092511361 | 1.795772643 | 1.762101913 |
| P00375 | Dhfr | Dihydrofolate reductase | 188.71 | 16.04 | 21606 | 3 | 4078497.25 | 3703753.5 | 3185696.75 | 2970118 | 3949301 | 2994072.75 | 3258373 | 2972876.75 | 3482712 | 2778173 | 2069364 | 3449049.75 | 2873216 | 2 | 3145160.5 | 2475836 | 1860836 | 1825632.62 | 2116349.5 | 2612738.5 | 3 | 0 | 8 | 0 | 8 | 0 | 0.644968741 | 0.744972082 | 0.60888743 | 0.703296312 |
| Q9D8N0 | Eef1g | Elongation factor 1-gamma | 1365.66 | 56.06 | 50061 | 21 | 170312690 | 175228626 | 184085762 | 184949970 | 168093346 | 179416610 | 168894690 | 178720290 | 162700370 | 170140674 | 211077890 | 225902082 | 177581234 | 178396434 | 182660818 | 204462722 | 203189874 | 206756370 | 203438162 | 195869618 | 6 | 0 | 4 | 0 | 6 | 0 | 1.105303253 | 1.134608103 | 1.148465784 | 1.178915 |
| P68101 | Eif2s1 | Eukaryotic translation initiation factor 2 subunit 1 | 1203.76 | 69.84 | 36108 | 21 | 76437002 | 83664978 | 76139722 | 82376266 | 76909202 | 78707594 | 74851866 | 74594354 | 70109626 | 78879202 | 89454978 | 82565130 | 74207770 | 71941394 | 79066874 | 88145082 | 86855866 | 84762954 | 89289274 | 92938602 | 12 | 0 | 8 | 0 | 12 | 0 | 1.004320755 | 1.053278261 | 1.117475136 | 1.171948565 |
| Q9R0P3 | Esd | S-formylglutathione hydrolase | 1127.09 | 71.63 | 31320 | 17 | 65296098 | 76205842 | 46468122 | 51129030 | 57718778 | 73324242 | 75570362 | 63368618 | 69002194 | 71205010 | 184859842 | 195729378 | 162199010 | 148743954 | 158067074 | 78615178 | 86860162 | 105089498 | 92808050 | 95206258 | 0 | 0 | 0 | 0 | 0 | 0 | 2.862358853 | 2.41041289 | 1.544984963 | 1.301042902 |
| Q8BTM8 | Flna | Filamin-A | 5224.061 | 41.9 | 281222 | 81 | 177941858 | 191113602 | 82794962 | 106899738 | 121273986 | 169158434 | 162421330 | 125754258 | 116163794 | 140355986 | 232483986 | 232977810 | 182502610 | 178620466 | 181363410 | 211277282 | 208845538 | 217285538 | 221223218 | 220296962 | 3 | 0 | 0 | 0 | 3 | 0 | 1.482224253 | 1.411981388 | 1.586603276 | 1.511413871 |
| P09528 | Fth1 | Ferritin heavy chain | 641.35 | 56.04 | 21067 | 11 | 21131550 | 18141730 | 19321128 | 34893818 | 29652728 | 27886138 | 25057054 | 55739530 | 54762290 | 55680038 | 171254306 | 172532530 | 283518574 | 253427186 | 282415842 | 384739938 | 554799298 | 602124482 | 627704770 | 567048514 | 0 | 0 | 0 | 0 | 0 | 0 | 9.445645013 | 5.308136783 | 22.22182721 | 12.48792414 |
| P29391 | Ftl1 | Ferritin light chain 1 | 1017.49 | 73.22 | 20802 | 15 | 35454822 | 27618676 | 45772886 | 53905598 | 52433102 | 40681886 | 38502082 | 74889490 | 81044098 | 82448042 | 105100170 | 116605522 | 150582738 | 154706914 | 153572834 | 551825986 | 664956482 | 658574466 | 716652034 | 665532546 | 0 | 0 | 0 | 0 | 0 | 0 | 3.162710748 | 2.143079043 | 15.13832396 | 10.25785392 |
| Q00612 | G6pdx | Glucose-6-phosphate 1-dehydrogenase X | 1459.48 | 53.01 | 59263 | 27 | 77125362 | 81088738 | 74658970 | 74655490 | 76736322 | 74286426 | 86203354 | 89308994 | 82921066 | 81973754 | 150968338 | 156430050 | 119277306 | 100537866 | 119809418 | 107510306 | 97650178 | 102922778 | 105539194 | 110128682 | 0 | 0 | 0 | 0 | 0 | 0 | 1.683794196 | 1.560243484 | 1.362995065 | 1.262983431 |
| Q6P6V1 | Galnt11 | Polypeptide N-acetylgalactosaminyltransferase 11 | 40.34 | 1.15 | 69039 | 1 | 4871399.5 | 4628804 | 6119983 | 6543670 | 5589983.5 | 5645836 | 4830788 | 4855836.5 | 4994786.5 | 4605390.5 | 2111794.5 | 1950528.5 | 2592086 | 3194829.25 | 2603771.5 | 3667474 | 4564209.5 | 4307746.5 | 4141163.25 | 4216927 | 0 | 0 | 0 | 0 | 0 | 0 | 0.448695018 | 0.499466202 | 0.752959599 | 0.83815923 |
| P14901 | Hmox1 | Heme oxygenase 1 | 559.13 | 51.21 | 32929 | 10 | 523223.28 | 333599.88 | 281882.22 | 486132.62 | 260096.28 | 396796.94 | 492680.75 | 417891.44 | 255481.94 | 966386.88 | 4228262 | 4485262.5 | 9708963 | 9171823 | 7049210 | 1636047.5 | 2272107 | 1785306.5 | 2238398.5 | 1883088.38 | 0 | 0 | 0 | 0 | 0 | 0 | 18.37916625 | 13.69721678 | 5.207050444 | 3.880594896 |
| Q64433 | Hspe1 | 10 kDa heat shock protein, mitochondrial | 474.69 | 67.65 | 10963 | 7 | 46814846 | 31449030 | 30110614 | 41931030 | 25769508 | 46977722 | 42066230 | 36473850 | 31469470 | 44223418 | 50963494 | 44858726 | 48755798 | 56888150 | 59396762 | 47254426 | 74640634 | 50619538 | 51834294 | 47250902 | 1 | 0 | 1 | 0 | 1 | 0 | 1.481544163 | 1.296466555 | 1.542523077 | 1.349827855 |
| Q9JL16 | Isg20 | Interferon-stimulated gene 20 kDa protein | 43.75 | 2.67 | 32503 | 1 | 289973.44 | 261718.09 | 297002.19 | 301518.16 | 240103.88 | 290711.06 | 262436.28 | 229108.72 | 386360.19 | 319806.16 | 184280.02 | 111135 | 90112.27 | 2 | 128588.41 | 169233.66 | 2 | 182507.41 | 147685.86 | 167307.08 | 0 | 0 | 0 | 0 | 0 | 0 | 0.369784847 | 0.345411152 | 0.479557255 | 0.447948113 |
| P49717 | Mcm4 | DNA replication licensing factor MCM4 | 823.1 | 23.43 | 96736 | 15 | 7513412.5 | 7577932 | 13372276 | 8609271 | 9802424 | 8576464 | 8042229 | 9391006 | 8865856 | 11103449 | 13865468 | 12985963 | 10329315 | 7463710 | 11657063 | 7404898 | 5140945 | 5711311.5 | 4927006.5 | 5384682 | 8 | 0 | 6 | 0 | 8 | 0 | 1.201090988 | 1.224504972 | 0.609464549 | 0.621345408 |
| D0QMC3 | Mndal | Myeloid cell nuclear differentiation antigen-like protein | 604.6 | 22.49 | 60526 | 10 | 6893806.5 | 6762796 | 6899946 | 8467844 | 7044165 | 7065211 | 6683493.5 | 7077912.5 | 7274583 | 7961776 | 1991161.12 | 2951066.75 | 3462431.5 | 910768.69 | 4816349 | 4785538.5 | 5024533 | 6334546.5 | 5169587.5 | 4217970 | 0 | 0 | 0 | 0 | 0 | 0 | 0.391803223 | 0.391863862 | 0.70787903 | 0.707988589 |
| Q9JK81 | Myg1 | UPF0160 protein MYG1, mitochondrial | 369.88 | 19.47 | 42723 | 7 | 8160355 | 8206388 | 8268758 | 8292585.5 | 8954494 | 8508474 | 8506411 | 7575559.5 | 9184911 | 9312169 | 9020308 | 7586060.5 | 6620637 | 9071594 | 5879706 | 9504076 | 11756131 | 9371294 | 11114906 | 9337223 | 10 | 0 | 7 | 0 | 10 | 0 | 0.911555712 | 0.886064028 | 1.219686786 | 1.185578206 |
| P11928 | Oas1a | 2'-5'-oligoadenylate synthase 1A | 631.77 | 37.87 | 42429 | 10 | 3303631 | 3524393.25 | 3408435.5 | 3377048.25 | 4853134.5 | 3590256.75 | 5291204 | 4240534 | 5202379.5 | 5104196 | 10773100 | 10577076 | 12851542 | 5960533.5 | 10445943 | 3179996.5 | 2571737.25 | 3008893.5 | 3019295.75 | 3160463.75 | 0 | 0 | 0 | 0 | 0 | 0 | 2.740519534 | 2.160105971 | 0.809047273 | 0.637699467 |
| Q4FK66 | Prpf38a | Pre-mRNA-splicing factor 38A | 123.39 | 7.05 | 37437 | 2 | 627606.25 | 803540.38 | 644271.75 | 470452.09 | 508239.97 | 569244.44 | 545125.69 | 777755.62 | 639800.56 | 530060.5 | 518593.28 | 424101.97 | 369074 | 580062.62 | 728454.44 | 2 | 2 | 428394 | 2 | 351444.94 | 8 | 0 | 7 | 0 | 8 | 0 | 0.857954014 | 0.855747093 | 0.255342744 | 0.254685924 |
| P62192 | Psmc1 | 26S proteasome regulatory subunit 4 | 879.95 | 37.27 | 49185 | 15 | 22715296 | 21359824 | 22409186 | 24681490 | 20494524 | 23904400 | 17986380 | 24827388 | 20429402 | 25615522 | 29614318 | 27341712 | 26399240 | 27098434 | 29712988 | 27664566 | 30281016 | 29915560 | 33298300 | 30705396 | 0 | 0 | 0 | 0 | 0 | 0 | 1.255295453 | 1.243019232 | 1.360060924 | 1.346760144 |
| Q91V41 | Rab14 | Ras-related protein Rab-14 | 819.96 | 55.35 | 23897 | 12 | 33951214 | 29001582 | 30542042 | 29548914 | 25610644 | 33451598 | 30019318 | 31561936 | 32439342 | 32963508 | 29221190 | 23350526 | 29704788 | 27778422 | 30628138 | 35356142 | 39071226 | 35948770 | 39369298 | 35218194 | 10 | 0 | 1 | 0 | 10 | 0 | 0.946376749 | 0.876881282 | 1.244252676 | 1.152883228 |
| P61314 | Rpl15 | 60S ribosomal protein L15 | 717.1 | 47.55 | 24146 | 12 | 47592318 | 62604774 | 70160354 | 52638130 | 55058354 | 54288130 | 54766918 | 73084754 | 68448202 | 68353186 | 94986746 | 106756218 | 85142002 | 86707466 | 84388634 | 34729338 | 27784160 | 39717230 | 32489202 | 45706570 | 0 | 0 | 0 | 0 | 0 | 0 | 1.589914312 | 1.435942049 | 0.626363612 | 0.565704605 |
| O09167 | Rpl21 | 60S ribosomal protein L21 | 303.88 | 38.12 | 18562 | 6 | 38344534 | 39227422 | 42229910 | 41982594 | 35936282 | 35935538 | 43926834 | 41244518 | 34839654 | 46160114 | 37975914 | 36074238 | 34238458 | 35705746 | 37217510 | 53003118 | 58739866 | 51015458 | 53441762 | 54538458 | 3 | 0 | 7 | 0 | 7 | 0 | 0.916504076 | 0.896615024 | 1.36929823 | 1.339583093 |
| P14115 | Rpl27a | 60S ribosomal protein L27a | 337.53 | 32.43 | 16605 | 6 | 61261942 | 65413842 | 63601414 | 55034730 | 54850222 | 59794706 | 68245474 | 53551498 | 50403722 | 54411162 | 68832978 | 62496970 | 51773478 | 50278622 | 46837058 | 24157912 | 23149892 | 15184820 | 12620028 | 13894153 | 8 | 0 | 10 | 0 | 10 | 0 | 0.933559098 | 0.978396249 | 0.296529076 | 0.31077083 |
| P47915 | Rpl29 | 60S ribosomal protein L29 | 95.98 | 11.88 | 17587 | 2 | 38707466 | 43819890 | 45158330 | 49810546 | 6586432 | 40824594 | 36131098 | 44362238 | 37447570 | 45243742 | 31242586 | 35148966 | 32799324 | 39393994 | 34224954 | 52324550 | 58511962 | 56768114 | 53750122 | 58263698 | 6 | 0 | 2 | 0 | 6 | 0 | 0.938762077 | 0.847068605 | 1.518983048 | 1.370616563 |
| P62242 | Rps8 | 40S ribosomal protein S8 | 811.89 | 55.29 | 24205 | 12 | 101469178 | 110569450 | 99636418 | 95703322 | 88469770 | 104153346 | 100354914 | 87081946 | 83625466 | 85556050 | 94070922 | 87682602 | 61028106 | 59951138 | 61165186 | 37705414 | 39662934 | 22164404 | 19233892 | 17156836 | 1 | 0 | 6 | 0 | 6 | 0 | 0.733889927 | 0.789757567 | 0.274123203 | 0.294990933 |
| Q9WU81 | Slc37a2 | Glucose-6-phosphate exchanger SLC37A2 | 290.42 | 11.18 | 55073 | 5 | 8284388 | 8088368 | 5438423.5 | 9965466 | 9921874 | 13788850 | 13205134 | 11421332 | 11904541 | 13245604 | 16187891 | 15760414 | 17926422 | 18540176 | 16922654 | 15151308 | 14198181 | 13826449 | 16089322 | 17467638 | 0 | 0 | 0 | 0 | 0 | 0 | 2.046536856 | 1.342514561 | 1.840182791 | 1.207147668 |
| P08228 | Sod1 | Superoxide dismutase [Cu-Zn] | 303.98 | 41.56 | 15943 | 5 | 4445996.5 | 3917995.75 | 12267304 | 13643726 | 9986887 | 5378816.5 | 3109375 | 9691028 | 4807349 | 8332123 | 4255234 | 5658031 | 9143916 | 14160305 | 10941188 | 16195545 | 18213628 | 21294936 | 16866326 | 21270154 | 12 | 0 | 7 | 0 | 12 | 0 | 0.997667628 | 1.409978255 | 2.120120677 | 2.996312569 |
| Q62351 | Tfrc | Transferrin receptor protein 1 | 1208.87 | 30.14 | 85731 | 21 | 18225534 | 17123336 | 17031192 | 11489983 | 15566815 | 14231232 | 16261950 | 15696246 | 17438198 | 16183347 | 18365210 | 19177910 | 17272828 | 14655296 | 21044256 | 4678022 | 2313503 | 6171230 | 4403883 | 4090596.25 | 5 | 0 | 5 | 0 | 5 | 0 | 1.139464727 | 1.1341235 | 0.272634571 | 0.2713566 |
| P05064 | Aldoa | Fructose-bisphosphate aldolase A | 2284.05 | 79.4 | 39356 | 32 | 873672706 | 874774786 | 840246722 | 1013832770 | 885816706 | 1009478018 | 963755266 | 975091330 | 904854978 | 1050313922 | 614135874 | 651428674 | 697288066 | 810012418 | 688416642 | 1032909826 | 1217700098 | 1095962370 | 1275317506 | 1156400898 | 0 | 0 | 0 | 1 | 0 | 1 | 0.771171263 | 0.705880749 | 1.287399339 | 1.178402843 |
| P15999 | Atp5f1a | ATP synthase subunit alpha, mitochondrial | 2944.79 | 63.47 | 59754 | 42 | 7766863.5 | 7417389.5 | 7649394 | 7779600.5 | 6668269.5 | 8189972 | 5242943 | 7285566.5 | 6824187 | 7559088 | 4754507.5 | 5047347.5 | 5240527.5 | 6344206.5 | 5071641.5 | 8549246 | 9591793 | 7875812.5 | 9648108 | 9336178 | 0 | 0 | 1 | 1 | 1 | 1 | 0.709687605 | 0.753758021 | 1.20706294 | 1.282019534 |
| Q9D287 | Bcas2 | Pre-mRNA-splicing factor SPF27 | 63.21 | 5.78 | 26131 | 1 | 2 | 2 | 923708.31 | 2 | 2 | 1089048.75 | 2 | 2 | 2 | 2 | 2 | 2 | 2 | 2 | 723536.56 | 1426194.12 | 1069288.75 | 1249903.62 | 1219625.5 | 1157161 | 12 | 0 | 12 | 1 | 12 | 1 | 0.783297374 | 0.664377279 | 6.627763225 | 5.62153716 |
| Q9Z0H7 | Bcl10 | B-cell lymphoma/leukemia 10 | 96.39 | 8.58 | 25948 | 2 | 876198.69 | 834370.25 | 991918.56 | 971190.5 | 1184364.25 | 806719.56 | 1213505.62 | 881183.56 | 1167156 | 920283.12 | 2 | 478889.97 | 2 | 2 | 2 | 875730.75 | 407561.66 | 720654.56 | 652777.75 | 713016.88 | 0 | 1 | 0 | 1 | 0 | 1 | 0.098578387 | 0.095993701 | 0.693641888 | 0.675454873 |
| P80313 | Cct7 | T-complex protein 1 subunit eta | 2304.58 | 64.71 | 59652 | 33 | 100202282 | 113481114 | 115497986 | 124026946 | 103497242 | 112411538 | 100522530 | 111683706 | 99555314 | 115008602 | 132479162 | 117021098 | 124916234 | 129594778 | 130966026 | 121111114 | 131870674 | 130732914 | 133505162 | 128493874 | 1 | 1 | 0 | 0 | 1 | 1 | 1.140598069 | 1.177668511 | 1.159883739 | 1.197580982 |
| P11240 | Cox5a | Cytochrome c oxidase subunit 5A, mitochondrial | 388.94 | 38.36 | 16130 | 7 | 44838462 | 35949570 | 40842742 | 44274914 | 39375850 | 47295454 | 38330634 | 34163878 | 39460162 | 50218982 | 33860022 | 39141578 | 47631746 | 56691922 | 54409438 | 48988986 | 67034246 | 56872070 | 60359082 | 57591114 | 9 | 0 | 9 | 1 | 9 | 1 | 1.128862869 | 1.106295367 | 1.416812738 | 1.388488728 |
| Q8VCN5 | Cth | Cystathionine gamma-lyase | 942.24 | 40.95 | 43567 | 15 | 38931342 | 39684586 | 22701240 | 39828798 | 34159694 | 48102042 | 41071706 | 38456990 | 35483394 | 43035890 | 105159770 | 82884922 | 94313754 | 95149810 | 99341730 | 46959098 | 56565026 | 57118294 | 58389214 | 59422318 | 0 | 0 | 0 | 1 | 0 | 1 | 2.720106048 | 2.313121199 | 1.588391099 | 1.350734515 |
| P24268 | Ctsd | Cathepsin D | 576.48 | 19.66 | 44681 | 10 | 28492142 | 21037556 | 15338439 | 10309086 | 49814782 | 10294403 | 23875046 | 19274632 | 27578042 | 10057534 | 1218670.5 | 4259972 | 1393277.75 | 545463 | 2177100.25 | 2 | 10114845 | 5529561 | 7100870 | 8432397 | 0 | 0 | 0 | 1 | 0 | 1 | 0.076760778 | 0.105341674 | 0.249437354 | 0.34231217 |
| Q6Q899 | Ddx58 | Antiviral innate immune response receptor RIG-I | 240.08 | 6.8 | 105975 | 5 | 3704273.25 | 3759491 | 3360024.5 | 2085483 | 4515250 | 2445025.5 | 3720996.75 | 5571987.5 | 4484877 | 3936143.5 | 648171.25 | 2362976.75 | 2345742.25 | 539298.12 | 1750258.75 | 2240114.5 | 697283.5 | 1960737.25 | 1394739.5 | 2075040.75 | 2 | 1 | 0 | 0 | 2 | 1 | 0.438832539 | 0.379306297 | 0.480237887 | 0.415095141 |
| O88455 | Dhcr7 | 7-dehydrocholesterol reductase | 250.55 | 13.38 | 53919 | 5 | 8420224 | 8326428.5 | 6577549 | 9156462 | 7734344 | 7811316.5 | 7582275.5 | 9103084 | 8379915 | 6764625 | 12156979 | 10105335 | 16013407 | 12918322 | 16240539 | 12110634 | 11161273 | 11089765 | 9033594 | 12011790 | 0 | 1 | 0 | 1 | 0 | 1 | 1.676851161 | 1.701122942 | 1.377770625 | 1.397713329 |
| O70251 | Eef1b | Elongation factor 1-beta | 586.81 | 39.56 | 24694 | 9 | 131771842 | 121854370 | 128707994 | 149759090 | 110387106 | 139726210 | 120997874 | 124049722 | 122448898 | 137379346 | 107920082 | 105154066 | 96249626 | 107846266 | 102002162 | 147016594 | 188359506 | 156442082 | 169184274 | 151086946 | 0 | 1 | 0 | 0 | 0 | 1 | 0.808074768 | 0.805415065 | 1.263990932 | 1.259830623 |
| P57776 | Eef1d | Elongation factor 1-delta | 848.2 | 54.09 | 31293 | 13 | 107040474 | 101984394 | 112340730 | 117554170 | 100273450 | 113836402 | 102780050 | 93984738 | 93946170 | 107623186 | 74352722 | 82442754 | 77204906 | 74214418 | 71849442 | 113664906 | 120213322 | 124013970 | 123229770 | 125278306 | 0 | 1 | 0 | 1 | 0 | 1 | 0.704875784 | 0.742065792 | 1.124643734 | 1.183981154 |
| B0BNA7 | Eif3i | Eukaryotic translation initiation factor 3 subunit I | 460.44 | 31.69 | 36461 | 9 | 2621145 | 2257842 | 2700000.5 | 2388992 | 2740422.5 | 2645388 | 2068384.62 | 2512597.75 | 3021868.5 | 1766340.38 | 7253555.5 | 5549140 | 6511433 | 6064594 | 6546310 | 3024335.5 | 3507737 | 3520692.5 | 3247407.5 | 2735716.5 | 0 | 1 | 0 | 1 | 0 | 1 | 2.512120131 | 2.657191054 | 1.261833628 | 1.334702503 |
| Q9DC16 | Ergic1 | Endoplasmic reticulum-Golgi intermediate compartment protein | 152.45 | 9.66 | 32562 | 3 | 3864363.75 | 4039279.5 | 4870981 | 4051686 | 3267568 | 4070570 | 4009455.5 | 4029218 | 4189291.5 | 4597938 | 6824986 | 7070171 | 8384725 | 7331553 | 8133665 | 4438117 | 4922859 | 5580743 | 5117602 | 5355347.5 | 0 | 1 | 0 | 1 | 0 | 1 | 1.878437777 | 1.806290468 | 1.264796581 | 1.216218091 |
| O09172 | Gclm | Glutamate--cysteine ligase regulatory subunit | 282.64 | 22.99 | 30535 | 5 | 2476578.25 | 2534911.5 | 2105851.75 | 2268303 | 3340085 | 3564871.75 | 2651398 | 2589788.75 | 4122152.5 | 2757284.5 | 13735158 | 15253114 | 12957263 | 17591150 | 16191247 | 4133617.25 | 3622444.25 | 6385710.5 | 5451630.5 | 4619630.5 | 0 | 0 | 0 | 1 | 0 | 1 | 5.950773353 | 4.827895427 | 1.9026833 | 1.543657515 |
| Q8CI11 | Gnl3 | Guanine nucleotide-binding protein-like 3 | 720.79 | 27.14 | 60786 | 12 | 5920561 | 5999730.5 | 4526905.5 | 4673007 | 6393163.5 | 5907068 | 5196668.5 | 4184427 | 6392547 | 4708152 | 6943035.5 | 6490127.5 | 6221106.5 | 5209271.5 | 8117401 | 4386107.5 | 2537562.5 | 3061264.75 | 2797507 | 2582216.25 | 4 | 0 | 3 | 1 | 4 | 1 | 1.198724293 | 1.249805368 | 0.558443382 | 0.582240254 |
| P38647 | Hspa9 | Stress-70 protein, mitochondrial | 2894.79 | 69.51 | 73461 | 41 | 121863890 | 138234162 | 96252466 | 109482770 | 92253346 | 130259778 | 126977170 | 106917298 | 88876530 | 110469658 | 231566034 | 220905202 | 220763714 | 230550002 | 226099922 | 141427714 | 170057570 | 140447042 | 137563682 | 144059698 | 0 | 1 | 0 | 0 | 0 | 1 | 2.024568956 | 2.005118019 | 1.314411888 | 1.301783746 |
| P63038 | Hspd1 | 60 kDa heat shock protein, mitochondrial | 4175.81 | 76.44 | 60955 | 54 | 405753922 | 420495266 | 382470690 | 430740898 | 326575970 | 425424674 | 362507586 | 404076610 | 339642530 | 453447074 | 916978498 | 866707522 | 918442114 | 920899522 | 978072386 | 431340546 | 524064386 | 484915778 | 470334530 | 492476642 | 0 | 0 | 0 | 1 | 0 | 1 | 2.340291987 | 2.317819545 | 1.222322974 | 1.210585728 |
| Q9ESY9 | Ifi30 | Gamma-interferon-inducible lysosomal thiol reductase | 322.3 | 28.23 | 27784 | 6 | 4818800 | 4628118 | 3605326.5 | 3213951 | 8805051 | 3610780.5 | 4337029.5 | 4440209.5 | 5575459.5 | 2720234.5 | 739986.69 | 933777.38 | 1160654.5 | 621738.62 | 257705.12 | 2571911.5 | 3272537.75 | 2392246.5 | 1288238.38 | 1255300.62 | 0 | 1 | 0 | 1 | 0 | 1 | 0.148132336 | 0.17955491 | 0.429983996 | 0.521194357 |
| P05555 | Itgam | Integrin alpha-M | 1456.28 | 27.93 | 127481 | 22 | 46727342 | 57519530 | 54105570 | 48778522 | 46165426 | 53299618 | 52724342 | 36114754 | 51656490 | 41754538 | 41694810 | 45241870 | 50168674 | 38972046 | 43008194 | 37767218 | 35145010 | 33982422 | 28642822 | 27613392 | 3 | 0 | 8 | 1 | 8 | 1 | 0.864937688 | 0.930103307 | 0.644110498 | 0.692638687 |
| P11835 | Itgb2 | Integrin beta-2 | 2152.3 | 50.58 | 85026 | 32 | 57889934 | 60608126 | 68087306 | 73967066 | 63443658 | 60340942 | 64724178 | 55744202 | 53610658 | 58985786 | 42107182 | 45060026 | 53500018 | 55665018 | 54271026 | 54387646 | 53489918 | 52105238 | 49769862 | 47157346 | 0 | 0 | 2 | 1 | 2 | 1 | 0.773476217 | 0.854118422 | 0.792941699 | 0.875613365 |
| Q7TNC4 | Luc7l2 | Putative RNA-binding protein Luc7-like 2 | 578.43 | 25.77 | 46583 | 10 | 5125886.5 | 5753026.5 | 5992173.5 | 6881297 | 7304687 | 5237472.5 | 5134608 | 6575725 | 5708022 | 4164100.5 | 3343101.5 | 3488291.5 | 3801143 | 3651439.25 | 3840224 | 4154419.25 | 3974542.5 | 3526939 | 3960842.25 | 4335961 | 0 | 0 | 0 | 1 | 0 | 1 | 0.583577232 | 0.675773598 | 0.642452868 | 0.743950692 |
| Q68FL6 | Mars | Methionine--tRNA ligase, cytoplasmic | 1008.35 | 24.83 | 101431 | 20 | 19082300 | 21104194 | 21305934 | 22471806 | 21906924 | 19830814 | 20959400 | 19159894 | 20666870 | 20406876 | 17946640 | 24897210 | 22859640 | 17894662 | 25523470 | 22518204 | 22456006 | 23029682 | 23750310 | 24122402 | 10 | 1 | 10 | 0 | 10 | 1 | 1.030702073 | 1.080156989 | 1.094505871 | 1.147022207 |
| P25206 | Mcm3 | DNA replication licensing factor MCM3 | 1799.12 | 39.29 | 91546 | 27 | 54164006 | 60851942 | 66252262 | 60250966 | 49907858 | 54236674 | 52595670 | 57907114 | 51514802 | 53644718 | 69823474 | 64819026 | 64087230 | 63686546 | 65313694 | 46892130 | 50817306 | 45727882 | 49117522 | 41931402 | 4 | 1 | 0 | 0 | 4 | 1 | 1.124569555 | 1.214269029 | 0.804613899 | 0.868792627 |
| E9Q7G0 | Numa1 | Nuclear mitotic apparatus protein 1 | 2466.11 | 22.21 | 235630 | 40 | 19486082 | 22440948 | 25189754 | 26811114 | 22685102 | 22959490 | 18977908 | 21892120 | 20031708 | 22809720 | 15317192 | 13045566 | 12584389 | 11650330 | 11715564 | 14569204 | 15813670 | 16976060 | 15772472 | 19687302 | 0 | 1 | 0 | 1 | 0 | 1 | 0.551508331 | 0.602910571 | 0.710201333 | 0.77639424 |
| P23298 | Prkch | Protein kinase C eta type | 67.3 | 1.46 | 77919 | 1 | 179823.39 | 159095.2 | 172126.92 | 292217.19 | 278111.72 | 180224.48 | 192252.2 | 142421.42 | 230204.97 | 177740.75 | 2 | 2 | 63347.53 | 2 | 2 | 2 | 2 | 2 | 2 | 144311.05 | 0 | 0 | 0 | 1 | 0 | 1 | 0.058587968 | 0.068652494 | 0.133458909 | 0.156385129 |
| P60892 | Prps1 | Ribose-phosphate pyrophosphokinase 1 | 521.28 | 29.87 | 34834 | 10 | 1908584.25 | 1872750.62 | 1363867 | 1354321.25 | 1080350.62 | 2218772.25 | 2201345.5 | 1545751.12 | 1090540.88 | 1616451.88 | 2717379.75 | 2741849.5 | 1994902 | 1949452.5 | 2065803.75 | 1042386.56 | 1337658.12 | 907363.25 | 1047229.88 | 558474.12 | 0 | 1 | 6 | 1 | 6 | 1 | 1.513137012 | 1.322445577 | 0.645540031 | 0.564186556 |
| Q62159 | Rhoc | Rho-related GTP-binding protein RhoC | 681.5 | 65.8 | 22006 | 10 | 5657037.5 | 4961224 | 4148133 | 6285029.5 | 5093144 | 7321600.5 | 6023679 | 5114305 | 4865852 | 6714911 | 6185593 | 5593749 | 6089082.5 | 6098007 | 6316691.5 | 2948077.5 | 4329709.5 | 3103206.25 | 2273900.75 | 2078614.5 | 5 | 1 | 11 | 0 | 11 | 1 | 1.158295023 | 1.008081648 | 0.563539948 | 0.490457326 |
| Q9CQS8 | Sec61b | Protein transport protein Sec61 subunit beta | 161.01 | 37.5 | 9958 | 3 | 14511030 | 13737012 | 13301174 | 17340382 | 12870361 | 14551708 | 14236868 | 16459350 | 13303834 | 15610830 | 10089208 | 9109832 | 9717178 | 9010318 | 11199416 | 17623090 | 20536134 | 17465326 | 16914198 | 19503906 | 0 | 1 | 0 | 0 | 0 | 1 | 0.684587236 | 0.662408797 | 1.282646413 | 1.241092767 |
| P10852 | Slc3a2 | Amino acid transporter heavy chain SLC3A2 | 1538.43 | 43.35 | 58337 | 23 | 19888242 | 22148812 | 20073252 | 21779348 | 18065094 | 23246186 | 27277712 | 22806582 | 22682570 | 25025252 | 74960418 | 88608218 | 90566370 | 74906730 | 89635922 | 27318094 | 29236768 | 27323670 | 25414426 | 28803020 | 0 | 0 | 0 | 1 | 0 | 1 | 4.10650476 | 3.459050987 | 1.35448305 | 1.140927919 |
| Q91ZW3 | Smarca5 | SWI/SNF-related matrix-associated actin-dependent regulator of chromatin subfamily A member 5 | 461.51 | 7.99 | 121627 | 9 | 6787082 | 8142618.5 | 8106559.5 | 8262912 | 9132291 | 7800210 | 7277738 | 7491777.5 | 7536482.5 | 6979846 | 7097229 | 5473063 | 7352491.5 | 6731863 | 5459363 | 6025054.5 | 4882014.5 | 5225097.5 | 7143520 | 5203866 | 2 | 1 | 3 | 1 | 3 | 1 | 0.794282648 | 0.865932232 | 0.704390848 | 0.767931592 |
| Q62376 | Snrnp70 | U1 small nuclear ribonucleoprotein 70 kDa | 523.92 | 23.21 | 51992 | 10 | 15061200 | 23722930 | 22027248 | 23325986 | 17034648 | 16335981 | 20248828 | 24318458 | 14511913 | 23108676 | 19284014 | 17584504 | 17837614 | 15476033 | 12664371 | 9595069 | 7143686 | 16311627 | 13795266 | 9226812 | 7 | 1 | 7 | 1 | 7 | 1 | 0.818868127 | 0.840877929 | 0.55422897 | 0.569125715 |
| Q9D975 | Srxn1 | Sulfiredoxin-1 | 45.05 | 6.62 | 14149 | 1 | 884547 | 1015408.5 | 419838.62 | 2 | 540690 | 1012313.19 | 1355964.38 | 424432.44 | 1356292.12 | 1661937.12 | 2668304.5 | 2834787.5 | 2518493 | 2501508.75 | 2235425.25 | 1430154.5 | 1663177.5 | 1931213.62 | 1946348.88 | 1901633 | 0 | 0 | 0 | 1 | 0 | 1 | 4.46026251 | 2.195603576 | 3.101755131 | 1.526866332 |
| Q04750 | Top1 | DNA topoisomerase 1 | 601.75 | 14.86 | 90876 | 12 | 8873592 | 8886410 | 7750781 | 7003899.5 | 7276418 | 8703637 | 9793704 | 4336913 | 7251859 | 6491784.5 | 5254741.5 | 8275957 | 5009715.5 | 4162399.5 | 4934715 | 3266837 | 3864007.5 | 4876597 | 2727665.5 | 3854328 | 3 | 0 | 6 | 1 | 6 | 1 | 0.694565573 | 0.755580019 | 0.467175694 | 0.508214968 |
| P23591 | Tsta3 | GDP-L-fucose synthase | 172.47 | 11.53 | 35878 | 4 | 3979640.5 | 3377112.25 | 3461272.5 | 4158720 | 3604465 | 3038053 | 3348999.5 | 3658084.5 | 3132918 | 4045029.75 | 2827480.25 | 2858639.75 | 3993813.5 | 3059543.5 | 2763826 | 4282214.5 | 4425437.5 | 4379113 | 4065330 | 4302162.5 | 4 | 1 | 5 | 0 | 5 | 1 | 0.83435378 | 0.900146706 | 1.154621104 | 1.245668695 |
| Q99KC8 | Vwa5a | von Willebrand factor A domain-containing protein 5A | 1672.17 | 37.45 | 87143 | 26 | 97203218 | 98990634 | 115614418 | 107912442 | 115783762 | 82159258 | 97843546 | 93395466 | 108455882 | 92003738 | 35362374 | 44539886 | 46947962 | 35819954 | 48290238 | 85062826 | 67895530 | 79923730 | 77929010 | 78708538 | 0 | 0 | 0 | 1 | 0 | 1 | 0.393947062 | 0.445197639 | 0.727388197 | 0.822017829 |
| Q9JIX8 | Acin1 | Apoptotic chromatin condensation inducer in the nucleus | 625.8 | 8.82 | 150719 | 13 | 5637156.5 | 4327341 | 5628949 | 5763103 | 6790344.5 | 5614522.5 | 7231470.5 | 7198500.5 | 6137494 | 5002033 | 3362996.25 | 2986162.75 | 2702165.5 | 3615508 | 3506767.5 | 3966059.75 | 4420869 | 3939507.25 | 4204008 | 4560664.5 | 0 | 2 | 0 | 0 | 0 | 2 | 0.574614023 | 0.518650249 | 0.749322767 | 0.676343466 |
| P63259 | Actg1 | Actin, cytoplasmic 2 | 4416.749 | 93.33 | 41793 | 59 | 3193824 | 4651257 | 2134905.75 | 2053409.38 | 1554199.38 | 3416460 | 4374765.5 | 2078096.12 | 2568780.75 | 2515026 | 4711988.5 | 4185462.25 | 2221636.5 | 1383974.88 | 2108570.75 | 1735729.88 | 1731756.5 | 1226768.5 | 946602 | 960163.19 | 11 | 2 | 11 | 0 | 11 | 2 | 1.075365606 | 0.977162271 | 0.48581223 | 0.441447429 |
| Q8JZQ2 | Afg3l2 | AFG3-like protein 2 | 410.61 | 12.97 | 89519 | 9 | 6845131.5 | 7766787 | 3979045 | 4973322.5 | 5312291 | 4622065.5 | 6322183 | 5467263 | 5756979 | 7902616 | 7953209.5 | 8205233 | 11657622 | 13026175 | 12836860 | 5787243.5 | 8783274 | 8861670 | 9259989 | 8774111 | 0 | 2 | 0 | 2 | 0 | 2 | 1.858914909 | 1.785072309 | 1.435983479 | 1.378941194 |
| Q9CVB6 | Arpc2 | Actin-related protein 2/3 complex subunit 2 | 1257.46 | 67.33 | 34357 | 18 | 54058158 | 54715686 | 57175410 | 46980326 | 53451254 | 38431922 | 49896970 | 48045818 | 60570462 | 50567038 | 30559312 | 51280198 | 35105486 | 20360430 | 41254186 | 45367218 | 31678088 | 39573426 | 33185108 | 35796450 | 1 | 0 | 5 | 2 | 5 | 2 | 0.670317039 | 0.721417388 | 0.696747912 | 0.749863168 |
| P29066 | Arrb1 | Beta-arrestin-1 | 120.39 | 6.7 | 47020 | 3 | 728971 | 702386.69 | 533680.19 | 327714.69 | 610151.88 | 328667.5 | 659856.88 | 543208.44 | 761962.44 | 349985.66 | 2 | 237990.5 | 297565.94 | 66171.43 | 175562.75 | 412344.41 | 167970.64 | 305472.34 | 297133.78 | 305457.84 | 0 | 1 | 0 | 2 | 0 | 2 | 0.267763763 | 0.294019075 | 0.512720634 | 0.562994951 |
| P50429 | Arsb | Arylsulfatase B | 499.22 | 18.16 | 59647 | 9 | 26900540 | 25801130 | 28698422 | 28841544 | 41645642 | 16307439 | 25303292 | 17455828 | 23083110 | 14600540 | 982859.5 | 2694667.25 | 2482796.75 | 1703945.5 | 3113993.75 | 16842870 | 12119315 | 10708555 | 11000842 | 12631576 | 0 | 0 | 0 | 2 | 0 | 2 | 0.072279014 | 0.113470171 | 0.416777223 | 0.654294793 |
| P29418 | Atp5f1e | ATP synthase subunit epsilon, mitochondrial | 49.29 | 15.69 | 5767 | 1 | 6547974.5 | 6319299 | 6488471 | 6383379.5 | 5801830 | 7712574.5 | 7063969 | 6951554 | 5727816 | 7624739 | 7462482 | 6368167.5 | 8342842.5 | 9833746 | 7747806.5 | 7195991 | 8685536 | 8201833 | 9197844 | 8701264 | 3 | 0 | 6 | 2 | 6 | 2 | 1.260426191 | 1.133247008 | 1.331046233 | 1.196741366 |
| Q8K354 | Cbr3 | Carbonyl reductase [NADPH] 3 | 41.84 | 3.97 | 30953 | 1 | 723297.19 | 797801.12 | 350954.75 | 431311.88 | 412297.69 | 943404.31 | 2 | 657069.69 | 554684.5 | 782372.25 | 942119.75 | 910504.94 | 823620.25 | 893761.69 | 704873.81 | 1026945.75 | 1060072.38 | 954834.44 | 812542 | 920030.38 | 2 | 0 | 6 | 2 | 6 | 2 | 1.574157406 | 1.455262223 | 1.758106805 | 1.625318033 |
| Q7TS56 | Cbr4 | Carbonyl reductase family member 4 | 102.59 | 8.05 | 25287 | 1 | 568247.06 | 780874.12 | 543262.12 | 695135.94 | 566350.06 | 2 | 717087.38 | 530505.44 | 453963.5 | 734855.19 | 479897.94 | 324699.06 | 516620.44 | 2 | 474756.72 | 756324.69 | 1213676.25 | 787501.94 | 793367.44 | 766045.75 | 0 | 2 | 7.5 | 0 | 7.5 | 2 | 0.569451676 | 0.737139304 | 1.368768221 | 1.771832266 |
| Q4VAA2 | Cdv3 | Protein CDV3 | 215.54 | 18.86 | 29729 | 3 | 3035774 | 3331059 | 2842529 | 3883012 | 2955565 | 3486896.5 | 2878235.25 | 3542792.5 | 3258618.25 | 4275869.5 | 3504389 | 3555693.25 | 2752858.75 | 2973024 | 2754446.75 | 3902453 | 5055901 | 4400552.5 | 3951605.5 | 4460441.5 | 10 | 0 | 8 | 2 | 10 | 2 | 0.968374303 | 0.890955434 | 1.356619906 | 1.248161865 |
| P28659 | Celf1 | CUGBP Elav-like family member 1 | 160.88 | 6.58 | 52107 | 3 | 865261.88 | 734913.44 | 745036.25 | 872370.88 | 600001.44 | 906230.38 | 581172.19 | 663452.06 | 734091.81 | 627396.31 | 1389668 | 939876.62 | 979234.5 | 899674.75 | 872242.56 | 880577.31 | 1094080.62 | 772794.38 | 978676.38 | 937467.69 | 1 | 2 | 2 | 2 | 2 | 2 | 1.330867003 | 1.446526376 | 1.221609404 | 1.327773715 |
| Q6NVF9 | Cpsf6 | Cleavage and polyadenylation specificity factor subunit 6 | 447.12 | 17.6 | 59153 | 7 | 7367215.5 | 8446803 | 7471223 | 8605520 | 6678760 | 9573927 | 8241998 | 7200472 | 6767321 | 8148053 | 8961622 | 7368817 | 7148781 | 8637070 | 8131952 | 6151530 | 6487039.5 | 6422705 | 7456730.5 | 4155834.5 | 9 | 2 | 12 | 2 | 12 | 2 | 1.043524535 | 1.007925293 | 0.795287012 | 0.768156251 |
| Q99LI7 | Cstf3 | Cleavage stimulation factor subunit 3 | 281.31 | 6.97 | 82877 | 4 | 1062641.5 | 746236.88 | 865839.5 | 843042.12 | 530829.88 | 973580.75 | 1103093 | 337176.41 | 961206 | 749848.25 | 1197545.38 | 969946.12 | 1115815.12 | 2 | 304738.56 | 1326596 | 2046384.5 | 1436053.25 | 1102324.5 | 1099629 | 11 | 0 | 12 | 2 | 12 | 2 | 0.886246147 | 0.869849777 | 1.731710906 | 1.69967266 |
| A6H8H2 | Dennd4c | DENN domain-containing protein 4C | 179.86 | 2.31 | 211458 | 4 | 1806835.75 | 1408024.75 | 1728051.38 | 1760529.25 | 1958200.5 | 1782875.88 | 1582826.62 | 1751823.5 | 2685984.75 | 2315327.25 | 2 | 390661 | 819668 | 2 | 804231.69 | 1718979.25 | 1521341.25 | 1183484.5 | 672851.38 | 1322616.5 | 0 | 2 | 0 | 1 | 0 | 2 | 0.232584627 | 0.199090517 | 0.74111504 | 0.634388344 |
| Q99J47 | Dhrs7b | Dehydrogenase/reductase SDR family member 7B | 311.79 | 19.81 | 34987 | 5 | 2487142.25 | 2193360.5 | 2090305.75 | 2643748 | 1884103.62 | 2774615 | 2580111.5 | 1696947.75 | 2079898.75 | 2242783.75 | 3696339.75 | 3548176.5 | 3308227.25 | 3226782 | 3673769 | 2843676.75 | 2885475 | 2441491.5 | 3737674.25 | 2831290.25 | 0 | 2 | 0 | 2 | 0 | 2 | 1.544722499 | 1.534442332 | 1.304544751 | 1.295862973 |
| Q9JHU4 | Dync1h1 | Cytoplasmic dynein 1 heavy chain 1 | 6142.521 | 27.07 | 532045 | 108 | 145004402 | 139568482 | 146092610 | 143011314 | 134834354 | 119510082 | 135156370 | 142668082 | 141465986 | 140054114 | 97889538 | 117178562 | 104135938 | 77970714 | 96465922 | 132771786 | 114973050 | 121877570 | 114409282 | 117620170 | 0 | 0 | 0 | 2 | 0 | 2 | 0.696729565 | 0.72716698 | 0.84917767 | 0.886274952 |
| Q8BH95 | Echs1 | Enoyl-CoA hydratase, mitochondrial | 468.06 | 35.86 | 31474 | 9 | 9979307 | 6286705 | 5104973 | 6703169 | 6592625.5 | 11275065 | 7426836 | 7789570 | 6736066 | 7310867 | 14751167 | 9621268 | 11057813 | 8953052 | 15725794 | 9080945 | 15836884 | 11572606 | 12389282 | 9650616 | 2 | 2 | 3 | 2 | 3 | 2 | 1.73391053 | 1.482769129 | 1.688369495 | 1.443824305 |
| Q8C0D5 | Efl1 | Elongation factor-like GTPase 1 | 204.8 | 3.55 | 125777 | 4 | 2273220 | 2338838.5 | 2245106 | 2594092.5 | 2189590.25 | 2409480.5 | 2201101.75 | 2588308.25 | 2680155.5 | 2408180.75 | 1922699.75 | 1855840.25 | 1744362 | 11608290 | 2277296.25 | 2436049.75 | 3479652 | 3438606.5 | 3432538.5 | 9249854 | 8 | 1 | 6 | 2 | 8 | 2 | 1.667274541 | 1.579566215 | 1.893049559 | 1.793464156 |
| O55135 | Eif6 | Eukaryotic translation initiation factor 6 | 345.48 | 26.94 | 26511 | 4 | 18018272 | 18007302 | 14240986 | 16138637 | 14202554 | 18133672 | 17437594 | 15082950 | 19283738 | 15859921 | 18379562 | 18082514 | 17013190 | 17601362 | 17987540 | 18055366 | 22712560 | 19711514 | 20161832 | 20063878 | 6 | 0 | 10 | 2 | 10 | 2 | 1.104908236 | 1.038069626 | 1.249323406 | 1.173748767 |
| Q8BPU7 | Elmo1 | Engulfment and cell motility protein 1 | 455.51 | 12.65 | 83936 | 9 | 10474585 | 11417404 | 12314479 | 12679575 | 11089144 | 9561596 | 10135867 | 11454115 | 11787528 | 11515365 | 5902627 | 8792404 | 5802504 | 2810125.75 | 8685426 | 10600521 | 8862784 | 8724269 | 8454968 | 6767100 | 0 | 1 | 0 | 2 | 0 | 2 | 0.551841027 | 0.587519926 | 0.748762432 | 0.797173147 |
| Q8CGC7 | Eprs | Bifunctional glutamate/proline--tRNA ligase | 2976.59 | 37.43 | 170079 | 47 | 69261714 | 79808250 | 73123498 | 78397066 | 72556914 | 75767354 | 73700258 | 73577666 | 83873530 | 83770346 | 154616690 | 150332530 | 132186002 | 144640738 | 126726594 | 79761218 | 94751962 | 91702866 | 86288674 | 89401194 | 0 | 1 | 0 | 2 | 0 | 2 | 1.898720115 | 1.813468705 | 1.184266229 | 1.131093376 |
| Q9EQH2 | Erap1 | Endoplasmic reticulum aminopeptidase 1 | 1195.16 | 27.85 | 106599 | 23 | 15928867 | 17380890 | 20719174 | 21131884 | 21456402 | 16762797 | 19238832 | 20854920 | 23770178 | 20499010 | 17647346 | 20107688 | 28858864 | 26812716 | 24815430 | 15239297 | 12707638 | 16510649 | 12291273 | 16813998 | 6 | 2 | 7 | 1 | 7 | 2 | 1.223819601 | 1.169257674 | 0.761384537 | 0.727439494 |
| P04117 | Fabp4 | Fatty acid-binding protein, adipocyte | 392.87 | 53.79 | 14650 | 6 | 3712428.25 | 2877863 | 2 | 2046855 | 1112137.38 | 5807552 | 1698319.62 | 2265547.5 | 3326754 | 4107701.75 | 4041946.75 | 1927676.75 | 4303404.5 | 3237206.75 | 3981523 | 4768674.5 | 7796943.5 | 5605300 | 6866908.5 | 6254466.5 | 4 | 0 | 12 | 2 | 12 | 2 | 1.794157892 | 1.016615422 | 3.209701119 | 1.818698162 |
| P05201 | Got1 | Aspartate aminotransferase, cytoplasmic | 1401.39 | 65.86 | 46248 | 23 | 113522818 | 117045410 | 126015354 | 141350386 | 134842066 | 109237498 | 105238154 | 119877010 | 116997346 | 125425698 | 70286122 | 76833890 | 76493218 | 81340842 | 81645114 | 131531642 | 147978738 | 141992338 | 148730754 | 151981010 | 0 | 2 | 0 | 0 | 0 | 2 | 0.610957377 | 0.670276473 | 1.141342976 | 1.25215829 |
| Q3TRM8 | Hk3 | Hexokinase-3 | 2205.37 | 45.34 | 100101 | 31 | 69005834 | 73097810 | 49343098 | 67001110 | 70139610 | 74696466 | 67914618 | 64077450 | 65527166 | 64659506 | 67456930 | 63003658 | 66717742 | 68092418 | 59135202 | 77777242 | 78632770 | 72473090 | 72917066 | 80681922 | 7 | 2 | 10 | 2 | 10 | 2 | 0.98727428 | 0.962985534 | 1.164019125 | 1.135382133 |
| Q61316 | Hspa4 | Heat shock 70 kDa protein 4 | 2016.81 | 42.33 | 94133 | 33 | 17570934 | 15673127 | 16665926 | 17431208 | 15477390 | 16731241 | 16995972 | 16069450 | 14895732 | 15951484 | 7164988 | 16827150 | 14625352 | 15983943 | 14966002 | 22349546 | 27430482 | 20621572 | 23909980 | 17387614 | 5 | 2 | 7 | 0 | 7 | 2 | 0.83999787 | 0.86264991 | 1.348721353 | 1.385092029 |
| Q66HG8 | Ik | Protein Red | 95.98 | 5.57 | 65588 | 2 | 2125162.75 | 2231188.5 | 2464802.25 | 1728169.88 | 2650804.5 | 1292077.75 | 2794753.5 | 1277337.38 | 2181089.5 | 1979706 | 401953.78 | 1063687.5 | 896554.88 | 2 | 542224.12 | 1898005.5 | 764302.25 | 844316.06 | 491318.97 | 985567.5 | 0 | 1 | 0 | 2 | 0 | 2 | 0.259320457 | 0.304927372 | 0.444951195 | 0.523205149 |
| Q9JKF1 | Iqgap1 | Ras GTPase-activating-like protein IQGAP1 | 6644.04 | 60.35 | 188742 | 102 | 374904002 | 375309954 | 392626946 | 363138082 | 411415170 | 331715074 | 376954690 | 376935202 | 397851842 | 388952322 | 282574434 | 360428866 | 378815394 | 322801570 | 364470850 | 324172866 | 301952354 | 320598626 | 346848162 | 342613826 | 4 | 0 | 5 | 2 | 5 | 2 | 0.891361388 | 0.912776533 | 0.853338282 | 0.873839914 |
| O89051 | Itm2b | Integral membrane protein 2B | 360.12 | 31.2 | 30260 | 6 | 2361586.5 | 1945130.25 | 2188844.25 | 1419264 | 4057121.75 | 1740583.75 | 2751589.75 | 2622307 | 5049737 | 1918818 | 3220586.25 | 4862778.5 | 5273106 | 4319535.5 | 4223757.5 | 1768867.75 | 2 | 1095932 | 1665588.25 | 1271955.12 | 1 | 2 | 4 | 1 | 4 | 2 | 1.829256695 | 1.555045697 | 0.484661788 | 0.41200955 |
| O88447 | Klc1 | Kinesin light chain 1 | 139.11 | 5.18 | 61450 | 3 | 1234729 | 1329136.75 | 1832212.75 | 1471942.75 | 2459430.75 | 1552733.25 | 1545023.12 | 1926138.25 | 2372504 | 1808694.25 | 869947.56 | 514727.22 | 1023695.62 | 1434415.62 | 1109551.75 | 2773553.25 | 2654447 | 1926038.25 | 2834221.75 | 3933038 | 2 | 1 | 0 | 2 | 2 | 2 | 0.594700248 | 0.537999732 | 1.695752584 | 1.534074418 |
| Q9EP89 | Lactb | Serine beta-lactamase-like protein LACTB, mitochondrial | 189.55 | 5.99 | 60705 | 3 | 1807807.38 | 1984280.25 | 1597271.25 | 1171047.75 | 2270608 | 1530411.38 | 1861333.88 | 1994968.75 | 1863822.88 | 1675037.75 | 395289.97 | 213890.73 | 2 | 143068.91 | 861726.12 | 1599788.88 | 2 | 921395.12 | 2 | 128094.3 | 0 | 2 | 0 | 1 | 0 | 2 | 0.182762434 | 0.180826198 | 0.299997499 | 0.296819242 |
| Q3KNM2 | March5 | E3 ubiquitin-protein ligase MARCH5 | 207.26 | 17.99 | 31232 | 4 | 1078629.88 | 1003967.44 | 970215 | 593966.62 | 1146369.88 | 1078736.12 | 838399.5 | 714066.56 | 1316340.12 | 1289057.12 | 582306.69 | 884668.62 | 1300189.75 | 1467569.5 | 1452474.38 | 741624.31 | 2 | 507526.31 | 335326.78 | 644340.19 | 9 | 2 | 9 | 1 | 9 | 2 | 1.186528763 | 1.086050027 | 0.465001124 | 0.425623465 |
| Q9CPR5 | Mrpl15 | 39S ribosomal protein L15, mitochondrial | 283.96 | 23.39 | 33542 | 5 | 1927839.88 | 3177078.75 | 2198457.75 | 1772618.12 | 2217582.5 | 2200819.75 | 2327372.25 | 1148367 | 1964178.25 | 1463587.12 | 1784919.12 | 2530076.25 | 2320614.5 | 1836002.88 | 1576287.38 | 1685108 | 954373.69 | 1133273 | 111953.42 | 543764.06 | 10 | 0 | 10 | 2 | 10 | 2 | 0.889700414 | 1.103640393 | 0.392123078 | 0.486414147 |
| Q7TPV4 | Mybbp1a | Myb-binding protein 1A | 1961.03 | 29.39 | 152037 | 30 | 16669950 | 16689174 | 10543772 | 15530982 | 18256520 | 17692290 | 16958520 | 15515895 | 15362147 | 17942284 | 15470809 | 19055898 | 24014454 | 18311890 | 22762764 | 22897156 | 19462508 | 17410934 | 19226136 | 20861750 | 4 | 1 | 4 | 2 | 4 | 2 | 1.282215275 | 1.193416309 | 1.285338814 | 1.196323529 |
| Q8VDK1 | Nit1 | Deaminated glutathione amidase | 92.23 | 5.57 | 35705 | 2 | 1453100.62 | 1793978.12 | 1165273.75 | 608631.81 | 1501805.5 | 593284.62 | 1475366.25 | 1434075.88 | 1432678 | 1180252 | 729199.69 | 1360792.5 | 450124.47 | 373706.66 | 862305.25 | 654782.75 | 560388.44 | 584645 | 190088.25 | 715878.5 | 4 | 2 | 4 | 2 | 4 | 2 | 0.578913116 | 0.617452667 | 0.414819889 | 0.442435384 |
| Q6P9R2 | Oxsr1 | Serine/threonine-protein kinase OSR1 | 84.35 | 3.8 | 58214 | 2 | 739846.31 | 899719.56 | 864014.62 | 503643.66 | 1333672.75 | 1056420.25 | 824048.88 | 157491.7 | 688381.56 | 501616.5 | 2539005.25 | 1556091.62 | 799448.56 | 571733.75 | 1090789.25 | 378975.88 | 2 | 2 | 128985.12 | 346824.44 | 8 | 0 | 5 | 2 | 8 | 2 | 1.510533095 | 2.031335793 | 0.196915398 | 0.264808032 |
| P60335 | Pcbp1 | Poly(rC)-binding protein 1 | 945.39 | 54.49 | 37498 | 13 | 45694862 | 57047054 | 42060306 | 28009404 | 47301782 | 39952702 | 51177298 | 40361566 | 47817650 | 26143302 | 63648918 | 52773410 | 31367120 | 34753930 | 42094018 | 34346298 | 27052316 | 18823846 | 21236138 | 23164248 | 12 | 1 | 10 | 2 | 12 | 2 | 1.020552987 | 1.093378646 | 0.566175623 | 0.60657736 |
| Q9CWM4 | Pfdn1 | Prefoldin subunit 1 | 102.38 | 14.75 | 14255 | 2 | 1026337.62 | 874228.75 | 769224.44 | 1162761 | 620966.75 | 1017767.12 | 885211.31 | 689704.31 | 733995.19 | 995674.88 | 929976.5 | 694433.06 | 854454.12 | 875374.06 | 1007660.94 | 1069790.88 | 1340700.5 | 1346995.75 | 1308532.25 | 1048482.75 | 12 | 2 | 12 | 0 | 12 | 2 | 0.979427529 | 1.009149154 | 1.37295984 | 1.414623562 |
| P67778 | Phb | Prohibitin | 1752.75 | 80.88 | 29820 | 25 | 215902386 | 216467602 | 194506034 | 206260498 | 199560674 | 227240322 | 230726114 | 205789042 | 213743362 | 234699170 | 300793474 | 278884802 | 369353890 | 357096834 | 342199362 | 234440322 | 267370482 | 246789874 | 258797250 | 232440418 | 0 | 0 | 0 | 2 | 0 | 2 | 1.596139092 | 1.482045775 | 1.200582662 | 1.114764039 |
| P35433 | Ppat | Amidophosphoribosyltransferase | 44.22 | 1.35 | 57437 | 1 | 2 | 1858454.25 | 1726975.5 | 2424481.5 | 1305613 | 2 | 1752391.62 | 1573035 | 1561889.62 | 1816592.12 | 1481712.25 | 1768407 | 1514191.88 | 1500761.75 | 1937698 | 2650655.75 | 2784098.5 | 2121447.75 | 2428061 | 2280356.25 | 12 | 2 | 12 | 0 | 12 | 2 | 1.121282407 | 1.223580036 | 1.676519068 | 1.829472441 |
| O89044 | Prim2 | DNA primase large subunit | 143.89 | 4.93 | 58603 | 2 | 551853.31 | 970609.88 | 809258.44 | 476590.53 | 1075688.5 | 557341.12 | 669469.12 | 592665.12 | 1209464.25 | 368183.78 | 270478.09 | 842869 | 2 | 143357.5 | 263835.72 | 557292.19 | 2 | 387330.41 | 67135.01 | 342822.53 | 3 | 2 | 4 | 2 | 4 | 2 | 0.391488685 | 0.447597021 | 0.348759503 | 0.398743874 |
| Q8BIW1 | Prune1 | Exopolyphosphatase PRUNE1 | 199.37 | 11.01 | 50239 | 4 | 518169.72 | 204134.19 | 297003.25 | 432005.25 | 290897.91 | 491538.25 | 241802.34 | 148216.84 | 136023.17 | 410617.19 | 365946.72 | 178574.67 | 218985.41 | 361571.25 | 99068.01 | 433242.38 | 752564.25 | 684863.25 | 521839.56 | 453871 | 7 | 2 | 10 | 2 | 10 | 2 | 0.702639656 | 0.857126421 | 1.633775445 | 1.992987568 |
| Q60692 | Psmb6 | Proteasome subunit beta type-6 | 171.85 | 12.61 | 25379 | 3 | 4689926 | 4987213 | 5375022 | 3604144 | 3616380 | 4484542 | 5613446.5 | 3007370.25 | 4166580.5 | 5315704 | 5705619 | 4755651 | 2875437.5 | 5380860 | 2969666.5 | 6018036 | 5338362 | 5935906 | 6972756.5 | 5346987.5 | 12 | 2 | 12 | 2 | 12 | 2 | 0.973714395 | 0.960137087 | 1.329523046 | 1.310984403 |
| O35593 | Psmd14 | 26S proteasome non-ATPase regulatory subunit 14 | 395.13 | 38.06 | 34577 | 6 | 8660083 | 9181106 | 7990924 | 10538366 | 8660145 | 8050112.5 | 9277148 | 8632354 | 7554669.5 | 8781094 | 12913041 | 11234559 | 11654591 | 13505668 | 11727562 | 9221145 | 10776824 | 10833144 | 11693914 | 9840708 | 0 | 2 | 0 | 1 | 0 | 2 | 1.35542028 | 1.443075435 | 1.162891613 | 1.238095921 |
| P53534 | Pygb | Glycogen phosphorylase, brain form | 871.84 | 19.09 | 96174 | 15 | 18244850 | 17799186 | 18200130 | 16737296 | 23066866 | 13948239 | 17028260 | 22343636 | 22001712 | 16215926 | 7046817.5 | 8231821 | 8822479 | 5368501 | 8261356 | 16399435 | 11798240 | 11463429 | 10741095 | 10378699 | 0 | 0 | 0 | 2 | 0 | 2 | 0.401187084 | 0.412190217 | 0.646273031 | 0.663997998 |
| Q99JI6 | Rap1b | Ras-related protein Rap-1b | 1218.61 | 83.15 | 20825 | 16 | 13629994 | 12422928 | 10544779 | 13016406 | 10829974 | 14608352 | 12932520 | 9714775 | 12313395 | 11269606 | 14003465 | 17803998 | 15466155 | 9798287 | 14401966 | 11017321 | 8555235 | 7833452.5 | 6050888 | 7311885 | 5 | 2 | 6 | 1 | 6 | 2 | 1.182479241 | 1.174810311 | 0.674487573 | 0.670113207 |
| Q9D0I9 | Rars | Arginine--tRNA ligase, cytoplasmic | 1695.8 | 44.85 | 75674 | 29 | 28191718 | 28314748 | 20498282 | 24726384 | 26574872 | 27343870 | 25145194 | 22467102 | 27191068 | 22871632 | 40809410 | 37054918 | 39739254 | 24907196 | 30945126 | 37023322 | 36738834 | 28191300 | 31265586 | 28915310 | 3 | 2 | 3 | 0 | 3 | 2 | 1.35189234 | 1.387437829 | 1.263653663 | 1.296879081 |
| B2RY56 | Rbm25 | RNA-binding protein 25 | 214.37 | 6.44 | 99552 | 5 | 6265755 | 7423930 | 6437427 | 4158522.75 | 6497223 | 862286.56 | 2727224.25 | 3084281 | 3615152.75 | 2623053.5 | 361820.56 | 950526.62 | 1198316.25 | 382179.44 | 1961176.25 | 852266 | 542825.19 | 887085.88 | 935398.06 | 648393.62 | 0 | 0 | 3 | 2 | 3 | 2 | 0.157685786 | 0.375930905 | 0.125588364 | 0.299409025 |
| Q9QYF1 | Rdh11 | Retinol dehydrogenase 11 | 470.53 | 26.9 | 35148 | 7 | 17384592 | 19125722 | 15907852 | 19009422 | 15508748 | 19923836 | 17336732 | 17727470 | 14947081 | 15625086 | 22761130 | 22865766 | 29008802 | 28993562 | 26978962 | 18373678 | 20837184 | 19972518 | 21413542 | 21018688 | 0 | 2 | 0 | 1 | 0 | 2 | 1.502343301 | 1.526506651 | 1.168850847 | 1.18765038 |
| P63325 | Rps10 | 40S ribosomal protein S10 | 348.69 | 24.24 | 18916 | 5 | 7829103 | 8679740 | 7061984 | 7032239 | 3822406.25 | 9502366 | 9222597 | 6733453.5 | 2889025 | 7618783 | 13789864 | 10730584 | 5890281 | 7654103 | 6812145 | 10777984 | 13390899 | 10459331 | 10745734 | 8948775 | 10 | 0 | 9 | 2 | 10 | 2 | 1.303598007 | 1.247753347 | 1.577980473 | 1.51038158 |
| P14131 | Rps16 | 40S ribosomal protein S16 | 769.18 | 64.38 | 16445 | 14 | 87129538 | 78053866 | 85512258 | 84388994 | 86635434 | 85250138 | 98477426 | 105833554 | 90691770 | 86924906 | 170714802 | 151698722 | 135426146 | 140187378 | 145977714 | 105846314 | 128481410 | 108636194 | 109695826 | 97306370 | 0 | 0 | 0 | 2 | 0 | 2 | 1.764214652 | 1.59255164 | 1.30410224 | 1.17720945 |
| Q63623 | Scaf8 | Protein SCAF8 | 75.3 | 0.95 | 139504 | 1 | 990841.06 | 982865.5 | 1003612.62 | 945580.38 | 1120100.5 | 869558.06 | 927541.56 | 920176.62 | 1256072.25 | 828604.81 | 667548.56 | 621888.12 | 784115.81 | 2 | 800262.06 | 894869.69 | 498700.78 | 762440.12 | 632900.06 | 618342.56 | 0 | 0 | 0 | 2 | 0 | 2 | 0.569862486 | 0.598468242 | 0.675640129 | 0.709555674 |
| Q923V8 | Selenof | Selenoprotein F | 144.31 | 17.28 | 17822 | 2 | 491594.06 | 532515.38 | 669065 | 845846.44 | 634432.25 | 902992.62 | 567793.69 | 851722.31 | 471901.03 | 875641.88 | 532241.44 | 445070.03 | 2 | 1589918.75 | 632996.5 | 392883.25 | 370041.88 | 519696.31 | 531933.31 | 371564.88 | 8 | 2 | 8 | 2 | 8 | 2 | 1.008437367 | 0.871984683 | 0.688877239 | 0.59566456 |
| Q6PEC4 | Skp1 | S-phase kinase-associated protein 1 | 357.08 | 30.06 | 18672 | 5 | 7714227 | 8018556 | 9415634 | 9755121 | 8676100 | 7041177.5 | 8738194 | 8569774 | 9685045 | 8540013 | 6860478 | 6549053 | 7013263.5 | 20096202 | 5289640.5 | 9893174 | 10753308 | 8727770 | 17146248 | 16563612 | 5 | 2 | 5 | 2 | 5 | 2 | 1.051147717 | 1.075971674 | 1.44755934 | 1.481744973 |
| P53986 | Slc16a1 | Monocarboxylate transporter 1 | 308.93 | 11.56 | 53267 | 5 | 1753786.12 | 1571419.25 | 1593456.25 | 1967701 | 1440187.62 | 1833914.38 | 1861837.38 | 2252977 | 1714045.88 | 2368603 | 5249233 | 3753128.75 | 4730612.5 | 5150874 | 4936248.5 | 2235543.5 | 2474423.25 | 2531022.5 | 2921630.25 | 2711864.25 | 0 | 0 | 0 | 2 | 0 | 2 | 2.86074017 | 2.37455887 | 1.54619661 | 1.283421302 |
| B1ARD6 | Slfn9 | Schlafen family member 9 | 151.73 | 3.85 | 103992 | 3 | 2271834.5 | 2426148.75 | 2726213.25 | 1914708.12 | 2006276 | 1143569.38 | 1881881.62 | 1823455.75 | 1888983.5 | 1557205.25 | 1691341.5 | 1848053.25 | 1163455.25 | 878350 | 1502683.38 | 1810381.62 | 642126.88 | 1114413.25 | 942001.19 | 1114537.5 | 0 | 0 | 7 | 2 | 7 | 2 | 0.624395822 | 0.853984548 | 0.495669538 | 0.677925943 |
| P62075 | Timm13 | Mitochondrial import inner membrane translocase subunit Tim13 | 91.99 | 14.74 | 10458 | 1 | 825765.62 | 931080.81 | 760500.81 | 718933.75 | 599405.88 | 1357459.12 | 689098.38 | 2 | 528331.56 | 899521.5 | 632144.56 | 1021881.5 | 904212.31 | 770224.25 | 669200 | 1499492.38 | 1122269.62 | 1370500 | 1617780.88 | 1237384.38 | 11 | 0 | 10 | 2 | 11 | 2 | 1.042228617 | 1.150601016 | 1.785189326 | 1.970815826 |
| Q8BH24 | Tm9sf4 | Transmembrane 9 superfamily member 4 | 188.46 | 8.4 | 74693 | 4 | 2 | 2 | 334859.97 | 451891.38 | 452252.06 | 688459.69 | 650141.75 | 676567.81 | 974322.56 | 1097224.12 | 6443011 | 1697644.38 | 5361137 | 5718504 | 2963633.75 | 938133.12 | 2243355.5 | 1233456.62 | 2210930.5 | 1528219.38 | 0 | 0 | 0 | 2 | 0 | 2 | 17.90459843 | 5.428302458 | 6.581151214 | 1.995268391 |
| Q9JMH6 | Txnrd1 | Thioredoxin reductase 1, cytoplasmic | 1007.85 | 35.73 | 67084 | 14 | 19807386 | 24552198 | 22245692 | 23546662 | 22307386 | 24420478 | 23366882 | 22487040 | 24123004 | 25967258 | 37900402 | 37285750 | 35741662 | 40957738 | 37186502 | 24680212 | 24894844 | 32046844 | 31508106 | 33738122 | 0 | 0 | 0 | 2 | 0 | 2 | 1.68124836 | 1.570826943 | 1.305966662 | 1.220193083 |
| Q09426 | Ugt8 | 2-hydroxyacylsphingosine 1-beta-galactosyltransferase | 42.84 | 1.29 | 61126 | 1 | 5231538.5 | 6088538.5 | 7125840.5 | 8207048.5 | 6179887.5 | 6730344 | 4833056.5 | 5598135 | 4795786 | 8426761 | 9113146 | 8816571 | 9385500 | 10784755 | 9414617 | 7063549 | 9171870 | 8665436 | 8791171 | 9978919 | 0 | 2 | 0 | 1 | 0 | 2 | 1.447165992 | 1.563798709 | 1.330098981 | 1.437296815 |
| Q9WV55 | Vapa | Vesicle-associated membrane protein-associated protein A | 640.84 | 61.85 | 27855 | 13 | 26413584 | 31079540 | 23345656 | 24419186 | 23578750 | 29828764 | 30623798 | 22647174 | 21246396 | 23947742 | 22142514 | 27667172 | 25932932 | 27838368 | 28562550 | 33945938 | 31952902 | 30099460 | 30510236 | 35705502 | 10 | 2 | 12 | 2 | 12 | 2 | 1.025666752 | 1.030006593 | 1.259066849 | 1.264394261 |
| Q9CZT5 | Vasn | Vasorin | 54.58 | 1.04 | 72261 | 1 | 1015842.25 | 1813874.88 | 637234.94 | 751756.94 | 507963.62 | 1474410.5 | 1170301.62 | 982576.12 | 1059798.62 | 1505575.5 | 13321597 | 10826836 | 10036818 | 9256282 | 7948387 | 1646626 | 1686999.25 | 2187867.25 | 2246833.25 | 1851863.12 | 0 | 2 | 0 | 0 | 0 | 2 | 10.87232479 | 8.29851799 | 2.035298321 | 1.553481897 |
| P68254 | Ywhaq | 14-3-3 protein theta | 1277.78 | 62.86 | 27778 | 20 | 30573510 | 38028930 | 31267838 | 29141440 | 38228498 | 24833428 | 31985984 | 34165622 | 50971466 | 25445520 | 17479840 | 20284504 | 15816199 | 13867849 | 15501930 | 26600342 | 17400452 | 20782536 | 19886744 | 22642332 | 0 | 0 | 0 | 2 | 0 | 2 | 0.495995066 | 0.495515657 | 0.641666272 | 0.641046064 |
| P10404 | | MLV-related proviral Env polyprotein | 181.57 | 3.43 | 69613 | 3 | 1736666.38 | 2479842 | 2844977.5 | 2408473.5 | 3305649.5 | 1980012.12 | 2119979.5 | 2445384 | 2694892 | 2655380 | 2447336.5 | 3195508.25 | 3503414 | 1943056 | 3789582 | 1875327.75 | 1445945.5 | 2133931.75 | 1683757 | 1725597.25 | 8 | 2 | 7 | 2 | 8 | 2 | 1.164633082 | 1.250784928 | 0.693865892 | 0.745193497 |
| P63168 | Dynll1 | Dynein light chain 1, cytoplasmic | 95.03 | 37.08 | 10366 | 2 | 846282.88 | 778651.75 | 657885.31 | 355434.91 | 484499.16 | 305563.5 | 1051397.38 | 344684.25 | 984296.94 | 2 | 2 | 842535.69 | 2 | 2 | 2 | 2 | 2 | 2 | 2 | 2 | 4 | 0 | 5 | 2.5 | 5 | 2.5 | 0.269807896 | 0.313686238 | 3.2023E-06 | 3.72309E-06 |

Supplementary Table 4: Results of the pathway analysis by the David tool on the proteins modulated in response to PR101

| | | | | | | |
|---|---|---|---|---|---|---|
| Annotation Cluster 1 | Enrichment Score: 16.74774209569559 | | | | | |
| Category | Term | Count | PValue | Genes | Fold Enrichment | FDR |
| GOTERM_BP_DIRECT | GO:0002181~cytoplasmic translation | 42 | 2.8E-33 | P62918, P35979, P99027, Q9D7S7, P62717, P62281, P12970, A2AGT5, P63323, P35980, P62855, P26638, P25444, P47955, P62852, P62830, P62754, P51410, P62849, P32233, P14206, P63276, O55142, P14131, P61255, Q9D8E6, Q3TIV5, P62702, P62900, P62889, P27659, P41105 | 12.19847507 | 1.1E-29 |
| GOTERM_CC_DIRECT | GO:0045202~synapse | 88 | 3.5E-27 | P62918, Q07076, P35979, P62717, Q8CGY8, Q99L45, P11688, P62281, P12970, P11881, P63001, O08553, O55131, P63323, Q80UU9, P35980, P63328, Q8QZY1, P61982, P25444, P47955, P62830, P28798, Q8BH43, Q9DBC7, O08917, P62849, P29341, P14206, P35922, P57776, O35678, P35293, P67778, O55142, P14131, P63213, Q9JLQ2, P62482, P47941, P62889, P01029, Q9CX86, P70280, P26883, O70435, Q9Z0N1, P99027, Q9ERS2, P10852, P70206, P53986, P68510, P62855, Q921J2, P62852, Q8C052, P62259, P51410, Q9CWZ7, Q9JMA1, Q62393, O08539, P68254, P14106, P35585, P12367, Q99J99, P63276, P58252, P61255, P62702, P62900, Q9D0E1, P27659, O88342, P62743, P41105 | 3.863763406 | 3.9E-25 |
| GOTERM_CC_DIRECT | GO:0005840~ribosome | 46 | 8E-27 | P62918, P11928, P35979, P99027, Q9D7S7, P62717, P62281, P12970, P63323, P35980, P62855, Q9CQF0, P25444, P47955, P62852, P62830, P62754, Q9CY73, P51410, P62849, P35564, P10126, P14206, P63276, O55142, P14131, P61255, P58252, Q9D8E6, P62702, Q9QUM9, P62900, P62889, P27659, Q6P542, P41105 | 7.736819172 | 7.7E-25 |
| GOTERM_CC_DIRECT | GO:1990904~ribonucleoprotein complex | 57 | 1.3E-25 | Q9CX86, Q9EQK5, P35979, Q9D7S7, P62717, Q501J6, P62281, P32067, Q8CGC7, P63323, P35980, Q99MR6, P28659, Q9CQF0, P25444, P62852, P62830, P62754, Q61686, Q9CY73, P51410, Q3UYV9, P29341, Q9WTM5, Q60591, P35922, Q01320, Q3UEB3, Q91VC3, Q05CL8, P63276, O55142, P14131, P60122, P58252, P61255, P57780, Q9D8E6, P62702, P62900, P62889, Q9D0E1, Q6Q899, P27659, P63017, P41105, Q8BVY0 | 5.609688581 | 1.1E-23 |
| GOTERM_CC_DIRECT | GO:0098793~presynapse | 54 | 6.4E-22 | P62918, P35979, P99027, P56399, O08529, P62281, P12970, P11881, P40124, O08553, P63323, P68037, Q6PIU9, P68510, O08992, P63087, O88712, P61028, P62855, P22892, P05132, P61982, P62830, P62754, P51410, Q8CI51, O08917, P62849, O08539, P35564, P61082, P35922, O35678, O55142, P14131, O54774, P61255, Q9JLQ2, Q9D8E6, Q920Q4, P62962, P63017, P62743, P41105 | 5.004561825 | 4.3E-20 |
| GOTERM_BP_DIRECT | GO:0140242~translation at postsynapse | 26 | 3.3E-21 | P62918, P35979, P51410, P99027, P62849, P62281, P12970, O55142, P14131, P63323, P61255, P58252, Q9D8E6, P62855, P62830, P41105 | 12.77935484 | 6.4E-18 |

| | | | | | | |
|---|---|---|---|---|---|---|
| UP_KW_MOLECULAR_FUNCTION | KW-0687~Ribonucleoprotein | 63 | 8.9E-21 | P62918, Q9EQK5, P35979, Q9D7S7, P62717, P62281, P12970, P63323, P62311, P35980, Q6DFW4, P62317, Q9D1N9, P25444, P47955, P62830, P62754, Q9CY73, P62849, Q91WM3, P14206, Q9JIK9, Q9D7A6, P35922, P62320, O55142, P14131, P62322, P62889, Q9CX86, P99027, P57784, P14576, Q9DB15, Q6ZWM4, P62855, Q9CQF0, P62852, P51410, Q9D6Z1, Q3UEB3, Q9ESX5, P63276, P16254, P61255, Q9D8E6, P62702, P62900, P62307, Q9D0E1, P27659, P41105 | 3.916743211 | 6.7E-19 |
| GOTERM_BP_DIRECT | GO:0140236~translation at presynapse | 25 | 4.1E-20 | P62918, P35979, P51410, P99027, P62849, P62281, P12970, O55142, P14131, P63323, P61255, Q9D8E6, P62855, P62830, P41105 | 12.47983871 | 5.4E-17 |
| GOTERM_MF_DIRECT | GO:0003735~structural constituent of ribosome | 42 | 4E-19 | P62918, P35979, P99027, Q9D7S7, P62717, P62281, P12970, Q9DB15, P63323, P35980, P62855, Q9D1N9, Q9CQF0, P25444, P47955, P62852, P62830, P62754, P51410, P62849, Q9JIK9, P14206, P63276, O55142, P14131, P61255, Q9D8E6, P62702, P62900, P62889, P27659, P41105 | 5.642920847 | 1.1E-16 |
| GOTERM_CC_DIRECT | GO:0032040~small-subunit processome | 27 | 1.3E-18 | Q8K224, P62849, Q91WM3, Q8BHY2, Q9D6Z1, O35130, P62281, P63276, Q8BU03, P14131, P63323, Q8C7V3, Q6DFW4, P62702, Q6NS46, P62754, Q9D0D4 | 9.753208556 | 5.7E-17 |
| GOTERM_CC_DIRECT | GO:0098794~postsynapse | 53 | 6.8E-18 | P62918, P35979, P99027, Q99LD9, O08529, P62281, P12970, P11881, P63001, P40124, A2AGT5, P61161, P63323, P42669, P63087, P62196, P62855, P05132, P25444, P47955, P62830, P51410, P62849, O08539, P14106, P14206, P61082, P35922, P63276, Q80TH2, O55142, P14131, O54774, P61255, Q9D8E6, P62900, P62889, Q8BPU7, P62962, P63017, P62743, P41105, P60766 | 4.14969574 | 2.9E-16 |
| GOTERM_BP_DIRECT | GO:0006412~translation | 40 | 8.8E-18 | P62918, P35979, Q9Z0N1, Q9D7S7, P62717, O70251, P62281, Q9DB15, Q8CGC7, P63323, P35980, Q9CQF0, P25444, P62830, P62754, P51410, P62849, P10126, P14206, P63276, O55142, P14131, Q8R050, Q8BP47, P61255, Q9D8E6, P62702, P62900, P27659, P41105 | 5.484701648 | 8.6E-15 |
| GOTERM_BP_DIRECT | GO:0042274~ribosomal small subunit biogenesis | 26 | 2.7E-17 | Q8K224, P62849, Q91WM3, Q9D6Z1, O35130, P62281, P63276, Q8BU03, P14131, P63323, Q8C7V3, Q6DFW4, P62702, P62852, P62754, Q9D0D4 | 9.22953405 | 2.1E-14 |
| GOTERM_CC_DIRECT | GO:0022627~cytosolic small ribosomal subunit | 21 | 1.2E-16 | P63276, P14131, P63323, P62849, P62855, P62702, P62281, P14206, P25444, P62852, P62754 | 12.1373262 | 4.8E-15 |
| GOTERM_CC_DIRECT | GO:0022626~cytosolic ribosome | 29 | 2.8E-16 | P62918, P35979, P62717, P62281, E9PZJ8, P12970, P63323, P61222, P35980, P25444, P47955, P62852, P62830, P62754, P51410, P62849, P10126, P14206, P63276, O55142, P14131, Q8R050, P61255, Q9D8E6, P62702, P62900, P62889, P27659, P41105 | 7.258730894 | 1E-14 |
| UP_KW_MOLECULAR_FUNCTION | KW-0689~Ribosomal protein | 43 | 1.8E-15 | P62918, P35979, P99027, Q9D7S7, P62717, P62281, P12970, Q9DB15, P63323, P35980, P62855, Q9D1N9, Q9CQF0, P25444, P47955, P62852, P62830, P62754, Q9CY73, P51410, P62849, Q9JIK9, P14206, P63276, O55142, P14131, P61255, Q9D8E6, P62702, P62900, P62889, P27659, P41105 | 4.234982444 | 6.7E-14 |
| INTERPRO | IPR000530:Ribosomal_eS12 | 11 | 1.9E-13 | P63323 | 23.56719577 | 2.8E-10 |
| INTERPRO | IPR047860:Ribosomal_eS12_CS | 11 | 1.9E-13 | P63323 | 23.56719577 | 2.8E-10 |

| | | | | | | |
|---|---|---|---|---|---|---|
| UP_SEQ_FEATURE | DOMAIN:Ribosomal protein eL8/eL30/eS12/Gadd45 | 13 | 3.6E-11 | P63323, P62889, P12970 | 13.20230924 | 2.5E-08 |
| INTERPRO | IPR004038:Ribosomal_eL8/eL30/eS12/Gad45 | 13 | 3E-10 | P63323, P62889, P12970 | 11.34716833 | 3E-07 |
| INTERPRO | IPR029064:Ribosomal_eL30-like_sf | 13 | 2.1E-09 | P63323, P62889, P12970 | 9.88301758 | 1.5E-06 |
| KEGG_PATHWAY | mmu05171:Coronavirus disease - COVID-19 | 52 | 2.6E-09 | P62918, P11928, P35979, Q9D7S7, P99027, P62717, P62281, P12970, P63323, Q9QUN7, P35980, P58681, Q9Z0F8, P62855, P25444, P47955, P62852, P62830, P62754, P47811, Q8VI93, P51410, P97333, P62849, P14106, P14206, P30993, P63276, O55142, P14131, E9Q9A9, P61255, Q61093, Q9D8E6, P62702, P62900, P62889, P01029, Q6Q899, P27659, P01027, P41105 | 2.461424259 | 1.1E-07 |
| GOTERM_CC_DIRECT | GO:0022625~cytosolic large ribosomal subunit | 17 | 6.9E-09 | P62918, P51410, P35979, P99027, P62717, P12970, O55142, P63323, P61255, P35980, Q9D8E6, P62889, P62900, P27659, P47955, P62830, P41105 | 6.433333333 | 1.5E-07 |
| KEGG_PATHWAY | mmu03010:Ribosome | 41 | 2E-07 | P62918, P35979, P99027, Q9D7S7, P62717, P62281, P12970, Q9DB15, P63323, P35980, P62855, Q9D1N9, Q9CQF0, P25444, P47955, P62852, P62830, P62754, P51410, P62849, P14206, P63276, O55142, P14131, P61255, Q9D8E6, P62702, P62900, P62889, P27659, P41105 | 2.446062685 | 5.5E-06 |
| Annotation Cluster 2 | Enrichment Score: 16.333330528653864 | | | | | |
| Category | Term | Count | PValue | Genes | Fold Enrichment | FDR |
| GOTERM_CC_DIRECT | GO:0005764~lysosome | 54 | 8.9E-23 | Q8BFR4, Q9Z0H7, O70172, O70370, P97821, P18242, P70665, Q9JHJ3, P41251, Q3UIR3, P61027, O09043, O88531, P28798, P10605, Q91VK4, Q9D7V9, Q61207, P50428, P17439, P70699, P12265, P16675, Q9CWK8, P50429, O89017, Q920Q4, P29416, Q9Z0J0, Q9WV54, Q8VCW4, Q80TY0, P20060, O08529, P35951, Q91WL8, P51569, Q9ESY9, O35350, Q9DC37, Q571E4, Q9WVJ3, P58681, Q9QWR8, Q8BMJ2, Q8BX70, Q8VEB4, P23780, P56542, Q9WUU7, O09159, Q3TCN2, Q8BG07 | 5.217521902 | 6.6E-21 |
| UP_KW_CELLULAR_COMPONENT | KW-0458~Lysosome | 62 | 3.2E-15 | P09528, Q8BFR4, O70172, O70370, P97821, P18242, P70665, Q9JHJ3, P41251, Q3UIR3, P61027, O88531, P28798, O35405, P10605, Q9Z127, Q91VK4, Q9D7V9, Q61207, P50428, P17439, Q99MR3, P70699, P12265, P16675, Q58A65, P50429, O89017, Q920Q4, P29416, P63017, P70280, Q9Z0J0, Q9WV54, Q8VCW4, Q80TY0, P20060, P35951, Q91WL8, P51569, Q9ESY9, P10852, P29391, Q9DC37, Q571E4, Q9WVJ3, P58681, Q9QWR8, Q8BPX9, Q8BX70, Q921J2, Q8VEB4, Q64337, P23780, Q9JIA7, P56542, Q9WUU7, O09159, Q3TCN2, O88384, Q8BG07 | 3.137517638 | 6.2E-14 |
| KEGG_PATHWAY | mmu04142:Lysosome | 36 | 3.5E-13 | Q8BFR4, Q9Z0J0, Q9WV54, Q8BVE3, P20060, O70370, P51569, P97821, P18242, P15920, Q571E4, O35643, Q9QWR8, P41251, O09043, P22892, O88531, Q8VEB4, P10605, P23780, Q7TN05, Q61207, P56542, P50428, P17439, P35585, Q9WUU7, P70699, P12265, O09159, P16675, O54774, P50429, O08585, O89017, P29416 | 4.184978123 | 9.9E-11 |

| Annotation Cluster 3 | Enrichment Score: 12.898496773729352 | | | | | |
|---|---|---|---|---|---|---|
| Category | Term | Count | PValue | Genes | Fold Enrichment | FDR |
| GOTERM_CC_DIRECT | GO:0005789~endoplasmic reticulum membrane | 98 | 5.8E-20 | O70378, Q07076, Q6GQT9, Q9CZW4, Q9R233, Q99J47, Q9R1J0, Q3U7R1, Q91VC9, Q9DCN2, O09005, Q8K2C9, P37040, O88736, Q64FW2, P61027, Q8BXZ1, O88455, Q9JJZ4, P47758, O35405, Q9R1S3, Q91YQ5, Q9CXR1, E2JF22, Q9DB25, Q9D6K9, O55143, O54734, Q9DBS1, Q9CQS8, P70280, Q91X58, Q8BYI6, Q8BU14, Q9CQB5, P58681, Q9QYF1, Q9D6J1, Q9D0F3, Q8BX70, Q3TDQ1, Q9Z2G6, Q6DID7, Q9D8V0, Q3U9G9, Q8R0X7, O70503, Q8CCJ3, Q91V01, O88384, Q3UVK0, Q9JLJ5, P11881, Q99L43, Q05769, Q8VCH8, Q80UM7, Q8BHF7, Q9DAV9, Q8BLN5, O08992, Q80WQ6, Q924Z4, Q9DC16, Q9DB73, P61620, Q99PL5, Q60953, P18572, P35564, Q9DBG7, Q9DBG6, P46978, P35293, Q9R0E1, Q8VHI3, Q8BGS7, P61804, Q8VCW4, O70152, Q91ZN5, Q6ZWQ7, Q9QUJ7, Q9CYN2, Q8C7X2, Q9WU81, Q921J2, Q8BM55, P14901, Q80W54, Q99J56, Q8VDP6, Q9CQE7, P08113, B9EJ86, Q8VE91, Q8BG07 | 2.806528882 | 2.8E-18 |
| UP_KW_CELLULAR_COMPONENT | KW-0256~Endoplasmic reticulum | 124 | 2.6E-13 | O70378, Q8BP92, Q6GQT9, Q9CZW4, Q9R233, Q99J47, Q9R1J0, Q3U7R1, Q9DCN2, O09005, O55013, Q8K2C9, P37040, O88736, Q64FW2, P61027, Q8BXZ1, O88455, Q9JJZ4, P47758, O35405, Q99LX0, Q9R1S3, Q91YQ5, Q9CXR1, E2JF22, P50428, P12265, Q9DB25, Q9D6K9, Q8BXQ2, O55143, O54734, Q9DBS1, Q9CQS8, P70280, Q9Z0J0, Q91X58, Q8BYI6, Q8BU14, Q9CQB5, Q9JKR6, P58681, Q9EPB4, Q9QYF1, Q9D6J1, Q9D0F3, Q8BX70, Q8K297, Q3TDQ1, P57759, Q9WTK3, Q9Z2G6, Q6DID7, Q9D8V0, O35381, P31786, Q3U9G9, Q8R0X7, O70503, Q8CCJ3, Q91V01, P11928, Q3UVK0, Q9JLJ5, P11881, Q99L43, Q99K23, Q05769, Q8VCH8, Q80UM7, Q9DAV9, Q8BLN5, Q80UU9, P35980, O08992, Q80WQ6, Q924Z4, Q9DC16, P08003, P61620, Q8BHN3, Q99PL5, Q64518, Q60953, P18572, P35564, Q9DBG7, Q9DBG6, P46978, P35293, Q9R0E1, Q8VHI3, P09103, Q8BGS7, P61804, Q8VCW4, O70152, P14576, Q6ZWQ7, Q9WVJ3, Q9QUJ7, P62855, Q9CYN2, Q8C7X2, Q9WU81, Q921J2, Q8BM55, Q64337, Q9JIA7, P14901, Q80W54, Q6PD26, Q99J56, Q8VDP6, Q6P5E4, P50247, Q9CQE7, Q6NZC7, P08113, B9EJ86, Q8VE91, Q8BG07 | 1.974328764 | 2.5E-12 |
| UP_SEQ_FEATURE | TOPO_DOM:Lumenal | 57 | 1.4E-07 | Q3UVK0, Q9R233, Q9CR60, Q99J47, Q61033, P11881, Q3U7R1, Q9JHJ3, Q80UM7, Q9DAV9, Q8K2C9, P37040, Q80WQ6, Q9DC16, Q924Z4, Q8BXZ1, P61620, Q9JJZ4, O35405, Q99PL5, P27046, Q64518, Q9R1S3, Q91YQ5, P35564, Q9DBG6, P46978, Q9DB25, Q9D6K9, O55143, Q8BXQ2, O54734, P61804, Q8BYI6, Q8BU14, Q9CQB5, Q6ZWQ7, P27808, Q80VA0, Q3UDR8, Q9CYN2, Q8C7X2, Q9D6J1, Q9D0F3, Q3TDQ1, Q9WTK3, Q80W54, Q9Z2G6, Q6PD26, Q6DID7, Q99J56, Q9D8V0, Q8VDP6, Q9CQE7, Q8R0X7, Q8VE91, Q8BG07 | 2.140661258 | 5.4E-05 |

| Annotation Cluster 4 | Enrichment Score: 12.479436827637414 | | | | | |
|---|---|---|---|---|---|---|
| Category | Term | Count | PValue | Genes | Fold Enrichment | FDR |
| GOTERM_MF_DIRECT | GO:0005524~ATP binding | 129 | 9.3E-24 | Q8K1R7, Q8CG47, Q9Z0P5, Q9CZW4, P38647, Q8K1J6, P70268, Q9DBL7, P11440, P68181, Q9WTP6, Q9EPU0, Q9JLV6, P62196, P43274, P49138, Q8BU30, Q91YR1, Q9JJZ4, P56480, Q8K224, Q91VK4, Q9CWX9, P58389, Q64430, Q6PDQ2, Q64433, Q91VC3, P17710, O55143, Q8K183, O88685, Q9JHU4, P32921, Q61655, Q9CZU3, E9PZJ8, Q5SUR0, P61161, Q8CGC7, Q8VDD5, Q9JKR6, Q9CZD3, Q8BMJ2, P04184, P26638, P23249, Q921N6, Q8CGK3, Q9D975, Q61024, P60122, Q04447, Q9DBQ7, Q9QZ08, Q99LE6, Q9D1G2, P11928, Q9Z277, P46471, Q61879, Q99LD9, O70172, Q9D2Y4, P11881, Q7TT37, O08901, Q8BHF7, Q80Y44, Q6PIU9, P61222, P46467, P05132, Q9WV85, Q64518, P70296, Q8JZQ2, O35286, Q9WTM5, Q9Z2I8, P61082, Q01320, Q920A7, Q5SUA5, E9Q9A9, Q8BP47, P09581, Q6Q899, Q9QXG4, P63017, Q61699, Q6PHZ2, Q99J62, Q91YL3, Q9ERS2, Q501J6, Q9R0N0, O08528, P11983, P70248, P68037, Q04899, Q9QYJ0, Q9QUJ7, P55096, P47811, Q9R061, P80314, Q8VI93, Q9ESL4, Q9CQN1, P80313, Q9JIA7, Q60930, Q9EQP2, P62192, Q8BYL4, Q9D0R4, Q9D0R2, Q9WUA2, O09110, P08113, Q9Z110, P23298, P63038, Q6P542, Q8BGQ7, Q9DBU3, E9PVX6 | 2.628417256 | 4.1E-21 |
| UP_KW_LIGAND | KW-0547~Nucleotide-binding | 162 | 7.2E-10 | Q8K1R7, A2AQ07, Q8CG47, Q9CZW4, P38647, Q8K1J6, P70268, Q9DBL7, P11440, P68181, Q9WTP6, Q8VEH6, O55131, Q99LH1, Q9EPU0, Q9JLV6, P62196, P61021, Q9CQD1, P61028, P61027, P49138, Q8BU30, Q9JJZ4, P56480, P47758, Q8K224, Q9DBC7, Q9CWX9, P58389, P10126, Q64430, Q6PDQ2, Q91VC3, P17710, Q01514, O55143, Q8K183, O88685, Q9JHU4, Q8BH04, P32921, Q61655, Q9CZU3, E9PZJ8, Q5SUR0, P61161, Q8CGC7, Q8VDD5, Q9JKR6, Q9CZD3, Q8BMJ2, P04184, P26638, P23249, Q921N6, Q8CGK3, Q9D975, Q921E2, P68372, Q9WUM5, P32233, P68373, P12367, Q61024, P60122, Q04447, Q9QZ08, Q99LE6, Q9D1G2, P08556, P60766, Q3UFY7, P11928, Q922F4, Q9Z277, P46471, Q61879, O70172, O08749, Q9D2Y4, P11881, O08901, P63001, Q8BHF7, Q80Y44, Q6PIU9, P61222, P46467, P05132, Q9WV85, Q60710, Q64518, Q99ME9, P47791, P70296, Q8JZQ2, O35286, Q9WTM5, Q9Z2I8, P61082, Q01320, Q9DBG7, Q920A7, Q5SUA5, P35293, Q8R050, E9Q9A9, Q8BP47, P09581, Q6Q899, Q9QXG4, P63017, P00493, Q9D0S9, Q61699, Q6PHZ2, Q99J62, Q9Z0N1, Q91YL3, Q501J6, Q9R0N0, O08528, P11983, P70248, P14576, P68037, Q04899, Q9QUJ7, P55096, Q921J2, P47811, Q9R061, P80314, Q8VI93, Q9ESL4, Q9CQN1, P80313, Q9JIA7, Q60930, Q9EQP2, P62192, Q8BYL4, Q9D0R4, O08810, Q9D892, Q9D0R2, Q9WUA2, P58252, O09110, P08113, Q9Z110, P23298, P63038, P08752, Q6P542, Q8BGQ7, Q9DBU3, E9PVX6 | 1.51275727 | 2.2E-08 |

| | | | | | | |
|---|---|---|---|---|---|---|
| UP_KW_LIGAND | KW-0067~ATP-binding | 119 | 5.5E-06 | Q8K1R7, Q8CG47, Q9CZW4, P38647, Q8K1J6, P70268, Q9DBL7, P11440, P68181, Q9WTP6, Q9EPU0, Q9JLV6, P62196, P49138, Q8BU30, Q9JJZ4, P56480, Q8K224, Q9CWX9, P58389, Q64430, Q6PDQ2, Q91VC3, P17710, O55143, Q8K183, O88685, Q9JHU4, P32921, Q61655, Q9CZU3, E9PZJ8, Q5SUR0, P61161, Q8CGC7, Q8VDD5, Q9JKR6, Q9CZD3, Q8BMJ2, P04184, P26638, P23249, Q921N6, Q8CGK3, Q9D975, Q61024, P60122, Q04447, Q9QZ08, Q99LE6, Q9D1G2, P11928, Q9Z277, P46471, Q61879, O70172, Q9D2Y4, P11881, O08901, Q8BHF7, Q80Y44, Q6PIU9, P61222, P46467, P05132, Q9WV85, Q64518, P70296, Q8JZQ2, O35286, Q9WTM5, Q9Z2I8, P61082, Q01320, Q920A7, Q5SUA5, E9Q9A9, Q8BP47, P09581, Q6Q899, Q9QXG4, P63017, Q61699, Q6PHZ2, Q99J62, Q91YL3, Q501J6, Q9R0N0, O08528, P11983, P70248, P68037, Q04899, Q9QUJ7, P55096, P47811, Q9R061, P80314, Q8VI93, Q9ESL4, Q9CQN1, P80313, Q9JIA7, Q60930, Q9EQP2, P62192, Q8BYL4, Q9D0R4, Q9D0R2, Q9WUA2, O09110, P08113, Q9Z110, P23298, P63038, Q6P542, Q8BGQ7, Q9DBU3, E9PVX6 | 1.451282051 | 8.2E-05 |
| Annotation Cluster 5 | Enrichment Score: 12.078966929415712 | | | | | |
| Category | Term | Count | PValue | Genes | Fold Enrichment | FDR |
| GOTERM_CC_DIRECT | GO:0005743~mitochondrial inner membrane | 59 | 5E-16 | Q9D0M3, Q9CR68, Q9CWU6, Q9CQQ7, Q9DCJ5, Q3URS9, Q8JZN5, Q9D880, Q9WTP6, Q91VC9, Q9DCN2, O35129, Q9CZN7, Q9CZ13, Q8BGH2, Q9D1N9, P56480, P08249, Q9DB77, Q9CY73, Q8JZQ2, Q9WTQ8, P19783, Q9JIK9, Q9CQX2, Q5U458, Q9CQ75, Q920A7, P67778, Q9DCM2, Q922Q1, Q8BMS1, P54071, Q8QZT1, Q9DB20, Q9QYA2, P51881, P03921, Q9ERS2, Q9CPQ1, Q9DB15, P42125, P09671, P55096, Q9CQF0, Q91VR2, Q99JB2, Q9CQN1, Q9JIA7, Q60930, Q791V5, Q9WUM5, Q99J99, Q9DC69, Q9CQA3, O88986, Q78IK4, Q9Z110, P63038 | 3.453970317 | 1.7E-14 |

| | | | | | | |
|---|---|---|---|---|---|---|
| UP_KW_CELLULAR_COMPONENT | KW-0496~Mitochondrion | 125 | 5.4E-14 | Q9CR68, Q9CZW5, Q9CZW4, P38647, Q8K1J6, Q9DBL7, P11440, Q8JZN5, Q9WTP6, Q9D880, Q91VC9, Q9DCN2, O09005, O35129, Q9CZN7, Q8BGH2, P56480, Q99LX0, Q9WTQ8, Q5U458, Q64433, P54987, P17710, P67778, Q8VEG4, Q9DCM2, Q922Q1, Q8BMS1, Q8BH04, P54071, Q9DB20, P51881, Q921F2, P97287, P03921, Q91WL8, Q9CPQ1, Q62465, Q9DB15, Q9CQB5, Q9CZD3, Q9EPB4, Q8BX70, P45952, P62897, Q8CGK3, Q91VR2, Q99JB2, Q9D6R2, Q921M7, P97333, Q8CHT0, Q8WTY4, O35143, Q9WUM5, P59017, Q99J99, Q9CQA3, Q04447, Q78IK4, Q9D0M3, P11928, Q9D6Y7, P50171, Q9CWU6, Q8CGY8, Q9CQQ7, Q9EP89, O08749, Q9DCJ5, Q3URS9, Q8BHF7, P61222, P63087, Q9CZ13, P05132, P61982, Q9D1N9, P08249, Q9DB77, Q8R5H1, Q9WV85, Q9D8T2, Q9CY73, P52760, P47791, Q9CPV4, Q8JZQ2, P35564, P19783, Q9Z2I8, Q9CQX2, Q9JIK9, Q8BMF4, Q9CQ75, Q920A7, Q9CRA5, Q9CYG7, Q8QZT1, Q9QYA2, Q9D0S9, Q9ERS2, O08528, P50637, Q9D8B6, Q9CPU4, P42125, P09671, Q9QYR9, Q9QUJ7, Q9CQF0, Q9JM90, Q9CQN1, Q9JIA7, Q60930, Q791V5, P18155, P39749, Q8BYL4, O35855, Q9DC69, O88986, Q9Z110, Q8BVW3, P63038 | 2.010061842 | 7E-13 |
| UP_KW_DOMAIN | KW-0809~Transit peptide | 57 | 3.6E-11 | Q9D0M3, Q9CR68, Q9D6Y7, Q9CQQ7, Q9EP89, P38647, O08749, Q8K1J6, Q3URS9, Q8JZN5, Q9D880, Q91VC9, Q8BHF7, Q9CZN7, Q9CZ13, Q9D1N9, P56480, P08249, Q9DB77, Q9CY73, P47791, Q8JZQ2, Q9Z2I8, P19783, Q8BMF4, Q920A7, Q922Q1, Q8BMS1, Q8BH04, P54071, Q8QZT1, Q9DB20, Q9D0S9, Q9D8B6, Q9DB15, P42125, Q9CZD3, P09671, Q9QYR9, Q9CQF0, P45952, Q8CGK3, Q91VR2, Q99JB2, Q9D6R2, Q9CQN1, Q8CHT0, O35143, Q9WUM5, P18155, Q8BYL4, O35855, Q9DC69, Q9CQA3, O88986, Q78IK4, P63038 | 2.66032504 | 1E-09 |
| UP_SEQ_FEATURE | TRANSIT:Mitochondrion | 56 | 4.9E-10 | Q9D0M3, Q9D6Y7, Q9CQQ7, Q9EP89, P38647, O08749, Q8K1J6, Q3URS9, Q8JZN5, Q9D880, Q91VC9, Q8BHF7, Q9CZN7, Q9CZ13, Q9D1N9, P56480, P08249, Q9DB77, Q9CY73, P47791, Q8JZQ2, Q9Z2I8, P19783, Q8BMF4, Q920A7, Q922Q1, Q8BMS1, Q8BH04, P54071, Q8QZT1, Q9DB20, Q9D0S9, Q9D8B6, Q9DB15, P42125, Q9CZD3, P09671, Q9QYR9, Q9CQF0, P45952, Q8CGK3, Q91VR2, Q99JB2, Q9D6R2, Q9CQN1, Q8CHT0, O35143, Q9WUM5, P18155, Q8BYL4, O35855, Q9DC69, Q9CQA3, O88986, Q78IK4, P63038 | 2.53701796 | 2.7E-07 |
| Annotation Cluster 6 | Enrichment Score: 11.666612444245178 | | | | | |
| Category | Term | Count | PValue | Genes | Fold Enrichment | FDR |

| | | | | | | |
|---|---|---|---|---|---|---|
| UP_SEQ_FEATURE | CROSSLNK:Glycyl lysine isopeptide (Lys-Gly) (interchain with G-Cter in SUMO2) | 106 | 9.5E-18 | P62918, Q9CZW5, O70251, Q9R0P4, Q61550, Q61033, P11440, Q9Z2D8, Q9JM76, Q8BHJ5, Q8C7V3, Q8VDF2, Q6DFW4, Q99MR6, Q52KI8, Q9JIX8, P62754, Q61686, Q91WM3, Q61164, Q6PDQ2, Q91VC3, Q3UMU9, P48678, Q9CX86, Q921F2, P83917, Q9CZU3, Q8CBY8, P10852, P51125, Q9JL35, O54941, P28659, A2RSY6, Q91XV3, Q3TEA8, Q3UYV9, Q9D6Z1, Q6PE01, Q9ESX5, P60122, Q8C2Q3, Q9D8E6, P62702, Q9D0E1, Q8BIQ5, P41105, Q9EQK5, Q62448, P35979, P23198, P62717, Q9Z277, Q8BG32, Q99L45, P97822, P12970, Q920B9, Q80Y44, P35980, Q9CYH6, P62317, Q8C156, P27546, P47713, Q99PL5, P46061, Q60710, Q60953, P62849, Q9CX56, Q99ME9, O35286, Q9WTM5, Q01320, Q9CYG7, Q9JLQ0, Q9WVA4, Q8BVY0, Q501J6, P57784, Q9ESU6, Q80X82, Q9QYJ0, Q6NS46, Q9CY57, Q61103, P80314, Q8BHS3, P80313, E9Q7G0, Q922K7, Q91VE6, Q3UEB3, Q05CL8, O88508, P61255, Q61191, Q8C9B9, P24547, P27659, P63038, A2APB8, Q9R190, E9PVX6 | 2.462907876 | 1.3E-14 |
| UP_KW_PTM | KW-1017~Isopeptide bond | 175 | 1.7E-12 | P62918, A2AQ07, Q60864, Q9CZW5, Q9Z0H7, O70251, Q9R0P4, Q61550, Q61033, P11440, Q6P5B0, Q9Z2D8, Q9JM76, Q8BHJ5, Q8C7V3, Q8VDF2, Q6DFW4, Q99MR6, Q52KI8, P61027, Q9JIX8, P49138, P25444, P62754, Q61686, Q99LX0, Q91YQ5, Q91WM3, P10126, P14206, Q61164, Q6PDQ2, Q91VC3, Q3UMU9, P48678, P05064, P62889, Q922Q1, Q9CX86, Q921F2, Q61655, P97287, P83917, Q9CZU3, Q8CBY8, P10852, P51125, Q9DB15, Q9JL35, P11031, Q9QUN7, O88712, Q9EPB4, O54941, P28659, A2RSY6, P62259, Q91XV3, Q921M7, Q8CI51, Q3TEA8, Q3UYV9, P68254, P68372, Q9D6Z1, Q6PE01, Q9ESX5, P60122, Q8C2Q3, Q9D8E6, Q3U0V1, O35892, Q04447, P62702, Q9D0E1, P08556, Q8BIQ5, P41105, Q61510, Q9EQK5, Q62448, P35979, P23198, P62717, Q922F4, Q9Z277, Q8BG32, Q99L45, P97822, P12970, P11881, P17751, Q920B9, P40124, P63001, Q80Y44, P35980, Q9CYH6, O35609, P62317, Q8C156, P27546, P47713, P47955, Q99PL5, P46061, Q80X90, Q60710, Q60953, P62849, Q9CX56, Q99ME9, P26350, O35286, Q9WTM5, Q9CQ71, Q01320, Q9DBG6, Q9DBG5, Q9CYG7, Q61093, Q9JLQ0, Q6Q899, P63017, Q9WVA3, P62962, Q9WVA4, Q8BVY0, P00493, Q99MD9, Q501J6, P57784, Q9R1P4, Q9ESU6, Q80X82, Q9QYJ0, Q5SUF2, Q6NS46, Q9CY57, Q921J2, Q61103, P80314, Q64337, Q8BHS3, Q3UZ39, P80313, Q60930, Q9CQV8, E9Q7G0, Q9WU84, Q922K7, P18155, P62192, P54726, Q8VDP4, Q9D2V7, Q91VE6, Q3UEB3, O08810, Q05CL8, P63276, O88508, P63158, P58252, P61255, Q61191, Q8C9B9, P24547, P27659, P63038, A2APB8, Q9R190, E9PVX6 | 1.674804886 | 1.1E-11 |

| | | | | | | |
|---|---|---|---|---|---|---|
| UP_KW_PTM | KW-0832~Ubl conjugation | 215 | 6.3E-07 | P62918, Q9CZW5, Q9JMH6, O70251, Q9DBD5, Q61550, P11440, Q9Z2D8, Q9JM76, Q52KI8, P61027, A1L314, P49138, Q61686, Q99LX0, Q60591, P10126, Q91VC3, Q9DB20, Q921F2, Q61655, Q9CZU3, Q8BUI3, P51125, Q9WTR6, Q9DB15, Q9JL35, P28659, Q8K298, P62259, Q91XV3, Q3TEA8, Q3UYV9, P68254, Q9D6Z1, P32233, Q8C2Q3, Q3U0V1, P62702, P22682, Q8BIQ5, Q62448, P46471, Q78IQ7, Q8BG32, P40124, P61222, Q9CYH6, P62317, Q8C156, Q80X90, Q60710, Q60953, Q9CX56, Q9WTM5, P35564, Q9CQ71, Q01320, Q9DBG6, Q9DBG5, Q9CYG7, Q9WVA3, Q9WVA4, P00493, Q91YL3, P57784, Q9ESU6, Q80X82, Q9D8B3, Q9QYJ0, Q9CY57, Q921J2, P02301, Q61103, P80314, Q8BHS3, P80313, Q9CQV8, Q60930, E9Q7G0, Q922K7, P54726, Q8VDP4, Q9D2V7, Q91VE6, Q3UEB3, Q9D0R2, O88508, P61255, Q8C9B9, P83940, P24547, Q8BGQ7, Q9R190, E9PVX6, Q60864, Q9Z0H7, Q9R0P4, Q61033, Q6P5B0, Q8BHJ5, Q8VDF2, Q8C7V3, Q6DFW4, Q99MR6, Q9JIX8, P25444, Q9JJZ4, P62754, O35405, Q91YQ5, Q91WM3, Q61164, P14206, Q6PDQ2, P35922, P13597, P16872, Q3UMU9, P01900, P48678, P05064, P62889, Q922Q1, Q9CX86, P97287, P83917, Q8CBY8, Q91WL8, P10852, P11031, Q8VDD5, Q9QUN7, O88712, Q9EPB4, P04184, O54941, P23249, A2RSY6, Q921M7, Q8CI51, Q6PE01, P11157, Q9ESX5, P60122, O35892, Q9D8E6, Q04447, Q9D0E1, P08556, Q8CCJ3, P41105, Q61510, Q9EQK5, P35979, P23198, P62717, Q9Z277, Q8CGY8, Q99L45, P97822, P12970, P11881, P17751, Q920B9, O08901, P63001, Q80Y44, P35980, Q61081, O35609, Q3UIR3, P47955, P27546, P47713, Q99PL5, Q8R5H1, P46061, P62849, Q99ME9, P26350, O35286, Q61093, P09581, Q9JLQ0, Q6Q899, P63017, P62962, Q8BVY0, Q99MD9, P35951, Q501J6, Q9R1P4, P68037, P09671, Q5SUF2, Q6NS46, P55096, P47811, Q64337, Q3UZ39, Q9WU84, P62192, P18155, O08810, Q05CL8, P63276, P58252, Q61191, P27659, Q8BVW3, P63038, A2APB8 | 1.355774884 | 3.2E-06 |
| Annotation Cluster 7 | Enrichment Score: 8.235762070418625 | | | | | |
| Category | Term | Count | PValue | Genes | Fold Enrichment | FDR |
| KEGG_PATHWAY | mmu05020:Prion disease | 48 | 1.9E-10 | P51881, Q9D0M3, A2AQ07, Q9CR68, O70435, Q60864, P99026, P46471, Q922F4, P03921, Q8BG32, Q9CQQ7, O35593, Q9ERS2, Q9R1P1, Q9DCJ5, Q9R1P4, P11881, Q9CPQ1, P68181, P63001, P62196, P63328, Q9Z2U1, Q9CZ13, P05132, P56480, Q9DB77, P62897, O09061, Q91VR2, P47811, Q60930, Q9CX56, P68372, P14106, P62192, P19783, P68373, Q9CQ75, Q9DC69, Q9CQA3, Q61093, O88447, Q9QUM9, O88685, P63017, Q9DB20 | 2.779765689 | 1.5E-08 |

| | | | | | | |
|---|---|---|---|---|---|---|
| KEGG_PATHWAY | mmu05012:Parkinson disease | 48 | 2.2E-10 | P51881, Q9D0M3, A2AQ07, Q9CR68, O70435, Q6PHZ2, P99026, P46471, Q922F4, P03921, Q8BG32, Q78IQ7, Q9CQQ7, O35593, Q9ERS2, Q9R1P1, Q9DCJ5, Q9R1P4, P11881, Q9CPQ1, P68181, P68037, P62196, Q9Z2U1, Q9CZ13, P05132, Q9JJZ4, P56480, Q9DB77, P62897, O09061, Q91VR2, Q9CQN1, Q60930, Q9CX56, Q99LX0, P68372, P62192, P19783, P68373, Q9CQ75, Q9DC69, Q9CQA3, O88447, Q9QUM9, O88685, P08752, Q9DB20 | 2.769620559 | 1.5E-08 |
| KEGG_PATHWAY | mmu05022:Pathways of neurodegeneration - multiple diseases | 66 | 2.8E-09 | Q9D0M3, Q9CR68, A2AQ07, P46471, Q922F4, Q9CQQ7, Q8BG32, Q9DCJ5, P11881, P63001, Q05769, P62196, Q9CQD1, P63328, Q9Z2U1, Q9CZ13, Q9JJZ4, P56480, Q9DB77, Q64518, Q9CX56, Q99LX0, P19783, Q9CQ75, O55143, Q61093, O88447, Q9QUM9, O88685, Q9DB20, Q9QYA2, P51881, Q921F2, O70435, Q6PHZ2, P03921, P99026, O35593, O08529, Q8CBY8, Q9ERS2, Q9R1P1, Q9R1P4, O35350, Q9CPQ1, P68037, P62897, Q91VR2, O09061, P47811, Q64337, Q9CQN1, Q60930, P24270, Q9WU84, P68372, P62192, P68373, Q99J56, Q9DC69, Q9CQA3, O09110, P08556, O89110 | 2.169344169 | 1.1E-07 |
| KEGG_PATHWAY | mmu05014:Amyotrophic lateral sclerosis | 54 | 1.9E-08 | Q9D0M3, Q07076, Q9CR68, A2AQ07, P46471, Q922F4, Q9CQQ7, Q8BG32, Q9DCJ5, F6ZDS4, P63001, P62196, Q9CQD1, P63328, Q9Z2U1, Q9CZ13, P56480, Q9DB77, Q9CX56, P19783, Q9CQ75, O88447, Q9QUM9, O88685, P62962, Q9DB20, Q9QYA2, Q921F2, O70435, P03921, P99026, O35593, Q8CBY8, Q9ERS2, Q9R1P1, Q9R1P4, Q9CPQ1, Q8VCB1, P62897, Q91VR2, O09061, P47811, Q64337, P24270, Q3UYV9, Q9WU84, P68372, P62192, P68373, Q99J56, Q9DC69, Q9CQA3, O08583, O09110 | 2.276628099 | 6.8E-07 |
| KEGG_PATHWAY | mmu05010:Alzheimer disease | 54 | 6.8E-08 | Q9D0M3, Q9CR68, A2AQ07, Q91ZX7, P46471, Q922F4, Q9CQQ7, Q8BG32, Q78IQ7, Q9DCJ5, P11881, Q05769, P62196, P63328, Q9Z2U1, Q9CZ13, P56480, Q9DB77, Q64518, Q9CX56, P19783, Q9CQ75, O55143, Q61093, O88447, Q9QUM9, O88685, Q9DB20, P51881, O70435, P03921, P99026, O35593, O08529, Q9ERS2, Q9R1P1, Q9R1P4, O35350, Q9CPQ1, P11152, Q9Z0F8, P62897, Q91VR2, O09061, Q60930, P68372, P62192, P68373, Q9DC69, Q9CQA3, P08556, O89110 | 2.194692898 | 2.1E-06 |
| KEGG_PATHWAY | mmu05016:Huntington disease | 45 | 2.4E-07 | P51881, Q9D0M3, A2AQ07, Q9CR68, O70435, P99026, P46471, Q922F4, P03921, Q8BG32, Q9CQQ7, Q8CBY8, O35593, Q9ERS2, Q9R1P1, Q9DCJ5, Q9R1P4, P11881, Q9CPQ1, P09671, P62196, Q9Z2U1, Q9CZ13, P56480, Q9DB77, P62897, O09061, Q91VR2, Q60930, Q9CX56, P68372, P62192, P19783, P68373, Q9CQ75, Q9DC69, Q9CQA3, O88447, O08585, Q9QUM9, O88685, P62488, P62743, O89110, Q9DB20 | 2.309890523 | 6.2E-06 |
| Annotation Cluster 8 | Enrichment Score: 7.734442457593948 | | | | | |
| Category | Term | Count | PValue | Genes | Fold Enrichment | FDR |

| | | | | | | |
|---|---|---|---|---|---|---|
| GOTERM_MF_DIRECT | GO:0051015~actin filament binding | 30 | 2E-10 | Q7TPR4, Q9QXZ0, Q61879, Q9Z0P5, P70248, Q61233, P61161, Q8VDD5, Q9JM76, P47753, P47754, Q91YR1, Q8C052, P26039, Q99K51, Q80X90, P24452, O70200, O08539, Q9WUM4, Q5SUA5, P58252, P57780, Q9JLQ0, Q6IRU2, Q920Q4, O88342, P26645, Q9WVA4, P28667 | 4.173335898 | 2.3E-08 |
| GOTERM_MF_DIRECT | GO:0003779~actin binding | 37 | 2.1E-09 | Q7TPR4, Q9QXZ0, P26041, Q61879, P26043, Q9Z0P5, P70248, P19973, P07091, P40124, Q9JHJ0, Q8VDD5, P68510, Q9JM76, P47753, P47754, P61982, Q91YR1, Q8K298, Q8C052, P26039, Q8BH43, Q8CI51, Q6P9Q6, Q9D2V7, Q5SUA5, Q01514, P57780, Q9JLQ0, Q6IRU2, Q6P549, P09103, Q920Q4, O88342, P62962, P26645, P28667 | 3.195735786 | 2.1E-07 |
| UP_KW_MOLECULAR_FUNCTION | KW-0009~Actin-binding | 32 | 1.5E-05 | Q7TPR4, Q9QXZ0, Q61879, P26043, Q9Z0P5, P70248, Q61233, P40124, P61161, Q9JHJ0, Q8VDD5, Q9JM76, P47753, P47754, Q91YR1, Q8K298, Q99K51, Q8BH43, Q80X90, P24452, Q6P9Q6, O70200, Q9WUM4, Q9D2V7, Q5SUA5, P57780, Q6IRU2, Q6P549, O88342, P62962, P26645, P28667 | 2.349174163 | 0.00015 |
| Annotation Cluster 9 | Enrichment Score: 7.508891541528185 | | | | | |
| Category | Term | Count | PValue | Genes | Fold Enrichment | FDR |
| GOTERM_MF_DIRECT | GO:0016491~oxidoreductase activity | 31 | 1.6E-11 | Q9CR68, P50171, Q9DBB8, Q9JMH6, Q91WL8, Q8BVI4, Q99J47, Q62465, Q571I9, P37040, P09671, Q64FW2, Q9DB73, Q9QYF1, O88455, Q9JI75, Q99LB2, Q00519, Q8CHT0, Q9CXR1, P11157, Q99MZ7, Q9CQA3, Q61093, P62482, Q9DCM2, Q9Z110, Q922Q1, O70503, Q8K385, Q8BMS1 | 4.470702034 | 2.1E-09 |
| UP_KW_MOLECULAR_FUNCTION | KW-0560~Oxidoreductase | 65 | 1.2E-07 | P09528, Q9D6Y7, Q923D2, Q922J9, P50171, Q9JMH6, O08749, Q8BVI4, Q99J47, Q9R1J0, Q8JZN5, Q9DCN2, O09005, Q05769, P37040, O88736, Q64FW2, Q9DB73, O88455, Q9JI75, P08249, Q00519, P47791, Q9CXR1, Q99MZ7, Q9R0E1, Q61093, P62482, Q61171, Q922Q1, Q8BMS1, Q8K385, P54071, P48999, P03921, Q9DBB8, Q9CY64, Q91WL8, Q9ESY9, Q8BKE6, Q9CPU4, Q62465, Q00612, P09671, O88712, Q9QYF1, P45952, Q99LB2, Q9D975, Q9D6R2, Q9CQV8, Q8CHT0, P24270, Q9WU84, P14901, P68254, P18155, Q99KP3, P11157, Q9CQA3, Q3U9G9, P24547, Q9Z110, O70503 | 1.998681553 | 1.8E-06 |
| UP_KW_LIGAND | KW-0521~NADP | 27 | 1.5E-05 | Q922J9, Q923D2, Q9DBB8, Q9CY64, Q9JMH6, Q91WL8, Q8BVI4, Q99J47, P37040, O88736, Q00612, Q64FW2, Q9QYF1, O88455, Q8K2T1, Q99LB2, P24270, P47791, Q9CXR1, P18155, Q99MZ7, Q61093, P62482, Q9Z110, O70503, P54071 | 2.555518395 | 0.00015 |
| Annotation Cluster 10 | Enrichment Score: 5.502289852572369 | | | | | |
| Category | Term | Count | PValue | Genes | Fold Enrichment | FDR |
| GOTERM_CC_DIRECT | GO:0000502~proteasome complex | 18 | 5.5E-13 | Q91X58, O70435, P99026, P46471, Q9CX56, Q9JMA1, Q8BG32, O35593, Q9R1P1, P28063, Q9R1P4, P62192, P54726, P62196, Q9Z2U1, Q9QUM9, O88685, O09061 | 10.40342246 | 1.5E-11 |

| | | | | | | |
|---|---|---|---|---|---|---|
| GOTERM_BP_DIRECT | GO:0051603~proteolysis involved in protein catabolic process | 19 | 6.5E-13 | P10605, O70435, P99026, O08529, O70370, P20918, P97821, Q9R1P1, P28063, Q9R1P4, Q9WUU7, O89017, Q9Z2U1, Q9QUM9, O09061, O35955, P28798, O89110, Q8CGK3 | 9.484677419 | 4.2E-10 |
| UP_KW_CELLULAR_COMPONENT | KW-0647~Proteasome | 20 | 1.1E-12 | P97372, O70435, P99026, P46471, Q9CX56, Q9JMA1, Q8BG32, O35593, Q9R1P1, P28063, Q9R1P4, P62192, P54726, P62196, Q9Z2U1, Q9QUM9, O88685, O09061, O35955, P97371 | 8.077356202 | 8.2E-12 |
| KEGG_PATHWAY | mmu03050:Proteasome | 18 | 1.5E-09 | P97372, O70435, P99026, P46471, Q9CX56, Q8BG32, O35593, Q9R1P1, P28063, Q9R1P4, P62192, P62196, Q9Z2U1, Q9QUM9, O88685, O09061, O35955, P97371 | 6.054861966 | 8.3E-08 |
| INTERPRO | IPR029055:Ntn_hydrolases_N | 12 | 2.9E-08 | O70435, P99026, P28063, Q9Z2U1, Q9QUM9, Q9R1P1, Q9R1P4, Q8C0M9, P47856, O09061, O35955, Q61024 | 9.122785458 | 1.7E-05 |
| GOTERM_CC_DIRECT | GO:0005839~proteasome core complex | 9 | 5.1E-08 | O70435, P99026, P28063, Q9Z2U1, Q9QUM9, Q9R1P1, Q9R1P4, O09061, O35955 | 15.05758514 | 1.1E-06 |
| INTERPRO | IPR001353:Proteasome_sua/b | 9 | 5E-07 | O70435, P99026, P28063, Q9Z2U1, Q9QUM9, Q9R1P1, Q9R1P4, O09061, O35955 | 11.16340852 | 0.00018 |
| KEGG_PATHWAY | mmu05017:Spinocerebellar ataxia | 24 | 2.3E-05 | P51881, O70435, Q60930, Q64518, P99026, P46471, Q9CX56, Q8BG32, O35593, Q8JZQ2, Q9R1P1, Q9R1P4, P62192, Q9D6Z1, P11881, P31266, Q920A7, O55143, P62196, Q9Z2U1, Q9QUM9, O88685, O09061, P62897 | 2.691049763 | 0.00045 |
| GOTERM_CC_DIRECT | GO:0019774~proteasome core complex, beta-subunit complex | 5 | 0.00027 | P99026, P28063, Q9R1P1, O09061, O35955 | 14.44919786 | 0.0025 |
| GOTERM_BP_DIRECT | GO:0043161~proteasome-mediated ubiquitin-dependent protein catabolic process | 24 | 0.00043 | Q91X58, O70435, Q60953, P99026, P46471, Q9CX56, Q9JMA1, O35593, Q9R1P1, P28063, Q9R1P4, P62192, P17439, P54726, Q99J56, Q8BHJ5, Q9JJU8, P62196, Q9JLQ0, Q9Z2U1, Q9QUM9, O88685, Q8CCH2, O35955 | 2.275256054 | 0.02382 |
| INTERPRO | IPR023333:Proteasome_suB-type | 5 | 0.00083 | P99026, P28063, Q9R1P1, O09061, O35955 | 10.71236171 | 0.09767 |
| INTERPRO | IPR016050:Proteasome_bsu_CS | 5 | 0.00121 | P99026, P28063, Q9R1P1, O09061, O35955 | 9.819664903 | 0.11063 |
| GOTERM_CC_DIRECT | GO:0019773~proteasome core complex, alpha-subunit complex | 4 | 0.00154 | O70435, Q9Z2U1, Q9QUM9, Q9R1P4 | 15.89411765 | 0.01098 |
| Annotation Cluster 11 | Enrichment Score: 5.219491526786178 | | | | | |
| Category | Term | Count | PValue | Genes | Fold Enrichment | FDR |
| GOTERM_BP_DIRECT | GO:0006457~protein folding | 19 | 1.2E-07 | P80314, Q9CQN1, Q61699, P80313, Q9WU28, P38647, P11983, P35564, Q8VCN9, Q64433, Q9JKR6, Q61081, Q9QYJ0, P08113, P08003, P09103, P63038, P48428, P63017 | 4.705576394 | 3.5E-05 |
| GOTERM_MF_DIRECT | GO:0140662~ATP-dependent protein folding chaperone | 10 | 2.5E-07 | P80314, Q61699, P80313, Q9CQN1, Q9JKR6, P38647, P08113, P11983, P63038, P63017 | 10.47971014 | 1.8E-05 |
| UP_KW_MOLECULAR_FUNCTION | KW-0143~Chaperone | 31 | 5.7E-07 | P38647, P11983, P97822, Q8VEH6, Q91YE6, Q9JKR6, P61222, Q61081, Q64152, Q9QYJ0, P48428, P80314, Q9CQN1, P80313, Q99LX0, Q9WU84, Q9WU28, Q8VCN9, Q9EST4, P35564, Q9EST5, Q5U458, Q64433, O08997, Q9CPP0, O08583, Q9CYG7, P08113, P09103, P63038, P63017 | 2.803325588 | 7.1E-06 |
| GOTERM_MF_DIRECT | GO:0051082~unfolded protein binding | 15 | 1.2E-06 | Q9R0U0, P80314, Q9CQN1, P80313, Q9WU28, P38647, P11983, P35564, O08915, Q64433, Q6P5E4, Q61081, Q9QYJ0, P08113, P63017 | 5.12594518 | 6E-05 |

| | | | | | | |
|---|---|---|---|---|---|---|
| GOTERM_MF_DIRECT | GO:0044183~protein folding chaperone | 10 | 7.3E-05 | P80314, P80313, Q3U0V1, P38647, P08113, Q9R233, P11983, P97822, Q64433, P63017 | 5.515636918 | 0.00252 |
| Annotation Cluster 12 | Enrichment Score: 5.217946081381777 | | | | | |
| Category | Term | Count | PValue | Genes | Fold Enrichment | FDR |
| GOTERM_BP_DIRECT | GO:0045087~innate immune response | 43 | 1.4E-08 | Q61510, Q9QZI9, P11928, B2RRE7, Q8VCW4, Q9Z0H7, P15307, P30681, Q8VC04, Q9JL16, Q80VI1, Q9QUN7, P58681, Q2EMV9, Q9EPB4, Q3UIR3, A1L314, Q8BPX9, Q99NH8, O35405, Q8VI94, Q8VI93, Q9D8T2, Q60710, Q60953, P20491, Q9JMA1, P14106, Q9R0Q8, P17710, E9Q9A9, P63158, Q8C2Q3, Q8C522, Q61093, P09581, Q9QZ08, P01029, Q8BVW3, Q6Q899, Q9DBS1, Q8BG07, Q9DBU3 | 2.698979657 | 5.5E-06 |
| UP_KW_BIOLOGICAL_PROCESS | KW-0399~Innate immunity | 44 | 2.3E-07 | Q61510, Q9QZI9, P11928, B2RRE7, Q8VCW4, Q9Z0H7, P30681, Q8VC04, Q9JL16, Q80VI1, Q9QUN7, P58681, Q2EMV9, Q9EPB4, Q3UIR3, A1L314, Q8BPX9, O35405, Q8VI94, Q8VI93, Q9D8T2, Q60710, Q60953, P20491, Q9JMA1, O35286, P14106, Q9R0Q8, P54987, P17710, Q01514, E9Q9A9, P63158, Q8C2Q3, Q8C522, P09581, Q9QZ08, P01029, P01027, Q8BVW3, Q6Q899, Q9DBS1, Q8BG07, Q9DBU3 | 2.361229606 | 6.6E-06 |
| UP_KW_BIOLOGICAL_PROCESS | KW-0391~Immunity | 58 | 0.06901 | Q9QZI9, Q61510, P11928, B2RRE7, Q9Z0H7, P30681, Q8VC04, Q9JL16, Q3UIR3, A1L314, O35405, Q9D8T2, Q60710, Q60953, P20491, O35286, Q9R0Q8, P54987, Q9ES52, Q5SUA5, P17710, Q01514, E9Q9A9, Q64281, P01902, Q61490, P09581, P01900, P01029, Q6Q899, P01027, Q9DBS1, Q8VCW4, Q501J6, P28063, Q9ESY9, Q9R1P4, Q80VI1, Q9QUN7, P58681, Q2EMV9, Q9EPB4, Q8BPX9, Q8VI94, Q8VI93, Q64337, Q9JMA1, P14106, P63158, Q8C2Q3, Q8C522, Q9QZ08, Q6P549, Q8BVW3, Q8BG07, Q9DBU3 | 1.234333264 | 0.25017 |
| Annotation Cluster 13 | Enrichment Score: 3.6500226919424854 | | | | | |
| Category | Term | Count | PValue | Genes | Fold Enrichment | FDR |
| GOTERM_BP_DIRECT | GO:0005975~carbohydrate metabolic process | 16 | 3.9E-06 | P23780, P27046, P20060, Q924M7, O08528, P51569, P10852, Q9CQ60, P70699, P12265, P70665, O88958, Q9QWR8, P29416, Q9D0F9, Q8BHN3 | 4.369010201 | 0.00056 |
| INTERPRO | IPR013780:Glyco_hydro_b | 7 | 0.00024 | O09159, P27046, Q9QWR8, P51569, P10852, P70699, Q8BHN3 | 7.498653199 | 0.04638 |
| INTERPRO | IPR017853:Glycoside_hydrolase_SF | 10 | 0.00047 | P23780, Q9QWR8, P20060, P51569, P10852, P17439, P70699, P29416, P12265, Q8BHN3 | 4.284944685 | 0.07608 |
| UP_KW_MOLECULAR_FUNCTION | KW-0326~Glycosidase | 13 | 0.00076 | P23780, P27046, P20060, P51569, P17439, P70699, P12265, O09159, Q80UM7, Q80VJ3, Q9QWR8, P29416, Q8BHN3 | 3.154016987 | 0.00568 |
| GOTERM_MF_DIRECT | GO:0004553~hydrolase activity, hydrolyzing O-glycosyl compounds | 5 | 0.00173 | P23780, Q9QWR8, P51569, P70699, P12265 | 9.246803069 | 0.03512 |

| Annotation Cluster 14 | Enrichment Score: 3.648494830930841 | | | | | |
|---|---|---|---|---|---|---|
| Category | Term | Count | PValue | Genes | Fold Enrichment | FDR |
| GOTERM_BP_DIRECT | GO:0018279~protein N-linked glycosylation via asparagine | 8 | 6.7E-07 | P46978, P61804, P27808, Q91YQ5, O54734, Q9JJZ4, Q3TDQ1, Q6P5E4 | 14.19928315 | 0.00012 |
| GOTERM_CC_DIRECT | GO:0008250~oligosaccharyltransferase complex | 6 | 4.8E-05 | P46978, P61804, Q91YQ5, O54734, Q3TDQ1, Q9DBG6 | 13.62352941 | 0.00054 |
| GOTERM_BP_DIRECT | GO:0006487~protein N-linked glycosylation | 10 | 9.7E-05 | P46978, Q80UM7, Q9DB25, P61804, P27808, Q91YQ5, O54734, Q3TDQ1, P47856, Q9DBG6 | 5.324731183 | 0.00789 |
| KEGG_PATHWAY | mmu00510:N-Glycan biosynthesis | 12 | 0.00046 | P46978, Q80UM7, Q9DB25, P61804, P27046, P27808, Q91YQ5, O54734, O70152, Q3TDQ1, Q8BHN3, Q9DBG6 | 3.513314968 | 0.00642 |
| KEGG_PATHWAY | mmu00513:Various types of N-glycan biosynthesis | 10 | 0.00122 | P46978, P61804, P27046, P27808, P20060, Q91YQ5, O54734, Q3TDQ1, P29416, Q9DBG6 | 3.676724966 | 0.01191 |
| Annotation Cluster 15 | Enrichment Score: 3.5506926701548456 | | | | | |
| Category | Term | Count | PValue | Genes | Fold Enrichment | FDR |
| GOTERM_MF_DIRECT | GO:0016887~ATP hydrolysis activity | 49 | 2.7E-14 | Q61655, Q8CG47, Q99J62, Q9QXZ0, P46471, Q9CZU3, Q9CQQ7, P38647, Q501J6, P11983, E9PZJ8, P14576, Q9EPU0, Q80Y44, P61222, P62196, P23249, P55096, P46467, Q921N6, P56480, Q8CGK3, Q91VR2, P80314, Q9CQN1, P80313, Q64518, Q9CWX9, Q8JZQ2, O35286, Q9WTM5, P62192, Q64430, Q6PDQ2, Q9D0R4, Q9DBG7, Q91VC3, Q920A7, O55143, P60122, P08113, Q99LE6, O88685, Q6Q899, Q9JHU4, P63017, Q6P542, Q9DB20 | 3.641885086 | 4E-12 |
| GOTERM_MF_DIRECT | GO:0003689~DNA clamp loader activity | 20 | 4.8E-07 | Q61655, Q99J62, Q9CZU3, Q9CWX9, Q501J6, Q8JZQ2, Q9WTM5, O35286, E9PZJ8, Q6PDQ2, Q9D0R4, Q91VC3, Q920A7, Q9EPU0, Q80Y44, P60122, P23249, P46467, Q6Q899, Q921N6 | 4.083003953 | 3.2E-05 |
| GOTERM_MF_DIRECT | GO:0140849~ATP-dependent H2AZ histone chaperone activity | 19 | 8.7E-07 | Q61655, Q9CZU3, Q9CWX9, Q501J6, Q8JZQ2, Q9WTM5, O35286, E9PZJ8, Q6PDQ2, Q9D0R4, Q91VC3, Q920A7, Q9EPU0, Q80Y44, P60122, P23249, P46467, Q6Q899, Q921N6 | 4.119610195 | 5E-05 |
| GOTERM_MF_DIRECT | GO:0140584~chromatin extrusion motor activity | 19 | 8.7E-07 | Q61655, Q9CZU3, Q9CWX9, Q501J6, Q8JZQ2, Q9WTM5, O35286, E9PZJ8, Q6PDQ2, Q9D0R4, Q91VC3, Q920A7, Q9EPU0, Q80Y44, P60122, P23249, P46467, Q6Q899, Q921N6 | 4.119610195 | 5E-05 |
| GOTERM_MF_DIRECT | GO:0140665~ATP-dependent H3-H4 histone complex chaperone activity | 19 | 8.7E-07 | Q61655, Q9CZU3, Q9CWX9, Q501J6, Q8JZQ2, Q9WTM5, O35286, E9PZJ8, Q6PDQ2, Q9D0R4, Q91VC3, Q920A7, Q9EPU0, Q80Y44, P60122, P23249, P46467, Q6Q899, Q921N6 | 4.119610195 | 5E-05 |
| GOTERM_MF_DIRECT | GO:0061775~cohesin loader activity | 19 | 1.1E-06 | Q61655, Q9CZU3, Q9CWX9, Q501J6, Q8JZQ2, Q9WTM5, O35286, E9PZJ8, Q6PDQ2, Q9D0R4, Q91VC3, Q920A7, Q9EPU0, Q80Y44, P60122, P23249, P46467, Q6Q899, Q921N6 | 4.063561077 | 5.6E-05 |
| GOTERM_MF_DIRECT | GO:0003724~RNA helicase activity | 11 | 7E-05 | Q61655, Q9EPU0, Q80Y44, Q9CZU3, Q9CWX9, Q501J6, O35286, Q6Q899, Q921N6, Q9D0R4, Q91VC3 | 4.940434783 | 0.00251 |

| | | | | | | |
|---|---|---|---|---|---|---|
| GOTERM_MF_DIRECT | GO:0003676~nucleic acid binding | 23 | 0.00018 | Q60710, Q9CZU3, Q9CWX9, Q501J6, Q9CYA6, O35286, E9PZJ8, Q9D0R4, Q91VC3, P63001, Q9JL16, P63158, Q8BP47, Q8VDF2, Q8C522, Q8VEG4, Q3U0V1, Q9D2E2, Q9JIX8, Q6NS46, Q6Q899, Q921N6, P62488 | 2.476369863 | 0.00576 |
| INTERPRO | IPR011545:DEAD/ DEAH_box_helicase_dom | 11 | 0.00098 | Q61655, Q80Y44, Q9CZU3, Q9CWX9, Q501J6, O35286, E9PZJ8, Q6Q899, Q921N6, Q9D0R4, Q91VC3 | 3.55122128 | 0.09928 |
| UP_KW_MOLECULAR_FUNCTION | KW-0347~Helicase | 16 | 0.00616 | Q61655, Q9CZU3, Q9CWX9, Q501J6, Q9WTM5, O35286, E9PZJ8, Q6PDQ2, Q9D0R4, Q91VC3, Q9EPU0, Q80Y44, P60122, P23249, Q6Q899, Q921N6 | 2.195262752 | 0.03299 |
| Annotation Cluster 16 | Enrichment Score: 3.527697970986361 | | | | | |
| Category | Term | Count | PValue | Genes | Fold Enrichment | FDR |
| SMART | SM00382:AAA | 21 | 2.5E-06 | Q99J62, P46471, Q8JZQ2, Q9WTM5, P62192, E9PZJ8, Q9DBG7, P14576, Q920A7, P61222, P60122, P62196, P55096, O88685, Q99LE6, P46467, Q9JHU4, P56480, Q6P542, Q8CGK3 | 3.451739366 | 0.00082 |
| INTERPRO | IPR003593:AAA+_ATPase | 21 | 3.7E-06 | Q99J62, P46471, Q8JZQ2, Q9WTM5, P62192, E9PZJ8, Q9DBG7, P14576, Q920A7, P61222, P60122, P62196, P55096, O88685, Q99LE6, P46467, Q9JHU4, P56480, Q6P542, Q8CGK3 | 3.389802131 | 0.00121 |
| GOTERM_CC_DIRECT | GO:0022624~proteasome accessory complex | 7 | 8.7E-06 | Q9CX56, P46471, P62196, Q8BG32, O35593, P62192, O88685 | 13.08927336 | 0.00012 |
| UP_SEQ_FEATURE | DOMAIN:AAA+ ATPase | 10 | 7.4E-05 | Q99J62, P60122, P46471, P62196, Q8JZQ2, Q9WTM5, P62192, O88685, P46467, Q9JHU4 | 5.416331995 | 0.02056 |
| GOTERM_MF_DIRECT | GO:0036402~proteasome-activating activity | 4 | 0.0006 | P46471, P62196, P62192, O88685 | 20.95942029 | 0.01704 |
| INTERPRO | IPR041569:AAA_lid_3 | 7 | 0.00063 | P46471, P62196, Q8JZQ2, P62192, O88685, P46467, Q920A7 | 6.345014245 | 0.08841 |
| INTERPRO | IPR003959:ATPase_AAA_core | 9 | 0.00073 | Q99J62, P46471, P62196, Q8JZQ2, P62192, O88685, P46467, Q8CGK3, Q920A7 | 4.512867275 | 0.09593 |
| INTERPRO | IPR003960:ATPase_AAA_CS | 7 | 0.00096 | P46471, P62196, Q8JZQ2, P62192, O88685, P46467, Q920A7 | 5.891798942 | 0.09928 |
| INTERPRO | IPR050221:26S_Proteasome_ATPase | 4 | 0.00138 | P46471, P62196, P62192, O88685 | 15.71146384 | 0.12243 |
| GOTERM_MF_DIRECT | GO:0017025~TBP-class protein binding | 6 | 0.00236 | P60122, P46471, P62196, Q9WTM5, P62192, Q6ZQ38 | 6.287826087 | 0.04313 |
| GOTERM_CC_DIRECT | GO:0008540~proteasome regulatory particle, base subcomplex | 4 | 0.0055 | P46471, P62196, P62192, O88685 | 10.59607843 | 0.02968 |
| Annotation Cluster 17 | Enrichment Score: 3.509550799125334 | | | | | |
| Category | Term | Count | PValue | Genes | Fold Enrichment | FDR |
| GOTERM_BP_DIRECT | GO:0045047~protein targeting to ER | 8 | 4.2E-07 | Q91X58, P16254, Q6ZWQ7, Q9CYN2, P47758, Q9DBG7, P14576 | 15.0345351 | 9.2E-05 |
| GOTERM_BP_DIRECT | GO:0006614~SRP-dependent cotranslational protein targeting to membrane | 6 | 3.1E-05 | P16254, P61620, Q9D7A6, Q9DBG7, P14576 | 14.74540943 | 0.00325 |
| KEGG_PATHWAY | mmu03060:Protein export | 11 | 0.00022 | Q9CQS8, Q8BU14, P16254, Q6ZWQ7, Q9CYN2, P61620, Q9D7A6, P47758, Q9DBG7, P14576 | 4.140692641 | 0.0032 |

| | | | | | | |
|---|---|---|---|---|---|---|
| UP_KW_CELLULAR_COMPONENT | KW-0733~Signal recognition particle | 4 | 0.00192 | P16254, Q9D7A6, P14576 | 14.00075075 | 0.00751 |
| GOTERM_CC_DIRECT | GO:0005786~signal recognition particle, endoplasmic reticulum targeting | 4 | 0.00225 | P16254, Q9D7A6, P14576 | 14.12810458 | 0.01483 |
| GOTERM_MF_DIRECT | GO:0008312~7S RNA binding | 4 | 0.00233 | P16254, Q9D7A6, P14576 | 13.97294686 | 0.04313 |
| GOTERM_MF_DIRECT | GO:0030942~endoplasmic reticulum signal peptide binding | 3 | 0.00946 | P16254, P14576 | 18.86347826 | 0.13397 |
| Annotation Cluster 18 | Enrichment Score: 3.467592314536267 | | | | | |
| Category | Term | Count | PValue | Genes | Fold Enrichment | FDR |
| UP_KW_DOMAIN | KW-0676~Redox-active center | 12 | 4.3E-06 | Q9D6Y7, Q9CQM5, Q9QUH0, P47791, Q61171, Q9JMH6, O08749, P08003, P09103, Q8BXZ1, Q9ESY9, Q99J99 | 5.886002173 | 6E-05 |
| UP_SEQ_FEATURE | DISULFID:Redox-active | 10 | 0.00024 | Q9CQM5, Q9QUH0, P47791, Q9JMH6, O08749, P08003, P09103, Q8BXZ1, Q9ESY9, Q9CQ75 | 4.68721038 | 0.05505 |
| Annotation Cluster 19 | Enrichment Score: 3.3162320066713202 | | | | | |
| Category | Term | Count | PValue | Genes | Fold Enrichment | FDR |
| GOTERM_MF_DIRECT | GO:0017124~SH3 domain binding | 17 | 3.9E-06 | Q8BH43, Q9D358, Q9WU78, Q3UIA2, Q62422, Q9ES52, Q8CBW3, P35831, Q9Z0F8, Q9JJU8, Q9JJ00, Q9JLQ0, Q8BIJ7, Q6P549, Q8BPU7, P55194, P22682 | 4.111270903 | 0.00017 |
| UP_KW_DOMAIN | KW-0729~SH3-binding | 9 | 0.00198 | Q9Z0F8, Q9JJU8, Q9JJ00, Q9JLQ0, Q3UIA2, Q6P549, P55194, Q8BPU7, Q9ES52 | 3.89947644 | 0.01849 |
| Annotation Cluster 20 | Enrichment Score: 3.1810523776832933 | | | | | |
| Category | Term | Count | PValue | Genes | Fold Enrichment | FDR |
| INTERPRO | IPR036013:Band_7/SPFH_dom_sf | 6 | 0.00013 | P67778, O35129, P54116, Q9EQK5, O08917, Q99JB2 | 10.87716728 | 0.032 |
| UP_SEQ_FEATURE | DOMAIN:Band 7 | 5 | 0.00107 | P67778, O35129, P54116, O08917, Q99JB2 | 10.15562249 | 0.19867 |
| SMART | SM00244:PHB | 5 | 0.00112 | P67778, O35129, P54116, O08917, Q99JB2 | 9.999086257 | 0.09172 |
| INTERPRO | IPR001107:Band_7 | 5 | 0.00121 | P67778, O35129, P54116, O08917, Q99JB2 | 9.819664903 | 0.11063 |
| Annotation Cluster 21 | Enrichment Score: 3.09038879539579 | | | | | |
| Category | Term | Count | PValue | Genes | Fold Enrichment | FDR |
| GOTERM_MF_DIRECT | GO:0005525~GTP binding | 39 | 1.1E-09 | A2AQ07, Q9Z0N1, Q922F4, Q99LD9, P14576, P63001, Q8VEH6, O55131, Q99LH1, P61021, P43274, Q9CQD1, P61028, P61027, Q921J2, P47758, Q60710, Q99ME9, Q921E2, P68372, Q9Z2I8, P32233, P10126, Q9EQP2, P68373, Q9DBG7, O08810, P35293, Q01514, Q8R050, P58252, Q6Q899, P08556, Q8BH04, P08752, P60766 | 3.143913043 | 1.3E-07 |

| | | | | | | |
|---|---|---|---|---|---|---|
| GOTERM_MF_DIRECT | GO:0003924~GTPase activity | 32 | 2.9E-08 | A2AQ07, Q9Z0N1, Q922F4, P14576, P63001, Q8VEH6, O55131, P61222, P61021, Q9CQD1, P61028, P61027, Q921J2, Q99ME9, Q921E2, Q8VCN9, P68372, P32233, P10126, Q9DBG7, O08810, P35293, Q01514, Q8R050, P63213, P58252, P08556, P08752, P60766 | 3.203987815 | 2.5E-06 |
| GOTERM_MF_DIRECT | GO:0019003~GDP binding | 15 | 3.3E-07 | P35293, Q01514, Q921E2, P61021, Q9CQD1, P61028, P61027, Q9WUM5, Q9Z2I8, Q921J2, P08556, P14576 | 5.681770561 | 2.3E-05 |
| GOTERM_MF_DIRECT | GO:0003925~G protein activity | 9 | 0.00042 | P63001, Q01514, P61021, Q9CQD1, P61027, P08556, P60766 | 4.964073227 | 0.01227 |
| INTERPRO | IPR005225:Small_GTP-bd | 17 | 0.00202 | Q99ME9, Q921E2, P32233, O08810, P63001, P35293, Q8R050, P58252, P61021, Q9CQD1, P61028, P61027, Q921J2, P08556, P60766 | 2.399055857 | 0.16901 |
| UP_KW_LIGAND | KW-0342~GTP-binding | 35 | 0.00343 | A2AQ07, Q9Z0N1, Q922F4, P14576, P63001, Q8VEH6, O55131, Q99LH1, P61021, Q9CQD1, P61028, P61027, Q921J2, P47758, Q60710, Q99ME9, Q921E2, P68372, Q9Z2I8, P32233, P10126, P68373, Q9DBG7, O08810, P35293, Q01514, Q8R050, P58252, P08556, Q8BH04, P08752, P60766 | 1.660867743 | 0.02056 |
| Annotation Cluster 22 | Enrichment Score: 2.9107338169007297 | | | | | |
| Category | Term | Count | PValue | Genes | Fold Enrichment | FDR |
| GOTERM_CC_DIRECT | GO:0005778~peroxisomal membrane | 10 | 0.0001 | Q922J9, P24270, Q9CZW4, P61028, Q9QUJ7, P55096, Q9Z210, Q99LB2, Q99MZ7 | 5.298039216 | 0.00107 |
| GOTERM_CC_DIRECT | GO:0005777~peroxisome | 15 | 0.00011 | P52760, Q922J9, P24270, Q99J47, Q99MZ7, Q9DCM2, Q9QUJ7, P55096, Q9Z210, Q922Q1, Q9D1G2, Q99LB2, P54071, Q00519 | 3.480463718 | 0.00115 |
| KEGG_PATHWAY | mmu04146:Peroxisome | 13 | 0.00822 | Q922J9, P24270, Q9CZW4, Q99MZ7, P09671, Q9DCM2, Q9QUJ7, P55096, Q9Z210, Q9D1G2, Q99LB2, P54071, Q00519 | 2.362401444 | 0.04882 |
| UP_KW_CELLULAR_COMPONENT | KW-0576~Peroxisome | 12 | 0.02495 | P52760, Q922J9, P24270, Q9CZW4, Q9QUJ7, P55096, Q9Z210, Q922Q1, Q99LB2, Q00519, Q99MZ7 | 2.135707742 | 0.07484 |
| Annotation Cluster 23 | Enrichment Score: 2.909229348992878 | | | | | |
| Category | Term | Count | PValue | Genes | Fold Enrichment | FDR |
| KEGG_PATHWAY | mmu05415:Diabetic cardiomyopathy | 30 | 9.7E-05 | P51881, Q9D0M3, Q9CR68, Q6PHZ2, P03921, Q9CQQ7, Q9ERS2, Q9DCJ5, P18242, Q9CPQ1, P63001, Q00612, P63087, Q9CZ13, P56480, P47856, Q9DB77, Q91VR2, P47811, Q60930, Q64518, P47791, P58389, P19783, Q9CQ75, Q9DC69, Q9CQA3, O55143, Q61093, Q9DB20 | 2.18570286 | 0.00168 |
| UP_KW_CELLULAR_COMPONENT | KW-0999~Mitochondrion inner membrane | 33 | 0.00022 | P51881, Q9D0M3, Q9CR68, P03921, Q9CWU6, Q9CQQ7, Q9ERS2, Q9DCJ5, Q3URS9, Q9CPQ1, Q8JZN5, Q9D880, Q91VC9, O35129, Q9CZN7, Q9CZ13, P56480, Q9DB77, Q91VR2, Q99JB2, Q9CQN1, Q9JIA7, Q8JZQ2, Q9WTQ8, P19783, Q9CQ75, Q920A7, P67778, Q9CQA3, Q78IK4, Q9Z110, Q8BMS1, Q9DB20 | 2.026424451 | 0.00105 |

| | | | | | | |
|---|---|---|---|---|---|---|
| KEGG_PATHWAY | mmu05208:Chemical carcinogenesis - reactive oxygen species | 29 | 0.00056 | P51881, Q9D0M3, Q9CR68, P03921, Q9CQQ7, Q9ERS2, Q9DCJ5, Q9CPQ1, Q9CPU4, P63001, P09671, Q9CZ13, P56480, Q9DB77, Q91VR2, P47811, Q9D358, Q60930, P24270, P14901, P19783, Q9CQ75, Q9DC69, Q9CQA3, P48774, P08556, Q9DB20 | 2.002129272 | 0.00698 |
| UP_KW_BIOLOGICAL_PROCESS | KW-0249~Electron transport | 15 | 0.00087 | Q9D0M3, Q9CR68, P03921, Q9ERS2, Q9DCJ5, Q9CQX2, Q9CQ75, Q9DC69, Q9CQA3, Q9QUH0, Q61093, Q9CZ13, Q8K385, Q9DB77, P62897 | 2.799567283 | 0.0078 |
| KEGG_PATHWAY | mmu04714:Thermogenesis | 29 | 0.00097 | Q9D0M3, Q9CR68, P03921, Q9CZW4, Q9CQQ7, Q9ERS2, Q9DCJ5, Q9CPQ1, P68181, Q9QUJ7, Q9CZ13, O54941, P05132, Q921J2, P56480, Q9DB77, P62754, Q91VR2, P47811, P19783, Q9CQ75, O35678, Q9DC69, Q9CQA3, O09110, P08556, Q9DB20 | 1.934546849 | 0.01041 |
| KEGG_PATHWAY | mmu00190:Oxidative phosphorylation | 20 | 0.00135 | Q9D0M3, Q9CR68, Q8BVE3, P03921, Q9CQQ7, Q9ERS2, Q9DCJ5, P19783, P15920, Q9CPQ1, Q9CQ75, Q9DC69, Q9CQA3, Q9CZ13, P56480, Q9DB77, P62897, Q91VR2, Q9DB20, Q9D819 | 2.242541469 | 0.01257 |
| UP_KW_BIOLOGICAL_PROCESS | KW-0679~Respiratory chain | 10 | 0.00455 | Q9D0M3, Q9DC69, Q9CR68, P03921, Q9ERS2, Q9DCJ5, Q9CZ13, Q9DB77, P62897, Q9CQ75 | 3.101481401 | 0.02776 |
| Annotation Cluster 24 | Enrichment Score: 2.8628092061038877 | | | | | |
| Category | Term | Count | PValue | Genes | Fold Enrichment | FDR |
| GOTERM_BP_DIRECT | GO:0045047~protein targeting to ER | 8 | 4.2E-07 | Q91X58, P16254, Q6ZWQ7, Q9CYN2, P47758, Q9DBG7, P14576 | 15.0345351 | 9.2E-05 |
| GOTERM_BP_DIRECT | GO:0006614~SRP-dependent cotranslational protein targeting to membrane | 6 | 3.1E-05 | P16254, P61620, Q9D7A6, Q9DBG7, P14576 | 14.74540943 | 0.00325 |
| Annotation Cluster 25 | Enrichment Score: 2.7707820188280143 | | | | | |
| Category | Term | Count | PValue | Genes | Fold Enrichment | FDR |
| GOTERM_BP_DIRECT | GO:0006099~tricarboxylic acid cycle | 7 | 0.00049 | Q9D6R2, Q9CQA3, Q9WUM5, Q9Z2I8, Q8BMF4, P08249, P54071 | 6.776930596 | 0.0264 |
| KEGG_PATHWAY | mmu00020:Citrate cycle (TCA cycle) | 9 | 0.00066 | Q9D6R2, Q9CQA3, O08749, Q9WUM5, Q9Z2I8, Q8BMF4, Q8BH04, P08249, P54071 | 4.446539256 | 0.00762 |
| UP_KW_BIOLOGICAL_PROCESS | KW-0816~Tricarboxylic acid cycle | 7 | 0.00138 | Q9D6R2, Q9CQA3, Q9WUM5, Q9Z2I8, Q8BMF4, P08249, P54071 | 5.467796841 | 0.01068 |
| Annotation Cluster 26 | Enrichment Score: 2.728723874464465 | | | | | |
| Category | Term | Count | PValue | Genes | Fold Enrichment | FDR |
| GOTERM_BP_DIRECT | GO:0008380~RNA splicing | 25 | 2.2E-08 | Q921F2, Q9CZU3, O35326, Q52KI8, Q5SUF2, Q9JIX8, P62317, P28659, Q8VE97, Q9CWX9, P29341, Q9CYA6, O35286, Q8VDP4, P35922, O08810, Q6PE01, Q3UEB3, Q91VC3, Q05CL8, P62322, O08583, Q3U0V1, P62307, P63017 | 3.973679987 | 7.8E-06 |

| | | | | | | |
|---|---|---|---|---|---|---|
| UP_KW_BIOLOGICAL_PROCESS | KW-0508~mRNA splicing | 35 | 1.8E-06 | Q921F2, Q9CZU3, Q501J6, P57784, O35326, Q6ZWM4, P62311, Q52KI8, Q5SUF2, Q9JIX8, P62317, P28659, Q9R0U0, Q8VE97, Q8BHS3, Q3UYV9, Q9CWX9, P29341, Q9CYA6, O35286, Q8VDP4, P35922, O08810, Q6PE01, Q3UEB3, Q91VC3, Q05CL8, P62320, P62322, O08583, Q3U0V1, Q7TNC4, P62307, Q9D0E1, P63017 | 2.468737704 | 4.2E-05 |
| GOTERM_BP_DIRECT | GO:0006397~mRNA processing | 24 | 6.9E-06 | Q921F2, Q8VE97, Q9CZU3, Q9CWX9, P29341, O35286, Q8VDP4, P35922, O08810, Q3UEB3, Q6PE01, Q80X82, Q05CL8, O35326, P62322, O08583, Q3U0V1, Q99MR6, Q52KI8, P62317, Q9JIX8, P28659, Q9D0E1, P05132 | 2.983506966 | 0.00094 |
| GOTERM_CC_DIRECT | GO:0071013~catalytic step 2 spliceosome | 15 | 1.9E-05 | Q8BHS3, Q9CZU3, P29341, Q9CYA6, P57784, O08810, Q6PE01, Q91VC3, P62320, P62311, O08583, Q52KI8, P62317, P62307, Q9D0E1 | 4.075414781 | 0.00025 |
| UP_KW_BIOLOGICAL_PROCESS | KW-0507~mRNA processing | 37 | 4.6E-05 | Q921F2, Q9CZU3, Q501J6, P57784, Q80X82, O35326, Q6ZWM4, P62311, Q52KI8, Q5SUF2, Q9JIX8, P62317, P28659, Q9R0U0, Q8VE97, Q8BHS3, Q3UYV9, Q9CWX9, P29341, Q9CYA6, O35286, Q8VDP4, P35922, O08810, Q6PE01, Q3UEB3, Q91VC3, Q05CL8, P62320, P62322, O08583, Q3U0V1, Q7TNC4, P62307, Q9D0E1, P63017, Q8BIQ5 | 2.069848065 | 0.0006 |
| GOTERM_CC_DIRECT | GO:0046540~U4/U6 x U5 tri-snRNP complex | 8 | 0.00013 | P62320, Q6ZWM4, P62322, P62311, P62317, P62307, O08810, Q6PE01 | 6.873131955 | 0.00136 |
| GOTERM_CC_DIRECT | GO:0005681~spliceosomal complex | 15 | 0.00027 | P29341, O35286, P57784, O08810, Q91VC3, P62320, O35841, P62322, O08583, Q52KI8, P62317, P62307, Q9D0E1, Q921J2, P63017 | 3.200157916 | 0.0025 |
| GOTERM_BP_DIRECT | GO:0000398~mRNA splicing, via spliceosome | 16 | 0.00027 | Q9R0U0, Q3UYV9, Q9EPV8, P57784, O08810, Q91VC3, O35326, P62320, Q6ZWM4, P62322, P62311, Q8C2Q3, Q52KI8, P62317, P62307, P63017 | 3.042703533 | 0.0176 |
| UP_SEQ_FEATURE | DOMAIN:Sm | 7 | 0.00066 | P62320, Q9D0R8, Q6ZWM4, P62322, P62311, P62317, P62307 | 6.319053994 | 0.13154 |
| INTERPRO | IPR047575:Sm | 7 | 0.00079 | P62320, Q9D0R8, Q6ZWM4, P62322, P62311, P62317, P62307 | 6.110013717 | 0.09593 |
| SMART | SM00651:Sm | 6 | 0.00087 | P62320, Q6ZWM4, P62322, P62311, P62317, P62307 | 7.578254848 | 0.09172 |
| UP_KW_CELLULAR_COMPONENT | KW-0747~Spliceosome | 18 | 0.00092 | Q8BHS3, Q9CZU3, P29341, Q9CYA6, P57784, O08810, Q6PE01, Q91VC3, P62320, Q6ZWM4, P62322, P62311, O08583, Q52KI8, P62317, P62307, Q9D0E1, P63017 | 2.486975462 | 0.004 |
| INTERPRO | IPR001163:Sm_dom_euk/arc | 6 | 0.00096 | P62320, Q6ZWM4, P62322, P62311, P62317, P62307 | 7.442272348 | 0.09928 |
| INTERPRO | IPR010920:LSM_dom_sf | 6 | 0.00195 | P62320, Q6ZWM4, P62322, P62311, P62317, P62307 | 6.427417027 | 0.16818 |
| GOTERM_CC_DIRECT | GO:0005685~U1 snRNP | 5 | 0.00258 | P62320, Q7TNC4, Q5SUF2, P62317, P62307 | 8.365325077 | 0.01678 |
| GOTERM_CC_DIRECT | GO:0071005~U2-type precatalytic spliceosome | 8 | 0.00268 | P62320, Q6ZWM4, P62322, P62311, P57784, P62317, P62307, O08810 | 4.238431373 | 0.01715 |
| GOTERM_CC_DIRECT | GO:0071007~U2-type catalytic step 2 spliceosome | 7 | 0.00339 | P62320, Q8BHS3, P57784, P62317, P62307, O08810, Q6PE01 | 4.734418023 | 0.02069 |
| GOTERM_CC_DIRECT | GO:0034715~pICln-Sm protein complex | 3 | 0.01865 | P62320, P62317, P62307 | 13.62352941 | 0.08233 |
| GOTERM_CC_DIRECT | GO:0120115~Lsm2-8 complex | 3 | 0.01865 | Q6ZWM4, P62322, P62311 | 13.62352941 | 0.08233 |
| GOTERM_CC_DIRECT | GO:0071011~precatalytic spliceosome | 4 | 0.02059 | P62320, Q6ZWM4, P62311, P62317 | 6.692260062 | 0.08912 |
| GOTERM_CC_DIRECT | GO:0005688~U6 snRNP | 3 | 0.02435 | Q6ZWM4, P62322, P62311 | 11.92058824 | 0.1015 |
| GOTERM_CC_DIRECT | GO:0005687~U4 snRNP | 3 | 0.04495 | P62320, P62317, P62307 | 8.669518717 | 0.16483 |

| | | | | | | |
|---|---|---|---|---|---|---|
| GOTERM_CC_DIRECT | GO:0005689~U12-type spliceosomal complex | 4 | 0.0566 | P62320, O35286, P62317, P62307 | 4.541176471 | 0.19678 |
| Annotation Cluster 27 | Enrichment Score: 2.556803035807914 | | | | | |
| Category | Term | Count | PValue | Genes | Fold Enrichment | FDR |
| GOTERM_BP_DIRECT | GO:0006605~protein targeting | 10 | 8.9E-06 | Q80TH2, O55101, Q9CQV8, Q7TN05, O54774, Q60953, Q3UVL4, P68254, P61982, P62259 | 7.099641577 | 0.00115 |
| UP_SEQ_FEATURE | SITE:Interaction with phosphoserine on interacting protein | 5 | 8.9E-05 | Q9CQV8, P68510, P68254, P61982, P62259 | 17.40963855 | 0.02259 |
| INTERPRO | IPR023409:14-3-3_CS | 5 | 0.0001 | Q9CQV8, P68510, P68254, P61982, P62259 | 16.83371126 | 0.02699 |
| SMART | SM00101:14_3_3 | 5 | 0.00018 | Q9CQV8, P68510, P68254, P61982, P62259 | 14.99862939 | 0.02963 |
| INTERPRO | IPR000308:14-3-3 | 5 | 0.0002 | Q9CQV8, P68510, P68254, P61982, P62259 | 14.72949735 | 0.04098 |
| INTERPRO | IPR023410:14-3-3_domain | 5 | 0.0002 | Q9CQV8, P68510, P68254, P61982, P62259 | 14.72949735 | 0.04098 |
| INTERPRO | IPR036815:14-3-3_dom_sf | 5 | 0.00034 | Q9CQV8, P68510, P68254, P61982, P62259 | 13.09288654 | 0.05871 |
| PIR_SUPERFAMILY | PIRSF000868:14-3-3 | 5 | 0.00077 | Q9CQV8, P68510, P68254, P61982, P62259 | 9.834791059 | 0.09738 |
| GOTERM_MF_DIRECT | GO:0140311~protein sequestering activity | 6 | 0.00471 | Q64337, Q9CQV8, O08992, P61982, Q9D0R4, P62259 | 5.389565217 | 0.07479 |
| Annotation Cluster 28 | Enrichment Score: 2.482997644631693 | | | | | |
| Category | Term | Count | PValue | Genes | Fold Enrichment | FDR |
| GOTERM_CC_DIRECT | GO:0005905~clathrin-coated pit | 10 | 0.00024 | Q7TN05, Q91ZX7, Q7TQH7, Q6PIU9, O08585, Q80TY0, P35951, P22892, P63038, P62743 | 4.74451273 | 0.0023 |
| UP_KW_CELLULAR_COMPONENT | KW-0168~Coated pit | 9 | 0.00336 | Q7TN05, Q91ZX7, Q7TQH7, Q6PIU9, O08585, Q80TY0, P35951, P22892, P62743 | 3.566228965 | 0.01191 |
| UP_KW_BIOLOGICAL_PROCESS | KW-0254~Endocytosis | 12 | 0.04409 | Q99KN9, Q91ZX7, Q7TQH7, Q6PIU9, Q9D8U8, Q6P9Q6, Q80TY0, Q8BIJ7, P35951, Q9CQD1, O08539, P62743 | 1.946776018 | 0.1705 |
| Annotation Cluster 29 | Enrichment Score: 2.430738991593397 | | | | | |
| Category | Term | Count | PValue | Genes | Fold Enrichment | FDR |
| UP_KW_MOLECULAR_FUNCTION | KW-0030~Aminoacyl-tRNA synthetase | 11 | 1.6E-05 | P32921, Q8CGC7, Q9WUA2, Q8BP47, Q9CZD3, Q8BMJ2, P26638, Q8BYL4, Q8BU30, Q8BGQ7, Q9D0R2 | 5.611288605 | 0.00015 |
| UP_KW_MOLECULAR_FUNCTION | KW-0436~Ligase | 19 | 0.00024 | P32921, Q61510, Q9CZW4, Q9WUM5, Q9Z2I8, Q8BYL4, Q61024, Q5SUR0, Q9D0R2, Q8CGC7, Q9WUA2, Q8BP47, Q9CZD3, Q8BMJ2, Q9QUJ7, P26638, Q8BU30, Q9QXG4, Q8BGQ7 | 2.680828405 | 0.002 |
| INTERPRO | IPR045864:aa-tRNA-synth_II/BPL/LPL | 7 | 0.00051 | Q8CGC7, Q9WUA2, Q8BP47, Q9CZD3, P26638, Q8BGQ7, Q9D0R2 | 6.598814815 | 0.07794 |
| GOTERM_BP_DIRECT | GO:0006418~tRNA aminoacylation for protein translation | 5 | 0.00128 | Q8CGC7, Q8BP47, Q9CZD3, Q8BYL4, Q8BU30 | 9.983870968 | 0.05307 |
| Annotation Cluster 30 | Enrichment Score: 2.4152115941926757 | | | | | |
| Category | Term | Count | PValue | Genes | Fold Enrichment | FDR |

| | | | | | | |
|---|---|---|---|---|---|---|
| UP_KW_BIOLOGICAL_PROCESS | KW-0028~Amino-acid biosynthesis | 7 | 0.00112 | O35855, Q99K85, Q9Z110, P24288, Q8VCN5, Q61024, P05201 | 5.678096719 | 0.00926 |
| KEGG_PATHWAY | mmu01230:Biosynthesis of amino acids | 14 | 0.00124 | Q99K85, Q9D6R2, Q99JW2, P17751, Q61024, Q8VCN5, O35855, Q9CZN7, P05063, P05064, Q9Z110, P24288, P54071, P05201 | 2.801757506 | 0.01191 |
| KEGG_PATHWAY | mmu00270:Cysteine and methionine metabolism | 11 | 0.00206 | O88508, O35855, Q99K85, P97355, O09172, P24288, P08249, Q8VCN5, P05201, P50247, Q99J99 | 3.161983471 | 0.01744 |
| UP_KW_MOLECULAR_FUNCTION | KW-0032~Aminotransferase | 5 | 0.01601 | O35855, Q99K85, P24288, P47856, P05201 | 4.973642173 | 0.07505 |
| UP_KW_LIGAND | KW-0663~Pyridoxal phosphate | 9 | 0.01836 | O35855, Q99K85, Q9CZN7, O88986, Q8R0X7, Q9JLI6, P24288, Q8VCN5, P05201 | 2.656584094 | 0.09182 |
| Annotation Cluster 31 | Enrichment Score: 2.413177829062143 | | | | | |
| Category | Term | Count | PValue | Genes | Fold Enrichment | FDR |
| GOTERM_BP_DIRECT | GO:1903232~melanosome assembly | 6 | 0.00024 | O35643, Q7TN05, O54774, P22892, P35585, P59438 | 10.08896435 | 0.01661 |
| GOTERM_CC_DIRECT | GO:0032588~trans-Golgi network membrane | 9 | 0.0003 | O35643, Q7TN05, Q9JLQ0, Q921E2, O35609, P22892, P35585, Q64430, Q8BG07 | 5.20171123 | 0.00275 |
| GOTERM_BP_DIRECT | GO:0060155~platelet dense granule organization | 6 | 0.0004 | O35643, Q7TN05, O54774, P22892, P35585, P59438 | 9.128110599 | 0.02239 |
| GOTERM_CC_DIRECT | GO:0030121~AP-1 adaptor complex | 4 | 0.00422 | O35643, Q7TN05, P22892, P35585 | 11.55935829 | 0.02444 |
| Annotation Cluster 32 | Enrichment Score: 2.3630332056771612 | | | | | |
| Category | Term | Count | PValue | Genes | Fold Enrichment | FDR |
| GOTERM_BP_DIRECT | GO:0042776~proton motive force-driven mitochondrial ATP synthesis | 11 | 2.8E-05 | Q9DC69, Q9CQA3, P03921, Q9CQQ7, Q9ERS2, Q9DCJ5, P56480, Q91VR2, Q9DB20, Q99JB2, Q9CQ75 | 5.491129032 | 0.00303 |
| GOTERM_BP_DIRECT | GO:0009060~aerobic respiration | 9 | 0.00198 | Q9DC69, Q9CQA3, P24270, P03921, Q9ERS2, Q9DCJ5, P08249, Q9CQ75 | 3.938842245 | 0.06894 |
| UP_KW_BIOLOGICAL_PROCESS | KW-0679~Respiratory chain | 10 | 0.00455 | Q9D0M3, Q9DC69, Q9CR68, P03921, Q9ERS2, Q9DCJ5, Q9CZ13, Q9DB77, P62897, Q9CQ75 | 3.101481401 | 0.02776 |
| Annotation Cluster 33 | Enrichment Score: 2.327825732384294 | | | | | |
| Category | Term | Count | PValue | Genes | Fold Enrichment | FDR |
| UP_KW_BIOLOGICAL_PROCESS | KW-0752~Steroid biosynthesis | 11 | 2.4E-05 | Q9DCN2, Q9D0S9, Q8BLN5, O88736, P50171, Q3U9G9, Q9DB73, O88455, Q9D1G2, Q9R1J0, O70503 | 5.395135089 | 0.00035 |
| GOTERM_BP_DIRECT | GO:0006695~cholesterol biosynthetic process | 8 | 0.00025 | Q9DCN2, Q8BLN5, O88736, Q00612, Q3U9G9, O88455, Q9D1G2, Q9R1J0 | 6.233831629 | 0.01705 |
| UP_KW_BIOLOGICAL_PROCESS | KW-1207~Sterol metabolism | 10 | 0.00455 | Q8VBT6, Q9DCN2, Q9Z0J0, Q3U9G9, P35951, Q9DB73, O88455, P17439, Q9D1G2, Q9R1J0 | 3.101481401 | 0.02776 |
| UP_KW_BIOLOGICAL_PROCESS | KW-0153~Cholesterol metabolism | 9 | 0.00772 | Q8VBT6, Q9DCN2, Q9Z0J0, Q3U9G9, P35951, O88455, P17439, Q9D1G2, Q9R1J0 | 3.111650193 | 0.04375 |
| UP_KW_BIOLOGICAL_PROCESS | KW-0756~Sterol biosynthesis | 6 | 0.00792 | Q9DCN2, Q3U9G9, Q9DB73, O88455, Q9D1G2, Q9R1J0 | 4.686683007 | 0.04375 |
| UP_KW_BIOLOGICAL_PROCESS | KW-0152~Cholesterol biosynthesis | 5 | 0.01562 | Q9DCN2, Q3U9G9, O88455, Q9D1G2, Q9R1J0 | 5.021446078 | 0.0788 |

| | | | | | | |
|---|---|---|---|---|---|---|
| UP_KW_BIOLOGICAL_PROCESS | KW-0753~Steroid metabolism | 10 | 0.03161 | Q8VBT6, Q9DCN2, Q9Z0J0, Q3U9G9, P35951, Q9DB73, O88455, P17439, Q9D1G2, Q9R1J0 | 2.267749842 | 0.13094 |
| KEGG_PATHWAY | mmu00100:Steroid biosynthesis | 5 | 0.03395 | Q8BLN5, O88736, Q3U9G9, O88455, Q9R1J0 | 3.952479339 | 0.14139 |
| Annotation Cluster 34 | Enrichment Score: 2.244173304316446 | | | | | |
| Category | Term | Count | PValue | Genes | Fold Enrichment | FDR |
| INTERPRO | IPR011012:Longin-like_dom_sf | 7 | 0.00096 | P70280, Q7TN05, P61924, P35585, Q9DBG7, Q5XJY5, P62743 | 5.891798942 | 0.09928 |
| Annotation Cluster 35 | Enrichment Score: 2.2232819282747704 | | | | | |
| Category | Term | Count | PValue | Genes | Fold Enrichment | FDR |
| GOTERM_MF_DIRECT | GO:0140662~ATP-dependent protein folding chaperone | 10 | 2.5E-07 | P80314, Q61699, P80313, Q9CQN1, Q9JKR6, P38647, P08113, P11983, P63038, P63017 | 10.47971014 | 1.8E-05 |
| Annotation Cluster 36 | Enrichment Score: 2.078362214028732 | | | | | |
| Category | Term | Count | PValue | Genes | Fold Enrichment | FDR |
| INTERPRO | IPR017853:Glycoside_hydrolase_SF | 10 | 0.00047 | P23780, Q9QWR8, P20060, P51569, P10852, P17439, P70699, P29416, P12265, Q8BHN3 | 4.284944685 | 0.07608 |
| GOTERM_BP_DIRECT | GO:0019915~lipid storage | 6 | 0.00499 | P20060, P51569, P17439, P43883, P29416, Q9DBG5 | 5.324731183 | 0.12097 |
| KEGG_PATHWAY | mmu00511:Other glycan degradation | 5 | 0.02369 | O09159, P23780, P20060, P17439, P29416 | 4.39164371 | 0.10491 |
| Annotation Cluster 37 | Enrichment Score: 2.0574225445819216 | | | | | |
| Category | Term | Count | PValue | Genes | Fold Enrichment | FDR |
| UP_SEQ_FEATURE | MOTIF:Prevents secretion from ER | 9 | 0.00541 | P57759, Q8BP92, Q9JKR6, P08003, P09103, P08113, Q9R1J0, Q8K297, Q6P5E4 | 3.323658269 | 0.42627 |
| GOTERM_CC_DIRECT | GO:0034663~endoplasmic reticulum chaperone complex | 4 | 0.0055 | Q9JKR6, P08003, P09103, P08113 | 10.59607843 | 0.02968 |
| GOTERM_CC_DIRECT | GO:0005790~smooth endoplasmic reticulum | 5 | 0.01386 | P57759, Q9JKR6, P08003, P08113, P35564 | 5.298039216 | 0.06415 |
| GOTERM_CC_DIRECT | GO:0005788~endoplasmic reticulum lumen | 10 | 0.01427 | P57759, Q8BP92, Q9R0E1, Q9JKR6, P08003, P09103, P08113, Q8K297, Q9DBS1, Q6P5E4 | 2.627126884 | 0.06557 |
| Annotation Cluster 38 | Enrichment Score: 2.042101983907897 | | | | | |
| Category | Term | Count | PValue | Genes | Fold Enrichment | FDR |
| GOTERM_CC_DIRECT | GO:0110158~calpain complex | 3 | 0.00567 | O88456, O08529, O35350 | 23.84117647 | 0.02998 |
| Annotation Cluster 39 | Enrichment Score: 2.0131373360585396 | | | | | |
| Category | Term | Count | PValue | Genes | Fold Enrichment | FDR |

| | | | | | | |
|---|---|---|---|---|---|---|
| GOTERM_BP_DIRECT | GO:0006164~purine nucleotide biosynthetic process | 7 | 3.3E-05 | P00493, Q8VI93, P11928, E9Q9A9, Q8R574, P24547, Q5SUR0 | 10.64946237 | 0.00342 |
| GOTERM_BP_DIRECT | GO:0140374~antiviral innate immune response | 11 | 6.2E-05 | Q8VI94, Q61510, Q9QZI9, P67778, O35129, Q8VI93, P11928, E9Q9A9, O35286, P14206, Q6Q899 | 5.020460829 | 0.00573 |
| GOTERM_BP_DIRECT | GO:0045071~negative regulation of viral genome replication | 8 | 0.00157 | Q8VI94, Q9JL16, Q8VI93, P11928, E9Q9A9, O54962, Q9JJ00, D0QMC3 | 4.647038123 | 0.05947 |
| GOTERM_BP_DIRECT | GO:0070106~interleukin-27-mediated signaling pathway | 4 | 0.01046 | Q8VI94, Q8VI93, P11928, E9Q9A9 | 8.519569892 | 0.19308 |
| Annotation Cluster 40 | Enrichment Score: 1.953020358411172 | | | | | |
| Category | Term | Count | PValue | Genes | Fold Enrichment | FDR |
| GOTERM_MF_DIRECT | GO:0030170~pyridoxal phosphate binding | 8 | 0.00154 | Q99K85, Q9CZN7, O88986, Q8R0X7, Q8K183, Q922Q1, Q8VCN5, P05201 | 4.657648953 | 0.03222 |
| UP_KW_LIGAND | KW-0663~Pyridoxal phosphate | 9 | 0.01836 | O35855, Q99K85, Q9CZN7, O88986, Q8R0X7, Q9JLI6, P24288, Q8VCN5, P05201 | 2.656584094 | 0.09182 |
| Annotation Cluster 41 | Enrichment Score: 1.9180632659405008 | | | | | |
| Category | Term | Count | PValue | Genes | Fold Enrichment | FDR |
| GOTERM_MF_DIRECT | GO:0017154~semaphorin receptor activity | 4 | 0.00895 | P97333, O35375, Q99NH8, P70206 | 8.982608696 | 0.12817 |
| GOTERM_BP_DIRECT | GO:0071526~semaphorin-plexin signaling pathway | 6 | 0.01065 | P63001, P97333, O35375, P55194, Q99NH8, P70206 | 4.457914479 | 0.19308 |
| Annotation Cluster 42 | Enrichment Score: 1.9113233409524002 | | | | | |
| Category | Term | Count | PValue | Genes | Fold Enrichment | FDR |
| KEGG_PATHWAY | mmu00531:Glycosaminoglycan degradation | 7 | 0.00148 | Q571E4, Q8BFR4, P23780, P50429, P20060, P29416, P12265 | 5.269972452 | 0.01332 |
| GOTERM_MF_DIRECT | GO:0008484~sulfuric ester hydrolase activity | 4 | 0.00436 | Q571E4, Q8BFR4, P50429, P50428 | 11.43241107 | 0.07085 |
| COG_ONTOLOGY | Inorganic ion transport and metabolism | 7 | 0.0087 | Q8BFR4, P37040, P09671, P24270, P50429, P50428, Q99J99 | 3.738676845 | 0.13047 |
| Annotation Cluster 43 | Enrichment Score: 1.8832143687928948 | | | | | |
| Category | Term | Count | PValue | Genes | Fold Enrichment | FDR |
| GOTERM_BP_DIRECT | GO:0006122~mitochondrial electron transport, ubiquinol to cytochrome c | 5 | 0.00054 | Q9D0M3, Q9CR68, Q9CZ13, Q9DB77, P62897 | 12.28784119 | 0.02779 |
| UP_KW_BIOLOGICAL_PROCESS | KW-0679~Respiratory chain | 10 | 0.00455 | Q9D0M3, Q9DC69, Q9CR68, P03921, Q9ERS2, Q9DCJ5, Q9CZ13, Q9DB77, P62897, Q9CQ75 | 3.101481401 | 0.02776 |
| GOTERM_CC_DIRECT | GO:0045275~respiratory chain complex III | 4 | 0.00699 | Q9D0M3, Q9CR68, Q9CZ13, Q9DB77 | 9.780995475 | 0.03579 |
| Annotation Cluster 44 | Enrichment Score: 1.8812959301122532 | | | | | |

| Category | Term | Count | PValue | Genes | Fold Enrichment | FDR |
|---|---|---|---|---|---|---|
| GOTERM_BP_DIRECT | GO:0006413~translational initiation | 10 | 0.00011 | Q62448, Q9Z0N1, P61222, Q99LD9, Q61749, Q9Z1D1, Q99L45, Q8QZY1, Q8R1B4, Q6P542 | 5.237440508 | 0.00847 |
| GOTERM_MF_DIRECT | GO:0003743~translation initiation factor activity | 9 | 0.00138 | Q62448, Q9Z0N1, Q99LD9, Q61749, Q9CQJ6, Q9Z1D1, Q99L45, Q8QZY1, Q8R1B4 | 4.161061381 | 0.03192 |
| GOTERM_BP_DIRECT | GO:0002183~cytoplasmic translational initiation | 4 | 0.00417 | Q9Z0N1, Q99LD9, Q61749, Q99L45 | 11.61759531 | 0.109 |
| UP_KW_MOLECULAR_FUNCTION | KW-0396~Initiation factor | 9 | 0.00462 | Q62448, Q9Z0N1, Q99LD9, Q61749, Q9CQJ6, Q9Z1D1, Q99L45, Q8QZY1, Q8R1B4 | 3.378322985 | 0.02663 |
| Annotation Cluster 45 | Enrichment Score: 1.8236640813477663 | | | | | |
| Category | Term | Count | PValue | Genes | Fold Enrichment | FDR |
| GOTERM_MF_DIRECT | GO:0005509~calcium ion binding | 40 | 0.00105 | Q07076, Q7TPR4, Q8BP92, Q91ZX7, Q9QXZ0, Q8BYI6, P35951, P34960, O08529, P11688, O35350, P11881, Q3U7R1, Q61233, P07091, P11152, P43274, O88456, Q60605, Q02819, P47713, P56480, P63028, Q99K51, Q64518, O70200, Q62393, P35564, P50428, Q9EQP2, P50543, Q9R0Q8, P14069, O55143, P57780, Q9JJ00, O35474, Q05910, P22682, O35639 | 1.734572714 | 0.02572 |
| Annotation Cluster 46 | Enrichment Score: 1.7510345638604603 | | | | | |
| Category | Term | Count | PValue | Genes | Fold Enrichment | FDR |
| GOTERM_MF_DIRECT | GO:0050660~flavin adenine dinucleotide binding | 9 | 0.002 | Q9DCN2, P37040, Q61093, P47791, Q9JMH6, O08749, P45952, Q8JZN5, Q00519 | 3.929891304 | 0.03849 |
| GOTERM_MF_DIRECT | GO:0071949~FAD binding | 6 | 0.01497 | Q9DCN2, Q61093, Q9JMH6, Q9DB73, Q9JI75, Q00519 | 4.100756144 | 0.18953 |
| UP_KW_LIGAND | KW-0274~FAD | 13 | 0.04845 | P47791, Q9JMH6, O08749, Q8JZN5, Q9DCN2, Q9DC69, P37040, Q61093, Q64FW2, Q9DB73, Q9JI75, P45952, Q00519 | 1.840650407 | 0.16151 |
| UP_KW_LIGAND | KW-0285~Flavoprotein | 13 | 0.06823 | P47791, Q9JMH6, O08749, Q8JZN5, Q9DCN2, Q9DC69, P37040, Q61093, Q64FW2, Q9DB73, Q9JI75, P45952, Q00519 | 1.741538462 | 0.18608 |
| Annotation Cluster 47 | Enrichment Score: 1.7464143993863241 | | | | | |
| Category | Term | Count | PValue | Genes | Fold Enrichment | FDR |
| GOTERM_BP_DIRECT | GO:0014823~response to activity | 8 | 0.00377 | P09671, P24270, O09172, Q9CZ13, P22682, P11440, P63017 | 3.993548387 | 0.10429 |
| Annotation Cluster 48 | Enrichment Score: 1.7092501018004969 | | | | | |
| Category | Term | Count | PValue | Genes | Fold Enrichment | FDR |
| GOTERM_BP_DIRECT | GO:0019430~removal of superoxide radicals | 5 | 0.00163 | P09671, Q99LX0, Q9WU84, Q61171, Q64430 | 9.39658444 | 0.05947 |

| | | | | | | |
|---|---|---|---|---|---|---|
| GOTERM_MF_DIRECT | GO:0016532~superoxide dismutase copper chaperone activity | 3 | 0.00296 | Q99LX0, Q9WU84, Q64430 | 31.43913043 | 0.05132 |
| Annotation Cluster 49 | Enrichment Score: 1.6883330826089125 | | | | | |
| Category | Term | Count | PValue | Genes | Fold Enrichment | FDR |
| GOTERM_CC_DIRECT | GO:0030126~COPI vesicle coat | 4 | 0.0055 | Q3USZ8, Q8CIE6, P61924, Q5XJY5 | 10.59607843 | 0.02968 |
| Annotation Cluster 50 | Enrichment Score: 1.6788183155444627 | | | | | |
| Category | Term | Count | PValue | Genes | Fold Enrichment | FDR |
| GOTERM_BP_DIRECT | GO:1904851~positive regulation of establishment of protein localization to telomere | 4 | 0.0031 | P80314, P80313, P11983, Q9ESX5 | 12.77935484 | 0.09091 |
| GOTERM_BP_DIRECT | GO:1904874~positive regulation of telomerase RNA localization to Cajal body | 3 | 0.00562 | P80314, P11983, Q9ESX5 | 23.96129032 | 0.12737 |
| GOTERM_BP_DIRECT | GO:0090666~scaRNA localization to Cajal body | 3 | 0.00917 | P80314, P11983, Q9ESX5 | 19.16903226 | 0.17614 |
| GOTERM_CC_DIRECT | GO:0005832~chaperonin-containing T-complex | 3 | 0.03755 | P80314, P80313, P11983 | 9.536470588 | 0.14154 |
| Annotation Cluster 51 | Enrichment Score: 1.67129149155184 | | | | | |
| Category | Term | Count | PValue | Genes | Fold Enrichment | FDR |
| KEGG_PATHWAY | mmu00531:Glycosaminoglycan degradation | 7 | 0.00148 | Q571E4, Q8BFR4, P23780, P50429, P20060, P29416, P12265 | 5.269972452 | 0.01332 |
| KEGG_PATHWAY | mmu00511:Other glycan degradation | 5 | 0.02369 | O09159, P23780, P20060, P17439, P29416 | 4.39164371 | 0.10491 |
| Annotation Cluster 52 | Enrichment Score: 1.6676318875553544 | | | | | |
| Category | Term | Count | PValue | Genes | Fold Enrichment | FDR |
| GOTERM_BP_DIRECT | GO:0032727~positive regulation of interferon-alpha production | 5 | 0.0107 | P63158, P58681, P34960, Q6Q899, P63038 | 5.705069124 | 0.19308 |
| Annotation Cluster 53 | Enrichment Score: 1.6525616868530644 | | | | | |
| Category | Term | Count | PValue | Genes | Fold Enrichment | FDR |
| GOTERM_BP_DIRECT | GO:0051017~actin filament bundle assembly | 8 | 0.0003 | Q99K51, Q7TPR4, Q8VDD5, P57780, Q61879, O70200, Q61233, P26645 | 6.085407066 | 0.01805 |
| INTERPRO | IPR001589:Actinin_actin-bd_CS | 6 | 0.00241 | Q99K51, Q80X90, Q7TPR4, Q9QXZ0, P57780, Q61233 | 6.147964113 | 0.19065 |
| GOTERM_CC_DIRECT | GO:0001725~stress fiber | 10 | 0.00553 | O55131, Q80X90, Q7TPR4, Q8VDD5, Q8CI51, P57780, Q61879, Q8CBY8, Q6IRU2, Q61233 | 3.056561086 | 0.02968 |

| | | | | | | |
|---|---|---|---|---|---|---|
| Annotation Cluster 54 | Enrichment Score: 1.6412086111288473 | | | | | |
| Category | Term | Count | PValue | Genes | Fold Enrichment | FDR |
| KEGG_PATHWAY | mmu00520:Amino sugar and nucleotide sugar metabolism | 10 | 0.00058 | P17710, O88958, P20060, Q924M7, Q9QZ08, Q9R0N0, O08528, P47856, P29416, Q9D0F9 | 4.053824963 | 0.00698 |
| GOTERM_BP_DIRECT | GO:0006002~fructose 6-phosphate metabolic process | 4 | 0.01046 | P17710, O88958, O08528, P47856 | 8.519569892 | 0.19308 |
| KEGG_PATHWAY | mmu00052:Galactose metabolism | 7 | 0.01369 | P17710, P23780, P51569, Q9R0N0, O08528, P70699, Q9D0F9 | 3.458419421 | 0.06945 |
| KEGG_PATHWAY | mmu01250:Biosynthesis of nucleotide sugars | 7 | 0.03034 | P17710, Q924M7, Q9QZ08, Q9R0N0, O08528, P47856, Q9D0F9 | 2.912353197 | 0.13023 |
| Annotation Cluster 55 | Enrichment Score: 1.6269166498380503 | | | | | |
| Category | Term | Count | PValue | Genes | Fold Enrichment | FDR |
| GOTERM_BP_DIRECT | GO:0006401~RNA catabolic process | 6 | 0.00188 | Q9JL16, Q9DAA6, E9Q9A9, Q3UYV9, Q9CZU3, Q8BTW3 | 6.610011123 | 0.06665 |
| Annotation Cluster 56 | Enrichment Score: 1.6252649520246683 | | | | | |
| Category | Term | Count | PValue | Genes | Fold Enrichment | FDR |
| INTERPRO | IPR009000:Transl_B-barrel_sf | 8 | 0.00032 | O55142, Q8R050, Q9Z0N1, P58252, P10126, P27659, O08810, Q8BGQ7 | 5.891798942 | 0.05845 |
| INTERPRO | IPR004161:EFTu-like_2 | 5 | 0.00228 | Q8R050, Q9Z0N1, P58252, P10126, O08810 | 8.416855631 | 0.18581 |
| GOTERM_MF_DIRECT | GO:0003746~translation elongation factor activity | 5 | 0.00648 | P58252, P10126, O70251, P57776, O08810 | 6.549818841 | 0.09922 |
| Annotation Cluster 57 | Enrichment Score: 1.6027871300450933 | | | | | |
| Category | Term | Count | PValue | Genes | Fold Enrichment | FDR |
| GOTERM_CC_DIRECT | GO:0000792~heterochromatin | 9 | 0.00096 | Q61686, O88508, Q9Z2D8, P23198, P83917, Q8VDF2, P43274, P05064, P11983 | 4.401447964 | 0.00737 |
| GOTERM_CC_DIRECT | GO:0090734~site of DNA damage | 7 | 0.00189 | Q61686, P23198, P83917, Q8VEG4, Q8CFE2, Q8CB77, Q6PDQ2 | 5.298039216 | 0.01321 |
| Annotation Cluster 58 | Enrichment Score: 1.5283038741031436 | | | | | |
| Category | Term | Count | PValue | Genes | Fold Enrichment | FDR |
| GOTERM_BP_DIRECT | GO:0032722~positive regulation of chemokine production | 7 | 0.00662 | P11152, P58681, Q9QUN7, Q9Z0F8, O70200, P09581, Q9EPB4 | 4.141457587 | 0.14586 |
| GOTERM_BP_DIRECT | GO:0002224~toll-like receptor signaling pathway | 5 | 0.0107 | P58681, Q9QUN7, Q8VCW4, Q9Z0H7, P49138 | 5.705069124 | 0.19308 |
| Annotation Cluster 59 | Enrichment Score: 1.5048462005876884 | | | | | |

| Category | Term | Count | PValue | Genes | Fold Enrichment | FDR |
|---|---|---|---|---|---|---|
| GOTERM_BP_DIRECT | GO:0006281~DNA repair | 20 | 0.00218 | P23492, Q9WV85, Q99J62, Q60710, Q99LX0, Q9WTM5, Q80W54, P39749, Q9CQ71, P35922, Q61550, Q920B9, Q3UMU9, Q9EPU0, P63158, Q9JLV6, P60122, Q8VDF2, Q8CFE2, Q8CCJ3 | 2.195765436 | 0.07102 |
| Annotation Cluster 60 | Enrichment Score: 1.5014643276310315 | | | | | |
| Category | Term | Count | PValue | Genes | Fold Enrichment | FDR |
| UP_KW_MOLECULAR_FUNCTION | KW-0788~Thiol protease | 17 | 0.00288 | P10605, Q8R5H1, B2RRE7, Q9JMA1, P56399, Q9JKB1, O08529, O70370, P97821, O35350, Q9WUU7, Q99K23, Q9EP97, Q8BY87, O89017, Q8R016, O89110 | 2.300732434 | 0.01797 |
| GOTERM_MF_DIRECT | GO:0004197~cysteine-type endopeptidase activity | 8 | 0.00377 | P10605, Q8R5H1, O70370, O89017, P97821, Q8R016, Q9WUU7, O89110 | 3.992270531 | 0.06279 |
| Annotation Cluster 61 | Enrichment Score: 1.4946926959206879 | | | | | |
| Category | Term | Count | PValue | Genes | Fold Enrichment | FDR |
| KEGG_PATHWAY | mmu03015:mRNA surveillance pathway | 14 | 0.0136 | Q921F2, Q61655, Q3UYV9, P29341, Q61151, Q80X82, Q91VC3, Q9EPU0, Q8R050, O08583, P63087, Q52KI8, Q9JIX8, Q8BIQ5 | 2.128258106 | 0.06945 |
| UP_KW_BIOLOGICAL_PROCESS | KW-0866~Nonsense-mediated mRNA decay | 5 | 0.06259 | Q9EPU0, Q8R050, Q3UYV9, P29341, Q91VC3 | 3.295323989 | 0.2342 |
| Annotation Cluster 62 | Enrichment Score: 1.483465388132444 | | | | | |
| Category | Term | Count | PValue | Genes | Fold Enrichment | FDR |
| GOTERM_CC_DIRECT | GO:0000793~condensed chromosome | 7 | 0.00146 | P63158, O54962, Q9Z277, Q61164, P30681, Q01320, E9PVX6 | 5.562941176 | 0.01076 |
| Annotation Cluster 63 | Enrichment Score: 1.455165395746659 | | | | | |
| Category | Term | Count | PValue | Genes | Fold Enrichment | FDR |
| GOTERM_BP_DIRECT | GO:0016255~attachment of GPI anchor to protein | 3 | 0.00917 | Q8BXQ2, Q9WTK3, Q6PD26 | 19.16903226 | 0.17614 |
| GOTERM_CC_DIRECT | GO:0042765~GPI-anchor transamidase complex | 3 | 0.00926 | Q8BXQ2, Q9WTK3, Q6PD26 | 19.07294118 | 0.04672 |
| Annotation Cluster 64 | Enrichment Score: 1.4299135333327888 | | | | | |
| Category | Term | Count | PValue | Genes | Fold Enrichment | FDR |
| GOTERM_MF_DIRECT | GO:0050660~flavin adenine dinucleotide binding | 9 | 0.002 | Q9DCN2, P37040, Q61093, P47791, Q9JMH6, O08749, P45952, Q8JZN5, Q00519 | 3.929891304 | 0.03849 |

| | | | | | | |
|---|---|---|---|---|---|---|
| Annotation Cluster 65 | Enrichment Score: 1.418086452836949 | | | | | |
| Category | Term | Count | PValue | Genes | Fold Enrichment | FDR |
| GOTERM_BP_DIRECT | GO:0061621~canonical glycolysis | 5 | 0.00253 | P17710, P05064, O08528, P59017, P17751 | 8.407470289 | 0.07888 |
| KEGG_PATHWAY | mmu00010:Glycolysis / Gluconeogenesis | 10 | 0.0242 | P17710, O08749, P05063, P05064, O08528, Q8BMF4, Q9D0F9, Q8BH04, P17751, Q9QXG4 | 2.359689158 | 0.10549 |
| Annotation Cluster 66 | Enrichment Score: 1.4166444555121378 | | | | | |
| Category | Term | Count | PValue | Genes | Fold Enrichment | FDR |
| KEGG_PATHWAY | mmu04664:Fc epsilon RI signaling pathway | 12 | 0.00259 | P63001, Q60787, P20491, O09110, Q9R0C8, P08556, P47713, P47811, P48999, Q9ES52 | 2.874530428 | 0.01986 |
| Annotation Cluster 67 | Enrichment Score: 1.4064752709245538 | | | | | |
| Category | Term | Count | PValue | Genes | Fold Enrichment | FDR |
| KEGG_PATHWAY | mmu05170:Human immunodeficiency virus 1 infection | 27 | 0.0051 | Q7TN05, Q60710, Q9R233, Q60591, P35585, P11881, P11440, P63001, O35643, P63213, Q9QUN7, P01902, O09110, P01900, P63328, P47941, P83940, P22892, P62869, P08556, P62897, O89110, P08752, P47811 | 1.771235554 | 0.0346 |
| KEGG_PATHWAY | mmu04218:Cellular senescence | 20 | 0.02207 | P51881, Q64337, Q60930, O08529, Q60591, O35350, P11881, P11440, P01902, O09110, P63087, P01900, P63328, P49138, Q921J2, P08556, P47811 | 1.72785982 | 0.10262 |
| Annotation Cluster 68 | Enrichment Score: 1.4012802532820392 | | | | | |
| Category | Term | Count | PValue | Genes | Fold Enrichment | FDR |
| GOTERM_BP_DIRECT | GO:0051058~negative regulation of small GTPase mediated signal transduction | 5 | 0.00054 | Q3TBD2, Q921M7, Q9JLQ0, Q3UIA2, P55194 | 12.28784119 | 0.02779 |
| GOTERM_MF_DIRECT | GO:0005096~GTPase activator activity | 17 | 0.00517 | P46061, P97333, Q5SSL4, Q69ZK0, Q3UIA2, Q60790, Q8C2K5, Q61210, Q6Y5D8, Q640N3, Q3TBD2, A3KGS3, P34022, Q9JLQ2, Q8BMJ2, Q8K1X4, P55194 | 2.190431219 | 0.08005 |
| Annotation Cluster 69 | Enrichment Score: 1.3820537141189329 | | | | | |
| Category | Term | Count | PValue | Genes | Fold Enrichment | FDR |
| GOTERM_MF_DIRECT | GO:0004128~cytochrome-b5 reductase activity, acting on NAD(P)H | 3 | 0.01389 | Q9DCN2, P37040, Q9DB73 | 15.71956522 | 0.17764 |
| GOTERM_MF_DIRECT | GO:0071949~FAD binding | 6 | 0.01497 | Q9DCN2, Q61093, Q9JMH6, Q9DB73, Q9JI75, Q00519 | 4.100756144 | 0.18953 |
| Annotation Cluster 70 | Enrichment Score: 1.30196011625609 | | | | | |
| Category | Term | Count | PValue | Genes | Fold Enrichment | FDR |

| | | | | | | |
|---|---|---|---|---|---|---|
| GOTERM_CC_DIRECT | GO:0005874~microtubule | 24 | 0.00059 | P80314, A2AQ07, Q61699, P80313, Q9QXZ0, Q8BVG4, Q922F4, O08539, P11983, P68372, P68373, Q61166, O08553, O88447, P62482, Q9EPB4, Q9JHU4, A2APB8, P48428, P27546, Q8C052, P63017, Q3TVI8, P26645 | 2.2242497 | 0.00496 |
| GOTERM_CC_DIRECT | GO:0015630~microtubule cytoskeleton | 14 | 0.03442 | A2AQ07, Q9QXZ0, Q922F4, P68372, P68373, P35922, Q61166, O55131, A2AGT5, Q9CZN7, A2APB8, P48428, P27546, Q8C052 | 1.901860231 | 0.13272 |
| Annotation Cluster 71 | Enrichment Score: 1.2963564869064537 | | | | | |
| Category | Term | Count | PValue | Genes | Fold Enrichment | FDR |
| UP_KW_LIGAND | KW-0001~2Fe-2S | 5 | 0.03411 | Q9CR68, Q9CQB5, Q9CQA3, Q8WTY4, Q00519 | 3.958041958 | 0.13861 |
| Annotation Cluster 72 | Enrichment Score: 1.2950374663179436 | | | | | |
| Category | Term | Count | PValue | Genes | Fold Enrichment | FDR |
| GOTERM_BP_DIRECT | GO:0008654~phospholipid biosynthetic process | 7 | 0.00331 | Q8BHF7, Q9DAV9, Q8BYI6, P20060, Q8BGS7, Q91V01, Q8VDP6 | 4.758270419 | 0.09485 |
| UP_KW_BIOLOGICAL_PROCESS | KW-1208~Phospholipid metabolism | 9 | 0.02 | Q8BHF7, Q8BYI6, Q8BGS7, Q99L43, P47713, Q91V01, O35405, Q8VDP6, Q8BG07 | 2.636259191 | 0.09668 |
| Annotation Cluster 73 | Enrichment Score: 1.2701781756421249 | | | | | |
| Category | Term | Count | PValue | Genes | Fold Enrichment | FDR |
| GOTERM_BP_DIRECT | GO:1900026~positive regulation of substrate adhesion-dependent cell spreading | 6 | 0.01065 | P63001, P97333, Q69ZK0, P47941, P09103, P60766 | 4.457914479 | 0.19308 |
| Annotation Cluster 74 | Enrichment Score: 1.2513495481399626 | | | | | |
| Category | Term | Count | PValue | Genes | Fold Enrichment | FDR |
| GOTERM_CC_DIRECT | GO:0044853~plasma membrane raft | 7 | 0.00214 | Q9DBC7, Q60787, O08917, P11835, Q99NH8, P05132, P12367 | 5.174829001 | 0.01451 |
| GOTERM_CC_DIRECT | GO:0005952~cAMP-dependent protein kinase complex | 4 | 0.00225 | Q9DBC7, P05132, P68181, P12367 | 14.12810458 | 0.01483 |
| GOTERM_CC_DIRECT | GO:0031588~nucleotide-activated protein kinase complex | 3 | 0.04495 | Q9DBC7, P05132, P12367 | 8.669518717 | 0.16483 |
| Annotation Cluster 75 | Enrichment Score: 1.231970065741046 | | | | | |
| Category | Term | Count | PValue | Genes | Fold Enrichment | FDR |
| GOTERM_BP_DIRECT | GO:0019370~leukotriene biosynthetic process | 4 | 0.00543 | P24527, P47713, P48999, Q9CPU4 | 10.64946237 | 0.12638 |
| UP_KW_BIOLOGICAL_PROCESS | KW-0434~Leukotriene biosynthesis | 3 | 0.04008 | P24527, P47713, P48999 | 9.038602941 | 0.16031 |

| | | | | | | |
|---|---|---|---|---|---|---|
| Annotation Cluster 76 | Enrichment Score: 1.2292690322512592 | | | | | |
| Category | Term | Count | PValue | Genes | Fold Enrichment | FDR |
| GOTERM_MF_DIRECT | GO:0046933~proton-transporting ATP synthase activity, rotational mechanism | 4 | 0.01558 | Q9CQQ7, P56480, Q91VR2, Q9DB20 | 7.397442455 | 0.19548 |
| GOTERM_CC_DIRECT | GO:0045259~proton-transporting ATP synthase complex | 4 | 0.03053 | Q9CQQ7, P56480, Q91VR2, Q9DB20 | 5.779679144 | 0.12176 |
| Annotation Cluster 77 | Enrichment Score: 1.2038629681313286 | | | | | |
| Category | Term | Count | PValue | Genes | Fold Enrichment | FDR |
| GOTERM_CC_DIRECT | GO:0000776~kinetochore | 12 | 0.01309 | Q61686, P46061, O55131, A2AGT5, P63087, Q9D8B3, Q8CBY8, Q8BUI3, O54941, F6ZDS4, Q9WVA3, O08901 | 2.369309463 | 0.06141 |
| UP_KW_CELLULAR_COMPONENT | KW-0137~Centromere | 12 | 0.0854 | Q61686, P46061, O55131, A2AGT5, P63087, Q8BUI3, Q61164, P35922, Q61550, F6ZDS4, Q9WVA3, O08901 | 1.738024231 | 0.16577 |
| Annotation Cluster 78 | Enrichment Score: 1.1907806845922653 | | | | | |
| Category | Term | Count | PValue | Genes | Fold Enrichment | FDR |
| GOTERM_MF_DIRECT | GO:0005200~structural constituent of cytoskeleton | 9 | 0.00282 | A2AQ07, P61161, Q9JM76, P57780, Q922F4, P48678, P68372, P68373, P26039 | 3.72305492 | 0.05025 |
| KEGG_PATHWAY | mmu04540:Gap junction | 12 | 0.01898 | A2AQ07, Q922F4, P68372, P68373, P05132, P08556, P11881, P11440, P08752, P68181 | 2.20603498 | 0.09129 |
| Annotation Cluster 79 | Enrichment Score: 1.1811093626566689 | | | | | |
| Category | Term | Count | PValue | Genes | Fold Enrichment | FDR |
| GOTERM_BP_DIRECT | GO:0015804~neutral amino acid transport | 5 | 0.00372 | Q9Z127, Q8K2P7, P10852, P51912, Q9WTR6 | 7.606758833 | 0.10349 |
| Annotation Cluster 80 | Enrichment Score: 1.1711457267905139 | | | | | |
| Category | Term | Count | PValue | Genes | Fold Enrichment | FDR |
| GOTERM_BP_DIRECT | GO:0007229~integrin-mediated signaling pathway | 11 | 0.00239 | P20491, Q9Z0F8, P97333, P24063, Q05910, P11835, Q9R0C8, P11688, Q9JHK5, P60766, P26039 | 3.19483871 | 0.07566 |
| GOTERM_MF_DIRECT | GO:0050839~cell adhesion molecule binding | 8 | 0.01046 | P26041, P97300, O08992, P26043, P24063, Q05910, P11835, P11688 | 3.309382151 | 0.14352 |
| Annotation Cluster 81 | Enrichment Score: 1.1630885047808155 | | | | | |
| Category | Term | Count | PValue | Genes | Fold Enrichment | FDR |

| | | | | | | |
|---|---|---|---|---|---|---|
| KEGG_PATHWAY | mmu00630:Glyoxylate and dicarboxylate metabolism | 7 | 0.01369 | Q9CZN7, P24270, O08749, P08249, Q9QXG4, Q8QZT1, Q8CAY6 | 3.458419421 | 0.06945 |
| Annotation Cluster 82 | Enrichment Score: 1.1625928816025548 | | | | | |
| Category | Term | Count | PValue | Genes | Fold Enrichment | FDR |
| GOTERM_BP_DIRECT | GO:0046513~ceramide biosynthetic process | 7 | 0.0008 | O09005, Q9WV54, Q9D6K9, Q924Z4, Q9D6J1, P17439, Q9JLJ5 | 6.21218638 | 0.0377 |
| Annotation Cluster 83 | Enrichment Score: 1.1536692318897048 | | | | | |
| Category | Term | Count | PValue | Genes | Fold Enrichment | FDR |
| GOTERM_BP_DIRECT | GO:0006869~lipid transport | 10 | 0.01015 | Q8VBT6, Q7TQH7, Q3UVL4, Q8BX94, P35951, B9EJ86, P50637, P55096, Q8BX70, Q3U7R1 | 2.778120617 | 0.19209 |
| Annotation Cluster 84 | Enrichment Score: 1.0727269680642555 | | | | | |
| Category | Term | Count | PValue | Genes | Fold Enrichment | FDR |
| GOTERM_BP_DIRECT | GO:0090611~ubiquitin-independent protein catabolic process via the multivesicular body sorting pathway | 3 | 0.00917 | Q9WU78, Q9D8B3, P46467 | 19.16903226 | 0.17614 |
| Annotation Cluster 85 | Enrichment Score: 1.0392081087612537 | | | | | |
| Category | Term | Count | PValue | Genes | Fold Enrichment | FDR |
| GOTERM_CC_DIRECT | GO:0030904~retromer complex | 4 | 0.02698 | Q9CWK8, Q9D8U8, A6H8H2, O70493 | 6.054901961 | 0.11106 |
| Annotation Cluster 86 | Enrichment Score: 0.9468157319337495 | | | | | |
| Category | Term | Count | PValue | Genes | Fold Enrichment | FDR |
| GOTERM_BP_DIRECT | GO:0032469~endoplasmic reticulum calcium ion homeostasis | 5 | 0.00308 | O55143, Q6PHZ2, Q64518, Q60953, P11881 | 7.987096774 | 0.09091 |
| GOTERM_CC_DIRECT | GO:0016529~sarcoplasmic reticulum | 7 | 0.01308 | P26883, O55143, Q6PHZ2, Q64518, O08528, P11881, Q00519 | 3.588994307 | 0.06141 |
| Annotation Cluster 87 | Enrichment Score: 0.9259284650944319 | | | | | |
| Category | Term | Count | PValue | Genes | Fold Enrichment | FDR |
| KEGG_PATHWAY | mmu01232:Nucleotide metabolism | 12 | 0.01613 | Q9WV85, P23492, P00493, Q91YL3, P04184, P24547, O08739, Q9D892, P11157, Q00519, Q9WTP6, Q3UFY7 | 2.258559622 | 0.07895 |
| Annotation Cluster 88 | Enrichment Score: 0.8623059953018872 | | | | | |

| Category | Term | Count | PValue | Genes | Fold Enrichment | FDR |
|---|---|---|---|---|---|---|
| GOTERM_CC_DIRECT | GO:0005952~cAMP-dependent protein kinase complex | 4 | 0.00225 | Q9DBC7, P05132, P68181, P12367 | 14.12810458 | 0.01483 |
| Annotation Cluster 89 | Enrichment Score: 0.8317467942316508 | | | | | |
| Category | Term | Count | PValue | Genes | Fold Enrichment | FDR |
| GOTERM_BP_DIRECT | GO:0001913~T cell mediated cytotoxicity | 6 | 0.00062 | P00493, P16872, P01902, P01900, P97821 | 8.334361851 | 0.03071 |
| GOTERM_BP_DIRECT | GO:0002485~antigen processing and presentation of endogenous peptide antigen via MHC class I via ER pathway, TAP-dependent | 3 | 0.00562 | P01902, P01900 | 23.96129032 | 0.12737 |
| Annotation Cluster 115 | Enrichment Score: 0.8040705028103422 | | | | | |
| Category | Term | Count | PValue | Genes | Fold Enrichment | FDR |
| GOTERM_MF_DIRECT | GO:0101005~deubiquitinase activity | 5 | 0.00393 | Q8BY87, Q8R5H1, Q9JMA1, P56399, Q9JKB1 | 7.485507246 | 0.06478 |
| Annotation Cluster 90 | Enrichment Score: 0.7062247461280503 | | | | | |
| Category | Term | Count | PValue | Genes | Fold Enrichment | FDR |
| GOTERM_MF_DIRECT | GO:0003925~G protein activity | 9 | 0.00042 | P63001, Q01514, P61021, Q9CQD1, P61027, P08556, P60766 | 4.964073227 | 0.01227 |
| KEGG_PATHWAY | mmu04370:VEGF signaling pathway | 12 | 0.00087 | P63001, Q05769, Q9JIA7, P63328, Q60591, P49138, P08556, P47713, P60766, P47811 | 3.271017384 | 0.0097 |
| GOTERM_BP_DIRECT | GO:0045445~myoblast differentiation | 6 | 0.0016 | P16045, Q501J6, P20918, P08556 | 6.846082949 | 0.05947 |
| KEGG_PATHWAY | mmu04664:Fc epsilon RI signaling pathway | 12 | 0.00259 | P63001, Q60787, P20491, O09110, Q9R0C8, P08556, P47713, P47811, P48999, Q9ES52 | 2.874530428 | 0.01986 |
| Annotation Cluster 91 | Enrichment Score: 0.6708982890127261 | | | | | |
| Category | Term | Count | PValue | Genes | Fold Enrichment | FDR |
| UP_KW_CELLULAR_COMPONENT | KW-0168~Coated pit | 9 | 0.00336 | Q7TN05, Q91ZX7, Q7TQH7, Q6PIU9, O08585, Q80TY0, P35951, P22892, P62743 | 3.566228965 | 0.01191 |
| Annotation Cluster 92 | Enrichment Score: 0.6202637043094084 | | | | | |
| Category | Term | Count | PValue | Genes | Fold Enrichment | FDR |

| | | | | | | |
|---|---|---|---|---|---|---|
| GOTERM_CC_DIRECT | GO:0016020~membrane | 219 | 0.00081 | Q9QXZ0, Q9CZW4, O70370, Q62086, O70251, O70493, Q99J47, Q9CR60, Q9R1J0, Q91VC9, O08553, P37040, O88456, P47753, P47754, P61027, A1L314, O88455, P56480, Q9D7N9, P47758, P28798, Q9Z127, Q61207, E2JF22, Q9ET30, P51912, Q8BVL3, Q64281, Q9JJ00, P62482, O88447, Q8VBZ3, P29416, Q9DB20, Q9CQS8, Q91X58, Q9WV54, P03921, P51125, Q8BKE6, Q9WTR6, P70206, Q9CQB5, Q7TQH7, P58681, P20934, Q9D6J1, Q99JB2, Q61543, P23780, P97333, Q6P9Q6, Q6DID7, Q3U9G9, O35375, Q91V01, P60766, Q9QZI9, P11928, Q8K2P7, P97300, Q5SSL4, Q69ZK0, Q9JLJ5, Q8VC04, Q80UU9, P40240, O08992, Q9DB73, Q9D8T2, Q64518, Q8JZQ2, Q9WTM5, P35564, Q9CQX2, Q9DBG7, O35678, Q58A65, Q8BIG7, Q9D270, Q05910, Q61490, Q8BGS7, Q9QXG4, P40237, Q99J62, Q8BRG8, P20060, P51569, O35350, Q91ZN5, Q9D8B6, Q61210, Q9CPU4, Q9D8B3, Q9QYJ0, Q9CY57, Q921J2, Q8VEB4, Q8VI94, Q09143, Q8VI93, Q9JIA7, Q60930, Q8BH24, O08539, Q80W54, Q9D2V7, P32037, Q99J56, Q8VDP6, B9EJ86, Q99KG5, P09528, Q8BVE3, Q9R233, Q61033, O55131, Q99KN9, Q8BXZ1, Q8BGH2, O35405, Q9D7V9, Q9R1S3, P50428, Q64430, Q99MR3, P14206, Q9R0Q8, P13597, P16872, O55143, P01902, Q8VEG4, P01900, P05064, Q3TVI8, O35639, P70280, P51881, P26883, Q8BH79, P97287, Q8BYI6, P10852, Q8BU14, Q9QWR8, Q9D0F3, Q921M7, Q8CI51, P11835, P59017, P11157, Q9EQ32, Q3TBD2, O09159, P60122, Q8R0X7, P08556, Q11011, P26645, O88384, Q9D0M3, Q91ZX7, Q9D6Y7, P50171, Q3UVK0, Q9EP89, P12970, P11881, Q99L43, P63001, Q9JHJ3, Q8CIM5, O35609, Q02819, P61620, Q99NH8, P08249, O08917, O08915, P70699, P46978, O55101, Q9CWK8, P62046, E9Q9A9, Q61093, P09581, Q920Q4, Q8K385, Q3U2C5, Q9QYA2, Q8VCW4, P35951, O70152, P50637, P70248, Q00612, Q9Z0F8, P08101, Q8C7X2, P55096, Q9CWZ7, Q3UVL4, Q791V5, Q62393, Q64695, P62192, Q9R1Q6, Q8C522, Q6P549, P63038 | 1.220053195 | 0.00648 |
| Annotation Cluster 93 | Enrichment Score: 0.35253415789006576 | | | | | |
| Category | Term | Count | PValue | Genes | Fold Enrichment | FDR |
| KEGG_PATHWAY | mmu04540:Gap junction | 12 | 0.01898 | A2AQ07, Q922F4, P68372, P68373, P05132, P08556, P11881, P11440, P08752, P68181 | 2.20603498 | 0.09129 |

Supplementary Table 5: Results of the pathway analysis by the David tool on the proteins modulated in response to PBk11

| | | | | | | |
|---|---|---|---|---|---|---|
| Annotation Cluster 1 | Enrichment Score: 6.4807314102770865 | | | | | |
| Category | Term | Count | PValue | Genes | Fold Enrichment | FDR |
| UP_SEQ_FEATURE | DISULFID:Interchain (between SU and TM chains, or C-337 with C-558) | | 1.435E-13 | P10404 | 2.8557E-11 | 2.7983E-11 |
| UP_SEQ_FEATURE | MOTIF:YXXL motif | | 1.435E-13 | P10404 | 2.8557E-11 | 2.7983E-11 |
| UP_SEQ_FEATURE | REGION:Immunosuppression | 7 | 1.707E-12 | P10404 | 144.232407 | 2.2191E-10 |
| UP_SEQ_FEATURE | REGION:Fusion peptide | 7 | 4.249E-12 | P10404 | 129.809167 | 3.3142E-10 |
| UP_SEQ_FEATURE | MOTIF:CX6CC | 7 | 4.249E-12 | P10404 | 129.809167 | 3.3142E-10 |
| UP_SEQ_FEATURE | MOTIF:CXXC | 7 | 3.427E-11 | P10404 | 99.8532051 | 2.2274E-09 |
| UP_KW_CELLULAR_COMPONENT | KW-0946~Virion | 7 | 2.561E-10 | P10404 | 73.7531073 | 3.201E-09 |
| UP_KW_CELLULAR_COMPONENT | KW-0261~Viral envelope protein | 7 | 2.561E-10 | P10404 | 73.7531073 | 3.201E-09 |
| INTERPRO | IPR008981:FMuLV_rcpt-bd | 7 | 9.589E-10 | P10404 | 62.3588 | 4.6218E-07 |
| INTERPRO | IPR018154:TLV/ENV_coat_polyprotein | 7 | 5.542E-09 | P10404 | 47.9683077 | 1.3357E-06 |
| UP_SEQ_FEATURE | SITE:Cleavage | 7 | 1.01E-05 | P10404 | 14.109692 | 0.0005625 |
| UP_SEQ_FEATURE | LIPID:S-palmitoyl cysteine | 10 | 3.11E-05 | Q9CQS8, P08228, P10404, Q62351 | 6.26492117 | 0.0015159 |
| UP_KW_PTM | KW-0449~Lipoprotein | 17 | 0.0048521 | Q9CQS8, Q62159, P62242, Q99JI6, P11928, P08228, E9Q7G0, P10404, Q91V41, P62192, Q62351 | 2.14913179 | 0.0274429 |
| UP_KW_PTM | KW-0564~Palmitate | 10 | 0.0074563 | Q9CQS8, P08228, P10404, Q62351 | 2.88735934 | 0.0274429 |
| UP_KW_PTM | KW-0165~Cleavage on pair of basic residues | 7 | 0.0252014 | P10404 | 3.0889297 | 0.07001 |
| Annotation Cluster 2 | Enrichment Score: 4.85806624607755 | | | | | |
| Category | Term | Count | PValue | Genes | Fold Enrichment | FDR |
| GOTERM_MF_DIRECT | GO:0005524~ATP binding | 26 | 5.506E-09 | Q9D975, P11928, O89051, P80313, B1ARD6, Q61316, Q3TRM8, Q04750, Q8JZQ2, P38647, P62192, Q64433, Q68FL6, Q8CGC7, P49717, Q6P9R2, P25206, Q9JHU4, Q6Q899, P23298, P63038, Q91ZW3, P60710, Q9D0I9 | 3.96242143 | 1.8223E-06 |
| GOTERM_MF_DIRECT | GO:0016887~ATP hydrolysis activity | 13 | 2.498E-07 | P80313, P49717, B1ARD6, P38647, Q8JZQ2, P25206, P62192, Q9JHU4, Q6Q899, Q91ZW3, P60710 | 7.22696957 | 2.7566E-05 |
| UP_KW_LIGAND | KW-0547~Nucleotide-binding | 28 | 0.0045327 | P11928, Q61316, P38647, Q8CGC7, P49717, Q6P9R2, P25206, P60710, Q62159, Q8CI11, Q9D975, P80313, B1ARD6, Q3TRM8, Q8JZQ2, Q8C0D5, P62192, Q99JI6, Q68FL6, Q91V41, Q9JHU4, Q6Q899, P23298, P63038, Q91ZW3, Q9D0I9 | 1.61858803 | 0.0355679 |
| UP_KW_LIGAND | KW-0067~ATP-binding | 23 | 0.005928 | Q9D975, P11928, P80313, B1ARD6, Q61316, Q3TRM8, P38647, Q8JZQ2, P62192, Q68FL6, Q8CGC7, P49717, Q6P9R2, P25206, Q9JHU4, Q6Q899, P23298, P63038, Q91ZW3, P60710, Q9D0I9 | 1.7364279 | 0.0355679 |
| Annotation Cluster 3 | Enrichment Score: 4.145763009116716 | | | | | |

| Category | Term | Count | PValue | Genes | Fold Enrichment | FDR |
|---|---|---|---|---|---|---|
| GOTERM_CC_DIRECT | GO:0005654~nucleoplasm | 36 | 8.683E-07 | P11928, P04117, Q923D2, Q9JMH6, Q9Z0H7, P10852, D0QMC3, P49717, Q9CVB6, O55135, Q6NVF9, Q9DC16, P60335, P25206, Q9JIX8, P28659, Q8K354, P60710, Q8CI11, P08228, B1ARD6, E9Q7G0, Q04750, P62192, Q62376, P57776, Q60692, P67778, Q99KC8, Q9JK81, Q91ZW3, Q4FK66, Q9D0I9, Q99LI7 | 2.39639866 | 3.2374E-05 |
| GOTERM_CC_DIRECT | GO:0005634~nucleus | 48 | 4.33E-06 | P11928, Q62426, P04117, Q61316, Q7TPV4, Q9JMH6, Q9Z0H7, O35593, P38647, Q8BTM8, Q9JKF1, Q99J47, D0QMC3, Q9D287, Q9JL16, P49717, Q9CVB6, O55135, Q64152, Q6NVF9, P60335, P25206, Q9JIX8, P28659, P60710, Q62159, P05555, Q8CI11, P08228, B1ARD6, E9Q7G0, Q8BIW1, Q9D8N0, Q04750, P14901, P68254, Q62376, P57776, Q60692, P67778, Q9JK81, Q91ZW3, Q4FK66, Q9D0I9, P00375, Q99LI7 | 1.89651844 | 0.0001027 |
| Annotation Cluster 4 | Enrichment Score: 4.137311070294695 | | | | | |
| Category | Term | Count | PValue | Genes | Fold Enrichment | FDR |
| GOTERM_CC_DIRECT | GO:1990904~ribonucleoprotein complex | 18 | 3.263E-14 | Q8C0D5, Q9JKF1, Q62376, P47915, P62242, Q8CGC7, P14131, Q6NVF9, O09167, P60335, P28659, Q6Q899, P63325, P60710 | 13.2506749 | 2.8385E-12 |
| GOTERM_BP_DIRECT | GO:0002181~cytoplasmic translation | 8 | 2.63E-07 | P47915, P62242, P14131, O09167, P14115, P63325 | 18.4689977 | 0.0001582 |
| GOTERM_BP_DIRECT | GO:0006412~translation | 10 | 3.538E-07 | P47915, P62242, Q8CGC7, P14131, Q9CPR5, O09167, O70251, P14115 | 10.8990866 | 0.0001582 |
| GOTERM_CC_DIRECT | GO:0045202~synapse | 16 | 3.738E-07 | P68254, P10852, P57776, P14115, P53986, P47915, P67778, P14131, Q9CVB6, O55135, O09167, P63325, P60710 | 5.25471823 | 1.951E-05 |
| GOTERM_CC_DIRECT | GO:0022626~cytosolic ribosome | 8 | 1.095E-06 | P47915, P62242, P14131, O09167, P14115, P63325 | 14.9780157 | 3.3288E-05 |
| GOTERM_CC_DIRECT | GO:0098794~postsynapse | 12 | 1.148E-06 | P47915, P14131, Q9CVB6, O09167, Q8BTM8, Q9JKF1, Q8BPU7, Q62351, P14115, P63325 | 7.02786207 | 3.3288E-05 |
| GOTERM_CC_DIRECT | GO:0005840~ribosome | 9 | 1.353E-06 | P47915, P62242, P14131, P11928, O09167, P14115, P63325 | 11.3226667 | 3.53E-05 |
| GOTERM_BP_DIRECT | GO:0140236~translation at presynapse | 6 | 5.276E-06 | P47915, P14131, P14115, P63325 | 23.8076923 | 0.0012734 |
| GOTERM_BP_DIRECT | GO:0140242~translation at postsynapse | 6 | 5.698E-06 | P47915, P14131, P14115, P63325 | 23.4414201 | 0.0012734 |
| GOTERM_MF_DIRECT | GO:0003735~structural constituent of ribosome | 9 | 7.098E-06 | P47915, P62242, P14131, Q9CPR5, O09167, P14115, P63325 | 9.04440275 | 0.0003916 |
| UP_KW_MOLECULAR_FUNCTION | KW-0687~Ribonucleoprotein | 11 | 6.843E-05 | P47915, P62242, P14131, Q9CPR5, O09167, P60335, P14115, Q62376, P63325 | 4.92076149 | 0.0015359 |
| UP_KW_MOLECULAR_FUNCTION | KW-0689~Ribosomal protein | 9 | 7.492E-05 | P47915, P62242, P14131, Q9CPR5, O09167, P14115, P63325 | 6.37794469 | 0.0015359 |
| UP_SEQ_FEATURE | REGION:2 X 8 AA tandem repeats of A-X-A-K-A-P-A-[KQ] | 3 | 8.477E-05 | P47915 | 185.441667 | 0.0036736 |
| INTERPRO | IPR002673:Ribosomal_eL29 | 3 | 0.0001832 | P47915 | 133.626 | 0.0220713 |

| | | | | | | |
|---|---|---|---|---|---|---|
| GOTERM_CC_DIRECT | GO:0022625~cytosolic large ribosomal subunit | 5 | 0.0004303 | P47915, O09167, P14115 | 14.1533333 | 0.0070192 |
| KEGG_PATHWAY | mmu03010:Ribosome | 9 | 0.0020449 | P47915, P62242, P14131, Q9CPR5, O09167, P14115, P63325 | 3.86725067 | 0.0777582 |
| KEGG_PATHWAY | mmu05171:Coronavirus disease - COVID-19 | 10 | 0.0023421 | P47915, P62242, P14131, P11928, O09167, Q6Q899, P14115, P63325 | 3.40925292 | 0.0777582 |
| GOTERM_CC_DIRECT | GO:0098793~presynapse | 7 | 0.003231 | P47915, P14131, Q9CVB6, P14115, P63325 | 4.85257143 | 0.0281097 |
| GOTERM_BP_DIRECT | GO:0031589~cell-substrate adhesion | 3 | 0.0046194 | P47915 | 29.3017751 | 0.1795535 |
| GOTERM_CC_DIRECT | GO:0022627~cytosolic small ribosomal subunit | 3 | 0.0222005 | P62242, P14131, P63325 | 12.9696 | 0.1148913 |
| Annotation Cluster 5 | Enrichment Score: 3.804778653986689 | | | | | |
| Category | Term | Count | PValue | Genes | Fold Enrichment | FDR |
| UP_KW_BIOLOGICAL_PROCESS | KW-0648~Protein biosynthesis | 8 | 1.337E-05 | Q68FL6, Q8CGC7, Q9D8N0, O55135, Q8C0D5, O70251, P57776, Q9D0I9 | 9.93333333 | 0.0007489 |
| GOTERM_MF_DIRECT | GO:0003746~translation elongation factor activity | 4 | 0.0001389 | Q9D8N0, Q8C0D5, O70251, P57776 | 39.1924119 | 0.0065675 |
| UP_KW_MOLECULAR_FUNCTION | KW-0251~Elongation factor | 4 | 0.0004453 | Q9D8N0, Q8C0D5, O70251, P57776 | 26.0271682 | 0.0045642 |
| GOTERM_BP_DIRECT | GO:0006414~translational elongation | 4 | 0.0007301 | Q9D8N0, Q8C0D5, O70251, P57776 | 22.5732194 | 0.0407923 |
| Annotation Cluster 6 | Enrichment Score: 3.0691661308611766 | | | | | |
| Category | Term | Count | PValue | Genes | Fold Enrichment | FDR |
| UP_KW_BIOLOGICAL_PROCESS | KW-0648~Protein biosynthesis | 8 | 1.337E-05 | Q68FL6, Q8CGC7, Q9D8N0, O55135, Q8C0D5, O70251, P57776, Q9D0I9 | 9.93333333 | 0.0007489 |
| INTERPRO | IPR036282:Glutathione-S-Trfase_C_sf | 4 | 0.0022846 | Q68FL6, Q8CGC7, Q9D8N0, O70251 | 15.163234 | 0.1348153 |
| Annotation Cluster 7 | Enrichment Score: 2.93034225106297 | | | | | |
| Category | Term | Count | PValue | Genes | Fold Enrichment | FDR |
| GOTERM_BP_DIRECT | GO:0006457~protein folding | 6 | 0.0001557 | P80313, Q61316, P38647, Q9CWM4, P63038, Q64433 | 11.8115683 | 0.0139185 |
| GOTERM_MF_DIRECT | GO:0140662~ATP-dependent protein folding chaperone | 4 | 0.0002735 | P80313, Q61316, P38647, P63038 | 31.3539295 | 0.0100589 |
| GOTERM_MF_DIRECT | GO:0044183~protein folding chaperone | 4 | 0.0018144 | P80313, P38647, Q9CWM4, Q64433 | 16.5020682 | 0.040038 |
| UP_KW_MOLECULAR_FUNCTION | KW-0143~Chaperone | 7 | 0.0041239 | P80313, P62075, Q64152, P38647, Q9CWM4, P63038, Q64433 | 4.55475444 | 0.0338163 |
| GOTERM_MF_DIRECT | GO:0051082~unfolded protein binding | 4 | 0.0069987 | P80313, P38647, Q9CWM4, Q64433 | 10.2241075 | 0.105299 |
| Annotation Cluster 8 | Enrichment Score: 2.816234029512564 | | | | | |
| Category | Term | Count | PValue | Genes | Fold Enrichment | FDR |
| UP_KW_MOLECULAR_FUNCTION | KW-0560~Oxidoreductase | 15 | 0.0001275 | P23591, Q9D975, P09528, P08228, Q923D2, Q9JMH6, P68254, P14901, Q9ESY9, Q99J47, Q00612, Q9QYF1, O88455, Q8K354, P00375 | 3.31876566 | 0.0017425 |
| UP_KW_LIGAND | KW-0521~NADP | 9 | 0.000241 | P23591, Q923D2, Q00612, Q9JMH6, Q9QYF1, Q99J47, O88455, Q8K354, P00375 | 5.27329193 | 0.0043377 |
| GOTERM_MF_DIRECT | GO:0016491~oxidoreductase activity | 6 | 0.0023586 | P23591, Q9JMH6, Q9QYF1, Q99J47, O88455, Q8K354 | 6.47214142 | 0.0478895 |

| | | | | | | |
|---|---|---|---|---|---|---|
| INTERPRO | IPR036291:NAD(P)-bd_dom_sf | 6 | 0.0025173 | P23591, Q923D2, Q00612, Q9QYF1, Q99J47, Q8K354 | 6.36314286 | 0.1348153 |
| Annotation Cluster 9 | Enrichment Score: 2.4677962984091995 | | | | | |
| Category | Term | Count | PValue | Genes | Fold Enrichment | FDR |
| UP_SEQ_FEATURE | CROSSLNK:Glycyl lysine isopeptide (Lys-Gly) (interchain with G-Cter in SUMO2) | 17 | 0.0001485 | Q8CI11, P80313, E9Q7G0, Q9D8N0, Q7TPV4, Q04750, P10852, O70251, Q62376, P62242, P49717, P60335, Q9JIX8, P28659, P63038, Q91ZW3, B2RY56 | 3.00525103 | 0.0057933 |
| UP_KW_PTM | KW-1017~Isopeptide bond | 24 | 0.0080714 | Q8CI11, P80313, E9Q7G0, Q9D8N0, Q7TPV4, Q9Z0H7, Q04750, Q8BTM8, P68254, P10852, P62192, O70251, Q62376, P62242, P49717, P60335, Q9JIX8, P05064, P28659, Q6Q899, P63038, P63325, Q91ZW3, B2RY56 | 1.73993383 | 0.0274429 |
| UP_KW_PTM | KW-0832~Ubl conjugation | 30 | 0.0329459 | Q3KNM2, Q7TPV4, Q9JMH6, Q9Z0H7, Q8BTM8, O70251, P10852, P49717, O55135, P60335, Q9JIX8, P28659, P63325, P60710, B2RY56, Q8CI11, P80313, E9Q7G0, Q9D8N0, Q04750, P68254, P62192, Q62376, P62242, P05064, Q6Q899, P63038, Q91ZW3 | 1.43306438 | 0.07001 |
| Annotation Cluster 10 | Enrichment Score: 2.30548862783743 | | | | | |
| Category | Term | Count | PValue | Genes | Fold Enrichment | FDR |
| KEGG_PATHWAY | mmu04216:Ferroptosis | 7 | 1.254E-06 | P09528, P60335, O09172, P14901, P29391, P10852, Q62351 | 19.4410569 | 0.0002082 |
| GOTERM_BP_DIRECT | GO:0006879~intracellular iron ion homeostasis | 5 | 0.0001751 | P09528, P08228, P14901, P29391, Q62351 | 17.8837125 | 0.0142346 |
| KEGG_PATHWAY | mmu00860:Porphyrin metabolism | 5 | 0.0007079 | P09528, Q8CGC7, Q923D2, P14901, P29391 | 12.1137285 | 0.0391704 |
| Annotation Cluster 11 | Enrichment Score: 2.184036762109446 | | | | | |
| Category | Term | Count | PValue | Genes | Fold Enrichment | FDR |
| UP_KW_BIOLOGICAL_PROCESS | KW-0648~Protein biosynthesis | 8 | 1.337E-05 | Q68FL6, Q8CGC7, Q9D8N0, O55135, Q8C0D5, O70251, P57776, Q9D0I9 | 9.93333333 | 0.0007489 |
| GOTERM_CC_DIRECT | GO:0017101~aminoacyl-tRNA synthetase multienzyme complex | 3 | 0.0011088 | Q68FL6, Q8CGC7, Q9D0I9 | 59.444 | 0.0131541 |
| INTERPRO | IPR001412:aa-tRNA-synth_I_CS | 3 | 0.0030793 | Q68FL6, Q8CGC7, Q9D0I9 | 35.6336 | 0.1484219 |
| UP_SEQ_FEATURE | MOTIF:'HIGH' region | 3 | 0.0036593 | Q68FL6, Q8CGC7, Q9D0I9 | 32.725 | 0.1297393 |
| UP_KW_MOLECULAR_FUNCTION | KW-0030~Aminoacyl-tRNA synthetase | 3 | 0.0296069 | Q68FL6, Q8CGC7, Q9D0I9 | 11.0114943 | 0.1517354 |
| Annotation Cluster 12 | Enrichment Score: 1.9517073987018518 | | | | | |
| Category | Term | Count | PValue | Genes | Fold Enrichment | FDR |
| GOTERM_CC_DIRECT | GO:0043209~myelin sheath | 8 | 1.565E-05 | P67778, P08228, P38647, P05064, P63038, P60710 | 10.0116211 | 0.0003404 |
| INTERPRO | IPR043129:ATPase_NBD | 6 | 2.679E-05 | Q61316, Q3TRM8, P38647, P60710 | 16.968381 | 0.0043035 |
| GOTERM_BP_DIRECT | GO:0045176~apical protein localization | 4 | 3.788E-05 | Q91V41, P60710 | 59.7526395 | 0.0056449 |
| GOTERM_BP_DIRECT | GO:0072749~cellular response to cytochalasin B | 3 | 4.522E-05 | P60710 | 253.948718 | 0.005775 |

| | | | | | | |
|---|---|---|---|---|---|---|
| GOTERM_BP_DIRECT | GO:0051621~regulation of norepinephrine uptake | 3 | 9.021E-05 | P60710 | 190.461538 | 0.0100805 |
| GOTERM_BP_DIRECT | GO:0150111~regulation of transepithelial transport | 3 | 0.00015 | P60710 | 152.369231 | 0.0139185 |
| GOTERM_CC_DIRECT | GO:0005925~focal adhesion | 7 | 0.0002274 | Q9CVB6, Q8BIW1, P11835, Q9JKF1, P60710 | 8.11918049 | 0.0042389 |
| GOTERM_BP_DIRECT | GO:0071896~protein localization to adherens junction | 3 | 0.0003133 | P60710 | 108.835165 | 0.0233412 |
| GOTERM_CC_DIRECT | GO:0030863~cortical cytoskeleton | 4 | 0.0003859 | Q8BTM8, P60710 | 27.9736471 | 0.0067139 |
| GOTERM_BP_DIRECT | GO:0001738~morphogenesis of a polarized epithelium | 3 | 0.0004167 | P60710 | 95.2307692 | 0.0286543 |
| GOTERM_MF_DIRECT | GO:0098973~structural constituent of postsynaptic actin cytoskeleton | 3 | 0.0004859 | P60710 | 88.1829268 | 0.0146226 |
| GOTERM_CC_DIRECT | GO:0036464~cytoplasmic ribonucleoprotein granule | 5 | 0.0005358 | P60335, Q9JKF1, P60710 | 13.3582022 | 0.0077697 |
| GOTERM_CC_DIRECT | GO:0097433~dense body | 3 | 0.0006098 | P60710 | 79.2586667 | 0.0083763 |
| GOTERM_BP_DIRECT | GO:0048870~cell motility | 4 | 0.0006385 | Q8BPU7, P60710 | 23.6231366 | 0.0380574 |
| GOTERM_MF_DIRECT | GO:0030957~Tat protein binding | 3 | 0.0009467 | P60710 | 64.1330377 | 0.0241034 |
| GOTERM_CC_DIRECT | GO:0015629~actin cytoskeleton | 7 | 0.0010467 | Q8BTM8, P05064, Q9JKF1, Q6Q899, P60710 | 6.07456934 | 0.0131541 |
| GOTERM_CC_DIRECT | GO:0140092~bBAF complex | 3 | 0.0011088 | P60710 | 59.444 | 0.0131541 |
| GOTERM_BP_DIRECT | GO:0034333~adherens junction assembly | 3 | 0.0015348 | P60710 | 50.7897436 | 0.0722179 |
| GOTERM_MF_DIRECT | GO:0019894~kinesin binding | 4 | 0.0015523 | O88447, P60710 | 17.4188497 | 0.0367005 |
| INTERPRO | IPR004001:Actin_CS | 3 | 0.0019568 | P60710 | 44.542 | 0.1348153 |
| GOTERM_CC_DIRECT | GO:0071564~npBAF complex | 3 | 0.0019941 | P60710 | 44.583 | 0.0216854 |
| GOTERM_CC_DIRECT | GO:0140288~GBAF complex | 3 | 0.0019941 | P60710 | 44.583 | 0.0216854 |
| GOTERM_CC_DIRECT | GO:0035060~brahma complex | 3 | 0.0022538 | P60710 | 41.9604706 | 0.0235295 |
| INTERPRO | IPR020902:Actin/actin-like_CS | 3 | 0.0023042 | P60710 | 41.1156923 | 0.1348153 |
| GOTERM_CC_DIRECT | GO:0016586~RSC-type complex | 3 | 0.0025286 | P60710 | 39.6293333 | 0.0244431 |
| GOTERM_CC_DIRECT | GO:0071565~nBAF complex | 3 | 0.0025286 | P60710 | 39.6293333 | 0.0244431 |
| GOTERM_CC_DIRECT | GO:0005911~cell-cell junction | 6 | 0.0028905 | Q99JI6, Q8BTM8, Q9JKF1, P23298, P60710 | 6.176 | 0.0269438 |
| GOTERM_CC_DIRECT | GO:0098871~postsynaptic actin cytoskeleton | 3 | 0.0034424 | P60710 | 33.968 | 0.0289825 |
| GOTERM_MF_DIRECT | GO:0050998~nitric-oxide synthase binding | 3 | 0.0045833 | P60710 | 29.3943089 | 0.0758535 |
| GOTERM_BP_DIRECT | GO:0070316~regulation of G0 to G1 transition | 3 | 0.0049763 | P60710 | 28.2165242 | 0.1853659 |
| KEGG_PATHWAY | mmu04145:Phagosome | 7 | 0.0050444 | Q9CQS8, P05555, P11835, Q9JHU4, Q62351, P60710 | 4.35564663 | 0.1395611 |
| GOTERM_BP_DIRECT | GO:1900242~regulation of synaptic vesicle endocytosis | 3 | 0.0053454 | P60710 | 27.2087912 | 0.1896219 |
| GOTERM_BP_DIRECT | GO:2000819~regulation of nucleotide-excision repair | 3 | 0.0061203 | P60710 | 25.3948718 | 0.1944695 |
| GOTERM_CC_DIRECT | GO:0005903~brush border | 4 | 0.0062001 | Q8BTM8, P60710 | 10.6865618 | 0.0476784 |
| GOTERM_CC_DIRECT | GO:0035267~NuA4 histone acetyltransferase complex | 3 | 0.006512 | P60710 | 24.5975172 | 0.0476784 |
| GOTERM_BP_DIRECT | GO:1903076~regulation of protein localization to plasma membrane | 3 | 0.0065258 | P60710 | 24.5756824 | 0.1944695 |
| GOTERM_BP_DIRECT | GO:2000779~regulation of double-strand break repair | 3 | 0.0065258 | P60710 | 24.5756824 | 0.1944695 |
| SMART | SM00268:ACTIN | 3 | 0.0069386 | P60710 | 23.3824786 | 0.1921944 |
| GOTERM_CC_DIRECT | GO:0016514~SWI/SNF complex | 3 | 0.0069582 | P60710 | 23.7776 | 0.0477916 |
| GOTERM_CC_DIRECT | GO:0043296~apical junction complex | 3 | 0.0069582 | P60710 | 23.7776 | 0.0477916 |
| GOTERM_CC_DIRECT | GO:0044305~calyx of Held | 3 | 0.0104528 | P60710 | 19.2791351 | 0.0699532 |

| | | | | | | |
|---|---|---|---|---|---|---|
| GOTERM_MF_DIRECT | GO:0019901~protein kinase binding | 8 | 0.0105753 | Q62159, P11835, Q6P9R2, Q9JKF1, Q62351, P60710 | 3.30621401 | 0.140017 |
| GOTERM_CC_DIRECT | GO:0098978~glutamatergic synapse | 10 | 0.0107537 | Q99JI6, P67778, Q9CVB6, Q8BTM8, Q8BPU7, Q62351, P57776, P60710 | 2.73620253 | 0.0700475 |
| GOTERM_CC_DIRECT | GO:0070160~tight junction | 3 | 0.0110036 | P60710 | 18.7717895 | 0.0700475 |
| GOTERM_CC_DIRECT | GO:0016363~nuclear matrix | 4 | 0.0202809 | E9Q7G0, P60710 | 6.89205797 | 0.1102774 |
| GOTERM_CC_DIRECT | GO:0098685~Schaffer collateral - CA1 synapse | 4 | 0.0263867 | Q9CVB6, P60710 | 6.21636601 | 0.1284353 |
| UP_KW_PTM | KW-0558~Oxidation | 3 | 0.0299333 | P60710 | 10.9629425 | 0.07001 |
| Annotation Cluster 13 | Enrichment Score: 1.8672017978113602 | | | | | |
| Category | Term | Count | PValue | Genes | Fold Enrichment | FDR |
| GOTERM_CC_DIRECT | GO:0005759~mitochondrial matrix | 6 | 0.0064905 | Q8BH95, P38647, P63038, Q64433, Q9JK81, P00375 | 5.0952 | 0.0476784 |
| UP_KW_DOMAIN | KW-0809~Transit peptide | 8 | 0.0123215 | Q9CPR5, Q8BH95, P38647, Q9EP89, Q8JZQ2, P63038, Q9JK81, Q8VDK1 | 3.14626677 | 0.0800899 |
| UP_KW_CELLULAR_COMPONENT | KW-0496~Mitochondrion | 16 | 0.0165646 | Q3KNM2, P11928, Q9CPR5, Q78IK2, Q8BH95, P38647, Q9EP89, Q8JZQ2, Q64433, Q8VDK1, P67778, P62075, Q9D8S9, P63038, Q9JK81, P00375 | 1.93620059 | 0.1035288 |
| Annotation Cluster 14 | Enrichment Score: 1.6517694145863193 | | | | | |
| Category | Term | Count | PValue | Genes | Fold Enrichment | FDR |
| GOTERM_MF_DIRECT | GO:0003729~mRNA binding | 8 | 0.0001919 | Q6NVF9, Q7TNC4, P60335, P28659, Q62376, P00375, Q99LI7, B2RY56 | 6.71869919 | 0.0079398 |
| GOTERM_BP_DIRECT | GO:0006397~mRNA processing | 7 | 0.00053 | Q6NVF9, P60335, Q9JIX8, P28659, Q9JK81, Q9D287, B2RY56 | 6.91689115 | 0.0338452 |
| UP_KW_BIOLOGICAL_PROCESS | KW-0507~mRNA processing | 8 | 0.0052896 | Q6NVF9, Q7TNC4, Q9JIX8, P28659, Q9D287, Q4FK66, Q99LI7, B2RY56 | 3.688771 | 0.1481089 |
| GOTERM_CC_DIRECT | GO:0016607~nuclear speck | 7 | 0.0137689 | Q6NVF9, Q7TNC4, P60335, Q9JIX8, Q62376, Q9D287, B2RY56 | 3.56409422 | 0.0781235 |
| UP_KW_MOLECULAR_FUNCTION | KW-0694~RNA-binding | 11 | 0.0253561 | Q9JL16, Q68FL6, Q8CGC7, P11928, Q6NVF9, P60335, P28659, Q6Q899, Q62376, P00375, B2RY56 | 2.19922301 | 0.1485145 |
| Annotation Cluster 15 | Enrichment Score: 1.1406656322628883 | | | | | |
| Category | Term | Count | PValue | Genes | Fold Enrichment | FDR |
| GOTERM_BP_DIRECT | GO:0009615~response to virus | 5 | 0.000895 | Q9JL16, P11928, Q9D8N0, Q6Q899, P60710 | 11.6490238 | 0.0470655 |
| Annotation Cluster 16 | Enrichment Score: 0.8232127172499263 | | | | | |
| Category | Term | Count | PValue | Genes | Fold Enrichment | FDR |
| KEGG_PATHWAY | mmu04145:Phagosome | 7 | 0.0050444 | Q9CQS8, P05555, P11835, Q9JHU4, Q62351, P60710 | 4.35564663 | 0.1395611 |
| Annotation Cluster 17 | Enrichment Score: 0.8024356248221765 | | | | | |

| Category | Term | Count | PValue | Genes | Fold Enrichment | FDR |
|---|---|---|---|---|---|---|
| GOTERM_CC_DIRECT | GO:0000502~proteasome complex | 3 | 0.0222005 | O35593, P62192, Q60692 | 12.9696 | 0.1148913 |
| Annotation Cluster 18 | Enrichment Score: 0.7188438045042296 | | | | | |
| Category | Term | Count | PValue | Genes | Fold Enrichment | FDR |
| GOTERM_MF_DIRECT | GO:0003697~single-stranded DNA binding | 4 | 0.0116695 | P49717, Q04750, P25206, P60335 | 8.47403501 | 0.1485622 |
| GOTERM_CC_DIRECT | GO:0005694~chromosome | 5 | 0.029374 | Q8CI11, P49717, E9Q7G0, Q04750, P25206 | 4.27654676 | 0.1393931 |
| Annotation Cluster 19 | Enrichment Score: 0.6779194659955587 | | | | | |
| Category | Term | Count | PValue | Genes | Fold Enrichment | FDR |
| UP_SEQ_FEATURE | LIPID:S-palmitoyl cysteine | 10 | 3.11E-05 | Q9CQS8, P08228, P10404, Q62351 | 6.26492117 | 0.0015159 |
| UP_KW_PTM | KW-0564~Palmitate | 10 | 0.0074563 | Q9CQS8, P08228, P10404, Q62351 | 2.88735934 | 0.0274429 |
| UP_KW_DOMAIN | KW-1133~Transmembrane helix | 30 | 0.2072463 | P05555, Q9CQS8, Q3KNM2, Q9EQH2, O89051, Q8BH24, Q9WV55, Q78IK2, P10404, P11835, Q9EP89, Q8JZQ2, P14901, O70251, P10852, Q99J47, Q62351, P53986, Q68FL6, Q9DC16, Q9QYF1, Q9CZT5, Q9WU81, O88455 | 1.16914512 | 0.553423 |
