## Supplemental Tables S6 to S10 for "Beyond the ink: cellular and molecular effects of iron-based pigments on macrophages"

Supplementary Table 6: List of proteins associated with iron homeostasis and modulated in response to iron-oxide pigments

| accession | name | Max U PR 101 | Max U PBK11 | PR101/ctrl | PBK11/ctrl |
|---|---|---|---|---|---|
| P09528 | Ferritin heavy chain | 0 | 0 | 9.45 | 22.22 |
| P10852 | Amino acid transporter heavy chain SLC3A2 | 0 | 1 | 4.11 | 1.35 |
| P14901 | Heme oxygenase 1 | 0 | 0 | 18.38 | 5.21 |
| P29391 | Ferritin light chain 1 | 0 | 0 | 3.16 | 15.14 |
| P60335 | Poly(rC)-binding protein 1 | 12 | 2 | 1.02 | 0.57 |
| Q62351 | Transferrin receptor protein 1 | 5 | 0 | 1.14 | 0.27 |
| Q9EQH2 | Endoplasmic reticulum aminopeptidase 1 | 7 | 2 | 1.22 | 0.76 |
| Q8K385 | Ferric-chelate reductase 1 | 0 | 5 | 0.4 | 0.77 |

Supplementary Table 7: List of mitochondrial proteins modulated in response to iron oxide pigments

| accession | name | Max U PR 101 | Max U PBK11 | PR101/ctrl | PBK11/ctrl |
|---|---|---|---|---|---|
| P50171 | (3R)-3-hydroxyacyl-CoA dehydrogenase | 2 | 10 | 0.65 | 0.95 |
| Q64433 | 10 kDa heat shock protein, mitochondrial | 1 | 0 | 1.48 | 1.3 |
| O88986 | 2-amino-3-ketobutyrate coenzyme A ligase, mitochondrial | 0 | 10 | 3.67 | 0.94 |
| Q9JIK9 | 28S ribosomal protein S34, mitochondrial | 0 | 12 | 2.44 | 0.95 |
| Q9CQF0 | 39S ribosomal protein L11, mitochondrial | 2 | 11 | 0.66 | 0.99 |
| Q9DB15 | 39S ribosomal protein L12, mitochondrial | 0 | 12.5 | ND Ctrl&ERM | ND PBK11 |
| Q9CPR5 | 39S ribosomal protein L15, mitochondrial | 10 | 2 | 0.89 | 0.39 |
| Q9D1N9 | 39S ribosomal protein L21, mitochondrial | 0 | 11 | 5.63 | 0.73 |
| Q9CY73 | 39S ribosomal protein L44, mitochondrial | 0 | 7 | 0.56 | 0.74 |
| P63038 | 60 kDa heat shock protein, mitochondrial | 0 | 1 | 2.34 | 1.22 |
| Q8QZT1 | Acetyl-CoA acetyltransferase, mitochondrial | 2 | 10 | 0.71 | 0.96 |
| Q99KI0 | Aconitate hydratase, mitochondrial | 6 | 3 | 0.91 | 1.17 |
| Q9QYR9 | Acyl-coenzyme A thioesterase 2, mitochondrial | 1 | 12 | 1.49 | 1.04 |
| Q9WTP6 | Adenylate kinase 2, mitochondrial | 0 | 7 | 0.5 | 1.11 |
| P51881 | ADP/ATP translocase 2 | 0 | 10 | 1.55 | 0.94 |
| P24483 | Adrenodoxin, mitochondrial | 2 | 12 | 1.88 | 0.8 |
| Q920A7 | AFG3-like protein 1 | 1 | 11 | 1.57 | 0.99 |
| Q8JZQ2 | AFG3-like protein 2 | 0 | 2 | 1.86 | 1.79 |
| Q9CQQ7 | ATP synthase F(0) complex subunit B1, mitochondrial | 1 | 4 | 1.49 | 1.39 |
| Q78IK2 | ATP synthase F(0) complex subunit k, mitochondrial | 6 | 0 | 0.8 | 1.3 |
| P56480 | ATP synthase subunit beta, mitochondrial | 2 | 12 | 1.36 | 1.03 |
| Q91VR2 | ATP synthase subunit gamma, mitochondrial | 2 | 12 | 1.66 | 1.19 |
| Q9DB20 | ATP synthase subunit O, mitochondrial | 0 | 4 | 1.42 | 1.14 |
| O35143 | ATPase inhibitor, mitochondrial | 0 | 0 | 0.48 | 1.08 |
| P59017 | Bcl-2-like protein 13 | 1 | 9 | 0.35 | 0.66 |
| Q9DBL7 | Bifunctional coenzyme A synthase | 0 | 8 | 1.29 | 1.06 |
| P18155 | Bifunctional methylenetetrahydrofolate dehydrogenase/cyclohydrolase, mitochondrial | 0 | 6 | 3 | 1.19 |
| Q9D8S9 | BolA-like protein 1 | 12 | 0 | 0.96 | 1.29 |
| O35855 | Branched-chain-amino-acid aminotransferase, mitochondrial | 1 | 12 | 0.8 | 0.98 |
| P05132 | cAMP-dependent protein kinase catalytic subunit alpha | 1 | 8 | 0.52 | 0.78 |
| Q8K1J6 | CCA tRNA nucleotidyltransferase 1, mitochondrial | 2 | 12 | 0.65 | 0.95 |
| B0K020 | CDGSH iron-sulfur domain-containing protein 1 | 0 | 12 | 4.45 | 1.6 |
| Q9CQB5 | CDGSH iron-sulfur domain-containing protein 2 | 0 | 5 | 3.58 | 1.76 |

| | | | | | |
|---|---|---|---|---|---|
| Q8BHF7 | CDP-diacylglycerol--glycerol-3-phosphate 3-phosphatidyltransferase, mitochondrial | 0 | 12 | 2.74 | 1.18 |
| P54987 | Cis-aconitate decarboxylase | 0 | 4 | 7.11 | 2.13 |
| Q8JZN5 | Complex I assembly factor ACAD9, mitochondrial, | 0 | 12 | 0.71 | 0.99 |
| Q921M7 | CYFIP-related Rac1 interactor B | 0 | 8 | 0.59 | 0.86 |
| Q9CZ13 | Cytochrome b-c1 complex subunit 1, mitochondrial | 0 | 12 | 1.65 | 1.02 |
| Q9DB77 | Cytochrome b-c1 complex subunit 2, mitochondrial | 0 | 12 | 1.66 | 1.02 |
| Q9CR68 | Cytochrome b-c1 complex subunit Rieske, mitochondrial | 0 | 11 | 1.84 | 1.06 |
| P19783 | Cytochrome c oxidase subunit 4 isoform 1, mitochondrial | 0 | 8 | 1.57 | 1.05 |
| Q9CPQ1 | Cytochrome c oxidase subunit 6C | 0 | 11 | 1.68 | 1.11 |
| P62897 | Cytochrome c, somatic | 1 | 6 | 0.76 | 1.14 |
| Q9D0M3 | Cytochrome c1, heme protein, mitochondrial | 1 | 8 | 1.25 | 1.08 |
| P28271 | Cytoplasmic aconitate hydratase | 3 | 11 | 0.84 | 1.17 |
| Q8VDK1 | Deaminated glutathione amidase | 4 | 2 | 0.58 | 0.42 |
| Q8CHT0 | Delta-1-pyrroline-5-carboxylate dehydrogenase, mitochondrial | 2 | 7 | 0.29 | 0.46 |
| Q9Z110 | Delta-1-pyrroline-5-carboxylate synthase | 0 | 9 | 2.48 | 1.17 |
| P00375 | Dihydrofolate reductase | 8 | 0 | 0.64 | 0.61 |
| O08749 | Dihydrolipoyl dehydrogenase, mitochondrial | 2 | 4 | 1.23 | 1.18 |
| Q8BMF4 | Dihydrolipoyllysine-residue acetyltransferase component of pyruvate dehydrogenase complex, mitochondrial | 0 | 11 | 1.54 | 1.05 |
| Q5U458 | DnaJ homolog subfamily C member 11 | 0 | 12 | 2.39 | 1.13 |
| Q3KNM2 | E3 ubiquitin-protein ligase MARCH5 | 9 | 2 | 1.18 | 0.47 |
| P42125 | Enoyl-CoA delta isomerase 1, mitochondrial | 0 | 11 | 2.4 | 1.3 |
| Q8BH95 | Enoyl-CoA hydratase, mitochondrial | 3 | 2 | 1.73 | 1.69 |
| Q8VEG4 | Exonuclease 3'-5' domain-containing protein 2 | 0 | 9 | 0.23 | 0.83 |
| P47791 | Glutathione reductase, mitochondrial | 0 | 10 | 1.76 | 1.14 |
| Q9DCM2 | Glutathione S-transferase kappa 1 | 2 | 7 | 0.53 | 0.68 |
| Q91VC9 | Growth hormone-inducible transmembrane protein | 0 | 10 | 2.68 | 1.22 |
| Q9CQN1 | Heat shock protein 75 kDa, mitochondrial | 0 | 10 | 1.52 | 0.99 |
| P17710 | Hexokinase-1 | 0 | 8 | 0.56 | 0.89 |
| P97287 | Induced myeloid leukemia cell differentiation protein Mcl-1 homolog | 0 | 10 | 3.35 | 1.7 |
| Q9D6R2 | Isocitrate dehydrogenase [NAD] subunit alpha, mitochondrial | 0 | 6 | 1.53 | 1.09 |
| P41565 | Isocitrate dehydrogenase [NAD] subunit gamma 1, mitochondrial | 0 | 9 | 1.39 | 0.96 |
| P54071 | Isocitrate dehydrogenase [NADP], mitochondrial | 0 | 12 | 0.55 | 0.82 |
| Q8CGK3 | Lon protease homolog, mitochondrial | 1 | 5 | 1.36 | 1.14 |
| P08249 | Malate dehydrogenase, mitochondrial | 0 | 7 | 1.47 | 1.22 |
| P45952 | Medium-chain specific acyl-CoA dehydrogenase, mitochondrial | 2 | 11 | 0.69 | 1.08 |
| Q9CRB9 | MICOS complex subunit Mic19 | 12 | 10 | 1.45 | 1.49 |

| | | | | | |
|---|---|---|---|---|---|
| Q91VN4 | MICOS complex subunit Mic25 | 7 | 12 | 1.56 | 1.22 |
| Q78IK4 | MICOS complex subunit Mic27 | 2 | 12 | 0.75 | 1.02 |
| Q8CAQ8 | MICOS complex subunit Mic60 | 7 | 8 | 1.24 | 1.2 |
| Q9CPU4 | Microsomal glutathione S-transferase 3 | 1 | 7 | 0.44 | 0.66 |
| Q922Q1 | Mitochondrial amidoxime reducing component 2 | 1 | 8 | 1.32 | 0.92 |
| Q791V5 | Mitochondrial carrier homolog 2 | 0 | 8 | 1.39 | 1.18 |
| P84817 | Mitochondrial fission 1 protein | 0 | 11 | 0.21 | 0.76 |
| P62075 | Mitochondrial import inner membrane translocase subunit Tim13 | 11 | 2 | 1.04 | 1.79 |
| Q9WTQ8 | Mitochondrial import inner membrane translocase subunit Tim23 | 1 | 6 | 1.43 | 1.13 |
| Q9D880 | Mitochondrial import inner membrane translocase subunit TIM50 | 1 | 7 | 0.65 | 0.8 |
| Q9CYG7 | Mitochondrial import receptor subunit TOM34 | 0 | 9 | 0.55 | 1.09 |
| Q9QYA2 | Mitochondrial import receptor subunit TOM40 homolog | 0 | 9 | 2.19 | 1.44 |
| Q9CZW5 | Mitochondrial import receptor subunit TOM70 | 2 | 12 | 0.75 | 0.97 |
| Q9D6Y7 | Mitochondrial peptide methionine sulfoxide reductase | 2 | 12 | 0.29 | 0.99 |
| Q3URS9 | Mitochondrial potassium channel | 2 | 10 | 2.2 | 1.8 |
| Q811U4 | Mitofusin-1 | 2.5 | 7 | ND PR101 | 0.34 |
| O08776 | NADH dehydrogenase [ubiquinone] 1 alpha subcomplex assembly factc | 2.5 | 12.5 | ND ctrl&ERM | ND PBk11 |
| Q99LC3 | NADH dehydrogenase [ubiquinone] 1 alpha subcomplex subunit 10, mit | 12 | 10 | 0.85 | 0.7 |
| Q9D8B4 | NADH dehydrogenase [ubiquinone] 1 alpha subcomplex subunit 11 | 8 | 10 | 0.83 | 0.92 |
| Q7TMF3 | NADH dehydrogenase [ubiquinone] 1 alpha subcomplex subunit 12 | 4 | 9 | 0.71 | 0.7 |
| Q9ERS2 | NADH dehydrogenase [ubiquinone] 1 alpha subcomplex subunit 13 | 0 | 11 | 0.58 | 0.92 |
| Q9CQ75 | NADH dehydrogenase [ubiquinone] 1 alpha subcomplex subunit 2 | 0 | 11 | 0.57 | 1.01 |
| Q9CPP6 | NADH dehydrogenase [ubiquinone] 1 alpha subcomplex subunit 5 | 9 | 12 | 1.07 | 0.74 |
| Q9DCJ5 | NADH dehydrogenase [ubiquinone] 1 alpha subcomplex subunit 8 | 1 | 5 | 0.6 | 0.678 |
| Q9DC69 | NADH dehydrogenase [ubiquinone] 1 alpha subcomplex subunit 9, mitochondrial | 0 | 12 | 1.35 | 1.02 |
| Q9DCS9 | NADH dehydrogenase [ubiquinone] 1 beta subcomplex subunit 10 | 5 | 9 | 1.5 | 1.32 |
| Q9CQC7 | NADH dehydrogenase [ubiquinone] 1 beta subcomplex subunit 4 | 7 | 10 | ND ctrl | ND ctrl |
| Q9CQH3 | NADH dehydrogenase [ubiquinone] 1 beta subcomplex subunit 5, mitoc | 11 | 7 | 0.7 | 1.16 |
| Q91YT0 | NADH dehydrogenase [ubiquinone] flavoprotein 1, mitochondrial | 3 | 7 | 0.66 | 0.93 |
| Q91WD5 | NADH dehydrogenase [ubiquinone] iron-sulfur protein 2, mitochondrial | 6 | 12 | 0.76 | 1.01 |
| Q9DCT2 | NADH dehydrogenase [ubiquinone] iron-sulfur protein 3, mitochondrial | 6 | 5 | 1.26 | 0.79 |
| Q9DC70 | NADH dehydrogenase [ubiquinone] iron-sulfur protein 7, mitochondrial | 6 | 11 | 0.76 | 1.02 |
| Q8K3J1 | NADH dehydrogenase [ubiquinone] iron-sulfur protein 8, mitochondrial | 8 | 12 | 0.29 | 0.55 |
| Q91VD9 | NADH-ubiquinone oxidoreductase 75 kDa subunit, mitochondrial | 11 | 11 | 0.96 | 0.93 |
| P03921 | NADH-ubiquinone oxidoreductase chain 5 | 0 | 10 | 2.06 | 0.78 |
| P97333 | Neuropilin-1 | 0 | 11 | 4.86 | 1.69 |

| | | | | | |
|---|---|---|---|---|---|
| Q8BH04 | Phosphoenolpyruvate carboxykinase [GTP], mitochondrial | 0 | 9 | 1.41 | 1.08 |
| P67778 | Prohibitin | 0 | 2 | 1.6 | 1.2 |
| O35129 | Prohibitin-2 | 0 | 9 | 5.93 | 1.14 |
| Q9D8B6 | Protein FAM210B, mitochondrial | 0 | 12.5 | ND Ctrl&ERM | ND PBK11 |
| Q99LX0 | Protein/nucleic acid deglycase DJ-1 | 0 | 3 | 0.57 | 1.26 |
| Q9EP89 | Serine beta-lactamase-like protein LACTB, mitochondrial | 0 | 2 | 0.18 | 0.3 |
| Q9CZN7 | Serine hydroxymethyltransferase, mitochondrial | 0 | 4 | 1.69 | 1.21 |
| P63087 | Serine/threonine-protein phosphatase PP1-gamma catalytic subunit | 0 | 12 | 0.74 | 1.01 |
| Q8BGH2 | Sorting and assembly machinery component 50 homolog | 0 | 10 | 1.95 | 1.18 |
| O09005 | Sphingolipid delta(4)-desaturase DES1 | 0 | 8 | 7.84 | 2.16 |
| Q9JIA7 | Sphingosine kinase 2 | 0 | 9 | 1.67 | 1.09 |
| Q99JB2 | Stomatin-like protein 2, mitochondrial | 0 | 12 | 2.83 | 1.17 |
| P38647 | Stress-70 protein, mitochondrial | 0 | 1 | 2.02 | 1.31 |
| Q8K2B3 | Succinate dehydrogenase [ubiquinone] flavoprotein subunit, mitochondr | 12 | 10 | 0.88 | 1.1 |
| Q9CQA3 | Succinate dehydrogenase [ubiquinone] iron-sulfur subunit, mitochondrial | 0 | 10 | 0.51 | 0.8 |
| Q9CZB0 | Succinate dehydrogenase cytochrome b560 subunit, mitochondrial | 12 | 11 | 1.01 | 0.85 |
| Q9WUM5 | Succinate--CoA ligase [ADP/GDP-forming] subunit alpha, mitochondrial | 0 | 6 | 9.56 | 3.08 |
| Q9Z2I8 | Succinate--CoA ligase [GDP-forming] subunit beta, mitochondrial | 0 | 12 | 1.41 | 0.96 |
| P09671 | Superoxide dismutase [Mn], mitochondrial | 0 | 5 | 2.01 | 1.32 |
| Q62465 | Synaptic vesicle membrane protein VAT-1 homolog | 0 | 5 | 0.5 | 1.16 |
| Q921F2 | TAR DNA-binding protein 43 | 0 | 8 | 1.53 | 1.08 |
| P50637 | Translocator protein | 0 | 7 | 5.14 | 2.27 |
| Q8BMS1 | Trifunctional enzyme subunit alpha, mitochondrial | 0 | 12 | 0.65 | 1.1 |
| Q8BVW3 | Tripartite motif-containing protein 14 | 0 | 11 | 3.14 | 0.69 |
| Q8BYL4 | Tyrosine--tRNA ligase, mitochondrial | 1 | 9 | 1.52 | 0.78 |
| Q9CWU6 | Ubiquinol-cytochrome-c reductase complex assembly factor 1 | 1 | 3 | 0.42 | 0.55 |
| Q9JK81 | UPF0160 protein MYG1, mitochondrial | 10 | 0 | 0.91 | 1.22 |
| Q8BX70 | Vacuolar protein sorting-associated protein 13C | 0 | 7 | 0.65 | 0.9 |
| Q60930 | Voltage-dependent anion-selective channel protein 2 | 1 | 10 | 1.34 | 1.14 |
| Q91WL8 | WW domain-containing oxidoreductase | 0 | 8 | 1.6 | 0.92 |

Supplementary Table 8: List of proteins associated with glutathione metabolism and modulated in response to iron-oxide pigments

| accession | name | Max U PR 101 | Max U PBK11 | PR101/ctrl | PBK11/ctrl |
|---|---|---|---|---|---|
| Q9CPU4 | Microsomal glutathione S-transferase 3 | 1 | 7 | 0.43 | 0.66 |
| Q9DCM2 | Glutathione S-transferase kappa 1 | 2 | 7 | 0.53 | 0.68 |
| O35952 | Hydroxyacylglutathione hydrolase, mitochondrial | 2 | 4 | 0.17 | 0.28 |
| Q9R0P3 | S-formylglutathione hydrolase | 0 | 0 | 2.86 | 1.54 |
| P23764 | Glutathione peroxidase 3 | 0 | 12.5 | ND ctrl (>30000) | ND PBk11 |
| P47791 | Glutathione reductase, mitochondrial | 0 | 10 | 1.76 | 1.14 |
| P48774 | Glutathione S-transferase Mu 5 | 0 | 9 | 0.38 | 0.73 |
| O09172 | Glutamate--cysteine ligase regulatory subunit | 0 | 1 | 5.95 | 1.9 |
| Q9CPU0 | Lactoylglutathione lyase | 7 (vs ERM) 0 vs ctrl | 11 | 0.63 | 0.76 |
| P46413 | Glutathione synthetase | 7 | 11 | 4 | 3.5 |
| P10649 | Glutathione S-transferase Mu 1 | 4 | 6 | 0.77 | 1.22 |
| P19468 | Glutamate--cysteine ligase catalytic subunit | 10 | 10 | 0.92 | 1.12 |

Supplementary Table 9: List of lysosomal proteins modulated in response to iron oxide pigments

| accession | name | Max U PR 101 | Max U PBK11 | PR101/ctrl | PBK11/ctrl |
|---|---|---|---|---|---|
| Q8BG07 | 5'-3' exonuclease PLD4 | 0 | 10 | 0.31 | 0.96 |
| Q9WV54 | Acid ceramidase | 2 | 10 | 0.72 | 0.83 |
| P51569 | Alpha-galactosidase A | 2 | 10 | 0.54 | 1.3 |
| Q9QWR8 | Alpha-N-acetylgalactosaminidase | 0 | 7 | 0.27 | 0.84 |
| P50428 | Arylsulfatase A | 0 | 9 | 0.06 | 1.09 |
| P50429 | Arylsulfatase B | 0 | 2 | 0.07 | 0.42 |
| P23780 | Beta-galactosidase | 0 | 11 | 0.17 | 0.78 |
| P12265 | Beta-glucuronidase | 0 | 7 | 0.19 | 0.6 |
| P29416 | Beta-hexosaminidase subunit alpha | 0 | 8 | 0.35 | 0.63 |
| P20060 | Beta-hexosaminidase subunit beta | 0 | 5 | 0.28 | 0.6 |
| Q9WVJ3 | Carboxypeptidase Q | 0 | 9 | 0.21 | 1.22 |
| P10605 | Cathepsin B | 0 | 8 | 0.29 | 0.83 |
| P18242 | Cathepsin D | 0 | 4 | 0.48 | 1.26 |
| O70370 | Cathepsin S | 0 | 10 | 0.29 | 0.9 |
| Q9WUU7 | Cathepsin Z | 0 | 12 | 0.19 | 0.74 |
| P56542 | Deoxyribonuclease-2-alpha | 0 | 5 | 0.24 | 0.61 |
| P97821 | Dipeptidyl peptidase 1 | 0 | 11 | 0.34 | 0.76 |
| Q80TY0 | Formin-binding protein 1 | 0 | 10 | 0.52 | 0.8 |
| Q9ESY9 | Gamma-interferon-inducible lysosomal thiol reductase | 0 | 1 | 0.15 | 0.43 |
| P17439 | Glucosylceramidase | 2 | 10 | 1.72 | 1.05 |
| Q9JHJ3 | Glycosylated lysosomal membrane protein | 0 | 12 | 13 | 0.29 |
| P28798 | Granulins | 1 | 10 | 0.49 | 0.75 |
| Q8VEB4 | Group XV phospholipase A2 | 0 | 9 | 0.22 | 0.71 |
| Q91VK4 | Integral membrane protein 2C | 0 | 11.5 | 7.81 | 0.79 |
| O89017 | Legumain | 1 | 6 | 0.04 | 0.4 |
| P35951 | Low-density lipoprotein receptor | 0 | 8 | 3.41 | 0.75 |
| P70699 | Lysosomal alpha-glucosidase | 2 | 4 | 0.66 | 1.35 |
| O09159 | Lysosomal alpha-mannosidase | 0 | 0 | 0.12 | 0.61 |
| P16675 | Lysosomal protective protein | 0 | 7 | 0.18 | 0.93 |
| Q9DC37 | Major facilitator superfamily domain-containing protein 1 | 0 | 9 | 12.33 | 3.57 |
| Q571E4 | N-acetylgalactosamine-6-sulfatase | 0 | 12 | 0.21 | 0.94 |
| Q8BFR4 | N-acetylglucosamine-6-sulfatase | 0 | 5 | 0.15 | 0.71 |
| O09043 | Napsin-A | 1 | 9 | 0.48 | 0.62 |
| P41251 | Natural resistance-associated macrophage protein 1 | 0 | 12 | 3.75 | 1.31 |
| Q9Z0J0 | NPC intracellular cholesterol transporter 2 | 0 | 5 | 0.14 | 0.66 |
| O88531 | Palmitoyl-protein thioesterase 1 | 0 | 10 | 0.17 | 0.95 |
| O70172 | Phosphatidylinositol 5-phosphate 4-kinase type-2 alpha | 1 | 12 | 0.62 | 0.99 |
| Q61207 | Prosaposin | 0 | 9 | 0.42 | 1.06 |
| Q8VCW4 | Protein unc-93 homolog B1 | 0 | 10 | 2.16 | 1.24 |
| P61027 | Ras-related protein Rab-10 | 2 | 7 | 0.66 | 0.82 |
| Q9CYN9 | Renin receptor | 6 | 12.5 | poorly detected protein | ND PBk11 |
| P70665 | Sialate O-acetylesterase | 1 | 5 | 0.07 | 1.41 |

| | | | | | |
|---|---|---|---|---|---|
| P25286 | V-type proton ATPase 116 kDa subunit a isoform 1 | 9 | 12 | 1.66 | 1.33 |
| P15920 | V-type proton ATPase 116 kDa subunit a isoform 2 | 0 | 12 | 15.83 | 6.3 vs ctrl, 0.97 vs ERM |
| P63081 | V-type proton ATPase 16 kDa proteolipid subunit | 6 | 11 | 0.78 | 0.98 |
| P50516 | V-type proton ATPase catalytic subunit A | 9 | 9 | 0.99 | 1.07 |
| P62814 | V-type proton ATPase subunit B, brain isoform | 5 | 6 | 0.89 | 1.13 |
| Q5FVI6 | V-type proton ATPase subunit C 1 | 0 | 10 | 0.54 | 0.98 |
| P57746 | V-type proton ATPase subunit D | 3 | 7 | 1.5 | 0.81 |
| P51863 | V-type proton ATPase subunit d 1 | 3 | 12 | 1.17 | 1.04 |
| P50518 | V-type proton ATPase subunit E 1 | 9 | 12 | 1.25 | 1.19 |
| P50408 | V-type proton ATPase subunit F | 11 | 3 | 0.84 | 1.67 |
| Q9CR51 | V-type proton ATPase subunit G 1 | 4 | 7 | 0.85 | 1.17 |
| Q8BVE3 | V-type proton ATPase subunit H | 1 | 7 | 1.45 | 0.81 |
| Q9R1Q9 | V-type proton ATPase subunit S1 | 10 | 12.5 | poorly detected protein | ND PBK11 |
| Q920Q4 | Vacuolar protein sorting-associated protein 16 homolog | 2 | 8 | 0.41 | 0.73 |

Supplementary Table 10: List of immunity-associated proteins modulated in response to iron-oxide pigments

| accession | name | Max U PR 101 | Max U PBK11 | PR101/ctrl | PBK11/ctrl |
|---|---|---|---|---|---|
| P11928 | 2'-5'-oligoadenylate synthase 1A Mus musculus | 0 | 0 | 2.74 | 0.8 |
| E9Q9A9 | 2'-5'-oligoadenylate synthase 2 Mus musculus | 0 | 6 | 4.22 | 1.65 |
| Q8VI93 | 2'-5'-oligoadenylate synthase 3 Mus musculus | 0 | 11 | 0.73 | 1 |
| Q8VI94 | 2'-5'-oligoadenylate synthase-like protein 1 Mus musculus | 1 | 6 | 2.65 | 0.67 |
| O35405 | 5'-3' exonuclease PLD3 | 1 | 3 | 2.56 | 2.46 |
| Q8BG07 | 5'-3' exonuclease PLD4 | 0 | 10 | 0.31 | 0.96 |
| Q6Q899 | Antiviral innate immune response receptor RIG-I | 2 | 1 | 0.43 | 0.48 |
| Q9EPB4 | Apoptosis-associated speck-like protein containing a CARD Mus musculus | 0 | 4 | 0.41 | 0.63 |
| Q9Z0H7 | B-cell lymphoma/leukemia 10 Mus musculus | 0 | 1 | 0.1 | 0.69 |
| Q9R0Q8 | C-type lectin domain family 4 member E Mus musculus | 1 | 10 | 21.81 | 7.31 |
| Q61490 | CD166 | 0 | 6 | 4 | 1.2 |
| P54987 | Cis-aconitate decarboxylase Mus musculus | 0 | 4 | 7.11 | 2.13 |
| P14106 | Complement C1q subcomponent subunit B Mus musculus | 0 | 8 | 0.16 | 0.6 |
| P01027 | Complement C3 Mus musculus | 0 | 9 | 3.08 | 1.15 |
| P01029 | Complement C4-B Mus musculus | 0 | 6 | ND in ctrl,17 vs. ERM | ND in ctrl,7 vs. ERM |
| Q60710 | Deoxynucleoside triphosphate triphosphohydrolase SAMHD1 Mus musculus | 0 | 10 | 0.24 | 0.81 |
| Q3UIR3 | E3 ubiquitin-protein ligase DTX3L Mus musculus | 0 | 11 | 0.2 | 0.94 |
| Q80VI1 | E3 ubiquitin-protein ligase TRIM56 Mus musculus | 2 | 6 | 0.54 | 0.83 |
| Q61510 | E3 ubiquitin/ISG15 ligase TRIM25 Mus musculus | 2 | 8 | 0.51 | 0.84 |
| Q8C522 | Endonuclease domain-containing 1 protein Mus musculus | 0 | 10 | ND in ctrl &ERM | poorly detected PBk11 |
| Q9EQH2 | Endoplasmic reticulum aminopeptidase | 7 | 2 | 1.22 | 0.76 |
| Q9ESY9 | Gamma-interferon-inducible lysosomal thiol reductase Mus musculus | 0 | 1 | 0.15 | 0.43 |
| Q9D8T2 | Gasdermin-D Mus musculus | 1 | 6 | 0.61 | 0.72 |
| Q01514 | Guanylate-binding protein 1 Mus musculus | 0 | 10 | 0.38 | 0.83 |
| P01900 | H-2 class I histocompatibility antigen, D-D alpha chain Mus musculus | 0 | 12 | 2.25 | 1.13 |
| P01902 | H-2 class I histocompatibility antigen, K-D alpha chain Mus musculus | 0 | 4 | 0.81 | 0.75 |
| P17710 | Hexokinase 1 | 0 | 8 | 0.56 | 0.89 |
| P20491 | High affinity immunoglobulin epsilon receptor subunit gamma Mus musculus | 0 | 9 | 5.02 | 1.32 |
| P63158 | High mobility group protein B1 Mus musculus | 0 | 10 | 0.61 | 1.06 |
| P30681 | High mobility group protein B2 Mus musculus | 0 | 12 | 0.5 | 0.83 |

| | | | | | |
|---|---|---|---|---|---|
| Q9JL16 | Interferon-stimulated gene 20 kDa protein Mus musculus | 0 | 0 | 0.37 | 0.48 |
| Q64281 | Leukocyte immunoglobulin-like receptor subfamily B member 4 Mus musculus | 0 | 5 | 3.02 | 1.38 |
| P09581 | Macrophage colony-stimulating factor 1 receptor Mus musculus | 0 | 5 | 0.53 | 0.61 |
| A1L314 | Macrophage-expressed gene 1 protein Mus musculus | 0 | 8 | 11.6 | 2.09 |
| Q9QZ08 | N-acetyl-D-glucosamine kinase Mus musculus | 0 | 9 | 0.32 | 0.67 |
| B2RRE7 | OTU domain-containing protein 4 Mus musculus | 0 | 11 | 0.4 | 0.96 |
| Q9ES52 | Phosphatidylinositol 3,4,5-trisphosphate 5-phosphatase 1 Mus musculus | 0 | 11 | 0.66 | 1.03 |
| Q6P549 | Phosphatidylinositol 3,4,5-trisphosphate 5-phosphatase 2 Mus musculus | 0 | 4 | 0.25 | 0.49 |
| Q2EMV9 | Poly [ADP-ribose] polymerase 14 Mus musculus | 2 | 10 | 0.66 | 0.98 |
| O35286 | Pre-mRNA-splicing factor ATP-dependent RNA helicase DHX15 Mus musculus | 0 | 11 | 0.87 | 0.97 |
| P28063 | Proteasome subunit beta type-8 Mus musculus | 0 | 5 | 0.63 | 1.17 |
| Q60953 | Protein PML Mus musculus | 2 | 4 | 0.36 | 0.65 |
| Q8VCW4 | Protein unc-93 homolog B1 Mus musculus | 0 | 10 | 2.16 | 1.24 |
| Q8C2Q3 | RNA-binding protein 14 Mus musculus | 0 | 6 | 2.76 | 1.22 |
| Q64337 | Sequestosome-1 Mus musculus | 0 | 12 | 12.75 | 1.46 |
| Q9QZI9 | Serine incorporator 3 Mus musculus | 0 | 12 | 2.62 | 0.99 |
| Q9DBU3 | Serine/threonine-protein kinase RIO3 Mus musculus | 1 | 11 | 2.37 | 0.83 |
| Q8BPX9 | Solute carrier family 15 member 3 Mus musculus | 0 | 9 | 3.73 | 1.37 |
| Q9QUN7 | Toll-like receptor 2 Mus musculus | 1 | 10 | 1.48 | 0.81 |
| P58681 | Toll-like receptor 7 Mus musculus | 1 | 3 | 0.36 | 0.41 |
| Q8VC04 | Transmembrane protein 106A Mus musculus | 0 | 5 | 1.91 | 1.36 |
| Q9DBS1 | Transmembrane protein 43 Mus musculus | 2 | 12 | 2.2 | 1.34 |
| Q8BVW3 | Tripartite motif-containing protein 14 Mus musculus | 0 | 11 | 3.14 | 0.69 |
| Q9JMA1 | Ubiquitin carboxyl-terminal hydrolase 14 Mus musculus | 0 | 9 | 0.78 | 1.04 |
| Q5SUA5 | Unconventional myosin-Ig Mus musculus | 0 | 6 | 2.2 | 0.56 |
